# Bioinformatic and Reactivity-Based Discovery of Linaridins

**DOI:** 10.1101/2020.07.09.196543

**Authors:** Matthew A. Georgiou, Shravan R. Dommaraju, Xiaorui Guo, Douglas A. Mitchell

## Abstract

Linaridins are members of the ribosomally synthesized and post-translationally modified peptide (RiPP) family of natural products. Five linaridins have been reported, which are defined by the presence of dehydrobutyrine, a dehydrated threonine residue. This work describes the development of a linaridin specific scoring module for Rapid ORF Description and Evaluation Online (RODEO), a genome-mining tool tailored towards RiPP discovery. Upon mining publicly accessible genomes available in the NCBI database, RODEO identified 561 (382 non-redundant) linaridin biosynthetic gene clusters (BGCs). Linaridin BGCs with unique gene architectures and precursor sequences markedly different from previous predictions were uncovered during these efforts. To aid in dataset validation, two new linaridins, pegvadin A and B, were detected through reactivity-based screening (RBS) and isolated from *Streptomyces noursei* and *Streptomyces auratus*, respectively. RBS involves the use of a reactive chemical probe that chemoselectively modifies a functional group present in the natural product. The dehydrated amino acids present in linaridins as α/β-unsaturated carbonyls were appropriate electrophiles for nucleophilic 1,4 addition using a thiol-functionalized probe. The data presented within significantly expands the number of predicted linaridin BGCs and serves as a road map for future work in the area. The combination of bioinformatics and RBS is a powerful approach to accelerate natural product discovery.

## Introduction

Genome sequencing continues to grow exponentially, providing numerous opportunities for microbial natural product discovery through genome-mining.^1,2^ Automated bioinformatics tools have accelerated the identification of putative natural product biosynthetic gene clusters (BGCs). As our understanding of these pathways improves, the chemical structure of many natural products can be predicted to varying degrees of accuracy through computational analysis.^3^ Such gene-to-structure methods permit a logical approach to prioritize the discovery of novel natural products from uncharacterized BGCs.^4–6^ One natural product family where a gene-to-structure approach is increasingly applied is with the ribosomally synthesized and post-translationally modified peptides (RiPPs).

RiPP natural products have attracted attention for several reasons. Biological activities associated with RiPPs include roles in intercellular signaling, cellular redox metabolism, and therapeutic applications ranging from antimicrobials and analgesics to the treatment of cystic fibrosis.^7^ Mature RiPPs have been shown to contain a diverse set of enzymatically installed post-translational modifications including, but not limited to: dehydration, heterocycle formation [i.e., thiazol(in)e, oxazol(in)e, (dehydro)piperidine, pyridine], various thioether crosslinks, glycosylations, and methylations.^7^ Enzymes involved in RiPP biosynthesis act on a ribosomally synthesized precursor peptide, generally composed of N-terminal “leader” and C-terminal “core” regions. The leader region facilitates the binding of enzymes involved in modifying the core and is removed during maturation of the natural product.^8^ The bipartite nature of RiPP precursor peptides, coupled with often permissive modifying enzymes, make RiPPs attractive targets for analog generation through genetic mutation.^9,10^

Knowledge of enzymes involved in the biosynthesis of RiPPs permits the genomic identification of new members using search algorithms, such as BLAST. Compared with other natural product classes, identification of RiPP BGCs presents a unique challenge: the precursor peptides are often short and hypervariable. Consequently, these precursors are frequently not predicted as protein-coding sequences by automated gene finders unless they are nearly identical to a known sequence.^11^ Without reliable precursor identification, the novelty of a hypothetical RiPP natural product cannot be confidently determined. Therefore, there has been a substantial demand for a computational method to accurately identify RiPP precursor peptides.

The bioinformatics program, Rapid ORF Description and Evaluation Online (RODEO), automates the precursor identification process.^11^ Prior to submission to RODEO, potential members of a RiPP class can be identified through iterative BLAST queries using a protein involved in a known biosynthetic pathway of interest.^12^ The NCBI accession identifiers for these proteins are then provided as input for RODEO, which queries NCBI databases to retrieve the local genomic neighborhood. Functional prediction of the coding sequences is provided by profile hidden Markov models (pHMMs) from the PFAM^13^ and TIGRFAM^14^ databases. The end-user can also supply a custom pHMM library to provide additional gene annotation. RODEO subsequently performs a six-frame translation within the intergenic regions of the genomic neighborhood to identify all potential open-reading frames (ORFs) that may encode the precursor peptide(s). The hypothetical ORFs are then scored through a combination of heuristic scoring, MEME motif analysis,^15^ and support vector machine (SVM) classification, each tailored toward a specific RiPP class. A predetermined scoring threshold separates valid precursors from hypothetical/non-coding sequences. Recent publications have shown that RODEO, alongside other bioinformatic tools, facilitates the identification of new BGCs from a variety of RiPP classes.^11,16–22^ Through the assembly of a comprehensive dataset of BGCs specific to a RiPP class, new insights into the natural product group are gleaned as well as information that can be used to prioritize the discovery of novel members of RiPP class.

Linaridins (linear arid peptides) are an understudied class of RiPPs, with only five characterized members: cypemycin, grisemycin, legonaridin, salinipeptin, and mononaridin (**Figure 1**).^23^ The linaridin RiPP class is defined by the presence of Thr-derived dehydrobutyrines (Dhb) in the final product, although additional tailoring modifications are known.^24^ The enzyme(s) responsible for linaridin Dhb installation remain unconfirmed but are known to be unrelated to those involved in lanthipeptide biosynthesis, where Dhb formation is well established.^25^ Previous work on cypemycin and legonaridin demonstrated that genetic deletion of *cypL, cypH,* or *legG* and *legE* (the two domains of LinH), prevented linaridin formation **(Figure 1**, **Table S1**).^26,27^ All characterized linaridins display *N*^*α*^,*N*^*α*^-dimethylation of the N-terminus through the action of a locally encoded methyltransferase (LinM). Some linaridins are also adorned with aminovinyl cysteine (AviCys) at the C-terminus. AviCys formation in the archetypal linaridin cypemycin involves decarboxylation by a flavin-dependent decarboxylase (CypD); however, enzymes responsible for ring formation have yet to be determined.^28,29^ Other studies have suggested that AviCys biosynthesis in linaridins parallel that found in lanthipeptides.^28^ Subsequent deletions of genes related to these ancillary modifications (i.e. *cypM* and *cypD* within the cypemycin BGC) yielded Dhb-containing linaridins lacking methylation and AviCys, respectively.

**Figure 1.**
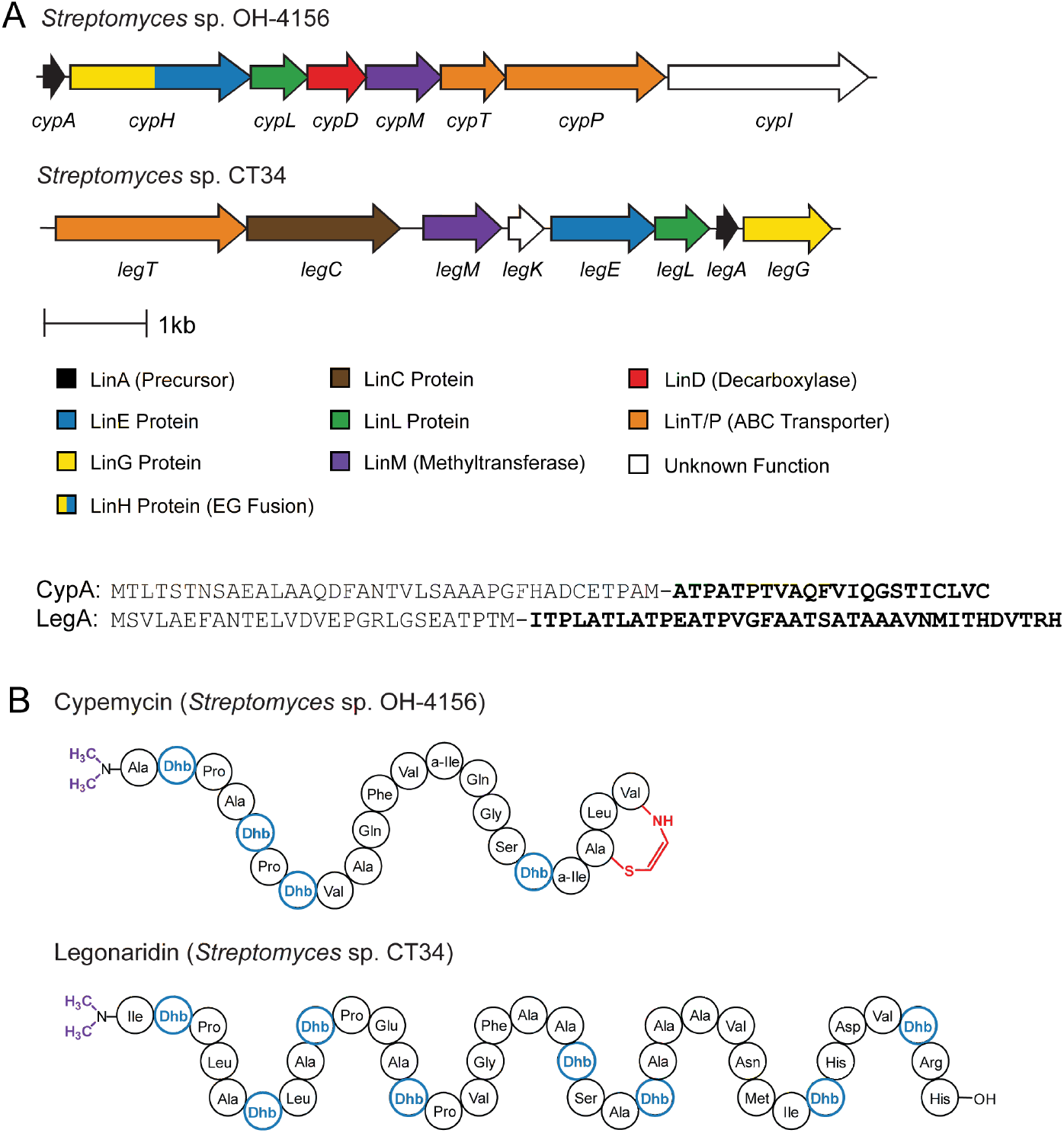
Biosynthetic gene clusters and structures of linaridins. (A) BGCs responsible for cypemycin and legonaridin. The gene names given unify the nomenclature for all linaridins with known functions indicated.^23^ (B) Abbreviated structures of cypemycin and legonaridin. Purple, N-terminal demethylation; Blue, Dhb; Red, AviCys; a-Ile, *allo*-isoleucine.

Linaridins remain underrepresented within the literature. Potential reasons include a poorly understood biosynthetic pathway and that only cypemycin and salinipeptin, exhibit a known bioactivity. Cypemycin shows activity against *Micrococcus luteus* and murine P388 leukemia cells.^30^ Salinipeptin shows modest antibacterial activity against *Streptococcus pyogenes* and is toxic towards U87 giloblastoma and HCT-116 colon carcinoma cells.^31^ Considering the less-studied status of linaridins, along with the class-defining modification, Dhb, being an ideal functional group for RBS-based discovery, we sought to create a comprehensive dataset of all linaridins. This goal was achieved by the development of an automated, linaridin-specific RODEO scoring module. A combination of various bioinformatic methods were used to construct the observable linaridin dataset (561 BGCs, see **Supplemental Methods**). The most-probable precursor peptide(s) from each linaridin BGC were identified using common features gleaned from characterized and high-confidence, predicted linaridins. As in previous reports,^11,16,17,22,23,32^ the RODEO enabled dataset was retrospectively analyzed to extract new insights into the linaridins, including the identification of previously unreported leader region motifs, assessment of the structural diversity of the core region, and broad-scale categorization of potential tailoring enzyme(s). Additionally, our analysis also highlights linaridin BGCs of unusual composition, both in precursor sequence and presence of biosynthetic enzymes, illuminating a subset that differ dramatically from characterized examples. Lastly, leveraging the dataset, an RBS-guided discovery campaign was conducted. A thiol-functionalized probe aided in the isolation and characterization of two new linaridins, pegvadin A and B, from *Streptomyces noursei* NRRL B-1714 and *Streptomyces auratus* NRRL B-8097 respectively, raising the total number of known linaridins to seven.

## Results and Discussion

### Linaridin genome-mining and precursor peptide scoring

A previous publication has suggested a standard nomenclature for linaridin biosynthetic proteins, which has been adopted for the current study **(Figure 1)**.^33^ Gene-deletion studies support that the class-defining Dhb modifications are installed by either LinE, LinG, LinH (a LinE-LinG fusion), or LinL (**Figure 1**).^27^ Therefore, a dataset of all observable homologs of these proteins encoded in RiPP-like genomic contexts was constructed. A multi-round PSI-BLAST^12^ was performed on recognized linaridin biosynthetic proteins until the number of sequences converged (**Supplemental Methods** and **Table S1**). The local genome context was then assessed to determine the presence of these key biosynthetic genes, which are strongly indicative of linaridin production.

The class-defining linaridin biosynthetic protein(s) remain to be experimentally validated and the existing HMMs that define these protein families were found to be insufficient for a broad-scale survey of all linaridins. To expedite the identification of linaridin BGCs, custom pHMMs were generated using HMMER3^34^ for the α/β hydrolase (LinE), transmembrane protein domain (LinG), the LinE-LinG fusion (LinH), and another protein of unknown function (LinL) (**Supplemental Methods** and **Supplemental Dataset 1**). Existing search models for ancillary proteins found within linaridin BGCs were adequate, including the *N*-methyltransferase (LinM, PF13649), ABC transporter (LinT, TIGR02204), and flavin-dependent decarboxylase (LinD, TIGR00521). The custom pHMMs for the core linaridin biosynthetic proteins were used to identify the presence or absence of each gene, aiding in the rapid identification of complete linaridin BGCs. A final dataset containing 561 linaridin BGCs with 382 non-redundant members was compiled (**Supplemental Datasets 2-3**). This dataset significantly expands upon a recent linaridin genome-mining study.^33^

A phylogenetic tree was produced for sequences encoding distinct LinE domains, theorized to be involved in Dhb formation (**Figures S1-S2**).^26,35^ Isolation of LinE-like domains was achieved by a computational script utilizing the LinE pHMM (**Supplemental Methods** and **Supplemental Note S1**). LinE domains from type A linaridin BGCs formed a distinct clade as did LinE domains from type B BGCs. Such co-evolution of biosynthetic enzymes and precursor peptides is well-documented within RiPP BGCs.^11^ Therefore, the precursors from BGCs from these two clades were adequate for use as a training set for a RODEO scoring module for linaridins.

Analysis of precursors from the type A and type B linaridin clades identified key features (**Figure S2**), including a PxxxTP motif used to determine the potential leader peptide cleavage site, noted in previous publications.^33,35^ Other features of interest included gene directionality, distance from key biosynthetic genes, and core hydrophobicity, where >55% of core residues were hydrophobic (considering Thr being converted to Dhb). A heuristic scoring scheme was devised using with these characteristics and applied to all potential precursor peptides (**Table S2**). When combined with a custom LinA pHMM and support vector machine (SVM) classification (**Supplemental Dataset 1** and **Supplemental Methods**), the linaridin scoring module identified probable precursor peptide(s) with maximum precision and recall statistics achieved at a threshold score of 12. Thus, sequences scoring 12 or higher are predicted to be valid linaridin precursor peptides (**Supplemental Figures S3-S4**).

### Comparison of RODEO to other bioinformatic tools

The final list of candidate linaridin precursor peptides found by RODEO (**Supplemental Datasets 2-3**) was cross-referenced against previous predictions reported in the literature.^19,21,23,36^ Comparison to the most recent linaridin-specific dataset published that contained 204 BGCs^33^ shows that the linaridin RODEO module identified 294 of the 303 predicted precursor peptides, returning an average score of 29.5 (**Table S3** and **Figure S5**). All nine of the reported precursor peptides that were not identified by RODEO contain highly irregular sequences. Many display a non-canonical RAVSTP motif near the predicted leader-core junction and/or a polybasic regions near the C-terminus (**Supplemental Dataset 4**). Of the nine, RODEO identified high-scoring linaridin precursor peptides for four cases, leading us to believe the previously reported peptides are misidentified.

Another report produced a generalized RiPP dataset utilizing the bioinformatic tool “DeepRiPP”, predicting a total of 135 linaridin precursor peptides (86 unique sequences).^19^ When compared to the RODEO-derived dataset, only 76 of the 135 (~56%) were confidently predicted as linaridin precursor peptides (**Table S3**). Of the 135 DeepRiPP-predicted linaridin precursor peptides, only 26 are reported to be encoded next to known linaridin biosynthetic proteins with the remaining 109 precursors being reported as distantly encoded (i.e. not within the BGC).^19^ Unfortunately, DeepRiPP concludes that the cypemycin and legonaridin precursor peptides are not encoded within the BGC; however, this is clearly erroneous (**Figure 1**). Furthermore, the 26 precursor peptides classified as within the BGC are actually not adjacent to linaridin biosynthetic genes, rather they occur near genes associated with other RiPP classes, including the thiopeptides. Contrary to the report, 93 of the 109 precursor peptides reported as distantly encoded are found within 3 kb of a linaridin biosynthetic gene, which was evident upon cross-referencing with the RODEO-derived dataset (**Supplemental Dataset 4**).

A final comparison was drawn against datasets from other RiPP discovery tools, namely NeuRiPP,^18^ MetaMiner,^36^ and RiPPMiner^37^ (**Table S3**). 94% of precursor peptides identified by NeuRiPP, and the single linaridin identified by MetaMiner, were also present in the RODEO dataset. Of the 13 precursor peptides found by RiPPMiner, 9 lacked key linaridin BGC features, such as omission of a LinE homolog (from the entire genome), the predicted precursor reported cannot be found in the local region, or the putative precursor peptide scored below the validity threshold. Only four members of the RiPPMiner dataset judged to be valid by the RODEO module. The sequences deemed invalid by RODEO we believe are misidentified (**Supplemental Dataset 4**). These comparisons show that the linaridin module of RODEO enabled the compilation of the most comprehensive and accurate dataset of linaridin BGCs, with each entry containing at least the minimum genes required for linaridin biosynthesis. Additionally, the module demonstrates greater efficacy in identifying precursor peptide sequences compared to automated gene finders, as nearly 10% of the linaridin precursor peptides identified by RODEO were not annotated as protein-coding sequences.

### Content analysis of linaridin precursor peptides and BGCs

Linaridin structural diversity is dependent on both primary sequence and modification by tailoring enzymes.^26,31,40,41^ A gene encoding an ABC transporter, LinT, co-occurs in 96% of linaridin BGCs and is presumed to be involved in compound export. The most widespread, co-occurring linaridin enzyme responsible for ancillary modification is the methyltransferase, LinM, found in 70% of BGCs (**Table S4**). LinM-dependent N-terminal dimethylation is vital for cypemycin bioactivity.^26^ Statistical analysis of the first amino acid of the precursor core region (+1 position) in LinM-containing BGCs most frequently appears as Ala, Gly, Leu, or Phe; however, when considering average codon usage, there is a clear enrichment for Ala, and to a lesser extent, Phe. Other large aromatic residues (Trp, Tyr), charged residues (Asp, Glu, Lys, Arg), and some polar residues (Asn, Gln) and have no representation at core position +1; further, three additional residues (i.e. His, Thr, and Val) are significantly depleted relative to the expected residue frequency (**Table S5**). Analysis of the residue identity for core position +1 in BGCs lacking a LinM homolog reveals a markedly different picture. Although Ala remains significantly enriched, Cys and Thr are also very common at position +1, comprising 86% of the total sequences.

Other co-occurring tailoring enzymes include the flavin-dependent decarboxylase, LinD, present in 9% of BGCs. The decarboxylase acts at the C-terminus of the core and plays an indirect role in the formation of the AviCys moiety.^26,28,29^ Genes encoding LinD co-occur exclusively with precursor peptides ending in a terminal CxxC motif. The final, prevalent co-occurring gene, present in 40% of BGCs, encodes a protein showing distant homology to a FAD-dependent desaturase (TIGR02734). Termed “*linC*” there is no identified function for this gene despite its widespread localization to linaridin BGCs, including those in cluster 1, although it is report as being essential for legonaridin expression.^27^

RODEO identified 568 high-confidence precursor peptides from 382 non-redundant linaridin BGCs. Several BGCs contain multiple precursor peptides (**Figure S6**), with a maximum of six unique sequences in *Leifsonia xyli* subsp. *xyli*. To assess diversity of the linaridin precursor peptides, a sequence similarity network (SSN) was generated (**Figure 2**). Of the 568 precursor peptides identified by RODEO, 457 had unique primary sequences. Within the dataset, only groups 1 (type A) and 2 (type B) contained characterized linaridins, meaning 87% of precursor peptides predicted by RODEO are substantially different compared to isolated linaridins. To better visualize the dominant sequence trends in each of these groups, an alignment sequence logo^42^ was produced for the leader and core and regions of each group of the SSN containing >4 members (**Figures S7-S8**).

**Figure 2.**
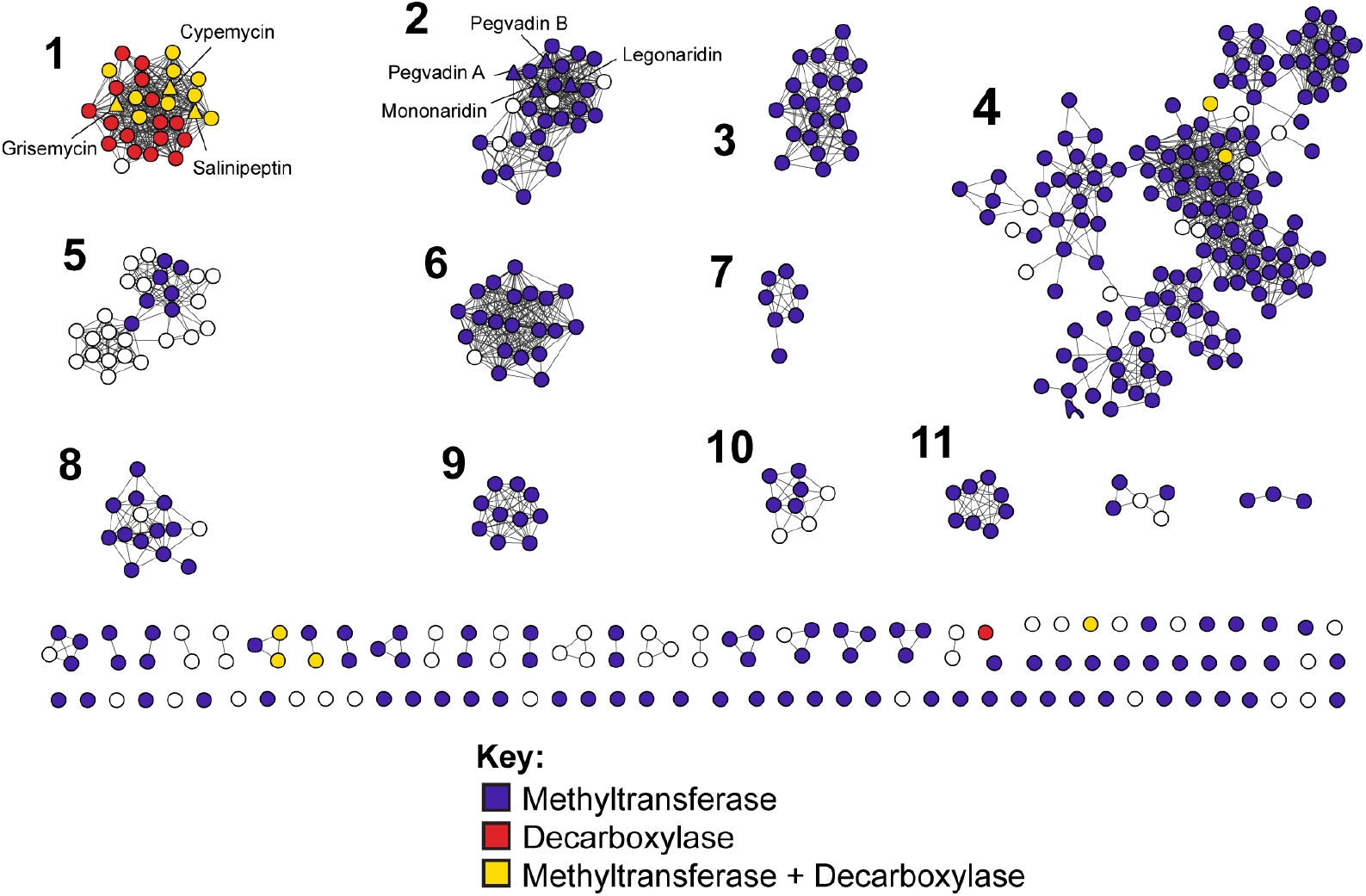
Sequence similarity network of linaridin precursors. Nodes within the SSN are color based on the co-occurrence of a LinM (purple) a, LinD (red), or both (yellow). Groups containing >5 members are numbered. Nodes representing known linaridins are shown as triangles and annotated. This SSN was generated using EFI-EST^38^ and visualized with Cytoscape^39^. Protein sequences are conflated at 100% identity (i.e. identical sequences are only represented once), resulting in 457 nodes. Edges indicate an alignment score of 7 (expectation value of <10^−7^). Phylogenetic tree information for LinE homologs is available in Supplemental Dataset 5.

Research on RiPP biosynthesis has demonstrated that conserved sequences within the leader region, correspond to key recognition motifs for biosynthetic proteins.^43,44^ Within linaridins, the foremost conserved sequence is found at the predicted cut-site (**Figure S9**).^35^ Due to this high level of conservation, this motif may play a role in leader peptidase binding. Further, various amino acid sequences are shown to be conserved within the leader region (**Figure S7**). Most notably, groups 1 and 2 display a conserved FANxxL motif, which had been observed in previous alignments but not specifically commented on in previous work.^35^ As LinM and LinD enzymes are predicted to be leader peptide-independent,^27,28^ this conserved motif may facilitate the recognition or binding of an enzyme(s) involved in the conversion of Thr to Dhb. Further experimental analysis will be required to test this hypothesis.

The core regions of linaridin precursor sequences also show varying levels of conservation, but there is a broad diversity between the groups (**Figure S9**). The length of the core region was assessed, ranging from 14 to 80 amino acids in length (**Figure S10**). This range is considerably wider than characterized linaridins, which vary from 19 to 37 core residues.^26,31,40,41^ Within the predicted linaridin core, the position and abundance of Thr residues is highly variable, yet all known examples convert all Thr to Dhb. While unmodified Ser has been observed in all characterized linaridins, mutational studies on cypemycin show that replacement of Thr with Ser led to dehydroalanine (Dha) formation,^35^ suggesting that dehydration is site-specific. To determine the maximum number of potential Dhb moieties in yet uncharacterized linaridins, Thr content was assessed and found to range from 1 to 13 (mean = 3.1 ± 1.9 Thr per core sequence, **Figure S11**). Factoring in the potential for Dha moieties to be found in future examples, Thr+Ser content was calculated, which ranged from 1 to 22 (mean = 4.2 ± 2.9). Taking these variables into account, the linaridin class of RiPPs contains much greater sequence diversity than what is represented by known members.

### Identification of BGCs with unusual architecture

The bioinformatic survey of linaridins revealed various BGCs containing unusual gene architectures. One striking observation was repeated with three bacteria: *Streptomyces* NRRL F-2890, *Streptomyces xiamensis*, and *Streptomyces atratus*. These BGCs contain the three key linaridin biosynthetic genes (LinE, LinG, and LinL) in an apparent hybrid BGC with a LanB dehydratase (PF04738) and LanC cyclase (PF05147), known from class I lanthipeptide biosynthesis (**Figure S12**). The precursors for these linaridin BGCs have leader and core regions that more closely resemble a class I lanthipeptide than any known than linaridin, with a polyanionic leader region and Cys residues present throughout the core.^22^ To determine the potential cut-sites, these precursor peptides were provided as input for RODEO with the module for lanthipeptide detection selected and manually adjusted.^22^ In combination with the lanthipeptide dehydratase, the plausible role for the LinE, LinG, and LinL homologs is Dha/Dhb formation.

All but two linaridin BGCs are encoded by Actinobacteria, despite this phylum accounting for less than 10% of genomes within NCBI (**Figures S1, S13-14**). The two exceptions are from *Kroppenstedtia sanguinis* (Firmicutes)^45^ and *Burkholderiales* bacterium PBB3 (Proteobacteria). The *K. sanguinis* BGC contains LinE, LinG, LinL, and LinT homologs, alongside another ABC transporter, and a gene of unknown function annotated as a metallopeptidase protein (PF01551). The most probable linaridin precursor peptide from *K. sanguinis* lacks several traits associated with known linaridins and thus was judged “invalid” by RODEO. Notably, the sequence contains 11 Ser residues. This abundance of Ser in the precursor may indicate a Dha-rich linaridin, which is without precedence. The BGC found in *Burkholderiales* bacterium PBB3 contains a LinH homolog, a potential decarboxylase, but lacks a LinL homolog (**Figure S13**). Future work is warranted on these non-actinobacterial linaridin BGCs.

Other precursors with unusual core sequences were also identified. A notable trend was the presence of core Cys residues without a co-occurring LinD homolog (**Supplemental Datasets 2-3**). Unmodified thiol groups on Cys are rare in mature natural products; therefore, it is plausible that such positions undergo additional post-translational modification. Cys content was calculated for all linaridin core regions from non-redundant precursor peptides ranging from 0-3, with a mean distribution of 0.4 ± 0.7 (**Figure S15**). A particularly unique subset of Cys-containing precursors was found in *Saccharomonospora* sp. which display core regions predicted to begin with Cys followed by a sequence rich in dehydratable Thr/Ser residues (**Table S6**). These unusual BGCs and precursors further suggest that linaridins characterized in the literature represent only a small subset of those encoded in nature.

### New linaridin discovery through reactivity-based screening

RBS involves the detection of natural products through the chemoselective targeting of a specific functional group. An RBS-based strategy for natural product discovery is associated with several advantages such as: prioritization of novelty through bioinformatic analysis, screening can be performed on cellular extracts, and compound detection is agnostic to biological activity. Covalent modification observed by comparative mass spectrometry enables the rapid and sensitive detection of natural products bearing the targeted functional group.^46^ The class-defining Dhb modifications in linaridins are Michael acceptors susceptible to 1,4-nucleophilic addition.^47^ This reactivity of dehydrated amino acids has been previously harnessed to map the location of amino acids within a target compound,^48^ identify lanthipeptides within cultures,^49^ and in semi-synthetic modification of thiostrepton.^16,46,50^ Dithiothreitol (DTT) was selected as the Michael donor owing to its commercial availability, reactivity, and literature precedence (**Supplemental Methods**)^46^ with labeling events yielding a 154 Da mass increase. Detection of labeling, along with high-resolution and tandem mass spectrometry (HR-MS/MS), allows identification of the corresponding parental mass, which can be readily cross-referenced with existing literature to assess novelty.

The compiled dataset identified 34 available strains harboring potentially new linaridin BGCs (**Table S7**). Target organisms were cultivated, the bacterial cells were harvested, and a methanolic cell-surface extraction was performed on all candidates. Cell lysis was avoided based on the assumption that the culture medium would be less complex than the bacterial cytosolic fraction and that the mature linaridins would be exported from the producing cell. The cell-surface extracts were then reacted with DTT supplemented with a hindered base (**Supplemental Methods**). Both unreacted and reacted extracts were analyzed by matrix-assisted laser desorption/ionization time-of-flight mass spectrometry (MALDI-TOF-MS) to identify compounds in the extract and subsequent labeling events. Upon completion of the screen, extracts from *Streptomyces noursei* NRRL B-1714 and *Streptomyces auratus* NRRL B-8097 contained masses corresponding to predicted linaridins. These natural products labeled up to three times upon reaction with DTT (**Figure 3**). With Dhb representing a sterically hindered Michael acceptor, incomplete labeling was observed under conditions that would show quantitative labeling for Dha residues.^46^

**Figure 3.**
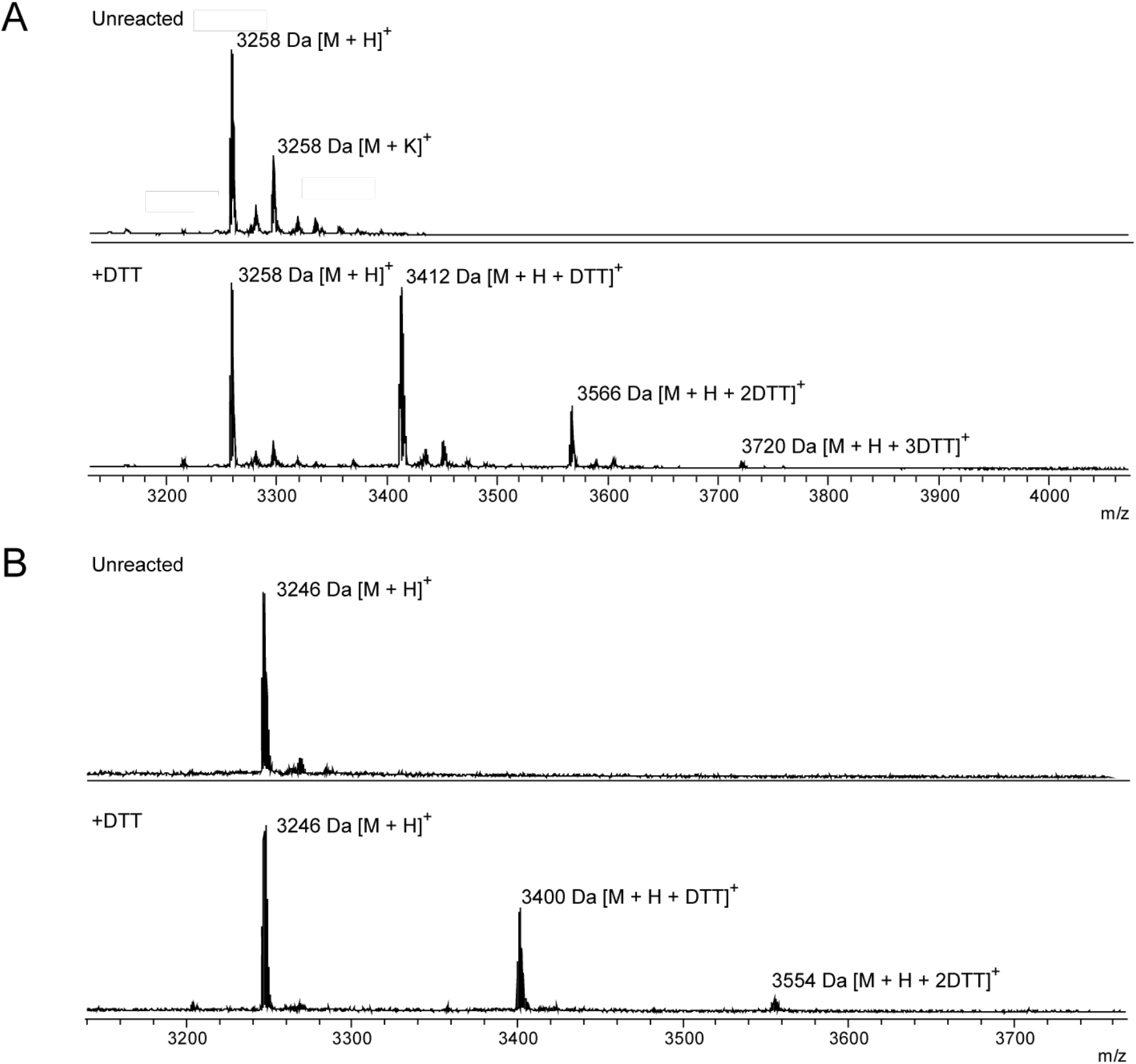
Reactivity-based screening of novel linaridins. MALDI-TOF mass spectra of unreacted (*top*) and DTT-labeled (*bottom*) for (A) *S. noursei* and (B) *S. auratus* extract.

The target compound from each strain was purified using reversed-phase high-performance liquid chromatography (HPLC) over several different gradients (**Supplemental Methods**). Each putative linaridin product was then analyzed by HR-MS/MS, which confirmed the molecular formula and peptidic sequence for both compounds. The product from *S. noursei* had an observed mass ([M + 2H]^2+^ = 1629.3778 Da) corresponding to C_150_H_239_N_39_O_42_^2+^ ([M + 2H]^2+^ = 1629.3804; error = 1.6 ppm) whereas the compound isolated from *S. auratus* had an observed mass ([M + 2H]^2+^ = 1623.3868 Da) corresponding to a molecular formula of C_149_H_239_N_39_O_42_^2+^ ([M + 2H]^2+^ = 1623.3882; error = 0.9 ppm, **Tables S8-S9**). These formulas correspond with bioinformatically predicted linaridin core peptides, where all Thr residues have been converted to Dhb and *N^α^,N^α^*-dimethylation of the N-terminus has taken place (**Figure 4**). Collision-induced dissociation revealed a series of daughter ions matching the expected linaridin core sequence. These data confirmed the presence and location of the post-translational modifications. (**Figure S16**, **Tables S8-S9**).

**Figure 4.**
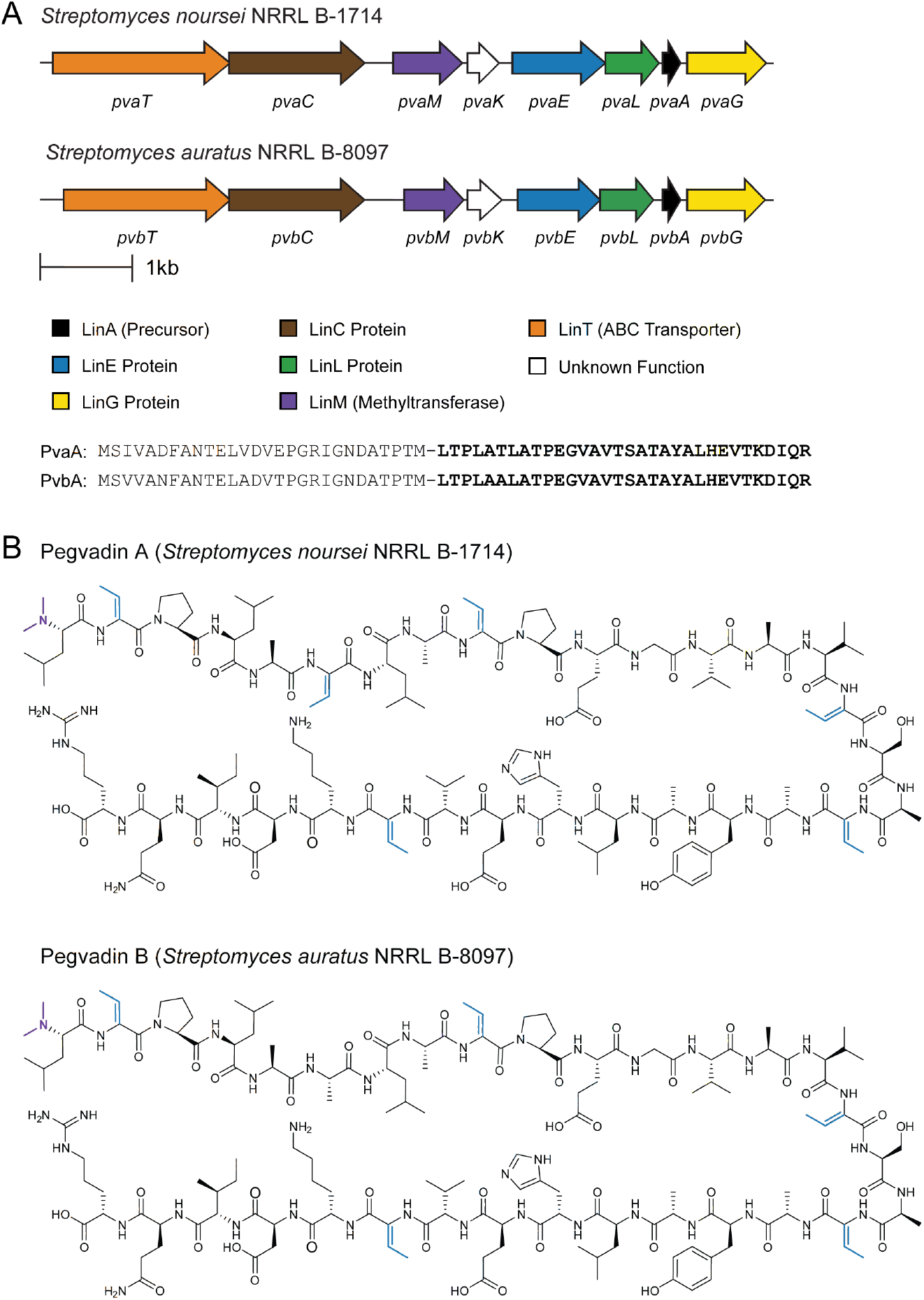
BGC and structure of pegvadin A and B. (A) BGCs for pegvadin A and B. Specific gene names are noted, derived from Pegvadin A (Pva) and Pegvadin B (Pvb) respectively. (B) Abbreviated structures of pegvadin A and B with the (Z) configuration for Dhb and natural stereochemistry for the proteinogenic amino acids shown.

### Comparison of pegvadin A & B to characterized linaridins

The two newly discovered compounds, pegvadin A and B, are type B linaridins (**Figures S2** and **4**)^35^. The overall architecture of the pegvadin BGCs resembles that of legonaridin and mononaridin. The leader regions of the known type B linaridins all share a conserved FANxxL motif with pegvadin A and B sharing the highest overall sequence similarity with legonaridin. The core regions also share some commonalities, as the N-terminal region of the pegvadin A and legonaridin core sequences only differ at position 1 (Leu versus Ile). However, the remainder of the pegvadin core regions are substantially different compared to other linaridins (**Figure 4**). Compared to legonaridin and mononaridin, the pegvadins have shorter (32-residue) core regions and fewer Dhb residues. Finally, the pegvadins display *N*^*α*^,*N*^*α*^-dimethylation of a Leu residue, which previously has only been observed on Ile (legonaridin), Ala (cypemycin), and Val (mononaridin). Leu at core position +1 is predicted to occur in <5% of linaridin BGCs that include a LinM homolog (**Table S5**).

## Conclusion

Despite a number of reports over the past decade, the linaridins remain an underexplored RiPP class. This dearth of knowledge is exemplified by the fact that there are only a few characterized members, the putative protein(s) responsible for the class-defining modification (i.e. Dhb installation) remain to be biochemically validated, and the biological role of the mature compounds remains speculative. To facilitate future work on the linaridins, we developed an automated genome-mining method specific for the molecular class. During the construction of the RODEO scoring module, pHMMs were created for each of the genes known to play a role in linaridin biosynthesis, allowing rapid identification of potential linaridin BGCs. Bioinformatic analysis of the expanded set of linaridin BGCs led to new insights, including rare linaridin BGCs encoded beyond the Actinobacteria, hybrid BGCs containing both lanthipeptide and linaridin biosynthetic enzymes, and trends relating methyltransferase co-occurrence to the identity of the first core position. Also, the linaridin sequence-function space was significantly expanded, indicating that isolated members are representative of only two precursor subclasses, containing less than a quarter of all currently identifiable linaridins.

Leveraging the dataset provided by the linaridin detection module of RODEO, a targeted set of potential linaridin producing organisms were cultivated. Using the inherent electrophilicity of its class-defining Dhb residues, RBS a thiol-functionalized probe enabled the discovery of two new linaridins, pegvadin A and B. The structure of these linaridins was confirmed through HR-MS/MS. Our data illustrate a plethora of undiscovered linaridins remain, which provide numerous organisms and enzymes for future biosynthetic, natural product isolation, and biological function studies.

## Supporting information

Supplemental Information Doc

Supplementary Dataset 1_HMMs

Supplementary Dataset 2_RODEO-tabular

Supplementary Dataset 3_RODEO-graphical

Supplementary Dataset 4_bioinformatics_comparison

Supplementary Dataset 5_LinE_phyloXML

## ASSOCIATED CONTENT

Additional experimental data, Figures S1−S16, and Tables S1−S11 (PDF).

- Supplemental Dataset 1: custom profile Hidden Markov Models (ZIP)
- Supplemental Dataset 2: tabular RODEO output (XLSX)
- Supplemental Dataset 3: graphical RODEO output (PDF)
- Supplemental Dataset 4: comparison of RODEO to other bioinformatic tools (XLSX)
- Supplemental Dataset 5: PhyloXML data for the phylogenetic tree generation (TXT)

## Notes

The authors declare no competing financial interest.

## ACKNOWLEDGEMENT

We would like to acknowledge G. Hudson, for his revision and editing suggestions for the manuscript. We further thank A. Kretsch for her compilation of data related to *Streptomyces* sp. codon frequency.

