## Supplemental Information Doc for "Bioinformatic and Reactivity-Based Discovery of Linaridins"

### Table of Contents

Experimental Methods……………………………………………………..………………………….S2
Table S1: NCBI accession list of linaridin biosynthetic genes ……………………………….……..S6
Figure S1: Phylogenetic distribution of LinE enzymes……………………………….……………..…S7
Figure S2: Phylogenetic relation of LinE proteins to known members..…………………………..S8
Note S1: Parsing script for pHMM-based sequence excision..………..…………………………..S9
Table S2: Features and weights used in RODEO linaridin module scoring……………………..S10
Figure S3: Precursor peptide scoring distributions. ……………………..…………………...……S11
Figure S4: Evaluation of scoring procedure for precision and recall……………………….……S12
Table S3: Comparison of RODEO scoring to other bioinformatic tools…..………….……….……S13
Figure S5: RODEO scoring of previously identified linaridin precursor peptides………….……S14
Table S4: Co-occurring genes that appear in linaridin BGCs………………………….…….…..…S15
Table S5: Identity of first amino acid in core region……………….………………………………S16
Figure S6: Number of precursor peptides per BGC………………………………………………...S17
Figure S7: Sequence analysis of linaridin leader region……………………………………………S18
Figure S8: Sequence analysis of linaridin core region………………………………..……………S19
Figure S9: Sequence analysis of linaridin leader/core cut-site………………………….…………S20
Figure S10: Linaridin core region length variability. ……..…………………………………..……S21
Figure S11: Thr and Ser content of predicted core region. ……….………………………..……S22
Figure S12: Hybrid Lanthipeptide and Linaridin BGCs. …………………………..………..…..S23
Figure S13: Non-Actinomycete BGCs……………………………………………………………S24
Figure S14: Phylogenetic distribution of NCBI sequences……….………………………………S25
Figure S15: Cys content of predicted core peptides……..……………..…………………….…...S26
Table S6: Linaridin core regions containing Cys.……………………………..……………………S27
Table S7: List of bacterial strains evaluated for linaridin production ……….……..…..…………S28
Table S8: Pegvadin A MS/MS ion assignments.……………….…..……………………..……....…S29
Table S9: Pegvadin B MS/MS ion assignments..……………………..…………………..………..…S30
Figure S16: High-resolution and tandem mass spectrometry of pegvadins A and B…………..S31
Table S10: Sequence identity/similarity of proteins in linaridin BGCs……….…………………...S32
Table S11: Screening media used and their components…………………………………………S33
Supplemental References…………………………………………………………………………S34

### Experimental Methods

**General.** Materials and reagents were purchased from Gold Biotechnology, Fisher Scientific, or Sigma Aldrich unless otherwise noted. Matrix-assisted laser desorption/ionization time-of-flight mass spectrometry (MALDI-TOF-MS) analysis was performed using a Bruker UltrafleXtreme MALDI-TOF-TOF mass spectrometer (Bruker Daltonics) in reflector positive mode at the University of Illinois School of Chemical Sciences Mass Spectrometry Laboratory. High-resolution and tandem MS analyses were performed using a ThermoFisher Scientific Orbitrap Fusion instrument equipped with an Advion TriVersa Nanomate 100.

**Bioinformatic mining of linaridin BGCs.** To identify potential linaridin biosynthetic enzymes, PSI-BLAST[1] (Position-Specific Iterative - Basic Local Alignment Search Tool) searches were conducted in May 2020 on all predicted key linaridin biosynthetic enzymes (**Table S1**) with a 0.05 expectation value threshold. Upon removing redundancies, four iterations retrieved 8,000 protein sequences, which were then submitted to the automated ribosomally synthesized post-translationally modified peptide (RiPP) genome-mining platform, Rapid ORF Description & Evaluation Online (RODEO; http://ripp.rodeo)[2]. Custom profile Hidden Markov Models (pHMMs) used are provided in **Supplemental Dataset 1** while tabular and graphical RODEO output are provided in **Supplemental Datasets 2-3.** These data were then further analyzed by determining co-localization to homologs of other genes found in the identified linaridin BGCs, particularly genes annotated as linaridin precursors. Initial compilation of accessions produced a dataset of 320 potential linaridin BGCs. Additional linaridin BGCs were identified using the HMMER[3] hmmsearch web tool provided by the EMBL-EBI website (https://www.ebi.ac.uk/).[3] This webtool enabled the use of custom pHMMs to search for LinA and LinL homologs, and results from the search were compared to the original dataset. After BGCs with identical genes and from the same species were removed, 382 linaridin BGCs were cataloged in the final dataset.

*Gene annotation using custom HMMs*. To aid in bioinformatic discovery, custom pHMM were utilized. Protein sequences were aligned using the Multiple Alignment using/by Fast Fourier Transform (MAFFT).[4] These alignments were converted into pHMMs using HMMER3.[3] pHMMs were generated for alignments of key biosynthetic genes that were not identified using pre-existing models within the PFAM and TIGRFAM collections.[5,6] Five custom pHMMs were thus generated for the linaridin-specific RODEO module: LinA, LinE, LinG, LinL, and LinH. To enable pHMM construction, a list of all non-redundant biosynthetic proteins was generated, including manually separating LinH fusion enzymes from their discretely encoded homologs. This information, when used in combination with RODEO, aided in identifying diverse gene clusters whose gene products retrieved no matches within the PFAM/TIGRFAM databases.

*Linaridin precursor scoring.* A set of heuristics for linaridin precursor peptide identification were created (**Table S2**) and then applied to the initial dataset with scoring determined by the relative importance of the trait and its frequency. Sequence motifs identified by the MEME-FIMO[7] tool were included in the list of heuristic traits.

Heuristics alone did not yield clear separation of manually predicted "valid" linaridin precursors from that of unrelated hypothetical sequences (**Figure S3**). The application of a support vector machine (SVM) module aided the separation of predicted linaridin precursors from non-linaridin sequences. The addition of a LinA custom pHMM homology metric, with an e-value cutoff of 0.001, also substantially aided separation between predicted linaridin precursors and non-linaridin sequences. The HMM metric and SVM classification captured overlapping but nonidentical subsets of linaridin precursors, so upon the combination of the two with the list of heuristics, the score distribution for predicted linaridins maximally separated from the scoring distribution for other hypothetical sequences (**Figure S3**).

**Sequence similarity network generation.** The sequences of precursor peptides within the dataset of potential linaridins were used to construct a sequence similarity network (SSN).[8] This network allowed the separation of series based on a pre-set alignment score cut off, enabling the assessment of trends within these groupings. All SSNs were generated using the Enzyme Function Initiative Enzyme Similarity Tool (EFI-EST)[9] (https://efi.igb.illinois.edu/). Visualization was performed using organic layout within Cytoscape.[10]

**Phylogenetic tree generation.** Phylogenetic trees were generated from the LinE domains of LinE and LinH protein sequences acquired from NCBI accession identifiers. LinE domains were selected from both LinE and LinH homologs with a newly developed parsing script **(Supplemental Note S1)**. The script uses a custom pHMM in combination with a FASTA file as input for an HMMER3 hmmsearch command,[3] and in this case, the LinE custom pHMM and a FASTA file with all LinE and LinH proteins were used as input. The script parses the hmmsearch output to identify the homologous region to the pHMM from each sequence in the FASTA file. The output is a multiple sequence alignment-ready FASTA file with the aforementioned homologous regions. Multiple sequence alignment was performed by MAFFT.[4] The tree was built using FastTree[11] software using the JTT+CAT amino acid revolution model. For tree visualization, we used the interactive Tree of Life (iTOL) website (http://itol.embl.de/).[12] **Supplemental dataset 5** includes a phyloXML formatted text file for regenerating the LinE phylogenetic tree.

**Biosynthetic gene analysis.** Interpro[13] (https://www.ebi.ac.uk/interpro) and HHPred[14] (https://toolkit.tuebingen.mpg.de/tools/hhpred) were used to identify discernable protein domains within the three key biosynthetic genes. As in previous studies, it was determined that the LinE protein resembled protein domains corresponding to an $\alpha/\beta$ hydrolase protein, the LinG protein resembles a transmembrane protein, and the LinL protein contains both a signal peptide domain and non-cytoplasmic domain.[15]

**Bacterial cultivation.** Eight media for Actinomycetes were screened, including: AGS, ATCC 172, Alternative Mannitol-Soy, C-Food, GUBC, ISP2, ISP4, and V8.[16] Recipes for each of medium can be found in **Table S11**. Previous studies have indicated that growth on different media containing different carbon sources and metal ion concentrations can elicit the production of secondary metabolites.[17] Target strains were initially grown on ATCC 172 media (pH 7.0). Plates were incubated at 30 °C until sporulation was evident or until 14 d had passed. Next, liquid cultures of ATCC 172 (pH 7.2) were inoculated with spores or single colonies and allowed to grow at 30 °C in a drum roller (40 rpm) for 7 d. Finally, the bacterial culture (100 µL) was inoculated onto each screening medium. Each sample was cultivated at 30 °C for 10 d and 18 d.

**Reactivity-based labeling of extracts.** Cells from plates were harvested using a sterile razor blade and placed into a 1.7 mL microfuge tube. MeOH (1 mL) was added to each extraction individually with gentle rocking at 25 °C for 18 h. The MeOH (cell-surface) extraction from each strain was subjected to reactivity-based labeling using a thiol-functionalized probe. Cellular debris was removed from the extractions by centrifugation (17,000 × g, 10 min, 25 °C). The soluble fraction was then reacted with 100 mM of dithiothreitol (DTT) and 10 mM *N,N*-

diisopropylethylamine for 4 h and 24 h at 25 °C. The samples were then analyzed by MALDI-TOF-MS.

**MALDI-TOF-MS.** Matrix-assisted laser desorption/ionization time-of-flight mass spectrometry (MALDI-TOF-MS) enabled the identification of peaks of interest when compared to predicted masses. Samples were spotted on a stainless-steel MALDI target with 50% MeCN saturated with α-cyano-4-hydroxycinnamic acid with 0.1% (*v/v*) formic acid. Sample analysis was performed using a Bruker Daltonics UltrafleXtreme MALDI-TOF mass spectrometer in reflector positive mode. After reactivity-based labeling of extracts, this technique allowed the identification of DTT adducts (addition of 154 Da per label).

**Large-scale production and purification of pegvadin A and B.** Pegvadin A: *Streptomyces noursei* NRRL B-1714 was grown on Alt-MS medium with 1.5% (0.6 mg/mL) agar on 10 cm sterilized dishes (n = 160) at 30 °C for 11 d. Pegvadin B: *Streptomyces auratus* NRRL B-8097 was grown on ATCC 172 medium with 1.5% (0.6 mg/mL) agar on 10 cm sterilized dishes (n=160) at 30 °C for 11 d. Sporulation was observed on the third day in both cases. Cells were harvested with a sterile razor blade and collectively extracted with 700 mL of MeOH with shaking at 25 °C for 3 h. The methanolic extract was vacuum-filtered, and the remaining cell extract was resuspended in 600 mL of MeOH. The solution then gently rocked at 25 °C for 3 h. This extraction process was repeated twice, the methanolic fractions were combined, and 10g of Celite 545 adsorbent was added. This mixture was dried using a rotary evaporator.

Pegvadin adsorbed onto the Celite was purified by using a Teledyne Isco Combiflash EZprep equipped with a RediSep Rf C18 cartridge (130 g media, 60 Å pore size, 40-63 μm particle size, 230-400 mesh) using a mobile phase of 10 mM aq. $NH_4HCO_3$ in MeCN at 75 mL $min^{-1}$ over a gradient from 20-70%. Wavelengths at 220 nm and 280 nm were monitored to determine which fractions contained peaks of interest. These fractions were further analyzed via MALDI-TOF-MS before being pooled and dried by rotary evaporation.

Reverse phase purification was performed using a Teledyne Isco Combiflash EZprep equipped with a RediSep Prep C18Aq column (100 Å pore size, 40-63 um particle size, 230-400 mesh) using a mobile phase of 10 mM aq. $NH_4HCO_3$/MeCN with a gradient from 20% to 60% MeCN over 25 min at 19 mL/min. Fractions containing pegvadin were determined through MALDI-TOF analysis. These fractions were combined and dried using a rotary evaporator. A semi-preparative HPLC was then performed using a Perkin Elmer Flexar HPLC equipped with a Betasil C18 (Thermo Scientific) reverse phase column. Pegvadin was dissolved in 8 mL of 10 mM aq. $NH_4HCO_3$/MeCN (70:30), the suspension was then clarified by centrifugation (17,000 x g for 20 min, 25 °C) before injection. Pegvadin was purified under the following conditions: solvent A ($NH_4HCO_3$), solvent B (MeCN) and the following gradient: t=0 min, 30% B, t=7 min, 35% B, t=40 min, 38% B, t=50 min, 95% B, t=55 min, 30% B. The column was equilibriated at starting conditions for 15 min before each injection. Following second purificaitons, samples containing pegvadin were once again combined. A final, analytical, purification was performed using the same equipment with an analytical Betasil C18 (Thermo Scientific) reverse phase column. Pegvadin was dissolved and clarified once again, as detailed above. Final purification was performed under the following conditions: solvent A ($NH_4HCO_3$), solvent B (MeCN) and the following gradient: t=0 min, 30% B, t=10 min, 39% B, t=15 min, 40% B, t=20 min, 45% B, t=26 min, 50% B, t=30 min, 50% B, t=35 min, 30% B. Purified pegvadin was intially dried under rotary evaporation at 25 °C and then dried to completion using a SpeedVac vacuum concentrator.

**HR-ESI-MS/MS analysis of pegvadin A and B.** Pegvadin A and B were desalted using a ZipTip and eluted into 75% aq MeCN supplemented with 0.1% acetic acid. Samples were directly infused onto a ThermoFisher Scientific Orbitrap Fusion ESI-MS using an Advion TriVersa Nanomate 100. MS calibration was performed with Pierce LTQ Velos ESI Positive Ion Calibration Solution (ThermoFisher). The MS was operated using the following parameters: 100,000 resolution, 2 *m/z* isolation width (MS/MS), 70 normalized collision energy (MS/MS), 0.4 activation q value (MS/MS), and 30 ms activation time (MS/MS). Fragmentation was performed using collision-induced dissociation (CID) at normalized collision energy. Data analysis was conducted using the Qualbrowser application of Xcalibur software (ThermoFisher Scientific).

**Table S1: NCBI accession list of linaridin biosynthetic genes.** The NCBI accession identifiers of all genes referenced directly in the main text. Associated PFAM and TIGRFAM models as referenced as well as any custom pHMM for sensitive detection of sequence homology.

| Protein | PFAM/TIGRFAM/pHMM | NCBI Accession ID |
|---|---|---|
| CypA | LinA | ADR72962.1 |
| CypH | LinH | ADR72963.1 |
| CypL | LinL | ADR72964.1 |
| CypD | Decarboxylase (TIGR00521) | ADR72965.1 |
| CypM | Methyltransferase (PF13649) | ADR72966.1 |
| CypT | ABC Transporter (TIGR02857) | ADR72967.1 |
| CypP | Unknown | ADR72968.1 |
| CypI | DUF255 (PF03190) | ADR72969.1 |
| LegT | ABC Transporter (TIGR02204) | WP_052230046.1 |
| LegC | Desaturase (TIGR02734) | WP_043270372.1 |
| LegM | Methyltransferase (PF13649) | WP_107068144.1 |
| LegE | LinE | WP_078894000.1 |
| LegF | LinL | WP_078894692.1 |
| LegA | LinA | WP_107068145.1 |
| LegH | LinG | WP_043265482.1 |
| GrmA | LinA | WP_003970682.1 |
| GrmH | LinH | BAG23194.1 |
| GrmL | LinL | BAG23193.1 |
| GrmD | Decarboxylase (TIGR02857) | BAG23192.1 |
| GrmM | Methyltransferase (PF13649) | BAG23191.1 |
| GrmT | ABC Transporter (TIGR02857) | BAG23190.1 |
| GrmP | Unknown | BAG23189.1 |
| SinA | LinA | WP_101256403.1 |
| SinH | LinH | WP_101256404.1 |
| SinL | LinL | WP_101256405.1 |
| SinD | Decarboxylase (TIGR00521) | WP_101256406.1 |
| SinM | Methyltransferase (PF13649) | WP_101256407.1 |
| SinT | ABC Transporter (TIGR02857) | WP_101256408.1 |
| SinP | Unknown | WP_101256409.1 |
| MonA | LinA | WP_030019228.1 |
| MonT | ABC Transporter (TIGR02204) | WP_078624127.1 |
| MonC | Desaturase (TIGR03467) | WP_050502300.1 |
| MonM | Methyltransferase (PF13649) | WP_030019231.1 |
| MonK | Helix-turn-helix (PF13518) | WP_078624119.1 |
| MonE | LinE | WP_078624126.1 |
| MonL | LinL | WP_050502306.1 |
| MonG | LinG | WP_030019227.1 |
| PvaA | LinA | ANZ17121.1 |
| PvaT | ABC Transporter (TIGR02204) | ANZ17127.1 |
| PvaC | Desaturase (TIGR03467) | ANZ17126.1 |
| PvaM | Methyltransferase (PF13649) | ANZ17125.1 |
| PvaK | Helix-turn-helix (PF13518) | ANZ17124.1 |
| PvaE | LinE | ANZ17123.1 |
| PvaL | LinL | ANZ17122.1 |
| PvaG | LinG | ANZ17120.1 |
| PvbA | LinA | WP_099053532.1 |
| PvbT | ABC Transporter (TIGR02204) | EJJ06683.1 |
| PvbC | Desaturase (TIGR02734) | EJJ06682.1 |
| PvbM | Methyltransferase (PF13649) | EJJ06681.1 |
| PvbK | Helix-turn-helix (PF13518) | EJJ06680.1 |
| PvbE | LinE | EJJ06679.1 |
| PvbL | LinL | EJJ06678.1 |
| PvbG | LinG | EJJ06677.1 |

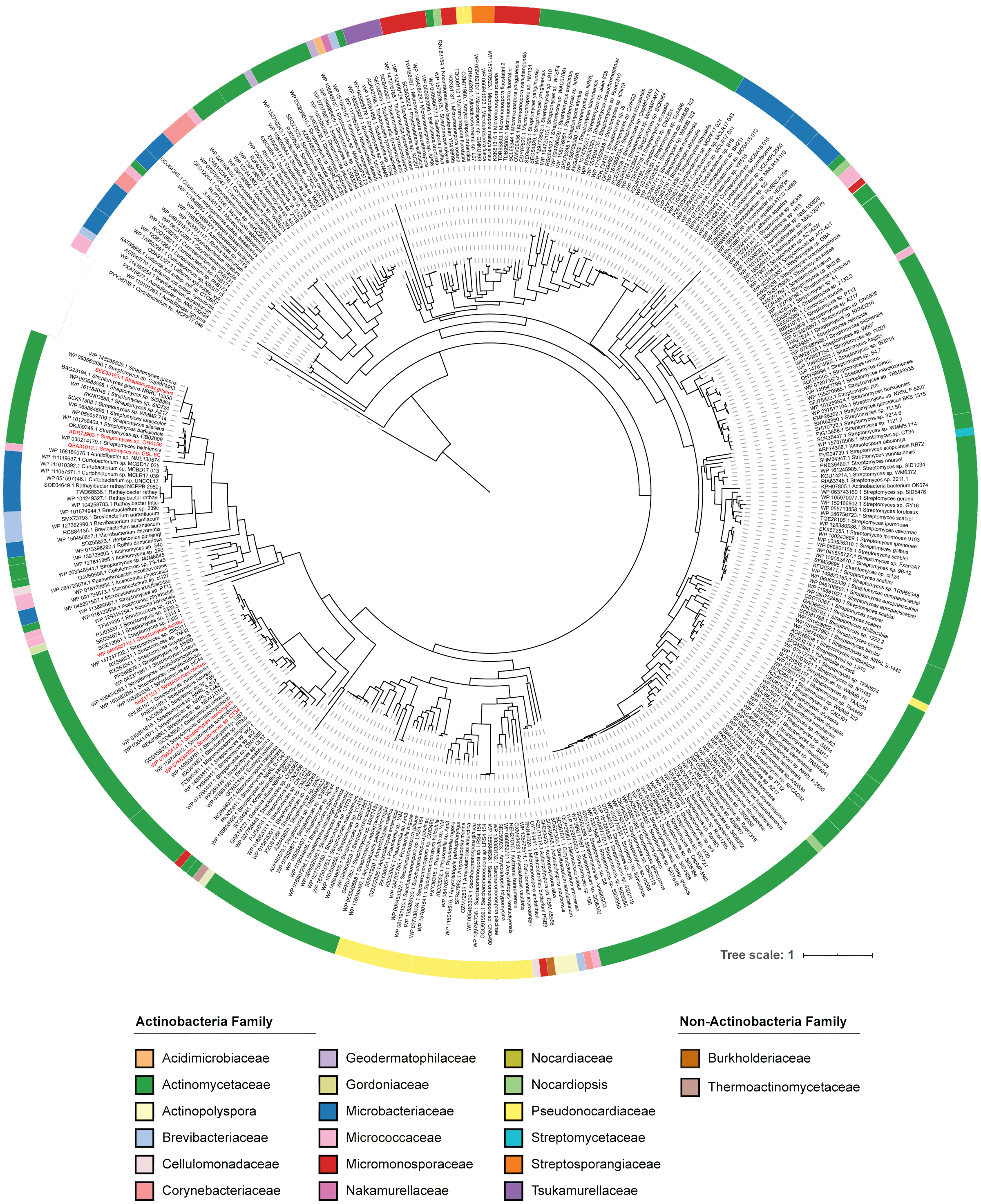


**Figure S1: Phylogenetic distribution of LinE protein domains.** Maximum likelihood tree of unique LinE sequences (n = 382; 100% identical sequences have been removed) colored by taxonomic family. Known linaridin producers are colored in red. The tree was rooted using a predicted SdpB homolog (PYY38796.1)[15] and visualized using interactive Tree of Life (iTOL). The phylogenetic classification of each identified LinE enzyme is listed in **Supplemental Dataset 2**. The data to regenerate this tree are available in phyloXML format (**Supplemental Dataset 5**).

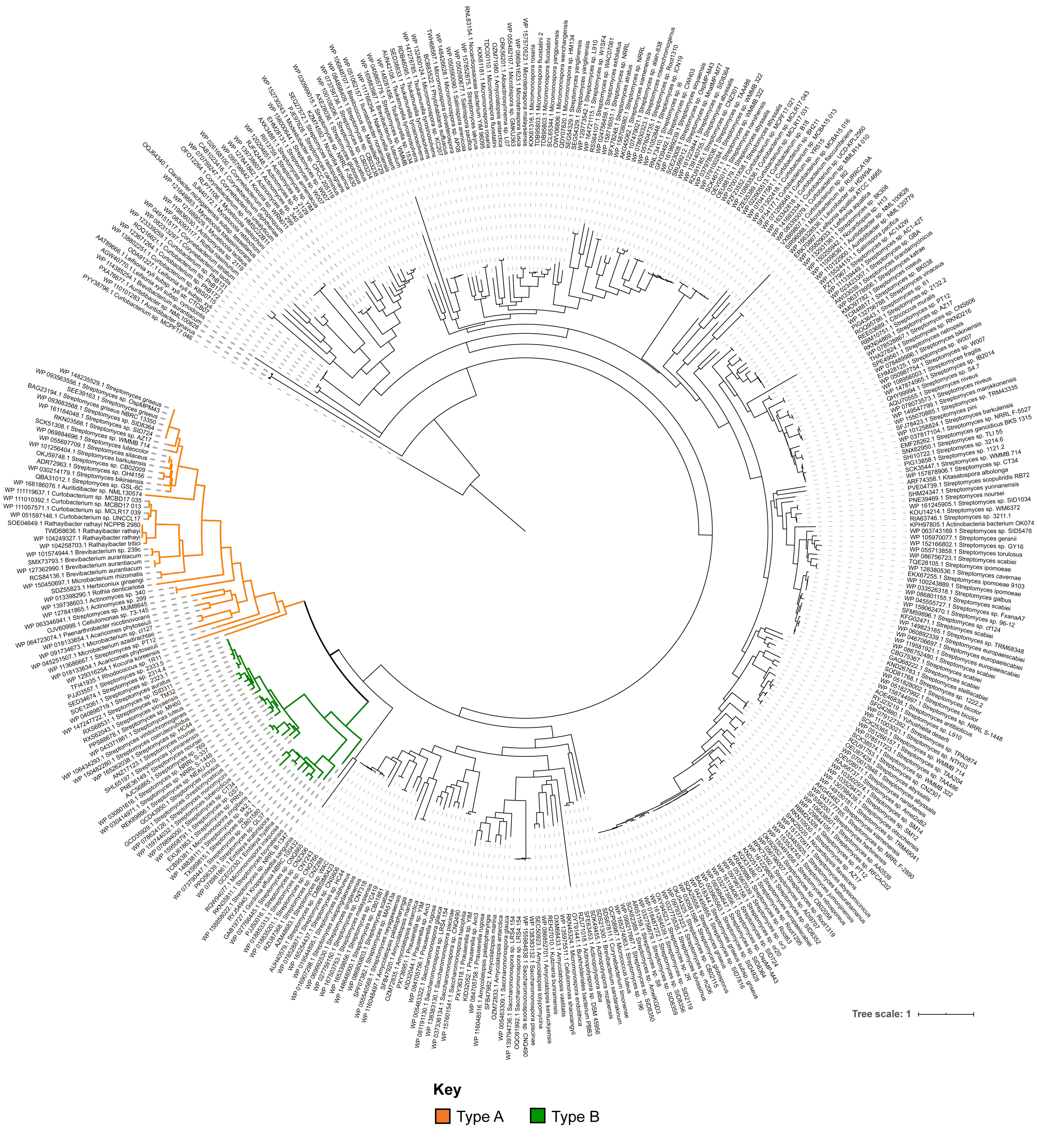


**Figure S2. Phylogenetic relation of LinE proteins to known members.** Maximum likelihood tree of unique LinE sequences (n = 382; 100% identical sequences have been removed). LinE domains were obtained using an alignment to the custom LinE pHMM. The tree is identical to that shown in Figure S1; however, rather than showing taxonomy, the clades with with type A and type B linaridins are highlighted.[15]

**Note S1: Parsing script for pHMM-based sequence excision.** The Python script, hmmexcise.py, takes a pHMM and a FASTA file as input. The input pHMM file must be pressed by hmmpress before use. The script then performs an hmmscan and prints a FASTA file with excised domains homologous to the pHMM. Optional '-bs' flag is provided to specify alternate inclusion thresholds (default BitScore is 25.0)

```
import argparse
import sys
import os
import re
import subprocess

def __main__():

# Run using the following command: python3 hmmexcise.py <hmm_file> <input_fasta_file>

    parser = argparse.ArgumentParser("hmmexicse.py Excises sequence with homology to HMM file")
    parser.add_argument('hmm_file', type=str)
    parser.add_argument('input_fasta', type=str)
    parser.add_argument('-bs', '--bitscore', type=float, default=25)
    # optional -bs flag if other bitscore cutoffs are wanted

    args = parser.parse_args()

# Subprocessing to run hmmscan within the program. Opens a text file that contains hmmscan output.

    process = subprocess.Popen(['hmmscan', '-o' + args.input_fasta.split('.')[0] + '_hmmscan.txt',
args.hmm_file, args.input_fasta])
    process.wait()
    fileoutput = args.input_fasta.split('.')[0] + '_excised.txt'
    hmmscan_output = args.input_fasta.split('.')[0] + '_hmmscan.txt'

# Opens hmmscan output file and reads the data. Splits the file by query and accesses
# the top hmm hit. Evaluates the BS of the match between each domain and the hmm file
# and prints each domain that hits the top hmm with Bit Score of at least 25

    hmmscanfile = open(hmmscan_output, 'r')
    FO = open(fileoutput, "w")
    HSO = re.split("Query:", hmmscanfile.read())[1:]
    i=0     # i used for optional counting in FASTA headers
    for query in HSO:
        lines = re.split("\n", query)
        accession = re.split(" +", lines[0])[1]
        hits = re.split(">+", query)
        organism = re.split("\[", lines[1])[1]
        if ">>" in query:
            HMMhit = re.split(" ", hits[1])[1]
            domains = re.split("==", hits[1])
            for domain in range(1, len(domains)):
                domainlines = re.split(accession[len(accession)-5:], domains[domain])
                domainBS = re.split(" ", domains[domain])[5]
                if float(domainBS) < args.bitscore:
                    continue
                sequence = ""
                for line in range(1, len(domainlines)):
                    sequence += re.split(" +", domainlines[line])[2]+"\n"
                # i+=1
                FO.write(">"+accession+"_"+organism+"\n"+sequence)
        else:
            print(re.split(" +", lines[0])[1])

if __name__=="__main__":
    __main__()
```

**Table S2: Features and weights used in the RODEO linaridin module scoring.** Heuristics analyzing hydrophobicity in the core region also include Thr because of the anticipated formation of Dhb.

| | Feature | Weight |
|---|---|---|
| Heuristic Scoring | ABC transporter (PF00005 or PF02163 or PF00664) present in BGC | +1 |
| | Methyltransferase (PF08241 or PF08242 or PF13489 or PF13649 or PF13847) present in BGC | +1 |
| | Flavin decarboxylase (TIGR00521 or TIGR02113 or PF02441) present in BGC | +1 |
| | CXXC sequence motif present in core region with flavin decarboxylase present in BGC | +1 |
| | GST sequence motif present in leader with flavin decarboxylase present in BGC | +1 |
| | LinE homolog present in BGC | +2 |
| | Precursor is <300 nucleotides of a LinE, LinL, LinG, or LinH homolog | +1 |
| | Precursor is <600 nucleotides of a LinE, LinL, LinG, or LinH homolog | +1 |
| | Precursor is >2200 nucleotides from a LinE, LinL, LinG, or LinH homolog | -1 |
| | Precursor length between 50 and 70 amino acids (inclusive) | +2 |
| | Precursor length between 71 and 100 amino acids (inclusive) | +1 |
| | Precursor length between 120 and 150 amino acids (inclusive) | -5 |
| | Precursor length >150 amino acids | -10 |
| | Leader region isoelectric point at pH 7 < +1 | +1 |
| | Core region isoelectric point at pH 7 > +6 or < -6 | -1 |
| | Core region contains >15% Ala | +1 |
| | Core region contains >20% Ala | +1 |
| | Core region contains >10% Thr | +1 |
| | Core region contains >15% Thr | +1 |
| | Core region contains 0 Thr | -2 |
| | Core region contains >7% Val | +1 |
| | Core region contains >12% Val | +1 |
| | Core region contains >2 Cys | -1 |
| | Core region contains 0 Cys | +1 |
| | Core region contains <50% hydrophobic residues (G, A, V, L, M, I, T) | -1 |
| | Core region contains >60% hydrophobic residues (G, A, V, L, M, I, T) | +1 |
| | Core region contains >75% hydrophobic residues (G, A, V, L, M, I, T) | +1 |
| | Core region begins with XTP sequence motif | +1 |
| | Leader region contains GxG motif | +1 |
| | Leader region contains LxD motif | +1 |
| | Leader region contains FAN motif | +1 |
| | Peptide contains any MEME/FIMO identified sequence motifs | +2 |
| | Peptide contains no MEME/FIMO identified sequence motifs | -1 |
| +HMM | Peptide has homology to LinA custom HMM at e-value < 0.001 | +10 |
| +SVM | SVM classifies as valid | +10 |

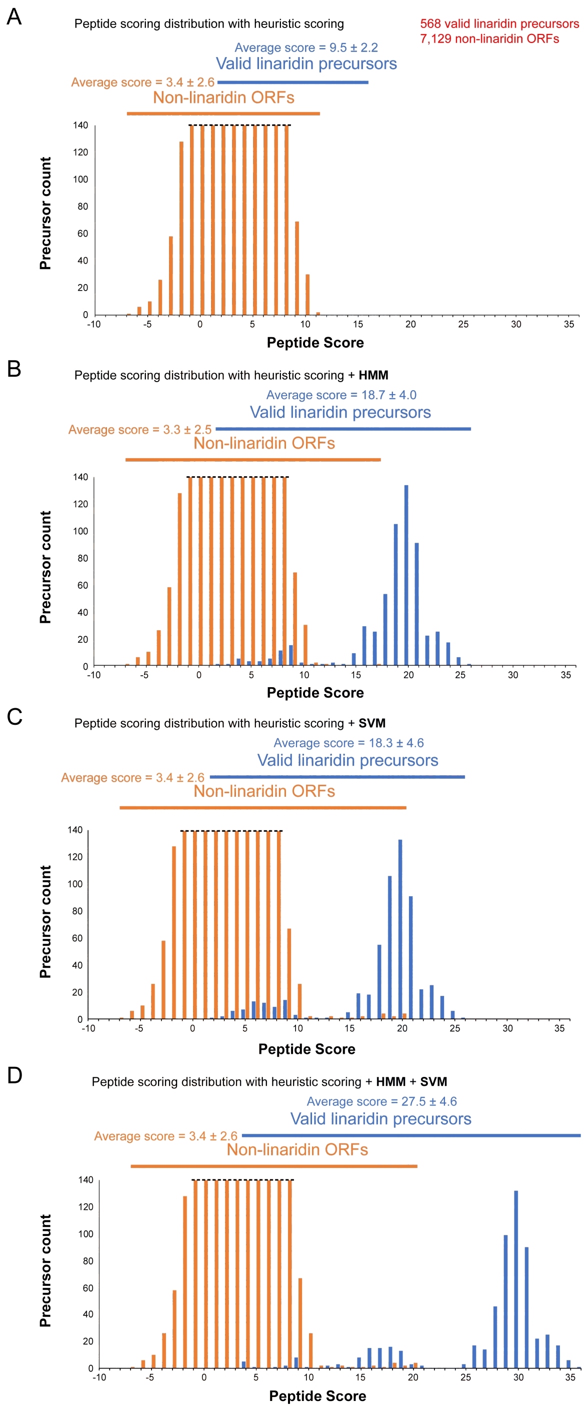


**Figure S3: Precursor peptide scoring distributions.** Precursor peptide score distributions are shown for the finalized linaridin dataset. (A) Score distribution is shown using heuristics only (including MEME/FIMO-identified motif scoring). (B) Score distribution is shown when distribution in Panel A is supplemented with support vector machine (SVM) learning classification. (C) Score distribution is shown when distribution in Panel A is supplemented with the LinA custom pHMM homology metric. (D) Score distribution is shown when distribution in Panel A is supplemented with SVM classification and the LinA pHMM homology metric simultaneously.

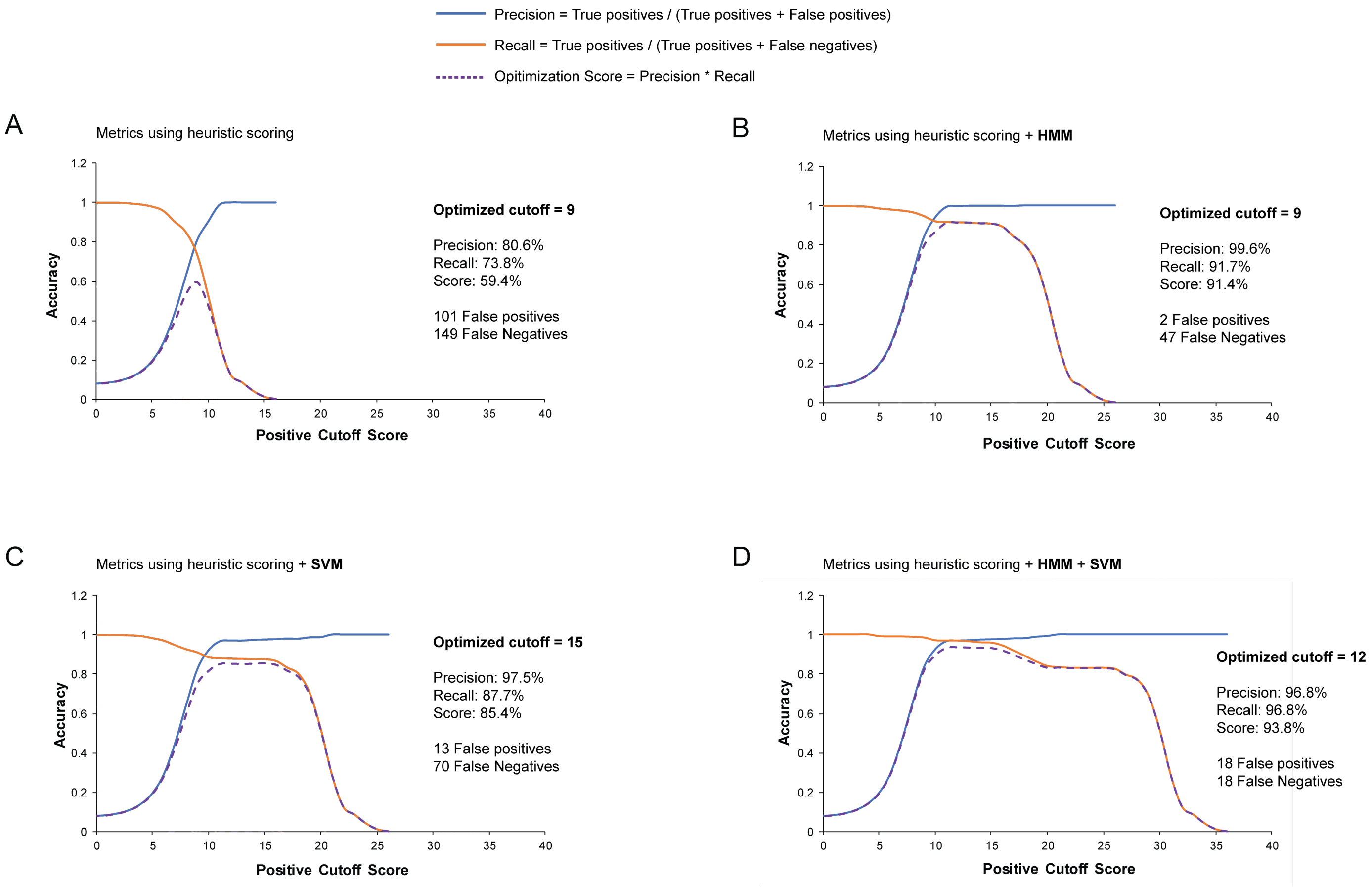


**Figure S4: Evaluation of precision and recall for linaridin precursor scoring.** Precision and recall statistics are shown for four scoring procedures used to separate identified linaridin precursor peptides from other non-linaridin predicted sequences. The optimization score describes how a given scoring cutoff minimizes false-negative and false-positive classifications. (A) Metrics using only heuristic scoring. (B) Metrics using HMM homology and heuristic scoring. (C) Parameters using support vector machine (SVM) classification and heuristic scoring. (D) Metrics using SVM classification, HMM homology, and heuristic scoring.

**Table S3: Comparison of RODEO scoring to linaridins identified from earlier bioinformatic studies.** Published datasets of other RiPP discovery tools were assessed for predicted linaridins. These were cross-referenced with the RODEO dataset produced in this paper. If common to both datasets, the average RODEO score was taken. Datasets with < 90% confirmed by RODEO include a number of predicted false-positives, addressed in the main text. **Supplemental Dataset 5** contains a full listing of previous bioinformatically identified linaridins.

| RiPP Mining Tool (publication year) | Precursor peptides identified | Confirmed by RODEO (%) | Average RODEO score of predicted precursors | Range of RODEO scores of predicted precursors |
|---|---|---|---|---|
| Ma and Zhang (2020)[15] | 303 | 294 (97%) | 29.5 | 14 - 36 |
| NeuRiPP (2019)[18] | 34 | 32 (94%) | 31.8 | 29 - 35 |
| DeepRiPP (2020)[19] | 135 | 76 (56%) | 31.2 | 17 - 36 |
| MetaMiner (2019)[20] | 1 | 1 (100%) | 34 | 34 |
| RiPPMiner (2020)[21] | 13 | 4 (31%) | 30.5 | 30 - 31 |
| Current work | 568 | 100% | 27.8 | 12 - 36 |

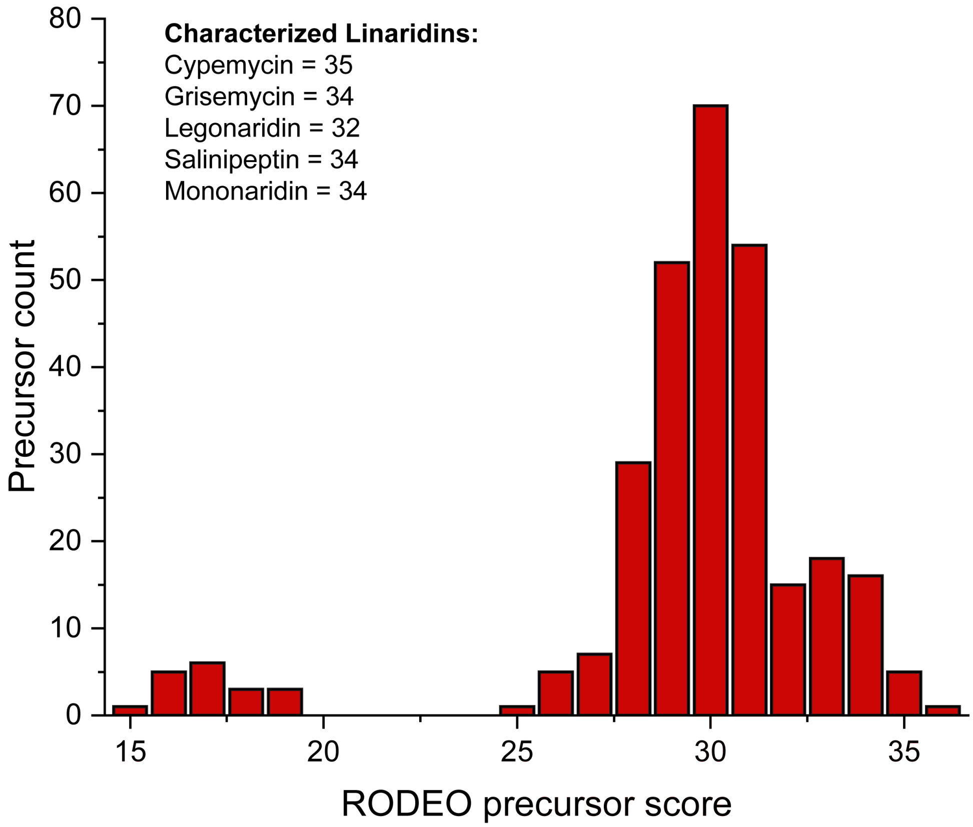


**Figure S5: RODEO scoring of previously predicted linaridin precursor peptides.** Histogram showing the score of predicted precursors from Ma and Zhang 2020 dataset,[15] using the linaridin scoring module. The score of known precursor peptides, which were used in the training set, are shown in the top left.

**Table S4: Co-occurring genes that appear in linaridin BGCs.** The most abundant genes appearing in linaridin BGCs are listed in descending order. Percent co-occurrence indicates the relative frequency a protein sequence within the BGC matches a pHMM. Eight genes were predicted in both directions from the query; only the best scoring (lowest e-value) pHMM was included. Proteins were compared against PFAM, TIGRFAM, and custom pHMMs (noted in red). Custom pHMMs are available in **Supplemental Dataset 1** as well as the RODEO webtool (http://ripp.rodeo) and RODEO Github (https://github.com/the-mitchell-lab/rodeo2). Co-occurrence statistics were obtained from the RODEO-derived dataset. Not all co-occurring protein families are necessarily involved in linaridin biosynthesis. However, some lower co-occurring entries, such as the decarboxylase (TIGR00521, yellow highlight), are associated with linaridin biosynthesis (AviCys formation).[22] Overlap between LinH and LinE/G pHMMs is observed in some cases. Additionally, some BGCs contain more than one predicted linaridin precursor, further increasing percent co-occurance.

| pHMM ID | pHMM Name | pHMM Description | Count | % co-occurrence |
|---|---|---|---|---|
| **LinA** | **LinA** | **LinA (linaridin precursor)** | **522** | **136%** |
| **LinL** | **LinL** | **LinL Homolog** | **386** | **101%** |
| **LinH** | **LinH** | **LinH (Fusion of LinE/LinG)** | **292** | **76%** |
| PF13649 | Methyltransf_25 | Methyltransferase domain (LinM homolog) | 268 | 70% |
| TIGR02204 | TIGR02204 | MsbA_rel: ABC transporter, permease/ATP-binding protein | 206 | 54% |
| **LinE** | **LinE** | **LinE Homolog** | **158** | **41%** |
| TIGR02734 | TIGR02734 | crtI_fam: phytoene desaturase | 154 | 40% |
| **LinG** | **LinG** | **LinG Homolog** | **113** | **30%** |
| PF00196 | GerE | Bacterial regulatory proteins, luxR family | 87 | 23% |
| PF00072 | Response_reg | Response regulator receiver domain | 68 | 18% |
| TIGR01188 | TIGR01188 | drrA: daunorubicin resistance ABC transporter, ATP-binding protein | 62 | 16% |
| PF07730 | HisKA_3 | Histidine kinase | 59 | 15% |
| PF01061 | ABC2_membrane | ABC-2 type transporter | 50 | 13% |
| PF01551 | Peptidase_M23 | Peptidase family M23 | 37 | 10% |
| **TIGR00521** | TIGR00521 | coaBC_dfp: phosphopantothenoylcysteine decarboxylase / phosphopantothenate--cysteine ligase | 36 | 9% |
| PF03704 | BTAD | Bacterial transcriptional activator domain | 33 | 9% |
| PF12698 | ABC2_membrane_3 | ABC-2 family transporter protein | 31 | 8% |
| PF12840 | HTH_20 | Helix-turn-helix domain | 30 | 8% |
| PF09335 | SNARE_assoc | SNARE associated Golgi protein | 29 | 8% |
| PF04134 | DUF393 | Protein of unknown function, DUF393 | 28 | 7% |
| PF06738 | ThrE | Putative threonine/serine exporter | 27 | 7% |
| PF00719 | Pyrophosphatase | Inorganic pyrophosphatase | 25 | 7% |
| PF12680 | SnoaL_2 | SnoaL-like domain | 25 | 7% |
| PF02113 | Peptidase_S13 | D-Ala-D-Ala carboxypeptidase 3 (S13) family | 23 | 6% |
| PF05719 | GPP34 | Golgi phosphoprotein 3 (GPP34) | 23 | 6% |
| PF00583 | Acetyltransf_1 | Acetyltransferase (GNAT) family | 21 | 5% |
| TIGR02857 | TIGR02857 | CydD: thiol reductant ABC exporter, CydD subunit | 21 | 5% |
| PF13560 | HTH_31 | Helix-turn-helix domain | 21 | 5% |
| TIGR02203 | TIGR02203 | MsbA_lipidA: lipid A export permease/ATP-binding protein MsbA | 21 | 5% |
| TIGR03883 | TIGR03883 | DUF2342_F420: uncharacterized protein, coenzyme F420 biosynthesis associated | 20 | 5% |
| PF01906 | YbjQ_1 | Putative heavy-metal-binding | 20 | 5% |
| TIGR01189 | TIGR01189 | ccmA: heme ABC exporter, ATP-binding protein CcmA | 20 | 5% |

**Table S5: Identity of first amino acid in core region.** The amino acid identity at position +1 of the core region of predicted linaridins is given as a function of whether the BGC encodes or omits a a linaridin methyltransferase (LinM) homolog. The last column indicates the expected frequency for each amino acid based on codon usage in *Streptomyces* sp. where the majority of linaridins are encoded. Values underlined in red text indicate a significant enrichment after factoring in codon usage frequency.

| | **LinM Present** | | **LinM Absent** | | **Expected Frequency in *Streptomyces* sp.** |
|---|---|---|---|---|---|
| **Position +1** | **# Cores** | **%** | **# Cores** | **%** | **%** |
| A | 345 | **72.9** | 29 | **30.5** | 13.7 |
| C | 1 | 0.2 | 15 | **15.8** | 0.8 |
| D | 0 | 0 | 0 | 0 | 5.9 |
| E | 0 | 0 | 0 | 0 | 5.7 |
| F | 22 | **4.7** | 1 | 1.1 | 2.7 |
| G | 44 | 9.3 | 2 | 2.1 | 9.6 |
| H | 1 | 0.2 | 0 | 0 | 2.3 |
| I | 5 | 1.1 | 1 | 1.1 | 3 |
| K | 0 | 0 | 0 | 0 | 2.1 |
| L | 22 | 4.7 | 3 | 3.2 | 10.3 |
| M | 6 | 1.3 | 0 | 0 | 1.6 |
| N | 0 | 0 | 0 | 0 | 1.7 |
| P | 0 | 0 | 0 | 0 | 6.3 |
| Q | 0 | 0 | 1 | 1.1 | 2.7 |
| R | 0 | 0 | 0 | 0 | 8.3 |
| S | 15 | 3.2 | 3 | 3.2 | 5.1 |
| T | 5 | 1.1 | 38 | **40.0** | 6.1 |
| V | 7 | 1.5 | 2 | 2.1 | 8.4 |
| W | 0 | 0 | 0 | 0 | 1.5 |
| Y | 0 | 0 | 0 | 0 | 2.0 |
| **Total** | **473** | **100** | **95** | **100** | **100** |

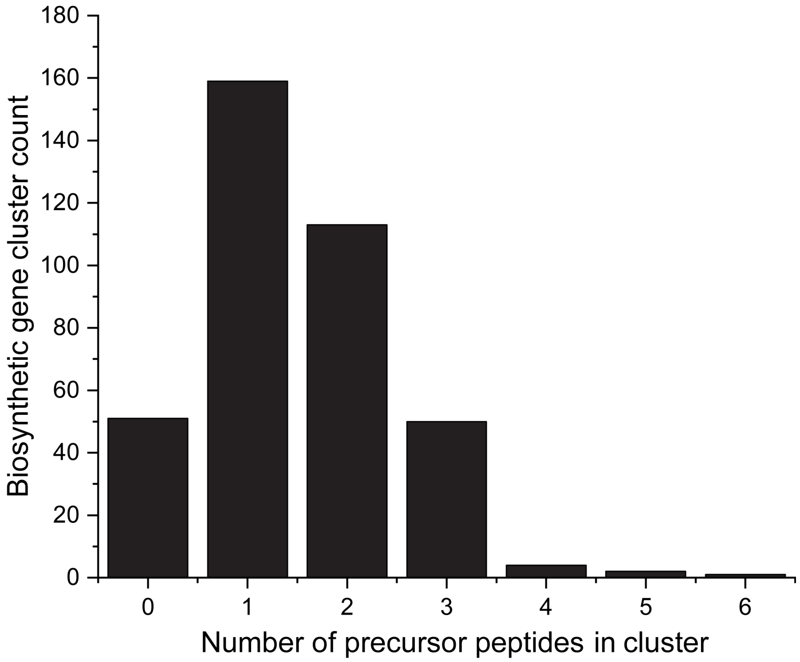


**Figure S6: Number of precursors per BGC.** Histogram of the number of linaridin precursor peptides detected within each non-redundant BGC (n = 382). Precursor peptides were identified by RODEO and deemed valid if receiving a score ≥12.

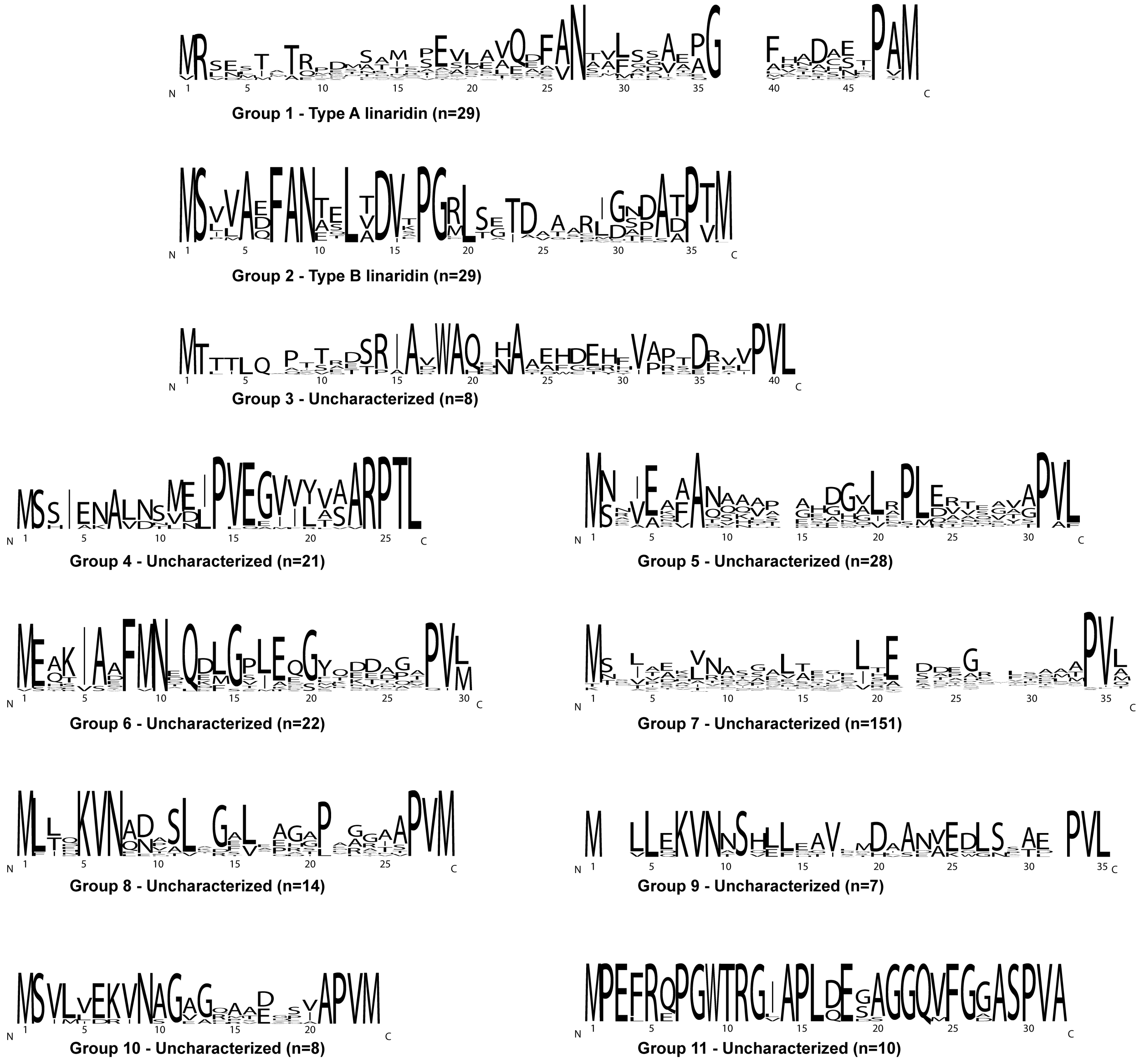


**Figure S7: Sequence analysis of RODEO-identified linaridin leader region.** All groups within the SSN containing greater than four members (Figure 2) were analyzed for leader sequence content using WebLogo.[23] Groups that include a known members are denoted with their type, with all other clusters classified as "Uncharacterized".

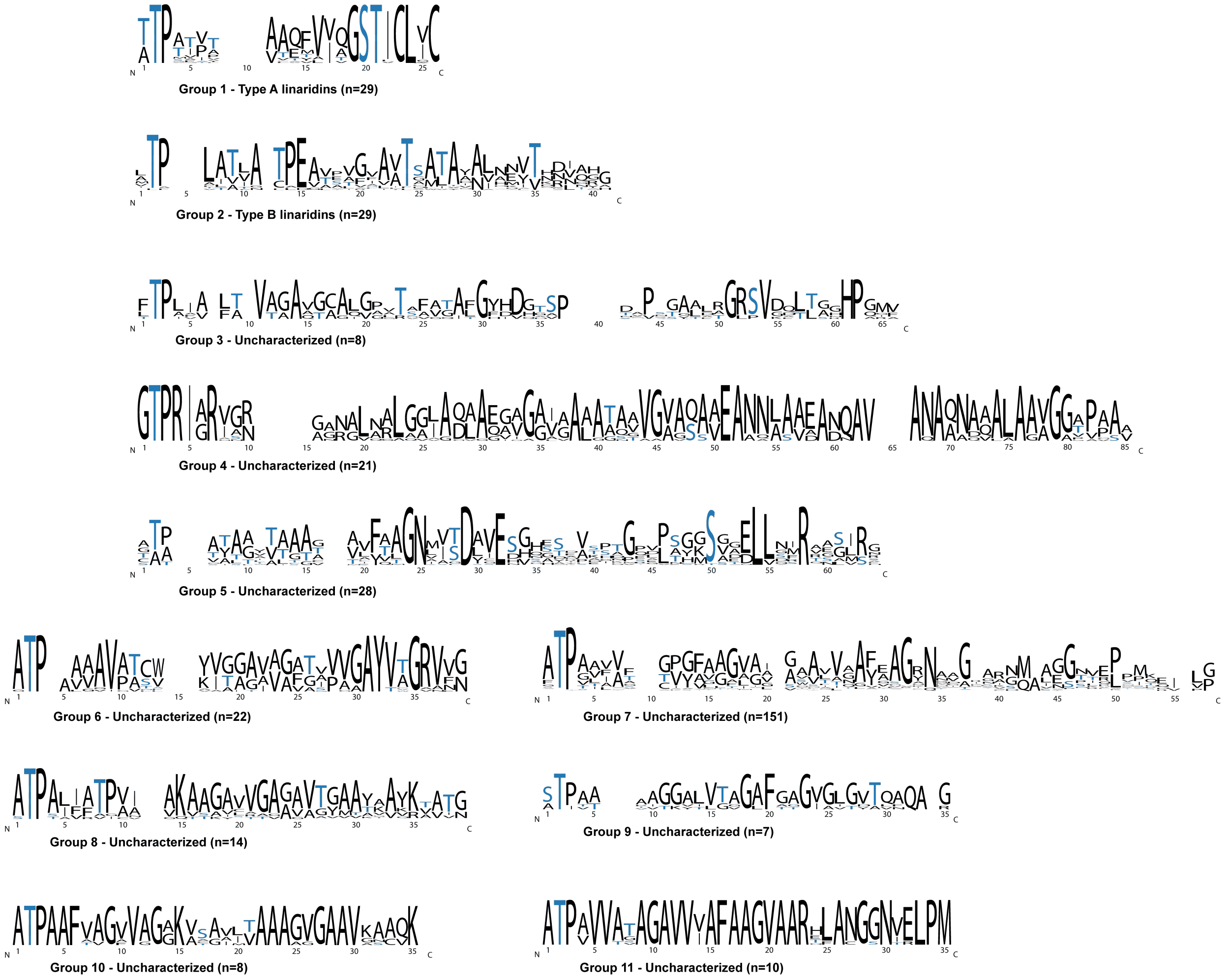


**Figure S8: Sequence analysis of RODEO identified linaridin core region.** All groups containing greater than four members on the SSN (Figure 2) were analyzed for core sequence content using WebLogo.[23] Groups that include a known members are denoted with their type, with all other clusters classified as "Uncharacterized". Blue, Ser and Thr (potential dehydration sites).

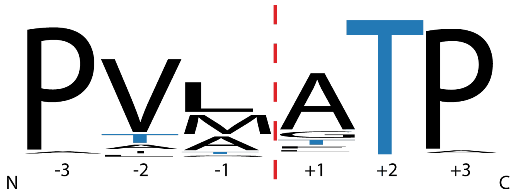


**Figure S9: Sequence analysis of linaridin leader/core proteolytic site.** A Logo of all predicted precursor proteolytic cleavage sites, manually curated to remove any potential misidentifications. A total of 549 predicted proteolytic sites were aligned using MAFFT[4] with visualization using WebLogo.[23] MAFFT website: https://mafft.cbrc.jp/alignment/server/

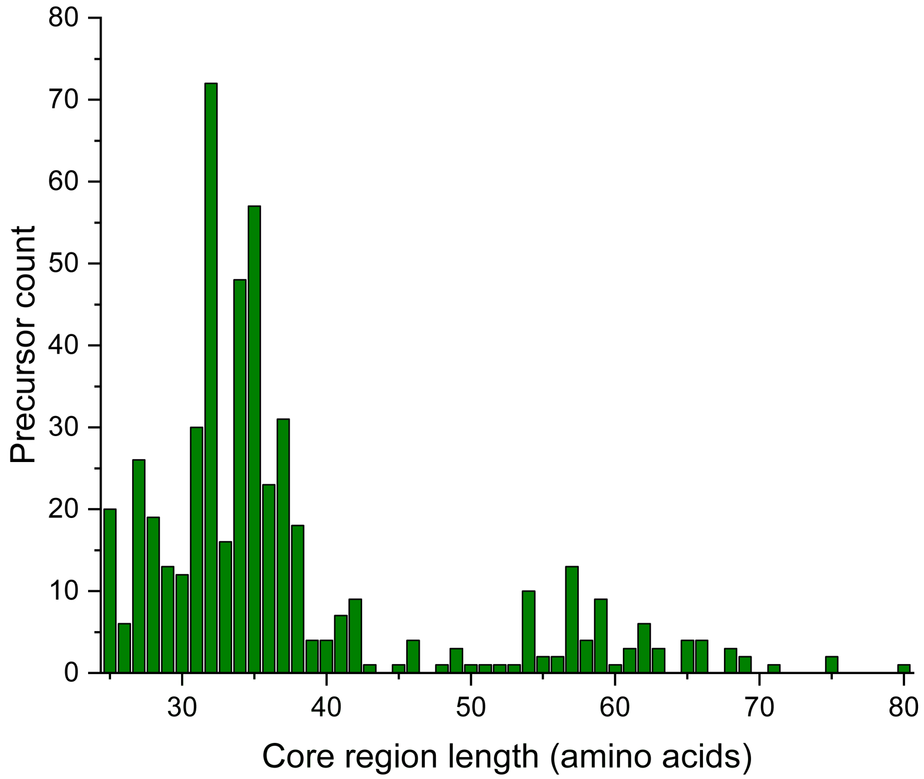


**Figure S10: Linaridin core region length variability.** Histogram of linaridin core region length identified by RODEO (n = 568). Sequences were identified/scored by RODEO. The populous group from 25-40 residues comprises all isolated linaridins, while a significant number of linaridins with longer core sequences remain uncharacterized.

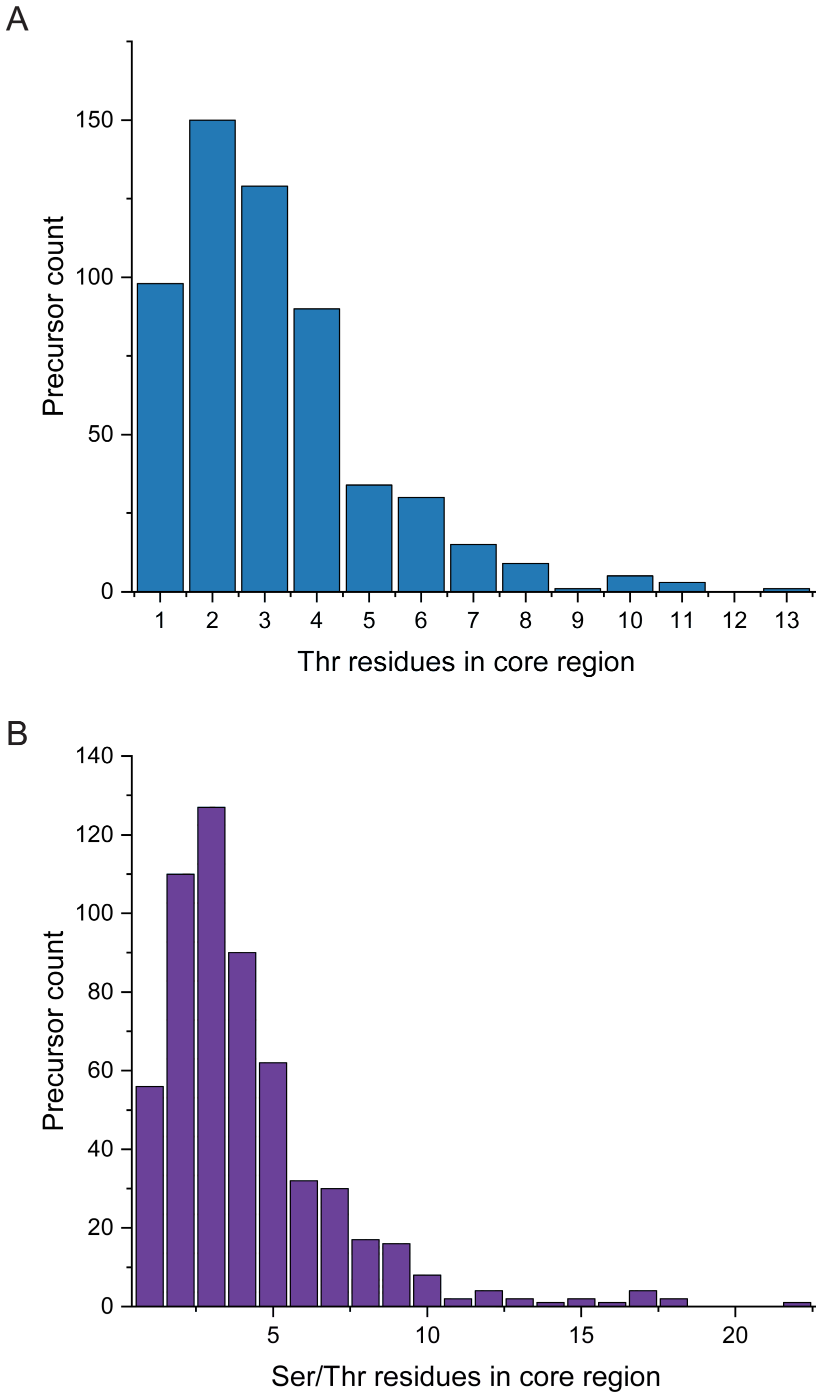


**Figure S11: Thr and Ser content of predicted core region.** Histogram showing the Thr content (A) and combined Ser/Thr content (B) of the linaridin core regions identified by RODEO (n = 568). The predicted leader/core cleavage site was predicted by motif analysis (Figure S9). Characterized linaridins contain 2-7 Thr in the core regions with all being converted to Dhb. Dehydroalanine formation (from Ser) has not yet been observed in wild-type linaridins but remains theoretically possible.

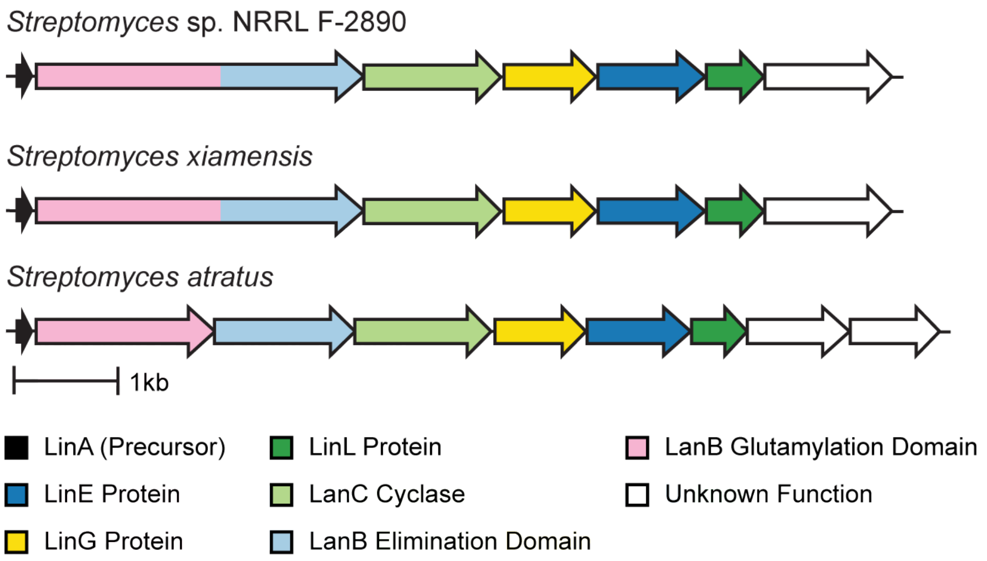


| | leader peptide | core peptide |
|---|---|---|
| NRRL F-2890: | MGADLITAAANEELSDLDLDLTFAPNDE | **NTHVMLACAVDSMGNNVNTSRGIIPQAPVCCA** |
| *S. xiamensis*: | MGADLITAAANEELSDLDLDLTFAPNDE | **NTHVMLACAVDSMGNNVNTSRGIIPQAPVCCA** |
| *S. atratus*: | MSTDLLIAAADEELSDLDLDVHFMSSGD | **VADQMYSCAVDSMGNGSSTSNGWRTCCA** |

**Figure S12: Hybrid Lanthipeptide and Linaridin BGCs.** Linaridin-lanthipeptide hybrids from various *Streptomyces* sp. The first two examples contain a fused LanB, whereas *Streptomyces atratus* contains a split LanB, like those found in thiopeptide biosynthetic gene clusters.[24] Type I Lanthipeptide RODEO module was used to identify precursors and their respective proteolytic cleavage sites in combination with manual curation.

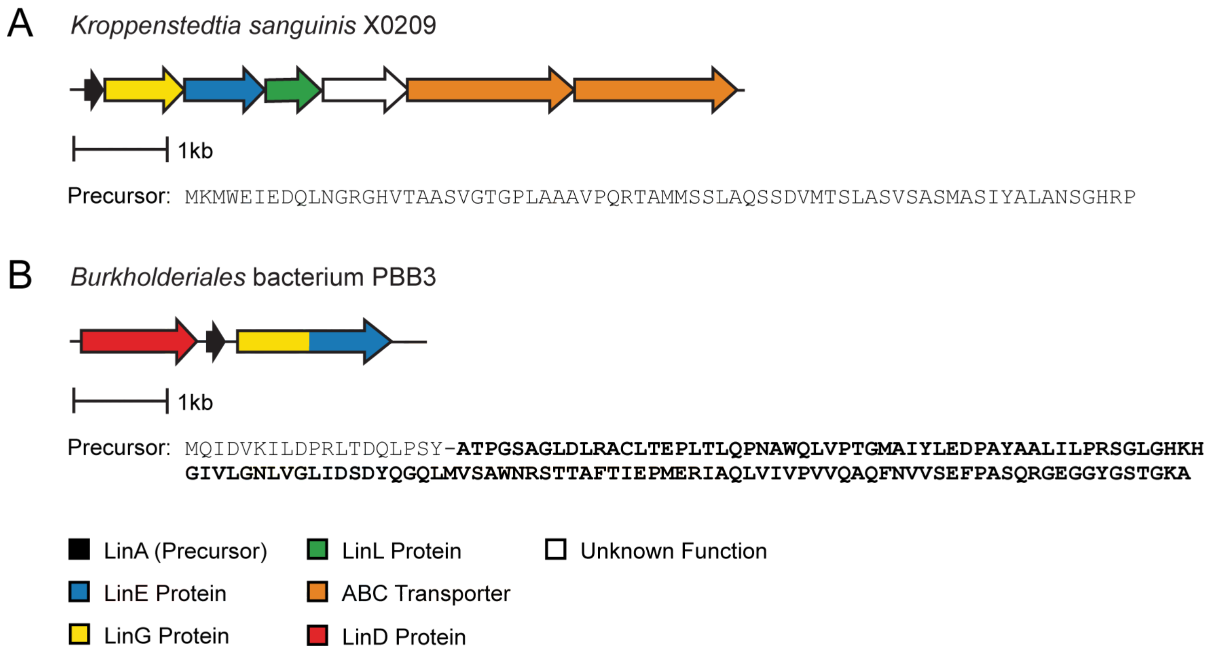


**Figure S13: Non-Actinomycete BGCs.** (A) BGC found in *Kroppenstadia sanguinis* X0209 (Firmicutes). The associated precursor peptide was not detected by the linaridin scoring module. (B) BGC found in *Burkholderiales* bacterium PBB3 (Proteobacteria). This cluster lacks a LinL homolog and contains an unusually long precursor peptide, scoring 12 on the linaridin module of RODEO.

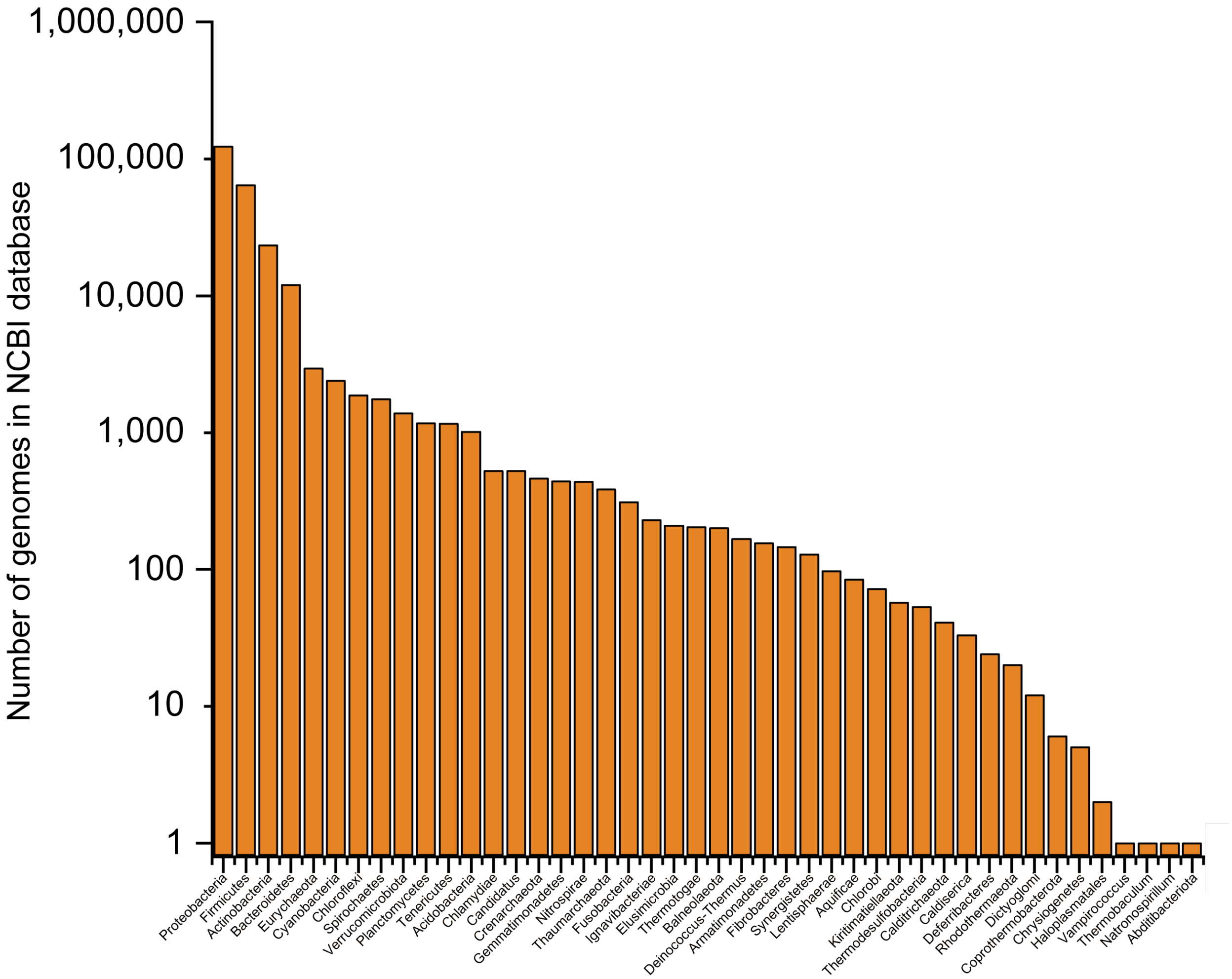


**Figure S14: Phylogenetic distribution of NCBI sequences.** Logarithmic histogram showing the number of sequenced prokaryotic genomes in NCBI database (Release 237) based on phylum.

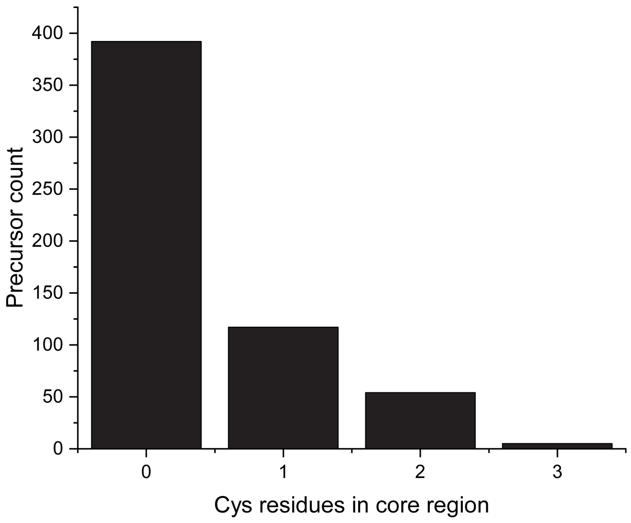


**Figure S15: Cys content of core peptides predicted by RODEO.** Histogram of linaridin core peptide Cys content (n = 568). Sequences were identified and scored by RODEO with the proteolytic sites predicted by motif analysis (**Figure S9**). Hybrid BGCs do not contain precursors validated by RODEO and are therefore not included in this analysis. Characterized linaridins contain either 0 or 2 Cys correlating with the presence of a gene encoding a decarboxylase (AviCys formation).

**Table S6: Linaridin core regions containing Cys.** Below is a subset of linaridin core sequences containing Cys (red) but lacking a putative AviCys-forming decarboxylase. *Saccharomonospora* sp. contain Cys at the first position of the core region (+1) followed by a Thr/Ser-rich region (blue).

| Genus / species | Strain | NCBI Accession | Truncated core peptide (+1 to +26) |
|---|---|---|---|
| *Saccharomonospora* sp. | LRS4.154 | OQO91990.1 | CTTSTTTATLAGTVGMVFTAGIISDA |
| *Saccharomonospora* sp. | CNQ490 | WP_024877745.1 | CTTTTTLTTTTATCTVFATGISSDAS |
| *Saccharomonospora* sp. | CNQ490 | WP_024877746.1 | CTTTTTTSTITGCVGTVFTMGVISDA |
| *Saccharomonospora* sp. | LRS4.154 | OQO91991.1 | CTATTLTTTTATCTVFATGIIGDAGE |
| *Saccharomonospora piscinae* | 06168H-1 | WP_138383134.1 | CTATTLTTTTATCTVFATGSIGDAGE |
| *Saccharomonospora piscinae* | 06168H-1 | WP_081191125.1 | CTTSTTTATLAGTVGMVFTAGIISDA |

**Table S7: List of bacterial strains evaluated for linaridin production**. The 34 strains listed below originated from the Agricultural Research Service Culture Collection (NRRL) in Peoria, IL (https://nrrl.ncaur.usda.gov/).

| Genus / species | Strain designation (NRRL) |
|---|---|
| *Streptomyces* sp. | B-1347 |
| *Salinispora pacifica* | B-24797 |
| *Streptomyces antibioticus* | B-1701, B-2032, B-2770, B-8002 |
| *Streptomyces auratus* | B-8097 |
| *Streptomyces bicolor* | B-5348 |
| *Streptomyces bottropensis* | ISP-5262 |
| *Streptomyces iakyrus* | ISP-5482 |
| *Streptomyces monomycini* | B-24309 |
| *Streptomyces neyagawaensis* | B-3092, B-16496, ISP-5588 |
| *Streptomyces noursei* | B-1714 |
| *Streptomyces scabrisporus* | B-24202 |
| *Streptomyces scopuliridis* | B-24574 |
| *Streptomyces sioyaensis* | B-5408 |
| *Streptomyces* sp. | F-5630 |
| *Streptomyces* sp. | S-337 |
| *Streptomyces* sp. | S-1448 |
| *Streptomyces stelliscabiei* | B-24447 |
| *Streptomyces sulphureus* | B-1331, B-2195 |
| *Streptomyces torulosus* | S-189, F-3039, B-3889 |
| *Streptomyces viridochromogenes* | B-3607, B-12033, S-256, S-452, S-453, S-635, S-636 |

**Table S8: Pegvadin A MS/MS ion assignments**. Masses assigned with calculated (calc.) and observed (obsv.) values given along with the error (ppm values given) in the assignment. Fragments were observed in the 1+ charge state unless otherwise noted.

| Ion | Calc. Mass | Obsv. Mass | Error |
|---|---|---|---|
| y3 | 416.2616 | 416.2611 | 1.1 |
| y4 | 531.2885 | 531.2879 | 1.1 |
| y5 | 659.3835 | 659.3827 | 1.3 |
| y6 | 742.4206 | 742.4196 | 1.4 |
| y7 | 841.4890 | 841.4875 | 1.8 |
| y8 | 970.5316 | 970.5301 | 1.6 |
| y9 | 1107.5905 | 1107.5890 | 1.3 |
| y10 | 1220.6746 | 1220.6726 | 1.7 |
| y11 | 1291.7117 | 1291.7096 | 1.6 |
| y12 | 1454.7750 | 1454.7728 | 1.5 |
| y13 | 1525.8122 | 1525.8099 | 1.5 |
| y14 | 1608.8493 | 1608.8463 | 1.9 |
| y15 | 1679.8864 | 1679.8828 | 2.1 |
| y16 | 1766.9184 | 1766.9136 | 2.7 |
| y17 | 1849.9555 | 1849.9517 | 2.1 |
| $y19^{2+}$ | 1010.5344 | 1010.5325 | 1.9 |
| $y23^{2+}$ | 1201.6271 | 1201.6255 | 1.3 |
| $y24^{2+}$ | 1243.1456 | 1243.1439 | 1.4 |
| $y25^{2+}$ | 1278.6642 | 1278.6621 | 1.6 |
| $y26^{2+}$ | 1335.2062 | 1335.2041 | 1.6 |
| $y27^{2+}$ | 1376.7248 | 1376.7227 | 1.5 |
| $y30^{2+}$ | 1517.3117 | 1517.3098 | 1.3 |
| $y31^{2+}$ | 1558.8303 | 1558.8278 | 1.6 |
| b4 | 435.2966 | 435.2961 | 1.1 |
| b5 | 506.3337 | 506.3331 | 1.3 |
| b6 | 589.3708 | 589.3700 | 1.3 |
| b7 | 702.4549 | 702.4539 | 1.4 |
| b8 | 773.4920 | 773.4909 | 1.5 |
| b9 | 856.5291 | 856.5277 | 1.6 |

**Table S9: Pegvadin B MS/MS ion assignments**. Masses assigned with calculated (calc.) and observed (obsv.) values given along with the error (ppm values given) in the assignment. Fragments were observed in the 1+ charge state unless otherwise noted.

| Ion | Calc. Mass | Obsv. Mass | Error |
|---|---|---|---|
| y3 | 416.2616 | 416.2611 | 1.2 |
| y4 | 531.2885 | 531.2879 | 1.1 |
| y5 | 659.3835 | 659.3825 | 1.5 |
| y6 | 742.4206 | 742.4195 | 1.5 |
| y8 | 970.5316 | 970.5299 | 1.8 |
| y9 | 1107.5905 | 1107.5884 | 2.0 |
| $y12^{2+}$ | 727.8914 | 727.8902 | 1.6 |
| $y13^{2+}$ | 763.4100 | 763.4087 | 1.6 |
| $y14^{2+}$ | 804.9285 | 804.9272 | 1.7 |
| $y15^{2+}$ | 840.4471 | 840.4459 | 1.5 |
| $y16^{2+}$ | 883.9631 | 883.9617 | 1.6 |
| $y17^{2+}$ | 925.4817 | 925.4804 | 1.4 |
| $y18^{2+}$ | 975.0159 | 975.0141 | 1.8 |
| $y19^{2+}$ | 1010.5344 | 1010.5326 | 1.8 |
| $y20^{2+}$ | 1060.0686 | 1060.0670 | 1.6 |
| $y21^{2+}$ | 1088.5794 | 1088.5774 | 1.8 |
| $y22^{2+}$ | 1153.1007 | 1153.0988 | 1.6 |
| $y23^{2+}$ | 1201.6271 | 1201.6248 | 1.9 |
| $y24^{2+}$ | 1243.1456 | 1243.1437 | 1.5 |
| b4 | 435.2966 | 435.2960 | 1.2 |
| b5 | 506.3337 | 506.3330 | 1.3 |
| b6 | 577.3708 | 577.3700 | 1.5 |
| b7 | 690.4549 | 690.4537 | 1.6 |
| b8 | 761.4920 | 761.4911 | 1.2 |
| b9 | 844.5291 | 844.5277 | 1.7 |
| $b12^{2+}$ | 564.3269 | 564.3261 | 1.5 |
| $b13^{2+}$ | 613.8611 | 613.8601 | 1.6 |
| $b14^{2+}$ | 649.3796 | 649.3785 | 1.7 |

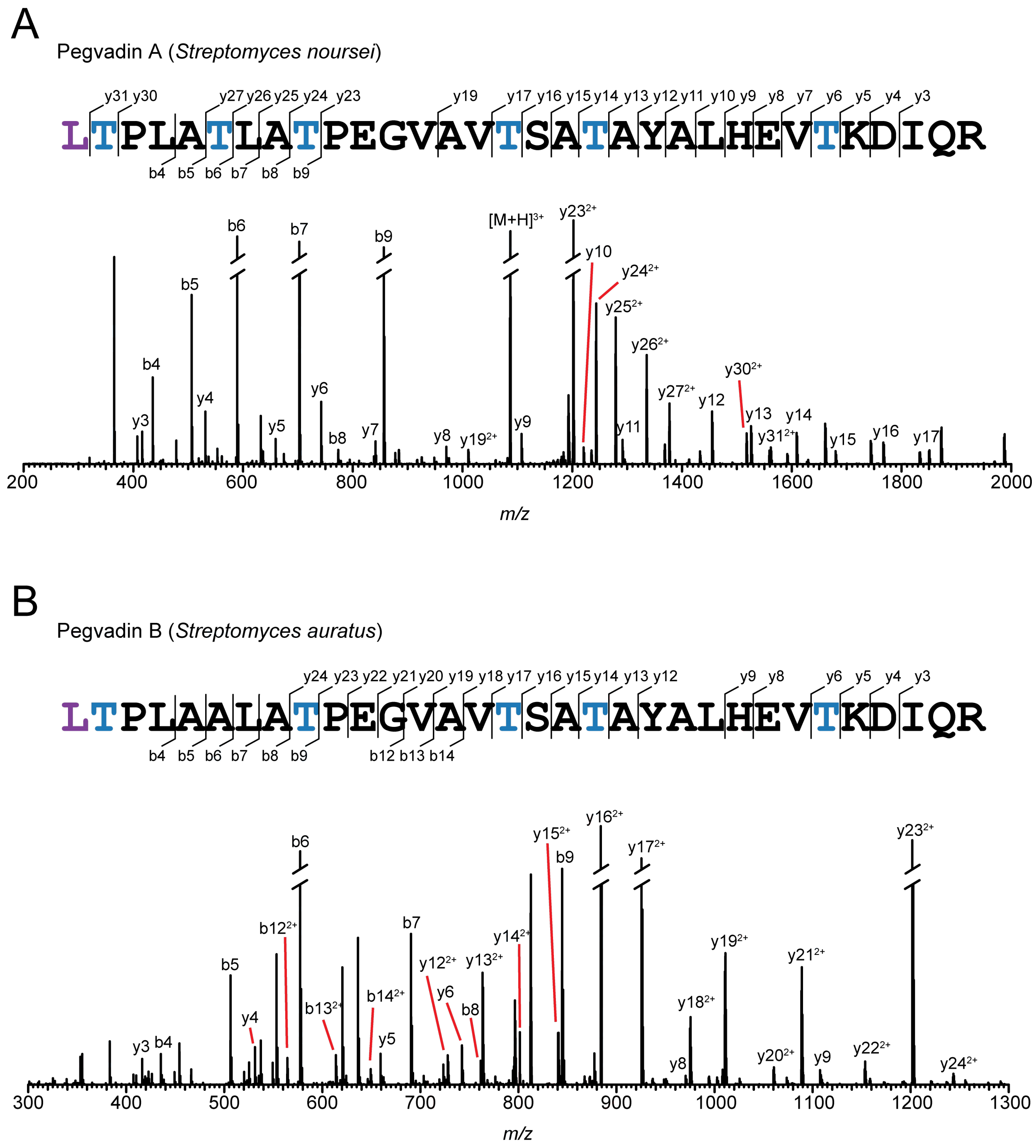


**Figure S16: High-resolution and tandem mass spectrometry of pegvadins A and B.** HRMS/MS data for pegvadin A (A) and pegvadin B (B). Fragmentation yielded b and y ions consistent with the sequence of the precursor peptides. Blue, Thr position was converted to Dhb; purple, dimethylated N-terminus.

**Table S10: Sequence identity/similarity comparison of enzymes in characterized linaridin BGCs.** Sequences were aligned with MAFFT and identity/similarity was assessed using the Sequence Identity And Similarity tool (http://imed.med.ucm.es/Tools/sias.html). Yellow, % identity; Green, % similarity.

| **LinE Protein** | **1** | **2** | **3** | **4** | **5** | **6** |
|---|---|---|---|---|---|---|
| 1. *Streptomyces noursei* (WP_073444435.1) | 100 | 79 | 69 | 31 | 31 | 28 |
| 2. *Streptomyces auratus* (WP_040898716.1) | 83 | 100 | 66 | 31 | 29 | 28 |
| 3. *Streptomyces* sp. CT34 (WP_078894000.1) | 76 | 74 | 100 | 31 | 30 | 29 |
| 4. *Streptomyces* sp. OH-4156 (ADR72963.1) | 44 | 43 | 43 | 100 | 82 | 68 |
| 5. *Streptomyces barkulensis* (WP_101256404.1) | 43 | 42 | 42 | 87 | 100 | 71 |
| 6. *Streptomyces griseus* (WP_003970681.1 ) | 40 | 39 | 39 | 77 | 77 | 100 |

| **LinG Protein** | **1** | **2** | **3** | **4** | **5** | **6** |
|---|---|---|---|---|---|---|
| 1. *Streptomyces noursei* (WP_067348450.1) | 100 | 92 | 84 | 33 | 31 | 31 |
| 2. *Streptomyces auratus* (WP_040898719.1) | 95 | 100 | 86 | 33 | 31 | 30 |
| 3. *Streptomyces* sp. CT34 (WP_043265482.1) | 88 | 88 | 100 | 31 | 30 | 30 |
| 4. *Streptomyces* sp. OH-4156 (ADR72963.1) | 45 | 44 | 42 | 100 | 84 | 84 |
| 5. *Streptomyces barkulensis* (WP_101256404.1) | 43 | 43 | 41 | 88 | 100 | 78 |
| 6. *Streptomyces griseus* (WP_003970681.1 ) | 45 | 44 | 44 | 87 | 83 | 100 |

| **LinL Protein** | **1** | **2** | **3** | **4** | **5** | **6** |
|---|---|---|---|---|---|---|
| 1. *Streptomyces noursei* (WP_099055705.1) | 100 | 29 | 28 | 26 | 27 | 26 |
| 2. *Streptomyces auratus* (WP_106430490.1) | 38 | 100 | 70 | 37 | 33 | 31 |
| 3. *Streptomyces* sp. CT34 (WP_078894692.1) | 38 | 76 | 100 | 36 | 33 | 35 |
| 4. *Streptomyces* sp. OH-4156 (ADR72964.1) | 37 | 46 | 46 | 100 | 87 | 74 |
| 5. *Streptomyces barkulensis* (WP_101256405.1) | 37 | 44 | 44 | 89 | 100 | 76 |
| 6. *Streptomyces griseus* (WP_003970680.1) | 37 | 40 | 44 | 81 | 81 | 100 |

| **LinM (Methyltransferase)** | **1** | **2** | **3** | **4** | **5** | **6** |
|---|---|---|---|---|---|---|
| 1. *Streptomyces noursei* (WP_067348459.1) | 100 | 85 | 87 | 25 | 22 | 21 |
| 2. *Streptomyces auratus* (WP_040898722.1) | 88 | 100 | 83 | 26 | 24 | 20 |
| 3. *Streptomyces* sp. CT34 (WP_107068144.1) | 89 | 87 | 100 | 28 | 25 | 23 |
| 4. *Streptomyces* sp. OH-4156 (ADR72966.1) | 32 | 34 | 33 | 100 | 85 | 76 |
| 5. *Streptomyces barkulensis* (WP_101256407.1) | 30 | 31 | 32 | 89 | 100 | 76 |
| 6. *Streptomyces griseus* (WP_012381886.1) | 30 | 30 | 32 | 83 | 84 | 100 |

| **LinT (ABC Transporter)** | **1** | **2** | **3** | **4** | **5** | **6** |
|---|---|---|---|---|---|---|
| 1. *Streptomyces noursei* (WP_102926632.1) | 100 | 74 | 77 | 21 | 22 | 23 |
| 2. *Streptomyces auratus* (WP_040900283.1) | 79 | 100 | 80 | 17 | 19 | 19 |
| 3. *Streptomyces* sp. CT34 (WP_052230046.1) | 83 | 85 | 100 | 18 | 20 | 20 |
| 4. *Streptomyces* sp. OH-4156 (ADR72968.1) | 30 | 26 | 28 | 100 | 78 | 61 |
| 5. *Streptomyces barkulensis* (WP_101256409.1) | 31 | 28 | 29 | 82 | 100 | 62 |
| 6. *Streptomyces griseus* (WP_012381885.1) | 30 | 25 | 27 | 70 | 70 | 100 |

**Table S11: Media recipes.** A list of components used for bacterial cultivation media to evaluate linaridin production.

| Media | Ingredients | Amount per L (g) |
|---|---|---|
| **ATCC 172** | Soluble Starch | 20 |
| | Glucose | 10 |
| | Yeast Extract | 5 |
| | N-Z Amine | 5 |
| | $CaCO_3$ | 1 |
| | pH | 7.2 |
| **ISP4** | Soluble Starch | 10 |
| | $K_2HPO_4$ | 1 |
| | $Mg_2SO_4.7H_2O$ | 1 |
| | NaCl | 1 |
| | Ammonium Sulfate | 2 |
| | $CaCO_3$ | 2 |
| | $FeSO_4.7H_2O$ | 0.001 |
| | $ZnSO_4.7H_2O$ | 0.001 |
| | $MnCl_2.2H_2O$ | 0.001 |
| | pH | 7.2 |
| **Alt-MS** | Mannitol | 10 |
| | Malt Extract | 10 |
| | Soy Flour | 10 |
| **GUBC** | Glycerol | 5 mL |
| | Sucrose | 10 |
| | Beef Extract | 5 |
| | Casamino Acids | 5 |
| | Solution A | 5 mL |
| | Solution B | 2 mL |
| | **After Autoclave** | |
| | Balch's Vitamins | 10 mL |
| **ISP2** | Yeast Extract | 4 |
| | Malt Extract | 10 |
| | Dextrose | 4 |
| **AGS** | Arginine HCl | 1 |
| | Glycerol | 12.5 |
| | K2HPO4 | 1 |
| | NaCl | 1 |
| | $Mg_2SO_4.7H_2O$ | 0.5 |
| | $Fe_2(SO_4)_3.6H_2O$ | 0.01 |
| | $CuSO_4$.5H2O | 0.001 |
| | MnSO4.H2O | 0.001 |
| | $ZnSO_4.7H_2O$ | 0.001 |
| | pH | 7 |
| **Adapted** | Glucose | 25 |
| **TVA1** | Fish Meal | 15 |
| **Medium** | Yeast Extract | 2 |
| | $CaCO_3$ | 4 |
| | pH | 7.2 |
| **V8** | V8 Juice | 200 mL |
| | $CaCO_3$ | 3 |
