## Supplementary Dataset 3_RODEO-graphical for "Bioinformatic and Reactivity-Based Discovery of Linaridins"

### RODEO2

#### Parameters

| | |
|---|---|
| **Run Time** | 07:03PM on May 26, 2020 |
| **Version** | 2.1.4 |
| **Gene Window** | +/-8 CDS |
| **Peptide Range** | 45-120 aa |
| **Fetch Distance** | 25bp |
| **Peptide Type** | linar |

##### Annotation Legend

| **Appearance** | **Accession/Name** |
|---|---|
| | PF08242 |
| | PF08241 |
| | PF13489 |
| | PF13847 |
| | PF13649 |
| | PF00664 |
| | PF00005 |
| | PF02441 |
| | TIGR02113 |
| | TIGR00521 |
| | LinH |
| | LinG |
| | LinL |
| | LinE |
| | LinA |
| | predicted linar peptide |

#### Input Queries (click to navigate)

- WP_018133634.1
- WP_018133654.1
- KPH97805.1
- RJF42448.1
- WP_119836030.1
- WP_120204651.1
- WP_127841862.1
- WP_127841865.1
- WP_139738603.1
- WP_139738607.1
- SFE63453.1
- SDK49455.1
- RZU71018.1
- CRK56201.1
- OZM71980.1
- OZM72833.1
- OZM72835.1
- WP_086852701.1
- SFB47920.1
- SFB47982.1
- WP_116048497.1
- WP_116048516.1
- SEC50923.1
- OXM68433.1
- WP_110101283.1
- WP_110098361.1
- PXA76677.1
- WP_110122900.1
- WP_168186076.1
- WP_106848707.1
- SMX73793.1
- RCS84136.1

WP_127362990.1
WP_165883987.1
SDS21300.1
WP_101574944.1
OYT91441.1
WP_135975511.1
OJV60998.1
REE03689.1
OQJ64340.1
CAB1023416.1
WP_082313297.1
WP_026168100.1
OFO12264.1
WP_049151517.1
SDS67811.1
WP_070759247.1
WP_163342818.1
VXB69807.1
WP_022905513.1
WP_070417581.1
WP_138802251.1
WP_071302418.1
WP_071295649.1
WP_111010392.1
WP_111119637.1
PZE55389.1
WP_111057571.1
WP_110863536.1
PZF23552.1
WP_083393079.1
WP_123339259.1
ROQ16621.1
WP_123671264.1
WP_141693334.1
WP_051597146.1
SFF54177.1
GCE02330.1
WP_078981861.1
GAB19727.1
SDZ55823.1
WP_051062157.1
ARF74358.1
WP_129316254.1
WP_095798642.1
RYJ54945.1
REH27010.1
ERK70867.1
WP_155829033.1
AGW40770.1
ODA91227.1
AAT89666.1
WP_166328536.1
WP_045251507.1
WP_150450697.1
WP_091734673.1
SIR96088.1
WP_055482107.1
QCP08987.1
RKN45324.1
TDB95803.1
TDC00110.1
RQW94077.1
TWH68587.1
KXK61181.1
KXK61316.1
WP_148426628.1
QDY07820.1
WP_132400124.1
OWV06806.1
SCL65344.1
TCB95381.1
WP_066941623.1
WP_157570523.1
WP_152730243.1
WP_121689029.1
RLP71106.1
SJN40172.1
WP_121649883.1
WP_111766294.1

KZM74592.1
RNL83154.1
SHK25020.1
WP_150255942.1
WP_064723074.1
BCB83522.1
WP_084705756.1
WP_084705758.1
KID32044.1
KID32052.1
PXY36318.1
PXY36951.1
TWD68636.1
WP_104249327.1
SOE04649.1
WP_104258703.1
TFI41935.1
WP_013398290.1
WP_083093117.1
WP_005463309.1
WP_005463322.1
WP_157601541.1
WP_138383130.1
WP_138383132.1
WP_037336134.1
WP_155984838.1
OQO91992.1
WP_081191130.1
WP_139794736.1
WP_050590090.1
WP_050590677.1
WP_155244151.1
OEU87428.1
OEU88179.1
WP_139140755.1
WP_070011838.1
WP_070012848.1
WP_151770911.1
AOE46838.1
AXK34881.1
SFX78248.1
WP_040898719.1
WP_101256404.1
WP_101258824.1
WP_051827992.1
WP_051828002.1
WP_078489996.1
WP_030214179.1
PJE97628.1
WP_128380536.1
GCD35929.1
WP_150482280.1
KND25157.1
WP_046706697.1
WP_060892339.1
WP_119581921.1
WP_024756459.1
WP_108956003.1
WP_032775535.1
WP_033526318.1
EMF28262.1
WP_105970077.1
WP_030581598.1
AXE28036.1
SQA25558.1
WP_148235529.1
SEE39163.1
BAG18344.1
BAG23194.1
SFS58200.1
RKN35811.1
WP_157852675.1
TQE28105.1
WP_100243889.1
EKX67255.1
WP_153524745.1
WP_063778956.1
WP_153483925.1
WP_069884696.1

WP_043371861.1
WP_167503753.1
WP_149547799.1
WP_078624126.1
OEV05017.1
OEV08011.1
SPE49561.1
WP_055540568.1
WP_150491658.1
AQU70555.1
WP_078073573.1
PNE36149.1
PNE39469.1
ANZ17123.1
WP_130798439.1
WP_100108206.1
SFJ78423.1
WP_069992530.1
SEQ27572.1
GCD43950.1
WP_078630323.1
KMO97782.1
GAQ68222.1
KFG02471.1
WP_086752480.1
WP_086756723.1
WP_086800803.1
WP_086801155.1
CBG75367.1
PVE04739.1
WP_055697709.1
RXS62043.1
WP_161989239.1
QBA31012.1
PIG13858.1
SOD81768.1
WP_165274063.1
ROQ95786.1
SED34674.1
SOE12061.1
PJJ03557.1
OXY94523.1
RIA63746.1
SHI10722.1
PIG43643.1
AJC56865.1
WP_132931361.1
WP_159062470.1
WP_106438501.1
WP_111339449.1
PZT77967.1
RPK73392.1
WP_107479021.1
RAJ66737.1
SCK62389.1
RKN03588.1
RKN04869.1
RKN10930.1
TCR48817.1
WP_073790447.1
WP_148648000.1
OKJ59748.1
OKI92952.1
OKJ54325.1
WP_073756736.1
WP_084898409.1
SFM59896.1
AUH40579.1
WP_018835201.1
WP_027766445.1
WP_078528867.1
WP_078528871.1
WP_027759150.1
WP_018853313.1
RZU87368.1
RZU91185.1
RZU91753.1
PJJ50016.1
WP_078894000.1

WP_157878906.1
GFH39492.1
WP_045555727.1
WP_023423357.1
WP_159508791.1
WP_152166802.1
WP_165254437.1
WP_165262038.1
RNL73410.1
WP_147874565.1
WP_171052848.1
WP_147247722.1
RYJ23219.1
WP_129773542.1
SPF07382.1
PPS88678.1
WP_063346941.1
SBU99212.1
REK89866.1
WP_158858022.1
WP_158716051.1
WP_043177785.1
WP_158708080.1
WP_037817104.1
WP_030999015.1
WP_030414971.1
WP_158744997.1
WP_030801616.1
WP_111003021.1
ADR72963.1
WP_097967158.1
WP_057667039.1
SCE23008.1
WP_032773956.1
WP_093563556.1
EXU61863.1
RBM10751.1
RBM24328.1
WP_113686667.1
PPQ56339.1
WP_129844105.1
WP_097937323.1
THA27824.1
KQX31486.1
WP_057604735.1
WP_056796007.1
KRA40986.1
QHY99994.1
ARI51159.1
WP_161245905.1
WP_161264539.1
WP_003965563.1
WP_063743169.1
WP_161184048.1
WP_161184067.1
WP_042495855.1
WP_099217667.1
WP_161229806.1
WP_018487272.1
WP_032756679.1
WP_093683568.1
WP_093684647.1
WP_161223844.1
TXS69815.1
WP_148838111.1
WP_103529472.1
WP_103502974.1
WP_051266157.1
WP_037978036.1
WP_078511723.1
SNX62950.1
RXS68531.1
WP_079127392.1
WP_155070885.1
WP_149180181.1
WP_149823165.1
EHM28125.1
EHM28131.1
WP_050987754.1

WP_158000644.1
WP_164721115.1
AZM48683.1
RSS64107.1
KOU14214.1
SCK38574.1
SCK46711.1
WP_079100284.1
SCK25365.1
SCK35447.1
SCK51308.1
WP_045865779.1
WP_165339856.1
RII13759.1
PWS41421.1
KND26783.1
WP_019544895.1
AWI33039.1
WP_055713858.1
WP_159744032.1
WP_132756788.1
WP_106434293.1
AKG45062.1
AKG45482.1
WP_016907298.1
SEG54329.1
SHL65197.1
SHM24347.1
WP_148281456.1
SED38833.1
WP_147276785.1
AUN42108.1
RDB46095.1
SFQ42880.1
WP_114385254.1

### Results for WP_018133634.1 [Acaricomes phytoseiuli DSM 14247] back to top

previous - next

#### Architecture

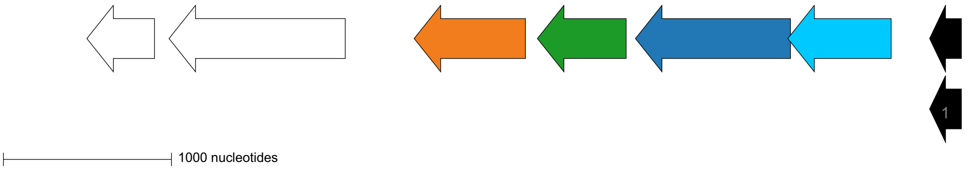

Link to nucleotide sequence

| Accession | start | end | direction | length (aa) | Pfam/HMM | name | description | E-value |
|---|---|---|---|---|---|---|---|---|
| WP_018133630.1 | 429 | 30 | - | 132 | NO PFAM MATCH | - | - | - |
| WP_018133631.1 | 1571 | 518 | - | 350 | LinG | LinG | LinG Homolog | 1.20E-49 |
| WP_083906470.1 | 2638 | 1981 | - | 218 | TIGR01189<br>TIGR02857<br>TIGR01188<br>TIGR02673<br>PF00005 | TIGR01189<br>TIGR02857<br>TIGR01188<br>TIGR02673<br>ABC_tran | ccmA: heme ABC exporter, ATP-binding protein CcmA<br>CydD: thiol reductant ABC exporter, CydD subunit<br>drrA: daunorubicin resistance ABC transporter, ATP-binding protein<br>FtsE: cell division ATP-binding protein FtsE<br>ABC transporter | 5.30E-26<br>3.60E-25<br>1.30E-24<br>3.40E-23<br>1.90E-22 |
| WP_156807538.1 | 3236 | 2708 | - | 175 | LinL | LinL | LinL | 4.40E-41 |
| WP_018133634.1 | 4216 | 3298 | - | 305 | LinE<br>LinH<br>PF12697<br>TIGR02427<br>TIGR03611 | LinE<br>LinH<br>Abhydrolase_6<br>TIGR02427<br>TIGR03611 | LinE Homolog<br>LinH (Fusion of LinE/LinG)<br>Alpha/beta hydrolase family<br>protocat_pcaD: 3-oxoadipate enol-lactonase<br>RutD: pyrimidine utilization protein D | 5.80E-76<br>2.50E-41<br>2.80E-10<br>2.10E-06<br>9.40E-05 |
| WP_083906472.1 | 4820 | 4205 | - | 204 | LinH<br>LinG<br>TIGR04033 | LinH<br>LinG<br>TIGR04033 | LinH (Fusion of LinE/LinG)<br>LinG Homolog<br>export_SdpB: antimicrobial peptide system protein, SdpB family | 5.00E-66<br>2.30E-64<br>5.80E-06 |
| WP_156807539.1 | 5234 | 5048 | - | 61 | LinA | LinA | LinA (linaridin precursor) | 5.10E-10 |

| index | leader | core | start | end | dir | score |
|---|---|---|---|---|---|---|
| 1 | MYAVGSSNFSSLQGAVNFAFSDVHPGYMGSNDSPVM | TTPTTALIAPAAAMAVTGSTLCLAC | 5234 | 5048 | - | 29 |

### Results for WP_018133654.1 [Acaricomes phytoseiuli DSM 14247] back to top

previous - next

#### Architecture

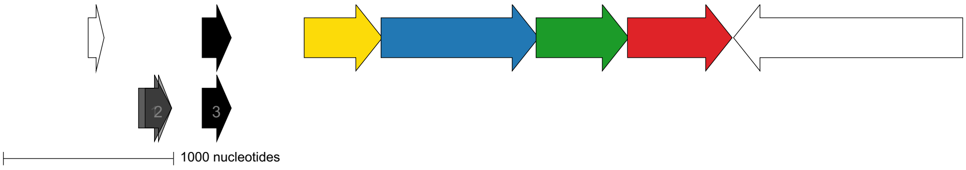

Link to nucleotide sequence

| Accession | start | end | direction | length (aa) | Pfam/HMM | name | description | E-value |
|---|---|---|---|---|---|---|---|---|
| WP_020500071.1 | 132 | 231 | + | 33 | INFERRED GENE | - | - | - |
| WP_156807541.1 | 803 | 974 | + | 56 | LinA | LinA | LinA (linaridin precursor) | 3.20E-09 |
| WP_156807543.1 | 1403 | 1859 | + | 151 | LinG<br>LinH<br>TIGR04033 | LinG<br>LinH<br>TIGR04033 | LinG Homolog<br>LinH (Fusion of LinE/LinG)<br>export_SdpB: antimicrobial peptide system protein, SdpB family | 5.50E-60<br>1.70E-56<br>3.30E-06 |
| WP_018133654.1 | 1855 | 2779 | + | 307 | LinE<br>LinH<br>PF12697<br>PF06821<br>TIGR02427 | LinE<br>LinH<br>Abhydrolase_6<br>Ser_hydrolase<br>TIGR02427 | LinE Homolog<br>LinH (Fusion of LinE/LinG)<br>Alpha/beta hydrolase family<br>Serine hydrolase<br>protocat_pcaD: 3-oxoadipate enol-lactonase | 2.70E-81<br>2.60E-46<br>1.50E-09<br>4.70E-06<br>9.00E-06 |
| WP_018133655.1 | 2765 | 3311 | + | 181 | LinL | LinL | LinL | 2.80E-48 |
| WP_018133656.1 | 3303 | 3918 | + | 204 | TIGR00521<br>PF02441<br>TIGR02113 | TIGR00521<br>Flavoprotein<br>TIGR02113 | coaBC_dfp: phosphopantothenoylcysteine decarboxylase / phosphopantothenate--cysteine ligase<br>Flavoprotein<br>coaC_strep: phosphopantothenoylcysteine decarboxylase | 2.90E-07<br>2.30E-06<br>3.00E-05 |
| WP_156807542.1 | 5271 | 3924 | - | 448 | NO PFAM MATCH | - | - | - |

| index | leader | core | start | end | dir | score |
|---|---|---|---|---|---|---|
| 1 | MRGDAQKLSLSPPLIGSGVTIIYAEPEEIAAASPKC | PTVRDMMHQVDPGGDRCVPSPASRVR | 432 | 621 | + | 8 |
| 2 | LIGSGVTIIYAEPEEIAAASPKC | PTVRDMMHQVDPGGDRCVPSPASRVR | 471 | 621 | + | 6 |
| 3 | MSSTLVQTPSVEDSVNAMFSHIDQGFVNAHNEPAM | TTPSAIVTQTLVVGSTICLVC | 803 | 974 | + | 31 |

### Results for KPH97805.1 [Actinobacteria bacterium OK074] back to top

previous - next

#### Architecture

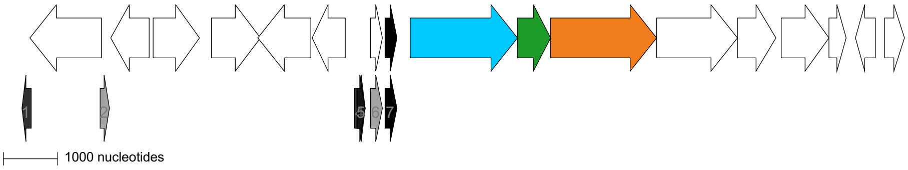

Link to nucleotide sequence

| Accession | start | end | direction | length (aa) | Pfam/HMM | name | description | E-value |
|---|---|---|---|---|---|---|---|---|
| KPH97797.1 | 8552 | 7235 | - | 438 | TIGR01356<br>PF00275<br>TIGR01072 | TIGR01356<br>EPSP_synthase<br>TIGR01072 | aroA: 3-phosphoshikimate 1-carboxyvinyltransferase<br>EPSP synthase (3-phosphoshikimate 1-carboxyvinyltransferase)<br>murA: UDP-N-acetylglucosamine 1-carboxyvinyltransferase | 1.20E-110<br>8.20E-105<br>1.80E-06 |
| KPH97798.1 | 9435 | 8712 | - | 240 | PF13398 | Peptidase_M50B | Peptidase M50B-like | 7.60E-62 |
| KPH97799.1 | 9497 | 10343 | + | 281 | PF02586 | SRAP | SOS response associated peptidase (SRAP) | 4.80E-66 |
| KPH97800.1 | 10575 | 11478 | + | 300 | PF00196<br>TIGR03541<br>TIGR03020<br>PF08281<br>TIGR02937 | GerE<br>TIGR03541<br>TIGR03020<br>Sigma70_r4_2<br>TIGR02937 | Bacterial regulatory proteins, luxR family<br>reg_near_HchA: LuxR family transcriptional regulatory, chaperone HchA-associated<br>EpsA: transcriptional regulator EpsA<br>Sigma-70, region 4<br>sigma70-ECF: RNA polymerase sigma factor, sigma-70 family | 2.40E-15<br>6.50E-10<br>5.10E-08<br>3.20E-04<br>3.40E-04 |
| KPH97801.1 | 12390 | 11430 | - | 319 | PF00248<br>TIGR01293 | Aldo_ket_red<br>TIGR01293 | Aldo/keto reductase family<br>Kv_beta: voltage-dependent potassium channel beta subunit | 8.30E-70<br>1.60E-37 |
| KPH97802.1 | 13036 | 12427 | - | 202 | PF00196<br>TIGR03541<br>PF00072<br>TIGR03020<br>TIGR02875 | GerE<br>TIGR03541<br>Response_reg<br>TIGR03020<br>TIGR02875 | Bacterial regulatory proteins, luxR family<br>reg_near_HchA: LuxR family transcriptional regulatory, chaperone HchA-associated<br>Response regulator receiver domain<br>EpsA: transcriptional regulator EpsA<br>spore_0_A: sporulation transcription factor Spo0A | 4.90E-13<br>1.20E-08<br>1.50E-06<br>6.00E-05<br>2.30E-04 |
| KPH97803.1 | 13486 | 13717 | + | 76 | NO PFAM MATCH | - | - | - |
| KPH97804.1 | 13757 | 13991 | + | 77 | LinA | LinA | LinA (linaridin precursor) | 4.10E-06 |
| KPH97805.1 | 14222 | 16211 | + | 662 | LinH<br>LinG<br>LinE<br>PF12697<br>PF12146 | LinH<br>LinG<br>LinE<br>Abhydrolase_6<br>Hydrolase_4 | LinH (Fusion of LinE/LinG)<br>LinG Homolog<br>LinE Homolog<br>Alpha/beta hydrolase family<br>Serine aminopeptidase, S33 | 7.70E-233<br>5.00E-122<br>4.00E-90<br>8.90E-16<br>6.30E-08 |
| KPH97806.1 | 16207 | 16816 | + | 202 | LinL | LinL | LinL | 9.80E-54 |

| | | | | | | | | |
|---|---|---|---|---|---|---|---|---|
| KPH97807.1 | 16812 | 18765 | + | 650 | TIGR02204<br>TIGR02203<br>TIGR01846<br>TIGR03796<br>TIGR00958 | TIGR02204<br>TIGR02203<br>TIGR01846<br>TIGR03796<br>TIGR00958 | MsbA_rel: ABC transporter, permease/ATP-binding protein<br>MsbA_lipidA: lipid A export permease/ATP-binding protein MsbA<br>type_I_sec_HlyB: type I secretion system ATPase<br>NHLM_micro_ABC1: NHLM bacteriocin system ABC transporter, peptidase/ATP-binding protein<br>3a01208: antigen peptide transporter 2 | 8.90E-97<br>2.70E-92<br>4.90E-85<br>5.50E-84<br>6.30E-83 |
| KPH97808.1 | 18761 | 20252 | + | 496 | TIGR02734<br>PF13450<br>TIGR00292<br>TIGR03467<br>TIGR02733 | TIGR02734<br>NAD_binding_8<br>TIGR00292<br>TIGR03467<br>TIGR02733 | crtI_fam: phytoene desaturase<br>NAD(P)-binding Rossmann-like domain<br>TIGR00292: thiazole biosynthesis enzyme<br>HpnE: squalene-associated FAD-dependent desaturase<br>desat_CrtD: C-3',4' desaturase CrtD | 1.20E-11<br>8.80E-09<br>4.30E-08<br>3.90E-07<br>1.40E-06 |
| KPH97809.1 | 20238 | 20949 | + | 236 | NO PFAM MATCH | - | - | - |
| KPH97810.1 | 21041 | 21926 | + | 294 | TIGR02947<br>TIGR02950<br>TIGR02939<br>TIGR02948<br>TIGR02937 | TIGR02947<br>TIGR02950<br>TIGR02939<br>TIGR02948<br>TIGR02937 | SigH_actino: RNA polymerase sigma-70 factor, TIGR02947 family<br>SigM_subfam: RNA polymerase sigma factor, SigM family<br>RpoE_Sigma70: RNA polymerase sigma factor RpoE<br>SigW_bacill: RNA polymerase sigma-W factor<br>sigma70-ECF: RNA polymerase sigma factor, sigma-70 family | 2.20E-108<br>5.40E-31<br>4.70E-29<br>6.20E-27<br>3.30E-26 |
| KPH97811.1 | 21922 | 22255 | + | 110 | TIGR02949<br>TIGR03988<br>PF13490 | TIGR02949<br>TIGR03988<br>zf-HC2 | anti_SigH_actin: anti-sigma factor, TIGR02949 family<br>antisig_RsrA: mycothiol system anti-sigma-R factor<br>Putative zinc-finger | 4.20E-36<br>3.90E-31<br>1.60E-08 |
| KPH97812.1 | 22789 | 22423 | - | 121 | NO PFAM MATCH | - | - | - |
| KPH97813.1 | 22954 | 23320 | + | 121 | NO PFAM MATCH | - | - | - |

| index | leader | core | start | end | dir | score |
|---|---|---|---|---|---|---|
| 1 | VDRDARGIGTPGTCAATA | STPTRTTSAAAPTPRATGRAPTSAPSTRTRPAAWSSPSTAAA | 7260 | 7077 | - | 9 |
| 2 | LGGVYGHAATLMASQWAGMRWFFGCGAVVAGRAHAAEPHN | VTAPRPYRGALAPRL | 8531 | 8699 | + | 6 |
| 3 | MPTGCAVCVPRHIFGCFLIDREYASPCPVCA | GPENLRVPLTDVIDVHDIGHAMNGSGIDIRV | 13207 | 13396 | + | 8 |
| 4 | VPRHIFGCFLIDREYASPCPVCAGPEN | LRVPLTDVIDVHDIGHAMNGSGIDIRV | 13231 | 13396 | + | 8 |
| 5 | LIDREYASPCPVCAGPENLRVP | LTDVIDVHDIGHAMNGSGIDIRV | 13258 | 13396 | + | 6 |
| 6 | MNAVAVHLNGLDLGSIRQGRAYDRQGHPVVSA | ATVVIGEVVGAVGGVATAVAGAVAASANDDACFDAADGGAGVCG | 13486 | 13717 | + | 6 |
| 7 | MNTAADHLNRLDLGPIRQSGVYAEPAAPIVSVAAAAVAVPVGGAVAGV | ATAFVASAIDAAQGDACFDAADGGAGVCG | 13757 | 13991 | + | 15 |

### Results for RJF42448.1 [Actinomyces sp. 2119] back to top

previous - next

#### Architecture

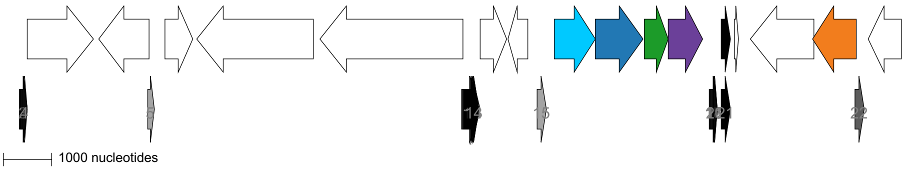

Link to nucleotide sequence

| Accession | start | end | direction | length (aa) | Pfam/HMM | name | description | E-value |
|---|---|---|---|---|---|---|---|---|
| RJF42441.1 | 40507 | 41893 | + | 461 | PF14501 | HATPase_c_5 | GHKL domain | 5.90E-22 |
| RJF42442.1 | 43034 | 41999 | - | 344 | PF00535<br>TIGR03937<br>PF13641<br>TIGR03965<br>TIGR04283 | Glycos_transf_2<br>TIGR03937<br>Glyco_tranf_2_3<br>TIGR03965<br>TIGR04283 | Glycosyl transferase family 2<br>PgaC_IcaA: poly-beta-1,6 N-acetyl-D-glucosamine synthase<br>Glycosyltransferase like family 2<br>mycofact_glyco: mycofactocin system glycosyltransferase<br>glyco_like_mftF: transferase 2, rSAM/selenodomain-associated | 1.60E-28<br>3.30E-19<br>5.30E-15<br>9.90E-15<br>5.10E-11 |
| WP_016461106.1 | 43354 | 43936 | + | 194 | INFERRED GENE | - | - | - |
| RJF42443.1 | 46434 | 44031 | - | 800 | PF00933<br>PF01915<br>PF14310 | Glyco_hydro_3<br>Glyco_hydro_3_C<br>Fn3-like | Glycosyl hydrolase family 3 N terminal domain<br>Glycosyl hydrolase family 3 C-terminal domain<br>Fibronectin type III-like domain | 9.10E-50<br>1.10E-47<br>4.80E-21 |
| RJF42444.1 | 49528 | 46582 | - | 981 | PF00933<br>PF01915<br>PF14310 | Glyco_hydro_3<br>Glyco_hydro_3_C<br>Fn3-like | Glycosyl hydrolase family 3 N terminal domain<br>Glycosyl hydrolase family 3 C-terminal domain<br>Fibronectin type III-like domain | 1.20E-33<br>1.70E-28<br>2.20E-14 |
| RJF42445.1 | 49903 | 50434 | + | 176 | PF09335 | SNARE_assoc | SNARE associated Golgi protein | 2.30E-10 |
| RJF42446.1 | 50891 | 50462 | - | 142 | PF09391 | DUF2000 | Protein of unknown function (DUF2000) | 3.10E-30 |

| | | | | | | | | |
|---|---|---|---|---|---|---|---|---|
| RJF42447.1 | 51426 | 52272 | + | 281 | LinH<br>LinG | LinH<br>LinG | LinH (Fusion of LinE/LinG)<br>LinG Homolog | 9.90E-95<br>3.10E-90 |
| RJF42448.1 | 52268 | 53252 | + | 327 | LinE<br>LinH<br>PF12697<br>PF12146<br>PF00561 | LinE<br>LinH<br>Abhydrolase_6<br>Hydrolase_4<br>Abhydrolase_1 | LinE Homolog<br>LinH (Fusion of LinE/LinG)<br>Alpha/beta hydrolase family<br>Serine aminopeptidase, S33<br>alpha/beta hydrolase fold | 3.90E-83<br>7.20E-67<br>1.10E-17<br>1.40E-10<br>1.10E-07 |
| RJF42449.1 | 53281 | 53788 | + | 168 | LinL | LinL | LinL | 5.00E-55 |
| RJF42450.1 | 53784 | 54507 | + | 240 | PF13649<br>PF08241<br>PF13489<br>PF08242<br>PF13847 | Methyltransf_25<br>Methyltransf_11<br>Methyltransf_23<br>Methyltransf_12<br>Methyltransf_31 | Methyltransferase domain<br>Methyltransferase domain<br>Methyltransferase domain<br>Methyltransferase domain<br>Methyltransferase domain | 4.10E-12<br>8.00E-11<br>5.80E-08<br>1.50E-07<br>1.00E-06 |
| RJF42451.1 | 54872 | 55061 | + | 62 | LinA | LinA | LinA (linaridin precursor) | 2.10E-12 |
| WP_003209438.1 | 55162 | 55249 | + | 29 | INFERRED GENE | - | - | - |
| RJF42452.1 | 56789 | 55493 | - | 431 | PF12698<br>PF12679 | ABC2_membrane_3<br>ABC2_membrane_2 | ABC-2 family transporter protein<br>ABC-2 family transporter protein | 5.20E-32<br>2.80E-08 |
| RJF42453.1 | 57666 | 56763 | - | 300 | TIGR01188<br>TIGR03522<br>TIGR04406<br>TIGR03864<br>TIGR03269 | TIGR01188<br>TIGR03522<br>TIGR04406<br>TIGR03864<br>TIGR03269 | drrA: daunorubicin resistance ABC transporter, ATP-binding protein<br>GldA_ABC_ATP: gliding motility-associated ABC transporter ATP-binding subunit GldA<br>LPS_export_lptB: LPS export ABC transporter ATP-binding protein<br>PQQ_ABC_ATP: ABC transporter, ATP-binding subunit, PQQ-dependent alcohol dehydrogenase system<br>met_CoM_red_A2: methyl coenzyme M reductase system, component A2 | 2.70E-50<br>1.30E-44<br>2.70E-43<br>1.70E-40<br>1.80E-38 |
| RJF42454.1 | 58602 | 57930 | - | 223 | PF00072<br>PF00196<br>TIGR02875<br>TIGR03020<br>TIGR01387 | Response_reg<br>GerE<br>TIGR02875<br>TIGR03020<br>TIGR01387 | Response regulator receiver domain<br>Bacterial regulatory proteins, luxR family<br>spore_0_A: sporulation transcription factor Spo0A<br>EpsA: transcriptional regulator EpsA<br>cztR_silR_copR: heavy metal response regulator | 8.00E-25<br>1.80E-18<br>2.80E-17<br>8.40E-13<br>1.50E-11 |

| index | leader | core | start | end | dir | score |
|---|---|---|---|---|---|---|
| 1 | VVGVHDQDCLMSTGEVLRVSRPRRRGLMSALADHVGGLPVSG | RPPGEAARTSRAST | 40340 | 40511 | + | 9 |
| 2 | MSTGEVLRVSRPRRRGLMSALADHVGGLPVSG | RPPGEAARTSRAST | 40370 | 40511 | + | 6 |
| 3 | VLRVSRPRRRGLMSALADHVGGLPVSG | RPPGEAARTSRAST | 40385 | 40511 | + | 6 |
| 4 | VSRPRRRGLMSALADHVGGLPVSG | RPPGEAARTSRAST | 40394 | 40511 | + | 6 |
| 5 | VPTGEGVPCTCPQSVRLSAS | CGVSCGAVRTRVTSCRPMPSP | 43009 | 43135 | + | 6 |
| 6 | LIDSMAEYLSVPLCTSVSVAAVPLP...AGPTCRFRDLFCTPPRPAHGTGPAP | LTAPNRHRTTRCRVVL | 49514 | 49862 | + | 9 |
| 7 | MAEYLSVPLCTSVSVAAVPLPAHGG...AGPTCRFRDLFCTPPRPAHGTGPAP | LTAPNRHRTTRCRVVL | 49526 | 49862 | + | 8 |
| 8 | LYLCVCGCCASPCARWEHGVGRAAC...HSQAGFPAAGPARRTNLSIPGPVLH | PAPPRSRHRARPAHGTKSS | 49552 | 49834 | + | 6 |
| 9 | VSVAAVPLPAHGGSTGWGGPPAAGG...AGPTCRFRDLFCTPPRPAHGTGPAP | LTAPNRHRTTRCRVVL | 49562 | 49862 | + | 9 |
| 10 | VAAVPLPAHGGSTGWGGPPAAGGTG...AGPTCRFRDLFCTPPRPAHGTGPAP | LTAPNRHRTTRCRVVL | 49568 | 49862 | + | 9 |
| 11 | VPLPAHGGSTGWGGPPAAGGTGMLA...AGPTCRFRDLFCTPPRPAHGTGPAP | LTAPNRHRTTRCRVVL | 49577 | 49862 | + | 9 |
| 12 | VGARGGAGRLPLAGRACWQCCPVLP...FPCGRASPQDQPVDSGTCSAPRPAP | LTAPGPPRSRHQIVIGQHGAVLSYDDYVALSRLS | 49596 | 49893 | + | 6 |
| 13 | MLAVLPGAAVVGHRQGGLGTVKPVS...AGPTCRFRDLFCTPPRPAHGTGPAP | LTAPNRHRTTRCRVVL | 49643 | 49862 | + | 9 |
| 14 | VLPGAAVVGHRQGGLGTVKPVSLRQ...AGPTCRFRDLFCTPPRPAHGTGPAP | LTAPNRHRTTRCRVVL | 49652 | 49862 | + | 10 |
| 15 | VVLWQSEHTDLDQPGDAPHP | LCPLQSRGRTVPGSPPGGGRAPQTTSHSRTGPGLR | 51082 | 51250 | + | 6 |
| 16 | VRMSRWYAFWLTTMFVGSVTYLTMFDHMDNPGRITSSVSLAITA | GTVVALFFVSSSDT | 54633 | 54810 | + | 7 |
| 17 | MSRWYAFWLTTMFVGSVTYLTMFDHMDNPGRITSSVSLAITA | GTVVALFFVSSSDT | 54639 | 54810 | + | 8 |
| 18 | MFVGSVTYLTMFDHMDNPGRITSSVSLAITA | GTVVALFFVSSSDT | 54672 | 54810 | + | 6 |
| 19 | MFDHMDNPGRITSSVSLAITA | GTVVALFFVSSSDT | 54702 | 54810 | + | 6 |

| | | | | | | |
|---|---|---|---|---|---|---|
| 20 | MDNPGRITSSVSLAITA | GTVVALFFVSSSDT | 54714 | 54810 | + | 6 |
| 21 | MPILLDKVNSDPALKEMLQLRASRSEAPVL | ATPAALAWAAGAALGGGSVGGAAYAAYRAVAN | 54872 | 55061 | + | 29 |
| 22 | VPNLQHHHLLAVCLSLPASVFL | PTTRLGATSQHLTLPGAGTSPCRGRRAHSPPPLPTKEQTQPHT | 57648 | 57846 | + | 8 |

### Results for WP_119836030.1 [Actinomyces sp. 2119] back to top

previous - next

#### Architecture

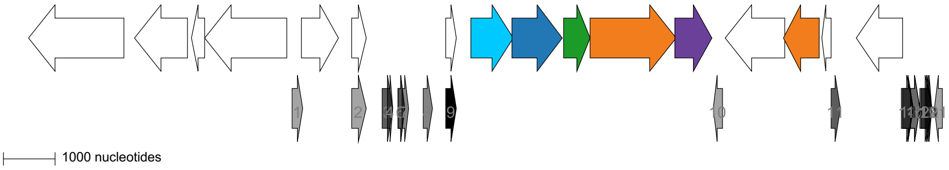

Link to nucleotide sequence

| Accession | start | end | direction | length (aa) | Pfam/HMM | name | description | E-value |
|---|---|---|---|---|---|---|---|---|
| WP_147392851.1 | 88718 | 86882 | - | 611 | NO PFAM MATCH | - | - | - |
| WP_119836023.1 | 89955 | 88935 | - | 339 | TIGR01484<br>TIGR00099<br>PF08282<br>TIGR01487<br>PF03332 | TIGR01484<br>TIGR00099<br>Hydrolase_3<br>TIGR01487<br>PMM | HAD-SF-IIB: HAD hydrolase, family IIB<br>Cof-subfamily: Cof-like hydrolase<br>haloacid dehalogenase-like hydrolase<br>Pglycolate_arch: phosphoglycolate phosphatase, TA0175-type<br>Eukaryotic phosphomannomutase | 1.80E-14<br>9.80E-09<br>1.60E-08<br>6.60E-07<br>6.80E-06 |
| WP_119836024.1 | 90277 | 90040 | - | 78 | PF16951 | MaAIMP_sms | Putative methionine and alanine importer, small subunit | 2.60E-06 |
| WP_119836025.1 | 91870 | 90280 | - | 529 | PF00209 | SNF | Sodium:neurotransmitter symporter family | 3.80E-95 |
| WP_119836026.1 | 92160 | 92871 | + | 236 | TIGR00107<br>PF01048<br>TIGR01718<br>TIGR01704<br>TIGR03468 | TIGR00107<br>PNP_UDP_1<br>TIGR01718<br>TIGR01704<br>TIGR03468 | deoD: purine nucleoside phosphorylase<br>Phosphorylase superfamily<br>Uridine-psphlse: uridine phosphorylase<br>MTA/SAH-Nsdase: MTA/SAH nucleosidase<br>HpnG: hopanoid-associated phosphorylase | 2.30E-78<br>5.10E-40<br>4.30E-36<br>5.40E-15<br>9.40E-14 |
| WP_119836027.1 | 93125 | 93407 | + | 93 | PF07561 | DUF1540 | Domain of Unknown Function (DUF1540) | 3.00E-08 |
| WP_119836028.1 | 94943 | 95153 | + | 69 | NO PFAM MATCH | - | - | - |
| WP_119836029.1 | 95446 | 96247 | + | 266 | LinH<br>LinG<br>PF05090 | LinH<br>LinG<br>VKG_Carbox | LinH (Fusion of LinE/LinG)<br>LinG Homolog<br>Vitamin K-dependent gamma-carboxylase | 8.00E-67<br>9.60E-66<br>4.00E-09 |
| WP_119836030.1 | 96243 | 97215 | + | 323 | LinE<br>LinH<br>PF12697<br>PF00561<br>TIGR02427 | LinE<br>LinH<br>Abhydrolase_6<br>Abhydrolase_1<br>TIGR02427 | LinE Homolog<br>LinH (Fusion of LinE/LinG)<br>Alpha/beta hydrolase family<br>alpha/beta hydrolase fold<br>protocat_pcaD: 3-oxoadipate enol-lactonase | 4.40E-57<br>3.80E-51<br>1.30E-16<br>1.30E-14<br>2.00E-12 |
| WP_147392852.1 | 97229 | 97748 | + | 172 | LinL | LinL | LinL | 1.50E-27 |
| WP_119836031.1 | 97744 | 99403 | + | 552 | TIGR03375<br>TIGR00958<br>TIGR02868<br>TIGR02204<br>TIGR02857 | TIGR03375<br>TIGR00958<br>TIGR02868<br>TIGR02204<br>TIGR02857 | type_I_sec_LssB: type I secretion system ATPase<br>3a01208: antigen peptide transporter 2<br>CydC: thiol reductant ABC exporter, CydC subunit<br>MsbA_rel: ABC transporter, permease/ATP-binding protein<br>CydD: thiol reductant ABC exporter, CydD subunit | 2.60E-57<br>1.80E-50<br>3.00E-50<br>1.00E-49<br>6.70E-49 |
| WP_119836032.1 | 99389 | 100106 | + | 238 | PF13649<br>PF13489<br>PF08241<br>TIGR02021<br>TIGR04345 | Methyltransf_25<br>Methyltransf_23<br>Methyltransf_11<br>TIGR02021<br>TIGR04345 | Methyltransferase domain<br>Methyltransferase domain<br>Methyltransferase domain<br>BchM-ChlM: magnesium protoporphyrin O-methyltransferase<br>ovoA_Cterm: putative 4-mercaptohistidine N1-methyltranferase | 7.50E-10<br>6.00E-09<br>1.10E-07<br>1.70E-06<br>1.70E-06 |
| WP_147392853.1 | 101515 | 100348 | - | 388 | NO PFAM MATCH | - | - | - |
| WP_119836034.1 | 102176 | 101489 | - | 228 | TIGR01188<br>TIGR04406<br>TIGR01288<br>TIGR02315<br>TIGR02857 | TIGR01188<br>TIGR04406<br>TIGR01288<br>TIGR02315<br>TIGR02857 | drrA: daunorubicin resistance ABC transporter, ATP-binding protein<br>LPS_export_lptB: LPS export ABC transporter ATP-binding protein<br>nodI: nodulation ABC transporter NodI<br>ABC_phnC: phosphonate ABC transporter, ATP-binding protein<br>CydD: thiol reductant ABC exporter, CydD subunit | 2.40E-21<br>3.00E-19<br>1.60E-18<br>2.00E-18<br>2.80E-18 |
| WP_147392854.1 | 102417 | 102225 | - | 63 | NO PFAM MATCH | - | - | - |
| WP_119836181.1 | 103790 | 102893 | - | 298 | NO PFAM MATCH | - | - | - |

| index | leader | core | start | end | dir | score |
|---|---|---|---|---|---|---|
| 1 | MWTAAWPAAGQTTARLPTWQPPATSLVVQAPCA | SCPVPARETQVDRGGLQKIIHRKVPHGHTAH | 91985 | 92180 | + | 6 |
| 2 | MTAVAQITSCSTTSCAFNNGGCTAFAITVGGDTGAPSC | GTFLSLDARGGLGVADAHVGACQRL...TDLMCTAEAVQVGGDTAACLSYQAR | 93125 | 93407 | + | 6 |
| 3 | VKGELRNAYAAFCLWHGIGRIRNFCWGEPDYAA | TMCNFDHQGYGPVGRSSGRRTTLTQGCTAGHHR | 93716 | 93917 | + | 8 |
| 4 | MCNFDHQGYGPVGRSS | GRRTTLTQGCTAGHHR | 93818 | 93917 | + | 7 |
| 5 | VRPAMRGSGTDPAGTFPCSLGQ | SGCDKHTHRPTAGTSSILMLSGT | 94023 | 94161 | + | 6 |
| 6 | MRGSGTDPAGTFPCSLGQSG | CDKHTHRPTAGTSSILMLSGT | 94035 | 94161 | + | 7 |
| 7 | VRQAYPQAHCRYEFNLDALRDLT | YIQTTGTTSSRLHTHEALLLRRDR | 94094 | 94238 | + | 7 |
| 8 | MHAVSVVLHQTTESATQTLWTKLESQ | ALPAQPPQTAQAARTPRVPCSYPHPLV | 94525 | 94687 | + | 7 |

| | | | | | | |
|---|---|---|---|---|---|---|
| 9 | MDRFIEMVNNDQTNNKIPTASDDFCDQGVMPMT | ATPVAARAAVATSRAVARTVTRRAVRRTQRGLPFPV | 94943 | 95153 | + | 21 |
| 10 | LPTSCCHCYHDGAGP | PCPASTAGPDLSGTALGKTSAGGVRTDILDTCRYSSTVLPILSGLVHSQR | 100313 | 100115 | - | 6 |
| 11 | VACLILGTILTFQFNARGRTADKAGCGKVAY | RTPFPVRNNCWLKRHADTLTHLSDRHLHE | 102394 | 102577 | + | 8 |
| 12 | VRSSMWGALALPAWGDVTRVPSCRVLAWRHGRPG | RWPRPGGEATLSGRFFLALVRDVDMEPVSGGQPRV | 103776 | 103986 | + | 6 |
| 13 | MWGALALPAWGDVTRVPSCRVLAWRHGRPGRW | PRPGGEATLSGRFFLALVRDVDMEPVSGGQPRV | 103788 | 103986 | + | 7 |
| 14 | LALPAWGDVTRVPSCRVLAWRHGRPGRWPR | PGGEATLSGRFFLALVRDVDMEPVSGGQPRV | 103800 | 103986 | + | 7 |
| 15 | VMWTWSPSQAVSPGCKNLAVQAVSRSRHLLV | VPPCAGPARGAYPGRHTHKAGRTSPA | 103939 | 104113 | + | 6 |
| 16 | MWTWSPSQAVSPGCKNLAVQAVSRSRHLLV | VPPCAGPARGAYPGRHTHKAGRTSPA | 103942 | 104113 | + | 6 |
| 17 | VELRGFEPLTSSMPWKRATNCAIA | PTPAAAPTQGQIPQVVELGGFEPPT...PWKRATNCAIAPSGRPPTRATQATV | 104122 | 104356 | + | 7 |
| 18 | MPWKRATNCAIA | PTPAAAPTQGQIPQVVELGGFEPPT...PWKRATNCAIAPSGRPPTRATQATV | 104158 | 104356 | + | 8 |
| 19 | VVELGGFEPPTSSMPWKRA | TNCAIAPSGRPPTRATQATV | 104236 | 104356 | + | 6 |
| 20 | VELGGFEPPTSSMPWKRAT | NCAIAPSGRPPTRATQATV | 104239 | 104356 | + | 7 |
| 21 | VAWATGTTRGCGPRTVSPGGRSWAG | TCSRRCTRREHPGTGSTESSVAAAGG | 104555 | 104399 | - | 6 |

### Results for WP_120204651.1 [Actinomyces sp. 2129] back to top

previous - next

#### Architecture

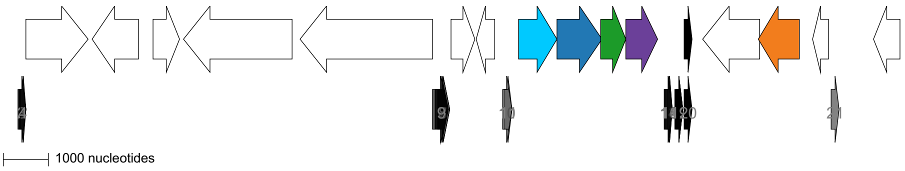

Link to nucleotide sequence

| Accession | start | end | direction | length (aa) | Pfam/HMM | name | description | E-value |
|---|---|---|---|---|---|---|---|---|
| WP_120204638.1 | 1837962 | 1839348 | + | 461 | PF14501 | HATPase_c_5 | GHKL domain | 5.90E-22 |
| WP_120204640.1 | 1840489 | 1839454 | - | 344 | PF00535<br>TIGR03937<br>PF13641<br>TIGR03965<br>TIGR04283 | Glycos_transf_2<br>TIGR03937<br>Glyco_tranf_2_3<br>TIGR03965<br>TIGR04283 | Glycosyl transferase family 2<br>PgaC_IcaA: poly-beta-1,6 N-acetyl-D-glucosamine synthase<br>Glycosyltransferase like family 2<br>mycofact_glyco: mycofactocin system glycosyltransferase<br>glyco_like_mftF: transferase 2, rSAM/selenodomain-associated | 1.60E-28<br>3.30E-19<br>5.00E-15<br>9.90E-15<br>5.10E-11 |
| WP_016461106.1 | 1840809 | 1841391 | + | 194 | INFERRED GENE | - | - | - |
| WP_120204642.1 | 1843889 | 1841486 | - | 800 | PF00933<br>PF01915<br>PF14310 | Glyco_hydro_3<br>Glyco_hydro_3_C<br>Fn3-like | Glycosyl hydrolase family 3 N terminal domain<br>Glycosyl hydrolase family 3 C-terminal domain<br>Fibronectin type III-like domain | 3.90E-50<br>1.10E-47<br>4.80E-21 |
| WP_120204644.1 | 1846996 | 1844050 | - | 981 | PF00933<br>PF01915<br>PF14310 | Glyco_hydro_3<br>Glyco_hydro_3_C<br>Fn3-like | Glycosyl hydrolase family 3 N terminal domain<br>Glycosyl hydrolase family 3 C-terminal domain<br>Fibronectin type III-like domain | 1.10E-33<br>1.70E-28<br>3.80E-15 |
| WP_120204645.1 | 1847412 | 1847943 | + | 176 | PF09335 | SNARE_assoc | SNARE associated Golgi protein | 2.30E-10 |
| WP_120204647.1 | 1848400 | 1847971 | - | 142 | PF09391 | DUF2000 | Protein of unknown function (DUF2000) | 8.80E-30 |
| WP_120204649.1 | 1848932 | 1849778 | + | 281 | LinH<br>LinG | LinH<br>LinG | LinH (Fusion of LinE/LinG)<br>LinG Homolog | 9.50E-96<br>9.40E-91 |
| WP_120204651.1 | 1849774 | 1850758 | + | 327 | LinE<br>LinH<br>PF12697<br>PF12146<br>PF00561 | LinE<br>LinH<br>Abhydrolase_6<br>Hydrolase_4<br>Abhydrolase_1 | LinE Homolog<br>LinH (Fusion of LinE/LinG)<br>Alpha/beta hydrolase family<br>Serine aminopeptidase, S33<br>alpha/beta hydrolase fold | 7.00E-82<br>1.10E-65<br>2.20E-17<br>2.30E-10<br>7.20E-08 |
| WP_147392849.1 | 1850754 | 1851294 | + | 179 | LinL | LinL | LinL | 2.00E-56 |

| | | | | | | | | |
|---|---|---|---|---|---|---|---|---|
| WP_120204652.1 | 1851290 | 1852013 | + | 240 | PF13649<br>PF08241<br>PF13489<br>PF08242<br>PF13847 | Methyltransf_25<br>Methyltransf_11<br>Methyltransf_23<br>Methyltransf_12<br>Methyltransf_31 | Methyltransferase domain<br>Methyltransferase domain<br>Methyltransferase domain<br>Methyltransferase domain<br>Methyltransferase domain | 1.30E-12<br>1.90E-11<br>3.00E-08<br>5.30E-08<br>5.80E-07 |
| WP_162933885.1 | 1852585 | 1852765 | + | 59 | LinA | LinA | LinA (linaridin precursor) | 2.20E-11 |
| WP_120204654.1 | 1854277 | 1852993 | - | 427 | PF12698<br>PF12679 | ABC2_membrane_3<br>ABC2_membrane_2 | ABC-2 family transporter protein<br>ABC-2 family transporter protein | 2.10E-32<br>3.70E-08 |
| WP_120204656.1 | 1855154 | 1854251 | - | 300 | TIGR01188<br>TIGR04406<br>TIGR03522<br>TIGR03864<br>TIGR03269 | TIGR01188<br>TIGR04406<br>TIGR03522<br>TIGR03864<br>TIGR03269 | drrA: daunorubicin resistance ABC transporter, ATP-binding protein<br>LPS_export_lptB: LPS export ABC transporter ATP-binding protein<br>GldA_ABC_ATP: gliding motility-associated ABC transporter ATP-binding subunit GldA<br>PQQ_ABC_ATP: ABC transporter, ATP-binding subunit, PQQ-dependent alcohol dehydrogenase system<br>met_CoM_red_A2: methyl coenzyme M reductase system, component A2 | 3.00E-50<br>4.00E-44<br>1.50E-43<br>3.90E-40<br>4.50E-38 |
| WP_120204658.1 | 1855806 | 1855437 | - | 122 | PF13630<br>TIGR04222 | SdpI<br>TIGR04222 | SdpI/YhfL protein family<br>near_uncomplex: TIGR04222 domain | 2.30E-06<br>2.60E-04 |
| WP_120204660.1 | 1857373 | 1856797 | - | 191 | PF17418<br>TIGR04034 | SdpA<br>TIGR04034 | Sporulation delaying protein SdpA<br>export_SdpA: antimicrobial peptide system protein, SdpA family | 4.70E-25<br>6.90E-21 |

| index | leader | core | start | end | dir | score |
|---|---|---|---|---|---|---|
| 1 | VVGVHDQDCLMSTGEVLRVSRPRRRGLMSALADHVGGLPVSG | RPPGEAARTSRAST | 1837795 | 1837966 | + | 9 |
| 2 | MSTGEVLRVSRPRRRGLMSALADHVGGLPVSG | RPPGEAARTSRAST | 1837825 | 1837966 | + | 6 |
| 3 | VLRVSRPRRRGLMSALADHVGGLPVSG | RPPGEAARTSRAST | 1837840 | 1837966 | + | 6 |
| 4 | VSRPRRRGLMSALADHVGGLPVSG | RPPGEAARTSRAST | 1837849 | 1837966 | + | 6 |
| 5 | LYLCVCGCCASPCARWEHGVGRAAC...APLTAPGPPRSRHRARPAHGTGPAP | LTAPNRHRTTRCRVVL | 1847020 | 1847371 | + | 8 |
| 6 | VGRAACRWRDGHAGSAARCCRRRSP...APLTAPGPPRSRHRARPAHGTGPAP | LTAPNRHRTTRCRVVL | 1847077 | 1847371 | + | 9 |
| 7 | MLAVLPGAAVVGHRQGGLGAVKPVS...FRDLFCTPPRSRHRARPAHGTGPAP | LTAPGPPRSRHQIVIGQHGAVLSYDDYVALSRLS | 1847111 | 1847402 | + | 6 |
| 8 | VLPGAAVVGHRQGGLGAVKPVSLRQ...FRDLFCTPPRSRHRARPAHGTGPAP | LTAPGPPRSRHQIVIGQHGAVLSYDDYVALSRLS | 1847120 | 1847402 | + | 6 |
| 9 | VLPSSVTVREAWGQSSRFPCGRASPQDQPVDSGTCSVPRPAHGTGPAP | LTAPGPPRSRHRARPAHGTKSS | 1847130 | 1847343 | + | 6 |
| 10 | MIITPAARGPVAVGAHRPRPARR | RTTPLCPLQSRGRTVPGSPPGGGRAPQTTSHSRTGPGLR | 1848568 | 1848757 | + | 6 |
| 11 | VAVGAHRPRPARR | RTTPLCPLQSRGRTVPGSPPGGGRAPQTTSHSRTGPGLR | 1848598 | 1848757 | + | 6 |
| 12 | VRMSRWYAFWLTTMFVGSVTYLTMFDHMDNPGRITSSVSLAITA | GTVVALFFVSSSDT | 1852139 | 1852316 | + | 7 |
| 13 | MSRWYAFWLTTMFVGSVTYLTMFDHMDNPGRITSSVSLAITA | GTVVALFFVSSSDT | 1852145 | 1852316 | + | 8 |
| 14 | MFVGSVTYLTMFDHMDNPGRITSSVSLAITA | GTVVALFFVSSSDT | 1852178 | 1852316 | + | 6 |
| 15 | MFDHMDNPGRITSSVSLAITA | GTVVALFFVSSSDT | 1852208 | 1852316 | + | 6 |
| 16 | MDNPGRITSSVSLAITA | GTVVALFFVSSSDT | 1852220 | 1852316 | + | 6 |
| 17 | MPILLDKVNSDPALKEMLQLRASRSEAPVL | ATPAALAWAAGAALGGGSVGGAAYAAYRAVAN | 1852378 | 1852567 | + | 29 |
| 18 | LKEMLQLRASRSEAPVL | ATPAALAWAAGAALGGGSVGGAAYAAYRAVAN | 1852417 | 1852567 | + | 15 |
| 19 | MLQLRASRSEAPVL | ATPAALAWAAGAALGGGSVGGAAYAAYRAVAN | 1852426 | 1852567 | + | 15 |
| 20 | MNELLDKVNSDTVVVVDRAQVPVL | ATPAAFVAGLGAGAKAAGALVAAAAGGAAVGAATR | 1852585 | 1852765 | + | 30 |
| 21 | MVRTQVIEQRKLHNPKVNAKTRCRTK | PNTISLILTCGENTAASAKDASTRSTI | 1855862 | 1856024 | + | 7 |

### Results for WP_127841862.1 [Actinomyces sp. 299] back to top

previous - next

#### Architecture

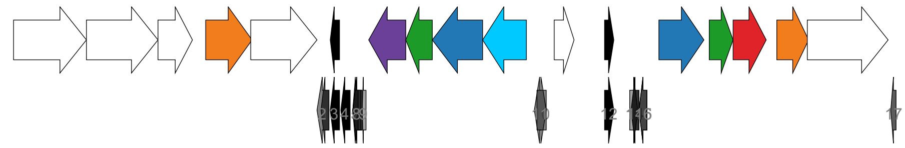

1000 nucleotides

Link to nucleotide sequence

| Accession | start | end | direction | length (aa) | Pfam/HMM | name | description | E-value |
|---|---|---|---|---|---|---|---|---|
| WP_127841855.1 | 1153119 | 1154547 | + | 475 | TIGR02734<br>PF13450<br>TIGR03467<br>PF01593<br>TIGR01317 | TIGR02734<br>NAD_binding_8<br>TIGR03467<br>Amino_oxidase<br>TIGR01317 | crtI_fam: phytoene desaturase<br>NAD(P)-binding Rossmann-like domain<br>HpnE: squalene-associated FAD-dependent desaturase<br>Flavin containing amine oxidoreductase<br>GOGAT_sm_gam: glutamate synthase, NADH/NADPH, small subunit | 4.50E-10<br>5.10E-10<br>1.40E-07<br>3.20E-07<br>5.30E-07 |
| WP_127841856.1 | 1154543 | 1155959 | + | 471 | PF07730 | HisKA_3 | Histidine kinase | 2.70E-10 |
| WP_127841857.1 | 1155955 | 1156627 | + | 223 | PF00072<br>PF00196<br>TIGR02875<br>TIGR03020<br>TIGR01387 | Response_reg<br>GerE<br>TIGR02875<br>TIGR03020<br>TIGR01387 | Response regulator receiver domain<br>Bacterial regulatory proteins, luxR family<br>spore_0_A: sporulation transcription factor Spo0A<br>EpsA: transcriptional regulator EpsA<br>cztR_silR_copR: heavy metal response regulator | 3.20E-25<br>1.80E-18<br>4.60E-18<br>1.30E-12<br>1.30E-11 |
| WP_127841858.1 | 1156893 | 1157796 | + | 300 | TIGR01188<br>TIGR04406<br>TIGR03522<br>TIGR03864<br>TIGR03740 | TIGR01188<br>TIGR04406<br>TIGR03522<br>TIGR03864<br>TIGR03740 | drrA: daunorubicin resistance ABC transporter, ATP-binding protein<br>LPS_export_lptB: LPS export ABC transporter ATP-binding protein<br>GldA_ABC_ATP: gliding motility-associated ABC transporter ATP-binding subunit GldA<br>PQQ_ABC_ATP: ABC transporter, ATP-binding subunit, PQQ-dependent alcohol dehydrogenase system<br>galliderm_ABC: lantibiotic protection ABC transporter, ATP-binding subunit | 4.60E-50<br>4.80E-44<br>5.60E-44<br>4.30E-40<br>1.20E-38 |
| WP_127841859.1 | 1157770 | 1159078 | + | 435 | PF12698<br>PF12679 | ABC2_membrane_3<br>ABC2_membrane_2 | ABC-2 family transporter protein<br>ABC-2 family transporter protein | 6.40E-28<br>7.10E-07 |
| WP_164860076.1 | 1159500 | 1159320 | - | 59 | LinA | LinA | LinA (linaridin precursor) | 1.20E-11 |
| WP_127841860.1 | 1160804 | 1160081 | - | 240 | PF13649<br>PF08241<br>PF08242<br>PF13489<br>TIGR01934 | Methyltransf_25<br>Methyltransf_11<br>Methyltransf_12<br>Methyltransf_23<br>TIGR01934 | Methyltransferase domain<br>Methyltransferase domain<br>Methyltransferase domain<br>Methyltransferase domain<br>MenG_MenH_UbiE: ubiquinone/menaquinone biosynthesis methyltransferase | 7.30E-12<br>2.70E-11<br>2.80E-08<br>6.40E-08<br>3.60E-06 |
| WP_127841861.1 | 1161340 | 1160800 | - | 179 | LinL | LinL | LinL | 1.10E-56 |
| WP_127841862.1 | 1162320 | 1161336 | - | 327 | LinE<br>LinH<br>PF12697<br>PF12146<br>PF00561 | LinE<br>LinH<br>Abhydrolase_6<br>Hydrolase_4<br>Abhydrolase_1 | LinE Homolog<br>LinH (Fusion of LinE/LinG)<br>Alpha/beta hydrolase family<br>Serine aminopeptidase, S33<br>alpha/beta hydrolase fold | 6.60E-82<br>3.60E-65<br>5.80E-16<br>1.40E-11<br>1.10E-07 |
| WP_127841863.1 | 1163162 | 1162316 | - | 281 | LinH<br>LinG | LinH<br>LinG | LinH (Fusion of LinE/LinG)<br>LinG Homolog | 8.80E-95<br>2.70E-90 |
| WP_127841864.1 | 1163721 | 1164102 | + | 126 | PF09391 | DUF2000 | Protein of unknown function (DUF2000) | 3.50E-21 |
| WP_139738604.1 | 1164697 | 1164886 | + | 62 | LinA | LinA | LinA (linaridin precursor) | 2.10E-10 |
| WP_127841865.1 | 1165762 | 1166659 | + | 298 | LinE<br>LinH<br>PF12697<br>TIGR02427 | LinE<br>LinH<br>Abhydrolase_6<br>TIGR02427 | LinE Homolog<br>LinH (Fusion of LinE/LinG)<br>Alpha/beta hydrolase family<br>protocat_pcaD: 3-oxoadipate enol-lactonase | 8.50E-68<br>7.90E-41<br>2.50E-06<br>5.80E-06 |
| WP_127841866.1 | 1166753 | 1167227 | + | 157 | LinL | LinL | LinL | 1.10E-36 |
| WP_164860077.1 | 1167223 | 1167874 | + | 216 | PF02441<br>TIGR00521<br>TIGR02113 | Flavoprotein<br>TIGR00521<br>TIGR02113 | Flavoprotein<br>coaBC_dfp: phosphopantothenoylcysteine decarboxylase / phosphopantothenate--cysteine ligase<br>coaC_strep: phosphopantothenoylcysteine decarboxylase | 1.10E-12<br>8.40E-11<br>3.60E-07 |

| | | | | | | | | |
|---|---|---|---|---|---|---|---|---|
| WP_164860078.1 | 1168079 | 1168688 | + | 202 | TIGR01189<br>TIGR02857<br>TIGR01277<br>TIGR01188<br>TIGR02868 | TIGR01189<br>TIGR02857<br>TIGR01277<br>TIGR01188<br>TIGR02868 | ccmA: heme ABC exporter, ATP-binding protein CcmA<br>CydD: thiol reductant ABC exporter, CydD subunit<br>thiQ: thiamine ABC transporter, ATP-binding protein<br>drrA: daunorubicin resistance ABC transporter, ATP-binding protein<br>CydC: thiol reductant ABC exporter, CydC subunit | 6.80E-30<br>9.10E-23<br>1.00E-22<br>1.10E-22<br>1.50E-21 |
| WP_127841869.1 | 1168684 | 1170250 | + | 521 | NO PFAM MATCH | - | - | - |

| index | leader | core | start | end | dir | score |
|---|---|---|---|---|---|---|
| 1 | VAETQHRTHGDIQAPGA | LTPQAVAPGACPAATFLVGDRVAAQPYLGQR | 1159299 | 1159152 | - | 8 |
| 2 | VACGGNSAPNPRRHTGAWSTDTAGSCSRRLSCCYLSR | WRQSCGSTLPRTTMTAKTTSGLPRNAKKKPCALPSEKA | 1159306 | 1159078 | - | 7 |
| 3 | MNELLDKVNSDTVVIVDRTQVPVL | ATPAAFLGGLAAGAKAAGALVGAAAGGAAVGAATR | 1159500 | 1159320 | - | 30 |
| 4 | MQILLDKVNSDPTLKGSLRLRASRVGTPVL | ATPAALAWAAGAAVGAGAVSGAAYGAYRVAAR | 1159715 | 1159526 | - | 30 |
| 5 | MFDHMANPGRAASSIVLA | VTAGVVVALLFVSSPDD | 1159886 | 1159778 | - | 6 |
| 6 | MYAGPVAYLLMFDHMANPGRAASSIVLA | VTAGVVVALLFVSSPDD | 1159916 | 1159778 | - | 6 |
| 7 | MSRWYAFWLTTMYAGPVAYLLMFDHMANPGRAASSIVLA | VTAGVVVALLFVSSPDD | 1159949 | 1159778 | - | 7 |
| 8 | VQMSRWYAFWLTTMYAGPVAYLLMFDHMANPGRAASSIVLA | VTAGVVVALLFVSSPDD | 1159955 | 1159778 | - | 7 |
| 9 | MRVTQCFRASQQPLK | DVRTKHLACLLGGADV | 1160037 | 1159941 | - | 6 |
| 10 | VGPFLLLAESLEPCAAGAGRALGALRCG | RGQRGCGAPPGWARSVCSDRHRSTGRRAL | 1163379 | 1163553 | + | 6 |
| 11 | VIITPAALCSCGGRSTPTSPSPAAHHTPSALVRNGEHPELDQPRQ | RTVPGSPPEGGKAPRTSSHSRTASGLW | 1163557 | 1163338 | - | 7 |
| 12 | MSDLGIAPTVEQLEKVANREFSGISEGFVNSHNLPTM | TTPVTPATPGALEVVVTGSTICLGC | 1164697 | 1164886 | + | 31 |
| 13 | VSSCGGGRTRINQRDRKLQVPLWQR | RGRPDVRRNMRLQGCYWPHTRSQPVR | 1165198 | 1165354 | + | 6 |
| 14 | LRVNSPEVDVIGQAVIWYEASNSPV | AAYYGGHLVCPVVAIGAPVTFDHAG | 1165382 | 1165229 | - | 9 |
| 15 | LIGQGNAVVAPQLHLIRTDLFQGPSE | ERCQKARNSKLTSLSQESSRPEMFAR | 1165530 | 1165371 | - | 7 |
| 16 | MLAPLIGQGNAVVAPQLHLIRTDLFQGP | SEERCQKARNSKLTSLSQESSRPEMFAR | 1165542 | 1165371 | - | 7 |
| 17 | VKRSKHPHQASTKALTD | ITDHRTASEAITMHAAYV | 1170422 | 1170314 | - | 8 |

### Results for WP_127841865.1 [Actinomyces sp. 299] back to top

previous - next

#### Architecture

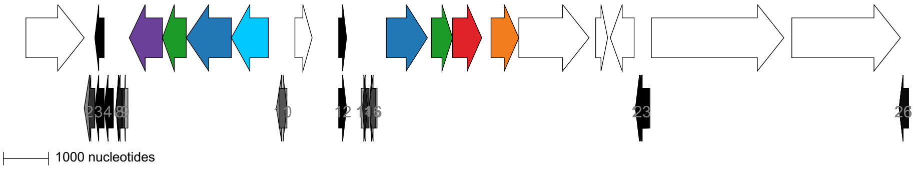

Link to nucleotide sequence

| Accession | start | end | direction | length (aa) | Pfam/HMM | name | description | E-value |
|---|---|---|---|---|---|---|---|---|
| WP_127841859.1 | 1157770 | 1159078 | + | 435 | PF12698<br>PF12679 | ABC2_membrane_3<br>ABC2_membrane_2 | ABC-2 family transporter protein<br>ABC-2 family transporter protein | 6.40E-28<br>7.10E-07 |
| WP_164860076.1 | 1159500 | 1159320 | - | 59 | LinA | LinA | LinA (linaridin precursor) | 1.20E-11 |
| WP_127841860.1 | 1160804 | 1160081 | - | 240 | PF13649<br>PF08241<br>PF08242<br>PF13489<br>TIGR01934 | Methyltransf_25<br>Methyltransf_11<br>Methyltransf_12<br>Methyltransf_23<br>TIGR01934 | Methyltransferase domain<br>Methyltransferase domain<br>Methyltransferase domain<br>Methyltransferase domain<br>MenG_MenH_UbiE: ubiquinone/menaquinone biosynthesis methyltransferase | 7.30E-12<br>2.70E-11<br>2.80E-08<br>6.40E-08<br>3.60E-06 |
| WP_127841861.1 | 1161340 | 1160800 | - | 179 | LinL | LinL | LinL | 1.10E-56 |

| | | | | | | | | |
|---|---|---|---|---|---|---|---|---|
| WP_127841862.1 | 1162320 | 1161336 | - | 327 | LinE<br>LinH<br>PF12697<br>PF12146<br>PF00561 | LinE<br>LinH<br>Abhydrolase_6<br>Hydrolase_4<br>Abhydrolase_1 | LinE Homolog<br>LinH (Fusion of LinE/LinG)<br>Alpha/beta hydrolase family<br>Serine aminopeptidase, S33<br>alpha/beta hydrolase fold | 6.60E-82<br>3.60E-65<br>5.80E-16<br>1.40E-11<br>1.10E-07 |
| WP_127841863.1 | 1163162 | 1162316 | - | 281 | LinH<br>LinG | LinH<br>LinG | LinH (Fusion of LinE/LinG)<br>LinG Homolog | 8.80E-95<br>2.70E-90 |
| WP_127841864.1 | 1163721 | 1164102 | + | 126 | PF09391 | DUF2000 | Protein of unknown function (DUF2000) | 3.50E-21 |
| WP_139738604.1 | 1164697 | 1164886 | + | 62 | LinA | LinA | LinA (linaridin precursor) | 2.10E-10 |
| WP_127841865.1 | 1165762 | 1166659 | + | 298 | LinE<br>LinH<br>PF12697<br>TIGR02427 | LinE<br>LinH<br>Abhydrolase_6<br>TIGR02427 | LinE Homolog<br>LinH (Fusion of LinE/LinG)<br>Alpha/beta hydrolase family<br>protocat_pcaD: 3-oxoadipate enol-lactonase | 8.50E-68<br>7.90E-41<br>2.50E-06<br>5.80E-06 |
| WP_127841866.1 | 1166753 | 1167227 | + | 157 | LinL | LinL | LinL | 1.10E-36 |
| WP_164860077.1 | 1167223 | 1167874 | + | 216 | PF02441<br>TIGR00521<br>TIGR02113 | Flavoprotein<br>TIGR00521<br>TIGR02113 | Flavoprotein<br>coaBC_dfp: phosphopantothenoylcysteine decarboxylase / phosphopantothenate--cysteine ligase<br>coaC_strep: phosphopantothenoylcysteine decarboxylase | 1.10E-12<br>8.40E-11<br>3.60E-07 |
| WP_164860078.1 | 1168079 | 1168688 | + | 202 | TIGR01189<br>TIGR02857<br>TIGR01277<br>TIGR01188<br>TIGR02868 | TIGR01189<br>TIGR02857<br>TIGR01277<br>TIGR01188<br>TIGR02868 | ccmA: heme ABC exporter, ATP-binding protein CcmA<br>CydD: thiol reductant ABC exporter, CydD subunit<br>thiQ: thiamine ABC transporter, ATP-binding protein<br>drrA: daunorubicin resistance ABC transporter, ATP-binding protein<br>CydC: thiol reductant ABC exporter, CydC subunit | 6.80E-30<br>9.10E-23<br>1.00E-22<br>1.10E-22<br>1.50E-21 |
| WP_127841869.1 | 1168684 | 1170250 | + | 521 | NO PFAM MATCH | - | - | - |
| WP_164860079.1 | 1170388 | 1170667 | + | 92 | PF09391 | DUF2000 | Protein of unknown function (DUF2000) | 3.70E-09 |
| WP_127841871.1 | 1171240 | 1170709 | - | 176 | PF09335 | SNARE_assoc | SNARE associated Golgi protein | 2.00E-10 |
| WP_127841872.1 | 1171616 | 1174562 | + | 981 | PF00933<br>PF01915<br>PF14310 | Glyco_hydro_3<br>Glyco_hydro_3_C<br>Fn3-like | Glycosyl hydrolase family 3 N terminal domain<br>Glycosyl hydrolase family 3 C-terminal domain<br>Fibronectin type III-like domain | 6.60E-34<br>1.50E-28<br>1.20E-16 |
| WP_127841873.1 | 1174718 | 1177121 | + | 800 | PF00933<br>PF01915<br>PF14310 | Glyco_hydro_3<br>Glyco_hydro_3_C<br>Fn3-like | Glycosyl hydrolase family 3 N terminal domain<br>Glycosyl hydrolase family 3 C-terminal domain<br>Fibronectin type III-like domain | 1.00E-48<br>5.40E-45<br>1.30E-20 |

| index | leader | core | start | end | dir | score |
|---|---|---|---|---|---|---|
| 1 | VAETQHRTHGDIQAPGA | LTPQAVAPGACPAATFLVGDRVAAQPYLGQR | 1159299 | 1159152 | - | 8 |
| 2 | VACGGNSAPNPRRHTGAWSTDTAGSCSRRLSCCYLSR | WRQSCGSTLPRTTMTAKTTSGLPRNAKKKPCALPSEKA | 1159306 | 1159078 | - | 7 |
| 3 | MNELLDKVNSDTVVIVDRTQVPVL | ATPAAFLGGLAAGAKAAGALVGAAAGGAAVGAATR | 1159500 | 1159320 | - | 30 |
| 4 | MQILLDKVNSDPTLKGSLRLRASRVGTPVL | ATPAALAWAAGAAVGAGAVSGAAYGAYRVAAR | 1159715 | 1159526 | - | 30 |
| 5 | MFDHMANPGRAASSIVLA | VTAGVVVALLFVSSPDD | 1159886 | 1159778 | - | 6 |
| 6 | MYAGPVAYLLMFDHMANPGRAASSIVLA | VTAGVVVALLFVSSPDD | 1159916 | 1159778 | - | 6 |
| 7 | MSRWYAFWLTTMYAGPVAYLLMFDHMANPGRAASSIVLA | VTAGVVVALLFVSSPDD | 1159949 | 1159778 | - | 7 |
| 8 | VQMSRWYAFWLTTMYAGPVAYLLMFDHMANPGRAASSIVLA | VTAGVVVALLFVSSPDD | 1159955 | 1159778 | - | 7 |
| 9 | MRVTQCFRASQQPLK | DVRTKHLACLLGGADV | 1160037 | 1159941 | - | 6 |
| 10 | VGPFLLLAESLEPCAAGAGRALGALRCG | RGQRGCGAPPGWARSVCSDRHRSTGRRAL | 1163379 | 1163553 | + | 6 |
| 11 | VIITPAALCSCGGRSTPTSPSPAAHHTPSALVRNGEHPELDQPRQ | RTVPGSPPEGGKAPRTSSHSRTASGLW | 1163557 | 1163338 | - | 7 |
| 12 | MSDLGIAPTVEQLEKVANREFSGISEGFVNSHNLPTM | TTPVTPATPGALEVVVTGSTICLGC | 1164697 | 1164886 | + | 31 |
| 13 | VSSCGGGRTRINQRDRKLQVPLWQR | RGRPDVRRNMRLQGCYWPHTRSQPVR | 1165198 | 1165354 | + | 6 |

| | | | | | | |
|---|---|---|---|---|---|---|
| 14 | LRVNSPEVDVIGQAVIWYEASNSPV | AAYYGGHLVCPVVAIGAPVTFDHAG | 1165382 | 1165229 | - | 9 |
| 15 | LIGQGNAVVAPQLHLIRTDLFQGPSE | ERCQKARNSKLTSLSQESSRPEMFAR | 1165530 | 1165371 | - | 7 |
| 16 | MLAPLIGQGNAVVAPQLHLIRTDLFQGP | SEERCQKARNSKLTSLSQESSRPEMFAR | 1165542 | 1165371 | - | 7 |
| 17 | LVTVREAWAQSSRFPCGRASPQDQPVDSRTCSAPRPRSRHRARPAR | LTAPNRHRTTRCRVVL | 1171470 | 1171281 | - | 10 |
| 18 | VGRERRRAPGTVGHRQGGLGTVKPVPLRQSQPAGPTCRFQDLFCA | QAPLTAPGPPRPAHGTKSS | 1171504 | 1171309 | - | 9 |
| 19 | VGRSTEPPLVGRERRRAPGTVGHRQ...TVKPVPLRQSQPAGPTCRFQDLFCA | QAPLTAPGPPRPAHGTKSS | 1171531 | 1171309 | - | 8 |
| 20 | VHEAEWAGARSRRWWDGSAGGRLAP...QDQPVDSRTCSAPRPRSRHRARPAR | LTAPNRHRTTRCRVVL | 1171545 | 1171281 | - | 9 |
| 21 | VCPADAWSVHEAEWAGARSRRWWDG...QDQPVDSRTCSAPRPRSRHRARPAR | LTAPNRHRTTRCRVVL | 1171569 | 1171281 | - | 9 |
| 22 | VAVCPADAWSVHEAEWAGARSRRWW...QDQPVDSRTCSAPRPRSRHRARPAR | LTAPNRHRTTRCRVVL | 1171575 | 1171281 | - | 9 |
| 23 | VSVAVCPADAWSVHEAEWAGARSRR...QDQPVDSRTCSAPRPRSRHRARPAR | LTAPNRHRTTRCRVVL | 1171581 | 1171281 | - | 9 |
| 24 | MNWARVSFLSSKRDHACLIPGTGSPCTVCRVYFAV | STLSVWWSACCCLAAPRRLGSRAA | 1177297 | 1177117 | - | 8 |
| 25 | MRKTMNWARVSFLSSKRDHACLIPGTGSPCTVCRVYFAV | STLSVWWSACCCLAAPRRLGSRAA | 1177309 | 1177117 | - | 8 |
| 26 | LDRMRKTMNWARVSFLSSKRDHACLIPGTGSPCTVCRVYFAV | STLSVWWSACCCLAAPRRLGSRAA | 1177318 | 1177117 | - | 8 |

### Results for WP_139738603.1 [Actinomyces sp. 340] back to top

previous - next

#### Architecture

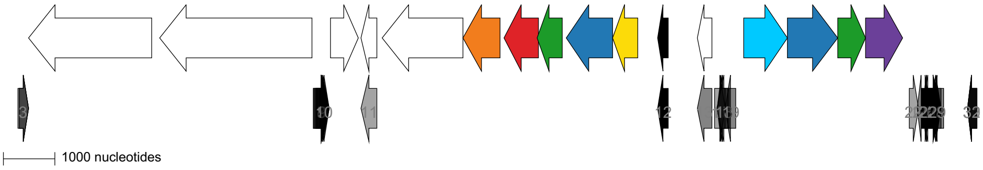

Link to nucleotide sequence

| Accession | start | end | direction | length (aa) | Pfam/HMM | name | description | E-value |
|---|---|---|---|---|---|---|---|---|
| WP_139738600.1 | 102104 | 99701 | - | 800 | PF00933<br>PF01915<br>PF14310 | Glyco_hydro_3<br>Glyco_hydro_3_C<br>Fn3-like | Glycosyl hydrolase family 3 N terminal domain<br>Glycosyl hydrolase family 3 C-terminal domain<br>Fibronectin type III-like domain | 1.00E-48<br>5.40E-45<br>6.10E-21 |
| WP_127841872.1 | 105206 | 102260 | - | 981 | PF00933<br>PF01915<br>PF14310 | Glyco_hydro_3<br>Glyco_hydro_3_C<br>Fn3-like | Glycosyl hydrolase family 3 N terminal domain<br>Glycosyl hydrolase family 3 C-terminal domain<br>Fibronectin type III-like domain | 6.60E-34<br>1.50E-28<br>1.20E-16 |
| WP_139738601.1 | 105582 | 106113 | + | 176 | PF09335 | SNARE_assoc | SNARE associated Golgi protein | 2.60E-10 |
| WP_127841870.1 | 106482 | 106155 | - | 108 | PF09391 | DUF2000 | Protein of unknown function (DUF2000) | 5.40E-09 |
| WP_139738602.1 | 108138 | 106572 | - | 521 | NO PFAM MATCH | - | - | - |
| WP_127841868.1 | 108875 | 108134 | - | 246 | TIGR01189<br>TIGR02857<br>TIGR02868<br>TIGR01188<br>TIGR01277 | TIGR01189<br>TIGR02857<br>TIGR02868<br>TIGR01188<br>TIGR01277 | ccmA: heme ABC exporter, ATP-binding protein CcmA<br>CydD: thiol reductant ABC exporter, CydD subunit<br>CydC: thiol reductant ABC exporter, CydC subunit<br>drrA: daunorubicin resistance ABC transporter, ATP-binding protein<br>thiQ: thiamine ABC transporter, ATP-binding protein | 1.10E-31<br>1.40E-26<br>4.00E-26<br>3.70E-24<br>5.80E-24 |
| WP_127841867.1 | 109614 | 108948 | - | 221 | PF02441<br>TIGR00521<br>TIGR02113 | Flavoprotein<br>TIGR00521<br>TIGR02113 | Flavoprotein<br>coaBC_dfp: phosphopantothenoylcysteine decarboxylase / phosphopantothenate--cysteine ligase<br>coaC_strep: phosphopantothenoylcysteine decarboxylase | 1.10E-12<br>8.90E-11<br>3.80E-07 |
| WP_127841866.1 | 110069 | 109595 | - | 157 | LinL | LinL | LinL | 1.10E-36 |
| WP_139738603.1 | 111060 | 110163 | - | 298 | LinE<br>LinH<br>PF12697<br>TIGR02427 | LinE<br>LinH<br>Abhydrolase_6<br>TIGR02427 | LinE Homolog<br>LinH (Fusion of LinE/LinG)<br>Alpha/beta hydrolase family<br>protocat_pcaD: 3-oxoadipate enol-lactonase | 2.20E-68<br>4.70E-41<br>2.10E-06<br>3.60E-06 |
| WP_127843254.1 | 111536 | 111056 | - | 159 | LinG<br>LinH<br>TIGR04033<br>PF06762 | LinG<br>LinH<br>TIGR04033<br>LMF1 | LinG Homolog<br>LinH (Fusion of LinE/LinG)<br>export_SdpB: antimicrobial peptide system protein, SdpB family<br>Lipase maturation factor | 7.10E-56<br>2.20E-52<br>2.10E-06<br>4.50E-04 |
| WP_139738604.1 | 112125 | 111936 | - | 62 | LinA | LinA | LinA (linaridin precursor) | 2.10E-10 |
| WP_139738605.1 | 112988 | 112703 | - | 94 | PF09391 | DUF2000 | Protein of unknown function (DUF2000) | 1.70E-11 |
| WP_139738606.1 | 113609 | 114455 | + | 281 | LinH<br>LinG | LinH<br>LinG | LinH (Fusion of LinE/LinG)<br>LinG Homolog | 6.10E-95<br>4.60E-90 |
| WP_139738607.1 | 114451 | 115435 | + | 327 | LinE<br>LinH<br>PF12697<br>PF12146<br>PF00561 | LinE<br>LinH<br>Abhydrolase_6<br>Hydrolase_4<br>Abhydrolase_1 | LinE Homolog<br>LinH (Fusion of LinE/LinG)<br>Alpha/beta hydrolase family<br>Serine aminopeptidase, S33<br>alpha/beta hydrolase fold | 1.70E-82<br>2.50E-65<br>3.50E-17<br>2.40E-11<br>3.70E-06 |
| WP_139738608.1 | 115431 | 115971 | + | 179 | LinL | LinL | LinL | 9.40E-57 |
| WP_139738609.1 | 115967 | 116687 | + | 239 | PF13649<br>PF08241<br>PF08242<br>PF13489<br>TIGR01934 | Methyltransf_25<br>Methyltransf_11<br>Methyltransf_12<br>Methyltransf_23<br>TIGR01934 | Methyltransferase domain<br>Methyltransferase domain<br>Methyltransferase domain<br>Methyltransferase domain<br>MenG_MenH_UbiE: ubiquinone/menaquinone biosynthesis methyltransferase | 3.00E-12<br>1.30E-11<br>8.70E-09<br>2.30E-07<br>1.00E-06 |

| index | leader | core | start | end | dir | score |
|---|---|---|---|---|---|---|
| 1 | LDRMRKTMNWARVSFLSSKRDHACLIPGTGSPCTVCRVYFAV | STLSVWWSACCCLAAPRRLGSRAA | 99504 | 99705 | + | 6 |
| 2 | MRKTMNWARVSFLSSKRDHACLIPGTGSPCTVCRVYFAV | STLSVWWSACCCLAAPRRLGSRAA | 99513 | 99705 | + | 6 |
| 3 | MNWARVSFLSSKRDHACLIPGTGSPCTVCRVYFAV | STLSVWWSACCCLAAPRRLGSRAA | 99525 | 99705 | + | 6 |
| 4 | VSVAVCPADAWSVHEAEWAGARSRR...QDQPVDSRTCSAPRPRSRHRARPAR | LTAPNRHRTTRCRVVL | 105241 | 105541 | + | 7 |
| 5 | VAVCPADAWSVHEAEWAGARSRRWW...QDQPVDSRTCSAPRPRSRHRARPAR | LTAPNRHRTTRCRVVL | 105247 | 105541 | + | 7 |
| 6 | VCPADAWSVHEAEWAGARSRRWWDG...QDQPVDSRTCSAPRPRSRHRARPAR | LTAPNRHRTTRCRVVL | 105253 | 105541 | + | 7 |
| 7 | VHEAEWAGARSRRWWDGSAGGRLAP...QDQPVDSRTCSAPRPRSRHRARPAR | LTAPNRHRTTRCRVVL | 105277 | 105541 | + | 7 |
| 8 | VGRSTEPPLVGRERRRAPGTVGHRQ...TVKPVPLRQSQPAGPTCRFQDLFCA | QAPLTAPGPPRPAHGTKSS | 105291 | 105513 | + | 6 |
| 9 | VGRERRRAPGTVGHRQGGLGTVKPVPLRQSQPAGPTCRFQDLFCA | QAPLTAPGPPRPAHGTKSS | 105318 | 105513 | + | 7 |
| 10 | LVTVREAWAQSSRFPCGRASPQDQPVDSRTCSAPRPRSRHRARPAR | LTAPNRHRTTRCRVVL | 105352 | 105541 | + | 8 |
| 11 | MLLTPCDDRLYLLGLWLTLGVGAWI...VSERWVLGTSITLEHPVRWAWADCY | ARLAQGCRTYEEYVSRLGSALTRDL...LGLRGPRRQVTSLTGDLPLYPAKDA | 106482 | 106155 | - | 6 |
| 12 | MSDLGIAPTVEQLEKVANREFSGISEGFVNSHNLPTM | TTPVTPATPGALEVVVTGSTICLGC | 112125 | 111936 | - | 33 |
| 13 | MITSACPQIMGPSV | RTADSELPGVVLTPLPVLAGDSGLL...EYVSRLGSALTRDLELAAHVTWSWL | 112988 | 112703 | - | 7 |
| 14 | VPVNHYHRLLLVGAHEYVPFSQEFGSQPWDPDRARASV | RPPEPGRSVQARPQPGQAGRAAASRTCAQTAE | 113032 | 113245 | + | 6 |
| 15 | MITQRSARRSVKPLP | CPPVLAGVLPARIARAPGDGQKHGRDPDPTAASRTPD | 113249 | 113090 | - | 6 |
| 16 | VLRPPQEHRAAGVMITQRSARRSVKPLP | CPPVLAGVLPARIARAPGDGQKHGRDPDPTAASRTPD | 113288 | 113090 | - | 6 |
| 17 | VWCAAGLGEVGVLRPPQEHRAAGVMITQRSARRSVKPLP | CPPVLAGVLPARIARAPGDGQKHGRDPDPTAASRTPD | 113321 | 113090 | - | 7 |
| 18 | VRCWGRTRAEGVWCAAGLGEVGVLRPPQEHRAAGVMITQRSARRSVKPLP | CPPVLAGVLPARIARAPGDGQKHGRDPDPTAASRTPD | 113354 | 113090 | - | 7 |
| 19 | LSAAPTRGRSLSCCGRRFVGPFLLLAESLKPCA | AGAGRGQRGCGAPPGWARSVCSDRHRSTGRRAL | 113446 | 113245 | - | 7 |
| 20 | VSRWFTFCLTAMFTGVITYLRIFDDLDN | PARAMSSASLAAMSGLVVALFFVSPSDD | 116816 | 116987 | + | 6 |
| 21 | MQILLDKVNSDPTLKGSLRLRASRVGTPVL | ATPAALAWAAGAAVGAGAVSGAAYGAYRVAAR | 117050 | 117239 | + | 18 |
| 22 | VASTGVPTLLARSRSDPFSVGSELTL | SSSICIRSLLSVPRGTICPVRTFLLEV | 117145 | 116983 | - | 7 |
| 23 | VGGWSSRRSGRCEWGRIRCLPGRGA...LMNELLDKVNSDTVVVVDRAQVPVL | ATPAAFYAGVVAGAKAAAITVAAASAGASVAR | 117160 | 117436 | + | 20 |
| 24 | LILSAPRPGRHRMRPHSQRPLLRLLQPP | TQGRRVWRVPAYLPCWPAVEAIPSVWDPS | 117246 | 117072 | - | 7 |
| 25 | MNELLDKVNSDTVVVVDRAQVPVL | ATPAAFYAGVVAGAKAAAITVAAASAGASVAR | 117265 | 117436 | + | 30 |
| 26 | MSPPSVLLILSAPRPGRHRMRPHSQRPLLRLL | QPPTQGRRVWRVPAYLPCWPAVEAIPSVWDPS | 117267 | 117072 | - | 7 |
| 27 | LSSSSFMSPPSVLLILSAPRPGRHRMRPHSQRPLL | RLLQPPTQGRRVWRVPAYLPCWPAVEAIPSVWDPS | 117285 | 117072 | - | 6 |
| 28 | LGLAGNRRTSAGCCNRDGRSFGACNHSRIKSSRCGEY | GNLCAVHHDHGVGVDLVKQLVHESSFCSVDPQRAATR | 117442 | 117217 | - | 7 |
| 29 | VYSRGFDDAGCCQRHPLGLGLAGNRRTSAGCCNRDGRSFGACNHSR | IKSSRCGEYGNLCAVHHDHGVGVDLVKQLVHESSFCSVDPQRAATR | 117496 | 117217 | - | 7 |
| 30 | MLAGSVGWWEPVLSLVITLAV | FTPLLVMLATAVYRRSLMQTQGRLTMRQALRLPAD | 118118 | 117947 | - | 8 |
| 31 | MPTRMLAGSVGWWEPVLSLVITLAV | FTPLLVMLATAVYRRSLMQTQGRLTMRQALRLPAD | 118130 | 117947 | - | 8 |
| 32 | VAAMPTRMLAGSVGWWEPVLSLVITLAV | FTPLLVMLATAVYRRSLMQTQGRLTMRQALRLPAD | 118139 | 117947 | - | 8 |

### Results for WP_139738607.1 [Actinomyces sp. 340] back to top

previous - next

#### Architecture

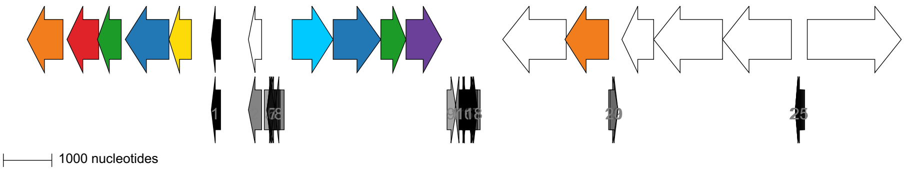

Link to nucleotide sequence

| Accession | start | end | direction | length (aa) | Pfam/HMM | name | description | E-value |
|---|---|---|---|---|---|---|---|---|
| WP_127841868.1 | 108875 | 108134 | - | 246 | TIGR01189<br>TIGR02857<br>TIGR02868<br>TIGR01188<br>TIGR01277 | TIGR01189<br>TIGR02857<br>TIGR02868<br>TIGR01188<br>TIGR01277 | ccmA: heme ABC exporter, ATP-binding protein CcmA<br>CydD: thiol reductant ABC exporter, CydD subunit<br>CydC: thiol reductant ABC exporter, CydC subunit<br>drrA: daunorubicin resistance ABC transporter, ATP-binding protein<br>thiQ: thiamine ABC transporter, ATP-binding protein | 1.10E-31<br>1.40E-26<br>4.00E-26<br>3.70E-24<br>5.80E-24 |
| WP_127841867.1 | 109614 | 108948 | - | 221 | PF02441<br>TIGR00521<br>TIGR02113 | Flavoprotein<br>TIGR00521<br>TIGR02113 | Flavoprotein<br>coaBC_dfp: phosphopantothenoylcysteine decarboxylase / phosphopantothenate--cysteine ligase<br>coaC_strep: phosphopantothenoylcysteine decarboxylase | 1.10E-12<br>8.90E-11<br>3.80E-07 |

| | | | | | | | | |
|---|---|---|---|---|---|---|---|---|
| WP_127841866.1 | 110069 | 109595 | - | 157 | LinL | LinL | LinL | 1.10E-36 |
| WP_139738603.1 | 111060 | 110163 | - | 298 | LinE<br>LinH<br>PF12697<br>TIGR02427 | LinE<br>LinH<br>Abhydrolase_6<br>TIGR02427 | LinE Homolog<br>LinH (Fusion of LinE/LinG)<br>Alpha/beta hydrolase family<br>protocat_pcaD: 3-oxoadipate enol-lactonase | 2.20E-68<br>4.70E-41<br>2.10E-06<br>3.60E-06 |
| WP_127843254.1 | 111536 | 111056 | - | 159 | LinG<br>LinH<br>TIGR04033<br>PF06762 | LinG<br>LinH<br>TIGR04033<br>LMF1 | LinG Homolog<br>LinH (Fusion of LinE/LinG)<br>export_SdpB: antimicrobial peptide system protein, SdpB family<br>Lipase maturation factor | 7.10E-56<br>2.20E-52<br>2.10E-06<br>4.50E-04 |
| WP_139738604.1 | 112125 | 111936 | - | 62 | LinA | LinA | LinA (linaridin precursor) | 2.10E-10 |
| WP_139738605.1 | 112988 | 112703 | - | 94 | PF09391 | DUF2000 | Protein of unknown function (DUF2000) | 1.70E-11 |
| WP_139738606.1 | 113609 | 114455 | + | 281 | LinH<br>LinG | LinH<br>LinG | LinH (Fusion of LinE/LinG)<br>LinG Homolog | 6.10E-95<br>4.60E-90 |
| WP_139738607.1 | 114451 | 115435 | + | 327 | LinE<br>LinH<br>PF12697<br>PF12146<br>PF00561 | LinE<br>LinH<br>Abhydrolase_6<br>Hydrolase_4<br>Abhydrolase_1 | LinE Homolog<br>LinH (Fusion of LinE/LinG)<br>Alpha/beta hydrolase family<br>Serine aminopeptidase, S33<br>alpha/beta hydrolase fold | 1.70E-82<br>2.50E-65<br>3.50E-17<br>2.40E-11<br>3.70E-06 |
| WP_139738608.1 | 115431 | 115971 | + | 179 | LinL | LinL | LinL | 9.40E-57 |
| WP_139738609.1 | 115967 | 116687 | + | 239 | PF13649<br>PF08241<br>PF08242<br>PF13489<br>TIGR01934 | Methyltransf_25<br>Methyltransf_11<br>Methyltransf_12<br>Methyltransf_23<br>TIGR01934 | Methyltransferase domain<br>Methyltransferase domain<br>Methyltransferase domain<br>Methyltransferase domain<br>MenG_MenH_UbiE: ubiquinone/menaquinone biosynthesis methyltransferase | 3.00E-12<br>1.30E-11<br>8.70E-09<br>2.30E-07<br>1.00E-06 |
| WP_127841859.1 | 119255 | 117947 | - | 435 | PF12698<br>PF12679 | ABC2_membrane_3<br>ABC2_membrane_2 | ABC-2 family transporter protein<br>ABC-2 family transporter protein | 6.40E-28<br>7.10E-07 |
| WP_139738610.1 | 120132 | 119229 | - | 300 | TIGR01188<br>TIGR03522<br>TIGR04406<br>TIGR03864<br>TIGR03740 | TIGR01188<br>TIGR03522<br>TIGR04406<br>TIGR03864<br>TIGR03740 | drrA: daunorubicin resistance ABC transporter, ATP-binding protein<br>GldA_ABC_ATP: gliding motility-associated ABC transporter ATP-binding subunit GldA<br>LPS_export_lptB: LPS export ABC transporter ATP-binding protein<br>PQQ_ABC_ATP: ABC transporter, ATP-binding subunit, PQQ-dependent alcohol dehydrogenase system<br>galliderm_ABC: lantibiotic protection ABC transporter, ATP-binding subunit | 7.10E-50<br>4.60E-44<br>8.30E-44<br>5.40E-40<br>1.30E-38 |
| WP_139738611.1 | 121078 | 120406 | - | 223 | PF00072<br>PF00196<br>TIGR02875<br>TIGR03020<br>TIGR01387 | Response_reg<br>GerE<br>TIGR02875<br>TIGR03020<br>TIGR01387 | Response regulator receiver domain<br>Bacterial regulatory proteins, luxR family<br>spore_0_A: sporulation transcription factor Spo0A<br>EpsA: transcriptional regulator EpsA<br>cztR_silR_copR: heavy metal response regulator | 3.00E-25<br>5.80E-18<br>8.50E-18<br>1.10E-12<br>6.20E-12 |
| WP_139738612.1 | 122490 | 121074 | - | 471 | PF07730 | HisKA_3 | Histidine kinase | 2.70E-10 |
| WP_127841855.1 | 123914 | 122486 | - | 475 | TIGR02734<br>PF13450<br>TIGR03467<br>PF01593<br>TIGR01317 | TIGR02734<br>NAD_binding_8<br>TIGR03467<br>Amino_oxidase<br>TIGR01317 | crtI_fam: phytoene desaturase<br>NAD(P)-binding Rossmann-like domain<br>HpnE: squalene-associated FAD-dependent desaturase<br>Flavin containing amine oxidoreductase<br>GOGAT_sm_gam: glutamate synthase, NADH/NADPH, small subunit | 4.50E-10<br>5.10E-10<br>1.40E-07<br>3.20E-07<br>5.30E-07 |
| WP_139738613.1 | 124256 | 126179 | + | 640 | TIGR00565<br>TIGR00564<br>TIGR01820<br>PF00425<br>TIGR01824 | TIGR00565<br>TIGR00564<br>TIGR01820<br>Chorismate_bind<br>TIGR01824 | trpE_proteo: anthranilate synthase component I<br>trpE_most: anthranilate synthase component I<br>TrpE-arch: anthranilate synthase component I<br>chorismate binding enzyme<br>PabB-clade2: aminodeoxychorismate synthase, component I | 3.00E-149<br>1.10E-110<br>2.90E-96<br>7.40E-83<br>2.30E-82 |

| index | leader | core | start | end | dir | score |
|---|---|---|---|---|---|---|
| 1 | MSDLGIAPTVEQLEKVANREFSGISEGFVNSHNLPTM | TTPVTPATPGALEVVVTGSTICLGC | 112125 | 111936 | - | 33 |
| 2 | MITSACPQIMGPSV | RTADSELPGVVLTPLPVLAGDSGLL...EYVSRLGSALTRDLELAAHVTWSWL | 112988 | 112703 | - | 7 |
| 3 | VPVNHYHRLLLVGAHEYVPFSQEFGSQPWDPDRARASV | RPPEPGRSVQARPQPGQAGRAAASRTCAQTAE | 113032 | 113245 | + | 6 |
| 4 | MITQRSARRSVKPLP | CPPVLAGVLPARIARAPGDGQKHGRDPDPTAASRTPD | 113249 | 113090 | - | 6 |
| 5 | VLRPPQEHRAAGVMITQRSARRSVKPLP | CPPVLAGVLPARIARAPGDGQKHGRDPDPTAASRTPD | 113288 | 113090 | - | 6 |
| 6 | VWCAAGLGEVGVLRPPQEHRAAGVMITQRSARRSVKPLP | CPPVLAGVLPARIARAPGDGQKHGRDPDPTAASRTPD | 113321 | 113090 | - | 7 |
| 7 | VRCWGRTRAEGVWCAAGLGEVGVLRPPQEHRAAGVMITQRSARRSVKPLP | CPPVLAGVLPARIARAPGDGQKHGRDPDPTAASRTPD | 113354 | 113090 | - | 7 |
| 8 | LSAAPTRGRSLSCCGRRFVGPFLLLAESLKPCA | AGAGRGQRGCGAPPGWARSVCSDRHRSTGRRAL | 113446 | 113245 | - | 7 |
| 9 | VSRWFTFCLTAMFTGVITYLRIFDDLDN | PARAMSSASLAAMSGLVVALFFVSPSDD | 116816 | 116987 | + | 6 |
| 10 | MQILLDKVNSDPTLKGSLRLRASRVGTPVL | ATPAALAWAAGAAVGAGAVSGAAYGAYRVAAR | 117050 | 117239 | + | 18 |
| 11 | VASTGVPTLLARSRSDPFSVGSELTL | SSSICIRSLLSVPRGTICPVRTFLLEV | 117145 | 116983 | - | 7 |
| 12 | VGGWSSRRSGRCEWGRIRCLPGRGA...LMNELLDKVNSDTVVVVDRAQVPVL | ATPAAFYAGVVAGAKAAAITVAAASAGASVAR | 117160 | 117436 | + | 20 |
| 13 | LILSAPRPGRHRMRPHSQRPLLRLLQPP | TQGRRVWRVPAYLPCWPAVEAIPSVWDPS | 117246 | 117072 | - | 7 |
| 14 | MNELLDKVNSDTVVVVDRAQVPVL | ATPAAFYAGVVAGAKAAAITVAAASAGASVAR | 117265 | 117436 | + | 30 |
| 15 | MSPPSVLLILSAPRPGRHRMRPHSQRPLLRLL | QPPTQGRRVWRVPAYLPCWPAVEAIPSVWDPS | 117267 | 117072 | - | 7 |
| 16 | LSSSSFMSPPSVLLILSAPRPGRHRMRPHSQRPLL | RLLQPPTQGRRVWRVPAYLPCWPAVEAIPSVWDPS | 117285 | 117072 | - | 6 |
| 17 | LGLAGNRRTSAGCCNRDGRSFGACNHSRIKSSRCGEY | GNLCAVHHDHGVGVDLVKQLVHESSFCSVDPQRAATR | 117442 | 117217 | - | 7 |
| 18 | VYSRGFDDAGCCQRHPLGLGLAGNRRTSAGCCNRDGRSFGACNHSR | IKSSRCGEYGNLCAVHHDHGVGVDLVKQLVHESSFCSVDPQRAATR | 117496 | 117217 | - | 7 |
| 19 | LLSACHCLSVSSCLLADYQARSNQPAPDPARS | RNLPLPRKESSQPTTPPHEGADPAAHLAGLVPT | 120142 | 120340 | + | 6 |

| | | | | | | |
|---|---|---|---|---|---|---|
| 20 | VSSCLLADYQARSNQPAPDPARSRNLPL | PRKESSQPTTPPHEGADPAAHLAGLVPT | 120169 | 120340 | + | 6 |
| 21 | VVMAEPYDGRGRAHK | RGATGVVASPAQRQGS | 124099 | 124003 | - | 7 |
| 22 | VRGVVMAEPYDGRGRAH | KRGATGVVASPAQRQGS | 124108 | 124003 | - | 6 |
| 23 | VARLLGSHHHWLVRGVVMAEPYD | GRGRAHKRGATGVVASPAQRQGS | 124144 | 124003 | - | 6 |
| 24 | VPTARSQRLRVARLLGSHHHWLVRGVVM | AEPYDGRGRAHKRGATGVVASPAQRQGS | 124174 | 124003 | - | 7 |
| 25 | VASSCGAVPTARSQRLRVARLLGSHHHWLVR | GVVMAEPYDGRGRAHKRGATGVVASPAQRQGS | 124195 | 124003 | - | 9 |

### Results for SFE63453.1 [Actinopolyspora alba] back to top

previous - next

#### Architecture

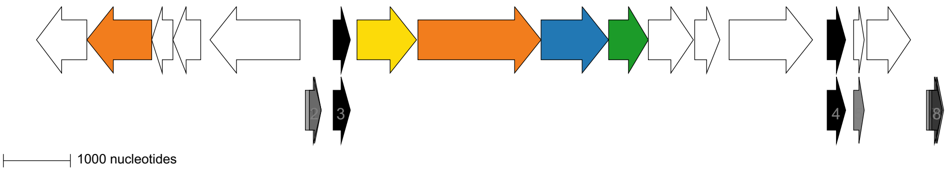

Link to nucleotide sequence

| Accession | start | end | direction | length (aa) | Pfam/HMM | name | description | E-value |
|---|---|---|---|---|---|---|---|---|
| SFE63304.1 | 25909 | 25153 | - | 251 | PF01061<br>PF12698<br>TIGR01247<br>PF12679<br>TIGR03861 | ABC2_membrane<br>ABC2_membrane_3<br>TIGR01247<br>ABC2_membrane_2<br>TIGR03861 | ABC-2 type transporter<br>ABC-2 family transporter protein<br>drrB: daunorubicin resistance ABC transporter membrane protein<br>ABC-2 family transporter protein<br>phenyl_ABC_PedC: alcohol ABC transporter, permease protein | 1.40E-15<br>1.80E-13<br>8.70E-12<br>1.20E-11<br>1.20E-05 |
| SFE63323.1 | 26875 | 25915 | - | 319 | TIGR01188<br>TIGR01288<br>TIGR03522<br>TIGR03864<br>TIGR04406 | TIGR01188<br>TIGR01288<br>TIGR03522<br>TIGR03864<br>TIGR04406 | drrA: daunorubicin resistance ABC transporter, ATP-binding protein<br>nodI: nodulation ABC transporter NodI<br>GldA_ABC_ATP: gliding motility-associated ABC transporter ATP-binding subunit GldA<br>PQQ_ABC_ATP: ABC transporter, ATP-binding subunit, PQQ-dependent alcohol dehydrogenase system<br>LPS_export_lptB: LPS export ABC transporter ATP-binding protein | 1.80E-64<br>1.50E-59<br>1.00E-48<br>2.10E-48<br>9.80E-41 |
| SFE63343.1 | 27198 | 26871 | - | 108 | PF13601<br>PF12840<br>PF01022<br>PF03551<br>PF02082 | HTH_34<br>HTH_20<br>HTH_5<br>PadR<br>Rrf2 | Winged helix DNA-binding domain<br>Helix-turn-helix domain<br>Bacterial regulatory protein, arsR family<br>Transcriptional regulator PadR-like family<br>Transcriptional regulator | 3.30E-26<br>7.10E-07<br>1.30E-05<br>4.30E-05<br>2.20E-04 |
| SFE63361.1 | 27617 | 27194 | - | 140 | PF06738 | ThrE | Putative threonine/serine exporter | 3.30E-04 |
| SFE63380.1 | 29090 | 27752 | - | 445 | PF05193<br>PF00675<br>TIGR02110 | Peptidase_M16_C<br>Peptidase_M16<br>TIGR02110 | Peptidase M16 inactive domain<br>Insulinase (Peptidase family M16)<br>PQQ_syn_pqqF: coenzyme PQQ biosynthesis protein PqqF | 7.90E-38<br>3.00E-34<br>5.00E-20 |
| SFE63394.1 | 29586 | 29850 | + | 87 | LinA | LinA | LinA (linaridin precursor) | 4.50E-07 |
| SFE63415.1 | 29943 | 30837 | + | 297 | LinG<br>LinH<br>TIGR04033 | LinG<br>LinH<br>TIGR04033 | LinG Homolog<br>LinH (Fusion of LinE/LinG)<br>export_SdpB: antimicrobial peptide system protein, SdpB family | 9.80E-<br>104<br>1.90E-92<br>1.60E-05 |
| SFE63435.1 | 30842 | 32681 | + | 612 | TIGR02204<br>TIGR02203<br>TIGR00958<br>TIGR01846<br>TIGR03796 | TIGR02204<br>TIGR02203<br>TIGR00958<br>TIGR01846<br>TIGR03796 | MsbA_rel: ABC transporter, permease/ATP-binding protein<br>MsbA_lipidA: lipid A export permease/ATP-binding protein MsbA<br>3a01208: antigen peptide transporter 2<br>type_I_sec_HlyB: type I secretion system ATPase<br>NHLM_micro_ABC1: NHLM bacteriocin system ABC transporter, peptidase/ATP-binding protein | 1.90E-<br>115<br>1.60E-<br>104<br>4.10E-93<br>6.40E-89<br>2.00E-83 |
| SFE63453.1 | 32681 | 33707 | + | 341 | LinE<br>LinH<br>PF12697<br>PF00561<br>TIGR02427 | LinE<br>LinH<br>Abhydrolase_6<br>Abhydrolase_1<br>TIGR02427 | LinE Homolog<br>LinH (Fusion of LinE/LinG)<br>Alpha/beta hydrolase family<br>alpha/beta hydrolase fold<br>protocat_pcaD: 3-oxoadipate enol-lactonase | 5.70E-<br>105<br>2.00E-81<br>3.30E-16<br>1.50E-13<br>8.50E-13 |
| SFE63470.1 | 33706 | 34297 | + | 196 | LinL | LinL | LinL | 1.00E-67 |
| SFE63488.1 | 34293 | 34974 | + | 226 | PF00196<br>TIGR03020<br>TIGR03541<br>TIGR02937 | GerE<br>TIGR03020<br>TIGR03541<br>TIGR02937 | Bacterial regulatory proteins, luxR family<br>EpsA: transcriptional regulator EpsA<br>reg_near_HchA: LuxR family transcriptional regulatory, chaperone HchA-associated<br>sigma70-ECF: RNA polymerase sigma factor, sigma-70 family | 3.50E-19<br>2.40E-14<br>2.00E-09<br>2.90E-04 |
| SFE63503.1 | 34975 | 35368 | + | 130 | PF04134 | DUF393 | Protein of unknown function, DUF393 | 3.60E-14 |
| SFE63521.1 | 35492 | 36752 | + | 419 | TIGR02032<br>PF04820<br>PF01494<br>TIGR02023<br>TIGR01988 | TIGR02032<br>Trp_halogenase<br>FAD_binding_3<br>TIGR02023<br>TIGR01988 | GG-red-SF: geranylgeranyl reductase family<br>Tryptophan halogenase<br>FAD binding domain<br>BchP-ChlP: geranylgeranyl reductase<br>Ubi-OHases: ubiquinone biosynthesis hydroxylase, UbiH/UbiF/VisC/COQ6 family | 3.30E-39<br>1.20E-38<br>1.30E-23<br>1.30E-16<br>6.20E-14 |
| SFE63544.1 | 36961 | 37231 | + | 89 | LinA | LinA | LinA (linaridin precursor) | 8.00E-09 |
| SFE63557.1 | 37357 | 37525 | + | 55 | NO PFAM MATCH | - | - | - |
| SFE63575.1 | 37561 | 38203 | + | 213 | PF05719 | GPP34 | Golgi phosphoprotein 3 (GPP34) | 7.60E-26 |

| index | leader | core | start | end | dir | score |
|---|---|---|---|---|---|---|

| | | | | | | |
|---|---|---|---|---|---|---|
| 1 | MKHESARRRNQETTNGKQRLHLVDL...PENATRNPRQVLCEIARTRRVSAVE | STTTILTVMYDTMPCSLLTMRLSLPGER | 29166 | 29406 | + | 6 |
| 2 | VDLRPENATRNPRQVLCEIARTRRVSAVE | STTTILTVMYDTMPCSLLTMRLSLPGER | 29232 | 29406 | + | 6 |
| 3 | MNRIESVANVQLPDAHDGAVRSLDTTGSTAPVL | TAAAITAAAVVCFAAGVASDAVESG...AASTGQDPTGMSSGELLQRRTALLG | 29586 | 29850 | + | 17 |
| 4 | MNRIENAATQQVTASAGAIEPMDSVEGVSPVL | TAAVAAGAVAATAAAFAAGNMVTDY...KHDGSELSGMSVGDLVNSRREAMSH | 36961 | 37231 | + | 16 |
| 5 | MSQSNNNAVIGGGAVLGAVVGGFVGIF | LGNPAIAAAIGLAVGIALATAMRAERKN | 37357 | 37525 | + | 7 |
| 6 | VSRKARRKVPPGVYRAGLHGSGSTVEAVAFDRVEERGP | RGAVNRESPTIAVLGVTDADHFACESDLNAIAAVSSAAA | 38443 | 38677 | + | 6 |
| 7 | VYRAGLHGSGSTVEAVAFDRVEERGPRGAVNR | ESPTIAVLGVTDADHFACESDLNAIAAVSSAAA | 38479 | 38677 | + | 7 |
| 8 | LIASRNEVHAVLLTARVPPSRSLVS | RTPITSRANPTSTQLPPFPPLRLDLRHTAP | 38531 | 38699 | + | 6 |

### Results for SDK49455.1 [Actinopolyspora mzabensis] back to top

previous - next

#### Architecture

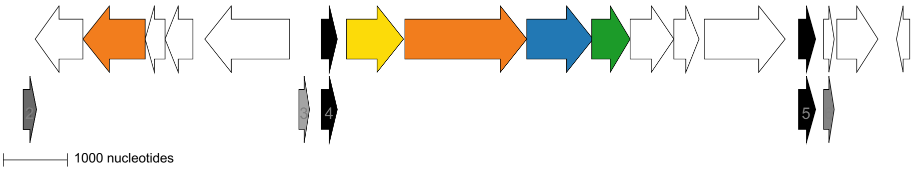

Link to nucleotide sequence

| Accession | start | end | direction | length (aa) | Pfam/HMM | name | description | E-value |
|---|---|---|---|---|---|---|---|---|
| SDK49274.1 | 263408 | 262652 | - | 251 | PF01061<br>PF12698<br>TIGR01247<br>PF12679<br>TIGR01291 | ABC2_membrane<br>ABC2_membrane_3<br>TIGR01247<br>ABC2_membrane_2<br>TIGR01291 | ABC-2 type transporter<br>ABC-2 family transporter protein<br>drrB: daunorubicin resistance ABC transporter membrane protein<br>ABC-2 family transporter protein<br>nodJ: ABC-2 type transporter, NodJ family | 2.00E-15<br>2.10E-14<br>4.60E-12<br>2.00E-10<br>1.80E-05 |
| SDK49294.1 | 264374 | 263414 | - | 319 | TIGR01188<br>TIGR01288<br>TIGR03522<br>TIGR03864<br>TIGR04406 | TIGR01188<br>TIGR01288<br>TIGR03522<br>TIGR03864<br>TIGR04406 | drrA: daunorubicin resistance ABC transporter, ATP-binding protein<br>nodI: nodulation ABC transporter NodI<br>GldA_ABC_ATP: gliding motility-associated ABC transporter ATP-binding subunit GldA<br>PQQ_ABC_ATP: ABC transporter, ATP-binding subunit, PQQ-dependent alcohol dehydrogenase system<br>LPS_export_lptB: LPS export ABC transporter ATP-binding protein | 5.10E-65<br>3.40E-59<br>3.70E-49<br>9.50E-49<br>5.90E-41 |
| SDK49313.1 | 264697 | 264370 | - | 108 | PF13601<br>PF12840<br>PF01022<br>PF03551<br>PF02082 | HTH_34<br>HTH_20<br>HTH_5<br>PadR<br>Rrf2 | Winged helix DNA-binding domain<br>Helix-turn-helix domain<br>Bacterial regulatory protein, arsR family<br>Transcriptional regulator PadR-like family<br>Transcriptional regulator | 2.40E-26<br>2.70E-06<br>1.90E-05<br>3.80E-05<br>2.30E-04 |
| SDK49340.1 | 265116 | 264693 | - | 140 | NO PFAM MATCH | - | - | - |
| SDK49364.1 | 266633 | 265301 | - | 443 | PF05193<br>PF00675<br>TIGR02110 | Peptidase_M16_C<br>Peptidase_M16<br>TIGR02110 | Peptidase M16 inactive domain<br>Insulinase (Peptidase family M16)<br>PQQ_syn_pqqF: coenzyme PQQ biosynthesis protein PqqF | 1.30E-35<br>4.70E-34<br>1.70E-21 |
| SDK49387.1 | 267128 | 267392 | + | 87 | LinA | LinA | LinA (linaridin precursor) | 4.10E-06 |
| SDK49412.1 | 267527 | 268424 | + | 298 | LinG<br>LinH<br>TIGR04033 | LinG<br>LinH<br>TIGR04033 | LinG Homolog<br>LinH (Fusion of LinE/LinG)<br>export_SdpB: antimicrobial peptide system protein, SdpB family | 2.20E-103<br>3.20E-92<br>1.80E-05 |
| SDK49434.1 | 268429 | 270343 | + | 637 | TIGR02204<br>TIGR02203<br>TIGR00958<br>TIGR01846<br>TIGR03796 | TIGR02204<br>TIGR02203<br>TIGR00958<br>TIGR01846<br>TIGR03796 | MsbA_rel: ABC transporter, permease/ATP-binding protein<br>MsbA_lipidA: lipid A export permease/ATP-binding protein MsbA<br>3a01208: antigen peptide transporter 2<br>type_I_sec_HlyB: type I secretion system ATPase<br>NHLM_micro_ABC1: NHLM bacteriocin system ABC transporter, peptidase/ATP-binding protein | 9.10E-113<br>8.20E-103<br>2.50E-93<br>7.50E-87<br>6.60E-84 |
| SDK49455.1 | 270343 | 271369 | + | 341 | LinE<br>LinH<br>PF12697<br>PF00561<br>TIGR02427 | LinE<br>LinH<br>Abhydrolase_6<br>Abhydrolase_1<br>TIGR02427 | LinE Homolog<br>LinH (Fusion of LinE/LinG)<br>Alpha/beta hydrolase family<br>alpha/beta hydrolase fold<br>protocat_pcaD: 3-oxoadipate enol-lactonase | 1.70E-104<br>5.60E-80<br>5.70E-16<br>8.00E-14<br>4.40E-13 |
| SDK49484.1 | 271368 | 271959 | + | 196 | LinL | LinL | LinL | 1.60E-66 |
| SDK49502.1 | 271955 | 272636 | + | 226 | PF00196<br>TIGR03020<br>TIGR03541<br>TIGR02937 | GerE<br>TIGR03020<br>TIGR03541<br>TIGR02937 | Bacterial regulatory proteins, luxR family<br>EpsA: transcriptional regulator EpsA<br>reg_near_HchA: LuxR family transcriptional regulatory, chaperone HchA-associated<br>sigma70-ECF: RNA polymerase sigma factor, sigma-70 family | 2.60E-19<br>2.30E-14<br>5.70E-09<br>3.00E-04 |
| SDK49525.1 | 272638 | 273031 | + | 130 | PF04134 | DUF393 | Protein of unknown function, DUF393 | 1.90E-14 |
| SDK49549.1 | 273111 | 274389 | + | 425 | TIGR02032<br>PF04820<br>PF01494<br>TIGR02023<br>TIGR01988 | TIGR02032<br>Trp_halogenase<br>FAD_binding_3<br>TIGR02023<br>TIGR01988 | GG-red-SF: geranylgeranyl reductase family<br>Tryptophan halogenase<br>FAD binding domain<br>BchP-ChlP: geranylgeranyl reductase<br>Ubi-OHases: ubiquinone biosynthesis hydroxylase, UbiH/UbiF/VisC/COQ6 family | 9.50E-38<br>1.40E-37<br>1.00E-23<br>2.00E-15<br>1.90E-13 |
| SDK49567.1 | 274580 | 274850 | + | 89 | LinA | LinA | LinA (linaridin precursor) | 5.80E-09 |
| SDK49593.1 | 274977 | 275145 | + | 55 | NO PFAM MATCH | - | - | - |

| | | | | | | | | |
|---|---|---|---|---|---|---|---|---|
| SDK49615.1 | 275180 | 275822 | + | 213 | PF05719 | GPP34 | Golgi phosphoprotein 3 (GPP34) | 3.80E-27 |
| SDK49642.1 | 276322 | 276124 | - | 65 | PF04149 | DUF397 | Domain of unknown function (DUF397) | 3.00E-18 |

| index | leader | core | start | end | dir | score |
|---|---|---|---|---|---|---|
| 1 | VQRVFPGQPCLVHGGQFRRSRRRGVTIGRTGSAGLLAGHVHT | VTTRRAARLSFPEETGSDAVSGCRCPS | 262468 | 262678 | + | 6 |
| 2 | VFPGQPCLVHGGQFRRSRRRGVTIGRTGSAGLLAGHVHT | VTTRRAARLSFPEETGSDAVSGCRCPS | 262477 | 262678 | + | 6 |
| 3 | VDAVSETVKQNPFPILCESIQTRRVSPL | GNRVKILNVTYDTMLCKRLTVRFCLPCER | 266774 | 266948 | + | 6 |
| 4 | MNRIESVANVQVPEAHAGAVRTLDTTGSTAPVL | TAAAITAAAVVSFAAGVASDAVESG...AASTGQDPTGMSSGELLQRRTALLG | 267128 | 267392 | + | 16 |
| 5 | MNRIENAATQQVTATAGAIEPMDSVEGVSPVL | TAAVAAGAVAATAAAFAAGNMVTDY...KHDGSELSGMSVGDLVNSRREAMSH | 274580 | 274850 | + | 16 |
| 6 | MSQSNNNAVIGGGAVLGAVVGGFVGIF | LGNPAIAAAIGLAVGIALATAMRAERKN | 274977 | 275145 | + | 7 |

### Results for CRK56201.1 [Alloactinosynnema sp. L-07] back to top

previous - next

#### Architecture

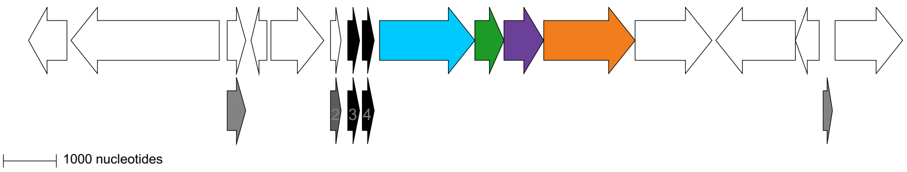

Link to nucleotide sequence

| Accession | start | end | direction | length (aa) | Pfam/HMM | name | description | E-value |
|---|---|---|---|---|---|---|---|---|
| CRK56193.1 | 1189640 | 1188938 | - | 233 | PF01925 | TauE | Sulfite exporter TauE/SafE | 1.10E-19 |
| CRK56194.1 | 1192527 | 1189716 | - | 936 | TIGR01341<br>TIGR02333<br>PF00330<br>TIGR01340<br>TIGR01342 | TIGR01341<br>TIGR02333<br>Aconitase<br>TIGR01340<br>TIGR01342 | aconitase_1: aconitate hydratase 1<br>2met_isocit_dHY: 2-methylisocitrate dehydratase, Fe/S-dependent<br>Aconitase family (aconitate hydratase)<br>aconitase_mito: aconitate hydratase, mitochondrial<br>acon_putative: putative aconitate hydratase | 0.00E+00<br>3.80E-252<br>7.60E-178<br>2.00E-73<br>8.90E-70 |
| CRK56195.1 | 1192673 | 1193033 | + | 119 | NO PFAM MATCH | - | - | - |
| CRK56196.1 | 1193428 | 1193140 | - | 95 | NO PFAM MATCH | - | - | - |
| CRK56197.1 | 1193515 | 1194496 | + | 326 | PF13384<br>PF13412 | HTH_23<br>HTH_24 | Homeodomain-like domain<br>Winged helix-turn-helix DNA-binding | 7.50E-04<br>8.10E-04 |
| CRK56198.1 | 1194624 | 1194840 | + | 71 | NO PFAM MATCH | - | - | - |
| CRK56199.1 | 1194967 | 1195189 | + | 73 | LinA | LinA | LinA (linaridin precursor) | 2.90E-13 |
| CRK56200.1 | 1195242 | 1195470 | + | 75 | LinA | LinA | LinA (linaridin precursor) | 2.20E-16 |
| CRK56201.1 | 1195548 | 1197369 | + | 606 | LinH<br>LinE<br>LinG<br>PF12697<br>TIGR03056 | LinH<br>LinE<br>LinG<br>Abhydrolase_6<br>TIGR03056 | LinH (Fusion of LinE/LinG)<br>LinE Homolog<br>LinG Homolog<br>Alpha/beta hydrolase family<br>bchO_mg_che_rel: putative magnesium chelatase accessory protein | 1.70E-253<br>3.50E-102<br>1.20E-96<br>3.80E-16<br>3.40E-14 |
| CRK56202.1 | 1197358 | 1197904 | + | 181 | LinL | LinL | LinL | 5.50E-65 |
| CRK56203.1 | 1197900 | 1198662 | + | 253 | PF13649<br>PF08241<br>PF13489<br>PF08242<br>PF13847 | Methyltransf_25<br>Methyltransf_11<br>Methyltransf_23<br>Methyltransf_12<br>Methyltransf_31 | Methyltransferase domain<br>Methyltransferase domain<br>Methyltransferase domain<br>Methyltransferase domain<br>Methyltransferase domain | 8.40E-16<br>3.90E-14<br>4.30E-11<br>1.40E-08<br>1.70E-05 |
| CRK56204.1 | 1198654 | 1200379 | + | 574 | TIGR02204<br>TIGR02203<br>TIGR00958<br>TIGR01846<br>TIGR03796 | TIGR02204<br>TIGR02203<br>TIGR00958<br>TIGR01846<br>TIGR03796 | MsbA_rel: ABC transporter, permease/ATP-binding protein<br>MsbA_lipidA: lipid A export permease/ATP-binding protein MsbA<br>3a01208: antigen peptide transporter 2<br>type_I_sec_HlyB: type I secretion system ATPase<br>NHLM_micro_ABC1: NHLM bacteriocin system ABC transporter, peptidase/ATP-binding protein | 6.60E-104<br>7.40E-102<br>6.90E-91<br>1.90E-86<br>1.00E-84 |
| CRK56205.1 | 1200392 | 1201832 | + | 479 | TIGR02734<br>TIGR02733<br>PF13450<br>TIGR03467<br>TIGR00292 | TIGR02734<br>TIGR02733<br>NAD_binding_8<br>TIGR03467<br>TIGR00292 | crtI_fam: phytoene desaturase<br>desat_CrtD: C-3',4' desaturase CrtD<br>NAD(P)-binding Rossmann-like domain<br>HpnE: squalene-associated FAD-dependent desaturase<br>TIGR00292: thiazole biosynthesis enzyme | 1.20E-17<br>5.60E-11<br>9.10E-10<br>6.30E-08<br>2.00E-07 |
| CRK56206.1 | 1203418 | 1201924 | - | 497 | NO PFAM MATCH | - | - | - |
| CRK56207.1 | 1203876 | 1203414 | - | 153 | NO PFAM MATCH | - | - | - |
| CRK56208.1 | 1204162 | 1205455 | + | 430 | PF00877<br>TIGR02168 | NLPC_P60<br>TIGR02168 | NlpC/P60 family<br>SMC_prok_B: chromosome segregation protein SMC | 3.90E-26<br>2.90E-06 |

| index | leader | core | start | end | dir | score |
|---|---|---|---|---|---|---|
| 1 | MIHHVQIACPEGAEDRMRAFYTGVL...LLAARGGCWFAVPGGGELHVGVESD | FRAARKAHPAFVVDIDAVGSALIQA...EIPGRGRFHTDDPVGNRLEFLEAER | 1192673 | 1193033 | + | 7 |
| 2 | MTFGLAAAALATLVGAGIASADEPAD | PTTPDGGGTTGAFMWGGCVCQGPVPVPDPVLPSVVATVPDFLVWH | 1194624 | 1194840 | + | 8 |
| 3 | MSTLVRHANEISPDLFGDLDAQTGARSLSGMTPVM | ATPTAVALGIAISAVAFHAGYMYGIARNMAEGSVVLPQ | 1194967 | 1195189 | + | 28 |
| 4 | MSSVLEKVNASAVFADIPLRADVPMTVGAASPVL | ATPATIVGIAAGVAVVAGVAGAFYAGYQIGRAIGHQNPLPQ | 1195242 | 1195470 | + | 29 |

| | | | | | | |
|---|---|---|---|---|---|---|
| 5 | LCARSGRSIDVSVTG | RVPDGKVAEGKYVPRVATGASGVQLAQSSCIQSVFRRFGGDRT | 1203937 | 1204114 | + | 7 |

### Results for RZU71018.1 [Actinopolyspora sp. DSM 45956] back to top

previous - next

#### Architecture

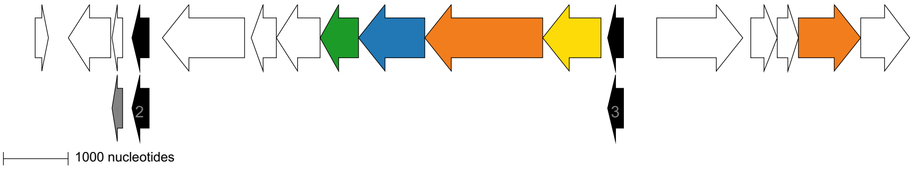

Link to nucleotide sequence

| Accession | start | end | direction | length (aa) | Pfam/HMM | name | description | E-value |
|---|---|---|---|---|---|---|---|---|
| RZU71010.1 | 4358423 | 4358624 | + | 66 | PF04149 | DUF397 | Domain of unknown function (DUF397) | 2.20E-21 |
| RZU71011.1 | 4359584 | 4358930 | - | 217 | PF05719 | GPP34 | Golgi phosphoprotein 3 (GPP34) | 1.90E-27 |
| RZU71012.1 | 4359784 | 4359619 | - | 54 | NO PFAM MATCH | - | - | - |
| RZU71013.1 | 4360180 | 4359910 | - | 89 | LinA | LinA | LinA (linaridin precursor) | 6.50E-09 |
| RZU71014.1 | 4361649 | 4360389 | - | 419 | TIGR02032<br>PF04820<br>PF01494<br>TIGR02023<br>TIGR01988 | TIGR02032<br>Trp_halogenase<br>FAD_binding_3<br>TIGR02023<br>TIGR01988 | GG-red-SF: geranylgeranyl reductase family<br>Tryptophan halogenase<br>FAD binding domain<br>BchP-ChlP: geranylgeranyl reductase<br>Ubi-OHases: ubiquinone biosynthesis hydroxylase, UbiH/UbiF/VisC/COQ6 family | 5.60E-40<br>8.10E-39<br>1.60E-24<br>6.70E-17<br>3.70E-15 |
| RZU71015.1 | 4362151 | 4361758 | - | 130 | PF04134 | DUF393 | Protein of unknown function, DUF393 | 2.30E-14 |
| RZU71016.1 | 4362833 | 4362152 | - | 226 | PF00196<br>TIGR03020<br>TIGR03541<br>TIGR02937 | GerE<br>TIGR03020<br>TIGR03541<br>TIGR02937 | Bacterial regulatory proteins, luxR family<br>EpsA: transcriptional regulator EpsA<br>reg_near_HchA: LuxR family transcriptional regulatory, chaperone HchA-associated<br>sigma70-ECF: RNA polymerase sigma factor, sigma-70 family | 3.50E-19<br>2.40E-14<br>2.10E-09<br>3.00E-04 |
| RZU71017.1 | 4363420 | 4362829 | - | 196 | LinL | LinL | LinL | 1.40E-67 |
| RZU71018.1 | 4364442 | 4363419 | - | 340 | LinE<br>LinH<br>PF12697<br>PF00561<br>TIGR02427 | LinE<br>LinH<br>Abhydrolase_6<br>Abhydrolase_1<br>TIGR02427 | LinE Homolog<br>LinH (Fusion of LinE/LinG)<br>Alpha/beta hydrolase family<br>alpha/beta hydrolase fold<br>protocat_pcaD: 3-oxoadipate enol-lactonase | 6.10E-104<br>7.70E-81<br>3.40E-16<br>2.40E-14<br>2.10E-13 |
| RZU71019.1 | 4366266 | 4364442 | - | 607 | TIGR02204<br>TIGR02203<br>TIGR00958<br>TIGR01846<br>TIGR03796 | TIGR02204<br>TIGR02203<br>TIGR00958<br>TIGR01846<br>TIGR03796 | MsbA_rel: ABC transporter, permease/ATP-binding protein<br>MsbA_lipidA: lipid A export permease/ATP-binding protein MsbA<br>3a01208: antigen peptide transporter 2<br>type_I_sec_HlyB: type I secretion system ATPase<br>NHLM_micro_ABC1: NHLM bacteriocin system ABC transporter, peptidase/ATP-binding protein | 3.10E-118<br>2.10E-107<br>3.30E-94<br>3.00E-90<br>1.30E-85 |
| RZU71020.1 | 4367165 | 4366271 | - | 297 | LinG<br>LinH<br>TIGR04033 | LinG<br>LinH<br>TIGR04033 | LinG Homolog<br>LinH (Fusion of LinE/LinG)<br>export_SdpB: antimicrobial peptide system protein, SdpB family | 7.10E-103<br>3.50E-92<br>1.20E-05 |
| RZU71021.1 | 4367522 | 4367258 | - | 87 | LinA | LinA | LinA (linaridin precursor) | 9.90E-08 |
| RZU71022.1 | 4368018 | 4369356 | + | 445 | PF05193<br>PF00675<br>TIGR02110 | Peptidase_M16_C<br>Peptidase_M16<br>TIGR02110 | Peptidase M16 inactive domain<br>Insulinase (Peptidase family M16)<br>PQQ_syn_pqqF: coenzyme PQQ biosynthesis protein PqqF | 1.00E-37<br>1.40E-33<br>1.00E-19 |
| RZU71023.1 | 4369469 | 4369892 | + | 140 | PF06738 | ThrE | Putative threonine/serine exporter | 3.60E-04 |
| RZU71024.1 | 4369888 | 4370215 | + | 108 | PF13601<br>PF12840<br>PF01022<br>PF03551<br>PF02082 | HTH_34<br>HTH_20<br>HTH_5<br>PadR<br>Rrf2 | Winged helix DNA-binding domain<br>Helix-turn-helix domain<br>Bacterial regulatory protein, arsR family<br>Transcriptional regulator PadR-like family<br>Transcriptional regulator | 3.70E-26<br>7.10E-07<br>1.30E-05<br>2.70E-05<br>2.10E-04 |
| RZU71025.1 | 4370211 | 4371171 | + | 319 | TIGR01188<br>TIGR01288<br>TIGR03522<br>TIGR03864<br>TIGR04406 | TIGR01188<br>TIGR01288<br>TIGR03522<br>TIGR03864<br>TIGR04406 | drrA: daunorubicin resistance ABC transporter, ATP-binding protein<br>nodI: nodulation ABC transporter NodI<br>GldA_ABC_ATP: gliding motility-associated ABC transporter ATP-binding subunit GldA<br>PQQ_ABC_ATP: ABC transporter, ATP-binding subunit, PQQ-dependent alcohol dehydrogenase system<br>LPS_export_lptB: LPS export ABC transporter ATP-binding protein | 1.10E-64<br>2.50E-59<br>4.30E-49<br>1.00E-48<br>4.10E-41 |
| RZU71026.1 | 4371177 | 4371933 | + | 251 | PF01061<br>PF12698<br>TIGR01247<br>PF12679<br>TIGR03861 | ABC2_membrane<br>ABC2_membrane_3<br>TIGR01247<br>ABC2_membrane_2<br>TIGR03861 | ABC-2 type transporter<br>ABC-2 family transporter protein<br>drrB: daunorubicin resistance ABC transporter membrane protein<br>ABC-2 family transporter protein<br>phenyl_ABC_PedC: alcohol ABC transporter, permease protein | 6.60E-16<br>3.80E-13<br>7.60E-12<br>5.60E-10<br>6.10E-06 |

| index | leader | core | start | end | dir | score |
|---|---|---|---|---|---|---|
| 1 | MSQNNNAVIGGGAVLGAVVGGFVGIFL | GNPAIAAAIGLAVGIALATAMRAERKN | 4359784 | 4359619 | - | 7 |
| 2 | MNRIENAATQQVTASAGAIEPMDSVEGVSPVL | TAAVAAGAVAATAAAFAAGNMVTDY...KHDGSELSDMSVGDLVNSRHEAMSR | 4360180 | 4359910 | - | 16 |
| 3 | MNRIESVANAQLPDAHDGAVRSLDTTGSTAPVL | TAAAITAAAVVCFAAGVASDAVESG...AASTGQDPTGMSSGELLQRRTALLG | 4367522 | 4367258 | - | 17 |

### Results for OZM71980.1 [Amycolatopsis antarctica] back to top

previous - next

#### Architecture

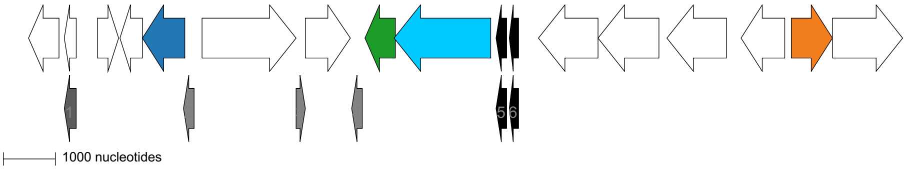

Link to nucleotide sequence

| Accession | start | end | direction | length (aa) | Pfam/HMM | name | description | E-value |
|---|---|---|---|---|---|---|---|---|
| OZM71975.1 | 182616 | 182046 | - | 189 | PF13302<br>PF00583 | Acetyltransf_3<br>Acetyltransf_1 | Acetyltransferase (GNAT) domain<br>Acetyltransferase (GNAT) family | 2.90E-26<br>4.40E-08 |
| OZM71976.1 | 182950 | 182740 | - | 69 | TIGR00638<br>PF03459<br>TIGR02142 | TIGR00638<br>TOBE<br>TIGR02142 | Mop: molybdenum-pterin binding domain<br>TOBE domain<br>modC_ABC: molybdate ABC transporter, ATP-binding protein | 5.90E-22<br>3.80E-20<br>3.40E-08 |
| OZM72050.1 | 183369 | 183786 | + | 138 | NO PFAM MATCH | - | - | - |
| OZM71977.1 | 184231 | 183793 | - | 145 | PF12802<br>TIGR00122<br>PF13551<br>PF01047<br>PF13518 | MarR_2<br>TIGR00122<br>HTH_29<br>MarR<br>HTH_28 | MarR family<br>birA_repr_reg: biotin operon repressor<br>Winged helix-turn helix<br>MarR family<br>Helix-turn-helix domain | 2.60E-08<br>1.90E-05<br>5.20E-05<br>5.50E-05<br>1.10E-04 |
| OZM71978.1 | 185037 | 184227 | - | 269 | TIGR02427<br>PF12697<br>LinE<br>LinH<br>PF12146 | TIGR02427<br>Abhydrolase_6<br>LinE<br>LinH<br>Hydrolase_4 | protocat_pcaD: 3-oxoadipate enol-lactonase<br>Alpha/beta hydrolase family<br>LinE Homolog<br>LinH (Fusion of LinE/LinG)<br>Serine aminopeptidase, S33 | 1.20E-26<br>5.30E-22<br>1.80E-19<br>1.30E-18<br>1.80E-18 |
| WP_004446014.1 | 185379 | 187197 | + | 606 | INFERRED GENE | - | - | - |
| OZM72051.1 | 187360 | 188233 | + | 290 | PF01551 | Peptidase_M23 | Peptidase family M23 | 1.10E-17 |
| OZM71979.1 | 189087 | 188520 | - | 188 | LinL | LinL | LinL | 6.80E-65 |
| OZM71980.1 | 190927 | 189067 | - | 619 | LinH<br>LinG<br>LinE<br>PF12697<br>PF12146 | LinH<br>LinG<br>LinE<br>Abhydrolase_6<br>Hydrolase_4 | LinH (Fusion of LinE/LinG)<br>LinG Homolog<br>LinE Homolog<br>Alpha/beta hydrolase family<br>Serine aminopeptidase, S33 | 1.80E-248<br>1.30E-102<br>1.00E-98<br>2.30E-17<br>5.00E-11 |
| OZM71981.1 | 191239 | 191038 | - | 66 | LinA | LinA | LinA (linaridin precursor) | 5.90E-18 |
| OZM71982.1 | 191477 | 191285 | - | 63 | LinA | LinA | LinA (linaridin precursor) | 7.10E-13 |
| OZM71983.1 | 192996 | 191838 | - | 385 | PF13556 | HTH_30 | PucR C-terminal helix-turn-helix domain | 1.10E-14 |
| OZM71984.1 | 194171 | 192992 | - | 392 | PF13556<br>PF02954 | HTH_30<br>HTH_8 | PucR C-terminal helix-turn-helix domain<br>Bacterial regulatory protein, Fis family | 1.90E-14<br>5.30E-04 |
| OZM71985.1 | 195470 | 194318 | - | 383 | PF08448<br>TIGR00229<br>PF13426<br>PF00196<br>PF08447 | PAS_4<br>TIGR00229<br>PAS_9<br>GerE<br>PAS_3 | PAS fold<br>sensory_box: PAS domain S-box protein<br>PAS domain<br>Bacterial regulatory proteins, luxR family<br>PAS fold | 4.70E-20<br>1.10E-17<br>7.90E-09<br>8.40E-09<br>1.10E-06 |
| OZM71986.1 | 196571 | 195752 | - | 272 | PF02909<br>PF00440 | TetR_C_1<br>TetR_N | Tetracyclin repressor-like, C-terminal domain<br>Bacterial regulatory proteins, tetR family | 2.10E-16<br>1.10E-06 |
| OZM71987.1 | 196712 | 197510 | + | 265 | TIGR02982<br>TIGR02211<br>TIGR02314<br>TIGR03608<br>TIGR02673 | TIGR02982<br>TIGR02211<br>TIGR02314<br>TIGR03608<br>TIGR02673 | heterocyst_DevA: ABC exporter ATP-binding subunit, DevA family<br>LolD_lipo_ex: lipoprotein releasing system, ATP-binding protein<br>ABC_MetN: D-methionine ABC transporter, ATP-binding protein<br>L_ocin_972_ABC: putative bacteriocin export ABC transporter, lactococcin 972 group<br>FtsE: cell division ATP-binding protein FtsE | 1.70E-56<br>5.20E-55<br>8.70E-54<br>1.90E-52<br>1.70E-51 |
| OZM71988.1 | 197506 | 198853 | + | 448 | PF02687<br>TIGR03434 | FtsX<br>TIGR03434 | FtsX-like permease family<br>ADOP: acidobacterial duplicated orphan permease | 2.40E-10<br>2.70E-04 |

| index | leader | core | start | end | dir | score |
|---|---|---|---|---|---|---|
| 1 | MRLSTRNQLTGTIASITEGSVMTVVTLDLDGGQQ | VTASITKEAAEDLGLAVGMPATALVKSTEVMLGVE | 182950 | 182740 | - | 8 |
| 2 | VGVPGRGGGAGHRASSPDFVVGVLVRQHSLC | RWPGPAQRLQEVATVVVDRRSPTGCFAIFRTC | 185222 | 185030 | - | 7 |
| 3 | LTGAGGAAGDGAPGQVPRGGRRSAPHAQLIHPPGD | GTVPTCRSTAGAVTAMVKAVR | 187193 | 187364 | + | 7 |
| 4 | LPCGTAHDAIPSPAKG | RTPLLPERESPYVDVLRWLPAQYVD...LRGSGTVWDGLPGTGWRVEPGLLVG | 188462 | 188258 | - | 7 |
| 5 | MSNLADKVNADRELARTPLLATEPQHVGALTPVL | ATPAAFGAGLAGAAAVAGTFGAGYVAGNAIGG | 191239 | 191038 | - | 28 |
| 6 | MDKLVSYTNAVSEDMFGELELAVDVRDMSAATPCM | ATPAAVAAGATLGAAAFGAGFGIGYAYG | 191477 | 191285 | - | 28 |

### Results for WP_086852701.1 [Amycolatopsis kentuckyensis] back to top

previous - next

#### Architecture

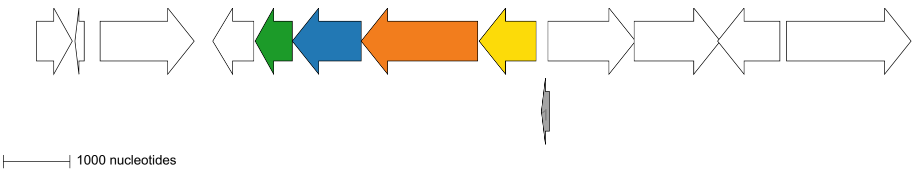

Link to nucleotide sequence

| Accession | start | end | direction | length (aa) | Pfam/HMM | name | description | E-value |
|---|---|---|---|---|---|---|---|---|
| WP_019532417.1 | 0 | 539 | + | 179 | INFERRED GENE | - | - | - |
| WP_158104608.1 | 735 | 588 | - | 48 | NO PFAM MATCH | - | - | - |
| WP_086852695.1 | 955 | 2383 | + | 475 | TIGR02734<br>TIGR03467<br>PF13450<br>TIGR01317<br>TIGR03315 | TIGR02734<br>TIGR03467<br>NAD_binding_8<br>TIGR01317<br>TIGR03315 | crtI_fam: phytoene desaturase<br>HpnE: squalene-associated FAD-dependent desaturase<br>NAD(P)-binding Rossmann-like domain<br>GOGAT_sm_gam: glutamate synthase, NADH/NADPH, small subunit<br>Se_ygfK: putative selenate reductase, YgfK subunit | 1.20E-11<br>3.90E-11<br>1.10E-10<br>8.50E-08<br>1.00E-06 |
| WP_143264388.1 | 3275 | 2651 | - | 207 | PF00196<br>TIGR03020<br>TIGR03541<br>PF08281<br>TIGR02937 | GerE<br>TIGR03020<br>TIGR03541<br>Sigma70_r4_2<br>TIGR02937 | Bacterial regulatory proteins, luxR family<br>EpsA: transcriptional regulator EpsA<br>reg_near_HchA: LuxR family transcriptional regulatory, chaperone HchA-associated<br>Sigma-70, region 4<br>sigma70-ECF: RNA polymerase sigma factor, sigma-70 family | 4.80E-19<br>1.70E-13<br>2.00E-09<br>1.50E-06<br>1.90E-06 |
| WP_086852699.1 | 3853 | 3301 | - | 183 | LinL | LinL | LinL | 2.00E-69 |
| WP_086852701.1 | 4887 | 3849 | - | 345 | LinE<br>LinH<br>TIGR02427<br>PF00561<br>PF12697 | LinE<br>LinH<br>TIGR02427<br>Abhydrolase_1<br>Abhydrolase_6 | LinE Homolog<br>LinH (Fusion of LinE/LinG)<br>protocat_pcaD: 3-oxoadipate enol-lactonase<br>alpha/beta hydrolase fold<br>Alpha/beta hydrolase family | 1.80E-97<br>6.20E-67<br>4.30E-15<br>2.30E-14<br>4.00E-14 |
| WP_086852703.1 | 6644 | 4883 | - | 586 | TIGR02204<br>TIGR02203<br>TIGR00958<br>TIGR01846<br>TIGR03796 | TIGR02204<br>TIGR02203<br>TIGR00958<br>TIGR01846<br>TIGR03796 | MsbA_rel: ABC transporter, permease/ATP-binding protein<br>MsbA_lipidA: lipid A export permease/ATP-binding protein MsbA<br>3a01208: antigen peptide transporter 2<br>type_I_sec_HlyB: type I secretion system ATPase<br>NHLM_micro_ABC1: NHLM bacteriocin system ABC transporter, peptidase/ATP-binding protein | 6.60E-97<br>8.40E-94<br>1.70E-83<br>2.10E-80<br>1.20E-77 |
| WP_086852705.1 | 7511 | 6653 | - | 285 | LinG<br>LinH<br>TIGR04033 | LinG<br>LinH<br>TIGR04033 | LinG Homolog<br>LinH (Fusion of LinE/LinG)<br>export_SdpB: antimicrobial peptide system protein, SdpB family | 9.60E-97<br>1.10E-87<br>2.60E-04 |
| WP_086852709.1 | 7698 | 9006 | + | 435 | TIGR02110<br>PF05193<br>PF00675 | TIGR02110<br>Peptidase_M16_C<br>Peptidase_M16 | PQQ_syn_pqqF: coenzyme PQQ biosynthesis protein PqqF<br>Peptidase M16 inactive domain<br>Insulinase (Peptidase family M16) | 1.70E-20<br>1.80E-20<br>1.40E-19 |
| WP_086852706.1 | 9002 | 10286 | + | 427 | PF05193<br>PF00675 | Peptidase_M16_C<br>Peptidase_M16 | Peptidase M16 inactive domain<br>Insulinase (Peptidase family M16) | 1.40E-21<br>4.60E-04 |
| WP_086852708.1 | 11191 | 10249 | - | 313 | TIGR03945<br>TIGR01136<br>TIGR01139<br>TIGR01137<br>PF00291 | TIGR03945<br>TIGR01136<br>TIGR01139<br>TIGR01137<br>PALP | PLP_SbnA_fam: 2,3-diaminopropionate biosynthesis protein SbnA<br>cysKM: cysteine synthase<br>cysK: cysteine synthase A<br>cysta_beta: cystathionine beta-synthase<br>Pyridoxal-phosphate dependent enzyme | 3.50E-113<br>7.70E-65<br>4.80E-58<br>9.90E-54<br>1.60E-52 |
| CCN39_RS44110 | 11279 | 13150 | + | 623 | INFERRED GENE | - | - | - |

| index | leader | core | start | end | dir | score |
|---|---|---|---|---|---|---|
| 1 | VLFMTVATVAVRGGRGRE | VITSADAASVFAGTASPPD | 7709 | 7595 | - | 6 |

### Results for OZM72833.1 [Amycolatopsis antarctica] back to top

previous - next

#### Architecture

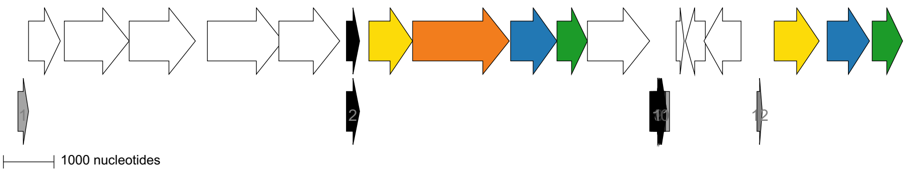

Link to nucleotide sequence

| Accession | start | end | direction | length (aa) | Pfam/HMM | name | description | E-value |
|---|---|---|---|---|---|---|---|---|
| OZM72619.1 | 101023 | 101668 | + | 214 | NO PFAM MATCH | - | - | - |
| OZM72620.1 | 101734 | 103033 | + | 432 | PF05193<br>PF00675<br>TIGR02110 | Peptidase_M16_C<br>Peptidase_M16<br>TIGR02110 | Peptidase M16 inactive domain<br>Insulinase (Peptidase family M16)<br>PQQ_syn_pqqF: coenzyme PQQ biosynthesis protein PqqF | 2.10E-20<br>2.40E-19<br>8.50E-10 |
| OZM72621.1 | 103029 | 104331 | + | 433 | PF05193<br>PF00675 | Peptidase_M16_C<br>Peptidase_M16 | Peptidase M16 inactive domain<br>Insulinase (Peptidase family M16) | 9.50E-20<br>8.80E-05 |
| OZM72622.1 | 104564 | 106070 | + | 501 | TIGR04022<br>PF08028<br>PF02771 | TIGR04022<br>Acyl-CoA_dh_2<br>Acyl-CoA_dh_N | sulfur_SfnB: sulfur acquisition oxidoreductase, SfnB family<br>Acyl-CoA dehydrogenase, C-terminal domain<br>Acyl-CoA dehydrogenase, N-terminal domain | 1.10E-14<br>2.60E-09<br>7.50E-07 |
| OZM72623.1 | 105984 | 107190 | + | 401 | PF08028<br>PF00441 | Acyl-CoA_dh_2<br>Acyl-CoA_dh_1 | Acyl-CoA dehydrogenase, C-terminal domain<br>Acyl-CoA dehydrogenase, C-terminal domain | 1.10E-16<br>3.50E-09 |
| OZM72624.1 | 107317 | 107578 | + | 86 | LinA | LinA | LinA (linaridin precursor) | 3.40E-14 |

| | | | | | | | | |
|---|---|---|---|---|---|---|---|---|
| OZM72625.1 | 107752 | 108619 | + | 288 | LinG<br>LinH | LinG<br>LinH | LinG Homolog<br>LinH (Fusion of LinE/LinG) | 3.90E-102<br>2.80E-89 |
| OZM72626.1 | 108621 | 110544 | + | 640 | TIGR02204<br>TIGR02203<br>TIGR00958<br>TIGR03796<br>TIGR03375 | TIGR02204<br>TIGR02203<br>TIGR00958<br>TIGR03796<br>TIGR03375 | MsbA_rel: ABC transporter, permease/ATP-binding protein<br>MsbA_lipidA: lipid A export permease/ATP-binding protein MsbA<br>3a01208: antigen peptide transporter 2<br>NHLM_micro_ABC1: NHLM bacteriocin system ABC transporter, peptidase/ATP-binding protein<br>type_I_sec_LssB: type I secretion system ATPase | 3.40E-106<br>1.90E-93<br>3.60E-89<br>3.10E-83<br>1.60E-75 |
| OZM72833.1 | 110549 | 111488 | + | 312 | LinE<br>LinH<br>TIGR02427<br>PF12697<br>PF00561 | LinE<br>LinH<br>TIGR02427<br>Abhydrolase_6<br>Abhydrolase_1 | LinE Homolog<br>LinH (Fusion of LinE/LinG)<br>protocat_pcaD: 3-oxoadipate enol-lactonase<br>Alpha/beta hydrolase family<br>alpha/beta hydrolase fold | 5.30E-84<br>8.10E-64<br>3.40E-18<br>9.00E-13<br>1.50E-11 |
| OZM72834.1 | 111480 | 112074 | + | 197 | LinL | LinL | LinL | 5.00E-60 |
| OZM72627.1 | 112070 | 113312 | + | 413 | PF04820<br>TIGR02032<br>PF01266<br>PF01494<br>TIGR02023 | Trp_halogenase<br>TIGR02032<br>DAO<br>FAD_binding_3<br>TIGR02023 | Tryptophan halogenase<br>GG-red-SF: geranylgeranyl reductase family<br>FAD dependent oxidoreductase<br>FAD binding domain<br>BchP-ChlP: geranylgeranyl reductase | 1.70E-21<br>1.70E-21<br>1.70E-11<br>1.00E-09<br>3.40E-07 |
| OZM72628.1 | 113825 | 113996 | + | 56 | NO PFAM MATCH | - | - | - |
| OZM72629.1 | 114401 | 114020 | - | 126 | PF04134 | DUF393 | Protein of unknown function, DUF393 | 4.10E-18 |
| OZM72630.1 | 115104 | 114393 | - | 236 | PF00196<br>TIGR03020<br>TIGR03541<br>TIGR02937<br>PF08281 | GerE<br>TIGR03020<br>TIGR03541<br>TIGR02937<br>Sigma70_r4_2 | Bacterial regulatory proteins, luxR family<br>EpsA: transcriptional regulator EpsA<br>reg_near_HchA: LuxR family transcriptional regulatory, chaperone HchA-associated<br>sigma70-ECF: RNA polymerase sigma factor, sigma-70 family<br>Sigma-70, region 4 | 2.20E-17<br>1.00E-12<br>4.50E-09<br>9.00E-06<br>2.60E-05 |
| OZM72631.1 | 115777 | 116671 | + | 297 | LinG<br>LinH<br>LinL<br>TIGR04033 | LinG<br>LinH<br>LinL<br>TIGR04033 | LinG Homolog<br>LinH (Fusion of LinE/LinG)<br>LinL<br>export_SdpB: antimicrobial peptide system protein, SdpB family | 1.40E-102<br>6.90E-94<br>7.30E-19<br>2.80E-05 |
| OZM72835.1 | 116817 | 117645 | + | 275 | LinE<br>LinH<br>PF12697<br>TIGR02427<br>PF00561 | LinE<br>LinH<br>Abhydrolase_6<br>TIGR02427<br>Abhydrolase_1 | LinE Homolog<br>LinH (Fusion of LinE/LinG)<br>Alpha/beta hydrolase family<br>protocat_pcaD: 3-oxoadipate enol-lactonase<br>alpha/beta hydrolase fold | 5.00E-87<br>4.50E-69<br>3.80E-14<br>3.00E-11<br>5.30E-09 |
| OZM72632.1 | 117697 | 118300 | + | 200 | LinL<br>LinG | LinL<br>LinG | LinL<br>LinG Homolog | 4.00E-71<br>2.00E-05 |

| index | leader | core | start | end | dir | score |
|---|---|---|---|---|---|---|
| 1 | VLTGTDPEWAPDGLIS | STVRDERTELLVHAEHCDAVLAEALRNGASVREVREVRDLPDAREVLPE | 100829 | 101027 | + | 6 |
| 2 | MSNVEAFANENADGVLNPLQVTTALGPVL | ATPAYAAYTAGAAAVFTAGLISDSV...VTPTGPVPAYGSVDELLGMRISSIN | 107317 | 107578 | + | 28 |
| 3 | MRDNMRPVGFPLINTIDRNGGIGMN...AQVAFQKTEGVLRPFERVSGNAPVL | ATPAAAAVVASATALYAAGNMVTDL...KNPGINLAGRSGAELLNIRAEGLRG | 113315 | 113654 | + | 26 |
| 4 | MRPVGFPLINTIDRNGGIGMNGIEK...AQVAFQKTEGVLRPFERVSGNAPVL | ATPAAAAVVASATALYAAGNMVTDL...KNPGINLAGRSGAELLNIRAEGLRG | 113327 | 113654 | + | 26 |
| 5 | MNGIEKAAQVAFQKTEGVLRPFERVSGNAPVL | ATPAAAAVVASATALYAAGNMVTDL...KNPGINLAGRSGAELLNIRAEGLRG | 113384 | 113654 | + | 26 |
| 6 | VSGNAPVLATPAAAAVVASATALYAAGNMVTD | LVGGQESAVKNPGINLAGRSGAELLNIRAEGLRG | 113456 | 113654 | + | 12 |
| 7 | LTADSWPTRSVTMLPAAYSAVAE | ATTAAAAGVARTGALPLTRSNGRRTPSVF | 113578 | 113419 | - | 6 |
| 8 | MPGFLTADSWPTRSVTMLPAAYSAVAEA | TTAAAAGVARTGALPLTRSNGRRTPSVF | 113590 | 113419 | - | 7 |
| 9 | LMPGFLTADSWPTRSVTMLPAAYSAVAE | ATTAAAAGVARTGALPLTRSNGRRTPSVF | 113593 | 113419 | - | 6 |
| 10 | MFSSSAPDLPARLMPGF | LTADSWPTRSVTMLPAAYSAVAEAT...AAAGVARTGALPLTRSNGRRTPSVF | 113629 | 113419 | - | 7 |
| 11 | MNSSIAVCPVGLLNQPRRPSARMFSSSAPDLPARLMPGF | LTADSWPTRSVTMLPAAYSAVAEAT...AAAGVARTGALPLTRSNGRRTPSVF | 113695 | 113419 | - | 6 |
| 12 | LEPACPPPLVQTVGMRPDG | LTPPTLVVGQRDRAA | 115425 | 115530 | + | 7 |

### Results for OZM72835.1 [Amycolatopsis antarctica] back to top

previous - next

#### Architecture

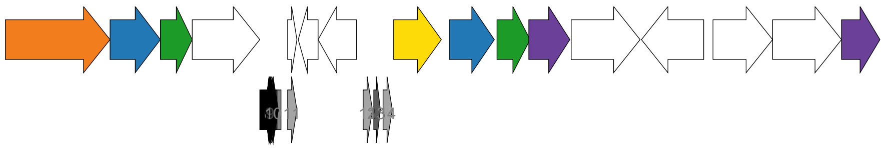

1000 nucleotides

Link to nucleotide sequence

| Accession | start | end | direction | length (aa) | Pfam/HMM | name | description | E-value |
|---|---|---|---|---|---|---|---|---|
| OZM72626.1 | 108621 | 110544 | + | 640 | TIGR02204<br>TIGR02203<br>TIGR00958<br>TIGR03796<br>TIGR03375 | TIGR02204<br>TIGR02203<br>TIGR00958<br>TIGR03796<br>TIGR03375 | MsbA_rel: ABC transporter, permease/ATP-binding protein<br>MsbA_lipidA: lipid A export permease/ATP-binding protein MsbA<br>3a01208: antigen peptide transporter 2<br>NHLM_micro_ABC1: NHLM bacteriocin system ABC transporter, peptidase/ATP-binding protein<br>type_I_sec_LssB: type I secretion system ATPase | 3.40E-106<br>1.90E-93<br>3.60E-89<br>3.10E-83<br>1.60E-75 |

| | | | | | | | | |
|---|---|---|---|---|---|---|---|---|
| OZM72833.1 | 110549 | 111488 | + | 312 | LinE<br>LinH<br>TIGR02427<br>PF12697<br>PF00561 | LinE<br>LinH<br>TIGR02427<br>Abhydrolase_6<br>Abhydrolase_1 | LinE Homolog<br>LinH (Fusion of LinE/LinG)<br>protocat_pcaD: 3-oxoadipate enol-lactonase<br>Alpha/beta hydrolase family<br>alpha/beta hydrolase fold | 5.30E-84<br>8.10E-64<br>3.40E-18<br>9.00E-13<br>1.50E-11 |
| OZM72834.1 | 111480 | 112074 | + | 197 | LinL | LinL | LinL | 5.00E-60 |
| OZM72627.1 | 112070 | 113312 | + | 413 | PF04820<br>TIGR02032<br>PF01266<br>PF01494<br>TIGR02023 | Trp_halogenase<br>TIGR02032<br>DAO<br>FAD_binding_3<br>TIGR02023 | Tryptophan halogenase<br>GG-red-SF: geranylgeranyl reductase family<br>FAD dependent oxidoreductase<br>FAD binding domain<br>BchP-ChlP: geranylgeranyl reductase | 1.70E-21<br>1.70E-21<br>1.70E-11<br>1.00E-09<br>3.40E-07 |
| OZM72628.1 | 113825 | 113996 | + | 56 | NO PFAM MATCH | - | - | - |
| OZM72629.1 | 114401 | 114020 | - | 126 | PF04134 | DUF393 | Protein of unknown function, DUF393 | 4.10E-18 |
| OZM72630.1 | 115104 | 114393 | - | 236 | PF00196<br>TIGR03020<br>TIGR03541<br>TIGR02937<br>PF08281 | GerE<br>TIGR03020<br>TIGR03541<br>TIGR02937<br>Sigma70_r4_2 | Bacterial regulatory proteins, luxR family<br>EpsA: transcriptional regulator EpsA<br>reg_near_HchA: LuxR family transcriptional regulatory, chaperone HchA-associated<br>sigma70-ECF: RNA polymerase sigma factor, sigma-70 family<br>Sigma-70, region 4 | 2.20E-17<br>1.00E-12<br>4.50E-09<br>9.00E-06<br>2.60E-05 |
| OZM72631.1 | 115777 | 116671 | + | 297 | LinG<br>LinH<br>LinL<br>TIGR04033 | LinG<br>LinH<br>LinL<br>TIGR04033 | LinG Homolog<br>LinH (Fusion of LinE/LinG)<br>LinL<br>export_SdpB: antimicrobial peptide system protein, SdpB family | 1.40E-102<br>6.90E-94<br>7.30E-19<br>2.80E-05 |
| OZM72835.1 | 116817 | 117645 | + | 275 | LinE<br>LinH<br>PF12697<br>TIGR02427<br>PF00561 | LinE<br>LinH<br>Abhydrolase_6<br>TIGR02427<br>Abhydrolase_1 | LinE Homolog<br>LinH (Fusion of LinE/LinG)<br>Alpha/beta hydrolase family<br>protocat_pcaD: 3-oxoadipate enol-lactonase<br>alpha/beta hydrolase fold | 5.00E-87<br>4.50E-69<br>3.80E-14<br>3.00E-11<br>5.30E-09 |
| OZM72632.1 | 117697 | 118300 | + | 200 | LinL<br>LinG | LinL<br>LinG | LinL<br>LinG Homolog | 4.00E-71<br>2.00E-05 |
| OZM72633.1 | 118284 | 119022 | + | 245 | PF13649<br>TIGR00755<br>TIGR02021<br>PF08241<br>PF13489 | Methyltransf_25<br>TIGR00755<br>TIGR02021<br>Methyltransf_11<br>Methyltransf_23 | Methyltransferase domain<br>ksgA: ribosomal RNA small subunit methyltransferase A<br>BchM-ChlM: magnesium protoporphyrin O-methyltransferase<br>Methyltransferase domain<br>Methyltransferase domain | 1.30E-11<br>1.50E-09<br>2.60E-09<br>9.10E-09<br>4.20E-08 |
| OZM72634.1 | 119052 | 120324 | + | 423 | TIGR01426<br>PF03033<br>PF00201 | TIGR01426<br>Glyco_transf_28<br>UDPGT | MGT: glycosyltransferase, MGT family<br>Glycosyltransferase family 28 N-terminal domain<br>UDP-glucoronosyl and UDP-glucosyl transferase | 7.00E-22<br>6.70E-17<br>5.10E-06 |
| OZM72635.1 | 121512 | 120354 | - | 385 | PF17843<br>PF13578 | MycE_N<br>Methyltransf_24 | MycE methyltransferase N-terminal<br>Methyltransferase domain | 9.60E-20<br>6.40E-05 |
| OZM72636.1 | 121676 | 122765 | + | 362 | NO PFAM MATCH | - | - | - |
| OZM72836.1 | 122779 | 124036 | + | 418 | TIGR01426<br>PF06722<br>PF00201<br>PF04101 | TIGR01426<br>DUF1205<br>UDPGT<br>Glyco_tran_28_C | MGT: glycosyltransferase, MGT family<br>Protein of unknown function (DUF1205)<br>UDP-glucoronosyl and UDP-glucosyl transferase<br>Glycosyltransferase family 28 C-terminal domain | 2.70E-29<br>1.70E-27<br>4.10E-10<br>2.10E-05 |
| OZM72637.1 | 124040 | 124751 | + | 236 | PF13649<br>PF08241<br>PF13847<br>PF13489<br>TIGR02021 | Methyltransf_25<br>Methyltransf_11<br>Methyltransf_31<br>Methyltransf_23<br>TIGR02021 | Methyltransferase domain<br>Methyltransferase domain<br>Methyltransferase domain<br>Methyltransferase domain<br>BchM-ChlM: magnesium protoporphyrin O-methyltransferase | 1.10E-15<br>2.20E-12<br>6.20E-10<br>7.80E-09<br>7.40E-08 |

| index | leader | core | start | end | dir | score |
|---|---|---|---|---|---|---|
| 1 | MRDNMRPVGFPLINTIDRNGGIGMN...AQVAFQKTEGVLRPFERVSGNAPVL | ATPAAAAVVASATALYAAGNMVTDL...KNPGINLAGRSGAELLNIRAEGLRG | 113315 | 113654 | + | 27 |
| 2 | MRPVGFPLINTIDRNGGIGMNGIEK...AQVAFQKTEGVLRPFERVSGNAPVL | ATPAAAAVVASATALYAAGNMVTDL...KNPGINLAGRSGAELLNIRAEGLRG | 113327 | 113654 | + | 27 |
| 3 | MNGIEKAAQVAFQKTEGVLRPFERVSGNAPVL | ATPAAAAVVASATALYAAGNMVTDL...KNPGINLAGRSGAELLNIRAEGLRG | 113384 | 113654 | + | 27 |
| 4 | VSGNAPVLATPAAAAVVASATALYAAGNMVTD | LVGQESAVKNPGINLAGRSGAELLNIRAEGLRG | 113456 | 113654 | + | 13 |
| 5 | VVTPLSWRPPRLPRWSPRLPRCTRRAT | WSPTWWARSPRSRTRASTWPAGPARSC | 113460 | 113625 | + | 6 |
| 6 | LTADSWPTRSVTMLPAAYSAVAE | ATTAAAAGVARTGALPLTRSNGRRTPSVF | 113578 | 113419 | - | 7 |
| 7 | MPGFLTADSWPTRSVTMLPAAYSAVAEA | TTAAAAGVARTGALPLTRSNGRRTPSVF | 113590 | 113419 | - | 8 |
| 8 | LMPGFLTADSWPTRSVTMLPAAYSAVAE | ATTAAAAGVARTGALPLTRSNGRRTPSVF | 113593 | 113419 | - | 7 |
| 9 | MFSSSAPDLPARLMPGF | LTADSWPTRSVTMLPAAYSAVAEAT...AAAGVARTGALPLTRSNGRRTPSVF | 113629 | 113419 | - | 8 |
| 10 | MNSSIAVCPVGLLNQPRRPSARMFSSSAPDLPARLMPGF | LTADSWPTRSVTMLPAAYSAVAEAT...AAAGVARTGALPLTRSNGRRTPSVF | 113695 | 113419 | - | 7 |
| 11 | MTSNEAGSKKHTVEY | FTAIGIALGFGAGMLFDQIIIGLAIGLAVGVILGFVVSGRK | 113825 | 113996 | + | 6 |
| 12 | MKRTTPHLDCCEPLAPEELRVSTRLRLGRDPAPPCR | SPPGDNALTSFFDSTF | 115231 | 115390 | + | 6 |
| 13 | LEPACPPPLVQTVGMRPDG | LTPPTLVVGQRDRAA | 115425 | 115530 | + | 8 |
| 14 | MTDDFDTGLAIGFEASDEGLSPALKIPRSKATSTSGSTVPRM | EPPTSSDPRSGCRTRR | 115578 | 115755 | + | 6 |

### Results for SFB47920.1 [Amycolatopsis marina] back to top

previous - next

#### Architecture

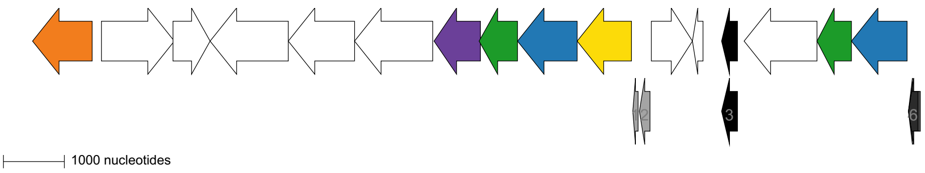

Link to nucleotide sequence

| Accession | start | end | direction | length (aa) | Pfam/HMM | name | description | E-value |
|---|---|---|---|---|---|---|---|---|
| SFB47846.1 | 68588 | 67619 | - | 322 | TIGR01188<br>TIGR03864<br>TIGR01288<br>TIGR03522<br>TIGR04406 | TIGR01188<br>TIGR03864<br>TIGR01288<br>TIGR03522<br>TIGR04406 | drrA: daunorubicin resistance ABC transporter, ATP-binding protein<br>PQQ_ABC_ATP: ABC transporter, ATP-binding subunit, PQQ-dependent alcohol dehydrogenase system<br>nodI: nodulation ABC transporter NodI<br>GldA_ABC_ATP: gliding motility-associated ABC transporter ATP-binding subunit GldA<br>LPS_export_lptB: LPS export ABC transporter ATP-binding protein | 8.20E-65<br>1.60E-50<br>3.90E-43<br>1.20E-37<br>6.20E-36 |
| SFB47854.1 | 68733 | 69924 | + | 396 | PF07730 | HisKA_3 | Histidine kinase | 3.50E-10 |
| SFB47864.1 | 69920 | 70523 | + | 200 | PF00072<br>TIGR02875<br>PF00196<br>TIGR02154<br>TIGR03020 | Response_reg<br>TIGR02875<br>GerE<br>TIGR02154<br>TIGR03020 | Response regulator receiver domain<br>spore_0_A: sporulation transcription factor Spo0A<br>Bacterial regulatory proteins, luxR family<br>PhoB: phosphate regulon transcriptional regulatory protein PhoB<br>EpsA: transcriptional regulator EpsA | 1.50E-21<br>3.10E-16<br>7.60E-11<br>4.90E-07<br>3.60E-06 |
| SFB47872.1 | 71795 | 70529 | - | 421 | TIGR01426<br>PF06722<br>PF00201<br>PF04101 | TIGR01426<br>DUF1205<br>UDPGT<br>Glyco_tran_28_C | MGT: glycosyltransferase, MGT family<br>Protein of unknown function (DUF1205)<br>UDP-glucoronosyl and UDP-glucosyl transferase<br>Glycosyltransferase family 28 C-terminal domain | 5.60E-28<br>4.20E-26<br>9.00E-07<br>3.10E-04 |
| SFB47881.1 | 72901 | 71809 | - | 363 | NO PFAM MATCH | - | - | - |
| SFB47891.1 | 74175 | 72888 | - | 428 | TIGR01426<br>PF03033<br>PF00201 | TIGR01426<br>Glyco_transf_28<br>UDPGT | MGT: glycosyltransferase, MGT family<br>Glycosyltransferase family 28 N-terminal domain<br>UDP-glucoronosyl and UDP-glucosyl transferase | 5.20E-18<br>7.20E-15<br>7.70E-05 |
| SFB47899.1 | 74955 | 74199 | - | 251 | PF13489<br>PF13649<br>PF08242<br>TIGR02021<br>PF08241 | Methyltransf_23<br>Methyltransf_25<br>Methyltransf_12<br>TIGR02021<br>Methyltransf_11 | Methyltransferase domain<br>Methyltransferase domain<br>Methyltransferase domain<br>BchM-ChlM: magnesium protoporphyrin O-methyltransferase<br>Methyltransferase domain | 1.20E-10<br>1.70E-10<br>8.20E-08<br>1.40E-07<br>3.30E-07 |
| SFB47906.1 | 75572 | 74939 | - | 210 | LinL<br>LinG | LinL<br>LinG | LinL<br>LinG Homolog | 1.40E-72<br>6.70E-07 |
| SFB47920.1 | 76540 | 75568 | - | 323 | LinE<br>LinH<br>TIGR02427<br>PF12697<br>PF00561 | LinE<br>LinH<br>TIGR02427<br>Abhydrolase_6<br>Abhydrolase_1 | LinE Homolog<br>LinH (Fusion of LinE/LinG)<br>protocat_pcaD: 3-oxoadipate enol-lactonase<br>Alpha/beta hydrolase family<br>alpha/beta hydrolase fold | 1.80E-96<br>4.90E-76<br>5.40E-15<br>4.90E-12<br>1.50E-10 |
| SFB47930.1 | 77433 | 76536 | - | 298 | LinG<br>LinH<br>LinL<br>TIGR04033<br>PF06762 | LinG<br>LinH<br>LinL<br>TIGR04033<br>LMF1 | LinG Homolog<br>LinH (Fusion of LinE/LinG)<br>LinL<br>export_SdpB: antimicrobial peptide system protein, SdpB family<br>Lipase maturation factor | 3.60E-102<br>7.80E-94<br>2.40E-16<br>8.90E-05<br>9.10E-04 |
| SFB47940.1 | 77758 | 78427 | + | 222 | PF00196<br>TIGR03020<br>TIGR03541<br>PF13384<br>PF08281 | GerE<br>TIGR03020<br>TIGR03541<br>HTH_23<br>Sigma70_r4_2 | Bacterial regulatory proteins, luxR family<br>EpsA: transcriptional regulator EpsA<br>reg_near_HchA: LuxR family transcriptional regulatory, chaperone HchA-associated<br>Homeodomain-like domain<br>Sigma-70, region 4 | 6.10E-19<br>4.90E-13<br>9.90E-10<br>2.30E-05<br>1.00E-04 |
| SFB47949.1 | 78607 | 78433 | - | 57 | PF04306 | DUF456 | Protein of unknown function (DUF456) | 2.30E-05 |
| SFB47958.1 | 79174 | 78913 | - | 86 | LinA | LinA | LinA (linaridin precursor) | 8.20E-14 |
| SFB47967.1 | 80476 | 79276 | - | 399 | PF04820<br>TIGR02032<br>PF01494<br>PF01266<br>TIGR03197 | Trp_halogenase<br>TIGR02032<br>FAD_binding_3<br>DAO<br>TIGR03197 | Tryptophan halogenase<br>GG-red-SF: geranylgeranyl reductase family<br>FAD binding domain<br>FAD dependent oxidoreductase<br>MnmC_Cterm: tRNA U-34 5-methylaminomethyl-2-thiouridine biosynthesis protein MnmC, C-terminal domain | 1.30E-17<br>1.10E-14<br>4.60E-10<br>6.10E-10<br>2.70E-08 |
| SFB47976.1 | 81054 | 80475 | - | 192 | LinL | LinL | LinL | 2.40E-62 |
| SFB47982.1 | 81968 | 81050 | - | 305 | LinE<br>LinH<br>TIGR02427<br>PF12697<br>PF00561 | LinE<br>LinH<br>TIGR02427<br>Abhydrolase_6<br>Abhydrolase_1 | LinE Homolog<br>LinH (Fusion of LinE/LinG)<br>protocat_pcaD: 3-oxoadipate enol-lactonase<br>Alpha/beta hydrolase family<br>alpha/beta hydrolase fold | 7.40E-90<br>4.40E-66<br>7.70E-14<br>3.00E-13<br>3.30E-12 |

| index | leader | core | start | end | dir | score |
|---|---|---|---|---|---|---|
| 1 | MPLIDGRVLSVAGCRF | VARSCGTAAPVRGDDHR | 77550 | 77448 | - | 6 |
| 2 | LALQDFAANAYPRPGTCAFSRCNTGIMAAQP | LTVTHLLWLWSSRRSRIRCPLSG | 77730 | 77565 | - | 6 |
| 3 | MNGIERAAQAAVPENGGTLRPLERANGTAPVL | GTPALVTAAGVLYVAGNMVTDLVGQ...TGKVRGLSGKSGAELLNIRAAGLRE | 79174 | 78913 | - | 26 |
| 4 | VVLEDGRVRETGTHRQLLRTSPTYQRL | TSIQFHADQTNDVSLTGRDSLPAVPGGH | 82141 | 81973 | - | 7 |
| 5 | VVDADKIVVLEDGRVRETGTHRQLLRTSPTY | QRLTSIQFHADQTNDVSLTGRDSLPAVPGGH | 82162 | 81973 | - | 6 |
| 6 | MSTVVDADKIVVLEDGRVRETGTHRQLLRTSP | TYQRLTSIQFHADQTNDVSLTGRDSLPAVPGGH | 82171 | 81973 | - | 7 |

### Results for SFB47982.1 [Amycolatopsis marina] back to top

previous - next

#### Architecture

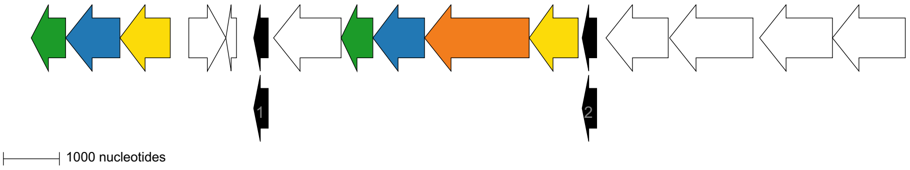

Link to nucleotide sequence

| Accession | start | end | direction | length (aa) | Pfam/HMM | name | description | E-value |
|---|---|---|---|---|---|---|---|---|
| SFB47906.1 | 75572 | 74939 | - | 210 | LinL | LinL | LinL | 1.40E-72 |
| | | | | | LinG | LinG | LinG Homolog | 6.70E-07 |
| SFB47920.1 | 76540 | 75568 | - | 323 | LinE | LinE | LinE Homolog | 1.80E-96 |
| | | | | | LinH | LinH | LinH (Fusion of LinE/LinG) | 4.90E-76 |
| | | | | | TIGR02427 | TIGR02427 | protocat_pcaD: 3-oxoadipate enol-lactonase | 5.40E-15 |
| | | | | | PF12697 | Abhydrolase_6 | Alpha/beta hydrolase family | 4.90E-12 |
| | | | | | PF00561 | Abhydrolase_1 | alpha/beta hydrolase fold | 1.50E-10 |
| SFB47930.1 | 77433 | 76536 | - | 298 | LinG | LinG | LinG Homolog | 3.60E-102 |
| | | | | | LinH | LinH | LinH (Fusion of LinE/LinG) | 7.80E-94 |
| | | | | | LinL | LinL | LinL | 2.40E-16 |
| | | | | | TIGR04033 | TIGR04033 | export_SdpB: antimicrobial peptide system protein, SdpB family | 8.90E-05 |
| | | | | | PF06762 | LMF1 | Lipase maturation factor | 9.10E-04 |
| SFB47940.1 | 77758 | 78427 | + | 222 | PF00196 | GerE | Bacterial regulatory proteins, luxR family | 6.10E-19 |
| | | | | | TIGR03020 | TIGR03020 | EpsA: transcriptional regulator EpsA | 4.90E-13 |
| | | | | | TIGR03541 | TIGR03541 | reg_near_HchA: LuxR family transcriptional regulatory, chaperone HchA-associated | 9.90E-10 |
| | | | | | PF13384 | HTH_23 | Homeodomain-like domain | 2.30E-05 |
| | | | | | PF08281 | Sigma70_r4_2 | Sigma-70, region 4 | 1.00E-04 |
| SFB47949.1 | 78607 | 78433 | - | 57 | PF04306 | DUF456 | Protein of unknown function (DUF456) | 2.30E-05 |
| SFB47958.1 | 79174 | 78913 | - | 86 | LinA | LinA | LinA (linaridin precursor) | 8.20E-14 |
| SFB47967.1 | 80476 | 79276 | - | 399 | PF04820 | Trp_halogenase | Tryptophan halogenase | 1.30E-17 |
| | | | | | TIGR02032 | TIGR02032 | GG-red-SF: geranylgeranyl reductase family | 1.10E-14 |
| | | | | | PF01494 | FAD_binding_3 | FAD binding domain | 4.60E-10 |
| | | | | | PF01266 | DAO | FAD dependent oxidoreductase | 6.10E-10 |
| | | | | | TIGR03197 | TIGR03197 | MnmC_Cterm: tRNA U-34 5-methylaminomethyl-2-thiouridine biosynthesis protein MnmC, C-terminal domain | 2.70E-08 |
| SFB47976.1 | 81054 | 80475 | - | 192 | LinL | LinL | LinL | 2.40E-62 |
| SFB47982.1 | 81968 | 81050 | - | 305 | LinE | LinE | LinE Homolog | 7.40E-90 |
| | | | | | LinH | LinH | LinH (Fusion of LinE/LinG) | 4.40E-66 |
| | | | | | TIGR02427 | TIGR02427 | protocat_pcaD: 3-oxoadipate enol-lactonase | 7.70E-14 |
| | | | | | PF12697 | Abhydrolase_6 | Alpha/beta hydrolase family | 3.00E-13 |
| | | | | | PF00561 | Abhydrolase_1 | alpha/beta hydrolase fold | 3.30E-12 |
| SFB47990.1 | 83833 | 81973 | - | 619 | TIGR02204 | TIGR02204 | MsbA_rel: ABC transporter, permease/ATP-binding protein | 1.00E-105 |
| | | | | | TIGR02203 | TIGR02203 | MsbA_lipidA: lipid A export permease/ATP-binding protein MsbA | 1.20E-96 |
| | | | | | TIGR00958 | TIGR00958 | 3a01208: antigen peptide transporter 2 | 1.00E-92 |
| | | | | | TIGR01846 | TIGR01846 | type_I_sec_HlyB: type I secretion system ATPase | 6.20E-82 |
| | | | | | TIGR03796 | TIGR03796 | NHLM_micro_ABC1: NHLM bacteriocin system ABC transporter, peptidase/ATP-binding protein | 2.20E-81 |
| SFB48000.1 | 84708 | 83835 | - | 290 | LinG | LinG | LinG Homolog | 1.50E-103 |
| | | | | | LinH | LinH | LinH (Fusion of LinE/LinG) | 3.80E-90 |
| | | | | | PF06762 | LMF1 | Lipase maturation factor | 5.70E-04 |
| SFB48010.1 | 85048 | 84787 | - | 86 | LinA | LinA | LinA (linaridin precursor) | 1.20E-13 |
| SFB48018.1 | 86316 | 85224 | - | 363 | PF08028 | Acyl-CoA_dh_2 | Acyl-CoA dehydrogenase, C-terminal domain | 3.80E-19 |
| | | | | | TIGR04022 | TIGR04022 | sulfur_SfnB: sulfur acquisition oxidoreductase, SfnB family | 3.00E-09 |
| | | | | | PF00441 | Acyl-CoA_dh_1 | Acyl-CoA dehydrogenase, C-terminal domain | 1.20E-06 |
| SFB48027.1 | 87848 | 86339 | - | 502 | TIGR04022 | TIGR04022 | sulfur_SfnB: sulfur acquisition oxidoreductase, SfnB family | 2.60E-17 |
| | | | | | PF08028 | Acyl-CoA_dh_2 | Acyl-CoA dehydrogenase, C-terminal domain | 7.90E-14 |
| | | | | | PF10604 | Polyketide_cyc2 | Polyketide cyclase / dehydrase and lipid transport | 4.50E-05 |
| | | | | | PF02771 | Acyl-CoA_dh_N | Acyl-CoA dehydrogenase, N-terminal domain | 2.30E-04 |
| | | | | | PF00441 | Acyl-CoA_dh_1 | Acyl-CoA dehydrogenase, C-terminal domain | 7.60E-04 |
| SFB48037.1 | 89263 | 87955 | - | 435 | PF05193 | Peptidase_M16_C | Peptidase M16 inactive domain | 5.70E-20 |
| | | | | | PF00675 | Peptidase_M16 | Insulinase (Peptidase family M16) | 1.10E-04 |
| SFB48046.1 | 90549 | 89259 | - | 429 | PF05193 | Peptidase_M16_C | Peptidase M16 inactive domain | 3.60E-23 |
| | | | | | PF00675 | Peptidase_M16 | Insulinase (Peptidase family M16) | 1.60E-20 |
| | | | | | TIGR02110 | TIGR02110 | PQQ_syn_pqqF: coenzyme PQQ biosynthesis protein PqqF | 5.50E-12 |

| index | leader | core | start | end | dir | score |
|---|---|---|---|---|---|---|
| 1 | MNGIERAAQAAVPENGGTLRPLERANGTAPVL | GTPALVTAAGVLYVAGNMVTDLVGQ...TGKVRGLSGKSGAELLNIRAAGLRE | 79174 | 78913 | - | 25 |
| 2 | MSNVEAFANQNADGVLNPLQVTSAVGPVL | ATPAYAGYAAGAGVVFTLGLISDAV...VSPTGPVPSYGSVDELLGMRVASIN | 85048 | 84787 | - | 28 |

### Results for WP_116048497.1 [Amycolatopsis palatopharyngis] back to top

previous - next

#### Architecture

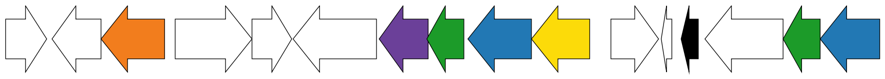

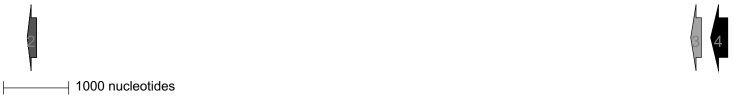

[Link to nucleotide sequence]

| Accession | start | end | direction | length (aa) | Pfam/HMM | name | description | E-value |
|---|---|---|---|---|---|---|---|---|
| WP_162830187.1 | 45380 | 46001 | + | 206 | PF00196<br>TIGR03020<br>TIGR03541<br>PF08281<br>PF13384 | GerE<br>TIGR03020<br>TIGR03541<br>Sigma70_r4_2<br>HTH_23 | Bacterial regulatory proteins, luxR family<br>EpsA: transcriptional regulator EpsA<br>reg_near_HchA: LuxR family transcriptional regulatory, chaperone HchA-associated<br>Sigma-70, region 4<br>Homeodomain-like domain | 1.10E-18<br>2.00E-10<br>3.00E-08<br>6.20E-06<br>2.00E-05 |
| WP_116048480.1 | 46840 | 46084 | - | 251 | PF01061<br>PF12698 | ABC2_membrane<br>ABC2_membrane_3 | ABC-2 type transporter<br>ABC-2 family transporter protein | 1.70E-06<br>2.00E-04 |
| WP_116048482.1 | 47817 | 46836 | - | 326 | TIGR01188<br>TIGR03864<br>TIGR01288<br>TIGR03522<br>TIGR04406 | TIGR01188<br>TIGR03864<br>TIGR01288<br>TIGR03522<br>TIGR04406 | drrA: daunorubicin resistance ABC transporter, ATP-binding protein<br>PQQ_ABC_ATP: ABC transporter, ATP-binding subunit, PQQ-dependent alcohol dehydrogenase system<br>nodI: nodulation ABC transporter NodI<br>GldA_ABC_ATP: gliding motility-associated ABC transporter ATP-binding subunit GldA<br>LPS_export_lptB: LPS export ABC transporter ATP-binding protein | 1.20E-64<br>9.20E-50<br>5.20E-43<br>3.80E-36<br>1.20E-34 |
| WP_162830188.1 | 47962 | 49150 | + | 395 | PF07730 | HisKA_3 | Histidine kinase | 5.20E-11 |
| WP_116048487.1 | 49149 | 49752 | + | 200 | PF00072<br>TIGR02875<br>PF00196<br>TIGR02154<br>TIGR03020 | Response_reg<br>TIGR02875<br>GerE<br>TIGR02154<br>TIGR03020 | Response regulator receiver domain<br>spore_0_A: sporulation transcription factor Spo0A<br>Bacterial regulatory proteins, luxR family<br>PhoB: phosphate regulon transcriptional regulatory protein PhoB<br>EpsA: transcriptional regulator EpsA | 2.20E-21<br>1.90E-15<br>5.50E-11<br>5.60E-07<br>7.50E-07 |
| WP_116048490.1 | 51061 | 49777 | - | 427 | TIGR01426<br>PF03033<br>PF00201 | TIGR01426<br>Glyco_transf_28<br>UDPGT | MGT: glycosyltransferase, MGT family<br>Glycosyltransferase family 28 N-terminal domain<br>UDP-glucoronosyl and UDP-glucosyl transferase | 9.00E-20<br>2.30E-14<br>3.60E-04 |
| WP_116048492.1 | 51836 | 51086 | - | 249 | PF13649<br>PF13489<br>PF08242<br>PF08241<br>TIGR02021 | Methyltransf_25<br>Methyltransf_23<br>Methyltransf_12<br>Methyltransf_11<br>TIGR02021 | Methyltransferase domain<br>Methyltransferase domain<br>Methyltransferase domain<br>Methyltransferase domain<br>BchM-ChlM: magnesium protoporphyrin O-methyltransferase | 6.90E-12<br>8.80E-12<br>1.50E-08<br>2.30E-08<br>4.00E-07 |
| WP_170161355.1 | 52390 | 51820 | - | 189 | LinL<br>LinG | LinL<br>LinG | LinL<br>LinG Homolog | 8.20E-72<br>6.40E-07 |
| WP_116048497.1 | 53418 | 52443 | - | 324 | LinE<br>LinH<br>TIGR02427<br>PF12697<br>PF00561 | LinE<br>LinH<br>TIGR02427<br>Abhydrolase_6<br>Abhydrolase_1 | LinE Homolog<br>LinH (Fusion of LinE/LinG)<br>protocat_pcaD: 3-oxoadipate enol-lactonase<br>Alpha/beta hydrolase family<br>alpha/beta hydrolase fold | 1.20E-98<br>9.00E-74<br>1.70E-14<br>8.20E-12<br>2.30E-10 |
| WP_116048500.1 | 54311 | 53414 | - | 298 | LinG<br>LinH<br>LinL<br>TIGR04033<br>PF06762 | LinG<br>LinH<br>LinL<br>TIGR04033<br>LMF1 | LinG Homolog<br>LinH (Fusion of LinE/LinG)<br>LinL<br>export_SdpB: antimicrobial peptide system protein, SdpB family<br>Lipase maturation factor | 1.40E-102<br>2.30E-95<br>1.00E-15<br>1.10E-05<br>1.30E-04 |
| WP_116048503.1 | 54632 | 55379 | + | 248 | PF00196<br>TIGR03020<br>TIGR03541<br>PF13384 | GerE<br>TIGR03020<br>TIGR03541<br>HTH_23 | Bacterial regulatory proteins, luxR family<br>EpsA: transcriptional regulator EpsA<br>reg_near_HchA: LuxR family transcriptional regulatory, chaperone HchA-associated<br>Homeodomain-like domain | 6.90E-17<br>4.00E-12<br>7.40E-09<br>1.10E-04 |
| WP_116048505.1 | 55575 | 55404 | - | 56 | PF04306 | DUF456 | Protein of unknown function (DUF456) | 2.70E-05 |
| WP_116048508.1 | 55977 | 55716 | - | 86 | LinA | LinA | LinA (linaridin precursor) | 5.90E-14 |
| WP_116048510.1 | 57272 | 56072 | - | 399 | TIGR02032<br>PF04820<br>PF01266<br>PF13450<br>TIGR03197 | TIGR02032<br>Trp_halogenase<br>DAO<br>NAD_binding_8<br>TIGR03197 | GG-red-SF: geranylgeranyl reductase family<br>Tryptophan halogenase<br>FAD dependent oxidoreductase<br>NAD(P)-binding Rossmann-like domain<br>MnmC_Cterm: tRNA U-34 5-methylaminomethyl-2-thiouridine biosynthesis protein MnmC, C-terminal domain | 1.30E-17<br>5.80E-17<br>1.80E-10<br>4.30E-07<br>4.50E-07 |
| WP_116048513.1 | 57850 | 57271 | - | 192 | LinL | LinL | LinL | 1.10E-62 |
| WP_116048516.1 | 58758 | 57846 | - | 303 | LinE<br>LinH<br>TIGR02427<br>PF12697<br>PF00561 | LinE<br>LinH<br>TIGR02427<br>Abhydrolase_6<br>Abhydrolase_1 | LinE Homolog<br>LinH (Fusion of LinE/LinG)<br>protocat_pcaD: 3-oxoadipate enol-lactonase<br>Alpha/beta hydrolase family<br>alpha/beta hydrolase fold | 1.20E-91<br>3.60E-64<br>1.20E-14<br>1.70E-13<br>3.00E-13 |

| index | leader | core | start | end | dir | score |
|---|---|---|---|---|---|---|
| 1 | LRSIAWIQDNRCSSPSNLPES | STNPMFTIESPVSCVVTQRVGD | 45391 | 45259 | - | 7 |
| 2 | MWNLLRSIAWIQDNRCSSPSNLP | ESSTNPMFTIESPVSCVVTQRVGD | 45403 | 45259 | - | 6 |
| 3 | MNSKETGARK | HTVEYLTSIGALVGFVTGMLFDQILVGLVVGMALGALIGFIMQKRG | 55575 | 55404 | - | 6 |
| 4 | MNGIERAAQAAVPENGGALRPLERANGTAPVL | GTPALVTAAGVLYVAGNMVTDLVGQ...NSKMRGLSGKSGAELLSIRTAGLRE | 55977 | 55716 | - | 26 |

### Results for WP_116048516.1 [Amycolatopsis palatopharyngis] back to top

previous - next

#### Architecture

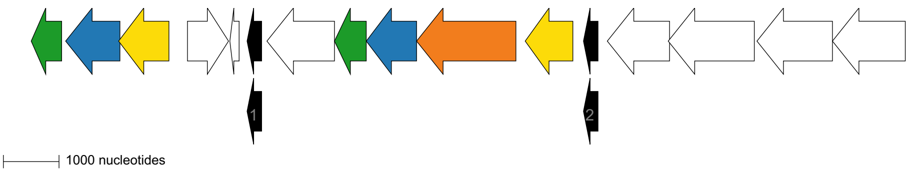

Link to nucleotide sequence

| Accession | start | end | direction | length (aa) | Pfam/HMM | name | description | E-value |
|---|---|---|---|---|---|---|---|---|
| WP_170161355.1 | 52390 | 51820 | - | 189 | LinL<br>LinG | LinL<br>LinG | LinL<br>LinG Homolog | 8.20E-72<br>6.40E-07 |
| WP_116048497.1 | 53418 | 52443 | - | 324 | LinE<br>LinH<br>TIGR02427<br>PF12697<br>PF00561 | LinE<br>LinH<br>TIGR02427<br>Abhydrolase_6<br>Abhydrolase_1 | LinE Homolog<br>LinH (Fusion of LinE/LinG)<br>protocat_pcaD: 3-oxoadipate enol-lactonase<br>Alpha/beta hydrolase family<br>alpha/beta hydrolase fold | 1.20E-98<br>9.00E-74<br>1.70E-14<br>8.20E-12<br>2.30E-10 |
| WP_116048500.1 | 54311 | 53414 | - | 298 | LinG<br>LinH<br>LinL<br>TIGR04033<br>PF06762 | LinG<br>LinH<br>LinL<br>TIGR04033<br>LMF1 | LinG Homolog<br>LinH (Fusion of LinE/LinG)<br>LinL<br>export_SdpB: antimicrobial peptide system protein, SdpB family<br>Lipase maturation factor | 1.40E-102<br>2.30E-95<br>1.00E-15<br>1.10E-05<br>1.30E-04 |
| WP_116048503.1 | 54632 | 55379 | + | 248 | PF00196<br>TIGR03020<br>TIGR03541<br>PF13384 | GerE<br>TIGR03020<br>TIGR03541<br>HTH_23 | Bacterial regulatory proteins, luxR family<br>EpsA: transcriptional regulator EpsA<br>reg_near_HchA: LuxR family transcriptional regulatory, chaperone HchA-associated<br>Homeodomain-like domain | 6.90E-17<br>4.00E-12<br>7.40E-09<br>1.10E-04 |
| WP_116048505.1 | 55575 | 55404 | - | 56 | PF04306 | DUF456 | Protein of unknown function (DUF456) | 2.70E-05 |
| WP_116048508.1 | 55977 | 55716 | - | 86 | LinA | LinA | LinA (linaridin precursor) | 5.90E-14 |
| WP_116048510.1 | 57272 | 56072 | - | 399 | TIGR02032<br>PF04820<br>PF01266<br>PF13450<br>TIGR03197 | TIGR02032<br>Trp_halogenase<br>DAO<br>NAD_binding_8<br>TIGR03197 | GG-red-SF: geranylgeranyl reductase family<br>Tryptophan halogenase<br>FAD dependent oxidoreductase<br>NAD(P)-binding Rossmann-like domain<br>MnmC_Cterm: tRNA U-34 5-methylaminomethyl-2-thiouridine biosynthesis protein MnmC, C-terminal domain | 1.30E-17<br>5.80E-17<br>1.80E-10<br>4.30E-07<br>4.50E-07 |
| WP_116048513.1 | 57850 | 57271 | - | 192 | LinL | LinL | LinL | 1.10E-62 |
| WP_116048516.1 | 58758 | 57846 | - | 303 | LinE<br>LinH<br>TIGR02427<br>PF12697<br>PF00561 | LinE<br>LinH<br>TIGR02427<br>Abhydrolase_6<br>Abhydrolase_1 | LinE Homolog<br>LinH (Fusion of LinE/LinG)<br>protocat_pcaD: 3-oxoadipate enol-lactonase<br>Alpha/beta hydrolase family<br>alpha/beta hydrolase fold | 1.20E-91<br>3.60E-64<br>1.20E-14<br>1.70E-13<br>3.00E-13 |
| WP_116048571.1 | 60554 | 58763 | - | 596 | TIGR02204<br>TIGR02203<br>TIGR00958<br>TIGR03796<br>TIGR01846 | TIGR02204<br>TIGR02203<br>TIGR00958<br>TIGR03796<br>TIGR01846 | MsbA_rel: ABC transporter, permease/ATP-binding protein<br>MsbA_lipidA: lipid A export permease/ATP-binding protein MsbA<br>3a01208: antigen peptide transporter 2<br>NHLM_micro_ABC1: NHLM bacteriocin system ABC transporter, peptidase/ATP-binding protein<br>type_I_sec_HlyB: type I secretion system ATPase | 1.00E-101<br>1.40E-91<br>2.70E-86<br>1.10E-78<br>8.90E-77 |
| WP_116048519.1 | 61579 | 60712 | - | 288 | LinG<br>LinH | LinG<br>LinH | LinG Homolog<br>LinH (Fusion of LinE/LinG) | 1.50E-104<br>4.50E-90 |
| WP_116048522.1 | 62013 | 61752 | - | 86 | LinA | LinA | LinA (linaridin precursor) | 1.40E-13 |
| WP_162830189.1 | 63299 | 62189 | - | 369 | PF08028<br>TIGR04022<br>PF00441 | Acyl-CoA_dh_2<br>TIGR04022<br>Acyl-CoA_dh_1 | Acyl-CoA dehydrogenase, C-terminal domain<br>sulfur_SfnB: sulfur acquisition oxidoreductase, SfnB family<br>Acyl-CoA dehydrogenase, C-terminal domain | 9.00E-20<br>1.40E-11<br>7.40E-06 |
| WP_116048527.1 | 64818 | 63291 | - | 508 | TIGR04022<br>PF08028<br>PF02771 | TIGR04022<br>Acyl-CoA_dh_2<br>Acyl-CoA_dh_N | sulfur_SfnB: sulfur acquisition oxidoreductase, SfnB family<br>Acyl-CoA dehydrogenase, C-terminal domain<br>Acyl-CoA dehydrogenase, N-terminal domain | 9.60E-15<br>9.50E-12<br>4.00E-05 |
| WP_162830190.1 | 66229 | 64885 | - | 447 | PF05193<br>PF00675 | Peptidase_M16_C<br>Peptidase_M16 | Peptidase M16 inactive domain<br>Insulinase (Peptidase family M16) | 1.60E-15<br>1.60E-06 |
| WP_116048573.1 | 67530 | 66225 | - | 434 | PF00675<br>PF05193<br>TIGR02110 | Peptidase_M16<br>Peptidase_M16_C<br>TIGR02110 | Insulinase (Peptidase family M16)<br>Peptidase M16 inactive domain<br>PQQ_syn_pqqF: coenzyme PQQ biosynthesis protein PqqF | 1.60E-21<br>3.50E-19<br>4.90E-14 |

| index | leader | core | start | end | dir | score |
|---|---|---|---|---|---|---|
| 1 | MNGIERAAQAAVPENGGALRPLERANGTAPVL | GTPALVTAAGVLYVAGNMVTDLVGQ...NSKMRGLSGKSGAELLSIRTAGLRE | 55977 | 55716 | - | 25 |
| 2 | MSNVEAFANQNVDGVLNPLQVTSAVGPVL | ATPAYAGYAAGAGVVFTLGLISDAV...VSPTGPVPSYGSVDELLGMRVASIS | 62013 | 61752 | - | 28 |

### Results for SEC50923.1 [Amycolatopsis tolypomycina] back to top

previous - next

#### Architecture

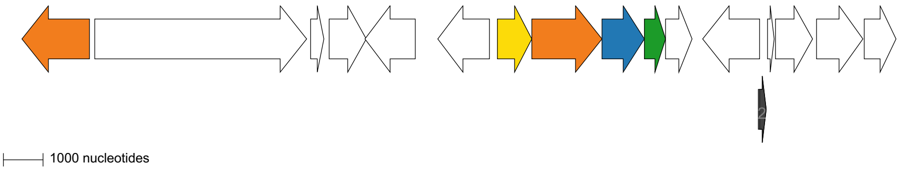

Link to nucleotide sequence

| Accession | start | end | direction | length (aa) | Pfam/HMM | name | description | E-value |
|---|---|---|---|---|---|---|---|---|
| SEC50700.1 | 2022138 | 2020458 | - | 559 | TIGR02203 | TIGR02203 | MsbA_lipidA: lipid A export permease/ATP-binding protein MsbA | 1.60E-93 |
| | | | | | TIGR02204 | TIGR02204 | MsbA_rel: ABC transporter, permease/ATP-binding protein | 9.10E-88 |
| | | | | | TIGR01846 | TIGR01846 | type_I_sec_HlyB: type I secretion system ATPase | 1.80E-79 |
| | | | | | TIGR02857 | TIGR02857 | CydD: thiol reductant ABC exporter, CydD subunit | 5.80E-78 |
| | | | | | TIGR00958 | TIGR00958 | 3a01208: antigen peptide transporter 2 | 3.40E-74 |
| SEC50725.1 | 2022268 | 2027617 | + | 1782 | TIGR01733 | TIGR01733 | AA-adenyl-dom: amino acid adenylation domain | 8.00E-124 |
| | | | | | PF00501 | AMP-binding | AMP-binding enzyme | 1.40E-94 |
| | | | | | TIGR01734 | TIGR01734 | D-ala-DACP-lig: D-alanine--poly(phosphoribitol) ligase, subunit 1 | 1.60E-76 |
| | | | | | TIGR03098 | TIGR03098 | ligase_PEP_1: acyl-CoA ligase (AMP-forming), exosortase A-associated | 8.00E-63 |
| | | | | | TIGR03443 | TIGR03443 | alpha_am_amid: L-aminoadipate-semialdehyde dehydrogenase | 1.30E-51 |
| SEC50755.1 | 2027736 | 2028039 | + | 100 | PF03459 | TOBE | TOBE domain | 1.90E-04 |
| SEC50787.1 | 2028175 | 2029108 | + | 310 | TIGR03945 | TIGR03945 | PLP_SbnA_fam: 2,3-diaminopropionate biosynthesis protein SbnA | 2.20E-111 |
| | | | | | TIGR01136 | TIGR01136 | cysKM: cysteine synthase | 6.80E-61 |
| | | | | | TIGR01139 | TIGR01139 | cysK: cysteine synthase A | 1.00E-52 |
| | | | | | PF00291 | PALP | Pyridoxal-phosphate dependent enzyme | 1.50E-51 |
| | | | | | TIGR01138 | TIGR01138 | cysM: cysteine synthase B | 3.10E-50 |
| SEC50817.1 | 2030358 | 2029083 | - | 424 | PF05193 | Peptidase_M16_C | Peptidase M16 inactive domain | 1.20E-23 |
| SEC50839.1 | 2032243 | 2030935 | - | 435 | TIGR02110 | TIGR02110 | PQQ_syn_pqqF: coenzyme PQQ biosynthesis protein PqqF | 8.10E-20 |
| | | | | | PF05193 | Peptidase_M16_C | Peptidase M16 inactive domain | 2.40E-19 |
| | | | | | PF00675 | Peptidase_M16 | Insulinase (Peptidase family M16) | 5.50E-19 |
| SEC50872.1 | 2032429 | 2033287 | + | 285 | LinG | LinG | LinG Homolog | 8.30E-98 |
| | | | | | LinH | LinH | LinH (Fusion of LinE/LinG) | 5.70E-90 |
| | | | | | TIGR04033 | TIGR04033 | export_SdpB: antimicrobial peptide system protein, SdpB family | 3.10E-04 |
| SEC50889.1 | 2033288 | 2035058 | + | 589 | TIGR02204 | TIGR02204 | MsbA_rel: ABC transporter, permease/ATP-binding protein | 1.20E-99 |
| | | | | | TIGR02203 | TIGR02203 | MsbA_lipidA: lipid A export permease/ATP-binding protein MsbA | 3.00E-92 |
| | | | | | TIGR00958 | TIGR00958 | 3a01208: antigen peptide transporter 2 | 1.40E-82 |
| | | | | | TIGR01846 | TIGR01846 | type_I_sec_HlyB: type I secretion system ATPase | 1.90E-82 |
| | | | | | TIGR03796 | TIGR03796 | NHLM_micro_ABC1: NHLM bacteriocin system ABC transporter, peptidase/ATP-binding protein | 2.30E-77 |
| SEC50923.1 | 2035054 | 2036128 | + | 357 | LinE | LinE | LinE Homolog | 1.40E-98 |
| | | | | | LinH | LinH | LinH (Fusion of LinE/LinG) | 8.80E-67 |
| | | | | | TIGR02427 | TIGR02427 | protocat_pcaD: 3-oxoadipate enol-lactonase | 1.80E-14 |
| | | | | | PF12697 | Abhydrolase_6 | Alpha/beta hydrolase family | 2.60E-14 |
| | | | | | PF00561 | Abhydrolase_1 | alpha/beta hydrolase fold | 6.10E-12 |
| SEC50939.1 | 2036124 | 2036676 | + | 183 | LinL | LinL | LinL | 3.00E-70 |
| SEC50972.1 | 2036677 | 2037331 | + | 217 | PF00196 | GerE | Bacterial regulatory proteins, luxR family | 3.50E-19 |
| | | | | | TIGR03020 | TIGR03020 | EpsA: transcriptional regulator EpsA | 2.70E-15 |
| | | | | | TIGR03541 | TIGR03541 | reg_near_HchA: LuxR family transcriptional regulatory, chaperone HchA-associated | 1.30E-09 |
| | | | | | PF08281 | Sigma70_r4_2 | Sigma-70, region 4 | 3.50E-07 |
| | | | | | PF13384 | HTH_23 | Homeodomain-like domain | 3.30E-06 |
| SEC51004.1 | 2039030 | 2037602 | - | 475 | TIGR02734 | TIGR02734 | crtI_fam: phytoene desaturase | 8.70E-12 |
| | | | | | TIGR03467 | TIGR03467 | HpnE: squalene-associated FAD-dependent desaturase | 7.10E-11 |
| | | | | | PF13450 | NAD_binding_8 | NAD(P)-binding Rossmann-like domain | 2.70E-10 |
| | | | | | TIGR01317 | TIGR01317 | GOGAT_sm_gam: glutamate synthase, NADH/NADPH, small subunit | 1.20E-06 |
| | | | | | TIGR03315 | TIGR03315 | Se_ygfK: putative selenate reductase, YgfK subunit | 3.50E-06 |
| SEC51026.1 | 2039251 | 2039398 | + | 48 | NO PFAM MATCH | - | - | - |
| SEC51056.1 | 2039441 | 2040386 | + | 314 | TIGR02957 | TIGR02957 | SigX4: RNA polymerase sigma-70 factor, TIGR02957 family | 1.30E-100 |
| | | | | | TIGR02960 | TIGR02960 | SigX5: RNA polymerase sigma-70 factor, TIGR02960 family | 2.80E-29 |
| | | | | | TIGR02937 | TIGR02937 | sigma70-ECF: RNA polymerase sigma factor, sigma-70 family | 3.60E-18 |
| | | | | | TIGR02985 | TIGR02985 | Sig70_bacteroi1: RNA polymerase sigma-70 factor, Bacteroides expansion family 1 | 2.20E-17 |
| | | | | | TIGR02983 | TIGR02983 | SigE-fam_strep: RNA polymerase sigma-70 factor, sigma-E family | 1.80E-15 |
| SEC51091.1 | 2040468 | 2041644 | + | 391 | PF07992 | Pyr_redox_2 | Pyridine nucleotide-disulphide oxidoreductase | 3.10E-37 |
| | | | | | TIGR03169 | TIGR03169 | Nterm_to_SelD: pyridine nucleotide-disulfide oxidoreductase family protein | 4.60E-28 |
| | | | | | TIGR02374 | TIGR02374 | nitri_red_nirB: nitrite reductase [NAD(P)H], large subunit | 5.80E-21 |
| | | | | | TIGR03385 | TIGR03385 | CoA_CoA_reduc: CoA-disulfide reductase | 2.50E-14 |
| | | | | | TIGR04369 | TIGR04369 | fusion_not_SelD: oxidoreductase/SelD-related fusion protein | 1.80E-10 |
| SEC51128.1 | 2041671 | 2042496 | + | 274 | TIGR03649 | TIGR03649 | ergot_EASG: ergot alkaloid biosynthesis protein, AFUA_2G17970 family | 6.40E-18 |
| | | | | | PF13460 | NAD_binding_10 | NAD(P)H-binding | 1.50E-14 |
| | | | | | PF01370 | Epimerase | NAD dependent epimerase/dehydratase family | 6.70E-05 |
| | | | | | TIGR01777 | TIGR01777 | yfcH: TIGR01777 family protein | 1.70E-04 |

| index | leader | core | start | end | dir | score |
|---|---|---|---|---|---|---|
| 1 | VLMVCSRSAGDHGRPVSAAGGVPRAERRRIGE | KVPRTADGPAAVVRDFPTRVRRAGRAGPRVVS | 2039022 | 2039217 | + | 7 |
| 2 | MVCSRSAGDHGRPVSAAGGVPRAERRRIGE | KVPRTADGPAAVVRDFPTRVRRAGRAGPRVVS | 2039028 | 2039217 | + | 7 |

### Results for OXM68433.1 [Amycolatopsis vastitatis] back to top

previous - next

#### Architecture

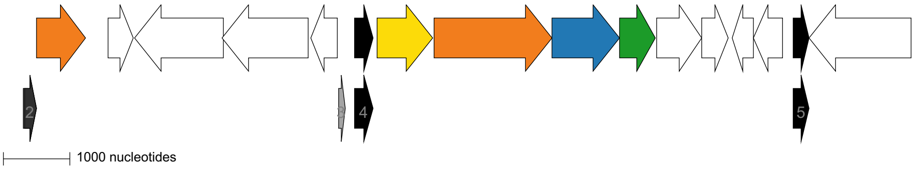

Link to nucleotide sequence

| Accession | start | end | direction | length (aa) | Pfam/HMM | name | description | E-value |
|---|---|---|---|---|---|---|---|---|
| OXM68426.1 | 111820 | 112573 | + | 250 | TIGR02633<br>TIGR02315<br>TIGR03265<br>TIGR03411<br>TIGR00968 | TIGR02633<br>TIGR02315<br>TIGR03265<br>TIGR03411<br>TIGR00968 | xylG: D-xylose ABC transporter, ATP-binding protein<br>ABC_phnC: phosphonate ABC transporter, ATP-binding protein<br>PhnT2: putative 2-aminoethylphosphonate ABC transporter, ATP-binding protein<br>urea_trans_UrtD: urea ABC transporter, ATP-binding protein UrtD<br>3a0106s01: sulfate ABC transporter, ATP-binding protein | 6.90E-48<br>2.50E-40<br>1.80E-38<br>2.00E-38<br>4.30E-38 |
| OXM68427.1 | 112904 | 113279 | + | 124 | NO PFAM MATCH | - | - | - |
| OXM68428.1 | 114631 | 113305 | - | 441 | PF05193 | Peptidase_M16_C | Peptidase M16 inactive domain | 1.60E-15 |
| OXM68556.1 | 115920 | 114627 | - | 430 | PF05193<br>PF00675<br>TIGR02110 | Peptidase_M16_C<br>Peptidase_M16<br>TIGR02110 | Peptidase M16 inactive domain<br>Insulinase (Peptidase family M16)<br>PQQ_syn_pqqF: coenzyme PQQ biosynthesis protein PqqF | 2.40E-26<br>9.70E-20<br>1.60E-18 |
| OXM68429.1 | 116347 | 115951 | - | 131 | NO PFAM MATCH | - | - | - |
| OXM68430.1 | 116624 | 116885 | + | 86 | LinA | LinA | LinA (linaridin precursor) | 2.00E-08 |
| OXM68431.1 | 116963 | 117788 | + | 274 | LinG<br>LinH | LinG<br>LinH | LinG Homolog<br>LinH (Fusion of LinE/LinG) | 3.20E-100<br>3.30E-94 |
| OXM68432.1 | 117809 | 119588 | + | 592 | TIGR02203<br>TIGR02204<br>TIGR00958<br>TIGR01846<br>TIGR03796 | TIGR02203<br>TIGR02204<br>TIGR00958<br>TIGR01846<br>TIGR03796 | MsbA_lipidA: lipid A export permease/ATP-binding protein MsbA<br>MsbA_rel: ABC transporter, permease/ATP-binding protein<br>3a01208: antigen peptide transporter 2<br>type_I_sec_HlyB: type I secretion system ATPase<br>NHLM_micro_ABC1: NHLM bacteriocin system ABC transporter, peptidase/ATP-binding protein | 4.70E-98<br>6.10E-96<br>1.00E-83<br>1.10E-76<br>7.60E-75 |
| OXM68433.1 | 119587 | 120610 | + | 340 | LinE<br>LinH<br>PF00561<br>TIGR02427<br>PF12697 | LinE<br>LinH<br>Abhydrolase_1<br>TIGR02427<br>Abhydrolase_6 | LinE Homolog<br>LinH (Fusion of LinE/LinG)<br>alpha/beta hydrolase fold<br>protocat_pcaD: 3-oxoadipate enol-lactonase<br>Alpha/beta hydrolase family | 1.10E-97<br>3.80E-76<br>1.90E-13<br>6.40E-13<br>1.20E-12 |
| OXM68434.1 | 120606 | 121143 | + | 178 | LinL | LinL | LinL | 1.10E-59 |
| OXM68435.1 | 121157 | 121832 | + | 224 | PF00196<br>TIGR03020<br>TIGR03541<br>PF04545<br>TIGR02937 | GerE<br>TIGR03020<br>TIGR03541<br>Sigma70_r4<br>TIGR02937 | Bacterial regulatory proteins, luxR family<br>EpsA: transcriptional regulator EpsA<br>reg_near_HchA: LuxR family transcriptional regulatory, chaperone HchA-associated<br>Sigma-70, region 4<br>sigma70-ECF: RNA polymerase sigma factor, sigma-70 family | 3.20E-19<br>7.70E-13<br>7.30E-09<br>1.60E-04<br>2.00E-04 |
| OXM68436.1 | 121838 | 122234 | + | 131 | PF04134 | DUF393 | Protein of unknown function, DUF393 | 2.70E-12 |
| OXM68437.1 | 122625 | 122301 | - | 107 | PF13601 | HTH_34 | Winged helix DNA-binding domain | 4.60E-24 |
| OXM68438.1 | 123053 | 122621 | - | 143 | NO PFAM MATCH | - | - | - |
| OXM68439.1 | 123208 | 123460 | + | 83 | LinA | LinA | LinA (linaridin precursor) | 7.60E-10 |
| OXM68440.1 | 124982 | 123449 | - | 510 | PF00561<br>PF08386 | Abhydrolase_1<br>Abhydrolase_4 | alpha/beta hydrolase fold<br>TAP-like protein | 3.60E-22<br>1.00E-15 |

| index | leader | core | start | end | dir | score |
|---|---|---|---|---|---|---|
| 1 | LLGGSGTVVGAFLGALLLGIVFDGFNLTGVSANAFNVVLGAAILVAMVLN | VTLIVVRKRAGSRELT | 111623 | 111824 | + | 8 |
| 2 | VVGAFLGALLLGIVFDGFNLTGVSANAFNVVLGAAILVAMVLN | VTLIVVRKRAGSRELT | 111644 | 111824 | + | 7 |
| 3 | MRAVDSDTIPQAGS | GTPTAIGLPVGPSQVATSR | 116370 | 116472 | + | 6 |
| 4 | MNRIECAAAKSTLEALEINHEIAAPVLTPVL | VTVTVAGFAGGAAVGYVQCARHGCI...GIRMTDIREMSVSELLQARQDAAVA | 116624 | 116885 | + | 17 |
| 5 | MNRIELAAEKLAGSMVPLQEVTPEQPVV | CTPVVTAGFAAFGIGYYVAKAFGPS...EASGAGIEAMSGNDLIQFRRTEVSA | 123208 | 123460 | + | 16 |

### Results for WP_110098361.1 [Auritidibacter sp. NML100628] back to top

previous - next

#### Architecture

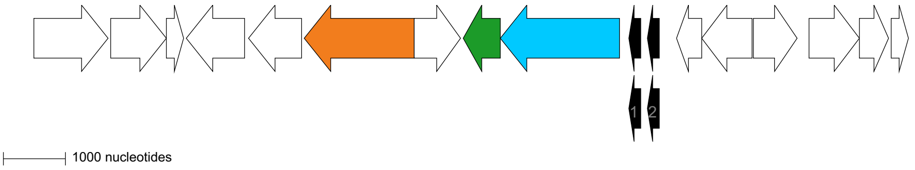

Link to nucleotide sequence

| Accession | start | end | direction | length (aa) | Pfam/HMM | name | description | E-value |
|---|---|---|---|---|---|---|---|---|
| WP_110098356.1 | 193377 | 194571 | + | 397 | TIGR02819<br>TIGR03989<br>TIGR03366<br>TIGR00692<br>PF08240 | TIGR02819<br>TIGR03989<br>TIGR03366<br>TIGR00692<br>ADH_N | fdhA_non_GSH: formaldehyde dehydrogenase, glutathione-independent<br>Rxyl_3153: NDMA-dependent alcohol dehydrogenase, Rxyl_3153 family<br>HpnZ_proposed: putative phosphonate catabolism associated alcohol dehydrogenase<br>tdh: L-threonine 3-dehydrogenase<br>Alcohol dehydrogenase GroES-like domain | 6.00E-35<br>3.40E-30<br>1.40E-29<br>4.90E-27<br>9.80E-25 |
| WP_013770884.1 | 194613 | 195509 | + | 298 | INFERRED GENE | - | - | - |
| WP_110098510.1 | 195514 | 195799 | + | 94 | PF07498 | Rho_N | Rho termination factor, N-terminal domain | 1.50E-07 |
| WP_158278463.1 | 196775 | 195833 | - | 313 | PF01551 | Peptidase_M23 | Peptidase family M23 | 2.30E-21 |
| WP_146206257.1 | 197689 | 196849 | - | 279 | PF12680 | SnoaL_2 | SnoaL-like domain | 7.00E-08 |
| WP_110098359.1 | 199511 | 197750 | - | 586 | TIGR02204<br>TIGR02203<br>TIGR00958<br>TIGR01846<br>TIGR03375 | TIGR02204<br>TIGR02203<br>TIGR00958<br>TIGR01846<br>TIGR03375 | MsbA_rel: ABC transporter, permease/ATP-binding protein<br>MsbA_lipidA: lipid A export permease/ATP-binding protein MsbA<br>3a01208: antigen peptide transporter 2<br>type_I_sec_HlyB: type I secretion system ATPase<br>type_I_sec_LssB: type I secretion system ATPase | 1.50E-102<br>5.70E-102<br>5.70E-88<br>7.40E-84<br>5.60E-82 |

| | | | | | | | | |
|---|---|---|---|---|---|---|---|---|
| WP_007452030.1 | 199507 | 200264 | + | 252 | INFERRED GENE | - | - | - |
| WP_110098360.1 | 200898 | 200310 | - | 195 | LinL | LinL | LinL | 1.80E-67 |
| WP_110098361.1 | 202835 | 200894 | - | 646 | LinH<br>LinG<br>LinE<br>PF12697<br>PF00561 | LinH<br>LinG<br>LinE<br>Abhydrolase_6<br>Abhydrolase_1 | LinH (Fusion of LinE/LinG)<br>LinG Homolog<br>LinE Homolog<br>Alpha/beta hydrolase family<br>alpha/beta hydrolase fold | 5.70E-230<br>2.40E-116<br>8.60E-92<br>3.20E-14<br>2.60E-11 |
| WP_110098362.1 | 203162 | 202973 | - | 62 | LinA | LinA | LinA (linaridin precursor) | 4.10E-13 |
| WP_146206258.1 | 203469 | 203268 | - | 66 | LinA | LinA | LinA (linaridin precursor) | 9.90E-16 |
| WP_110098363.1 | 204160 | 203743 | - | 138 | PF07100 | ASRT | Anabaena sensory rhodopsin transducer | 1.10E-40 |
| WP_110098364.1 | 204966 | 204159 | - | 268 | TIGR01679<br>PF04030<br>TIGR01678<br>TIGR01677 | TIGR01679<br>ALO<br>TIGR01678<br>TIGR01677 | bact_FAD_ox: FAD-linked oxidoreductase<br>D-arabinono-1,4-lactone oxidase<br>FAD_lactone_ox: sugar 1,4-lactone oxidases<br>pln_FAD_oxido: plant-specific FAD-dependent oxidoreductase | 7.60E-59<br>7.20E-46<br>2.70E-24<br>2.30E-12 |
| WP_015400475.1 | 204992 | 205692 | + | 233 | INFERRED GENE | - | - | - |
| WP_158278462.1 | 205879 | 206701 | + | 273 | PF01487<br>TIGR01093 | DHquinase_I<br>TIGR01093 | Type I 3-dehydroquinase<br>aroD: 3-dehydroquinate dehydratase, type I | 1.20E-51<br>3.20E-51 |
| WP_110098366.1 | 206697 | 207156 | + | 152 | PF11255 | DUF3054 | Protein of unknown function (DUF3054) | 2.10E-20 |
| WP_110098367.1 | 207205 | 207484 | + | 92 | PF01037 | AsnC_trans_reg | Lrp/AsnC ligand binding domain | 1.90E-16 |

| index | leader | core | start | end | dir | score |
|---|---|---|---|---|---|---|
| 1 | MEHIIEKVERETEPQLSEHSSMRNPGLDPVM | ATPAAFVGAAGYAWITLQAYDMGREHAGANG | 203162 | 202973 | - | 27 |
| 2 | MSTRVADFVNSQDLGELKSGFRREASPVM | ATPAAVVGALLGKKVVGAAAAGVTGVGAAYTVGRVVN | 203469 | 203268 | - | 29 |

### Results for WP_110122900.1 [Auritidibacter sp. NML120779] back to top

previous - next

#### Architecture

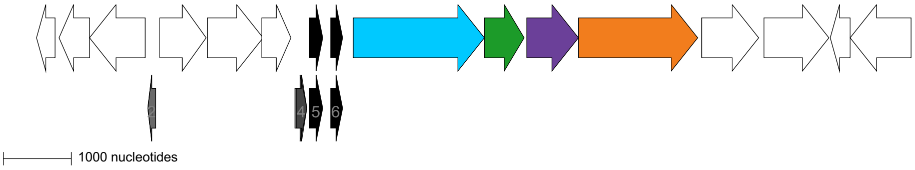

Link to nucleotide sequence

| Accession | start | end | direction | length (aa) | Pfam/HMM | name | description | E-value |
|---|---|---|---|---|---|---|---|---|
| WP_110098367.1 | 1767 | 1488 | - | 92 | PF01037 | AsnC_trans_reg | Lrp/AsnC ligand binding domain | 1.90E-16 |
| WP_110098366.1 | 2275 | 1816 | - | 152 | PF11255 | DUF3054 | Protein of unknown function (DUF3054) | 2.10E-20 |
| WP_158278462.1 | 3093 | 2271 | - | 273 | PF01487<br>TIGR01093 | DHquinase_I<br>TIGR01093 | Type I 3-dehydroquinase<br>aroD: 3-dehydroquinate dehydratase, type I | 1.20E-51<br>3.20E-51 |
| WP_009493164.1 | 3292 | 3980 | + | 229 | INFERRED GENE | - | - | - |
| WP_110122898.1 | 4006 | 4813 | + | 268 | TIGR01679<br>PF04030<br>TIGR01678<br>TIGR01677 | TIGR01679<br>ALO<br>TIGR01678<br>TIGR01677 | bact_FAD_ox: FAD-linked oxidoreductase<br>D-arabinono-1,4-lactone oxidase<br>FAD_lactone_ox: sugar 1,4-lactone oxidases<br>pln_FAD_oxido: plant-specific FAD-dependent oxidoreductase | 6.40E-59<br>1.60E-45<br>7.40E-24<br>7.90E-12 |
| WP_110098363.1 | 4812 | 5229 | + | 138 | PF07100 | ASRT | Anabaena sensory rhodopsin transducer | 1.10E-40 |
| WP_146206258.1 | 5505 | 5706 | + | 66 | LinA | LinA | LinA (linaridin precursor) | 9.90E-16 |
| WP_110122899.1 | 5812 | 6001 | + | 62 | LinA | LinA | LinA (linaridin precursor) | 8.90E-13 |
| WP_110122900.1 | 6139 | 8080 | + | 646 | LinH<br>LinG<br>LinE<br>PF12697<br>PF00561 | LinH<br>LinG<br>LinE<br>Abhydrolase_6<br>Abhydrolase_1 | LinH (Fusion of LinE/LinG)<br>LinG Homolog<br>LinE Homolog<br>Alpha/beta hydrolase family<br>alpha/beta hydrolase fold | 1.70E-229<br>2.10E-116<br>1.90E-91<br>7.70E-14<br>2.70E-11 |
| WP_110098360.1 | 8076 | 8664 | + | 195 | LinL | LinL | LinL | 1.80E-67 |
| WP_110122901.1 | 8710 | 9469 | + | 252 | PF13649<br>PF08241<br>PF13847<br>PF13489<br>TIGR01983 | Methyltransf_25<br>Methyltransf_11<br>Methyltransf_31<br>Methyltransf_23<br>TIGR01983 | Methyltransferase domain<br>Methyltransferase domain<br>Methyltransferase domain<br>Methyltransferase domain<br>UbiG: 3-demethylubiquinone-9 3-O-methyltransferase | 7.50E-14<br>2.30E-09<br>1.30E-07<br>4.30E-07<br>6.60E-06 |
| WP_110122902.1 | 9465 | 11226 | + | 586 | TIGR02204<br>TIGR02203<br>TIGR00958<br>TIGR01846<br>TIGR03375 | TIGR02204<br>TIGR02203<br>TIGR00958<br>TIGR01846<br>TIGR03375 | MsbA_rel: ABC transporter, permease/ATP-binding protein<br>MsbA_lipidA: lipid A export permease/ATP-binding protein MsbA<br>3a01208: antigen peptide transporter 2<br>type_I_sec_HlyB: type I secretion system ATPase<br>type_I_sec_LssB: type I secretion system ATPase | 5.60E-102<br>1.90E-101<br>3.00E-87<br>2.10E-83<br>1.50E-81 |
| WP_146206753.1 | 11288 | 12128 | + | 279 | PF12680 | SnoaL_2 | SnoaL-like domain | 5.20E-08 |
| WP_158278463.1 | 12202 | 13144 | + | 313 | PF01551 | Peptidase_M23 | Peptidase family M23 | 2.30E-21 |
| WP_110098510.1 | 13463 | 13178 | - | 94 | PF07498 | Rho_N | Rho termination factor, N-terminal domain | 1.50E-07 |

| | | | | | | | | |
|---|---|---|---|---|---|---|---|---|
| WP_110122904.1 | 14365 | 13468 | - | 298 | PF13561<br>TIGR03971<br>TIGR01830<br>PF00106<br>TIGR01829 | adh_short_C2<br>TIGR03971<br>TIGR01830<br>adh_short<br>TIGR01829 | Enoyl-(Acyl carrier protein) reductase<br>SDR_subfam_1: SDR family mycofactocin-dependent oxidoreductase<br>3oxo_ACP_reduc: 3-oxoacyl-[acyl-carrier-protein] reductase<br>short chain dehydrogenase<br>AcAcCoA_reduct: acetoacetyl-CoA reductase | 6.90E-58<br>8.60E-45<br>2.00E-43<br>2.70E-41<br>7.50E-37 |

| index | leader | core | start | end | dir | score |
|---|---|---|---|---|---|---|
| 1 | MDECGRAGQHLAAGSG | SAGAQKTASLGHTFNMT | 3233 | 3131 | - | 6 |
| 2 | VDLMDECGRAGQHLAAGS | GSAGAQKTASLGHTFNMT | 3242 | 3131 | - | 6 |
| 3 | LACVCSERFCELNATSTHSKSLRRKLRC | ILAPVQVLCNVLCVATGSIMQLHLPQETS | 5292 | 5466 | + | 7 |
| 4 | VCSERFCELNATSTHSKSLRRKLRCIL | APVQVLCNVLCVATGSIMQLHLPQETS | 5301 | 5466 | + | 7 |
| 5 | MSTRVADFVNSQDLGELKSGFRREASPVM | ATPAAVVGALLGKKVVGAAAAGVTGVGAAYTVGRVVN | 5505 | 5706 | + | 30 |
| 6 | MEHIIEKVERETEPQLSEHSSMRNPGLDPVM | ATPAAFVGAAGYGWITLQAYDMGREHAGANG | 5812 | 6001 | + | 28 |

### Results for WP_110101283.1 [Auritidibacter ignavus] back to top

previous - next

#### Architecture

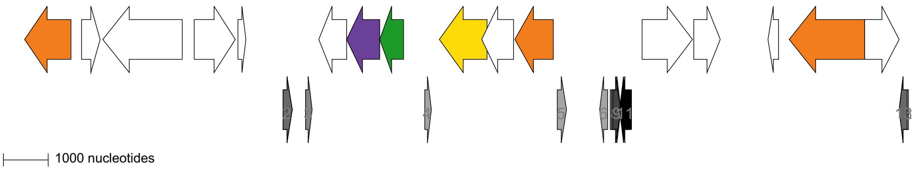

Link to nucleotide sequence

| Accession | start | end | direction | length (aa) | Pfam/HMM | name | description | E-value |
|---|---|---|---|---|---|---|---|---|
| WP_110101275.1 | 26638 | 25630 | - | 335 | TIGR02314<br>TIGR02315<br>TIGR02673<br>TIGR01186<br>TIGR03005 | TIGR02314<br>TIGR02315<br>TIGR02673<br>TIGR01186<br>TIGR03005 | ABC_MetN: D-methionine ABC transporter, ATP-binding protein<br>ABC_phnC: phosphonate ABC transporter, ATP-binding protein<br>FtsE: cell division ATP-binding protein FtsE<br>proV: glycine betaine/L-proline transport ATP binding subunit<br>ectoine_ehuA: ectoine/hydroxyectoine ABC transporter, ATP-binding protein EhuA | 1.70E-92<br>6.50E-59<br>6.50E-59<br>1.00E-58<br>2.70E-57 |
| WP_110101346.1 | 26880 | 27294 | + | 137 | PF11349 | DUF3151 | Protein of unknown function (DUF3151) | 5.50E-55 |
| WP_110101276.1 | 29136 | 27369 | - | 588 | PF00755 | Carn_acyltransf | Choline/Carnitine o-acyltransferase | 1.20E-88 |
| WP_110101277.1 | 29414 | 30332 | + | 305 | PF00877<br>PF01476<br>TIGR02899<br>PF05382 | NLPC_P60<br>LysM<br>TIGR02899<br>Amidase_5 | NlpC/P60 family<br>LysM domain<br>spore_safA: spore coat assembly protein SafA<br>Bacteriophage peptidoglycan hydrolase | 5.10E-22<br>4.90E-15<br>5.20E-08<br>2.60E-07 |
| WP_005290314.1 | 30369 | 30561 | + | 64 | INFERRED GENE | - | - | - |
| WP_110101279.1 | 32795 | 32183 | - | 203 | PF00196<br>TIGR03541<br>TIGR03020<br>TIGR01884<br>PF08279 | GerE<br>TIGR03541<br>TIGR03020<br>TIGR01884<br>HTH_11 | Bacterial regulatory proteins, luxR family<br>reg_near_HchA: LuxR family transcriptional regulatory, chaperone HchA-associated<br>EpsA: transcriptional regulator EpsA<br>cas_HTH: CRISPR locus-related DNA-binding protein<br>HTH domain | 5.80E-16<br>4.00E-12<br>4.40E-10<br>6.70E-05<br>6.60E-04 |
| WP_110101280.1 | 33535 | 32791 | - | 247 | PF13649<br>PF13489<br>PF08241<br>TIGR02072<br>PF08242 | Methyltransf_25<br>Methyltransf_23<br>Methyltransf_11<br>TIGR02072<br>Methyltransf_12 | Methyltransferase domain<br>Methyltransferase domain<br>Methyltransferase domain<br>BioC: malonyl-acyl carrier protein O-methyltransferase BioC<br>Methyltransferase domain | 2.20E-12<br>3.80E-11<br>6.30E-11<br>2.70E-09<br>6.10E-08 |
| WP_122548721.1 | 34083 | 33531 | - | 183 | LinL | LinL | LinL | 2.80E-50 |
| WP_110101283.1 | 35947 | 34870 | - | 358 | LinG<br>LinH<br>PF05090<br>PF06762<br>TIGR04033 | LinG<br>LinH<br>VKG_Carbox<br>LMF1<br>TIGR04033 | LinG Homolog<br>LinH (Fusion of LinE/LinG)<br>Vitamin K-dependent gamma-carboxylase<br>Lipase maturation factor<br>export_SdpB: antimicrobial peptide system protein, SdpB family | 2.10E-87<br>1.50E-76<br>1.30E-06<br>6.70E-05<br>1.10E-04 |
| WP_110101284.1 | 36540 | 35832 | - | 235 | PF01061<br>PF12698 | ABC2_membrane<br>ABC2_membrane_3 | ABC-2 type transporter<br>ABC-2 family transporter protein | 3.00E-09<br>7.70E-04 |
| WP_110110510.1 | 37428 | 36549 | - | 292 | TIGR01188<br>TIGR03864<br>TIGR01288<br>TIGR03740<br>TIGR03265 | TIGR01188<br>TIGR03864<br>TIGR01288<br>TIGR03740<br>TIGR03265 | drrA: daunorubicin resistance ABC transporter, ATP-binding protein<br>PQQ_ABC_ATP: ABC transporter, ATP-binding subunit, PQQ-dependent alcohol dehydrogenase system<br>nodI: nodulation ABC transporter NodI<br>galliderm_ABC: lantibiotic protection ABC transporter, ATP-binding subunit<br>PhnT2: putative 2-aminoethylphosphonate ABC transporter, ATP-binding protein | 8.00E-58<br>1.30E-44<br>2.70E-41<br>1.20E-37<br>5.20E-36 |
| WP_110101287.1 | 39391 | 40528 | + | 378 | PF07730 | HisKA_3 | Histidine kinase | 1.10E-10 |
| WP_110101288.1 | 40542 | 41145 | + | 200 | PF00072<br>TIGR02875<br>PF00196<br>TIGR01818<br>TIGR03020 | Response_reg<br>TIGR02875<br>GerE<br>TIGR01818<br>TIGR03020 | Response regulator receiver domain<br>spore_0_A: sporulation transcription factor Spo0A<br>Bacterial regulatory proteins, luxR family<br>ntrC: nitrogen regulation protein NR(I)<br>EpsA: transcriptional regulator EpsA | 8.30E-25<br>2.80E-18<br>1.60E-14<br>2.60E-11<br>1.70E-08 |
| WP_110101289.1 | 42452 | 42191 | - | 86 | NO PFAM MATCH | - | - | - |

| | | | | | | | | |
|---|---|---|---|---|---|---|---|---|
| WP_110101290.1 | 44375 | 42692 | - | 560 | TIGR02770<br>TIGR02769<br>TIGR01186<br>TIGR00968<br>TIGR03265 | TIGR02770<br>TIGR02769<br>TIGR01186<br>TIGR00968<br>TIGR03265 | nickel_nikD: nickel import ATP-binding protein NikD<br>nickel_nikE: nickel import ATP-binding protein NikE<br>proV: glycine betaine/L-proline transport ATP binding subunit<br>3a0106s01: sulfate ABC transporter, ATP-binding protein<br>PhnT2: putative 2-aminoethylphosphonate ABC transporter, ATP-binding protein | 8.10E-108<br>1.10E-102<br>9.60E-78<br>2.10E-74<br>3.30E-74 |
| WP_019175934.1 | 44371 | 45144 | + | 257 | INFERRED GENE | - | - | - |

| index | leader | core | start | end | dir | score |
|---|---|---|---|---|---|---|
| 1 | VLPRGVAQGVAETCLRPDLRSMPTLHMPAITAFSANDLSRELSSAVG | RTFKPATVGKTCSHGRW | 31390 | 31585 | + | 6 |
| 2 | LPRGVAQGVAETCLRPDLRSMPTLHMPAITAFSANDLSRELSSAVG | RTFKPATVGKTCSHGRW | 31393 | 31585 | + | 6 |
| 3 | MVGHGCWWFGNCGHSEPFH | VTVSVKVQCSKRSAPVEAKSVNPQEVWLREVAIGT | 31879 | 32044 | + | 7 |
| 4 | VLEESTSANPLTHCVIWATKCASNTPRKEWFEK | ITTVLVRRRSRNRSAAL | 34544 | 34697 | + | 6 |
| 5 | LCAAFATHNPEPLASSYGRNLTDASRETDRKT | SWPGRFTGRSLTPRVRMGGVASRLICGPLLKV | 37506 | 37701 | + | 6 |
| 6 | LGHRPNAVSNFSGHRDAGPGRSRSIRGGG | VLVAPIHRYVSGPVTTGYRWRQPNATSLL | 38620 | 38443 | - | 6 |
| 7 | VIPELSLCLPMSATGSLSNKLPYSL...AYWNAYVFHVFAVADIDNPRPIALG | VTPTGLSSVQEESKVLREHFSVALRHLVE | 38673 | 38922 | + | 6 |
| 8 | LSNKLPYSLNALAYWNAYVFHVFAVADIDNPRPIALG | VTPTGLSSVQEESKVLREHFSVALRHLVE | 38721 | 38922 | + | 6 |
| 9 | LNALAYWNAYVFHVFAVADIDNPRPIALG | VTPTGLSSVQEESKVLREHFSVALRHLVE | 38745 | 38922 | + | 6 |
| 10 | MPTLATPATTATTATTAIFGFTDSNPVTVSAEV | APYGSVNELIRARESALQPNGTQLTLQGGGGPR | 39091 | 38890 | - | 13 |
| 11 | MRNDLTHFADTLNIGEVGHAVTAMPTL | ATPATTATTATTAIFGFTDSNPVTV...LIRARESALQPNGTQLTLQGGGGPR | 39160 | 38890 | - | 25 |
| 12 | VIEYLFRYPGLGQALIDAVTYRDVQVVQAI | SLLIAVVYVLVNLAADIVGILANPKLRSAS | 45323 | 45140 | - | 6 |
| 13 | VVVIEYLFRYPGLGQALIDAVTYRDVQVVQA | ISLLIAVVYVLVNLAADIVGILANPKLRSAS | 45329 | 45140 | - | 6 |

### Results for PXA76677.1 [Auritidibacter sp. NML100628] back to top

previous - next

#### Architecture

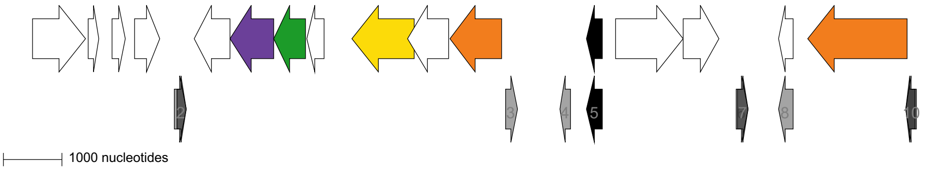

Link to nucleotide sequence

| Accession | start | end | direction | length (aa) | Pfam/HMM | name | description | E-value |
|---|---|---|---|---|---|---|---|---|
| PXA76670.1 | 28163 | 29081 | + | 305 | PF00877<br>PF01476<br>TIGR02899<br>PF05382<br>TIGR02594 | NLPC_P60<br>LysM<br>TIGR02899<br>Amidase_5<br>TIGR02594 | NlpC/P60 family<br>LysM domain<br>spore_safA: spore coat assembly protein SafA<br>Bacteriophage peptidoglycan hydrolase<br>TIGR02594: TIGR02594 family protein | 3.30E-22<br>4.30E-15<br>5.20E-08<br>7.60E-07<br>9.70E-04 |
| WP_005290314.1 | 29118 | 29310 | + | 64 | INFERRED GENE | - | - | - |
| PXA76671.1 | 29527 | 29761 | + | 77 | PF01370<br>TIGR03466<br>PF04321<br>PF01113<br>PF05368 | Epimerase<br>TIGR03466<br>RmlD_sub_bind<br>DapB_N<br>NmrA | NAD dependent epimerase/dehydratase family<br>HpnA: hopanoid-associated sugar epimerase<br>RmlD substrate binding domain<br>Dihydrodipicolinate reductase, N-terminus<br>NmrA-like family | 7.40E-05<br>2.30E-04<br>2.40E-04<br>3.50E-04<br>5.40E-04 |
| PXA76672.1 | 29918 | 30335 | + | 138 | NO PFAM MATCH | - | - | - |
| PXA76673.1 | 31545 | 30933 | - | 203 | PF00196<br>TIGR03541<br>TIGR03020<br>TIGR01884<br>PF08279 | GerE<br>TIGR03541<br>TIGR03020<br>TIGR01884<br>HTH_11 | Bacterial regulatory proteins, luxR family<br>reg_near_HchA: LuxR family transcriptional regulatory, chaperone HchA-associated<br>EpsA: transcriptional regulator EpsA<br>cas_HTH: CRISPR locus-related DNA-binding protein<br>HTH domain | 2.90E-16<br>2.30E-12<br>6.40E-11<br>1.60E-05<br>1.80E-04 |
| PXA76674.1 | 32285 | 31541 | - | 247 | PF13649<br>PF13489<br>PF08241<br>TIGR02072<br>PF08242 | Methyltransf_25<br>Methyltransf_23<br>Methyltransf_11<br>TIGR02072<br>Methyltransf_12 | Methyltransferase domain<br>Methyltransferase domain<br>Methyltransferase domain<br>BioC: malonyl-acyl carrier protein O-methyltransferase BioC<br>Methyltransferase domain | 4.40E-13<br>3.00E-12<br>6.20E-12<br>2.00E-10<br>5.10E-09 |
| PXA76675.1 | 32833 | 32281 | - | 183 | LinL | LinL | LinL | 2.90E-52 |
| PXA76676.1 | 33149 | 32855 | - | 97 | NO PFAM MATCH | - | - | - |
| PXA76677.1 | 34696 | 33619 | - | 358 | LinG<br>LinH<br>PF05090<br>PF06762<br>TIGR04033 | LinG<br>LinH<br>VKG_Carbox<br>LMF1<br>TIGR04033 | LinG Homolog<br>LinH (Fusion of LinE/LinG)<br>Vitamin K-dependent gamma-carboxylase<br>Lipase maturation factor<br>export_SdpB: antimicrobial peptide system protein, SdpB family | 4.50E-87<br>1.40E-76<br>1.20E-06<br>3.20E-05<br>1.30E-04 |

| | | | | | | | | |
|---|---|---|---|---|---|---|---|---|
| PXA76678.1 | 35289 | 34581 | - | 235 | PF01061<br>PF12698 | ABC2_membrane<br>ABC2_membrane_3 | ABC-2 type transporter<br>ABC-2 family transporter protein | 3.00E-09<br>8.90E-04 |
| PXA76679.1 | 36177 | 35298 | - | 292 | TIGR01188<br>TIGR03864<br>TIGR01288<br>TIGR03740<br>TIGR03265 | TIGR01188<br>TIGR03864<br>TIGR01288<br>TIGR03740<br>TIGR03265 | drrA: daunorubicin resistance ABC transporter, ATP-binding protein<br>PQQ_ABC_ATP: ABC transporter, ATP-binding subunit, PQQ-dependent alcohol dehydrogenase<br>system<br>nodI: nodulation ABC transporter NodI<br>galliderm_ABC: lantibiotic protection ABC transporter, ATP-binding subunit<br>PhnT2: putative 2-aminoethylphosphonate ABC transporter, ATP-binding protein | 4.70E-57<br>3.10E-44<br>2.70E-41<br>9.00E-38<br>1.70E-35 |
| PXA76680.1 | 37909 | 37639 | - | 89 | LinA | LinA | LinA (linaridin precursor) | 7.10E-07 |
| PXA76681.1 | 38140 | 39277 | + | 378 | PF07730 | HisKA_3 | Histidine kinase | 1.10E-10 |
| PXA76682.1 | 39291 | 39894 | + | 200 | PF00072<br>TIGR02875<br>PF00196<br>TIGR01818<br>TIGR03020 | Response_reg<br>TIGR02875<br>GerE<br>TIGR01818<br>TIGR03020 | Response regulator receiver domain<br>spore_0_A: sporulation transcription factor Spo0A<br>Bacterial regulatory proteins, luxR family<br>ntrC: nitrogen regulation protein NR(I)<br>EpsA: transcriptional regulator EpsA | 1.50E-25<br>1.90E-18<br>1.60E-14<br>8.70E-12<br>1.60E-08 |
| PXA76683.1 | 41174 | 40913 | - | 86 | NO PFAM<br>MATCH | - | - | - |
| PXA76684.1 | 43103 | 41420 | - | 560 | TIGR02770<br>TIGR02769<br>TIGR01186<br>TIGR00968<br>TIGR03265 | TIGR02770<br>TIGR02769<br>TIGR01186<br>TIGR00968<br>TIGR03265 | nickel_nikD: nickel import ATP-binding protein NikD<br>nickel_nikE: nickel import ATP-binding protein NikE<br>proV: glycine betaine/L-proline transport ATP binding subunit<br>3a0106s01: sulfate ABC transporter, ATP-binding protein<br>PhnT2: putative 2-aminoethylphosphonate ABC transporter, ATP-binding protein | 1.80E-<br>110<br>2.00E-<br>105<br>1.50E-79<br>1.80E-76<br>7.20E-76 |

| index | leader | core | start | end | dir | score |
|---|---|---|---|---|---|---|
| 1 | VLSLRRGGMVGHGRWRLGNCDHSEPFH | VTVSVKVQCSKPSAPIEVISVSPQEVWLREVGTGT | 30605 | 30794 | + | 6 |
| 2 | MVGHGRWRLGNCDHSEPFH | VTVSVKVQCSKPSAPIEVISVSPQEVWLREVGTGT | 30629 | 30794 | + | 7 |
| 3 | LCAAFATHNPEPLASSYGRNLTDASRETDRKT | SWPGRFTGRSLIPRVRMGGVASRLTCGPLLKV | 36255 | 36450 | + | 6 |
| 4 | LGHRRNAVSNFSGHRDAGPGRYRGIRGGG | VLVAPIHRYVSGPVTTGYRRRQPNATSLL | 37369 | 37192 | - | 6 |
| 5 | MRNDLTHFADTLNIGEVGHAVTAMPTL | ATPATTATTATTAIFGFTDSNPVTV...LIRARESALQPNGTQLTLQGGGEPR | 37909 | 37639 | - | 25 |
| 6 | VGGSYCQFTRWACVVRSGGCFSVWTGTGLGSEGDSPRP | ATITGTISFPSGLGLAREPTPSAGTSGELCV | 40187 | 40397 | + | 6 |
| 7 | VVRSGGCFSVWTGTGLGSEGDSPRP | ATITGTISFPSGLGLAREPTPSAGTSGELCV | 40226 | 40397 | + | 7 |
| 8 | MQAVRLVSLVVMIIALVVAVFVVLNNVMDPSSLPDP | IAPAIEALPEQGSGAAIVLTVVVVGVGILASVVHKNAKKALEKQDSKKNL | 41174 | 40913 | - | 6 |
| 9 | VIAIAVLTIGTNLLADALAR | ATATTTLPVKASRLRSTKNTPETTAEKVANR | 43255 | 43099 | - | 7 |
| 10 | VILPVIAIAVLTIGTNLLADALAR | ATATTTLPVKASRLRSTKNTPETTAEKVANR | 43267 | 43099 | - | 7 |

### Results for WP_168186076.1 [Auritidibacter sp. NML130574] back to top

previous - next

#### Architecture

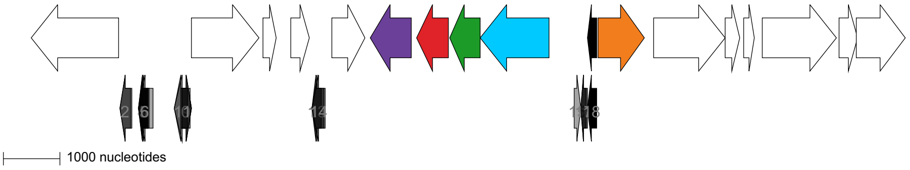

Link to nucleotide sequence

| Accession | start | end | direction | length (aa) | Pfam/HMM | name | description | E-value |
|---|---|---|---|---|---|---|---|---|
| WP_110110875.1 | 1162457 | 1160903 | - | 517 | TIGR00835<br>PF01235 | TIGR00835<br>Na_Ala_symp | agcS: amino acid carrier protein<br>Sodium:alanine symporter family | 2.20E-154<br>7.90E-144 |
| WP_117328665.1 | 1163737 | 1164937 | + | 399 | PF13936<br>PF00665<br>TIGR01884 | HTH_38<br>rve<br>TIGR01884 | Helix-turn-helix domain<br>Integrase core domain<br>cas_HTH: CRISPR locus-related DNA-binding protein | 2.50E-19<br>1.40E-06<br>8.50E-04 |
| WP_076611468.1 | 1164998 | 1165246 | + | 82 | INFERRED GENE | - | - | - |
| WP_148707508.1 | 1165491 | 1165827 | + | 111 | NO PFAM<br>MATCH | - | - | - |
| WP_017832712.1 | 1166221 | 1166816 | + | 198 | INFERRED GENE | - | - | - |
| WP_110111035.1 | 1167623 | 1166915 | - | 235 | PF13649<br>PF08241<br>TIGR02072<br>PF13847<br>TIGR01934 | Methyltransf_25<br>Methyltransf_11<br>TIGR02072<br>Methyltransf_31<br>TIGR01934 | Methyltransferase domain<br>Methyltransferase domain<br>BioC: malonyl-acyl carrier protein O-methyltransferase BioC<br>Methyltransferase domain<br>MenG_MenH_UbiE: ubiquinone/menaquinone biosynthesis methyltransferase | 6.30E-15<br>3.80E-12<br>6.70E-12<br>4.00E-11<br>2.80E-10 |
| WP_110111036.1 | 1168291 | 1167715 | - | 191 | TIGR00521<br>PF02441<br>TIGR02113 | TIGR00521<br>Flavoprotein<br>TIGR02113 | coaBC_dfp: phosphopantothenoylcysteine decarboxylase / phosphopantothenate--cysteine<br>ligase<br>Flavoprotein<br>coaC_strep: phosphopantothenoylcysteine decarboxylase | 2.10E-09<br>2.10E-06<br>1.50E-04 |
| WP_110111037.1 | 1168854 | 1168314 | - | 179 | LinL | LinL | LinL | 6.10E-45 |

| | | | | | | | | |
|---|---|---|---|---|---|---|---|---|
| WP_168186076.1 | 1170073 | 1168846 | - | 408 | LinH<br>LinE<br>LinG<br>PF12697<br>TIGR01738 | LinH<br>LinE<br>LinG<br>Abhydrolase_6<br>TIGR01738 | LinH (Fusion of LinE/LinG)<br>LinE Homolog<br>LinG Homolog<br>Alpha/beta hydrolase family<br>bioH: pimelyl-[acyl-carrier protein] methyl ester esterase | 4.00E-104<br>9.00E-58<br>1.50E-28<br>1.40E-08<br>2.90E-04 |
| WP_148707509.1 | 1170922 | 1170745 | - | 58 | LinA | LinA | LinA (linaridin precursor) | 1.20E-12 |
| WP_110111039.1 | 1170943 | 1171759 | + | 271 | TIGR01189<br>TIGR02673<br>TIGR02868<br>PF00005<br>TIGR02857 | TIGR01189<br>TIGR02673<br>TIGR02868<br>ABC_tran<br>TIGR02857 | ccmA: heme ABC exporter, ATP-binding protein CcmA<br>FtsE: cell division ATP-binding protein FtsE<br>CydC: thiol reductant ABC exporter, CydC subunit<br>ABC transporter<br>CydD: thiol reductant ABC exporter, CydD subunit | 4.60E-33<br>7.50E-26<br>1.10E-25<br>1.10E-24<br>1.80E-24 |
| WP_148707510.1 | 1171914 | 1173180 | + | 421 | NO PFAM MATCH | - | - | - |
| WP_110111041.1 | 1173180 | 1173435 | + | 84 | PF02796<br>PF13936<br>PF13384<br>PF00239<br>PF13551 | HTH_7<br>HTH_38<br>HTH_23<br>Resolvase<br>HTH_29 | Helix-turn-helix domain of resolvase<br>Helix-turn-helix domain<br>Homeodomain-like domain<br>Resolvase, N terminal domain<br>Winged helix-turn helix | 1.90E-09<br>4.70E-07<br>1.10E-06<br>2.00E-05<br>4.60E-05 |
| WP_010147305.1 | 1173518 | 1173701 | + | 61 | INFERRED GENE | - | - | - |
| WP_117328628.1 | 1173850 | 1175158 | + | 435 | PF01610<br>PF14690<br>PF13542 | DDE_Tnp_ISL3<br>zf-ISL3<br>HTH_Tnp_ISL3 | Transposase<br>zinc-finger of transposase IS204/IS1001/IS1096/IS1165<br>Helix-turn-helix domain of transposase family ISL3 | 1.60E-67<br>4.30E-09<br>5.20E-05 |
| WP_117328629.1 | 1175199 | 1175520 | + | 106 | PF01527 | HTH_Tnp_1 | Transposase | 1.60E-06 |
| WP_117328630.1 | 1175516 | 1176368 | + | 283 | PF00665<br>PF13276<br>PF13683 | rve<br>HTH_21<br>rve_3 | Integrase core domain<br>HTH-like domain<br>Integrase core domain | 6.10E-22<br>1.40E-14<br>4.70E-08 |

| index | leader | core | start | end | dir | score |
|---|---|---|---|---|---|---|
| 1 | VRHDLSYALNINKVRRYIDLDGSVARIT | GVSADEVVGMARLLADSPRTQISVGPDNR | 1162661 | 1162487 | - | 7 |
| 2 | MASRSRYVRHDLSYALNINKVRRYIDLDGSVA | RITGVSADEVVGMARLLADSPRTQISVGPDNR | 1162682 | 1162487 | - | 7 |
| 3 | MPINRHTRINCFFAFAEYFVGLCFRRSM | RTTFLTPMVCEVRYRLAVDRPNPVEAQN | 1162968 | 1162797 | - | 9 |
| 4 | LPEGMPINRHTRINCFFAFAEYFVGLCFRR | SMRTTFLTPMVCEVRYRLAVDRPNPVEAQN | 1162980 | 1162797 | - | 8 |
| 5 | MYLPEGMPINRHTRINCFFAFAEYFVGLCFR | RSMRTTFLTPMVCEVRYRLAVDRPNPVEAQN | 1162986 | 1162797 | - | 8 |
| 6 | VGLYPERPNANSSWASNASNQSSITGFMYLPEGMPINRHTRINC | FFAFAEYFVGLCFRRSMRTTFLTPMVCEVRYRLAVDRPNPVEAQN | 1163067 | 1162797 | - | 6 |
| 7 | MRRQWRADRALRPAMRSPGRPQPSRAVQRQFWRLIAAG | VTTAEAALAVGCRCRSDRGGSVTLAVCHR | 1163546 | 1163750 | + | 6 |
| 8 | LRPAMRSPGRPQPSRAVQRQFWRLIAAG | VTTAEAALAVGCRCRSDRGGSVTLAVCHR | 1163576 | 1163750 | + | 6 |
| 9 | MRSPGRPQPSRAVQRQFWRLIAAG | VTTAEAALAVGCRCRSDRGGSVTLAVCHR | 1163588 | 1163750 | + | 6 |
| 10 | VTPAAIRRQNCRCTALDGCGLPGERIAGRNARSARHC | RRIPSGTSLIFMLQSLVTILEHLSHLKVLRRPVESAL | 1163665 | 1163440 | - | 7 |
| 11 | VTEPPRSDRHRHPTASAASAVVTPAAIRRQNCRCTALDGCGLPGERI | AGRNARSARHCRRIPSGTSLIFMLQSLVTILEHLSHLKVLRRPVESAL | 1163728 | 1163440 | - | 6 |
| 12 | MSRNGKWRLACALFNSRCRRGFTALLLAA | GTPFRSFGLVSRARAAQQLAGWRSERRE | 1166047 | 1165873 | - | 7 |
| 13 | LRDATPVVHRHSMSRNGKWRLACALFNSRCRRGFTALLLAA | GTPFRSFGLVSRARAAQQLAGWRSERRE | 1166083 | 1165873 | - | 8 |
| 14 | VRRLASHPLRDATPVVHRHSMSRNGKWRLACALFNSRCRRGFTALLLAA | GTPFRSFGLVSRARAAQQLAGWRSERRE | 1166107 | 1165873 | - | 7 |
| 15 | LIMILRQPGSAQSQVVEMF | VAKLSAELALLATFSVFYV | 1170504 | 1170621 | + | 6 |
| 16 | LHDPMRVTSCLIGL | VTLIQSRQSRRITPHYTQ | 1170736 | 1170637 | - | 8 |
| 17 | LLHDPMRVTSCLIGL | VTLIQSRQSRRITPHYTQ | 1170739 | 1170637 | - | 8 |
| 18 | MSIEIDIVQQFANETFKEATPGFVTAAETPAM | ATPGVTPATPHAAQYVVQGSTICLIC | 1170922 | 1170745 | - | 34 |

### Results for SMX73793.1 [Brevibacterium aurantiacum] back to top

previous - next

#### Architecture

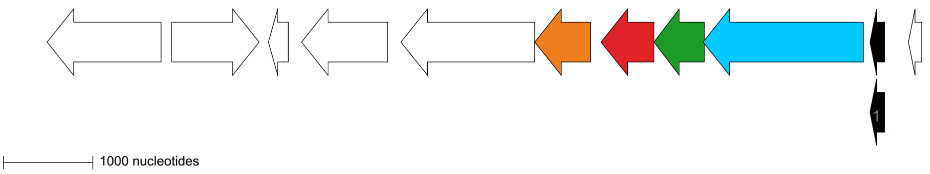

Link to nucleotide sequence

| Accession | start | end | direction | length (aa) | Pfam/HMM | name | description | E-value |
|---|---|---|---|---|---|---|---|---|
| SMX73705.1 | 619364 | 618089 | - | 424 | PF01385<br>PF07282<br>PF12323<br>TIGR01766 | OrfB_IS605<br>OrfB_Zn_ribbon<br>HTH_OrfB_IS605<br>TIGR01766 | Probable transposase<br>Putative transposase DNA-binding domain<br>Helix-turn-helix domain<br>tspaseT_teng_C: transposase, IS605 OrfB family | 1.40E-22<br>2.50E-18<br>2.20E-14<br>4.60E-08 |
| SMX73716.1 | 619473 | 620463 | + | 329 | PF07751 | Abi_2 | Abi-like protein | 2.10E-07 |

| | | | | | | | | |
|---|---|---|---|---|---|---|---|---|
| SMX73727.1 | 620785 | 620566 | - | 72 | PF02796<br>PF01527<br>PF13518<br>PF13551<br>PF13384 | HTH_7<br>HTH_Tnp_1<br>HTH_28<br>HTH_29<br>HTH_23 | Helix-turn-helix domain of resolvase<br>Transposase<br>Helix-turn-helix domain<br>Winged helix-turn helix<br>Homeodomain-like domain | 7.40E-10<br>1.20E-06<br>5.20E-06<br>1.10E-04<br>3.00E-04 |
| SMX73738.1 | 621900 | 620937 | - | 320 | PF03372 | Exo_endo_phos | Endonuclease/Exonuclease/phosphatase family | 1.00E-10 |
| SMX73753.1 | 623538 | 622038 | - | 499 | NO PFAM MATCH | - | - | - |
| SMX73761.1 | 624164 | 623534 | - | 209 | TIGR02673<br>TIGR03873<br>TIGR01189<br>TIGR01166<br>TIGR02857 | TIGR02673<br>TIGR03873<br>TIGR01189<br>TIGR01166<br>TIGR02857 | FtsE: cell division ATP-binding protein FtsE<br>F420-0_ABC_ATP: proposed F420-0 ABC transporter, ATP-binding protein<br>ccmA: heme ABC exporter, ATP-binding protein CcmA<br>cbiO: cobalt ABC transporter, ATP-binding protein<br>CydD: thiol reductant ABC exporter, CydD subunit | 1.10E-29<br>3.60E-27<br>3.80E-27<br>4.90E-26<br>3.10E-25 |
| SMX73771.1 | 624878 | 624284 | - | 197 | TIGR00521<br>PF02441 | TIGR00521<br>Flavoprotein | coaBC_dfp: phosphopantothenoylcysteine decarboxylase / phosphopantothenate--cysteine ligase<br>Flavoprotein | 2.60E-10<br>1.80E-08 |
| SMX73782.1 | 625432 | 624874 | - | 185 | LinL | LinL | LinL | 1.60E-45 |
| SMX73793.1 | 627215 | 625421 | - | 597 | LinH<br>LinE<br>LinG<br>PF12697<br>TIGR02427 | LinH<br>LinE<br>LinG<br>Abhydrolase_6<br>TIGR02427 | LinH (Fusion of LinE/LinG)<br>LinE Homolog<br>LinG Homolog<br>Alpha/beta hydrolase family<br>protocat_pcaD: 3-oxoadipate enol-lactonase | 7.40E-161<br>2.00E-99<br>5.60E-70<br>1.20E-08<br>3.50E-08 |
| SMX73804.1 | 627448 | 627292 | - | 51 | LinA | LinA | LinA (linaridin precursor) | 1.80E-10 |
| SMX73815.1 | 627868 | 627718 | - | 49 | NO PFAM MATCH | - | - | - |

| index | leader | core | start | end | dir | score |
|---|---|---|---|---|---|---|
| 1 | MDTEVMDVQNFANSAMGSAEAGLKTDASAPAM | TTPTAVQFVIQGSTICLIC | 627448 | 627292 | - | 32 |

### Results for RCS84136.1 [Brevibacterium aurantiacum] back to top

previous - next

#### Architecture

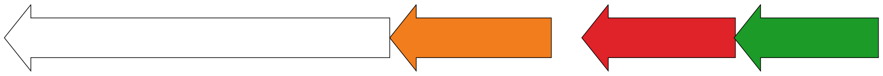

1000 nucleotides

Link to nucleotide sequence

| Accession | start | end | direction | length (aa) | Pfam/HMM | name | description | E-value |
|---|---|---|---|---|---|---|---|---|
| RCS84132.1 | 1703 | 203 | - | 499 | NO PFAM MATCH | - | - | - |
| RCS84133.1 | 2329 | 1699 | - | 209 | TIGR02673<br>TIGR01189<br>TIGR03873<br>TIGR02857<br>TIGR01166 | TIGR02673<br>TIGR01189<br>TIGR03873<br>TIGR02857<br>TIGR01166 | FtsE: cell division ATP-binding protein FtsE<br>ccmA: heme ABC exporter, ATP-binding protein CcmA<br>F420-0_ABC_ATP: proposed F420-0 ABC transporter, ATP-binding protein<br>CydD: thiol reductant ABC exporter, CydD subunit<br>cbiO: cobalt ABC transporter, ATP-binding protein | 1.30E-30<br>1.20E-26<br>1.40E-26<br>1.40E-25<br>7.10E-25 |
| RCS84134.1 | 3043 | 2449 | - | 197 | PF02441<br>TIGR00521<br>TIGR02113 | Flavoprotein<br>TIGR00521<br>TIGR02113 | Flavoprotein<br>coaBC_dfp: phosphopantothenoylcysteine decarboxylase / phosphopantothenate--cysteine ligase<br>coaC_strep: phosphopantothenoylcysteine decarboxylase | 8.00E-10<br>8.60E-10<br>4.80E-04 |
| RCS84135.1 | 3597 | 3039 | - | 185 | LinL | LinL | LinL | 1.10E-45 |

| index | leader | core | start | end | dir | score |
|---|---|---|---|---|---|---|

### Results for WP_106848707.1 [Blastococcus sp. Marseille-P5729] back to top

previous - next

#### Architecture

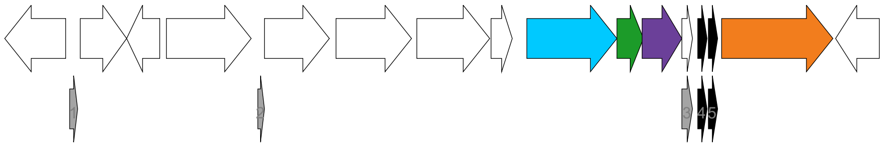

1000 nucleotides

Link to nucleotide sequence

| Accession | start | end | direction | length (aa) | Pfam/HMM | name | description | E-value |
|---|---|---|---|---|---|---|---|---|
| WP_106849284.1 | 1175256 | 1173975 | - | 426 | TIGR03449<br>TIGR02149<br>PF00534<br>TIGR03088<br>PF13579 | TIGR03449<br>TIGR02149<br>Glycos_transf_1<br>TIGR03088<br>Glyco_trans_4_4 | mycothiol_MshA: D-inositol-3-phosphate glycosyltransferase<br>glgA_Coryne: glycogen synthase, Corynebacterium family<br>Glycosyl transferases group 1<br>stp2: sugar transferase, PEP-CTERM/EpsH1 system associated<br>Glycosyl transferase 4-like domain | 4.20E-31<br>2.40E-30<br>5.40E-30<br>1.90E-25<br>5.50E-25 |

| | | | | | | | | |
|---|---|---|---|---|---|---|---|---|
| WP_158716262.1 | 1175568 | 1176525 | + | 318 | NO PFAM MATCH | - | - | - |
| WP_106848702.1 | 1177223 | 1176521 | - | 233 | PF13472 | Lipase_GDSL_2 | GDSL-like Lipase/Acylhydrolase family | 1.20E-08 |
| WP_106848703.1 | 1177341 | 1179123 | + | 593 | PF12806<br>PF00441<br>PF02770<br>TIGR03207<br>PF02771 | Acyl-CoA_dh_C<br>Acyl-CoA_dh_1<br>Acyl-CoA_dh_M<br>TIGR03207<br>Acyl-CoA_dh_N | Acetyl-CoA dehydrogenase C-terminal like<br>Acyl-CoA dehydrogenase, C-terminal domain<br>Acyl-CoA dehydrogenase, middle domain<br>cyc_hxne_CoA_dh: cyclohexanecarboxyl-CoA dehydrogenase<br>Acyl-CoA dehydrogenase, N-terminal domain | 1.60E-24<br>1.60E-19<br>1.40E-14<br>2.30E-10<br>3.00E-09 |
| WP_106848704.1 | 1179392 | 1180730 | + | 445 | TIGR02279<br>TIGR02437<br>TIGR02441<br>TIGR02440<br>PF02737 | TIGR02279<br>TIGR02437<br>TIGR02441<br>TIGR02440<br>3HCDH_N | PaaC-3OHAcCoADH: 3-hydroxyacyl-CoA dehydrogenase PaaC<br>FadB: fatty oxidation complex, alpha subunit FadB<br>fa_ox_alpha_mit: fatty acid oxidation complex, alpha subunit, mitochondrial<br>FadJ: fatty oxidation complex, alpha subunit FadJ<br>3-hydroxyacyl-CoA dehydrogenase, NAD binding domain | 6.00E-82<br>3.00E-80<br>1.30E-77<br>2.60E-73<br>1.50E-65 |
| WP_158716263.1 | 1180871 | 1182458 | + | 528 | PF18202<br>PF08341<br>TIGR03934 | TQ<br>TED<br>TIGR03934 | T-Q ester bond containing domain<br>Thioester domain<br>TQXA_dom: TQXA domain | 5.40E-21<br>2.40E-17<br>8.00E-15 |
| WP_106848706.1 | 1182571 | 1184068 | + | 498 | PF00743<br>PF13738<br>PF07992<br>PF13450<br>TIGR04018 | FMO-like<br>Pyr_redox_3<br>Pyr_redox_2<br>NAD_binding_8<br>TIGR04018 | Flavin-binding monooxygenase-like<br>Pyridine nucleotide-disulphide oxidoreductase<br>Pyridine nucleotide-disulphide oxidoreductase<br>NAD(P)-binding Rossmann-like domain<br>Bthiol_YpdA: putative bacillithiol system oxidoreductase, YpdA family | 1.30E-13<br>3.80E-11<br>3.40E-10<br>1.40E-09<br>2.20E-09 |
| WP_106849285.1 | 1184117 | 1184552 | + | 144 | PF00355<br>TIGR01416<br>TIGR02694<br>PF10518<br>TIGR01409 | Rieske<br>TIGR01416<br>TIGR02694<br>TAT_signal<br>TIGR01409 | Rieske [2Fe-2S] domain<br>Rieske_proteo: ubiquinol-cytochrome c reductase, iron-sulfur subunit<br>arsenite_ox_S: arsenite oxidase, small subunit<br>TAT (twin-arginine translocation) pathway signal sequence<br>TAT_signal_seq: Tat (twin-arginine translocation) pathway signal sequence | 8.80E-15<br>2.90E-11<br>1.60E-05<br>1.20E-04<br>1.40E-04 |
| WP_106848707.1 | 1184855 | 1186730 | + | 624 | LinH<br>LinG<br>LinE<br>PF12697<br>TIGR02427 | LinH<br>LinG<br>LinE<br>Abhydrolase_6<br>TIGR02427 | LinH (Fusion of LinE/LinG)<br>LinG Homolog<br>LinE Homolog<br>Alpha/beta hydrolase family<br>protocat_pcaD: 3-oxoadipate enol-lactonase | 6.30E-223<br>8.00E-94<br>3.80E-93<br>2.00E-13<br>4.90E-10 |
| WP_106848708.1 | 1186730 | 1187270 | + | 179 | LinL | LinL | LinL | 3.20E-62 |
| WP_106848709.1 | 1187262 | 1188072 | + | 269 | PF13649<br>PF08241<br>PF08242<br>PF13847<br>PF13489 | Methyltransf_25<br>Methyltransf_11<br>Methyltransf_12<br>Methyltransf_31<br>Methyltransf_23 | Methyltransferase domain<br>Methyltransferase domain<br>Methyltransferase domain<br>Methyltransferase domain<br>Methyltransferase domain | 3.20E-15<br>5.00E-13<br>1.50E-12<br>6.40E-11<br>7.10E-11 |
| WP_106848710.1 | 1188076 | 1188286 | + | 69 | NO PFAM MATCH | - | - | - |
| WP_158716264.1 | 1188394 | 1188598 | + | 67 | LinA | LinA | LinA (linaridin precursor) | 7.00E-17 |
| WP_158716265.1 | 1188632 | 1188827 | + | 64 | LinA | LinA | LinA (linaridin precursor) | 1.80E-14 |
| WP_158716266.1 | 1188894 | 1191219 | + | 774 | TIGR02204<br>TIGR00958<br>TIGR02203<br>TIGR03796<br>TIGR01846 | TIGR02204<br>TIGR00958<br>TIGR02203<br>TIGR03796<br>TIGR01846 | MsbA_rel: ABC transporter, permease/ATP-binding protein<br>3a01208: antigen peptide transporter 2<br>MsbA_lipidA: lipid A export permease/ATP-binding protein MsbA<br>NHLM_micro_ABC1: NHLM bacteriocin system ABC transporter, peptidase/ATP-binding protein<br>type_I_sec_HlyB: type I secretion system ATPase | 1.20E-98<br>2.50E-97<br>2.80E-96<br>4.70E-84<br>8.80E-81 |
| WP_106848712.1 | 1192175 | 1191263 | - | 303 | PF00196<br>PF11495<br>PF13384<br>TIGR03541<br>PF13518 | GerE<br>Regulator_TrmB<br>HTH_23<br>TIGR03541<br>HTH_28 | Bacterial regulatory proteins, luxR family<br>Archaeal transcriptional regulator TrmB<br>Homeodomain-like domain<br>reg_near_HchA: LuxR family transcriptional regulatory, chaperone HchA-associated<br>Helix-turn-helix domain | 3.00E-05<br>4.60E-05<br>1.70E-04<br>4.40E-04<br>8.80E-04 |

| index | leader | core | start | end | dir | score |
|---|---|---|---|---|---|---|
| 1 | LSSWAGMSRDYSPVPARVGHGAVPPPR | RPGESGPGAGARHWNQIVTEPVPAPAL | 1175331 | 1175496 | + | 6 |
| 2 | LRRSCRNRGQLVQTCGRRHRSGRC | SISRTAGAARAGPPFAITPVDGVE | 1179249 | 1179396 | + | 6 |
| 3 | MQSKQMLAVTLTFIIAAVA | LTFAFMQFFDSAAIAIGPAIAVAVGIAAAVYFAMRDKQGSATGDQREMRR | 1188076 | 1188286 | + | 6 |
| 4 | MRNLARNVNENPSFDLGDLTPTTEGRKLDPHAPVL | ATPAAAAAATVGAAATVGAGVAGFVAEEAADG | 1188394 | 1188598 | + | 31 |
| 5 | MNRIEKINNSAELAELPIDADQPTKLGDMPPVT | ATPGALAGGVAAGAALVGAAAGGAAVGEAID | 1188632 | 1188827 | + | 30 |

### Results for WP_101574944.1 [Brevibacterium sp. 239c] back to top

previous - next

#### Architecture

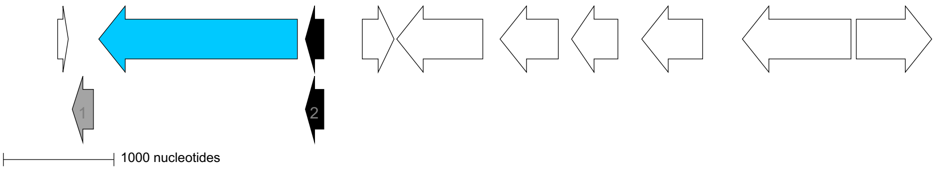

Link to nucleotide sequence

| Accession | start | end | direction | length (aa) | Pfam/HMM | name | description | E-value |
|---|---|---|---|---|---|---|---|---|
| WP_145997438.1 | 0 | 98 | + | 31 | PF13683 | rve_3 | Integrase core domain | 6.80E-05 |

| | | | | | | | | |
|---|---|---|---|---|---|---|---|---|
| WP_101574944.1 | 2160 | 366 | - | 597 | LinH<br>LinE<br>LinG<br>PF12697<br>TIGR03611 | LinH<br>LinE<br>LinG<br>Abhydrolase_6<br>TIGR03611 | LinH (Fusion of LinE/LinG)<br>LinE Homolog<br>LinG Homolog<br>Alpha/beta hydrolase family<br>RutD: pyrimidine utilization protein D | 3.30E-161<br>3.10E-97<br>8.10E-69<br>9.40E-09<br>1.50E-07 |
| WP_145997504.1 | 2411 | 2237 | - | 57 | LinA | LinA | LinA (linaridin precursor) | 1.60E-10 |
| WP_101574966.1 | 2750 | 3041 | + | 96 | PF02586 | SRAP | SOS response associated peptidase (SRAP) | 1.90E-11 |
| WP_145997505.1 | 3841 | 3064 | - | 258 | NO PFAM MATCH | - | - | - |
| WP_145997506.1 | 4532 | 3995 | - | 178 | NO PFAM MATCH | - | - | - |
| WP_101574947.1 | 5065 | 4651 | - | 137 | PF00239<br>PF02796<br>PF01527<br>PF13551<br>PF13518 | Resolvase<br>HTH_7<br>HTH_Tnp_1<br>HTH_29<br>HTH_28 | Resolvase, N terminal domain<br>Helix-turn-helix domain of resolvase<br>Transposase<br>Winged helix-turn helix<br>Helix-turn-helix domain | 6.10E-18<br>7.30E-09<br>2.20E-06<br>1.10E-04<br>1.50E-04 |
| WP_101574948.1 | 5827 | 5281 | - | 181 | PF02517 | CPBP | CPBP intramembrane metalloprotease | 8.40E-11 |
| WP_101574949.1 | 7175 | 6194 | - | 326 | PF01370<br>PF13460<br>PF04321<br>TIGR01746<br>TIGR01777 | Epimerase<br>NAD_binding_10<br>RmlD_sub_bind<br>TIGR01746<br>TIGR01777 | NAD dependent epimerase/dehydratase family<br>NAD(P)H-binding<br>RmlD substrate binding domain<br>Thioester-redct: thioester reductase domain<br>yfcH: TIGR01777 family protein | 6.30E-14<br>6.20E-09<br>9.10E-08<br>5.00E-07<br>2.00E-06 |
| WP_145997507.1 | 7224 | 7908 | + | 227 | NO PFAM MATCH | - | - | - |

| index | leader | core | start | end | dir | score |
|---|---|---|---|---|---|---|
| 1 | LEPMSQRQHCKIHQILTIWICVLTWEDGQCRAG | GFSRPTQATRTYTFWCVLAMLKEHRSAIGKTSP | 325 | 124 | - | 6 |
| 2 | MSERNAMDTEVMDVQNFANSAMGSAEAGLKADASAPAM | TTPTAVQFVIQGSTICLIC | 2411 | 2237 | - | 28 |

### Results for WP_165883987.1 [Brevibacterium luteolum] back to top

previous - next

#### Architecture

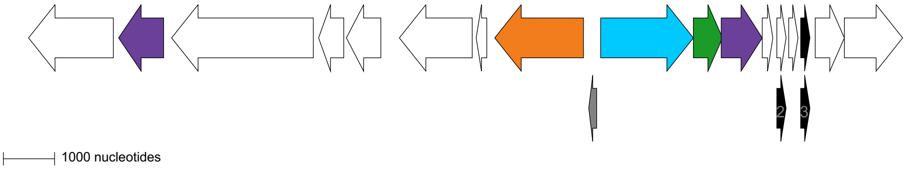

Link to nucleotide sequence

| Accession | start | end | direction | length (aa) | Pfam/HMM | name | description | E-value |
|---|---|---|---|---|---|---|---|---|
| WP_165883979.1 | 2189589 | 2187942 | - | 548 | PF02720<br>PF01844 | DUF222<br>HNH | Domain of unknown function (DUF222)<br>HNH endonuclease | 3.20E-21<br>8.40E-08 |
| WP_165883980.1 | 2190583 | 2189716 | - | 288 | TIGR00755<br>PF00398<br>PF13649 | TIGR00755<br>RrnaAD<br>Methyltransf_25 | ksgA: ribosomal RNA small subunit methyltransferase A<br>Ribosomal RNA adenine dimethylase<br>Methyltransferase domain | 1.50E-42<br>1.10E-37<br>2.10E-04 |
| WP_165883981.1 | 2193516 | 2190741 | - | 924 | PF00723<br>TIGR00685<br>PF02358<br>TIGR01577<br>TIGR01484 | Glyco_hydro_15<br>TIGR00685<br>Trehalose_PPase<br>TIGR01577<br>TIGR01484 | Glycosyl hydrolases family 15<br>T6PP: trehalose-phosphatase<br>Trehalose-phosphatase<br>oligosac_amyl: oligosaccharide amylase<br>HAD-SF-IIB: HAD hydrolase, family IIB | 1.20E-36<br>1.70E-29<br>1.00E-23<br>2.20E-12<br>1.20E-09 |
| WP_165883982.1 | 2194100 | 2193620 | - | 159 | PF03061<br>TIGR02286<br>TIGR00369 | 4HBT<br>TIGR02286<br>TIGR00369 | Thioesterase superfamily<br>PaaD: phenylacetic acid degradation protein PaaD<br>unchar_dom_1: uncharacterized domain 1 | 3.40E-13<br>5.60E-04<br>8.10E-04 |
| WP_165883983.1 | 2194821 | 2194161 | - | 219 | NO PFAM MATCH | - | - | - |
| WP_165883984.1 | 2196611 | 2195192 | - | 472 | TIGR02734<br>TIGR03467<br>PF13450<br>TIGR00292<br>PF01266 | TIGR02734<br>TIGR03467<br>NAD_binding_8<br>TIGR00292<br>DAO | crtI_fam: phytoene desaturase<br>HpnE: squalene-associated FAD-dependent desaturase<br>NAD(P)-binding Rossmann-like domain<br>TIGR00292: thiazole biosynthesis enzyme<br>FAD dependent oxidoreductase | 1.40E-12<br>2.10E-12<br>5.30E-11<br>1.40E-07<br>1.60E-07 |
| WP_165883985.1 | 2196894 | 2196675 | - | 72 | NO PFAM MATCH | - | - | - |
| WP_165883986.1 | 2198793 | 2197056 | - | 578 | TIGR02203<br>TIGR02204<br>TIGR00958<br>TIGR03375<br>TIGR03796 | TIGR02203<br>TIGR02204<br>TIGR00958<br>TIGR03375<br>TIGR03796 | MsbA_lipidA: lipid A export permease/ATP-binding protein MsbA<br>MsbA_rel: ABC transporter, permease/ATP-binding protein<br>3a01208: antigen peptide transporter 2<br>type_I_sec_LssB: type I secretion system ATPase<br>NHLM_micro_ABC1: NHLM bacteriocin system ABC transporter, peptidase/ATP-binding protein | 2.00E-89<br>2.80E-86<br>7.00E-82<br>1.40E-75<br>6.80E-71 |
| WP_165883987.1 | 2199108 | 2200938 | + | 609 | LinH<br>LinG<br>LinE<br>PF12146<br>PF12697 | LinH<br>LinG<br>LinE<br>Hydrolase_4<br>Abhydrolase_6 | LinH (Fusion of LinE/LinG)<br>LinG Homolog<br>LinE Homolog<br>Serine aminopeptidase, S33<br>Alpha/beta hydrolase family | 3.30E-177<br>3.50E-87<br>9.80E-65<br>5.00E-13<br>6.30E-13 |
| WP_165883988.1 | 2200934 | 2201486 | + | 183 | LinL | LinL | LinL | 9.20E-52 |

| | | | | | | | | |
|---|---|---|---|---|---|---|---|---|
| WP_165883989.1 | 2201482 | 2202277 | + | 264 | PF13649<br>PF13847<br>PF13489<br>PF03848<br>TIGR00477 | Methyltransf_25<br>Methyltransf_31<br>Methyltransf_23<br>TehB<br>TIGR00477 | Methyltransferase domain<br>Methyltransferase domain<br>Methyltransferase domain<br>Tellurite resistance protein TehB<br>tehB: tellurite resistance protein TehB | 2.70E-14<br>1.60E-10<br>1.90E-08<br>3.90E-08<br>5.50E-08 |
| WP_165883990.1 | 2202263 | 2202491 | + | 75 | NO PFAM MATCH | - | - | - |
| WP_165883991.1 | 2202569 | 2202740 | + | 56 | NO PFAM MATCH | - | - | - |
| WP_165883992.1 | 2202792 | 2202972 | + | 59 | NO PFAM MATCH | - | - | - |
| WP_165883993.1 | 2203028 | 2203205 | + | 58 | LinA | LinA | LinA (linaridin precursor) | 6.80E-06 |
| WP_165883994.1 | 2203306 | 2203876 | + | 189 | PF07931<br>PF13238<br>PF13671 | CPT<br>AAA_18<br>AAA_33 | Chloramphenicol phosphotransferase-like protein<br>AAA domain<br>AAA domain | 6.80E-30<br>2.20E-07<br>5.00E-07 |
| WP_165883995.1 | 2203872 | 2205009 | + | 378 | PF00144 | Beta-lactamase | Beta-lactamase | 3.30E-40 |

| index | leader | core | start | end | dir | score |
|---|---|---|---|---|---|---|
| 1 | MGVGWEDRVMGVGWDRSRERIGCGTCT | DGSTITHHRFAGLVEDGGQTLISDRVHA | 2199052 | 2198884 | - | 7 |
| 2 | MSIADSLNATKSVDIKAAGSTYSSPEAFVG | ATPGVAIGSAMVVAAIKGHQMAQADD | 2202569 | 2202740 | + | 19 |
| 3 | MSRFTDTLAQKSAPLTVLSR | STAIDPAGQMISATPVAAGAAVVGSAVVAYAIEEAQDD | 2203028 | 2203205 | + | 18 |

### Results for SDS21300.1 [Brevibacterium sandarakinum] back to top

previous - next

#### Architecture

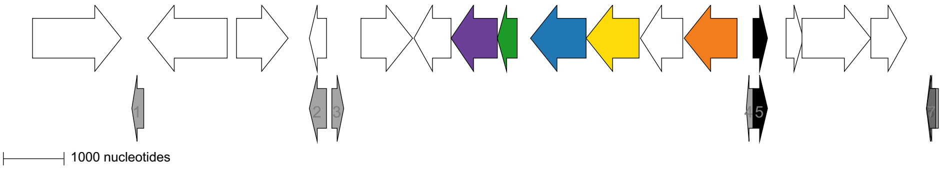

Link to nucleotide sequence

| Accession | start | end | direction | length (aa) | Pfam/HMM | name | description | E-value |
|---|---|---|---|---|---|---|---|---|
| SDS20963.1 | 1558578 | 1560033 | + | 484 | TIGR00909<br>PF13520<br>PF00324<br>TIGR00906<br>TIGR00908 | TIGR00909<br>AA_permease_2<br>AA_permease<br>TIGR00906<br>TIGR00908 | 2A0306: amino acid transporter<br>Amino acid permease<br>Amino acid permease<br>2A0303: cationic amino acid transport permease<br>2A0305: ethanolamine permease | 1.60E-52<br>4.30E-40<br>4.10E-32<br>1.70E-29<br>8.10E-29 |
| SDS21004.1 | 1561781 | 1560476 | - | 434 | PF01571<br>TIGR00528<br>TIGR01372<br>PF08669 | GCV_T<br>TIGR00528<br>TIGR01372<br>GCV_T_C | Aminomethyltransferase folate-binding domain<br>gcvT: glycine cleavage system T protein<br>soxA: sarcosine oxidase, alpha subunit family<br>Glycine cleavage T-protein C-terminal barrel domain | 4.10E-43<br>1.80E-41<br>1.00E-34<br>1.00E-10 |
| SDS21039.1 | 1561938 | 1562790 | + | 283 | TIGR00676<br>PF02219<br>TIGR00677 | TIGR00676<br>MTHFR<br>TIGR00677 | fadh2: methylenetetrahydrofolate reductase [NAD(P)H]<br>Methylenetetrahydrofolate reductase<br>fadh2_euk: methylenetetrahydrofolate reductase | 2.90E-17<br>1.10E-13<br>1.80E-11 |
| SDS21076.1 | 1563407 | 1563134 | - | 90 | NO PFAM MATCH | - | - | - |
| SDS21131.1 | 1563988 | 1564843 | + | 284 | PF00144 | Beta-lactamase | Beta-lactamase | 1.50E-23 |
| SDS21181.1 | 1565464 | 1564861 | - | 200 | PF00196<br>TIGR03020<br>TIGR03541<br>PF08281 | GerE<br>TIGR03020<br>TIGR03541<br>Sigma70_r4_2 | Bacterial regulatory proteins, luxR family<br>EpsA: transcriptional regulator EpsA<br>reg_near_HchA: LuxR family transcriptional regulatory, chaperone HchA-associated<br>Sigma-70, region 4 | 3.50E-15<br>2.40E-09<br>9.00E-09<br>8.20E-04 |

| | | | | | | | | |
|---|---|---|---|---|---|---|---|---|
| SDS21228.1 | 1566231 | 1565460 | - | 256 | PF13649<br>PF08241<br>PF13489<br>TIGR01983<br>TIGR02021 | Methyltransf_25<br>Methyltransf_11<br>Methyltransf_23<br>TIGR01983<br>TIGR02021 | Methyltransferase domain<br>Methyltransferase domain<br>Methyltransferase domain<br>UbiG: 3-demethylubiquinone-9 3-O-methyltransferase<br>BchM-ChlM: magnesium protoporphyrin O-methyltransferase | 1.30E-12<br>4.80E-11<br>1.20E-10<br>7.90E-08<br>1.20E-07 |
| SDS21265.1 | 1566569 | 1566227 | - | 113 | LinL | LinL | LinL | 6.90E-32 |
| SDS21300.1 | 1567696 | 1566769 | - | 308 | LinE<br>LinH<br>TIGR02427<br>PF12697<br>PF00561 | LinE<br>LinH<br>TIGR02427<br>Abhydrolase_6<br>Abhydrolase_1 | LinE Homolog<br>LinH (Fusion of LinE/LinG)<br>protocat_pcaD: 3-oxoadipate enol-lactonase<br>Alpha/beta hydrolase family<br>alpha/beta hydrolase fold | 1.20E-70<br>4.00E-55<br>5.70E-13<br>3.40E-12<br>6.80E-11 |
| SDS21339.1 | 1568560 | 1567699 | - | 286 | LinG<br>LinH<br>TIGR04033<br>PF05090 | LinG<br>LinH<br>TIGR04033<br>VKG_Carbox | LinG Homolog<br>LinH (Fusion of LinE/LinG)<br>export_SdpB: antimicrobial peptide system protein, SdpB family<br>Vitamin K-dependent gamma-carboxylase | 1.70E-89<br>7.30E-78<br>1.50E-04<br>1.90E-04 |
| SDS21380.1 | 1569296 | 1568591 | - | 234 | PF01061<br>PF12698 | ABC2_membrane<br>ABC2_membrane_3 | ABC-2 type transporter<br>ABC-2 family transporter protein | 8.50E-08<br>5.40E-05 |
| SDS21421.1 | 1570191 | 1569309 | - | 293 | TIGR01188<br>TIGR03864<br>TIGR01288<br>TIGR03740<br>TIGR00968 | TIGR01188<br>TIGR03864<br>TIGR01288<br>TIGR03740<br>TIGR00968 | drrA: daunorubicin resistance ABC transporter, ATP-binding protein<br>PQQ_ABC_ATP: ABC transporter, ATP-binding subunit, PQQ-dependent alcohol dehydrogenase system<br>nodI: nodulation ABC transporter NodI<br>galliderm_ABC: lantibiotic protection ABC transporter, ATP-binding subunit<br>3a0106s01: sulfate ABC transporter, ATP-binding protein | 3.50E-57<br>9.50E-43<br>1.60E-42<br>9.80E-32<br>3.20E-31 |
| SDS21459.1 | 1570449 | 1570689 | + | 79 | LinA | LinA | LinA (linaridin precursor) | 1.50E-10 |
| SDS21509.1 | 1570999 | 1571269 | + | 89 | NO PFAM MATCH | - | - | - |
| SDS21579.1 | 1571265 | 1572387 | + | 373 | PF07730 | HisKA_3 | Histidine kinase | 3.30E-11 |
| SDS21616.1 | 1572383 | 1572989 | + | 201 | PF00072<br>TIGR02875<br>PF00196<br>TIGR02154<br>TIGR01818 | Response_reg<br>TIGR02875<br>GerE<br>TIGR02154<br>TIGR01818 | Response regulator receiver domain<br>spore_0_A: sporulation transcription factor Spo0A<br>Bacterial regulatory proteins, luxR family<br>PhoB: phosphate regulon transcriptional regulatory protein PhoB<br>ntrC: nitrogen regulation protein NR(I) | 1.30E-24<br>3.70E-17<br>3.40E-14<br>2.50E-10<br>4.80E-09 |

| index | leader | core | start | end | dir | score |
|---|---|---|---|---|---|---|
| 1 | MPVAALASAPSPFNCAPRTRSSIFYPHHYF | LPAASMTTGGFTRGTGSANSYMCTGVICCSW | 1560397 | 1560211 | - | 6 |
| 2 | MKVRTTGLMISMLLAAGVVVLVAIE...TGDSFLLPTAVLDNAWWVLVPLIAW | ITISALLVGRAQRSKRTAK | 1563407 | 1563134 | - | 6 |
| 3 | MSYPWFRAPTGTDCDPSRNIQKNRDCVGSFAVPSLDIQPSK | PTPYQKLRVQELLCFATTTIRPIG | 1563502 | 1563700 | + | 6 |
| 4 | LSVSCYYMCDLNGRIH | YLPDTTTVSKREGSLNP | 1570447 | 1570345 | - | 6 |
| 5 | MQNRITRFADSAEITRIGRNIDAQPVL | ATPVTTTATTAVTCITAAFGSHDSE...DVTSSFESDETVDGLLKARVVAMSS | 1570449 | 1570689 | + | 25 |
| 6 | VYGVDGRVVAAVGISGPTMRIADHH | ETVGKLLVDEAGLLSRSLKRKVTHR | 1573452 | 1573299 | - | 6 |
| 7 | VHEEFEEGLDALAAPVYGVDGRVVAAVGISGP | TMRIADHHETVGKLLVDEAGLLSRSLKRKVTHR | 1573497 | 1573299 | - | 6 |

### Results for WP_127362990.1 [Brevibacterium aurantiacum] back to top

previous - next

#### Architecture

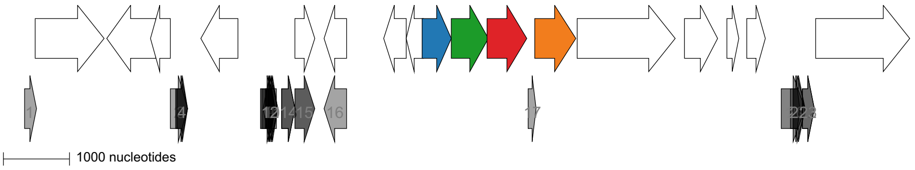

Link to nucleotide sequence

| Accession | start | end | direction | length (aa) | Pfam/HMM | name | description | E-value |
|---|---|---|---|---|---|---|---|---|
| WP_096147163.1 | 2053667 | 2054693 | + | 341 | TIGR02224<br>TIGR02225<br>PF00589<br>TIGR02249<br>PF02899 | TIGR02224<br>TIGR02225<br>Phage_integrase<br>TIGR02249<br>Phage_int_SAM_1 | recomb_XerC: tyrosine recombinase XerC<br>recomb_XerD: tyrosine recombinase XerD<br>Phage integrase family<br>integrase_gron: integron integrase<br>Phage integrase, N-terminal SAM-like domain | 4.30E-47<br>3.00E-44<br>1.50E-30<br>6.80E-12<br>1.30E-11 |
| WP_127362988.1 | 2055474 | 2054739 | - | 244 | PF13276<br>PF00665 | HTH_21<br>rve | HTH-like domain<br>Integrase core domain | 4.00E-17<br>2.10E-12 |
| WP_009883477.1 | 2055710 | 2055398 | - | 103 | PF01527 | HTH_Tnp_1 | Transposase | 5.70E-14 |
| WP_125178125.1 | 2056726 | 2056168 | - | 185 | NO PFAM MATCH | - | - | - |
| WP_127362989.1 | 2057582 | 2057885 | + | 100 | PF02861<br>TIGR02639 | Clp_N<br>TIGR02639 | Clp amino terminal domain, pathogenicity island component<br>ClpA: ATP-dependent Clp protease ATP-binding subunit ClpA | 2.20E-04<br>4.80E-04 |
| WP_125178127.1 | 2058370 | 2058022 | - | 115 | NO PFAM MATCH | - | - | - |
| WP_009885501.1 | 2059271 | 2058938 | - | 110 | NO PFAM MATCH | - | - | - |
| WP_081448679.1 | 2059528 | 2059267 | - | 86 | PF13936<br>PF13384<br>PF01381<br>PF13542<br>TIGR04111 | HTH_38<br>HTH_23<br>HTH_3<br>HTH_Tnp_ISL3<br>TIGR04111 | Helix-turn-helix domain<br>Homeodomain-like domain<br>Helix-turn-helix<br>Helix-turn-helix domain of transposase family ISL3<br>BcepMu_gp16: phage-associated protein, BcepMu gp16 family | 1.70E-12<br>1.80E-10<br>2.00E-08<br>8.20E-08<br>2.20E-07 |
| WP_127362990.1 | 2059520 | 2059961 | + | 146 | LinE<br>LinH | LinE<br>LinH | LinE Homolog<br>LinH (Fusion of LinE/LinG) | 1.00E-30<br>5.30E-21 |
| WP_114385254.1 | 2059950 | 2060508 | + | 185 | LinL | LinL | LinL | 1.10E-45 |
| WP_114385253.1 | 2060504 | 2061098 | + | 197 | PF02441<br>TIGR00521<br>TIGR02113 | Flavoprotein<br>TIGR00521<br>TIGR02113 | Flavoprotein<br>coaBC_dfp: phosphopantothenoylcysteine decarboxylase / phosphopantothenate--cysteine ligase<br>coaC_strep: phosphopantothenoylcysteine decarboxylase | 8.00E-10<br>8.60E-10<br>4.80E-04 |
| WP_101621248.1 | 2061218 | 2061848 | + | 209 | TIGR02673<br>TIGR01189<br>TIGR03873<br>TIGR02857<br>TIGR01166 | TIGR02673<br>TIGR01189<br>TIGR03873<br>TIGR02857<br>TIGR01166 | FtsE: cell division ATP-binding protein FtsE<br>ccmA: heme ABC exporter, ATP-binding protein CcmA<br>F420-0_ABC_ATP: proposed F420-0 ABC transporter, ATP-binding protein<br>CydD: thiol reductant ABC exporter, CydD subunit<br>cbiO: cobalt ABC transporter, ATP-binding protein | 1.30E-30<br>1.20E-26<br>1.40E-26<br>1.40E-25<br>7.10E-25 |
| WP_127362991.1 | 2061868 | 2063344 | + | 491 | NO PFAM MATCH | - | - | - |
| WP_127362992.1 | 2063477 | 2063978 | + | 166 | PF03372 | Exo_endo_phos | Endonuclease/Exonuclease/phosphatase family | 9.20E-06 |
| WP_009212852.1 | 2064121 | 2064313 | + | 64 | INFERRED GENE | - | - | - |
| WP_005961783.1 | 2064435 | 2064714 | + | 93 | INFERRED GENE | - | - | - |
| WP_009883329.1 | 2065466 | 2066900 | + | 477 | TIGR04416<br>PF00078<br>PF08388 | TIGR04416<br>RVT_1<br>GIIM | group_II_RT_mat: group II intron reverse transcriptase/maturase<br>Reverse transcriptase (RNA-dependent DNA polymerase)<br>Group II intron, maturase-specific domain | 2.80E-113<br>1.30E-38<br>1.70E-23 |

| index | leader | core | start | end | dir | score |
|---|---|---|---|---|---|---|
| 1 | MREGTDVNTMLPYLSAYMGHAGIDSTLYY | VHTSPDFMDSYAELAGKTANTVIPEPDAR | 2053494 | 2053671 | + | 6 |
| 2 | MRESPSSNLPASIIPGAIHSAQYRLGDMLMRCEADCRQVRV | PPACTQGLLDSSQVKIFKYVLVSSIFSLGIYVASTSCKFPWP | 2055708 | 2055960 | + | 6 |
| 3 | MLMRCEADCRQVRVPPACTQGLLDSSQV | KIFKYVLVSSIFSLGIYVASTSCKFPWP | 2055789 | 2055960 | + | 8 |
| 4 | MRCEADCRQVRVPPACTQGLLDSSQVK | IFKYVLVSSIFSLGIYVASTSCKFPWP | 2055795 | 2055960 | + | 7 |
| 5 | VSQLGCHLSVYQSLTSAQYRLGDML...DGGVEQTVDKLGECRSKNFCGVPHV | KIPGRWNLVTVGEQEYAQSPN | 2057068 | 2057287 | + | 7 |
| 6 | VYQSLTSAQYRLGDMLVDGGVEQTVDKLGECRSKNFCGVPHV | KIPGRWNLVTVGEQEYAQSPN | 2057095 | 2057287 | + | 8 |
| 7 | MLVDGGVEQTVDKLGECRSKNFCGVPHV | KIPGRWNLVTVGEQEYAQSPN | 2057137 | 2057287 | + | 6 |
| 8 | MVVWSRLSISLESADPKTSAECLMSRSPAD | GTLSRSESRSMLSRPISLAFWIWPGTRIGE | 2057147 | 2057330 | + | 6 |
| 9 | VVWSRLSISLESADPKTSAECLMSRSPAD | GTLSRSESRSMLSRPISLAFWIWPGTRIGE | 2057150 | 2057330 | + | 6 |
| 10 | VWSRLSISLESADPKTSAECLMSRSPAD | GTLSRSESRSMLSRPISLAFWIWPGTRIGE | 2057153 | 2057330 | + | 6 |
| 11 | LESADPKTSAECLMSRSPAD | GTLSRSESRSMLSRPISLAFWIWPGTRIGE | 2057177 | 2057330 | + | 6 |
| 12 | VPGQIQKARLIGRLSILLLSDRDKV...GDLDMRHSAEVFGSALSKLIDSLLH | TTIDEHVAEAVLRGG | 2057314 | 2057107 | - | 6 |
| 13 | VAVVALGVLGV | AAPASASASPSHEQQEVVALDPYVA...CRNGGGYPLNGICFGGRWDGAIIIV | 2057385 | 2057586 | + | 7 |
| 14 | VALGVLGVAAPASASASPSHEQQEVVALDPY | VAVTEATCRNGGGYPLNGICFGGRWDGAIIIV | 2057394 | 2057586 | + | 6 |
| 15 | MSLPSGELQVGRLCIDGWVTPRLEA...GVEHVMLALIAQGDSPTLVDRVVPV | ETVREWRDELSKVIASTKPEIASDSINVSVERE | 2057582 | 2057885 | + | 8 |
| 16 | MDEDQMVQFYERRRIYPECQSGWYP...ERVDKLVDAAEAKAYTTCEVCGVSD | STVTTSGPSWVKTLCTTHRGKIR | 2058370 | 2058022 | - | 6 |
| 17 | LHIFIVMVCF | STLVRTQWDTFCHNETFRMVEQSLYDRTQ | 2061115 | 2061235 | + | 6 |
| 18 | MGARAGSSPDNAETIPVLPDGSGSANTTERYTHRNQCPNAPQV | RPPTRNLVDRGRVASQTEAGVPVEG...AGVKLGTDPVNRLNSERGNPAAGPC | 2064946 | 2065300 | + | 7 |

| | | | | | | |
|---|---|---|---|---|---|---|
| 19 | VDRGRVASQTEAGVPVEGTSGSENVAGPGGHSE | SLRRSYGEDAGVKLGTDPVNRLNSERGNPAAGPC | 2065096 | 2065300 | + | 6 |
| 20 | VCQSRELPDRRMSPVR | EAIVKACGVAMARMQE | 2065133 | 2065232 | + | 6 |
| 21 | VPVEGTSGSENVAGPGGHSESLRRSYG | EDAGVKLGTDPVNRLNSERGNPAAGPC | 2065135 | 2065300 | + | 7 |
| 22 | VNVGTPPLVPADHGFHAWSAGKVRRRLI | PVVWGRRVRSSPRSGKPTTWRRDSVPIPQ | 2065265 | 2065439 | + | 6 |
| 23 | VGTPPLVPADHGFHAWSAGKVRRRLIP | VVWGRRVRSSPRSGKPTTWRRDSVPIPQ | 2065271 | 2065439 | + | 6 |

### Results for OYT91441.1 [Burkholderiales bacterium PBB3] back to top

previous - next

#### Architecture

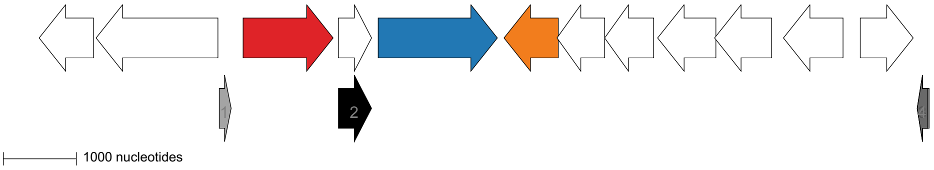

Link to nucleotide sequence

| Accession | start | end | direction | length (aa) | Pfam/HMM | name | description | E-value |
|---|---|---|---|---|---|---|---|---|
| OYT91437.1 | 742 | 0 | - | 247 | TIGR01362<br>PF00793<br>TIGR01361 | TIGR01362<br>DAHP_synth_1<br>TIGR01361 | KDO8P_synth: 3-deoxy-8-phosphooctulonate synthase<br>DAHP synthetase I family<br>DAHP_synth_Bsub: 3-deoxy-7-phosphoheptulonate synthase | 1.50E-112<br>1.60E-63<br>4.20E-28 |
| OYT91438.1 | 2450 | 782 | - | 555 | TIGR00337<br>PF06418<br>PF00117<br>TIGR01368<br>TIGR00888 | TIGR00337<br>CTP_synth_N<br>GATase<br>TIGR01368<br>TIGR00888 | PyrG: CTP synthase<br>CTP synthase N-terminus<br>Glutamine amidotransferase class-I<br>CPSaseIIsmall: carbamoyl-phosphate synthase, small subunit<br>guaA_Nterm: GMP synthase (glutamine-hydrolyzing), N-terminal domain | 7.00E-235<br>1.20E-122<br>8.30E-54<br>5.50E-10<br>3.20E-09 |
| OYT91439.1 | 2794 | 4024 | + | 409 | TIGR00521<br>PF04127<br>TIGR02113<br>PF02441<br>TIGR02114 | TIGR00521<br>DFP<br>TIGR02113<br>Flavoprotein<br>TIGR02114 | coaBC_dfp: phosphopantothenoylcysteine decarboxylase / phosphopantothenate--cysteine ligase<br>DNA / pantothenate metabolism flavoprotein<br>coaC_strep: phosphopantothenoylcysteine decarboxylase<br>Flavoprotein<br>coaB_strep: phosphopantothenate--cysteine ligase | 3.10E-121<br>2.00E-62<br>5.50E-46<br>8.10E-42<br>4.50E-12 |
| OYT91440.1 | 4110 | 4557 | + | 148 | TIGR00576<br>PF00692<br>TIGR02274 | TIGR00576<br>dUTPase<br>TIGR02274 | dut: dUTP diphosphatase<br>dUTPase<br>dCTP_deam: deoxycytidine triphosphate deaminase | 4.90E-46<br>8.00E-37<br>6.20E-08 |
| OYT91441.1 | 4644 | 6279 | + | 544 | LinE<br>LinH<br>PF12697<br>TIGR02427<br>PF00561 | LinE<br>LinH<br>Abhydrolase_6<br>TIGR02427<br>Abhydrolase_1 | LinE Homolog<br>LinH (Fusion of LinE/LinG)<br>Alpha/beta hydrolase family<br>protocat_pcaD: 3-oxoadipate enol-lactonase<br>alpha/beta hydrolase fold | 2.80E-35<br>4.40E-29<br>4.60E-13<br>9.70E-11<br>1.20E-10 |
| OYT91442.1 | 7109 | 6374 | - | 244 | TIGR03005<br>TIGR00972<br>TIGR03265<br>TIGR02315<br>TIGR00968 | TIGR03005<br>TIGR00972<br>TIGR03265<br>TIGR02315<br>TIGR00968 | ectoine_ehuA: ectoine/hydroxyectoine ABC transporter, ATP-binding protein EhuA<br>3a0107s01c2: phosphate ABC transporter, ATP-binding protein<br>PhnT2: putative 2-aminoethylphosphonate ABC transporter, ATP-binding protein<br>ABC_phnC: phosphonate ABC transporter, ATP-binding protein<br>3a0106s01: sulfate ABC transporter, ATP-binding protein | 1.80E-84<br>2.50E-66<br>1.70E-62<br>3.80E-60<br>6.20E-60 |
| OYT91443.1 | 7746 | 7095 | - | 216 | TIGR03004<br>TIGR03003<br>TIGR01726<br>PF00528 | TIGR03004<br>TIGR03003<br>TIGR01726<br>BPD_transp_1 | ectoine_ehuC: ectoine/hydroxyectoine ABC transporter, permease protein EhuC<br>ectoine_ehuD: ectoine/hydroxyectoine ABC transporter, permease protein EhuD<br>HEQRo_perm_3TM: amino ABC transporter, permease protein, 3-TM region, His/Glu/Gln/Arg/opine family<br>Binding-protein-dependent transport system inner membrane component | 6.90E-38<br>4.50E-36<br>9.20E-27<br>7.60E-14 |
| OYT91444.1 | 8433 | 7749 | - | 227 | TIGR03003<br>TIGR03004<br>TIGR01726<br>PF00528<br>TIGR03416 | TIGR03003<br>TIGR03004<br>TIGR01726<br>BPD_transp_1<br>TIGR03416 | ectoine_ehuD: ectoine/hydroxyectoine ABC transporter, permease protein EhuD<br>ectoine_ehuC: ectoine/hydroxyectoine ABC transporter, permease protein EhuC<br>HEQRo_perm_3TM: amino ABC transporter, permease protein, 3-TM region, His/Glu/Gln/Arg/opine family<br>Binding-protein-dependent transport system inner membrane component<br>ABC_choXWV_perm: choline ABC transporter, permease protein | 5.00E-39<br>5.70E-32<br>1.60E-26<br>9.10E-17<br>2.40E-06 |
| OYT91445.1 | 9271 | 8470 | - | 266 | PF00497<br>TIGR01096<br>TIGR02995<br>TIGR04262<br>PF09084 | SBP_bac_3<br>TIGR01096<br>TIGR02995<br>TIGR04262<br>NMT1 | Bacterial extracellular solute-binding proteins, family 3<br>3A0103s03R: lysine-arginine-ornithine-binding periplasmic protein<br>ectoine_ehuB: ectoine/hydroxyectoine ABC transporter solute-binding protein EhuB<br>orph_peri_GRRM: extracellular substrate-binding orphan protein, GRRM family<br>NMT1/THI5 like | 3.50E-48<br>1.60E-37<br>4.20E-13<br>3.70E-07<br>1.20E-05 |
| OYT91446.1 | 10041 | 9267 | - | 257 | TIGR03338<br>PF07729<br>PF00392<br>TIGR03337<br>TIGR02325 | TIGR03338<br>FCD<br>GntR<br>TIGR03337<br>TIGR02325 | phnR_burk: phosphonate utilization associated transcriptional regulator<br>FCD domain<br>Bacterial regulatory proteins, gntR family<br>phnR: phosphonate utilization transcriptional regulator PhnR<br>C_P_lyase_phnF: phosphonate metabolism transcriptional regulator PhnF | 1.60E-27<br>8.10E-20<br>1.00E-15<br>1.80E-10<br>4.40E-08 |
| OYT91447.1 | 11030 | 10199 | - | 276 | PF00990<br>TIGR00254 | GGDEF<br>TIGR00254 | Diguanylate cyclase, GGDEF domain<br>GGDEF: diguanylate cyclase (GGDEF) domain | 5.40E-51<br>8.70E-46 |
| OYT91448.1 | 11258 | 11984 | + | 241 | TIGR00530<br>PF01553 | TIGR00530<br>Acyltransferase | AGP_acyltrn: 1-acylglycerol-3-phosphate O-acyltransferases<br>Acyltransferase | 4.80E-43<br>1.50E-29 |

| index | leader | core | start | end | dir | score |
|---|---|---|---|---|---|---|
| 1 | VVNGDYTGALSPNPGAFLPTPPNP | LAPLGEGGALTALFGLAFEVVMYTLNC | 2477 | 2633 | + | 6 |
| 2 | MQIDVKILDPRLTDQLPSY | ATPGSAGLDLRACLTEPLTLQPNAW...AQFNVVSEFPASQRGEGGYGSTGKA | 4110 | 4557 | + | 12 |
| 3 | VPDTAKGGDAQRLQINAQNCVHCKTCDIKDPTQNIVW | VTPEGGGGPNYAGM | 12192 | 12036 | - | 6 |
| 4 | VYEFVPDTAKGGDAQRLQINAQNCVHCKTCDIKDPTQNIVW | VTPEGGGGPNYAGM | 12204 | 12036 | - | 6 |

### Results for WP_135975511.1 [Cellulomonas shaoxiangyii] back to top

previous - next

#### Architecture

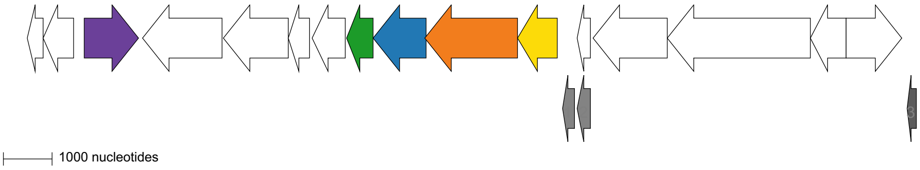

Link to nucleotide sequence

| Accession | start | end | direction | length (aa) | Pfam/HMM | name | description | E-value |
|---|---|---|---|---|---|---|---|---|
| WP_135975000.1 | 3749722 | 3749386 | - | 111 | PF13601<br>PF01022<br>PF12840<br>PF17782<br>PF12802 | HTH_34<br>HTH_5<br>HTH_20<br>DprA_WH<br>MarR_2 | Winged helix DNA-binding domain<br>Bacterial regulatory protein, arsR family<br>Helix-turn-helix domain<br>DprA winged helix domain<br>MarR family | 4.80E-17<br>3.70E-07<br>7.10E-07<br>3.80E-05<br>1.40E-04 |
| WP_135975002.1 | 3750333 | 3749718 | - | 204 | NO PFAM MATCH | - | - | - |
| WP_135975004.1 | 3750558 | 3751674 | + | 371 | PF00891<br>TIGR02716<br>PF13649<br>PF13489 | Methyltransf_2<br>TIGR02716<br>Methyltransf_25<br>Methyltransf_23 | O-methyltransferase domain<br>C20_methyl_CrtF: C-20 methyltransferase BchU<br>Methyltransferase domain<br>Methyltransferase domain | 1.50E-37<br>1.40E-13<br>2.20E-05<br>1.70E-04 |
| WP_135975009.1 | 3753393 | 3751749 | - | 547 | PF00561<br>PF08386<br>TIGR01249 | Abhydrolase_1<br>Abhydrolase_4<br>TIGR01249 | alpha/beta hydrolase fold<br>TAP-like protein<br>pro_imino_pep_1: prolyl aminopeptidase | 1.90E-20<br>2.50E-15<br>5.80E-04 |
| WP_135975007.1 | 3754760 | 3753431 | - | 442 | PF04820<br>TIGR02032<br>PF01494<br>PF01266<br>PF12831 | Trp_halogenase<br>TIGR02032<br>FAD_binding_3<br>DAO<br>FAD_oxidored | Tryptophan halogenase<br>GG-red-SF: geranylgeranyl reductase family<br>FAD binding domain<br>FAD dependent oxidoreductase<br>FAD dependent oxidoreductase | 2.60E-38<br>2.30E-35<br>2.00E-18<br>1.60E-11<br>1.50E-10 |
| WP_136225574.1 | 3755185 | 3754756 | - | 142 | PF04134 | DUF393 | Protein of unknown function, DUF393 | 2.80E-11 |
| WP_135975507.1 | 3755925 | 3755235 | - | 229 | PF00196<br>TIGR03020<br>TIGR03541<br>TIGR02983 | GerE<br>TIGR03020<br>TIGR03541<br>TIGR02983 | Bacterial regulatory proteins, luxR family<br>EpsA: transcriptional regulator EpsA<br>reg_near_HchA: LuxR family transcriptional regulatory, chaperone HchA-associated<br>SigE-fam_strep: RNA polymerase sigma-70 factor, sigma-E family | 7.50E-19<br>6.40E-13<br>3.30E-11<br>1.40E-04 |
| WP_135975509.1 | 3756499 | 3755947 | - | 183 | LinL | LinL | LinL | 5.80E-59 |
| WP_135975511.1 | 3757572 | 3756495 | - | 358 | LinE<br>LinH<br>PF12697<br>PF00561<br>TIGR02427 | LinE<br>LinH<br>Abhydrolase_6<br>Abhydrolase_1<br>TIGR02427 | LinE Homolog<br>LinH (Fusion of LinE/LinG)<br>Alpha/beta hydrolase family<br>alpha/beta hydrolase fold<br>protocat_pcaD: 3-oxoadipate enol-lactonase | 8.60E-92<br>3.20E-73<br>1.30E-13<br>4.00E-13<br>5.00E-12 |
| WP_167306028.1 | 3759464 | 3757568 | - | 631 | TIGR02203<br>TIGR02204<br>TIGR01846<br>TIGR00958<br>TIGR03796 | TIGR02203<br>TIGR02204<br>TIGR01846<br>TIGR00958<br>TIGR03796 | MsbA_lipidA: lipid A export permease/ATP-binding protein MsbA<br>MsbA_rel: ABC transporter, permease/ATP-binding protein<br>type_I_sec_HlyB: type I secretion system ATPase<br>3a01208: antigen peptide transporter 2<br>NHLM_micro_ABC1: NHLM bacteriocin system ABC transporter, peptidase/ATP-binding protein | 6.40E-108<br>2.20E-104<br>1.10E-95<br>4.20E-91<br>3.20E-87 |
| WP_135975530.1 | 3760288 | 3759469 | - | 272 | LinG<br>LinH<br>TIGR04033 | LinG<br>LinH<br>TIGR04033 | LinG Homolog<br>LinH (Fusion of LinE/LinG)<br>export_SdpB: antimicrobial peptide system protein, SdpB family | 9.60E-82<br>1.30E-78<br>8.70E-04 |
| WP_135975532.1 | 3760968 | 3760695 | - | 90 | NO PFAM MATCH | - | - | - |
| WP_135975534.1 | 3762547 | 3761023 | - | 507 | TIGR01733<br>PF00501<br>TIGR01734<br>TIGR03098<br>TIGR03443 | TIGR01733<br>AMP-binding<br>TIGR01734<br>TIGR03098<br>TIGR03443 | AA-adenyl-dom: amino acid adenylation domain<br>AMP-binding enzyme<br>D-ala-DACP-lig: D-alanine--poly(phosphoribitol) ligase, subunit 1<br>ligase_PEP_1: acyl-CoA ligase (AMP-forming), exosortase A-associated<br>alpha_am_amid: L-aminoadipate-semialdehyde dehydrogenase | 5.30E-112<br>6.40E-88<br>1.90E-59<br>2.50E-54<br>2.20E-42 |
| WP_136225527.1 | 3765483 | 3762543 | - | 979 | TIGR01746<br>TIGR03443<br>PF07993<br>PF00668<br>PF00550 | TIGR01746<br>TIGR03443<br>NAD_binding_4<br>Condensation<br>PP-binding | Thioester-redct: thioester reductase domain<br>alpha_am_amid: L-aminoadipate-semialdehyde dehydrogenase<br>Male sterility protein<br>Condensation domain<br>Phosphopantetheine attachment site | 3.20E-74<br>1.00E-59<br>1.90E-55<br>1.70E-53<br>2.50E-12 |
| WP_135973116.1 | 3766217 | 3765479 | - | 245 | PF00975<br>PF12697<br>PF00561<br>TIGR03056 | Thioesterase<br>Abhydrolase_6<br>Abhydrolase_1<br>TIGR03056 | Thioesterase domain<br>Alpha/beta hydrolase family<br>alpha/beta hydrolase fold<br>bchO_mg_che_rel: putative magnesium chelatase accessory protein | 3.40E-33<br>1.60E-07<br>1.20E-05<br>1.40E-04 |
| WP_015897483.1 | 3766213 | 3767350 | + | 379 | INFERRED GENE | - | - | - |

| index | leader | core | start | end | dir | score |
|---|---|---|---|---|---|---|
| 1 | VRPTSCCVRSGAPRAGPAWHRACAPAPACGPSHHGSRPCV | GVPRSGGARPPARSATAAVAGQAAAGALVVIPGNAHDGRGRR | 3760629 | 3760380 | - | 7 |
| 2 | MNSIENAAQRLVPAEGEAVIAPLEPVAVAAPVL | TPVAVAWAAGMAAGAAYHCYKNGCT...TMVEDNGADLSADELLHGRAELLRG | 3760968 | 3760695 | - | 7 |
| 3 | VAAADVDAAGAAASALAAALTDLAAG | CTAADGTAGVTAGDFPMSGLDDDELTGLLAALTQDGAR | 3767649 | 3767454 | - | 8 |

### Results for REE03689.1 [Citricoccus muralis] back to top

previous - next

#### Architecture

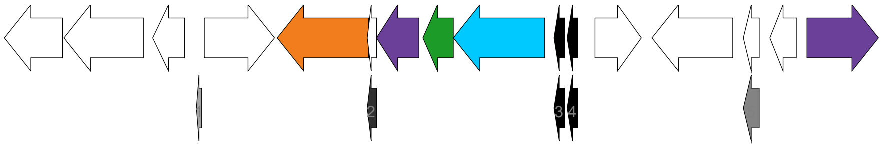

1000 nucleotides

Link to nucleotide sequence

| Accession | start | end | direction | length (aa) | Pfam/HMM | name | description | E-value |
|---|---|---|---|---|---|---|---|---|
| REE03681.1 | 1698685 | 1697548 | - | 378 | TIGR01304<br>PF00478<br>TIGR01302<br>TIGR01303<br>PF01070 | TIGR01304<br>IMPDH<br>TIGR01302<br>TIGR01303<br>FMN_dh | IMP_DH_rel_2: IMP dehydrogenase family protein<br>IMP dehydrogenase / GMP reductase domain<br>IMP_dehydrog: inosine-5'-monophosphate dehydrogenase<br>IMP_DH_rel_1: IMP dehydrogenase family protein<br>FMN-dependent dehydrogenase | 9.10E-170<br>1.40E-43<br>4.30E-37<br>3.10E-14<br>3.60E-14 |
| REE03682.1 | 1700279 | 1698734 | - | 514 | TIGR01302<br>PF00478<br>TIGR01303<br>TIGR01305<br>TIGR01306 | TIGR01302<br>IMPDH<br>TIGR01303<br>TIGR01305<br>TIGR01306 | IMP_dehydrog: inosine-5'-monophosphate dehydrogenase<br>IMP dehydrogenase / GMP reductase domain<br>IMP_DH_rel_1: IMP dehydrogenase family protein<br>GMP_reduct_1: guanosine monophosphate reductase<br>GMP_reduct_2: guanosine monophosphate reductase | 2.60E-172<br>8.80E-152<br>3.20E-78<br>7.60E-48<br>8.80E-42 |
| REE03683.1 | 1701075 | 1700472 | - | 200 | PF04203<br>TIGR01076<br>TIGR03784 | Sortase<br>TIGR01076<br>TIGR03784 | Sortase domain<br>sortase_fam: sortase<br>marine_sortase: sortase, marine proteobacterial type | 5.40E-25<br>1.20E-14<br>2.90E-09 |
| REE03684.1 | 1701472 | 1702849 | + | 458 | PF00375 | SDF | Sodium:dicarboxylate symporter family | 5.30E-112 |
| REE03685.1 | 1704687 | 1702908 | - | 592 | TIGR02204<br>TIGR02203<br>TIGR00958<br>TIGR03796<br>TIGR03375 | TIGR02204<br>TIGR02203<br>TIGR00958<br>TIGR03796<br>TIGR03375 | MsbA_rel: ABC transporter, permease/ATP-binding protein<br>MsbA_lipidA: lipid A export permease/ATP-binding protein MsbA<br>3a01208: antigen peptide transporter 2<br>NHLM_micro_ABC1: NHLM bacteriocin system ABC transporter, peptidase/ATP-binding protein<br>type_I_sec_LssB: type I secretion system ATPase | 5.40E-91<br>4.30E-88<br>3.20E-81<br>5.10E-80<br>4.20E-75 |
| REE03686.1 | 1704845 | 1704683 | - | 53 | NO PFAM MATCH | - | - | - |
| REE03687.1 | 1705690 | 1704841 | - | 282 | PF13649<br>PF08241<br>TIGR00536<br>TIGR03533<br>TIGR01983 | Methyltransf_25<br>Methyltransf_11<br>TIGR00536<br>TIGR03533<br>TIGR01983 | Methyltransferase domain<br>Methyltransferase domain<br>hemK_fam: methyltransferase, HemK family<br>L3_gln_methyl: protein-(glutamine-N5) methyltransferase, ribosomal protein L3-specific<br>UbiG: 3-demethylubiquinone-9 3-O-methyltransferase | 1.70E-10<br>2.70E-08<br>6.90E-08<br>1.70E-05<br>2.30E-05 |
| REE03688.1 | 1706355 | 1705761 | - | 197 | LinL | LinL | LinL | 7.90E-58 |
| REE03689.1 | 1708160 | 1706351 | - | 602 | LinH<br>LinG<br>LinE<br>PF12697<br>PF00561 | LinH<br>LinG<br>LinE<br>Abhydrolase_6<br>Abhydrolase_1 | LinH (Fusion of LinE/LinG)<br>LinG Homolog<br>LinE Homolog<br>Alpha/beta hydrolase family<br>alpha/beta hydrolase fold | 7.50E-192<br>1.70E-90<br>2.90E-82<br>8.40E-13<br>4.40E-12 |
| REE03690.1 | 1708537 | 1708327 | - | 69 | LinA | LinA | LinA (linaridin precursor) | 4.30E-08 |
| REE03691.1 | 1708793 | 1708583 | - | 69 | LinA | LinA | LinA (linaridin precursor) | 1.90E-08 |
| REE03692.1 | 1709133 | 1710060 | + | 308 | PF01551 | Peptidase_M23 | Peptidase family M23 | 8.10E-19 |
| REE03693.1 | 1711845 | 1710249 | - | 531 | TIGR02348<br>PF00118<br>TIGR02339<br>TIGR02345<br>TIGR02347 | TIGR02348<br>Cpn60_TCP1<br>TIGR02339<br>TIGR02345<br>TIGR02347 | GroEL: chaperonin GroL<br>TCP-1/cpn60 chaperonin family<br>thermosome_arch: thermosome, various subunits, archaeal<br>chap_CCT_eta: T-complex protein 1, eta subunit<br>chap_CCT_zeta: T-complex protein 1, zeta subunit | 4.70E-237<br>4.80E-95<br>6.80E-27<br>2.30E-21<br>5.50E-21 |
| REE03694.1 | 1712346 | 1712049 | - | 98 | PF00166 | Cpn10 | Chaperonin 10 Kd subunit | 1.70E-34 |
| REE03695.1 | 1713089 | 1712561 | - | 175 | NO PFAM MATCH | - | - | - |
| REE03696.1 | 1713283 | 1714690 | + | 468 | PF18096<br>PF13649 | Thump_like<br>Methyltransf_25 | THUMP domain-like<br>Methyltransferase domain | 4.20E-21<br>2.70E-04 |

| index | leader | core | start | end | dir | score |
|---|---|---|---|---|---|---|
| 1 | VRWWRKSPMKSSARLPQC | CIGVSSDDVTVTRDSAVRV | 1701437 | 1701323 | - | 6 |
| 2 | MKWNDLVVGGLMGIVSAVVLAQFMD | IIPAVVIAIIAGVSVAFASRTATSKSEG | 1704845 | 1704683 | - | 9 |
| 3 | MKTVTEVMNEKASYDLAYGAVAPSSAARINPVL | ATPGLVAFGAAFAKGAAAGGAVVAAGIAAYTVTGDS | 1708537 | 1708327 | - | 29 |
| 4 | MTAVAEFMNAQEINFEGGSMSTAGAPVL | GTPAAAAPVAVATCWKVAGGVAGFGAVVGAAAALFTVANGD | 1708793 | 1708583 | - | 30 |
| 5 | MSVSIKPLEDRIVVRPLEAEQTTASGLVIP | DTAKEKPQEGQVVAVGAGRIDDKGN...GTEVKVGGEEYLVLSARDVLAIVTK | 1712346 | 1712049 | - | 7 |

### Results for OJV60998.1 [Cellulomonas sp. 73-145] back to top

previous - next

#### Architecture

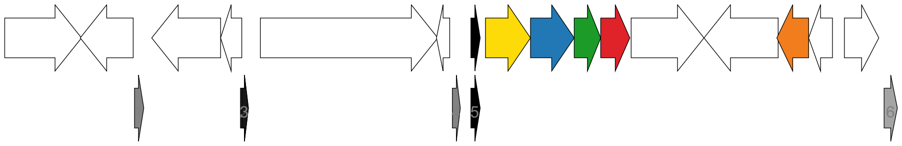

1000 nucleotides

Link to nucleotide sequence

| Accession | start | end | direction | length (aa) | Pfam/HMM | name | description | E-value |
|---|---|---|---|---|---|---|---|---|
| OJV60990.1 | 190504 | 192070 | + | 521 | TIGR00546<br>PF00795 | TIGR00546<br>CN_hydrolase | Int: apolipoprotein N-acyltransferase<br>Carbon-nitrogen hydrolase | 2.30E-72<br>1.80E-13 |
| OJV60991.1 | 193117 | 192046 | - | 356 | PF01790<br>TIGR00544 | LGT<br>TIGR00544 | Prolipoprotein diacylglyceryl transferase<br>lgt: prolipoprotein diacylglyceryl transferase | 1.10E-19<br>1.30E-06 |
| OJV60992.1 | 194906 | 193511 | - | 464 | NO PFAM MATCH | - | - | - |
| OJV60993.1 | 195328 | 194902 | - | 141 | NO PFAM MATCH | - | - | - |
| OJV60994.1 | 195731 | 199331 | + | 1199 | PF08751<br>TIGR02768<br>PF13604<br>TIGR02760<br>TIGR02686 | TrwC<br>TIGR02768<br>AAA_30<br>TIGR02760<br>TIGR02686 | TrwC relaxase<br>TraA_Ti: Ti-type conjugative transfer relaxase TraA<br>AAA domain<br>TraI_TIGR: conjugative transfer relaxase protein TraI<br>relax_trwC: conjugative relaxase domain | 5.50E-81<br>1.10E-57<br>1.00E-54<br>4.90E-30<br>1.20E-27 |
| OJV60995.1 | 199571 | 199322 | - | 82 | NO PFAM MATCH | - | - | - |
| OJV60996.1 | 200015 | 200198 | + | 60 | LinA | LinA | LinA (linaridin precursor) | 2.00E-13 |
| OJV60997.1 | 200317 | 201235 | + | 305 | LinG<br>LinH<br>TIGR04033 | LinG<br>LinH<br>TIGR04033 | LinG Homolog<br>LinH (Fusion of LinE/LinG)<br>export_SdpB: antimicrobial peptide system protein, SdpB family | 4.20E-86<br>2.20E-75<br>6.40E-05 |
| OJV60998.1 | 201224 | 202136 | + | 303 | LinE<br>LinH<br>PF12697<br>PF12146<br>PF06821 | LinE<br>LinH<br>Abhydrolase_6<br>Hydrolase_4<br>Ser_hydrolase | LinE Homolog<br>LinH (Fusion of LinE/LinG)<br>Alpha/beta hydrolase family<br>Serine aminopeptidase, S33<br>Serine hydrolase | 3.40E-79<br>4.40E-48<br>2.00E-09<br>3.90E-05<br>5.30E-04 |
| OJV60999.1 | 202132 | 202675 | + | 180 | LinL | LinL | LinL | 1.80E-47 |
| OJV61000.1 | 202667 | 203261 | + | 197 | TIGR00521<br>PF02441<br>TIGR02113 | TIGR00521<br>Flavoprotein<br>TIGR02113 | coaBC_dfp: phosphopantothenoylcysteine decarboxylase / phosphopantothenate--cysteine ligase<br>Flavoprotein<br>coaC_strep: phosphopantothenoylcysteine decarboxylase | 4.60E-12<br>4.70E-08<br>2.90E-06 |
| OJV61001.1 | 203282 | 204770 | + | 495 | TIGR02734<br>PF13450<br>TIGR03997<br>TIGR03467<br>TIGR00292 | TIGR02734<br>NAD_binding_8<br>TIGR03997<br>TIGR03467<br>TIGR00292 | crtI_fam: phytoene desaturase<br>NAD(P)-binding Rossmann-like domain<br>mycofact_OYE_2: mycofactocin system FadH/OYE family oxidoreductase 2<br>HpnE: squalene-associated FAD-dependent desaturase<br>TIGR00292: thiazole biosynthesis enzyme | 2.00E-11<br>1.40E-08<br>7.40E-08<br>1.90E-07<br>2.20E-07 |
| OJV61002.1 | 206287 | 204766 | - | 506 | NO PFAM MATCH | - | - | - |
| OJV61003.1 | 206912 | 206258 | - | 217 | TIGR01189<br>TIGR01188<br>TIGR02857<br>TIGR01166<br>PF00005 | TIGR01189<br>TIGR01188<br>TIGR02857<br>TIGR01166<br>ABC_tran | ccmA: heme ABC exporter, ATP-binding protein CcmA<br>drrA: daunorubicin resistance ABC transporter, ATP-binding protein<br>CydD: thiol reductant ABC exporter, CydD subunit<br>cbiO: cobalt ABC transporter, ATP-binding protein<br>ABC transporter | 5.10E-30<br>7.10E-27<br>7.50E-26<br>2.30E-25<br>5.10E-25 |
| OJV61061.1 | 207397 | 206908 | - | 162 | PF12840 | HTH_20 | Helix-turn-helix domain | 2.40E-08 |
| WP_013347800.1 | 207646 | 208338 | + | 230 | INFERRED GENE | - | - | - |

| index | leader | core | start | end | dir | score |
|---|---|---|---|---|---|---|
| 1 | VRGGGTETFWSVLTRGADPPVGPRSGDAFGAP | WRKKEAPGPFCVRSGPGGVGTERTGPERAVIA | 193148 | 193343 | + | 7 |
| 2 | MVTSDPPLVPGSSGASVQGAGSRCSVDD | TAPHKRVCGIGERRDAEVRCAA | 195326 | 195479 | + | 9 |
| 3 | VTSDPPLVPGSSGASVQGAGSRCSVDD | TAPHKRVCGIGERRDAEVRCAA | 195329 | 195479 | + | 7 |
| 4 | VARTVRHRPRPHRRLGRPTRMPTG | QPPGTASLTAREPVRSRTACRRSWATVTNGATPMPLV | 199624 | 199810 | + | 7 |
| 5 | MAQANTLAPDSSIAQAVNEVFADVQAGFVNRHNEPAM | TTPAIVYEAAVTVVAGSTICLIC | 200015 | 200198 | + | 33 |
| 6 | LCASTCDDVYRRLPPAQESRSRLPTRGQHRLTTPREGFTGSDVATG | STPLAAVDSIGTAQPLSQDHARRPGVGSLAMNAMRGGRVPAGRGRRDLR | 208439 | 208727 | + | 6 |

### Results for OQJ64340.1 [Clavibacter michiganensis subsp. tessellarius] back to top

previous - next

#### Architecture

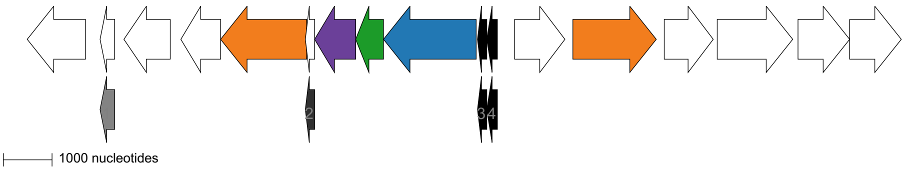

Link to nucleotide sequence

| Accession | start | end | direction | length (aa) | Pfam/HMM | name | description | E-value |
|---|---|---|---|---|---|---|---|---|
| OQJ62162.1 | 779579 | 778373 | - | 401 | PF01590<br>PF13472 | GAF<br>Lipase_GDSL_2 | GAF domain<br>GDSL-like Lipase/Acylhydrolase family | 1.90E-08<br>1.70E-05 |
| OQJ62163.1 | 780181 | 779881 | - | 99 | NO PFAM MATCH | - | - | - |
| OQJ62164.1 | 781328 | 780371 | - | 318 | PF05067 | Mn_catalase | Manganese containing catalase | 2.40E-64 |
| OQJ62165.1 | 782352 | 781533 | - | 272 | PF01551 | Peptidase_M23 | Peptidase family M23 | 5.90E-14 |

| | | | | | | | | |
|---|---|---|---|---|---|---|---|---|
| OQJ62166.1 | 784106 | 782348 | - | 585 | TIGR03375<br>TIGR02868<br>TIGR02203<br>TIGR02857<br>TIGR02204 | TIGR03375<br>TIGR02868<br>TIGR02203<br>TIGR02857<br>TIGR02204 | type_I_sec_LssB: type I secretion system ATPase<br>CydC: thiol reductant ABC exporter, CydC subunit<br>MsbA_lipidA: lipid A export permease/ATP-binding protein MsbA<br>CydD: thiol reductant ABC exporter, CydD subunit<br>MsbA_rel: ABC transporter, permease/ATP-binding protein | 4.00E-67<br>2.60E-60<br>1.10E-59<br>8.90E-59<br>1.50E-57 |
| OQJ62167.1 | 784288 | 784102 | - | 61 | NO PFAM MATCH | - | - | - |
| OQJ62168.1 | 785133 | 784290 | - | 280 | PF13649<br>PF08241<br>PF08242<br>PF13489<br>PF13847 | Methyltransf_25<br>Methyltransf_11<br>Methyltransf_12<br>Methyltransf_23<br>Methyltransf_31 | Methyltransferase domain<br>Methyltransferase domain<br>Methyltransferase domain<br>Methyltransferase domain<br>Methyltransferase domain | 1.90E-20<br>2.30E-17<br>3.70E-14<br>2.90E-12<br>4.10E-12 |
| OQJ62169.1 | 785705 | 785129 | - | 191 | LinL | LinL | LinL | 2.60E-51 |
| OQJ64340.1 | 787594 | 785701 | - | 630 | LinE<br>LinH<br>PF12697<br>PF12146<br>PF00561 | LinE<br>LinH<br>Abhydrolase_6<br>Hydrolase_4<br>Abhydrolase_1 | LinE Homolog<br>LinH (Fusion of LinE/LinG)<br>Alpha/beta hydrolase family<br>Serine aminopeptidase, S33<br>alpha/beta hydrolase fold | 6.30E-61<br>3.30E-53<br>6.70E-11<br>5.30E-06<br>4.60E-05 |
| OQJ64341.1 | 787812 | 787620 | - | 63 | LinA | LinA | LinA (linaridin precursor) | 6.90E-12 |
| OQJ62170.1 | 788048 | 787823 | - | 74 | LinA | LinA | LinA (linaridin precursor) | 4.20E-04 |
| OQJ64342.1 | 788400 | 789423 | + | 340 | TIGR01481<br>PF13377<br>TIGR02417<br>TIGR02405<br>PF00356 | TIGR01481<br>Peripla_BP_3<br>TIGR02417<br>TIGR02405<br>LacI | ccpA: catabolite control protein A<br>Periplasmic binding protein-like domain<br>fruct_sucro_rep: D-fructose-responsive transcription factor<br>trehalos_R_Ecol: trehalose operon repressor<br>Bacterial regulatory proteins, lacI family | 1.20E-42<br>1.90E-36<br>1.30E-29<br>1.50E-20<br>4.10E-19 |
| OQJ62171.1 | 789592 | 791314 | + | 573 | TIGR02633<br>TIGR03410<br>TIGR01188<br>TIGR03265<br>TIGR03864 | TIGR02633<br>TIGR03410<br>TIGR01188<br>TIGR03265<br>TIGR03864 | xylG: D-xylose ABC transporter, ATP-binding protein<br>urea_trans_UrtE: urea ABC transporter, ATP-binding protein UrtE<br>drrA: daunorubicin resistance ABC transporter, ATP-binding protein<br>PhnT2: putative 2-aminoethylphosphonate ABC transporter, ATP-binding protein<br>PQQ_ABC_ATP: ABC transporter, ATP-binding subunit, PQQ-dependent alcohol dehydrogenase system | 9.30E-106<br>4.90E-63<br>5.10E-63<br>2.50E-61<br>8.20E-59 |
| OQJ62172.1 | 791465 | 792470 | + | 334 | PF13407<br>TIGR02634<br>PF13377<br>PF00532<br>TIGR01481 | Peripla_BP_4<br>TIGR02634<br>Peripla_BP_3<br>Peripla_BP_1<br>TIGR01481 | Periplasmic binding protein domain<br>xylF: D-xylose ABC transporter, D-xylose-binding protein<br>Periplasmic binding protein-like domain<br>Periplasmic binding proteins and sugar binding domain of LacI family<br>ccpA: catabolite control protein A | 4.70E-47<br>2.80E-19<br>1.60E-10<br>5.90E-09<br>2.70E-07 |
| WP_016464282.1 | 792563 | 794111 | + | 516 | INFERRED GENE | - | - | - |
| OQJ62173.1 | 794203 | 795280 | + | 358 | PF02653<br>TIGR03409<br>TIGR03408 | BPD_transp_2<br>TIGR03409<br>TIGR03408 | Branched-chain amino acid transport system / permease component<br>urea_trans_UrtB: urea ABC transporter, permease protein UrtB<br>urea_trans_UrtC: urea ABC transporter, permease protein UrtC | 2.30E-36<br>1.40E-05<br>1.60E-05 |
| OQJ62174.1 | 795276 | 796335 | + | 352 | PF02653<br>TIGR03408<br>TIGR03727<br>TIGR03409 | BPD_transp_2<br>TIGR03408<br>TIGR03727<br>TIGR03409 | Branched-chain amino acid transport system / permease component<br>urea_trans_UrtC: urea ABC transporter, permease protein UrtC<br>urea_t_UrtC_arc: urea ABC transporter, permease protein UrtC<br>urea_trans_UrtB: urea ABC transporter, permease protein UrtB | 6.50E-49<br>5.50E-09<br>1.90E-08<br>4.50E-04 |

| index | leader | core | start | end | dir | score |
|---|---|---|---|---|---|---|
| 1 | MSAGPSRPPA | TTTALDASASGWLTALVVVQGLATI...AVALVCVLVFGFQASVTQEVSVVRQ | 780181 | 779881 | - | 7 |
| 2 | MAFTTGIVTAMMVFSIYFLMF | DAPAGTGQAQHGAVAAAIALVAGAVAYAVVRPRRRRGSSR | 784288 | 784102 | - | 9 |
| 3 | MSATHRLLGLVNSDDVRVDGVTSPVM | ATPAAFAAGVVAGGKATAVVVGAAGVGAAVGNAVGGG | 787812 | 787620 | - | 30 |
| 4 | MDLLTHVNADAQLRSHLDGTGSEVRTPIM | ASPATVAAVPAAAGAGAVAAKAAGAVVGGGAVVGAAYAAYRAVVR | 788048 | 787823 | - | 19 |

### Results for CAB1023416.1 [Corynebacterium diphtheriae] back to top

previous - next

#### Architecture

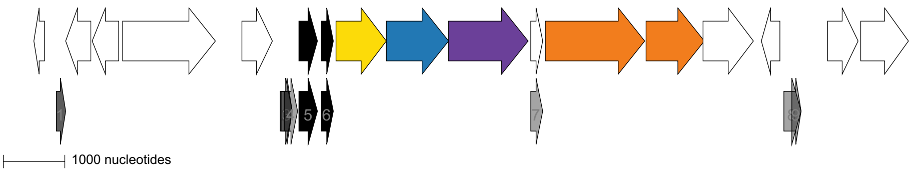

Link to nucleotide sequence

| Accession | start | end | direction | length (aa) | Pfam/HMM | name | description | E-value |
|---|---|---|---|---|---|---|---|---|
| CAB1023325.1 | 707 | 521 | - | 61 | PF13333<br>PF13683 | rve_2<br>rve_3 | Integrase core domain<br>Integrase core domain | 1.30E-13<br>4.60E-05 |
| CAB1023329.1 | 1450 | 1042 | - | 135 | PF13276 | HTH_21 | HTH-like domain | 5.50E-15 |
| CAB1023335.1 | 1902 | 1476 | - | 141 | PF06056<br>PF13384 | Terminase_5<br>HTH_23 | Putative ATPase subunit of terminase (gpP-like)<br>Homeodomain-like domain | 1.00E-13<br>7.10E-04 |

| | | | | | | | | |
|---|---|---|---|---|---|---|---|---|
| CAB1023347.1 | 1981 | 3484 | + | 500 | PF00239<br>PF07508<br>PF13408 | Resolvase<br>Recombinase<br>Zn_ribbon_recom | Resolvase, N terminal domain<br>Recombinase<br>Recombinase zinc beta ribbon domain | 5.80E-31<br>2.50E-05<br>6.60E-04 |
| CAB1023359.1 | 3920 | 4409 | + | 162 | PF00239 | Resolvase | Resolvase, N terminal domain | 2.70E-38 |
| CAB1023361.1 | 4853 | 5153 | + | 99 | LinA | LinA | LinA (linaridin precursor) | 3.00E-06 |
| CAB1023373.1 | 5219 | 5399 | + | 59 | LinA | LinA | LinA (linaridin precursor) | 6.50E-11 |
| CAB1023385.1 | 5440 | 6274 | + | 277 | LinG<br>LinH | LinG<br>LinH | LinG Homolog<br>LinH (Fusion of LinE/LinG) | 7.80E-83<br>4.60E-78 |
| CAB1023416.1 | 6270 | 7275 | + | 334 | LinE<br>LinH<br>PF12697<br>PF12146<br>PF00561 | LinE<br>LinH<br>Abhydrolase_6<br>Hydrolase_4<br>Abhydrolase_1 | LinE Homolog<br>LinH (Fusion of LinE/LinG)<br>Alpha/beta hydrolase family<br>Serine aminopeptidase, S33<br>alpha/beta hydrolase fold | 4.50E-69<br>5.50E-51<br>4.90E-12<br>7.80E-10<br>5.90E-09 |
| CAB1023429.1 | 7274 | 8585 | + | 436 | LinL<br>TIGR02469<br>PF13649<br>PF08242<br>TIGR02072 | LinL<br>TIGR02469<br>Methyltransf_25<br>Methyltransf_12<br>TIGR02072 | LinL<br>CbiT: precorrin-6Y C5,15-methyltransferase (decarboxylating), CbiT subunit<br>Methyltransferase domain<br>Methyltransferase domain<br>BioC: malonyl-acyl carrier protein O-methyltransferase BioC | 4.90E-46<br>2.50E-07<br>2.40E-06<br>2.60E-06<br>3.20E-06 |
| CAB1023438.1 | 8629 | 8803 | + | 57 | NO PFAM MATCH | - | - | - |
| CAB1023450.1 | 8857 | 10474 | + | 538 | TIGR03375<br>TIGR00958<br>TIGR02203<br>TIGR02857<br>TIGR02204 | TIGR03375<br>TIGR00958<br>TIGR02203<br>TIGR02857<br>TIGR02204 | type_I_sec_LssB: type I secretion system ATPase<br>3a01208: antigen peptide transporter 2<br>MsbA_lipidA: lipid A export permease/ATP-binding protein MsbA<br>CydD: thiol reductant ABC exporter, CydD subunit<br>MsbA_rel: ABC transporter, permease/ATP-binding protein | 1.20E-52<br>1.90E-47<br>4.20E-47<br>5.80E-45<br>2.00E-44 |
| CAB1023462.1 | 10498 | 11443 | + | 314 | TIGR03522<br>TIGR03740<br>TIGR04406<br>TIGR01188<br>TIGR03864 | TIGR03522<br>TIGR03740<br>TIGR04406<br>TIGR01188<br>TIGR03864 | GldA_ABC_ATP: gliding motility-associated ABC transporter ATP-binding subunit GldA<br>galliderm_ABC: lantibiotic protection ABC transporter, ATP-binding subunit<br>LPS_export_lptB: LPS export ABC transporter ATP-binding protein<br>drrA: daunorubicin resistance ABC transporter, ATP-binding protein<br>PQQ_ABC_ATP: ABC transporter, ATP-binding subunit, PQQ-dependent alcohol dehydrogenase system | 9.40E-46<br>5.00E-45<br>1.10E-44<br>1.80E-44<br>8.80E-36 |
| CAB1023476.1 | 11429 | 12239 | + | 269 | PF12698<br>PF12679 | ABC2_membrane_3<br>ABC2_membrane_2 | ABC-2 family transporter protein<br>ABC-2 family transporter protein | 5.10E-09<br>2.00E-07 |
| CAB1023480.1 | 12669 | 12372 | - | 98 | PF00872 | Transposase_mut | Transposase, Mutator family | 2.80E-08 |
| CAB1023483.1 | 13445 | 13949 | + | 167 | PF14690<br>PF13542 | zf-ISL3<br>HTH_Tnp_ISL3 | zinc-finger of transposase IS204/IS1001/IS1096/IS1165<br>Helix-turn-helix domain of transposase family ISL3 | 4.90E-08<br>1.00E-07 |
| CAB1023494.1 | 13995 | 14760 | + | 254 | PF01610<br>PF18047 | DDE_Tnp_ISL3<br>PatG_D | Transposase<br>PatG Domain | 3.10E-60<br>7.40E-04 |

| index | leader | core | start | end | dir | score |
|---|---|---|---|---|---|---|
| 1 | MNTCFCCHDLPV | KTVDDDRQIHATLVSSENTDVSQPQPMWMLSMKITVHQVR | 891 | 1050 | + | 8 |
| 2 | VVCVDLYGNQIQLYAGVEPSFCVEQTQG | RRLTAVSASGGKHPIIVSSGFSHLSKIKA | 4551 | 4725 | + | 6 |
| 3 | VCVDLYGNQIQLYAGVEPSFCVEQTQGR | RLTAVSASGGKHPIIVSSGFSHLSKIKA | 4554 | 4725 | + | 7 |
| 4 | LCGTDTRAETDSGVSEWGKAPYNRIVGI | FTLEQNQGINSNAAPPPQDACGCRRRRIKRMPFAVGNIVHLA | 4613 | 4826 | + | 6 |
| 5 | MQPFSGFSLSSDERLAYFSIGHFIL...QLVNDNTAFRGRLDGMAETALPPVT | ATPAIAVTAAKAAGAVIGGGAVSGAAYAAYRAVIK | 4853 | 5153 | + | 28 |
| 6 | MSRLSTLVNNDQITLEQVHSPVM | ATPAAFGAGVVAGAKACGALVAAAGVGAAIASATKK | 5219 | 5399 | + | 29 |
| 7 | MTRYFTALIAFMAVLSIAGIMFDGIDSK | QGIAITVIVSLASAAVIYWLYPFFTDLEK | 8629 | 8803 | + | 6 |
| 8 | LARSLSKARTQSRRLWYTTRCGDCFHERGPLGLSPYGRKARILRIV | ETPDLDWWCLWCRVVGILHRMWWNSCVTWFQAVIVLVVRRWNWGLA | 12741 | 13020 | + | 6 |
| 9 | VETPDLDWWCLWCRVVGILHRMW | WNSCVTWFQAVIVLVVRRWNWGLA | 12876 | 13020 | + | 6 |

### Results for WP_026168100.1 [Corynebacterium propinquum] back to top

previous - next

#### Architecture

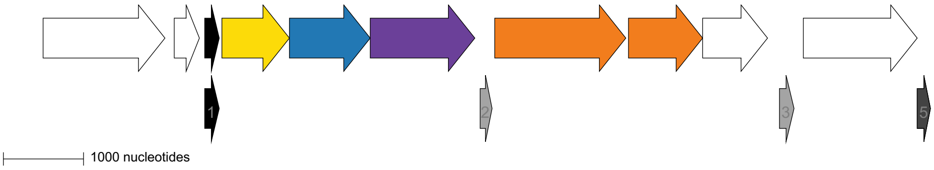

Link to nucleotide sequence

| Accession | start | end | direction | length (aa) | Pfam/HMM | name | description | E-value |
|---|---|---|---|---|---|---|---|---|
| WP_005329153.1 | 114 | 1644 | + | 510 | INFERRED GENE | - | - | - |
| WP_014301133.1 | 1747 | 2069 | + | 107 | INFERRED GENE | - | - | - |
| WP_026168099.1 | 2135 | 2315 | + | 59 | LinA | LinA | LinA (linaridin precursor) | 5.40E-11 |
| WP_018121646.1 | 2356 | 3190 | + | 277 | LinG<br>LinH<br>PF05090 | LinG<br>LinH<br>VKG_Carbox | LinG Homolog<br>LinH (Fusion of LinE/LinG)<br>Vitamin K-dependent gamma-carboxylase | 2.00E-85<br>1.90E-81<br>1.60E-04 |
| WP_026168100.1 | 3186 | 4191 | + | 334 | LinE<br>LinH<br>PF12697<br>PF12146<br>PF00561 | LinE<br>LinH<br>Abhydrolase_6<br>Hydrolase_4<br>Abhydrolase_1 | LinE Homolog<br>LinH (Fusion of LinE/LinG)<br>Alpha/beta hydrolase family<br>Serine aminopeptidase, S33<br>alpha/beta hydrolase fold | 5.80E-68<br>4.50E-49<br>3.40E-11<br>8.10E-11<br>8.80E-10 |
| WP_026168101.1 | 4190 | 5501 | + | 436 | LinL<br>TIGR02469<br>PF13649<br>PF08242<br>TIGR02072 | LinL<br>TIGR02469<br>Methyltransf_25<br>Methyltransf_12<br>TIGR02072 | LinL<br>CbiT: precorrin-6Y C5,15-methyltransferase (decarboxylating), CbiT subunit<br>Methyltransferase domain<br>Methyltransferase domain<br>BioC: malonyl-acyl carrier protein O-methyltransferase BioC | 8.70E-47<br>2.60E-06<br>8.30E-06<br>2.40E-05<br>5.00E-05 |
| WP_026168102.1 | 5749 | 7390 | + | 546 | TIGR03375<br>TIGR00958<br>TIGR02857<br>TIGR02203<br>TIGR02204 | TIGR03375<br>TIGR00958<br>TIGR02857<br>TIGR02203<br>TIGR02204 | type_I_sec_LssB: type I secretion system ATPase<br>3a01208: antigen peptide transporter 2<br>CydD: thiol reductant ABC exporter, CydD subunit<br>MsbA_lipidA: lipid A export permease/ATP-binding protein MsbA<br>MsbA_rel: ABC transporter, permease/ATP-binding protein | 2.80E-51<br>2.90E-47<br>3.30E-47<br>2.40E-45<br>5.60E-44 |
| WP_018121651.1 | 7414 | 8350 | + | 311 | TIGR03522<br>TIGR01188<br>TIGR03740<br>TIGR04406<br>TIGR03864 | TIGR03522<br>TIGR01188<br>TIGR03740<br>TIGR04406<br>TIGR03864 | GldA_ABC_ATP: gliding motility-associated ABC transporter ATP-binding subunit GldA<br>drrA: daunorubicin resistance ABC transporter, ATP-binding protein<br>galliderm_ABC: lantibiotic protection ABC transporter, ATP-binding subunit<br>LPS_export_lptB: LPS export ABC transporter ATP-binding protein<br>PQQ_ABC_ATP: ABC transporter, ATP-binding subunit, PQQ-dependent alcohol dehydrogenase system | 5.80E-48<br>9.10E-47<br>1.30E-46<br>2.50E-46<br>3.70E-37 |
| WP_018121652.1 | 8346 | 9156 | + | 269 | PF12698<br>PF12679 | ABC2_membrane_3<br>ABC2_membrane_2 | ABC-2 family transporter protein<br>ABC-2 family transporter protein | 1.30E-13<br>6.10E-08 |
| WP_049148872.1 | 9601 | 11017 | + | 471 | TIGR02734<br>TIGR03467<br>PF13450<br>PF01266<br>TIGR01317 | TIGR02734<br>TIGR03467<br>NAD_binding_8<br>DAO<br>TIGR01317 | crtI_fam: phytoene desaturase<br>HpnE: squalene-associated FAD-dependent desaturase<br>NAD(P)-binding Rossmann-like domain<br>FAD dependent oxidoreductase<br>GOGAT_sm_gam: glutamate synthase, NADH/NADPH, small subunit | 2.90E-10<br>3.40E-10<br>8.30E-10<br>3.80E-09<br>4.60E-08 |

| index | leader | core | start | end | dir | score |
|---|---|---|---|---|---|---|
| 1 | MSRLSTLVNNDQITLEQVHSPVM | ATPAAFGAGVVAGAKACGALVAAAGVGAAIATATKE | 2135 | 2315 | + | 29 |
| 2 | LIAFMVVLSIAGIMFNGIDSKQGIA | ITIIVSLASAAVIYWLYPFFTNPEK | 5566 | 5719 | + | 6 |

| | | | | | | |
|---|---|---|---|---|---|---|
| 3 | MDCFHPVAFAASMSSSVSRRLPLGTIWGANVPARSLGTITW | GCPQIMNITSLTLFLYGHTY | 9297 | 9483 | + | 6 |
| 4 | MLDGNPGFGMYITLNCAAPFFRNLPL | PTLNRKCLVMATEPNHLDLTAHFDESNHSN | 11018 | 11189 | + | 7 |
| 5 | LDGNPGFGMYITLNCAAPFFRNLPL | PTLNRKCLVMATEPNHLDLTAHFDESNHSN | 11021 | 11189 | + | 7 |

### Results for OFO12264.1 [Corynebacterium sp. HMSC22B11] back to top

previous - next

#### Architecture

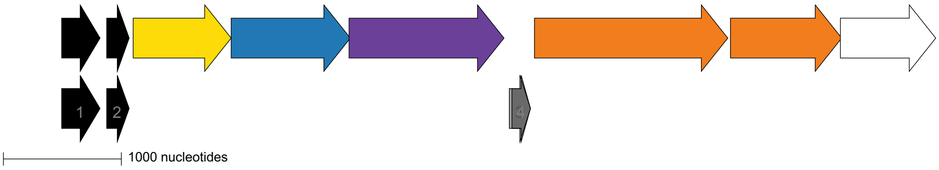

Link to nucleotide sequence

| Accession | start | end | direction | length (aa) | Pfam/HMM | name | description | E-value |
|---|---|---|---|---|---|---|---|---|
| OFO12261.1 | 152 | 473 | + | 106 | LinA | LinA | LinA (linaridin precursor) | 2.70E-06 |
| OFO12262.1 | 539 | 719 | + | 59 | LinA | LinA | LinA (linaridin precursor) | 6.50E-11 |
| OFO12263.1 | 760 | 1594 | + | 277 | LinG<br>LinH<br>PF05090 | LinG<br>LinH<br>VKG_Carbox | LinG Homolog<br>LinH (Fusion of LinE/LinG)<br>Vitamin K-dependent gamma-carboxylase | 2.00E-84<br>1.60E-78<br>1.60E-04 |
| OFO12264.1 | 1590 | 2595 | + | 334 | LinE<br>LinH<br>PF12697<br>PF12146<br>PF00561 | LinE<br>LinH<br>Abhydrolase_6<br>Hydrolase_4<br>Abhydrolase_1 | LinE Homolog<br>LinH (Fusion of LinE/LinG)<br>Alpha/beta hydrolase family<br>Serine aminopeptidase, S33<br>alpha/beta hydrolase fold | 7.90E-69<br>1.40E-50<br>3.10E-12<br>8.10E-10<br>1.70E-08 |
| OFO12265.1 | 2594 | 3905 | + | 436 | LinL<br>TIGR02469<br>PF08242<br>TIGR02072<br>PF13649 | LinL<br>TIGR02469<br>Methyltransf_12<br>TIGR02072<br>Methyltransf_25 | LinL<br>CbiT: precorrin-6Y C5,15-methyltransferase (decarboxylating), CbiT subunit<br>Methyltransferase domain<br>BioC: malonyl-acyl carrier protein O-methyltransferase BioC<br>Methyltransferase domain | 3.10E-46<br>8.20E-07<br>2.80E-06<br>6.00E-06<br>7.40E-06 |
| OFO12266.1 | 4158 | 5799 | + | 546 | TIGR03375<br>TIGR00958<br>TIGR02203<br>TIGR02857<br>TIGR02204 | TIGR03375<br>TIGR00958<br>TIGR02203<br>TIGR02857<br>TIGR02204 | type_I_sec_LssB: type I secretion system ATPase<br>3a01208: antigen peptide transporter 2<br>MsbA_lipidA: lipid A export permease/ATP-binding protein MsbA<br>CydD: thiol reductant ABC exporter, CydD subunit<br>MsbA_rel: ABC transporter, permease/ATP-binding protein | 4.30E-52<br>1.20E-47<br>1.20E-47<br>6.60E-46<br>6.90E-45 |
| OFO12267.1 | 5823 | 6768 | + | 314 | TIGR03522<br>TIGR04406<br>TIGR01188<br>TIGR03740<br>TIGR03864 | TIGR03522<br>TIGR04406<br>TIGR01188<br>TIGR03740<br>TIGR03864 | GldA_ABC_ATP: gliding motility-associated ABC transporter ATP-binding subunit GldA<br>LPS_export_lptB: LPS export ABC transporter ATP-binding protein<br>drrA: daunorubicin resistance ABC transporter, ATP-binding protein<br>galliderm_ABC: lantibiotic protection ABC transporter, ATP-binding subunit<br>PQQ_ABC_ATP: ABC transporter, ATP-binding subunit, PQQ-dependent alcohol dehydrogenase system | 2.60E-47<br>3.60E-46<br>4.60E-46<br>2.40E-45<br>1.30E-36 |
| OFO12268.1 | 6754 | 7564 | + | 269 | PF12698<br>PF12679 | ABC2_membrane_3<br>ABC2_membrane_2 | ABC-2 family transporter protein<br>ABC-2 family transporter protein | 2.90E-11<br>2.10E-07 |

| index | leader | core | start | end | dir | score |
|---|---|---|---|---|---|---|
| 1 | MLDQRLTVQPFSGFSLSSDESLAYF...QLVNDNPAFRGRLDGMAETALPPVT | ATPAIAVTAAKAAGAVIGGGAVSGAAYAAYRAVIK | 152 | 473 | + | 27 |
| 2 | MSRLSTLVNNDQITLEQVHSPVM | ATPAAFGAGVVAGAKACGALVAAAGVGAAIASATKK | 539 | 719 | + | 29 |
| 3 | MTRYFTALIAFMAVLSIAGIMFDGIDSK | QGIAITVIVSLASAAVIYWLYPFFTDLEK | 3949 | 4123 | + | 6 |
| 4 | LIAFMAVLSIAGIMFDGIDSKQGIA | ITVIVSLASAAVIYWLYPFFTDLEK | 3970 | 4123 | + | 6 |

### Results for WP_082313297.1 [Corynebacterium lactis RW2-5] back to top

previous - next

#### Architecture

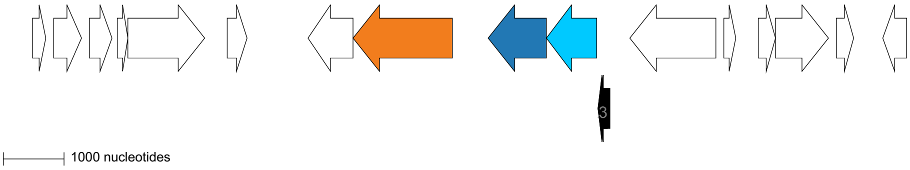

Link to nucleotide sequence

| Accession | start | end | direction | length (aa) | Pfam/HMM | name | description | E-value |
|---|---|---|---|---|---|---|---|---|
| WP_141764566.1 | 2068166 | 2068382 | + | 71 | NO PFAM MATCH | - | - | - |
| WP_053412643.1 | 2068501 | 2068960 | + | 152 | NO PFAM MATCH | - | - | - |
| WP_053412644.1 | 2069097 | 2069472 | + | 124 | NO PFAM MATCH | - | - | - |
| WP_156324815.1 | 2069558 | 2069732 | + | 57 | PF18495 | VbhA | Antitoxin VbhA | 3.30E-08 |

| | | | | | | | | |
|---|---|---|---|---|---|---|---|---|
| WP_053412645.1 | 2069728 | 2070988 | + | 419 | PF02661<br>TIGR02613 | Fic<br>TIGR02613 | Fic/DOC family<br>mob_myst_B: mobile mystery protein B | 2.10E-17<br>3.60E-07 |
| WP_053412646.1 | 2071376 | 2071688 | + | 103 | NO PFAM MATCH | - | - | - |
| WP_082313293.1 | 2073441 | 2072709 | - | 243 | PF13561<br>PF00106<br>TIGR01830<br>TIGR01829<br>TIGR04316 | adh_short_C2<br>adh_short<br>TIGR01830<br>TIGR01829<br>TIGR04316 | Enoyl-(Acyl carrier protein) reductase<br>short chain dehydrogenase<br>3oxo_ACP_reduc: 3-oxoacyl-[acyl-carrier-protein] reductase<br>AcAcCoA_reduct: acetoacetyl-CoA reductase<br>dhbA_paeA: 2,3-dihydro-2,3-dihydroxybenzoate dehydrogenase | 1.60E-46<br>3.00E-39<br>9.10E-38<br>3.60E-34<br>4.00E-34 |
| WP_082313295.1 | 2075086 | 2073433 | - | 550 | TIGR02204<br>TIGR03375<br>TIGR02203<br>TIGR00958<br>TIGR02857 | TIGR02204<br>TIGR03375<br>TIGR02203<br>TIGR00958<br>TIGR02857 | MsbA_rel: ABC transporter, permease/ATP-binding protein<br>type_I_sec_LssB: type I secretion system ATPase<br>MsbA_lipidA: lipid A export permease/ATP-binding protein MsbA<br>3a01208: antigen peptide transporter 2<br>CydD: thiol reductant ABC exporter, CydD subunit | 5.70E-81<br>3.60E-78<br>9.40E-76<br>1.10E-71<br>2.90E-70 |
| WP_082313297.1 | 2076641 | 2075666 | - | 324 | LinE<br>LinH<br>PF12697<br>PF12146<br>TIGR01250 | LinE<br>LinH<br>Abhydrolase_6<br>Hydrolase_4<br>TIGR01250 | LinE Homolog<br>LinH (Fusion of LinE/LinG)<br>Alpha/beta hydrolase family<br>Serine aminopeptidase, S33<br>pro_imino_pep_2: proline-specific peptidase | 5.60E-38<br>3.70E-34<br>7.80E-13<br>1.40E-10<br>1.80E-09 |
| WP_156324816.1 | 2077461 | 2076633 | - | 275 | LinH<br>LinG<br>PF05090<br>TIGR04033 | LinH<br>LinG<br>VKG_Carbox<br>TIGR04033 | LinH (Fusion of LinE/LinG)<br>LinG Homolog<br>Vitamin K-dependent gamma-carboxylase<br>export_SdpB: antimicrobial peptide system protein, SdpB family | 3.30E-65<br>8.70E-62<br>1.20E-07<br>2.00E-04 |
| WP_082313298.1 | 2079427 | 2078005 | - | 473 | TIGR02734<br>PF13450<br>TIGR03467<br>TIGR02733<br>PF01266 | TIGR02734<br>NAD_binding_8<br>TIGR03467<br>TIGR02733<br>DAO | crtI_fam: phytoene desaturase<br>NAD(P)-binding Rossmann-like domain<br>HpnE: squalene-associated FAD-dependent desaturase<br>desat_CrtD: C-3',4' desaturase CrtD<br>FAD dependent oxidoreductase | 5.90E-14<br>8.30E-11<br>1.60E-09<br>3.60E-08<br>3.90E-08 |
| WP_014311594.1 | 2079559 | 2079751 | + | 64 | INFERRED GENE | - | - | - |
| WP_013170013.1 | 2080133 | 2080415 | + | 93 | PF01527<br>PF13384<br>PF13022<br>PF13551<br>PF13518 | HTH_Tnp_1<br>HTH_23<br>HTH_Tnp_1_2<br>HTH_29<br>HTH_28 | Transposase<br>Homeodomain-like domain<br>Helix-turn-helix of insertion element transposase<br>Winged helix-turn helix<br>Helix-turn-helix domain | 2.60E-15<br>6.60E-07<br>8.30E-06<br>6.40E-05<br>3.90E-04 |
| WP_156324817.1 | 2080408 | 2081290 | + | 293 | PF00665<br>PF13683<br>PF13276 | rve<br>rve_3<br>HTH_21 | Integrase core domain<br>Integrase core domain<br>HTH-like domain | 9.30E-23<br>7.90E-20<br>5.50E-10 |
| WP_156324818.1 | 2081415 | 2081706 | + | 96 | NO PFAM MATCH | - | - | - |
| WP_156324819.1 | 2082579 | 2082174 | - | 134 | NO PFAM MATCH | - | - | - |

| index | leader | core | start | end | dir | score |
|---|---|---|---|---|---|---|
| 1 | MGSAPMTCTPTVTVRITVRIMR | ATKRTVRGAEPAASDRVETLVAV | 2077612 | 2077474 | - | 7 |
| 2 | LIPANYDGFLSMGSAPMT | CTPTVTVRITVRIMRATKRTVRGAEPAASDRVETLVAV | 2077645 | 2077474 | - | 19 |
| 3 | MLATAVNSAQDRGLIPANYDGFLSMGSAPMT | CTPTVTVRITVRIMRATKRTVRGAEPAASDRVETLVAV | 2077684 | 2077474 | - | 19 |

### Results for WP_070759247.1 [Corynebacterium urealyticum] back to top

previous - next

#### Architecture

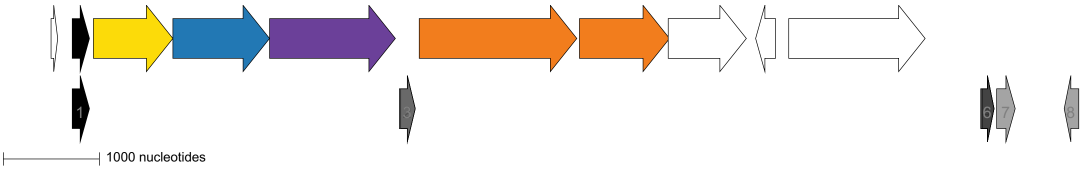

Link to nucleotide sequence

| Accession | start | end | direction | length (aa) | Pfam/HMM | name | description | E-value |
|---|---|---|---|---|---|---|---|---|
| WP_006770492.1 | 0 | 77 | + | 25 | INFERRED GENE | - | - | - |
| WP_014301132.1 | 220 | 400 | + | 59 | LinA | LinA | LinA (linaridin precursor) | 6.50E-11 |
| WP_070759244.1 | 441 | 1275 | + | 277 | LinG<br>LinH<br>PF05090 | LinG<br>LinH<br>VKG_Carbox | LinG Homolog<br>LinH (Fusion of LinE/LinG)<br>Vitamin K-dependent gamma-carboxylase | 2.00E-84<br>1.60E-78<br>1.60E-04 |

| | | | | | | | | |
|---|---|---|---|---|---|---|---|---|
| WP_070759247.1 | 1271 | 2276 | + | 334 | LinE<br>LinH<br>PF12697<br>PF12146<br>PF00561 | LinE<br>LinH<br>Abhydrolase_6<br>Hydrolase_4<br>Abhydrolase_1 | LinE Homolog<br>LinH (Fusion of LinE/LinG)<br>Alpha/beta hydrolase family<br>Serine aminopeptidase, S33<br>alpha/beta hydrolase fold | 7.90E-69<br>1.40E-50<br>3.10E-12<br>8.10E-10<br>1.70E-08 |
| WP_070759249.1 | 2275 | 3586 | + | 436 | LinL<br>TIGR02469<br>PF08242<br>TIGR02072<br>PF13649 | LinL<br>TIGR02469<br>Methyltransf_12<br>TIGR02072<br>Methyltransf_25 | LinL<br>CbiT: precorrin-6Y C5,15-methyltransferase (decarboxylating), CbiT subunit<br>Methyltransferase domain<br>BioC: malonyl-acyl carrier protein O-methyltransferase BioC<br>Methyltransferase domain | 3.10E-46<br>8.20E-07<br>2.80E-06<br>6.00E-06<br>7.40E-06 |
| WP_149122014.1 | 3839 | 5480 | + | 546 | TIGR03375<br>TIGR00958<br>TIGR02203<br>TIGR02857<br>TIGR02204 | TIGR03375<br>TIGR00958<br>TIGR02203<br>TIGR02857<br>TIGR02204 | type_I_sec_LssB: type I secretion system ATPase<br>3a01208: antigen peptide transporter 2<br>MsbA_lipidA: lipid A export permease/ATP-binding protein MsbA<br>CydD: thiol reductant ABC exporter, CydD subunit<br>MsbA_rel: ABC transporter, permease/ATP-binding protein | 4.40E-52<br>6.80E-48<br>1.10E-47<br>2.90E-45<br>6.90E-45 |
| WP_070759253.1 | 5504 | 6449 | + | 314 | TIGR03522<br>TIGR04406<br>TIGR01188<br>TIGR03740<br>TIGR03864 | TIGR03522<br>TIGR04406<br>TIGR01188<br>TIGR03740<br>TIGR03864 | GldA_ABC_ATP: gliding motility-associated ABC transporter ATP-binding subunit GldA<br>LPS_export_lptB: LPS export ABC transporter ATP-binding protein<br>drrA: daunorubicin resistance ABC transporter, ATP-binding protein<br>galliderm_ABC: lantibiotic protection ABC transporter, ATP-binding subunit<br>PQQ_ABC_ATP: ABC transporter, ATP-binding subunit, PQQ-dependent alcohol dehydrogenase system | 2.60E-47<br>3.60E-46<br>4.60E-46<br>2.40E-45<br>1.30E-36 |
| WP_148791634.1 | 6435 | 7245 | + | 269 | PF12698<br>PF12679 | ABC2_membrane_3<br>ABC2_membrane_2 | ABC-2 family transporter protein<br>ABC-2 family transporter protein | 2.30E-11<br>8.20E-08 |
| WP_141759455.1 | 7544 | 7337 | - | 68 | PF00872 | Transposase_mut | Transposase, Mutator family | 5.00E-04 |
| WP_083317374.1 | 7690 | 9106 | + | 471 | TIGR03467<br>TIGR02734<br>PF13450<br>PF01266<br>TIGR01317 | TIGR03467<br>TIGR02734<br>NAD_binding_8<br>DAO<br>TIGR01317 | HpnE: squalene-associated FAD-dependent desaturase<br>crtI_fam: phytoene desaturase<br>NAD(P)-binding Rossmann-like domain<br>FAD dependent oxidoreductase<br>GOGAT_sm_gam: glutamate synthase, NADH/NADPH, small subunit | 1.60E-10<br>2.30E-10<br>1.10E-09<br>1.40E-08<br>8.10E-08 |

| index | leader | core | start | end | dir | score |
|---|---|---|---|---|---|---|
| 1 | MSRLSTLVNNDQITLEQVHSPVM | ATPAAFGAGVVAGAKACGALVAAAGVGAAIASATKK | 220 | 400 | + | 29 |
| 2 | MTRYFTALIAFMAVLSIAGIMFDGIDSK | QGIAITVIVSLASAAVIYWLYPFFTDLEK | 3630 | 3804 | + | 6 |
| 3 | LIAFMAVLSIAGIMFDGIDSKQGIA | ITVIVSLASAAVIYWLYPFFTDLEK | 3651 | 3804 | + | 6 |
| 4 | MLLARHPYLEERLRCGDWLSSSD | FPGQVIVSKYQVMRLQPQVDTLQS | 9685 | 9829 | + | 6 |
| 5 | LLARHPYLEERLRCGDWLSSSDF | PGQVIVSKYQVMRLQPQVDTLQS | 9688 | 9829 | + | 6 |
| 6 | LEERLRCGDWLSSSDFPGQ | VIVSKYQVMRLQPQVDTLQS | 9709 | 9829 | + | 6 |
| 7 | VNELELGNVPANAVGAIATDAKTVVI | ATPRRLIFFKIFSFHFESDVGGNRECAIRVNYALKH | 9855 | 10044 | + | 6 |
| 8 | LAHFRTEIKKSIADILQTWPQAGIE | ASTDGVTVTRTAPSVQQKPPRPEID | 10715 | 10562 | - | 6 |

### Results for WP_049151517.1 [Corynebacterium striatum] back to top

previous - next

#### Architecture

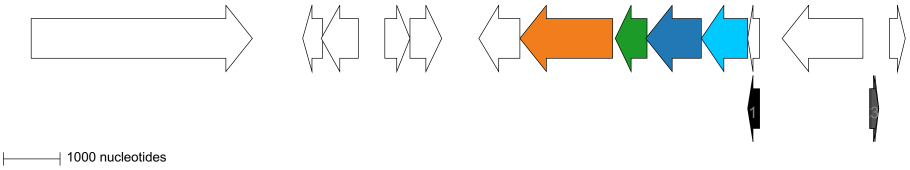

1000 nucleotides

Link to nucleotide sequence

| Accession | start | end | direction | length (aa) | Pfam/HMM | name | description | E-value |
|---|---|---|---|---|---|---|---|---|

| | | | | | | | | |
|---|---|---|---|---|---|---|---|---|
| WP_152681333.1 | 9519 | 13416 | + | 1298 | PF08751<br>PF13604<br>TIGR02768<br>TIGR02760<br>TIGR02686 | TrwC<br>AAA_30<br>TIGR02768<br>TIGR02760<br>TIGR02686 | TrwC relaxase<br>AAA domain<br>TraA_Ti: Ti-type conjugative transfer relaxase TraA<br>TraI_TIGR: conjugative transfer relaxase protein TraI<br>relax_trwC: conjugative relaxase domain | 1.40E-86<br>2.80E-56<br>2.50E-55<br>7.80E-38<br>3.20E-26 |
| WP_049151512.1 | 14654 | 14324 | - | 109 | NO PFAM MATCH | - | - | - |
| WP_049151513.1 | 15288 | 14640 | - | 215 | PF13614<br>PF09140<br>PF01656<br>TIGR01969<br>TIGR03371 | AAA_31<br>MipZ<br>CbiA<br>TIGR01969<br>TIGR03371 | AAA domain<br>ATPase MipZ<br>CobQ/CobB/MinD/ParA nucleotide binding domain<br>minD_arch: cell division ATPase MinD<br>cellulose_yhjQ: cellulose synthase operon protein YhjQ | 3.80E-16<br>1.10E-14<br>1.30E-14<br>5.70E-13<br>2.40E-12 |
| WP_065420383.1 | 15765 | 16215 | + | 149 | TIGR01764<br>PF12728 | TIGR01764<br>HTH_17 | excise: DNA binding domain, excisionase family<br>Helix-turn-helix domain | 3.80E-18<br>1.80E-11 |
| WP_080969961.1 | 16211 | 16766 | + | 184 | PF13470 | PIN_3 | PIN domain | 8.00E-14 |
| WP_049151514.1 | 18150 | 17418 | - | 243 | PF13561<br>PF00106<br>TIGR01830<br>TIGR04316<br>TIGR01829 | adh_short_C2<br>adh_short<br>TIGR01830<br>TIGR04316<br>TIGR01829 | Enoyl-(Acyl carrier protein) reductase<br>short chain dehydrogenase<br>3oxo_ACP_reduc: 3-oxoacyl-[acyl-carrier-protein] reductase<br>dhbA_paeA: 2,3-dihydro-2,3-dihydroxybenzoate dehydrogenase<br>AcAcCoA_reduct: acetoacetyl-CoA reductase | 4.40E-49<br>3.90E-41<br>2.60E-40<br>6.10E-37<br>1.10E-36 |
| WP_080969963.1 | 19795 | 18142 | - | 550 | TIGR02204<br>TIGR03375<br>TIGR02203<br>TIGR02857<br>TIGR00958 | TIGR02204<br>TIGR03375<br>TIGR02203<br>TIGR02857<br>TIGR00958 | MsbA_rel: ABC transporter, permease/ATP-binding protein<br>type_I_sec_LssB: type I secretion system ATPase<br>MsbA_lipidA: lipid A export permease/ATP-binding protein MsbA<br>CydD: thiol reductant ABC exporter, CydD subunit<br>3a01208: antigen peptide transporter 2 | 9.80E-79<br>3.20E-78<br>7.80E-77<br>3.60E-71<br>8.70E-70 |
| WP_152681334.1 | 20386 | 19837 | - | 182 | LinL | LinL | LinL | 1.80E-29 |
| WP_049151517.1 | 21350 | 20375 | - | 324 | LinE<br>LinH<br>PF12697<br>PF12146<br>TIGR01250 | LinE<br>LinH<br>Abhydrolase_6<br>Hydrolase_4<br>TIGR01250 | LinE Homolog<br>LinH (Fusion of LinE/LinG)<br>Alpha/beta hydrolase family<br>Serine aminopeptidase, S33<br>pro_imino_pep_2: proline-specific peptidase | 1.90E-38<br>1.10E-34<br>9.40E-15<br>3.30E-11<br>1.10E-09 |
| WP_049151518.1 | 22170 | 21342 | - | 275 | LinH<br>LinG<br>PF05090<br>TIGR04033 | LinH<br>LinG<br>VKG_Carbox<br>TIGR04033 | LinH (Fusion of LinE/LinG)<br>LinG Homolog<br>Vitamin K-dependent gamma-carboxylase<br>export_SdpB: antimicrobial peptide system protein, SdpB family | 1.40E-66<br>1.40E-62<br>3.10E-07<br>1.70E-04 |
| WP_049151519.1 | 22387 | 22177 | - | 69 | NO PFAM MATCH | - | - | - |
| WP_080969965.1 | 24209 | 22787 | - | 473 | TIGR02734<br>PF13450<br>TIGR03467<br>TIGR02733<br>PF01266 | TIGR02734<br>NAD_binding_8<br>TIGR03467<br>TIGR02733<br>DAO | crtI_fam: phytoene desaturase<br>NAD(P)-binding Rossmann-like domain<br>HpnE: squalene-associated FAD-dependent desaturase<br>desat_CrtD: C-3',4' desaturase CrtD<br>FAD dependent oxidoreductase | 3.60E-14<br>6.00E-12<br>5.90E-10<br>1.20E-08<br>3.40E-08 |
| WP_080969966.1 | 24671 | 24947 | + | 91 | PF01022<br>PF13518<br>PF13412<br>PF00392<br>PF13730 | HTH_5<br>HTH_28<br>HTH_24<br>GntR<br>HTH_36 | Bacterial regulatory protein, arsR family<br>Helix-turn-helix domain<br>Winged helix-turn-helix DNA-binding<br>Bacterial regulatory proteins, gntR family<br>Helix-turn-helix domain | 7.40E-08<br>3.70E-06<br>3.90E-06<br>4.20E-06<br>8.50E-06 |

| index | leader | core | start | end | dir | score |
|---|---|---|---|---|---|---|
| 1 | MLATAVNSAQDRGLIPADRDGFLAIGSAPMT | CTPTVTVRITVRIMRAVTRKQRGAEPAATDHKETLVAV | 22387 | 22177 | - | 19 |
| 2 | MRRRVGGGTALALKCSAKQRDFEDKSYTGK | YRAKTSTSATTVERVVGSMTSRTSKGAPNV | 24308 | 24491 | + | 7 |
| 3 | LALKCSAKQRDFEDKSYTGKYRAKT | STSATTVERVVGSMTSRTSKGAPNV | 24338 | 24491 | + | 6 |

### Results for WP_022905513.1 [Curtobacterium sp. B18] back to top

previous - next

#### Architecture

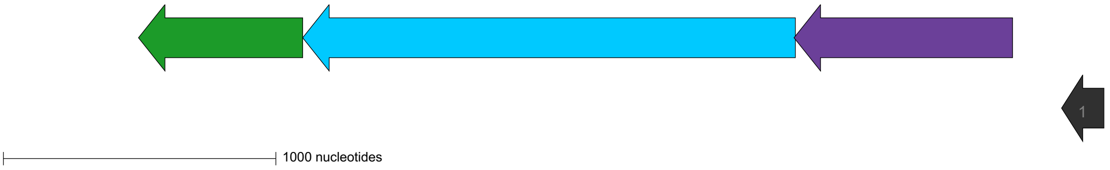

Link to nucleotide sequence

| Accession | start | end | direction | length (aa) | Pfam/HMM | name | description | E-value |
|---|---|---|---|---|---|---|---|---|
| WP_022905512.1 | 868 | 265 | - | 200 | LinL | LinL | LinL | 5.60E-55 |
| WP_022905513.1 | 2673 | 864 | - | 602 | LinH<br>LinG<br>LinE<br>PF12697<br>TIGR02427 | LinH<br>LinG<br>LinE<br>Abhydrolase_6<br>TIGR02427 | LinH (Fusion of LinE/LinG)<br>LinG Homolog<br>LinE Homolog<br>Alpha/beta hydrolase family<br>protocat_pcaD: 3-oxoadipate enol-lactonase | 6.80E-197<br>3.40E-83<br>4.70E-62<br>3.60E-12<br>2.80E-10 |
| WP_083857832.1 | 3473 | 2669 | - | 267 | PF13649<br>PF08241<br>PF13847<br>PF08242<br>PF13489 | Methyltransf_25<br>Methyltransf_11<br>Methyltransf_31<br>Methyltransf_12<br>Methyltransf_23 | Methyltransferase domain<br>Methyltransferase domain<br>Methyltransferase domain<br>Methyltransferase domain<br>Methyltransferase domain | 3.20E-18<br>1.10E-14<br>7.20E-12<br>2.70E-11<br>4.50E-11 |

| index | leader | core | start | end | dir | score |
|---|---|---|---|---|---|---|

| | | | | | | |
|---|---|---|---|---|---|---|
| 1 | VVASGEIRCGHDEDLYLRTRSEYVT | IGPTSGDSTVGWYAATYSDLAATAAD | 3805 | 3649 | - | 9 |

### Results for SDS67811.1 [Corynebacterium timonense] back to top

previous - next

#### Architecture

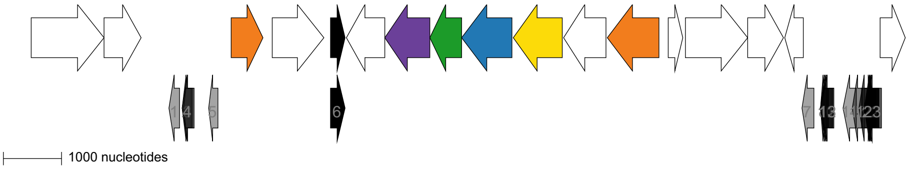

Link to nucleotide sequence

| Accession | start | end | direction | length (aa) | Pfam/HMM | name | description | E-value |
|---|---|---|---|---|---|---|---|---|
| SDS67360.1 | 2197616 | 2198855 | + | 412 | PF07730 | HisKA_3 | Histidine kinase | 3.60E-16 |
| SDS67411.1 | 2198851 | 2199490 | + | 212 | PF00072<br>PF00196<br>TIGR01818<br>TIGR02875<br>TIGR02956 | Response_reg<br>GerE<br>TIGR01818<br>TIGR02875<br>TIGR02956 | Response regulator receiver domain<br>Bacterial regulatory proteins, luxR family<br>ntrC: nitrogen regulation protein NR(I)<br>spore_0_A: sporulation transcription factor Spo0A<br>TMAO_torS: TMAO reductase sytem sensor TorS | 1.20E-26<br>7.90E-15<br>5.10E-14<br>2.10E-11<br>6.80E-11 |
| SDS67524.1 | 2201043 | 2201580 | + | 178 | TIGR01188<br>TIGR03864<br>TIGR04406<br>TIGR03522<br>TIGR01189 | TIGR01188<br>TIGR03864<br>TIGR04406<br>TIGR03522<br>TIGR01189 | drrA: daunorubicin resistance ABC transporter, ATP-binding protein<br>PQQ_ABC_ATP: ABC transporter, ATP-binding subunit, PQQ-dependent alcohol dehydrogenase system<br>LPS_export_lptB: LPS export ABC transporter ATP-binding protein<br>GldA_ABC_ATP: gliding motility-associated ABC transporter ATP-binding subunit GldA<br>ccmA: heme ABC exporter, ATP-binding protein CcmA | 3.00E-38<br>1.00E-33<br>3.90E-29<br>9.00E-28<br>7.10E-27 |
| SDS67565.1 | 2201751 | 2202645 | + | 297 | PF00144 | Beta-lactamase | Beta-lactamase | 1.00E-25 |
| SDS67612.1 | 2202753 | 2202999 | + | 81 | LinA | LinA | LinA (linaridin precursor) | 4.00E-10 |
| SDS67656.1 | 2203693 | 2203033 | - | 219 | PF00196<br>TIGR03020<br>TIGR03541 | GerE<br>TIGR03020<br>TIGR03541 | Bacterial regulatory proteins, luxR family<br>EpsA: transcriptional regulator EpsA<br>reg_near_HchA: LuxR family transcriptional regulatory, chaperone HchA-associated | 1.60E-17<br>6.90E-11<br>1.40E-10 |
| SDS67703.1 | 2204454 | 2203689 | - | 254 | PF13649<br>PF08241<br>PF13489<br>TIGR01983<br>TIGR02021 | Methyltransf_25<br>Methyltransf_11<br>Methyltransf_23<br>TIGR01983<br>TIGR02021 | Methyltransferase domain<br>Methyltransferase domain<br>Methyltransferase domain<br>UbiG: 3-demethylubiquinone-9 3-O-methyltransferase<br>BchM-ChlM: magnesium protoporphyrin O-methyltransferase | 5.40E-12<br>1.90E-10<br>4.50E-09<br>3.00E-08<br>1.30E-07 |
| SDS67752.1 | 2205002 | 2204450 | - | 183 | LinL | LinL | LinL | 7.80E-54 |
| SDS67811.1 | 2205880 | 2204998 | - | 293 | LinE<br>LinH<br>PF12697<br>TIGR02427<br>PF12146 | LinE<br>LinH<br>Abhydrolase_6<br>TIGR02427<br>Hydrolase_4 | LinE Homolog<br>LinH (Fusion of LinE/LinG)<br>Alpha/beta hydrolase family<br>protocat_pcaD: 3-oxoadipate enol-lactonase<br>Serine aminopeptidase, S33 | 1.10E-76<br>3.90E-59<br>2.40E-13<br>2.60E-10<br>1.40E-08 |
| SDS67867.1 | 2206747 | 2205886 | - | 286 | LinG<br>LinH<br>PF05090<br>PF06762<br>TIGR04033 | LinG<br>LinH<br>VKG_Carbox<br>LMF1<br>TIGR04033 | LinG Homolog<br>LinH (Fusion of LinE/LinG)<br>Vitamin K-dependent gamma-carboxylase<br>Lipase maturation factor<br>export_SdpB: antimicrobial peptide system protein, SdpB family | 3.60E-88<br>1.70E-75<br>5.90E-06<br>2.80E-05<br>3.40E-05 |
| SDS67915.1 | 2207477 | 2206769 | - | 235 | LinG | LinG | LinG Homolog | 2.00E-19 |
| SDS67980.1 | 2208388 | 2207500 | - | 295 | TIGR01188<br>TIGR03864<br>TIGR01288<br>TIGR03265<br>TIGR03740 | TIGR01188<br>TIGR03864<br>TIGR01288<br>TIGR03265<br>TIGR03740 | drrA: daunorubicin resistance ABC transporter, ATP-binding protein<br>PQQ_ABC_ATP: ABC transporter, ATP-binding subunit, PQQ-dependent alcohol dehydrogenase system<br>nodI: nodulation ABC transporter NodI<br>PhnT2: putative 2-aminoethylphosphonate ABC transporter, ATP-binding protein<br>galliderm_ABC: lantibiotic protection ABC transporter, ATP-binding subunit | 8.20E-58<br>7.00E-44<br>6.80E-43<br>5.70E-38<br>4.80E-35 |
| SDS68029.1 | 2208555 | 2208807 | + | 83 | NO PFAM MATCH | - | - | - |
| SDS68083.1 | 2208856 | 2209921 | + | 354 | PF07730 | HisKA_3 | Histidine kinase | 1.80E-11 |
| SDS68129.1 | 2209924 | 2210530 | + | 201 | PF00072<br>TIGR02875<br>PF00196<br>PF08281<br>TIGR03020 | Response_reg<br>TIGR02875<br>GerE<br>Sigma70_r4_2<br>TIGR03020 | Response regulator receiver domain<br>spore_0_A: sporulation transcription factor Spo0A<br>Bacterial regulatory proteins, luxR family<br>Sigma-70, region 4<br>EpsA: transcriptional regulator EpsA | 1.70E-24<br>6.30E-17<br>5.80E-15<br>6.60E-08<br>2.10E-07 |
| SDS68175.1 | 2210885 | 2210552 | - | 110 | NO PFAM MATCH | - | - | - |
| SDS68277.1 | 2212204 | 2212645 | + | 146 | NO PFAM MATCH | - | - | - |

| index | leader | core | start | end | dir | score |
|---|---|---|---|---|---|---|
| 1 | VGDAYDNALTESIVDVGLATMPPVQCHNNRRSHFTLRM | LPPAEYKRPTGLGALQSLSPVKAATPI | 2200163 | 2199965 | - | 6 |
| 2 | VCRLHRRLSCPDLSRGGTGARP | AAPGVVGMRPNPPLGCRRLSTRRWNSPTT | 2200351 | 2200195 | - | 7 |
| 3 | MSPPKPGNRMCVAFIVGCHAQTFRG | VELVLAPLRLALWECDRTHHSGAGG | 2200380 | 2200227 | - | 6 |
| 4 | VSAPGLPKRDVTAQARQPDVCRLHRRLSCPDLSRGGTGARP | AAPGVVGMRPNPPLGCRRLSTRRWNSPTT | 2200408 | 2200195 | - | 7 |
| 5 | VRLVRQRRVDSRLLAAASMAMVIRWEF | FRETLRNWVQQAEVDADHAPGVTTEEE | 2200814 | 2200649 | - | 6 |

| | | | | | | |
|---|---|---|---|---|---|---|
| 6 | MRNNITKFADEADLQQVGSLYGDSEPVL | ATIITTTITATTATTATTAAGYHSS...EVMAGIDEDAPVSEMLAARKDAMLV | 2202753 | 2202999 | + | 16 |
| 7 | VTPWCSANADDCGAAVPGASCGINAQSTSGS | CRLSASISWTRTASVSTNRPHDGEERAWICSS | 2211058 | 2210866 | - | 6 |
| 8 | VTECWISSTSPRRMSRTR | RGQSQVLVDTVVPVPSTSR | 2211285 | 2211171 | - | 6 |
| 9 | MGGSQSIRFNVTECWISSTSPRR | MSRTRRGQSQVLVDTVVPVPSTSR | 2211315 | 2211171 | - | 6 |
| 10 | MASAVFAHAPYSRSSASIVMGGSQSIRFNVTEC | WISSTSPRRMSRTRRGQSQVLVDTVVPVPSTSR | 2211372 | 2211171 | - | 8 |
| 11 | LIWYSMASAVFAHAPYSRSSASIVMGGSQSIRFNV | TECWISSTSPRRMSRTRRGQSQVLVDTVVPVPSTSR | 2211387 | 2211171 | - | 6 |
| 12 | MARWRLLIWYSMASAVFAHAPYSRSSASIVMGGSQSIR | FNVTECWISSTSPRRMSRTRRGQSQVLVDTVVPVPSTSR | 2211405 | 2211171 | - | 6 |
| 13 | VMARWRLLIWYSMASAVFAHAPYSRSSASIVMGGSQSIR | FNVTECWISSTSPRRMSRTRRGQSQVLVDTVVPVPSTSR | 2211408 | 2211171 | - | 6 |
| 14 | LPLPTTRKRQSGATTRRTGTHRSAT...DMRCVTVVVILPLPGVGAEVTWPRR | STPFPSPMPSHAPPP | 2211782 | 2211557 | - | 6 |
| 15 | VSPLSDCKEGLYGQLQRHHGDVFERSELCSHLCCLGVLQQGYRPDASF | PVPYQPVGHGFEVTLDELSHPRVTSVSVVYL | 2211874 | 2212114 | + | 6 |
| 16 | LQSESGDTPTQIEPTSSTKMTEKEDPYHL | SGKPLPTTQIAPTNYTETAVRCHDPQDGHS | 2211897 | 2211717 | - | 7 |
| 17 | VFERSELCSHLCCLGVLQQGYRPDASF | PVPYQPVGHGFEVTLDELSHPRVTSVSVVYL | 2211937 | 2212114 | + | 6 |
| 18 | MGPGSWHRGDIPVGALQGSRDGCIARSAQTH | RHDVSEVGHKGPPCSQKVVTPRHRLSRRHLQK | 2212028 | 2211836 | - | 8 |
| 19 | MRPLTDSHRYTTLTDVTLGCDSSSSVTSKPCPTGWYGTGKLASGRYPCW | STPRQQRWLHSSLRSNTSP | 2212134 | 2211927 | - | 9 |
| 20 | LIRKVLRWPPHTFMRPLTDSHRYTT...SVTSKPCPTGWYGTGKLASGRYPCW | STPRQQRWLHSSLRSNTSP | 2212173 | 2211927 | - | 8 |
| 21 | LELIRKVLRWPPHTFMRPLTDSHRY...SVTSKPCPTGWYGTGKLASGRYPCW | STPRQQRWLHSSLRSNTSP | 2212179 | 2211927 | - | 8 |
| 22 | MLELIRKVLRWPPHTFMRPLTDSHR...SVTSKPCPTGWYGTGKLASGRYPCW | STPRQQRWLHSSLRSNTSP | 2212182 | 2211927 | - | 8 |
| 23 | LGPAAIAVIPFEEMLELIRKVLRWP...SVTSKPCPTGWYGTGKLASGRYPCW | STPRQQRWLHSSLRSNTSP | 2212221 | 2211927 | - | 8 |

### Results for WP_163342818.1 [Curtobacterium flaccumfaciens pv. flaccumfaciens] back to top

previous - next

#### Architecture

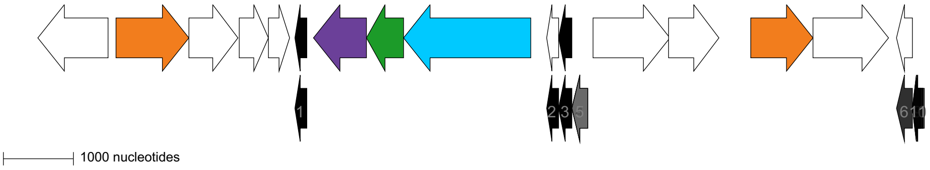

Link to nucleotide sequence

| Accession | start | end | direction | length (aa) | Pfam/HMM | name | description | E-value |
|---|---|---|---|---|---|---|---|---|
| WP_163342812.1 | 720081 | 719073 | - | 335 | PF07730 | HisKA_3 | Histidine kinase | 6.30E-10 |
| WP_163342813.1 | 720187 | 721237 | + | 349 | TIGR01188<br>TIGR03864<br>TIGR01288<br>TIGR03873<br>TIGR02142 | TIGR01188<br>TIGR03864<br>TIGR01288<br>TIGR03873<br>TIGR02142 | drrA: daunorubicin resistance ABC transporter, ATP-binding protein<br>PQQ_ABC_ATP: ABC transporter, ATP-binding subunit, PQQ-dependent alcohol dehydrogenase system<br>nodI: nodulation ABC transporter NodI<br>F420-0_ABC_ATP: proposed F420-0 ABC transporter, ATP-binding protein<br>modC_ABC: molybdate ABC transporter, ATP-binding protein | 8.00E-61<br>1.40E-52<br>1.50E-44<br>6.00E-42<br>8.80E-38 |
| WP_163342814.1 | 721233 | 721929 | + | 231 | PF01061<br>PF12698 | ABC2_membrane<br>ABC2_membrane_3 | ABC-2 type transporter<br>ABC-2 family transporter protein | 5.90E-06<br>8.80E-05 |
| WP_163342815.1 | 721957 | 722368 | + | 136 | NO PFAM MATCH | - | - | - |
| WP_152376805.1 | 722364 | 722676 | + | 103 | PF13601<br>PF12840 | HTH_34<br>HTH_20 | Winged helix DNA-binding domain<br>Helix-turn-helix domain | 1.30E-17<br>6.00E-06 |
| WP_163342816.1 | 722911 | 722743 | - | 55 | LinA | LinA | LinA (linaridin precursor) | 2.90E-07 |
| WP_152376806.1 | 723772 | 723013 | - | 252 | PF13649<br>PF13489<br>PF13847<br>PF08241<br>TIGR03534 | Methyltransf_25<br>Methyltransf_23<br>Methyltransf_31<br>Methyltransf_11<br>TIGR03534 | Methyltransferase domain<br>Methyltransferase domain<br>Methyltransferase domain<br>Methyltransferase domain<br>RF_mod_PrmC: protein-(glutamine-N5) methyltransferase, release factor-specific | 9.30E-15<br>2.60E-09<br>4.70E-09<br>5.60E-09<br>1.80E-08 |
| WP_163342817.1 | 724308 | 723768 | - | 179 | LinL | LinL | LinL | 1.10E-51 |
| WP_163342818.1 | 726119 | 724304 | - | 604 | LinH<br>LinG<br>LinE<br>PF12697<br>PF12146 | LinH<br>LinG<br>LinE<br>Abhydrolase_6<br>Hydrolase_4 | LinH (Fusion of LinE/LinG)<br>LinG Homolog<br>LinE Homolog<br>Alpha/beta hydrolase family<br>Serine aminopeptidase, S33 | 4.40E-171<br>6.30E-81<br>1.50E-80<br>4.30E-12<br>4.90E-10 |
| WP_152376810.1 | 726522 | 726339 | - | 60 | NO PFAM MATCH | - | - | - |
| WP_152376811.1 | 726707 | 726521 | - | 61 | LinA | LinA | LinA (linaridin precursor) | 6.50E-08 |

| | | | | | | | | |
|---|---|---|---|---|---|---|---|---|
| WP_152378122.1 | 726999 | 728106 | + | 368 | PF07730<br>PF02518<br>PF13581 | HisKA_3<br>HATPase_c<br>HATPase_c_2 | Histidine kinase<br>Histidine kinase-, DNA gyrase B-, and HSP90-like ATPase<br>Histidine kinase-like ATPase domain | 3.30E-19<br>5.00E-06<br>1.40E-04 |
| WP_163342819.1 | 728087 | 728813 | + | 241 | PF00072<br>PF00196<br>TIGR02875<br>TIGR03020<br>TIGR02154 | Response_reg<br>GerE<br>TIGR02875<br>TIGR03020<br>TIGR02154 | Response regulator receiver domain<br>Bacterial regulatory proteins, luxR family<br>spore_0_A: sporulation transcription factor Spo0A<br>EpsA: transcriptional regulator EpsA<br>PhoB: phosphate regulon transcriptional regulatory protein PhoB | 5.40E-28<br>3.00E-18<br>3.40E-17<br>1.10E-13<br>7.30E-12 |
| WP_027465984.1 | 729267 | 730149 | + | 293 | TIGR03522<br>TIGR01188<br>TIGR03864<br>TIGR04406<br>TIGR03740 | TIGR03522<br>TIGR01188<br>TIGR03864<br>TIGR04406<br>TIGR03740 | GldA_ABC_ATP: gliding motility-associated ABC transporter ATP-binding subunit GldA<br>drrA: daunorubicin resistance ABC transporter, ATP-binding protein<br>PQQ_ABC_ATP: ABC transporter, ATP-binding subunit, PQQ-dependent alcohol dehydrogenase system<br>LPS_export_lptB: LPS export ABC transporter ATP-binding protein<br>galliderm_ABC: lantibiotic protection ABC transporter, ATP-binding subunit | 1.50E-51<br>1.60E-50<br>6.40E-48<br>5.90E-43<br>5.10E-41 |
| WP_111030589.1 | 730145 | 731222 | + | 358 | PF12698<br>PF12679 | ABC2_membrane_3<br>ABC2_membrane_2 | ABC-2 family transporter protein<br>ABC-2 family transporter protein | 5.10E-24<br>1.60E-08 |
| WP_071245067.1 | 731569 | 731338 | - | 76 | PF13443<br>PF01381<br>PF13560<br>PF12844<br>TIGR02612 | HTH_26<br>HTH_3<br>HTH_31<br>HTH_19<br>TIGR02612 | Cro/C1-type HTH DNA-binding domain<br>Helix-turn-helix<br>Helix-turn-helix domain<br>Helix-turn-helix domain<br>mob_myst_A: mobile mystery protein A | 3.70E-17<br>5.60E-08<br>3.90E-06<br>3.00E-05<br>3.70E-04 |

| index | leader | core | start | end | dir | score |
|---|---|---|---|---|---|---|
| 1 | MTLIEIITEQASVHEARTSRDRAALVDSSADVVM | GTPTFFLAGVGVAQLIVTTLD | 722911 | 722743 | - | 31 |
| 2 | MSVQMKITEILDETEWDTDNASIEPTPRRPEL | GLPAVVATLGVANALVAAYNKGKADCGG | 726522 | 726339 | - | 19 |
| 3 | MKDDRVLHMVAQRLATMPAPDTSRAVIDVDRNHPVL | GTPAIAFWGGATLVAAFYDAGRQAG | 726707 | 726521 | - | 28 |
| 4 | VSASAAAKPTARTASLRSSVGPTGNRLHCSAPTV | DTLADRGDCFRPFVTFVTTPSRSWATDSRWGHDNQTEKG | 726932 | 726710 | - | 6 |
| 5 | VVSASAAAKPTARTASLRSSVGPTGNRLHCSAPTV | DTLADRGDCFRPFVTFVTTPSRSWATDSRWGHDNQTEKG | 726935 | 726710 | - | 6 |
| 6 | MPPEQEDGHAVVCHLDDLLAERGMT...LADRVGVTVVNLSVLKNNRARAIRF | STLTRLCEVLGCQPGDVFSVRTD | 731569 | 731338 | - | 9 |
| 7 | VDHQFLPVYLGLLVVVCF | RFALTASQRMVRDQAGLV | 731679 | 731568 | - | 6 |
| 8 | VVDHQFLPVYLGLLVVVC | FRFALTASQRMVRDQAGLV | 731682 | 731568 | - | 6 |
| 9 | VSGLGWEGYLPTPVVDHQFLPVYLG | LLVVVCFRFALTASQRMVRDQAGLV | 731721 | 731568 | - | 8 |
| 10 | VVSGLGWEGYLPTPVVDHQFLPVYL | GLLVVVCFRFALTASQRMVRDQAGLV | 731724 | 731568 | - | 8 |
| 11 | MGWNWVVSGLGWEGYLPTPVVDHQFLPV | YLGLLVVVCFRFALTASQRMVRDQAGLV | 731739 | 731568 | - | 8 |

### Results for VXB69807.1 [Curtobacterium sp. 8I-2] back to top

previous - next

#### Architecture

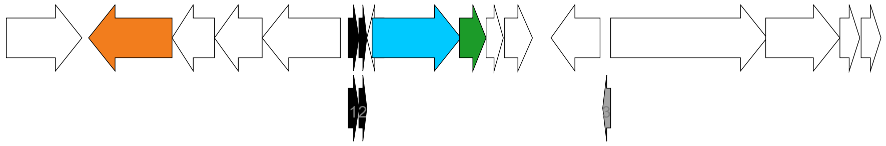

1000 nucleotides

Link to nucleotide sequence

| Accession | start | end | direction | length (aa) | Pfam/HMM | name | description | E-value |
|---|---|---|---|---|---|---|---|---|
| VXB69592.1 | 2186397 | 2187963 | + | 521 | TIGR00883<br>PF00083<br>PF07690<br>TIGR00891<br>TIGR00895 | TIGR00883<br>Sugar_tr<br>MFS_1<br>TIGR00891<br>TIGR00895 | 2A0106: MFS transporter, metabolite:H+ symporter (MHS) family protein<br>Sugar (and other) transporter<br>Major Facilitator Superfamily<br>2A0112: MFS transporter, sialate:H+ symporter (SHS) family<br>2A0115: MFS transporter, aromatic acid:H+ symporter (AAHS) family | 1.80E-123<br>1.40E-45<br>1.20E-40<br>1.20E-24<br>1.50E-24 |
| VXB69613.1 | 2189861 | 2188112 | - | 582 | TIGR02770<br>TIGR02769<br>TIGR00968<br>TIGR03265<br>TIGR02142 | TIGR02770<br>TIGR02769<br>TIGR00968<br>TIGR03265<br>TIGR02142 | nickel_nikD: nickel import ATP-binding protein NikD<br>nickel_nikE: nickel import ATP-binding protein NikE<br>3a0106s01: sulfate ABC transporter, ATP-binding protein<br>PhnT2: putative 2-aminoethylphosphonate ABC transporter, ATP-binding protein<br>modC_ABC: molybdate ABC transporter, ATP-binding protein | 1.50E-105<br>4.40E-101<br>2.60E-69<br>4.20E-68<br>2.20E-66 |
| VXB69665.1 | 2190757 | 2189857 | - | 299 | TIGR02790<br>PF00528<br>TIGR01097 | TIGR02790<br>BPD_transp_1<br>TIGR01097 | nickel_nikC: nickel ABC transporter, permease subunit NikC<br>Binding-protein-dependent transport system inner membrane component<br>PhnE: phosphonate ABC transporter, permease protein PhnE | 3.90E-55<br>5.30E-20<br>1.80E-05 |
| VXB69672.1 | 2191749 | 2190753 | - | 331 | TIGR02789<br>PF00528 | TIGR02789<br>BPD_transp_1 | nickel_nikB: nickel ABC transporter, permease subunit NikB<br>Binding-protein-dependent transport system inner membrane component | 9.90E-59<br>1.30E-23 |
| VXB69679.1 | 2193395 | 2191754 | - | 546 | PF00496<br>TIGR02294<br>TIGR04028 | SBP_bac_5<br>TIGR02294<br>TIGR04028 | Bacterial extracellular solute-binding proteins, family 5 Middle<br>nickel_nikA: nickel ABC transporter, nickel/metallophore periplasmic binding protein<br>SBP_KPN_01854: ABC transporter substrate binding protein, KPN_01854 family | 1.80E-79<br>1.40E-65<br>9.30E-65 |
| VXB69700.1 | 2193559 | 2193781 | + | 73 | LinA | LinA | LinA (linaridin precursor) | 7.60E-09 |
| VXB69718.1 | 2193777 | 2193963 | + | 61 | LinA | LinA | LinA (linaridin precursor) | 4.60E-08 |
| VXB69741.1 | 2194312 | 2193949 | - | 120 | NO PFAM MATCH | - | - | - |
| VXB69807.1 | 2194076 | 2195918 | + | 613 | LinH<br>LinG<br>LinE | LinH<br>LinG<br>LinE | LinH (Fusion of LinE/LinG)<br>LinG Homolog<br>LinE Homolog | 6.10E-132<br>1.60E-67<br>3.20E-56 |

| | | | | | | | | |
|---|---|---|---|---|---|---|---|---|
| VXB69808.1 | 2195893 | 2196469 | + | 191 | LinL | LinL | LinL | 1.50E-48 |
| VXB69855.1 | 2196465 | 2196804 | + | 112 | NO PFAM MATCH | - | - | - |
| VXB69861.1 | 2196841 | 2197441 | + | 199 | TIGR03968<br>PF17754<br>PF00440<br>TIGR03384<br>TIGR03613 | TIGR03968<br>TetR_C_14<br>TetR_N<br>TIGR03384<br>TIGR03613 | mycofact_TetR: mycofactocin system transcriptional regulator<br>MftR C-terminal domain<br>Bacterial regulatory proteins, tetR family<br>betaine_BetI: transcriptional repressor BetI<br>RutR: pyrimidine utilization regulatory protein R | 1.30E-40<br>3.90E-24<br>1.20E-16<br>5.90E-07<br>6.50E-07 |
| VXB69928.1 | 2198839 | 2197816 | - | 340 | TIGR01297<br>PF01545 | TIGR01297<br>Cation_efflux | CDF: cation diffusion facilitator family transporter<br>Cation efflux family | 1.20E-37<br>2.70E-28 |
| VXB69944.1 | 2199083 | 2202341 | + | 1085 | TIGR00392<br>PF00133<br>TIGR00422<br>TIGR00395<br>TIGR00396 | TIGR00392<br>tRNA-synt_1<br>TIGR00422<br>TIGR00395<br>TIGR00396 | ileS: isoleucine--tRNA ligase<br>tRNA synthetases class I (I, L, M and V)<br>valS: valine--tRNA ligase<br>leuS_arch: leucine--tRNA ligase<br>leuS_bact: leucine--tRNA ligase | 7.10E-240<br>1.70E-167<br>5.10E-97<br>3.70E-34<br>4.20E-29 |
| VXB69977.1 | 2202337 | 2203891 | + | 517 | TIGR01499<br>TIGR01085<br>PF08245<br>TIGR02068<br>TIGR01143 | TIGR01499<br>TIGR01085<br>Mur_ligase_M<br>TIGR02068<br>TIGR01143 | folC: bifunctional protein FolC<br>murE: UDP-N-acetylmuramyl-tripeptide synthetase<br>Mur ligase middle domain<br>cya_phycin_syn: cyanophycin synthetase<br>murF: UDP-N-acetylmuramoyl-tripeptide--D-alanyl-D-alanine ligase | 4.90E-100<br>8.60E-24<br>2.40E-15<br>7.70E-12<br>1.70E-09 |
| VXB70019.1 | 2203887 | 2204304 | + | 138 | PF14017 | DUF4233 | Protein of unknown function (DUF4233) | 8.00E-23 |
| VXB70051.1 | 2204339 | 2204759 | + | 139 | PF00334 | NDK | Nucleoside diphosphate kinase | 1.20E-49 |

| index | leader | core | start | end | dir | score |
|---|---|---|---|---|---|---|
| 1 | MSSTLDQRVNDQLRLVSSEHEPLAAGTAQPVL | TTPATVALGYAAVIGYAAEGAAFTAGATAVVGAYTVGRAAG | 2193559 | 2193781 | + | 29 |
| 2 | MTGLDTAVNRQIAAVLAESRASAVVPGEWNAQPTL | TTPAFLAGVAISSAVINAYNAGINAG | 2193777 | 2193963 | + | 28 |
| 3 | VGSLDLHEDESRFRGTTSFAAQRALCDHS | VSGVVTVPNPPGSTDGGAGAPRVLPEAPR | 2199072 | 2198895 | - | 6 |

### Results for WP_070417581.1 [Curtobacterium sp. BH-2-1-1] back to top

previous - next

#### Architecture

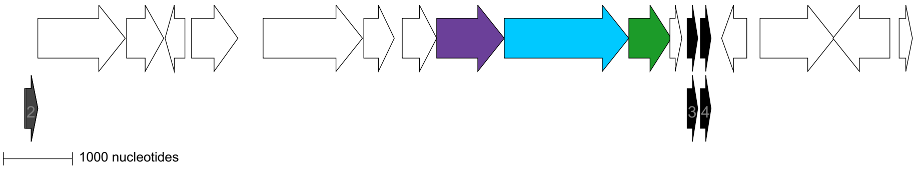

Link to nucleotide sequence

| Accession | start | end | direction | length (aa) | Pfam/HMM | name | description | E-value |
|---|---|---|---|---|---|---|---|---|
| WP_070417573.1 | 1837241 | 1838519 | + | 425 | TIGR03434<br>TIGR02212<br>PF02687<br>PF12704<br>TIGR02213 | TIGR03434<br>TIGR02212<br>FtsX<br>MacB_PCD<br>TIGR02213 | ADOP: acidobacterial duplicated orphan permease<br>lolCE: lipoprotein releasing system, transmembrane protein, LolC/E family<br>FtsX-like permease family<br>MacB-like periplasmic core domain<br>lolE_release: lipoprotein releasing system, transmembrane protein LolE | 8.50E-23<br>2.20E-20<br>6.80E-15<br>3.00E-14<br>8.70E-12 |
| WP_070417574.1 | 1838545 | 1839073 | + | 175 | PF13302<br>PF00583<br>PF13420 | Acetyltransf_3<br>Acetyltransf_1<br>Acetyltransf_4 | Acetyltransferase (GNAT) domain<br>Acetyltransferase (GNAT) family<br>Acetyltransferase (GNAT) domain | 3.40E-12<br>3.60E-10<br>4.40E-07 |
| WP_070417575.1 | 1839381 | 1839105 | - | 91 | PF13828 | DUF4190 | Domain of unknown function (DUF4190) | 4.60E-04 |
| WP_070417576.1 | 1839484 | 1840156 | + | 223 | NO PFAM MATCH | - | - | - |
| WP_070417577.1 | 1840521 | 1841967 | + | 481 | TIGR02734<br>PF13450<br>TIGR03467<br>PF01593<br>PF12831 | TIGR02734<br>NAD_binding_8<br>TIGR03467<br>Amino_oxidase<br>FAD_oxidored | crtI_fam: phytoene desaturase<br>NAD(P)-binding Rossmann-like domain<br>HpnE: squalene-associated FAD-dependent desaturase<br>Flavin containing amine oxidoreductase<br>FAD dependent oxidoreductase | 1.10E-15<br>4.60E-11<br>6.60E-11<br>4.60E-09<br>1.90E-08 |
| WP_070417578.1 | 1841969 | 1842413 | + | 147 | PF10604 | Polyketide_cyc2 | Polyketide cyclase / dehydrase and lipid transport | 4.50E-07 |
| WP_156794814.1 | 1842535 | 1843045 | + | 169 | NO PFAM MATCH | - | - | - |
| WP_083295152.1 | 1843031 | 1844024 | + | 330 | PF13649<br>PF08241<br>PF13847<br>PF08242<br>PF13489 | Methyltransf_25<br>Methyltransf_11<br>Methyltransf_31<br>Methyltransf_12<br>Methyltransf_23 | Methyltransferase domain<br>Methyltransferase domain<br>Methyltransferase domain<br>Methyltransferase domain<br>Methyltransferase domain | 5.80E-18<br>2.90E-14<br>1.80E-12<br>2.10E-11<br>4.10E-11 |
| WP_070417581.1 | 1844020 | 1845829 | + | 602 | LinH<br>LinG<br>LinE<br>PF12697<br>TIGR02427 | LinH<br>LinG<br>LinE<br>Abhydrolase_6<br>TIGR02427 | LinH (Fusion of LinE/LinG)<br>LinG Homolog<br>LinE Homolog<br>Alpha/beta hydrolase family<br>protocat_pcaD: 3-oxoadipate enol-lactonase | 4.70E-195<br>1.00E-81<br>9.40E-62<br>2.50E-12<br>3.30E-10 |
| WP_070417582.1 | 1845825 | 1846428 | + | 200 | LinL | LinL | LinL | 1.90E-54 |
| WP_022905510.1 | 1846424 | 1846592 | + | 55 | NO PFAM MATCH | - | - | - |
| WP_156794815.1 | 1846663 | 1846822 | + | 52 | LinA | LinA | LinA (linaridin precursor) | 2.70E-13 |
| WP_156794816.1 | 1846852 | 1847017 | + | 54 | LinA | LinA | LinA (linaridin precursor) | 1.20E-11 |

| | | | | | | | | |
|---|---|---|---|---|---|---|---|---|
| WP_070419333.1 | 1847525 | 1847162 | - | 120 | NO PFAM MATCH | - | - | - |
| WP_070417584.1 | 1847726 | 1848797 | + | 356 | TIGR00233<br>PF00579<br>TIGR00234 | TIGR00233<br>tRNA-synt_1b<br>TIGR00234 | trpS: tryptophan--tRNA ligase<br>tRNA synthetases class I (W and Y)<br>tyrS: tyrosine--tRNA ligase | 1.10E-71<br>8.10E-49<br>7.10E-09 |
| WP_070417585.1 | 1849599 | 1848777 | - | 273 | TIGR00507<br>TIGR01809<br>PF08501 | TIGR00507<br>TIGR01809<br>Shikimate_dh_N | aroE: shikimate dehydrogenase<br>Shik-DH-AROM: shikimate-5-dehydrogenase<br>Shikimate dehydrogenase substrate binding domain | 2.30E-42<br>5.10E-25<br>7.10E-17 |
| WP_070419334.1 | 1849734 | 1849926 | + | 63 | PF04024<br>TIGR02978 | PspC<br>TIGR02978 | PspC domain<br>phageshock_pspC: phage shock protein C | 5.80E-20<br>2.60E-13 |

| index | leader | core | start | end | dir | score |
|---|---|---|---|---|---|---|
| 1 | VVEDMLFGLQRDRGITLVVVTHDEELAERC | DRRVNVRDGLIVGGSDDAVDAPATHGRHSA | 1837062 | 1837245 | + | 7 |
| 2 | MLFGLQRDRGITLVVVTHDEELAERCDR | RVNVRDGLIVGGSDDAVDAPATHGRHSA | 1837074 | 1837245 | + | 7 |
| 3 | MTTIADIAFDSDRGQRFAPEQPLA | ATPLAALAATSVFTGAVAAYAVEEAVGN | 1846663 | 1846822 | + | 30 |
| 4 | MQQTRQLDAVPFDSDEATTFRGLRPVA | ATPAAVAWAGAVVAAAGVGAAIGDAVD | 1846852 | 1847017 | + | 29 |

### Results for WP_071295649.1 [Curtobacterium sp. MCBA15_016] back to top

previous - next

#### Architecture

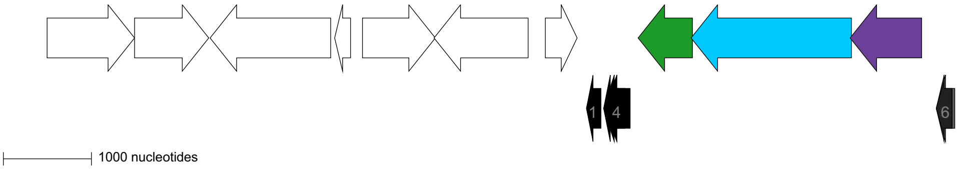

Link to nucleotide sequence

| Accession | start | end | direction | length (aa) | Pfam/HMM | name | description | E-value |
|---|---|---|---|---|---|---|---|---|
| WP_071295643.1 | 13843 | 14845 | + | 333 | TIGR03168<br>TIGR03828<br>TIGR01231<br>PF00294<br>TIGR04382 | TIGR03168<br>TIGR03828<br>TIGR01231<br>PfkB<br>TIGR04382 | 1-PFK: hexose kinase, 1-phosphofructokinase family<br>pfkB: 1-phosphofructokinase<br>lacC: tagatose-6-phosphate kinase<br>pfkB family carbohydrate kinase<br>myo_inos_iolC_N: 5-dehydro-2-deoxygluconokinase | 1.90E-61<br>7.20E-58<br>9.70E-35<br>1.20E-29<br>1.70E-09 |
| WP_071295644.1 | 14841 | 15678 | + | 278 | PF01116<br>TIGR01859<br>TIGR00167<br>TIGR01858<br>TIGR01521 | F_bP_aldolase<br>TIGR01859<br>TIGR00167<br>TIGR01858<br>TIGR01521 | Fructose-bisphosphate aldolase class-II<br>fruc_bis_ald_: fructose-1,6-bisphosphate aldolase, class II<br>cbbA: ketose-bisphosphate aldolase<br>tag_bisphos_ald: class II aldolase, tagatose bisphosphate family<br>FruBisAldo_II_B: fructose-bisphosphate aldolase, class II, Calvin cycle subtype | 5.10E-75<br>2.70E-66<br>2.40E-60<br>8.30E-57<br>7.60E-33 |
| WP_071295645.1 | 17070 | 15696 | - | 457 | TIGR00720<br>PF03313<br>PF03315<br>TIGR00718<br>TIGR00719 | TIGR00720<br>SDH_alpha<br>SDH_beta<br>TIGR00718<br>TIGR00719 | sda_mono: L-serine ammonia-lyase<br>Serine dehydratase alpha chain<br>Serine dehydratase beta chain<br>sda_alpha: L-serine dehydratase, iron-sulfur-dependent, alpha subunit<br>sda_beta: L-serine dehydratase, iron-sulfur-dependent, beta subunit | 4.60E-195<br>1.10E-97<br>2.00E-54<br>2.70E-53<br>5.10E-13 |
| WP_071295653.1 | 17296 | 17104 | - | 63 | PF04024<br>TIGR02978 | PspC<br>TIGR02978 | PspC domain<br>phageshock_pspC: phage shock protein C | 1.70E-19<br>2.30E-13 |
| WP_071295646.1 | 17431 | 18253 | + | 273 | TIGR00507<br>TIGR01809<br>PF08501 | TIGR00507<br>TIGR01809<br>Shikimate_dh_N | aroE: shikimate dehydrogenase<br>Shik-DH-AROM: shikimate-5-dehydrogenase<br>Shikimate dehydrogenase substrate binding domain | 2.00E-41<br>4.60E-24<br>7.00E-17 |
| WP_071295647.1 | 19304 | 18233 | - | 356 | TIGR00233<br>PF00579<br>TIGR00234 | TIGR00233<br>tRNA-synt_1b<br>TIGR00234 | trpS: tryptophan--tRNA ligase<br>tRNA synthetases class I (W and Y)<br>tyrS: tyrosine--tRNA ligase | 1.10E-70<br>3.40E-49<br>7.80E-09 |
| WP_070419333.1 | 19505 | 19868 | + | 120 | NO PFAM MATCH | - | - | - |
| WP_071295648.1 | 21167 | 20564 | - | 200 | LinL | LinL | LinL | 3.60E-54 |
| WP_071295649.1 | 22972 | 21163 | - | 602 | LinH<br>LinG<br>LinE<br>PF12697<br>TIGR02427 | LinH<br>LinG<br>LinE<br>Abhydrolase_6<br>TIGR02427 | LinH (Fusion of LinE/LinG)<br>LinG Homolog<br>LinE Homolog<br>Alpha/beta hydrolase family<br>protocat_pcaD: 3-oxoadipate enol-lactonase | 6.70E-198<br>4.50E-83<br>7.60E-63<br>1.50E-12<br>2.40E-10 |
| WP_083403352.1 | 23772 | 22968 | - | 267 | PF13649<br>PF08241<br>PF13847<br>PF08242<br>PF13489 | Methyltransf_25<br>Methyltransf_11<br>Methyltransf_31<br>Methyltransf_12<br>Methyltransf_23 | Methyltransferase domain<br>Methyltransferase domain<br>Methyltransferase domain<br>Methyltransferase domain<br>Methyltransferase domain | 1.40E-18<br>1.80E-14<br>2.40E-14<br>2.40E-12<br>1.60E-11 |

| index | leader | core | start | end | dir | score |
|---|---|---|---|---|---|---|
| 1 | MQQTRQLDAVPFDSDEATTFRGLRPVA | ATPAAVAWAGAVVAAAGVGAAIGDAVD | 20140 | 19975 | - | 29 |
| 2 | MTTIADIAFDSDRGQRFAPEQPLA | ATPLAALAATSVFTGAVAAYAVEEAVGN | 20329 | 20170 | - | 30 |
| 3 | VTSAHRAGNRAQRPTITIAERKESSMTTIADIAFDSDRGQRFAPEQPLA | ATPLAALAATSVFTGAVAAYAVEEAVGN | 20404 | 20170 | - | 28 |
| 4 | VAVGPGRARHRSRSLRRLGTTRVTS...SMTTIADIAFDSDRGQRFAPEQPLA | ATPLAALAATSVFTGAVAAYAVEEAVGN | 20470 | 20170 | - | 29 |
| 5 | VDGAENGKRLASGEIRCGHDEDLYLRTRS | QYVTIGPTSGDSTVGWYAATYSDLAATAAD | 24128 | 23948 | - | 8 |

| | | | | | | |
|---|---|---|---|---|---|---|
| 6 | VWAGRIVDGAENGKRLASGEIRCGHDEDLYLR | TRSQYVTIGPTSGDSTVGWYAATYSDLAATAAD | 24146 | 23948 | - | 8 |

### Results for WP_138802251.1 [Curtobacterium sp. KBS0715] back to top

previous - next

#### Architecture

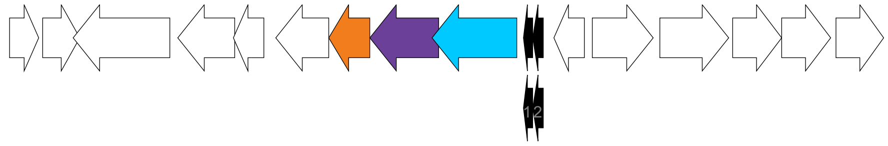

1000 nucleotides

Link to nucleotide sequence

| Accession | start | end | direction | length (aa) | Pfam/HMM | name | description | E-value |
|---|---|---|---|---|---|---|---|---|
| WP_138802244.1 | 681547 | 682189 | + | 213 | TIGR00554<br>PF00485<br>PF13604<br>PF06414<br>TIGR00750 | TIGR00554<br>PRK<br>AAA_30<br>Zeta_toxin<br>TIGR00750 | panK_bact: pantothenate kinase<br>Phosphoribulokinase / Uridine kinase family<br>AAA domain<br>Zeta toxin<br>lao: LAO/AO transport system ATPase | 6.90E-05<br>1.10E-04<br>2.60E-04<br>2.70E-04<br>3.00E-04 |
| WP_138802245.1 | 682295 | 683129 | + | 277 | PF13460<br>PF05368<br>TIGR03649<br>TIGR03466 | NAD_binding_10<br>NmrA<br>TIGR03649<br>TIGR03466 | NAD(P)H-binding<br>NmrA-like family<br>ergot_EASG: ergot alkaloid biosynthesis protein, AFUA_2G17970 family<br>HpnA: hopanoid-associated sugar epimerase | 1.20E-23<br>2.00E-23<br>4.10E-19<br>8.00E-11 |
| WP_138802246.1 | 685181 | 682970 | - | 736 | PF13231 | PMT_2 | Dolichyl-phosphate-mannose-protein mannosyltransferase | 6.50E-12 |
| WP_138802247.1 | 686626 | 685345 | - | 426 | TIGR00896<br>PF07690<br>TIGR00881 | TIGR00896<br>MFS_1<br>TIGR00881 | CynX: cyanate transporter<br>Major Facilitator Superfamily<br>2A0104: phosphoglycerate transporter family protein | 5.10E-71<br>1.50E-30<br>7.00E-04 |
| WP_138802248.1 | 687297 | 686622 | - | 224 | TIGR01449<br>PF13419<br>TIGR01454<br>TIGR02009<br>PF00702 | TIGR01449<br>HAD_2<br>TIGR01454<br>TIGR02009<br>Hydrolase | PGP_bact: phosphoglycolate phosphatase, bacterial<br>Haloacid dehalogenase-like hydrolase<br>AHBA_synth_RP: AHBA synthesis associated protein<br>PGMB-YQAB-SF: beta-phosphoglucomutase family hydrolase<br>haloacid dehalogenase-like hydrolase | 8.50E-25<br>1.90E-21<br>4.30E-14<br>2.90E-12<br>8.00E-10 |
| WP_138802249.1 | 688776 | 687579 | - | 398 | PF12698 | ABC2_membrane_3 | ABC-2 family transporter protein | 1.30E-24 |
| WP_138802250.1 | 689704 | 688765 | - | 312 | TIGR03522<br>TIGR03864<br>TIGR01188<br>TIGR04406<br>TIGR03740 | TIGR03522<br>TIGR03864<br>TIGR01188<br>TIGR04406<br>TIGR03740 | GldA_ABC_ATP: gliding motility-associated ABC transporter ATP-binding subunit GldA<br>PQQ_ABC_ATP: ABC transporter, ATP-binding subunit, PQQ-dependent alcohol dehydrogenase system<br>drrA: daunorubicin resistance ABC transporter, ATP-binding protein<br>LPS_export_lptB: LPS export ABC transporter ATP-binding protein<br>galliderm_ABC: lantibiotic protection ABC transporter, ATP-binding subunit | 2.70E-54<br>9.60E-45<br>1.80E-44<br>5.00E-39<br>2.00E-38 |
| WP_138803410.1 | 691242 | 689700 | - | 513 | PF13649<br>PF13489<br>PF08241<br>PF08242<br>TIGR02021 | Methyltransf_25<br>Methyltransf_23<br>Methyltransf_11<br>Methyltransf_12<br>TIGR02021 | Methyltransferase domain<br>Methyltransferase domain<br>Methyltransferase domain<br>Methyltransferase domain<br>BchM-ChlM: magnesium protoporphyrin O-methyltransferase | 1.10E-14<br>1.00E-12<br>3.90E-11<br>8.90E-10<br>2.50E-07 |
| WP_138802251.1 | 693013 | 691099 | - | 637 | LinH<br>LinG<br>LinE<br>PF12697<br>TIGR03611 | LinH<br>LinG<br>LinE<br>Abhydrolase_6<br>TIGR03611 | LinH (Fusion of LinE/LinG)<br>LinG Homolog<br>LinE Homolog<br>Alpha/beta hydrolase family<br>RutD: pyrimidine utilization protein D | 1.90E-210<br>1.40E-93<br>1.50E-89<br>2.50E-15<br>4.50E-10 |
| WP_138802252.1 | 693391 | 693172 | - | 72 | LinA | LinA | LinA (linaridin precursor) | 1.00E-12 |
| WP_123339258.1 | 693639 | 693396 | - | 80 | LinA | LinA | LinA (linaridin precursor) | 2.10E-11 |
| WP_027466993.1 | 694559 | 693863 | - | 231 | PF01939 | NucS | Endonuclease NucS | 2.00E-73 |
| WP_138802253.1 | 694734 | 696102 | + | 455 | TIGR00800<br>TIGR02358<br>PF02133 | TIGR00800<br>TIGR02358<br>Transp_cyt_pur | ncs1: NCS1 nucleoside transporter family<br>thia_cytX: putative hydroxymethylpyrimidine transporter CytX<br>Permease for cytosine/purines, uracil, thiamine, allantoin | 2.60E-32<br>1.10E-27<br>2.60E-23 |
| WP_138802254.1 | 696269 | 697826 | + | 518 | PF05378<br>PF01968<br>PF01869<br>TIGR03123 | Hydant_A_N<br>Hydantoinase_A<br>BcrAD_BadFG<br>TIGR03123 | Hydantoinase/oxoprolinase N-terminal region<br>Hydantoinase/oxoprolinase<br>BadF/BadG/BcrA/BcrD ATPase family<br>one_C_unchar_1: probable H4MPT-linked C1 transfer pathway protein | 6.00E-37<br>8.60E-32<br>1.90E-05<br>4.70E-04 |
| WP_138803411.1 | 697924 | 699022 | + | 365 | PF06032 | DUF917 | Protein of unknown function (DUF917) | 1.10E-122 |
| WP_138802255.1 | 699018 | 700143 | + | 374 | PF06032 | DUF917 | Protein of unknown function (DUF917) | 6.40E-124 |
| WP_138802256.1 | 700241 | 701336 | + | 364 | PF06032 | DUF917 | Protein of unknown function (DUF917) | 8.40E-74 |

| index | leader | core | start | end | dir | score |
|---|---|---|---|---|---|---|
| 1 | MTMLLEKVNADAALATLPFADVSSRGLRASVPVL | ATPATAAAFCPGFVAAFKGAAAGAAAFTAGFTIGQAIG | 693391 | 693172 | - | 28 |
| 2 | MGKTMTSLASRVNSDRSIANLQMDRTNWSGAPGELPVL | ATPATVTAAAAFVAANKGAVAGAAAFAGLFGAGVWVGVATAD | 693639 | 693396 | - | 28 |

### Results for WP_071302418.1 [Curtobacterium sp. MCBA15_013] back to top

previous - next

#### Architecture

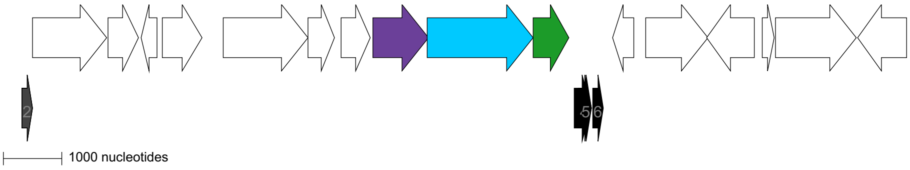

Link to nucleotide sequence

| Accession | start | end | direction | length (aa) | Pfam/HMM | name | description | E-value |
|---|---|---|---|---|---|---|---|---|
| WP_071302412.1 | 23090 | 24368 | + | 425 | TIGR03434<br>TIGR02212<br>PF02687<br>PF12704<br>TIGR02213 | TIGR03434<br>TIGR02212<br>FtsX<br>MacB_PCD<br>TIGR02213 | ADOP: acidobacterial duplicated orphan permease<br>lolCE: lipoprotein releasing system, transmembrane protein, LolC/E family<br>FtsX-like permease family<br>MacB-like periplasmic core domain<br>lolE_release: lipoprotein releasing system, transmembrane protein LolE | 1.00E-22<br>6.30E-20<br>6.80E-15<br>3.40E-14<br>1.50E-11 |
| WP_070417574.1 | 24393 | 24921 | + | 175 | PF13302<br>PF00583<br>PF13420 | Acetyltransf_3<br>Acetyltransf_1<br>Acetyltransf_4 | Acetyltransferase (GNAT) domain<br>Acetyltransferase (GNAT) family<br>Acetyltransferase (GNAT) domain | 3.40E-12<br>3.60E-10<br>4.40E-07 |
| WP_071302413.1 | 25229 | 24953 | - | 91 | PF13828 | DUF4190 | Domain of unknown function (DUF4190) | 1.70E-04 |
| WP_071294162.1 | 25332 | 26004 | + | 223 | NO PFAM MATCH | - | - | - |
| WP_071302414.1 | 26368 | 27814 | + | 481 | TIGR02734<br>TIGR03467<br>PF13450<br>PF01593<br>PF12831 | TIGR02734<br>TIGR03467<br>NAD_binding_8<br>Amino_oxidase<br>FAD_oxidored | crtI_fam: phytoene desaturase<br>HpnE: squalene-associated FAD-dependent desaturase<br>NAD(P)-binding Rossmann-like domain<br>Flavin containing amine oxidoreductase<br>FAD dependent oxidoreductase | 1.20E-15<br>6.10E-11<br>6.80E-11<br>2.70E-09<br>2.00E-08 |
| WP_071302415.1 | 27816 | 28260 | + | 147 | PF10604 | Polyketide_cyc2 | Polyketide cyclase / dehydrase and lipid transport | 1.90E-07 |
| WP_071302416.1 | 28382 | 28892 | + | 169 | NO PFAM MATCH | - | - | - |
| WP_083413857.1 | 28923 | 29871 | + | 315 | PF13649<br>PF08241<br>PF13847<br>PF08242<br>PF13489 | Methyltransf_25<br>Methyltransf_11<br>Methyltransf_31<br>Methyltransf_12<br>Methyltransf_23 | Methyltransferase domain<br>Methyltransferase domain<br>Methyltransferase domain<br>Methyltransferase domain<br>Methyltransferase domain | 9.00E-18<br>5.60E-14<br>3.60E-12<br>2.80E-11<br>8.60E-11 |
| WP_071302418.1 | 29867 | 31676 | + | 602 | LinH<br>LinG<br>LinE<br>PF12697<br>TIGR02427 | LinH<br>LinG<br>LinE<br>Abhydrolase_6<br>TIGR02427 | LinH (Fusion of LinE/LinG)<br>LinG Homolog<br>LinE Homolog<br>Alpha/beta hydrolase family<br>protocat_pcaD: 3-oxoadipate enol-lactonase | 9.40E-198<br>5.60E-83<br>1.10E-62<br>2.40E-12<br>4.20E-10 |
| WP_071302419.1 | 31672 | 32275 | + | 200 | LinL | LinL | LinL | 3.30E-54 |
| WP_070419333.1 | 33386 | 33023 | - | 120 | NO PFAM MATCH | - | - | - |
| WP_071302421.1 | 33594 | 34665 | + | 356 | TIGR00233<br>PF00579<br>TIGR00234 | TIGR00233<br>tRNA-synt_1b<br>TIGR00234 | trpS: tryptophan--tRNA ligase<br>tRNA synthetases class I (W and Y)<br>tyrS: tyrosine--tRNA ligase | 7.60E-70<br>1.90E-48<br>7.20E-09 |
| WP_071302422.1 | 35467 | 34645 | - | 273 | TIGR00507<br>TIGR01809<br>PF08501 | TIGR00507<br>TIGR01809<br>Shikimate_dh_N | aroE: shikimate dehydrogenase<br>Shik-DH-AROM: shikimate-5-dehydrogenase<br>Shikimate dehydrogenase substrate binding domain | 9.00E-41<br>2.20E-25<br>1.50E-16 |
| WP_071302423.1 | 35602 | 35794 | + | 63 | PF04024<br>TIGR02978 | PspC<br>TIGR02978 | PspC domain<br>phageshock_pspC: phage shock protein C | 9.40E-20<br>9.80E-13 |
| WP_071302424.1 | 35828 | 37202 | + | 457 | TIGR00720<br>PF03313<br>PF03315<br>TIGR00718<br>TIGR00719 | TIGR00720<br>SDH_alpha<br>SDH_beta<br>TIGR00718<br>TIGR00719 | sda_mono: L-serine ammonia-lyase<br>Serine dehydratase alpha chain<br>Serine dehydratase beta chain<br>sda_alpha: L-serine dehydratase, iron-sulfur-dependent, alpha subunit<br>sda_beta: L-serine dehydratase, iron-sulfur-dependent, beta subunit | 4.10E-196<br>8.10E-98<br>1.60E-55<br>1.10E-53<br>2.40E-13 |
| WP_070417587.1 | 38057 | 37220 | - | 278 | PF01116<br>TIGR01859<br>TIGR00167<br>TIGR01858<br>TIGR01521 | F_bP_aldolase<br>TIGR01859<br>TIGR00167<br>TIGR01858<br>TIGR01521 | Fructose-bisphosphate aldolase class-II<br>fruc_bis_ald_: fructose-1,6-bisphosphate aldolase, class II<br>cbbA: ketose-bisphosphate aldolase<br>tag_bisphos_ald: class II aldolase, tagatose bisphosphate family<br>FruBisAldo_II_B: fructose-bisphosphate aldolase, class II, Calvin cycle subtype | 2.10E-75<br>1.80E-66<br>1.50E-60<br>7.10E-57<br>6.00E-33 |

| index | leader | core | start | end | dir | score |
|---|---|---|---|---|---|---|
| 1 | VVEDMLFGLQRDRGITLVVVTHDEELAERC | DRRVNVRDGLIVGGSDDAVDAPATHGRHSA | 22911 | 23094 | + | 7 |
| 2 | MLFGLQRDRGITLVVVTHDEELAERCDR | RVNVRDGLIVGGSDDAVDAPATHGRHSA | 22923 | 23094 | + | 7 |
| 3 | VAVGPGRARHRSRSLRRLGTTRVTS...SMTTIADIAFDSDRGQRFAPEQPLA | ATPLAALAATSVFTGAVAAYAVEEAVGN | 32369 | 32669 | + | 29 |
| 4 | VTSAHRAGNRAQRPTSTIAERKESSMTTIADIAFDSDRGQRFAPEQPLA | ATPLAALAATSVFTGAVAAYAVEEAVGN | 32435 | 32669 | + | 28 |
| 5 | MTTIADIAFDSDRGQRFAPEQPLA | ATPLAALAATSVFTGAVAAYAVEEAVGN | 32510 | 32669 | + | 30 |
| 6 | MQQTRQLDAVPFDSDEATTFRGLRPVA | ATPAAVAWAGAVVAAAGVGAAIGDAVD | 32699 | 32864 | + | 29 |

### Results for WP_111010392.1 [Curtobacterium sp. MCBD17_013] back to top

previous - next

#### Architecture

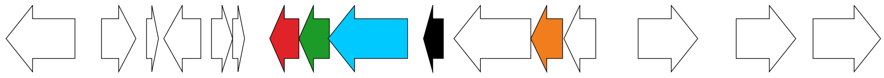

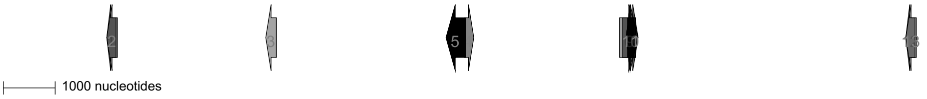

Link to nucleotide sequence

| Accession | start | end | direction | length (aa) | Pfam/HMM | name | description | E-value |
|---|---|---|---|---|---|---|---|---|
| WP_111010289.1 | 44965 | 43639 | - | 441 | NO PFAM MATCH | - | - | - |
| WP_111010290.1 | 45483 | 46137 | + | 217 | PF08378 | NERD | Nuclease-related domain | 1.90E-10 |
| WP_017888443.1 | 46342 | 46594 | + | 84 | INFERRED GENE | - | - | - |
| WP_111010391.1 | 47399 | 46682 | - | 238 | PF02909<br>PF00440<br>TIGR03384<br>PF13305 | TetR_C_1<br>TetR_N<br>TIGR03384<br>TetR_C_33 | Tetracyclin repressor-like, C-terminal domain<br>Bacterial regulatory proteins, tetR family<br>betaine_BetI: transcriptional repressor BetI<br>Tetracyclin repressor-like, C-terminal domain | 5.30E-14<br>7.80E-07<br>1.50E-05<br>3.70E-04 |
| WP_146244053.1 | 47600 | 48014 | + | 137 | NO PFAM MATCH | - | - | - |
| WP_146244054.1 | 48010 | 48238 | + | 75 | NO PFAM MATCH | - | - | - |
| WP_111010292.1 | 49307 | 48728 | - | 192 | TIGR00521<br>TIGR02113<br>PF02441 | TIGR00521<br>TIGR02113<br>Flavoprotein | coaBC_dfp: phosphopantothenoylcysteine decarboxylase / phosphopantothenate--cysteine ligase<br>coaC_strep: phosphopantothenoylcysteine decarboxylase<br>Flavoprotein | 1.40E-08<br>1.30E-07<br>1.40E-06 |
| WP_111010293.1 | 49879 | 49303 | - | 191 | LinL | LinL | LinL | 2.10E-52 |
| WP_111010392.1 | 51398 | 49865 | - | 510 | LinH<br>LinE<br>LinG<br>PF12697<br>PF00561 | LinH<br>LinE<br>LinG<br>Abhydrolase_6<br>Abhydrolase_1 | LinH (Fusion of LinE/LinG)<br>LinE Homolog<br>LinG Homolog<br>Alpha/beta hydrolase family<br>alpha/beta hydrolase fold | 5.60E-157<br>7.40E-72<br>3.50E-67<br>7.30E-10<br>1.20E-04 |
| WP_146244055.1 | 52088 | 51710 | - | 125 | LinA | LinA | LinA (linaridin precursor) | 1.60E-10 |
| WP_111010294.1 | 53771 | 52292 | - | 492 | NO PFAM MATCH | - | - | - |
| WP_111010295.1 | 54394 | 53767 | - | 208 | TIGR02857<br>TIGR01189<br>TIGR02673<br>TIGR01166<br>TIGR03608 | TIGR02857<br>TIGR01189<br>TIGR02673<br>TIGR01166<br>TIGR03608 | CydD: thiol reductant ABC exporter, CydD subunit<br>ccmA: heme ABC exporter, ATP-binding protein CcmA<br>FtsE: cell division ATP-binding protein FtsE<br>cbiO: cobalt ABC transporter, ATP-binding protein<br>L_ocin_972_ABC: putative bacteriocin export ABC transporter, lactococcin 972 group | 1.30E-29<br>1.50E-29<br>7.30E-28<br>2.90E-25<br>2.50E-24 |
| WP_111010296.1 | 55035 | 54423 | - | 203 | PF13026 | DUF3887 | Protein of unknown function (DUF3887) | 4.20E-13 |
| WP_017824271.1 | 55838 | 57001 | + | 387 | INFERRED GENE | - | - | - |
| WP_146244056.1 | 57729 | 58881 | + | 383 | NO PFAM MATCH | - | - | - |
| WP_069599718.1 | 59220 | 60532 | + | 437 | INFERRED GENE | - | - | - |

| index | leader | core | start | end | dir | score |
|---|---|---|---|---|---|---|
| 1 | MSSANGTGRSLADLTFMALMGAKA | RTLADAVIDEGPDSNYVLGRGITGDPSDRGAQRCHAAR | 45330 | 45141 | - | 6 |
| 2 | LCRSMSSANGTGRSLADLTFMALMGAKA | RTLADAVIDEGPDSNYVLGRGITGDPSDRGAQRCHAAR | 45342 | 45141 | - | 7 |
| 3 | LAPTVSTILILLAVFDLLLAPHRNGAWLPLL | VVMPCILWRSPSSDRRDRPRLLVFPLRLSFT | 48419 | 48230 | - | 6 |
| 4 | VRDDEHARVRSARLQSRSLGTEHHSKPPLDMFSDAGP | WTAVVHEPRAFGTTPP | 52070 | 52232 | + | 7 |
| 5 | MLVISHSATPSVEPQRLTSRRTARS...VQAFANTALSSAEAGARADASIPAM | TTPTIVAAAEFVVAGSTICLIC | 52088 | 51710 | - | 16 |
| 6 | MPVRQGAWTGFPEAQPVIVSDSTRA...AGLLDPGRASQRVRRRREARCGLAA | RLPGHVDAARTAVLLGAAMVPTSAVVAVFQPVGRTH | 55057 | 55411 | + | 6 |
| 7 | VSDSTRAETDLKLCEDSGQSFVTCA...AGLLDPGRASQRVRRRREARCGLAA | RLPGHVDAARTAVLLGAAMVPTSAVVAVFQPVGRTH | 55111 | 55411 | + | 7 |
| 8 | VTARRPGAGLLDPGRASQRVRRRREARCGLAA | RLPGHVDAARTAVLLGAAMVPTSAVVAVFQPVGRTH | 55204 | 55411 | + | 8 |
| 9 | VRRRRGPGGALLARTAPPVCDARAG | WLGQGLTGRRRVDVPSLTTPTEEHK | 55328 | 55175 | - | 9 |
| 10 | LRRRAGPPYVRRRRGPGGALLARTAPPVC | DARAGWLGQGLTGRRRVDVPSLTTPTEEHK | 55355 | 55175 | - | 7 |
| 11 | MWAPLRRRAGPPYVRRRRGPGGALLARTAPP | VCDARAGWLGQGLTGRRRVDVPSLTTPTEEHK | 55367 | 55175 | - | 7 |
| 12 | MEMKAAAVDAVVFVTNDNKKDWSRNGVAHP | VLAAEARAVIGKPFYLWSLEEFQRFATEQMG | 60771 | 60585 | - | 6 |
| 13 | LFGAMEMKAAAVDAVVFVTNDNKKDWSRNGVA | HPVLAAEARAVIGKPFYLWSLEEFQRFATEQMG | 60783 | 60585 | - | 6 |

### Results for PZE55389.1 [Curtobacterium sp. MCLR17_031] back to top

previous - next

#### Architecture

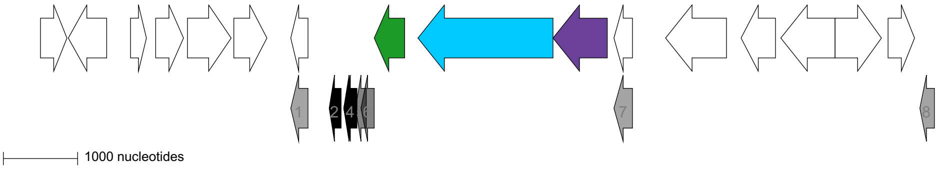

Link to nucleotide sequence

| Accession | start | end | direction | length (aa) | Pfam/HMM | name | description | E-value |
|---|---|---|---|---|---|---|---|---|

| | | | | | | | | |
|---|---|---|---|---|---|---|---|---|
| PZE55381.1 | 64192 | 64552 | + | 119 | PF02148 | zf-UBP | Zn-finger in ubiquitin-hydrolases and other protein | 2.20E-11 |
| PZE55382.1 | 65092 | 64570 | - | 173 | PF01625<br>TIGR00401 | PMSR<br>TIGR00401 | Peptide methionine sulfoxide reductase<br>msrA: peptide-methionine (S)-S-oxide reductase | 4.80E-58<br>2.90E-53 |
| PZE55383.1 | 65408 | 65624 | + | 71 | NO PFAM MATCH | - | - | - |
| PZE55384.1 | 65760 | 66132 | + | 123 | NO PFAM MATCH | - | - | - |
| PZE55385.1 | 66175 | 66763 | + | 195 | PF13646<br>PF03130 | HEAT_2<br>HEAT_PBS | HEAT repeats<br>PBS lyase HEAT-like repeat | 2.30E-16<br>1.50E-05 |
| PZE55386.1 | 66795 | 67257 | + | 153 | PF14085 | DUF4265 | Domain of unknown function (DUF4265) | 1.40E-09 |
| PZE55387.1 | 67798 | 67579 | - | 72 | PF10049 | DUF2283 | Protein of unknown function (DUF2283) | 2.00E-08 |
| PZE55388.1 | 69109 | 68686 | - | 140 | LinL | LinL | LinL | 2.00E-31 |
| PZE55389.1 | 71097 | 69282 | - | 604 | LinH<br>LinG<br>LinE<br>PF12697<br>TIGR02427 | LinH<br>LinG<br>LinE<br>Abhydrolase_6<br>TIGR02427 | LinH (Fusion of LinE/LinG)<br>LinG Homolog<br>LinE Homolog<br>Alpha/beta hydrolase family<br>protocat_pcaD: 3-oxoadipate enol-lactonase | 1.00E-191<br>4.10E-80<br>8.20E-62<br>2.20E-13<br>2.40E-10 |
| PZE55390.1 | 71828 | 71096 | - | 243 | PF13649<br>PF08241<br>PF13847<br>PF13489<br>PF08242 | Methyltransf_25<br>Methyltransf_11<br>Methyltransf_31<br>Methyltransf_23<br>Methyltransf_12 | Methyltransferase domain<br>Methyltransferase domain<br>Methyltransferase domain<br>Methyltransferase domain<br>Methyltransferase domain | 5.50E-18<br>2.30E-14<br>2.80E-12<br>2.80E-10<br>3.70E-10 |
| PZE55391.1 | 72174 | 71922 | - | 83 | NO PFAM MATCH | - | - | - |
| PZE55392.1 | 73432 | 72610 | - | 273 | PF00248<br>TIGR01293 | Aldo_ket_red<br>TIGR01293 | Aldo/keto reductase family<br>Kv_beta: voltage-dependent potassium channel beta subunit | 1.10E-43<br>1.30E-05 |
| PZE55393.1 | 74095 | 73633 | - | 153 | PF14325 | DUF4383 | Domain of unknown function (DUF4383) | 5.00E-41 |
| PZE55394.1 | 74908 | 74173 | - | 244 | TIGR03384<br>TIGR03613<br>PF00440 | TIGR03384<br>TIGR03613<br>TetR_N | betaine_BetI: transcriptional repressor BetI<br>RutR: pyrimidine utilization regulatory protein R<br>Bacterial regulatory proteins, tetR family | 1.60E-10<br>1.80E-09<br>2.30E-08 |
| PZE55395.1 | 74906 | 75521 | + | 204 | PF00583<br>PF13508<br>PF13673<br>TIGR01575<br>PF13527 | Acetyltransf_1<br>Acetyltransf_7<br>Acetyltransf_10<br>TIGR01575<br>Acetyltransf_9 | Acetyltransferase (GNAT) family<br>Acetyltransferase (GNAT) domain<br>Acetyltransferase (GNAT) domain<br>rimI: ribosomal-protein-alanine acetyltransferase<br>Acetyltransferase (GNAT) domain | 7.20E-09<br>2.60E-06<br>7.60E-05<br>2.40E-04<br>4.30E-04 |
| PZE55396.1 | 75605 | 75962 | + | 118 | PF00903<br>PF18029<br>TIGR03211<br>TIGR02295 | Glyoxalase<br>Glyoxalase_6<br>TIGR03211<br>TIGR02295 | Glyoxalase/Bleomycin resistance protein/Dioxygenase superfamily<br>Glyoxalase-like domain<br>catechol_2_3: catechol 2,3 dioxygenase<br>HpaD: 3,4-dihydroxyphenylacetate 2,3-dioxygenase | 3.50E-11<br>9.60E-06<br>2.20E-04<br>2.30E-04 |

| index | leader | core | start | end | dir | score |
|---|---|---|---|---|---|---|
| 1 | MRITYDQTVEAAYSSVDEIADGAGARQLHSVATPGELGE | VTLDFDENGLLLGIEVLGATKVLPKEVLAAAQV | 67798 | 67579 | - | 6 |
| 2 | MQQIPQLNDIPFDTDDATAFRGLRPVA | ATPAAVAWAAAVVAAGGVGAAIGDAVD | 68257 | 68092 | - | 30 |
| 3 | MTTIADIAFDSDRGQRFAPEQPLA | ATPLAALAATSVFTGAVAAYAVEETVGN | 68446 | 68287 | - | 30 |
| 4 | MQRKDTSMTTIADIAFDSDRGQRFAPEQPLA | ATPLAALAATSVFTGAVAAYAVEETVGN | 68467 | 68287 | - | 30 |
| 5 | VALGTGRPRHRRRSLRRLSPTR | VTPVRRTGGRTTVVRPSTQCNGRTPA | 68589 | 68442 | - | 7 |
| 6 | MNVNSRATLWFAAPIVLIALVVAILLFDLPGLVWL | SVPVVLAIAAAVFVVSARRE | 68687 | 68519 | - | 7 |
| 7 | MAKFFNSPHPVAEMGLGAVFVIMGVVVGPVV | AAPEDRVNQGPLNMIPTPVLVGVLV...GVVVFAQGLGTWIKRKRERTEASSQ | 72174 | 71922 | - | 6 |
| 8 | VVGFDDSAWALRCDPQL | STVRQPARLLGEQAAHQVLEQIGGTDDGPRGLLLPTEVVWRGSA | 76232 | 76046 | - | 6 |

### Results for WP_111119637.1 [Curtobacterium sp. MCBD17_035] back to top

previous - next

#### Architecture

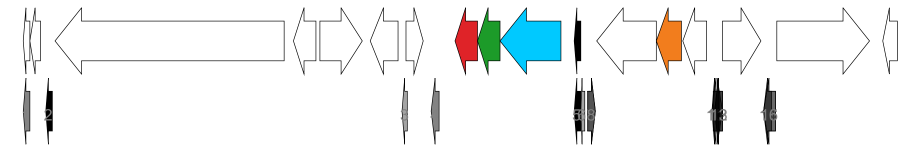

1000 nucleotides

Link to nucleotide sequence

| Accession | start | end | direction | length (aa) | Pfam/HMM | name | description | E-value |
|---|---|---|---|---|---|---|---|---|
| WP_146244374.1 | 16472 | 16280 | - | 63 | NO PFAM MATCH | - | - | - |
| WP_111119635.1 | 16714 | 16474 | - | 79 | NO PFAM MATCH | - | - | - |
| WP_111119541.1 | 22803 | 17103 | - | 1899 | TIGR01643<br>PF05593<br>TIGR03696 | TIGR01643<br>RHS_repeat<br>TIGR03696 | YD_repeat_2x: YD repeat (two copies)<br>RHS Repeat<br>Rhs_assc_core: RHS repeat-associated core domain | 3.00E-11<br>1.20E-10<br>4.20E-06 |
| WP_111119542.1 | 23604 | 23040 | - | 187 | PF16859<br>PF00440 | TetR_C_11<br>TetR_N | Tetracyclin repressor-like, C-terminal domain<br>Bacterial regulatory proteins, tetR family | 3.20E-19<br>7.30E-06 |

| | | | | | | | | |
|---|---|---|---|---|---|---|---|---|
| WP_111119543.1 | 23695 | 24769 | + | 357 | PF07690 | MFS_1 | Major Facilitator Superfamily | 5.00E-21 |
| | | | | | TIGR00895 | TIGR00895 | 2A0115: MFS transporter, aromatic acid:H+ symporter (AAHS) family | 7.40E-16 |
| | | | | | TIGR00710 | TIGR00710 | efflux_Bcr_CflA: drug resistance transporter, Bcr/CflA subfamily | 2.50E-15 |
| | | | | | TIGR00880 | TIGR00880 | 2_A_01_02: multidrug resistance protein | 2.20E-12 |
| | | | | | PF00083 | Sugar_tr | Sugar (and other) transporter | 1.10E-11 |
| WP_111119636.1 | 25669 | 24952 | - | 238 | PF02909 | TetR_C_1 | Tetracyclin repressor-like, C-terminal domain | 3.80E-14 |
| | | | | | PF00440 | TetR_N | Bacterial regulatory proteins, tetR family | 3.30E-06 |
| | | | | | TIGR03384 | TIGR03384 | betaine_BetI: transcriptional repressor BetI | 8.40E-05 |
| | | | | | PF13305 | TetR_C_33 | Tetracyclin repressor-like, C-terminal domain | 3.90E-04 |
| WP_111119544.1 | 25870 | 26284 | + | 137 | NO PFAM MATCH | - | - | - |
| WP_111010292.1 | 27654 | 27075 | - | 192 | TIGR00521 | TIGR00521 | coaBC_dfp: phosphopantothenoylcysteine decarboxylase / phosphopantothenate--cysteine ligase | 1.40E-08 |
| | | | | | TIGR02113 | TIGR02113 | coaC_strep: phosphopantothenoylcysteine decarboxylase | 1.30E-07 |
| | | | | | PF02441 | Flavoprotein | Flavoprotein | 1.40E-06 |
| WP_111119547.1 | 28226 | 27650 | - | 191 | LinL | LinL | LinL | 1.30E-52 |
| WP_111119637.1 | 29745 | 28212 | - | 510 | LinH | LinH | LinH (Fusion of LinE/LinG) | 1.00E-157 |
| | | | | | LinE | LinE | LinE Homolog | 1.50E-72 |
| | | | | | LinG | LinG | LinG Homolog | 4.10E-67 |
| | | | | | PF12697 | Abhydrolase_6 | Alpha/beta hydrolase family | 4.90E-10 |
| | | | | | PF00561 | Abhydrolase_1 | alpha/beta hydrolase fold | 1.20E-04 |
| WP_146244375.1 | 30222 | 30057 | - | 54 | LinA | LinA | LinA (linaridin precursor) | 1.30E-11 |
| WP_111119548.1 | 32118 | 30639 | - | 492 | NO PFAM MATCH | - | - | - |
| WP_111010295.1 | 32741 | 32114 | - | 208 | TIGR02857 | TIGR02857 | CydD: thiol reductant ABC exporter, CydD subunit | 1.30E-29 |
| | | | | | TIGR01189 | TIGR01189 | ccmA: heme ABC exporter, ATP-binding protein CcmA | 1.50E-29 |
| | | | | | TIGR02673 | TIGR02673 | FtsE: cell division ATP-binding protein FtsE | 7.30E-28 |
| | | | | | TIGR01166 | TIGR01166 | cbiO: cobalt ABC transporter, ATP-binding protein | 2.90E-25 |
| | | | | | TIGR03608 | TIGR03608 | L_ocin_972_ABC: putative bacteriocin export ABC transporter, lactococcin 972 group | 2.50E-24 |
| WP_111010296.1 | 33382 | 32770 | - | 203 | PF13026 | DUF3887 | Protein of unknown function (DUF3887) | 4.20E-13 |
| WP_019738739.1 | 33780 | 34718 | + | 312 | INFERRED GENE | - | - | - |
| WP_111119549.1 | 35142 | 37437 | + | 764 | PF13671 | AAA_33 | AAA domain | 1.00E-04 |
| | | | | | PF05729 | NACHT | NACHT domain | 1.40E-04 |
| | | | | | PF04471 | Mrr_cat | Restriction endonuclease | 3.30E-04 |
| | | | | | PF13604 | AAA_30 | AAA domain | 4.20E-04 |
| | | | | | PF13191 | AAA_16 | AAA ATPase domain | 6.00E-04 |
| WP_146244376.1 | 38127 | 37773 | - | 117 | NO PFAM MATCH | - | - | - |

| index | leader | core | start | end | dir | score |
|---|---|---|---|---|---|---|
| 1 | MKITDLDQPVAFGTRTFPLSTLIEDWAARVLKFAED | PTTETERPWTFDDYLGALHLRDVSTPA | 16472 | 16280 | - | 7 |
| 2 | VRRSWSICGDAGRAPGVVAHHRRLTDGIEIEVEIVT | DTVPRSGRGCSVSLT | 17028 | 16872 | - | 10 |
| 3 | MPLLSITCFSLSTAPSSE | RRCGVSRCAQPGALVRLTA | 25890 | 25776 | - | 6 |
| 4 | VAPTGRGHALHPSAFAIERPARPEGADLSSGVPVD | GPPRFAATRVPHSRRRLLVFPLRLSFT | 26689 | 26500 | - | 7 |
| 5 | MNSEVLEVQAFANTALSSAEAGARADASIPAM | TTPTIVAAAEFVVAGSTICLIC | 30222 | 30057 | - | 34 |
| 6 | MPVAHAACSDCEALPHCHL | LTRCTMPGRPFRHTRKGIR | 30335 | 30218 | - | 6 |
| 7 | LRLNARRGGVRDDEHARVRSARLQSRSLGTEHHSKPPLDMFSDAGP | WTAVVHEPRAFGTTPP | 30390 | 30579 | + | 6 |
| 8 | VRDDEHARVRSARLQSRSLGTEHHSKPPLDMFSDAGP | WTAVVHEPRAFGTTPP | 30417 | 30579 | + | 7 |
| 9 | VTARRPGAGRLDPGRASQRVRRRREARCGLAA | RLPGYVDAARTAVLLGGRNGVPTSAVVAVFRPVGRTH | 33551 | 33761 | + | 6 |
| 10 | VRRRRNPEGALPARTAPPVCDARAG | WLGQGLTGRRRVDVPSLTTPTEEHK | 33675 | 33522 | - | 8 |
| 11 | MWAHHCGRRAGPPYVRRRRNPEGALPARTAPP | VCDARAGWLGQGLTGRRRVDVPSLTTPTEEHK | 33717 | 33522 | - | 7 |
| 12 | VRSADRPEHRDDGRCGHTIAAAEQDRRTCGVDVTRKARCQPAP | RLPSATHALAGSARV | 33758 | 33581 | - | 7 |
| 13 | MARLVGPESVRSADRPEHRDDGRCG...IAAAEQDRRTCGVDVTRKARCQPAP | RLPSATHALAGSARV | 33785 | 33581 | - | 7 |
| 14 | VAARSFGISPALRLTRPYSEKGKNGAGPGETQ | TTTPSEELGIRSTARTAETCKWPASGAPNETP | 34992 | 34797 | - | 7 |
| 15 | VPSTKATEVANDFGCASPTAVSRRLATVPAVAARSFGISPALRLTRP | YSEKGKNGAGPGETQTTTPSEELGIRSTARTAETCKWPASGAPNETP | 35082 | 34797 | - | 6 |
| 16 | VTGFAPVPSTKATEVANDFGCASPTAVSRRLATVPAVAARSFGISPALRL | TRPYSEKGKNGAGPGETQTTTPSEELGIRSTARTAETCKWPASGAPNETP | 35100 | 34797 | - | 6 |

### Results for WP_111057571.1 [Curtobacterium sp. MCLR17_039] back to top

previous - next

#### Architecture

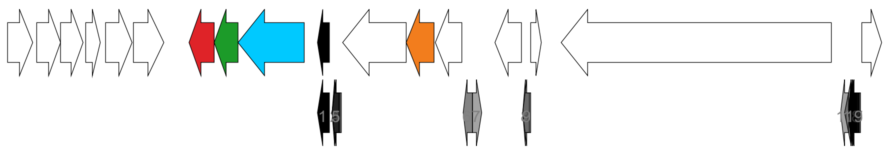

1000 nucleotides

Link to nucleotide sequence

| Accession | start | end | direction | length (aa) | Pfam/HMM | name | description | E-value |
|---|---|---|---|---|---|---|---|---|

| | | | | | | | | |
|---|---|---|---|---|---|---|---|---|
| WP_071245214.1 | 72632 | 73226 | + | 197 | PF02632 | BioY | BioY family | 1.60E-38 |
| WP_146247958.1 | 73311 | 73872 | + | 186 | NO PFAM MATCH | - | - | - |
| WP_111057538.1 | 73868 | 74399 | + | 176 | NO PFAM MATCH | - | - | - |
| WP_051597132.1 | 74448 | 74790 | + | 113 | NO PFAM MATCH | - | - | - |
| WP_111057539.1 | 74893 | 75523 | + | 209 | PF00440<br>TIGR03968<br>TIGR03613 | TetR_N<br>TIGR03968<br>TIGR03613 | Bacterial regulatory proteins, tetR family<br>mycofact_TetR: mycofactocin system transcriptional regulator<br>RutR: pyrimidine utilization regulatory protein R | 3.60E-15<br>5.60E-06<br>1.60E-04 |
| WP_111057540.1 | 75550 | 76264 | + | 237 | PF03807<br>TIGR01915<br>PF02826 | F420_oxidored<br>TIGR01915<br>2-Hacid_dh_C | NADP oxidoreductase coenzyme F420-dependent<br>npdG: NADPH-dependent F420 reductase<br>D-isomer specific 2-hydroxyacid dehydrogenase, NAD binding domain | 1.00E-17<br>2.30E-10<br>8.00E-05 |
| WP_111057541.1 | 77423 | 76850 | - | 190 | TIGR00521<br>TIGR02113<br>PF02441 | TIGR00521<br>TIGR02113<br>Flavoprotein | coaBC_dfp: phosphopantothenoylcysteine decarboxylase / phosphopantothenate--cysteine ligase<br>coaC_strep: phosphopantothenoylcysteine decarboxylase<br>Flavoprotein | 1.10E-08<br>1.50E-07<br>4.60E-07 |
| WP_111057542.1 | 77983 | 77425 | - | 185 | LinL | LinL | LinL | 5.20E-51 |
| WP_111057571.1 | 79502 | 77969 | - | 510 | LinH<br>LinE<br>LinG<br>PF12697<br>TIGR02427 | LinH<br>LinE<br>LinG<br>Abhydrolase_6<br>TIGR02427 | LinH (Fusion of LinE/LinG)<br>LinE Homolog<br>LinG Homolog<br>Alpha/beta hydrolase family<br>protocat_pcaD: 3-oxoadipate enol-lactonase | 2.80E-156<br>7.00E-73<br>8.00E-66<br>8.20E-09<br>9.30E-05 |
| WP_146247959.1 | 80093 | 79814 | - | 92 | LinA | LinA | LinA (linaridin precursor) | 6.00E-11 |
| WP_111057543.1 | 81880 | 80401 | - | 492 | NO PFAM MATCH | - | - | - |
| WP_027467258.1 | 82506 | 81876 | - | 209 | TIGR02857<br>TIGR02673<br>TIGR01189<br>TIGR03608<br>TIGR01188 | TIGR02857<br>TIGR02673<br>TIGR01189<br>TIGR03608<br>TIGR01188 | CydD: thiol reductant ABC exporter, CydD subunit<br>FtsE: cell division ATP-binding protein FtsE<br>ccmA: heme ABC exporter, ATP-binding protein CcmA<br>L_ocin_972_ABC: putative bacteriocin export ABC transporter, lactococcin 972 group<br>drrA: daunorubicin resistance ABC transporter, ATP-binding protein | 3.50E-30<br>6.50E-30<br>7.00E-30<br>1.20E-25<br>2.30E-25 |
| WP_111057544.1 | 83147 | 82535 | - | 203 | PF13026 | DUF3887 | Protein of unknown function (DUF3887) | 2.50E-14 |
| WP_146247960.1 | 84532 | 83911 | - | 206 | NO PFAM MATCH | - | - | - |
| WP_111057546.1 | 84752 | 84998 | + | 81 | NO PFAM MATCH | - | - | - |
| WP_111057548.1 | 91704 | 85449 | - | 2084 | TIGR01643<br>TIGR03696<br>PF05593 | TIGR01643<br>TIGR03696<br>RHS_repeat | YD_repeat_2x: YD repeat (two copies)<br>Rhs_assc_core: RHS repeat-associated core domain<br>RHS Repeat | 6.00E-19<br>2.90E-16<br>9.40E-12 |
| WP_146247961.1 | 92423 | 92876 | + | 150 | NO PFAM MATCH | - | - | - |

| index | leader | core | start | end | dir | score |
|---|---|---|---|---|---|---|
| 1 | MPVDARNVLRLRGASPLPPPHEVHH...VQAFANAALSSAEAGARADASIPAM | TTPTIVAAAEFVVAGSTICLIC | 80093 | 79814 | - | 33 |
| 2 | LWTMAVHGPKPGTCRSRGASSVVQY | RAVSAAAALIGAARARRLAPRHVER | 80309 | 80156 | - | 6 |
| 3 | VPKARDLWTMAVHGPKPGTCRSRGASSV | VQYRAVSAAAALIGAARARRLAPRHVER | 80327 | 80156 | - | 6 |
| 4 | VVPKARDLWTMAVHGPKPGTCRSRGASS | VVQYRAVSAAAALIGAARARRLAPRHVER | 80330 | 80156 | - | 7 |
| 5 | VTAPFRGGSGAEGARLVDHGGPRPEAWNMSIEGGF | ECCSVPSGLSCRRSHRSRACSPSRTPSRRALNCNA | 80355 | 80142 | - | 6 |
| 6 | VSAGTGPHRCGTPVRGRAGTDRKQGTFRSGRASA | GRVHRAATQRRCCLTASPEAAGAQASLPRLPSIRP | 83390 | 83180 | - | 7 |
| 7 | LNLLLQHPGKTGHEWGGAGWVIRVP...WKRGLLQRLHSEPKIAEDTIYPAIT | ETARLAKHIVKQAVTQQ | 83415 | 83625 | + | 6 |
| 8 | MAMGYSTPHVYL | ATANGSDYSAGTFPEERLASGIAFACGNPRRSSSCRNP | 84734 | 84581 | - | 6 |
| 9 | VAPGVMAMGYSTPHVYL | ATANGSDYSAGTFPEERLASGIAFACGNPRRSSSCRNP | 84749 | 84581 | - | 6 |
| 10 | LAAASSPSAAAPALMLVTDPRTSTDVRIEMS | ADLPTGIRPPKSLLKRSRAAVTLATIAEAIR | 92105 | 91916 | - | 6 |
| 11 | VPSGFATTDTVGVSEEETM | TSWSYVSATSSAALASSTA | 92243 | 92126 | - | 7 |
| 12 | VRSSVPSGFATTDTVGVSEEE | TMTSWSYVSATSSAALASSTA | 92255 | 92126 | - | 7 |
| 13 | VSSVRSSVPSGFATTDTVGVSE | EETMTSWSYVSATSSAALASSTA | 92264 | 92126 | - | 7 |
| 14 | VPSCWVRTGTVSSVRS | SVPSGFATTDTVGVSEEETMTSWSYVSATSSAALASSTA | 92294 | 92126 | - | 9 |
| 15 | VSTGTQVPSCWVRTGTVSSVRS | SVPSGFATTDTVGVSEEETMTSWSYVSATSSAALASSTA | 92312 | 92126 | - | 9 |
| 16 | VAEGVSHVPSVAREVSTGTQVPSCWVRTGTVSSVRS | SVPSGFATTDTVGVSEEETMTSWSYVSATSSAALASSTA | 92354 | 92126 | - | 8 |
| 17 | VSVAEGVSHVPSVAREVSTGTQVPSCWVRTGTVSSVRS | SVPSGFATTDTVGVSEEETMTSWSYVSATSSAALASSTA | 92360 | 92126 | - | 8 |
| 18 | VRVGGGRSQPRSVCREGGVDGHPSA...DCVLGALERAVWLCHHRHRRGQRGG | DDDVMVVRLRHLLSSARIVNGLNVG...RVLTVRCRASADAGDGSQDEHGREN | 92364 | 92019 | - | 6 |
| 19 | VSEPPRENRTCVSVAEGVSHVPSVAREVSTGTQVPSCWVRTGTVSSVRS | SVPSGFATTDTVGVSEEETMTSWSYVSATSSAALASSTA | 92393 | 92126 | - | 8 |

### Results for WP_110863536.1 [Curtobacterium sp. MCLR17_043] back to top

previous - next

#### Architecture

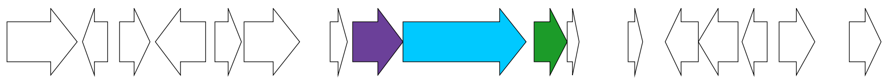

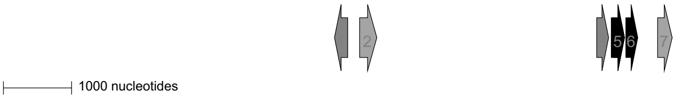

Link to nucleotide sequence

| Accession | start | end | direction | length (aa) | Pfam/HMM | name | description | E-value |
|---|---|---|---|---|---|---|---|---|
| WP_110863528.1 | 44910 | 45933 | + | 340 | TIGR01481<br>PF13377<br>TIGR02417<br>TIGR02405<br>PF00532 | TIGR01481<br>Peripla_BP_3<br>TIGR02417<br>TIGR02405<br>Peripla_BP_1 | ccpA: catabolite control protein A<br>Periplasmic binding protein-like domain<br>fruct_sucro_rep: D-fructose-responsive transcription factor<br>trehalos_R_Ecol: trehalose operon repressor<br>Periplasmic binding proteins and sugar binding domain of LacI family | 1.40E-38<br>5.50E-31<br>1.00E-22<br>6.30E-20<br>5.40E-19 |
| WP_110863529.1 | 46375 | 46018 | - | 118 | PF00903<br>PF18029<br>TIGR03211<br>TIGR02295 | Glyoxalase<br>Glyoxalase_6<br>TIGR03211<br>TIGR02295 | Glyoxalase/Bleomycin resistance protein/Dioxygenase superfamily<br>Glyoxalase-like domain<br>catechol_2_3: catechol 2,3 dioxygenase<br>HpaD: 3,4-dihydroxyphenylacetate 2,3-dioxygenase | 3.50E-11<br>9.60E-06<br>2.20E-04<br>2.30E-04 |
| WP_110863530.1 | 46548 | 47010 | + | 153 | PF14325 | DUF4383 | Domain of unknown function (DUF4383) | 3.10E-42 |
| WP_110863531.1 | 47819 | 47087 | - | 243 | PF06884 | DUF1264 | Protein of unknown function (DUF1264) | 1.10E-56 |
| WP_146237637.1 | 47952 | 48342 | + | 129 | NO PFAM MATCH | - | - | - |
| WP_110863533.1 | 48377 | 49199 | + | 273 | PF00248<br>TIGR01293 | Aldo_ket_red<br>TIGR01293 | Aldo/keto reductase family<br>Kv_beta: voltage-dependent potassium channel beta subunit | 5.10E-43<br>9.20E-06 |
| WP_110863534.1 | 49634 | 49898 | + | 87 | NO PFAM MATCH | - | - | - |
| WP_110863535.1 | 49980 | 50712 | + | 243 | PF13649<br>PF08241<br>PF13847<br>PF13489<br>PF08242 | Methyltransf_25<br>Methyltransf_11<br>Methyltransf_31<br>Methyltransf_23<br>Methyltransf_12 | Methyltransferase domain<br>Methyltransferase domain<br>Methyltransferase domain<br>Methyltransferase domain<br>Methyltransferase domain | 5.40E-18<br>1.90E-14<br>1.00E-12<br>9.30E-11<br>1.00E-10 |
| WP_110863536.1 | 50708 | 52520 | + | 603 | LinH<br>LinG<br>LinE<br>PF12697<br>TIGR02427 | LinH<br>LinG<br>LinE<br>Abhydrolase_6<br>TIGR02427 | LinH (Fusion of LinE/LinG)<br>LinG Homolog<br>LinE Homolog<br>Alpha/beta hydrolase family<br>protocat_pcaD: 3-oxoadipate enol-lactonase | 6.70E-192<br>3.30E-80<br>2.20E-62<br>2.10E-13<br>1.60E-10 |
| WP_146237638.1 | 52633 | 53116 | + | 160 | LinL | LinL | LinL | 2.60E-42 |
| WP_146237639.1 | 53115 | 53283 | + | 55 | NO PFAM MATCH | - | - | - |
| WP_110863538.1 | 54003 | 54222 | + | 72 | PF10049 | DUF2283 | Protein of unknown function (DUF2283) | 3.80E-09 |
| WP_110863539.1 | 55042 | 54544 | - | 165 | PF14085 | DUF4265 | Domain of unknown function (DUF4265) | 4.70E-09 |
| WP_110863540.1 | 55626 | 55038 | - | 195 | PF13646<br>PF03130 | HEAT_2<br>HEAT_PBS | HEAT repeats<br>PBS lyase HEAT-like repeat | 3.20E-16<br>1.40E-05 |
| WP_110863541.1 | 56047 | 55669 | - | 125 | NO PFAM MATCH | - | - | - |
| WP_110863542.1 | 56228 | 56750 | + | 173 | PF01625<br>TIGR00401 | PMSR<br>TIGR00401 | Peptide methionine sulfoxide reductase<br>msrA: peptide-methionine (S)-S-oxide reductase | 1.80E-57<br>3.30E-53 |
| WP_018191724.1 | 57255 | 57720 | + | 155 | INFERRED GENE | - | - | - |

| index | leader | core | start | end | dir | score |
|---|---|---|---|---|---|---|
| 1 | VKGCRTLHATPGDEGQGTRADILLDTGIA | VTVTDAPDDNLEHEDRNSVAEITVRPDDG | 49457 | 49280 | - | 7 |
| 2 | MAKFFNSPHPVAEMGLGAVFVIMGVVVGPVV | AAPEDRVNQGPLNMIPTPVLVGVLV...FAQGLGTWIKRKRERTEASSQETEN | 49634 | 49898 | + | 6 |
| 3 | MNVNSRATLWFAAPIVLIALVVAILLFDLPGLVWL | SVPVVLAIAAAVFVVSARRE | 53115 | 53283 | + | 7 |
| 4 | MQRKDTSMTTIADIAFDSDRGQRFAPEQPLA | ATPLAALAATSVFTGAVAAYAVEETVGN | 53335 | 53515 | + | 30 |
| 5 | MTTIADIAFDSDRGQRFAPEQPLA | ATPLAALAATSVFTGAVAAYAVEETVGN | 53356 | 53515 | + | 30 |
| 6 | MQQIPQLNDIPFDTDDATAFRGLRPVA | ATPAAVAWAAAVVAAGGVGAAIGDAVD | 53545 | 53710 | + | 30 |
| 7 | MRITYDQTVEAAYSSVDEIADVAGARQLHSVATPGELGE | VTLDFDENGLLLGIEVLGATKVLPKEVLAAAQA | 54003 | 54222 | + | 6 |

### Results for WP_083393079.1 [Curtobacterium sp. MMLR14_010] back to top

previous - next

#### Architecture

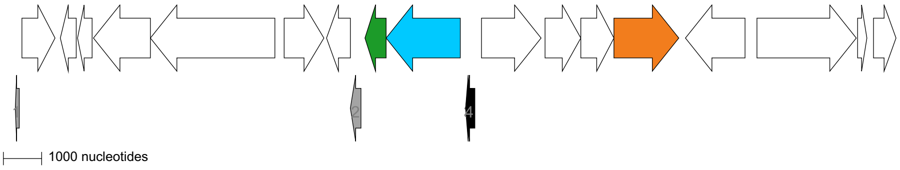

Link to nucleotide sequence

| Accession | start | end | direction | length (aa) | Pfam/HMM | name | description | E-value |
|---|---|---|---|---|---|---|---|---|
| WP_071274220.1 | 36676 | 37543 | + | 288 | PF13462<br>PF01323 | Thioredoxin_4<br>DSBA | Thioredoxin<br>DSBA-like thioredoxin domain | 3.20E-24<br>3.50E-07 |
| WP_071274223.1 | 38108 | 37688 | - | 139 | PF00334 | NDK | Nucleoside diphosphate kinase | 8.10E-49 |

| | | | | | | | | |
|---|---|---|---|---|---|---|---|---|
| WP_083393076.1 | 38514 | 38142 | - | 123 | PF14017 | DUF4233 | Protein of unknown function (DUF4233) | 6.50E-23 |
| WP_083393077.1 | 40034 | 38555 | - | 492 | TIGR01499<br>TIGR01085<br>PF08245<br>TIGR02068<br>TIGR01143 | TIGR01499<br>TIGR01085<br>Mur_ligase_M<br>TIGR02068<br>TIGR01143 | folC: bifunctional protein FolC<br>murE: UDP-N-acetylmuramyl-tripeptide synthetase<br>Mur ligase middle domain<br>cya_phycin_syn: cyanophycin synthetase<br>murF: UDP-N-acetylmuramoyl-tripeptide--D-alanyl-D-alanine ligase | 3.20E-103<br>4.00E-20<br>2.20E-16<br>1.10E-12<br>9.40E-12 |
| WP_071274229.1 | 43285 | 40030 | - | 1084 | TIGR00392<br>PF00133<br>TIGR00422<br>TIGR00395<br>TIGR00396 | TIGR00392<br>tRNA-synt_1<br>TIGR00422<br>TIGR00395<br>TIGR00396 | ileS: isoleucine--tRNA ligase<br>tRNA synthetases class I (I, L, M and V)<br>valS: valine--tRNA ligase<br>leuS_arch: leucine--tRNA ligase<br>leuS_bact: leucine--tRNA ligase | 1.10E-237<br>1.20E-164<br>1.70E-94<br>2.60E-34<br>2.20E-30 |
| WP_071274232.1 | 43532 | 44564 | + | 343 | TIGR01297<br>PF01545 | TIGR01297<br>Cation_efflux | CDF: cation diffusion facilitator family transporter<br>Cation efflux family | 4.90E-37<br>7.00E-29 |
| WP_139200616.1 | 45260 | 44639 | - | 206 | TIGR03968<br>PF17754<br>PF00440<br>TIGR03384<br>TIGR03613 | TIGR03968<br>TetR_C_14<br>TetR_N<br>TIGR03384<br>TIGR03613 | mycofact_TetR: mycofactocin system transcriptional regulator<br>MftR C-terminal domain<br>Bacterial regulatory proteins, tetR family<br>betaine_BetI: transcriptional repressor BetI<br>RutR: pyrimidine utilization regulatory protein R | 1.50E-38<br>2.60E-20<br>3.80E-16<br>1.60E-07<br>6.10E-07 |
| WP_139200617.1 | 46202 | 45629 | - | 190 | LinL | LinL | LinL | 3.20E-48 |
| WP_083393079.1 | 48130 | 46198 | - | 643 | LinH<br>LinG<br>LinE<br>PF12146<br>PF05090 | LinH<br>LinG<br>LinE<br>Hydrolase_4<br>VKG_Carbox | LinH (Fusion of LinE/LinG)<br>LinG Homolog<br>LinE Homolog<br>Serine aminopeptidase, S33<br>Vitamin K-dependent gamma-carboxylase | 1.30E-132<br>1.00E-67<br>6.10E-53<br>1.60E-05<br>2.20E-04 |
| WP_071274303.1 | 48686 | 50252 | + | 521 | PF00496<br>TIGR02294<br>TIGR04028 | SBP_bac_5<br>TIGR02294<br>TIGR04028 | Bacterial extracellular solute-binding proteins, family 5 Middle<br>nickel_nikA: nickel ABC transporter, nickel/metallophore periplasmic binding protein<br>SBP_KPN_01854: ABC transporter substrate binding protein, KPN_01854 family | 5.20E-79<br>5.50E-66<br>3.40E-65 |
| WP_071274306.1 | 50340 | 51264 | + | 307 | TIGR02789<br>PF00528 | TIGR02789<br>BPD_transp_1 | nickel_nikB: nickel ABC transporter, permease subunit NikB<br>Binding-protein-dependent transport system inner membrane component | 1.40E-56<br>7.70E-24 |
| WP_071274244.1 | 51260 | 52130 | + | 289 | TIGR02790<br>PF00528<br>TIGR01097<br>TIGR01726 | TIGR02790<br>BPD_transp_1<br>TIGR01097<br>TIGR01726 | nickel_nikC: nickel ABC transporter, permease subunit NikC<br>Binding-protein-dependent transport system inner membrane component<br>PhnE: phosphonate ABC transporter, permease protein PhnE<br>HEQRo_perm_3TM: amino ABC transporter, permease protein, 3-TM region, His/Glu/Gln/Arg/opine family | 4.70E-59<br>5.80E-19<br>4.80E-08<br>6.20E-04 |
| WP_071274247.1 | 52126 | 53833 | + | 568 | TIGR02770<br>TIGR02769<br>TIGR00968<br>TIGR03265<br>TIGR02673 | TIGR02770<br>TIGR02769<br>TIGR00968<br>TIGR03265<br>TIGR02673 | nickel_nikD: nickel import ATP-binding protein NikD<br>nickel_nikE: nickel import ATP-binding protein NikE<br>3a0106s01: sulfate ABC transporter, ATP-binding protein<br>PhnT2: putative 2-aminoethylphosphonate ABC transporter, ATP-binding protein<br>FtsE: cell division ATP-binding protein FtsE | 7.10E-106<br>8.50E-98<br>1.10E-70<br>8.40E-70<br>4.90E-68 |
| WP_071274309.1 | 55558 | 54016 | - | 513 | TIGR00883<br>PF00083<br>PF07690<br>TIGR00891<br>TIGR00895 | TIGR00883<br>Sugar_tr<br>MFS_1<br>TIGR00891<br>TIGR00895 | 2A0106: MFS transporter, metabolite:H+ symporter (MHS) family protein<br>Sugar (and other) transporter<br>Major Facilitator Superfamily<br>2A0112: MFS transporter, sialate:H+ symporter (SHS) family<br>2A0115: MFS transporter, aromatic acid:H+ symporter (AAHS) family | 7.00E-123<br>3.20E-45<br>8.10E-37<br>2.30E-25<br>8.00E-25 |
| WP_071274250.1 | 55859 | 58457 | + | 865 | TIGR00422<br>PF00133<br>TIGR00392<br>TIGR00396<br>TIGR00395 | TIGR00422<br>tRNA-synt_1<br>TIGR00392<br>TIGR00396<br>TIGR00395 | valS: valine--tRNA ligase<br>tRNA synthetases class I (I, L, M and V)<br>ileS: isoleucine--tRNA ligase<br>leuS_bact: leucine--tRNA ligase<br>leuS_arch: leucine--tRNA ligase | 1.10E-165<br>1.30E-95<br>2.70E-91<br>1.40E-42<br>6.30E-42 |
| WP_071274253.1 | 58505 | 58757 | + | 83 | NO PFAM MATCH | - | - | - |
| WP_071274256.1 | 58918 | 59506 | + | 195 | PF00440<br>TIGR03384<br>TIGR03968<br>TIGR03613 | TetR_N<br>TIGR03384<br>TIGR03968<br>TIGR03613 | Bacterial regulatory proteins, tetR family<br>betaine_BetI: transcriptional repressor BetI<br>mycofact_TetR: mycofactocin system transcriptional regulator<br>RutR: pyrimidine utilization regulatory protein R | 5.90E-11<br>8.80E-07<br>1.20E-06<br>3.70E-05 |

| index | leader | core | start | end | dir | score |
|---|---|---|---|---|---|---|
| 1 | VTCHRVRPGLRDTERGRTR | HRNDRTVGNLTRIVTTMKR | 36608 | 36491 | - | 6 |
| 2 | VTSCATEERTGLRGLAA | GAPVGVAAVVGAFSFWCLRVDGSWA...RLVREKANEPPARDVGRGHRDDADA | 45526 | 45256 | - | 6 |
| 3 | VGAAQPVLTTPLAVAVGYAAVVGY | AAEGAAFTAGAAAVVGAYTVGRAAG | 48430 | 48280 | - | 15 |
| 4 | MYSELDRRVNIQLERSPQKEFQVAVGAAQPVL | TTPLAVAVGYAAVVGYAAEGAAFTAGAAAVVGAYTVGRAAG | 48502 | 48280 | - | 28 |

### Results for PZF23552.1 [Curtobacterium sp. MCPF17_021] back to top

previous - next

#### Architecture

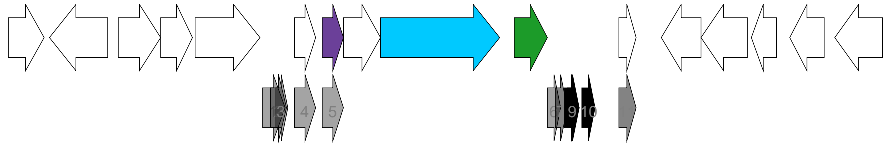

1000 nucleotides

Link to nucleotide sequence

| Accession | start | end | direction | length (aa) | Pfam/HMM | name | description | E-value |
|---|---|---|---|---|---|---|---|---|
| PZF23544.1 | 159293 | 159755 | + | 153 | PF14325 | DUF4383 | Domain of unknown function (DUF4383) | 4.90E-41 |
| PZF23545.1 | 160548 | 159816 | - | 243 | PF06884 | DUF1264 | Protein of unknown function (DUF1264) | 1.70E-56 |
| PZF23546.1 | 160688 | 161207 | + | 172 | NO PFAM MATCH | - | - | - |
| PZF23547.1 | 161209 | 161608 | + | 132 | NO PFAM MATCH | - | - | - |
| PZF23548.1 | 161643 | 162465 | + | 273 | PF00248<br>TIGR01293 | Aldo_ket_red<br>TIGR01293 | Aldo/keto reductase family<br>Kv_beta: voltage-dependent potassium channel beta subunit | 2.10E-43<br>1.50E-05 |
| PZF23549.1 | 162901 | 163165 | + | 87 | NO PFAM MATCH | - | - | - |
| PZF23550.1 | 163247 | 163508 | + | 86 | PF13649<br>PF08241<br>TIGR02021<br>TIGR00537<br>TIGR03534 | Methyltransf_25<br>Methyltransf_11<br>TIGR02021<br>TIGR00537<br>TIGR03534 | Methyltransferase domain<br>Methyltransferase domain<br>BchM-ChlM: magnesium protoporphyrin O-methyltransferase<br>hemK_rel_arch: putative methylase<br>RF_mod_PrmC: protein-(glutamine-N5) methyltransferase, release factor-specific | 2.00E-10<br>2.40E-08<br>7.10E-07<br>1.70E-06<br>1.70E-06 |
| PZF23551.1 | 163504 | 163978 | + | 157 | NO PFAM MATCH | - | - | - |
| PZF23552.1 | 163977 | 165480 | + | 500 | LinH<br>LinE<br>LinG<br>PF12697<br>TIGR02427 | LinH<br>LinE<br>LinG<br>Abhydrolase_6<br>TIGR02427 | LinH (Fusion of LinE/LinG)<br>LinE Homolog<br>LinG Homolog<br>Alpha/beta hydrolase family<br>protocat_pcaD: 3-oxoadipate enol-lactonase | 1.70E-148<br>1.90E-60<br>3.00E-53<br>1.40E-13<br>1.10E-10 |
| PZF23553.1 | 165653 | 166076 | + | 140 | LinL | LinL | LinL | 2.00E-31 |
| PZF23554.1 | 166963 | 167182 | + | 72 | PF10049 | DUF2283 | Protein of unknown function (DUF2283) | 2.00E-08 |
| PZF23555.1 | 168002 | 167504 | - | 165 | PF14085 | DUF4265 | Domain of unknown function (DUF4265) | 4.70E-09 |
| PZF23556.1 | 168586 | 167998 | - | 195 | PF13646<br>PF03130 | HEAT_2<br>HEAT_PBS | HEAT repeats<br>PBS lyase HEAT-like repeat | 5.90E-18<br>8.10E-06 |
| PZF23557.1 | 168943 | 168640 | - | 100 | NO PFAM MATCH | - | - | - |
| PZF23558.1 | 169556 | 169124 | - | 143 | PF14085 | DUF4265 | Domain of unknown function (DUF4265) | 2.70E-13 |
| PZF23559.1 | 170274 | 169686 | - | 195 | PF13646<br>PF03130 | HEAT_2<br>HEAT_PBS | HEAT repeats<br>PBS lyase HEAT-like repeat | 1.70E-17<br>4.40E-05 |

| index | leader | core | start | end | dir | score |
|---|---|---|---|---|---|---|
| 1 | VPAGTAPPVGTRPSHGRGSATVRTHRDFATKFRPSCSRLSSGRVRDR | HGHAGVEKDVGAGSLAFVARGGVERLVPLHRAFLRDRYTGIPLVRRQ | 162493 | 162778 | + | 6 |
| 2 | VLKVVVGARPRPSRPCRCREGCWRG...GLRRPGWRGAFGTPSPGIFAGQVYG | NTPGQTAVGNGRCRR | 162597 | 162801 | + | 6 |
| 3 | LARVPWPSSPGVAWSVWYPFTGHFC | GTGIREYPWSDGSRQRKVSKVTAVV | 162662 | 162815 | + | 6 |
| 4 | MAKFFNSPHPVAEMGLGAVFVIMGVVVGPVV | AAPEDRVNQGPLNMIPTPVLVGVLV...FAQGLGTWIKRKRERTEARSQEPEN | 162901 | 163165 | + | 6 |
| 5 | MQETYGQQFAAIYDDVFPPDPDLVGLLTQSQPTTALELGVGTG | RIAVPLAHAGVAVTGVDLSPEMLERCRRGAQDAVVDVELIQGS | 163247 | 163508 | + | 6 |
| 6 | MNVNSRATLWFAAPIVLIALVVAILLFDLPGLVWL | SVPVVLAIAAAVFVVSVRRE | 166075 | 166243 | + | 6 |
| 7 | VALGTGRPRHRRRSLRRLSPTR | VTPVRRTGGRTTVVRPSTQCNGRTPA | 166173 | 166320 | + | 7 |
| 8 | MQRKDTSMTTIADIAFDSDRGQRFAPEQPLA | ATPLAALAATSVFTGAVAAYAVEETVGN | 166295 | 166475 | + | 30 |
| 9 | MTTIADIAFDSDRGQRFAPEQPLA | ATPLAALAATSVFTGAVAAYAVEETVGN | 166316 | 166475 | + | 30 |
| 10 | MQQIPQLNDIPFDTDDATAFRGLRPVA | ATPAAVAWAAAVVAAGGVGAAIGDAVD | 166505 | 166670 | + | 30 |
| 11 | MRITYDQTVEAAYSSVDEIADGAGARQLHSVATPGELGE | VTLDFDENGLLLGIEVLGATKVLPKEVLAAAQA | 166963 | 167182 | + | 7 |

### Results for WP_123339259.1 [Curtobacterium sp. PhB131] back to top

previous - next

#### Architecture

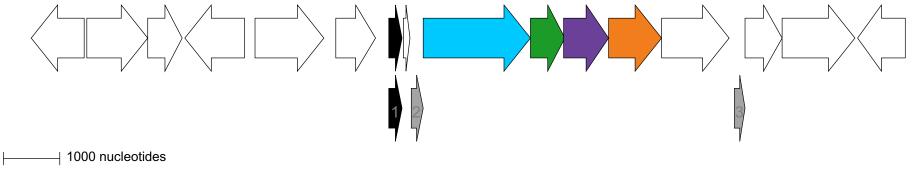

1000 nucleotides

Link to nucleotide sequence

| Accession | start | end | direction | length (aa) | Pfam/HMM | name | description | E-value |
|---|---|---|---|---|---|---|---|---|
| WP_123339255.1 | 224062 | 223123 | - | 312 | PF04909 | Amidohydro_2 | Amidohydrolase | 1.50E-38 |
| WP_123339256.1 | 224110 | 225202 | + | 363 | PF01979<br>PF07969<br>TIGR02318<br>TIGR03178<br>TIGR03583 | Amidohydro_1<br>Amidohydro_3<br>TIGR02318<br>TIGR03178<br>TIGR03583 | Amidohydrolase family<br>Amidohydrolase family<br>phosphono_phnM: phosphonate metabolism protein PhnM<br>allantoinase: allantoinase<br>EF_0837: putative amidohydrolase, EF_0837/AHA_3915 family | 3.50E-19<br>1.20E-15<br>1.40E-09<br>8.00E-08<br>1.10E-07 |
| WP_123339257.1 | 225198 | 225795 | + | 198 | PF16859<br>PF00440<br>TIGR03384 | TetR_C_11<br>TetR_N<br>TIGR03384 | Tetracyclin repressor-like, C-terminal domain<br>Bacterial regulatory proteins, tetR family<br>betaine_BetI: transcriptional repressor BetI | 4.50E-18<br>1.00E-10<br>5.80E-04 |

| | | | | | | | | |
|---|---|---|---|---|---|---|---|---|
| WP_123339532.1 | 226910 | 225854 | - | 351 | PF13377<br>TIGR01481<br>PF13407<br>TIGR02417<br>PF00532 | Peripla_BP_3<br>TIGR01481<br>Peripla_BP_4<br>TIGR02417<br>Peripla_BP_1 | Periplasmic binding protein-like domain<br>ccpA: catabolite control protein A<br>Periplasmic binding protein domain<br>fruct_sucro_rep: D-fructose-responsive transcription factor<br>Periplasmic binding proteins and sugar binding domain of LacI family | 1.10E-22<br>2.00E-20<br>2.00E-11<br>3.20E-10<br>9.60E-10 |
| WP_017887747.1 | 227109 | 228324 | + | 405 | INFERRED GENE | - | - | - |
| WP_111083752.1 | 228531 | 229227 | + | 231 | PF01939 | NucS | Endonuclease NucS | 4.90E-75 |
| WP_148053811.1 | 229463 | 229694 | + | 76 | LinA | LinA | LinA (linaridin precursor) | 1.40E-11 |
| WP_012775716.1 | 229730 | 229844 | + | 38 | INFERRED GENE | - | - | - |
| WP_123339259.1 | 230077 | 231991 | + | 637 | LinH<br>LinG<br>LinE<br>PF12697<br>TIGR03611 | LinH<br>LinG<br>LinE<br>Abhydrolase_6<br>TIGR03611 | LinH (Fusion of LinE/LinG)<br>LinG Homolog<br>LinE Homolog<br>Alpha/beta hydrolase family<br>RutD: pyrimidine utilization protein D | 2.20E-211<br>1.20E-92<br>1.50E-89<br>9.70E-15<br>3.20E-10 |
| WP_123339260.1 | 231990 | 232587 | + | 198 | LinL | LinL | LinL | 2.00E-67 |
| WP_123339261.1 | 232580 | 233378 | + | 265 | PF13649<br>PF13489<br>PF08241<br>PF08242<br>TIGR02021 | Methyltransf_25<br>Methyltransf_23<br>Methyltransf_11<br>Methyltransf_12<br>TIGR02021 | Methyltransferase domain<br>Methyltransferase domain<br>Methyltransferase domain<br>Methyltransferase domain<br>BchM-ChlM: magnesium protoporphyrin O-methyltransferase | 3.90E-15<br>2.10E-13<br>1.20E-11<br>2.90E-10<br>7.50E-08 |
| WP_123339262.1 | 233374 | 234313 | + | 312 | TIGR03522<br>TIGR03864<br>TIGR01188<br>TIGR04406<br>TIGR01288 | TIGR03522<br>TIGR03864<br>TIGR01188<br>TIGR04406<br>TIGR01288 | GldA_ABC_ATP: gliding motility-associated ABC transporter ATP-binding subunit GldA<br>PQQ_ABC_ATP: ABC transporter, ATP-binding subunit, PQQ-dependent alcohol dehydrogenase system<br>drrA: daunorubicin resistance ABC transporter, ATP-binding protein<br>LPS_export_lptB: LPS export ABC transporter ATP-binding protein<br>nodI: nodulation ABC transporter NodI | 3.80E-54<br>6.70E-45<br>2.00E-44<br>6.00E-39<br>8.60E-38 |
| WP_123339263.1 | 234302 | 235499 | + | 398 | PF12698 | ABC2_membrane_3 | ABC-2 family transporter protein | 6.60E-23 |
| WP_111057287.1 | 235781 | 236456 | + | 224 | TIGR01449<br>PF13419<br>TIGR01454<br>TIGR02009<br>PF00702 | TIGR01449<br>HAD_2<br>TIGR01454<br>TIGR02009<br>Hydrolase | PGP_bact: phosphoglycolate phosphatase, bacterial<br>Haloacid dehalogenase-like hydrolase<br>AHBA_synth_RP: AHBA synthesis associated protein<br>PGMB-YQAB-SF: beta-phosphoglucomutase family hydrolase<br>haloacid dehalogenase-like hydrolase | 3.50E-25<br>3.20E-22<br>1.00E-14<br>7.50E-13<br>1.10E-09 |
| WP_123339264.1 | 236452 | 237742 | + | 429 | TIGR00896<br>PF07690<br>TIGR00895<br>TIGR00881<br>PF00083 | TIGR00896<br>MFS_1<br>TIGR00895<br>TIGR00881<br>Sugar_tr | CynX: cyanate transporter<br>Major Facilitator Superfamily<br>2A0115: MFS transporter, aromatic acid:H+ symporter (AAHS) family<br>2A0104: phosphoglycerate transporter family protein<br>Sugar (and other) transporter | 1.70E-72<br>1.00E-30<br>3.60E-12<br>4.10E-07<br>2.10E-04 |
| WP_123339265.1 | 238626 | 237792 | - | 277 | PF13460<br>PF05368<br>TIGR03649<br>TIGR03466<br>TIGR01777 | NAD_binding_10<br>NmrA<br>TIGR03649<br>TIGR03466<br>TIGR01777 | NAD(P)H-binding<br>NmrA-like family<br>ergot_EASG: ergot alkaloid biosynthesis protein, AFUA_2G17970 family<br>HpnA: hopanoid-associated sugar epimerase<br>yfcH: TIGR01777 family protein | 9.90E-26<br>1.50E-24<br>2.30E-19<br>9.00E-13<br>3.40E-04 |

| index | leader | core | start | end | dir | score |
|---|---|---|---|---|---|---|
| 1 | MTSLASRVNSDRSIANLQMDRTNWSGAPGELPVL | ATPATVTAAAAFVAANKGAVAGAAAFAGLFGAGVWVGVATAD | 229463 | 229694 | + | 29 |
| 2 | VPPPSRPGSPSVRPSADHLCRPTVP...STAQRSGHVRPRPDGSHKRLPRPGG | HTPMHQRPASGDRA | 229868 | 230081 | + | 6 |
| 3 | LVLAETPLIGERRPSTSPSGHYEDRPVP | GLEARRRSTPSPPRPRVVARYGPARVDRT | 235611 | 235785 | + | 6 |

### Results for ROQ16621.1 [Curtobacterium sp. PhB171] back to top

previous - next

#### Architecture

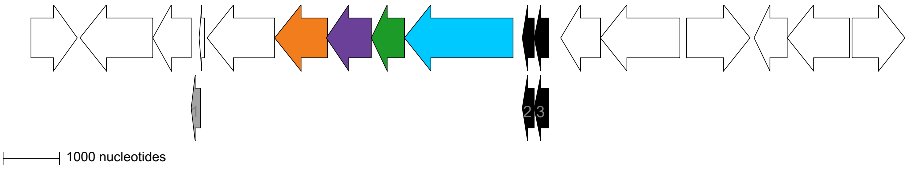

1000 nucleotides

Link to nucleotide sequence

| Accession | start | end | direction | length (aa) | Pfam/HMM | name | description | E-value |
|---|---|---|---|---|---|---|---|---|
| ROQ16613.1 | 381303 | 382137 | + | 277 | PF13460<br>PF05368<br>TIGR03649<br>TIGR03466<br>TIGR01777 | NAD_binding_10<br>NmrA<br>TIGR03649<br>TIGR03466<br>TIGR01777 | NAD(P)H-binding<br>NmrA-like family<br>ergot_EASG: ergot alkaloid biosynthesis protein, AFUA_2G17970 family<br>HpnA: hopanoid-associated sugar epimerase<br>yfcH: TIGR01777 family protein | 9.90E-26<br>1.50E-24<br>2.30E-19<br>9.00E-13<br>3.40E-04 |
| ROQ16614.1 | 383477 | 382187 | - | 429 | TIGR00896<br>PF07690<br>TIGR00895<br>TIGR00881<br>PF00083 | TIGR00896<br>MFS_1<br>TIGR00895<br>TIGR00881<br>Sugar_tr | CynX: cyanate transporter<br>Major Facilitator Superfamily<br>2A0115: MFS transporter, aromatic acid:H+ symporter (AAHS) family<br>2A0104: phosphoglycerate transporter family protein<br>Sugar (and other) transporter | 1.70E-72<br>1.00E-30<br>3.60E-12<br>4.10E-07<br>2.10E-04 |

| | | | | | | | | |
|---|---|---|---|---|---|---|---|---|
| ROQ16615.1 | 384148 | 383473 | - | 224 | TIGR01449<br>PF13419<br>TIGR01454<br>TIGR02009<br>PF00702 | TIGR01449<br>HAD_2<br>TIGR01454<br>TIGR02009<br>Hydrolase | PGP_bact: phosphoglycolate phosphatase, bacterial<br>Haloacid dehalogenase-like hydrolase<br>AHBA_synth_RP: AHBA synthesis associated protein<br>PGMB-YQAB-SF: beta-phosphoglucomutase family hydrolase<br>haloacid dehalogenase-like hydrolase | 3.50E-25<br>3.20E-22<br>1.00E-14<br>7.50E-13<br>1.10E-09 |
| ROQ16616.1 | 384389 | 384293 | - | 31 | NO PFAM MATCH | - | - | - |
| ROQ16617.1 | 385627 | 384430 | - | 398 | PF12698 | ABC2_membrane_3 | ABC-2 family transporter protein | 6.60E-23 |
| ROQ16618.1 | 386555 | 385616 | - | 312 | TIGR03522<br>TIGR03864<br>TIGR01188<br>TIGR04406<br>TIGR01288 | TIGR03522<br>TIGR03864<br>TIGR01188<br>TIGR04406<br>TIGR01288 | GldA_ABC_ATP: gliding motility-associated ABC transporter ATP-binding subunit GldA<br>PQQ_ABC_ATP: ABC transporter, ATP-binding subunit, PQQ-dependent alcohol dehydrogenase system<br>drrA: daunorubicin resistance ABC transporter, ATP-binding protein<br>LPS_export_lptB: LPS export ABC transporter ATP-binding protein<br>nodI: nodulation ABC transporter NodI | 3.80E-54<br>6.70E-45<br>2.00E-44<br>6.00E-39<br>8.60E-38 |
| ROQ16619.1 | 387346 | 386551 | - | 264 | PF13649<br>PF13489<br>PF08241<br>PF08242<br>TIGR02021 | Methyltransf_25<br>Methyltransf_23<br>Methyltransf_11<br>Methyltransf_12<br>TIGR02021 | Methyltransferase domain<br>Methyltransferase domain<br>Methyltransferase domain<br>Methyltransferase domain<br>BchM-ChlM: magnesium protoporphyrin O-methyltransferase | 3.90E-15<br>2.10E-13<br>1.20E-11<br>2.90E-10<br>7.50E-08 |
| ROQ16620.1 | 387939 | 387342 | - | 198 | LinL | LinL | LinL | 2.00E-67 |
| ROQ16621.1 | 389852 | 387938 | - | 637 | LinH<br>LinG<br>LinE<br>PF12697<br>TIGR03611 | LinH<br>LinG<br>LinE<br>Abhydrolase_6<br>TIGR03611 | LinH (Fusion of LinE/LinG)<br>LinG Homolog<br>LinE Homolog<br>Alpha/beta hydrolase family<br>RutD: pyrimidine utilization protein D | 2.20E-211<br>1.20E-92<br>1.50E-89<br>9.70E-15<br>3.20E-10 |
| ROQ16622.1 | 390224 | 390011 | - | 70 | LinA | LinA | LinA (linaridin precursor) | 5.30E-12 |
| ROQ16623.1 | 390478 | 390235 | - | 80 | LinA | LinA | LinA (linaridin precursor) | 2.10E-11 |
| ROQ16624.1 | 391398 | 390702 | - | 231 | PF01939 | NucS | Endonuclease NucS | 4.90E-75 |
| ROQ16625.1 | 392820 | 391407 | - | 470 | PF07690<br>PF00083<br>TIGR00880<br>TIGR00879<br>TIGR00895 | MFS_1<br>Sugar_tr<br>TIGR00880<br>TIGR00879<br>TIGR00895 | Major Facilitator Superfamily<br>Sugar (and other) transporter<br>2_A_01_02: multidrug resistance protein<br>SP: MFS transporter, sugar porter (SP) family<br>2A0115: MFS transporter, aromatic acid:H+ symporter (AAHS) family | 8.60E-27<br>7.50E-06<br>8.50E-05<br>2.00E-04<br>4.40E-04 |
| ROQ16626.1 | 392938 | 394066 | + | 375 | PF13377<br>TIGR01481<br>PF13407<br>TIGR02417<br>PF00356 | Peripla_BP_3<br>TIGR01481<br>Peripla_BP_4<br>TIGR02417<br>LacI | Periplasmic binding protein-like domain<br>ccpA: catabolite control protein A<br>Periplasmic binding protein domain<br>fruct_sucro_rep: D-fructose-responsive transcription factor<br>Bacterial regulatory proteins, lacI family | 1.30E-22<br>1.10E-20<br>2.40E-11<br>1.30E-10<br>5.60E-10 |
| ROQ16627.1 | 394722 | 394125 | - | 198 | PF16859<br>PF00440<br>TIGR03384 | TetR_C_11<br>TetR_N<br>TIGR03384 | Tetracyclin repressor-like, C-terminal domain<br>Bacterial regulatory proteins, tetR family<br>betaine_BetI: transcriptional repressor BetI | 4.50E-18<br>1.00E-10<br>5.80E-04 |
| ROQ16628.1 | 395810 | 394718 | - | 363 | PF01979<br>PF07969<br>TIGR02318<br>TIGR03178<br>TIGR03583 | Amidohydro_1<br>Amidohydro_3<br>TIGR02318<br>TIGR03178<br>TIGR03583 | Amidohydrolase family<br>Amidohydrolase family<br>phosphono_phnM: phosphonate metabolism protein PhnM<br>allantoinase: allantoinase<br>EF_0837: putative amidohydrolase, EF_0837/AHA_3915 family | 3.50E-19<br>1.20E-15<br>1.40E-09<br>8.00E-08<br>1.10E-07 |
| ROQ16629.1 | 395858 | 396797 | + | 312 | PF04909 | Amidohydro_2 | Amidohydrolase | 1.50E-38 |

| index | leader | core | start | end | dir | score |
|---|---|---|---|---|---|---|
| 1 | LVLAETPLIGERRPSTSPSGHYEDRPVP | GLEARRRSTPSPPRPRVVARYGPARVDRT | 384318 | 384144 | - | 6 |
| 2 | MLLEKVNADAALATLPFADVSSRGLSASVPVL | ATPATAAAFCPGFVAAFKGAAAGAAAFTAGFTIGQAIG | 390224 | 390011 | - | 29 |
| 3 | MGKTMTSLASRVNSDRSIANLQMDRTNWSGAPGELPVL | ATPATVTAAAAFVAANKGAVAGAAAFAGLFGAGVWVGVATAD | 390478 | 390235 | - | 28 |

### Results for WP_123671264.1 [Curtobacterium sp. PhB172] back to top

previous - next

#### Architecture

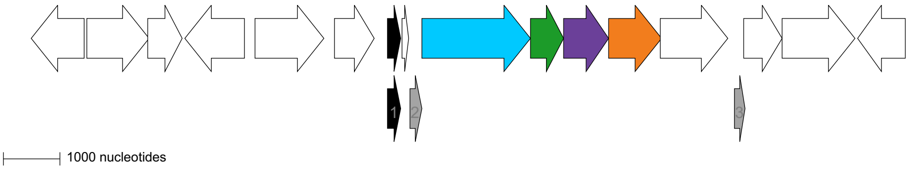

Link to nucleotide sequence

| Accession | start | end | direction | length (aa) | Pfam/HMM | name | description | E-value |
|---|---|---|---|---|---|---|---|---|
| WP_123671261.1 | 91627 | 90688 | - | 312 | PF04909 | Amidohydro_2 | Amidohydrolase | 1.20E-38 |
| WP_123671262.1 | 91675 | 92767 | + | 363 | PF01979<br>PF07969<br>TIGR02318<br>TIGR03178<br>TIGR03583 | Amidohydro_1<br>Amidohydro_3<br>TIGR02318<br>TIGR03178<br>TIGR03583 | Amidohydrolase family<br>Amidohydrolase family<br>phosphono_phnM: phosphonate metabolism protein PhnM<br>allantoinase: allantoinase<br>EF_0837: putative amidohydrolase, EF_0837/AHA_3915 family | 4.20E-19<br>4.50E-15<br>1.50E-09<br>8.20E-08<br>1.10E-07 |
| WP_123671263.1 | 92763 | 93360 | + | 198 | PF16859<br>PF00440<br>TIGR03384 | TetR_C_11<br>TetR_N<br>TIGR03384 | Tetracyclin repressor-like, C-terminal domain<br>Bacterial regulatory proteins, tetR family<br>betaine_BetI: transcriptional repressor BetI | 1.90E-17<br>9.90E-11<br>5.60E-04 |

| | | | | | | | | |
|---|---|---|---|---|---|---|---|---|
| WP_123671559.1 | 94475 | 93419 | - | 351 | PF13377<br>TIGR01481<br>PF13407<br>TIGR02417<br>PF00532 | Peripla_BP_3<br>TIGR01481<br>Peripla_BP_4<br>TIGR02417<br>Peripla_BP_1 | Periplasmic binding protein-like domain<br>ccpA: catabolite control protein A<br>Periplasmic binding protein domain<br>fruct_sucro_rep: D-fructose-responsive transcription factor<br>Periplasmic binding proteins and sugar binding domain of LacI family | 1.10E-22<br>2.50E-20<br>6.70E-11<br>4.40E-10<br>1.20E-09 |
| WP_017887747.1 | 94653 | 95877 | + | 408 | INFERRED GENE | - | - | - |
| WP_017887754.1 | 96057 | 96753 | + | 231 | PF01939 | NucS | Endonuclease NucS | 2.40E-73 |
| WP_148053811.1 | 96989 | 97220 | + | 76 | LinA | LinA | LinA (linaridin precursor) | 1.40E-11 |
| WP_012775716.1 | 97256 | 97370 | + | 38 | INFERRED GENE | - | - | - |
| WP_123671264.1 | 97603 | 99517 | + | 637 | LinH<br>LinG<br>LinE<br>PF12697<br>TIGR03611 | LinH<br>LinG<br>LinE<br>Abhydrolase_6<br>TIGR03611 | LinH (Fusion of LinE/LinG)<br>LinG Homolog<br>LinE Homolog<br>Alpha/beta hydrolase family<br>RutD: pyrimidine utilization protein D | 7.70E-<br>211<br>5.30E-93<br>2.00E-89<br>9.80E-15<br>1.90E-10 |
| WP_123339260.1 | 99516 | 100113 | + | 198 | LinL | LinL | LinL | 2.00E-67 |
| WP_123339261.1 | 100106 | 100904 | + | 265 | PF13649<br>PF13489<br>PF08241<br>PF08242<br>TIGR02021 | Methyltransf_25<br>Methyltransf_23<br>Methyltransf_11<br>Methyltransf_12<br>TIGR02021 | Methyltransferase domain<br>Methyltransferase domain<br>Methyltransferase domain<br>Methyltransferase domain<br>BchM-ChlM: magnesium protoporphyrin O-methyltransferase | 3.90E-15<br>2.10E-13<br>1.20E-11<br>2.90E-10<br>7.50E-08 |
| WP_123671265.1 | 100900 | 101839 | + | 312 | TIGR03522<br>TIGR01188<br>TIGR03864<br>TIGR04406<br>TIGR03740 | TIGR03522<br>TIGR01188<br>TIGR03864<br>TIGR04406<br>TIGR03740 | GldA_ABC_ATP: gliding motility-associated ABC transporter ATP-binding subunit GldA<br>drrA: daunorubicin resistance ABC transporter, ATP-binding protein<br>PQQ_ABC_ATP: ABC transporter, ATP-binding subunit, PQQ-dependent alcohol dehydrogenase system<br>LPS_export_lptB: LPS export ABC transporter ATP-binding protein<br>galliderm_ABC: lantibiotic protection ABC transporter, ATP-binding subunit | 4.00E-54<br>3.90E-44<br>4.20E-44<br>4.50E-39<br>1.10E-38 |
| WP_123671266.1 | 101828 | 103025 | + | 398 | PF12698 | ABC2_membrane_3 | ABC-2 family transporter protein | 6.90E-23 |
| WP_111057287.1 | 103307 | 103982 | + | 224 | TIGR01449<br>PF13419<br>TIGR01454<br>TIGR02009<br>PF00702 | TIGR01449<br>HAD_2<br>TIGR01454<br>TIGR02009<br>Hydrolase | PGP_bact: phosphoglycolate phosphatase, bacterial<br>Haloacid dehalogenase-like hydrolase<br>AHBA_synth_RP: AHBA synthesis associated protein<br>PGMB-YQAB-SF: beta-phosphoglucomutase family hydrolase<br>haloacid dehalogenase-like hydrolase | 3.50E-25<br>3.20E-22<br>1.00E-14<br>7.50E-13<br>1.10E-09 |
| WP_123339264.1 | 103978 | 105268 | + | 429 | TIGR00896<br>PF07690<br>TIGR00895<br>TIGR00881<br>PF00083 | TIGR00896<br>MFS_1<br>TIGR00895<br>TIGR00881<br>Sugar_tr | CynX: cyanate transporter<br>Major Facilitator Superfamily<br>2A0115: MFS transporter, aromatic acid:H+ symporter (AAHS) family<br>2A0104: phosphoglycerate transporter family protein<br>Sugar (and other) transporter | 1.70E-72<br>1.00E-30<br>3.60E-12<br>4.10E-07<br>2.10E-04 |
| WP_123671267.1 | 106152 | 105318 | - | 277 | PF13460<br>PF05368<br>TIGR03649<br>TIGR03466<br>TIGR01777 | NAD_binding_10<br>NmrA<br>TIGR03649<br>TIGR03466<br>TIGR01777 | NAD(P)H-binding<br>NmrA-like family<br>ergot_EASG: ergot alkaloid biosynthesis protein, AFUA_2G17970 family<br>HpnA: hopanoid-associated sugar epimerase<br>yfcH: TIGR01777 family protein | 2.60E-25<br>1.20E-23<br>1.30E-19<br>2.20E-12<br>4.20E-04 |

| index | leader | core | start | end | dir | score |
|---|---|---|---|---|---|---|
| 1 | MTSLASRVNSDRSIANLQMDRTNWSGAPGELPVL | ATPATVTAAAAFVAANKGAVAGAAAFAGLFGAGVWVGVATAD | 96989 | 97220 | + | 29 |
| 2 | VPPPSRPGSPSVRPSADHLCRPTVP...STAQRSGHVRPRPDGSHKRLPRPGG | HTPMHQRPASGDRA | 97394 | 97607 | + | 6 |
| 3 | LVLAETPLIGERRPSTSPSGHHEDRPVP | GLEARRRSTPSPPRPRVVARYGPARVDRT | 103137 | 103311 | + | 6 |

### Results for WP_141693334.1 [Curtobacterium sp. UCD-KPL2560] back to top

previous - next

#### Architecture

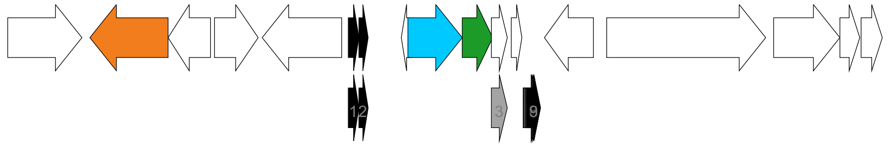

1000 nucleotides

Link to nucleotide sequence

| Accession | start | end | direction | length (aa) | Pfam/HMM | name | description | E-value |
|---|---|---|---|---|---|---|---|---|
| WP_065960197.1 | 48180 | 49716 | + | 511 | TIGR00883<br>PF00083<br>PF07690<br>TIGR00891<br>TIGR00895 | TIGR00883<br>Sugar_tr<br>MFS_1<br>TIGR00891<br>TIGR00895 | 2A0106: MFS transporter, metabolite:H+ symporter (MHS) family protein<br>Sugar (and other) transporter<br>Major Facilitator Superfamily<br>2A0112: MFS transporter, sialate:H+ symporter (SHS) family<br>2A0115: MFS transporter, aromatic acid:H+ symporter (AAHS) family | 9.30E-124<br>6.20E-45<br>2.40E-36<br>4.10E-27<br>6.60E-26 |
| WP_065960199.1 | 51492 | 49887 | - | 534 | TIGR02770<br>TIGR02769<br>TIGR00968<br>TIGR02142<br>TIGR03265 | TIGR02770<br>TIGR02769<br>TIGR00968<br>TIGR02142<br>TIGR03265 | nickel_nikD: nickel import ATP-binding protein NikD<br>nickel_nikE: nickel import ATP-binding protein NikE<br>3a0106s01: sulfate ABC transporter, ATP-binding protein<br>modC_ABC: molybdate ABC transporter, ATP-binding protein<br>PhnT2: putative 2-aminoethylphosphonate ABC transporter, ATP-binding protein | 1.70E-105<br>7.70E-98<br>5.60E-71<br>3.10E-67<br>4.20E-67 |
| WP_083230031.1 | 52370 | 51488 | - | 293 | TIGR02790<br>PF00528<br>TIGR01097 | TIGR02790<br>BPD_transp_1<br>TIGR01097 | nickel_nikC: nickel ABC transporter, permease subunit NikC<br>Binding-protein-dependent transport system inner membrane component<br>PhnE: phosphonate ABC transporter, permease protein PhnE | 4.00E-56<br>9.30E-17<br>8.70E-06 |
| WP_017885991.1 | 52447 | 53353 | + | 302 | INFERRED GENE | - | - | - |

| | | | | | | | | |
|---|---|---|---|---|---|---|---|---|
| WP_065960203.1 | 55068 | 53427 | - | 546 | PF00496<br>TIGR04028<br>TIGR02294 | SBP_bac_5<br>TIGR04028<br>TIGR02294 | Bacterial extracellular solute-binding proteins, family 5 Middle<br>SBP_KPN_01854: ABC transporter substrate binding protein, KPN_01854 family<br>nickel_nikA: nickel ABC transporter, nickel/metallophore periplasmic binding protein | 4.70E-77<br>8.20E-65<br>1.50E-59 |
| WP_141693338.1 | 55211 | 55421 | + | 69 | LinA | LinA | LinA (linaridin precursor) | 3.40E-08 |
| WP_065960206.1 | 55417 | 55606 | + | 62 | LinA | LinA | LinA (linaridin precursor) | 8.50E-10 |
| WP_141693333.1 | 56532 | 56292 | - | 79 | NO PFAM MATCH | - | - | - |
| WP_141693334.1 | 56441 | 57551 | + | 369 | LinH<br>LinE<br>LinG<br>PF12697 | LinH<br>LinE<br>LinG<br>Abhydrolase_6 | LinH (Fusion of LinE/LinG)<br>LinE Homolog<br>LinG Homolog<br>Alpha/beta hydrolase family | 2.10E-65<br>8.70E-49<br>2.30E-13<br>1.60E-06 |
| WP_141693335.1 | 57547 | 58174 | + | 208 | LinL | LinL | LinL | 8.30E-48 |
| WP_141693336.1 | 58160 | 58484 | + | 107 | NO PFAM MATCH | - | - | - |
| WP_018171101.1 | 58575 | 58776 | + | 67 | INFERRED GENE | - | - | - |
| WP_065960215.1 | 60277 | 59242 | - | 344 | TIGR01297<br>PF01545 | TIGR01297<br>Cation_efflux | CDF: cation diffusion facilitator family transporter<br>Cation efflux family | 1.10E-38<br>8.80E-30 |
| WP_065960219.1 | 60535 | 63805 | + | 1089 | TIGR00392<br>PF00133<br>TIGR00422<br>TIGR00395<br>TIGR00396 | TIGR00392<br>tRNA-synt_1<br>TIGR00422<br>TIGR00395<br>TIGR00396 | ileS: isoleucine--tRNA ligase<br>tRNA synthetases class I (I, L, M and V)<br>valS: valine--tRNA ligase<br>leuS_arch: leucine--tRNA ligase<br>leuS_bact: leucine--tRNA ligase | 1.40E-241<br>2.30E-166<br>7.80E-95<br>1.00E-34<br>1.00E-31 |
| WP_065960272.1 | 63987 | 65349 | + | 453 | TIGR01499<br>TIGR01085<br>PF08245<br>TIGR01143<br>TIGR02068 | TIGR01499<br>TIGR01085<br>Mur_ligase_M<br>TIGR01143<br>TIGR02068 | folC: bifunctional protein FolC<br>murE: UDP-N-acetylmuramyl-tripeptide synthetase<br>Mur ligase middle domain<br>murF: UDP-N-acetylmuramoyl-tripeptide--D-alanyl-D-alanine ligase<br>cya_phycin_syn: cyanophycin synthetase | 3.10E-103<br>1.30E-22<br>3.70E-17<br>1.40E-13<br>5.90E-12 |
| WP_065960221.1 | 65345 | 65762 | + | 138 | PF14017 | DUF4233 | Protein of unknown function (DUF4233) | 1.60E-21 |
| WP_065960223.1 | 65795 | 66215 | + | 139 | PF00334 | NDK | Nucleoside diphosphate kinase | 2.00E-50 |

| index | leader | core | start | end | dir | score |
|---|---|---|---|---|---|---|
| 1 | MDAAVNRELARIETERAYVSHREPEPVL | TTPAAVAIGYAAVVGYAAQGAAFTAGATAVVGAYAVGKAVG | 55211 | 55421 | + | 29 |
| 2 | MTNLDAAVNRALQAFPQTQREFSAQLWAREAVPVL | TTPTAFMAGVAFSSAIINAYNAGANAG | 55417 | 55606 | + | 28 |
| 3 | MGIADALGTVAVVECVAATLVVL | HTVSGLVDSFRRWVTDCAEPGSGAA...SLVVLFAGIFGEPPLHRRIARSKRH | 58160 | 58484 | + | 6 |
| 4 | VRAVEQALLATARSHDPERVPWAIA...VTAQHAAVAAFVGGRTGEHPASLWP | QTVSGAMLGAAAAAFGVWVADGVARRPLVEYVGAALTPVAAGLDAR | 58818 | 59163 | + | 7 |
| 5 | VEQALLATARSHDPERVPWAIAQAE...VTAQHAAVAAFVGGRTGEHPASLWP | QTVSGAMLGAAAAAFGVWVADGVARRPLVEYVGAALTPVAAGLDAR | 58827 | 59163 | + | 7 |
| 6 | VPWAIAQAEVMGIGDELVASVAARVTAQHAAVAAFVGGRTGEHPASLWP | QTVSGAMLGAAAAAFGVWVADGVARRPLVEYVGAALTPVAAGLDAR | 58875 | 59163 | + | 8 |
| 7 | VMGIGDELVASVAARVTAQHAAVAAFVGGRTGEHPASLWP | QTVSGAMLGAAAAAFGVWVADGVARRPLVEYVGAALTPVAAGLDAR | 58902 | 59163 | + | 8 |
| 8 | MGIGDELVASVAARVTAQHAAVAAFVGGRTGEHPASLWP | QTVSGAMLGAAAAAFGVWVADGVARRPLVEYVGAALTPVAAGLDAR | 58905 | 59163 | + | 8 |
| 9 | VAARVTAQHAAVAAFVGGRTGEHPASLWP | QTVSGAMLGAAAAAFGVWVADGVARRPLVEYVGAALTPVAAGLDAR | 58935 | 59163 | + | 6 |

### Results for WP_051597146.1 [Curtobacterium sp. UNCCL17] back to top

previous - next

#### Architecture

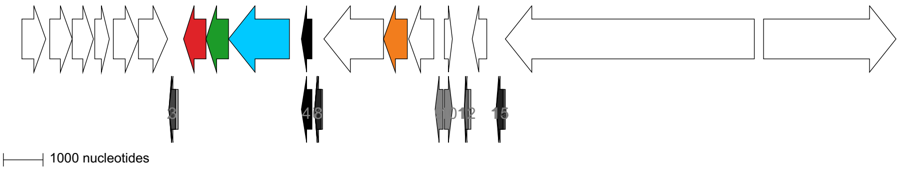

Link to nucleotide sequence

| Accession | start | end | direction | length (aa) | Pfam/HMM | name | description | E-value |
|---|---|---|---|---|---|---|---|---|
| WP_027467252.1 | 147058 | 147652 | + | 197 | PF02632 | BioY | BioY family | 1.60E-38 |
| WP_155896961.1 | 147737 | 148319 | + | 193 | NO PFAM MATCH | - | - | - |
| WP_051597131.1 | 148315 | 148846 | + | 176 | NO PFAM MATCH | - | - | - |
| WP_051597132.1 | 148895 | 149237 | + | 113 | NO PFAM MATCH | - | - | - |
| WP_027467253.1 | 149341 | 149971 | + | 209 | PF00440<br>TIGR03968<br>TIGR03613 | TetR_N<br>TIGR03968<br>TIGR03613 | Bacterial regulatory proteins, tetR family<br>mycofact_TetR: mycofactocin system transcriptional regulator<br>RutR: pyrimidine utilization regulatory protein R | 1.60E-14<br>2.30E-06<br>2.00E-04 |
| WP_027467254.1 | 149998 | 150712 | + | 237 | PF03807<br>TIGR01915<br>PF02826 | F420_oxidored<br>TIGR01915<br>2-Hacid_dh_C | NADP oxidoreductase coenzyme F420-dependent<br>npdG: NADPH-dependent F420 reductase<br>D-isomer specific 2-hydroxyacid dehydrogenase, NAD binding domain | 1.00E-17<br>8.90E-10<br>6.30E-05 |
| WP_035809673.1 | 151699 | 151126 | - | 190 | TIGR00521<br>TIGR02113<br>PF02441 | TIGR00521<br>TIGR02113<br>Flavoprotein | coaBC_dfp: phosphopantothenoylcysteine decarboxylase / phosphopantothenate--cysteine ligase<br>coaC_strep: phosphopantothenoylcysteine decarboxylase<br>Flavoprotein | 2.20E-08<br>2.40E-07<br>1.30E-06 |
| WP_027467256.1 | 152259 | 151701 | - | 185 | LinL | LinL | LinL | 2.60E-51 |

| | | | | | | | | |
|---|---|---|---|---|---|---|---|---|
| WP_051597146.1 | 153778 | 152245 | - | 510 | LinH<br>LinE<br>LinG<br>PF12697<br>TIGR02427 | LinH<br>LinE<br>LinG<br>Abhydrolase_6<br>TIGR02427 | LinH (Fusion of LinE/LinG)<br>LinE Homolog<br>LinG Homolog<br>Alpha/beta hydrolase family<br>protocat_pcaD: 3-oxoadipate enol-lactonase | 1.60E-155<br>3.10E-72<br>2.50E-65<br>6.70E-09<br>2.60E-04 |
| WP_146247959.1 | 154369 | 154090 | - | 92 | LinA | LinA | LinA (linaridin precursor) | 6.00E-11 |
| WP_027467257.1 | 156156 | 154677 | - | 492 | NO PFAM MATCH | - | - | - |
| WP_027467258.1 | 156782 | 156152 | - | 209 | TIGR02857<br>TIGR02673<br>TIGR01189<br>TIGR03608<br>TIGR01188 | TIGR02857<br>TIGR02673<br>TIGR01189<br>TIGR03608<br>TIGR01188 | CydD: thiol reductant ABC exporter, CydD subunit<br>FtsE: cell division ATP-binding protein FtsE<br>ccmA: heme ABC exporter, ATP-binding protein CcmA<br>L_ocin_972_ABC: putative bacteriocin export ABC transporter, lactococcin 972 group<br>drrA: daunorubicin resistance ABC transporter, ATP-binding protein | 3.50E-30<br>6.50E-30<br>7.00E-30<br>1.20E-25<br>2.30E-25 |
| WP_051597133.1 | 157423 | 156811 | - | 203 | PF13026 | DUF3887 | Protein of unknown function (DUF3887) | 4.40E-14 |
| WP_155896962.1 | 157691 | 157901 | + | 69 | NO PFAM MATCH | - | - | - |
| WP_051597134.1 | 158765 | 158402 | - | 120 | NO PFAM MATCH | - | - | - |
| WP_081827479.1 | 165536 | 159248 | - | 2095 | TIGR01643<br>TIGR03696<br>PF05593 | TIGR01643<br>TIGR03696<br>RHS_repeat | YD_repeat_2x: YD repeat (two copies)<br>Rhs_assc_core: RHS repeat-associated core domain<br>RHS Repeat | 4.30E-18<br>2.70E-14<br>3.60E-11 |
| WP_027467260.1 | 165752 | 169112 | + | 1119 | NO PFAM MATCH | - | - | - |

| index | leader | core | start | end | dir | score |
|---|---|---|---|---|---|---|
| 1 | MPGVGDLPAVEQQRRADRLRGLQPRSVASPMG | ETARSSYCSTGATRGPPRSQG | 150915 | 150753 | - | 6 |
| 2 | VMPGVGDLPAVEQQRRADRLRGLQPRSVASPMG | ETARSSYCSTGATRGPPRSQG | 150918 | 150753 | - | 6 |
| 3 | MATSICCGVTVGPALRLAFAALPLL...PAVEQQRRADRLRGLQPRSVASPMG | ETARSSYCSTGATRGPPRSQG | 150999 | 150753 | - | 6 |
| 4 | MPVDARNVLRLRGASPLPPPHEVHH...VQAFANAALSSAEAGARADASIPAM | TTPTIVAAAEFVVAGSTICLIC | 154369 | 154090 | - | 33 |
| 5 | LDHGGPGPEAWNMSIEGGFECCSVPSG | LSWRRSHRSRACSSSRTPSRRALNCNA | 154583 | 154418 | - | 6 |
| 6 | VPKARGFWTMAVQGPKPGTCRSRGASNV | VQYRAVSAGAALIGAARARRLAPRHVER | 154603 | 154432 | - | 6 |
| 7 | VVPKARGFWTMAVQGPKPGTCRSRGASN | VVQYRAVSAGAALIGAARARRLAPRHVER | 154606 | 154432 | - | 7 |
| 8 | VTAPFRGGSGAEGARLLDHGGPGPEAWNMSIEGGF | ECCSVPSGLSWRRSHRSRACSSSRTPSRRALNCNA | 154631 | 154418 | - | 6 |
| 9 | VSAGTGPHRCGTPVRGRAGTDRKQGTFRSGRASA | GRVHRAAMQRRCCLTASPEAAEAQASLPRLPSIRP | 157666 | 157456 | - | 7 |
| 10 | MNLLLQHPEKTGHEWGGAGWVIRVP...WKRGLLQRLHSEPKIAEDTIYPAIT | ETARLAKHIVKQAVTQQ | 157691 | 157901 | + | 7 |
| 11 | MRLGGGEGRDFGASLVDDRRL | GKEATALTKRAREGGEEPPRT | 158316 | 158187 | - | 6 |
| 12 | LTRERGEFDEDPIYFLKRIMRLGGGEGRDF | GASLVDDRRLGKEATALTKRAREGGEEPPRT | 158373 | 158187 | - | 6 |
| 13 | VELHDDANEVVAEVFEDDETGACIFSMPN | GASVPVSHIRELLRRAALEFPGAVDCPAD | 159186 | 159009 | - | 7 |
| 14 | MAVELHDDANEVVAEVFEDDETGACIFSMP | NGASVPVSHIRELLRRAALEFPGAVDCPAD | 159192 | 159009 | - | 7 |
| 15 | LTDYSFVLASFVGERDGMAVELHDDANEVVAEVFEDDE | TGACIFSMPNGASVPVSHIRELLRRAALEFPGAVDCPAD | 159243 | 159009 | - | 7 |

### Results for SFF54177.1 [Curtobacterium sp. YR515] back to top

previous - next

#### Architecture

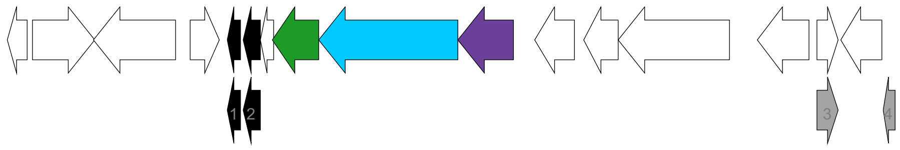

1000 nucleotides

Link to nucleotide sequence

| Accession | start | end | direction | length (aa) | Pfam/HMM | name | description | E-value |
|---|---|---|---|---|---|---|---|---|
| SFF54079.1 | 432360 | 432099 | - | 86 | PF04024<br>TIGR02978 | PspC<br>TIGR02978 | PspC domain<br>phageshock_pspC: phage shock protein C | 1.50E-19<br>6.90E-13 |
| SFF54089.1 | 432426 | 433248 | + | 273 | TIGR00507<br>TIGR01809<br>PF08501 | TIGR00507<br>TIGR01809<br>Shikimate_dh_N | aroE: shikimate dehydrogenase<br>Shik-DH-AROM: shikimate-5-dehydrogenase<br>Shikimate dehydrogenase substrate binding domain | 1.30E-41<br>6.80E-25<br>7.00E-17 |
| SFF54101.1 | 434299 | 433228 | - | 356 | TIGR00233<br>PF00579<br>TIGR00234 | TIGR00233<br>tRNA-synt_1b<br>TIGR00234 | trpS: tryptophan--tRNA ligase<br>tRNA synthetases class I (W and Y)<br>tyrS: tyrosine--tRNA ligase | 1.20E-70<br>8.20E-49<br>7.00E-09 |
| SFF54114.1 | 434476 | 434863 | + | 128 | NO PFAM MATCH | - | - | - |
| SFF54125.1 | 435135 | 434970 | - | 54 | LinA | LinA | LinA (linaridin precursor) | 3.30E-11 |
| SFF54139.1 | 435399 | 435165 | - | 77 | LinA | LinA | LinA (linaridin precursor) | 3.50E-15 |
| SFF54151.1 | 435563 | 435395 | - | 55 | NO PFAM MATCH | - | - | - |
| SFF54165.1 | 436162 | 435559 | - | 200 | LinL | LinL | LinL | 1.90E-54 |

| | | | | | | | | |
|---|---|---|---|---|---|---|---|---|
| SFF54177.1 | 437967 | 436158 | - | 602 | LinH<br>LinG<br>LinE<br>PF12697<br>TIGR02427 | LinH<br>LinG<br>LinE<br>Abhydrolase_6<br>TIGR02427 | LinH (Fusion of LinE/LinG)<br>LinG Homolog<br>LinE Homolog<br>Alpha/beta hydrolase family<br>protocat_pcaD: 3-oxoadipate enol-lactonase | 7.10E-197<br>6.30E-83<br>9.40E-63<br>3.20E-13<br>2.70E-10 |
| SFF54191.1 | 438695 | 437963 | - | 243 | PF13649<br>PF08241<br>PF13847<br>PF08242<br>PF13489 | Methyltransf_25<br>Methyltransf_11<br>Methyltransf_31<br>Methyltransf_12<br>Methyltransf_23 | Methyltransferase domain<br>Methyltransferase domain<br>Methyltransferase domain<br>Methyltransferase domain<br>Methyltransferase domain | 2.10E-18<br>2.30E-14<br>1.40E-12<br>1.50E-11<br>2.20E-11 |
| SFF54203.1 | 439478 | 438968 | - | 169 | NO PFAM MATCH | - | - | - |
| SFF54211.1 | 440044 | 439600 | - | 147 | PF10604<br>PF08327 | Polyketide_cyc2<br>AHSA1 | Polyketide cyclase / dehydrase and lipid transport<br>Activator of Hsp90 ATPase homolog 1-like protein | 1.00E-07<br>6.30E-04 |
| SFF54221.1 | 441492 | 440046 | - | 481 | TIGR02734<br>PF13450<br>TIGR03467<br>PF01593<br>PF12831 | TIGR02734<br>NAD_binding_8<br>TIGR03467<br>Amino_oxidase<br>FAD_oxidored | crtI_fam: phytoene desaturase<br>NAD(P)-binding Rossmann-like domain<br>HpnE: squalene-associated FAD-dependent desaturase<br>Flavin containing amine oxidoreductase<br>FAD dependent oxidoreductase | 1.00E-15<br>2.50E-11<br>7.10E-11<br>5.90E-09<br>2.00E-08 |
| SFF54230.1 | 442528 | 441856 | - | 223 | NO PFAM MATCH | - | - | - |
| SFF54240.1 | 442631 | 442907 | + | 91 | PF13828 | DUF4190 | Domain of unknown function (DUF4190) | 7.50E-05 |
| SFF54250.1 | 443467 | 442939 | - | 175 | PF13302<br>PF00583<br>PF13420 | Acetyltransf_3<br>Acetyltransf_1<br>Acetyltransf_4 | Acetyltransferase (GNAT) domain<br>Acetyltransferase (GNAT) family<br>Acetyltransferase (GNAT) domain | 3.40E-12<br>3.60E-10<br>4.40E-07 |

| index | leader | core | start | end | dir | score |
|---|---|---|---|---|---|---|
| 1 | MQQTRQLDAVPFDSDEATTFCGLRPVA | ATPAAVAWAGAVVAAAGVGAAIGDAVD | 435135 | 434970 | - | 30 |
| 2 | MTSAHRAGNRAQRPTITIAERKESSMTTIADIAFDSDRGQRFAPEQPLA | ATPLAALAATSVFTGAVAAYAVEEAVGN | 435399 | 435165 | - | 28 |
| 3 | MVTYSESPRTGLGTASLVLGVCSLLAG | WTFVAPIVGLCLGVASRSREPFARG...VWILLLVLLVVGGAFAVWSGATQGR | 442631 | 442907 | + | 6 |
| 4 | VLKDLPGLQIMQFAPSSVITIILVVMFIAFLA | GTLPARRAARQNPIEALRYE | 443651 | 443492 | - | 6 |

### Results for GCE02330.1 [Embleya hyalina] back to top

previous - next

#### Architecture

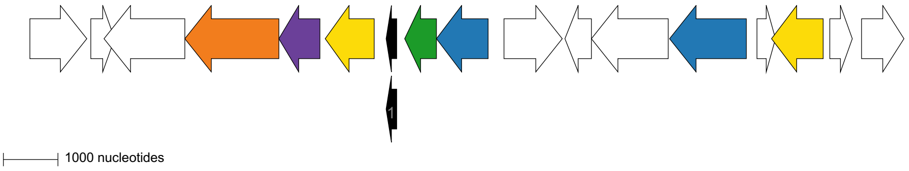

1000 nucleotides

Link to nucleotide sequence

| Accession | start | end | direction | length (aa) | Pfam/HMM | name | description | E-value |
|---|---|---|---|---|---|---|---|---|
| GCE02322.1 | 44498 | 45530 | + | 343 | PF12833<br>PF14525<br>TIGR04094<br>PF00165 | HTH_18<br>AraC_binding_2<br>TIGR04094<br>HTH_AraC | Helix-turn-helix domain<br>AraC-binding-like domain<br>adjacent_YSIRK: YSIRK-targeted surface antigen transcriptional regulator<br>Bacterial regulatory helix-turn-helix proteins, AraC family | 2.60E-18<br>2.70E-10<br>6.60E-06<br>5.30E-04 |
| GCE02323.1 | 45623 | 46043 | + | 139 | NO PFAM MATCH | - | - | - |
| GCE02324.1 | 47335 | 45865 | - | 489 | TIGR02734<br>TIGR02733<br>TIGR03467<br>TIGR01317<br>TIGR00292 | TIGR02734<br>TIGR02733<br>TIGR03467<br>TIGR01317<br>TIGR00292 | crtI_fam: phytoene desaturase<br>desat_CrtD: C-3',4' desaturase CrtD<br>HpnE: squalene-associated FAD-dependent desaturase<br>GOGAT_sm_gam: glutamate synthase, NADH/NADPH, small subunit<br>TIGR00292: thiazole biosynthesis enzyme | 7.40E-14<br>7.70E-09<br>7.70E-09<br>9.30E-08<br>2.00E-07 |
| GCE02325.1 | 49065 | 47331 | - | 577 | TIGR02204<br>TIGR02203<br>TIGR00958<br>TIGR03796<br>TIGR01846 | TIGR02204<br>TIGR02203<br>TIGR00958<br>TIGR03796<br>TIGR01846 | MsbA_rel: ABC transporter, permease/ATP-binding protein<br>MsbA_lipidA: lipid A export permease/ATP-binding protein MsbA<br>3a01208: antigen peptide transporter 2<br>NHLM_micro_ABC1: NHLM bacteriocin system ABC transporter, peptidase/ATP-binding protein<br>type_I_sec_HlyB: type I secretion system ATPase | 1.20E-102<br>1.80E-95<br>2.40E-84<br>8.30E-82<br>7.70E-80 |
| GCE02326.1 | 49817 | 49061 | - | 251 | PF13649<br>PF08241<br>PF13489<br>PF08242<br>PF13847 | Methyltransf_25<br>Methyltransf_11<br>Methyltransf_23<br>Methyltransf_12<br>Methyltransf_31 | Methyltransferase domain<br>Methyltransferase domain<br>Methyltransferase domain<br>Methyltransferase domain<br>Methyltransferase domain | 1.10E-13<br>2.70E-09<br>9.10E-08<br>5.50E-07<br>9.20E-07 |
| GCE02327.1 | 50799 | 49899 | - | 299 | LinG<br>LinH<br>TIGR04033 | LinG<br>LinH<br>TIGR04033 | LinG Homolog<br>LinH (Fusion of LinE/LinG)<br>export_SdpB: antimicrobial peptide system protein, SdpB family | 2.50E-102<br>1.80E-73<br>6.90E-04 |
| GCE02328.1 | 51226 | 51028 | - | 65 | LinA | LinA | LinA (linaridin precursor) | 4.10E-12 |
| GCE02329.1 | 51945 | 51378 | - | 188 | LinL | LinL | LinL | 9.20E-55 |
| GCE02330.1 | 52889 | 51941 | - | 315 | LinE<br>LinH<br>PF12697<br>PF00561<br>TIGR02427 | LinE<br>LinH<br>Abhydrolase_6<br>Abhydrolase_1<br>TIGR02427 | LinE Homolog<br>LinH (Fusion of LinE/LinG)<br>Alpha/beta hydrolase family<br>alpha/beta hydrolase fold<br>protocat_pcaD: 3-oxoadipate enol-lactonase | 1.30E-91<br>2.10E-41<br>2.80E-15<br>1.40E-09<br>7.80E-08 |
| GCE02331.1 | 53189 | 54263 | + | 357 | PF12840<br>PF01022<br>PF12802 | HTH_20<br>HTH_5<br>MarR_2 | Helix-turn-helix domain<br>Bacterial regulatory protein, arsR family<br>MarR family | 3.20E-08<br>5.90E-06<br>2.30E-04 |

| | | | | | | | | |
|---|---|---|---|---|---|---|---|---|
| GCE02332.1 | 54802 | 54295 | - | 168 | PF12770 | CHAT | CHAT domain | 4.40E-26 |
| GCE02333.1 | 56199 | 54798 | - | 466 | PF17874<br>PF13424<br>PF14559<br>PF13428 | TPR_MalT<br>TPR_12<br>TPR_19<br>TPR_14 | MalT-like TPR region<br>Tetratricopeptide repeat<br>Tetratricopeptide repeat<br>Tetratricopeptide repeat | 1.30E-08<br>2.60E-07<br>2.60E-04<br>5.20E-04 |
| GCE02334.1 | 57631 | 56227 | - | 467 | PF02450<br>PF07819<br>LinE | LCAT<br>PGAP1<br>LinE | Lecithin:cholesterol acyltransferase<br>PGAP1-like protein<br>LinE Homolog | 8.40E-10<br>9.60E-08<br>5.00E-06 |
| GCE02335.1 | 57816 | 58164 | + | 115 | NO PFAM MATCH | - | - | - |
| GCE02336.1 | 59033 | 58103 | - | 309 | PF00561<br>PF12697<br>LinH<br>TIGR02427<br>LinE | Abhydrolase_1<br>Abhydrolase_6<br>LinH<br>TIGR02427<br>LinE | alpha/beta hydrolase fold<br>Alpha/beta hydrolase family<br>LinH (Fusion of LinE/LinG)<br>protocat_pcaD: 3-oxoadipate enol-lactonase<br>LinE Homolog | 1.80E-23<br>2.80E-20<br>6.20E-19<br>2.00E-18<br>2.60E-18 |
| GCE02337.1 | 59170 | 59581 | + | 136 | PF12840 | HTH_20 | Helix-turn-helix domain | 8.10E-12 |
| GCE02338.1 | 59757 | 60525 | + | 255 | NO PFAM MATCH | - | - | - |

| index | leader | core | start | end | dir | score |
|---|---|---|---|---|---|---|
| 1 | MSPVAQFANASLTDVSPGMLTGIDAARLDSPADPVM | ATPLITAVFAVATDMLTQNIANYVNRNAH | 51226 | 51028 | - | 32 |

### Results for WP_078981861.1 [Embleya scabrispora] back to top

previous - next

#### Architecture

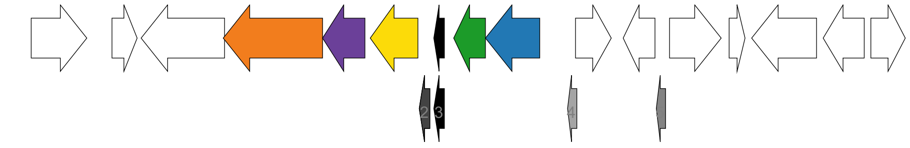

1000 nucleotides

Link to nucleotide sequence

| Accession | start | end | direction | length (aa) | Pfam/HMM | name | description | E-value |
|---|---|---|---|---|---|---|---|---|
| WP_078981856.1 | 281872 | 282859 | + | 328 | PF01978<br>PF12840<br>PF12802<br>PF13384<br>PF13551 | TrmB<br>HTH_20<br>MarR_2<br>HTH_23<br>HTH_29 | Sugar-specific transcriptional regulator TrmB<br>Helix-turn-helix domain<br>MarR family<br>Homeodomain-like domain<br>Winged helix-turn helix | 3.50E-08<br>2.70E-07<br>3.90E-07<br>2.60E-06<br>2.20E-05 |
| WP_101897533.1 | 283289 | 283727 | + | 145 | NO PFAM MATCH | - | - | - |
| WP_078981858.1 | 285267 | 283806 | - | 486 | TIGR02734<br>TIGR02733<br>TIGR01317<br>TIGR00292<br>TIGR03467 | TIGR02734<br>TIGR02733<br>TIGR01317<br>TIGR00292<br>TIGR03467 | crtI_fam: phytoene desaturase<br>desat_CrtD: C-3',4' desaturase CrtD<br>GOGAT_sm_gam: glutamate synthase, NADH/NADPH, small subunit<br>TIGR00292: thiazole biosynthesis enzyme<br>HpnE: squalene-associated FAD-dependent desaturase | 5.30E-11<br>1.60E-08<br>5.00E-08<br>5.90E-08<br>1.70E-07 |
| WP_078981859.1 | 286997 | 285263 | - | 577 | TIGR02204<br>TIGR02203<br>TIGR00958<br>TIGR03796<br>TIGR01846 | TIGR02204<br>TIGR02203<br>TIGR00958<br>TIGR03796<br>TIGR01846 | MsbA_rel: ABC transporter, permease/ATP-binding protein<br>MsbA_lipidA: lipid A export permease/ATP-binding protein MsbA<br>3a01208: antigen peptide transporter 2<br>NHLM_micro_ABC1: NHLM bacteriocin system ABC transporter, peptidase/ATP-binding protein<br>type_I_sec_HlyB: type I secretion system ATPase | 2.10E-104<br>2.40E-97<br>1.00E-83<br>3.70E-83<br>5.70E-80 |
| WP_078982366.1 | 287749 | 286993 | - | 251 | PF13649<br>PF08241<br>PF13489<br>PF08242<br>PF13847 | Methyltransf_25<br>Methyltransf_11<br>Methyltransf_23<br>Methyltransf_12<br>Methyltransf_31 | Methyltransferase domain<br>Methyltransferase domain<br>Methyltransferase domain<br>Methyltransferase domain<br>Methyltransferase domain | 1.60E-13<br>6.80E-09<br>3.30E-08<br>3.70E-07<br>6.80E-06 |
| WP_078982367.1 | 288680 | 287831 | - | 282 | LinG<br>LinH<br>TIGR04033 | LinG<br>LinH<br>TIGR04033 | LinG Homolog<br>LinH (Fusion of LinE/LinG)<br>export_SdpB: antimicrobial peptide system protein, SdpB family | 3.90E-102<br>1.90E-73<br>5.10E-05 |
| WP_161501002.1 | 289154 | 288956 | - | 65 | LinA | LinA | LinA (linaridin precursor) | 2.60E-12 |
| WP_078981860.1 | 289873 | 289306 | - | 188 | LinL | LinL | LinL | 6.70E-56 |
| WP_078981861.1 | 290817 | 289869 | - | 315 | LinE<br>LinH<br>PF12697<br>PF00561<br>TIGR02427 | LinE<br>LinH<br>Abhydrolase_6<br>Abhydrolase_1<br>TIGR02427 | LinE Homolog<br>LinH (Fusion of LinE/LinG)<br>Alpha/beta hydrolase family<br>alpha/beta hydrolase fold<br>protocat_pcaD: 3-oxoadipate enol-lactonase | 2.10E-88<br>5.80E-41<br>9.20E-17<br>3.00E-11<br>9.70E-08 |
| WP_101897565.1 | 291452 | 292085 | + | 210 | NO PFAM MATCH | - | - | - |
| WP_078982368.1 | 292853 | 292295 | - | 185 | PF09346 | SMI1_KNR4 | SMI1 / KNR4 family (SUKH-1) | 6.30E-06 |
| WP_078981863.1 | 293105 | 294011 | + | 301 | TIGR00654<br>PF02567 | TIGR00654<br>PhzC-PhzF | PhzF_family: phenazine biosynthesis protein, PhzF family<br>Phenazine biosynthesis-like protein | 2.70E-37<br>4.40E-32 |
| WP_107504212.1 | 294168 | 294438 | + | 89 | PF14200 | RicinB_lectin_2 | Ricin-type beta-trefoil lectin domain-like | 2.00E-06 |
| WP_161501003.1 | 295704 | 294555 | - | 382 | PF07690<br>TIGR00880<br>TIGR00710<br>TIGR00900<br>TIGR00893 | MFS_1<br>TIGR00880<br>TIGR00710<br>TIGR00900<br>TIGR00893 | Major Facilitator Superfamily<br>2_A_01_02: multidrug resistance protein<br>efflux_Bcr_CflA: drug resistance transporter, Bcr/CflA subfamily<br>2A0121: H+ Antiporter protein<br>2A0114: D-galactonate transporter | 2.30E-33<br>1.10E-15<br>3.00E-14<br>1.00E-11<br>4.00E-10 |
| WP_161501004.1 | 296534 | 295823 | - | 236 | PF04954<br>PF08021 | SIP<br>FAD_binding_9 | Siderophore-interacting protein<br>Siderophore-interacting FAD-binding domain | 3.00E-21<br>7.20E-19 |

| WP_020548778.1 | 296662 | 297258 | + | 198 | INFERRED GENE | - | - | - |
|---|---|---|---|---|---|---|---|---|

| index | leader | core | start | end | dir | score |
|---|---|---|---|---|---|---|
| 1 | VPRRRRGRRRSQRRRAARARRAMFGVRRTAPRR | PTPSQPGTGALPVPGPHPHRKETPCSVA | 288883 | 288697 | - | 7 |
| 2 | VTVPAVPRRRRGRRRSQRRRAARARRAMFGVRRTAPRR | PTPSQPGTGALPVPGPHPHRKETPCSVA | 288898 | 288697 | - | 7 |
| 3 | MSPVAQFANASLTDVSPGMLSGIDAARLDSPADPVM | ATPLITAVFAVATDMLAQNIANYVNRNAH | 289154 | 288956 | - | 32 |
| 4 | VHGRRPRAVGRREGARCGWGVSARLRA | VGGRRSRRTGGSPGRRRTGASGPTPPSP | 291471 | 291303 | - | 6 |
| 5 | MKRSGSPPRAFAASMRHETGCAPPAS | ASKATSTGGLSVSWRYGDTVRVPSGRR | 293035 | 292873 | - | 7 |

### Results for GAB19727.1 [Gordonia effusa NBRC 100432] back to top

previous - next

#### Architecture

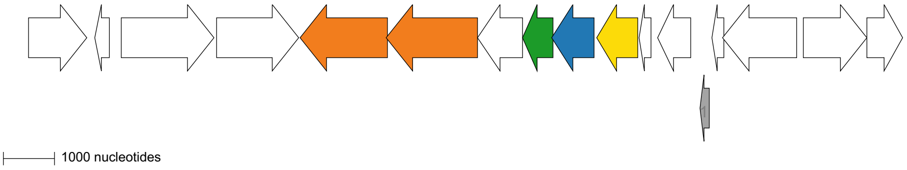

Link to nucleotide sequence

| Accession | start | end | direction | length (aa) | Pfam/HMM | name | description | E-value |
|---|---|---|---|---|---|---|---|---|
| GAB19719.1 | 14616 | 15774 | + | 385 | TIGR01304<br>PF00478<br>TIGR01302<br>PF01070<br>TIGR01303 | TIGR01304<br>IMPDH<br>TIGR01302<br>FMN_dh<br>TIGR01303 | IMP_DH_rel_2: IMP dehydrogenase family protein<br>IMP dehydrogenase / GMP reductase domain<br>IMP_dehydrog: inosine-5'-monophosphate dehydrogenase<br>FMN-dependent dehydrogenase<br>IMP_DH_rel_1: IMP dehydrogenase family protein | 2.20E-171<br>1.10E-27<br>2.10E-24<br>5.10E-09<br>6.10E-08 |
| GAB19720.1 | 16197 | 15921 | - | 91 | NO PFAM MATCH | - | - | - |
| GAB19721.1 | 16443 | 18240 | + | 598 | PF05199<br>PF01266<br>PF00732<br>TIGR03970<br>PF00890 | GMC_oxred_C<br>DAO<br>GMC_oxred_N<br>TIGR03970<br>FAD_binding_2 | GMC oxidoreductase<br>FAD dependent oxidoreductase<br>GMC oxidoreductase<br>Rv0697: dehydrogenase, Rv0697 family<br>FAD binding domain | 7.30E-12<br>1.70E-11<br>1.40E-10<br>2.20E-09<br>4.80E-08 |
| GAB19722.1 | 18310 | 19903 | + | 530 | TIGR00884<br>TIGR00888<br>PF00958<br>PF00117<br>TIGR01815 | TIGR00884<br>TIGR00888<br>GMP_synt_C<br>GATase<br>TIGR01815 | guaA_Cterm: GMP synthase (glutamine-hydrolyzing), C-terminal domain<br>guaA_Nterm: GMP synthase (glutamine-hydrolyzing), N-terminal domain<br>GMP synthase C terminal domain<br>Glutamine amidotransferase class-I<br>TrpE-clade3: anthranilate synthase | 1.20E-141<br>1.10E-68<br>2.80E-38<br>3.20E-35<br>1.20E-16 |
| GAB19723.1 | 21632 | 19922 | - | 569 | TIGR02857<br>TIGR03375<br>TIGR02204<br>TIGR02203<br>TIGR03796 | TIGR02857<br>TIGR03375<br>TIGR02204<br>TIGR02203<br>TIGR03796 | CydD: thiol reductant ABC exporter, CydD subunit<br>type_I_sec_LssB: type I secretion system ATPase<br>MsbA_rel: ABC transporter, permease/ATP-binding protein<br>MsbA_lipidA: lipid A export permease/ATP-binding protein MsbA<br>NHLM_micro_ABC1: NHLM bacteriocin system ABC transporter, peptidase/ATP-binding protein | 4.10E-67<br>1.10E-66<br>1.20E-64<br>3.60E-62<br>4.50E-62 |
| GAB19724.1 | 23404 | 21628 | - | 591 | TIGR03796<br>TIGR03375<br>TIGR02868<br>TIGR02857<br>TIGR02203 | TIGR03796<br>TIGR03375<br>TIGR02868<br>TIGR02857<br>TIGR02203 | NHLM_micro_ABC1: NHLM bacteriocin system ABC transporter, peptidase/ATP-binding protein<br>type_I_sec_LssB: type I secretion system ATPase<br>CydC: thiol reductant ABC exporter, CydC subunit<br>CydD: thiol reductant ABC exporter, CydD subunit<br>MsbA_lipidA: lipid A export permease/ATP-binding protein MsbA | 1.80E-45<br>1.10E-42<br>1.20E-42<br>6.00E-40<br>1.90E-33 |
| GAB19725.1 | 24294 | 23400 | - | 297 | PF01551 | Peptidase_M23 | Peptidase family M23 | 3.50E-14 |
| GAB19726.1 | 24872 | 24290 | - | 193 | LinL | LinL | LinL | 1.00E-27 |
| GAB19727.1 | 25674 | 24864 | - | 269 | LinE<br>LinH | LinE<br>LinH | LinE Homolog<br>LinH (Fusion of LinE/LinG) | 1.40E-19<br>7.00E-16 |
| GAB19728.1 | 26546 | 25727 | - | 272 | LinG<br>LinH<br>TIGR04033 | LinG<br>LinH<br>TIGR04033 | LinG Homolog<br>LinH (Fusion of LinE/LinG)<br>export_SdpB: antimicrobial peptide system protein, SdpB family | 2.60E-51<br>9.60E-45<br>3.20E-04 |
| GAB19729.1 | 26791 | 26575 | - | 71 | NO PFAM MATCH | - | - | - |
| GAB19730.1 | 27570 | 26919 | - | 216 | PF09203 | MspA | MspA | 3.60E-34 |
| GAB19731.1 | 28212 | 27999 | - | 70 | NO PFAM MATCH | - | - | - |
| GAB19732.1 | 29635 | 28204 | - | 476 | PF04024<br>TIGR02978<br>PF09922 | PspC<br>TIGR02978<br>DUF2154 | PspC domain<br>phageshock_pspC: phage shock protein C<br>Cell wall-active antibiotics response 4TMS YvqF | 2.80E-14<br>6.90E-11<br>2.90E-06 |
| GAB19733.1 | 29767 | 31021 | + | 417 | PF04024<br>TIGR02978<br>PF13581<br>TIGR01924<br>PF02518 | PspC<br>TIGR02978<br>HATPase_c_2<br>TIGR01924<br>HATPase_c | PspC domain<br>phageshock_pspC: phage shock protein C<br>Histidine kinase-like ATPase domain<br>rsbW_low_gc: anti-sigma B factor<br>Histidine kinase-, DNA gyrase B-, and HSP90-like ATPase | 7.50E-15<br>1.80E-10<br>6.80E-06<br>6.80E-04<br>7.00E-04 |
| GAB19734.1 | 31028 | 31715 | + | 228 | PF00196<br>PF00072<br>TIGR02875<br>TIGR02154<br>TIGR02956 | GerE<br>Response_reg<br>TIGR02875<br>TIGR02154<br>TIGR02956 | Bacterial regulatory proteins, luxR family<br>Response regulator receiver domain<br>spore_0_A: sporulation transcription factor Spo0A<br>PhoB: phosphate regulon transcriptional regulatory protein PhoB<br>TMAO_torS: TMAO reductase sytem sensor TorS | 1.20E-18<br>3.00E-17<br>1.70E-11<br>3.60E-09<br>3.30E-08 |

| index | leader | core | start | end | dir | score |
|---|---|---|---|---|---|---|
| 1 | VRSADSGSRSRVAMASHARPRPCARQSAI | ALHANDIEQNRSVDRSWRATFRQVTAANMV | 27929 | 27749 | - | 6 |

### Results for SDZ55823.1 [Herbiconiux ginsengi] back to top

previous - next

#### Architecture

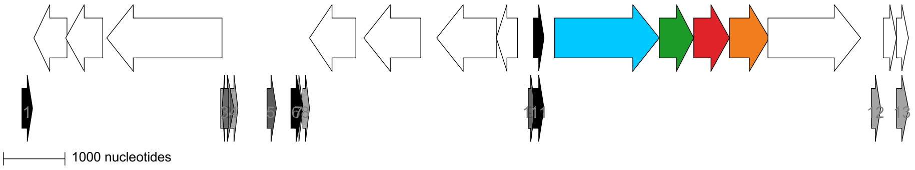

Link to nucleotide sequence

| Accession | start | end | direction | length (aa) | Pfam/HMM | name | description | E-value |
|---|---|---|---|---|---|---|---|---|
| SDZ55740.1 | 53888 | 53354 | - | 177 | TIGR02227<br>PF10502<br>TIGR02771<br>PF00717<br>TIGR02754 | TIGR02227<br>Peptidase_S26<br>TIGR02771<br>Peptidase_S24<br>TIGR02754 | sigpep_I_bact: signal peptidase I<br>Signal peptidase, peptidase S26<br>TraF_Ti: conjugative transfer signal peptidase TraF<br>Peptidase S24-like<br>sod_Ni_protease: nickel-type superoxide dismutase maturation protease | 1.40E-27<br>2.60E-16<br>6.70E-12<br>5.30E-11<br>1.10E-07 |
| SDZ55753.1 | 54484 | 53884 | - | 199 | PF03713<br>PF13628 | DUF305<br>DUF4142 | Domain of unknown function (DUF305)<br>Domain of unknown function (DUF4142) | 9.80E-37<br>8.30E-04 |
| SDZ55761.1 | 56412 | 54552 | - | 619 | PF01179<br>TIGR01167 | Cu_amine_oxid<br>TIGR01167 | Copper amine oxidase, enzyme domain<br>LPXTG_anchor: LPXTG cell wall anchor domain | 1.10E-31<br>7.10E-04 |
| SDZ55775.1 | 58584 | 57828 | - | 251 | PF13350 | Y_phosphatase3 | Tyrosine phosphatase family | 2.00E-56 |
| SDZ55786.1 | 59650 | 58708 | - | 313 | PF06742<br>PF06863 | DUF1214<br>DUF1254 | Protein of unknown function (DUF1214)<br>Protein of unknown function (DUF1254) | 2.00E-10<br>7.70E-10 |
| SDZ55797.1 | 60870 | 59895 | - | 324 | NO PFAM MATCH | - | - | - |
| SDZ55804.1 | 61213 | 60871 | - | 113 | PF03551<br>TIGR03433<br>TIGR02719 | PadR<br>TIGR03433<br>TIGR02719 | Transcriptional regulator PadR-like family<br>padR_acidobact: transcriptional regulator, Acidobacterial, PadR-family<br>repress_PhaQ: poly-beta-hydroxybutyrate-responsive repressor | 3.20E-19<br>4.00E-14<br>1.00E-07 |
| SDZ55813.1 | 61484 | 61649 | + | 54 | LinA | LinA | LinA (linaridin precursor) | 2.70E-13 |
| SDZ55823.1 | 61818 | 63525 | + | 568 | LinH<br>LinG<br>LinE<br>TIGR04033 | LinH<br>LinG<br>LinE<br>TIGR04033 | LinH (Fusion of LinE/LinG)<br>LinG Homolog<br>LinE Homolog<br>export_SdpB: antimicrobial peptide system protein, SdpB family | 9.10E-158<br>2.20E-76<br>3.50E-67<br>3.50E-04 |
| SDZ55834.1 | 63514 | 64075 | + | 186 | LinL | LinL | LinL | 8.20E-52 |
| SDZ55844.1 | 64071 | 64668 | + | 198 | TIGR00521<br>PF02441<br>TIGR02113 | TIGR00521<br>Flavoprotein<br>TIGR02113 | coaBC_dfp: phosphopantothenoylcysteine decarboxylase / phosphopantothenate--cysteine ligase<br>Flavoprotein<br>coaC_strep: phosphopantothenoylcysteine decarboxylase | 2.40E-09<br>5.00E-07<br>1.00E-04 |
| SDZ55854.1 | 64664 | 65294 | + | 209 | TIGR01166<br>TIGR02857<br>TIGR02673<br>TIGR03873<br>TIGR01189 | TIGR01166<br>TIGR02857<br>TIGR02673<br>TIGR03873<br>TIGR01189 | cbiO: cobalt ABC transporter, ATP-binding protein<br>CydD: thiol reductant ABC exporter, CydD subunit<br>FtsE: cell division ATP-binding protein FtsE<br>F420-0_ABC_ATP: proposed F420-0 ABC transporter, ATP-binding protein<br>ccmA: heme ABC exporter, ATP-binding protein CcmA | 3.70E-28<br>2.00E-27<br>4.10E-27<br>1.40E-26<br>2.10E-26 |
| SDZ55867.1 | 65290 | 66790 | + | 499 | NO PFAM MATCH | - | - | - |
| SDZ55878.1 | 67147 | 67366 | + | 72 | NO PFAM MATCH | - | - | - |
| SDZ55887.1 | 67362 | 67558 | + | 65 | NO PFAM MATCH | - | - | - |

| index | leader | core | start | end | dir | score |
|---|---|---|---|---|---|---|
| 1 | VAVPIKAPQGELLACIALAVRGPEPESALR | HVPTLLGAAHKLAHATSTGTVSAIR | 53168 | 53336 | + | 10 |
| 2 | VSLSLDADHGAIGQVYDERRGVEVSTTRDAELHLDAAVNLTH | ITADRAGVGTECTSRHTELVDAAQFPAGD | 56399 | 56615 | + | 6 |
| 3 | VCQVDRCVEMQFRVPGGGY | LDTSSFVVDLSNRTVIGIQ | 56525 | 56408 | - | 6 |
| 4 | VLASGLNAPPDTPSWSMLLSSPPATEVYSARTPAL | AVASARKTRKA | 56538 | 56679 | + | 6 |
| 5 | VRSAPDLLMNSCLMAQFARTARPVR | SYPARVSQFTKHHFRQITAVGTLAWA | 57145 | 57301 | + | 8 |
| 6 | VERLDHVLGDPAPLGEVLRVVGGAE...LRALPRDEHLTPPDVRRALCRGATA | MTTVPSTPPAPGDR | 57532 | 57730 | + | 10 |
| 7 | VVGGAELLRALPRDEHLTPPDVRRALCRGATA | MTTVPSTPPAPGDR | 57589 | 57730 | + | 6 |
| 8 | VIADTRGGCGARRSWGLVNS | VASCPSGKTTVPGEAVPMRC | 57719 | 57842 | + | 6 |
| 9 | MMGRESIKTPDLQLNSC | VARARATPALSNGECYAE | 61387 | 61495 | + | 6 |
| 10 | MGRESIKTPDLQLNSCV | ARARATPALSNGECYAE | 61390 | 61495 | + | 6 |
| 11 | MQSEVLEVQEFANQMMRHAEPGMRGDASNPAM | TTPAFVTAAQFVIQGSTICLIC | 61484 | 61649 | + | 32 |
| 12 | MNFIESVSPETIDSEQGHGGAVRRP | RIRGSSDAQKACRNPHLLKTGDGGF | 66954 | 67107 | + | 6 |
| 13 | MSVDDKYTYPGSGGVLVNHYNIRDAAKLDQAL | NAVASVRWAAMSREPLPERFNVAQLQAIHRRLF | 67362 | 67558 | + | 6 |

### Results for WP_051062157.1 [Ilumatobacter nonamiensis YM16-303] back to top

previous - next

#### Architecture

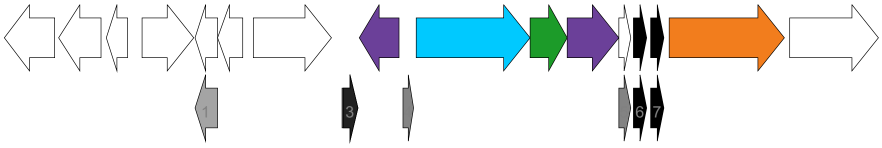

1000 nucleotides

Link to nucleotide sequence

| Accession | start | end | direction | length (aa) | Pfam/HMM | name | description | E-value |
|---|---|---|---|---|---|---|---|---|
| WP_051062152.1 | 9037 | 8206 | - | 276 | PF05721<br>TIGR02408<br>TIGR01762 | PhyH<br>TIGR02408<br>TIGR01762 | Phytanoyl-CoA dioxygenase (PhyH)<br>ectoine_ThpD: ectoine hydroxylase<br>chlorin-enz: chlorinating enzyme | 1.90E-32<br>1.80E-07<br>1.00E-06 |
| WP_051062153.1 | 9785 | 9086 | - | 232 | PF00578 | AhpC-TSA | AhpC/TSA family | 2.90E-04 |
| WP_051062154.1 | 10208 | 9863 | - | 114 | PF00578<br>PF08534 | AhpC-TSA<br>Redoxin | AhpC/TSA family<br>Redoxin | 3.00E-05<br>3.40E-04 |
| WP_083913940.1 | 10444 | 11293 | + | 282 | PF02668 | TauD | Taurine catabolism dioxygenase TauD, TfdA family | 1.80E-54 |
| WP_040490962.1 | 11674 | 11314 | - | 119 | PF02656 | DUF202 | Domain of unknown function (DUF202) | 5.60E-08 |
| WP_040490963.1 | 12072 | 11670 | - | 133 | PF02656 | DUF202 | Domain of unknown function (DUF202) | 6.30E-14 |
| WP_040490964.1 | 12251 | 13532 | + | 426 | NO PFAM MATCH | - | - | - |
| WP_051062156.1 | 14611 | 13969 | - | 213 | PF08241<br>PF13649<br>TIGR02072<br>TIGR01934<br>PF13489 | Methyltransf_11<br>Methyltransf_25<br>TIGR02072<br>TIGR01934<br>Methyltransf_23 | Methyltransferase domain<br>Methyltransferase domain<br>BioC: malonyl-acyl carrier protein O-methyltransferase BioC<br>MenG_MenH_UbiE: ubiquinone/menaquinone biosynthesis methyltransferase<br>Methyltransferase domain | 2.50E-11<br>1.70E-09<br>1.60E-05<br>4.10E-05<br>2.30E-04 |
| WP_051062157.1 | 14892 | 16743 | + | 616 | LinH<br>LinE<br>LinG<br>PF12697<br>TIGR02427 | LinH<br>LinE<br>LinG<br>Abhydrolase_6<br>TIGR02427 | LinH (Fusion of LinE/LinG)<br>LinE Homolog<br>LinG Homolog<br>Alpha/beta hydrolase family<br>protocat_pcaD: 3-oxoadipate enol-lactonase | 7.80E-210<br>5.00E-118<br>5.30E-91<br>1.50E-15<br>1.50E-10 |
| WP_051062158.1 | 16739 | 17351 | + | 203 | LinL | LinL | LinL | 4.90E-54 |
| WP_040490965.1 | 17347 | 18184 | + | 278 | PF13489<br>PF13649<br>TIGR02021<br>TIGR03534<br>TIGR00755 | Methyltransf_23<br>Methyltransf_25<br>TIGR02021<br>TIGR03534<br>TIGR00755 | Methyltransferase domain<br>Methyltransferase domain<br>BchM-ChlM: magnesium protoporphyrin O-methyltransferase<br>RF_mod_PrmC: protein-(glutamine-N5) methyltransferase, release factor-specific<br>ksgA: ribosomal RNA small subunit methyltransferase A | 2.30E-10<br>9.80E-09<br>2.20E-07<br>5.70E-05<br>7.30E-05 |
| WP_040490966.1 | 18180 | 18378 | + | 65 | NO PFAM MATCH | - | - | - |
| WP_040490967.1 | 18429 | 18642 | + | 70 | LinA | LinA | LinA (linaridin precursor) | 9.20E-15 |
| WP_040490968.1 | 18711 | 18921 | + | 69 | LinA | LinA | LinA (linaridin precursor) | 1.30E-17 |
| WP_148288453.1 | 19014 | 20883 | + | 622 | TIGR02203<br>TIGR02204<br>TIGR00958<br>TIGR01846<br>TIGR03796 | TIGR02203<br>TIGR02204<br>TIGR00958<br>TIGR01846<br>TIGR03796 | MsbA_lipidA: lipid A export permease/ATP-binding protein MsbA<br>MsbA_rel: ABC transporter, permease/ATP-binding protein<br>3a01208: antigen peptide transporter 2<br>type_I_sec_HlyB: type I secretion system ATPase<br>NHLM_micro_ABC1: NHLM bacteriocin system ABC transporter, peptidase/ATP-binding protein | 2.10E-89<br>7.80E-88<br>2.60E-84<br>8.70E-73<br>1.50E-72 |
| WP_040490969.1 | 20957 | 22397 | + | 479 | TIGR02734<br>TIGR03467<br>PF13450<br>PF01593<br>PF01266 | TIGR02734<br>TIGR03467<br>NAD_binding_8<br>Amino_oxidase<br>DAO | crtl_fam: phytoene desaturase<br>HpnE: squalene-associated FAD-dependent desaturase<br>NAD(P)-binding Rossmann-like domain<br>Flavin containing amine oxidoreductase<br>FAD dependent oxidoreductase | 3.30E-11<br>4.50E-10<br>1.70E-09<br>3.20E-08<br>6.00E-07 |

| index | leader | core | start | end | dir | score |
|---|---|---|---|---|---|---|
| 1 | MTSSGGSKEPRPTVVFDRMLQHERT...HYEHLHAPLRAGENPAHPRAARLVG | ITTIGFTGVATVVAVAALVT | 11674 | 11314 | - | 6 |
| 2 | LLGVLRPSPEGVLEGVDSVPQGLGH...CGAGGDDRVGRRDGRIPSGRVRPDG | STPVPRWPPTRAKWVATSVDRLAAGVKPPY | 13684 | 13957 | + | 8 |
| 3 | VLEGVDSVPQGLGHRRCRTRTGPRCGAGGDDRVGRRDGRIPSGRVRPDG | STPVPRWPPTRAKWVATSVDRLAAGVKPPY | 13717 | 13957 | + | 8 |
| 4 | MDHPFGRCVAEPASVMSRSEVDRPRCVHRRPPDPKVQR | AVPTSSPLPLAAGSFGVD | 14681 | 14852 | + | 7 |
| 5 | MKPTNLAIMIVGSVLVALVVFALLMVAFDRVLFAIGPAI | ATAFSTAAAIYLILRDSERPEQTSDT | 18180 | 18378 | + | 7 |
| 6 | MSTPTQPTPSDDLGSDSPLGDLDESTEESVMNEADPTM | ATPAAAAAVAAGVAATAAGFAAGYAAEEAADG | 18429 | 18642 | + | 29 |
| 7 | MNDETIPLADQANSDPRVDEIEMTAEEAEELDGTEPVT | ATPAAFAAGVAAAGAVAGAAAAGAAIGEAVD | 18711 | 18921 | + | 30 |

### Results for ARF74358.1 [Kitasatospora albolonga] back to top

previous - next

#### Architecture

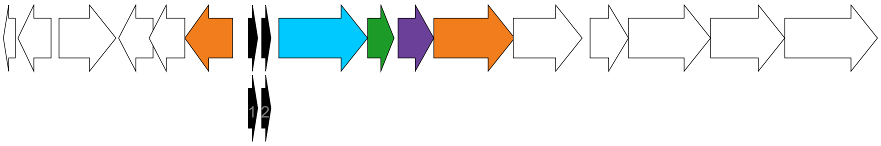

1000 nucleotides

Link to nucleotide sequence

| Accession | start | end | direction | length (aa) | Pfam/HMM | name | description | E-value |
|---|---|---|---|---|---|---|---|---|
| ARF74350.1 | 4669079 | 4668836 | - | 80 | PF01402 | RHH_1 | Ribbon-helix-helix protein, copG family | 6.40E-08 |
| | | | | | PF12651 | RHH_3 | Ribbon-helix-helix domain | 4.70E-07 |
| | | | | | PF07878 | RHH_5 | CopG-like RHH_1 or ribbon-helix-helix domain, RHH_5 | 8.30E-04 |
| ARF74351.1 | 4669851 | 4669137 | - | 237 | PF00293 | NUDIX | NUDIX domain | 5.40E-17 |
| | | | | | TIGR00586 | TIGR00586 | mutt: mutator mutT protein | 1.90E-10 |
| | | | | | TIGR02705 | TIGR02705 | nudix_YtkD: nucleoside triphosphatase YtkD | 6.40E-08 |
| ARF74352.1 | 4670017 | 4671235 | + | 405 | PF07730 | HisKA_3 | Histidine kinase | 1.50E-15 |
| ARF74353.1 | 4672017 | 4671303 | - | 237 | PF00072 | Response_reg | Response regulator receiver domain | 4.60E-24 |
| | | | | | TIGR02875 | TIGR02875 | spore_0_A: sporulation transcription factor Spo0A | 4.10E-15 |
| | | | | | PF00196 | GerE | Bacterial regulatory proteins, luxR family | 5.40E-12 |
| | | | | | TIGR01818 | TIGR01818 | ntrC: nitrogen regulation protein NR(I) | 9.10E-09 |
| | | | | | TIGR02956 | TIGR02956 | TMAO_torS: TMAO reductase sytem sensor TorS | 6.20E-08 |
| ARF74354.1 | 4672721 | 4671956 | - | 254 | PF01061 | ABC2_membrane | ABC-2 type transporter | 8.50E-14 |
| | | | | | PF12698 | ABC2_membrane_3 | ABC-2 family transporter protein | 1.30E-08 |
| | | | | | TIGR03062 | TIGR03062 | pip_yhgE_Cterm: YhgE/Pip C-terminal domain | 1.60E-07 |
| | | | | | TIGR01291 | TIGR01291 | nodJ: ABC-2 type transporter, NodJ family | 6.70E-04 |
| | | | | | TIGR01247 | TIGR01247 | drrB: daunorubicin resistance ABC transporter membrane protein | 9.40E-04 |
| ARF74355.1 | 4673728 | 4672717 | - | 336 | TIGR01188 | TIGR01188 | drrA: daunorubicin resistance ABC transporter, ATP-binding protein | 7.20E-63 |
| | | | | | TIGR01288 | TIGR01288 | nodI: nodulation ABC transporter NodI | 5.40E-44 |
| | | | | | TIGR03864 | TIGR03864 | PQQ_ABC_ATP: ABC transporter, ATP-binding subunit, PQQ-dependent alcohol dehydrogenase system | 1.60E-40 |
| | | | | | TIGR03740 | TIGR03740 | galliderm_ABC: lantibiotic protection ABC transporter, ATP-binding subunit | 1.10E-33 |
| | | | | | TIGR03522 | TIGR03522 | GldA_ABC_ATP: gliding motility-associated ABC transporter ATP-binding subunit GldA | 2.20E-33 |
| ARF74356.1 | 4674072 | 4674273 | + | 66 | LinA | LinA | LinA (linaridin precursor) | 4.30E-11 |
| ARF74357.1 | 4674348 | 4674558 | + | 69 | LinA | LinA | LinA (linaridin precursor) | 7.10E-08 |
| ARF74358.1 | 4674709 | 4676614 | + | 634 | LinH | LinH | LinH (Fusion of LinE/LinG) | 1.30E-245 |
| | | | | | LinG | LinG | LinG Homolog | 6.10E-119 |
| | | | | | LinE | LinE | LinE Homolog | 3.30E-98 |
| | | | | | PF12697 | Abhydrolase_6 | Alpha/beta hydrolase family | 2.30E-16 |
| | | | | | PF12146 | Hydrolase_4 | Serine aminopeptidase, S33 | 5.90E-09 |
| ARF74359.1 | 4676610 | 4677177 | + | 188 | LinL | LinL | LinL | 9.90E-66 |
| ARF77293.1 | 4677268 | 4678027 | + | 252 | PF13649 | Methyltransf_25 | Methyltransferase domain | 3.40E-16 |
| | | | | | PF08241 | Methyltransf_11 | Methyltransferase domain | 4.00E-12 |
| | | | | | PF13489 | Methyltransf_23 | Methyltransferase domain | 1.10E-09 |
| | | | | | PF13847 | Methyltransf_31 | Methyltransferase domain | 4.70E-09 |
| | | | | | PF08242 | Methyltransf_12 | Methyltransferase domain | 2.30E-08 |
| ARF74360.1 | 4678023 | 4679742 | + | 572 | TIGR02204 | TIGR02204 | MsbA_rel: ABC transporter, permease/ATP-binding protein | 7.90E-103 |
| | | | | | TIGR02203 | TIGR02203 | MsbA_lipidA: lipid A export permease/ATP-binding protein MsbA | 1.10E-99 |
| | | | | | TIGR00958 | TIGR00958 | 3a01208: antigen peptide transporter 2 | 2.70E-89 |
| | | | | | TIGR01846 | TIGR01846 | type_I_sec_HlyB: type I secretion system ATPase | 1.90E-87 |
| | | | | | TIGR03375 | TIGR03375 | type_I_sec_LssB: type I secretion system ATPase | 9.60E-87 |
| ARF74361.1 | 4679738 | 4681202 | + | 487 | PF13450 | NAD_binding_8 | NAD(P)-binding Rossmann-like domain | 9.80E-10 |
| | | | | | TIGR02734 | TIGR02734 | crtI_fam: phytoene desaturase | 1.70E-09 |
| | | | | | PF01593 | Amino_oxidase | Flavin containing amine oxidoreductase | 1.10E-08 |
| | | | | | TIGR03467 | TIGR03467 | HpnE: squalene-associated FAD-dependent desaturase | 1.30E-08 |
| | | | | | TIGR01317 | TIGR01317 | GOGAT_sm_gam: glutamate synthase, NADH/NADPH, small subunit | 4.90E-08 |
| ARF74362.1 | 4681376 | 4682189 | + | 270 | TIGR03084 | TIGR03084 | TIGR03084: TIGR03084 family protein | 9.20E-105 |
| | | | | | TIGR03083 | TIGR03083 | TIGR03083: uncharacterized Actinobacterial protein TIGR03083 | 8.30E-49 |
| | | | | | PF11716 | MDMPI_N | Mycothiol maleylpyruvate isomerase N-terminal domain | 2.70E-32 |
| | | | | | PF12867 | DinB_2 | DinB superfamily | 7.70E-10 |
| | | | | | PF08608 | Wyosine_form | Wyosine base formation | 7.40E-06 |
| ARF74363.1 | 4682185 | 4683931 | + | 581 | PF07287 | AtuA | Acyclic terpene utilisation family protein AtuA | 4.80E-120 |
| ARF74364.1 | 4683927 | 4685526 | + | 532 | PF01039 | Carboxyl_trans | Carboxyl transferase domain | 2.60E-134 |
| | | | | | TIGR01117 | TIGR01117 | mmdA: methylmalonyl-CoA decarboxylase alpha subunit | 5.60E-82 |
| | | | | | TIGR00513 | TIGR00513 | accA: acetyl-CoA carboxylase, carboxyl transferase, alpha subunit | 5.90E-11 |
| ARF74365.1 | 4685533 | 4687510 | + | 658 | TIGR02712 | TIGR02712 | urea_carbox: urea carboxylase | 2.90E-150 |
| | | | | | TIGR01235 | TIGR01235 | pyruv_carbox: pyruvate carboxylase | 5.30E-139 |
| | | | | | TIGR00514 | TIGR00514 | accC: acetyl-CoA carboxylase, biotin carboxylase subunit | 1.30E-135 |
| | | | | | PF02786 | CPSase_L_D2 | Carbamoyl-phosphate synthase L chain, ATP binding domain | 8.60E-60 |
| | | | | | PF00289 | Biotin_carb_N | Biotin carboxylase, N-terminal domain | 3.80E-42 |

| index | leader | core | start | end | dir | score |
|---|---|---|---|---|---|---|
| 1 | MSDIAAYFNARQDLGGIGSAHVPAADAPLL | ATPATITTIAGWKAVGAFAAGAGCVVGAYTAGKANG | 4674072 | 4674273 | + | 31 |
| 2 | MTNVITRLHATGQLPVTAIADTESGGLTAAAYPVL | GTPAFAVPFTAKAFSAGIAFGAATVAAYDAGAGK | 4674348 | 4674558 | + | 28 |

### Results for WP_129316254.1 [Kocuria koreensis] back to top

previous - next

#### Architecture

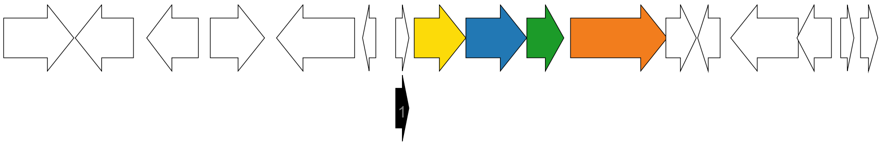

1000 nucleotides

Link to nucleotide sequence

| Accession | start | end | direction | length (aa) | Pfam/HMM | name | description | E-value |
|---|---|---|---|---|---|---|---|---|
| WP_129316246.1 | 17979 | 19065 | + | 361 | PF02317<br>PF02558<br>PF03807<br>PF01210<br>TIGR00745 | Octopine_DH<br>ApbA<br>F420_oxidored<br>NAD_Gly3P_dh_N<br>TIGR00745 | NAD/NADP octopine/nopaline dehydrogenase, alpha-helical domain<br>Ketopantoate reductase PanE/ApbA<br>NADP oxidoreductase coenzyme F420-dependent<br>NAD-dependent glycerol-3-phosphate dehydrogenase N-terminus<br>apbA_panE: 2-dehydropantoate 2-reductase | 2.90E-29<br>1.00E-09<br>1.70E-09<br>5.90E-09<br>2.60E-08 |
| WP_129316247.1 | 19986 | 19077 | - | 302 | PF01925 | TauE | Sulfite exporter TauE/SafE | 1.60E-33 |
| WP_129316248.1 | 20979 | 20190 | - | 262 | TIGR00099<br>PF08282<br>TIGR01484<br>TIGR01482<br>TIGR01487 | TIGR00099<br>Hydrolase_3<br>TIGR01484<br>TIGR01482<br>TIGR01487 | Cof-subfamily: Cof-like hydrolase<br>haloacid dehalogenase-like hydrolase<br>HAD-SF-IIB: HAD hydrolase, family IIB<br>SPP-subfamily: sucrose-phosphate phosphatase subfamily<br>Pglycolate_arch: phosphoglycolate phosphatase, TA0175-type | 1.20E-30<br>3.50E-27<br>1.10E-15<br>4.20E-11<br>2.70E-10 |
| WP_129316249.1 | 21156 | 21987 | + | 276 | PF13622 | 4HBT_3 | Thioesterase-like superfamily | 1.50E-29 |
| WP_129316250.1 | 23388 | 22170 | - | 405 | PF13450<br>TIGR02734<br>TIGR02730<br>TIGR03467<br>TIGR03315 | NAD_binding_8<br>TIGR02734<br>TIGR02730<br>TIGR03467<br>TIGR03315 | NAD(P)-binding Rossmann-like domain<br>crtI_fam: phytoene desaturase<br>carot_isom: carotene isomerase<br>HpnE: squalene-associated FAD-dependent desaturase<br>Se_ygfK: putative selenate reductase, YgfK subunit | 3.00E-10<br>3.90E-10<br>2.60E-08<br>1.40E-07<br>6.40E-07 |
| WP_129316251.1 | 23706 | 23514 | - | 63 | PF13567 | DUF4131 | Domain of unknown function (DUF4131) | 8.60E-04 |
| WP_129316252.1 | 24004 | 24226 | + | 73 | NO PFAM MATCH | - | - | - |
| WP_129316253.1 | 24290 | 25103 | + | 270 | LinG<br>LinH<br>TIGR04033 | LinG<br>LinH<br>TIGR04033 | LinG Homolog<br>LinH (Fusion of LinE/LinG)<br>export_SdpB: antimicrobial peptide system protein, SdpB family | 2.60E-66<br>3.60E-61<br>3.30E-04 |
| WP_129316254.1 | 25099 | 26038 | + | 312 | LinE<br>LinH<br>PF12697 | LinE<br>LinH<br>Abhydrolase_6 | LinE Homolog<br>LinH (Fusion of LinE/LinG)<br>Alpha/beta hydrolase family | 3.20E-40<br>3.00E-28<br>3.40E-09 |
| WP_156265487.1 | 26037 | 26610 | + | 190 | LinL | LinL | LinL | 8.10E-33 |
| WP_129316256.1 | 26699 | 28193 | + | 497 | TIGR02868<br>TIGR02857<br>TIGR02204<br>TIGR03608<br>TIGR01166 | TIGR02868<br>TIGR02857<br>TIGR02204<br>TIGR03608<br>TIGR01166 | CydC: thiol reductant ABC exporter, CydC subunit<br>CydD: thiol reductant ABC exporter, CydD subunit<br>MsbA_rel: ABC transporter, permease/ATP-binding protein<br>L_ocin_972_ABC: putative bacteriocin export ABC transporter, lactococcin 972 group<br>cbiO: cobalt ABC transporter, ATP-binding protein | 5.30E-19<br>2.90E-16<br>2.00E-14<br>3.70E-14<br>6.90E-13 |
| WP_129316257.1 | 28177 | 28648 | + | 156 | NO PFAM MATCH | - | - | - |
| WP_156265488.1 | 29014 | 28666 | - | 115 | PF08310 | LGFP | LGFP repeat | 5.00E-16 |
| WP_129315891.1 | 30217 | 29167 | - | 349 | PF00589<br>TIGR02224<br>TIGR02225<br>TIGR02249 | Phage_integrase<br>TIGR02224<br>TIGR02225<br>TIGR02249 | Phage integrase family<br>recomb_XerC: tyrosine recombinase XerC<br>recomb_XerD: tyrosine recombinase XerD<br>integrase_gron: integron integrase | 5.00E-21<br>6.50E-18<br>3.30E-14<br>5.50E-14 |
| WP_129315892.1 | 30722 | 30194 | - | 175 | PF01381 | HTH_3 | Helix-turn-helix | 3.90E-04 |
| WP_129315893.1 | 30858 | 31065 | + | 68 | PF12728 | HTH_17 | Helix-turn-helix domain | 5.70E-05 |
| WP_129315894.1 | 31166 | 31436 | + | 89 | NO PFAM MATCH | - | - | - |

| index | leader | core | start | end | dir | score |
|---|---|---|---|---|---|---|
| 1 | MHAVRHKSQDLDSAVASAFANVEQGVAPTQEGMPSM | STPTAVTVATTPEDVSAVAATVSAATSVAQLSVATQK | 24004 | 24226 | + | 31 |

### Results for WP_095798642.1 [Kocuria sp. WRN011] back to top

previous - next

#### Architecture

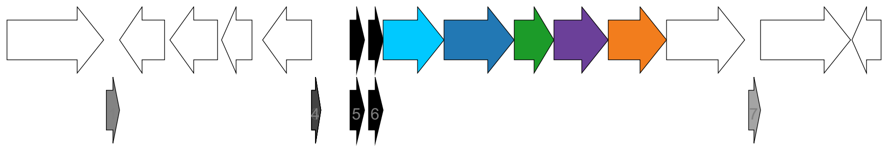

1000 nucleotides

Link to nucleotide sequence

| Accession | start | end | direction | length (aa) | Pfam/HMM | name | description | E-value |
|---|---|---|---|---|---|---|---|---|
| WP_095798634.1 | 84430 | 85834 | + | 467 | PF08310<br>PF10503<br>TIGR01840 | LGFP<br>Esterase_phd<br>TIGR01840 | LGFP repeat<br>Esterase PHB depolymerase<br>esterase_phb: esterase, PHB depolymerase family | 2.10E-23<br>3.00E-04<br>3.60E-04 |
| WP_095798636.1 | 86725 | 86059 | - | 221 | NO PFAM MATCH | - | - | - |
| WP_095798637.1 | 87501 | 86793 | - | 235 | PF05988<br>PF00578 | DUF899<br>AhpC-TSA | Bacterial protein of unknown function (DUF899)<br>AhpC/TSA family | 3.00E-61<br>2.30E-04 |

| | | | | | | | | |
|---|---|---|---|---|---|---|---|---|
| WP_095798638.1 | 87990 | 87555 | - | 144 | PF13302<br>PF00583 | Acetyltransf_3<br>Acetyltransf_1 | Acetyltransferase (GNAT) domain<br>Acetyltransferase (GNAT) family | 9.10E-13<br>7.90E-05 |
| WP_095798660.1 | 88861 | 88156 | - | 234 | NO PFAM MATCH | - | - | - |
| WP_095798639.1 | 89435 | 89642 | + | 68 | LinA | LinA | LinA (linaridin precursor) | 4.20E-09 |
| WP_095798640.1 | 89705 | 89909 | + | 67 | LinA | LinA | LinA (linaridin precursor) | 6.80E-10 |
| WP_143470006.1 | 89918 | 90797 | + | 292 | LinH<br>LinG<br>PF05090 | LinH<br>LinG<br>VKG_Carbox | LinH (Fusion of LinE/LinG)<br>LinG Homolog<br>Vitamin K-dependent gamma-carboxylase | 2.20E-88<br>9.60E-85<br>6.90E-04 |
| WP_095798642.1 | 90793 | 91825 | + | 343 | LinE<br>LinH<br>PF12697<br>PF12146<br>PF00561 | LinE<br>LinH<br>Abhydrolase_6<br>Hydrolase_4<br>Abhydrolase_1 | LinE Homolog<br>LinH (Fusion of LinE/LinG)<br>Alpha/beta hydrolase family<br>Serine aminopeptidase, S33<br>alpha/beta hydrolase fold | 7.70E-61<br>1.80E-54<br>7.60E-16<br>1.30E-14<br>1.70E-14 |
| WP_143470007.1 | 91821 | 92409 | + | 195 | LinL | LinL | LinL | 3.10E-56 |
| WP_143470008.1 | 92405 | 93203 | + | 265 | PF13489<br>PF13649<br>TIGR02021<br>PF08242<br>PF08241 | Methyltransf_23<br>Methyltransf_25<br>TIGR02021<br>Methyltransf_12<br>Methyltransf_11 | Methyltransferase domain<br>Methyltransferase domain<br>BchM-ChlM: magnesium protoporphyrin O-methyltransferase<br>Methyltransferase domain<br>Methyltransferase domain | 4.40E-10<br>3.90E-09<br>1.80E-07<br>4.00E-07<br>6.10E-07 |
| WP_095798644.1 | 93187 | 94042 | + | 284 | TIGR03522<br>TIGR01188<br>TIGR03740<br>TIGR04406<br>TIGR03864 | TIGR03522<br>TIGR01188<br>TIGR03740<br>TIGR04406<br>TIGR03864 | GldA_ABC_ATP: gliding motility-associated ABC transporter ATP-binding subunit GldA<br>drrA: daunorubicin resistance ABC transporter, ATP-binding protein<br>galliderm_ABC: lantibiotic protection ABC transporter, ATP-binding subunit<br>LPS_export_lptB: LPS export ABC transporter ATP-binding protein<br>PQQ_ABC_ATP: ABC transporter, ATP-binding subunit, PQQ-dependent alcohol dehydrogenase system | 2.20E-53<br>5.20E-48<br>5.60E-41<br>2.90E-40<br>3.10E-40 |
| WP_095798645.1 | 94038 | 95193 | + | 384 | PF12698 | ABC2_membrane_3 | ABC-2 family transporter protein | 1.30E-13 |
| WP_095798647.1 | 95415 | 96723 | + | 435 | TIGR02734<br>TIGR03467<br>PF13450<br>TIGR02730<br>TIGR01316 | TIGR02734<br>TIGR03467<br>NAD_binding_8<br>TIGR02730<br>TIGR01316 | crtI_fam: phytoene desaturase<br>HpnE: squalene-associated FAD-dependent desaturase<br>NAD(P)-binding Rossmann-like domain<br>carot_isom: carotene isomerase<br>gltA: glutamate synthase (NADPH), homotetrameric | 3.20E-15<br>1.40E-10<br>1.20E-08<br>5.70E-08<br>1.50E-06 |
| WP_143470009.1 | 97181 | 96755 | - | 141 | TIGR03083 | TIGR03083 | TIGR03083: uncharacterized Actinobacterial protein TIGR03083 | 2.20E-05 |

| index | leader | core | start | end | dir | score |
|---|---|---|---|---|---|---|
| 1 | VPALFRATSGPSHLTDWWDKRGLIIKIVTDFL | TNMLDRKFALQGVAFHFQVAGNCVVTRWDGGG | 85868 | 86063 | + | 7 |
| 2 | MGMMGPRNERCFFDPSSIAVL | FNVRTSFGWVQKNCPGNEKVSR | 88865 | 88997 | + | 6 |
| 3 | MMGPRNERCFFDPSSIAVLF | NVRTSFGWVQKNCPGNEKVSR | 88871 | 88997 | + | 6 |
| 4 | MGPRNERCFFDPSSIAVLFN | VRTSFGWVQKNCPGNEKVSR | 88874 | 88997 | + | 6 |
| 5 | MSVLQAVNVEPRLRGDIESGPLSYSDAPVL | ACPLAAVGAAGVAKAAGVVVGGGAVTGVAYVAYKTVAN | 89435 | 89642 | + | 29 |
| 6 | MQPTAKLLEHVNHTEISIHERQGVPVL | ATPAAFGAGVVLGAKVSAGGIAVAGGGAALCTATVLGCGN | 89705 | 89909 | + | 30 |
| 7 | MLLRTFVAIL | GTVAILFVFLGVLQSWDVSPVVRGMLAVGFSAAFAYIVFKKPSRAASQRR | 95236 | 95419 | + | 6 |

### Results for RYJ54945.1 [Kroppenstedtia sanguinis] back to top

previous - next

#### Architecture

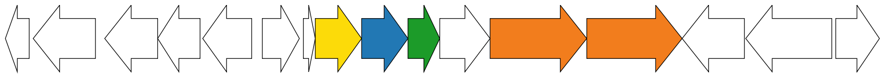

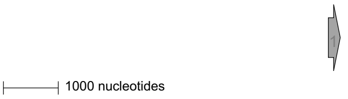

[Link to nucleotide sequence]

| Accession | start | end | direction | length (aa) | Pfam/HMM | name | description | E-value |
|---|---|---|---|---|---|---|---|---|
| RYJ54937.1 | 90517 | 90079 | - | 145 | PF13577<br>PF12680<br>TIGR02246<br>PF13474 | SnoaL_4<br>SnoaL_2<br>TIGR02246<br>SnoaL_3 | SnoaL-like domain<br>SnoaL-like domain<br>TIGR02246: conserved hypothetical protein<br>SnoaL-like domain | 6.10E-24<br>2.80E-06<br>7.40E-06<br>2.50E-05 |
| RYJ54938.1 | 91724 | 90575 | - | 382 | PF02515<br>TIGR03253<br>TIGR04253 | CoA_transf_3<br>TIGR03253<br>TIGR04253 | CoA-transferase family III<br>oxalate_frc: formyl-CoA transferase<br>mesacon_CoA_iso: mesaconyl-CoA isomerase | 8.50E-86<br>7.20E-37<br>2.30E-07 |
| RYJ54939.1 | 92848 | 91891 | - | 318 | PF03466<br>TIGR02424<br>TIGR03339<br>PF00126<br>TIGR03298 | LysR_substrate<br>TIGR02424<br>TIGR03339<br>HTH_1<br>TIGR03298 | LysR substrate binding domain<br>TF_pcaQ: pca operon transcription factor PcaQ<br>phn_lysR: aminoethylphosphonate catabolism associated LysR family transcriptional regulator<br>Bacterial regulatory helix-turn-helix protein, lysR family<br>argP: transcriptional regulator, ArgP family | 3.30E-32<br>3.30E-19<br>1.30E-16<br>7.70E-12<br>2.00E-11 |
| RYJ54940.1 | 93632 | 92858 | - | 257 | PF00378<br>TIGR02280<br>PF16113<br>TIGR01929<br>TIGR03210 | ECH_1<br>TIGR02280<br>ECH_2<br>TIGR01929<br>TIGR03210 | Enoyl-CoA hydratase/isomerase<br>PaaB1: phenylacetate degradation probable enoyl-CoA hydratase PaaB<br>Enoyl-CoA hydratase/isomerase<br>menB: naphthoate synthase<br>badI: 2-ketocyclohexanecarboxyl-CoA hydrolase | 1.50E-53<br>6.50E-50<br>2.60E-33<br>3.10E-32<br>9.60E-26 |
| RYJ54941.1 | 94569 | 93663 | - | 301 | TIGR01289<br>PF00106<br>TIGR04316<br>PF13561<br>TIGR01963 | TIGR01289<br>adh_short<br>TIGR04316<br>adh_short_C2<br>TIGR01963 | LPOR: light-dependent protochlorophyllide reductase<br>short chain dehydrogenase<br>dhbA_paeA: 2,3-dihydro-2,3-dihydroxybenzoate dehydrogenase<br>Enoyl-(Acyl carrier protein) reductase<br>PHB_DH: 3-hydroxybutyrate dehydrogenase | 4.30E-26<br>1.10E-20<br>8.80E-12<br>8.00E-11<br>1.00E-10 |
| RYJ54942.1 | 94774 | 95431 | + | 218 | PF09203 | MspA | MspA | 5.60E-33 |
| RYJ54943.1 | 95505 | 95718 | + | 70 | NO PFAM MATCH | - | - | - |
| RYJ54944.1 | 95723 | 96569 | + | 281 | LinG<br>LinH<br>TIGR04033<br>PF05090 | LinG<br>LinH<br>TIGR04033<br>VKG_Carbox | LinG Homolog<br>LinH (Fusion of LinE/LinG)<br>export_SdpB: antimicrobial peptide system protein, SdpB family<br>Vitamin K-dependent gamma-carboxylase | 3.60E-50<br>4.80E-45<br>7.50E-07<br>3.10E-04 |
| RYJ54945.1 | 96565 | 97420 | + | 284 | LinE<br>LinH<br>PF12697 | LinE<br>LinH<br>Abhydrolase_6 | LinE Homolog<br>LinH (Fusion of LinE/LinG)<br>Alpha/beta hydrolase family | 1.50E-26<br>2.00E-21<br>3.00E-06 |
| RYJ54946.1 | 97416 | 98010 | + | 197 | LinL | LinL | LinL | 7.50E-28 |
| RYJ54947.1 | 98006 | 98918 | + | 303 | PF01551 | Peptidase_M23 | Peptidase family M23 | 5.90E-16 |
| RYJ54948.1 | 98914 | 100690 | + | 591 | TIGR03796<br>TIGR02868<br>TIGR02203<br>TIGR03375<br>TIGR02857 | TIGR03796<br>TIGR02868<br>TIGR02203<br>TIGR03375<br>TIGR02857 | NHLM_micro_ABC1: NHLM bacteriocin system ABC transporter, peptidase/ATP-binding protein<br>CydC: thiol reductant ABC exporter, CydC subunit<br>MsbA_lipidA: lipid A export permease/ATP-binding protein MsbA<br>type_I_sec_LssB: type I secretion system ATPase<br>CydD: thiol reductant ABC exporter, CydD subunit | 3.90E-44<br>1.80E-43<br>2.60E-43<br>5.90E-42<br>6.70E-42 |
| RYJ54949.1 | 100686 | 102411 | + | 574 | TIGR02857<br>TIGR03796<br>TIGR03375<br>TIGR02868<br>TIGR02204 | TIGR02857<br>TIGR03796<br>TIGR03375<br>TIGR02868<br>TIGR02204 | CydD: thiol reductant ABC exporter, CydD subunit<br>NHLM_micro_ABC1: NHLM bacteriocin system ABC transporter, peptidase/ATP-binding protein<br>type_I_sec_LssB: type I secretion system ATPase<br>CydC: thiol reductant ABC exporter, CydC subunit<br>MsbA_rel: ABC transporter, permease/ATP-binding protein | 1.20E-65<br>6.30E-65<br>2.50E-63<br>3.30E-61<br>1.50E-58 |
| RYJ54950.1 | 103550 | 102416 | - | 377 | PF03060<br>TIGR03151<br>PF01070<br>TIGR03966<br>PF00478 | NMO<br>TIGR03151<br>FMN_dh<br>TIGR03966<br>IMPDH | Nitronate monooxygenase<br>enACPred_II: putative enoyl-[acyl-carrier-protein] reductase II<br>FMN-dependent dehydrogenase<br>actino_HemFlav: heme/flavin dehydrogenase, mycofactocin system<br>IMP dehydrogenase / GMP reductase domain | 2.00E-47<br>4.10E-44<br>1.70E-09<br>1.20E-08<br>2.70E-05 |
| RYJ54951.1 | 105160 | 103582 | - | 525 | PF00501<br>TIGR03098<br>TIGR01733<br>TIGR02275<br>TIGR03205 | AMP-binding<br>TIGR03098<br>TIGR01733<br>TIGR02275<br>TIGR03205 | AMP-binding enzyme<br>ligase_PEP_1: acyl-CoA ligase (AMP-forming), exosortase A-associated<br>AA-adenyl-dom: amino acid adenylation domain<br>DHB_AMP_lig: (2,3-dihydroxybenzoyl)adenylate synthase<br>pimA: dicarboxylate--CoA ligase PimA | 7.20E-61<br>1.90E-47<br>5.40E-47<br>5.40E-39<br>6.80E-36 |
| RYJ55056.1 | 105221 | 106028 | + | 268 | PF00378<br>TIGR02280<br>PF16113<br>TIGR01929<br>TIGR02441 | ECH_1<br>TIGR02280<br>ECH_2<br>TIGR01929<br>TIGR02441 | Enoyl-CoA hydratase/isomerase<br>PaaB1: phenylacetate degradation probable enoyl-CoA hydratase PaaB<br>Enoyl-CoA hydratase/isomerase<br>menB: naphthoate synthase<br>fa_ox_alpha_mit: fatty acid oxidation complex, alpha subunit, mitochondrial | 3.40E-56<br>1.50E-38<br>1.80E-29<br>2.70E-27<br>1.20E-25 |

| index | leader | core | start | end | dir | score |
|---|---|---|---|---|---|---|
| 1 | MKMWEIEDQLNGRGHVTAASVGTGPLAAAVPQ | RTAMMSSLAQSSDVMTSLASVSASMASIYALANSGHRP | 95505 | 95718 | + | 6 |

### Results for ERK70867.1 [Leifsonia aquatica ATCC 14665] back to top

previous - next

#### Architecture

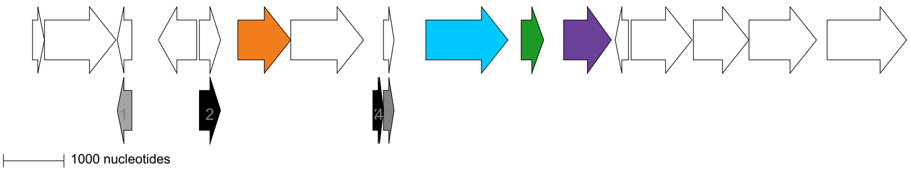

Link to nucleotide sequence

| Accession | start | end | direction | length (aa) | Pfam/HMM | name | description | E-value |
|---|---|---|---|---|---|---|---|---|
| ERK70859.1 | 0 | 185 | + | 60 | NO PFAM MATCH | - | - | - |
| ERK70860.1 | 181 | 1369 | + | 395 | PF01266<br>TIGR02352<br>TIGR03197<br>TIGR01373<br>PF07992 | DAO<br>TIGR02352<br>TIGR03197<br>TIGR01373<br>Pyr_redox_2 | FAD dependent oxidoreductase<br>thiamin_ThiO: glycine oxidase ThiO<br>MnmC_Cterm: tRNA U-34 5-methylaminomethyl-2-thiouridine biosynthesis protein MnmC, C-terminal domain<br>soxB: sarcosine oxidase, beta subunit family<br>Pyridine nucleotide-disulphide oxidoreductase | 1.90E-39<br>5.10E-22<br>2.10E-12<br>6.30E-08<br>3.80E-07 |
| ERK70861.1 | 1624 | 1393 | - | 76 | NO PFAM MATCH | - | - | - |
| ERK70862.1 | 2709 | 2073 | - | 211 | NO PFAM MATCH | - | - | - |
| ERK70863.1 | 2754 | 3096 | + | 113 | NO PFAM MATCH | - | - | - |
| ERK70864.1 | 3381 | 4260 | + | 292 | TIGR01188<br>TIGR01288<br>TIGR03522<br>TIGR03864<br>TIGR01186 | TIGR01188<br>TIGR01288<br>TIGR03522<br>TIGR03864<br>TIGR01186 | drrA: daunorubicin resistance ABC transporter, ATP-binding protein<br>nodI: nodulation ABC transporter NodI<br>GldA_ABC_ATP: gliding motility-associated ABC transporter ATP-binding subunit GldA<br>PQQ_ABC_ATP: ABC transporter, ATP-binding subunit, PQQ-dependent alcohol dehydrogenase system<br>proV: glycine betaine/L-proline transport ATP binding subunit | 5.00E-53<br>2.80E-42<br>2.40E-41<br>1.90E-39<br>4.50E-37 |
| ERK70865.1 | 4256 | 5453 | + | 398 | PF12698 | ABC2_membrane_3 | ABC-2 family transporter protein | 4.80E-28 |
| ERK70866.1 | 5789 | 5972 | + | 60 | NO PFAM MATCH | - | - | - |
| ERK70867.1 | 6485 | 7844 | + | 452 | LinH<br>LinE<br>LinG<br>PF12146<br>PF00561 | LinH<br>LinE<br>LinG<br>Hydrolase_4<br>Abhydrolase_1 | LinH (Fusion of LinE/LinG)<br>LinE Homolog<br>LinG Homolog<br>Serine aminopeptidase, S33<br>alpha/beta hydrolase fold | 3.80E-113<br>2.60E-59<br>1.70E-49<br>5.90E-15<br>3.10E-13 |
| ERK70868.1 | 8058 | 8439 | + | 126 | LinL | LinL | LinL | 7.60E-26 |
| ERK70869.1 | 8749 | 9553 | + | 267 | PF13649<br>PF08242<br>PF13847<br>PF08241<br>PF13489 | Methyltransf_25<br>Methyltransf_12<br>Methyltransf_31<br>Methyltransf_11<br>Methyltransf_23 | Methyltransferase domain<br>Methyltransferase domain<br>Methyltransferase domain<br>Methyltransferase domain<br>Methyltransferase domain | 7.50E-13<br>1.30E-11<br>1.90E-10<br>8.60E-09<br>3.10E-08 |
| ERK70870.1 | 9839 | 9623 | - | 71 | NO PFAM MATCH | - | - | - |
| ERK70871.1 | 9881 | 10883 | + | 333 | TIGR01135<br>PF01380<br>TIGR03127<br>TIGR00393<br>TIGR02128 | TIGR01135<br>SIS<br>TIGR03127<br>TIGR00393<br>TIGR02128 | glmS: glutamine-fructose-6-phosphate transaminase (isomerizing)<br>SIS domain<br>RuMP_HxlB: 6-phospho 3-hexuloisomerase<br>kpsF: sugar isomerase, KpsF/GutQ family<br>G6PI_arch: bifunctional phosphoglucose/phosphomannose isomerase | 1.70E-17<br>2.20E-15<br>1.10E-05<br>3.30E-05<br>1.00E-04 |
| ERK70872.1 | 10893 | 11817 | + | 307 | PF00294<br>TIGR04382<br>TIGR03168<br>TIGR03828<br>TIGR02152 | PfkB<br>TIGR04382<br>TIGR03168<br>TIGR03828<br>TIGR02152 | pfkB family carbohydrate kinase<br>myo_inos_iolC_N: 5-dehydro-2-deoxygluconokinase<br>1-PFK: hexose kinase, 1-phosphofructokinase family<br>pfkB: 1-phosphofructokinase<br>D_ribokin_bact: ribokinase | 1.10E-26<br>6.60E-15<br>9.90E-14<br>3.60E-13<br>4.30E-10 |
| ERK70873.1 | 11813 | 12944 | + | 376 | TIGR01481<br>TIGR02417<br>PF13377<br>PF00532<br>TIGR02405 | TIGR01481<br>TIGR02417<br>Peripla_BP_3<br>Peripla_BP_1<br>TIGR02405 | ccpA: catabolite control protein A<br>fruct_sucro_rep: D-fructose-responsive transcription factor<br>Periplasmic binding protein-like domain<br>Periplasmic binding proteins and sugar binding domain of LacI family<br>trehalos_R_Ecol: trehalose operon repressor | 3.80E-52<br>1.10E-40<br>5.80E-33<br>5.80E-27<br>3.10E-19 |
| ERK70874.1 | 13102 | 14431 | + | 442 | PF01547<br>TIGR03851<br>TIGR03850 | SBP_bac_1<br>TIGR03851<br>TIGR03850 | Bacterial extracellular solute-binding protein<br>chitin_NgcE: carbohydrate ABC transporter, N-acetylglucosamine/diacetylchitobiose-binding protein<br>bind_CPR_0540: carbohydrate ABC transporter substrate-binding protein, CPR_0540 family | 9.10E-25<br>5.80E-16<br>1.20E-13 |

| index | leader | core | start | end | dir | score |
|---|---|---|---|---|---|---|
| 1 | MRITAGTCACPLGRRALEDLETLIP...LYESFYPGDELTARTAAIIQAVLDQ | GAPKPTAPARPDLG | 1624 | 1393 | - | 6 |
| 2 | MTEQILDRIEEFAPGARDVILSITA...GGDILAGTTRGLAFARRPTLHPAPW | RTPVRGVYHCSSAVAPGPGVHGMVGHLAALDALRTHFNLAAPALGSTRT | 2754 | 3096 | + | 16 |
| 3 | MSNVLGYVNANAKSTAGGSRVQVMPSESAPVL | ACPVVAFAAIGFGMYTVFNKCFG | 5609 | 5777 | + | 17 |
| 4 | MPSESAPVLACPVVAF | AAIGFGMYTVFNKCFG | 5678 | 5777 | + | 14 |
| 5 | MLLQAIEDQTSRAGVSLERMTITRKPDAAVL | ATPTLAMAFGALQGGWFVGEMVKNVIRGG | 5789 | 5972 | + | 7 |

### Results for WP_155829033.1 [Leifsonia aquatica ATCC 14665] back to top

previous - next

#### Architecture

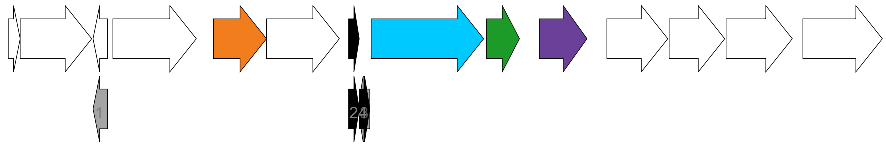

1000 nucleotides

Link to nucleotide sequence

| Accession | start | end | direction | length (aa) | Pfam/HMM | name | description | E-value |
|---|---|---|---|---|---|---|---|---|
| WP_020075996.1 | 0 | 185 | + | 61 | INFERRED GENE | - | - | - |
| WP_021763227.1 | 181 | 1369 | + | 395 | PF01266<br>TIGR02352<br>TIGR03197<br>TIGR01373<br>PF07992 | DAO<br>TIGR02352<br>TIGR03197<br>TIGR01373<br>Pyr_redox_2 | FAD dependent oxidoreductase<br>thiamin_ThiO: glycine oxidase ThiO<br>MnmC_Cterm: tRNA U-34 5-methylaminomethyl-2-thiouridine biosynthesis protein MnmC, C-terminal domain<br>soxB: sarcosine oxidase, beta subunit family<br>Pyridine nucleotide-disulphide oxidoreductase | 1.90E-39<br>5.10E-22<br>2.10E-12<br>6.30E-08<br>3.80E-07 |
| WP_021763228.1 | 1624 | 1393 | - | 76 | NO PFAM MATCH | - | - | - |
| WP_169730337.1 | 1710 | 3096 | + | 461 | PF01593<br>TIGR03467 | Amino_oxidase<br>TIGR03467 | Flavin containing amine oxidoreductase<br>HpnE: squalene-associated FAD-dependent desaturase | 3.00E-06<br>5.70E-04 |
| WP_021763231.1 | 3381 | 4260 | + | 292 | TIGR01188<br>TIGR01288<br>TIGR03522<br>TIGR03864<br>TIGR01186 | TIGR01188<br>TIGR01288<br>TIGR03522<br>TIGR03864<br>TIGR01186 | drrA: daunorubicin resistance ABC transporter, ATP-binding protein<br>nodI: nodulation ABC transporter NodI<br>GldA_ABC_ATP: gliding motility-associated ABC transporter ATP-binding subunit GldA<br>PQQ_ABC_ATP: ABC transporter, ATP-binding subunit, PQQ-dependent alcohol dehydrogenase system<br>proV: glycine betaine/L-proline transport ATP binding subunit | 5.00E-53<br>2.80E-42<br>2.40E-41<br>1.90E-39<br>4.50E-37 |
| WP_021763232.1 | 4256 | 5453 | + | 398 | PF12698 | ABC2_membrane_3 | ABC-2 family transporter protein | 4.80E-28 |
| WP_155829032.1 | 5609 | 5777 | + | 55 | LinA | LinA | LinA (linaridin precursor) | 2.90E-08 |
| WP_155829033.1 | 5990 | 7844 | + | 617 | LinH<br>LinG<br>LinE<br>PF12146<br>PF00561 | LinH<br>LinG<br>LinE<br>Hydrolase_4<br>Abhydrolase_1 | LinH (Fusion of LinE/LinG)<br>LinG Homolog<br>LinE Homolog<br>Serine aminopeptidase, S33<br>alpha/beta hydrolase fold | 4.40E-143<br>2.10E-76<br>5.50E-59<br>1.20E-14<br>4.20E-13 |
| WP_155829034.1 | 7884 | 8439 | + | 184 | LinL | LinL | LinL | 2.10E-44 |
| WP_021763236.1 | 8749 | 9553 | + | 267 | PF13649<br>PF08242<br>PF13847<br>PF08241<br>PF13489 | Methyltransf_25<br>Methyltransf_12<br>Methyltransf_31<br>Methyltransf_11<br>Methyltransf_23 | Methyltransferase domain<br>Methyltransferase domain<br>Methyltransferase domain<br>Methyltransferase domain<br>Methyltransferase domain | 7.50E-13<br>1.30E-11<br>1.90E-10<br>8.60E-09<br>3.10E-08 |
| WP_021763238.1 | 9881 | 10883 | + | 333 | TIGR01135<br>PF01380<br>TIGR03127<br>TIGR00393<br>TIGR02128 | TIGR01135<br>SIS<br>TIGR03127<br>TIGR00393<br>TIGR02128 | glmS: glutamine-fructose-6-phosphate transaminase (isomerizing)<br>SIS domain<br>RuMP_HxlB: 6-phospho 3-hexuloisomerase<br>kpsF: sugar isomerase, KpsF/GutQ family<br>G6PI_arch: bifunctional phosphoglucose/phosphomannose isomerase | 1.70E-17<br>2.20E-15<br>1.10E-05<br>3.30E-05<br>1.00E-04 |
| WP_039922394.1 | 10902 | 11817 | + | 304 | PF00294<br>TIGR04382<br>TIGR03168<br>TIGR03828<br>TIGR02152 | PfkB<br>TIGR04382<br>TIGR03168<br>TIGR03828<br>TIGR02152 | pfkB family carbohydrate kinase<br>myo_inos_iolC_N: 5-dehydro-2-deoxygluconokinase<br>1-PFK: hexose kinase, 1-phosphofructokinase family<br>pfkB: 1-phosphofructokinase<br>D_ribokin_bact: ribokinase | 1.10E-26<br>6.60E-15<br>9.70E-14<br>3.50E-13<br>4.20E-10 |
| WP_039922389.1 | 11852 | 12944 | + | 363 | TIGR01481<br>TIGR02417<br>PF13377<br>PF00532<br>TIGR02405 | TIGR01481<br>TIGR02417<br>Peripla_BP_3<br>Peripla_BP_1<br>TIGR02405 | ccpA: catabolite control protein A<br>fruct_sucro_rep: D-fructose-responsive transcription factor<br>Periplasmic binding protein-like domain<br>Periplasmic binding proteins and sugar binding domain of LacI family<br>trehalos_R_Ecol: trehalose operon repressor | 3.20E-52<br>9.50E-41<br>5.10E-33<br>5.20E-27<br>2.80E-19 |
| WP_021763241.1 | 13102 | 14431 | + | 442 | PF01547<br>TIGR03851<br>TIGR03850 | SBP_bac_1<br>TIGR03851<br>TIGR03850 | Bacterial extracellular solute-binding protein<br>chitin_NgcE: carbohydrate ABC transporter, N-acetylglucosamine/diacetylchitobiose-binding protein<br>bind_CPR_0540: carbohydrate ABC transporter substrate-binding protein, CPR_0540 family | 9.10E-25<br>5.80E-16<br>1.20E-13 |

| index | leader | core | start | end | dir | score |
|---|---|---|---|---|---|---|
| 1 | MRITAGTCACPLGRRALEDLETLIP...LYESFYPGDELTARTAAIIQAVLDQ | GAPKPTAPARPDLG | 1624 | 1393 | - | 6 |
| 2 | MSNVLGYVNANAKSTAGGSRVQVMPSESAPVL | ACPVVAFAAIGFGMYTVFNKCFG | 5609 | 5777 | + | 17 |
| 3 | MLLQAIEDQTSRAGVSLERMTITRKPDAAVL | ATPTLAMAFGALQGGWFVGEMVKNVIRGG | 5789 | 5972 | + | 17 |
| 4 | MTFLTISPTN | HPPWRAPKAMARVGVARTAASGFRV...VIRSRETPARDVWSSIACSSMSVVS | 5959 | 5773 | - | 6 |

### Results for REH27010.1 [Kutzneria buriramensis] back to top

previous - next

#### Architecture

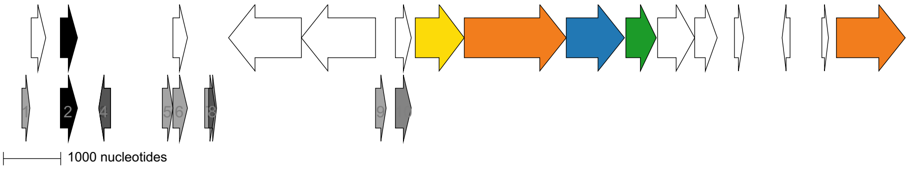

Link to nucleotide sequence

| Accession | start | end | direction | length (aa) | Pfam/HMM | name | description | E-value |
|---|---|---|---|---|---|---|---|---|
| REH27002.1 | 59976 | 60225 | + | 82 | PF14690 | zf-ISL3 | zinc-finger of transposase IS204/IS1001/IS1096/IS1165 | 1.60E-06 |
| REH27003.1 | 60470 | 60773 | + | 100 | LinA | LinA | LinA (linaridin precursor) | 2.70E-05 |

| | | | | | | | | |
|---|---|---|---|---|---|---|---|---|
| REH27004.1 | 62432 | 62699 | + | 88 | NO PFAM MATCH | - | - | - |
| REH27005.1 | 64689 | 63417 | - | 423 | PF05193 | Peptidase_M16_C | Peptidase M16 inactive domain | 3.80E-22 |
| REH27006.1 | 65987 | 64685 | - | 433 | PF00675<br>PF05193<br>TIGR02110 | Peptidase_M16<br>Peptidase_M16_C<br>TIGR02110 | Insulinase (Peptidase family M16)<br>Peptidase M16 inactive domain<br>PQQ_syn_pqqF: coenzyme PQQ biosynthesis protein PqqF | 1.10E-21<br>5.70E-21<br>3.20E-20 |
| REH27007.1 | 66317 | 66593 | + | 91 | NO PFAM MATCH | - | - | - |
| REH27008.1 | 66675 | 67518 | + | 280 | LinG<br>LinH | LinG<br>LinH | LinG Homolog<br>LinH (Fusion of LinE/LinG) | 1.80E-96<br>2.10E-92 |
| REH27009.1 | 67521 | 69297 | + | 591 | TIGR02204<br>TIGR02203<br>TIGR00958<br>TIGR01846<br>TIGR03796 | TIGR02204<br>TIGR02203<br>TIGR00958<br>TIGR01846<br>TIGR03796 | MsbA_rel: ABC transporter, permease/ATP-binding protein<br>MsbA_lipidA: lipid A export permease/ATP-binding protein MsbA<br>3a01208: antigen peptide transporter 2<br>type_I_sec_HlyB: type I secretion system ATPase<br>NHLM_micro_ABC1: NHLM bacteriocin system ABC transporter, peptidase/ATP-binding protein | 1.30E-98<br>5.00E-94<br>3.30E-91<br>2.30E-85<br>2.90E-83 |
| REH27010.1 | 69293 | 70322 | + | 342 | LinE<br>LinH<br>PF12697<br>TIGR02427<br>PF00561 | LinE<br>LinH<br>Abhydrolase_6<br>TIGR02427<br>Abhydrolase_1 | LinE Homolog<br>LinH (Fusion of LinE/LinG)<br>Alpha/beta hydrolase family<br>protocat_pcaD: 3-oxoadipate enol-lactonase<br>alpha/beta hydrolase fold | 1.10E-96<br>1.10E-69<br>9.10E-13<br>1.40E-11<br>9.30E-11 |
| REH27011.1 | 70333 | 70876 | + | 180 | LinL | LinL | LinL | 4.10E-69 |
| REH27012.1 | 70887 | 71541 | + | 217 | PF00196<br>TIGR03020<br>TIGR03541<br>PF08281<br>TIGR02937 | GerE<br>TIGR03020<br>TIGR03541<br>Sigma70_r4_2<br>TIGR02937 | Bacterial regulatory proteins, luxR family<br>EpsA: transcriptional regulator EpsA<br>reg_near_HchA: LuxR family transcriptional regulatory, chaperone HchA-associated<br>Sigma-70, region 4<br>sigma70-ECF: RNA polymerase sigma factor, sigma-70 family | 9.70E-18<br>8.00E-14<br>9.50E-09<br>3.20E-07<br>7.20E-06 |
| REH27013.1 | 71552 | 71939 | + | 128 | PF04134 | DUF393 | Protein of unknown function, DUF393 | 9.80E-15 |
| REH27014.1 | 72232 | 72385 | + | 50 | PF01609 | DDE_Tnp_1 | Transposase DDE domain | 3.70E-05 |
| REH27015.1 | 73211 | 73073 | - | 45 | TIGR01773<br>PF00324 | TIGR01773<br>AA_permease | GABAperm: GABA permease<br>Amino acid permease | 4.40E-06<br>6.00E-06 |
| REH27016.1 | 73760 | 73880 | + | 39 | NO PFAM MATCH | - | - | - |
| REH27017.1 | 74017 | 75217 | + | 399 | TIGR02633<br>TIGR01188<br>TIGR03410<br>TIGR00968<br>TIGR04406 | TIGR02633<br>TIGR01188<br>TIGR03410<br>TIGR00968<br>TIGR04406 | xylG: D-xylose ABC transporter, ATP-binding protein<br>drrA: daunorubicin resistance ABC transporter, ATP-binding protein<br>urea_trans_UrtE: urea ABC transporter, ATP-binding protein UrtE<br>3a0106s01: sulfate ABC transporter, ATP-binding protein<br>LPS_export_lptB: LPS export ABC transporter ATP-binding protein | 1.40E-31<br>1.10E-22<br>1.10E-21<br>1.80E-21<br>3.70E-19 |

| index | leader | core | start | end | dir | score |
|---|---|---|---|---|---|---|
| 1 | VPTIRVFELDRGLMTTRWQSGR | QTPVWALGEAVALLVIAIVYAG | 59819 | 59954 | + | 6 |
| 2 | MNHVEAAVAHAGVSLDSVLGPFEVSSPVA | CTPAAAWRAVTALVTAARVVGEREY...EMAHRTAPAGSLSLEALLDAVVQYG | 60470 | 60773 | + | 17 |
| 3 | LSRRFGFSAVQKIDCLAGVDVEQYGAVHVSLA | GSASGQALAVQWGAEAPTRTSAVTVPTKRRRRR | 61352 | 61154 | - | 6 |
| 4 | LQPLSRRFGFSAVQKIDCLAGVDVEQYGAVHVSL | AGSASGQALAVQWGAEAPTRTSAVTVPTKRRRRR | 61361 | 61154 | - | 7 |
| 5 | VRVVHPPIGGYLADFSQAASEFSLLGV | STARVDGFDVYDRGVHYLWMDRRLPGSLL | 62265 | 62436 | + | 6 |
| 6 | MNIVERAVAASDVLSPLTVANPGE | PTACLVEGLEGIARYAGEVVNETAQ...RSGSASGDGSVLSVVQLTEALVAAV | 62432 | 62699 | + | 6 |
| 7 | LRPVTAQSGSRHSVRRWRFDALWRSVGSVI | APLELGSLFACTRAAVWRAVIRLATAERVG | 62993 | 63176 | + | 6 |
| 8 | MEICGFGDRPARTGFALRVYPGGRL | EGGDQAGDRRARWVTRVRLRQRARAP | 63058 | 63214 | + | 7 |
| 9 | MVTVESVGATESGRLTDPVTASVAQPEGS | LTRFGSMGPTSAHVSKKTSFYCGIPSPSY | 65985 | 66162 | + | 6 |
| 10 | MNRIELIAQDCTADIAALDLAGLAV...YVQCKNHGCVLNAPEDYQVDRDLAG | LTVGDMVSMRAEVASLR | 66317 | 66593 | + | 7 |

### Results for AGW40770.1 [Leifsonia xyli subsp. cynodontis DSM 46306] back to top

previous - next

#### Architecture

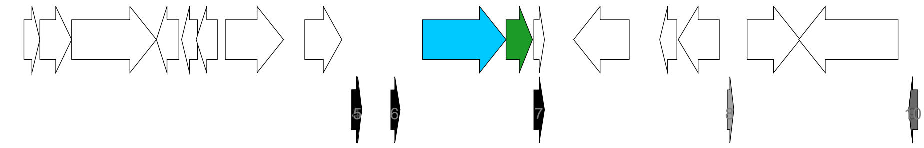

1000 nucleotides

Link to nucleotide sequence

| Accession | start | end | direction | length (aa) | Pfam/HMM | name | description | E-value |
|---|---|---|---|---|---|---|---|---|
| AGW40762.1 | 540950 | 541283 | + | 110 | PF00072<br>TIGR02875<br>TIGR01387<br>TIGR02154<br>TIGR01818 | Response_reg<br>TIGR02875<br>TIGR01387<br>TIGR02154<br>TIGR01818 | Response regulator receiver domain<br>spore_0_A: sporulation transcription factor Spo0A<br>cztR_silR_copR: heavy metal response regulator<br>PhoB: phosphate regulon transcriptional regulatory protein PhoB<br>ntrC: nitrogen regulation protein NR(I) | 1.50E-18<br>6.70E-08<br>2.20E-06<br>2.20E-06<br>4.80E-06 |
| AGW40763.1 | 541287 | 541968 | + | 226 | PF00196<br>PF00072<br>TIGR03541<br>TIGR03020<br>PF08281 | GerE<br>Response_reg<br>TIGR03541<br>TIGR03020<br>Sigma70_r4_2 | Bacterial regulatory proteins, luxR family<br>Response regulator receiver domain<br>reg_near_HchA: LuxR family transcriptional regulatory, chaperone HchA-associated<br>EpsA: transcriptional regulator EpsA<br>Sigma-70, region 4 | 2.40E-17<br>1.30E-15<br>1.90E-11<br>1.20E-09<br>2.60E-08 |

| | | | | | | | | |
|---|---|---|---|---|---|---|---|---|
| AGW40764.1 | 541972 | 543736 | + | 587 | PF07730<br>PF02518 | HisKA_3<br>HATPase_c | Histidine kinase<br>Histidine kinase-, DNA gyrase B-, and HSP90-like ATPase | 6.80E-10<br>5.80E-04 |
| AGW40765.1 | 544207 | 543736 | - | 156 | NO PFAM MATCH | - | - | - |
| AGW40766.1 | 544599 | 544260 | - | 112 | PF01035<br>TIGR00589 | DNA_binding_1<br>TIGR00589 | 6-O-methylguanine DNA methyltransferase, DNA binding domain<br>ogt: methylated-DNA--[protein]-cysteine S-methyltransferase | 4.40E-19<br>1.50E-08 |
| AGW40767.1 | 545011 | 544582 | - | 142 | PF00583<br>TIGR02406 | Acetyltransf_1<br>TIGR02406 | Acetyltransferase (GNAT) family<br>ectoine_EctA: diaminobutyrate acetyltransferase | 1.10E-08<br>6.50E-04 |
| AGW40768.1 | 545183 | 546398 | + | 404 | TIGR00127<br>PF00180<br>TIGR00175<br>TIGR02088<br>TIGR02924 | TIGR00127<br>Iso_dh<br>TIGR00175<br>TIGR02088<br>TIGR02924 | nadp_idh_euk: isocitrate dehydrogenase, NADP-dependent<br>Isocitrate/isopropylmalate dehydrogenase<br>mito_nad_idh: isocitrate dehydrogenase, NAD-dependent<br>LEU3_arch: isopropylmalate/isohomocitrate dehydrogenases<br>ICDH_alpha: isocitrate dehydrogenase | 3.90E-210<br>5.20E-75<br>2.30E-11<br>3.00E-07<br>2.00E-06 |
| AGW40769.1 | 546853 | 547639 | + | 261 | TIGR00502<br>PF01182<br>TIGR01198 | TIGR00502<br>Glucosamine_iso<br>TIGR01198 | nagB: glucosamine-6-phosphate deaminase<br>Glucosamine-6-phosphate isomerases/6-phosphogluconolactonase<br>pgl: 6-phosphogluconolactonase | 2.40E-70<br>5.20E-21<br>2.70E-20 |
| AGW40770.1 | 549318 | 551061 | + | 580 | LinH<br>LinG<br>LinE<br>PF12697<br>PF06821 | LinH<br>LinG<br>LinE<br>Abhydrolase_6<br>Ser_hydrolase | LinH (Fusion of LinE/LinG)<br>LinG Homolog<br>LinE Homolog<br>Alpha/beta hydrolase family<br>Serine hydrolase | 1.50E-113<br>1.10E-72<br>2.80E-31<br>6.90E-09<br>8.90E-04 |
| AGW40771.1 | 551057 | 551633 | + | 191 | LinL | LinL | LinL | 1.70E-36 |
| AGW40772.1 | 551646 | 551859 | + | 70 | NO PFAM MATCH | - | - | - |
| AGW40773.1 | 553651 | 552481 | - | 389 | PF00665<br>PF13683 | rve<br>rve_3 | Integrase core domain<br>Integrase core domain | 1.60E-40<br>1.90E-19 |
| AGW40774.1 | 554650 | 554284 | - | 121 | PF10823 | DUF2568 | Protein of unknown function (DUF2568) | 2.80E-20 |
| AGW40775.1 | 555536 | 554663 | - | 290 | TIGR00655<br>PF00551<br>TIGR00639<br>TIGR00460<br>PF01842 | TIGR00655<br>Formyl_trans_N<br>TIGR00639<br>TIGR00460<br>ACT | PurU: formyltetrahydrofolate deformylase<br>Formyl transferase<br>PurN: phosphoribosylglycinamide formyltransferase<br>fmt: methionyl-tRNA formyltransferase<br>ACT domain | 2.90E-91<br>2.50E-36<br>3.20E-33<br>3.30E-14<br>4.90E-08 |
| AGW40776.1 | 556111 | 557221 | + | 369 | PF07690<br>TIGR00880<br>TIGR00893<br>TIGR00881<br>TIGR00895 | MFS_1<br>TIGR00880<br>TIGR00893<br>TIGR00881<br>TIGR00895 | Major Facilitator Superfamily<br>2_A_01_02: multidrug resistance protein<br>2A0114: D-galactonate transporter<br>2A0104: phosphoglycerate transporter family protein<br>2A0115: MFS transporter, aromatic acid:H+ symporter (AAHS) family | 7.20E-45<br>3.30E-21<br>2.10E-13<br>6.30E-13<br>1.30E-12 |
| AGW40777.1 | 559271 | 557207 | - | 687 | TIGR02937<br>TIGR02983<br>TIGR02960<br>TIGR02985<br>PF13490 | TIGR02937<br>TIGR02983<br>TIGR02960<br>TIGR02985<br>zf-HC2 | sigma70-ECF: RNA polymerase sigma factor, sigma-70 family<br>SigE-fam_strep: RNA polymerase sigma-70 factor, sigma-E family<br>SigX5: RNA polymerase sigma-70 factor, TIGR02960 family<br>Sig70_bacteroi1: RNA polymerase sigma-70 factor, Bacteroides expansion family 1<br>Putative zinc-finger | 6.10E-16<br>2.10E-12<br>4.90E-09<br>6.60E-09<br>9.20E-09 |

| index | leader | core | start | end | dir | score |
|---|---|---|---|---|---|---|
| 1 | LVSLRALVIAIARATSAVIVEAVRSCDACLDPTTT | QVTADAAVVVSLIGVEDNRAASGPTAADALDRGDVV | 547825 | 548041 | + | 7 |
| 2 | VSLRALVIAIARATSAVIVEAVRSCDACLDPTTTQ | VTADAAVVVSLIGVEDNRAASGPTAADALDRGDVV | 547828 | 548041 | + | 8 |
| 3 | LRALVIAIARATSAVIVEAVRSCDACLDPTTTQV | TADAAVVVSLIGVEDNRAASGPTAADALDRGDVV | 547834 | 548041 | + | 8 |
| 4 | VIAIARATSAVIVEAVRSCDACLDPTTTQVTA | DAAVVVSLIGVEDNRAASGPTAADALDRGDVV | 547846 | 548041 | + | 8 |
| 5 | VIVEAVRSCDACLDPTTTQVTADAAVV | VSLIGVEDNRAASGPTAADALDRGDVV | 547876 | 548041 | + | 7 |
| 6 | MSLQQTGQTLNDRAVTGLDTDATASFSAQHHEVT | CFVMSIFVVAATADIAAYDMAIGASALAATAFAV | 548646 | 548853 | + | 17 |
| 7 | MTMSQTAETVNGLLSSRLKT | TTPAEGFRSQSEVTCLSVAAVMAYAAFLGAEATVDVAVGVAASAATALAV | 551646 | 551859 | + | 17 |
| 8 | VPTTCSGGGPPSQSPCSGGACGSFPPEFGL | VLPAGGACAALTGGAFSTR | 555691 | 555841 | + | 6 |
| 9 | VGIGVEHAVLALT | ETPATAIGRGDEFGRLAAGFAADAVLLDDDFRVVQVWAAGSPVARS | 559693 | 559513 | - | 6 |
| 10 | VTEVGIGVEHAVLALT | ETPATAIGRGDEFGRLAAGFAADAVLLDDDFRVVQVWAAGSPVARS | 559702 | 559513 | - | 6 |

### Results for ODA91227.1 [Leifsonia xyli subsp. xyli] back to top

previous - next

#### Architecture

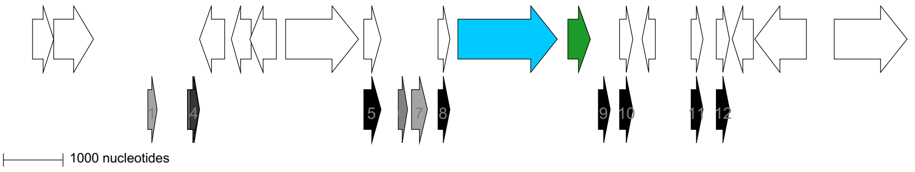

Link to nucleotide sequence

| Accession | start | end | direction | length (aa) | Pfam/HMM | name | description | E-value |
|---|---|---|---|---|---|---|---|---|
| ODA91220.1 | 94975 | 95323 | + | 115 | PF00072<br>TIGR02875<br>TIGR02154<br>TIGR01387<br>TIGR01818 | Response_reg<br>TIGR02875<br>TIGR02154<br>TIGR01387<br>TIGR01818 | Response regulator receiver domain<br>spore_0_A: sporulation transcription factor Spo0A<br>PhoB: phosphate regulon transcriptional regulatory protein PhoB<br>cztR_silR_copR: heavy metal response regulator<br>ntrC: nitrogen regulation protein NR(I) | 7.40E-19<br>2.20E-09<br>7.10E-08<br>3.40E-07<br>1.10E-06 |

| | | | | | | | | |
|---|---|---|---|---|---|---|---|---|
| ODA91221.1 | 95336 | 96008 | + | 223 | PF00196<br>PF00072<br>TIGR03541<br>TIGR03020<br>PF08281 | GerE<br>Response_reg<br>TIGR03541<br>TIGR03020<br>Sigma70_r4_2 | Bacterial regulatory proteins, luxR family<br>Response regulator receiver domain<br>reg_near_HchA: LuxR family transcriptional regulatory, chaperone HchA-associated<br>EpsA: transcriptional regulator EpsA<br>Sigma-70, region 4 | 1.10E-16<br>6.70E-14<br>6.80E-11<br>2.30E-09<br>3.20E-08 |
| ODA91402.1 | 98202 | 97776 | - | 141 | NO PFAM MATCH | - | - | - |
| ODA91222.1 | 98640 | 98301 | - | 112 | PF01035<br>TIGR00589 | DNA_binding_1<br>TIGR00589 | 6-O-methylguanine DNA methyltransferase, DNA binding domain<br>ogt: methylated-DNA--[protein]-cysteine S-methyltransferase | 4.80E-18<br>9.70E-07 |
| ODA91223.1 | 99076 | 98623 | - | 150 | PF00583<br>TIGR01575 | Acetyltransf_1<br>TIGR01575 | Acetyltransferase (GNAT) family<br>rimI: ribosomal-protein-alanine acetyltransferase | 2.90E-09<br>7.70E-04 |
| ODA91224.1 | 99222 | 100437 | + | 404 | TIGR00127<br>PF00180<br>TIGR00175<br>TIGR02924<br>TIGR02088 | TIGR00127<br>Iso_dh<br>TIGR00175<br>TIGR02924<br>TIGR02088 | nadp_idh_euk: isocitrate dehydrogenase, NADP-dependent<br>Isocitrate/isopropylmalate dehydrogenase<br>mito_nad_idh: isocitrate dehydrogenase, NAD-dependent<br>ICDH_alpha: isocitrate dehydrogenase<br>LEU3_arch: isopropylmalate/isohomocitrate dehydrogenases | 1.70E-211<br>3.90E-75<br>4.60E-11<br>7.70E-07<br>8.20E-07 |
| ODA91225.1 | 100542 | 100812 | + | 89 | NO PFAM MATCH | - | - | - |
| ODA91226.1 | 101777 | 101984 | + | 68 | NO PFAM MATCH | - | - | - |
| ODA91227.1 | 102107 | 103775 | + | 555 | LinH<br>LinG<br>LinE<br>PF12697<br>PF00975 | LinH<br>LinG<br>LinE<br>Abhydrolase_6<br>Thioesterase | LinH (Fusion of LinE/LinG)<br>LinG Homolog<br>LinE Homolog<br>Alpha/beta hydrolase family<br>Thioesterase domain | 2.30E-107<br>9.70E-69<br>4.20E-30<br>9.70E-07<br>9.90E-04 |
| ODA91228.1 | 103955 | 104345 | + | 129 | LinL | LinL | LinL | 1.20E-20 |
| ODA91229.1 | 104833 | 105058 | + | 74 | NO PFAM MATCH | - | - | - |
| ODA91230.1 | 105430 | 105214 | - | 71 | NO PFAM MATCH | - | - | - |
| ODA91231.1 | 106025 | 106232 | + | 68 | NO PFAM MATCH | - | - | - |
| ODA91232.1 | 106449 | 106668 | + | 72 | NO PFAM MATCH | - | - | - |
| ODA91233.1 | 107079 | 106713 | - | 121 | PF10823 | DUF2568 | Protein of unknown function (DUF2568) | 2.10E-20 |
| ODA91234.1 | 107965 | 107092 | - | 290 | TIGR00655<br>PF00551<br>TIGR00639<br>TIGR00460<br>PF01842 | TIGR00655<br>Formyl_trans_N<br>TIGR00639<br>TIGR00460<br>ACT | PurU: formyltetrahydrofolate deformylase<br>Formyl transferase<br>PurN: phosphoribosylglycinamide formyltransferase<br>fmt: methionyl-tRNA formyltransferase<br>ACT domain | 1.30E-90<br>2.20E-35<br>1.10E-32<br>5.60E-15<br>5.20E-07 |
| ODA91235.1 | 108432 | 109650 | + | 405 | PF07690<br>TIGR00880<br>TIGR00881<br>TIGR00893<br>TIGR00895 | MFS_1<br>TIGR00880<br>TIGR00881<br>TIGR00893<br>TIGR00895 | Major Facilitator Superfamily<br>2_A_01_02: multidrug resistance protein<br>2A0104: phosphoglycerate transporter family protein<br>2A0114: D-galactonate transporter<br>2A0115: MFS transporter, aromatic acid:H+ symporter (AAHS) family | 2.30E-41<br>1.10E-20<br>1.90E-14<br>5.00E-14<br>1.20E-10 |

| index | leader | core | start | end | dir | score |
|---|---|---|---|---|---|---|
| 1 | VGGEVARLRDQRRGAGAGCDGEPPG | PLDPGDRLRPGTHRADRARYRAEPG | 96914 | 97067 | + | 6 |
| 2 | VVKRDRDAVAITVKDDGRGGARIEAGGGLEG | LRDRVETVGGRVEVYSEAGLGTIVELLIPVPA | 97584 | 97776 | + | 6 |
| 3 | VKRDRDAVAITVKDDGRGGARIEAGGGLEGL | RDRVETVGGRVEVYSEAGLGTIVELLIPVPA | 97587 | 97776 | + | 6 |
| 4 | VKDDGRGGARIEAGGGLEGLRDRVE | TVGGRVEVYSEAGLGTIVELLIPVPA | 97620 | 97776 | + | 7 |
| 5 | MTAIKATLNEGIAAAVLVFALTVAA...SIAGRDGLPFLASLALSRRISLVPE | RTPVAALEAVLGGGTVG | 100542 | 100812 | + | 18 |
| 6 | MCPTALSMASSTRAPTTSGRSPSPAGWTCRSLVSA | LTATSASTSPAPRSPRSRA | 101106 | 101271 | + | 7 |
| 7 | MHCVTQGLSTILKARHLLLLAFGEG...EAVAGAVEGPVSASNPGSAIQLPPH | VTVLVDEAAASRLRNIAYYRYAYDHKPSWQRL | 101336 | 101591 | + | 6 |
| 8 | MSLQRTAQALSDLVATELNTDVTAGSSVPLHEVTCLGVIAVVA | ATAMDIAAYDVASGAAALAVTTLSV | 101777 | 101984 | + | 19 |
| 9 | MSLQKTAQTLNDRVTTEIDPDATASTSVLHRENT | CIGVVTIIVGAAADVAAFDVATGASALAATALAI | 104464 | 104671 | + | 18 |
| 10 | MKRNMTMSQTAETVNGVLSSRLETTTPAGCSRSQSEI | TCLSVAAVMAYAAFLGAEATVDVAVGSAAVAATALAV | 104833 | 105058 | + | 16 |
| 11 | MSLQRTAQALSDLVATELNTDVTAGSSVPLHEVTCLGVIAVVA | ATAMDIAAYDVASGAAALAVTTLSV | 106025 | 106232 | + | 19 |
| 12 | MNLIADTQARLAETTANEPLEPETLSAFARRPGETTCAAIFVA | ATVELVIPAAVEVAAADIAVTAAATALAV | 106449 | 106668 | + | 18 |

### Results for AAT89666.1 [Leifsonia xyli subsp. xyli str. CTCB07] back to top

previous - next

#### Architecture

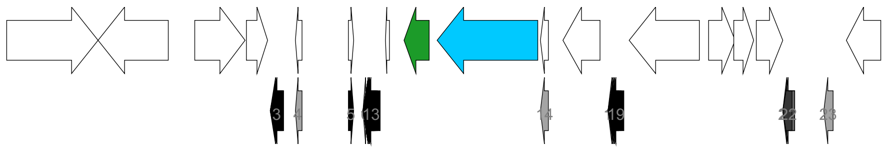

1000 nucleotides

Link to nucleotide sequence

| Accession | start | end | direction | length (aa) | Pfam/HMM | name | description | E-value |
|---|---|---|---|---|---|---|---|---|

| | | | | | | | | |
|---|---|---|---|---|---|---|---|---|
| AAT89658.1 | 1997316 | 1998900 | + | 527 | PF13490<br>PF17936<br>TIGR02949 | zf-HC2<br>Big_6<br>TIGR02949 | Putative zinc-finger<br>Bacterial Ig domain<br>anti_SigH_actin: anti-sigma factor, TIGR02949 family | 7.00E-10<br>1.70E-06<br>9.80E-05 |
| AAT89659.1 | 2000104 | 1998886 | - | 405 | PF07690<br>TIGR00880<br>TIGR00881<br>TIGR00893<br>TIGR00895 | MFS_1<br>TIGR00880<br>TIGR00881<br>TIGR00893<br>TIGR00895 | Major Facilitator Superfamily<br>2_A_01_02: multidrug resistance protein<br>2A0104: phosphoglycerate transporter family protein<br>2A0114: D-galactonate transporter<br>2A0115: MFS transporter, aromatic acid:H+ symporter (AAHS) family | 2.30E-41<br>1.10E-20<br>1.90E-14<br>5.00E-14<br>1.20E-10 |
| AAT89660.1 | 2000571 | 2001444 | + | 290 | TIGR00655<br>PF00551<br>TIGR00639<br>TIGR00460<br>PF01842 | TIGR00655<br>Formyl_trans_N<br>TIGR00639<br>TIGR00460<br>ACT | PurU: formyltetrahydrofolate deformylase<br>Formyl transferase<br>PurN: phosphoribosylglycinamide formyltransferase<br>fmt: methionyl-tRNA formyltransferase<br>ACT domain | 1.30E-90<br>2.20E-35<br>1.10E-32<br>5.60E-15<br>5.20E-07 |
| AAT89661.1 | 2001457 | 2001823 | + | 121 | PF10823 | DUF2568 | Protein of unknown function (DUF2568) | 2.10E-20 |
| AAT89662.1 | 2002430 | 2002304 | - | 41 | NO PFAM MATCH | - | - | - |
| AAT89663.1 | 2003217 | 2003322 | + | 34 | NO PFAM MATCH | - | - | - |
| AAT89664.1 | 2003946 | 2003865 | - | 26 | NO PFAM MATCH | - | - | - |
| AAT89665.1 | 2004614 | 2004191 | - | 140 | LinL | LinL | LinL | 3.00E-29 |
| AAT89666.1 | 2006507 | 2004761 | - | 581 | LinH<br>LinG<br>LinE<br>PF12697 | LinH<br>LinG<br>LinE<br>Abhydrolase_6 | LinH (Fusion of LinE/LinG)<br>LinG Homolog<br>LinE Homolog<br>Alpha/beta hydrolase family | 1.40E-112<br>5.40E-72<br>4.30E-30<br>1.10E-06 |
| AAT89667.1 | 2006678 | 2006552 | - | 41 | NO PFAM MATCH | - | - | - |
| AAT89668.1 | 2007578 | 2006945 | - | 210 | TIGR00502<br>PF01182<br>TIGR01198 | TIGR00502<br>Glucosamine_iso<br>TIGR01198 | nagB: glucosamine-6-phosphate deaminase<br>Glucosamine-6-phosphate isomerases/6-phosphogluconolactonase<br>pgl: 6-phosphogluconolactonase | 4.70E-58<br>5.20E-14<br>4.30E-13 |
| AAT89669.1 | 2009314 | 2008099 | - | 404 | TIGR00127<br>PF00180<br>TIGR00175<br>TIGR02924<br>TIGR02088 | TIGR00127<br>Iso_dh<br>TIGR00175<br>TIGR02924<br>TIGR02088 | nadp_idh_euk: isocitrate dehydrogenase, NADP-dependent<br>Isocitrate/isopropylmalate dehydrogenase<br>mito_nad_idh: isocitrate dehydrogenase, NAD-dependent<br>ICDH_alpha: isocitrate dehydrogenase<br>LEU3_arch: isopropylmalate/isohomocitrate dehydrogenases | 1.70E-211<br>3.90E-75<br>4.60E-11<br>7.70E-07<br>8.20E-07 |
| AAT89670.1 | 2009460 | 2009913 | + | 150 | PF00583<br>TIGR01575 | Acetyltransf_1<br>TIGR01575 | Acetyltransferase (GNAT) family<br>rimI: ribosomal-protein-alanine acetyltransferase | 2.90E-09<br>7.70E-04 |
| AAT89671.1 | 2009896 | 2010235 | + | 112 | PF01035<br>TIGR00589 | DNA_binding_1<br>TIGR00589 | 6-O-methylguanine DNA methyltransferase, DNA binding domain<br>ogt: methylated-DNA--[protein]-cysteine S-methyltransferase | 4.80E-18<br>9.70E-07 |
| AAT89672.1 | 2010289 | 2010760 | + | 156 | NO PFAM MATCH | - | - | - |
| AAT89673.1 | 2012453 | 2011841 | - | 203 | NO PFAM MATCH | - | - | - |

| index | leader | core | start | end | dir | score |
|---|---|---|---|---|---|---|
| 1 | MNLIADTQARLAETTANEPLEPETLSAFARRPGETTCAAIFVA | ATVELVIPAAVEVAAADIAVTAAATALAV | 2002087 | 2001868 | - | 18 |
| 2 | MRGMNLIADTQARLAETTANEPLEPETLSAFARRPGETTCAAIFVA | ATVELVIPAAVEVAAADIAVTAAATALAV | 2002096 | 2001868 | - | 18 |
| 3 | MKKEMRGMNLIADTQARLAETTANEPLEPETLSAFARRPGETTCAAIFVA | ATVELVIPAAVEVAAADIAVTAAATALAV | 2002108 | 2001868 | - | 18 |
| 4 | MPLHEVTCLGVIAVVA | ATAMDIAAYDVASGAAALAVTTLSV | 2002430 | 2002304 | - | 6 |
| 5 | MIIVAIAAAEAAGDFAA | IDIADAAVIAAATTVAG | 2003217 | 2003322 | + | 13 |
| 6 | LSVAAVMAYAAFLGAEA | TVDVAVGSAAVAATALAV | 2003586 | 2003478 | - | 6 |
| 7 | LSSRLETTTPAGCSRSQSEITCLSVAAV | MAYAAFLGAEATVDVAVGSAAVAATALAV | 2003652 | 2003478 | - | 8 |
| 8 | VNGVLSSRLETTTPAGCSRSQSEITCLSVA | AVMAYAAFLGAEATVDVAVGSAAVAATALAV | 2003664 | 2003478 | - | 8 |
| 9 | MSQTAETVNGVLSSRLETTTPAGCSRSQSEITCL | SVAAVMAYAAFLGAEATVDVAVGSAAVAATALAV | 2003685 | 2003478 | - | 18 |
| 10 | MTMSQTAETVNGVLSSRLETTTPAGCSRSQSEITC | LSVAAVMAYAAFLGAEATVDVAVGSAAVAATALAV | 2003691 | 2003478 | - | 18 |
| 11 | MKRNMTMSQTAETVNGVLSSRLETTTPAGCSRSQSEI | TCLSVAAVMAYAAFLGAEATVDVAVGSAAVAATALAV | 2003703 | 2003478 | - | 16 |
| 12 | MKKEDEMKRNMTMSQTAETVNGVLSSRLETTTPAGCSRSQ | SEITCLSVAAVMAYAAFLGAEATVDVAVGSAAVAATALAV | 2003721 | 2003478 | - | 16 |
| 13 | MYRLRADRASTSPQNAARRMKKEDEMKRNMTMSQTAETVNGVLSSRLET | TTPAGCSRSQSEITCLSVAAVMAYAAFLGAEATVDVAVGSAAVAATALAV | 2003778 | 2003478 | - | 16 |
| 14 | MPLHEVTCLGVIAVVA | ATAMDIAAYDVASGAAALAVTTLSV | 2006678 | 2006552 | - | 6 |
| 15 | MSTRCSIAGRDGLPFLASLALSRRISLVPE | RTPVAALEAVLGGGTVG | 2007868 | 2007724 | - | 6 |
| 16 | VIGGVSPSGTGGSAMSTRCSIAGRDGLPFLASLALSRRISLVPE | RTPVAALEAVLGGGTVG | 2007910 | 2007724 | - | 9 |
| 17 | VVIGGVSPSGTGGSAMSTRCSIAGRDGLPFLASLALSRRISLVPE | RTPVAALEAVLGGGTVG | 2007913 | 2007724 | - | 9 |
| 18 | VLVFALTVAADTVVIGGVSPSGTGG...SIAGRDGLPFLASLALSRRISLVPE | RTPVAALEAVLGGGTVG | 2007949 | 2007724 | - | 8 |
| 19 | VTAIKATLNEGIAAAVLVFALTVAA...SIAGRDGLPFLASLALSRRISLVPE | RTPVAALEAVLGGGTVG | 2007994 | 2007724 | - | 18 |
| 20 | VKDDGRGGARIEAGGGLEGLRDRVE | TVGGRVEVYSEAGLGTIVELLIPVPA | 2010916 | 2010760 | - | 7 |
| 21 | VKRDRDAVAITVKDDGRGGARIEAGGGLEGL | RDRVETVGGRVEVYSEAGLGTIVELLIPVPA | 2010949 | 2010760 | - | 6 |
| 22 | VVKRDRDAVAITVKDDGRGGARIEAGGGLEG | LRDRVETVGGRVEVYSEAGLGTIVELLIPVPA | 2010952 | 2010760 | - | 6 |
| 23 | VGGEVARLRDQRRGAGAGCDGEPPG | PLDPGDRLRPGTHRADRARYRAEPG | 2011622 | 2011469 | - | 6 |

### Results for WP_166328536.1 [Leucobacter sp. HDW9A] back to top

previous - next

### Architecture

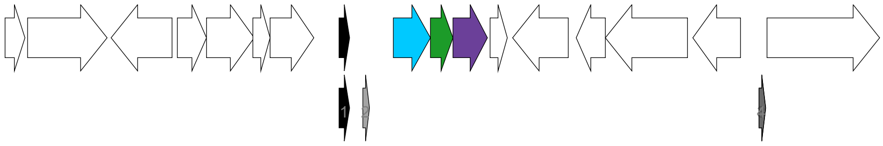

1000 nucleotides

Link to nucleotide sequence

| Accession | start | end | direction | length (aa) | Pfam/HMM | name | description | E-value |
|---|---|---|---|---|---|---|---|---|
| WP_166328524.1 | 333568 | 334048 | + | 159 | PF13396 | PLDc_N | Phospholipase_D-nuclease N-terminal | 1.20E-10 |
| WP_166331849.1 | 334083 | 335925 | + | 613 | TIGR00173<br>PF02776<br>TIGR02418<br>TIGR03457<br>TIGR00118 | TIGR00173<br>TPP_enzyme_N<br>TIGR02418<br>TIGR03457<br>TIGR00118 | menD: 2-succinyl-5-enolpyruvyl-6-hydroxy-3-cyclohexene-1-carboxylic-acid synthase<br>Thiamine pyrophosphate enzyme, N-terminal TPP binding domain<br>acolac_catab: acetolactate synthase, catabolic<br>sulphoacet_xsc: sulfoacetaldehyde acetyltransferase<br>acolac_lg: acetolactate synthase, large subunit, biosynthetic type | 4.20E-95<br>4.40E-25<br>4.10E-11<br>2.30E-08<br>4.70E-07 |
| WP_166328526.1 | 337428 | 336027 | - | 466 | TIGR00883<br>PF00083<br>PF07690<br>TIGR00891<br>TIGR00898 | TIGR00883<br>Sugar_tr<br>MFS_1<br>TIGR00891<br>TIGR00898 | 2A0106: MFS transporter, metabolite:H+ symporter (MHS) family protein<br>Sugar (and other) transporter<br>Major Facilitator Superfamily<br>2A0112: MFS transporter, sialate:H+ symporter (SHS) family<br>2A0119: cation transport protein | 1.90E-102<br>1.90E-35<br>3.30E-28<br>2.10E-19<br>1.90E-17 |
| WP_166328527.1 | 337569 | 338238 | + | 222 | TIGR02154<br>TIGR01387<br>TIGR03787<br>PF00072<br>PF00486 | TIGR02154<br>TIGR01387<br>TIGR03787<br>Response_reg<br>Trans_reg_C | PhoB: phosphate regulon transcriptional regulatory protein PhoB<br>cztR_silR_copR: heavy metal response regulator<br>marine_sort_RR: proteobacterial dedicated sortase system response regulator<br>Response regulator receiver domain<br>Transcriptional regulatory protein, C terminal | 5.30E-49<br>2.80E-42<br>6.20E-29<br>5.00E-24<br>3.40E-19 |
| WP_166328528.1 | 338234 | 339335 | + | 366 | TIGR01386<br>PF00512<br>PF00672<br>TIGR03785 | TIGR01386<br>HisKA<br>HAMP<br>TIGR03785 | cztS_silS_copS: heavy metal sensor kinase<br>His Kinase A (phospho-acceptor) domain<br>HAMP domain<br>marine_sort_HK: proteobacterial dedicated sortase system histidine kinase | 9.40E-10<br>6.40E-09<br>1.10E-07<br>6.40E-04 |
| WP_166328530.1 | 339319 | 339697 | + | 125 | TIGR02966<br>TIGR01386<br>PF02518<br>TIGR02956<br>TIGR03785 | TIGR02966<br>TIGR01386<br>HATPase_c<br>TIGR02956<br>TIGR03785 | phoR_proteo: phosphate regulon sensor kinase PhoR<br>cztS_silS_copS: heavy metal sensor kinase<br>Histidine kinase-, DNA gyrase B-, and HSP90-like ATPase<br>TMAO_torS: TMAO reductase sytem sensor TorS<br>marine_sort_HK: proteobacterial dedicated sortase system histidine kinase | 3.20E-18<br>7.70E-16<br>3.10E-15<br>2.50E-11<br>1.30E-08 |
| WP_166328532.1 | 339693 | 340725 | + | 343 | TIGR02122<br>PF16868<br>TIGR01728 | TIGR02122<br>NMT1_3<br>TIGR01728 | TRAP_TAXI: TRAP transporter solute receptor, TAXI family<br>NMT1-like family<br>SsuA_fam: ABC transporter, substrate-binding protein, aliphatic sulfonates family | 2.10E-45<br>4.60E-42<br>1.10E-04 |
| WP_166328534.1 | 341296 | 341539 | + | 80 | LinA | LinA | LinA (linaridin precursor) | 1.10E-07 |
| WP_166328536.1 | 342543 | 343407 | + | 287 | LinH<br>LinE<br>PF12697<br>TIGR02427<br>TIGR03611 | LinH<br>LinE<br>Abhydrolase_6<br>TIGR02427<br>TIGR03611 | LinH (Fusion of LinE/LinG)<br>LinE Homolog<br>Alpha/beta hydrolase family<br>protocat_pcaD: 3-oxoadipate enol-lactonase<br>RutD: pyrimidine utilization protein D | 4.90E-47<br>1.70E-43<br>8.20E-10<br>8.80E-06<br>1.80E-05 |
| WP_166328538.1 | 343418 | 343931 | + | 170 | LinL | LinL | LinL | 7.50E-37 |
| WP_166328540.1 | 343945 | 344740 | + | 264 | PF13649<br>PF13847<br>PF08241<br>PF08242<br>TIGR03438 | Methyltransf_25<br>Methyltransf_31<br>Methyltransf_11<br>Methyltransf_12<br>TIGR03438 | Methyltransferase domain<br>Methyltransferase domain<br>Methyltransferase domain<br>Methyltransferase domain<br>egtD_ergothio: dimethylhistidine N-methyltransferase | 6.60E-13<br>9.00E-10<br>1.10E-09<br>7.60E-08<br>5.70E-06 |
| WP_166328542.1 | 344786 | 345179 | + | 130 | NO PFAM MATCH | - | - | - |
| WP_166331850.1 | 346617 | 345318 | - | 432 | TIGR00543<br>PF00425<br>TIGR00564<br>TIGR01820<br>TIGR01824 | TIGR00543<br>Chorismate_bind<br>TIGR00564<br>TIGR01820<br>TIGR01824 | isochor_syn: isochorismate synthase<br>chorismate binding enzyme<br>trpE_most: anthranilate synthase component I<br>TrpE-arch: anthranilate synthase component I<br>PabB-clade2: aminodeoxychorismate synthase, component I | 1.80E-75<br>1.30E-65<br>6.40E-40<br>1.30E-38<br>4.50E-38 |
| WP_166328544.1 | 347469 | 346785 | - | 227 | PF13166<br>TIGR03411<br>TIGR02857<br>TIGR00972 | AAA_13<br>TIGR03411<br>TIGR02857<br>TIGR00972 | AAA domain<br>urea_trans_UrtD: urea ABC transporter, ATP-binding protein UrtD<br>CydD: thiol reductant ABC exporter, CydD subunit<br>3a0107s01c2: phosphate ABC transporter, ATP-binding protein | 1.00E-06<br>8.10E-06<br>1.70E-04<br>2.20E-04 |
| WP_166328545.1 | 349358 | 347471 | - | 628 | NO PFAM MATCH | - | - | - |
| WP_166328547.1 | 350600 | 349496 | - | 367 | PF13588<br>PF04313 | HSDR_N_2<br>HSDR_N | Type I restriction enzyme R protein N terminus (HSDR_N)<br>Type I restriction enzyme R protein N terminus (HSDR_N) | 7.60E-11<br>2.90E-08 |
| WP_166328549.1 | 351202 | 353800 | + | 865 | TIGR00376<br>PF13087<br>PF13086<br>PF13245<br>PF13604 | TIGR00376<br>AAA_12<br>AAA_11<br>AAA_19<br>AAA_30 | TIGR00376: putative DNA helicase<br>AAA domain<br>AAA domain<br>AAA domain<br>AAA domain | 1.10E-46<br>2.60E-27<br>3.10E-24<br>1.20E-14<br>2.00E-13 |

| index | leader | core | start | end | dir | score |
|---|---|---|---|---|---|---|
| 1 | MTSQRLAAVLSEFDFLGKDEMHIGSSGTKGIL | ATPAITVIGMTVGYFAACWQAGAAVCQVMNAYGRVAHDRLEEHKQTLL | 341296 | 341539 | + | 26 |
| 2 | MVGQGGPVTSFEGCRGCVNCITYGG | WNSQTSWRRWVGPFKFPSSSRYFHS | 341849 | 342002 | + | 6 |
| 3 | VAHLRIRLEKSKSTFTCMLRVRSKSN | TLGGRNGTATRWRILHRCKTQLVWRTA | 351012 | 351174 | + | 6 |
| 4 | LRIRLEKSKSTFTCMLRVRSKSNTL | GGRNGTATRWRILHRCKTQLVWRTA | 351021 | 351174 | + | 6 |

### Results for WP_045251507.1 [Microbacterium azadirachtae] back to top

previous - next

#### Architecture

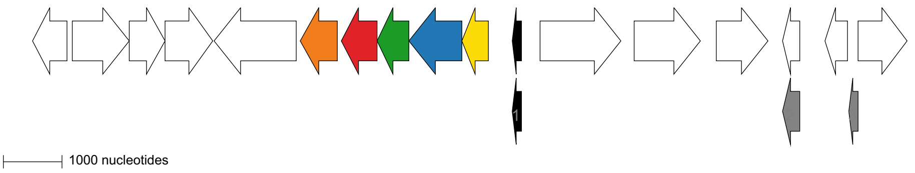

Link to nucleotide sequence

| Accession | start | end | direction | length (aa) | Pfam/HMM | name | description | E-value |
|---|---|---|---|---|---|---|---|---|
| WP_169748242.1 | 24697 | 24091 | - | 201 | PF16859<br>PF00440 | TetR_C_11<br>TetR_N | Tetracyclin repressor-like, C-terminal domain<br>Bacterial regulatory proteins, tetR family | 5.40E-20<br>6.30E-08 |
| WP_082072407.1 | 24782 | 25754 | + | 323 | PF05655 | AvrD | Pseudomonas avirulence D protein (AvrD) | 7.00E-30 |
| WP_045251504.1 | 25750 | 26365 | + | 204 | NO PFAM MATCH | - | - | - |
| WP_169748243.1 | 26361 | 27168 | + | 268 | PF13561<br>TIGR01830<br>PF00106<br>TIGR01829<br>TIGR02415 | adh_short_C2<br>TIGR01830<br>adh_short<br>TIGR01829<br>TIGR02415 | Enoyl-(Acyl carrier protein) reductase<br>3oxo_ACP_reduc: 3-oxoacyl-[acyl-carrier-protein] reductase<br>short chain dehydrogenase<br>AcAcCoA_reduct: acetoacetyl-CoA reductase<br>23BDH: acetoin reductases | 3.10E-45<br>6.20E-38<br>1.40E-37<br>2.70E-33<br>1.40E-30 |
| WP_156156804.1 | 28606 | 27193 | - | 470 | NO PFAM MATCH | - | - | - |
| WP_045251649.1 | 29310 | 28677 | - | 210 | TIGR01189<br>TIGR02857<br>TIGR03873<br>PF00005<br>TIGR01188 | TIGR01189<br>TIGR02857<br>TIGR03873<br>ABC_tran<br>TIGR01188 | ccmA: heme ABC exporter, ATP-binding protein CcmA<br>CydD: thiol reductant ABC exporter, CydD subunit<br>F420-0_ABC_ATP: proposed F420-0 ABC transporter, ATP-binding protein<br>ABC transporter<br>drrA: daunorubicin resistance ABC transporter, ATP-binding protein | 3.70E-31<br>1.30E-26<br>2.50E-26<br>1.00E-24<br>2.10E-24 |
| WP_082072410.1 | 29989 | 29368 | - | 206 | TIGR00521<br>PF02441<br>TIGR02113 | TIGR00521<br>Flavoprotein<br>TIGR02113 | coaBC_dfp: phosphopantothenoylcysteine decarboxylase / phosphopantothenate--cysteine ligase<br>Flavoprotein<br>coaC_strep: phosphopantothenoylcysteine decarboxylase | 7.70E-13<br>1.70E-11<br>3.40E-08 |
| WP_045251506.1 | 30522 | 29985 | - | 178 | LinL | LinL | LinL | 8.60E-50 |
| WP_045251507.1 | 31430 | 30518 | - | 303 | LinE<br>LinH<br>PF12697<br>TIGR02427<br>TIGR03611 | LinE<br>LinH<br>Abhydrolase_6<br>TIGR02427<br>TIGR03611 | LinE Homolog<br>LinH (Fusion of LinE/LinG)<br>Alpha/beta hydrolase family<br>protocat_pcaD: 3-oxoadipate enol-lactonase<br>RutD: pyrimidine utilization protein D | 1.90E-80<br>1.30E-48<br>2.40E-09<br>5.20E-06<br>1.50E-04 |
| WP_052674424.1 | 31882 | 31426 | - | 151 | LinG<br>LinH<br>TIGR04033<br>PF06762 | LinG<br>LinH<br>TIGR04033<br>LMF1 | LinG Homolog<br>LinH (Fusion of LinE/LinG)<br>export_SdpB: antimicrobial peptide system protein, SdpB family<br>Lipase maturation factor | 1.60E-61<br>1.30E-55<br>2.00E-06<br>3.80E-05 |
| WP_169748244.1 | 32453 | 32279 | - | 57 | LinA | LinA | LinA (linaridin precursor) | 1.50E-11 |
| WP_082072412.1 | 32753 | 34127 | + | 457 | TIGR02734<br>PF01593<br>PF13450<br>TIGR03467<br>TIGR02730 | TIGR02734<br>Amino_oxidase<br>NAD_binding_8<br>TIGR03467<br>TIGR02730 | crtI_fam: phytoene desaturase<br>Flavin containing amine oxidoreductase<br>NAD(P)-binding Rossmann-like domain<br>HpnE: squalene-associated FAD-dependent desaturase<br>carot_isom: carotene isomerase | 9.80E-09<br>8.90E-07<br>4.20E-05<br>7.50E-05<br>4.30E-04 |
| WP_076611781.1 | 34359 | 35492 | + | 377 | INFERRED GENE | - | - | - |
| WP_013807217.1 | 35775 | 36637 | + | 287 | INFERRED GENE | - | - | - |
| WP_156156806.1 | 37196 | 36893 | - | 100 | PF13830 | DUF4192 | Domain of unknown function (DUF4192) | 1.00E-04 |
| WP_045251511.1 | 38007 | 37623 | - | 127 | PF04237 | YjbR | YjbR | 5.10E-07 |
| WP_045251512.1 | 38188 | 39013 | + | 274 | PF01261<br>TIGR04379 | AP_endonuc_2<br>TIGR04379 | Xylose isomerase-like TIM barrel<br>myo_inos_iolE: myo-inosose-2 dehydratase | 2.60E-15<br>1.60E-11 |

| index | leader | core | start | end | dir | score |
|---|---|---|---|---|---|---|
| 1 | MSTTLTASRALEAEVNAIFGDVEAGFAETSEMPVM | TTPMIVPAATMVIQGSTICLVC | 32453 | 32279 | - | 32 |
| 2 | MARSTMGTVAAIIVGDAARPDPARL...WLSWALGRSTRAAFFTEQATRTLGA | PTFTATLAEIIDRGPVPAWAFAG | 37196 | 36893 | - | 7 |
| 3 | VTFLLCKGTGTPCPCGQSIRNLHLEV | YVPSPQDARAAITRGPTEIRDDSVRVRGGA | 38187 | 38016 | - | 7 |

### Results for SIR96088.1 [Microbacterium sp. RURRCA19A] back to top

previous - next

#### Architecture

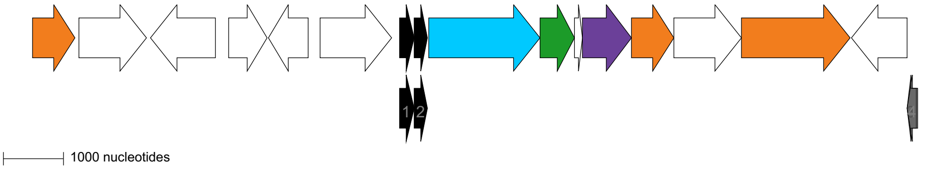

Link to nucleotide sequence

| Accession | start | end | direction | length (aa) | Pfam/HMM | name | description | E-value |
|---|---|---|---|---|---|---|---|---|

| | | | | | | | | |
|---|---|---|---|---|---|---|---|---|
| SIR95953.1 | 466019 | 466724 | + | 234 | TIGR03608<br>TIGR02211<br>TIGR02673<br>TIGR02982<br>TIGR00968 | TIGR03608<br>TIGR02211<br>TIGR02673<br>TIGR02982<br>TIGR00968 | L_ocin_972_ABC: putative bacteriocin export ABC transporter, lactococcin 972 group<br>LolD_lipo_ex: lipoprotein releasing system, ATP-binding protein<br>FtsE: cell division ATP-binding protein FtsE<br>heterocyst_DevA: ABC exporter ATP-binding subunit, DevA family<br>3a0106s01: sulfate ABC transporter, ATP-binding protein | 9.90E-57<br>5.10E-55<br>8.30E-54<br>1.80E-50<br>9.10E-45 |
| SIR95967.1 | 466792 | 467908 | + | 371 | TIGR03434<br>PF02687 | TIGR03434<br>FtsX | ADOP: acidobacterial duplicated orphan permease<br>FtsX-like permease family | 6.90E-15<br>5.20E-07 |
| SIR95982.1 | 469065 | 467988 | - | 358 | PF00724<br>TIGR03997<br>TIGR03996 | Oxidored_FMN<br>TIGR03997<br>TIGR03996 | NADH:flavin oxidoreductase / NADH oxidase family<br>mycofact_OYE_2: mycofactocin system FadH/OYE family oxidoreductase 2<br>mycofact_OYE_1: mycofactocin system FadH/OYE family oxidoreductase 1 | 4.90E-76<br>2.20E-44<br>2.50E-27 |
| SIR96018.1 | 469275 | 469917 | + | 213 | TIGR00558<br>PF01243<br>PF10590<br>TIGR04026<br>TIGR03618 | TIGR00558<br>Putative_PNPOx<br>PNP_phzG_C<br>TIGR04026<br>TIGR03618 | pdxH: pyridoxamine 5'-phosphate oxidase<br>Pyridoxamine 5'-phosphate oxidase<br>Pyridoxine 5'-phosphate oxidase C-terminal dimerisation region<br>PPOX_FMN_cyano: PPOX class probable FMN-dependent enzyme, alr4036 family<br>Rv1155_F420: PPOX class probable F420-dependent enzyme | 4.90E-72<br>2.80E-20<br>2.60E-17<br>6.50E-10<br>3.80E-09 |
| SIR96032.1 | 470593 | 469933 | - | 219 | PF00072<br>TIGR02875<br>PF00196<br>TIGR01818<br>TIGR02154 | Response_reg<br>TIGR02875<br>GerE<br>TIGR01818<br>TIGR02154 | Response regulator receiver domain<br>spore_0_A: sporulation transcription factor Spo0A<br>Bacterial regulatory proteins, luxR family<br>ntrC: nitrogen regulation protein NR(I)<br>PhoB: phosphate regulon transcriptional regulatory protein PhoB | 2.20E-26<br>1.50E-18<br>4.00E-17<br>2.60E-13<br>3.00E-12 |
| SIR96047.1 | 470804 | 471992 | + | 395 | PF07730 | HisKA_3 | Histidine kinase | 3.30E-12 |
| SIR96061.1 | 472125 | 472371 | + | 81 | LinA | LinA | LinA (linaridin precursor) | 2.70E-07 |
| SIR96076.1 | 472373 | 472583 | + | 69 | LinA | LinA | LinA (linaridin precursor) | 4.10E-09 |
| SIR96088.1 | 472620 | 474462 | + | 613 | LinH<br>LinG<br>LinE<br>PF12697<br>PF00561 | LinH<br>LinG<br>LinE<br>Abhydrolase_6<br>Abhydrolase_1 | LinH (Fusion of LinE/LinG)<br>LinG Homolog<br>LinE Homolog<br>Alpha/beta hydrolase family<br>alpha/beta hydrolase fold | 5.00E-<br>171<br>6.10E-86<br>2.00E-72<br>6.10E-10<br>4.00E-09 |
| SIR96101.1 | 474461 | 475031 | + | 189 | LinL | LinL | LinL | 1.10E-52 |
| SIR96118.1 | 475042 | 475171 | + | 42 | NO PFAM MATCH | - | - | - |
| SIR96132.1 | 475167 | 475986 | + | 272 | PF13649<br>PF13489<br>PF08241<br>PF08242 | Methyltransf_25<br>Methyltransf_23<br>Methyltransf_11<br>Methyltransf_12 | Methyltransferase domain<br>Methyltransferase domain<br>Methyltransferase domain<br>Methyltransferase domain | 2.10E-10<br>2.60E-07<br>1.30E-06<br>1.90E-06 |
| SIR96149.1 | 475982 | 476687 | + | 234 | TIGR03864<br>TIGR01188<br>TIGR03522<br>TIGR03740<br>TIGR04406 | TIGR03864<br>TIGR01188<br>TIGR03522<br>TIGR03740<br>TIGR04406 | PQQ_ABC_ATP: ABC transporter, ATP-binding subunit, PQQ-dependent alcohol dehydrogenase system<br>drrA: daunorubicin resistance ABC transporter, ATP-binding protein<br>GldA_ABC_ATP: gliding motility-associated ABC transporter ATP-binding subunit GldA<br>gallidermin_ABC: lantibiotic protection ABC transporter, ATP-binding subunit<br>LPS_export_lptB: LPS export ABC transporter ATP-binding protein | 1.50E-45<br>1.60E-45<br>7.20E-44<br>4.40E-43<br>7.60E-37 |
| SIR96164.1 | 476683 | 477823 | + | 379 | PF12698 | ABC2_membrane_3 | ABC-2 family transporter protein | 2.70E-04 |
| SIR96179.1 | 477819 | 479625 | + | 601 | TIGR02204<br>TIGR02203<br>TIGR00958<br>TIGR03796<br>TIGR03797 | TIGR02204<br>TIGR02203<br>TIGR00958<br>TIGR03796<br>TIGR03797 | MsbA_rel: ABC transporter, permease/ATP-binding protein<br>MsbA_lipidA: lipid A export permease/ATP-binding protein MsbA<br>3a01208: antigen peptide transporter 2<br>NHLM_micro_ABC1: NHLM bacteriocin system ABC transporter, peptidase/ATP-binding protein<br>NHLM_micro_ABC2: NHLM bacteriocin system ABC transporter, ATP-binding protein | 4.60E-97<br>1.70E-90<br>2.90E-84<br>1.70E-78<br>1.50E-75 |
| SIR96192.1 | 480574 | 479644 | - | 309 | PF06966<br>PF02544<br>PF04140 | DUF1295<br>Steroid_dh<br>ICMT | Protein of unknown function (DUF1295)<br>3-oxo-5-alpha-steroid 4-dehydrogenase<br>Isoprenylcysteine carboxyl methyltransferase (ICMT) family | 2.40E-68<br>1.30E-07<br>2.50E-04 |

| index | leader | core | start | end | dir | score |
|---|---|---|---|---|---|---|
| 1 | MMTMTMMSRDAALSARVGAAVAAIPDDGPFGNAVSAPVL | GTPATVAAAPAAVATWKLAAAAVGGGVSVVGAYVAGRAVGTW | 472125 | 472371 | + | 29 |
| 2 | MATLEARVGDAVTVFSSRPAIGEAVRVRSDADGPVL | ATPLALAGAFAGGAAIGGGLVAAFEAGRAVAGG | 472373 | 472583 | + | 28 |
| 3 | VSRSEQVRRFELVVADLDDRELVT | PTLKIKRTVVVDRAAETIASLYRGAPS | 480726 | 480570 | - | 6 |
| 4 | VDAANALVSRSEQVRRFELVVADLDDRELVT | PTLKIKRTVVVDRAAETIASLYRGAPS | 480747 | 480570 | - | 6 |

### Results for WP_091734673.1 [Microbacterium sp. cl127] back to top

previous - next

#### Architecture

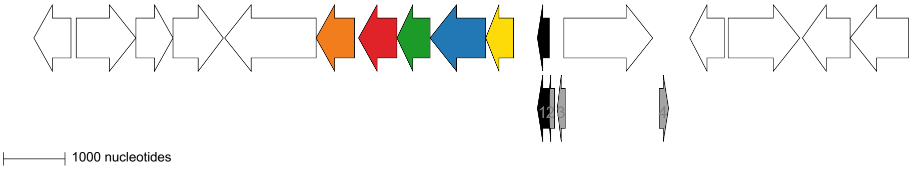

Link to nucleotide sequence

| Accession | start | end | direction | length (aa) | Pfam/HMM | name | description | E-value |
|---|---|---|---|---|---|---|---|---|
| WP_091734660.1 | 383207 | 382601 | - | 201 | PF16859<br>PF00440 | TetR_C_11<br>TetR_N | Tetracyclin repressor-like, C-terminal domain<br>Bacterial regulatory proteins, tetR family | 1.80E-21<br>6.30E-08 |
| WP_139232187.1 | 383292 | 384264 | + | 323 | PF05655 | AvrD | Pseudomonas avirulence D protein (AvrD) | 5.10E-29 |
| WP_091734663.1 | 384260 | 384875 | + | 204 | PF16859 | TetR_C_11 | Tetracyclin repressor-like, C-terminal domain | 6.80E-04 |

| | | | | | | | | |
|---|---|---|---|---|---|---|---|---|
| WP_091734665.1 | 384865 | 385681 | + | 271 | PF13561<br>PF00106<br>TIGR01830<br>TIGR01829<br>TIGR01963 | adh_short_C2<br>adh_short<br>TIGR01830<br>TIGR01829<br>TIGR01963 | Enoyl-(Acyl carrier protein) reductase<br>short chain dehydrogenase<br>3oxo_ACP_reduc: 3-oxoacyl-[acyl-carrier-protein] reductase<br>AcAcCoA_reduct: acetoacetyl-CoA reductase<br>PHB_DH: 3-hydroxybutyrate dehydrogenase | 2.40E-45<br>9.00E-38<br>1.80E-37<br>9.70E-33<br>7.10E-31 |
| WP_139232188.1 | 387194 | 385706 | - | 495 | NO PFAM MATCH | - | - | - |
| WP_091737084.1 | 387823 | 387190 | - | 210 | TIGR01189<br>TIGR03873<br>TIGR02857<br>TIGR02673<br>TIGR01166 | TIGR01189<br>TIGR03873<br>TIGR02857<br>TIGR02673<br>TIGR01166 | ccmA: heme ABC exporter, ATP-binding protein CcmA<br>F420-0_ABC_ATP: proposed F420-0 ABC transporter, ATP-binding protein<br>CydD: thiol reductant ABC exporter, CydD subunit<br>FtsE: cell division ATP-binding protein FtsE<br>cbiO: cobalt ABC transporter, ATP-binding protein | 5.40E-31<br>9.30E-28<br>1.30E-26<br>7.80E-25<br>1.70E-24 |
| WP_091734669.1 | 388502 | 387881 | - | 206 | TIGR00521<br>PF02441<br>TIGR02113 | TIGR00521<br>Flavoprotein<br>TIGR02113 | coaBC_dfp: phosphopantothenoylcysteine decarboxylase / phosphopantothenate--cysteine ligase<br>Flavoprotein<br>coaC_strep: phosphopantothenoylcysteine decarboxylase | 5.00E-13<br>8.70E-11<br>5.70E-09 |
| WP_091734671.1 | 389035 | 388498 | - | 178 | LinL | LinL | LinL | 3.80E-48 |
| WP_091734673.1 | 389943 | 389031 | - | 303 | LinE<br>LinH<br>PF12697<br>TIGR02427<br>TIGR03611 | LinE<br>LinH<br>Abhydrolase_6<br>TIGR02427<br>TIGR03611 | LinE Homolog<br>LinH (Fusion of LinE/LinG)<br>Alpha/beta hydrolase family<br>protocat_pcaD: 3-oxoadipate enol-lactonase<br>RutD: pyrimidine utilization protein D | 8.20E-82<br>1.90E-50<br>9.40E-09<br>1.10E-05<br>3.10E-04 |
| WP_091734675.1 | 390395 | 389939 | - | 151 | LinG<br>LinH<br>TIGR04033<br>PF06762 | LinG<br>LinH<br>TIGR04033<br>LMF1 | LinG Homolog<br>LinH (Fusion of LinE/LinG)<br>export_SdpB: antimicrobial peptide system protein, SdpB family<br>Lipase maturation factor | 9.60E-61<br>1.80E-55<br>2.60E-05<br>5.30E-05 |
| WP_091734677.1 | 390978 | 390792 | - | 61 | LinA | LinA | LinA (linaridin precursor) | 8.90E-12 |
| WP_139232189.1 | 391213 | 392644 | + | 476 | TIGR02734<br>TIGR03467<br>PF13450<br>PF01593<br>TIGR02733 | TIGR02734<br>TIGR03467<br>NAD_binding_8<br>Amino_oxidase<br>TIGR02733 | crtI_fam: phytoene desaturase<br>HpnE: squalene-associated FAD-dependent desaturase<br>NAD(P)-binding Rossmann-like domain<br>Flavin containing amine oxidoreductase<br>desat_CrtD: C-3',4' desaturase CrtD | 5.30E-16<br>4.50E-12<br>1.10E-11<br>1.70E-10<br>2.60E-09 |
| WP_091734682.1 | 393816 | 393249 | - | 188 | PF01327<br>TIGR00079 | Pep_deformylase<br>TIGR00079 | Polypeptide deformylase<br>pept_deformyl: peptide deformylase | 9.80E-42<br>1.80E-34 |
| WP_139232217.1 | 393883 | 395032 | + | 382 | PF05653 | Mg_trans_NIPA | Magnesium transporter NIPA | 1.50E-07 |
| WP_091734684.1 | 395863 | 395083 | - | 259 | PF01182<br>TIGR01198<br>TIGR00502 | Glucosamine_iso<br>TIGR01198<br>TIGR00502 | Glucosamine-6-phosphate isomerases/6-phosphogluconolactonase<br>pgl: 6-phosphogluconolactonase<br>nagB: glucosamine-6-phosphate deaminase | 1.30E-72<br>2.70E-67<br>1.50E-07 |
| WP_091734686.1 | 396800 | 395864 | - | 311 | PF10128<br>TIGR00534 | OpcA_G6PD_assem<br>TIGR00534 | Glucose-6-phosphate dehydrogenase subunit<br>OpcA: glucose-6-phosphate dehydrogenase assembly protein OpcA | 2.30E-83<br>3.50E-66 |

| index | leader | core | start | end | dir | score |
|---|---|---|---|---|---|---|
| 1 | MEINMSTTLIASRALEAEVNAIFGDVEAGFAETSEMPVM | TTPMIVPAATMVIQGSTICLVC | 390978 | 390792 | - | 32 |
| 2 | LSRRMAASSLSLDLRY | SAVCNVTDSTNEWRST | 391061 | 390962 | - | 6 |
| 3 | MRRVSRGAARRKRVGD | TPPGRAVAQAIATATRRPVYVTDHP | 391229 | 391103 | - | 6 |
| 4 | VAPFVSVWAPTLSDFGTGQISDAPGKS | VLEAINGALRGSRASRSSTRSQTTRSTS | 392753 | 392921 | + | 6 |

### Results for WP_150450697.1 [Microbacterium rhizomatis] back to top

previous - next

#### Architecture

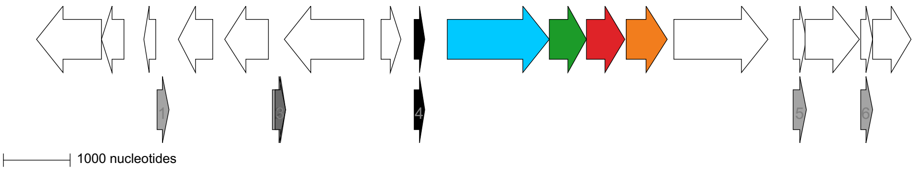

Link to nucleotide sequence

| Accession | start | end | direction | length (aa) | Pfam/HMM | name | description | E-value |
|---|---|---|---|---|---|---|---|---|
| WP_150450683.1 | 5461 | 4480 | - | 326 | NO PFAM MATCH | - | - | - |
| WP_150450684.1 | 5804 | 5462 | - | 113 | PF03551<br>TIGR03433<br>TIGR02719 | PadR<br>TIGR03433<br>TIGR02719 | Transcriptional regulator PadR-like family<br>padR_acidobact: transcriptional regulator, Acidobacterial, PadR-family<br>repress_PhaQ: poly-beta-hydroxybutyrate-responsive repressor | 2.40E-19<br>1.60E-14<br>6.30E-09 |
| WP_150450686.1 | 6276 | 6090 | - | 61 | NO PFAM MATCH | - | - | - |
| WP_150450688.1 | 7122 | 6615 | - | 168 | NO PFAM MATCH | - | - | - |
| WP_150450690.1 | 7970 | 7316 | - | 217 | NO PFAM MATCH | - | - | - |
| WP_150450692.1 | 9388 | 8197 | - | 396 | PF07690<br>TIGR00881 | MFS_1<br>TIGR00881 | Major Facilitator Superfamily<br>2A0104: phosphoglycerate transporter family protein | 1.40E-22<br>1.60E-07 |
| WP_170285758.1 | 9641 | 9953 | + | 103 | PF04134 | DUF393 | Protein of unknown function, DUF393 | 2.50E-10 |
| WP_150450695.1 | 10137 | 10302 | + | 54 | LinA | LinA | LinA (linaridin precursor) | 1.70E-13 |

| | | | | | | | | |
|---|---|---|---|---|---|---|---|---|
| WP_150450697.1 | 10636 | 12178 | + | 513 | LinH<br>LinG<br>LinE<br>PF12697<br>TIGR04033 | LinH<br>LinG<br>LinE<br>Abhydrolase_6<br>TIGR04033 | LinH (Fusion of LinE/LinG)<br>LinG Homolog<br>LinE Homolog<br>Alpha/beta hydrolase family<br>export_SdpB: antimicrobial peptide system protein, SdpB family | 4.80E-150<br>4.60E-68<br>8.80E-67<br>8.50E-08<br>3.30E-04 |
| WP_150450698.1 | 12167 | 12728 | + | 186 | LinL | LinL | LinL | 3.50E-51 |
| WP_150450699.1 | 12724 | 13306 | + | 193 | TIGR00521<br>PF02441<br>TIGR02113 | TIGR00521<br>Flavoprotein<br>TIGR02113 | coaBC_dfp: phosphopantothenoylcysteine decarboxylase / phosphopantothenate--cysteine ligase<br>Flavoprotein<br>coaC_strep: phosphopantothenoylcysteine decarboxylase | 3.10E-09<br>3.60E-07<br>2.40E-05 |
| WP_150450701.1 | 13317 | 13947 | + | 209 | TIGR01166<br>TIGR02857<br>TIGR01189<br>TIGR00968<br>TIGR02673 | TIGR01166<br>TIGR02857<br>TIGR01189<br>TIGR00968<br>TIGR02673 | cbiO: cobalt ABC transporter, ATP-binding protein<br>CydD: thiol reductant ABC exporter, CydD subunit<br>ccmA: heme ABC exporter, ATP-binding protein CcmA<br>3a0106s01: sulfate ABC transporter, ATP-binding protein<br>FtsE: cell division ATP-binding protein FtsE | 2.70E-28<br>2.10E-27<br>4.20E-27<br>6.00E-27<br>6.00E-27 |
| WP_150450703.1 | 14039 | 15443 | + | 467 | NO PFAM MATCH | - | - | - |
| WP_150450777.1 | 15822 | 16017 | + | 64 | NO PFAM MATCH | - | - | - |
| WP_150450704.1 | 16013 | 16817 | + | 267 | PF02661<br>TIGR02613 | Fic<br>TIGR02613 | Fic/DOC family<br>mob_myst_B: mobile mystery protein B | 3.70E-12<br>1.10E-07 |
| WP_170285764.1 | 16837 | 17011 | + | 57 | PF18495 | VbhA | Antitoxin VbhA | 1.40E-09 |
| WP_150450707.1 | 17007 | 17586 | + | 192 | PF00239<br>PF08279 | Resolvase<br>HTH_11 | Resolvase, N terminal domain<br>HTH domain | 7.90E-37<br>7.80E-04 |

| index | leader | core | start | end | dir | score |
|---|---|---|---|---|---|---|
| 1 | MDRRKQNRHSSFQRVTTLGGGHLHVPPA | YVPNQGTHPPGLTMNVEVANTSRHRFGLEF | 6296 | 6473 | + | 6 |
| 2 | MSARSQTVHSVHRRKRSLSGRVIRGALRGARNGCRLDIRF | GTIGFGAGFDRLQFRVGAVSRGGGR | 8022 | 8220 | + | 6 |
| 3 | VHRRKRSLSGRVIRGALRGARNGCRLDIRF | GTIGFGAGFDRLQFRVGAVSRGGGR | 8052 | 8220 | + | 6 |
| 4 | MQSEVLEVQEFANQMLRHAEPGMRGDASNPAM | TTPAFVTAAQFVIQGSTICLIC | 10137 | 10302 | + | 32 |
| 5 | MAVVAPGLERPIDVENVMAEIEKGQQLAGHFP | DADALVRARRVLVGEISEEDAMREVRQAFRPA | 15822 | 16017 | + | 6 |
| 6 | MRATVVADAAHSSAMSNLQTGTEYRADA | AAYVDGLIDVDELGRRTRRRYGVPEEDGQ | 16837 | 17011 | + | 6 |

### Results for WP_055482107.1 [Microbispora sp. GMKU363] back to top

previous - next

#### Architecture

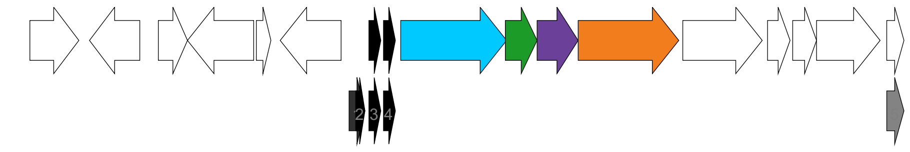

1000 nucleotides

Link to nucleotide sequence

| Accession | start | end | direction | length (aa) | Pfam/HMM | name | description | E-value |
|---|---|---|---|---|---|---|---|---|
| WP_055482206.1 | 50696 | 51617 | + | 306 | TIGR00274<br>PF13580<br>PF01380<br>PF14555<br>TIGR03127 | TIGR00274<br>SIS_2<br>SIS<br>UBA_4<br>TIGR03127 | TIGR00274: N-acetylmuramic acid 6-phosphate etherase<br>SIS domain<br>SIS domain<br>UBA-like domain<br>RuMP_HxlB: 6-phospho 3-hexuloisomerase | 7.20E-107<br>2.20E-10<br>3.40E-07<br>1.10E-06<br>3.60E-04 |
| WP_083502399.1 | 52767 | 51813 | - | 317 | PF01380<br>TIGR03127<br>PF01418<br>TIGR01135<br>TIGR00274 | SIS<br>TIGR03127<br>HTH_6<br>TIGR01135<br>TIGR00274 | SIS domain<br>RuMP_HxlB: 6-phospho 3-hexuloisomerase<br>Helix-turn-helix domain, rpiR family<br>glmS: glutamine-fructose-6-phosphate transaminase (isomerizing)<br>TIGR00274: N-acetylmuramic acid 6-phosphate etherase | 3.20E-19<br>7.00E-17<br>4.90E-15<br>3.30E-10<br>1.40E-09 |
| WP_010988148.1 | 53103 | 53637 | + | 178 | INFERRED GENE | - | - | - |
| WP_055482102.1 | 54887 | 53654 | - | 410 | PF07282<br>PF01385<br>PF12323<br>TIGR01766 | OrfB_Zn_ribbon<br>OrfB_IS605<br>HTH_OrfB_IS605<br>TIGR01766 | Putative transposase DNA-binding domain<br>Probable transposase<br>Helix-turn-helix domain<br>tspaseT_teng_C: transposase, IS605 OrfB family | 4.00E-18<br>2.10E-17<br>2.20E-10<br>1.80E-05 |
| WP_012892167.1 | 54919 | 55195 | + | 92 | INFERRED GENE | - | - | - |
| WP_083502400.1 | 56521 | 55366 | - | 384 | PF00553<br>PF17936 | CBM_2<br>Big_6 | Cellulose binding domain<br>Bacterial Ig domain | 8.80E-26<br>7.40E-04 |
| WP_055482105.1 | 57024 | 57258 | + | 77 | LinA | LinA | LinA (linaridin precursor) | 2.30E-15 |
| WP_055482106.1 | 57292 | 57535 | + | 80 | LinA | LinA | LinA (linaridin precursor) | 4.60E-18 |
| WP_055482107.1 | 57622 | 59593 | + | 656 | LinH<br>LinE<br>LinG<br>PF12697<br>PF00561 | LinH<br>LinE<br>LinG<br>Abhydrolase_6<br>Abhydrolase_1 | LinH (Fusion of LinE/LinG)<br>LinE Homolog<br>LinG Homolog<br>Alpha/beta hydrolase family<br>alpha/beta hydrolase fold | 2.90E-261<br>3.30E-104<br>3.10E-100<br>5.70E-16<br>7.50E-15 |
| WP_055482108.1 | 59589 | 60174 | + | 194 | LinL | LinL | LinL | 1.80E-68 |

| | | | | | | | | |
|---|---|---|---|---|---|---|---|---|
| WP_055482109.1 | 60166 | 60934 | + | 255 | PF13649<br>PF08241<br>PF08242<br>PF13489<br>PF13847 | Methyltransf_25<br>Methyltransf_11<br>Methyltransf_12<br>Methyltransf_23<br>Methyltransf_31 | Methyltransferase domain<br>Methyltransferase domain<br>Methyltransferase domain<br>Methyltransferase domain<br>Methyltransferase domain | 4.50E-15<br>1.90E-10<br>2.10E-08<br>4.80E-08<br>7.20E-08 |
| WP_152990600.1 | 60930 | 62823 | + | 630 | TIGR02204<br>TIGR02203<br>TIGR00958<br>TIGR03796<br>TIGR01846 | TIGR02204<br>TIGR02203<br>TIGR00958<br>TIGR03796<br>TIGR01846 | MsbA_rel: ABC transporter, permease/ATP-binding protein<br>MsbA_lipidA: lipid A export permease/ATP-binding protein MsbA<br>3a01208: antigen peptide transporter 2<br>NHLM_micro_ABC1: NHLM bacteriocin system ABC transporter, peptidase/ATP-binding protein<br>type_I_sec_HlyB: type I secretion system ATPase | 8.90E-107<br>3.90E-98<br>3.70E-92<br>5.20E-88<br>3.20E-85 |
| WP_055482209.1 | 62902 | 64381 | + | 492 | TIGR02734<br>PF13450<br>PF01593<br>TIGR02730<br>TIGR01317 | TIGR02734<br>NAD_binding_8<br>Amino_oxidase<br>TIGR02730<br>TIGR01317 | crtI_fam: phytoene desaturase<br>NAD(P)-binding Rossmann-like domain<br>Flavin containing amine oxidoreductase<br>carot_isom: carotene isomerase<br>GOGAT_sm_gam: glutamate synthase, NADH/NADPH, small subunit | 8.90E-13<br>3.20E-09<br>6.00E-09<br>9.80E-07<br>4.60E-06 |
| WP_055482110.1 | 64464 | 64905 | + | 146 | PF13560<br>PF00046 | HTH_31<br>Homeodomain | Helix-turn-helix domain<br>Homeodomain | 4.50E-06<br>2.20E-04 |
| WP_055482210.1 | 64933 | 65398 | + | 154 | NO PFAM MATCH | - | - | - |
| WP_055482111.1 | 65401 | 66565 | + | 387 | NO PFAM MATCH | - | - | - |
| WP_055482112.1 | 66690 | 67014 | + | 107 | TIGR03433<br>PF03551<br>TIGR02719 | TIGR03433<br>PadR<br>TIGR02719 | padR_acidobact: transcriptional regulator, Acidobacterial, PadR-family<br>Transcriptional regulator PadR-like family<br>repress_PhaQ: poly-beta-hydroxybutyrate-responsive repressor | 2.80E-19<br>2.80E-18<br>1.00E-05 |

| index | leader | core | start | end | dir | score |
|---|---|---|---|---|---|---|
| 1 | VKIPSGSVHHGSRQACGAAAGRPEP...SSLVAPLLSSSSALRCTALDTAPSG | QTLAVTALATGVLS | 56661 | 56970 | + | 9 |
| 2 | VPLPQTFAPSRRPGQHLAALSSSLVAPLLSSSSALRCTALDTAPSG | QTLAVTALATGVLS | 56787 | 56970 | + | 11 |
| 3 | MTSLVDYTNAVSPEMFGDLRETAEGHGLTAASPVL | ATPTAVALGVAIGAVAFHAGYMYGIARNMAEGKYPQPKQLPQ | 57024 | 57258 | + | 30 |
| 4 | MPLLAEKVNADPALGAISIHSTEPCHEIGGAPVL | ATPATIVGVAAGVGLVGSVVGLFAAGYAMGRNFGGTRPPKSEQLPS | 57292 | 57535 | + | 29 |
| 5 | MTEQAFLVLTALVDESRHGYGIAQEVERLSEGRVRLKI | GTLYGVIDRLAADELVALDREEAHR...EAVRLAEKARYATERLRARHAGGTA | 66690 | 67014 | + | 7 |

### Results for TDB95803.1 [Micromonospora fluostatini] back to top

previous - next

#### Architecture

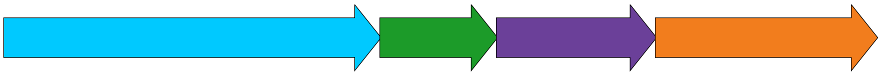

1000 nucleotides

Link to nucleotide sequence

| Accession | start | end | direction | length (aa) | Pfam/HMM | name | description | E-value |
|---|---|---|---|---|---|---|---|---|
| TDB95803.1 | 18 | 1827 | + | 602 | LinH<br>LinE<br>LinG<br>PF12697<br>PF00561 | LinH<br>LinE<br>LinG<br>Abhydrolase_6<br>Abhydrolase_1 | LinH (Fusion of LinE/LinG)<br>LinE Homolog<br>LinG Homolog<br>Alpha/beta hydrolase family<br>alpha/beta hydrolase fold | 1.90E-260<br>2.70E-104<br>3.50E-100<br>1.00E-18<br>2.00E-14 |
| TDB95804.1 | 1823 | 2387 | + | 187 | LinL | LinL | LinL | 3.90E-64 |
| TDB95805.1 | 2379 | 3150 | + | 256 | PF13649<br>PF08241<br>PF13489<br>PF13847<br>PF08242 | Methyltransf_25<br>Methyltransf_11<br>Methyltransf_23<br>Methyltransf_31<br>Methyltransf_12 | Methyltransferase domain<br>Methyltransferase domain<br>Methyltransferase domain<br>Methyltransferase domain<br>Methyltransferase domain | 1.60E-16<br>5.50E-11<br>6.20E-09<br>7.40E-09<br>1.80E-08 |
| TDB95806.1 | 3146 | 4214 | + | 356 | PF00664<br>TIGR02204<br>TIGR03796<br>TIGR00958<br>TIGR02203 | ABC_membrane<br>TIGR02204<br>TIGR03796<br>TIGR00958<br>TIGR02203 | ABC transporter transmembrane region<br>MsbA_rel: ABC transporter, permease/ATP-binding protein<br>NHLM_micro_ABC1: NHLM bacteriocin system ABC transporter, peptidase/ATP-binding protein<br>3a01208: antigen peptide transporter 2<br>MsbA_lipidA: lipid A export permease/ATP-binding protein MsbA | 3.20E-33<br>1.60E-29<br>1.20E-24<br>1.40E-23<br>9.80E-23 |

| index | leader | core | start | end | dir | score |
|---|---|---|---|---|---|---|

### Results for TDC00110.1 [Micromonospora fluostatini] back to top

previous - next

#### Architecture

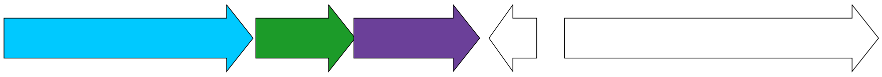

1000 nucleotides

Link to nucleotide sequence

| Accession | start | end | direction | length (aa) | Pfam/HMM | name | description | E-value |
|---|---|---|---|---|---|---|---|---|
| TDC00110.1 | 0 | 1500 | + | 499 | LinH<br>LinE<br>LinG<br>PF00561<br>PF12697 | LinH<br>LinE<br>LinG<br>Abhydrolase_1<br>Abhydrolase_6 | LinH (Fusion of LinE/LinG)<br>LinE Homolog<br>LinG Homolog<br>alpha/beta hydrolase fold<br>Alpha/beta hydrolase family | 7.90E-177<br>7.20E-86<br>3.40E-61<br>3.60E-13<br>7.00E-12 |
| TDC00111.1 | 1521 | 2115 | + | 197 | LinL | LinL | LinL | 5.20E-54 |
| TDC00112.1 | 2111 | 2870 | + | 252 | PF13649<br>PF08241<br>PF13847<br>TIGR02072<br>PF13489 | Methyltransf_25<br>Methyltransf_11<br>Methyltransf_31<br>TIGR02072<br>Methyltransf_23 | Methyltransferase domain<br>Methyltransferase domain<br>Methyltransferase domain<br>BioC: malonyl-acyl carrier protein O-methyltransferase BioC<br>Methyltransferase domain | 1.10E-12<br>5.40E-11<br>1.90E-09<br>3.00E-09<br>7.60E-09 |
| TDC00113.1 | 3213 | 2922 | - | 96 | PF07561 | DUF1540 | Domain of Unknown Function (DUF1540) | 8.30E-09 |
| WP_015038567.1 | 3382 | 5273 | + | 630 | INFERRED GENE | - | - | - |

| index | leader | core | start | end | dir | score |
|---|---|---|---|---|---|---|

### Results for RKN45324.1 [Micromonospora endolithica] back to top

previous - next

#### Architecture

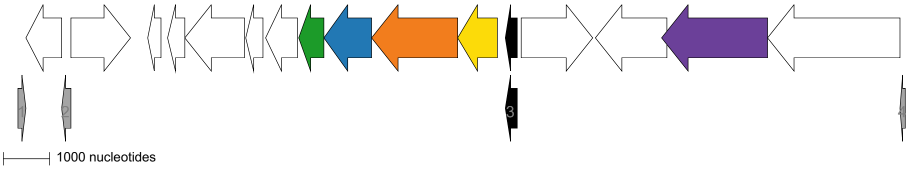

Link to nucleotide sequence

| Accession | start | end | direction | length (aa) | Pfam/HMM | name | description | E-value |
|---|---|---|---|---|---|---|---|---|
| RKN45317.1 | 123870 | 123081 | - | 262 | NO PFAM MATCH | - | - | - |
| RKN45318.1 | 124053 | 125352 | + | 432 | PF05193<br>PF00675<br>TIGR02110<br>TIGR00640 | Peptidase_M16_C<br>Peptidase_M16<br>TIGR02110<br>TIGR00640 | Peptidase M16 inactive domain<br>Insulinase (Peptidase family M16)<br>PQQ_syn_pqqF: coenzyme PQQ biosynthesis protein PqqF<br>acid_CoA_mut_C: methylmalonyl-CoA mutase C-terminal domain | 2.00E-35<br>2.00E-29<br>3.50E-21<br>2.10E-04 |
| RKN45319.1 | 126007 | 125725 | - | 93 | NO PFAM MATCH | - | - | - |
| RKN45320.1 | 126532 | 126145 | - | 128 | NO PFAM MATCH | - | - | - |
| RKN45321.1 | 127818 | 126528 | - | 429 | PF04820<br>TIGR02032<br>PF01494<br>TIGR02023<br>PF01266 | Trp_halogenase<br>TIGR02032<br>FAD_binding_3<br>TIGR02023<br>DAO | Tryptophan halogenase<br>GG-red-SF: geranylgeranyl reductase family<br>FAD binding domain<br>BchP-ChlP: geranylgeranyl reductase<br>FAD dependent oxidoreductase | 2.50E-36<br>5.20E-35<br>4.50E-17<br>3.40E-12<br>7.90E-12 |
| RKN45514.1 | 128215 | 127843 | - | 123 | PF04134 | DUF393 | Protein of unknown function, DUF393 | 1.80E-18 |
| RKN45322.1 | 128963 | 128261 | - | 233 | PF00196<br>TIGR03020<br>TIGR03541<br>PF13384<br>TIGR02937 | GerE<br>TIGR03020<br>TIGR03541<br>HTH_23<br>TIGR02937 | Bacterial regulatory proteins, luxR family<br>EpsA: transcriptional regulator EpsA<br>reg_near_HchA: LuxR family transcriptional regulatory, chaperone HchA-associated<br>Homeodomain-like domain<br>sigma70-ECF: RNA polymerase sigma factor, sigma-70 family | 1.30E-21<br>6.70E-14<br>1.50E-10<br>6.40E-04<br>8.60E-04 |
| RKN45323.1 | 129541 | 128989 | - | 183 | LinL | LinL | LinL | 6.20E-60 |
| RKN45324.1 | 130563 | 129537 | - | 341 | LinE<br>LinH<br>TIGR02427<br>PF00561<br>PF12697 | LinE<br>LinH<br>TIGR02427<br>Abhydrolase_1<br>Abhydrolase_6 | LinE Homolog<br>LinH (Fusion of LinE/LinG)<br>protocat_pcaD: 3-oxoadipate enol-lactonase<br>alpha/beta hydrolase fold<br>Alpha/beta hydrolase family | 1.10E-96<br>1.30E-74<br>6.50E-12<br>7.40E-12<br>2.30E-10 |
| RKN45325.1 | 132429 | 130578 | - | 616 | TIGR02204<br>TIGR02203<br>TIGR00958<br>TIGR03796<br>TIGR01846 | TIGR02204<br>TIGR02203<br>TIGR00958<br>TIGR03796<br>TIGR01846 | MsbA_rel: ABC transporter, permease/ATP-binding protein<br>MsbA_lipidA: lipid A export permease/ATP-binding protein MsbA<br>3a01208: antigen peptide transporter 2<br>NHLM_micro_ABC1: NHLM bacteriocin system ABC transporter, peptidase/ATP-binding protein<br>type_I_sec_HlyB: type I secretion system ATPase | 3.70E-108<br>5.20E-101<br>4.30E-91<br>9.80E-86<br>4.40E-84 |
| RKN45326.1 | 133291 | 132421 | - | 289 | LinG<br>LinH | LinG<br>LinH | LinG Homolog<br>LinH (Fusion of LinE/LinG) | 5.40E-94<br>5.60E-88 |
| RKN45327.1 | 133708 | 133453 | - | 84 | LinA | LinA | LinA (linaridin precursor) | 1.20E-07 |
| RKN45328.1 | 133812 | 135345 | + | 510 | PF00999<br>TIGR00932 | Na_H_Exchanger<br>TIGR00932 | Sodium/hydrogen exchanger family<br>2a37: transporter, monovalent cation:proton antiporter-2 (CPA2) family | 1.80E-48<br>4.60E-40 |
| RKN45329.1 | 136969 | 135409 | - | 519 | TIGR01733<br>PF00501<br>TIGR01734<br>TIGR03098<br>TIGR03443 | TIGR01733<br>AMP-binding<br>TIGR01734<br>TIGR03098<br>TIGR03443 | AA-adenyl-dom: amino acid adenylation domain<br>AMP-binding enzyme<br>D-ala-DACP-lig: D-alanine--poly(phosphoribitol) ligase, subunit 1<br>ligase_PEP_1: acyl-CoA ligase (AMP-forming), exosortase A-associated<br>alpha_am_amid: L-aminoadipate-semialdehyde dehydrogenase | 3.80E-113<br>2.20E-86<br>4.50E-62<br>3.50E-59<br>2.20E-41 |
| RKN45330.1 | 139134 | 136857 | - | 758 | TIGR01733<br>PF00501<br>PF08241<br>PF08242<br>PF13649 | TIGR01733<br>AMP-binding<br>Methyltransf_11<br>Methyltransf_12<br>Methyltransf_25 | AA-adenyl-dom: amino acid adenylation domain<br>AMP-binding enzyme<br>Methyltransferase domain<br>Methyltransferase domain<br>Methyltransferase domain | 1.70E-24<br>5.40E-21<br>1.70E-13<br>9.80E-13<br>6.00E-12 |

| | | | | | | | | |
|---|---|---|---|---|---|---|---|---|
| RKN45331.1 | 142004 | 139130 | - | 957 | TIGR01746<br>PF00668<br>TIGR03443<br>PF07993<br>TIGR03466 | TIGR01746<br>Condensation<br>TIGR03443<br>NAD_binding_4<br>TIGR03466 | Thioester-redct: thioester reductase domain<br>Condensation domain<br>alpha_am_amid: L-aminoadipate-semialdehyde dehydrogenase<br>Male sterility protein<br>HpnA: hopanoid-associated sugar epimerase | 9.80E-75<br>5.30E-65<br>1.60E-54<br>6.80E-48<br>2.90E-09 |

| index | leader | core | start | end | dir | score |
|---|---|---|---|---|---|---|
| 1 | VGWLRTVAPVVGAGNRPRSRPAMGRLLA | DAGLRVVGWRAVSRRGPVPILHAVAATR | 122907 | 123078 | + | 6 |
| 2 | MPPSLPHSTPSHTGCPDGPCPSGDPDR | RCPIGQVGWSAPGNRCRTRIRADGPDRGRTEGRKS | 124055 | 123866 | - | 6 |
| 3 | MEAAAERDIPAVGEGVLEPLELVGVAAPVL | TPFAVGVAVGAGLYAYHKWGGAEET...RLDSETADQMTVSELVQSRRQAVTR | 133708 | 133453 | - | 17 |
| 4 | MFPGGHFFLRTARDELLS | RLAGTLAPIAGPTPLRRRA | 142114 | 142000 | - | 6 |

### Results for QCP08987.1 [Micrococcus luteus] back to top

previous - next

#### Architecture

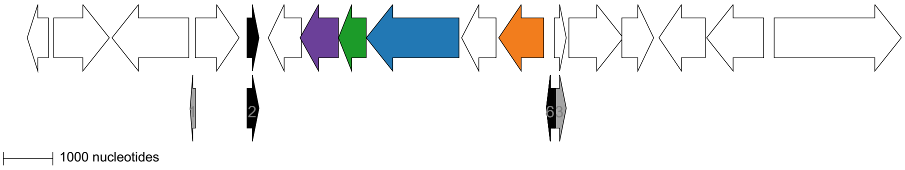

Link to nucleotide sequence

| Accession | start | end | direction | length (aa) | Pfam/HMM | name | description | E-value |
|---|---|---|---|---|---|---|---|---|
| QCP08729.1 | 2739150 | 2738730 | - | 139 | PF02656 | DUF202 | Domain of unknown function (DUF202) | 2.00E-14 |
| QCP08730.1 | 2739255 | 2740386 | + | 376 | PF01988 | VIT1 | VIT family | 1.00E-53 |
| QCP08731.1 | 2741995 | 2740450 | - | 514 | TIGR02734<br>PF13450<br>TIGR02733<br>TIGR03467<br>PF01593 | TIGR02734<br>NAD_binding_8<br>TIGR02733<br>TIGR03467<br>Amino_oxidase | crtI_fam: phytoene desaturase<br>NAD(P)-binding Rossmann-like domain<br>desat_CrtD: C-3',4' desaturase CrtD<br>HpnE: squalene-associated FAD-dependent desaturase<br>Flavin containing amine oxidoreductase | 1.10E-13<br>4.30E-12<br>2.40E-10<br>3.90E-10<br>1.90E-08 |
| QCP08732.1 | 2742116 | 2743022 | + | 301 | PF00144 | Beta-lactamase | Beta-lactamase | 2.60E-24 |
| QCP08733.1 | 2743184 | 2743421 | + | 78 | LinA | LinA | LinA (linaridin precursor) | 1.30E-06 |
| QCP08734.1 | 2744270 | 2743592 | - | 225 | PF00196<br>TIGR03020<br>TIGR03541<br>TIGR02937<br>PF08281 | GerE<br>TIGR03020<br>TIGR03541<br>TIGR02937<br>Sigma70_r4_2 | Bacterial regulatory proteins, luxR family<br>EpsA: transcriptional regulator EpsA<br>reg_near_HchA: LuxR family transcriptional regulatory, chaperone HchA-associated<br>sigma70-ECF: RNA polymerase sigma factor, sigma-70 family<br>Sigma-70, region 4 | 8.10E-15<br>2.00E-10<br>1.10E-09<br>3.30E-05<br>3.90E-05 |
| QCP08735.1 | 2745009 | 2744256 | - | 250 | PF13489<br>PF13649<br>PF08241<br>TIGR01983<br>PF13847 | Methyltransf_23<br>Methyltransf_25<br>Methyltransf_11<br>TIGR01983<br>Methyltransf_31 | Methyltransferase domain<br>Methyltransferase domain<br>Methyltransferase domain<br>UbiG: 3-demethylubiquinone-9 3-O-methyltransferase<br>Methyltransferase domain | 5.50E-11<br>6.20E-11<br>3.50E-10<br>8.80E-08<br>5.20E-07 |
| QCP08736.1 | 2745581 | 2745008 | - | 190 | LinL | LinL | LinL | 6.60E-49 |
| QCP08987.1 | 2747458 | 2745595 | - | 620 | LinE<br>LinH<br>TIGR02427<br>PF00561<br>PF12146 | LinE<br>LinH<br>TIGR02427<br>Abhydrolase_1<br>Hydrolase_4 | LinE Homolog<br>LinH (Fusion of LinE/LinG)<br>protocat_pcaD: 3-oxoadipate enol-lactonase<br>alpha/beta hydrolase fold<br>Serine aminopeptidase, S33 | 1.00E-52<br>4.50E-42<br>7.10E-12<br>1.50E-11<br>1.70E-07 |
| QCP08737.1 | 2748208 | 2747500 | - | 235 | PF01061 | ABC2_membrane | ABC-2 type transporter | 1.10E-05 |
| QCP08738.1 | 2749175 | 2748254 | - | 306 | TIGR01188<br>TIGR03864<br>TIGR01288<br>TIGR03265<br>TIGR00968 | TIGR01188<br>TIGR03864<br>TIGR01288<br>TIGR03265<br>TIGR00968 | drrA: daunorubicin resistance ABC transporter, ATP-binding protein<br>PQQ_ABC_ATP: ABC transporter, ATP-binding subunit, PQQ-dependent alcohol dehydrogenase system<br>nodI: nodulation ABC transporter NodI<br>PhnT2: putative 2-aminoethylphosphonate ABC transporter, ATP-binding protein<br>3a0106s01: sulfate ABC transporter, ATP-binding protein | 3.30E-63<br>1.70E-47<br>2.80E-44<br>7.80E-42<br>1.50E-39 |
| QCP08739.1 | 2749386 | 2749635 | + | 82 | NO PFAM MATCH | - | - | - |
| QCP08740.1 | 2749675 | 2750785 | + | 369 | PF07730 | HisKA_3 | Histidine kinase | 7.80E-11 |
| QCP08741.1 | 2750754 | 2751390 | + | 211 | PF00072<br>TIGR02875<br>TIGR01818<br>TIGR02154<br>PF00196 | Response_reg<br>TIGR02875<br>TIGR01818<br>TIGR02154<br>GerE | Response regulator receiver domain<br>spore_0_A: sporulation transcription factor Spo0A<br>ntrC: nitrogen regulation protein NR(I)<br>PhoB: phosphate regulon transcriptional regulatory protein PhoB<br>Bacterial regulatory proteins, luxR family | 3.90E-27<br>2.40E-18<br>3.40E-15<br>8.30E-13<br>1.90E-12 |
| QCP08742.1 | 2752429 | 2751508 | - | 306 | TIGR01019<br>PF02629<br>TIGR02717<br>PF00549<br>PF13607 | TIGR01019<br>CoA_binding<br>TIGR02717<br>Ligase_CoA<br>Succ_CoA_lig | sucCoAalpha: succinate-CoA ligase, alpha subunit<br>CoA binding domain<br>AcCoA-syn-alpha: acetyl coenzyme A synthetase (ADP forming), alpha domain<br>CoA-ligase<br>Succinyl-CoA ligase like flavodoxin domain | 2.10E-126<br>8.10E-25<br>1.40E-24<br>5.00E-15<br>2.50E-12 |

| | | | | | | | | |
|---|---|---|---|---|---|---|---|---|
| QCP08743.1 | 2753626 | 2752456 | - | 389 | TIGR01016<br>PF08442<br>PF00549<br>PF13549<br>TIGR00877 | TIGR01016<br>ATP-grasp_2<br>Ligase_CoA<br>ATP-grasp_5<br>TIGR00877 | sucCoAbeta: succinate-CoA ligase, beta subunit<br>ATP-grasp domain<br>CoA-ligase<br>ATP-grasp domain<br>purD: phosphoribosylamine--glycine ligase | 1.60E-125<br>3.20E-69<br>1.30E-12<br>3.20E-10<br>4.80E-07 |
| WP_017488278.1 | 2753828 | 2756410 | + | 860 | INFERRED GENE | - | - | - |

| index | leader | core | start | end | dir | score |
|---|---|---|---|---|---|---|
| 1 | MAILLEVGRDQHRHSVC | LTDRWHRSSRGLAWAEAA | 2742118 | 2742010 | - | 6 |
| 2 | MSTIAQFADAAALPYGAKADMAAEPTL | ATIATTATTALTCITAAFGHHDSER...DVMSTVEADASVDTMVAARRDALQG | 2743184 | 2743421 | + | 17 |
| 3 | MKTIRTIAFILFLCATAFGVMLTLHYLPIPEILPSEVTSWVPAR | DTADATWLIVGMVVAVVATAVGFSASHKAVERETQVTS | 2749386 | 2749635 | + | 6 |
| 4 | MGASRVQVDGWFSSAVPIRRGCRGIGTTSTFPSPAR | PAPVTQRHACAPTLSCRAVT | 2749388 | 2749217 | - | 9 |
| 5 | MVFMGASRVQVDGWFSSAVPIRRGCRGIGTTSTFPSPAR | PAPVTQRHACAPTLSCRAVT | 2749397 | 2749217 | - | 9 |
| 6 | MVRMVFMGASRVQVDGWFSSAVPIRRGCRGIGTTSTFPSPAR | PAPVTQRHACAPTLSCRAVT | 2749406 | 2749217 | - | 9 |

### Results for RQW94077.1 [Micromonospora inaquosa] back to top

previous - next

#### Architecture

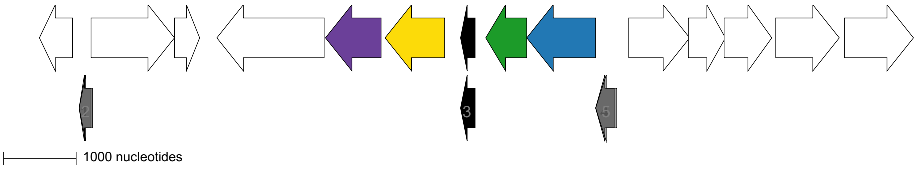

Link to nucleotide sequence

| Accession | start | end | direction | length (aa) | Pfam/HMM | name | description | E-value |
|---|---|---|---|---|---|---|---|---|
| RQW94082.1 | 24975 | 24516 | - | 152 | PF04978<br>PF12867<br>PF11716<br>TIGR03083 | DUF664<br>DinB_2<br>MDMPI_N<br>TIGR03083 | Protein of unknown function (DUF664)<br>DinB superfamily<br>Mycothiol maleylpyruvate isomerase N-terminal domain<br>TIGR03083: uncharacterized Actinobacterial protein TIGR03083 | 7.70E-36<br>2.10E-11<br>1.60E-08<br>3.00E-05 |
| RQW94071.1 | 25236 | 26385 | + | 382 | PF13191 | AAA_16 | AAA ATPase domain | 7.10E-04 |
| RQW94072.1 | 26381 | 26732 | + | 116 | NO PFAM MATCH | - | - | - |
| RQW94073.1 | 28439 | 26966 | - | 490 | TIGR02734<br>TIGR00292<br>TIGR03467<br>TIGR02733<br>TIGR01317 | TIGR02734<br>TIGR00292<br>TIGR03467<br>TIGR02733<br>TIGR01317 | crtI_fam: phytoene desaturase<br>TIGR00292: thiazole biosynthesis enzyme<br>HpnE: squalene-associated FAD-dependent desaturase<br>desat_CrtD: C-3',4' desaturase CrtD<br>GOGAT_sm_gam: glutamate synthase, NADH/NADPH, small subunit | 5.60E-09<br>2.20E-08<br>2.60E-08<br>8.60E-08<br>2.10E-07 |
| RQW94074.1 | 29220 | 28464 | - | 251 | PF13649<br>PF08241<br>PF08242<br>PF13489<br>TIGR01983 | Methyltransf_25<br>Methyltransf_11<br>Methyltransf_12<br>Methyltransf_23<br>TIGR01983 | Methyltransferase domain<br>Methyltransferase domain<br>Methyltransferase domain<br>Methyltransferase domain<br>UbiG: 3-demethylubiquinone-9 3-O-methyltransferase | 8.40E-14<br>2.50E-09<br>4.00E-08<br>6.90E-08<br>8.90E-06 |
| RQW94083.1 | 30113 | 29279 | - | 277 | LinG<br>LinH<br>TIGR04033 | LinG<br>LinH<br>TIGR04033 | LinG Homolog<br>LinH (Fusion of LinE/LinG)<br>export_SdpB: antimicrobial peptide system protein, SdpB family | 3.50E-102<br>1.30E-75<br>3.20E-04 |
| RQW94075.1 | 30520 | 30319 | - | 66 | LinA | LinA | LinA (linaridin precursor) | 1.70E-12 |
| RQW94076.1 | 31244 | 30680 | - | 187 | LinL | LinL | LinL | 2.60E-55 |
| RQW94077.1 | 32194 | 31240 | - | 317 | LinE<br>LinH<br>PF12697<br>PF00561<br>TIGR02427 | LinE<br>LinH<br>Abhydrolase_6<br>Abhydrolase_1<br>TIGR02427 | LinE Homolog<br>LinH (Fusion of LinE/LinG)<br>Alpha/beta hydrolase family<br>alpha/beta hydrolase fold<br>protocat_pcaD: 3-oxoadipate enol-lactonase | 1.70E-91<br>7.10E-43<br>3.60E-16<br>7.70E-09<br>6.80E-08 |
| RQW94078.1 | 32644 | 33475 | + | 276 | NO PFAM MATCH | - | - | - |
| RQW94079.1 | 33467 | 33968 | + | 166 | TIGR02983<br>TIGR02937<br>TIGR02954<br>TIGR02985<br>PF08281 | TIGR02983<br>TIGR02937<br>TIGR02954<br>TIGR02985<br>Sigma70_r4_2 | SigE-fam_strep: RNA polymerase sigma-70 factor, sigma-E family<br>sigma70-ECF: RNA polymerase sigma factor, sigma-70 family<br>Sig70_famx3: RNA polymerase sigma-70 factor, TIGR02954 family<br>Sig70_bacteroi1: RNA polymerase sigma-70 factor, Bacteroides expansion family 1<br>Sigma-70, region 4 | 6.60E-47<br>6.60E-20<br>1.20E-16<br>3.50E-12<br>1.70E-11 |
| RQW94080.1 | 33964 | 34624 | + | 219 | NO PFAM MATCH | - | - | - |
| RQW94081.1 | 34673 | 35540 | + | 288 | PF13354<br>PF00768 | Beta-lactamase2<br>Peptidase_S11 | Beta-lactamase enzyme family<br>D-alanyl-D-alanine carboxypeptidase | 1.20E-08<br>3.50E-04 |
| WP_003083047.1 | 35616 | 36567 | + | 317 | INFERRED GENE | - | - | - |

| index | leader | core | start | end | dir | score |
|---|---|---|---|---|---|---|
| 1 | VRILRKQAHRNAPPTVRPIAASAEGSGRAADASNRQAVP | VTVPGECRSRLLTCRIR | 25236 | 25065 | - | 6 |
| 2 | LHVRILRKQAHRNAPPTVRPIAASAEGSGRAADASNRQAVP | VTVPGECRSRLLTCRIR | 25242 | 25065 | - | 6 |
| 3 | MSLVAQFANESLTDVSPGMLSETDAAARLDTPADPVM | ATPLIVETVFLATGMLAANIANYVDRNNY | 30520 | 30319 | - | 30 |

| | | | | | | |
|---|---|---|---|---|---|---|
| 4 | VRASATRTGEIGGGRLSPGSPATCS...SDLFRPVPYSDFPGPIEACWDRLER | RTTVGDLPVGAAAIWPKSGPALGAYEER | 32436 | 32190 | - | 6 |
| 5 | LLTRRVGQPEPTVRASATRTGEIGG...SDLFRPVPYSDFPGPIEACWDRLER | RTTVGDLPVGAAAIWPKSGPALGAYEER | 32472 | 32190 | - | 6 |

### Results for TWH68587.1 [Micromonospora olivasterospora] back to top

previous - next

#### Architecture

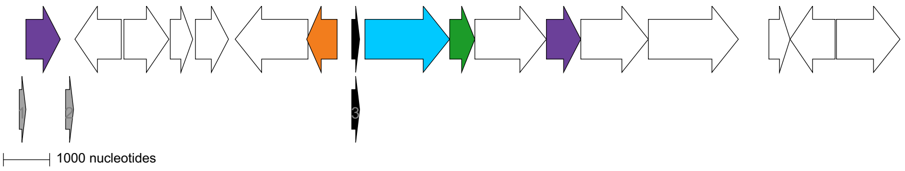

Link to nucleotide sequence

| Accession | start | end | direction | length (aa) | Pfam/HMM | name | description | E-value |
|---|---|---|---|---|---|---|---|---|
| TWH68579.1 | 3853994 | 3854750 | + | 251 | PF13649<br>PF08241<br>PF13489<br>PF08242<br>TIGR00537 | Methyltransf_25<br>Methyltransf_11<br>Methyltransf_23<br>Methyltransf_12<br>TIGR00537 | Methyltransferase domain<br>Methyltransferase domain<br>Methyltransferase domain<br>Methyltransferase domain<br>hemK_rel_arch: putative methylase | 4.90E-12<br>2.80E-11<br>2.30E-07<br>1.50E-05<br>5.90E-05 |
| TWH68580.1 | 3856042 | 3855052 | - | 329 | PF00248<br>TIGR01293 | Aldo_ket_red<br>TIGR01293 | Aldo/keto reductase family<br>Kv_beta: voltage-dependent potassium channel beta subunit | 6.30E-66<br>6.00E-49 |
| TWH68581.1 | 3856125 | 3857094 | + | 322 | PF00248<br>TIGR01293 | Aldo_ket_red<br>TIGR01293 | Aldo/keto reductase family<br>Kv_beta: voltage-dependent potassium channel beta subunit | 1.70E-59<br>1.00E-15 |
| TWH68582.1 | 3857101 | 3857599 | + | 165 | PF01814 | Hemerythrin | Hemerythrin HHE cation binding domain | 1.20E-10 |
| TWH68583.1 | 3857666 | 3858371 | + | 234 | NO PFAM MATCH | - | - | - |
| TWH68584.1 | 3860085 | 3858516 | - | 522 | NO PFAM MATCH | - | - | - |
| TWH68585.1 | 3860722 | 3860071 | - | 216 | TIGR01189<br>TIGR02857<br>TIGR00968<br>TIGR03873<br>TIGR01277 | TIGR01189<br>TIGR02857<br>TIGR00968<br>TIGR03873<br>TIGR01277 | ccmA: heme ABC exporter, ATP-binding protein CcmA<br>CydD: thiol reductant ABC exporter, CydD subunit<br>3a0106s01: sulfate ABC transporter, ATP-binding protein<br>F420-0_ABC_ATP: proposed F420-0 ABC transporter, ATP-binding protein<br>thiQ: thiamine ABC transporter, ATP-binding protein | 4.60E-34<br>3.70E-31<br>2.40E-30<br>1.80E-28<br>1.90E-28 |
| TWH68586.1 | 3861042 | 3861210 | + | 55 | LinA | LinA | LinA (linaridin precursor) | 5.30E-10 |
| TWH68587.1 | 3861307 | 3863149 | + | 613 | LinH<br>LinE<br>LinG<br>PF12697<br>PF12146 | LinH<br>LinE<br>LinG<br>Abhydrolase_6<br>Hydrolase_4 | LinH (Fusion of LinE/LinG)<br>LinE Homolog<br>LinG Homolog<br>Alpha/beta hydrolase family<br>Serine aminopeptidase, S33 | 8.50E-225<br>6.80E-99<br>5.90E-95<br>5.30E-15<br>2.70E-09 |
| TWH68588.1 | 3863145 | 3863691 | + | 181 | LinL | LinL | LinL | 2.20E-55 |
| TWH68589.1 | 3863687 | 3865226 | + | 512 | PF13450<br>TIGR02733<br>TIGR03467<br>PF01593<br>TIGR02734 | NAD_binding_8<br>TIGR02733<br>TIGR03467<br>Amino_oxidase<br>TIGR02734 | NAD(P)-binding Rossmann-like domain<br>desat_CrtD: C-3',4' desaturase CrtD<br>HpnE: squalene-associated FAD-dependent desaturase<br>Flavin containing amine oxidoreductase<br>crtI_fam: phytoene desaturase | 2.80E-11<br>5.50E-11<br>1.10E-10<br>1.40E-09<br>1.90E-09 |
| TWH68590.1 | 3865222 | 3865972 | + | 249 | PF13649<br>PF08241<br>TIGR02021<br>PF13489<br>PF08242 | Methyltransf_25<br>Methyltransf_11<br>TIGR02021<br>Methyltransf_23<br>Methyltransf_12 | Methyltransferase domain<br>Methyltransferase domain<br>BchM-ChlM: magnesium protoporphyrin O-methyltransferase<br>Methyltransferase domain<br>Methyltransferase domain | 7.40E-12<br>1.70E-09<br>6.40E-09<br>3.40E-08<br>3.70E-08 |
| TWH68591.1 | 3865982 | 3867434 | + | 483 | TIGR02030<br>TIGR02031<br>TIGR02442<br>PF01078<br>PF00158 | TIGR02030<br>TIGR02031<br>TIGR02442<br>Mg_chelatase<br>Sigma54_activat | BchI-ChlI: magnesium chelatase ATPase subunit I<br>BchD-ChlD: magnesium chelatase ATPase subunit D<br>Cob-chelat-sub: cobaltochelatase subunit<br>Magnesium chelatase, subunit ChlI<br>Sigma-54 interaction domain | 5.40E-10<br>8.70E-10<br>6.40E-08<br>7.40E-07<br>1.10E-05 |
| TWH68592.1 | 3867436 | 3869389 | + | 650 | PF13519<br>TIGR02442 | VWA_2<br>TIGR02442 | von Willebrand factor type A domain<br>Cob-chelat-sub: cobaltochelatase subunit | 2.10E-08<br>4.00E-08 |
| TWH68593.1 | 3870031 | 3870499 | + | 155 | NO PFAM MATCH | - | - | - |
| TWH68594.1 | 3871484 | 3870509 | - | 324 | PF01965<br>PF12833<br>PF00165<br>TIGR04094<br>TIGR01383 | DJ-1_PfpI<br>HTH_18<br>HTH_AraC<br>TIGR04094<br>TIGR01383 | DJ-1/PfpI family<br>Helix-turn-helix domain<br>Bacterial regulatory helix-turn-helix proteins, AraC family<br>adjacent_YSIRK: YSIRK-targeted surface antigen transcriptional regulator<br>not_thiJ: DJ-1 family protein | 1.90E-22<br>3.10E-21<br>7.30E-18<br>5.30E-10<br>1.30E-08 |
| TWH68595.1 | 3871503 | 3872871 | + | 455 | PF07690<br>TIGR00892<br>TIGR00890<br>TIGR00893<br>TIGR04259 | MFS_1<br>TIGR00892<br>TIGR00890<br>TIGR00893<br>TIGR04259 | Major Facilitator Superfamily<br>2A0113: monocarboxylate transporter<br>2A0111: oxalate/formate antiporter family transporter<br>2A0114: D-galactonate transporter<br>oxa_formateAnti: oxalate/formate antiporter | 4.80E-33<br>4.20E-17<br>1.10E-16<br>7.60E-16<br>6.80E-15 |

| index | leader | core | start | end | dir | score |
|---|---|---|---|---|---|---|
| 1 | LRKSPGAGSNPPTRAFLHKSGTIRPV | LAVWATRAPIPLMNGAISGSRAGSRP | 3853839 | 3853998 | + | 6 |
| 2 | LKRPGEGHDSRQGVCRVDQTVPVGRWC | AGLLLLRRGPTIHPGPTPGAGSGVPGGL | 3854855 | 3855023 | + | 6 |
| 3 | MRNVARHANRVSTARFGSLTAGAPAKKMADSAPVL | FTPAVAVLGAAALITAVGSI | 3861042 | 3861210 | + | 29 |

### Results for KXK61181.1 [Micromonospora rosaria] back to top

previous - next

#### Architecture

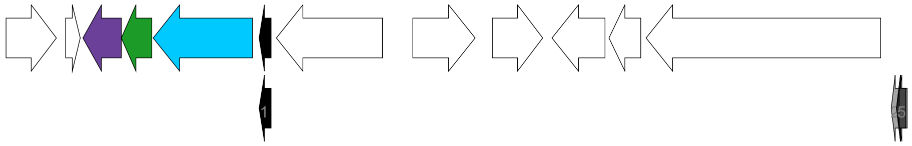

1000 nucleotides

Link to nucleotide sequence

| Accession | start | end | direction | length (aa) | Pfam/HMM | name | description | E-value |
|---|---|---|---|---|---|---|---|---|
| AWW66_14800 | 0 | 986 | + | 328 | INFERRED GENE | - | - | - |
| KXK61178.1 | 1156 | 1447 | + | 96 | PF07561 | DUF1540 | Domain of Unknown Function (DUF1540) | 7.30E-09 |
| KXK61179.1 | 2257 | 1498 | - | 252 | PF13649<br>PF08241<br>PF13489<br>TIGR02072<br>PF13847 | Methyltransf_25<br>Methyltransf_11<br>Methyltransf_23<br>TIGR02072<br>Methyltransf_31 | Methyltransferase domain<br>Methyltransferase domain<br>Methyltransferase domain<br>BioC: malonyl-acyl carrier protein O-methyltransferase BioC<br>Methyltransferase domain | 9.70E-13<br>4.80E-11<br>2.40E-09<br>3.80E-09<br>4.40E-09 |
| KXK61180.1 | 2847 | 2253 | - | 197 | LinL | LinL | LinL | 2.10E-54 |
| KXK61181.1 | 4822 | 2869 | - | 650 | LinH<br>LinG<br>LinE<br>PF00561<br>PF12697 | LinH<br>LinG<br>LinE<br>Abhydrolase_1<br>Abhydrolase_6 | LinH (Fusion of LinE/LinG)<br>LinG Homolog<br>LinE Homolog<br>alpha/beta hydrolase fold<br>Alpha/beta hydrolase family | 5.40E-223<br>1.10E-95<br>1.70E-87<br>7.50E-13<br>1.90E-12 |
| KXK61182.1 | 5177 | 4940 | - | 78 | LinA | LinA | LinA (linaridin precursor) | 1.60E-09 |
| KXK61183.1 | 7360 | 5272 | - | 695 | PF03704<br>PF13191<br>PF00486<br>TIGR02928 | BTAD<br>AAA_16<br>Trans_reg_C<br>TIGR02928 | Bacterial transcriptional activator domain<br>AAA ATPase domain<br>Transcriptional regulatory protein, C terminal<br>TIGR02928: orc1/cdc6 family replication initiation protein | 5.60E-34<br>1.50E-10<br>1.10E-07<br>1.00E-04 |
| KXK61184.1 | 7942 | 9160 | + | 405 | PF03704<br>PF00486 | BTAD<br>Trans_reg_C | Bacterial transcriptional activator domain<br>Transcriptional regulatory protein, C terminal | 5.40E-31<br>1.60E-04 |
| WP_015621840.1 | 9492 | 10473 | + | 327 | INFERRED GENE | - | - | - |
| KXK61185.1 | 11703 | 10662 | - | 346 | PF06280 | fn3_5 | Fn3-like domain | 5.40E-04 |
| KXK61186.1 | 12392 | 11768 | - | 207 | NO PFAM MATCH | - | - | - |
| KXK61206.1 | 17075 | 12497 | - | 1525 | PF00754<br>PF07532 | F5_F8_type_C<br>Big_4 | F5/8 type C domain<br>Bacterial Ig-like domain (group 4) | 1.00E-11<br>5.10E-11 |

| index | leader | core | start | end | dir | score |
|---|---|---|---|---|---|---|
| 1 | MSDPLASVNSSRYLKTVPVRAERSTWLGRKDGSPVM | LLPTIVTPAIVTGTVNVATQVVAQVTGVRQPDPNATDPSNLA | 5177 | 4940 | - | 16 |
| 2 | VGQPGALDDAGLAADDRLTAAGRRGGRH | RHMLTLTDGNGQLGPARAARVTTPEVPGR | 17451 | 17277 | - | 6 |
| 3 | VRCRRIRPREDEWPYR | GTPAPPDPVDADLDEVALVAYHAWANRGPSTMRVWLPTTD | 17567 | 17396 | - | 6 |
| 4 | VTVRCRRIRPREDEWPYR | GTPAPPDPVDADLDEVALVAYHAWANRGPSTMRVWLPTTD | 17573 | 17396 | - | 6 |
| 5 | VVDGRVTVRCRRIRPREDEWPYR | GTPAPPDPVDADLDEVALVAYHAWANRGPSTMRVWLPTTD | 17588 | 17396 | - | 6 |

### Results for KXK61316.1 [Micromonospora rosaria] back to top

previous - next

#### Architecture

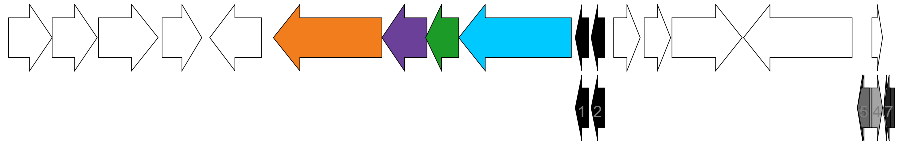

1000 nucleotides

Link to nucleotide sequence

| Accession | start | end | direction | length (aa) | Pfam/HMM | name | description | E-value |
|---|---|---|---|---|---|---|---|---|
| KXK61326.1 | 30357 | 31098 | + | 246 | PF00528 | BPD_transp_1 | Binding-protein-dependent transport system inner membrane component | 2.70E-11 |
| KXK61310.1 | 31094 | 31874 | + | 259 | PF00528<br>TIGR01183<br>TIGR03416<br>TIGR01097<br>TIGR03226 | BPD_transp_1<br>TIGR01183<br>TIGR03416<br>TIGR01097<br>TIGR03226 | Binding-protein-dependent transport system inner membrane component<br>ntrB: nitrate ABC transporter, permease protein<br>ABC_choXWV_perm: choline ABC transporter, permease protein<br>PhnE: phosphonate ABC transporter, permease protein PhnE<br>PhnU: 2-aminoethylphosphonate ABC transporter, permease protein | 4.50E-23<br>6.50E-11<br>1.50E-05<br>4.50E-04<br>6.80E-04 |
| WP_011607820.1 | 31896 | 32919 | + | 341 | INFERRED GENE | - | - | - |
| KXK61311.1 | 32990 | 33665 | + | 224 | PF02698 | DUF218 | DUF218 domain | 2.60E-21 |

| | | | | | | | | |
|---|---|---|---|---|---|---|---|---|
| KXK61312.1 | 34685 | 33812 | - | 290 | NO PFAM MATCH | - | - | - |
| KXK61313.1 | 36764 | 34901 | - | 620 | TIGR02204<br>TIGR02203<br>TIGR00958<br>TIGR03796<br>TIGR01846 | TIGR02204<br>TIGR02203<br>TIGR00958<br>TIGR03796<br>TIGR01846 | MsbA_rel: ABC transporter, permease/ATP-binding protein<br>MsbA_lipidA: lipid A export permease/ATP-binding protein MsbA<br>3a01208: antigen peptide transporter 2<br>NHLM_micro_ABC1: NHLM bacteriocin system ABC transporter, peptidase/ATP-binding protein<br>type_I_sec_HlyB: type I secretion system ATPase | 7.00E-101<br>9.70E-95<br>3.90E-92<br>4.40E-87<br>4.00E-83 |
| KXK61314.1 | 37531 | 36760 | - | 256 | PF13649<br>PF08241<br>PF13489<br>PF13847<br>PF08242 | Methyltransf_25<br>Methyltransf_11<br>Methyltransf_23<br>Methyltransf_31<br>Methyltransf_12 | Methyltransferase domain<br>Methyltransferase domain<br>Methyltransferase domain<br>Methyltransferase domain<br>Methyltransferase domain | 6.40E-17<br>1.90E-11<br>3.00E-09<br>5.40E-09<br>1.90E-08 |
| KXK61315.1 | 38087 | 37523 | - | 187 | LinL | LinL | LinL | 1.30E-64 |
| KXK61316.1 | 40033 | 38083 | - | 649 | LinH<br>LinE<br>LinG<br>PF12697<br>PF00561 | LinH<br>LinE<br>LinG<br>Abhydrolase_6<br>Abhydrolase_1 | LinH (Fusion of LinE/LinG)<br>LinE Homolog<br>LinG Homolog<br>Alpha/beta hydrolase family<br>alpha/beta hydrolase fold | 1.40E-262<br>2.70E-104<br>3.60E-101<br>1.60E-18<br>8.80E-15 |
| KXK61317.1 | 40315 | 40084 | - | 76 | LinA | LinA | LinA (linaridin precursor) | 1.30E-15 |
| KXK61318.1 | 40588 | 40366 | - | 73 | LinA | LinA | LinA (linaridin precursor) | 4.90E-12 |
| KXK61319.1 | 40744 | 41215 | + | 156 | PF13560 | HTH_31 | Helix-turn-helix domain | 9.00E-06 |
| KXK61327.1 | 41265 | 41727 | + | 153 | NO PFAM MATCH | - | - | - |
| KXK61320.1 | 41745 | 42954 | + | 402 | NO PFAM MATCH | - | - | - |
| KXK61321.1 | 44852 | 42977 | - | 624 | NO PFAM MATCH | - | - | - |
| KXK61322.1 | 45178 | 45382 | + | 67 | NO PFAM MATCH | - | - | - |

| index | leader | core | start | end | dir | score |
|---|---|---|---|---|---|---|
| 1 | MSVLATRVNTDPDLAAIPIDSGEACSQVGGMPVL | ATPATIVSVAAGVGLVGSVAALFGAGYVAGRNLGGTRPPVQN | 40315 | 40084 | - | 29 |
| 2 | MTALVAHTNEISAEMFGELTEDRTAEGLAGLTPVL | ATPTAVALGVAVGAVAFHAGYMVGIARNMAEGTRPRPQ | 40588 | 40366 | - | 29 |
| 3 | MGGRQSAGGNRLSTPRRRNSRSCVAGSNVER | YPVEARLVYPFIESPSSRDERLGDTATGVAP | 45138 | 44949 | - | 6 |
| 4 | MRRNRRGSGSAYDDEPRHRPLLDLTALVVLALAGGAGWLAYREGGWPVA | LTVGFAVAAGLHRLLRGG | 45178 | 45382 | + | 6 |
| 5 | MKLLPVPVLREFLGMGGRQSAGGNRLSTPRRRNSRSCV | AGSNVERYPVEARLVYPFIESPSSRDERLGDTATGVAP | 45180 | 44949 | - | 6 |
| 6 | VLRDIDPATAEADRPAGGGLP | ETAYLTHTNDPDVGAPMNPPR | 45515 | 45386 | - | 6 |
| 7 | VDGVDEAGNSWNGPGWGTQVLRDIDP | ATAEADRPAGGGLPETAYLTHTNDPDVGAPMNPPR | 45572 | 45386 | - | 9 |

### Results for WP_148426628.1 [Micromonospora sp. AP08] back to top

previous - next

#### Architecture

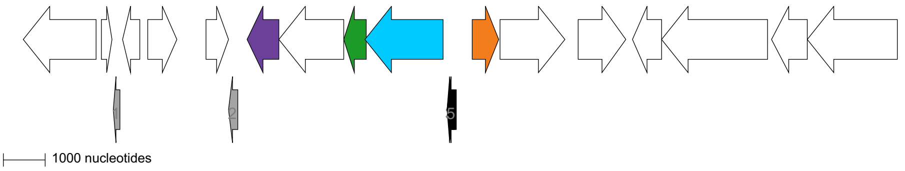

Link to nucleotide sequence

| Accession | start | end | direction | length (aa) | Pfam/HMM | name | description | E-value |
|---|---|---|---|---|---|---|---|---|
| WP_148426621.1 | 402054 | 400311 | - | 580 | NO PFAM MATCH | - | - | - |
| WP_148426622.1 | 402177 | 402441 | + | 87 | PF01022<br>PF12840 | HTH_5<br>HTH_20 | Bacterial regulatory protein, arsR family<br>Helix-turn-helix domain | 6.70E-08<br>1.40E-06 |
| WP_148426623.1 | 403096 | 402688 | - | 135 | NO PFAM MATCH | - | - | - |
| WP_148426624.1 | 403292 | 403970 | + | 225 | NO PFAM MATCH | - | - | - |
| WP_148426665.1 | 404683 | 405220 | + | 178 | PF01551 | Peptidase_M23 | Peptidase family M23 | 4.00E-34 |
| WP_148426625.1 | 406426 | 405670 | - | 251 | PF13649<br>PF08241<br>PF13489<br>PF08242<br>PF13847 | Methyltransf_25<br>Methyltransf_11<br>Methyltransf_23<br>Methyltransf_12<br>Methyltransf_31 | Methyltransferase domain<br>Methyltransferase domain<br>Methyltransferase domain<br>Methyltransferase domain<br>Methyltransferase domain | 3.30E-13<br>4.20E-11<br>3.50E-10<br>8.20E-10<br>1.00E-07 |
| WP_148426626.1 | 407973 | 406422 | - | 516 | PF13450<br>TIGR02734<br>TIGR02733<br>TIGR03467<br>TIGR00292 | NAD_binding_8<br>TIGR02734<br>TIGR02733<br>TIGR03467<br>TIGR00292 | NAD(P)-binding Rossmann-like domain<br>crtI_fam: phytoene desaturase<br>desat_CrtD: C-3',4' desaturase CrtD<br>HpnE: squalene-associated FAD-dependent desaturase<br>TIGR00292: thiazole biosynthesis enzyme | 1.40E-11<br>8.30E-11<br>1.00E-10<br>4.90E-09<br>3.90E-08 |
| WP_148426627.1 | 408512 | 407969 | - | 180 | LinL | LinL | LinL | 6.60E-56 |
| WP_148426628.1 | 410352 | 408492 | - | 619 | LinH<br>LinE<br>LinG<br>PF12697<br>TIGR02427 | LinH<br>LinE<br>LinG<br>Abhydrolase_6<br>TIGR02427 | LinH (Fusion of LinE/LinG)<br>LinE Homolog<br>LinG Homolog<br>Alpha/beta hydrolase family<br>protocat_pcaD: 3-oxoadipate enol-lactonase | 7.40E-234<br>1.10E-106<br>2.50E-97<br>3.90E-13<br>2.00E-12 |

| | | | | | | | | |
|---|---|---|---|---|---|---|---|---|
| WP_148426629.1 | 411032 | 411683 | + | 216 | TIGR01189 | TIGR01189 | ccmA: heme ABC exporter, ATP-binding protein CcmA | 1.10E-33 |
| | | | | | TIGR02857 | TIGR02857 | CydD: thiol reductant ABC exporter, CydD subunit | 1.90E-30 |
| | | | | | TIGR00968 | TIGR00968 | 3a0106s01: sulfate ABC transporter, ATP-binding protein | 1.10E-27 |
| | | | | | TIGR03740 | TIGR03740 | galliderm_ABC: lantibiotic protection ABC transporter, ATP-binding subunit | 4.70E-27 |
| | | | | | PF00005 | ABC_tran | ABC transporter | 4.00E-26 |
| WP_148426630.1 | 411723 | 413253 | + | 509 | NO PFAM MATCH | - | - | - |
| WP_014692506.1 | 413589 | 414708 | + | 373 | INFERRED GENE | - | - | - |
| WP_148426631.1 | 415578 | 414879 | - | 232 | TIGR02154 | TIGR02154 | PhoB: phosphate regulon transcriptional regulatory protein PhoB | 5.00E-47 |
| | | | | | TIGR01387 | TIGR01387 | cztR_silR_copR: heavy metal response regulator | 4.90E-39 |
| | | | | | TIGR03787 | TIGR03787 | marine_sort_RR: proteobacterial dedicated sortase system response regulator | 2.40E-29 |
| | | | | | PF00072 | Response_reg | Response regulator receiver domain | 1.80E-28 |
| | | | | | PF00486 | Trans_reg_C | Transcriptional regulatory protein, C terminal | 2.60E-22 |
| WP_148426632.1 | 418130 | 415574 | - | 851 | PF02702 | KdpD | Osmosensitive K+ channel His kinase sensor domain | 3.70E-101 |
| | | | | | TIGR02956 | TIGR02956 | TMAO_torS: TMAO reductase sytem sensor TorS | 2.30E-30 |
| | | | | | PF13493 | DUF4118 | Domain of unknown function (DUF4118) | 9.50E-29 |
| | | | | | TIGR02966 | TIGR02966 | phoR_proteo: phosphate regulon sensor kinase PhoR | 1.10E-26 |
| | | | | | TIGR01386 | TIGR01386 | cztS_silS_copS: heavy metal sensor kinase | 1.80E-24 |
| WP_148426633.1 | 419080 | 418198 | - | 293 | PF02669 | KdpC | K+-transporting ATPase, c chain | 6.80E-59 |
| | | | | | TIGR00681 | TIGR00681 | kdpC: K+-transporting ATPase, C subunit | 5.60E-48 |
| WP_148426666.1 | 421230 | 419079 | - | 716 | TIGR01497 | TIGR01497 | kdpB: K+-transporting ATPase, B subunit | 0.00E+00 |
| | | | | | TIGR01525 | TIGR01525 | ATPase-IB_hvy: heavy metal translocating P-type ATPase | 5.40E-82 |
| | | | | | TIGR01494 | TIGR01494 | ATPase_P-type: HAD ATPase, P-type, family IC | 5.70E-67 |
| | | | | | TIGR01512 | TIGR01512 | ATPase-IB2_Cd: cadmium-translocating P-type ATPase | 3.50E-59 |
| | | | | | TIGR01511 | TIGR01511 | ATPase-IB1_Cu: copper-translocating P-type ATPase | 8.00E-57 |

| index | leader | core | start | end | dir | score |
|---|---|---|---|---|---|---|
| 1 | VATGMTGAEPVSC | QCPQAKRTAAVCRPGPPRLGWTRRRVPGRRVLLPKPH | 402612 | 402459 | - | 6 |
| 2 | LPCRLVRFPCCWLRRHKQVGVPESPWRSRSSERSG | TPGLGIGSAGAAQGPGCPFGGPRRPSRIPAEDGAA | 405429 | 405216 | - | 6 |
| 3 | MRNVARHANRVSNVRFGSLSEATPAKKMSDSAPVL | FTPAVAVFGAAALITAVGSI | 410617 | 410449 | - | 29 |
| 4 | MSTSALEGCMRNVARHANRVSNVRFGSLSEATPAKKMSDSAPVL | FTPAVAVFGAAALITAVGSI | 410644 | 410449 | - | 30 |
| 5 | LNAMSTSALEGCMRNVARHANRVSNVRFGSLSEATPAKKMSDSAPVL | FTPAVAVFGAAALITAVGSI | 410653 | 410449 | - | 30 |

### Results for QDY07820.1 [Micromonospora sp. HM134] back to top

previous - next

#### Architecture

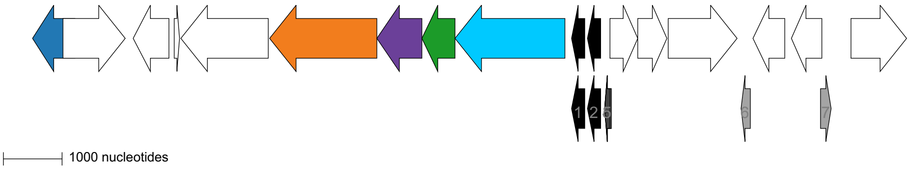

Link to nucleotide sequence

| Accession | start | end | direction | length (aa) | Pfam/HMM | name | description | E-value |
|---|---|---|---|---|---|---|---|---|
| QDY07814.1 | 2727816 | 2727084 | - | 243 | TIGR02427 | TIGR02427 | protocat_pcaD: 3-oxoadipate enol-lactonase | 2.70E-22 |
| | | | | | PF00561 | Abhydrolase_1 | alpha/beta hydrolase fold | 4.50E-19 |
| | | | | | PF12697 | Abhydrolase_6 | Alpha/beta hydrolase family | 2.30E-17 |
| | | | | | TIGR03056 | TIGR03056 | bchO_mg_che_rel: putative magnesium chelatase accessory protein | 5.00E-13 |
| | | | | | TIGR01738 | TIGR01738 | bioH: pimelyl-[acyl-carrier protein] methyl ester esterase | 1.60E-10 |
| QDY07815.1 | 2727607 | 2728678 | + | 356 | PF17765 | MLTR_LBD | MmyB-like transcription regulator ligand binding domain | 2.70E-31 |
| | | | | | PF13560 | HTH_31 | Helix-turn-helix domain | 1.00E-17 |
| QDY11584.1 | 2729427 | 2728815 | - | 203 | PF13673 | Acetyltransf_10 | Acetyltransferase (GNAT) domain | 8.40E-09 |
| | | | | | PF00583 | Acetyltransf_1 | Acetyltransferase (GNAT) family | 1.80E-07 |
| | | | | | PF13508 | Acetyltransf_7 | Acetyltransferase (GNAT) domain | 2.60E-07 |
| WP_003881694.1 | 2729506 | 2729587 | + | 27 | INFERRED GENE | - | - | - |
| QDY07816.1 | 2731112 | 2729630 | - | 493 | TIGR02734 | TIGR02734 | crtI_fam: phytoene desaturase | 4.30E-13 |
| | | | | | PF01593 | Amino_oxidase | Flavin containing amine oxidoreductase | 7.90E-08 |
| | | | | | TIGR02733 | TIGR02733 | desat_CrtD: C-3',4' desaturase CrtD | 3.00E-07 |
| | | | | | TIGR03467 | TIGR03467 | HpnE: squalene-associated FAD-dependent desaturase | 4.80E-07 |
| | | | | | PF13450 | NAD_binding_8 | NAD(P)-binding Rossmann-like domain | 9.50E-07 |
| QDY07817.1 | 2732959 | 2731129 | - | 609 | TIGR02204 | TIGR02204 | MsbA_rel: ABC transporter, permease/ATP-binding protein | 3.60E-106 |
| | | | | | TIGR02203 | TIGR02203 | MsbA_lipidA: lipid A export permease/ATP-binding protein MsbA | 3.90E-96 |
| | | | | | TIGR00958 | TIGR00958 | 3a01208: antigen peptide transporter 2 | 8.00E-91 |
| | | | | | TIGR03796 | TIGR03796 | NHLM_micro_ABC1: NHLM bacteriocin system ABC transporter, peptidase/ATP-binding protein | 1.00E-86 |
| | | | | | TIGR01846 | TIGR01846 | type_I_sec_HlyB: type I secretion system ATPase | 3.60E-85 |
| QDY07818.1 | 2733726 | 2732955 | - | 256 | PF13649 | Methyltransf_25 | Methyltransferase domain | 3.60E-18 |
| | | | | | PF08241 | Methyltransf_11 | Methyltransferase domain | 5.20E-12 |
| | | | | | PF13489 | Methyltransf_23 | Methyltransferase domain | 3.10E-09 |
| | | | | | PF08242 | Methyltransf_12 | Methyltransferase domain | 1.50E-08 |
| | | | | | PF13847 | Methyltransf_31 | Methyltransferase domain | 1.70E-07 |
| QDY07819.1 | 2734289 | 2733722 | - | 188 | LinL | LinL | LinL | 8.90E-65 |

| | | | | | | | | |
|---|---|---|---|---|---|---|---|---|
| QDY07820.1 | 2736172 | 2734285 | - | 628 | LinH<br>LinG<br>LinE<br>PF12697<br>PF00561 | LinH<br>LinG<br>LinE<br>Abhydrolase_6<br>Abhydrolase_1 | LinH (Fusion of LinE/LinG)<br>LinG Homolog<br>LinE Homolog<br>Alpha/beta hydrolase family<br>alpha/beta hydrolase fold | 1.60E-261<br>5.00E-103<br>1.20E-101<br>1.80E-18<br>8.00E-14 |
| QDY07821.1 | 2736505 | 2736277 | - | 75 | LinA | LinA | LinA (linaridin precursor) | 3.40E-17 |
| QDY07822.1 | 2736768 | 2736543 | - | 74 | LinA | LinA | LinA (linaridin precursor) | 4.70E-12 |
| QDY07823.1 | 2736927 | 2737404 | + | 158 | PF13560 | HTH_31 | Helix-turn-helix domain | 2.20E-05 |
| QDY07824.1 | 2737393 | 2737906 | + | 170 | PF06114 | Peptidase_M78 | IrrE N-terminal-like domain | 8.50E-04 |
| QDY07825.1 | 2737921 | 2739100 | + | 392 | NO PFAM MATCH | - | - | - |
| QDY07826.1 | 2739915 | 2739357 | - | 185 | PF01872<br>TIGR01508 | RibD_C<br>TIGR01508 | RibD C-terminal domain<br>rib_reduct_arch: diaminohydroxyphosphoribosylaminopyrimidine reductase | 2.00E-17<br>4.80E-04 |
| QDY07827.1 | 2740533 | 2740023 | - | 169 | PF13238<br>PF13671 | AAA_18<br>AAA_33 | AAA domain<br>AAA domain | 1.80E-08<br>9.20E-05 |
| QDY07828.1 | 2741034 | 2741976 | + | 313 | PF03547<br>TIGR00946 | Mem_trans<br>TIGR00946 | Membrane transport protein<br>2a69: auxin efflux carrier | 2.90E-33<br>5.80E-24 |

| index | leader | core | start | end | dir | score |
|---|---|---|---|---|---|---|
| 1 | MSVLANRVNADPALATIPIHSDEARNHVGGMPVL | ATPATIVSVAAGVGLVGSVAALFGAGYVAGRNLGGTRPVPK | 2736505 | 2736277 | - | 28 |
| 2 | MTALVDHTNDISLEMFGELTEADRTAHGLAGLTPVL | ATPTAVALGVAVGAVAFHAGYMVGIARNMAEGTRPQPK | 2736768 | 2736543 | - | 30 |
| 3 | MWCPPLVGSGGPRFPW | LTAATPSDILAIHDF | 2736929 | 2736833 | - | 6 |
| 4 | VLGWSMWCPPLVGSGGPRFPW | LTAATPSDILAIHDF | 2736944 | 2736833 | - | 6 |
| 5 | VPVLGWSMWCPPLVGSGGPRFPW | LTAATPSDILAIHDF | 2736950 | 2736833 | - | 6 |
| 6 | MPSQEEVRSQGVTSRCWVSAGSRRWRC | VLGSGSHTGIPANRAVGSGRCLGAADG | 2739322 | 2739157 | - | 6 |
| 7 | VPRPYGSGQRDRRQELLAIHRAVRGSHDTRPRPTT | QTVAVVSPLLTEPTKARRDLRPD | 2740526 | 2740703 | + | 6 |

### Results for OWV06806.1 [Micromonospora wenchangensis] back to top

previous - next

#### Architecture

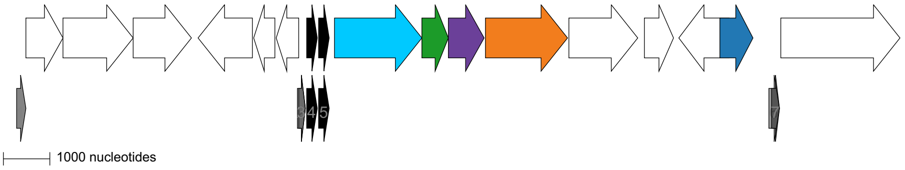

Link to nucleotide sequence

| Accession | start | end | direction | length (aa) | Pfam/HMM | name | description | E-value |
|---|---|---|---|---|---|---|---|---|
| OWV06799.1 | 36347 | 37157 | + | 269 | TIGR01694<br>PF01048<br>TIGR01698<br>TIGR01697<br>TIGR01700 | TIGR01694<br>PNP_UDP_1<br>TIGR01698<br>TIGR01697<br>TIGR01700 | MTAP: methylthioadenosine phosphorylase<br>Phosphorylase superfamily<br>PUNP: purine nucleotide phosphorylase<br>PNPH-PUNA-XAPA: inosine/guanosine/xanthosine phosphorylase family<br>PNPH: purine nucleoside phosphorylase I, inosine and guanosine-specific | 6.50E-88<br>5.80E-41<br>4.40E-19<br>2.20E-15<br>9.80E-12 |
| OWV06800.1 | 37153 | 38665 | + | 503 | TIGR00895<br>TIGR00898<br>PF00083<br>PF07690<br>TIGR00891 | TIGR00895<br>TIGR00898<br>Sugar_tr<br>MFS_1<br>TIGR00891 | 2A0115: MFS transporter, aromatic acid:H+ symporter (AAHS) family<br>2A0119: cation transport protein<br>Sugar (and other) transporter<br>Major Facilitator Superfamily<br>2A0112: MFS transporter, sialate:H+ symporter (SHS) family | 8.70E-80<br>5.80E-56<br>1.20E-42<br>1.90E-42<br>8.40E-39 |
| OWV06801.1 | 38648 | 39899 | + | 416 | PF01979<br>PF07969<br>TIGR02967<br>TIGR01224<br>TIGR03121 | Amidohydro_1<br>Amidohydro_3<br>TIGR02967<br>TIGR01224<br>TIGR03121 | Amidohydrolase family<br>Amidohydrolase family<br>guan_deamin: guanine deaminase<br>hutI: imidazolonepropionase<br>one_C_dehyd_A: formylmethanofuran dehydrogenase subunit A | 1.20E-40<br>6.10E-14<br>2.70E-09<br>1.10E-08<br>1.50E-07 |
| OWV06802.1 | 41230 | 40051 | - | 392 | NO PFAM MATCH | - | - | - |
| OWV06851.1 | 41707 | 41245 | - | 153 | PF06114 | Peptidase_M78 | IrrE N-terminal-like domain | 4.80E-04 |
| OWV06803.1 | 42224 | 41747 | - | 158 | PF13560 | HTH_31 | Helix-turn-helix domain | 2.20E-05 |
| OWV06804.1 | 42376 | 42610 | + | 77 | LinA | LinA | LinA (linaridin precursor) | 2.30E-12 |
| OWV06805.1 | 42643 | 42874 | + | 76 | LinA | LinA | LinA (linaridin precursor) | 3.70E-17 |
| OWV06806.1 | 42979 | 44869 | + | 629 | LinH<br>LinG<br>LinE<br>PF12697<br>PF00561 | LinH<br>LinG<br>LinE<br>Abhydrolase_6<br>Abhydrolase_1 | LinH (Fusion of LinE/LinG)<br>LinG Homolog<br>LinE Homolog<br>Alpha/beta hydrolase family<br>alpha/beta hydrolase fold | 5.90E-261<br>1.30E-102<br>1.20E-100<br>4.20E-18<br>8.10E-14 |
| OWV06807.1 | 44865 | 45432 | + | 188 | LinL | LinL | LinL | 1.00E-65 |
| OWV06808.1 | 45428 | 46199 | + | 256 | PF13649<br>PF08241<br>PF08242<br>PF13489<br>PF13847 | Methyltransf_25<br>Methyltransf_11<br>Methyltransf_12<br>Methyltransf_23<br>Methyltransf_31 | Methyltransferase domain<br>Methyltransferase domain<br>Methyltransferase domain<br>Methyltransferase domain<br>Methyltransferase domain | 3.30E-17<br>1.10E-11<br>3.90E-09<br>6.70E-09<br>1.70E-07 |

| | | | | | | | | |
|---|---|---|---|---|---|---|---|---|
| OWV06852.1 | 46234 | 48013 | + | 592 | TIGR02204<br>TIGR02203<br>TIGR00958<br>TIGR03796<br>TIGR01846 | TIGR02204<br>TIGR02203<br>TIGR00958<br>TIGR03796<br>TIGR01846 | MsbA_rel: ABC transporter, permease/ATP-binding protein<br>MsbA_lipidA: lipid A export permease/ATP-binding protein MsbA<br>3a01208: antigen peptide transporter 2<br>NHLM_micro_ABC1: NHLM bacteriocin system ABC transporter, peptidase/ATP-binding protein<br>type_I_sec_HlyB: type I secretion system ATPase | 1.00E-103<br>7.00E-95<br>1.50E-90<br>3.80E-86<br>1.40E-84 |
| OWV06809.1 | 48029 | 49511 | + | 493 | TIGR02734<br>PF13450<br>TIGR02733<br>PF01593<br>TIGR03467 | TIGR02734<br>NAD_binding_8<br>TIGR02733<br>Amino_oxidase<br>TIGR03467 | crtI_fam: phytoene desaturase<br>NAD(P)-binding Rossmann-like domain<br>desat_CrtD: C-3',4' desaturase CrtD<br>Flavin containing amine oxidoreductase<br>HpnE: squalene-associated FAD-dependent desaturase | 2.70E-12<br>1.00E-07<br>3.00E-07<br>3.50E-07<br>5.80E-07 |
| OWV06810.1 | 49667 | 50279 | + | 203 | PF13673<br>PF00583<br>PF13508 | Acetyltransf_10<br>Acetyltransf_1<br>Acetyltransf_7 | Acetyltransferase (GNAT) domain<br>Acetyltransferase (GNAT) family<br>Acetyltransferase (GNAT) domain | 5.50E-08<br>1.60E-07<br>4.10E-06 |
| OWV06811.1 | 51486 | 50415 | - | 356 | PF17765<br>PF13560 | MLTR_LBD<br>HTH_31 | MmyB-like transcription regulator ligand binding domain<br>Helix-turn-helix domain | 5.50E-30<br>4.60E-16 |
| OWV06853.1 | 51277 | 52009 | + | 243 | TIGR02427<br>PF00561<br>PF12697<br>TIGR03056<br>TIGR01738 | TIGR02427<br>Abhydrolase_1<br>Abhydrolase_6<br>TIGR03056<br>TIGR01738 | protocat_pcaD: 3-oxoadipate enol-lactonase<br>alpha/beta hydrolase fold<br>Alpha/beta hydrolase family<br>bchO_mg_che_rel: putative magnesium chelatase accessory protein<br>bioH: pimelyl-[acyl-carrier protein] methyl ester esterase | 2.80E-20<br>2.80E-18<br>1.30E-15<br>2.90E-11<br>2.00E-09 |
| OWV06812.1 | 52587 | 55164 | + | 858 | PF00196<br>TIGR03541<br>PF13191<br>TIGR03020<br>PF08281 | GerE<br>TIGR03541<br>AAA_16<br>TIGR03020<br>Sigma70_r4_2 | Bacterial regulatory proteins, luxR family<br>reg_near_HchA: LuxR family transcriptional regulatory, chaperone HchA-associated<br>AAA ATPase domain<br>EpsA: transcriptional regulator EpsA<br>Sigma-70, region 4 | 2.40E-15<br>2.60E-09<br>3.40E-07<br>5.90E-07<br>5.40E-06 |

| index | leader | core | start | end | dir | score |
|---|---|---|---|---|---|---|
| 1 | VADGIRFCRELPHRVGVAAIPSAVFYDDPAD | GLPIVRFVFCKEPQVLDEAARRLADLTTPYHQEAL | 36150 | 36351 | + | 7 |
| 2 | VPVPGWSMWCPSPRRIGWTPFPVVDG | GDPLGHPSDSRFLAEMNGARAAFHTG | 42201 | 42360 | + | 6 |
| 3 | VPGWSMWCPSPRRIGWTPFPVVDGG | DPLGHPSDSRFLAEMNGARAAFHTG | 42207 | 42360 | + | 6 |
| 4 | MNQMTALVDHTNDISLEMFGELTEADRNAHGLAGLTPVL | ATPTAVALGVAVGAVAFHAGYMVGIARNMAEGTRPLPK | 42376 | 42610 | + | 30 |
| 5 | MSVLANRVNADPALATIPIHSDEARNHVGGMPVL | ATPATIVSVAAGVGLVGSVAALFGAGYVAGRNLGGTRPPAPK | 42643 | 42874 | + | 29 |
| 6 | MPGPQPRRPREPLWTYPTVCGRGDGHCGQPRRPVGIPM | EAHCWRTGSSRPQCRQHYAATTTSGPAARFMHHQGMGR | 52349 | 52580 | + | 6 |
| 7 | VCGRGDGHCGQPRRPVGIPMEAHCWRTGS | SRPQCRQHYAATTTSGPAARFMHHQGMGR | 52403 | 52580 | + | 7 |

### Results for WP_132400124.1 [Micromonospora sp. KC207] back to top

previous - next

#### Architecture

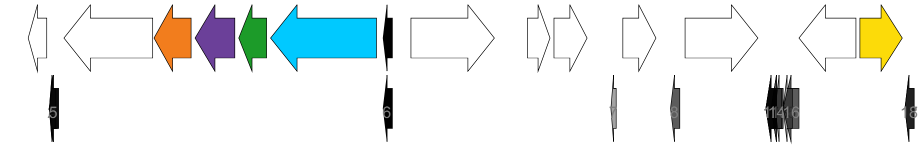

1000 nucleotides

Link to nucleotide sequence

| Accession | start | end | direction | length (aa) | Pfam/HMM | name | description | E-value |
|---|---|---|---|---|---|---|---|---|
| WP_165949750.1 | 306 | 0 | - | 102 | PF05598 | DUF772 | Transposase domain (DUF772) | 3.70E-16 |
| WP_132400115.1 | 2151 | 612 | - | 512 | NO PFAM MATCH | - | - | - |
| WP_132400118.1 | 2816 | 2165 | - | 216 | TIGR01189<br>TIGR02857<br>TIGR00968<br>TIGR01166<br>TIGR03873 | TIGR01189<br>TIGR02857<br>TIGR00968<br>TIGR01166<br>TIGR03873 | ccmA: heme ABC exporter, ATP-binding protein CcmA<br>CydD: thiol reductant ABC exporter, CydD subunit<br>3a0106s01: sulfate ABC transporter, ATP-binding protein<br>cbiO: cobalt ABC transporter, ATP-binding protein<br>F420-0_ABC_ATP: proposed F420-0 ABC transporter, ATP-binding protein | 2.90E-31<br>7.30E-30<br>3.70E-27<br>1.20E-26<br>2.30E-26 |
| WP_132400165.1 | 3561 | 2883 | - | 225 | PF13649<br>PF08241<br>PF13489<br>PF08242<br>TIGR02021 | Methyltransf_25<br>Methyltransf_11<br>Methyltransf_23<br>Methyltransf_12<br>TIGR02021 | Methyltransferase domain<br>Methyltransferase domain<br>Methyltransferase domain<br>Methyltransferase domain<br>BchM-ChlM: magnesium protoporphyrin O-methyltransferase | 6.50E-12<br>9.80E-10<br>1.20E-09<br>2.50E-09<br>1.10E-08 |
| WP_165949751.1 | 4121 | 3647 | - | 157 | LinL | LinL | LinL | 6.70E-52 |
| WP_132400124.1 | 6031 | 4186 | - | 614 | LinH<br>LinE<br>LinG<br>PF12697<br>TIGR03056 | LinH<br>LinE<br>LinG<br>Abhydrolase_6<br>TIGR03056 | LinH (Fusion of LinE/LinG)<br>LinE Homolog<br>LinG Homolog<br>Alpha/beta hydrolase family<br>bchO_mg_che_rel: putative magnesium chelatase accessory protein | 3.00E-217<br>5.50E-97<br>1.90E-90<br>4.70E-13<br>3.00E-09 |
| WP_165949752.1 | 6299 | 6128 | - | 56 | LinA | LinA | LinA (linaridin precursor) | 8.60E-09 |
| WP_132400127.1 | 6616 | 8068 | + | 483 | TIGR02734<br>PF13450<br>TIGR03467<br>PF01593<br>TIGR00292 | TIGR02734<br>NAD_binding_8<br>TIGR03467<br>Amino_oxidase<br>TIGR00292 | crtI_fam: phytoene desaturase<br>NAD(P)-binding Rossmann-like domain<br>HpnE: squalene-associated FAD-dependent desaturase<br>Flavin containing amine oxidoreductase<br>TIGR00292: thiazole biosynthesis enzyme | 1.80E-14<br>2.60E-10<br>1.10E-08<br>2.80E-08<br>1.70E-07 |
| WP_132400130.1 | 8634 | 9021 | + | 128 | PF14690 | zf-ISL3 | zinc-finger of transposase IS204/IS1001/IS1096/IS1165 | 1.20E-08 |

| | | | | | | | | |
|---|---|---|---|---|---|---|---|---|
| WP_165949753.1 | 9095 | 9671 | + | 191 | PF01610<br>PF13384<br>PF13551<br>TIGR03879 | DDE_Tnp_ISL3<br>HTH_23<br>HTH_29<br>TIGR03879 | Transposase<br>Homeodomain-like domain<br>Winged helix-turn helix<br>near_KaiC_dom: probable regulatory domain | 4.10E-15<br>1.30E-04<br>1.40E-04<br>4.70E-04 |
| WP_132400136.1 | 10293 | 10860 | + | 188 | NO PFAM MATCH | - | - | - |
| WP_076611994.1 | 11371 | 12615 | + | 414 | INFERRED GENE | - | - | - |
| WP_132400139.1 | 14315 | 13343 | - | 323 | PF13280<br>PF08279<br>PF02082<br>TIGR00122<br>TIGR02944 | WYL<br>HTH_11<br>Rrf2<br>TIGR00122<br>TIGR02944 | WYL domain<br>HTH domain<br>Transcriptional regulator<br>birA_repr_reg: biotin operon repressor<br>suf_reg_Xantho: FeS assembly SUF system regulator | 3.20E-32<br>3.10E-14<br>6.20E-08<br>5.60E-05<br>1.00E-04 |
| WP_132400142.1 | 14396 | 15131 | + | 244 | PF12697<br>LinH<br>LinE | Abhydrolase_6<br>LinH<br>LinE | Alpha/beta hydrolase family<br>LinH (Fusion of LinE/LinG)<br>LinE Homolog | 1.60E-12<br>2.90E-09<br>8.20E-08 |

| index | leader | core | start | end | dir | score |
|---|---|---|---|---|---|---|
| 1 | VSSGWGASLPELRASRCWGS | GMSMGRGSGSVVPVETARVA | 474 | 351 | - | 6 |
| 2 | LVVSSGWGASLPELRASRCWG | SGMSMGRGSGSVVPVETARVA | 480 | 351 | - | 6 |
| 3 | VILVVSSGWGASLPELRASRCW | GSGMSMGRGSGSVVPVETARVA | 486 | 351 | - | 6 |
| 4 | LVILVVSSGWGASLPELRASRC | WGSGMSMGRGSGSVVPVETARVA | 489 | 351 | - | 6 |
| 5 | VGRFRLCLTFEMLLVILVVSSGWGASLPE | LRASRCWGSGMSMGRGSGSVVPVETARVA | 528 | 351 | - | 9 |
| 6 | MYDVVSHANGISEEKFGKLSADSPAKKMTATTPVL | FTPAAVTVLVPAALITAVGSV | 6299 | 6128 | - | 31 |
| 7 | VASSFGSDPLGCGVVRD | AQACRMRCRSSRTWTVRR | 10179 | 10071 | - | 6 |
| 8 | MGVVLFVGDGSLVDPGPKPVRQ | GVGQASPRAASARSVAPRRTWRC | 11261 | 11123 | - | 8 |
| 9 | LGRGGNRRYGPRLRPPSNRPEDAPVR | VAPDGAVSSVRSGIPATPGTRRLLGFAGVSTAAAV | 12939 | 12753 | - | 8 |
| 10 | LDPLGRGGNRRYGPRLRPPSNRPEDAPVR | VAPDGAVSSVRSGIPATPGTRRLLGFAGVSTAAAV | 12948 | 12753 | - | 8 |
| 11 | LLLDPLGRGGNRRYGPRLRPPSNRPEDAPVR | VAPDGAVSSVRSGIPATPGTRRLLGFAGVSTAAAV | 12954 | 12753 | - | 9 |
| 12 | VRHRRSLPAEGASTRTRSTTRPYAGCYW | TRWAGAVIVATDRAFGHHQTVRRTRQCE | 13028 | 12857 | - | 7 |
| 13 | MASSSHEPCDTGAACLRKGLL | PGLVRLRGPTRVAIGPVGPGR | 13051 | 12922 | - | 6 |
| 14 | VIRVGVDGFQQPRAVRHRRSLPAEGASTRTRSTTR | PYAGCYWTRWAGAVIVATDRAFGHHQTVRRTRQCE | 13070 | 12857 | - | 6 |
| 15 | LTNELVERVATRRSTHLLRVAI | DAPDTASKTTLAGEAADKLTAQAAR | 13210 | 13066 | - | 6 |
| 16 | MPRPAHARALARFDHQGAGARTESR...TGGLTNELVERVATRRSTHLLRVAI | DAPDTASKTTLAGEAADKLTAQAAR | 13339 | 13066 | - | 8 |
| 17 | MSRPRARMPRSAAVASCAWMAKTRRT | TAAAESSRTPSASACARARIRRTRFW | 15319 | 15160 | - | 8 |
| 18 | VMSRPRARMPRSAAVASCAWMAKTRR | TTAAAESSRTPSASACARARIRRTRFW | 15322 | 15160 | - | 8 |

### Results for SCL65344.1 [Micromonospora yangpuensis] back to top

previous - next

#### Architecture

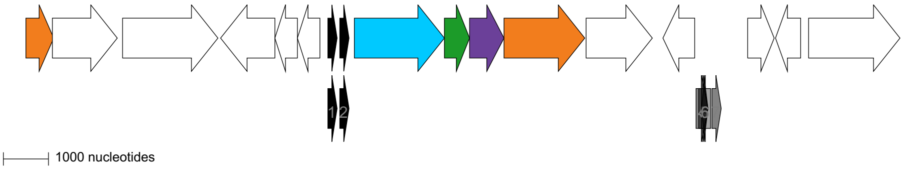

Link to nucleotide sequence

| Accession | start | end | direction | length (aa) | Pfam/HMM | name | description | E-value |
|---|---|---|---|---|---|---|---|---|
| SCL65317.1 | 6247041 | 6247659 | + | 205 | TIGR01189<br>TIGR01166<br>TIGR03265<br>TIGR02868<br>TIGR02857 | TIGR01189<br>TIGR01166<br>TIGR03265<br>TIGR02868<br>TIGR02857 | ccmA: heme ABC exporter, ATP-binding protein CcmA<br>cbiO: cobalt ABC transporter, ATP-binding protein<br>PhnT2: putative 2-aminoethylphosphonate ABC transporter, ATP-binding protein<br>CydC: thiol reductant ABC exporter, CydC subunit<br>CydD: thiol reductant ABC exporter, CydD subunit | 2.10E-38<br>6.40E-32<br>4.40E-31<br>8.10E-30<br>2.20E-29 |
| SCL65320.1 | 6247655 | 6249083 | + | 475 | NO PFAM MATCH | - | - | - |
| SCL65325.1 | 6249192 | 6251304 | + | 703 | PF18448<br>PF00150<br>PF00553<br>PF03442 | CBM46<br>Cellulase<br>CBM_2<br>CBM_X2 | Carbohydrate binding domain<br>Cellulase (glycosyl hydrolase family 5)<br>Cellulose binding domain<br>Carbohydrate binding domain X2 | 2.80E-53<br>1.80E-52<br>1.10E-29<br>9.10E-19 |
| SCL65326.1 | 6252561 | 6251382 | - | 392 | NO PFAM MATCH | - | - | - |
| SCL65331.1 | 6253089 | 6252576 | - | 170 | PF06114 | Peptidase_M78 | IrrE N-terminal-like domain | 3.90E-04 |
| SCL65333.1 | 6253579 | 6253078 | - | 166 | PF13560 | HTH_31 | Helix-turn-helix domain | 3.10E-05 |
| SCL65337.1 | 6253741 | 6253969 | + | 75 | LinA | LinA | LinA (linaridin precursor) | 2.00E-12 |
| SCL65341.1 | 6254002 | 6254230 | + | 75 | LinA | LinA | LinA (linaridin precursor) | 6.00E-17 |

| | | | | | | | | |
|---|---|---|---|---|---|---|---|---|
| SCL65344.1 | 6254326 | 6256333 | + | 668 | LinH | LinH | LinH (Fusion of LinE/LinG) | 7.90E-262 |
| | | | | | LinE | LinE | LinE Homolog | 7.30E-103 |
| | | | | | LinG | LinG | LinG Homolog | 6.80E-102 |
| | | | | | PF12697 | Abhydrolase_6 | Alpha/beta hydrolase family | 1.90E-19 |
| | | | | | PF00561 | Abhydrolase_1 | alpha/beta hydrolase fold | 8.20E-13 |
| SCL65348.1 | 6256329 | 6256899 | + | 189 | LinL | LinL | LinL | 8.50E-67 |
| SCL65350.1 | 6256891 | 6257662 | + | 256 | PF13649 | Methyltransf_25 | Methyltransferase domain | 9.30E-17 |
| | | | | | PF08241 | Methyltransf_11 | Methyltransferase domain | 3.00E-11 |
| | | | | | PF13489 | Methyltransf_23 | Methyltransferase domain | 1.40E-09 |
| | | | | | PF08242 | Methyltransf_12 | Methyltransferase domain | 5.60E-09 |
| | | | | | PF13847 | Methyltransf_31 | Methyltransferase domain | 8.90E-07 |
| SCL65355.1 | 6257658 | 6259461 | + | 600 | TIGR02204 | TIGR02204 | MsbA_rel: ABC transporter, permease/ATP-binding protein | 9.90E-101 |
| | | | | | TIGR02203 | TIGR02203 | MsbA_lipidA: lipid A export permease/ATP-binding protein MsbA | 3.00E-94 |
| | | | | | TIGR00958 | TIGR00958 | 3a01208: antigen peptide transporter 2 | 4.30E-89 |
| | | | | | TIGR03796 | TIGR03796 | NHLM_micro_ABC1: NHLM bacteriocin system ABC transporter, peptidase/ATP-binding protein | 4.20E-86 |
| | | | | | TIGR01846 | TIGR01846 | type_I_sec_HlyB: type I secretion system ATPase | 1.20E-79 |
| SCL65366.1 | 6259472 | 6260930 | + | 485 | TIGR02734 | TIGR02734 | crtI_fam: phytoene desaturase | 3.10E-13 |
| | | | | | PF13450 | NAD_binding_8 | NAD(P)-binding Rossmann-like domain | 5.80E-09 |
| | | | | | TIGR03467 | TIGR03467 | HpnE: squalene-associated FAD-dependent desaturase | 5.80E-09 |
| | | | | | PF01593 | Amino_oxidase | Flavin containing amine oxidoreductase | 2.00E-08 |
| | | | | | TIGR02733 | TIGR02733 | desat_CrtD: C-3',4' desaturase CrtD | 1.80E-07 |
| SCL65369.1 | 6261882 | 6261180 | - | 233 | PF02698 | DUF218 | DUF218 domain | 5.70E-20 |
| SCL65372.1 | 6263064 | 6263655 | + | 196 | PF00440 | TetR_N | Bacterial regulatory proteins, tetR family | 8.50E-07 |
| | | | | | TIGR03384 | TIGR03384 | betaine_BetI: transcriptional repressor BetI | 1.50E-04 |
| SCL65376.1 | 6264246 | 6263688 | - | 185 | NO PFAM MATCH | - | - | - |
| SCL65379.1 | 6264400 | 6266422 | + | 673 | TIGR01991 | TIGR01991 | HscA: Fe-S protein assembly chaperone HscA | 8.80E-25 |
| | | | | | PF00012 | HSP70 | Hsp70 protein | 2.40E-19 |
| | | | | | TIGR02350 | TIGR02350 | prok_dnaK: chaperone protein DnaK | 3.20E-18 |

| index | leader | core | start | end | dir | score |
|---|---|---|---|---|---|---|
| 1 | MTALVDHTNDVSLEMFGELNEADRTAQGLAGLTPVL | ATPTAVALGVAVGAVAFHAGYMVGIVRNMAEGTVTPRPK | 6253741 | 6253969 | + | 31 |
| 2 | MSVLATKVNADPALAAIPIHSDEACNQVGGMPVL | ATPATIVGVAAGVGLVGSVAALFGAGYVVGRNLGGTTPRPK | 6254002 | 6254230 | + | 29 |
| 3 | VPAQQRAGSPGRGPGAGGRVTWQQS...WVRGGSVSRSPPPTPPRRQPPPPPS | STADALALSVPDRL | 6261909 | 6262173 | + | 7 |
| 4 | VTWQQSPQGPFSSPSPGLLPTPTET...WVRGGSVSRSPPPTPPRRQPPPPPS | STADALALSVPDRL | 6261966 | 6262173 | + | 7 |
| 5 | LPTPTETRRDSWVRGGSVSRSPPPTPPRRQPPPPPS | STADALALSVPDRL | 6262020 | 6262173 | + | 7 |
| 6 | VNLGHLPFYLDGKCPRSTADRVRREQAHRLW | RRAGAEAVDGAGLVVVTGKPNHLAPSCPAGSP | 6262222 | 6262030 | - | 6 |
| 7 | VAQRSWKQRTPGGAVCFQYLAAHPAGTIPWGTPRG | HTPRFVERAGPPEPVGGRGAQRVVRVRTDHGCLVGHG | 6262254 | 6262473 | + | 7 |

### Results for TCB95381.1 [Micromonospora zingiberis] back to top

previous - next

#### Architecture

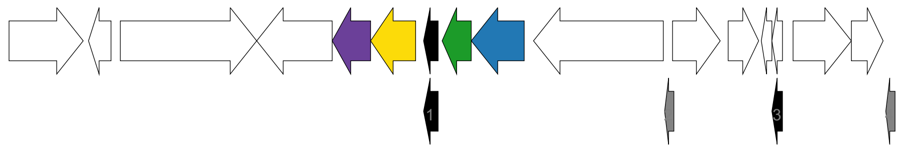

1000 nucleotides

Link to nucleotide sequence

| Accession | start | end | direction | length (aa) | Pfam/HMM | name | description | E-value |
|---|---|---|---|---|---|---|---|---|
| TCB95407.1 | 121664 | 123128 | + | 487 | TIGR00838 | TIGR00838 | argH: argininosuccinate lyase | 1.10E-171 |
| | | | | | PF00206 | Lyase_1 | Lyase | 2.20E-67 |
| | | | | | PF14698 | ASL_C2 | Argininosuccinate lyase C-terminal | 5.40E-23 |
| | | | | | TIGR00928 | TIGR00928 | purB: adenylosuccinate lyase | 2.80E-22 |
| | | | | | TIGR02426 | TIGR02426 | protocat_pcaB: 3-carboxy-cis,cis-muconate cycloisomerase | 4.60E-22 |
| TCB95374.1 | 123695 | 123245 | - | 149 | NO PFAM MATCH | - | - | - |
| TCB95375.1 | 123859 | 126577 | + | 905 | PF13424 | TPR_12 | Tetratricopeptide repeat | 1.80E-18 |
| | | | | | PF00931 | NB-ARC | NB-ARC domain | 4.70E-15 |
| | | | | | PF07719 | TPR_2 | Tetratricopeptide repeat | 5.50E-07 |
| | | | | | PF13191 | AAA_16 | AAA ATPase domain | 6.70E-07 |
| | | | | | PF13176 | TPR_7 | Tetratricopeptide repeat | 1.20E-05 |
| TCB95376.1 | 128052 | 126564 | - | 495 | TIGR02734 | TIGR02734 | crtI_fam: phytoene desaturase | 4.90E-10 |
| | | | | | TIGR00292 | TIGR00292 | TIGR00292: thiazole biosynthesis enzyme | 1.20E-08 |
| | | | | | TIGR02733 | TIGR02733 | desat_CrtD: C-3',4' desaturase CrtD | 2.20E-08 |
| | | | | | TIGR03467 | TIGR03467 | HpnE: squalene-associated FAD-dependent desaturase | 7.60E-08 |
| | | | | | PF13450 | NAD_binding_8 | NAD(P)-binding Rossmann-like domain | 1.10E-07 |
| TCB95377.1 | 128804 | 128048 | - | 251 | PF13649 | Methyltransf_25 | Methyltransferase domain | 7.80E-14 |
| | | | | | PF08241 | Methyltransf_11 | Methyltransferase domain | 9.30E-10 |
| | | | | | PF13489 | Methyltransf_23 | Methyltransferase domain | 2.00E-07 |
| | | | | | PF08242 | Methyltransf_12 | Methyltransferase domain | 2.40E-07 |
| | | | | | PF13847 | Methyltransf_31 | Methyltransferase domain | 6.40E-07 |
| TCB95378.1 | 129707 | 128816 | - | 296 | LinG | LinG | LinG Homolog | 1.80E-100 |
| | | | | | LinH | LinH | LinH (Fusion of LinE/LinG) | 6.30E-78 |

| | | | | | | | | |
|---|---|---|---|---|---|---|---|---|
| TCB95379.1 | 130156 | 129868 | - | 95 | LinA | LinA | LinA (linaridin precursor) | 1.10E-08 |
| TCB95380.1 | 130806 | 130242 | - | 187 | LinL | LinL | LinL | 4.30E-54 |
| TCB95381.1 | 131852 | 130802 | - | 349 | LinE<br>LinH<br>PF12697<br>PF00561<br>TIGR02427 | LinE<br>LinH<br>Abhydrolase_6<br>Abhydrolase_1<br>TIGR02427 | LinE Homolog<br>LinH (Fusion of LinE/LinG)<br>Alpha/beta hydrolase family<br>alpha/beta hydrolase fold<br>protocat_pcaD: 3-oxoadipate enol-lactonase | 3.50E-87<br>4.90E-43<br>5.20E-18<br>6.00E-10<br>1.70E-07 |
| TCB95382.1 | 134601 | 132039 | - | 853 | PF00501<br>TIGR03098<br>TIGR01733<br>TIGR01923<br>TIGR02262 | AMP-binding<br>TIGR03098<br>TIGR01733<br>TIGR01923<br>TIGR02262 | AMP-binding enzyme<br>ligase_PEP_1: acyl-CoA ligase (AMP-forming), exosortase A-associated<br>AA-adenyl-dom: amino acid adenylation domain<br>menE: O-succinylbenzoate-CoA ligase<br>benz_CoA_lig: benzoate-CoA ligase family | 1.60E-48<br>5.80E-44<br>1.80E-41<br>3.80E-26<br>3.60E-23 |
| TCB95383.1 | 134784 | 135726 | + | 313 | PF18164<br>PF18082 | GNAT_C<br>DUF5596 | GNAT-like C-terminal domain<br>Domain of unknown function (DUF5596) | 7.70E-40<br>1.70E-19 |
| TCB95384.1 | 135885 | 136491 | + | 201 | PF02245<br>TIGR00567 | Pur_DNA_glyco<br>TIGR00567 | Methylpurine-DNA glycosylase (MPG)<br>3mg: DNA-3-methyladenine glycosylase | 1.10E-56<br>3.50E-46 |
| TCB95385.1 | 136751 | 136526 | - | 74 | PF07927 | HicA_toxin | HicA toxin of bacterial toxin-antitoxin, | 4.40E-14 |
| TCB95386.1 | 136960 | 136747 | - | 70 | PF15919 | HicB_lk_antitox | HicB_like antitoxin of bacterial toxin-antitoxin system | 7.50E-04 |
| TCB95387.1 | 137174 | 138311 | + | 378 | TIGR04036<br>PF00296<br>TIGR03858<br>TIGR03619<br>TIGR03558 | TIGR04036<br>Bac_luciferase<br>TIGR03858<br>TIGR03619<br>TIGR03558 | LLM_CE1758_fam: putative luciferase-like monooxygenase, FMN-dependent, CE1758 family<br>Luciferase-like monooxygenase<br>LLM_2I7G: probable oxidoreductase, LLM family<br>F420_Rv2161c: probable F420-dependent oxidoreductase, Rv2161c family<br>oxido_grp_1: luciferase family oxidoreductase, group 1 | 4.00E-217<br>2.50E-56<br>1.60E-51<br>2.20E-22<br>5.40E-22 |
| TCB95388.1 | 138307 | 138931 | + | 207 | TIGR04037<br>TIGR03566<br>PF03358<br>TIGR03567<br>PF02525 | TIGR04037<br>TIGR03566<br>FMN_red<br>TIGR03567<br>Flavodoxin_2 | LLM_duo_CE1759: LLM-partnered FMN reductase, CE1759 family<br>FMN_reduc_MsuE: FMN reductase<br>NADPH-dependent FMN reductase<br>FMN_reduc_SsuE: FMN reductase<br>Flavodoxin-like fold | 1.10E-83<br>1.80E-41<br>3.90E-35<br>3.40E-24<br>4.60E-05 |

| index | leader | core | start | end | dir | score |
|---|---|---|---|---|---|---|
| 1 | MGSGRCAAGRITRRSKKDHGEGKQM...TDVPPGMLTQTDVTSPLDTPADPVM | ATFLTVTCPEVAVLFGAATGALAANVYNYVARNG | 130156 | 129868 | - | 32 |
| 2 | VSSRSTPTTLPTDGAAATGPDSRQPLD | LIMRDRRPATGRHDAALVADVIRAARS | 134801 | 134636 | - | 7 |
| 3 | MVHTFTAAVHQEEDWYVARCLEVEV...LDEALPNLREAVEIYLEEVSQPAIE | ATPLVTSFQIGRAA | 136960 | 136747 | - | 20 |
| 4 | VVTAPKPDEALIEASGEAALVVAGSRGRGGF | AGLLLGSVSQALVHHARCPVLVAHPYAPEHAA | 139176 | 138984 | - | 7 |

### Results for WP_066941623.1 [Microtetraspora fusca] back to top

previous - next

#### Architecture

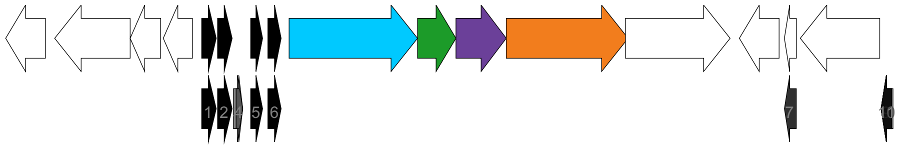

1000 nucleotides

Link to nucleotide sequence

| Accession | start | end | direction | length (aa) | Pfam/HMM | name | description | E-value |
|---|---|---|---|---|---|---|---|---|
| WP_066941610.1 | 197370 | 196764 | - | 201 | PF00574<br>TIGR00493<br>TIGR00706<br>TIGR01705 | CLP_protease<br>TIGR00493<br>TIGR00706<br>TIGR01705 | Clp protease<br>clpP: ATP-dependent Clp endopeptidase, proteolytic subunit ClpP<br>SppA_dom: signal peptide peptidase SppA, 36K type<br>MTA/SAH-nuc-hyp: putative 5'-methylthioadenosine/S-adenosylhomocysteine nucleosidase | 3.10E-67<br>7.80E-62<br>3.60E-06<br>1.80E-04 |
| WP_066941612.1 | 198662 | 197498 | - | 387 | NO PFAM MATCH | - | - | - |
| WP_066941739.1 | 199118 | 198665 | - | 150 | NO PFAM MATCH | - | - | - |
| WP_066941614.1 | 199599 | 199158 | - | 146 | PF13560 | HTH_31 | Helix-turn-helix domain | 1.30E-06 |
| WP_066941616.1 | 199753 | 199969 | + | 71 | LinA | LinA | LinA (linaridin precursor) | 2.10E-15 |
| WP_066941618.1 | 199994 | 200210 | + | 71 | LinA | LinA | LinA (linaridin precursor) | 5.00E-18 |
| WP_066941620.1 | 200488 | 200677 | + | 62 | LinA | LinA | LinA (linaridin precursor) | 9.30E-14 |
| WP_066941621.1 | 200762 | 200954 | + | 63 | LinA | LinA | LinA (linaridin precursor) | 4.20E-09 |
| WP_066941623.1 | 201086 | 203036 | + | 649 | LinH<br>LinE<br>LinG<br>PF12697<br>PF00561 | LinH<br>LinE<br>LinG<br>Abhydrolase_6<br>Abhydrolase_1 | LinH (Fusion of LinE/LinG)<br>LinE Homolog<br>LinG Homolog<br>Alpha/beta hydrolase family<br>alpha/beta hydrolase fold | 8.10E-258<br>9.00E-103<br>5.30E-98<br>1.20E-19<br>3.20E-16 |
| WP_066941625.1 | 203035 | 203620 | + | 194 | LinL | LinL | LinL | 2.50E-67 |
| WP_066941627.1 | 203612 | 204377 | + | 254 | PF13649<br>PF08241<br>PF13847<br>PF13489<br>TIGR01983 | Methyltransf_25<br>Methyltransf_11<br>Methyltransf_31<br>Methyltransf_23<br>TIGR01983 | Methyltransferase domain<br>Methyltransferase domain<br>Methyltransferase domain<br>Methyltransferase domain<br>UbiG: 3-demethylubiquinone-9 3-O-methyltransferase | 3.30E-15<br>4.00E-10<br>6.10E-09<br>2.70E-07<br>6.60E-07 |

| | | | | | | | | |
|---|---|---|---|---|---|---|---|---|
| WP_084464122.1 | 204388 | 206239 | + | 616 | TIGR02204<br>TIGR02203<br>TIGR00958<br>TIGR03796<br>TIGR01846 | TIGR02204<br>TIGR02203<br>TIGR00958<br>TIGR03796<br>TIGR01846 | MsbA_rel: ABC transporter, permease/ATP-binding protein<br>MsbA_lipidA: lipid A export permease/ATP-binding protein MsbA<br>3a01208: antigen peptide transporter 2<br>NHLM_micro_ABC1: NHLM bacteriocin system ABC transporter, peptidase/ATP-binding protein<br>type_I_sec_HlyB: type I secretion system ATPase | 5.40E-110<br>1.60E-99<br>2.20E-97<br>6.10E-90<br>8.10E-90 |
| WP_169808938.1 | 206192 | 207791 | + | 532 | TIGR02734<br>PF01593<br>PF13450<br>TIGR02733<br>TIGR01317 | TIGR02734<br>Amino_oxidase<br>NAD_binding_8<br>TIGR02733<br>TIGR01317 | crtI_fam: phytoene desaturase<br>Flavin containing amine oxidoreductase<br>NAD(P)-binding Rossmann-like domain<br>desat_CrtD: C-3',4' desaturase CrtD<br>GOGAT_sm_gam: glutamate synthase, NADH/NADPH, small subunit | 2.60E-14<br>8.60E-11<br>3.30E-09<br>1.50E-07<br>1.00E-06 |
| WP_066941630.1 | 208538 | 207938 | - | 199 | PF01381<br>PF13560<br>PF07883<br>PF12844<br>PF13443 | HTH_3<br>HTH_31<br>Cupin_2<br>HTH_19<br>HTH_26 | Helix-turn-helix<br>Helix-turn-helix domain<br>Cupin domain<br>Helix-turn-helix domain<br>Cro/C1-type HTH DNA-binding domain | 6.80E-17<br>4.80E-16<br>2.40E-13<br>1.90E-12<br>6.30E-10 |
| WP_066941632.1 | 208815 | 208620 | - | 64 | PF13370<br>PF13459<br>PF06902 | Fer4_13<br>Fer4_15<br>Fer4_19 | 4Fe-4S single cluster domain of Ferredoxin I<br>4Fe-4S single cluster domain<br>Divergent 4Fe-4S mono-cluster | 5.70E-13<br>8.20E-13<br>4.50E-07 |
| WP_066941634.1 | 210082 | 208864 | - | 405 | TIGR04458<br>PF00067 | TIGR04458<br>p450 | CYP450_TxtE: 4-nitrotryptophan synthase<br>Cytochrome P450 | 5.70E-39<br>7.50E-24 |

| index | leader | core | start | end | dir | score |
|---|---|---|---|---|---|---|
| 1 | MSTLADHTNTISPGRFGDLTETTRGRGLTAATPVL | ATPSAVALGVAIGMAAFHAGYMYGIARSMAEGTVAN | 199753 | 199969 | + | 30 |
| 2 | MPLLAEKANADPALAAISIYSTEPCHEIGGAPVL | ATPATIVGVAAGVGLVGSVVGLFAAGYAIGHSGTLPK | 199994 | 200210 | + | 29 |
| 3 | LPGHRRPWHPGNGPMPQPSRPAAH | PTPSASRPRKEAVRPAASVRA | 200225 | 200363 | + | 7 |
| 4 | MPQPSRPAAH | PTPSASRPRKEAVRPAASVRA | 200267 | 200363 | + | 6 |
| 5 | MSTLADYTNAVSPETFGDLTETTAGRGLTAGTPVL | ATPAVVVTMVAAFAAGYAMGRSGTVPK | 200488 | 200677 | + | 33 |
| 6 | MTLLAEKVNADPAFDAISIHSTYPVHTSVEVPVL | CVTATMGAASGLSTANVGAVNGGTVLTIA | 200762 | 200954 | + | 31 |
| 7 | MKIVIDQDRCIASGQCVTAVGEVFDQRDEDGVVVLLD | PTPPAELASAVRQAAAVCPALAIRVEE | 208815 | 208620 | - | 9 |
| 8 | VHICAERERALAEGRPTPRSTWYDA | ATGLIDAIVGLWAAPVTPTPLRADPS | 210251 | 210095 | - | 7 |
| 9 | MVHICAERERALAEGRPTPRSTWYDA | ATGLIDAIVGLWAAPVTPTPLRADPS | 210254 | 210095 | - | 7 |
| 10 | LMVHICAERERALAEGRPTPRSTWYD | AATGLIDAIVGLWAAPVTPTPLRADPS | 210257 | 210095 | - | 8 |
| 11 | MTSQLMVHICAERERALAEGRPTPRSTW | YDAATGLIDAIVGLWAAPVTPTPLRADPS | 210269 | 210095 | - | 6 |

### Results for WP_157570523.1 [Microtetraspora malaysiensis] back to top

previous - next

#### Architecture

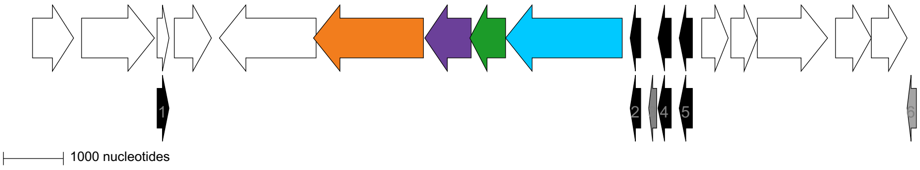

Link to nucleotide sequence

| Accession | start | end | direction | length (aa) | Pfam/HMM | name | description | E-value |
|---|---|---|---|---|---|---|---|---|
| WP_067137185.1 | 31890 | 32562 | + | 223 | PF17939<br>PF00440<br>TIGR03384<br>TIGR03968<br>TIGR03613 | TetR_C_30<br>TetR_N<br>TIGR03384<br>TIGR03968<br>TIGR03613 | Tetracyclin repressor-like, C-terminal domain<br>Bacterial regulatory proteins, tetR family<br>betaine_BetI: transcriptional repressor BetI<br>mycofact_TetR: mycofactocin system transcriptional regulator<br>RutR: pyrimidine utilization regulatory protein R | 5.30E-10<br>3.80E-08<br>1.20E-05<br>2.10E-05<br>7.10E-04 |
| WP_067136941.1 | 32695 | 33913 | + | 405 | TIGR04458<br>PF00067 | TIGR04458<br>p450 | CYP450_TxtE: 4-nitrotryptophan synthase<br>Cytochrome P450 | 2.20E-38<br>8.80E-24 |
| WP_067136943.1 | 33962 | 34157 | + | 64 | PF13370<br>PF13459<br>PF06902 | Fer4_13<br>Fer4_15<br>Fer4_19 | 4Fe-4S single cluster domain of Ferredoxin I<br>4Fe-4S single cluster domain<br>Divergent 4Fe-4S mono-cluster | 1.80E-13<br>2.00E-12<br>2.80E-06 |
| WP_067136945.1 | 34262 | 34862 | + | 199 | PF01381<br>PF13560<br>PF07883<br>PF12844<br>PF13443 | HTH_3<br>HTH_31<br>Cupin_2<br>HTH_19<br>HTH_26 | Helix-turn-helix<br>Helix-turn-helix domain<br>Cupin domain<br>Helix-turn-helix domain<br>Cro/C1-type HTH DNA-binding domain | 6.70E-17<br>2.20E-16<br>5.30E-14<br>6.60E-12<br>6.20E-10 |
| WP_169806611.1 | 36612 | 35013 | - | 532 | TIGR02734<br>PF13450<br>PF01593<br>TIGR02733<br>TIGR03467 | TIGR02734<br>NAD_binding_8<br>Amino_oxidase<br>TIGR02733<br>TIGR03467 | crtI_fam: phytoene desaturase<br>NAD(P)-binding Rossmann-like domain<br>Flavin containing amine oxidoreductase<br>desat_CrtD: C-3',4' desaturase CrtD<br>HpnE: squalene-associated FAD-dependent desaturase | 4.20E-14<br>1.10E-09<br>2.70E-09<br>1.50E-07<br>6.30E-07 |
| WP_084259215.1 | 38416 | 36565 | - | 616 | TIGR02204<br>TIGR02203<br>TIGR00958<br>TIGR03796<br>TIGR01846 | TIGR02204<br>TIGR02203<br>TIGR00958<br>TIGR03796<br>TIGR01846 | MsbA_rel: ABC transporter, permease/ATP-binding protein<br>MsbA_lipidA: lipid A export permease/ATP-binding protein MsbA<br>3a01208: antigen peptide transporter 2<br>NHLM_micro_ABC1: NHLM bacteriocin system ABC transporter, peptidase/ATP-binding protein<br>type_I_sec_HlyB: type I secretion system ATPase | 2.50E-110<br>6.50E-100<br>2.20E-98<br>2.40E-90<br>4.20E-90 |

| | | | | | | | | |
|---|---|---|---|---|---|---|---|---|
| WP_067136949.1 | 39192 | 38427 | - | 254 | PF13649 | Methyltransf_25 | Methyltransferase domain | 3.20E-15 |
| | | | | | PF08241 | Methyltransf_11 | Methyltransferase domain | 8.10E-10 |
| | | | | | PF13847 | Methyltransf_31 | Methyltransferase domain | 4.20E-09 |
| | | | | | PF08242 | Methyltransf_12 | Methyltransferase domain | 1.00E-07 |
| | | | | | PF13489 | Methyltransf_23 | Methyltransferase domain | 1.20E-07 |
| WP_067136950.1 | 39769 | 39184 | - | 194 | LinL | LinL | LinL | 2.00E-66 |
| WP_157570523.1 | 41718 | 39768 | - | 649 | LinH | LinH | LinH (Fusion of LinE/LinG) | 4.60E-256 |
| | | | | | LinE | LinE | LinE Homolog | 3.10E-102 |
| | | | | | LinG | LinG | LinG Homolog | 6.20E-99 |
| | | | | | PF12697 | Abhydrolase_6 | Alpha/beta hydrolase family | 1.90E-18 |
| | | | | | PF00561 | Abhydrolase_1 | alpha/beta hydrolase fold | 1.00E-14 |
| WP_067136954.1 | 42039 | 41847 | - | 63 | LinA | LinA | LinA (linaridin precursor) | 1.20E-13 |
| WP_067136956.1 | 42534 | 42318 | - | 71 | LinA | LinA | LinA (linaridin precursor) | 2.60E-17 |
| WP_067136958.1 | 42896 | 42680 | - | 71 | LinA | LinA | LinA (linaridin precursor) | 2.20E-14 |
| WP_067136960.1 | 43049 | 43490 | + | 146 | PF13560 | HTH_31 | Helix-turn-helix domain | 1.60E-06 |
| WP_067137187.1 | 43530 | 43983 | + | 150 | NO PFAM MATCH | - | - | - |
| WP_067136962.1 | 43986 | 45150 | + | 387 | NO PFAM MATCH | - | - | - |
| WP_067136964.1 | 45277 | 45883 | + | 201 | PF00574 | CLP_protease | Clp protease | 3.80E-66 |
| | | | | | TIGR00493 | TIGR00493 | clpP: ATP-dependent Clp endopeptidase, proteolytic subunit ClpP | 1.10E-60 |
| | | | | | TIGR00706 | TIGR00706 | SppA_dom: signal peptide peptidase SppA, 36K type | 3.10E-05 |
| | | | | | TIGR01705 | TIGR01705 | MTA/SAH-nuc-hyp: putative 5'-methylthioadenosine/S-adenosylhomocysteine nucleosidase | 4.50E-04 |
| WP_067136967.1 | 45879 | 46476 | + | 198 | TIGR00493 | TIGR00493 | clpP: ATP-dependent Clp endopeptidase, proteolytic subunit ClpP | 4.50E-76 |
| | | | | | PF00574 | CLP_protease | Clp protease | 4.80E-73 |
| | | | | | TIGR00706 | TIGR00706 | SppA_dom: signal peptide peptidase SppA, 36K type | 1.60E-06 |

| index | leader | core | start | end | dir | score |
|---|---|---|---|---|---|---|
| 1 | MKIVIDQDRCIASGQCVSAVGEVFDQRDEDGVVVLLD | PTPPADLAVAVRQAAAVCPARAIHVEE | 33962 | 34157 | + | 20 |
| 2 | MSTLVDYTNAVSPGMFGDLTETTGGRGLTAETPVL | ATPAVVVTMVAAFAAGYAMGRGAGTLPN | 42039 | 41847 | - | 33 |
| 3 | LPGCRRPWHPGNGPMSRPARPAAN | PTPSASRPRNETVRPAASVRA | 42303 | 42165 | - | 7 |
| 4 | MTLLAEKVNADPALTAISIYSTEPCHEIGGAPVL | ATPATIVGVAAGVGLVGSVVGLFAAGYAFGHSGTLPK | 42534 | 42318 | - | 29 |
| 5 | MSTLADYTNTISPGKFGDLTETTGGRGLTAATPVL | ATPSAVALGVAIGMAAFHAGYMYGIARSMAEGTVAN | 42896 | 42680 | - | 30 |
| 6 | MLHSVCDALGLPIEHLLTRAVRR | ITASRRIATHRPAGTPRVELLAA | 46620 | 46479 | - | 6 |

### Results for SJN40172.1 [Mycetocola reblochoni REB411] back to top

previous - next

#### Architecture

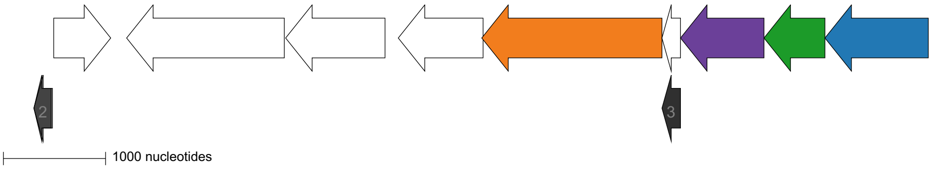

Link to nucleotide sequence

| Accession | start | end | direction | length (aa) | Pfam/HMM | name | description | E-value |
|---|---|---|---|---|---|---|---|---|
| SJN40118.1 | 1262 | 1820 | + | 185 | PF01872 | RibD_C | RibD C-terminal domain | 2.30E-18 |
| | | | | | TIGR00227 | TIGR00227 | ribD_Cterm: riboflavin-specific deaminase C-terminal domain | 1.20E-05 |
| | | | | | PF00186 | DHFR_1 | Dihydrofolate reductase | 9.20E-05 |
| SJN40126.1 | 3514 | 1978 | - | 511 | TIGR01312 | TIGR01312 | XylB: xylulokinase | 2.40E-92 |
| | | | | | PF00370 | FGGY_N | FGGY family of carbohydrate kinases, N-terminal domain | 6.20E-45 |
| | | | | | TIGR01311 | TIGR01311 | glycerol_kin: glycerol kinase | 2.80E-40 |
| | | | | | TIGR01314 | TIGR01314 | gntK_FGGY: gluconate kinase | 2.80E-39 |
| | | | | | TIGR01234 | TIGR01234 | L-ribulokinase: ribulokinase | 7.10E-39 |
| SJN40135.1 | 4505 | 3524 | - | 326 | PF04198 | Sugar-bind | Putative sugar-binding domain | 1.40E-39 |
| | | | | | PF12802 | MarR_2 | MarR family | 9.60E-07 |
| | | | | | PF04545 | Sigma70_r4 | Sigma-70, region 4 | 9.60E-06 |
| | | | | | TIGR02937 | TIGR02937 | sigma70-ECF: RNA polymerase sigma factor, sigma-70 family | 1.50E-05 |
| | | | | | PF13730 | HTH_36 | Helix-turn-helix domain | 3.70E-05 |
| SJN40142.1 | 5457 | 4632 | - | 274 | PF01551 | Peptidase_M23 | Peptidase family M23 | 1.40E-14 |
| SJN40149.1 | 7217 | 5453 | - | 587 | TIGR03375 | TIGR03375 | type_I_sec_LssB: type I secretion system ATPase | 5.80E-69 |
| | | | | | TIGR02868 | TIGR02868 | CydC: thiol reductant ABC exporter, CydC subunit | 9.10E-66 |
| | | | | | TIGR02204 | TIGR02204 | MsbA_rel: ABC transporter, permease/ATP-binding protein | 3.60E-65 |
| | | | | | TIGR02857 | TIGR02857 | CydD: thiol reductant ABC exporter, CydD subunit | 9.40E-60 |
| | | | | | TIGR02203 | TIGR02203 | MsbA_lipidA: lipid A export permease/ATP-binding protein MsbA | 3.00E-58 |
| SJN40154.1 | 7396 | 7213 | - | 60 | NO PFAM MATCH | - | - | - |
| SJN40160.1 | 8217 | 7392 | - | 274 | PF13649 | Methyltransf_25 | Methyltransferase domain | 2.50E-18 |
| | | | | | PF08241 | Methyltransf_11 | Methyltransferase domain | 4.60E-16 |
| | | | | | PF08242 | Methyltransf_12 | Methyltransferase domain | 1.00E-12 |
| | | | | | PF13847 | Methyltransf_31 | Methyltransferase domain | 2.40E-12 |
| | | | | | TIGR02072 | TIGR02072 | BioC: malonyl-acyl carrier protein O-methyltransferase BioC | 3.70E-12 |
| SJN40166.1 | 8804 | 8213 | - | 196 | LinL | LinL | LinL | 5.50E-50 |

| | | | | | | | | |
|---|---|---|---|---|---|---|---|---|
| SJN40172.1 | 9820 | 8803 | - | 338 | LinE<br>LinH<br>PF12697<br>PF12146<br>PF00561 | LinE<br>LinH<br>Abhydrolase_6<br>Hydrolase_4<br>Abhydrolase_1 | LinE Homolog<br>LinH (Fusion of LinE/LinG)<br>Alpha/beta hydrolase family<br>Serine aminopeptidase, S33<br>alpha/beta hydrolase fold | 2.30E-66<br>5.80E-58<br>2.00E-10<br>1.30E-06<br>2.60E-06 |

| index | leader | core | start | end | dir | score |
|---|---|---|---|---|---|---|
| 1 | VRRTGSEAAGGPAGGGRDPAAARM | AAPVRASKSTSAAATGSPSRGSPASASSAAR | 1235 | 1067 | - | 7 |
| 2 | VSSVRRTGSEAAGGPAGGGRDPAAARM | AAPVRASKSTSAAATGSPSRGSPASASSAAR | 1244 | 1067 | - | 7 |
| 3 | MRFTTAIVTALMVFSIYFIMF | ATPDGADMTRQGVVAAIVAVVGGAIAYAITRPRQKDSSQ | 7396 | 7213 | - | 9 |

### Results for WP_121689029.1 [Mycetocola lacteus] back to top

previous - next

#### Architecture

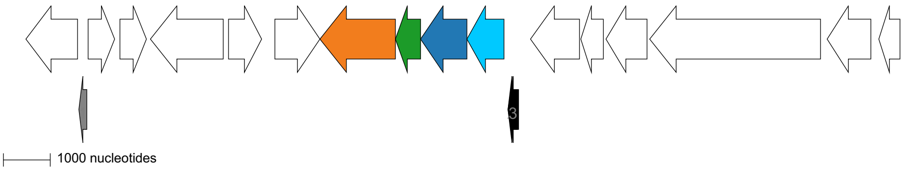

Link to nucleotide sequence

| Accession | start | end | direction | length (aa) | Pfam/HMM | name | description | E-value |
|---|---|---|---|---|---|---|---|---|
| WP_121689023.1 | 43673 | 42584 | - | 362 | NO PFAM MATCH | - | - | - |
| WP_121689024.1 | 43913 | 44465 | + | 183 | NO PFAM MATCH | - | - | - |
| WP_121689025.1 | 44575 | 45157 | + | 193 | NO PFAM MATCH | - | - | - |
| WP_121689026.1 | 46796 | 45239 | - | 518 | TIGR00959<br>TIGR01425<br>PF00448<br>TIGR00064<br>PF02978 | TIGR00959<br>TIGR01425<br>SRP54<br>TIGR00064<br>SRP_SPB | ffh: signal recognition particle protein<br>SRP54_euk: signal recognition particle protein SRP54<br>SRP54-type protein, GTPase domain<br>ftsY: signal recognition particle-docking protein FtsY<br>Signal peptide binding domain | 3.10E-172<br>2.10E-91<br>6.40E-69<br>6.60E-62<br>6.50E-33 |
| WP_121689027.1 | 46907 | 47618 | + | 236 | TIGR00214<br>PF03099 | TIGR00214<br>BPL_LplA_LipB | lipB: lipoyl(octanoyl) transferase<br>Biotin/lipoate A/B protein ligase family | 3.50E-37<br>7.90E-09 |
| WP_121689093.1 | 47899 | 48865 | + | 321 | TIGR00510<br>PF04055<br>TIGR00433<br>PF16881<br>TIGR03910 | TIGR00510<br>Radical_SAM<br>TIGR00433<br>LIAS_N<br>TIGR03910 | lipA: lipoyl synthase<br>Radical SAM superfamily<br>bioB: biotin synthase<br>N-terminal domain of lipoyl synthase of Radical_SAM family<br>pyrrolys_PylB: pyrrolysine biosynthesis radical SAM protein | 4.60E-95<br>3.20E-10<br>3.70E-09<br>1.90E-06<br>7.20E-05 |
| WP_121689028.1 | 50472 | 48861 | - | 536 | TIGR03375<br>TIGR02868<br>TIGR02204<br>TIGR02203<br>TIGR02857 | TIGR03375<br>TIGR02868<br>TIGR02204<br>TIGR02203<br>TIGR02857 | type_I_sec_LssB: type I secretion system ATPase<br>CydC: thiol reductant ABC exporter, CydC subunit<br>MsbA_rel: ABC transporter, permease/ATP-binding protein<br>MsbA_lipidA: lipid A export permease/ATP-binding protein MsbA<br>CydD: thiol reductant ABC exporter, CydD subunit | 6.50E-61<br>4.50E-59<br>3.90E-58<br>1.00E-57<br>1.50E-53 |
| WP_147441693.1 | 51010 | 50464 | - | 181 | LinL | LinL | LinL | 7.00E-37 |
| WP_121689029.1 | 51986 | 50999 | - | 328 | LinE<br>LinH<br>PF12697<br>PF12146<br>PF00561 | LinE<br>LinH<br>Abhydrolase_6<br>Hydrolase_4<br>Abhydrolase_1 | LinE Homolog<br>LinH (Fusion of LinE/LinG)<br>Alpha/beta hydrolase family<br>Serine aminopeptidase, S33<br>alpha/beta hydrolase fold | 1.20E-57<br>4.10E-49<br>8.80E-14<br>8.60E-13<br>1.70E-10 |
| WP_147441694.1 | 52789 | 51982 | - | 268 | LinH<br>LinG<br>PF05090 | LinH<br>LinG<br>VKG_Carbox | LinH (Fusion of LinE/LinG)<br>LinG Homolog<br>Vitamin K-dependent gamma-carboxylase | 8.20E-63<br>4.00E-61<br>4.10E-05 |
| WP_121689030.1 | 54399 | 53358 | - | 346 | PF00583<br>PF13508<br>PF13673<br>TIGR01575<br>PF08445 | Acetyltransf_1<br>Acetyltransf_7<br>Acetyltransf_10<br>TIGR01575<br>FR47 | Acetyltransferase (GNAT) family<br>Acetyltransferase (GNAT) domain<br>Acetyltransferase (GNAT) domain<br>rimI: ribosomal-protein-alanine acetyltransferase<br>FR47-like protein | 1.70E-23<br>6.50E-19<br>6.60E-16<br>3.50E-10<br>8.20E-07 |
| WP_121689031.1 | 54906 | 54408 | - | 165 | PF09406 | DUF2004 | Protein of unknown function (DUF2004) | 1.50E-09 |
| WP_121689032.1 | 55844 | 54968 | - | 291 | TIGR00064<br>PF00448<br>TIGR00959<br>TIGR01425<br>TIGR03499 | TIGR00064<br>SRP54<br>TIGR00959<br>TIGR01425<br>TIGR03499 | ftsY: signal recognition particle-docking protein FtsY<br>SRP54-type protein, GTPase domain<br>ffh: signal recognition particle protein<br>SRP54_euk: signal recognition particle protein SRP54<br>FlhF: flagellar biosynthesis protein FlhF | 2.00E-92<br>9.20E-69<br>3.80E-61<br>1.60E-45<br>1.40E-21 |
| WP_121689033.1 | 59542 | 55888 | - | 1217 | TIGR02168<br>TIGR02169<br>PF02463<br>PF13304<br>PF13476 | TIGR02168<br>TIGR02169<br>SMC_N<br>AAA_21<br>AAA_23 | SMC_prok_B: chromosome segregation protein SMC<br>SMC_prok_A: chromosome segregation protein SMC<br>RecF/RecN/SMC N terminal domain<br>AAA domain, putative AbiEii toxin, Type IV TA system<br>AAA domain | 2.90E-252<br>8.10E-129<br>2.30E-71<br>2.10E-17<br>7.40E-17 |
| WP_121689034.1 | 60610 | 59665 | - | 314 | TIGR02824<br>TIGR02823<br>PF00107<br>PF13602<br>PF08240 | TIGR02824<br>TIGR02823<br>ADH_zinc_N<br>ADH_zinc_N_2<br>ADH_N | quinone_pig3: putative NAD(P)H quinone oxidoreductase, PIG3 family<br>oxido_YhdH: putative quinone oxidoreductase, YhdH/YhfP family<br>Zinc-binding dehydrogenase<br>Zinc-binding dehydrogenase<br>Alcohol dehydrogenase GroES-like domain | 1.50E-24<br>6.40E-10<br>5.50E-09<br>1.20E-08<br>3.10E-06 |
| WP_121689035.1 | 61238 | 60776 | - | 153 | NO PFAM MATCH | - | - | - |

| index | leader | core | start | end | dir | score |
|---|---|---|---|---|---|---|

| | | | | | | |
|---|---|---|---|---|---|---|
| 1 | LPPIHGARRRRLGPCYREWGAGGPSTTSVIKLTRASV | ATTLRPCPRGSGTQPGV | 43883 | 43718 | - | 7 |
| 2 | VNQDQRSGKIVCNSSKLSSGRSVPIM | ATPATIVTVALVAARAASRTVTRRAVRRTQRGAEPLI | 53061 | 52869 | - | 21 |
| 3 | MDVLTHSVNQDQRSGKIVCNSSKLSSGRSVPIM | ATPATIVTVALVAARAASRTVTRRAVRRTQRGAEPLI | 53082 | 52869 | - | 21 |

### Results for WP_152730243.1 [Modestobacter sp. CPCC 205119] back to top

previous - next

#### Architecture

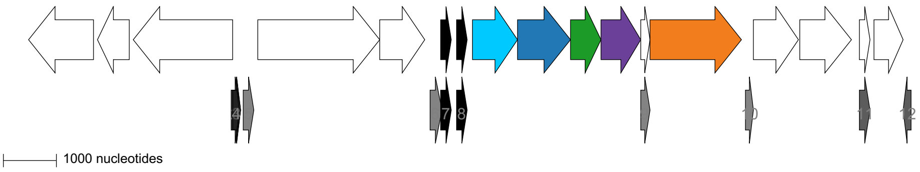

Link to nucleotide sequence

| Accession | start | end | direction | length (aa) | Pfam/HMM | name | description | E-value |
|---|---|---|---|---|---|---|---|---|
| WP_152730236.1 | 11982 | 10773 | - | 402 | TIGR02349<br>PF01556<br>PF00226<br>TIGR03835<br>PF00684 | TIGR02349<br>DnaJ_C<br>DnaJ<br>TIGR03835<br>DnaJ_CXXCXGXG | DnaJ_bact: chaperone protein DnaJ<br>DnaJ C terminal domain<br>DnaJ domain<br>termin_org_DnaJ: terminal organelle assembly protein TopJ<br>DnaJ central domain | 6.40E-116<br>9.10E-37<br>6.10E-24<br>6.70E-23<br>5.20E-13 |
| WP_152730237.1 | 12661 | 12064 | - | 198 | PF01025 | GrpE | GrpE | 4.60E-29 |
| WP_152730238.1 | 14600 | 12734 | - | 621 | TIGR02350<br>PF00012<br>TIGR01991<br>PF06723<br>TIGR00904 | TIGR02350<br>HSP70<br>TIGR01991<br>MreB_Mbl<br>TIGR00904 | prok_dnaK: chaperone protein DnaK<br>Hsp70 protein<br>HscA: Fe-S protein assembly chaperone HscA<br>MreB/Mbl protein<br>mreB: cell shape determining protein, MreB/Mrl family | 4.20E-274<br>3.80E-243<br>6.00E-169<br>1.30E-17<br>2.60E-17 |
| WP_162393318.1 | 15078 | 17379 | + | 766 | PF02754<br>PF13183<br>TIGR03379<br>PF13237<br>TIGR03290 | CCG<br>Fer4_8<br>TIGR03379<br>Fer4_10<br>TIGR03290 | Cysteine-rich domain<br>4Fe-4S dicluster domain<br>glycerol3P_GlpC: glycerol-3-phosphate dehydrogenase, anaerobic, C subunit<br>4Fe-4S dicluster domain<br>CoB_CoM_SS_C: CoB--CoM heterodisulfide reductase, subunit C | 1.90E-35<br>3.00E-12<br>4.20E-11<br>3.70E-10<br>6.10E-10 |
| WP_152730239.1 | 17392 | 18235 | + | 280 | PF01551 | Peptidase_M23 | Peptidase family M23 | 1.60E-10 |
| WP_152730240.1 | 18537 | 18747 | + | 69 | LinA | LinA | LinA (linaridin precursor) | 1.30E-09 |
| WP_152730241.1 | 18829 | 19024 | + | 64 | LinA | LinA | LinA (linaridin precursor) | 1.50E-09 |
| WP_152730242.1 | 19132 | 19978 | + | 281 | LinH<br>LinG | LinH<br>LinG | LinH (Fusion of LinE/LinG)<br>LinG Homolog | 3.50E-103<br>2.80E-98 |
| WP_152730243.1 | 19974 | 20985 | + | 336 | LinE<br>LinH<br>PF12697<br>PF12146<br>PF00561 | LinE<br>LinH<br>Abhydrolase_6<br>Hydrolase_4<br>Abhydrolase_1 | LinE Homolog<br>LinH (Fusion of LinE/LinG)<br>Alpha/beta hydrolase family<br>Serine aminopeptidase, S33<br>alpha/beta hydrolase fold | 9.00E-105<br>8.30E-73<br>1.20E-14<br>3.20E-10<br>7.00E-08 |
| WP_152730244.1 | 20981 | 21563 | + | 193 | LinL | LinL | LinL | 6.10E-58 |
| WP_152730245.1 | 21559 | 22315 | + | 251 | PF13649<br>PF08241<br>PF13489<br>PF08242<br>TIGR01934 | Methyltransf_25<br>Methyltransf_11<br>Methyltransf_23<br>Methyltransf_12<br>TIGR01934 | Methyltransferase domain<br>Methyltransferase domain<br>Methyltransferase domain<br>Methyltransferase domain<br>MenG_MenH_UbiE: ubiquinone/menaquinone biosynthesis methyltransferase | 8.60E-13<br>2.00E-10<br>6.50E-07<br>2.20E-06<br>2.80E-06 |
| WP_162393317.1 | 22308 | 22485 | + | 58 | NO PFAM MATCH | - | - | - |
| WP_152730246.1 | 22481 | 24206 | + | 574 | TIGR02203<br>TIGR02204<br>TIGR03375<br>TIGR00958<br>TIGR01846 | TIGR02203<br>TIGR02204<br>TIGR03375<br>TIGR00958<br>TIGR01846 | MsbA_lipidA: lipid A export permease/ATP-binding protein MsbA<br>MsbA_rel: ABC transporter, permease/ATP-binding protein<br>type_I_sec_LssB: type I secretion system ATPase<br>3a01208: antigen peptide transporter 2<br>type_I_sec_HlyB: type I secretion system ATPase | 1.30E-102<br>6.60E-99<br>3.60E-88<br>1.30E-87<br>2.30E-81 |
| WP_152730247.1 | 24436 | 25294 | + | 285 | PF01551 | Peptidase_M23 | Peptidase family M23 | 6.50E-13 |
| WP_152730248.1 | 25321 | 26290 | + | 322 | PF00196<br>TIGR03541 | GerE<br>TIGR03541 | Bacterial regulatory proteins, luxR family<br>reg_near_HchA: LuxR family transcriptional regulatory, chaperone HchA-associated | 1.50E-05<br>4.00E-04 |
| WP_152730249.1 | 26433 | 26634 | + | 66 | NO PFAM MATCH | - | - | - |
| WP_152730250.1 | 26710 | 27274 | + | 187 | NO PFAM MATCH | - | - | - |

| index | leader | core | start | end | dir | score |
|---|---|---|---|---|---|---|
| 1 | MPTARAMGDGLSSVCVVVLAGELGGRGHSPGVPGHCDPVP | RAPCADRAQLSCPP | 14580 | 14745 | + | 7 |
| 2 | VPWVMASLPCAWWCWPGSWAAGGTPLGSRGTATLSPGR | LAPTALNFPAHRNGMP | 14593 | 14758 | + | 6 |
| 3 | VMASLPCAWWCWPGSWAAGGTPLGSRGTATLSPGR | LAPTALNFPAHRNGMP | 14602 | 14758 | + | 6 |
| 4 | MASLPCAWWCWPGSWAAGGTPLGSRGTATLSPGR | LAPTALNFPAHRNGMP | 14605 | 14758 | + | 6 |
| 5 | LLARPAQPPAPRSPAVSPHRTHRRRPRDPR | ATLTRTPGDVPEPGGAARLTGSARHEVVTGE | 14815 | 15001 | + | 7 |
| 6 | VRQPAGAARAHTRRPRALRRCPGPDTDGRPCPDPDVDRRPRCPWGVSP | STVEVTATVATDPDSETEKDTCCWTR | 18331 | 18556 | + | 7 |
| 7 | MLLDKVNAEPALSAALSPATATGTLVPVM | ATPAAFAAIGAAGAAKAAGAVVGAGAVTGAAYVAYKVATG | 18537 | 18747 | + | 31 |

| | | | | | | |
|---|---|---|---|---|---|---|
| 8 | MLLNKVNAEPSLRRSLSPGIPVLTASPVM | ATPAAFAAGVVAGAKASAILTGAAGVGVAVKTAQK | 18829 | 19024 | + | 29 |
| 9 | MLTRRRSTSFVVAFFFALPVL | AVVLRDESSGAVPVTAVLGALGVAVVLAAGLGGQERS | 22308 | 22485 | + | 7 |
| 10 | VPTIDTADLDDAISSAFQELRTAMR | SFNRDSIQLYSAALEALVKLRQVTP | 24287 | 24440 | + | 7 |
| 11 | MSRTAMLLVSGLGAGVLGLTGELRP | ETAQHSSSAAVVTSAAPAAPGVPPAAPRLAPARQLCLVCWL | 26433 | 26634 | + | 8 |
| 12 | VRAAAVGGEQLPHGLGPVGGHGDPQ | VEQHAVTVPDRAARPGDAGCAVTPAS | 27413 | 27257 | - | 8 |

### Results for RLP71106.1 [Mycetocola reblochoni] back to top

previous - next

#### Architecture

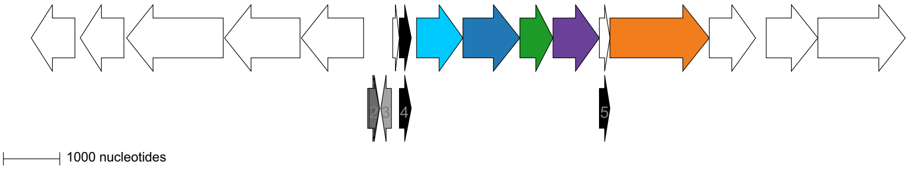

Link to nucleotide sequence

| Accession | start | end | direction | length (aa) | Pfam/HMM | name | description | E-value |
|---|---|---|---|---|---|---|---|---|
| RLP71100.1 | 277060 | 276280 | - | 259 | PF13561 | adh_short_C2 | Enoyl-(Acyl carrier protein) reductase | 3.60E-62 |
| | | | | | TIGR01830 | TIGR01830 | 3oxo_ACP_reduc: 3-oxoacyl-[acyl-carrier-protein] reductase | 1.00E-54 |
| | | | | | TIGR01963 | TIGR01963 | PHB_DH: 3-hydroxybutyrate dehydrogenase | 1.20E-54 |
| | | | | | PF00106 | adh_short | short chain dehydrogenase | 1.80E-51 |
| | | | | | TIGR01832 | TIGR01832 | kduD: 2-deoxy-D-gluconate 3-dehydrogenase | 1.10E-49 |
| RLP71101.1 | 277934 | 277154 | - | 259 | PF13561 | adh_short_C2 | Enoyl-(Acyl carrier protein) reductase | 1.60E-62 |
| | | | | | TIGR01963 | TIGR01963 | PHB_DH: 3-hydroxybutyrate dehydrogenase | 7.70E-56 |
| | | | | | TIGR01830 | TIGR01830 | 3oxo_ACP_reduc: 3-oxoacyl-[acyl-carrier-protein] reductase | 2.50E-53 |
| | | | | | PF00106 | adh_short | short chain dehydrogenase | 1.70E-52 |
| | | | | | TIGR02415 | TIGR02415 | 23BDH: acetoin reductases | 1.00E-47 |
| RLP71102.1 | 279679 | 277969 | - | 569 | PF02733 | Dak1 | Dak1 domain | 3.10E-98 |
| | | | | | TIGR02361 | TIGR02361 | dak_ATP: dihydroxyacetone kinase | 2.90E-95 |
| | | | | | TIGR02363 | TIGR02363 | dhaK1: dihydroxyacetone kinase, DhaK subunit | 9.20E-83 |
| | | | | | TIGR02362 | TIGR02362 | dhaK1b: probable dihydroxyacetone kinase DhaK1b subunit | 3.30E-59 |
| | | | | | PF02734 | Dak2 | DAK2 domain | 5.90E-42 |
| RLP71103.1 | 281045 | 279710 | - | 444 | TIGR00897 | TIGR00897 | 2A0118: polyol permease | 2.50E-92 |
| | | | | | PF07690 | MFS_1 | Major Facilitator Superfamily | 1.60E-24 |
| | | | | | TIGR00880 | TIGR00880 | 2_A_01_02: multidrug resistance protein | 8.70E-07 |
| | | | | | TIGR00881 | TIGR00881 | 2A0104: phosphoglycerate transporter family protein | 1.30E-04 |
| RLP71282.1 | 282180 | 281079 | - | 366 | TIGR00692 | TIGR00692 | tdh: L-threonine 3-dehydrogenase | 8.90E-64 |
| | | | | | TIGR03451 | TIGR03451 | mycoS_dep_FDH: S-(hydroxymethyl)mycothiol dehydrogenase | 1.60E-35 |
| | | | | | TIGR03989 | TIGR03989 | Rxyl_3153: NDMA-dependent alcohol dehydrogenase, Rxyl_3153 family | 1.50E-32 |
| | | | | | TIGR03366 | TIGR03366 | HpnZ_proposed: putative phosphonate catabolism associated alcohol dehydrogenase | 9.70E-31 |
| | | | | | PF08240 | ADH_N | Alcohol dehydrogenase GroES-like domain | 8.60E-24 |
| WP_014850790.1 | 282695 | 282809 | + | 38 | INFERRED GENE | - | - | - |
| RLP71104.1 | 282813 | 283020 | + | 68 | LinA | LinA | LinA (linaridin precursor) | 5.30E-12 |
| RLP71105.1 | 283116 | 283941 | + | 274 | LinH | LinH | LinH (Fusion of LinE/LinG) | 4.90E-84 |
| | | | | | LinG | LinG | LinG Homolog | 4.00E-79 |
| | | | | | PF05090 | VKG_Carbox | Vitamin K-dependent gamma-carboxylase | 2.60E-04 |
| RLP71106.1 | 283937 | 284954 | + | 338 | LinE | LinE | LinE Homolog | 2.30E-66 |
| | | | | | LinH | LinH | LinH (Fusion of LinE/LinG) | 5.80E-58 |
| | | | | | PF12697 | Abhydrolase_6 | Alpha/beta hydrolase family | 2.00E-10 |
| | | | | | PF12146 | Hydrolase_4 | Serine aminopeptidase, S33 | 1.30E-06 |
| | | | | | PF00561 | Abhydrolase_1 | alpha/beta hydrolase fold | 2.60E-06 |
| RLP71107.1 | 284953 | 285544 | + | 196 | LinL | LinL | LinL | 5.50E-50 |
| RLP71108.1 | 285540 | 286365 | + | 274 | PF13649 | Methyltransf_25 | Methyltransferase domain | 2.50E-18 |
| | | | | | PF08241 | Methyltransf_11 | Methyltransferase domain | 4.60E-16 |
| | | | | | PF08242 | Methyltransf_12 | Methyltransferase domain | 1.00E-12 |
| | | | | | PF13847 | Methyltransf_31 | Methyltransferase domain | 2.40E-12 |
| | | | | | TIGR02072 | TIGR02072 | BioC: malonyl-acyl carrier protein O-methyltransferase BioC | 3.70E-12 |
| RLP71109.1 | 286361 | 286544 | + | 60 | NO PFAM MATCH | - | - | - |
| RLP71110.1 | 286540 | 288304 | + | 587 | TIGR03375 | TIGR03375 | type_I_sec_LssB: type I secretion system ATPase | 5.80E-69 |
| | | | | | TIGR02868 | TIGR02868 | CydC: thiol reductant ABC exporter, CydC subunit | 9.10E-66 |
| | | | | | TIGR02204 | TIGR02204 | MsbA_rel: ABC transporter, permease/ATP-binding protein | 3.60E-65 |
| | | | | | TIGR02857 | TIGR02857 | CydD: thiol reductant ABC exporter, CydD subunit | 9.40E-60 |
| | | | | | TIGR02203 | TIGR02203 | MsbA_lipidA: lipid A export permease/ATP-binding protein MsbA | 3.00E-58 |
| RLP71111.1 | 288300 | 289125 | + | 274 | PF01551 | Peptidase_M23 | Peptidase family M23 | 1.40E-14 |
| RLP71283.1 | 289327 | 290233 | + | 301 | PF04198 | Sugar-bind | Putative sugar-binding domain | 1.10E-39 |
| | | | | | PF12802 | MarR_2 | MarR family | 5.90E-06 |
| | | | | | PF04545 | Sigma70_r4 | Sigma-70, region 4 | 1.60E-05 |
| | | | | | PF13730 | HTH_36 | Helix-turn-helix domain | 3.10E-05 |
| | | | | | PF13545 | HTH_Crp_2 | Crp-like helix-turn-helix domain | 1.80E-04 |
| RLP71112.1 | 290243 | 291779 | + | 511 | TIGR01312 | TIGR01312 | XylB: xylulokinase | 2.40E-92 |
| | | | | | PF00370 | FGGY_N | FGGY family of carbohydrate kinases, N-terminal domain | 6.20E-45 |
| | | | | | TIGR01311 | TIGR01311 | glycerol_kin: glycerol kinase | 2.80E-40 |
| | | | | | TIGR01314 | TIGR01314 | gntK_FGGY: gluconate kinase | 2.80E-39 |
| | | | | | TIGR01234 | TIGR01234 | L-ribulokinase: ribulokinase | 7.10E-39 |

| index | leader | core | start | end | dir | score |
|---|---|---|---|---|---|---|
| 1 | MLAHPEHLCSTHVQLDGVETRSIIDAHEMARFTSCICIQFTI | PTIPQLPPQCPRDYLGKASKNSASAESP | 282247 | 282460 | + | 6 |
| 2 | VQLDGVETRSIIDAHEMARFTSCICIQFTI | PTIPQLPPQCPRDYLGKASKNSASAESP | 282283 | 282460 | + | 6 |
| 3 | VCVPANYPHRGANEGACLHLLGSRDPSGLPF | VVLSEFESSREDRPALVFPIRTATCAAISPEA | 282671 | 282479 | - | 6 |
| 4 | MSVAELSATNRLLGMVNSDTVRTDAVVSPVM | ATPAAFAAGIVAGGKATAVVVGAAGVGAAIGNAVGGK | 282813 | 283020 | + | 30 |
| 5 | MRFTTAIVTALMVFSIYFIMF | ATPDGADMTRQGVVAAIVAVVGGAIAYAITRPRQKDSSQ | 286361 | 286544 | + | 19 |

### Results for WP_121649883.1 [Mycetocola tolaasinivorans] back to top

previous - next

#### Architecture

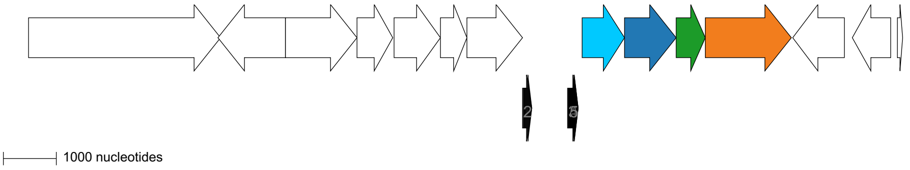

Link to nucleotide sequence

| Accession | start | end | direction | length (aa) | Pfam/HMM | name | description | E-value |
|---|---|---|---|---|---|---|---|---|
| WP_121649876.1 | 10034 | 13640 | + | 1201 | TIGR02168<br>TIGR02169<br>PF02463<br>PF13304<br>PF13476 | TIGR02168<br>TIGR02169<br>SMC_N<br>AAA_21<br>AAA_23 | SMC_prok_B: chromosome segregation protein SMC<br>SMC_prok_A: chromosome segregation protein SMC<br>RecF/RecN/SMC N terminal domain<br>AAA domain, putative AbiEii toxin, Type IV TA system<br>AAA domain | 5.20E-257<br>6.70E-131<br>6.90E-71<br>7.50E-17<br>7.80E-17 |
| WP_121649877.1 | 14885 | 13589 | - | 431 | PF13439<br>PF13579<br>PF13692<br>TIGR03088<br>TIGR04063 | Glyco_transf_4<br>Glyco_trans_4_4<br>Glyco_trans_1_4<br>TIGR03088<br>TIGR04063 | Glycosyltransferase Family 4<br>Glycosyl transferase 4-like domain<br>Glycosyl transferases group 1<br>stp2: sugar transferase, PEP-CTERM/EpsH1 system associated<br>stp3: PEP-CTERM/exosortase A-associated glycosyltransferase, Daro_2409 family | 1.70E-20<br>1.90E-13<br>2.20E-09<br>8.60E-07<br>7.80E-05 |
| WP_121649878.1 | 14865 | 16233 | + | 455 | PF07730 | HisKA_3 | Histidine kinase | 8.90E-16 |
| WP_121649879.1 | 16229 | 16892 | + | 220 | PF00072<br>TIGR02875<br>PF00196<br>TIGR03020<br>TIGR02154 | Response_reg<br>TIGR02875<br>GerE<br>TIGR03020<br>TIGR02154 | Response regulator receiver domain<br>spore_0_A: sporulation transcription factor Spo0A<br>Bacterial regulatory proteins, luxR family<br>EpsA: transcriptional regulator EpsA<br>PhoB: phosphate regulon transcriptional regulatory protein PhoB | 1.70E-23<br>3.70E-23<br>1.10E-16<br>2.00E-11<br>8.90E-10 |
| WP_121649880.1 | 16932 | 17808 | + | 291 | TIGR00064<br>PF00448<br>TIGR00959<br>TIGR01425<br>TIGR03499 | TIGR00064<br>SRP54<br>TIGR00959<br>TIGR01425<br>TIGR03499 | ftsY: signal recognition particle-docking protein FtsY<br>SRP54-type protein, GTPase domain<br>ffh: signal recognition particle protein<br>SRP54_euk: signal recognition particle protein SRP54<br>FlhF: flagellar biosynthesis protein FlhF | 8.30E-93<br>3.40E-69<br>1.10E-61<br>2.00E-46<br>1.60E-21 |
| WP_121649881.1 | 17810 | 18308 | + | 165 | PF09406 | DUF2004 | Protein of unknown function (DUF2004) | 1.10E-09 |
| WP_121649882.1 | 18317 | 19358 | + | 346 | PF00583<br>PF13508<br>PF13673<br>TIGR01575<br>PF13527 | Acetyltransf_1<br>Acetyltransf_7<br>Acetyltransf_10<br>TIGR01575<br>Acetyltransf_9 | Acetyltransferase (GNAT) family<br>Acetyltransferase (GNAT) domain<br>Acetyltransferase (GNAT) domain<br>rimI: ribosomal-protein-alanine acetyltransferase<br>Acetyltransferase (GNAT) domain | 5.70E-26<br>7.70E-20<br>6.40E-16<br>4.40E-10<br>1.70E-06 |
| WP_147440512.1 | 20479 | 21286 | + | 268 | LinH<br>LinG<br>PF05090 | LinH<br>LinG<br>VKG_Carbox | LinH (Fusion of LinE/LinG)<br>LinG Homolog<br>Vitamin K-dependent gamma-carboxylase | 1.10E-62<br>1.20E-60<br>2.00E-04 |
| WP_121649883.1 | 21282 | 22269 | + | 328 | LinE<br>LinH<br>PF00561<br>PF12697<br>PF12146 | LinE<br>LinH<br>Abhydrolase_1<br>Abhydrolase_6<br>Hydrolase_4 | LinE Homolog<br>LinH (Fusion of LinE/LinG)<br>alpha/beta hydrolase fold<br>Alpha/beta hydrolase family<br>Serine aminopeptidase, S33 | 8.80E-60<br>1.90E-54<br>3.70E-14<br>1.00E-13<br>1.30E-13 |
| WP_147440513.1 | 22258 | 22804 | + | 181 | LinL | LinL | LinL | 1.30E-37 |
| WP_121649884.1 | 22800 | 24426 | + | 541 | TIGR02204<br>TIGR03375<br>TIGR02868<br>TIGR02857<br>TIGR00958 | TIGR02204<br>TIGR03375<br>TIGR02868<br>TIGR02857<br>TIGR00958 | MsbA_rel: ABC transporter, permease/ATP-binding protein<br>type_I_sec_LssB: type I secretion system ATPase<br>CydC: thiol reductant ABC exporter, CydC subunit<br>CydD: thiol reductant ABC exporter, CydD subunit<br>3a01208: antigen peptide transporter 2 | 5.40E-58<br>9.50E-58<br>8.90E-57<br>2.80E-52<br>1.60E-46 |
| WP_121649888.1 | 25428 | 24465 | - | 320 | TIGR00510<br>PF04055<br>TIGR00433<br>PF16881<br>TIGR03910 | TIGR00510<br>Radical_SAM<br>TIGR00433<br>LIAS_N<br>TIGR03910 | lipA: lipoyl synthase<br>Radical SAM superfamily<br>bioB: biotin synthase<br>N-terminal domain of lipoyl synthase of Radical_SAM family<br>pyrrolys_PylB: pyrrolysine biosynthesis radical SAM protein | 1.20E-95<br>1.20E-09<br>2.90E-09<br>1.50E-06<br>3.60E-05 |
| WP_121649885.1 | 26315 | 25589 | - | 241 | TIGR00214<br>PF03099 | TIGR00214<br>BPL_LplA_LipB | lipB: lipoyl(octanoyl) transferase<br>Biotin/lipoate A/B protein ligase family | 1.10E-35<br>2.40E-08 |
| WP_147440514.1 | 26426 | 26550 | + | 41 | PF02881<br>TIGR00959 | SRP54_N<br>TIGR00959 | SRP54-type protein, helical bundle domain<br>ffh: signal recognition particle protein | 2.80E-06<br>1.30E-05 |

| index | leader | core | start | end | dir | score |
|---|---|---|---|---|---|---|
| 1 | LLAAGAGAGAGVDAACVDVGADANVGFLAGFEQGEASM | STPVSISLLGPGPVF | 19368 | 19530 | + | 9 |
| 2 | LAAGAGAGAGVDAACVDVGADANVGFLAGFEQGEASM | STPVSISLLGPGPVF | 19371 | 19530 | + | 9 |

| | | | | | | |
|---|---|---|---|---|---|---|
| 3 | MDNLTRLVNEDELAGKLPRHEYDQAGRRSVPIM | ATPATIITVALVAARAASRTVTRRAVARTQRGLETAV | 20195 | 20408 | + | 31 |
| 4 | VPIMATPATIITVALVAARA | ASRTVTRRAVARTQRGLETAV | 20282 | 20408 | + | 16 |
| 5 | MATPATIITVALVAARAAS | RTVTRRAVARTQRGLETAV | 20291 | 20408 | + | 6 |

### Results for WP_111766294.1 [Nakamurella deserti] back to top

previous - next

#### Architecture

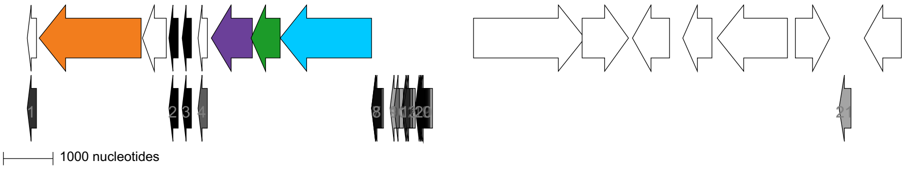

Link to nucleotide sequence

| Accession | start | end | direction | length (aa) | Pfam/HMM | name | description | E-value |
|---|---|---|---|---|---|---|---|---|
| WP_157970581.1 | 1853269 | 1853098 | - | 56 | PF05433<br>PF13441 | Rick_17kDa_Anti<br>Gly-zipper_YMGG | Glycine zipper 2TM domain<br>YMGG-like Gly-zipper | 1.80E-05<br>4.80E-05 |
| WP_111766288.1 | 1855376 | 1853324 | - | 683 | TIGR02204<br>TIGR00958<br>TIGR02203<br>TIGR03796<br>TIGR01846 | TIGR02204<br>TIGR00958<br>TIGR02203<br>TIGR03796<br>TIGR01846 | MsbA_rel: ABC transporter, permease/ATP-binding protein<br>3a01208: antigen peptide transporter 2<br>MsbA_lipidA: lipid A export permease/ATP-binding protein MsbA<br>NHLM_micro_ABC1: NHLM bacteriocin system ABC transporter, peptidase/ATP-binding protein<br>type_I_sec_HlyB: type I secretion system ATPase | 4.50E-100<br>1.20E-96<br>1.10E-94<br>8.90E-84<br>2.00E-77 |
| WP_111766289.1 | 1855884 | 1855413 | - | 156 | PF04134 | DUF393 | Protein of unknown function, DUF393 | 1.20E-15 |
| WP_111766290.1 | 1856132 | 1855949 | - | 60 | LinA | LinA | LinA (linaridin precursor) | 1.90E-11 |
| WP_111766291.1 | 1856413 | 1856221 | - | 63 | LinA | LinA | LinA (linaridin precursor) | 1.50E-17 |
| WP_111766292.1 | 1856712 | 1856526 | - | 61 | NO PFAM MATCH | - | - | - |
| WP_157970582.1 | 1857622 | 1856812 | - | 269 | PF13649<br>PF08241<br>TIGR01983<br>PF13847<br>PF13489 | Methyltransf_25<br>Methyltransf_11<br>TIGR01983<br>Methyltransf_31<br>Methyltransf_23 | Methyltransferase domain<br>Methyltransferase domain<br>UbiG: 3-demethylubiquinone-9 3-O-methyltransferase<br>Methyltransferase domain<br>Methyltransferase domain | 1.30E-14<br>1.90E-12<br>5.50E-08<br>1.90E-05<br>5.80E-05 |
| WP_157970583.1 | 1858182 | 1857615 | - | 188 | LinL | LinL | LinL | 4.10E-62 |
| WP_111766294.1 | 1860023 | 1858178 | - | 614 | LinH<br>LinG<br>LinE<br>PF12697<br>PF00561 | LinH<br>LinG<br>LinE<br>Abhydrolase_6<br>Abhydrolase_1 | LinH (Fusion of LinE/LinG)<br>LinG Homolog<br>LinE Homolog<br>Alpha/beta hydrolase family<br>alpha/beta hydrolase fold | 3.90E-227<br>1.40E-97<br>2.30E-90<br>5.70E-15<br>2.10E-13 |
| WP_170315549.1 | 1862095 | 1864330 | + | 744 | TIGR03705<br>PF17941<br>PF02503<br>PF13090<br>PF13089 | TIGR03705<br>PP_kinase_C_1<br>PP_kinase<br>PP_kinase_C<br>PP_kinase_N | poly_P_kin: polyphosphate kinase 1<br>Polyphosphate kinase C-terminal domain 1<br>Polyphosphate kinase middle domain<br>Polyphosphate kinase C-terminal domain 2<br>Polyphosphate kinase N-terminal domain | 2.50E-244<br>2.80E-75<br>4.80E-69<br>3.60E-58<br>9.70E-37 |
| WP_111766295.1 | 1864277 | 1865216 | + | 312 | PF00293<br>PF00300<br>TIGR03848<br>TIGR03162 | NUDIX<br>His_Phos_1<br>TIGR03848<br>TIGR03162 | NUDIX domain<br>Histidine phosphatase superfamily (branch 1)<br>MSMEG_4193: probable phosphomutase, MSMEG_4193 family<br>ribazole_cobC: alpha-ribazole phosphatase | 3.60E-14<br>9.00E-13<br>3.90E-12<br>1.60E-08 |
| WP_111766296.1 | 1866045 | 1865292 | - | 250 | PF00216<br>TIGR00988<br>TIGR00987<br>PF18291 | Bac_DNA_binding<br>TIGR00988<br>TIGR00987<br>HU-HIG | Bacterial DNA-binding protein<br>hip: integration host factor, beta subunit<br>himA: integration host factor, alpha subunit<br>HU domain fused to wHTH, Ig, or Glycine-rich motif | 5.50E-26<br>1.90E-17<br>4.50E-16<br>6.70E-05 |
| WP_111766297.1 | 1866904 | 1866316 | - | 195 | TIGR00171<br>PF00694<br>TIGR02087<br>TIGR02084<br>TIGR00139 | TIGR00171<br>Aconitase_C<br>TIGR02087<br>TIGR02084<br>TIGR00139 | leuD: 3-isopropylmalate dehydratase, small subunit<br>Aconitase C-terminal domain<br>LEUD_arch: 3-isopropylmalate dehydratase, small subunit<br>leud: 3-isopropylmalate dehydratase, small subunit<br>h_aconitase: homoaconitase | 6.70E-56<br>2.60E-23<br>1.60E-19<br>1.30E-16<br>4.10E-10 |
| WP_111766298.1 | 1868411 | 1866992 | - | 472 | TIGR00170<br>PF00330<br>TIGR01343<br>TIGR02086<br>TIGR02083 | TIGR00170<br>Aconitase<br>TIGR01343<br>TIGR02086<br>TIGR02083 | leuC: 3-isopropylmalate dehydratase, large subunit<br>Aconitase family (aconitate hydratase)<br>hacA_fam: homoaconitate hydratase family protein<br>IPMI_arch: 3-isopropylmalate dehydratase, large subunit<br>LEU2: 3-isopropylmalate dehydratase, large subunit | 4.00E-224<br>3.70E-157<br>1.10E-101<br>3.80E-89<br>2.70E-81 |
| WP_111766299.1 | 1868565 | 1869258 | + | 230 | TIGR02431<br>PF01614<br>PF09339 | TIGR02431<br>IclR<br>HTH_IclR | pcaR_pcaU: beta-ketoadipate pathway transcriptional regulators, PcaR/PcaU/PobR family<br>Bacterial transcriptional regulator<br>IclR helix-turn-helix domain | 4.60E-23<br>4.20E-18<br>2.20E-12 |
| WP_157970584.1 | 1870716 | 1869969 | - | 248 | TIGR02253<br>TIGR02254<br>PF13419<br>TIGR02252<br>TIGR01549 | TIGR02253<br>TIGR02254<br>HAD_2<br>TIGR02252<br>TIGR01549 | CTE7: HAD hydrolase, TIGR02253 family<br>YjjG/YfnB: noncanonical pyrimidine nucleotidase, YjjG family<br>Haloacid dehalogenase-like hydrolase<br>DREG-2: HAD hydrolase, REG-2-like, family IA<br>HAD-SF-IA-v1: HAD hydrolase, family IA, variant 1 | 7.60E-28<br>4.80E-24<br>1.70E-21<br>4.20E-21<br>3.10E-17 |

| index | leader | core | start | end | dir | score |
|---|---|---|---|---|---|---|
| 1 | MSSGQLIGLLVGAAVGALIGLRFFRGNGRNRATALIAPAV | FTVSAAAVGYLLGGAL | 1853269 | 1853098 | - | 9 |

| | | | | | | |
|---|---|---|---|---|---|---|
| 2 | MTLQDRVNDNPMMADVPMDAERIHKFSDLNPVL | GTPMGLAVVGGMAAAAVAGSEIGKNAD | 1856132 | 1855949 | - | 30 |
| 3 | MSDLSARINAASADLFGPLTPETPAARLGSAMPVL | ATPAGVAAGVATAAAFVAGYAVEEAGDK | 1856413 | 1856221 | - | 30 |
| 4 | MIAKSSTGAVVGFVIGAVLGYLFFNDADDVLLGTVVPTAILAI | STAVVGYLMVRYTARQKQ | 1856712 | 1856526 | - | 8 |
| 5 | MWCTGTRRAQAGNGEVAIP | LCPVVILCSFSGNRPDRVSRSGNVFLWDERTAAGDRTAVATSAVSAG | 1860220 | 1860019 | - | 8 |
| 6 | LMWCTGTRRAQAGNGEVAIP | LCPVVILCSFSGNRPDRVSRSGNVFLWDERTAAGDRTAVATSAVSAG | 1860223 | 1860019 | - | 8 |
| 7 | VLRSFTPALRALMWCTGTRRAQAGNGEVAIPLCPVVILC | SFSGNRPDRVSRSGNVFLWDERTAAGDRTAVATSAVSAG | 1860256 | 1860019 | - | 7 |
| 8 | MPVLRSFTPALRALMWCTGTRRAQAGNGEVAIPLCPVVIL | CSFSGNRPDRVSRSGNVFLWDERTAAGDRTAVATSAVSAG | 1860262 | 1860019 | - | 6 |
| 9 | VFPGNHFRAVGATGPGRPTGRCRSARRRR | MGDEVLAGGRRPGCSDRSRARRTAGKGSG | 1860580 | 1860403 | - | 6 |
| 10 | MAGGGPVNSRRTISVCGRPVRPIRCRRHIP | GVSRQPLPGGGCHRSRPPDGALPVGAATANG | 1860671 | 1860485 | - | 6 |
| 11 | VVAPTGARRRPRVGRRRTHRTARDGRP | GTPPTVWCGSGGRATSGVVDEWPAVDR | 1860814 | 1860649 | - | 8 |
| 12 | VPQRRPGGTVPRAVVAPTGARRRPRVGRRRTHR | TARDGRPGTPPTVWCGSGGRATSGVVDEWPAVDR | 1860853 | 1860649 | - | 6 |
| 13 | VPGDPPCARLTSRPGGGRRRHRAAATTRRDRSAGRGRADRRAPPAPRR | TTTDAPDGPRRAPRNTADRVVRIRRPGNERCRR | 1860917 | 1860671 | - | 6 |
| 14 | VRVRRARGAGA | VAPFVASGAGGVCATVAARRRAGAR...AVVFSVVVTRSFSPGRRRSGASGAR | 1861113 | 1860906 | - | 8 |
| 15 | MAAVRVRRARGAGA | VAPFVASGAGGVCATVAARRRAGAR...AVVFSVVVTRSFSPGRRRSGASGAR | 1861122 | 1860906 | - | 7 |
| 16 | VGMAAVRVRRARGAGA | VAPFVASGAGGVCATVAARRRAGAR...AVVFSVVVTRSFSPGRRRSGASGAR | 1861128 | 1860906 | - | 7 |
| 17 | VGAPSAVGMAAVRVRRARGAGA | VAPFVASGAGGVCATVAARRRAGAR...AVVFSVVVTRSFSPGRRRSGASGAR | 1861146 | 1860906 | - | 7 |
| 18 | VRRVPPGVGAPSAVGMAAVRVRRARGAGAVAPFVASGAGGVCA | TVAARRRAGARRGRGAASAVVFSVVVTRSFSPGRRRSGASGAR | 1861167 | 1860906 | - | 8 |
| 19 | VSVRAPAGVGAVGVRRVPPGV | GAPSAVGMAAVRVRRARGAGAVAPF...AVVFSVVVTRSFSPGRRRSGASGAR | 1861206 | 1860906 | - | 7 |
| 20 | VTAVRVRRGVFPAAAVSVRAPAGVGAVGVRRVPPGV | GAPSAVGMAAVRVRRARGAGAVAPF...AVVFSVVVTRSFSPGRRRSGASGAR | 1861251 | 1860906 | - | 6 |
| 21 | MAPSSSGLGRCPLTAVTRVRIPSGLHRQQTPGLRSGGLLRFRLTTSAPPP | APPRSSVAAVASAASGAAAVGRAGCS | 1869683 | 1869452 | - | 6 |

### Results for RNL83154.1 [Nocardiopsaceae bacterium YIM 96095] back to top

previous - next

#### Architecture

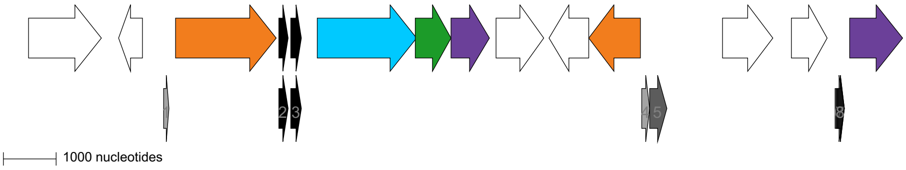

Link to nucleotide sequence

| Accession | start | end | direction | length (aa) | Pfam/HMM | name | description | E-value |
|---|---|---|---|---|---|---|---|---|
| RNL83149.1 | 615 | 1974 | + | 452 | TIGR00804<br>PF07662<br>PF01773<br>PF07670 | TIGR00804<br>Nucleos_tra2_C<br>Nucleos_tra2_N<br>Gate | nupC: nucleoside transporter, NupC family<br>Na+ dependent nucleoside transporter C-terminus<br>Na+ dependent nucleoside transporter N-terminus<br>Nucleoside recognition | 1.20E-136<br>3.20E-78<br>6.00E-20<br>1.10E-06 |
| RNL83150.1 | 2763 | 2301 | - | 153 | NO PFAM MATCH | - | - | - |
| RNL83151.1 | 3402 | 5298 | + | 631 | TIGR02203<br>TIGR02204<br>TIGR03796<br>TIGR00958<br>TIGR01846 | TIGR02203<br>TIGR02204<br>TIGR03796<br>TIGR00958<br>TIGR01846 | MsbA_lipidA: lipid A export permease/ATP-binding protein MsbA<br>MsbA_rel: ABC transporter, permease/ATP-binding protein<br>NHLM_micro_ABC1: NHLM bacteriocin system ABC transporter, peptidase/ATP-binding protein<br>3a01208: antigen peptide transporter 2<br>type_I_sec_HlyB: type I secretion system ATPase | 1.00E-103<br>2.00E-99<br>2.50E-83<br>1.40E-79<br>1.70E-75 |
| RNL83152.1 | 5344 | 5536 | + | 63 | LinA | LinA | LinA (linaridin precursor) | 1.90E-15 |
| RNL83153.1 | 5575 | 5782 | + | 68 | LinA | LinA | LinA (linaridin precursor) | 2.20E-14 |
| RNL83154.1 | 6083 | 7949 | + | 621 | LinH<br>LinE<br>LinG<br>PF12146<br>PF12697 | LinH<br>LinE<br>LinG<br>Hydrolase_4<br>Abhydrolase_6 | LinH (Fusion of LinE/LinG)<br>LinE Homolog<br>LinG Homolog<br>Serine aminopeptidase, S33<br>Alpha/beta hydrolase family | 1.70E-236<br>1.70E-99<br>2.10E-92<br>3.00E-18<br>3.40E-16 |
| RNL83155.1 | 7945 | 8614 | + | 222 | LinL | LinL | LinL | 1.60E-66 |
| RNL83156.1 | 8610 | 9345 | + | 244 | PF13649<br>PF08241<br>PF13847<br>PF13489<br>TIGR00755 | Methyltransf_25<br>Methyltransf_11<br>Methyltransf_31<br>Methyltransf_23<br>TIGR00755 | Methyltransferase domain<br>Methyltransferase domain<br>Methyltransferase domain<br>Methyltransferase domain<br>ksgA: ribosomal RNA small subunit methyltransferase A | 3.60E-15<br>1.90E-14<br>4.40E-10<br>4.40E-07<br>5.60E-06 |
| RNL83157.1 | 9468 | 10365 | + | 298 | PF01551 | Peptidase_M23 | Peptidase family M23 | 2.30E-14 |

| | | | | | | | | |
|---|---|---|---|---|---|---|---|---|
| RNL83158.1 | 11229 | 10473 | - | 251 | PF01061<br>PF12698<br>TIGR00025<br>TIGR01247<br>PF12679 | ABC2_membrane<br>ABC2_membrane_3<br>TIGR00025<br>TIGR01247<br>ABC2_membrane_2 | ABC-2 type transporter<br>ABC-2 family transporter protein<br>Mtu_efflux: ABC transporter efflux protein, DrrB family<br>drrB: daunorubicin resistance ABC transporter membrane protein<br>ABC-2 family transporter protein | 2.00E-16<br>3.80E-15<br>1.70E-09<br>3.00E-09<br>7.60E-09 |
| RNL83159.1 | 12200 | 11225 | - | 324 | TIGR01188<br>TIGR01288<br>TIGR03522<br>TIGR03864<br>TIGR03740 | TIGR01188<br>TIGR01288<br>TIGR03522<br>TIGR03864<br>TIGR03740 | drrA: daunorubicin resistance ABC transporter, ATP-binding protein<br>nodI: nodulation ABC transporter NodI<br>GldA_ABC_ATP: gliding motility-associated ABC transporter ATP-binding subunit GldA<br>PQQ_ABC_ATP: ABC transporter, ATP-binding subunit, PQQ-dependent alcohol dehydrogenase system<br>galliderm_ABC: lantibiotic protection ABC transporter, ATP-binding subunit | 1.70E-63<br>4.10E-49<br>7.40E-48<br>4.40E-41<br>4.50E-38 |
| RNL83160.1 | 13753 | 14719 | + | 321 | NO PFAM MATCH | - | - | - |
| RNL83239.1 | 15058 | 15754 | + | 231 | PF01596<br>PF13578<br>TIGR04188 | Methyltransf_3<br>Methyltransf_24<br>TIGR04188 | O-methyltransferase<br>Methyltransferase domain<br>methyltr_grsp: methyltransferase, ATP-grasp peptide maturase system | 2.90E-49<br>6.20E-16<br>3.80E-04 |
| RNL83161.1 | 16162 | 17185 | + | 340 | PF00891<br>TIGR02716<br>PF16864<br>PF13489<br>TIGR02469 | Methyltransf_2<br>TIGR02716<br>Dimerisation2<br>Methyltransf_23<br>TIGR02469 | O-methyltransferase domain<br>C20_methyl_CrtF: C-20 methyltransferase BchU<br>Dimerisation domain<br>Methyltransferase domain<br>CbiT: precorrin-6Y C5,15-methyltransferase (decarboxylating), CbiT subunit | 7.30E-41<br>4.20E-19<br>5.00E-14<br>2.60E-05<br>9.30E-05 |

| index | leader | core | start | end | dir | score |
|---|---|---|---|---|---|---|
| 1 | MNENAFAQGITRAPIKT | NTYHQVVTCDFVTKKGH | 3162 | 3267 | + | 6 |
| 2 | MEELLDRTNEQSDDLFGVLTPQTQVEFDTGNTPVL | ATPAAAAAGAGATVGAFAAGVAVGYAVG | 5344 | 5536 | + | 31 |
| 3 | MVVDTLLEKVNGSSDLVRVSIREEQATTMVGLGPVL | ATPGAFAAGAGAACAVAGAFGAGYAIGQDLAG | 5575 | 5782 | + | 29 |
| 4 | VGSYDRVARSTCLTPFRHSRIRAVFRA | QSNLFRRGITNRGPMTRLDLARNTKNPR | 12223 | 12391 | + | 6 |
| 5 | VKGLPRSHPTRLGDPRIRVVSGGHP...TDGATVGPTDPPEFRWGPGLPKLLR | GVPAAFVASSWLSTGAHEDWRRQGPARAARHATEGTPT | 12387 | 12702 | + | 8 |
| 6 | MRPPTPLNPAEPDPPTRSSSSMNPPKPPSPCATASAAE | RTTYAPHELPADFTKRRE | 15892 | 16063 | + | 8 |
| 7 | LNPAEPDPPTRSSSSMNPPKPPSPCATASAAE | RTTYAPHELPADFTKRRE | 15910 | 16063 | + | 8 |
| 8 | MNPPKPPSPCATASAAE | RTTYAPHELPADFTKRRE | 15955 | 16063 | + | 6 |

### Results for SHK25020.1 [Nocardiopsis flavescens] back to top

previous - next

#### Architecture

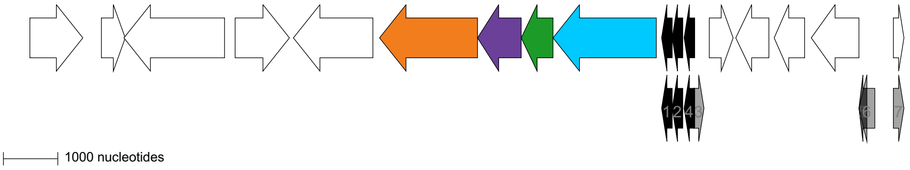

Link to nucleotide sequence

| Accession | start | end | direction | length (aa) | Pfam/HMM | name | description | E-value |
|---|---|---|---|---|---|---|---|---|
| SHK24772.1 | 64738 | 65713 | + | 324 | TIGR00186<br>PF00588<br>PF08032<br>TIGR00050<br>TIGR00185 | TIGR00186<br>SpoU_methylase<br>SpoU_sub_bind<br>TIGR00050<br>TIGR00185 | rRNA_methyl_3: RNA methyltransferase, TrmH family, group 3<br>SpoU rRNA Methylase family<br>RNA 2'-O ribose methyltransferase substrate binding<br>rRNA_methyl_1: RNA methyltransferase, TrmH family, group 1<br>tRNA_yibK_trmL: tRNA (cytidine(34)-2'-O)-methyltransferase | 6.50E-56<br>3.90E-33<br>3.90E-14<br>2.10E-06<br>1.90E-05 |
| SHK24815.1 | 66059 | 66497 | + | 145 | NO PFAM MATCH | - | - | - |
| SHK24844.1 | 68307 | 66468 | - | 612 | NO PFAM MATCH | - | - | - |
| SHK24873.1 | 68501 | 69509 | + | 335 | NO PFAM MATCH | - | - | - |
| SHK24891.1 | 71046 | 69576 | - | 489 | TIGR02734<br>TIGR03467<br>PF13450<br>TIGR01317<br>TIGR00292 | TIGR02734<br>TIGR03467<br>NAD_binding_8<br>TIGR01317<br>TIGR00292 | crtI_fam: phytoene desaturase<br>HpnE: squalene-associated FAD-dependent desaturase<br>NAD(P)-binding Rossmann-like domain<br>GOGAT_sm_gam: glutamate synthase, NADH/NADPH, small subunit<br>TIGR00292: thiazole biosynthesis enzyme | 2.00E-10<br>7.90E-09<br>9.80E-09<br>2.10E-08<br>1.40E-07 |
| SHK24926.1 | 72962 | 71159 | - | 600 | TIGR02204<br>TIGR02203<br>TIGR00958<br>TIGR03796<br>TIGR01846 | TIGR02204<br>TIGR02203<br>TIGR00958<br>TIGR03796<br>TIGR01846 | MsbA_rel: ABC transporter, permease/ATP-binding protein<br>MsbA_lipidA: lipid A export permease/ATP-binding protein MsbA<br>3a01208: antigen peptide transporter 2<br>NHLM_micro_ABC1: NHLM bacteriocin system ABC transporter, peptidase/ATP-binding protein<br>type_I_sec_HlyB: type I secretion system ATPase | 5.10E-103<br>5.90E-94<br>1.00E-90<br>9.50E-84<br>1.20E-81 |
| SHK24957.1 | 73771 | 72958 | - | 270 | PF13649<br>PF08241<br>PF13489<br>PF08242<br>PF13847 | Methyltransf_25<br>Methyltransf_11<br>Methyltransf_23<br>Methyltransf_12<br>Methyltransf_31 | Methyltransferase domain<br>Methyltransferase domain<br>Methyltransferase domain<br>Methyltransferase domain<br>Methyltransferase domain | 1.50E-14<br>4.30E-10<br>1.60E-09<br>5.30E-09<br>1.10E-04 |
| SHK24989.1 | 74358 | 73767 | - | 196 | LinL | LinL | LinL | 9.30E-59 |
| SHK25020.1 | 76247 | 74354 | - | 630 | LinH<br>LinG<br>LinE<br>PF12697<br>LinL | LinH<br>LinG<br>LinE<br>Abhydrolase_6<br>LinL | LinH (Fusion of LinE/LinG)<br>LinG Homolog<br>LinE Homolog<br>Alpha/beta hydrolase family<br>LinL | 5.20E-235<br>6.20E-111<br>9.40E-92<br>1.80E-15<br>7.80E-13 |

| | | | | | | | | |
|---|---|---|---|---|---|---|---|---|
| SHK25069.1 | 76535 | 76334 | - | 66 | LinA | LinA | LinA (linaridin precursor) | 1.00E-08 |
| SHK25096.1 | 76743 | 76551 | - | 63 | LinA | LinA | LinA (linaridin precursor) | 7.30E-09 |
| SHK25126.1 | 76966 | 76759 | - | 68 | LinA | LinA | LinA (linaridin precursor) | 1.30E-09 |
| SHK25156.1 | 77233 | 77671 | + | 145 | PF13380<br>PF02629 | CoA_binding_2<br>CoA_binding | CoA binding domain<br>CoA binding domain | 1.20E-34<br>5.00E-06 |
| SHK25188.1 | 78311 | 77708 | - | 200 | PF00440<br>TIGR03968<br>PF16859 | TetR_N<br>TIGR03968<br>TetR_C_11 | Bacterial regulatory proteins, tetR family<br>mycofact_TetR: mycofactocin system transcriptional regulator<br>Tetracyclin repressor-like, C-terminal domain | 2.90E-10<br>5.80E-05<br>1.70E-04 |
| SHK25220.1 | 78971 | 78404 | - | 188 | PF01872 | RibD_C | RibD C-terminal domain | 2.80E-34 |
| SHK25280.1 | 79966 | 79099 | - | 288 | PF05653 | Mg_trans_NIPA | Magnesium transporter NIPA | 2.40E-06 |
| SHK25312.1 | 80597 | 80801 | + | 67 | NO PFAM MATCH | - | - | - |

| index | leader | core | start | end | dir | score |
|---|---|---|---|---|---|---|
| 1 | MNDVSFLSRPGSMADIDIRDPHGRSVSGQAPVT | ATPGVAAGVAIGYALIYAFNAGLNSTHPTLPHA | 76535 | 76334 | - | 28 |
| 2 | MNPLTPPASLVGSGALTETDGGSVRLGSAPVT | ATPLAVTAGLAAGVALGYALANSTHPTLPHA | 76743 | 76551 | - | 30 |
| 3 | VRSDLCPHARSSRPRCRGPMWFRDGPGRGRPS | RRGSAGAVPTIATHGAEVKADRRHEAAARGAA | 76942 | 77137 | + | 6 |
| 4 | MRTQVRPHWDIVPGALEEDAEGQVFESGLPVT | ATPVVVGAAGAIVVAYAAGALANALGGGTHPTLPHA | 76966 | 76759 | - | 30 |
| 5 | LPPARHPHLRGSGHRPGRTPRHRGHARSRPGHAGSSPGM | TTPPPSPGEDR | 80120 | 79967 | - | 8 |
| 6 | VSGPGVPRPARGRTAGTGPPAPNTA...RPGRTPRHRGHARSRPGHAGSSPGM | TTPPPSPGEDR | 80276 | 79967 | - | 6 |
| 7 | MNTASIVSWVLFVAVFALTRALTRQPYADLFGGPGLAATLASF | GTAAALGLLVLLAVQRFAGRPPRE | 80597 | 80801 | + | 6 |

### Results for WP_150255942.1 [Nocardiopsis sp. H13] back to top

previous - next

#### Architecture

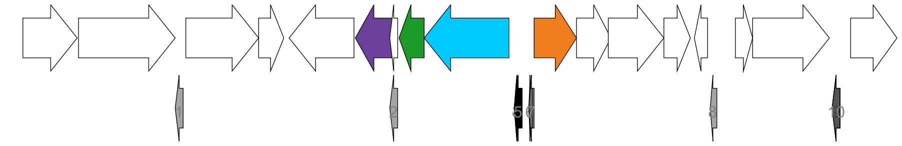

1000 nucleotides

Link to nucleotide sequence

| Accession | start | end | direction | length (aa) | Pfam/HMM | name | description | E-value |
|---|---|---|---|---|---|---|---|---|
| WP_150255925.1 | 4844 | 6074 | + | 409 | TIGR03253<br>PF02515<br>TIGR04253 | TIGR03253<br>CoA_transf_3<br>TIGR04253 | oxalate_frc: formyl-CoA transferase<br>CoA-transferase family III<br>mesacon_CoA_iso: mesaconyl-CoA isomerase | 4.10E-195<br>1.20E-103<br>2.10E-21 |
| WP_150255927.1 | 6078 | 8238 | + | 719 | TIGR02717<br>PF13549<br>PF13607<br>PF13380<br>TIGR01019 | TIGR02717<br>ATP-grasp_5<br>Succ_CoA_lig<br>CoA_binding_2<br>TIGR01019 | AcCoA-syn-alpha: acetyl coenzyme A synthetase (ADP forming), alpha domain<br>ATP-grasp domain<br>Succinyl-CoA ligase like flavodoxin domain<br>CoA binding domain<br>sucCoAalpha: succinate-CoA ligase, alpha subunit | 2.80E-137<br>1.00E-72<br>1.90E-49<br>6.50E-25<br>6.70E-11 |
| WP_150255929.1 | 8474 | 10073 | + | 532 | NO PFAM MATCH | - | - | - |
| WP_150255931.1 | 10072 | 10636 | + | 187 | NO PFAM MATCH | - | - | - |
| WP_150255933.1 | 12203 | 10751 | - | 483 | PF13450<br>TIGR02734<br>TIGR03467<br>TIGR02733<br>TIGR01317 | NAD_binding_8<br>TIGR02734<br>TIGR03467<br>TIGR02733<br>TIGR01317 | NAD(P)-binding Rossmann-like domain<br>crtI_fam: phytoene desaturase<br>HpnE: squalene-associated FAD-dependent desaturase<br>desat_CrtD: C-3',4' desaturase CrtD<br>GOGAT_sm_gam: glutamate synthase, NADH/NADPH, small subunit | 9.20E-11<br>8.70E-10<br>1.60E-08<br>7.40E-08<br>1.90E-07 |
| WP_150255935.1 | 13050 | 12222 | - | 275 | PF13649<br>PF08241<br>PF08242<br>TIGR01983<br>PF13489 | Methyltransf_25<br>Methyltransf_11<br>Methyltransf_12<br>TIGR01983<br>Methyltransf_23 | Methyltransferase domain<br>Methyltransferase domain<br>Methyltransferase domain<br>UbiG: 3-demethylubiquinone-9 3-O-methyltransferase<br>Methyltransferase domain | 4.60E-15<br>1.50E-10<br>1.90E-07<br>1.40E-06<br>1.50E-06 |
| WP_150255937.1 | 13175 | 12995 | - | 59 | NO PFAM MATCH | - | - | - |
| WP_150255939.1 | 13753 | 13192 | - | 186 | LinL | LinL | LinL | 7.50E-48 |
| WP_150255942.1 | 15639 | 13749 | - | 629 | LinH<br>LinG<br>LinE<br>PF12697<br>PF00975 | LinH<br>LinG<br>LinE<br>Abhydrolase_6<br>Thioesterase | LinH (Fusion of LinE/LinG)<br>LinG Homolog<br>LinE Homolog<br>Alpha/beta hydrolase family<br>Thioesterase domain | 9.50E-214<br>1.00E-101<br>3.00E-89<br>5.10E-15<br>3.40E-07 |

| | | | | | | | | |
|---|---|---|---|---|---|---|---|---|
| WP_150255944.1 | 16182 | 17148 | + | 321 | TIGR01188<br>TIGR01288<br>TIGR03864<br>TIGR03740<br>TIGR03522 | TIGR01188<br>TIGR01288<br>TIGR03864<br>TIGR03740<br>TIGR03522 | drrA: daunorubicin resistance ABC transporter, ATP-binding protein<br>nodI: nodulation ABC transporter NodI<br>PQQ_ABC_ATP: ABC transporter, ATP-binding subunit, PQQ-dependent alcohol dehydrogenase system<br>gallidermABC: lantibiotic protection ABC transporter, ATP-binding subunit<br>GldA_ABC_ATP: gliding motility-associated ABC transporter ATP-binding subunit GldA | 2.10E-63<br>6.90E-45<br>4.20E-43<br>1.10E-37<br>1.50E-36 |
| WP_150255946.1 | 17144 | 17906 | + | 253 | PF01061<br>PF12698<br>TIGR03062<br>TIGR01291<br>TIGR01247 | ABC2_membrane<br>ABC2_membrane_3<br>TIGR03062<br>TIGR01291<br>TIGR01247 | ABC-2 type transporter<br>ABC-2 family transporter protein<br>pip_yhgE_Cterm: YhgE/Pip C-terminal domain<br>nodJ: ABC-2 type transporter, NodJ family<br>drrB: daunorubicin resistance ABC transporter membrane protein | 2.90E-20<br>9.30E-12<br>2.10E-10<br>2.40E-04<br>4.10E-04 |
| WP_150255948.1 | 17845 | 19072 | + | 408 | PF07730 | HisKA_3 | Histidine kinase | 2.50E-14 |
| WP_150255950.1 | 19068 | 19674 | + | 201 | PF00072<br>TIGR02875<br>PF00196<br>TIGR02154<br>TIGR02956 | Response_reg<br>TIGR02875<br>GerE<br>TIGR02154<br>TIGR02956 | Response regulator receiver domain<br>spore_0_A: sporulation transcription factor Spo0A<br>Bacterial regulatory proteins, luxR family<br>PhoB: phosphate regulon transcriptional regulatory protein PhoB<br>TMAO_torS: TMAO reductase sytem sensor TorS | 1.50E-24<br>3.60E-15<br>7.10E-15<br>1.20E-12<br>2.40E-09 |
| WP_150255952.1 | 20052 | 19737 | - | 104 | PF00893 | Multi_Drug_Res | Small Multidrug Resistance protein | 3.60E-22 |
| WP_150255954.1 | 20653 | 21052 | + | 132 | PF04341 | DUF485 | Protein of unknown function, DUF485 | 3.30E-27 |
| WP_150255956.1 | 21048 | 22734 | + | 561 | TIGR02711<br>PF00474<br>TIGR03648<br>TIGR00813<br>TIGR02121 | TIGR02711<br>SSF<br>TIGR03648<br>TIGR00813<br>TIGR02121 | symport_actP: cation/acetate symporter ActP<br>Sodium:solute symporter family<br>Na_symport_lg: probable sodium:solute symporter, VC_2705 subfamily<br>sss: transporter, solute:sodium symporter (SSS) family<br>Na_Pro_sym: sodium/proline symporter | 2.20E-133<br>2.40E-91<br>3.20E-79<br>1.40E-56<br>2.90E-34 |
| WP_150255958.1 | 23214 | 24231 | + | 338 | PF06580<br>PF02518 | His_kinase<br>HATPase_c | Histidine kinase<br>Histidine kinase-, DNA gyrase B-, and HSP90-like ATPase | 1.40E-22<br>5.10E-06 |

| index | leader | core | start | end | dir | score |
|---|---|---|---|---|---|---|
| 1 | MAPPGGADSSLPDARARSPRRSGCVARWGTRR | PCGGGSMAAGFRGGARWYGARSGPEHRPGTGSE | 8427 | 8229 | - | 6 |
| 2 | MKSQARSVIAVVAAVAVAVVLVALALTEV | LSVAATLGLGAVLWVGLLVIFVPRKKEGSR | 13175 | 12995 | - | 6 |
| 3 | MKEIIEHFDALEFDAEEGMVAFDQAPVM | ATVVTTATTATTSITGIFG | 15887 | 15743 | - | 27 |
| 4 | VMKEIIEHFDALEFDAEEGMVAFDQAPVM | ATVVTTATTATTSITGIFG | 15890 | 15743 | - | 27 |
| 5 | LVRRSIYPKENVMKEIIEHFDALEFDAEEGMVAFDQAPVM | ATVVTTATTATTSITGIFG | 15923 | 15743 | - | 29 |
| 6 | LTLGDLSPVLLAMDS | YCMCHSEFVTLVMSES | 16171 | 16075 | - | 6 |
| 7 | LSPYGHCLTLGDLSPVLLA | MDSYCMCHSEFVTLVMSES | 16192 | 16075 | - | 6 |
| 8 | VGSRLHRARNPRPVPGVVACAPSPPS | RAPASARSATPPDEVRRTR | 20244 | 20106 | - | 6 |
| 9 | VPGEGWAPGWSGPPTVPSWWFGSPPGLRVE | RRPGGAVLVRASTGSWRCRRRVREERANPP | 22977 | 22794 | - | 6 |
| 10 | VLGAAAVPGEGWAPGWSGPPTVPSWWFGSPPGL | RVERRPGGAVLVRASTGSWRCRRRVREERANPP | 22995 | 22794 | - | 7 |

### Results for KZM74592.1 [Nocardia terpenica] back to top

previous - next

#### Architecture

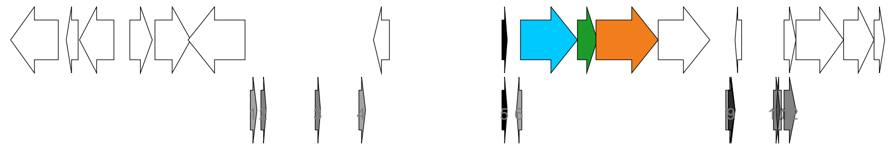

1000 nucleotides

Link to nucleotide sequence

| Accession | start | end | direction | length (aa) | Pfam/HMM | name | description | E-value |
|---|---|---|---|---|---|---|---|---|
| KZM74585.1 | 18937 | 17569 | - | 455 | TIGR02587 | TIGR02587 | TIGR02587: putative integral membrane protein TIGR02587 | 3.90E-04 |
| KZM75116.1 | 19483 | 19144 | - | 112 | NO PFAM MATCH | - | - | - |
| KZM74586.1 | 20490 | 19539 | - | 316 | NO PFAM MATCH | - | - | - |
| KZM74587.1 | 20956 | 21607 | + | 216 | NO PFAM MATCH | - | - | - |
| KZM74588.1 | 21684 | 22695 | + | 336 | TIGR01177<br>PF01555<br>PF01170<br>TIGR03704 | TIGR01177<br>N6_N4_Mtase<br>UPF0020<br>TIGR03704 | TIGR01177: putative methyltransferase, TIGR01177 family<br>DNA methylase<br>Putative RNA methylase family UPF0020<br>PrmC_rel_meth: putative protein-(glutamine-N5) methyltransferase, unknown substrate-specific | 3.40E-14<br>1.00E-13<br>5.60E-13<br>4.70E-08 |
| KZM74589.1 | 24229 | 22627 | - | 533 | PF01610<br>PF14690<br>PF02796 | DDE_Tnp_ISL3<br>zf-ISL3<br>HTH_7 | Transposase<br>zinc-finger of transposase IS204/IS1001/IS1096/IS1165<br>Helix-turn-helix domain of resolvase | 1.30E-28<br>2.50E-08<br>4.90E-04 |
| KZM74590.1 | 28302 | 27873 | - | 142 | PF06114 | Peptidase_M78 | IrrE N-terminal-like domain | 2.40E-05 |
| KZM74591.1 | 31493 | 31676 | + | 60 | LinA | LinA | LinA (linaridin precursor) | 4.30E-09 |

| | | | | | | | | |
|---|---|---|---|---|---|---|---|---|
| KZM74592.1 | 32038 | 33661 | + | 540 | LinH<br>LinE<br>LinG<br>PF12697<br>TIGR02427 | LinH<br>LinE<br>LinG<br>Abhydrolase_6<br>TIGR02427 | LinH (Fusion of LinE/LinG)<br>LinE Homolog<br>LinG Homolog<br>Alpha/beta hydrolase family<br>protocat_pcaD: 3-oxoadipate enol-lactonase | 5.20E-170<br>3.00E-109<br>7.30E-78<br>2.60E-14<br>6.20E-11 |
| KZM74593.1 | 33657 | 34200 | + | 180 | LinL | LinL | LinL | 2.20E-58 |
| KZM74594.1 | 34196 | 35948 | + | 583 | TIGR02204<br>TIGR02203<br>TIGR00958<br>TIGR01846<br>TIGR03375 | TIGR02204<br>TIGR02203<br>TIGR00958<br>TIGR01846<br>TIGR03375 | MsbA_rel: ABC transporter, permease/ATP-binding protein<br>MsbA_lipidA: lipid A export permease/ATP-binding protein MsbA<br>3a01208: antigen peptide transporter 2<br>type_I_sec_HlyB: type I secretion system ATPase<br>type_I_sec_LssB: type I secretion system ATPase | 3.20E-109<br>3.30E-103<br>1.30E-90<br>3.40E-88<br>3.80E-85 |
| KZM74595.1 | 35944 | 37411 | + | 488 | TIGR02734<br>TIGR03467<br>TIGR01372<br>PF13450<br>PF12831 | TIGR02734<br>TIGR03467<br>TIGR01372<br>NAD_binding_8<br>FAD_oxidored | crtI_fam: phytoene desaturase<br>HpnE: squalene-associated FAD-dependent desaturase<br>soxA: sarcosine oxidase, alpha subunit family<br>NAD(P)-binding Rossmann-like domain<br>FAD dependent oxidoreductase | 2.90E-10<br>1.90E-08<br>2.50E-07<br>8.40E-07<br>8.90E-07 |
| KZM74596.1 | 38325 | 38127 | - | 65 | NO PFAM MATCH | - | - | - |
| KZM74597.1 | 39512 | 39833 | + | 106 | PF10824 | T7SS_ESX_EspC | Excreted virulence factor EspC, type VII ESX diderm | 9.90E-04 |
| KZM74598.1 | 39838 | 41215 | + | 458 | TIGR03930<br>PF06013 | TIGR03930<br>WXG100 | WXG100_ESAT6: WXG100 family type VII secretion target<br>Proteins of 100 residues with WXG | 2.70E-07<br>1.20E-06 |
| KZM74599.1 | 41211 | 42060 | + | 282 | NO PFAM MATCH | - | - | - |
| KZM74600.1 | 42056 | 42359 | + | 100 | NO PFAM MATCH | - | - | - |

| index | leader | core | start | end | dir | score |
|---|---|---|---|---|---|---|
| 1 | VVIGVVSAVRGSGWYGTGRASGMSPTHRCAA | WQSLGKVLRSPRPTGRDRIAAPPDTAARVLA | 24390 | 24579 | + | 6 |
| 2 | MYWPRVAGDPPELPDPTGCRSPTLWV | VECGEDRGVLGERHSGSAGGGTRRSS | 24680 | 24839 | + | 7 |
| 3 | MICPVKLSGVVFPSRRLVVERAPSA | PTTRSALASFAVTAVLK | 26230 | 26359 | + | 7 |
| 4 | LKSTTAGSAKPYGLEVGCVLRSAASN | NTTETAMGIRLMARTRLTTSRRCDAIRKNAAVKYL | 27464 | 27650 | + | 6 |
| 5 | MFQLMEEVMLLEKVNADNTLVGELKAAALPGPATPVM | STPTLVRVIANVAQATKATRAAG | 31493 | 31676 | + | 31 |
| 6 | VHALSHPLASNEVSDRLHRFGLCWYLAPG | MTPVIQTPQLTVGSTG | 32056 | 31918 | - | 6 |
| 7 | VTLGTLIGTVVGAGIGCLAGGSVGA...VLSIPGFLGGCLVTGAVGAGIGATV | GTIVVGGPAAVIGGLLFLDALNRQNTN | 37850 | 38111 | + | 6 |
| 8 | VGTIGVLSIPGFLGGCLVTGAVGAGIGATV | GTIVVGGPAAVIGGLLFLDALNRQNTN | 37937 | 38111 | + | 7 |
| 9 | VLSIPGFLGGCLVTGAVGAGIGATV | GTIVVGGPAAVIGGLLFLDALNRQNTN | 37952 | 38111 | + | 7 |
| 10 | VVDGLGSVFS | VTPPLDHSRRPGTGAALVKEYLGST...QQRSLLYGCGDRRDRTGHWFRPVVE | 39196 | 39403 | + | 7 |
| 11 | MQHLPPHSLHDWAEPVPGP | VTPIATAVEEGPLLAWTTQDHPCTA | 39424 | 39289 | - | 6 |
| 12 | MVCVLVDPAILRAFAVQVGEASKAV...GLKDKVFGSGDGVGGSALQWAAHGL | GVNLDGPLAKFATEVGWMGDAVRGA...FEVSDEELAGRFAMLYTDPPKAKGK | 39512 | 39833 | + | 7 |

### Results for WP_064723074.1 [Paenarthrobacter nicotinovorans] back to top

previous - next

#### Architecture

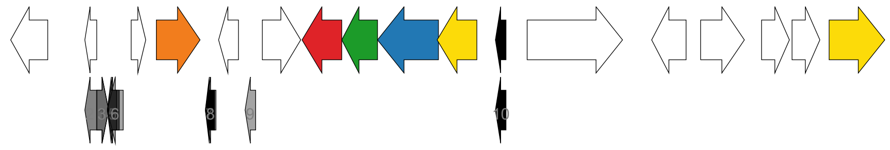

1000 nucleotides

Link to nucleotide sequence

| Accession | start | end | direction | length (aa) | Pfam/HMM | name | description | E-value |
|---|---|---|---|---|---|---|---|---|
| WP_082924713.1 | 58523 | 57980 | - | 180 | PF13302<br>PF00583<br>TIGR03585 | Acetyltransf_3<br>Acetyltransf_1<br>TIGR03585 | Acetyltransferase (GNAT) domain<br>Acetyltransferase (GNAT) family<br>PseH: pseudaminic acid biosynthesis N-acetyl transferase | 1.80E-22<br>1.50E-12<br>5.70E-04 |
| WP_156524806.1 | 59271 | 59091 | - | 59 | NO PFAM MATCH | - | - | - |
| WP_064723071.1 | 59802 | 60015 | + | 70 | NO PFAM MATCH | - | - | - |
| WP_064723134.1 | 60183 | 60834 | + | 216 | TIGR01189<br>TIGR02857<br>TIGR01166<br>TIGR02673<br>TIGR01188 | TIGR01189<br>TIGR02857<br>TIGR01166<br>TIGR02673<br>TIGR01188 | ccmA: heme ABC exporter, ATP-binding protein CcmA<br>CydD: thiol reductant ABC exporter, CydD subunit<br>cbiO: cobalt ABC transporter, ATP-binding protein<br>FtsE: cell division ATP-binding protein FtsE<br>drrA: daunorubicin resistance ABC transporter, ATP-binding protein | 6.80E-30<br>1.10E-25<br>3.10E-25<br>1.60E-24<br>8.00E-24 |
| WP_156524807.1 | 61429 | 61120 | - | 102 | NO PFAM MATCH | - | - | - |
| WP_156524808.1 | 61778 | 62363 | + | 194 | NO PFAM MATCH | - | - | - |
| WP_082924714.1 | 62992 | 62377 | - | 204 | TIGR00521<br>PF02441<br>TIGR02113 | TIGR00521<br>Flavoprotein<br>TIGR02113 | coaBC_dfp: phosphopantothenoylcysteine decarboxylase / phosphopantothenate--cysteine ligase<br>Flavoprotein<br>coaC_strep: phosphopantothenoylcysteine decarboxylase | 4.70E-13<br>1.90E-10<br>2.90E-08 |
| WP_064723073.1 | 63531 | 62988 | - | 180 | LinL | LinL | LinL | 9.40E-48 |

| | | | | | | | | |
|---|---|---|---|---|---|---|---|---|
| WP_064723074.1 | 64441 | 63517 | - | 307 | LinE<br>LinH<br>PF12697<br>TIGR02427<br>PF06821 | LinE<br>LinH<br>Abhydrolase_6<br>TIGR02427<br>Ser_hydrolase | LinE Homolog<br>LinH (Fusion of LinE/LinG)<br>Alpha/beta hydrolase family<br>protocat_pcaD: 3-oxoadipate enol-lactonase<br>Serine hydrolase | 3.10E-80<br>1.40E-46<br>1.20E-09<br>3.30E-06<br>5.20E-05 |
| WP_082924724.1 | 65034 | 64437 | - | 198 | LinG<br>LinH<br>TIGR04033 | LinG<br>LinH<br>TIGR04033 | LinG Homolog<br>LinH (Fusion of LinE/LinG)<br>export_SdpB: antimicrobial peptide system protein, SdpB family | 1.40E-74<br>8.10E-70<br>5.70E-05 |
| WP_156524809.1 | 65477 | 65309 | - | 55 | LinA | LinA | LinA (linaridin precursor) | 5.50E-09 |
| WP_082924715.1 | 65789 | 67229 | + | 479 | TIGR02734<br>PF13450<br>TIGR03467<br>TIGR02733<br>PF01266 | TIGR02734<br>NAD_binding_8<br>TIGR03467<br>TIGR02733<br>DAO | crtI_fam: phytoene desaturase<br>NAD(P)-binding Rossmann-like domain<br>HpnE: squalene-associated FAD-dependent desaturase<br>desat_CrtD: C-3',4' desaturase CrtD<br>FAD dependent oxidoreductase | 1.60E-11<br>4.40E-11<br>1.90E-09<br>5.90E-09<br>1.60E-08 |
| WP_064723076.1 | 68204 | 67670 | - | 177 | NO PFAM MATCH | - | - | - |
| WP_012630978.1 | 68420 | 69082 | + | 220 | INFERRED GENE | - | - | - |
| WP_064723136.1 | 69344 | 69770 | + | 141 | NO PFAM MATCH | - | - | - |
| WP_064723077.1 | 69807 | 70221 | + | 137 | NO PFAM MATCH | - | - | - |
| WP_064723078.1 | 70362 | 71217 | + | 284 | PF00561<br>TIGR03056<br>LinH<br>TIGR03695<br>LinE | Abhydrolase_1<br>TIGR03056<br>LinH<br>TIGR03695<br>LinE | alpha/beta hydrolase fold<br>bchO_mg_che_rel: putative magnesium chelatase accessory protein<br>LinH (Fusion of LinE/LinG)<br>menH_SHCHC: 2-succinyl-6-hydroxy-2,4-cyclohexadiene-1-carboxylate synthase<br>LinE Homolog | 1.70E-22<br>3.50E-15<br>3.20E-13<br>5.00E-13<br>1.10E-12 |

| index | leader | core | start | end | dir | score |
|---|---|---|---|---|---|---|
| 1 | MVNGPEVTWYVPEPSH | STVPSNAGRGTGFWRATSTWFFEVGGKMSTVGTQTVTIWKTI | 59269 | 59446 | + | 6 |
| 2 | MTVWVAVAAPRCWNTTSVFVAELDGANEM | MPLTLKSTEGSPVPTHLITVFEMAPSAPLE | 59271 | 59091 | - | 7 |
| 3 | VNGPEVTWYVPEPSH | STVPSNAGRGTGFWRATSTWFFEVGGKMSTVGTQTVTIWKTI | 59272 | 59446 | + | 6 |
| 4 | LAKFPLFRNSLICPKINSVTWIPID | SVYITRVNNLWFHRYSPAKTRLSSR | 59579 | 59426 | - | 6 |
| 5 | MLVGPQEHLSDVCDLAKFPLFRNSLICPKINS | VTWIPIDSVYITRVNNLWFHRYSPAKTRLSSR | 59621 | 59426 | - | 8 |
| 6 | MPVFVRKLCINNKRPTRGMLVGPQEHLSDVCDLAKFPLFRN | SLICPKINSVTWIPIDSVYITRVNNLWFHRYSPAKTRLSSR | 59675 | 59426 | - | 6 |
| 7 | MTPQALTICEEAGFRRTWRDSG | LTAVDTRVVTTGAATSKAVRIAI | 61060 | 60922 | - | 10 |
| 8 | MPAKKSKIKRMTPQA | LTICEEAGFRRTWRDSGLTAVDTRVVTTGAATSKAVRIAI | 61090 | 60922 | - | 7 |
| 9 | VTGFGSENSQSCGKLTRRCGNACR | LPPLALFHQGPGPKETEQGWRQRRKCSS | 61673 | 61514 | - | 6 |
| 10 | MSSILVEPSVEDSVNARFSSVDQGFINAHNEPAM | TTPSAIVTQTLVVGSTICLIC | 65477 | 65309 | - | 32 |

### Results for BCB83522.1 [Phytohabitans suffuscus] back to top

previous - next

#### Architecture

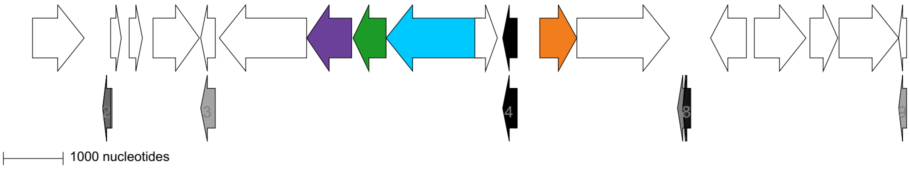

Link to nucleotide sequence

| Accession | start | end | direction | length (aa) | Pfam/HMM | name | description | E-value |
|---|---|---|---|---|---|---|---|---|
| BCB83514.1 | 930552 | 931422 | + | 289 | NO PFAM MATCH | - | - | - |
| BCB83515.1 | 931850 | 932033 | + | 60 | NO PFAM MATCH | - | - | - |
| BCB83516.1 | 932173 | 932383 | + | 69 | NO PFAM MATCH | - | - | - |
| BCB83517.1 | 932577 | 933348 | + | 256 | PF13359<br>PF13613<br>PF01609 | DDE_Tnp_4<br>HTH_Tnp_4<br>DDE_Tnp_1 | DDE superfamily endonuclease<br>Helix-turn-helix of DDE superfamily endonuclease<br>Transposase DDE domain | 6.80E-25<br>2.30E-15<br>1.80E-10 |
| BCB83518.1 | 933619 | 933361 | - | 85 | NO PFAM MATCH | - | - | - |
| BCB83519.1 | 935151 | 933678 | - | 490 | TIGR02734<br>PF13450<br>PF01593<br>TIGR00292<br>TIGR02733 | TIGR02734<br>NAD_binding_8<br>Amino_oxidase<br>TIGR00292<br>TIGR02733 | crtI_fam: phytoene desaturase<br>NAD(P)-binding Rossmann-like domain<br>Flavin containing amine oxidoreductase<br>TIGR00292: thiazole biosynthesis enzyme<br>desat_CrtD: C-3',4' desaturase CrtD | 1.40E-14<br>8.40E-10<br>1.10E-09<br>5.30E-08<br>3.10E-07 |
| BCB83520.1 | 935891 | 935147 | - | 247 | PF13649<br>PF08241<br>PF13489<br>PF08242<br>PF13847 | Methyltransf_25<br>Methyltransf_11<br>Methyltransf_23<br>Methyltransf_12<br>Methyltransf_31 | Methyltransferase domain<br>Methyltransferase domain<br>Methyltransferase domain<br>Methyltransferase domain<br>Methyltransferase domain | 6.90E-14<br>1.10E-11<br>1.90E-11<br>3.80E-11<br>6.80E-08 |
| BCB83521.1 | 936468 | 935916 | - | 183 | LinL | LinL | LinL | 1.60E-57 |

| | | | | | | | | |
|---|---|---|---|---|---|---|---|---|
| BCB83522.1 | 938060 | 936464 | - | 531 | LinH<br>LinE<br>LinG<br>PF12697<br>PF00561 | LinH<br>LinE<br>LinG<br>Abhydrolase_6<br>Abhydrolase_1 | LinH (Fusion of LinE/LinG)<br>LinE Homolog<br>LinG Homolog<br>Alpha/beta hydrolase family<br>alpha/beta hydrolase fold | 3.00E-201<br>6.20E-104<br>4.50E-73<br>3.40E-17<br>7.20E-11 |
| BCB83523.1 | 937950 | 938343 | + | 130 | NO PFAM MATCH | - | - | - |
| BCB83524.1 | 938659 | 938419 | - | 79 | LinA | LinA | LinA (linaridin precursor) | 2.00E-10 |
| BCB83525.1 | 939036 | 939672 | + | 211 | TIGR01189<br>TIGR02857<br>TIGR00968<br>TIGR03873<br>TIGR01166 | TIGR01189<br>TIGR02857<br>TIGR00968<br>TIGR03873<br>TIGR01166 | ccmA: heme ABC exporter, ATP-binding protein CcmA<br>CydD: thiol reductant ABC exporter, CydD subunit<br>3a0106s01: sulfate ABC transporter, ATP-binding protein<br>F420-0_ABC_ATP: proposed F420-0 ABC transporter, ATP-binding protein<br>cbiO: cobalt ABC transporter, ATP-binding protein | 2.80E-33<br>3.80E-31<br>1.50E-27<br>6.00E-27<br>9.60E-26 |
| BCB83526.1 | 939671 | 941207 | + | 511 | NO PFAM MATCH | - | - | - |
| BCB83527.1 | 942492 | 941907 | - | 194 | PF07920 | DUF1684 | Protein of unknown function (DUF1684) | 2.70E-38 |
| BCB83528.1 | 942641 | 943502 | + | 286 | PF01513 | NAD_kinase | ATP-NAD kinase | 2.20E-48 |
| BCB83529.1 | 943571 | 944036 | + | 154 | PF12802<br>PF01047<br>TIGR02337<br>PF13463<br>PF13412 | MarR_2<br>MarR<br>TIGR02337<br>HTH_27<br>HTH_24 | MarR family<br>MarR family<br>HpaR: homoprotocatechuate degradation operon regulator, HpaR<br>Winged helix DNA-binding domain<br>Winged helix-turn-helix DNA-binding | 7.60E-15<br>3.00E-12<br>2.50E-07<br>2.20E-06<br>2.80E-04 |
| BCB83530.1 | 944047 | 945040 | + | 330 | PF02678<br>PF05726<br>PF07883 | Pirin<br>Pirin_C<br>Cupin_2 | Pirin<br>Pirin C-terminal cupin domain<br>Cupin domain | 9.80E-31<br>2.10E-29<br>2.30E-05 |
| BCB83531.1 | 945184 | 945055 | - | 42 | NO PFAM MATCH | - | - | - |

| index | leader | core | start | end | dir | score |
|---|---|---|---|---|---|---|
| 1 | VTISDKQRPQPADVRAAQCKN | PATSDMPHTVTVGKRGSGVDNR | 931867 | 931735 | - | 6 |
| 2 | VVTISDKQRPQPADVRAAQCKN | PATSDMPHTVTVGKRGSGVDNR | 931870 | 931735 | - | 6 |
| 3 | MSQRLRVASWLAAGPTALTMLVMGQ...SAAYGEATASGTFTRSVRTVTDILT | ITTVQIEGELVNRSSTCHTFQ | 933619 | 933361 | - | 6 |
| 4 | MVTRLPGAWIGHEMHVSPALLGGNM...VSEARFGRLSVETPAKKMSASAPVL | FTPAVVVVTAAALITAVGSV | 938659 | 938419 | - | 29 |
| 5 | VGCRTLRTRLRSVSDRDPPIARSRPPRGRQ | RIRGFHDQHTGVHGAVLCGAPAGPVQHGNAV | 941540 | 941354 | - | 7 |
| 6 | MAVARCGLQDSAHQAAIGVR | PRPADREITPAAWPAADTWLP | 941556 | 941430 | - | 6 |
| 7 | MAWTMAVARCGLQDSAHQAAIG | VRPRPADREITPAAWPAADTWLP | 941568 | 941430 | - | 7 |
| 8 | MMAWTMAVARCGLQDSAHQAAIG | VRPRPADREITPAAWPAADTWLP | 941571 | 941430 | - | 7 |
| 9 | MTTESIATTEQAQAPVEAGRG | LLLTMLLMVLLGVAGMYLVTL | 945184 | 945055 | - | 6 |

### Results for WP_084705756.1 [Prauserella rugosa] back to top

previous - next

#### Architecture

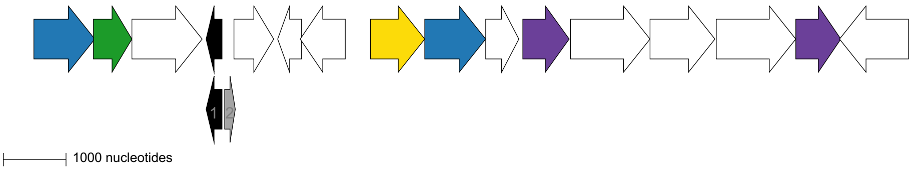

Link to nucleotide sequence

| Accession | start | end | direction | length (aa) | Pfam/HMM | name | description | E-value |
|---|---|---|---|---|---|---|---|---|
| WP_084705758.1 | 2743484 | 2744462 | + | 325 | LinE<br>LinH<br>TIGR02427<br>PF12697<br>PF00561 | LinE<br>LinH<br>TIGR02427<br>Abhydrolase_6<br>Abhydrolase_1 | LinE Homolog<br>LinH (Fusion of LinE/LinG)<br>protocat_pcaD: 3-oxoadipate enol-lactonase<br>Alpha/beta hydrolase family<br>alpha/beta hydrolase fold | 5.30E-86<br>1.10E-58<br>3.10E-13<br>2.10E-12<br>4.40E-12 |
| WP_084705757.1 | 2744424 | 2745045 | + | 206 | LinL | LinL | LinL | 2.00E-62 |
| WP_030531701.1 | 2745037 | 2746171 | + | 377 | TIGR02032<br>PF01494<br>PF01266<br>PF04820<br>TIGR01988 | TIGR02032<br>FAD_binding_3<br>DAO<br>Trp_halogenase<br>TIGR01988 | GG-red-SF: geranylgeranyl reductase family<br>FAD binding domain<br>FAD dependent oxidoreductase<br>Tryptophan halogenase<br>Ubi-OHases: ubiquinone biosynthesis hydroxylase, UbiH/UbiF/VisC/COQ6 family | 1.50E-14<br>1.00E-10<br>1.60E-10<br>2.20E-10<br>3.50E-07 |
| WP_030531700.1 | 2746484 | 2746226 | - | 85 | LinA | LinA | LinA (linaridin precursor) | 1.10E-14 |
| WP_030531699.1 | 2746671 | 2747319 | + | 215 | PF05719 | GPP34 | Golgi phosphoprotein 3 (GPP34) | 5.80E-21 |
| WP_030531698.1 | 2747755 | 2747371 | - | 127 | PF04134 | DUF393 | Protein of unknown function, DUF393 | 1.70E-18 |
| WP_145600453.1 | 2748443 | 2747744 | - | 232 | PF00196<br>TIGR03020<br>TIGR03541<br>TIGR02937<br>PF13384 | GerE<br>TIGR03020<br>TIGR03541<br>TIGR02937<br>HTH_23 | Bacterial regulatory proteins, luxR family<br>EpsA: transcriptional regulator EpsA<br>reg_near_HchA: LuxR family transcriptional regulatory, chaperone HchA-associated<br>sigma70-ECF: RNA polymerase sigma factor, sigma-70 family<br>Homeodomain-like domain | 1.80E-17<br>5.20E-16<br>4.50E-11<br>3.80E-05<br>3.10E-04 |

| | | | | | | | | |
|---|---|---|---|---|---|---|---|---|
| WP_145600456.1 | 2748848 | 2749718 | + | 289 | LinG<br>LinH<br>LinL<br>PF06762<br>TIGR04033 | LinG<br>LinH<br>LinL<br>LMF1<br>TIGR04033 | LinG Homolog<br>LinH (Fusion of LinE/LinG)<br>LinL<br>Lipase maturation factor<br>export_SdpB: antimicrobial peptide system protein, SdpB family | 3.80E-103<br>3.80E-96<br>1.00E-16<br>1.30E-04<br>1.70E-04 |
| WP_084705756.1 | 2749714 | 2750704 | + | 329 | LinE<br>LinH<br>PF12697<br>TIGR02427<br>PF00561 | LinE<br>LinH<br>Abhydrolase_6<br>TIGR02427<br>Abhydrolase_1 | LinE Homolog<br>LinH (Fusion of LinE/LinG)<br>Alpha/beta hydrolase family<br>protocat_pcaD: 3-oxoadipate enol-lactonase<br>alpha/beta hydrolase fold | 3.90E-99<br>3.80E-73<br>5.10E-12<br>5.40E-12<br>5.60E-11 |
| WP_018835202.1 | 2750700 | 2751231 | + | 177 | INFERRED GENE | - | - | - |
| WP_051757635.1 | 2751296 | 2752037 | + | 246 | PF13649<br>PF13489<br>PF08241<br>PF08242<br>PF13847 | Methyltransf_25<br>Methyltransf_23<br>Methyltransf_11<br>Methyltransf_12<br>Methyltransf_31 | Methyltransferase domain<br>Methyltransferase domain<br>Methyltransferase domain<br>Methyltransferase domain<br>Methyltransferase domain | 1.40E-13<br>8.90E-13<br>2.60E-11<br>7.30E-10<br>1.10E-08 |
| WP_030531692.1 | 2752040 | 2753327 | + | 428 | PF03033<br>TIGR01426 | Glyco_transf_28<br>TIGR01426 | Glycosyltransferase family 28 N-terminal domain<br>MGT: glycosyltransferase, MGT family | 1.00E-14<br>3.10E-14 |
| WP_030531691.1 | 2753314 | 2754358 | + | 347 | PF00067<br>TIGR04458 | p450<br>TIGR04458 | Cytochrome P450<br>CYP450_TxtE: 4-nitrotryptophan synthase | 3.60E-04<br>6.50E-04 |
| WP_030531690.1 | 2754378 | 2755641 | + | 420 | TIGR01426<br>PF06722<br>PF00201 | TIGR01426<br>DUF1205<br>UDPGT | MGT: glycosyltransferase, MGT family<br>Protein of unknown function (DUF1205)<br>UDP-glucoronosyl and UDP-glucosyl transferase | 3.30E-27<br>8.40E-24<br>1.10E-09 |
| WP_030531689.1 | 2755637 | 2756357 | + | 239 | PF13649<br>PF13847<br>PF08241<br>PF13489<br>PF05401 | Methyltransf_25<br>Methyltransf_31<br>Methyltransf_11<br>Methyltransf_23<br>NodS | Methyltransferase domain<br>Methyltransferase domain<br>Methyltransferase domain<br>Methyltransferase domain<br>Nodulation protein S (NodS) | 4.00E-15<br>2.00E-11<br>7.40E-11<br>1.00E-09<br>1.70E-07 |
| WP_030531688.1 | 2757447 | 2756337 | - | 369 | PF01041<br>TIGR03588<br>TIGR04427<br>TIGR04181<br>TIGR02379 | DegT_DnrJ_EryC1<br>TIGR03588<br>TIGR04427<br>TIGR04181<br>TIGR02379 | DegT/DnrJ/EryC1/StrS aminotransferase family<br>PseC: UDP-4-amino-4,6-dideoxy-N-acetyl-beta-L-altrosamine transaminase<br>PLP_DesI: dTDP-4-dehydro-6-deoxyglucose aminotransferase<br>NHT_00031: aminotransferase, LLPSF_NHT_00031 family<br>ECA_wecE: TDP-4-keto-6-deoxy-D-glucose transaminase | 2.90E-121<br>2.30E-65<br>2.60E-55<br>7.40E-55<br>7.00E-41 |

| index | leader | core | start | end | dir | score |
|---|---|---|---|---|---|---|
| 1 | MNPIEQAAGAATSGEGGLRPLERVEHCAPVL | GTPAWVTAAGVLYVAGNMVTDLIGQ...DGQGQDLTGKSGAELLDIRRSGMRA | 2746484 | 2746226 | - | 15 |
| 2 | MNVTNDFGGNIHDVAHFSL | HTTRFTAYCGMFVGVRPRDSLGEEAEPCRPTTTP | 2746534 | 2746696 | + | 6 |

### Results for WP_084705758.1 [Prauserella rugosa] back to top

previous - next

#### Architecture

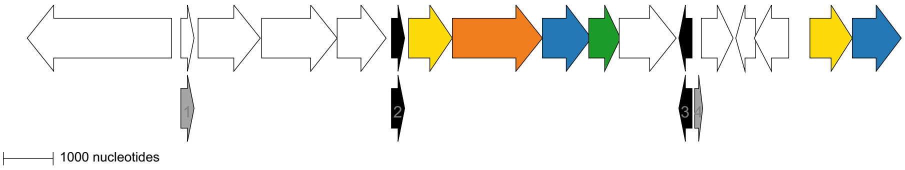

Link to nucleotide sequence

| Accession | start | end | direction | length (aa) | Pfam/HMM | name | description | E-value |
|---|---|---|---|---|---|---|---|---|
| WP_157575509.1 | 2736013 | 2733130 | - | 960 | PF17132 | Glyco_hydro_106 | alpha-L-rhamnosidase | 5.70E-78 |
| WP_030531710.1 | 2736215 | 2736488 | + | 90 | NO PFAM MATCH | - | - | - |
| WP_145600449.1 | 2736558 | 2737797 | + | 412 | TIGR04458<br>PF00067 | TIGR04458<br>p450 | CYP450_TxtE: 4-nitrotryptophan synthase<br>Cytochrome P450 | 2.60E-28<br>2.70E-15 |
| WP_145600451.1 | 2737837 | 2739352 | + | 504 | TIGR04022<br>PF08028<br>PF00441 | TIGR04022<br>Acyl-CoA_dh_2<br>Acyl-CoA_dh_1 | sulfur_SfnB: sulfur acquisition oxidoreductase, SfnB family<br>Acyl-CoA dehydrogenase, C-terminal domain<br>Acyl-CoA dehydrogenase, C-terminal domain | 1.70E-13<br>8.30E-09<br>2.50E-04 |
| WP_030531707.1 | 2739348 | 2740332 | + | 327 | PF08028<br>TIGR04022<br>PF00441 | Acyl-CoA_dh_2<br>TIGR04022<br>Acyl-CoA_dh_1 | Acyl-CoA dehydrogenase, C-terminal domain<br>sulfur_SfnB: sulfur acquisition oxidoreductase, SfnB family<br>Acyl-CoA dehydrogenase, C-terminal domain | 1.90E-18<br>1.20E-06<br>6.30E-05 |
| WP_030531706.1 | 2740452 | 2740713 | + | 86 | LinA | LinA | LinA (linaridin precursor) | 8.30E-15 |
| WP_030531705.1 | 2740793 | 2741666 | + | 290 | LinG<br>LinH | LinG<br>LinH | LinG Homolog<br>LinH (Fusion of LinE/LinG) | 1.80E-105<br>1.80E-89 |
| WP_051757684.1 | 2741670 | 2743488 | + | 605 | TIGR02204<br>TIGR02203<br>TIGR00958<br>TIGR03796<br>TIGR01846 | TIGR02204<br>TIGR02203<br>TIGR00958<br>TIGR03796<br>TIGR01846 | MsbA_rel: ABC transporter, permease/ATP-binding protein<br>MsbA_lipidA: lipid A export permease/ATP-binding protein MsbA<br>3a01208: antigen peptide transporter 2<br>NHLM_micro_ABC1: NHLM bacteriocin system ABC transporter, peptidase/ATP-binding protein<br>type_I_sec_HlyB: type I secretion system ATPase | 4.20E-103<br>4.00E-94<br>6.30E-85<br>1.40E-81<br>1.60E-78 |
| WP_084705758.1 | 2743484 | 2744462 | + | 325 | LinE<br>LinH<br>TIGR02427<br>PF12697<br>PF00561 | LinE<br>LinH<br>TIGR02427<br>Abhydrolase_6<br>Abhydrolase_1 | LinE Homolog<br>LinH (Fusion of LinE/LinG)<br>protocat_pcaD: 3-oxoadipate enol-lactonase<br>Alpha/beta hydrolase family<br>alpha/beta hydrolase fold | 5.30E-86<br>1.10E-58<br>3.10E-13<br>2.10E-12<br>4.40E-12 |

| | | | | | | | | |
|---|---|---|---|---|---|---|---|---|
| WP_084705757.1 | 2744424 | 2745045 | + | 206 | LinL | LinL | LinL | 2.00E-62 |
| WP_030531701.1 | 2745037 | 2746171 | + | 377 | TIGR02032 | TIGR02032 | GG-red-SF: geranylgeranyl reductase family | 1.50E-14 |
| | | | | | PF01494 | FAD_binding_3 | FAD binding domain | 1.00E-10 |
| | | | | | PF01266 | DAO | FAD dependent oxidoreductase | 1.60E-10 |
| | | | | | PF04820 | Trp_halogenase | Tryptophan halogenase | 2.20E-10 |
| | | | | | TIGR01988 | TIGR01988 | Ubi-OHases: ubiquinone biosynthesis hydroxylase, UbiH/UbiF/VisC/COQ6 family | 3.50E-07 |
| WP_030531700.1 | 2746484 | 2746226 | - | 85 | LinA | LinA | LinA (linaridin precursor) | 1.10E-14 |
| WP_030531699.1 | 2746671 | 2747319 | + | 215 | PF05719 | GPP34 | Golgi phosphoprotein 3 (GPP34) | 5.80E-21 |
| WP_030531698.1 | 2747755 | 2747371 | - | 127 | PF04134 | DUF393 | Protein of unknown function, DUF393 | 1.70E-18 |
| WP_145600453.1 | 2748443 | 2747744 | - | 232 | PF00196 | GerE | Bacterial regulatory proteins, luxR family | 1.80E-17 |
| | | | | | TIGR03020 | TIGR03020 | EpsA: transcriptional regulator EpsA | 5.20E-16 |
| | | | | | TIGR03541 | TIGR03541 | reg_near_HchA: LuxR family transcriptional regulatory, chaperone HchA-associated | 4.50E-11 |
| | | | | | TIGR02937 | TIGR02937 | sigma70-ECF: RNA polymerase sigma factor, sigma-70 family | 3.80E-05 |
| | | | | | PF13384 | HTH_23 | Homeodomain-like domain | 3.10E-04 |
| WP_145600456.1 | 2748848 | 2749718 | + | 289 | LinG | LinG | LinG Homolog | 3.80E-103 |
| | | | | | LinH | LinH | LinH (Fusion of LinE/LinG) | 3.80E-96 |
| | | | | | LinL | LinL | LinL | 1.00E-16 |
| | | | | | PF06762 | LMF1 | Lipase maturation factor | 1.30E-04 |
| | | | | | TIGR04033 | TIGR04033 | export_SdpB: antimicrobial peptide system protein, SdpB family | 1.70E-04 |
| WP_084705756.1 | 2749714 | 2750704 | + | 329 | LinE | LinE | LinE Homolog | 3.90E-99 |
| | | | | | LinH | LinH | LinH (Fusion of LinE/LinG) | 3.80E-73 |
| | | | | | PF12697 | Abhydrolase_6 | Alpha/beta hydrolase family | 5.10E-12 |
| | | | | | TIGR02427 | TIGR02427 | protocat_pcaD: 3-oxoadipate enol-lactonase | 5.40E-12 |
| | | | | | PF00561 | Abhydrolase_1 | alpha/beta hydrolase fold | 5.60E-11 |

| index | leader | core | start | end | dir | score |
|---|---|---|---|---|---|---|
| 1 | MPTPTPTATPAALAANASDHVEPGAISSAAPAQIDIAQVTTGVLT | ATTYQAIVDAAGGSAGGTSHIAAIEIHEHASVTELTAARAGVLTD | 2736215 | 2736488 | + | 6 |
| 2 | MSNVEAFANQHADNVLSPLEVTSAVGPVL | ATPAYAGYTAAAAAVFTAGLISDAV...VTPTGPVPNYGSVGDLLDMRVSSIN | 2740452 | 2740713 | + | 27 |
| 3 | MNPIEQAAGAATSGEGGLRPLERVEHCAPVL | GTPAWVTAAGVLYVAGNMVTDLIGQ...DGQGQDLTGKSGAELLDIRRSGMRA | 2746484 | 2746226 | - | 15 |
| 4 | MNVTNDFGGNIHDVAHFSL | HTTRFTAYCGMFVGVRPRDSLGEEAEPCRPTTTP | 2746534 | 2746696 | + | 6 |

### Results for KID32044.1 [Prauserella sp. Am3] back to top

previous - next

#### Architecture

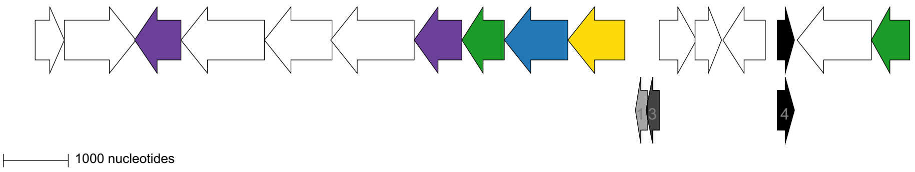

Link to nucleotide sequence

| Accession | start | end | direction | length (aa) | Pfam/HMM | name | description | E-value |
|---|---|---|---|---|---|---|---|---|
| KID32036.1 | 422417 | 422861 | + | 147 | PF05523 | FdtA | WxcM-like, C-terminal | 7.20E-47 |
| KID32037.1 | 422857 | 423967 | + | 369 | PF01041 | DegT_DnrJ_EryC1 | DegT/DnrJ/EryC1/StrS aminotransferase family | 3.80E-118 |
| | | | | | TIGR03588 | TIGR03588 | PseC: UDP-4-amino-4,6-dideoxy-N-acetyl-beta-L-altrosamine transaminase | 4.80E-64 |
| | | | | | TIGR04427 | TIGR04427 | PLP_DesI: dTDP-4-dehydro-6-deoxyglucose aminotransferase | 4.00E-54 |
| | | | | | TIGR04181 | TIGR04181 | NHT_00031: aminotransferase, LLPSF_NHT_00031 family | 3.30E-52 |
| | | | | | TIGR02379 | TIGR02379 | ECA_wecE: TDP-4-keto-6-deoxy-D-glucose transaminase | 4.30E-39 |
| KID32038.1 | 424667 | 423947 | - | 239 | PF13649 | Methyltransf_25 | Methyltransferase domain | 4.00E-15 |
| | | | | | PF13847 | Methyltransf_31 | Methyltransferase domain | 2.10E-11 |
| | | | | | PF08241 | Methyltransf_11 | Methyltransferase domain | 7.40E-11 |
| | | | | | PF13489 | Methyltransf_23 | Methyltransferase domain | 7.00E-10 |
| | | | | | PF05401 | NodS | Nodulation protein S (NodS) | 1.70E-07 |
| KID32039.1 | 425950 | 424663 | - | 428 | TIGR01426 | TIGR01426 | MGT: glycosyltransferase, MGT family | 3.40E-27 |
| | | | | | PF06722 | DUF1205 | Protein of unknown function (DUF1205) | 8.50E-24 |
| | | | | | PF00201 | UDPGT | UDP-glucoronosyl and UDP-glucosyl transferase | 1.20E-09 |
| KID32040.1 | 426990 | 425946 | - | 347 | NO PFAM MATCH | - | - | - |
| KID32041.1 | 428252 | 426977 | - | 424 | TIGR01426 | TIGR01426 | MGT: glycosyltransferase, MGT family | 1.80E-13 |
| | | | | | PF03033 | Glyco_transf_28 | Glycosyltransferase family 28 N-terminal domain | 2.50E-12 |
| | | | | | PF00201 | UDPGT | UDP-glucoronosyl and UDP-glucosyl transferase | 1.20E-04 |
| KID32042.1 | 429007 | 428266 | - | 246 | PF13489 | Methyltransf_23 | Methyltransferase domain | 1.10E-13 |
| | | | | | PF13649 | Methyltransf_25 | Methyltransferase domain | 1.10E-13 |
| | | | | | PF08241 | Methyltransf_11 | Methyltransferase domain | 1.70E-11 |
| | | | | | PF08242 | Methyltransf_12 | Methyltransferase domain | 4.70E-10 |
| | | | | | PF13847 | Methyltransf_31 | Methyltransferase domain | 2.50E-08 |
| KID32043.1 | 429654 | 428991 | - | 220 | LinL | LinL | LinL | 1.70E-73 |
| KID32044.1 | 430640 | 429650 | - | 329 | LinE | LinE | LinE Homolog | 1.60E-100 |
| | | | | | LinH | LinH | LinH (Fusion of LinE/LinG) | 4.00E-75 |
| | | | | | TIGR02427 | TIGR02427 | protocat_pcaD: 3-oxoadipate enol-lactonase | 8.10E-13 |
| | | | | | PF12697 | Abhydrolase_6 | Alpha/beta hydrolase family | 1.60E-12 |
| | | | | | PF00561 | Abhydrolase_1 | alpha/beta hydrolase fold | 2.90E-11 |

| | | | | | | | | |
|---|---|---|---|---|---|---|---|---|
| KID32045.1 | 431503 | 430636 | - | 288 | LinG<br>LinH<br>LinL<br>TIGR04033<br>PF06762 | LinG<br>LinH<br>LinL<br>TIGR04033<br>LMF1 | LinG Homolog<br>LinH (Fusion of LinE/LinG)<br>LinL<br>export_SdpB: antimicrobial peptide system protein, SdpB family<br>Lipase maturation factor | 1.80E-102<br>2.20E-95<br>1.90E-16<br>4.50E-06<br>4.00E-05 |
| KID32046.1 | 432045 | 432609 | + | 187 | PF00196<br>TIGR03020<br>TIGR03541<br>TIGR02937<br>PF08281 | GerE<br>TIGR03020<br>TIGR03541<br>TIGR02937<br>Sigma70_r4_2 | Bacterial regulatory proteins, luxR family<br>EpsA: transcriptional regulator EpsA<br>reg_near_HchA: LuxR family transcriptional regulatory, chaperone HchA-associated<br>sigma70-ECF: RNA polymerase sigma factor, sigma-70 family<br>Sigma-70, region 4 | 1.80E-17<br>3.60E-16<br>1.90E-11<br>3.40E-05<br>4.00E-04 |
| KID32047.1 | 432601 | 432994 | + | 130 | PF04134 | DUF393 | Protein of unknown function, DUF393 | 6.20E-18 |
| KID32048.1 | 433676 | 433028 | - | 215 | PF05719 | GPP34 | Golgi phosphoprotein 3 (GPP34) | 4.40E-23 |
| KID32049.1 | 433859 | 434117 | + | 85 | LinA | LinA | LinA (linaridin precursor) | 2.10E-14 |
| KID32050.1 | 435317 | 434168 | - | 382 | TIGR02032<br>PF01494<br>PF04820<br>PF01266<br>TIGR01988 | TIGR02032<br>FAD_binding_3<br>Trp_halogenase<br>DAO<br>TIGR01988 | GG-red-SF: geranylgeranyl reductase family<br>FAD binding domain<br>Tryptophan halogenase<br>FAD dependent oxidoreductase<br>Ubi-OHases: ubiquinone biosynthesis hydroxylase, UbiH/UbiF/VisC/COQ6 family | 1.90E-15<br>6.80E-12<br>9.40E-12<br>1.60E-08<br>1.90E-07 |
| KID32051.1 | 435903 | 435309 | - | 197 | LinL | LinL | LinL | 6.60E-63 |

| index | leader | core | start | end | dir | score |
|---|---|---|---|---|---|---|
| 1 | VPTVAEKSHRHPDSARNRRKHCVPIPM | PTVTLRCTEQRLRSTRRDTATAPTGDLCDSDRS | 431856 | 431673 | - | 6 |
| 2 | MVCDTCCIRNSEPVEIHEVQPREHGFYRPRRSIFDDPQSNHATSLSVER | LTIRTQDFCVRCPQ | 432035 | 431843 | - | 7 |
| 3 | MASSMVCDTCCIRNSEPVEIHEVQP...GFYRPRRSIFDDPQSNHATSLSVER | LTIRTQDFCVRCPQ | 432047 | 431843 | - | 7 |
| 4 | MNPIEEAAGAATSGEGGLRPLEQVEHCAPVL | GTPAWVTAAGVLYVAGNMVTDLIGQ...DGQGQDLTGKSGAELLDIRRSGMRA | 433859 | 434117 | + | 15 |

### Results for KID32052.1 [Prauserella sp. Am3] back to top

previous - next

#### Architecture

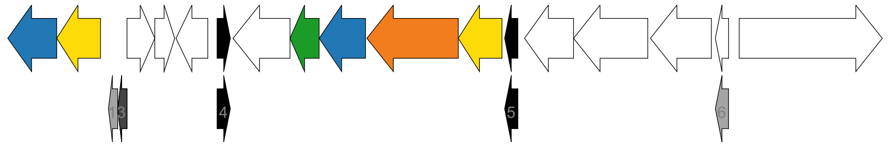

1000 nucleotides

Link to nucleotide sequence

| Accession | start | end | direction | length (aa) | Pfam/HMM | name | description | E-value |
|---|---|---|---|---|---|---|---|---|
| KID32044.1 | 430640 | 429650 | - | 329 | LinE<br>LinH<br>TIGR02427<br>PF12697<br>PF00561 | LinE<br>LinH<br>TIGR02427<br>Abhydrolase_6<br>Abhydrolase_1 | LinE Homolog<br>LinH (Fusion of LinE/LinG)<br>protocat_pcaD: 3-oxoadipate enol-lactonase<br>Alpha/beta hydrolase family<br>alpha/beta hydrolase fold | 1.60E-100<br>4.00E-75<br>8.10E-13<br>1.60E-12<br>2.90E-11 |
| KID32045.1 | 431503 | 430636 | - | 288 | LinG<br>LinH<br>LinL<br>TIGR04033<br>PF06762 | LinG<br>LinH<br>LinL<br>TIGR04033<br>LMF1 | LinG Homolog<br>LinH (Fusion of LinE/LinG)<br>LinL<br>export_SdpB: antimicrobial peptide system protein, SdpB family<br>Lipase maturation factor | 1.80E-102<br>2.20E-95<br>1.90E-16<br>4.50E-06<br>4.00E-05 |
| KID32046.1 | 432045 | 432609 | + | 187 | PF00196<br>TIGR03020<br>TIGR03541<br>TIGR02937<br>PF08281 | GerE<br>TIGR03020<br>TIGR03541<br>TIGR02937<br>Sigma70_r4_2 | Bacterial regulatory proteins, luxR family<br>EpsA: transcriptional regulator EpsA<br>reg_near_HchA: LuxR family transcriptional regulatory, chaperone HchA-associated<br>sigma70-ECF: RNA polymerase sigma factor, sigma-70 family<br>Sigma-70, region 4 | 1.80E-17<br>3.60E-16<br>1.90E-11<br>3.40E-05<br>4.00E-04 |
| KID32047.1 | 432601 | 432994 | + | 130 | PF04134 | DUF393 | Protein of unknown function, DUF393 | 6.20E-18 |
| KID32048.1 | 433676 | 433028 | - | 215 | PF05719 | GPP34 | Golgi phosphoprotein 3 (GPP34) | 4.40E-23 |
| KID32049.1 | 433859 | 434117 | + | 85 | LinA | LinA | LinA (linaridin precursor) | 2.10E-14 |
| KID32050.1 | 435317 | 434168 | - | 382 | TIGR02032<br>PF01494<br>PF04820<br>PF01266<br>TIGR01988 | TIGR02032<br>FAD_binding_3<br>Trp_halogenase<br>DAO<br>TIGR01988 | GG-red-SF: geranylgeranyl reductase family<br>FAD binding domain<br>Tryptophan halogenase<br>FAD dependent oxidoreductase<br>Ubi-OHases: ubiquinone biosynthesis hydroxylase, UbiH/UbiF/VisC/COQ6 family | 1.90E-15<br>6.80E-12<br>9.40E-12<br>1.60E-08<br>1.90E-07 |
| KID32051.1 | 435903 | 435309 | - | 197 | LinL | LinL | LinL | 6.60E-63 |
| KID32052.1 | 436828 | 435892 | - | 311 | LinE<br>LinH<br>TIGR02427<br>PF12697<br>PF00561 | LinE<br>LinH<br>TIGR02427<br>Abhydrolase_6<br>Abhydrolase_1 | LinE Homolog<br>LinH (Fusion of LinE/LinG)<br>protocat_pcaD: 3-oxoadipate enol-lactonase<br>Alpha/beta hydrolase family<br>alpha/beta hydrolase fold | 3.70E-85<br>3.30E-58<br>2.80E-13<br>1.50E-12<br>4.80E-12 |
| KID32053.1 | 438684 | 436866 | - | 605 | TIGR02204<br>TIGR02203<br>TIGR00958<br>TIGR03796<br>TIGR01846 | TIGR02204<br>TIGR02203<br>TIGR00958<br>TIGR03796<br>TIGR01846 | MsbA_rel: ABC transporter, permease/ATP-binding protein<br>MsbA_lipidA: lipid A export permease/ATP-binding protein MsbA<br>3a01208: antigen peptide transporter 2<br>NHLM_micro_ABC1: NHLM bacteriocin system ABC transporter, peptidase/ATP-binding protein<br>type_I_sec_HlyB: type I secretion system ATPase | 3.00E-103<br>1.20E-93<br>5.10E-85<br>5.20E-81<br>1.60E-78 |
| KID32054.1 | 439561 | 438688 | - | 290 | LinG<br>LinH | LinG<br>LinH | LinG Homolog<br>LinH (Fusion of LinE/LinG) | 6.30E-106<br>8.00E-90 |
| KID32055.1 | 439898 | 439637 | - | 86 | LinA | LinA | LinA (linaridin precursor) | 1.10E-14 |

| | | | | | | | | |
|---|---|---|---|---|---|---|---|---|
| KID32056.1 | 441002 | 440018 | - | 327 | PF08028<br>TIGR04022<br>PF00441 | Acyl-CoA_dh_2<br>TIGR04022<br>Acyl-CoA_dh_1 | Acyl-CoA dehydrogenase, C-terminal domain<br>sulfur_SfnB: sulfur acquisition oxidoreductase, SfnB family<br>Acyl-CoA dehydrogenase, C-terminal domain | 1.90E-18<br>1.10E-06<br>6.20E-05 |
| KID32057.1 | 442513 | 440998 | - | 504 | TIGR04022<br>PF08028<br>PF00441 | TIGR04022<br>Acyl-CoA_dh_2<br>Acyl-CoA_dh_1 | sulfur_SfnB: sulfur acquisition oxidoreductase, SfnB family<br>Acyl-CoA dehydrogenase, C-terminal domain<br>Acyl-CoA dehydrogenase, C-terminal domain | 2.10E-13<br>1.90E-08<br>5.30E-04 |
| KID32058.1 | 443791 | 442549 | - | 413 | TIGR04458<br>PF00067 | TIGR04458<br>p450 | CYP450_TxtE: 4-nitrotryptophan synthase<br>Cytochrome P450 | 4.70E-27<br>7.40E-15 |
| KID32059.1 | 444134 | 443861 | - | 90 | NO PFAM MATCH | - | - | - |
| KID32060.1 | 444342 | 447222 | + | 959 | PF17132 | Glyco_hydro_106 | alpha-L-rhamnosidase | 1.20E-74 |

| index | leader | core | start | end | dir | score |
|---|---|---|---|---|---|---|
| 1 | VPTVAEKSHRHPDSARNRRKHCVPIPM | PTVTLRCTEQRLRSTRRDTATAPTGDLCDSDRS | 431856 | 431673 | - | 6 |
| 2 | MVCDTCCIRNSEPVEIHEVQPREHGFYRPRRSIFDDPQSNHATSLSVER | LTIRTQDFCVRCPQ | 432035 | 431843 | - | 7 |
| 3 | MASSMVCDTCCIRNSEPVEIHEVQP...GFYRPRRSIFDDPQSNHATSLSVER | LTIRTQDFCVRCPQ | 432047 | 431843 | - | 7 |
| 4 | MNPIEEAAGAATSGEGGLRPLEQVEHCAPVL | GTPAWVTAAGVLYVAGNMVTDLIGQ...DGQGQDLTGKSGAELLDIRRSGMRA | 433859 | 434117 | + | 15 |
| 5 | MSNVEAFANQHADNVLSPLEVTSAVGPVL | ATPAYAGYTAAAAAVFTAGLISDAV...VTPTGPVPNYGSVGDLLDMRVSSIN | 439898 | 439637 | - | 27 |
| 6 | MPTPTPTATPAALAANASDHVEPGAISSAAPAQIDIAQVTTGVLT | ATTYQAIVDAAGGSAGGTSHIAAIEIHEHASVTELTAARAGALTD | 444134 | 443861 | - | 6 |

### Results for PXY36318.1 [Prauserella sp. YIM 121212] back to top

previous - next

#### Architecture

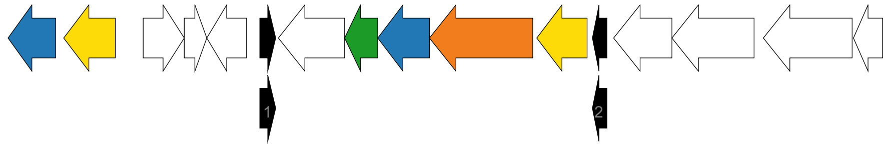

1000 nucleotides

Link to nucleotide sequence

| Accession | start | end | direction | length (aa) | Pfam/HMM | name | description | E-value |
|---|---|---|---|---|---|---|---|---|
| PXY36951.1 | 1183736 | 1182908 | - | 275 | LinE<br>LinH<br>PF12697<br>TIGR02427<br>PF00561 | LinE<br>LinH<br>Abhydrolase_6<br>TIGR02427<br>Abhydrolase_1 | LinE Homolog<br>LinH (Fusion of LinE/LinG)<br>Alpha/beta hydrolase family<br>protocat_pcaD: 3-oxoadipate enol-lactonase<br>alpha/beta hydrolase fold | 2.00E-88<br>6.60E-67<br>1.80E-09<br>2.50E-09<br>5.70E-09 |
| PXY36311.1 | 1184764 | 1183870 | - | 297 | LinG<br>LinH<br>LinL<br>TIGR04033<br>PF06762 | LinG<br>LinH<br>LinL<br>TIGR04033<br>LMF1 | LinG Homolog<br>LinH (Fusion of LinE/LinG)<br>LinL<br>export_SdpB: antimicrobial peptide system protein, SdpB family<br>Lipase maturation factor | 1.50E-103<br>1.10E-95<br>1.40E-16<br>1.70E-05<br>3.40E-05 |
| PXY36312.1 | 1185236 | 1185947 | + | 236 | PF00196<br>TIGR03020<br>TIGR03541 | GerE<br>TIGR03020<br>TIGR03541 | Bacterial regulatory proteins, luxR family<br>EpsA: transcriptional regulator EpsA<br>reg_near_HchA: LuxR family transcriptional regulatory, chaperone HchA-associated | 7.90E-18<br>4.10E-15<br>2.40E-11 |
| PXY36313.1 | 1185939 | 1186314 | + | 124 | PF04134 | DUF393 | Protein of unknown function, DUF393 | 1.60E-20 |
| PXY36314.1 | 1187000 | 1186331 | - | 222 | PF05719 | GPP34 | Golgi phosphoprotein 3 (GPP34) | 9.90E-26 |
| PXY36315.1 | 1187238 | 1187505 | + | 88 | LinA | LinA | LinA (linaridin precursor) | 9.20E-13 |
| PXY36316.1 | 1188689 | 1187552 | - | 378 | PF04820<br>TIGR02032<br>PF01494<br>PF12831<br>TIGR03197 | Trp_halogenase<br>TIGR02032<br>FAD_binding_3<br>FAD_oxidored<br>TIGR03197 | Tryptophan halogenase<br>GG-red-SF: geranylgeranyl reductase family<br>FAD binding domain<br>FAD dependent oxidoreductase<br>MnmC_Cterm: tRNA U-34 5-methylaminomethyl-2-thiouridine biosynthesis protein MnmC, C-terminal domain | 9.60E-12<br>4.10E-10<br>4.60E-07<br>3.20E-06<br>5.30E-05 |
| PXY36317.1 | 1189258 | 1188685 | - | 190 | LinL | LinL | LinL | 1.40E-60 |
| PXY36318.1 | 1190150 | 1189247 | - | 300 | LinE<br>LinH<br>TIGR02427<br>PF00561<br>PF12697 | LinE<br>LinH<br>TIGR02427<br>Abhydrolase_1<br>Abhydrolase_6 | LinE Homolog<br>LinH (Fusion of LinE/LinG)<br>protocat_pcaD: 3-oxoadipate enol-lactonase<br>alpha/beta hydrolase fold<br>Alpha/beta hydrolase family | 8.90E-88<br>1.40E-60<br>1.50E-14<br>1.10E-13<br>1.50E-11 |
| PXY36952.1 | 1191921 | 1190142 | - | 592 | TIGR02204<br>TIGR02203<br>TIGR00958<br>TIGR03796<br>TIGR01846 | TIGR02204<br>TIGR02203<br>TIGR00958<br>TIGR03796<br>TIGR01846 | MsbA_rel: ABC transporter, permease/ATP-binding protein<br>MsbA_lipidA: lipid A export permease/ATP-binding protein MsbA<br>3a01208: antigen peptide transporter 2<br>NHLM_micro_ABC1: NHLM bacteriocin system ABC transporter, peptidase/ATP-binding protein<br>type_I_sec_HlyB: type I secretion system ATPase | 3.00E-105<br>1.00E-94<br>1.60E-84<br>7.70E-82<br>1.40E-74 |
| PXY36319.1 | 1192841 | 1191986 | - | 284 | LinG<br>LinH | LinG<br>LinH | LinG Homolog<br>LinH (Fusion of LinE/LinG) | 3.70E-103<br>6.70E-90 |
| PXY36320.1 | 1193185 | 1192924 | - | 86 | LinA | LinA | LinA (linaridin precursor) | 4.00E-14 |
| PXY36321.1 | 1194289 | 1193287 | - | 333 | PF08028<br>TIGR04022<br>PF00441 | Acyl-CoA_dh_2<br>TIGR04022<br>Acyl-CoA_dh_1 | Acyl-CoA dehydrogenase, C-terminal domain<br>sulfur_SfnB: sulfur acquisition oxidoreductase, SfnB family<br>Acyl-CoA dehydrogenase, C-terminal domain | 1.90E-20<br>7.50E-09<br>1.20E-07 |
| PXY36322.1 | 1195710 | 1194285 | - | 474 | TIGR04022<br>PF08028 | TIGR04022<br>Acyl-CoA_dh_2 | sulfur_SfnB: sulfur acquisition oxidoreductase, SfnB family<br>Acyl-CoA dehydrogenase, C-terminal domain | 1.60E-16<br>8.70E-08 |

| | | | | | | | | |
|---|---|---|---|---|---|---|---|---|
| PXY36323.1 | 1197379 | 1195858 | - | 506 | PF01970 | TctA | Tripartite tricarboxylate transporter TctA family | 4.50E-137 |
| PXY36324.1 | 1197895 | 1197400 | - | 164 | PF07331 | TctB | Tripartite tricarboxylate transporter TctB family | 3.80E-15 |

| index | leader | core | start | end | dir | score |
|---|---|---|---|---|---|---|
| 1 | MNSVEKAADAAMSREDWLQPLERADKVAPVL | GTPGIAAVVACGTALFAAGNMVTDL...DLPGEGVSGKSGTELLALRREGLRG | 1187238 | 1187505 | + | 13 |
| 2 | MSNVEAFANEHVDGVLSPLEVTTAVGPVL | ATPAYAGYAAGAGAVFTLGLISDAI...VTPTGPVPSYGSVDQLLGMRVASIS | 1193185 | 1192924 | - | 27 |

### Results for PXY36951.1 [Prauserella sp. YIM 121212] back to top

previous - next

#### Architecture

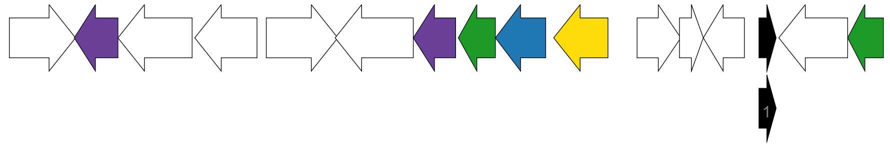

1000 nucleotides

Link to nucleotide sequence

| Accession | start | end | direction | length (aa) | Pfam/HMM | name | description | E-value |
|---|---|---|---|---|---|---|---|---|
| PXY36305.1 | 1174957 | 1176055 | + | 365 | PF01041<br>TIGR03588<br>TIGR04427<br>TIGR04181<br>TIGR02379 | DegT_DnrJ_EryC1<br>TIGR03588<br>TIGR04427<br>TIGR04181<br>TIGR02379 | DegT/DnrJ/EryC1/StrS aminotransferase family<br>PseC: UDP-4-amino-4,6-dideoxy-N-acetyl-beta-L-altrosamine transaminase<br>PLP_DesI: dTDP-4-dehydro-6-deoxyglucose aminotransferase<br>NHT_00031: aminotransferase, LLPSF_NHT_00031 family<br>ECA_wecE: TDP-4-keto-6-deoxy-D-glucose transaminase | 2.10E-123<br>1.60E-65<br>2.40E-59<br>3.70E-51<br>1.30E-39 |
| PXY36306.1 | 1176734 | 1176035 | - | 232 | PF13649<br>PF08241<br>PF13847<br>PF13489<br>PF08242 | Methyltransf_25<br>Methyltransf_11<br>Methyltransf_31<br>Methyltransf_23<br>Methyltransf_12 | Methyltransferase domain<br>Methyltransferase domain<br>Methyltransferase domain<br>Methyltransferase domain<br>Methyltransferase domain | 1.00E-17<br>8.20E-14<br>7.80E-11<br>2.60E-09<br>3.00E-09 |
| PXY36949.1 | 1177948 | 1176736 | - | 403 | PF06722<br>TIGR01426<br>PF00201<br>PF04101 | DUF1205<br>TIGR01426<br>UDPGT<br>Glyco_tran_28_C | Protein of unknown function (DUF1205)<br>MGT: glycosyltransferase, MGT family<br>UDP-glucoronosyl and UDP-glucosyl transferase<br>Glycosyltransferase family 28 C-terminal domain | 1.10E-28<br>1.70E-23<br>9.60E-06<br>6.30E-05 |
| PXY36307.1 | 1179027 | 1178010 | - | 338 | NO PFAM MATCH | - | - | - |
| PXY36308.1 | 1179178 | 1180336 | + | 385 | PF17843<br>PF13578 | MycE_N<br>Methyltransf_24 | MycE methyltransferase N-terminal<br>Methyltransferase domain | 5.30E-15<br>6.30E-05 |
| PXY36309.1 | 1181579 | 1180304 | - | 424 | TIGR01426<br>PF03033<br>PF00201 | TIGR01426<br>Glyco_transf_28<br>UDPGT | MGT: glycosyltransferase, MGT family<br>Glycosyltransferase family 28 N-terminal domain<br>UDP-glucoronosyl and UDP-glucosyl transferase | 5.30E-18<br>3.50E-14<br>7.80E-07 |
| PXY36950.1 | 1182262 | 1181575 | - | 228 | PF13649<br>PF13489<br>PF08241<br>PF08242<br>TIGR02021 | Methyltransf_25<br>Methyltransf_23<br>Methyltransf_11<br>Methyltransf_12<br>TIGR02021 | Methyltransferase domain<br>Methyltransferase domain<br>Methyltransferase domain<br>Methyltransferase domain<br>BchM-ChlM: magnesium protoporphyrin O-methyltransferase | 7.50E-13<br>1.10E-11<br>2.90E-10<br>1.20E-09<br>9.50E-09 |
| PXY36310.1 | 1182909 | 1182306 | - | 200 | LinL | LinL | LinL | 4.80E-67 |
| PXY36951.1 | 1183736 | 1182908 | - | 275 | LinE<br>LinH<br>PF12697<br>TIGR02427<br>PF00561 | LinE<br>LinH<br>Abhydrolase_6<br>TIGR02427<br>Abhydrolase_1 | LinE Homolog<br>LinH (Fusion of LinE/LinG)<br>Alpha/beta hydrolase family<br>protocat_pcaD: 3-oxoadipate enol-lactonase<br>alpha/beta hydrolase fold | 2.00E-88<br>6.60E-67<br>1.80E-09<br>2.50E-09<br>5.70E-09 |
| PXY36311.1 | 1184764 | 1183870 | - | 297 | LinG<br>LinH<br>LinL<br>TIGR04033<br>PF06762 | LinG<br>LinH<br>LinL<br>TIGR04033<br>LMF1 | LinG Homolog<br>LinH (Fusion of LinE/LinG)<br>LinL<br>export_SdpB: antimicrobial peptide system protein, SdpB family<br>Lipase maturation factor | 1.50E-103<br>1.10E-95<br>1.40E-16<br>1.70E-05<br>3.40E-05 |
| PXY36312.1 | 1185236 | 1185947 | + | 236 | PF00196<br>TIGR03020<br>TIGR03541 | GerE<br>TIGR03020<br>TIGR03541 | Bacterial regulatory proteins, luxR family<br>EpsA: transcriptional regulator EpsA<br>reg_near_HchA: LuxR family transcriptional regulatory, chaperone HchA-associated | 7.90E-18<br>4.10E-15<br>2.40E-11 |
| PXY36313.1 | 1185939 | 1186314 | + | 124 | PF04134 | DUF393 | Protein of unknown function, DUF393 | 1.60E-20 |
| PXY36314.1 | 1187000 | 1186331 | - | 222 | PF05719 | GPP34 | Golgi phosphoprotein 3 (GPP34) | 9.90E-26 |
| PXY36315.1 | 1187238 | 1187505 | + | 88 | LinA | LinA | LinA (linaridin precursor) | 9.20E-13 |
| PXY36316.1 | 1188689 | 1187552 | - | 378 | PF04820<br>TIGR02032<br>PF01494<br>PF12831<br>TIGR03197 | Trp_halogenase<br>TIGR02032<br>FAD_binding_3<br>FAD_oxidored<br>TIGR03197 | Tryptophan halogenase<br>GG-red-SF: geranylgeranyl reductase family<br>FAD binding domain<br>FAD dependent oxidoreductase<br>MnmC_Cterm: tRNA U-34 5-methylaminomethyl-2-thiouridine biosynthesis protein MnmC, C-terminal domain | 9.60E-12<br>4.10E-10<br>4.60E-07<br>3.20E-06<br>5.30E-05 |
| PXY36317.1 | 1189258 | 1188685 | - | 190 | LinL | LinL | LinL | 1.40E-60 |

| index | leader | core | start | end | dir | score |
|---|---|---|---|---|---|---|

| | | | | | | |
|---|---|---|---|---|---|---|
| 1 | MNSVEKAADAAMSREDWLQPLERADKVAPVL | GTPGIAAVVACGTALFAAGNMVTDL...DLPGEGVSGKSGTELLALRREGLRG | 1187238 | 1187505 | + | 13 |

### Results for TWD68636.1 [Rathayibacter rathayi] back to top

previous - next

#### Architecture

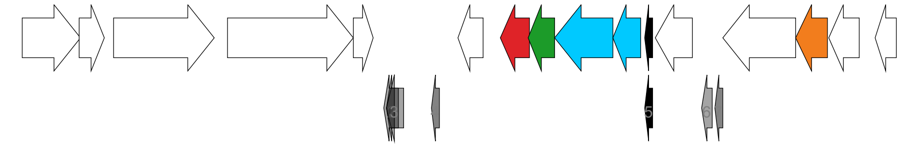

1000 nucleotides

Link to nucleotide sequence

| Accession | start | end | direction | length (aa) | Pfam/HMM | name | description | E-value |
|---|---|---|---|---|---|---|---|---|
| TWD68628.1 | 325072 | 326269 | + | 398 | TIGR00611<br>PF02463<br>PF13476<br>PF13175<br>TIGR00634 | TIGR00611<br>SMC_N<br>AAA_23<br>AAA_15<br>TIGR00634 | recf: DNA replication and repair protein RecF<br>RecF/RecN/SMC N terminal domain<br>AAA domain<br>AAA ATPase domain<br>recN: DNA repair protein RecN | 1.00E-104<br>1.20E-15<br>1.00E-05<br>6.90E-05<br>7.60E-04 |
| TWD68629.1 | 326261 | 326777 | + | 171 | PF05258 | DciA | Dna[CI] antecedent, DciA | 4.00E-21 |
| TWD68630.1 | 326954 | 329024 | + | 689 | TIGR01059<br>TIGR01058<br>TIGR01055<br>PF00204<br>PF00986 | TIGR01059<br>TIGR01058<br>TIGR01055<br>DNA_gyraseB<br>DNA_gyraseB_C | gyrB: DNA gyrase, B subunit<br>parE_Gpos: DNA topoisomerase IV, B subunit<br>parE_Gneg: DNA topoisomerase IV, B subunit<br>DNA gyrase B<br>DNA gyrase B subunit, carboxyl terminus | 2.80E-287<br>2.30E-229<br>9.70E-177<br>1.90E-62<br>6.10E-29 |
| TWD68631.1 | 329293 | 331894 | + | 866 | TIGR01063<br>TIGR01061<br>TIGR01062<br>PF00521<br>PF03989 | TIGR01063<br>TIGR01061<br>TIGR01062<br>DNA_topoisoIV<br>DNA_gyraseA_C | gyrA: DNA gyrase, A subunit<br>parC_Gpos: DNA topoisomerase IV, A subunit<br>parC_Gneg: DNA topoisomerase IV, A subunit<br>DNA gyrase/topoisomerase IV, subunit A<br>DNA gyrase C-terminal domain, beta-propeller | 0.00E+00<br>1.60E-182<br>1.30E-154<br>1.10E-142<br>3.60E-67 |
| TWD68632.1 | 331886 | 332297 | + | 136 | PF12089<br>PF07670 | DUF3566<br>Gate | Transmembrane domain of unknown function (DUF3566)<br>Nucleoside recognition | 3.50E-31<br>1.30E-04 |
| TWD68633.1 | 334566 | 334050 | - | 171 | NO PFAM MATCH | - | - | - |
| TWD68634.1 | 335504 | 334925 | - | 192 | TIGR00521<br>TIGR02113<br>PF02441 | TIGR00521<br>TIGR02113<br>Flavoprotein | coaBC_dfp: phosphopantothenoylcysteine decarboxylase / phosphopantothenate--cysteine ligase<br>coaC_strep: phosphopantothenoylcysteine decarboxylase<br>Flavoprotein | 2.20E-07<br>5.80E-06<br>7.00E-06 |
| TWD68635.1 | 336025 | 335500 | - | 174 | LinL | LinL | LinL | 1.00E-45 |
| TWD68636.1 | 337235 | 336044 | - | 396 | LinH<br>LinE<br>LinG<br>PF12697<br>TIGR02427 | LinH<br>LinE<br>LinG<br>Abhydrolase_6<br>TIGR02427 | LinH (Fusion of LinE/LinG)<br>LinE Homolog<br>LinG Homolog<br>Alpha/beta hydrolase family<br>protocat_pcaD: 3-oxoadipate enol-lactonase | 5.30E-98<br>9.60E-75<br>8.70E-26<br>6.50E-12<br>1.80E-04 |
| TWD68637.1 | 337818 | 337227 | - | 196 | LinH<br>LinG | LinH<br>LinG | LinH (Fusion of LinE/LinG)<br>LinG Homolog | 2.60E-57<br>6.50E-47 |
| TWD68638.1 | 338059 | 337894 | - | 54 | LinA | LinA | LinA (linaridin precursor) | 1.70E-10 |
| TWD68639.1 | 338875 | 338095 | - | 259 | PF13359<br>PF13613<br>PF01609<br>TIGR02937<br>PF13586 | DDE_Tnp_4<br>HTH_Tnp_4<br>DDE_Tnp_1<br>TIGR02937<br>DDE_Tnp_1_2 | DDE superfamily endonuclease<br>Helix-turn-helix of DDE superfamily endonuclease<br>Transposase DDE domain<br>sigma70-ECF: RNA polymerase sigma factor, sigma-70 family<br>Transposase DDE domain | 4.70E-21<br>7.60E-10<br>3.10E-09<br>6.70E-04<br>6.80E-04 |
| TWD68640.1 | 340987 | 339490 | - | 498 | NO PFAM MATCH | - | - | - |
| TWD68641.1 | 341661 | 340983 | - | 225 | TIGR01189<br>TIGR02673<br>TIGR02857<br>TIGR03258<br>TIGR01166 | TIGR01189<br>TIGR02673<br>TIGR02857<br>TIGR03258<br>TIGR01166 | ccmA: heme ABC exporter, ATP-binding protein CcmA<br>FtsE: cell division ATP-binding protein FtsE<br>CydD: thiol reductant ABC exporter, CydD subunit<br>PhnT: 2-aminoethylphosphonate ABC transport system, ATP-binding component PhnT<br>cbiO: cobalt ABC transporter, ATP-binding protein | 6.00E-33<br>1.70E-32<br>1.80E-30<br>2.40E-28<br>4.70E-28 |
| TWD68642.1 | 342305 | 341681 | - | 207 | PF13026 | DUF3887 | Protein of unknown function (DUF3887) | 8.40E-10 |
| TWD68643.1 | 343061 | 342632 | - | 142 | PF00293<br>TIGR02705<br>TIGR00586 | NUDIX<br>TIGR02705<br>TIGR00586 | NUDIX domain<br>nudix_YtkD: nucleoside triphosphatase YtkD<br>mutt: mutator mutT protein | 1.50E-20<br>1.90E-06<br>1.20E-05 |

| index | leader | core | start | end | dir | score |
|---|---|---|---|---|---|---|
| 1 | VRRLRKSTGETRWCETRNTPRRTPQGRDTQRPAIS | ETAISDTLWSLGETVEPRRFELLTSCLQSRRSTN | 332724 | 332514 | - | 6 |
| 2 | VCTSTQLWDSDRHHICIDPCSCLRRRSRRSDRRSAGETAEEID | GRDSMVRDPEHTSSDPAGPRYAKAGDLGDGDLGHTVEPRRDCGA | 332830 | 332566 | - | 6 |
| 3 | LSSTVQARRADDVKRCCAHSGTGQD...STQLWDSDRHHICIDPCSCLRRRSR | RSDRRSAGETAEEIDGRDSMVRDPE...RYAKAGDLGDGDLGHTVEPRRDCGA | 332923 | 332566 | - | 6 |
| 4 | MVAETSARRPAETGTETVGVTPRCR | SADCASARPVRPFPSVTGWMVSNCA | 333653 | 333500 | - | 7 |
| 5 | MTSEVLEVQTFANSALWSVEAGAQTDGSIPAM | TTPTVVVATEYVVAGSTICLIC | 338059 | 337894 | - | 33 |
| 6 | LGGEECLSGSERCDELVSIARNSVV...ALFEPPRPSGRLHAIPVIETDHFRA | QTAMPVARSPISGTFILRCTR | 339277 | 339046 | - | 6 |
| 7 | MKKLGPNKHKGWTGRCTSAILR | DTTSDRTEPNGDRTDRYRTVGLVGERTPAARP | 339484 | 339319 | - | 7 |

### Results for WP_104249327.1 [Rathayibacter rathayi] back to top

previous - next

#### Architecture

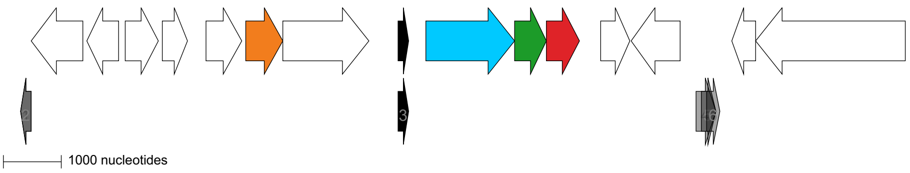

Link to nucleotide sequence

| Accession | start | end | direction | length (aa) | Pfam/HMM | name | description | E-value |
|---|---|---|---|---|---|---|---|---|
| WP_097167734.1 | 14546 | 13667 | - | 292 | PF01694<br>TIGR04239<br>TIGR03902 | Rhomboid<br>TIGR04239<br>TIGR03902 | Rhomboid family<br>rhombo_GlpG: rhomboid family protease GlpG<br>rhom_GG_sort: rhombosortase | 1.40E-30<br>4.70E-14<br>4.50E-11 |
| WP_097167733.1 | 15159 | 14619 | - | 179 | PF00160 | Pro_isomerase | Cyclophilin type peptidyl-prolyl cis-trans isomerase/CLD | 4.70E-42 |
| WP_097167732.1 | 15293 | 15854 | + | 186 | NO PFAM MATCH | - | - | - |
| WP_097167731.1 | 15924 | 16353 | + | 142 | PF00293<br>TIGR02705<br>TIGR00586 | NUDIX<br>TIGR02705<br>TIGR00586 | NUDIX domain<br>nudix_YtkD: nucleoside triphosphatase YtkD<br>mutt: mutator mutT protein | 1.50E-20<br>1.90E-06<br>1.20E-05 |
| WP_104249156.1 | 16680 | 17304 | + | 207 | PF13026 | DUF3887 | Protein of unknown function (DUF3887) | 2.10E-10 |
| WP_097167729.1 | 17372 | 18002 | + | 209 | TIGR01189<br>TIGR02673<br>TIGR02857<br>TIGR01166<br>TIGR01188 | TIGR01189<br>TIGR02673<br>TIGR02857<br>TIGR01166<br>TIGR01188 | ccmA: heme ABC exporter, ATP-binding protein CcmA<br>FtsE: cell division ATP-binding protein FtsE<br>CydD: thiol reductant ABC exporter, CydD subunit<br>cbiO: cobalt ABC transporter, ATP-binding protein<br>drrA: daunorubicin resistance ABC transporter, ATP-binding protein | 4.70E-33<br>1.30E-32<br>6.00E-30<br>3.70E-28<br>5.40E-28 |
| WP_104268301.1 | 17998 | 19495 | + | 498 | NO PFAM MATCH | - | - | - |
| WP_133115533.1 | 20004 | 20169 | + | 54 | LinA | LinA | LinA (linaridin precursor) | 1.70E-10 |
| WP_104249327.1 | 20485 | 22018 | + | 510 | LinH<br>LinE<br>LinG<br>PF12697<br>TIGR02427 | LinH<br>LinE<br>LinG<br>Abhydrolase_6<br>TIGR02427 | LinH (Fusion of LinE/LinG)<br>LinE Homolog<br>LinG Homolog<br>Alpha/beta hydrolase family<br>protocat_pcaD: 3-oxoadipate enol-lactonase | 3.60E-148<br>2.00E-74<br>3.90E-67<br>1.20E-11<br>2.50E-04 |
| WP_097166898.1 | 22004 | 22562 | + | 185 | LinL | LinL | LinL | 2.20E-47 |
| WP_104275120.1 | 22558 | 23137 | + | 192 | TIGR00521<br>PF02441<br>TIGR02113 | TIGR00521<br>Flavoprotein<br>TIGR02113 | coaBC_dfp: phosphopantothenoylcysteine decarboxylase / phosphopantothenate--cysteine ligase<br>Flavoprotein<br>coaC_strep: phosphopantothenoylcysteine decarboxylase | 4.80E-07<br>8.50E-06<br>1.00E-05 |
| WP_104275118.1 | 23496 | 24012 | + | 171 | NO PFAM MATCH | - | - | - |
| WP_104268303.1 | 24867 | 24015 | - | 283 | PF02661 | Fic | Fic/DOC family | 1.10E-14 |
| WP_097166724.1 | 26176 | 25765 | - | 136 | PF12089<br>PF07670 | DUF3566<br>Gate | Transmembrane domain of unknown function (DUF3566)<br>Nucleoside recognition | 3.50E-31<br>1.30E-04 |
| WP_097166725.1 | 28769 | 26168 | - | 866 | TIGR01063<br>TIGR01061<br>TIGR01062<br>PF00521<br>PF03989 | TIGR01063<br>TIGR01061<br>TIGR01062<br>DNA_topoisoIV<br>DNA_gyraseA_C | gyrA: DNA gyrase, A subunit<br>parC_Gpos: DNA topoisomerase IV, A subunit<br>parC_Gneg: DNA topoisomerase IV, A subunit<br>DNA gyrase/topoisomerase IV, subunit A<br>DNA gyrase C-terminal domain, beta-propeller | 0.00E+00<br>1.60E-182<br>1.30E-154<br>1.10E-142<br>3.60E-67 |

| index | leader | core | start | end | dir | score |
|---|---|---|---|---|---|---|
| 1 | VAGSEVLSALSGPLSRCALSASLRCSSPSC | PRRLSTTGEDVIHRVVNTGDNASNVLPGRT | 13650 | 13467 | - | 6 |
| 2 | VVAGSEVLSALSGPLSRCALSASLRCSSPS | CPRRLSTTGEDVIHRVVNTGDNASNVLPGRT | 13653 | 13467 | - | 6 |
| 3 | MTSEVLEVQTFANSALWSVEAGAQTDGSIPAM | TTPTVVVATEYVVAGSTICLIC | 20004 | 20169 | + | 33 |
| 4 | LSSTVQARRADDVKRCCAHSGTGQD...RRSRRSDRRSAGETAEEIDGRDPMV | RAPEHTSSDPAGQRYAKAGDLGDGDLGHTVEPRRDCGA | 25139 | 25496 | + | 6 |
| 5 | VCTSTQLWDSDRHHICIDPCSCLRRRSRRSDRRSAGETAEEIDGRDPMV | RAPEHTSSDPAGQRYAKAGDLGDGDLGHTVEPRRDCGA | 25232 | 25496 | + | 6 |
| 6 | VRRLRKSTGETRWCEPRNTPRRTPQGSDTQRPAIS | ETAISDTLWSLGETVEPRRFELLTSCLQSRRSTN | 25338 | 25548 | + | 6 |

### Results for SOE04649.1 [Rathayibacter rathayi NCPPB 2980 = VKM Ac-1601]

back to top

previous - next

#### Architecture

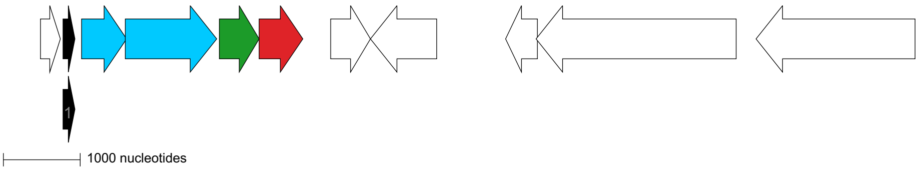

Link to nucleotide sequence

| Accession | start | end | direction | length (aa) | Pfam/HMM | name | description | E-value |
|---|---|---|---|---|---|---|---|---|

| | | | | | | | | |
|---|---|---|---|---|---|---|---|---|
| SOE04646.1 | 0 | 243 | + | 80 | PF13359<br>PF13586 | DDE_Tnp_4<br>DDE_Tnp_1_2 | DDE superfamily endonuclease<br>Transposase DDE domain | 4.70E-05<br>1.50E-04 |
| SOE04647.1 | 279 | 444 | + | 54 | LinA | LinA | LinA (linaridin precursor) | 1.70E-10 |
| SOE04648.1 | 520 | 1111 | + | 196 | LinH<br>LinG | LinH<br>LinG | LinH (Fusion of LinE/LinG)<br>LinG Homolog | 2.60E-57<br>6.50E-47 |
| SOE04649.1 | 1103 | 2294 | + | 396 | LinH<br>LinE<br>LinG<br>PF12697<br>TIGR02427 | LinH<br>LinE<br>LinG<br>Abhydrolase_6<br>TIGR02427 | LinH (Fusion of LinE/LinG)<br>LinE Homolog<br>LinG Homolog<br>Alpha/beta hydrolase family<br>protocat_pcaD: 3-oxoadipate enol-lactonase | 5.30E-98<br>9.60E-75<br>8.70E-26<br>6.50E-12<br>1.80E-04 |
| SOE04650.1 | 2313 | 2838 | + | 174 | LinL | LinL | LinL | 1.00E-45 |
| SOE04651.1 | 2834 | 3413 | + | 192 | TIGR00521<br>TIGR02113<br>PF02441 | TIGR00521<br>TIGR02113<br>Flavoprotein | coaBC_dfp: phosphopantothenoylcysteine decarboxylase / phosphopantothenate--cysteine ligase<br>coaC_strep: phosphopantothenoylcysteine decarboxylase<br>Flavoprotein | 2.20E-07<br>5.80E-06<br>7.00E-06 |
| SOE04652.1 | 3772 | 4288 | + | 171 | NO PFAM MATCH | - | - | - |
| SOE04653.1 | 5143 | 4291 | - | 283 | PF02661 | Fic | Fic/DOC family | 6.20E-15 |
| SOE04654.1 | 6452 | 6041 | - | 136 | PF12089<br>PF07670 | DUF3566<br>Gate | Transmembrane domain of unknown function (DUF3566)<br>Nucleoside recognition | 3.50E-31<br>1.30E-04 |
| SOE04655.1 | 9045 | 6444 | - | 866 | TIGR01063<br>TIGR01061<br>TIGR01062<br>PF00521<br>PF03989 | TIGR01063<br>TIGR01061<br>TIGR01062<br>DNA_topoisoIV<br>DNA_gyraseA_C | gyrA: DNA gyrase, A subunit<br>parC_Gpos: DNA topoisomerase IV, A subunit<br>parC_Gneg: DNA topoisomerase IV, A subunit<br>DNA gyrase/topoisomerase IV, subunit A<br>DNA gyrase C-terminal domain, beta-propeller | 0.00E+00<br>1.60E-182<br>1.30E-154<br>1.10E-142<br>3.60E-67 |
| SOE04656.1 | 11384 | 9314 | - | 689 | TIGR01059<br>TIGR01058<br>TIGR01055<br>PF00204<br>PF00986 | TIGR01059<br>TIGR01058<br>TIGR01055<br>DNA_gyraseB<br>DNA_gyraseB_C | gyrB: DNA gyrase, B subunit<br>parE_Gpos: DNA topoisomerase IV, B subunit<br>parE_Gneg: DNA topoisomerase IV, B subunit<br>DNA gyrase B<br>DNA gyrase B subunit, carboxyl terminus | 2.80E-287<br>2.30E-229<br>9.70E-177<br>1.90E-62<br>6.10E-29 |

| index | leader | core | start | end | dir | score |
|---|---|---|---|---|---|---|
| 1 | MTSEVLEVQTFANSALWSVEAGAQTDGSIPAM | TTPTVVVATEYVVAGSTICLIC | 279 | 444 | + | 32 |

### Results for WP_083093117.1 [Rothia nasimurium] back to top

previous - next

#### Architecture

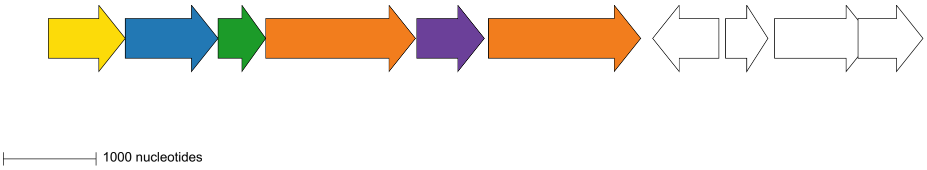

Link to nucleotide sequence

| Accession | start | end | direction | length (aa) | Pfam/HMM | name | description | E-value |
|---|---|---|---|---|---|---|---|---|
| WP_143539221.1 | 225 | 1044 | + | 272 | LinG<br>LinH<br>PF05090 | LinG<br>LinH<br>VKG_Carbox | LinG Homolog<br>LinH (Fusion of LinE/LinG)<br>Vitamin K-dependent gamma-carboxylase | 1.70E-67<br>3.20E-64<br>3.40E-07 |
| WP_083093117.1 | 1050 | 2052 | + | 333 | LinE<br>LinH<br>PF12146<br>PF12697<br>PF00561 | LinE<br>LinH<br>Hydrolase_4<br>Abhydrolase_6<br>Abhydrolase_1 | LinE Homolog<br>LinH (Fusion of LinE/LinG)<br>Serine aminopeptidase, S33<br>Alpha/beta hydrolase family<br>alpha/beta hydrolase fold | 9.20E-39<br>1.10E-31<br>2.30E-11<br>6.90E-10<br>5.90E-08 |
| WP_143539222.1 | 2044 | 2569 | + | 174 | LinL | LinL | LinL | 1.70E-36 |
| WP_083093120.1 | 2570 | 4181 | + | 536 | TIGR03375<br>TIGR02857<br>TIGR02868<br>TIGR02204<br>TIGR02203 | TIGR03375<br>TIGR02857<br>TIGR02868<br>TIGR02204<br>TIGR02203 | type_I_sec_LssB: type I secretion system ATPase<br>CydD: thiol reductant ABC exporter, CydD subunit<br>CydC: thiol reductant ABC exporter, CydC subunit<br>MsbA_rel: ABC transporter, permease/ATP-binding protein<br>MsbA_lipidA: lipid A export permease/ATP-binding protein MsbA | 1.40E-56<br>5.00E-50<br>7.00E-44<br>2.50E-42<br>2.40E-41 |
| WP_083093123.1 | 4199 | 4919 | + | 239 | PF13649<br>PF13847<br>PF08241<br>TIGR01934<br>PF08242 | Methyltransf_25<br>Methyltransf_31<br>Methyltransf_11<br>TIGR01934<br>Methyltransf_12 | Methyltransferase domain<br>Methyltransferase domain<br>Methyltransferase domain<br>MenG_MenH_UbiE: ubiquinone/menaquinone biosynthesis methyltransferase<br>Methyltransferase domain | 4.00E-16<br>3.10E-13<br>3.00E-12<br>8.30E-10<br>3.30E-09 |
| WP_083093126.1 | 4959 | 6609 | + | 549 | TIGR03719<br>TIGR02857<br>PF00005<br>TIGR01189<br>TIGR02868 | TIGR03719<br>TIGR02857<br>ABC_tran<br>TIGR01189<br>TIGR02868 | ABC_ABC_ChvD: ATP-binding cassette protein, ChvD family<br>CydD: thiol reductant ABC exporter, CydD subunit<br>ABC transporter<br>ccmA: heme ABC exporter, ATP-binding protein CcmA<br>CydC: thiol reductant ABC exporter, CydC subunit | 3.40E-56<br>4.00E-41<br>6.00E-37<br>1.40E-35<br>5.60E-35 |
| WP_083093128.1 | 7457 | 6746 | - | 236 | TIGR02325<br>PF07702<br>TIGR02018<br>TIGR03337<br>TIGR02404 | TIGR02325<br>UTRA<br>TIGR02018<br>TIGR03337<br>TIGR02404 | C_P_lyase_phnF: phosphonate metabolism transcriptional regulator PhnF<br>UTRA domain<br>his_ut_repres: histidine utilization repressor<br>phnR: phosphonate utilization transcriptional regulator PhnR<br>trehalos_R_Bsub: trehalose operon repressor | 3.10E-33<br>9.30E-32<br>5.70E-28<br>6.10E-28<br>2.50E-23 |
| WP_083093130.1 | 7522 | 7984 | + | 153 | NO PFAM MATCH | - | - | - |
| WP_083093133.1 | 8054 | 9113 | + | 352 | TIGR00233<br>PF00579<br>TIGR00234 | TIGR00233<br>tRNA-synt_1b<br>TIGR00234 | trpS: tryptophan--tRNA ligase<br>tRNA synthetases class I (W and Y)<br>tyrS: tyrosine--tRNA ligase | 2.30E-105<br>5.70E-81<br>8.20E-12 |

| | | | | | | | | |
|---|---|---|---|---|---|---|---|---|
| WP_143539223.1 | 8952 | 9663 | + | 236 | PF13563 | 2_5_RNA_ligase2 | 2'-5' RNA ligase superfamily | 4.10E-11 |

| index | leader | core | start | end | dir | score |
|---|---|---|---|---|---|---|

### Results for WP_104258703.1 [Rathayibacter tritici] back to top

previous - next

#### Architecture

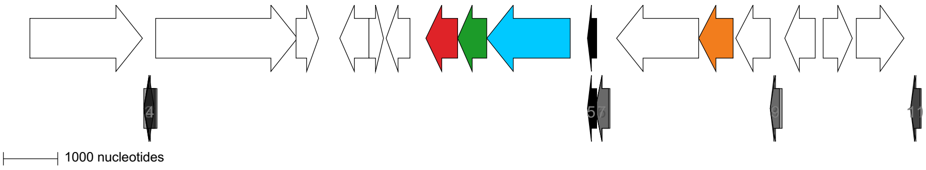

Link to nucleotide sequence

| Accession | start | end | direction | length (aa) | Pfam/HMM | name | description | E-value |
|---|---|---|---|---|---|---|---|---|
| WP_104258667.1 | 38130 | 40200 | + | 689 | TIGR01059<br>TIGR01058<br>TIGR01055<br>PF00204<br>PF00986 | TIGR01059<br>TIGR01058<br>TIGR01055<br>DNA_gyraseB<br>DNA_gyraseB_C | gyrB: DNA gyrase, B subunit<br>parE_Gpos: DNA topoisomerase IV, B subunit<br>parE_Gneg: DNA topoisomerase IV, B subunit<br>DNA gyrase B<br>DNA gyrase B subunit, carboxyl terminus | 3.50E-287<br>6.70E-229<br>2.80E-176<br>4.70E-63<br>6.20E-29 |
| WP_068250044.1 | 40442 | 43043 | + | 866 | TIGR01063<br>TIGR01061<br>TIGR01062<br>PF00521<br>PF03989 | TIGR01063<br>TIGR01061<br>TIGR01062<br>DNA_topoisoIV<br>DNA_gyraseA_C | gyrA: DNA gyrase, A subunit<br>parC_Gpos: DNA topoisomerase IV, A subunit<br>parC_Gneg: DNA topoisomerase IV, A subunit<br>DNA gyrase/topoisomerase IV, subunit A<br>DNA gyrase C-terminal domain, beta-propeller | 0.00E+00<br>1.50E-183<br>1.00E-154<br>1.10E-142<br>3.60E-67 |
| WP_068250046.1 | 43035 | 43446 | + | 136 | PF12089 | DUF3566 | Transmembrane domain of unknown function (DUF3566) | 2.50E-32 |
| WP_104258666.1 | 44355 | 43836 | - | 172 | NO PFAM MATCH | - | - | - |
| WP_017828088.1 | 44374 | 44640 | + | 88 | INFERRED GENE | - | - | - |
| WP_068250056.1 | 45113 | 44684 | - | 142 | PF00293<br>TIGR02705<br>TIGR00586 | NUDIX<br>TIGR02705<br>TIGR00586 | NUDIX domain<br>nudix_YtkD: nucleoside triphosphatase YtkD<br>mutt: mutator mutT protein | 5.20E-21<br>1.30E-07<br>1.50E-05 |
| WP_104258665.1 | 45991 | 45412 | - | 192 | TIGR00521<br>TIGR02113<br>PF02441 | TIGR00521<br>TIGR02113<br>Flavoprotein | coaBC_dfp: phosphopantothenoylcysteine decarboxylase / phosphopantothenate--cysteine ligase<br>coaC_strep: phosphopantothenoylcysteine decarboxylase<br>Flavoprotein | 2.50E-06<br>4.30E-05<br>6.30E-05 |
| WP_104258664.1 | 46545 | 45987 | - | 185 | LinL | LinL | LinL | 2.40E-46 |
| WP_104258703.1 | 48064 | 46531 | - | 510 | LinH<br>LinE<br>LinG<br>PF12697<br>TIGR02427 | LinH<br>LinE<br>LinG<br>Abhydrolase_6<br>TIGR02427 | LinH (Fusion of LinE/LinG)<br>LinE Homolog<br>LinG Homolog<br>Alpha/beta hydrolase family<br>protocat_pcaD: 3-oxoadipate enol-lactonase | 5.60E-150<br>4.60E-74<br>3.10E-68<br>4.20E-11<br>5.90E-04 |
| WP_146076708.1 | 48546 | 48381 | - | 54 | LinA | LinA | LinA (linaridin precursor) | 1.30E-10 |
| WP_104258663.1 | 50426 | 48929 | - | 498 | NO PFAM MATCH | - | - | - |
| WP_104258702.1 | 51052 | 50422 | - | 209 | TIGR02673<br>TIGR01189<br>TIGR02857<br>TIGR01166<br>TIGR03258 | TIGR02673<br>TIGR01189<br>TIGR02857<br>TIGR01166<br>TIGR03258 | FtsE: cell division ATP-binding protein FtsE<br>ccmA: heme ABC exporter, ATP-binding protein CcmA<br>CydD: thiol reductant ABC exporter, CydD subunit<br>cbiO: cobalt ABC transporter, ATP-binding protein<br>PhnT: 2-aminoethylphosphonate ABC transport system, ATP-binding component PhnT | 2.20E-33<br>2.40E-32<br>2.00E-29<br>8.60E-29<br>1.40E-28 |
| WP_104258662.1 | 51744 | 51120 | - | 207 | PF13026 | DUF3887 | Protein of unknown function (DUF3887) | 1.30E-10 |
| WP_068250059.1 | 52581 | 52020 | - | 186 | NO PFAM MATCH | - | - | - |
| WP_068250062.1 | 52715 | 53255 | + | 179 | PF00160 | Pro_isomerase | Cyclophilin type peptidyl-prolyl cis-trans isomerase/CLD | 3.50E-42 |
| WP_068250064.1 | 53328 | 54207 | + | 292 | PF01694<br>TIGR04239<br>TIGR03902 | Rhomboid<br>TIGR04239<br>TIGR03902 | Rhomboid family<br>rhombo_GlpG: rhomboid family protease GlpG<br>rhom_GG_sort: rhombosortase | 1.70E-30<br>3.70E-16<br>8.40E-11 |

| index | leader | core | start | end | dir | score |
|---|---|---|---|---|---|---|
| 1 | LKGGSPAIAPRSERRQPRGTRPRTRPRGEGDGGPQWAARPPPNR | HPPTTDPLTPSATAALAPRGHGPT | 40222 | 40429 | + | 6 |
| 2 | VSPRCEGGGRAGCQGVGCRWVAVRRWSCGPLRAAVTFSTWPGPRPCAAG | LPPFGSRCDRGAAAFQRT | 40417 | 40213 | - | 7 |
| 3 | MVSGLSGGSVSPRCEGGGRAGCQGV...RWSCGPLRAAVTFSTWPGPRPCAAG | LPPFGSRCDRGAAAFQRT | 40444 | 40213 | - | 6 |
| 4 | VSASSAMVSGLSGGSVSPRCEGGGR...RWSCGPLRAAVTFSTWPGPRPCAAG | LPPFGSRCDRGAAAFQRT | 40462 | 40213 | - | 6 |
| 5 | MNSEVLEVQTFANAALSSVEAGARTDGSIPAM | TTPTVVVATEYVVAGSTVCLIC | 48546 | 48381 | - | 33 |
| 6 | LPSLSPNRHVPSVSSGYYRSFRPIL...WHDLVSLAPLFFVAPAGARQPPPRR | GTPHRSDCSGTRERESR | 48770 | 48542 | - | 6 |
| 7 | VTTSLASLPSLSPNRHVPSVSSGYY...WHDLVSLAPLFFVAPAGARQPPPRR | GTPHRSDCSGTRERESR | 48791 | 48542 | - | 6 |
| 8 | VLFHAARSGPARPRDQRTHGRSPAP | PLDAQAGVRAVGAESHTHQDSPPVSR | 51909 | 51753 | - | 6 |
| 9 | VETGHRCACPSTGALRAIHVLFHAARSGPARPRDQ | RTHGRSPAPPLDAQAGVRAVGAESHTHQDSPPVSR | 51966 | 51753 | - | 6 |
| 10 | VENCADCRRRASSRGGRQGSAPVRWRSRT | SLLLPGPPGEDTRGVVPSVNNSVDNFLPSC | 54505 | 54325 | - | 7 |
| 11 | VESPVENCADCRRRASSRGGRQGSAPVRWRS | RTSLLLPGPPGEDTRGVVPSVNNSVDNFLPSC | 54517 | 54325 | - | 7 |

### Results for TFI41935.1 [Rhodococcus sp. 1R11] back to top

previous - next

#### Architecture

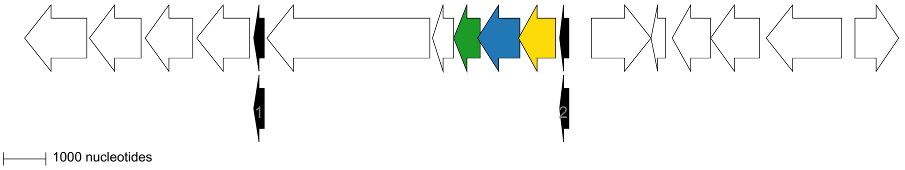

Link to nucleotide sequence

| Accession | start | end | direction | length (aa) | Pfam/HMM | name | description | E-value |
|---|---|---|---|---|---|---|---|---|
| TFI41927.1 | 73304 | 71834 | - | 489 | NO PFAM MATCH | - | - | - |
| TFI41928.1 | 74585 | 73376 | - | 402 | PF17270 | DUF5336 | Family of unknown function (DUF5336) | 6.80E-32 |
| TFI41929.1 | 75803 | 74678 | - | 374 | TIGR04021<br>PF00296<br>TIGR03565<br>TIGR03612<br>TIGR03619 | TIGR04021<br>Bac_luciferase<br>TIGR03565<br>TIGR03612<br>TIGR03619 | LLM_DMSO2_sfnG: dimethyl sulfone monooxygenase SfnG<br>Luciferase-like monooxygenase<br>alk_sulf_monoox: alkanesulfonate monooxygenase, FMNH(2)-dependent<br>RutA: pyrimidine utilization protein A<br>F420_Rv2161c: probable F420-dependent oxidoreductase, Rv2161c family | 3.50E-192<br>1.20E-72<br>3.10E-49<br>2.10E-40<br>4.60E-33 |
| TFI41930.1 | 77135 | 75911 | - | 407 | TIGR04022<br>PF08028<br>PF00441<br>PF02770<br>TIGR03207 | TIGR04022<br>Acyl-CoA_dh_2<br>Acyl-CoA_dh_1<br>Acyl-CoA_dh_M<br>TIGR03207 | sulfur_SfnB: sulfur acquisition oxidoreductase, SfnB family<br>Acyl-CoA dehydrogenase, C-terminal domain<br>Acyl-CoA dehydrogenase, C-terminal domain<br>Acyl-CoA dehydrogenase, middle domain<br>cyc_hxne_CoA_dh: cyclohexanecarboxyl-CoA dehydrogenase | 4.00E-80<br>3.40E-18<br>1.90E-08<br>7.60E-08<br>8.00E-07 |
| TFI41931.1 | 77508 | 77241 | - | 88 | LinA | LinA | LinA (linaridin precursor) | 2.10E-05 |
| TFI41932.1 | 81403 | 77560 | - | 1280 | PF00400<br>PF12894<br>PF16529<br>PF13191<br>TIGR00955 | WD40<br>ANAPC4_WD40<br>Ge1_WD40<br>AAA_16<br>TIGR00955 | WD domain, G-beta repeat<br>Anaphase-promoting complex subunit 4 WD40 domain<br>WD40 region of Ge1, enhancer of mRNA-decapping protein<br>AAA ATPase domain<br>3a01204: pigment precursor permease | 7.30E-58<br>1.00E-24<br>4.80E-16<br>4.90E-05<br>4.60E-04 |
| TFI41933.1 | 81956 | 81467 | - | 162 | NO PFAM MATCH | - | - | - |
| TFI41934.1 | 82578 | 81948 | - | 209 | LinL | LinL | LinL | 3.70E-48 |
| TFI41935.1 | 83506 | 82516 | - | 329 | LinE<br>LinH<br>TIGR02427 | LinE<br>LinH<br>TIGR02427 | LinE Homolog<br>LinH (Fusion of LinE/LinG)<br>protocat_pcaD: 3-oxoadipate enol-lactonase | 9.30E-58<br>9.00E-39<br>3.10E-06 |
| TFI41936.1 | 84347 | 83495 | - | 283 | LinG<br>LinH<br>TIGR04033 | LinG<br>LinH<br>TIGR04033 | LinG Homolog<br>LinH (Fusion of LinE/LinG)<br>export_SdpB: antimicrobial peptide system protein, SdpB family | 2.60E-77<br>4.50E-69<br>1.00E-04 |
| TFI41937.1 | 84658 | 84445 | - | 70 | LinA | LinA | LinA (linaridin precursor) | 7.80E-09 |
| TFI41938.1 | 85189 | 86605 | + | 471 | TIGR02734<br>TIGR03467<br>PF13450<br>TIGR00292<br>PF01593 | TIGR02734<br>TIGR03467<br>NAD_binding_8<br>TIGR00292<br>Amino_oxidase | crtI_fam: phytoene desaturase<br>HpnE: squalene-associated FAD-dependent desaturase<br>NAD(P)-binding Rossmann-like domain<br>TIGR00292: thiazole biosynthesis enzyme<br>Flavin containing amine oxidoreductase | 5.70E-13<br>1.40E-10<br>2.10E-09<br>1.30E-06<br>5.20E-06 |
| TFI41939.1 | 86953 | 86611 | - | 113 | PF09966 | DUF2200 | Uncharacterized protein conserved in bacteria (DUF2200) | 1.70E-50 |
| TFI41940.1 | 87997 | 87085 | - | 303 | TIGR01019<br>PF02629<br>TIGR02717<br>PF00549<br>PF13607 | TIGR01019<br>CoA_binding<br>TIGR02717<br>Ligase_CoA<br>Succ_CoA_lig | sucCoAalpha: succinate-CoA ligase, alpha subunit<br>CoA binding domain<br>AcCoA-syn-alpha: acetyl coenzyme A synthetase (ADP forming), alpha domain<br>CoA-ligase<br>Succinyl-CoA ligase like flavodoxin domain | 2.50E-127<br>4.90E-25<br>7.30E-23<br>1.10E-14<br>1.60E-11 |
| TFI41941.1 | 89169 | 88005 | - | 387 | TIGR01016<br>PF08442<br>PF00549<br>PF13549<br>TIGR02068 | TIGR01016<br>ATP-grasp_2<br>Ligase_CoA<br>ATP-grasp_5<br>TIGR02068 | sucCoAbeta: succinate-CoA ligase, beta subunit<br>ATP-grasp domain<br>CoA-ligase<br>ATP-grasp domain<br>cya_phycin_syn: cyanophycin synthetase | 4.30E-118<br>6.60E-65<br>4.90E-13<br>2.80E-08<br>6.70E-07 |
| TFI42092.1 | 91088 | 89315 | - | 590 | NO PFAM MATCH | - | - | - |
| TFI41942.1 | 91410 | 92415 | + | 334 | PF01551 | Peptidase_M23 | Peptidase family M23 | 3.50E-31 |

| index | leader | core | start | end | dir | score |
|---|---|---|---|---|---|---|
| 1 | MTIRLAEYRVRAAPSTPSDYLRKKE...GRDVARGINELWTEGTHAASAQPVF | TTPATVTTPTVSVCDYMVVATPARQVSVG | 77508 | 77241 | - | 26 |
| 2 | MQTIKIANRDVAAAANQHFTDVVTGVDTKGEIPVL | TTPVTVVTTPEDLALLTATLNATTAVVNATNAHRG | 84658 | 84445 | - | 29 |

### Results for WP_013398290.1 [Rothia dentocariosa] back to top

previous - next

#### Architecture

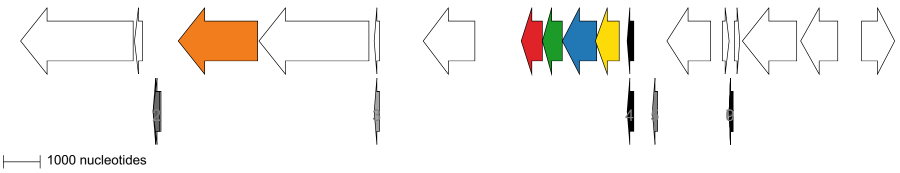

Link to nucleotide sequence

| Accession | start | end | direction | length (aa) | Pfam/HMM | name | description | E-value |
|---|---|---|---|---|---|---|---|---|
| WP_041321664.1 | 311551 | 308509 | - | 1013 | TIGR03897<br>PF13575<br>PF05147 | TIGR03897<br>DUF4135<br>LANC_like | lanti_2_LanM: type 2 lantibiotic biosynthesis protein LanM<br>Domain of unknown function (DUF4135)<br>Lanthionine synthetase C-like protein | 2.20E-122<br>1.50E-52<br>8.50E-33 |
| WP_013398297.1 | 311793 | 311592 | - | 66 | NO PFAM MATCH | - | - | - |
| WP_013398295.1 | 314963 | 312797 | - | 721 | TIGR03796<br>TIGR03797<br>TIGR01846<br>TIGR03375<br>TIGR01193 | TIGR03796<br>TIGR03797<br>TIGR01846<br>TIGR03375<br>TIGR01193 | NHLM_micro_ABC1: NHLM bacteriocin system ABC transporter, peptidase/ATP-binding protein<br>NHLM_micro_ABC2: NHLM bacteriocin system ABC transporter, ATP-binding protein<br>type_I_sec_HlyB: type I secretion system ATPase<br>type_I_sec_LssB: type I secretion system ATPase<br>bacteriocin_ABC: ABC-type bacteriocin transporter | 3.30E-132<br>6.10E-96<br>3.10E-95<br>4.90E-90<br>7.80E-89 |
| WP_041321663.1 | 317980 | 314974 | - | 1001 | TIGR03897<br>PF13575<br>PF05147 | TIGR03897<br>DUF4135<br>LANC_like | lanti_2_LanM: type 2 lantibiotic biosynthesis protein LanM<br>Domain of unknown function (DUF4135)<br>Lanthionine synthetase C-like protein | 3.20E-135<br>9.00E-61<br>3.40E-39 |
| WP_080556864.1 | 318267 | 318111 | - | 51 | PF14867 | Lantibiotic_a | Lantibiotic alpha | 4.20E-11 |
| WP_013398292.1 | 320863 | 319453 | - | 469 | TIGR02734<br>TIGR03467<br>PF13450<br>TIGR02730<br>TIGR00292 | TIGR02734<br>TIGR03467<br>NAD_binding_8<br>TIGR02730<br>TIGR00292 | crtI_fam: phytoene desaturase<br>HpnE: squalene-associated FAD-dependent desaturase<br>NAD(P)-binding Rossmann-like domain<br>carot_isom: carotene isomerase<br>TIGR00292: thiazole biosynthesis enzyme | 5.00E-13<br>2.50E-11<br>1.60E-09<br>4.90E-09<br>1.40E-07 |
| WP_013398291.1 | 322706 | 322133 | - | 190 | TIGR00521<br>PF02441<br>TIGR02113 | TIGR00521<br>Flavoprotein<br>TIGR02113 | coaBC_dfp: phosphopantothenoylcysteine decarboxylase / phosphopantothenate--cysteine ligase<br>Flavoprotein<br>coaC_strep: phosphopantothenoylcysteine decarboxylase | 4.90E-12<br>9.60E-10<br>5.40E-08 |
| WP_141744383.1 | 323245 | 322705 | - | 179 | LinL | LinL | LinL | 7.20E-33 |
| WP_013398290.1 | 324155 | 323234 | - | 306 | LinE<br>LinH<br>TIGR02427 | LinE<br>LinH<br>TIGR02427 | LinE Homolog<br>LinH (Fusion of LinE/LinG)<br>protocat_pcaD: 3-oxoadipate enol-lactonase | 3.70E-62<br>6.10E-35<br>1.30E-05 |
| WP_141744384.1 | 324777 | 324147 | - | 209 | LinG<br>LinH<br>TIGR04033 | LinG<br>LinH<br>TIGR04033 | LinG Homolog<br>LinH (Fusion of LinE/LinG)<br>export_SdpB: antimicrobial peptide system protein, SdpB family | 4.00E-72<br>6.50E-65<br>7.40E-04 |
| WP_141744385.1 | 325170 | 325002 | - | 55 | LinA | LinA | LinA (linaridin precursor) | 5.60E-10 |
| WP_041321662.1 | 327277 | 326080 | - | 398 | PF00067<br>TIGR04458 | p450<br>TIGR04458 | Cytochrome P450<br>CYP450_TxtE: 4-nitrotryptophan synthase | 3.90E-27<br>2.20E-21 |
| WP_013398286.1 | 327607 | 327772 | + | 54 | NO PFAM MATCH | - | - | - |
| WP_013398285.1 | 327902 | 328058 | + | 51 | NO PFAM MATCH | - | - | - |
| WP_013398284.1 | 329605 | 328123 | - | 493 | PF00083<br>TIGR00879<br>PF07690<br>TIGR00895<br>TIGR00898 | Sugar_tr<br>TIGR00879<br>MFS_1<br>TIGR00895<br>TIGR00898 | Sugar (and other) transporter<br>SP: MFS transporter, sugar porter (SP) family<br>Major Facilitator Superfamily<br>2A0115: MFS transporter, aromatic acid:H+ symporter (AAHS) family<br>2A0119: cation transport protein | 8.00E-102<br>2.30E-98<br>3.70E-37<br>3.00E-33<br>1.90E-32 |
| WP_013398283.1 | 330727 | 329725 | - | 333 | PF00248<br>TIGR01293 | Aldo_ket_red<br>TIGR01293 | Aldo/keto reductase family<br>Kv_beta: voltage-dependent potassium channel beta subunit | 2.00E-64<br>3.70E-37 |
| WP_041321661.1 | 331380 | 332292 | + | 303 | TIGR00676<br>PF02219<br>TIGR00677 | TIGR00676<br>MTHFR<br>TIGR00677 | fadh2: methylenetetrahydrofolate reductase [NAD(P)H]<br>Methylenetetrahydrofolate reductase<br>fadh2_euk: methylenetetrahydrofolate reductase | 1.30E-78<br>4.50E-69<br>4.70E-54 |

| index | leader | core | start | end | dir | score |
|---|---|---|---|---|---|---|
| 1 | LTPRLHLETRLSSCYCRVTDFNREKTDSDKDFCS | MTLPHDENRRYISPPLYVAARDESS | 312291 | 312111 | - | 6 |
| 2 | VHDKFLTPRLHLETRLSSCYCRVTDFNREKTDSDKDFCS | MTLPHDENRRYISPPLYVAARDESS | 312306 | 312111 | - | 6 |
| 3 | MPQDITLLEEISEQDLSQVNGGTAF | PGWYSKVIGNRGRVCTVTVECMSVCQ | 318267 | 318111 | - | 6 |
| 4 | MMMNVSDLANSIEIDTTPGAADYAMNSAQPSM | TTPAGVYAVAEMAVAGSTICLVC | 325170 | 325002 | - | 33 |
| 5 | VNKASKPTLLYLHQNHAEKDFGTPQ | FTIIHTLVRLKTAPLKILQPTVSSH | 325827 | 325674 | - | 7 |
| 6 | MENACVSLNISSQSY | LRTAEDSAAATTEPSW | 327876 | 327780 | - | 8 |
| 7 | LMENACVSLNISSQSY | LRTAEDSAAATTEPSW | 327879 | 327780 | - | 8 |
| 8 | MLLMENACVSLNISSQS | YLRTAEDSAAATTEPSW | 327885 | 327780 | - | 8 |
| 9 | VMLLMENACVSLNISSQ | SYLRTAEDSAAATTEPSW | 327888 | 327780 | - | 8 |

### Results for WP_005463322.1 [Saccharomonospora glauca K62] back to top

previous - next

#### Architecture

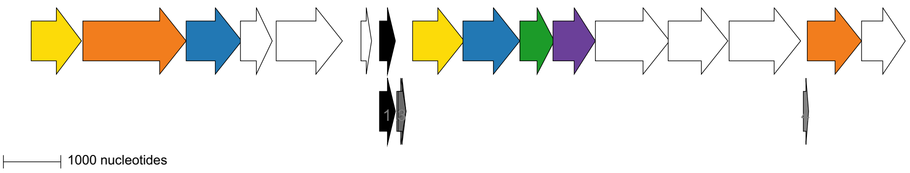

Link to nucleotide sequence

| Accession | start | end | direction | length (aa) | Pfam/HMM | name | description | E-value |
|---|---|---|---|---|---|---|---|---|
| WP_005463305.1 | 1600801 | 1601671 | + | 289 | LinG<br>LinH | LinG<br>LinH | LinG Homolog<br>LinH (Fusion of LinE/LinG) | 2.50E-<br>104<br>1.20E-92 |
| WP_157608908.1 | 1601707 | 1603495 | + | 595 | TIGR02204<br>TIGR02203<br>TIGR00958<br>TIGR03796<br>TIGR03375 | TIGR02204<br>TIGR02203<br>TIGR00958<br>TIGR03796<br>TIGR03375 | MsbA_rel: ABC transporter, permease/ATP-binding protein<br>MsbA_lipidA: lipid A export permease/ATP-binding protein MsbA<br>3a01208: antigen peptide transporter 2<br>NHLM_micro_ABC1: NHLM bacteriocin system ABC transporter, peptidase/ATP-binding protein<br>type_I_sec_LssB: type I secretion system ATPase | 4.30E-<br>108<br>1.40E-<br>103<br>8.80E-89<br>3.60E-83<br>2.20E-82 |
| WP_005463309.1 | 1603498 | 1604464 | + | 321 | LinE<br>LinH<br>TIGR02427<br>PF12697<br>PF00561 | LinE<br>LinH<br>TIGR02427<br>Abhydrolase_6<br>Abhydrolase_1 | LinE Homolog<br>LinH (Fusion of LinE/LinG)<br>protocat_pcaD: 3-oxoadipate enol-lactonase<br>Alpha/beta hydrolase family<br>alpha/beta hydrolase fold | 1.80E-92<br>2.40E-66<br>1.10E-15<br>3.80E-13<br>1.20E-10 |
| WP_019806232.1 | 1604453 | 1604987 | + | 178 | INFERRED GENE | - | - | - |
| WP_005463314.1 | 1605055 | 1606231 | + | 391 | TIGR02032<br>PF04820<br>PF01494<br>TIGR02023<br>PF00890 | TIGR02032<br>Trp_halogenase<br>FAD_binding_3<br>TIGR02023<br>FAD_binding_2 | GG-red-SF: geranylgeranyl reductase family<br>Tryptophan halogenase<br>FAD binding domain<br>BchP-ChlP: geranylgeranyl reductase<br>FAD binding domain | 3.00E-17<br>2.80E-16<br>9.40E-09<br>9.40E-09<br>1.20E-06 |
| WP_005463316.1 | 1606535 | 1606718 | + | 60 | PF09527 | ATPase_gene1 | Putative F0F1-ATPase subunit Ca2+/Mg2+ transporter | 3.00E-04 |
| WP_005463318.1 | 1606875 | 1607136 | + | 86 | LinA | LinA | LinA (linaridin precursor) | 2.90E-15 |
| WP_005463320.1 | 1607444 | 1608335 | + | 296 | LinG<br>LinH<br>LinL<br>TIGR04033<br>PF06762 | LinG<br>LinH<br>LinL<br>TIGR04033<br>LMF1 | LinG Homolog<br>LinH (Fusion of LinE/LinG)<br>LinL<br>export_SdpB: antimicrobial peptide system protein, SdpB family<br>Lipase maturation factor | 7.40E-<br>100<br>3.20E-95<br>1.10E-34<br>5.40E-06<br>1.30E-04 |
| WP_005463322.1 | 1608331 | 1609309 | + | 325 | LinE<br>LinH<br>TIGR02427<br>PF00561<br>PF12697 | LinE<br>LinH<br>TIGR02427<br>Abhydrolase_1<br>Abhydrolase_6 | LinE Homolog<br>LinH (Fusion of LinE/LinG)<br>protocat_pcaD: 3-oxoadipate enol-lactonase<br>alpha/beta hydrolase fold<br>Alpha/beta hydrolase family | 1.60E-98<br>4.60E-77<br>1.80E-13<br>6.70E-11<br>2.00E-10 |
| WP_005463324.1 | 1609317 | 1609905 | + | 195 | LinL<br>LinG | LinL<br>LinG | LinL<br>LinG Homolog | 2.00E-68<br>1.20E-15 |
| WP_005463326.1 | 1609889 | 1610630 | + | 246 | PF13649<br>PF13489<br>PF08242<br>PF08241<br>TIGR02021 | Methyltransf_25<br>Methyltransf_23<br>Methyltransf_12<br>Methyltransf_11<br>TIGR02021 | Methyltransferase domain<br>Methyltransferase domain<br>Methyltransferase domain<br>Methyltransferase domain<br>BchM-ChlM: magnesium protoporphyrin O-methyltransferase | 2.00E-13<br>6.50E-11<br>1.60E-10<br>1.80E-10<br>4.90E-09 |
| WP_005463328.1 | 1610639 | 1611911 | + | 423 | TIGR01426<br>PF03033 | TIGR01426<br>Glyco_transf_28 | MGT: glycosyltransferase, MGT family<br>Glycosyltransferase family 28 N-terminal domain | 5.10E-17<br>9.00E-14 |
| WP_005463330.1 | 1611889 | 1612942 | + | 350 | NO PFAM MATCH | - | - | - |
| WP_040919718.1 | 1612956 | 1614210 | + | 417 | PF06722<br>TIGR01426<br>PF00201 | DUF1205<br>TIGR01426<br>UDPGT | Protein of unknown function (DUF1205)<br>MGT: glycosyltransferase, MGT family<br>UDP-glucoronosyl and UDP-glucosyl transferase | 5.10E-27<br>1.30E-21<br>2.70E-05 |
| WP_005463335.1 | 1614314 | 1615277 | + | 320 | TIGR01188<br>TIGR03864<br>TIGR01288<br>TIGR03522<br>TIGR04406 | TIGR01188<br>TIGR03864<br>TIGR01288<br>TIGR03522<br>TIGR04406 | drrA: daunorubicin resistance ABC transporter, ATP-binding protein<br>PQQ_ABC_ATP: ABC transporter, ATP-binding subunit, PQQ-dependent alcohol dehydrogenase system<br>nodI: nodulation ABC transporter NodI<br>GldA_ABC_ATP: gliding motility-associated ABC transporter ATP-binding subunit GldA<br>LPS_export_lptB: LPS export ABC transporter ATP-binding protein | 1.60E-62<br>2.50E-50<br>6.20E-46<br>2.60E-37<br>1.30E-35 |
| WP_005463337.1 | 1615273 | 1616032 | + | 252 | PF01061 | ABC2_membrane | ABC-2 type transporter | 1.10E-07 |

| index | leader | core | start | end | dir | score |
|---|---|---|---|---|---|---|
| 1 | MNNIEKAAQAALPEQGGVLRPLERANGAAPVL | GTPALVTATGVFFVAGNMVLDLVGH...KHDGAVPTGKSGAELLSMRANGLRG | 1606875 | 1607136 | + | 26 |
| 2 | MTPESTRSPVGSGHCRAPRAAATRCAAALFHVHIGL | TAPELLPNDQQHDGSALRGNAVG | 1607158 | 1607338 | + | 6 |
| 3 | VGSGHCRAPRAAATRCAAALFHVHIGL | TAPELLPNDQQHDGSALRGNAVG | 1607185 | 1607338 | + | 6 |
| 4 | MTNTHDSLTVVPTSG | PVLHSTVDLCTVLWST | 1614243 | 1614339 | + | 7 |

### Results for WP_157601541.1 [Saccharomonospora halophila 8] back to top

previous - next

#### Architecture

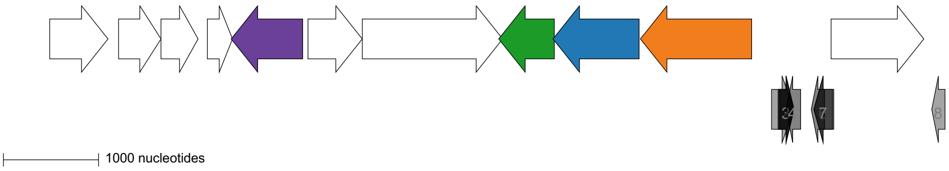

Link to nucleotide sequence

| Accession | start | end | direction | length (aa) | Pfam/HMM | name | description | E-value |
|---|---|---|---|---|---|---|---|---|
| WP_019806225.1 | 837 | 1440 | + | 200 | PF00072<br>TIGR02875<br>PF00196<br>TIGR03020<br>TIGR02154 | Response_reg<br>TIGR02875<br>GerE<br>TIGR03020<br>TIGR02154 | Response regulator receiver domain<br>spore_0_A: sporulation transcription factor Spo0A<br>Bacterial regulatory proteins, luxR family<br>EpsA: transcriptional regulator EpsA<br>PhoB: phosphate regulon transcriptional regulatory protein PhoB | 3.50E-23<br>2.00E-14<br>3.00E-12<br>2.60E-09<br>1.40E-08 |
| WP_019806226.1 | 1551 | 2001 | + | 149 | PF05523 | FdtA | WxcM-like, C-terminal | 6.90E-47 |
| WP_019806227.1 | 1997 | 2389 | + | 131 | PF01041<br>TIGR04181<br>TIGR04427<br>TIGR03588<br>TIGR02379 | DegT_DnrJ_EryC1<br>TIGR04181<br>TIGR04427<br>TIGR03588<br>TIGR02379 | DegT/DnrJ/EryC1/StrS aminotransferase family<br>NHT_00031: aminotransferase, LLPSF_NHT_00031 family<br>PLP_DesI: dTDP-4-dehydro-6-deoxyglucose aminotransferase<br>PseC: UDP-4-amino-4,6-dideoxy-N-acetyl-beta-L-altrosamine transaminase<br>ECA_wecE: TDP-4-keto-6-deoxy-D-glucose transaminase | 1.40E-43<br>1.90E-22<br>1.20E-18<br>3.20E-18<br>4.00E-14 |
| SACHA_RS0100155 | 2489 | 2744 | + | 85 | INFERRED GENE | - | - | - |
| WP_019806228.1 | 3490 | 2740 | - | 249 | PF13649<br>PF08241<br>PF13489<br>PF08242<br>TIGR02021 | Methyltransf_25<br>Methyltransf_11<br>Methyltransf_23<br>Methyltransf_12<br>TIGR02021 | Methyltransferase domain<br>Methyltransferase domain<br>Methyltransferase domain<br>Methyltransferase domain<br>BchM-ChlM: magnesium protoporphyrin O-methyltransferase | 2.80E-14<br>5.40E-13<br>9.40E-12<br>1.50E-10<br>5.20E-10 |
| WP_018835202.1 | 3549 | 4113 | + | 188 | INFERRED GENE | - | - | - |
| SACHA_RS24220 | 4112 | 5594 | + | 494 | INFERRED GENE | - | - | - |
| WP_039737341.1 | 6127 | 5551 | - | 191 | LinL | LinL | LinL | 4.90E-62 |
| WP_157601541.1 | 7025 | 6113 | - | 303 | LinE<br>LinH<br>TIGR02427<br>PF12697<br>PF00561 | LinE<br>LinH<br>TIGR02427<br>Abhydrolase_6<br>Abhydrolase_1 | LinE Homolog<br>LinH (Fusion of LinE/LinG)<br>protocat_pcaD: 3-oxoadipate enol-lactonase<br>Alpha/beta hydrolase family<br>alpha/beta hydrolase fold | 2.50E-90<br>3.00E-66<br>9.90E-14<br>1.00E-13<br>2.50E-11 |
| WP_019806234.1 | 8205 | 7030 | - | 390 | TIGR02204<br>TIGR02203<br>TIGR00958<br>TIGR03375<br>TIGR03796 | TIGR02204<br>TIGR02203<br>TIGR00958<br>TIGR03375<br>TIGR03796 | MsbA_rel: ABC transporter, permease/ATP-binding protein<br>MsbA_lipidA: lipid A export permease/ATP-binding protein MsbA<br>3a01208: antigen peptide transporter 2<br>type_I_sec_LssB: type I secretion system ATPase<br>NHLM_micro_ABC1: NHLM bacteriocin system ABC transporter, peptidase/ATP-binding protein | 1.50E-82<br>3.80E-78<br>9.30E-70<br>9.70E-68<br>1.40E-66 |
| WP_157601540.1 | 9044 | 10019 | + | 324 | TIGR03971<br>PF00106<br>TIGR01963<br>TIGR01830<br>TIGR02632 | TIGR03971<br>adh_short<br>TIGR01963<br>TIGR01830<br>TIGR02632 | SDR_subfam_1: SDR family mycofactocin-dependent oxidoreductase<br>short chain dehydrogenase<br>PHB_DH: 3-hydroxybutyrate dehydrogenase<br>3oxo_ACP_reduc: 3-oxoacyl-[acyl-carrier-protein] reductase<br>RhaD_aldol-ADH: rhamnulose-1-phosphate aldolase/alcohol dehydrogenase | 1.10E-14<br>1.60E-12<br>4.30E-12<br>3.50E-08<br>5.10E-08 |

| index | leader | core | start | end | dir | score |
|---|---|---|---|---|---|---|
| 1 | VSANAGRAQRPRSTPGSSSCPVLRS...VDVADGGGVPASPSTSHSAPARPSC | STAKYTGPPPAASSAR | 8418 | 8655 | + | 6 |
| 2 | VLRSSEVPSGRSSTVQVDVADGGGVPASPSTSHSAPARPSC | STAKYTGPPPAASSAR | 8481 | 8655 | + | 8 |
| 3 | VPSGRSSTVQVDVADGGGVPASPSTSHSAPARPSC | STAKYTGPPPAASSAR | 8499 | 8655 | + | 8 |
| 4 | VAVRCGPRGRFPPPGRGACVLPRRR | RRRGRARVLRGGARRAGGRTVARRR | 8715 | 8562 | - | 7 |
| 5 | VHGLRSVSGQCHRGASTRGHGTS | TRGHGVATRGDSVELLRSGWVAAA | 8975 | 8831 | - | 7 |
| 6 | MKPPFPETLPPH | GTVSRAVSVGFPCMVSAPFRVSVTVAPRLAGMARRLAGMGWRLAGIPLNC | 9046 | 8857 | - | 7 |
| 7 | VSVSFAAMKPPFPETLPPH | GTVSRAVSVGFPCMVSAPFRVSVTVAPRLAGMARRLAGMGWRLAGIPLNC | 9067 | 8857 | - | 7 |
| 8 | VVVALEQPEIAAGQLLCLGTEVE | ILGPDEVRAAFADTARRMADRHG | 10241 | 10100 | - | 6 |

### Results for WP_005463309.1 [Saccharomonospora glauca K62] back to top

previous - next

#### Architecture

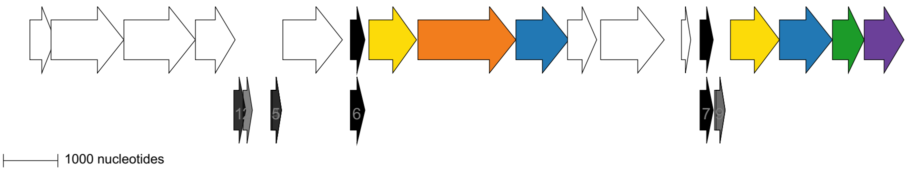

1000 nucleotides

Link to nucleotide sequence

| Accession | start | end | direction | length (aa) | Pfam/HMM | name | description | E-value |
|---|---|---|---|---|---|---|---|---|
| WP_157608779.1 | 1594575 | 1595025 | + | 149 | NO PFAM MATCH | - | - | - |
| WP_083852938.1 | 1594966 | 1596310 | + | 447 | PF05193<br>PF00675<br>TIGR02110 | Peptidase_M16_C<br>Peptidase_M16<br>TIGR02110 | Peptidase M16 inactive domain<br>Insulinase (Peptidase family M16)<br>PQQ_syn_pqqF: coenzyme PQQ biosynthesis protein PqqF | 1.70E-19<br>2.70E-19<br>3.60E-15 |
| WP_005463296.1 | 1596306 | 1597611 | + | 434 | PF05193 | Peptidase_M16_C | Peptidase M16 inactive domain | 2.00E-20 |
| SACGLDRAFT_RS22280 | 1597623 | 1598346 | + | 241 | INFERRED GENE | - | - | - |
| WP_005463301.1 | 1599224 | 1600325 | + | 366 | PF08028<br>TIGR04022<br>PF00441 | Acyl-CoA_dh_2<br>TIGR04022<br>Acyl-CoA_dh_1 | Acyl-CoA dehydrogenase, C-terminal domain<br>sulfur_SfnB: sulfur acquisition oxidoreductase, SfnB family<br>Acyl-CoA dehydrogenase, C-terminal domain | 8.30E-24<br>2.10E-11<br>7.50E-09 |

| | | | | | | | | |
|---|---|---|---|---|---|---|---|---|
| WP_157608781.1 | 1600465 | 1600717 | + | 83 | LinA | LinA | LinA (linaridin precursor) | 9.10E-14 |
| WP_005463305.1 | 1600801 | 1601671 | + | 289 | LinG<br>LinH | LinG<br>LinH | LinG Homolog<br>LinH (Fusion of LinE/LinG) | 2.50E-<br>104<br>1.20E-92 |
| WP_157608908.1 | 1601707 | 1603495 | + | 595 | TIGR02204<br>TIGR02203<br>TIGR00958<br>TIGR03796<br>TIGR03375 | TIGR02204<br>TIGR02203<br>TIGR00958<br>TIGR03796<br>TIGR03375 | MsbA_rel: ABC transporter, permease/ATP-binding protein<br>MsbA_lipidA: lipid A export permease/ATP-binding protein MsbA<br>3a01208: antigen peptide transporter 2<br>NHLM_micro_ABC1: NHLM bacteriocin system ABC transporter, peptidase/ATP-binding protein<br>type_I_sec_LssB: type I secretion system ATPase | 4.30E-<br>108<br>1.40E-<br>103<br>8.80E-89<br>3.60E-83<br>2.20E-82 |
| WP_005463309.1 | 1603498 | 1604464 | + | 321 | LinE<br>LinH<br>TIGR02427<br>PF12697<br>PF00561 | LinE<br>LinH<br>TIGR02427<br>Abhydrolase_6<br>Abhydrolase_1 | LinE Homolog<br>LinH (Fusion of LinE/LinG)<br>protocat_pcaD: 3-oxoadipate enol-lactonase<br>Alpha/beta hydrolase family<br>alpha/beta hydrolase fold | 1.80E-92<br>2.40E-66<br>1.10E-15<br>3.80E-13<br>1.20E-10 |
| WP_019806232.1 | 1604453 | 1604987 | + | 178 | INFERRED GENE | - | - | - |
| WP_005463314.1 | 1605055 | 1606231 | + | 391 | TIGR02032<br>PF04820<br>PF01494<br>TIGR02023<br>PF00890 | TIGR02032<br>Trp_halogenase<br>FAD_binding_3<br>TIGR02023<br>FAD_binding_2 | GG-red-SF: geranylgeranyl reductase family<br>Tryptophan halogenase<br>FAD binding domain<br>BchP-ChlP: geranylgeranyl reductase<br>FAD binding domain | 3.00E-17<br>2.80E-16<br>9.40E-09<br>9.40E-09<br>1.20E-06 |
| WP_005463316.1 | 1606535 | 1606718 | + | 60 | PF09527 | ATPase_gene1 | Putative F0F1-ATPase subunit Ca2+/Mg2+ transporter | 3.00E-04 |
| WP_005463318.1 | 1606875 | 1607136 | + | 86 | LinA | LinA | LinA (linaridin precursor) | 2.90E-15 |
| WP_005463320.1 | 1607444 | 1608335 | + | 296 | LinG<br>LinH<br>LinL<br>TIGR04033<br>PF06762 | LinG<br>LinH<br>LinL<br>TIGR04033<br>LMF1 | LinG Homolog<br>LinH (Fusion of LinE/LinG)<br>LinL<br>export_SdpB: antimicrobial peptide system protein, SdpB family<br>Lipase maturation factor | 7.40E-<br>100<br>3.20E-95<br>1.10E-34<br>5.40E-06<br>1.30E-04 |
| WP_005463322.1 | 1608331 | 1609309 | + | 325 | LinE<br>LinH<br>TIGR02427<br>PF00561<br>PF12697 | LinE<br>LinH<br>TIGR02427<br>Abhydrolase_1<br>Abhydrolase_6 | LinE Homolog<br>LinH (Fusion of LinE/LinG)<br>protocat_pcaD: 3-oxoadipate enol-lactonase<br>alpha/beta hydrolase fold<br>Alpha/beta hydrolase family | 1.60E-98<br>4.60E-77<br>1.80E-13<br>6.70E-11<br>2.00E-10 |
| WP_005463324.1 | 1609317 | 1609905 | + | 195 | LinL<br>LinG | LinL<br>LinG | LinL<br>LinG Homolog | 2.00E-68<br>1.20E-15 |
| WP_005463326.1 | 1609889 | 1610630 | + | 246 | PF13649<br>PF13489<br>PF08242<br>PF08241<br>TIGR02021 | Methyltransf_25<br>Methyltransf_23<br>Methyltransf_12<br>Methyltransf_11<br>TIGR02021 | Methyltransferase domain<br>Methyltransferase domain<br>Methyltransferase domain<br>Methyltransferase domain<br>BchM-ChlM: magnesium protoporphyrin O-methyltransferase | 2.00E-13<br>6.50E-11<br>1.60E-10<br>1.80E-10<br>4.90E-09 |

| index | leader | core | start | end | dir | score |
|---|---|---|---|---|---|---|
| 1 | VASSATRTPTRRGCRPTRSRAWWGRTAPSSWRDRRRCWARPNRPAT | AATASTGGGGSTADACTPTGSWPG | 1598329 | 1598542 | + | 9 |
| 2 | VAVQQRMPARRLDHGRGDGARRGRDT | GAPAGVRPVATSHRARHLARRRTGGHG | 1598489 | 1598651 | + | 7 |
| 3 | LALVLRHAVDTAKHVVESVSAIAEHGGDAGAARTL | RVLLRDVYGAGAHFACSDDVYLRNATRFAGRAGDP | 1598994 | 1599207 | + | 7 |
| 4 | VLRHAVDTAKHVVESVSAIAEHGGDAGAARTLR | VLLRDVYGAGAHFACSDDVYLRNATRFAGRAGDP | 1599003 | 1599207 | + | 7 |
| 5 | VDTAKHVVESVSAIAEHGGDAGAARTLRVLL | RDVYGAGAHFACSDDVYLRNATRFAGRAGDP | 1599018 | 1599207 | + | 6 |
| 6 | MSIVEKFANSHVDETLRPLQVGSVAGPVL | ATPAYAGYAAGAFTLGLITDLIENH...VTPTGPVPESGSADELLGMRVESIR | 1600465 | 1600717 | + | 27 |
| 7 | MNNIEKAAQAALPEQGGVLRPLERANGAAPVL | GTPALVTATGVFFVAGNMVLDLVGH...KHDGAVPTGKSGAELLSMRANGLRG | 1606875 | 1607136 | + | 26 |
| 8 | MTPESTRSPVGSGHCRAPRAAATRCAAALFHVHIGL | TAPELLPNDQQHDGSALRGNAVG | 1607158 | 1607338 | + | 6 |
| 9 | VGSGHCRAPRAAATRCAAALFHVHIGL | TAPELLPNDQQHDGSALRGNAVG | 1607185 | 1607338 | + | 6 |

### Results for WP_138383130.1 [Saccharomonospora piscinae] back to top

previous - next

#### Architecture

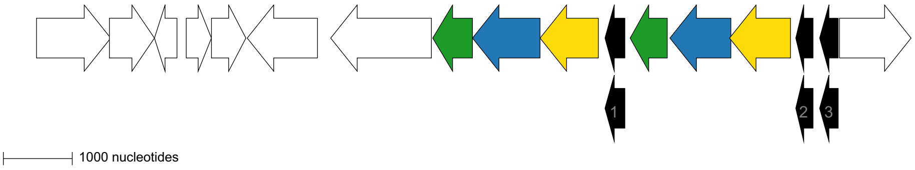

Link to nucleotide sequence

| Accession | start | end | direction | length (aa) | Pfam/HMM | name | description | E-value |
|---|---|---|---|---|---|---|---|---|
| WP_081191136.1 | 995228 | 996302 | + | 357 | NO PFAM MATCH | - | - | - |
| WP_081191135.1 | 996268 | 996940 | + | 223 | PF00072<br>PF00196 | Response_reg<br>GerE | Response regulator receiver domain<br>Bacterial regulatory proteins, luxR family | 1.40E-07<br>5.00E-07 |
| WP_081191134.1 | 997265 | 996941 | - | 107 | TIGR03954<br>PF12823 | TIGR03954<br>DUF3817 | integ_memb_HG: integral membrane protein<br>Domain of unknown function (DUF3817) | 7.90E-27<br>2.50E-25 |

| | | | | | | | | |
|---|---|---|---|---|---|---|---|---|
| WP_081191133.1 | 997396 | 997759 | + | 120 | PF12840<br>PF01022<br>PF12802<br>PF13412<br>PF08279 | HTH_20<br>HTH_5<br>MarR_2<br>HTH_24<br>HTH_11 | Helix-turn-helix domain<br>Bacterial regulatory protein, arsR family<br>MarR family<br>Winged helix-turn-helix DNA-binding<br>HTH domain | 2.50E-15<br>4.10E-14<br>2.50E-08<br>3.50E-06<br>5.90E-06 |
| WP_081191132.1 | 997755 | 998262 | + | 168 | PF08327 | AHSA1 | Activator of Hsp90 ATPase homolog 1-like protein | 4.60E-12 |
| WP_138383236.1 | 999291 | 998283 | - | 335 | TIGR03451<br>TIGR00692<br>TIGR03989<br>TIGR02818<br>TIGR02819 | TIGR03451<br>TIGR00692<br>TIGR03989<br>TIGR02818<br>TIGR02819 | mycoS_dep_FDH: S-(hydroxymethyl)mycothiol dehydrogenase<br>tdh: L-threonine 3-dehydrogenase<br>Rxyl_3153: NDMA-dependent alcohol dehydrogenase, Rxyl_3153 family<br>adh_III_F_hyde: S-(hydroxymethyl)glutathione dehydrogenase/class III alcohol dehydrogenase<br>fdhA_non_GSH: formaldehyde dehydrogenase, glutathione-independent | 4.90E-39<br>1.20E-38<br>6.00E-31<br>1.10E-28<br>1.50E-28 |
| WP_138383237.1 | 1000956 | 999489 | - | 488 | PF00171<br>TIGR02299<br>TIGR01804<br>TIGR03216<br>TIGR01780 | Aldedh<br>TIGR02299<br>TIGR01804<br>TIGR03216<br>TIGR01780 | Aldehyde dehydrogenase family<br>HpaE: 5-carboxymethyl-2-hydroxymuconate semialdehyde dehydrogenase<br>BADH: betaine-aldehyde dehydrogenase<br>OH_muco_semi_DH: 2-hydroxymuconic semialdehyde dehydrogenase<br>SSADH: succinate-semialdehyde dehydrogenase | 1.50E-151<br>3.90E-122<br>1.50E-121<br>5.90E-106<br>1.40E-100 |
| WP_081191131.1 | 1001548 | 1000975 | - | 190 | LinL | LinL | LinL | 2.60E-68 |
| WP_138383130.1 | 1002528 | 1001547 | - | 326 | LinE<br>LinH<br>PF12697<br>TIGR02427<br>TIGR03695 | LinE<br>LinH<br>Abhydrolase_6<br>TIGR02427<br>TIGR03695 | LinE Homolog<br>LinH (Fusion of LinE/LinG)<br>Alpha/beta hydrolase family<br>protocat_pcaD: 3-oxoadipate enol-lactonase<br>menH_SHCHC: 2-succinyl-6-hydroxy-2,4-cyclohexadiene-1-carboxylate synthase | 2.30E-86<br>4.80E-67<br>6.80E-15<br>9.30E-10<br>1.10E-09 |
| WP_138383238.1 | 1003379 | 1002524 | - | 284 | LinG<br>LinH<br>PF06762<br>TIGR04033 | LinG<br>LinH<br>LMF1<br>TIGR04033 | LinG Homolog<br>LinH (Fusion of LinE/LinG)<br>Lipase maturation factor<br>export_SdpB: antimicrobial peptide system protein, SdpB family | 9.30E-102<br>1.50E-95<br>1.00E-06<br>4.00E-04 |
| WP_138383131.1 | 1003760 | 1003481 | - | 92 | LinA | LinA | LinA (linaridin precursor) | 5.10E-10 |
| WP_081192168.1 | 1004377 | 1003843 | - | 177 | LinL | LinL | LinL | 8.50E-59 |
| WP_138383132.1 | 1005296 | 1004411 | - | 294 | LinE<br>LinH<br>PF12697<br>TIGR02427<br>TIGR03611 | LinE<br>LinH<br>Abhydrolase_6<br>TIGR02427<br>TIGR03611 | LinE Homolog<br>LinH (Fusion of LinE/LinG)<br>Alpha/beta hydrolase family<br>protocat_pcaD: 3-oxoadipate enol-lactonase<br>RutD: pyrimidine utilization protein D | 2.00E-58<br>2.00E-43<br>7.20E-11<br>3.10E-10<br>3.10E-09 |
| WP_138383133.1 | 1006165 | 1005292 | - | 290 | LinG<br>LinH | LinG<br>LinH | LinG Homolog<br>LinH (Fusion of LinE/LinG) | 1.10E-99<br>1.20E-87 |
| WP_138383134.1 | 1006495 | 1006237 | - | 85 | LinA | LinA | LinA (linaridin precursor) | 6.30E-09 |
| WP_081191125.1 | 1006861 | 1006597 | - | 87 | LinA | LinA | LinA (linaridin precursor) | 3.80E-08 |
| WP_138383239.1 | 1006880 | 1007930 | + | 349 | PF13561<br>TIGR01830<br>TIGR02415<br>PF00106<br>TIGR01829 | adh_short_C2<br>TIGR01830<br>TIGR02415<br>adh_short<br>TIGR01829 | Enoyl-(Acyl carrier protein) reductase<br>3oxo_ACP_reduc: 3-oxoacyl-[acyl-carrier-protein] reductase<br>23BDH: acetoin reductases<br>short chain dehydrogenase<br>AcAcCoA_reduct: acetoacetyl-CoA reductase | 6.30E-62<br>1.50E-55<br>8.30E-51<br>1.60E-49<br>2.40E-48 |

| index | leader | core | start | end | dir | score |
|---|---|---|---|---|---|---|
| 1 | MNKVEQAAQAAQAGHEGQLLPLRRDEPAAPALSAAWVAATAGVTLA | ATAVFQAGNAVTDYVGNHTNPEINPEALSGKSGTELLSIRQNGVRR | 1003760 | 1003481 | - | 13 |
| 2 | MSAAAAFANSKAGQGLAPLEANQVYAPVL | CTATTLTTTTATCTVFATGSIGDAG...DVSSAGLPAHGSAEELLNFRAGSIG | 1006495 | 1006237 | - | 16 |
| 3 | MSVIAAFANSKADHGLAPLEASPVYTPVF | CTTSTTTATLAGTVGMVFTAGIISD...DVSSAGLPAHGSAQDLLNFRAGSIG | 1006861 | 1006597 | - | 14 |

### Results for WP_037336134.1 [Saccharomonospora sp. CNQ490] back to top

previous - next

#### Architecture

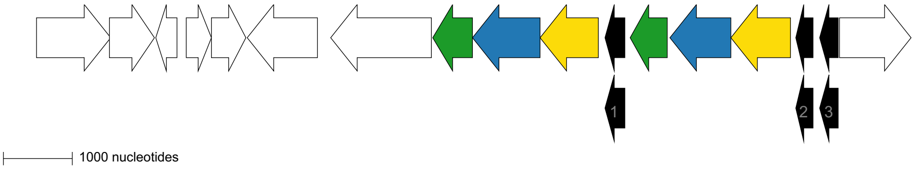

Link to nucleotide sequence

| Accession | start | end | direction | length (aa) | Pfam/HMM | name | description | E-value |
|---|---|---|---|---|---|---|---|---|
| WP_024877733.1 | 74290 | 75364 | + | 357 | NO PFAM MATCH | - | - | - |
| WP_037336131.1 | 75330 | 76002 | + | 223 | PF00072<br>PF00196 | Response_reg<br>GerE | Response regulator receiver domain<br>Bacterial regulatory proteins, luxR family | 3.70E-07<br>5.50E-07 |
| WP_024877735.1 | 76327 | 76003 | - | 107 | TIGR03954<br>PF12823 | TIGR03954<br>DUF3817 | integ_memb_HG: integral membrane protein<br>Domain of unknown function (DUF3817) | 2.30E-27<br>9.20E-26 |
| WP_024877736.1 | 76458 | 76821 | + | 120 | PF12840<br>PF01022<br>PF12802<br>PF13412<br>PF08279 | HTH_20<br>HTH_5<br>MarR_2<br>HTH_24<br>HTH_11 | Helix-turn-helix domain<br>Bacterial regulatory protein, arsR family<br>MarR family<br>Winged helix-turn-helix DNA-binding<br>HTH domain | 2.50E-15<br>4.10E-14<br>2.40E-08<br>3.50E-06<br>6.00E-06 |
| WP_024877737.1 | 76817 | 77324 | + | 168 | PF08327 | AHSA1 | Activator of Hsp90 ATPase homolog 1-like protein | 5.00E-12 |

| | | | | | | | | |
|---|---|---|---|---|---|---|---|---|
| WP_037336182.1 | 78353 | 77345 | - | 335 | TIGR03451<br>TIGR00692<br>TIGR03989<br>TIGR02819<br>TIGR03366 | TIGR03451<br>TIGR00692<br>TIGR03989<br>TIGR02819<br>TIGR03366 | mycoS_dep_FDH: S-(hydroxymethyl)mycothiol dehydrogenase<br>tdh: L-threonine 3-dehydrogenase<br>Rxyl_3153: NDMA-dependent alcohol dehydrogenase, Rxyl_3153 family<br>fdhA_non_GSH: formaldehyde dehydrogenase, glutathione-independent<br>HpnZ_proposed: putative phosphonate catabolism associated alcohol dehydrogenase | 5.20E-38<br>9.30E-38<br>6.20E-31<br>1.30E-29<br>3.90E-28 |
| WP_051473488.1 | 80018 | 78551 | - | 488 | PF00171<br>TIGR02299<br>TIGR01804<br>TIGR03216<br>TIGR01780 | Aldedh<br>TIGR02299<br>TIGR01804<br>TIGR03216<br>TIGR01780 | Aldehyde dehydrogenase family<br>HpaE: 5-carboxymethyl-2-hydroxymuconate semialdehyde dehydrogenase<br>BADH: betaine-aldehyde dehydrogenase<br>OH_muco_semi_DH: 2-hydroxymuconic semialdehyde dehydrogenase<br>SSADH: succinate-semialdehyde dehydrogenase | 5.80E-151<br>1.30E-122<br>2.50E-121<br>2.40E-104<br>4.50E-100 |
| WP_024877740.1 | 80610 | 80037 | - | 190 | LinL | LinL | LinL | 2.00E-66 |
| WP_037336134.1 | 81590 | 80609 | - | 326 | LinE<br>LinH<br>PF12697<br>TIGR02427<br>PF12146 | LinE<br>LinH<br>Abhydrolase_6<br>TIGR02427<br>Hydrolase_4 | LinE Homolog<br>LinH (Fusion of LinE/LinG)<br>Alpha/beta hydrolase family<br>protocat_pcaD: 3-oxoadipate enol-lactonase<br>Serine aminopeptidase, S33 | 1.90E-89<br>2.40E-69<br>9.80E-17<br>4.70E-12<br>3.10E-10 |
| WP_037336136.1 | 82441 | 81586 | - | 284 | LinG<br>LinH<br>PF06762<br>TIGR04033 | LinG<br>LinH<br>LMF1<br>TIGR04033 | LinG Homolog<br>LinH (Fusion of LinE/LinG)<br>Lipase maturation factor<br>export_SdpB: antimicrobial peptide system protein, SdpB family | 1.60E-101<br>1.20E-94<br>4.80E-07<br>1.00E-04 |
| WP_024877741.1 | 82820 | 82541 | - | 92 | LinA | LinA | LinA (linaridin precursor) | 7.10E-09 |
| WP_051473489.1 | 83437 | 82903 | - | 177 | LinL | LinL | LinL | 2.40E-58 |
| WP_155984838.1 | 84356 | 83471 | - | 294 | LinE<br>LinH<br>PF12697<br>TIGR02427<br>TIGR03611 | LinE<br>LinH<br>Abhydrolase_6<br>TIGR02427<br>TIGR03611 | LinE Homolog<br>LinH (Fusion of LinE/LinG)<br>Alpha/beta hydrolase family<br>protocat_pcaD: 3-oxoadipate enol-lactonase<br>RutD: pyrimidine utilization protein D | 5.80E-58<br>3.20E-43<br>3.10E-10<br>3.40E-09<br>1.50E-08 |
| WP_155984839.1 | 85225 | 84352 | - | 290 | LinG<br>LinH | LinG<br>LinH | LinG Homolog<br>LinH (Fusion of LinE/LinG) | 9.30E-100<br>1.00E-87 |
| WP_024877745.1 | 85560 | 85299 | - | 86 | LinA | LinA | LinA (linaridin precursor) | 3.10E-08 |
| WP_024877746.1 | 85916 | 85652 | - | 87 | LinA | LinA | LinA (linaridin precursor) | 6.50E-07 |
| WP_081761953.1 | 85935 | 86985 | + | 349 | PF13561<br>TIGR01830<br>TIGR02415<br>PF00106<br>TIGR01829 | adh_short_C2<br>TIGR01830<br>TIGR02415<br>adh_short<br>TIGR01829 | Enoyl-(Acyl carrier protein) reductase<br>3oxo_ACP_reduc: 3-oxoacyl-[acyl-carrier-protein] reductase<br>23BDH: acetoin reductases<br>short chain dehydrogenase<br>AcAcCoA_reduct: acetoacetyl-CoA reductase | 9.40E-62<br>1.80E-55<br>5.00E-51<br>7.00E-49<br>3.80E-48 |

| index | leader | core | start | end | dir | score |
|---|---|---|---|---|---|---|
| 1 | MNKVEQAAQAAQAGHEGQLLPLRRNQAAAPALSAAWVAATSAVTLA | ATAIYQAGNAVTDYVGNHTNPEINPEALSGKTGAELLSIRQNGVRS | 82820 | 82541 | - | 11 |
| 2 | MSAVAAFANSKADHGLAPLEASQVYTPVL | CTTTTTLTTTTATCTVFATGISSDA...DVSPAGLPAHGSAEELLNFRTGSIG | 85560 | 85299 | - | 15 |
| 3 | MSAVAAFANSKADHGLAPLEASQVYTPVF | CTTTTTTSTITGCVGTVFTMGVISD...DVSPAGLPAHGSAEELLNFRAGSIG | 85916 | 85652 | - | 15 |

### Results for WP_138383132.1 [Saccharomonospora piscinae] back to top

previous - next

#### Architecture

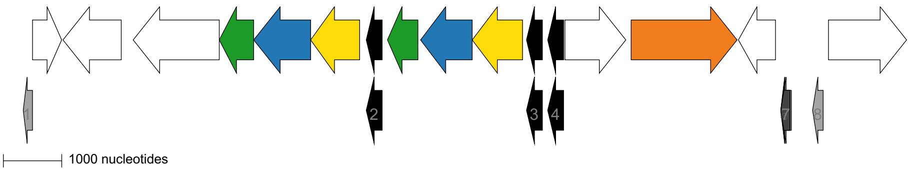

Link to nucleotide sequence

| Accession | start | end | direction | length (aa) | Pfam/HMM | name | description | E-value |
|---|---|---|---|---|---|---|---|---|
| WP_081191132.1 | 997755 | 998262 | + | 168 | PF08327 | AHSA1 | Activator of Hsp90 ATPase homolog 1-like protein | 4.60E-12 |
| WP_138383236.1 | 999291 | 998283 | - | 335 | TIGR03451<br>TIGR00692<br>TIGR03989<br>TIGR02818<br>TIGR02819 | TIGR03451<br>TIGR00692<br>TIGR03989<br>TIGR02818<br>TIGR02819 | mycoS_dep_FDH: S-(hydroxymethyl)mycothiol dehydrogenase<br>tdh: L-threonine 3-dehydrogenase<br>Rxyl_3153: NDMA-dependent alcohol dehydrogenase, Rxyl_3153 family<br>adh_III_F_hyde: S-(hydroxymethyl)glutathione dehydrogenase/class III alcohol dehydrogenase<br>fdhA_non_GSH: formaldehyde dehydrogenase, glutathione-independent | 4.90E-39<br>1.20E-38<br>6.00E-31<br>1.10E-28<br>1.50E-28 |
| WP_138383237.1 | 1000956 | 999489 | - | 488 | PF00171<br>TIGR02299<br>TIGR01804<br>TIGR03216<br>TIGR01780 | Aldedh<br>TIGR02299<br>TIGR01804<br>TIGR03216<br>TIGR01780 | Aldehyde dehydrogenase family<br>HpaE: 5-carboxymethyl-2-hydroxymuconate semialdehyde dehydrogenase<br>BADH: betaine-aldehyde dehydrogenase<br>OH_muco_semi_DH: 2-hydroxymuconic semialdehyde dehydrogenase<br>SSADH: succinate-semialdehyde dehydrogenase | 1.50E-151<br>3.90E-122<br>1.50E-121<br>5.90E-106<br>1.40E-100 |
| WP_081191131.1 | 1001548 | 1000975 | - | 190 | LinL | LinL | LinL | 2.60E-68 |
| WP_138383130.1 | 1002528 | 1001547 | - | 326 | LinE<br>LinH<br>PF12697<br>TIGR02427<br>TIGR03695 | LinE<br>LinH<br>Abhydrolase_6<br>TIGR02427<br>TIGR03695 | LinE Homolog<br>LinH (Fusion of LinE/LinG)<br>Alpha/beta hydrolase family<br>protocat_pcaD: 3-oxoadipate enol-lactonase<br>menH_SHCHC: 2-succinyl-6-hydroxy-2,4-cyclohexadiene-1-carboxylate synthase | 2.30E-86<br>4.80E-67<br>6.80E-15<br>9.30E-10<br>1.10E-09 |
| WP_138383238.1 | 1003379 | 1002524 | - | 284 | LinG<br>LinH<br>PF06762<br>TIGR04033 | LinG<br>LinH<br>LMF1<br>TIGR04033 | LinG Homolog<br>LinH (Fusion of LinE/LinG)<br>Lipase maturation factor<br>export_SdpB: antimicrobial peptide system protein, SdpB family | 9.30E-102<br>1.50E-95<br>1.00E-06<br>4.00E-04 |
| WP_138383131.1 | 1003760 | 1003481 | - | 92 | LinA | LinA | LinA (linaridin precursor) | 5.10E-10 |

| | | | | | | | | |
|---|---|---|---|---|---|---|---|---|
| WP_081192168.1 | 1004377 | 1003843 | - | 177 | LinL | LinL | LinL | 8.50E-59 |
| WP_138383132.1 | 1005296 | 1004411 | - | 294 | LinE | LinE | LinE Homolog | 2.00E-58 |
| | | | | | LinH | LinH | LinH (Fusion of LinE/LinG) | 2.00E-43 |
| | | | | | PF12697 | Abhydrolase_6 | Alpha/beta hydrolase family | 7.20E-11 |
| | | | | | TIGR02427 | TIGR02427 | protocat_pcaD: 3-oxoadipate enol-lactonase | 3.10E-10 |
| | | | | | TIGR03611 | TIGR03611 | RutD: pyrimidine utilization protein D | 3.10E-09 |
| WP_138383133.1 | 1006165 | 1005292 | - | 290 | LinG | LinG | LinG Homolog | 1.10E-99 |
| | | | | | LinH | LinH | LinH (Fusion of LinE/LinG) | 1.20E-87 |
| WP_138383134.1 | 1006495 | 1006237 | - | 85 | LinA | LinA | LinA (linaridin precursor) | 6.30E-09 |
| WP_081191125.1 | 1006861 | 1006597 | - | 87 | LinA | LinA | LinA (linaridin precursor) | 3.80E-08 |
| WP_138383239.1 | 1006880 | 1007930 | + | 349 | PF13561 | adh_short_C2 | Enoyl-(Acyl carrier protein) reductase | 6.30E-62 |
| | | | | | TIGR01830 | TIGR01830 | 3oxo_ACP_reduc: 3-oxoacyl-[acyl-carrier-protein] reductase | 1.50E-55 |
| | | | | | TIGR02415 | TIGR02415 | 23BDH: acetoin reductases | 8.30E-51 |
| | | | | | PF00106 | adh_short | short chain dehydrogenase | 1.60E-49 |
| | | | | | TIGR01829 | TIGR01829 | AcAcCoA_reduct: acetoacetyl-CoA reductase | 2.40E-48 |
| WP_138383135.1 | 1008016 | 1009846 | + | 609 | TIGR02204 | TIGR02204 | MsbA_rel: ABC transporter, permease/ATP-binding protein | 2.30E-105 |
| | | | | | TIGR02203 | TIGR02203 | MsbA_lipidA: lipid A export permease/ATP-binding protein MsbA | 2.70E-96 |
| | | | | | TIGR00958 | TIGR00958 | 3a01208: antigen peptide transporter 2 | 2.60E-89 |
| | | | | | TIGR03796 | TIGR03796 | NHLM_micro_ABC1: NHLM bacteriocin system ABC transporter, peptidase/ATP-binding protein | 8.90E-85 |
| | | | | | TIGR03375 | TIGR03375 | type_I_sec_LssB: type I secretion system ATPase | 1.90E-81 |
| WP_138383136.1 | 1010489 | 1009856 | - | 210 | PF00196 | GerE | Bacterial regulatory proteins, luxR family | 4.10E-18 |
| | | | | | TIGR03020 | TIGR03020 | EpsA: transcriptional regulator EpsA | 5.40E-13 |
| | | | | | TIGR03541 | TIGR03541 | reg_near_HchA: LuxR family transcriptional regulatory, chaperone HchA-associated | 1.70E-09 |
| | | | | | TIGR02937 | TIGR02937 | sigma70-ECF: RNA polymerase sigma factor, sigma-70 family | 4.60E-05 |
| | | | | | PF13384 | HTH_23 | Homeodomain-like domain | 3.00E-04 |
| WP_167305385.1 | 1011411 | 1012734 | + | 440 | PF06863 | DUF1254 | Protein of unknown function (DUF1254) | 2.10E-42 |
| | | | | | PF06742 | DUF1214 | Protein of unknown function (DUF1214) | 5.00E-32 |

| index | leader | core | start | end | dir | score |
|---|---|---|---|---|---|---|
| 1 | VGTGLMRCVPLRSRWGHVRGVRGAYD | GLRSPARRAGAASRAARRTARATRRGR | 997772 | 997610 | - | 6 |
| 2 | MNKVEQAAQAAQAGHEGQLLPLRRDEPAAPALSAAWVAATAGVTLA | ATAVFQAGNAVTDYVGNHTNPEINPEALSGKSGTELLSIRQNGVRR | 1003760 | 1003481 | - | 14 |
| 3 | MSAAAAFANSKAGQGLAPLEANQVYAPVL | CTATTLTTTTATCTVFATGSIGDAG...DVSSAGLPAHGSAEELLNFRAGSIG | 1006495 | 1006237 | - | 17 |
| 4 | MSVIAAFANSKADHGLAPLEASPVYTPVF | CTTSTTTATLAGTVGMVFTAGIISD...DVSSAGLPAHGSAQDLLNFRAGSIG | 1006861 | 1006597 | - | 15 |
| 5 | LLSPVGATIRLSSETQSGMRIIRQQA | VNWRGATRADDAPRWACRTAGETAAR | 1010751 | 1010592 | - | 6 |
| 6 | LLLSPVGATIRLSSETQSGMRIIRQQ | AVNWRGATRADDAPRWACRTAGETAAR | 1010754 | 1010592 | - | 6 |
| 7 | VRLLLSPVGATIRLSSETQSGMRIIRQ | QAVNWRGATRADDAPRWACRTAGETAAR | 1010760 | 1010592 | - | 6 |
| 8 | VWPATIDNGSTACHRWVAARAVLDLGPGV | LLPVRTHSSSRALRAGPGNRPRPVTAPRISRPV | 1011323 | 1011134 | - | 6 |

### Results for WP_155984838.1 [Saccharomonospora sp. CNQ490] back to top

previous - next

#### Architecture

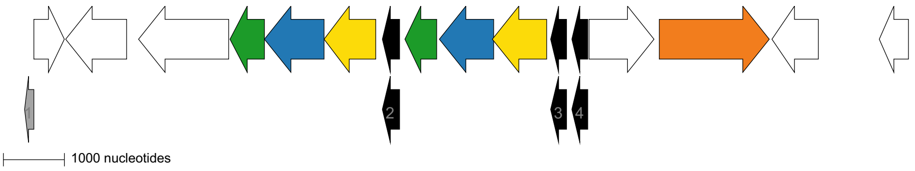

Link to nucleotide sequence

| Accession | start | end | direction | length (aa) | Pfam/HMM | name | description | E-value |
|---|---|---|---|---|---|---|---|---|
| WP_024877737.1 | 76817 | 77324 | + | 168 | PF08327 | AHSA1 | Activator of Hsp90 ATPase homolog 1-like protein | 5.00E-12 |
| WP_037336182.1 | 78353 | 77345 | - | 335 | TIGR03451 | TIGR03451 | mycoS_dep_FDH: S-(hydroxymethyl)mycothiol dehydrogenase | 5.20E-38 |
| | | | | | TIGR00692 | TIGR00692 | tdh: L-threonine 3-dehydrogenase | 9.30E-38 |
| | | | | | TIGR03989 | TIGR03989 | Rxyl_3153: NDMA-dependent alcohol dehydrogenase, Rxyl_3153 family | 6.20E-31 |
| | | | | | TIGR02819 | TIGR02819 | fdhA_non_GSH: formaldehyde dehydrogenase, glutathione-independent | 1.30E-29 |
| | | | | | TIGR03366 | TIGR03366 | HpnZ_proposed: putative phosphonate catabolism associated alcohol dehydrogenase | 3.90E-28 |
| WP_051473488.1 | 80018 | 78551 | - | 488 | PF00171 | Aldedh | Aldehyde dehydrogenase family | 5.80E-151 |
| | | | | | TIGR02299 | TIGR02299 | HpaE: 5-carboxymethyl-2-hydroxymuconate semialdehyde dehydrogenase | 1.30E-122 |
| | | | | | TIGR01804 | TIGR01804 | BADH: betaine-aldehyde dehydrogenase | 2.50E-121 |
| | | | | | TIGR03216 | TIGR03216 | OH_muco_semi_DH: 2-hydroxymuconic semialdehyde dehydrogenase | 2.40E-104 |
| | | | | | TIGR01780 | TIGR01780 | SSADH: succinate-semialdehyde dehydrogenase | 4.50E-100 |
| WP_024877740.1 | 80610 | 80037 | - | 190 | LinL | LinL | LinL | 2.00E-66 |
| WP_037336134.1 | 81590 | 80609 | - | 326 | LinE | LinE | LinE Homolog | 1.90E-89 |
| | | | | | LinH | LinH | LinH (Fusion of LinE/LinG) | 2.40E-69 |
| | | | | | PF12697 | Abhydrolase_6 | Alpha/beta hydrolase family | 9.80E-17 |
| | | | | | TIGR02427 | TIGR02427 | protocat_pcaD: 3-oxoadipate enol-lactonase | 4.70E-12 |
| | | | | | PF12146 | Hydrolase_4 | Serine aminopeptidase, S33 | 3.10E-10 |
| WP_037336136.1 | 82441 | 81586 | - | 284 | LinG | LinG | LinG Homolog | 1.60E-101 |
| | | | | | LinH | LinH | LinH (Fusion of LinE/LinG) | 1.20E-94 |
| | | | | | PF06762 | LMF1 | Lipase maturation factor | 4.80E-07 |
| | | | | | TIGR04033 | TIGR04033 | export_SdpB: antimicrobial peptide system protein, SdpB family | 1.00E-04 |
| WP_024877741.1 | 82820 | 82541 | - | 92 | LinA | LinA | LinA (linaridin precursor) | 7.10E-09 |

| | | | | | | | | |
|---|---|---|---|---|---|---|---|---|
| WP_051473489.1 | 83437 | 82903 | - | 177 | LinL | LinL | LinL | 2.40E-58 |
| WP_155984838.1 | 84356 | 83471 | - | 294 | LinE<br>LinH<br>PF12697<br>TIGR02427<br>TIGR03611 | LinE<br>LinH<br>Abhydrolase_6<br>TIGR02427<br>TIGR03611 | LinE Homolog<br>LinH (Fusion of LinE/LinG)<br>Alpha/beta hydrolase family<br>protocat_pcaD: 3-oxoadipate enol-lactonase<br>RutD: pyrimidine utilization protein D | 5.80E-58<br>3.20E-43<br>3.10E-10<br>3.40E-09<br>1.50E-08 |
| WP_155984839.1 | 85225 | 84352 | - | 290 | LinG<br>LinH | LinG<br>LinH | LinG Homolog<br>LinH (Fusion of LinE/LinG) | 9.30E-100<br>1.00E-87 |
| WP_024877745.1 | 85560 | 85299 | - | 86 | LinA | LinA | LinA (linaridin precursor) | 3.10E-08 |
| WP_024877746.1 | 85916 | 85652 | - | 87 | LinA | LinA | LinA (linaridin precursor) | 6.50E-07 |
| WP_081761953.1 | 85935 | 86985 | + | 349 | PF13561<br>TIGR01830<br>TIGR02415<br>PF00106<br>TIGR01829 | adh_short_C2<br>TIGR01830<br>TIGR02415<br>adh_short<br>TIGR01829 | Enoyl-(Acyl carrier protein) reductase<br>3oxo_ACP_reduc: 3-oxoacyl-[acyl-carrier-protein] reductase<br>23BDH: acetoin reductases<br>short chain dehydrogenase<br>AcAcCoA_reduct: acetoacetyl-CoA reductase | 9.40E-62<br>1.80E-55<br>5.00E-51<br>7.00E-49<br>3.80E-48 |
| WP_024877748.1 | 87072 | 88884 | + | 603 | TIGR02204<br>TIGR02203<br>TIGR00958<br>TIGR03796<br>TIGR01846 | TIGR02204<br>TIGR02203<br>TIGR00958<br>TIGR03796<br>TIGR01846 | MsbA_rel: ABC transporter, permease/ATP-binding protein<br>MsbA_lipidA: lipid A export permease/ATP-binding protein MsbA<br>3a01208: antigen peptide transporter 2<br>NHLM_micro_ABC1: NHLM bacteriocin system ABC transporter, peptidase/ATP-binding protein<br>type_I_sec_HlyB: type I secretion system ATPase | 6.80E-106<br>3.50E-96<br>2.60E-89<br>2.20E-86<br>3.40E-81 |
| WP_081761954.1 | 89695 | 88918 | - | 258 | PF00196<br>TIGR03020<br>TIGR03541<br>TIGR02937<br>PF13384 | GerE<br>TIGR03020<br>TIGR03541<br>TIGR02937<br>HTH_23 | Bacterial regulatory proteins, luxR family<br>EpsA: transcriptional regulator EpsA<br>reg_near_HchA: LuxR family transcriptional regulatory, chaperone HchA-associated<br>sigma70-ECF: RNA polymerase sigma factor, sigma-70 family<br>Homeodomain-like domain | 5.90E-18<br>7.40E-13<br>2.20E-09<br>5.10E-05<br>2.90E-04 |
| WP_024877750.1 | 91156 | 90697 | - | 152 | TIGR03618<br>PF01243<br>TIGR03667<br>TIGR03666<br>TIGR03668 | TIGR03618<br>Putative_PNPOx<br>TIGR03667<br>TIGR03666<br>TIGR03668 | Rv1155_F420: PPOX class probable F420-dependent enzyme<br>Pyridoxamine 5'-phosphate oxidase<br>Rv3369: PPOX class probable F420-dependent enzyme, Rv3369 family<br>Rv2061_F420: PPOX class probable F420-dependent enzyme, Rv2061 family<br>Rv0121_F420: PPOX class probable F420-dependent enzyme, Rv0121 family | 1.10E-36<br>3.10E-12<br>2.70E-08<br>2.40E-05<br>4.50E-05 |

| index | leader | core | start | end | dir | score |
|---|---|---|---|---|---|---|
| 1 | VGTGLMRCVPLRSRWGHVRGVRGAYD | GPRSPARRAGAASRAALRTARATRRGR | 76834 | 76672 | - | 6 |
| 2 | MNKVEQAAQAAQAGHEGQLLPLRRNQAAAPALSAAWVAATSAVTLA | ATAIYQAGNAVTDYVGNHTNPEINPEALSGKTGAELLSIRQNGVRS | 82820 | 82541 | - | 12 |
| 3 | MSAVAAFANSKADHGLAPLEASQVYTPVL | CTTTTTLTTTTATCTVFATGISSDA...DVSPAGLPAHGSAEELLNFRTGSIG | 85560 | 85299 | - | 16 |
| 4 | MSAVAAFANSKADHGLAPLEASQVYTPVF | CTTTTTTSTITGCVGTVFTMGVISD...DVSPAGLPAHGSAEELLNFRAGSIG | 85916 | 85652 | - | 16 |

### Results for OQO91992.1 [Saccharomonospora sp. LRS4.154] back to top

previous - next

#### Architecture

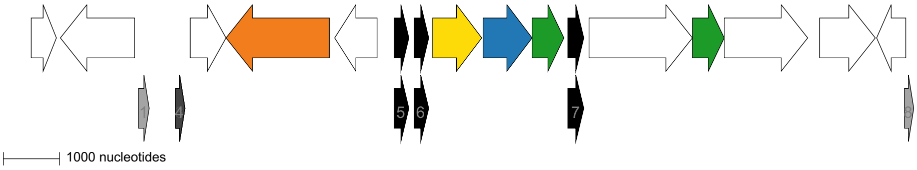

Link to nucleotide sequence

| Accession | start | end | direction | length (aa) | Pfam/HMM | name | description | E-value |
|---|---|---|---|---|---|---|---|---|
| OQO91985.1 | 86546 | 87005 | + | 152 | TIGR03618<br>PF01243<br>TIGR03667<br>TIGR03668<br>TIGR03666 | TIGR03618<br>Putative_PNPOx<br>TIGR03667<br>TIGR03668<br>TIGR03666 | Rv1155_F420: PPOX class probable F420-dependent enzyme<br>Pyridoxamine 5'-phosphate oxidase<br>Rv3369: PPOX class probable F420-dependent enzyme, Rv3369 family<br>Rv0121_F420: PPOX class probable F420-dependent enzyme, Rv0121 family<br>Rv2061_F420: PPOX class probable F420-dependent enzyme, Rv2061 family | 5.70E-37<br>1.60E-11<br>3.50E-08<br>2.30E-05<br>3.10E-05 |
| OQO91986.1 | 88385 | 87062 | - | 440 | PF06863<br>PF06742 | DUF1254<br>DUF1214 | Protein of unknown function (DUF1254)<br>Protein of unknown function (DUF1214) | 1.90E-43<br>6.30E-32 |
| OQO91987.1 | 89387 | 90020 | + | 210 | PF00196<br>TIGR03020<br>TIGR03541<br>TIGR02937<br>PF13384 | GerE<br>TIGR03020<br>TIGR03541<br>TIGR02937<br>HTH_23 | Bacterial regulatory proteins, luxR family<br>EpsA: transcriptional regulator EpsA<br>reg_near_HchA: LuxR family transcriptional regulatory, chaperone HchA-associated<br>sigma70-ECF: RNA polymerase sigma factor, sigma-70 family<br>Homeodomain-like domain | 3.50E-18<br>5.30E-13<br>1.70E-09<br>4.00E-05<br>2.90E-04 |
| OQO91988.1 | 91860 | 90030 | - | 609 | TIGR02204<br>TIGR02203<br>TIGR00958<br>TIGR03796<br>TIGR03375 | TIGR02204<br>TIGR02203<br>TIGR00958<br>TIGR03796<br>TIGR03375 | MsbA_rel: ABC transporter, permease/ATP-binding protein<br>MsbA_lipidA: lipid A export permease/ATP-binding protein MsbA<br>3a01208: antigen peptide transporter 2<br>NHLM_micro_ABC1: NHLM bacteriocin system ABC transporter, peptidase/ATP-binding protein<br>type_I_sec_LssB: type I secretion system ATPase | 3.60E-105<br>5.30E-96<br>5.40E-89<br>8.70E-85<br>5.90E-81 |
| OQO91989.1 | 92711 | 91946 | - | 254 | PF13561<br>TIGR01830<br>TIGR02415<br>PF00106<br>TIGR01829 | adh_short_C2<br>TIGR01830<br>TIGR02415<br>adh_short<br>TIGR01829 | Enoyl-(Acyl carrier protein) reductase<br>3oxo_ACP_reduc: 3-oxoacyl-[acyl-carrier-protein] reductase<br>23BDH: acetoin reductases<br>short chain dehydrogenase<br>AcAcCoA_reduct: acetoacetyl-CoA reductase | 2.80E-62<br>5.30E-56<br>2.60E-51<br>6.70E-50<br>8.60E-49 |
| OQO91990.1 | 93015 | 93279 | + | 87 | LinA | LinA | LinA (linaridin precursor) | 3.80E-08 |
| OQO91991.1 | 93381 | 93639 | + | 85 | LinA | LinA | LinA (linaridin precursor) | 4.00E-09 |
| OQO93074.1 | 93711 | 94584 | + | 290 | LinG<br>LinH | LinG<br>LinH | LinG Homolog<br>LinH (Fusion of LinE/LinG) | 7.20E-100<br>1.20E-87 |

| | | | | | | | | |
|---|---|---|---|---|---|---|---|---|
| OQO91992.1 | 94604 | 95465 | + | 286 | LinE<br>LinH<br>PF12697<br>TIGR02427<br>TIGR03611 | LinE<br>LinH<br>Abhydrolase_6<br>TIGR02427<br>TIGR03611 | LinE Homolog<br>LinH (Fusion of LinE/LinG)<br>Alpha/beta hydrolase family<br>protocat_pcaD: 3-oxoadipate enol-lactonase<br>RutD: pyrimidine utilization protein D | 7.60E-58<br>7.50E-43<br>6.60E-11<br>2.40E-09<br>9.70E-09 |
| OQO91993.1 | 95466 | 96033 | + | 188 | LinL | LinL | LinL | 2.60E-61 |
| OQO91994.1 | 96116 | 96395 | + | 92 | LinA | LinA | LinA (linaridin precursor) | 4.60E-10 |
| WP_016907298.1 | 96496 | 98328 | + | 610 | INFERRED GENE | - | - | - |
| OQO91995.1 | 98327 | 98900 | + | 190 | LinL | LinL | LinL | 2.60E-68 |
| OQO91996.1 | 98904 | 100386 | + | 493 | PF00171<br>TIGR01804<br>TIGR02299<br>TIGR03216<br>TIGR01780 | Aldedh<br>TIGR01804<br>TIGR02299<br>TIGR03216<br>TIGR01780 | Aldehyde dehydrogenase family<br>BADH: betaine-aldehyde dehydrogenase<br>HpaE: 5-carboxymethyl-2-hydroxymuconate semialdehyde dehydrogenase<br>OH_muco_semi_DH: 2-hydroxymuconic semialdehyde dehydrogenase<br>SSADH: succinate-semialdehyde dehydrogenase | 1.60E-152<br>4.00E-123<br>6.60E-123<br>4.30E-105<br>1.80E-101 |
| OQO93075.1 | 100584 | 101592 | + | 335 | TIGR03451<br>TIGR00692<br>TIGR03989<br>TIGR02818<br>TIGR02819 | TIGR03451<br>TIGR00692<br>TIGR03989<br>TIGR02818<br>TIGR02819 | mycoS_dep_FDH: S-(hydroxymethyl)mycothiol dehydrogenase<br>tdh: L-threonine 3-dehydrogenase<br>Rxyl_3153: NDMA-dependent alcohol dehydrogenase, Rxyl_3153 family<br>adh_III_F_hyde: S-(hydroxymethyl)glutathione dehydrogenase/class III alcohol dehydrogenase<br>fdhA_non_GSH: formaldehyde dehydrogenase, glutathione-independent | 5.20E-39<br>1.40E-38<br>6.80E-31<br>1.70E-28<br>2.10E-28 |
| OQO91997.1 | 102120 | 101613 | - | 168 | PF08327 | AHSA1 | Activator of Hsp90 ATPase homolog 1-like protein | 4.60E-12 |

| index | leader | core | start | end | dir | score |
|---|---|---|---|---|---|---|
| 1 | VWPATIDNGSTACHRWVAARAVLDLGPGV | LLPVRTHSSSRALRAGPGNRPRPVTAPRISRPV | 88473 | 88662 | + | 6 |
| 2 | VRLLLSPVGATIRLSSETQSGMRIIRQ | QAVNWRGATRADDAPRWACRTAGETAAR | 89116 | 89284 | + | 6 |
| 3 | LLLSPVGATIRLSSETQSGMRIIRQQ | AVNWRGATRADDAPRWACRTAGETAAR | 89122 | 89284 | + | 6 |
| 4 | LLSPVGATIRLSSETQSGMRIIRQQA | VNWRGATRADDAPRWACRTAGETAAR | 89125 | 89284 | + | 6 |
| 5 | MSVIAAFANSKADHGLAPLEASPVYTPVF | CTTSTTTATLAGTVGMVFTAGIISD...DVSSAGLPAHGSAQDLLNFRAGSIG | 93015 | 93279 | + | 15 |
| 6 | MSAAAAFANSKAGHGLAPLEANQVYAPVL | CTATTLTTTTATCTVFATGIIGDAG...DVSSAGLPAHGSAEELLNFRAGSIG | 93381 | 93639 | + | 17 |
| 7 | MNKVEQAAQAAQAGHEGQLLPLRRDEPAAPALSAAWVAATAGVTLA | ATAIYQAGNAVTDYVGNHTNPEINPEALSGKSGTELLSIRQNGVRR | 96116 | 96395 | + | 13 |
| 8 | VGTGLMRCVPLRSRWGHVRGVRGAYD | GLRSPARRAGAASRAARRTARATRRGR | 102103 | 102265 | + | 6 |

### Results for WP_050590090.1 [Salinispora arenicola CNH905] back to top

previous - next

#### Architecture

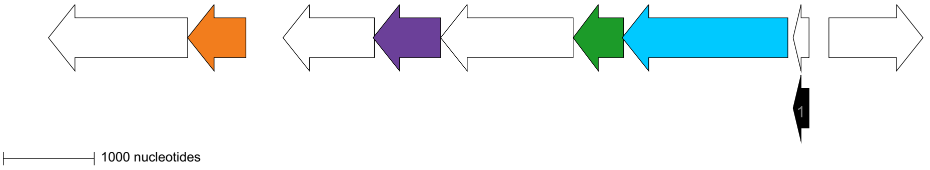

Link to nucleotide sequence

| Accession | start | end | direction | length (aa) | Pfam/HMM | name | description | E-value |
|---|---|---|---|---|---|---|---|---|
| WP_029024475.1 | 1798 | 262 | - | 511 | NO PFAM MATCH | - | - | - |
| WP_029024476.1 | 2432 | 1799 | - | 210 | TIGR01189<br>TIGR00968<br>TIGR03873<br>TIGR01188<br>TIGR03740 | TIGR01189<br>TIGR00968<br>TIGR03873<br>TIGR01188<br>TIGR03740 | ccmA: heme ABC exporter, ATP-binding protein CcmA<br>3a0106s01: sulfate ABC transporter, ATP-binding protein<br>F420-0_ABC_ATP: proposed F420-0 ABC transporter, ATP-binding protein<br>drrA: daunorubicin resistance ABC transporter, ATP-binding protein<br>gallidem_ABC: lantibiotic protection ABC transporter, ATP-binding subunit | 4.00E-34<br>1.50E-30<br>1.00E-27<br>1.30E-27<br>2.10E-27 |
| WP_050590088.1 | 3848 | 2843 | - | 334 | PF00144<br>PF13354 | Beta-lactamase<br>Beta-lactamase2 | Beta-lactamase<br>Beta-lactamase enzyme family | 7.00E-48<br>3.90E-05 |
| WP_029024477.1 | 4588 | 3844 | - | 247 | PF13649<br>PF08241<br>PF13489<br>PF08242<br>PF13847 | Methyltransf_25<br>Methyltransf_11<br>Methyltransf_23<br>Methyltransf_12<br>Methyltransf_31 | Methyltransferase domain<br>Methyltransferase domain<br>Methyltransferase domain<br>Methyltransferase domain<br>Methyltransferase domain | 1.10E-12<br>2.70E-10<br>5.40E-10<br>1.00E-08<br>7.50E-08 |
| WP_032724752.1 | 6048 | 4584 | - | 487 | TIGR02734<br>PF13450<br>TIGR03467<br>TIGR00292<br>PF01946 | TIGR02734<br>NAD_binding_8<br>TIGR03467<br>TIGR00292<br>Thi4 | crtI_fam: phytoene desaturase<br>NAD(P)-binding Rossmann-like domain<br>HpnE: squalene-associated FAD-dependent desaturase<br>TIGR00292: thiazole biosynthesis enzyme<br>Thi4 family | 5.50E-11<br>1.20E-10<br>1.60E-09<br>3.80E-08<br>1.60E-07 |
| WP_050590089.1 | 6590 | 6044 | - | 181 | LinL | LinL | LinL | 1.40E-60 |
| WP_050590090.1 | 8401 | 6586 | - | 604 | LinH<br>LinG<br>LinE<br>PF12697<br>TIGR02427 | LinH<br>LinG<br>LinE<br>Abhydrolase_6<br>TIGR02427 | LinH (Fusion of LinE/LinG)<br>LinG Homolog<br>LinE Homolog<br>Alpha/beta hydrolase family<br>protocat_pcaD: 3-oxoadipate enol-lactonase | 1.50E-221<br>1.30E-94<br>6.50E-91<br>2.20E-18<br>3.40E-11 |
| WP_155254174.1 | 8645 | 8468 | - | 58 | NO PFAM MATCH | - | - | - |
| WP_019030235.1 | 8860 | 9890 | + | 343 | INFERRED GENE | - | - | - |

| index | leader | core | start | end | dir | score |
|---|---|---|---|---|---|---|
| 1 | MPNIAQHANMVSETQFQGLSEESPSRKINDGEPLG | LLPVIVAAAAITAIGSVLGIGSV | 8645 | 8468 | - | 19 |

### Results for WP_081191130.1 [Saccharomonospora sp. LRS4.154] back to top

previous - next

#### Architecture

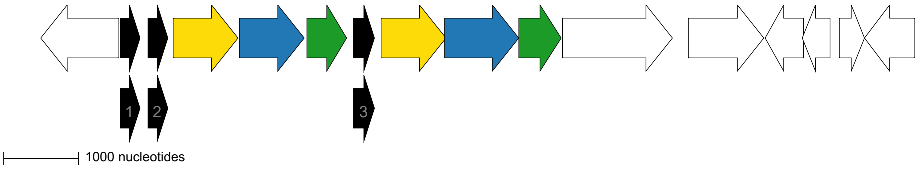

Link to nucleotide sequence

| Accession | start | end | direction | length (aa) | Pfam/HMM | name | description | E-value |
|---|---|---|---|---|---|---|---|---|
| WP_138383239.1 | 92996 | 91946 | - | 349 | PF13561<br>TIGR01830<br>TIGR02415<br>PF00106<br>TIGR01829 | adh_short_C2<br>TIGR01830<br>TIGR02415<br>adh_short<br>TIGR01829 | Enoyl-(Acyl carrier protein) reductase<br>3oxo_ACP_reduc: 3-oxoacyl-[acyl-carrier-protein] reductase<br>23BDH: acetoin reductases<br>short chain dehydrogenase<br>AcAcCoA_reduct: acetoacetyl-CoA reductase | 6.30E-62<br>1.50E-55<br>8.30E-51<br>1.60E-49<br>2.40E-48 |
| WP_081191125.1 | 93015 | 93279 | + | 87 | LinA | LinA | LinA (linaridin precursor) | 3.80E-08 |
| WP_081191126.1 | 93381 | 93639 | + | 85 | LinA | LinA | LinA (linaridin precursor) | 4.00E-09 |
| WP_081192166.1 | 93711 | 94584 | + | 290 | LinG<br>LinH | LinG<br>LinH | LinG Homolog<br>LinH (Fusion of LinE/LinG) | 7.20E-100<br>1.20E-87 |
| WP_139794736.1 | 94592 | 95465 | + | 290 | LinE<br>LinH<br>PF12697<br>TIGR02427<br>TIGR03611 | LinE<br>LinH<br>Abhydrolase_6<br>TIGR02427<br>TIGR03611 | LinE Homolog<br>LinH (Fusion of LinE/LinG)<br>Alpha/beta hydrolase family<br>protocat_pcaD: 3-oxoadipate enol-lactonase<br>RutD: pyrimidine utilization protein D | 5.50E-58<br>5.70E-43<br>6.80E-11<br>2.50E-09<br>1.00E-08 |
| WP_081192168.1 | 95499 | 96033 | + | 177 | LinL | LinL | LinL | 8.50E-59 |
| WP_081191128.1 | 96116 | 96395 | + | 92 | LinA | LinA | LinA (linaridin precursor) | 4.60E-10 |
| WP_081191129.1 | 96496 | 97351 | + | 284 | LinG<br>LinH<br>PF06762<br>TIGR04033 | LinG<br>LinH<br>LMF1<br>TIGR04033 | LinG Homolog<br>LinH (Fusion of LinE/LinG)<br>Lipase maturation factor<br>export_SdpB: antimicrobial peptide system protein, SdpB family | 9.70E-102<br>4.50E-95<br>2.90E-07<br>1.20E-04 |
| WP_081191130.1 | 97347 | 98328 | + | 326 | LinE<br>LinH<br>PF12697<br>TIGR03695<br>PF12146 | LinE<br>LinH<br>Abhydrolase_6<br>TIGR03695<br>Hydrolase_4 | LinE Homolog<br>LinH (Fusion of LinE/LinG)<br>Alpha/beta hydrolase family<br>menH_SHCHC: 2-succinyl-6-hydroxy-2,4-cyclohexadiene-1-carboxylate synthase<br>Serine aminopeptidase, S33 | 1.40E-85<br>6.20E-66<br>8.60E-14<br>4.40E-09<br>7.60E-09 |
| WP_081191131.1 | 98327 | 98900 | + | 190 | LinL | LinL | LinL | 2.60E-68 |
| WP_081192170.1 | 98919 | 100386 | + | 488 | PF00171<br>TIGR01804<br>TIGR02299<br>TIGR03216<br>TIGR01780 | Aldedh<br>TIGR01804<br>TIGR02299<br>TIGR03216<br>TIGR01780 | Aldehyde dehydrogenase family<br>BADH: betaine-aldehyde dehydrogenase<br>HpaE: 5-carboxymethyl-2-hydroxymuconate semialdehyde dehydrogenase<br>OH_muco_semi_DH: 2-hydroxymuconic semialdehyde dehydrogenase<br>SSADH: succinate-semialdehyde dehydrogenase | 1.50E-152<br>3.80E-123<br>6.40E-123<br>4.10E-105<br>1.70E-101 |
| WP_081192172.1 | 100584 | 101592 | + | 335 | TIGR03451<br>TIGR00692<br>TIGR03989<br>TIGR02818<br>TIGR02819 | TIGR03451<br>TIGR00692<br>TIGR03989<br>TIGR02818<br>TIGR02819 | mycoS_dep_FDH: S-(hydroxymethyl)mycothiol dehydrogenase<br>tdh: L-threonine 3-dehydrogenase<br>Rxyl_3153: NDMA-dependent alcohol dehydrogenase, Rxyl_3153 family<br>adh_III_F_hyde: S-(hydroxymethyl)glutathione dehydrogenase/class III alcohol dehydrogenase<br>fdhA_non_GSH: formaldehyde dehydrogenase, glutathione-independent | 5.20E-39<br>1.40E-38<br>6.80E-31<br>1.70E-28<br>2.10E-28 |
| WP_081191132.1 | 102120 | 101613 | - | 168 | PF08327 | AHSA1 | Activator of Hsp90 ATPase homolog 1-like protein | 4.60E-12 |
| WP_081191133.1 | 102479 | 102116 | - | 120 | PF12840<br>PF01022<br>PF12802<br>PF13412<br>PF08279 | HTH_20<br>HTH_5<br>MarR_2<br>HTH_24<br>HTH_11 | Helix-turn-helix domain<br>Bacterial regulatory protein, arsR family<br>MarR family<br>Winged helix-turn-helix DNA-binding<br>HTH domain | 2.50E-15<br>4.10E-14<br>2.50E-08<br>3.50E-06<br>5.90E-06 |
| WP_081191134.1 | 102610 | 102934 | + | 107 | TIGR03954<br>PF12823 | TIGR03954<br>DUF3817 | integ_memb_HG: integral membrane protein<br>Domain of unknown function (DUF3817) | 7.90E-27<br>2.50E-25 |
| WP_081191135.1 | 103607 | 102935 | - | 223 | PF00072<br>PF00196 | Response_reg<br>GerE | Response regulator receiver domain<br>Bacterial regulatory proteins, luxR family | 1.40E-07<br>5.00E-07 |

| index | leader | core | start | end | dir | score |
|---|---|---|---|---|---|---|
| 1 | MSVIAAFANSKADHGLAPLEASPVYTPVF | CTTSTTTATLAGTVGMVFTAGIISD...DVSSAGLPAHGSAQDLLNFRAGSIG | 93015 | 93279 | + | 14 |
| 2 | MSAAAAFANSKAGHGLAPLEANQVYAPVL | CTATTLTTTTATCTVFATGIIGDAG...DVSSAGLPAHGSAEELLNFRAGSIG | 93381 | 93639 | + | 16 |
| 3 | MNKVEQAAQAAQAGHEGQLLPLRRDEPAAPALSAAWVAATAGVTLA | ATAIYQAGNAVTDYVGNHTNPEINPEALSGKSGTELLSIRQNGVRR | 96116 | 96395 | + | 12 |

### Results for WP_139794736.1 [Saccharomonospora sp. LRS4.154] back to top

previous - next

#### Architecture

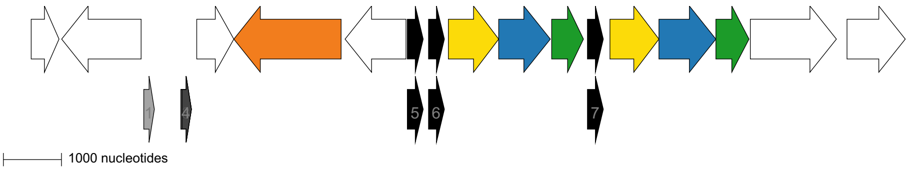

Link to nucleotide sequence

| Accession | start | end | direction | length (aa) | Pfam/HMM | name | description | E-value |
|---|---|---|---|---|---|---|---|---|
| WP_081191120.1 | 86546 | 87005 | + | 152 | TIGR03618 | TIGR03618 | Rv1155_F420: PPOX class probable F420-dependent enzyme | 5.70E-37 |
| | | | | | PF01243 | Putative_PNPOx | Pyridoxamine 5'-phosphate oxidase | 1.60E-11 |
| | | | | | TIGR03667 | TIGR03667 | Rv3369: PPOX class probable F420-dependent enzyme, Rv3369 family | 3.50E-08 |
| | | | | | TIGR03668 | TIGR03668 | Rv0121_F420: PPOX class probable F420-dependent enzyme, Rv0121 family | 2.30E-05 |
| | | | | | TIGR03666 | TIGR03666 | Rv2061_F420: PPOX class probable F420-dependent enzyme, Rv2061 family | 3.10E-05 |
| WP_081191121.1 | 88427 | 87062 | - | 454 | PF06863 | DUF1254 | Protein of unknown function (DUF1254) | 2.10E-43 |
| | | | | | PF06742 | DUF1214 | Protein of unknown function (DUF1214) | 6.70E-32 |
| WP_081191122.1 | 89387 | 90020 | + | 210 | PF00196 | GerE | Bacterial regulatory proteins, luxR family | 3.50E-18 |
| | | | | | TIGR03020 | TIGR03020 | EpsA: transcriptional regulator EpsA | 5.30E-13 |
| | | | | | TIGR03541 | TIGR03541 | reg_near_HchA: LuxR family transcriptional regulatory, chaperone HchA-associated | 1.70E-09 |
| | | | | | TIGR02937 | TIGR02937 | sigma70-ECF: RNA polymerase sigma factor, sigma-70 family | 4.00E-05 |
| | | | | | PF13384 | HTH_23 | Homeodomain-like domain | 2.90E-04 |
| WP_081191123.1 | 91860 | 90030 | - | 609 | TIGR02204 | TIGR02204 | MsbA_rel: ABC transporter, permease/ATP-binding protein | 3.60E-105 |
| | | | | | TIGR02203 | TIGR02203 | MsbA_lipidA: lipid A export permease/ATP-binding protein MsbA | 5.30E-96 |
| | | | | | TIGR00958 | TIGR00958 | 3a01208: antigen peptide transporter 2 | 5.40E-89 |
| | | | | | TIGR03796 | TIGR03796 | NHLM_micro_ABC1: NHLM bacteriocin system ABC transporter, peptidase/ATP-binding protein | 8.70E-85 |
| | | | | | TIGR03375 | TIGR03375 | type_I_sec_LssB: type I secretion system ATPase | 5.90E-81 |
| WP_138383239.1 | 92996 | 91946 | - | 349 | PF13561 | adh_short_C2 | Enoyl-(Acyl carrier protein) reductase | 6.30E-62 |
| | | | | | TIGR01830 | TIGR01830 | 3oxo_ACP_reduc: 3-oxoacyl-[acyl-carrier-protein] reductase | 1.50E-55 |
| | | | | | TIGR02415 | TIGR02415 | 23BDH: acetoin reductases | 8.30E-51 |
| | | | | | PF00106 | adh_short | short chain dehydrogenase | 1.60E-49 |
| | | | | | TIGR01829 | TIGR01829 | AcAcCoA_reduct: acetoacetyl-CoA reductase | 2.40E-48 |
| WP_081191125.1 | 93015 | 93279 | + | 87 | LinA | LinA | LinA (linaridin precursor) | 3.80E-08 |
| WP_081191126.1 | 93381 | 93639 | + | 85 | LinA | LinA | LinA (linaridin precursor) | 4.00E-09 |
| WP_081192166.1 | 93711 | 94584 | + | 290 | LinG | LinG | LinG Homolog | 7.20E-100 |
| | | | | | LinH | LinH | LinH (Fusion of LinE/LinG) | 1.20E-87 |
| WP_139794736.1 | 94592 | 95465 | + | 290 | LinE | LinE | LinE Homolog | 5.50E-58 |
| | | | | | LinH | LinH | LinH (Fusion of LinE/LinG) | 5.70E-43 |
| | | | | | PF12697 | Abhydrolase_6 | Alpha/beta hydrolase family | 6.80E-11 |
| | | | | | TIGR02427 | TIGR02427 | protocat_pcaD: 3-oxoadipate enol-lactonase | 2.50E-09 |
| | | | | | TIGR03611 | TIGR03611 | RutD: pyrimidine utilization protein D | 1.00E-08 |
| WP_081192168.1 | 95499 | 96033 | + | 177 | LinL | LinL | LinL | 8.50E-59 |
| WP_081191128.1 | 96116 | 96395 | + | 92 | LinA | LinA | LinA (linaridin precursor) | 4.60E-10 |
| WP_081191129.1 | 96496 | 97351 | + | 284 | LinG | LinG | LinG Homolog | 9.70E-102 |
| | | | | | LinH | LinH | LinH (Fusion of LinE/LinG) | 4.50E-95 |
| | | | | | PF06762 | LMF1 | Lipase maturation factor | 2.90E-07 |
| | | | | | TIGR04033 | TIGR04033 | export_SdpB: antimicrobial peptide system protein, SdpB family | 1.20E-04 |
| WP_081191130.1 | 97347 | 98328 | + | 326 | LinE | LinE | LinE Homolog | 1.40E-85 |
| | | | | | LinH | LinH | LinH (Fusion of LinE/LinG) | 6.20E-66 |
| | | | | | PF12697 | Abhydrolase_6 | Alpha/beta hydrolase family | 8.60E-14 |
| | | | | | TIGR03695 | TIGR03695 | menH_SHCHC: 2-succinyl-6-hydroxy-2,4-cyclohexadiene-1-carboxylate synthase | 4.40E-09 |
| | | | | | PF12146 | Hydrolase_4 | Serine aminopeptidase, S33 | 7.60E-09 |
| WP_081191131.1 | 98327 | 98900 | + | 190 | LinL | LinL | LinL | 2.60E-68 |
| WP_081192170.1 | 98919 | 100386 | + | 488 | PF00171 | Aldedh | Aldehyde dehydrogenase family | 1.50E-152 |
| | | | | | TIGR01804 | TIGR01804 | BADH: betaine-aldehyde dehydrogenase | 3.80E-123 |
| | | | | | TIGR02299 | TIGR02299 | HpaE: 5-carboxymethyl-2-hydroxymuconate semialdehyde dehydrogenase | 6.40E-123 |
| | | | | | TIGR03216 | TIGR03216 | OH_muco_semi_DH: 2-hydroxymuconic semialdehyde dehydrogenase | 4.10E-105 |
| | | | | | TIGR01780 | TIGR01780 | SSADH: succinate-semialdehyde dehydrogenase | 1.70E-101 |
| WP_081192172.1 | 100584 | 101592 | + | 335 | TIGR03451 | TIGR03451 | mycoS_dep_FDH: S-(hydroxymethyl)mycothiol dehydrogenase | 5.20E-39 |
| | | | | | TIGR00692 | TIGR00692 | tdh: L-threonine 3-dehydrogenase | 1.40E-38 |
| | | | | | TIGR03989 | TIGR03989 | Rxyl_3153: NDMA-dependent alcohol dehydrogenase, Rxyl_3153 family | 6.80E-31 |
| | | | | | TIGR02818 | TIGR02818 | adh_III_F_hyde: S-(hydroxymethyl)glutathione dehydrogenase/class III alcohol dehydrogenase | 1.70E-28 |
| | | | | | TIGR02819 | TIGR02819 | fdhA_non_GSH: formaldehyde dehydrogenase, glutathione-independent | 2.10E-28 |

| index | leader | core | start | end | dir | score |
|---|---|---|---|---|---|---|
| 1 | VWPATIDNGSTACHRWVAARAVLDLGPGV | LLPVRTHSSSRALRAGPGNRPRPVTAPRISRPV | 88473 | 88662 | + | 6 |
| 2 | VRLLLSPVGATIRLSSETQSGMRIIRQ | QAVNWRGATRADDAPRWACRTAGETAAR | 89116 | 89284 | + | 6 |
| 3 | LLLSPVGATIRLSSETQSGMRIIRQQ | AVNWRGATRADDAPRWACRTAGETAAR | 89122 | 89284 | + | 6 |
| 4 | LLSPVGATIRLSSETQSGMRIIRQQA | VNWRGATRADDAPRWACRTAGETAAR | 89125 | 89284 | + | 6 |
| 5 | MSVIAAFANSKADHGLAPLEASPVYTPVF | CTTSTTTATLAGTVGMVFTAGIISD...DVSSAGLPAHGSAQDLLNFRAGSIG | 93015 | 93279 | + | 15 |
| 6 | MSAAAAFANSKAGHGLAPLEANQVYAPVL | CTATTLTTTTATCTVFATGIIGDAG...DVSSAGLPAHGSAEELLNFRAGSIG | 93381 | 93639 | + | 17 |
| 7 | MNKVEQAAQAAQAGHEGQLLPLRRDEPAAPALSAAWVAATAGVTLA | ATAIYQAGNAVTDYVGNHTNPEINPEALSGKSGTELLSIRQNGVRR | 96116 | 96395 | + | 13 |

### Results for WP_050590677.1 [Salinispora pacifica CNT131] back to top

previous - next

#### Architecture

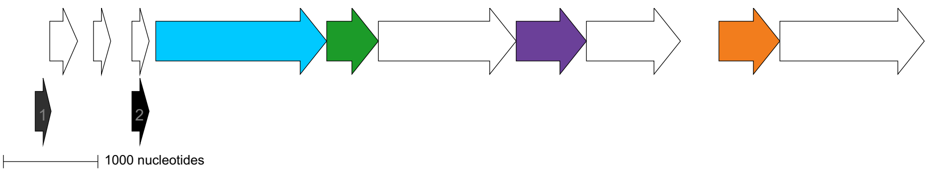

Link to nucleotide sequence

| Accession | start | end | direction | length (aa) | Pfam/HMM | name | description | E-value |
|---|---|---|---|---|---|---|---|---|
| WP_018825617.1 | 168 | 456 | + | 96 | INFERRED GENE | - | - | - |
| WP_011133493.1 | 625 | 808 | + | 61 | INFERRED GENE | - | - | - |
| WP_155254361.1 | 1041 | 1218 | + | 58 | NO PFAM MATCH | - | - | - |
| WP_050590677.1 | 1285 | 3100 | + | 604 | LinH<br>LinG<br>LinE<br>PF12697<br>TIGR02427 | LinH<br>LinG<br>LinE<br>Abhydrolase_6<br>TIGR02427 | LinH (Fusion of LinE/LinG)<br>LinG Homolog<br>LinE Homolog<br>Alpha/beta hydrolase family<br>protocat_pcaD: 3-oxoadipate enol-lactonase | 1.90E-221<br>2.10E-95<br>2.20E-89<br>1.20E-16<br>3.80E-10 |
| WP_050590679.1 | 3096 | 3642 | + | 181 | LinL | LinL | LinL | 1.30E-60 |
| WP_033660744.1 | 3638 | 5102 | + | 487 | TIGR02734<br>PF13450<br>TIGR03467<br>TIGR00292<br>PF01946 | TIGR02734<br>NAD_binding_8<br>TIGR03467<br>TIGR00292<br>Thi4 | crtI_fam: phytoene desaturase<br>NAD(P)-binding Rossmann-like domain<br>HpnE: squalene-associated FAD-dependent desaturase<br>TIGR00292: thiazole biosynthesis enzyme<br>Thi4 family | 4.90E-11<br>1.90E-10<br>1.80E-09<br>3.60E-08<br>1.50E-07 |
| WP_027658662.1 | 5098 | 5842 | + | 247 | PF13649<br>PF13489<br>PF08241<br>PF08242<br>PF13847 | Methyltransf_25<br>Methyltransf_23<br>Methyltransf_11<br>Methyltransf_12<br>Methyltransf_31 | Methyltransferase domain<br>Methyltransferase domain<br>Methyltransferase domain<br>Methyltransferase domain<br>Methyltransferase domain | 7.70E-13<br>1.30E-10<br>9.10E-10<br>5.10E-09<br>4.30E-08 |
| WP_050590681.1 | 5838 | 6843 | + | 334 | PF00144<br>PF13354 | Beta-lactamase<br>Beta-lactamase2 | Beta-lactamase<br>Beta-lactamase enzyme family | 9.30E-48<br>4.40E-05 |
| WP_027658663.1 | 7253 | 7886 | + | 210 | TIGR01189<br>TIGR00968<br>TIGR03873<br>TIGR03265<br>TIGR01188 | TIGR01189<br>TIGR00968<br>TIGR03873<br>TIGR03265<br>TIGR01188 | ccmA: heme ABC exporter, ATP-binding protein CcmA<br>3a0106s01: sulfate ABC transporter, ATP-binding protein<br>F420-0_ABC_ATP: proposed F420-0 ABC transporter, ATP-binding protein<br>PhnT2: putative 2-aminoethylphosphonate ABC transporter, ATP-binding protein<br>drrA: daunorubicin resistance ABC transporter, ATP-binding protein | 3.00E-34<br>5.50E-31<br>3.80E-28<br>1.30E-27<br>1.40E-27 |
| WP_027658664.1 | 7887 | 9423 | + | 511 | NO PFAM MATCH | - | - | - |

| index | leader | core | start | end | dir | score |
|---|---|---|---|---|---|---|
| 1 | VLDWLHEQDDKRGRRLEYSLGWSIGGPERAAIT | ALPAKAWSPASAVTTNVHGVG | 14 | 179 | + | 9 |
| 2 | MPNIAQHANMVSETQFQGLSEESPSRKINEGEPLG | LLPVIVAAAAVTAVGAVLGIGSV | 1041 | 1218 | + | 19 |

### Results for WP_155244151.1 [Salinispora pacifica CNT851] back to top

previous - next

#### Architecture

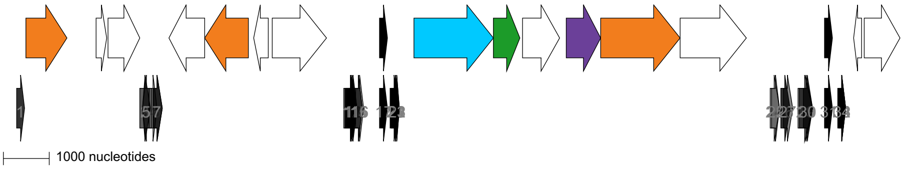

Link to nucleotide sequence

| Accession | start | end | direction | length (aa) | Pfam/HMM | name | description | E-value |
|---|---|---|---|---|---|---|---|---|
| WP_018819928.1 | 2540 | 3428 | + | 295 | TIGR03522<br>TIGR01188<br>TIGR03864<br>TIGR03265<br>TIGR02857 | TIGR03522<br>TIGR01188<br>TIGR03864<br>TIGR03265<br>TIGR02857 | GldA_ABC_ATP: gliding motility-associated ABC transporter ATP-binding subunit GldA<br>drrA: daunorubicin resistance ABC transporter, ATP-binding protein<br>PQQ_ABC_ATP: ABC transporter, ATP-binding subunit, PQQ-dependent alcohol dehydrogenase system<br>PhnT2: putative 2-aminoethylphosphonate ABC transporter, ATP-binding protein<br>CydD: thiol reductant ABC exporter, CydD subunit | 8.40E-58<br>5.00E-54<br>3.00E-45<br>1.40E-37<br>2.50E-36 |
| WP_009739003.1 | 4071 | 4311 | + | 80 | INFERRED GENE | - | - | - |
| WP_018805652.1 | 4338 | 5043 | + | 235 | INFERRED GENE | - | - | - |
| WP_080643112.1 | 6437 | 5672 | - | 254 | PF01061<br>PF12698 | ABC2_membrane<br>ABC2_membrane_3 | ABC-2 type transporter<br>ABC-2 family transporter protein | 1.70E-15<br>4.70E-07 |
| WP_050563922.1 | 7387 | 6433 | - | 317 | TIGR01188<br>TIGR03864<br>TIGR01288<br>TIGR03740<br>TIGR02673 | TIGR01188<br>TIGR03864<br>TIGR01288<br>TIGR03740<br>TIGR02673 | drrA: daunorubicin resistance ABC transporter, ATP-binding protein<br>PQQ_ABC_ATP: ABC transporter, ATP-binding subunit, PQQ-dependent alcohol dehydrogenase system<br>nodI: nodulation ABC transporter NodI<br>galliderm_ABC: lantibiotic protection ABC transporter, ATP-binding subunit<br>FtsE: cell division ATP-binding protein FtsE | 2.40E-59<br>3.80E-42<br>3.00E-40<br>2.80E-35<br>5.40E-35 |

| | | | | | | | | |
|---|---|---|---|---|---|---|---|---|
| WP_155244150.1 | 7837 | 7522 | - | 104 | NO PFAM MATCH | - | - | - |
| WP_013161536.1 | 7911 | 9112 | + | 400 | INFERRED GENE | - | - | - |
| WP_018824030.1 | 10251 | 10443 | + | 63 | LinA | LinA | LinA (linaridin precursor) | 5.90E-13 |
| WP_155244151.1 | 11020 | 12733 | + | 570 | LinH<br>LinG<br>LinE<br>PF12697<br>PF12146 | LinH<br>LinG<br>LinE<br>Abhydrolase_6<br>Hydrolase_4 | LinH (Fusion of LinE/LinG)<br>LinG Homolog<br>LinE Homolog<br>Alpha/beta hydrolase family<br>Serine aminopeptidase, S33 | 2.00E-227<br>6.60E-97<br>1.30E-96<br>2.30E-16<br>1.90E-08 |
| WP_018824032.1 | 12741 | 13317 | + | 191 | LinL | LinL | LinL | 9.40E-65 |
| WP_018824033.1 | 13375 | 14200 | + | 274 | PF12680 | SnoaL_2 | SnoaL-like domain | 8.10E-07 |
| WP_050563924.1 | 14333 | 15095 | + | 253 | PF13649<br>PF08241<br>TIGR01983<br>PF08242<br>PF13489 | Methyltransf_25<br>Methyltransf_11<br>TIGR01983<br>Methyltransf_12<br>Methyltransf_23 | Methyltransferase domain<br>Methyltransferase domain<br>UbiG: 3-demethylubiquinone-9 3-O-methyltransferase<br>Methyltransferase domain<br>Methyltransferase domain | 1.00E-10<br>5.00E-08<br>2.00E-06<br>1.10E-05<br>2.00E-05 |
| WP_018824035.1 | 15091 | 16810 | + | 572 | TIGR02204<br>TIGR02203<br>TIGR03375<br>TIGR01846<br>TIGR00958 | TIGR02204<br>TIGR02203<br>TIGR03375<br>TIGR01846<br>TIGR00958 | MsbA_rel: ABC transporter, permease/ATP-binding protein<br>MsbA_lipidA: lipid A export permease/ATP-binding protein MsbA<br>type_I_sec_LssB: type I secretion system ATPase<br>type_I_sec_HlyB: type I secretion system ATPase<br>3a01208: antigen peptide transporter 2 | 1.80E-101<br>1.60E-96<br>8.30E-83<br>3.60E-82<br>5.30E-82 |
| WP_080643113.1 | 16802 | 18251 | + | 482 | TIGR02734<br>PF13450<br>TIGR03467<br>PF01593<br>TIGR01317 | TIGR02734<br>NAD_binding_8<br>TIGR03467<br>Amino_oxidase<br>TIGR01317 | crtI_fam: phytoene desaturase<br>NAD(P)-binding Rossmann-like domain<br>HpnE: squalene-associated FAD-dependent desaturase<br>Flavin containing amine oxidoreductase<br>GOGAT_sm_gam: glutamate synthase, NADH/NADPH, small subunit | 8.60E-13<br>1.50E-10<br>2.00E-09<br>1.50E-08<br>6.40E-08 |
| WP_033671195.1 | 19951 | 20143 | + | 63 | LinA | LinA | LinA (linaridin precursor) | 8.40E-13 |
| WP_018824037.1 | 20759 | 20603 | - | 51 | NO PFAM MATCH | - | - | - |
| WP_155244152.1 | 20833 | 21604 | + | 256 | PF13276 | HTH_21 | HTH-like domain | 4.80E-05 |

| index | leader | core | start | end | dir | score |
|---|---|---|---|---|---|---|
| 1 | VTLLLTTTHFGSGRSLLLTVMGQAA | VSVVAVVALVALALARRRFDEEFTAI | 2354 | 2510 | + | 9 |
| 2 | VQAVQGDRAPWRPCTFDPRALRRPS...INPRRYRSPVARWSERRNPAKIPAR | SPPTAPAPERVPAHPRTAAKHRYHAQKRHATTT | 5019 | 5274 | + | 8 |
| 3 | VFGGGAWMSGNSLGCGGGWRTSGGD | LRGVAALGPAGDGAAVSAGVDAGRPA | 5235 | 5079 | - | 6 |
| 4 | MLKSDTPQRPNKALLGRNRCAGPALPADALGCRPGC | CPPSFAVHAVHPPRTPL | 5242 | 5404 | + | 6 |
| 5 | LLGRCGVSLLSMVAVFGGGAWMSGNSLGCGGG | WRTSGGDLRGVAALGPAGDGAAVSAGVDAGRPA | 5277 | 5079 | - | 7 |
| 6 | MHLGAGRAAARPASRSTLYTRRVRRCRGGHRSTPPRPAVA | NTAGACTTPRDRAGPVLPSTP | 5324 | 5510 | + | 8 |
| 7 | LGAGRAAARPASRSTLYTRRVRRCRGGHRSTPPRPAVA | NTAGACTTPRDRAGPVLPSTP | 5330 | 5510 | + | 8 |
| 8 | MITGRLGRIPTMSIRLSPAALMERWLPGCRLAPGVAIGGFGWPGL | ADVTGGLVLARAEATESLLDAAGGVSALECCMLAANVDTKRRESR | 9484 | 9757 | + | 7 |
| 9 | LGRIPTMSIRLSPAALMERWLPGCRLAPGVAIGGFGWPGLAD | VTGGLVLARAEATESLLDAAGGVSALECCMLAANVDTKRRESR | 9499 | 9757 | + | 8 |
| 10 | MSIRLSPAALMERWLPGCRLAPGVAIGGFGWPGLADVTG | GLVLARAEATESLLDAAGGVSALECCMLAANVDTKRRESR | 9517 | 9757 | + | 8 |
| 11 | LMERWLPGCRLAPGVAIGGFGWPGLADVTGGLVLA | RAEATESLLDAAGGVSALECCMLAANVDTKRRESR | 9544 | 9757 | + | 9 |
| 12 | MERWLPGCRLAPGVAIGGFGWPGLADVTGGLVLA | RAEATESLLDAAGGVSALECCMLAANVDTKRRESR | 9547 | 9757 | + | 9 |
| 13 | VAIGGFGWPGLADVTGGLVLARAEATES | LLDAAGGVSALECCMLAANVDTKRRESR | 9586 | 9757 | + | 9 |
| 14 | VLHVSRECRHQAPRVAVTIRL | PTPTSNGRVSGSRVDPDTNRPCIEPQIRKSVKWVIRQPN | 9705 | 9888 | + | 7 |
| 15 | LHVSRECRHQAPRVAVTIRL | PTPTSNGRVSGSRVDPDTNRPCIEPQIRKSVKWVIRQPN | 9708 | 9888 | + | 7 |
| 16 | MSTPSAESRGDYPTANSDLQRPRV | GVPCRPGYQSSMHRTPNQKVGQVGDPATELTENL | 9725 | 9902 | + | 6 |
| 17 | MASIADFMNAQLLGPIEQGMTKGNGVPVM | ATPAVAVPASVKITAAVVAFTVPVAAYVAGRAVN | 10251 | 10443 | + | 31 |
| 18 | MSRVVESIRAATVPVLTETEGDMVSPMMPVA | FTPAVLVTCPYAAGGLAAGVTIVNAYEAGRAAAG | 10487 | 10685 | + | 29 |
| 19 | VPVLTETEGDMVSPMMPVA | FTPAVLVTCPYAAGGLAAGVTIVNAYEAGRAAAG | 10523 | 10685 | + | 29 |
| 20 | VLTETEGDMVSPMMPVA | FTPAVLVTCPYAAGGLAAGVTIVNAYEAGRAAAG | 10529 | 10685 | + | 29 |
| 21 | MVSPMMPVAFTPAVLVTCPYA | AGGLAAGVTIVNAYEAGRAAAG | 10553 | 10685 | + | 17 |
| 22 | MMPVAFTPAVLVTCPYAAG | GLAAGVTIVNAYEAGRAAAG | 10565 | 10685 | + | 17 |
| 23 | MPVAFTPAVLVTCPYAAGG | LAAGVTIVNAYEAGRAAAG | 10568 | 10685 | + | 17 |
| 24 | MTAIILIAMLGTVAPAGAIPPDAPAG | PTVVLWSATHGDATASGGRWLELPNGSLSIDLVAEGEL | 18789 | 18984 | + | 6 |
| 25 | MLGTVAPAGAIPPDAPAG | PTVVLWSATHGDATASGGRWLELPNGSLSIDLVAEGEL | 18813 | 18984 | + | 6 |
| 26 | VQWWRDPAPLPWTKVASQCDVGAVL | VTTGYLPTVTGRLKVCRGENTLQDGGADTLLTTWSVTGRSLG | 19016 | 19220 | + | 9 |
| 27 | VRRRRCACDHRLPTHGDGSPESLPRREHSAGRRRGHLVDHLVGD | GTFARVIHGDRRDAPFIGM | 19069 | 19261 | + | 7 |
| 28 | VWCRRSVRRYCGCWSESSHSCQTAA...IPYPPPTRKIVYRHKGGSLRWPISA | LAPDSRLTAAVVENQRSISEALAH | 19398 | 19707 | + | 7 |

| | | | | | | |
|---|---|---|---|---|---|---|
| 29 | VRRYCGCWSESSHSCQTAAIASPSM...IPYPPPTRKIVYRHKGGSLRWPISA | LAPDSRLTAAVVENQRSISEALAH | 19416 | 19707 | + | 8 |
| 30 | MYVQARGRGSTCRNWPAIPRHQVIPYPPPTRKIVYRHKGGSLRWPISA | LAPDSRLTAAVVENQRSISEALAH | 19488 | 19707 | + | 8 |
| 31 | MASIADFMNAQRLGPIEQGVTKENGVPVM | ATPAVAVPASFAATITVVAFTVPAAAYVAGRVVN | 19951 | 20143 | + | 31 |
| 32 | MVDPRMPLGFTPAVLVTCPYAAGGMAAGA | AAVAAYEAGRAARRRLSNRHCRPPGYRPGQ | 20254 | 20434 | + | 16 |
| 33 | MPLGFTPAVLVTCPYAAGGMAAGAAAV | AAYEAGRAARRRLSNRHCRPPGYRPGQ | 20269 | 20434 | + | 15 |
| 34 | LGFTPAVLVTCPYAAGGMAAGAAAVA | AYEAGRAARRRLSNRHCRPPGYRPGQ | 20275 | 20434 | + | 15 |

### Results for WP_139140755.1 [Streptomyces abyssalis] back to top

previous - next

#### Architecture

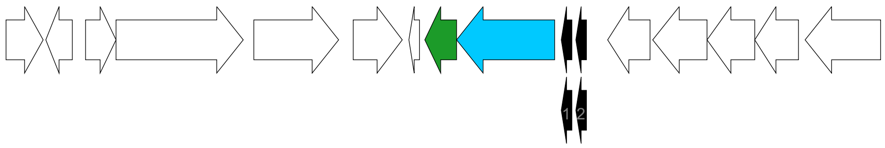

1000 nucleotides

Link to nucleotide sequence

| Accession | start | end | direction | length (aa) | Pfam/HMM | name | description | E-value |
|---|---|---|---|---|---|---|---|---|
| WP_070011776.1 | 759614 | 760313 | + | 232 | NO PFAM MATCH | - | - | - |
| WP_070011777.1 | 760894 | 760366 | - | 175 | PF13302<br>TIGR03585<br>PF13420<br>PF00583<br>PF13523 | Acetyltransf_3<br>TIGR03585<br>Acetyltransf_4<br>Acetyltransf_1<br>Acetyltransf_8 | Acetyltransferase (GNAT) domain<br>PseH: pseudaminic acid biosynthesis N-acetyl transferase<br>Acetyltransferase (GNAT) domain<br>Acetyltransferase (GNAT) family<br>Acetyltransferase (GNAT) domain | 9.60E-27<br>7.90E-17<br>3.80E-14<br>1.90E-13<br>7.40E-13 |
| WP_070011778.1 | 761136 | 761727 | + | 196 | NO PFAM MATCH | - | - | - |
| WP_079123166.1 | 761731 | 764182 | + | 816 | PF01435<br>PF13828<br>PF05569 | Peptidase_M48<br>DUF4190<br>Peptidase_M56 | Peptidase family M48<br>Domain of unknown function (DUF4190)<br>BlaR1 peptidase M56 | 2.60E-08<br>1.30E-06<br>1.50E-05 |
| WP_070011779.1 | 764382 | 765996 | + | 537 | TIGR02937<br>TIGR02950<br>TIGR02985<br>TIGR02948 | TIGR02937<br>TIGR02950<br>TIGR02985<br>TIGR02948 | sigma70-ECF: RNA polymerase sigma factor, sigma-70 family<br>SigM_subfam: RNA polymerase sigma factor, SigM family<br>Sig70_bacteroi1: RNA polymerase sigma-70 factor, Bacteroides expansion family 1<br>SigW_bacill: RNA polymerase sigma-W factor | 5.70E-08<br>8.00E-07<br>1.00E-05<br>8.80E-05 |
| WP_070011780.1 | 766296 | 767220 | + | 307 | PF00089<br>PF13365 | Trypsin<br>Trypsin_2 | Trypsin<br>Trypsin-like peptidase domain | 6.20E-51<br>1.20E-06 |
| WP_107374392.1 | 767561 | 767354 | - | 68 | NO PFAM MATCH | - | - | - |
| WP_070011781.1 | 768276 | 767658 | - | 205 | LinL | LinL | LinL | 1.90E-74 |
| WP_139140755.1 | 770153 | 768272 | - | 626 | LinH<br>LinE<br>LinG<br>PF12697<br>PF00561 | LinH<br>LinE<br>LinG<br>Abhydrolase_6<br>Abhydrolase_1 | LinH (Fusion of LinE/LinG)<br>LinE Homolog<br>LinG Homolog<br>Alpha/beta hydrolase family<br>alpha/beta hydrolase fold | 1.00E-251<br>7.10E-117<br>5.80E-107<br>1.20E-19<br>4.40E-13 |
| WP_139140756.1 | 770496 | 770289 | - | 68 | LinA<br>TIGR03789 | LinA<br>TIGR03789 | LinA (linaridin precursor)<br>pdsO: proteobacterial sortase system peptidoglycan-associated protein | 1.10E-16<br>7.30E-04 |
| WP_070011784.1 | 770759 | 770558 | - | 66 | LinA | LinA | LinA (linaridin precursor) | 7.60E-14 |
| WP_070011785.1 | 771977 | 771170 | - | 268 | PF00455<br>PF08220 | DeoRC<br>HTH_DeoR | DeoR C terminal sensor domain<br>DeoR-like helix-turn-helix domain | 2.10E-37<br>2.00E-09 |
| WP_070011786.1 | 773086 | 772030 | - | 351 | TIGR00692<br>TIGR03366<br>TIGR03989<br>TIGR03451<br>PF08240 | TIGR00692<br>TIGR03366<br>TIGR03989<br>TIGR03451<br>ADH_N | tdh: L-threonine 3-dehydrogenase<br>HpnZ_proposed: putative phosphonate catabolism associated alcohol dehydrogenase<br>Rxyl_3153: NDMA-dependent alcohol dehydrogenase, Rxyl_3153 family<br>mycoS_dep_FDH: S-(hydroxymethyl)mycothiol dehydrogenase<br>Alcohol dehydrogenase GroES-like domain | 5.40E-56<br>2.20E-35<br>3.50E-34<br>1.20E-32<br>1.00E-31 |
| WP_079123167.1 | 773994 | 773082 | - | 303 | PF00528<br>TIGR02141<br>TIGR03226<br>TIGR03262<br>TIGR01581 | BPD_transp_1<br>TIGR02141<br>TIGR03226<br>TIGR03262<br>TIGR01581 | Binding-protein-dependent transport system inner membrane component<br>modB_ABC: molybdate ABC transporter, permease protein<br>PhnU: 2-aminoethylphosphonate ABC transporter, permease protein<br>PhnU2: putative 2-aminoethylphosphonate ABC transporter, permease protein<br>Mo_ABC_porter: NifC-like ABC-type porter | 3.10E-21<br>6.60E-15<br>9.00E-14<br>4.70E-13<br>5.00E-13 |
| WP_070012084.1 | 774830 | 773990 | - | 279 | PF00528<br>TIGR02141<br>TIGR02140<br>TIGR03226<br>TIGR01581 | BPD_transp_1<br>TIGR02141<br>TIGR02140<br>TIGR03226<br>TIGR01581 | Binding-protein-dependent transport system inner membrane component<br>modB_ABC: molybdate ABC transporter, permease protein<br>permease_CysW: sulfate ABC transporter, permease protein CysW<br>PhnU: 2-aminoethylphosphonate ABC transporter, permease protein<br>Mo_ABC_porter: NifC-like ABC-type porter | 4.40E-15<br>1.50E-10<br>1.80E-09<br>8.20E-08<br>1.90E-07 |
| WP_070011787.1 | 776425 | 774958 | - | 488 | PF13416<br>PF01547 | SBP_bac_8<br>SBP_bac_1 | Bacterial extracellular solute-binding protein<br>Bacterial extracellular solute-binding protein | 3.00E-24<br>4.90E-24 |

| index | leader | core | start | end | dir | score |
|---|---|---|---|---|---|---|
| 1 | MSALLEKVNASAELDALAVQVDEGEWARPTAPVL | ATPAVIGTAVGAAGAVVGALVGGFIVADNVLGGS | 770496 | 770289 | - | 28 |
| 2 | MLLEKVNNSEELSTLPLMLDEDKGAAPTEPVL | ATPAAVAAGIAGVTLVATAVAGGIGLAVTVNGNS | 770759 | 770558 | - | 29 |

### Results for OEU87428.1 [Streptomyces abyssalis] back to top

### Architecture

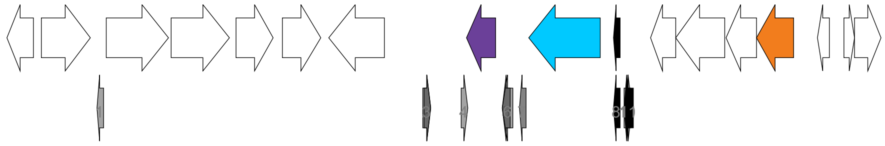

1000 nucleotides

Link to nucleotide sequence

| Accession | start | end | direction | length (aa) | Pfam/HMM | name | description | E-value |
|---|---|---|---|---|---|---|---|---|
| OEU87426.1 | 63255 | 62550 | - | 234 | PF18183 | SLATT_2 | SMODS and SLOG-associating 2TM effector domain 2 | 2.10E-87 |
| OEU86534.1 | 63453 | 64734 | + | 426 | TIGR00877<br>PF01071<br>PF02844<br>PF02843<br>TIGR02068 | TIGR00877<br>GARS_A<br>GARS_N<br>GARS_C<br>TIGR02068 | purD: phosphoribosylamine--glycine ligase<br>Phosphoribosylglycinamide synthetase, ATP-grasp (A) domain<br>Phosphoribosylglycinamide synthetase, N domain<br>Phosphoribosylglycinamide synthetase, C domain<br>cya_phycin_syn: cyanophycin synthetase | 1.50E-143<br>1.40E-64<br>3.70E-30<br>4.30E-27<br>1.30E-06 |
| OEU86535.1 | 65158 | 66778 | + | 539 | NO PFAM MATCH | - | - | - |
| OEU86536.1 | 66861 | 68388 | + | 508 | NO PFAM MATCH | - | - | - |
| OEU86537.1 | 68557 | 69511 | + | 317 | PF01259<br>TIGR00081<br>TIGR02735 | SAICAR_synt<br>TIGR00081<br>TIGR02735 | SAICAR synthetase<br>purC: phosphoribosylaminoimidazolesuccinocarboxamide synthase<br>purC_vibrio: phosphoribosylaminoimidazole-succinocarboxamide synthase | 3.30E-99<br>1.60E-69<br>7.00E-21 |
| OEU87427.1 | 69770 | 70781 | + | 336 | PF00196 | GerE | Bacterial regulatory proteins, luxR family | 4.00E-05 |
| OEU86538.1 | 72435 | 70968 | - | 488 | TIGR02734<br>TIGR03467<br>PF13450<br>TIGR00292<br>TIGR02733 | TIGR02734<br>TIGR03467<br>NAD_binding_8<br>TIGR00292<br>TIGR02733 | crtI_fam: phytoene desaturase<br>HpnE: squalene-associated FAD-dependent desaturase<br>NAD(P)-binding Rossmann-like domain<br>TIGR00292: thiazole biosynthesis enzyme<br>desat_CrtD: C-3',4' desaturase CrtD | 2.20E-09<br>3.00E-09<br>1.00E-07<br>1.80E-07<br>9.80E-07 |
| OEU86539.1 | 75363 | 74604 | - | 252 | PF13649<br>PF08241<br>PF13847<br>PF13489<br>PF08242 | Methyltransf_25<br>Methyltransf_11<br>Methyltransf_31<br>Methyltransf_23<br>Methyltransf_12 | Methyltransferase domain<br>Methyltransferase domain<br>Methyltransferase domain<br>Methyltransferase domain<br>Methyltransferase domain | 2.00E-18<br>3.90E-13<br>5.20E-12<br>1.80E-11<br>3.00E-10 |
| OEU87428.1 | 78096 | 76212 | - | 627 | LinH<br>LinG<br>LinE<br>PF12697<br>TIGR02427 | LinH<br>LinG<br>LinE<br>Abhydrolase_6<br>TIGR02427 | LinH (Fusion of LinE/LinG)<br>LinG Homolog<br>LinE Homolog<br>Alpha/beta hydrolase family<br>protocat_pcaD: 3-oxoadipate enol-lactonase | 5.40E-244<br>5.00E-121<br>3.50E-96<br>2.90E-13<br>5.00E-09 |
| OEU86540.1 | 78617 | 78437 | - | 59 | LinA | LinA | LinA (linaridin precursor) | 2.80E-16 |
| OEU87429.1 | 80075 | 79418 | - | 218 | PF00072<br>PF00196<br>TIGR02875<br>TIGR02154<br>TIGR02956 | Response_reg<br>GerE<br>TIGR02875<br>TIGR02154<br>TIGR02956 | Response regulator receiver domain<br>Bacterial regulatory proteins, luxR family<br>spore_0_A: sporulation transcription factor Spo0A<br>PhoB: phosphate regulon transcriptional regulatory protein PhoB<br>TMAO_torS: TMAO reductase sytem sensor TorS | 4.30E-25<br>1.00E-13<br>3.70E-13<br>4.80E-12<br>2.30E-10 |
| OEU87430.1 | 81358 | 80074 | - | 427 | PF07730 | HisKA_3 | Histidine kinase | 2.70E-15 |
| OEU86541.1 | 82205 | 81392 | - | 270 | PF01061<br>PF12698<br>TIGR03062<br>TIGR01247 | ABC2_membrane<br>ABC2_membrane_3<br>TIGR03062<br>TIGR01247 | ABC-2 type transporter<br>ABC-2 family transporter protein<br>pip_yhgE_Cterm: YhgE/Pip C-terminal domain<br>drrB: daunorubicin resistance ABC transporter membrane protein | 6.60E-20<br>2.40E-11<br>6.00E-08<br>6.80E-05 |
| OEU86542.1 | 83170 | 82201 | - | 322 | TIGR01188<br>TIGR03864<br>TIGR01288<br>TIGR04406<br>TIGR03522 | TIGR01188<br>TIGR03864<br>TIGR01288<br>TIGR04406<br>TIGR03522 | drrA: daunorubicin resistance ABC transporter, ATP-binding protein<br>PQQ_ABC_ATP: ABC transporter, ATP-binding subunit, PQQ-dependent alcohol dehydrogenase system<br>nodI: nodulation ABC transporter NodI<br>LPS_export_lptB: LPS export ABC transporter ATP-binding protein<br>GldA_ABC_ATP: gliding motility-associated ABC transporter ATP-binding subunit GldA | 1.10E-69<br>1.50E-49<br>4.60E-48<br>1.60E-39<br>9.60E-37 |
| OEU87431.1 | 84101 | 83783 | - | 105 | PF11774 | Lsr2 | Lsr2 | 7.70E-36 |
| OEU86543.1 | 84483 | 84768 | + | 94 | PF02700<br>TIGR00302 | PurS<br>TIGR00302 | Phosphoribosylformylglycinamidine (FGAM) synthase<br>TIGR00302: phosphoribosylformylglycinamidine synthase, purS protein | 3.40E-28<br>1.30E-20 |
| OEU86544.1 | 84764 | 85451 | + | 228 | TIGR01737<br>PF13507<br>TIGR01735<br>TIGR01857<br>PF07685 | TIGR01737<br>GATase_5<br>TIGR01735<br>TIGR01857<br>GATase_3 | FGAM_synth_I: phosphoribosylformylglycinamidine synthase I<br>CobB/CobQ-like glutamine amidotransferase domain<br>FGAM_synt: phosphoribosylformylglycinamidine synthase<br>FGAM-synthase: phosphoribosylformylglycinamidine synthase<br>CobB/CobQ-like glutamine amidotransferase domain | 9.70E-81<br>7.60E-34<br>9.70E-17<br>1.10E-15<br>2.00E-08 |

| index | leader | core | start | end | dir | score |
|---|---|---|---|---|---|---|
| 1 | VAPRAATATCAGPAGTRQGPGAVPKSLPQ | TPSGRGSPRSRQGECPGPALRPDDAGSGHP | 65099 | 64919 | - | 6 |
| 2 | VKRTERNSTASWSSESDDLTGGFGRCRILS | STPPSWGRVVSSSTRRIAARPCWNWPMARKNSTRGETM | 73446 | 73653 | + | 6 |
| 3 | LTGGFGRCRILS | STPPSWGRVVSSSTRRIAARPCWNWPMARKNSTRGETM | 73500 | 73653 | + | 6 |
| 4 | VASRQIPHGGRGGAAAPPARVCRTQMLER | PVHVGSCGVRRVLRLQGAAHAGVTSPCRW | 74450 | 74627 | + | 6 |
| 5 | VKGFQFTLVRATGFEESDPEVVFVSP | YTPMTGPGETAGGDGGASGWGPSYDSHIRDDSRGEVPAQPA | 75745 | 75541 | - | 6 |
| 6 | MLCAFLRHRIREDGRAEGVKGFQFTLVRATGFEESDPEVVFVSP | YTPMTGPGETAGGDGGASGWGPSYDSHIRDDSRGEVPAQPA | 75799 | 75541 | - | 6 |

| | | | | | | |
|---|---|---|---|---|---|---|
| 7 | VPRGRPCCSARSRRSTRTRCSTGRRRS | TSSRGSSPTGGSSRPLRLSTTTTSSTGR | 76146 | 75978 | - | 7 |
| 8 | MNDVTSHINASAELASVEITDAEGVVFTDYTPVM | GTPGAFLGGVAIASATVAAFEQGQG | 78617 | 78437 | - | 29 |
| 9 | MCQWSAHGLAGGVRARHDGRAADEGTRERGGGEDR | SRLFGDLAGVLLVELVLNLKVQLLFQHGFLRACET | 78708 | 78921 | + | 6 |
| 10 | MLEEKLNLQIQDQLHEEHAGEIAEEAAPVL | ATPALAGAFVGGAAVVTGAYAAGKAVG | 78894 | 78720 | - | 30 |
| 11 | VTTLSFLASTTFARNSQSRLTSSEE...LNLQIQDQLHEEHAGEIAEEAAPVL | ATPALAGAFVGGAAVVTGAYAAGKAVG | 78972 | 78720 | - | 29 |

### Results for OEU88179.1 [Streptomyces abyssalis] back to top

previous - next

#### Architecture

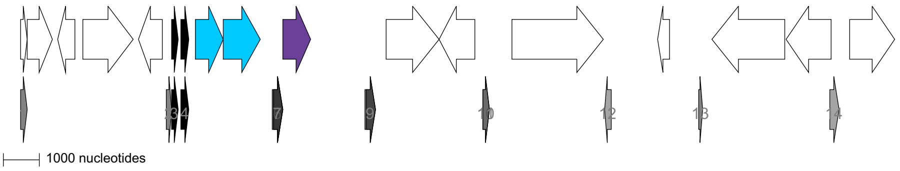

Link to nucleotide sequence

| Accession | start | end | direction | length (aa) | Pfam/HMM | name | description | E-value |
|---|---|---|---|---|---|---|---|---|
| OEU88171.1 | 847208 | 847388 | + | 59 | NO PFAM MATCH | - | - | - |
| OEU88172.1 | 847398 | 848091 | + | 230 | PF13829 | DUF4191 | Domain of unknown function (DUF4191) | 4.30E-84 |
| OEU88173.1 | 848725 | 848254 | - | 156 | PF06271 | RDD | RDD family | 5.50E-10 |
| OEU88174.1 | 848926 | 850342 | + | 471 | TIGR00653<br>PF00120<br>TIGR03105<br>PF03951 | TIGR00653<br>Gln-synt_C<br>TIGR03105<br>Gln-synt_N | GlnA: glutamine synthetase, type I<br>Glutamine synthetase, catalytic domain<br>gln_synth_III: glutamine synthetase, type III<br>Glutamine synthetase, beta-Grasp domain | 2.10E-199<br>6.10E-137<br>5.70E-71<br>1.20E-23 |
| OEU88175.1 | 851156 | 850493 | - | 220 | PF05719 | GPP34 | Golgi phosphoprotein 3 (GPP34) | 9.10E-43 |
| OEU88176.1 | 851412 | 851607 | + | 64 | LinA | LinA | LinA (linaridin precursor) | 8.40E-12 |
| OEU88177.1 | 851661 | 851874 | + | 70 | LinA | LinA | LinA (linaridin precursor) | 1.20E-15 |
| OEU88178.1 | 852079 | 852850 | + | 256 | LinH<br>LinG | LinH<br>LinG | LinH (Fusion of LinE/LinG)<br>LinG Homolog | 2.20E-114<br>7.60E-108 |
| OEU88179.1 | 852846 | 853887 | + | 346 | LinH<br>LinE<br>PF12697<br>PF12146<br>PF00561 | LinH<br>LinE<br>Abhydrolase_6<br>Hydrolase_4<br>Abhydrolase_1 | LinH (Fusion of LinE/LinG)<br>LinE Homolog<br>Alpha/beta hydrolase family<br>Serine aminopeptidase, S33<br>alpha/beta hydrolase fold | 2.10E-134<br>1.50E-121<br>1.80E-18<br>5.10E-09<br>9.10E-09 |
| OEU88180.1 | 854503 | 855274 | + | 256 | PF13649<br>PF08241<br>PF13489<br>PF08242<br>PF13847 | Methyltransf_25<br>Methyltransf_11<br>Methyltransf_23<br>Methyltransf_12<br>Methyltransf_31 | Methyltransferase domain<br>Methyltransferase domain<br>Methyltransferase domain<br>Methyltransferase domain<br>Methyltransferase domain | 3.30E-16<br>2.50E-10<br>9.90E-09<br>1.40E-08<br>1.70E-06 |
| OEU88181.1 | 857384 | 858863 | + | 492 | TIGR02734<br>TIGR03467<br>PF01593<br>PF13450<br>TIGR01317 | TIGR02734<br>TIGR03467<br>Amino_oxidase<br>NAD_binding_8<br>TIGR01317 | crtI_fam: phytoene desaturase<br>HpnE: squalene-associated FAD-dependent desaturase<br>Flavin containing amine oxidoreductase<br>NAD(P)-binding Rossmann-like domain<br>GOGAT_sm_gam: glutamate synthase, NADH/NADPH, small subunit | 4.40E-12<br>8.20E-09<br>1.80E-08<br>1.90E-08<br>4.80E-08 |
| OEU88182.1 | 859845 | 858870 | - | 324 | PF01978<br>PF00196 | TrmB<br>GerE | Sugar-specific transcriptional regulator TrmB<br>Bacterial regulatory proteins, luxR family | 2.00E-05<br>4.10E-04 |
| OEU88183.1 | 860898 | 863406 | + | 835 | PF00196<br>PF13424<br>PF13176<br>TIGR03020<br>PF13384 | GerE<br>TPR_12<br>TPR_7<br>TIGR03020<br>HTH_23 | Bacterial regulatory proteins, luxR family<br>Tetratricopeptide repeat<br>Tetratricopeptide repeat<br>EpsA: transcriptional regulator EpsA<br>Homeodomain-like domain | 1.10E-13<br>1.00E-10<br>1.40E-07<br>5.90E-07<br>1.50E-06 |
| OEU88647.1 | 865271 | 864938 | - | 110 | PF00932 | LTD | Lamin Tail Domain | 2.70E-16 |
| OEU88184.1 | 868465 | 866434 | - | 676 | NO PFAM MATCH | - | - | - |
| OEU88185.1 | 869748 | 868506 | - | 413 | PF13535 | ATP-grasp_4 | ATP-grasp domain | 6.90E-05 |
| OEU88186.1 | 870271 | 871534 | + | 420 | TIGR02155<br>PF00501<br>TIGR03335<br>TIGR01733 | TIGR02155<br>AMP-binding<br>TIGR03335<br>TIGR01733 | PA_CoA_ligase: phenylacetate-CoA ligase<br>AMP-binding enzyme<br>F390_ftsA: coenzyme F390 synthetase<br>AA-adenyl-dom: amino acid adenylation domain | 6.00E-13<br>2.10E-09<br>5.10E-04<br>8.80E-04 |

| index | leader | core | start | end | dir | score |
|---|---|---|---|---|---|---|
| 1 | MRTTLRALESFESLLMGRGHRTACRNAWT | AVLEDRRRATARTEAQRVLEAAAERRPAAT | 847208 | 847388 | + | 7 |
| 2 | LTHAIVCDAGSELPRNPITILVTGFQVAISAHSAPESGAER | ETTKARERRRHTHVA | 851252 | 851423 | + | 7 |
| 3 | MLLEKVNNSHLLEAVSMDVANVEDLSSAEPVL | STPAAAAGGALVTAGAFGAGVGLGVTQAQADG | 851412 | 851607 | + | 28 |
| 4 | MTAISIKVNASQELDGVSIHAGENEWNGRMSPVL | ATPAAVGAGLAGAAAVAGAGAAGIAVGNAAGQAAGG | 851661 | 851874 | + | 28 |
| 5 | VCGELGRLIVRKDVKVTDSAAYRLL...QFLVVLYSGHDDSEDPQYSFISPFI | RLPEPDPTDHADDADHAGAADRLEPAA | 854210 | 854507 | + | 7 |
| 6 | LIVRKDVKVTDSAAYRLLRDFVTRH...QFLVVLYSGHDDSEDPQYSFISPFI | RLPEPDPTDHADDADHAGAADRLEPAA | 854231 | 854507 | + | 6 |

| | | | | | | |
|---|---|---|---|---|---|---|
| 7 | VKVTDSAAYRLLRDFVTRHLDESDP...QFLVVLYSGHDDSEDPQYSFISPFI | RLPEPDPTDHADDADHAGAADRLEPAA | 854249 | 854507 | + | 6 |
| 8 | MPQRPTSARPWRWPVSRMSSKVCRK...GSGCASPARCCRSPRSCFWTRPPPI | WTPTRSGISGKCSTGSPGTVRSSPSRTASPR | 856783 | 857080 | + | 7 |
| 9 | MSSKVCRKDSIPIWAIRAAGSPAVSGSGCASPARCCRSPRSCFWTRPPPI | WTPTRSGISGKCSTGSPGTVRSSPSRTASPR | 856834 | 857080 | + | 7 |
| 10 | LVSDPGGLWWKVRCSHCSWARAPMRRVSPD | ATGLVVIEVHRICGGVSGGLSSATGGPPPVR | 860064 | 860250 | + | 6 |
| 11 | VSDPGGLWWKVRCSHCSWARAPMRRVSPDA | TGLVVIEVHRICGGVSGGLSSATGGPPPVR | 860067 | 860250 | + | 6 |
| 12 | VHTPFAGRRLERQPIFRRTSGQGLPGRQRLLGRRPGPVPSAAC | GAPVPCVLSEHTARGTET | 863647 | 863461 | - | 6 |
| 13 | MSLFGTCSQASG | GTAVLAGAVRDVPASGRVIAASPVGR | 866069 | 866186 | + | 7 |
| 14 | VAWTVTPSACALVDGVAGIPYPART...AQAPDSRRGVRPSPDLRRDELLARH | GLPRLVPLRSREVPVAVRTITPP | 869741 | 869975 | + | 6 |

### Results for WP_070011838.1 [Streptomyces abyssalis] back to top

previous - next

#### Architecture

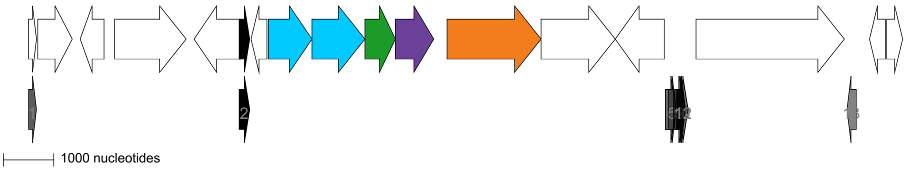

Link to nucleotide sequence

| Accession | start | end | direction | length (aa) | Pfam/HMM | name | description | E-value |
|---|---|---|---|---|---|---|---|---|
| WP_070011830.1 | 847208 | 847388 | + | 59 | NO PFAM MATCH | - | - | - |
| WP_070011831.1 | 847398 | 848091 | + | 230 | PF13829 | DUF4191 | Domain of unknown function (DUF4191) | 4.30E-84 |
| WP_070011832.1 | 848725 | 848254 | - | 156 | PF06271 | RDD | RDD family | 5.50E-10 |
| WP_070011833.1 | 848926 | 850342 | + | 471 | TIGR00653<br>PF00120<br>TIGR03105<br>PF03951 | TIGR00653<br>Gln-synt_C<br>TIGR03105<br>Gln-synt_N | GlnA: glutamine synthetase, type I<br>Glutamine synthetase, catalytic domain<br>gln_synth_III: glutamine synthetase, type III<br>Glutamine synthetase, beta-Grasp domain | 2.10E-199<br>6.10E-137<br>5.70E-71<br>1.20E-23 |
| WP_079123177.1 | 851414 | 850493 | - | 306 | PF05719 | GPP34 | Golgi phosphoprotein 3 (GPP34) | 2.50E-42 |
| WP_070011835.1 | 851412 | 851607 | + | 64 | LinA | LinA | LinA (linaridin precursor) | 8.40E-12 |
| WP_107374393.1 | 851939 | 851648 | - | 96 | NO PFAM MATCH | - | - | - |
| WP_139140757.1 | 851977 | 852850 | + | 290 | LinH<br>LinG | LinH<br>LinG | LinH (Fusion of LinE/LinG)<br>LinG Homolog | 2.50E-120<br>7.10E-114 |
| WP_070011838.1 | 852846 | 853887 | + | 346 | LinH<br>LinE<br>PF12697<br>PF12146<br>PF00561 | LinH<br>LinE<br>Abhydrolase_6<br>Hydrolase_4<br>Abhydrolase_1 | LinH (Fusion of LinE/LinG)<br>LinE Homolog<br>Alpha/beta hydrolase family<br>Serine aminopeptidase, S33<br>alpha/beta hydrolase fold | 2.10E-134<br>1.50E-121<br>1.80E-18<br>5.10E-09<br>9.10E-09 |
| WP_070011839.1 | 853886 | 854507 | + | 206 | LinL | LinL | LinL | 1.10E-72 |
| WP_070011840.1 | 854503 | 855274 | + | 256 | PF13649<br>PF08241<br>PF13489<br>PF08242<br>PF13847 | Methyltransf_25<br>Methyltransf_11<br>Methyltransf_23<br>Methyltransf_12<br>Methyltransf_31 | Methyltransferase domain<br>Methyltransferase domain<br>Methyltransferase domain<br>Methyltransferase domain<br>Methyltransferase domain | 3.30E-16<br>2.50E-10<br>9.90E-09<br>1.40E-08<br>1.70E-06 |
| WP_079123266.1 | 855528 | 857388 | + | 619 | TIGR02204<br>TIGR02203<br>TIGR00958<br>TIGR03796<br>TIGR03375 | TIGR02204<br>TIGR02203<br>TIGR00958<br>TIGR03796<br>TIGR03375 | MsbA_rel: ABC transporter, permease/ATP-binding protein<br>MsbA_lipidA: lipid A export permease/ATP-binding protein MsbA<br>3a01208: antigen peptide transporter 2<br>NHLM_micro_ABC1: NHLM bacteriocin system ABC transporter, peptidase/ATP-binding protein<br>type_I_sec_LssB: type I secretion system ATPase | 1.80E-101<br>1.90E-100<br>1.90E-90<br>4.10E-84<br>1.10E-80 |
| WP_070011841.1 | 857384 | 858863 | + | 492 | TIGR02734<br>TIGR03467<br>PF01593<br>PF13450<br>TIGR01317 | TIGR02734<br>TIGR03467<br>Amino_oxidase<br>NAD_binding_8<br>TIGR01317 | crtI_fam: phytoene desaturase<br>HpnE: squalene-associated FAD-dependent desaturase<br>Flavin containing amine oxidoreductase<br>NAD(P)-binding Rossmann-like domain<br>GOGAT_sm_gam: glutamate synthase, NADH/NADPH, small subunit | 4.40E-12<br>8.20E-09<br>1.80E-08<br>1.90E-08<br>4.80E-08 |
| WP_070011842.1 | 859845 | 858870 | - | 324 | PF01978<br>PF00196 | TrmB<br>GerE | Sugar-specific transcriptional regulator TrmB<br>Bacterial regulatory proteins, luxR family | 2.00E-05<br>4.10E-04 |
| WP_107374394.1 | 860460 | 863406 | + | 981 | PF00196<br>PF13424<br>PF13176<br>PF13384<br>TIGR03020 | GerE<br>TPR_12<br>TPR_7<br>HTH_23<br>TIGR03020 | Bacterial regulatory proteins, luxR family<br>Tetratricopeptide repeat<br>Tetratricopeptide repeat<br>Homeodomain-like domain<br>EpsA: transcriptional regulator EpsA | 1.40E-13<br>3.20E-09<br>2.90E-07<br>4.20E-07<br>6.90E-07 |
| WP_139140758.1 | 864224 | 863903 | - | 106 | NO PFAM MATCH | - | - | - |
| WP_139140759.1 | 864260 | 864554 | + | 97 | NO PFAM MATCH | - | - | - |

| index | leader | core | start | end | dir | score |
|---|---|---|---|---|---|---|
| 1 | MRTTLRALESFESLLMGRGHRTACRNAWT | AVLEDRRRATARTEAQRVLEAAAERRPAAT | 847208 | 847388 | + | 8 |
| 2 | MLLEKVNNSHLLEAVSMDVANVEDLSSAEPVL | STPAAAAGGALVTAGAFGAGVGLGVTQAQADG | 851412 | 851607 | + | 29 |
| 3 | LVSNPRPRCSWVGWGWRPALMRAGRRGGVGA | GTLISTSRMGVGFRAPHSRARFTWTGHTAHRCRPSVNSLGSW | 859849 | 860071 | + | 6 |

| | | | | | | |
|---|---|---|---|---|---|---|
| 4 | VSNPRPRCSWVGWGWRPALMRAGRRGGVGA | GTLISTSRMGVGFRAPHSRARFTWTGHTAHRCRPSVNSLGSW | 859852 | 860071 | + | 6 |
| 5 | MRAGRRGGVGA | GTLISTSRMGVGFRAPHSRARFTWTGHTAHRCRPSVNSLGSW | 859909 | 860071 | + | 6 |
| 6 | VDRPHRSPVPAVRELSWQLVSDPGGLWWKVRCSHCSWAR | APMRRVSPDATGLVVIEVHRICGGVSGGLSSATGGPPPVR | 860010 | 860250 | + | 6 |
| 7 | VPAVRELSWQLVSDPGGLWWKVRCSHCSWARAPMR | RVSPDATGLVVIEVHRICGGVSGGLSSATGGPPPVR | 860034 | 860250 | + | 6 |
| 8 | LVSDPGGLWWKVRCSHCSWARAPMRRVSPD | ATGLVVIEVHRICGGVSGGLSSATGGPPPVR | 860064 | 860250 | + | 7 |
| 9 | VSDPGGLWWKVRCSHCSWARAPMRRVSPDA | TGLVVIEVHRICGGVSGGLSSATGGPPPVR | 860067 | 860250 | + | 7 |
| 10 | VRCSHCSWARAPMRRVSPDATGLVV | IEVHRICGGVSGGLSSATGGPPPVR | 860097 | 860250 | + | 6 |
| 11 | VLALLVGPRAHASGIARRDRPGRYRGSQNLWRRFRWAVFGNWRSAPSP | VTPNAFDLLAALVRPLCRGSE | 860102 | 860312 | + | 6 |
| 12 | VGPRAHASGIARRDRPGRYRGSQNLWRRFRWAVFGNWRSAPSP | VTPNAFDLLAALVRPLCRGSE | 860117 | 860312 | + | 6 |
| 13 | VHTPFAGRRLERQPIFRRTSGQGLPGRQRLLGRRPGPVPSAAC | GAPVPCVLSEHTARGTET | 863647 | 863461 | - | 7 |

### Results for WP_151770911.1 [Streptomyces abyssomicinicus] back to top

previous - next

#### Architecture

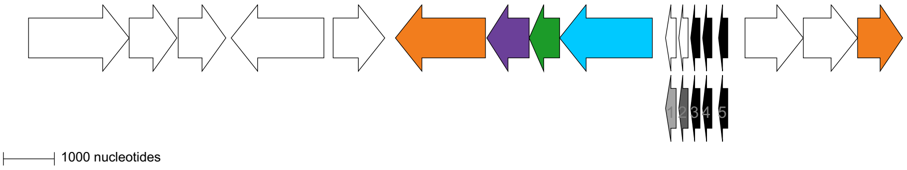

Link to nucleotide sequence

| Accession | start | end | direction | length (aa) | Pfam/HMM | name | description | E-value |
|---|---|---|---|---|---|---|---|---|
| WP_151770064.1 | 776569 | 778558 | + | 662 | TIGR01217<br>TIGR02188<br>TIGR02316<br>PF00501<br>TIGR01733 | TIGR01217<br>TIGR02188<br>TIGR02316<br>AMP-binding<br>TIGR01733 | ac_ac_CoA_syn: acetoacetate-CoA ligase<br>Ac_CoA_lig_AcsA: acetate--CoA ligase<br>propion_prpE: propionate--CoA ligase<br>AMP-binding enzyme<br>AA-adenyl-dom: amino acid adenylation domain | 4.00E-219<br>6.70E-74<br>1.40E-54<br>4.90E-52<br>8.20E-29 |
| WP_151770065.1 | 778554 | 779490 | + | 311 | PF02737<br>TIGR02279<br>TIGR02437<br>PF00725<br>TIGR02440 | 3HCDH_N<br>TIGR02279<br>TIGR02437<br>3HCDH<br>TIGR02440 | 3-hydroxyacyl-CoA dehydrogenase, NAD binding domain<br>PaaC-3OHAcCoADH: 3-hydroxyacyl-CoA dehydrogenase PaaC<br>FadB: fatty oxidation complex, alpha subunit FadB<br>3-hydroxyacyl-CoA dehydrogenase, C-terminal domain<br>FadJ: fatty oxidation complex, alpha subunit FadJ | 1.60E-41<br>6.80E-25<br>3.90E-13<br>4.70E-13<br>4.50E-10 |
| WP_151770066.1 | 779524 | 780448 | + | 307 | PF04909 | Amidohydro_2 | Amidohydrolase | 3.90E-53 |
| WP_151770067.1 | 782376 | 780543 | - | 610 | PF01078 | Mg_chelatase | Magnesium chelatase, subunit ChlI | 4.70E-04 |
| WP_151770068.1 | 782560 | 783577 | + | 338 | NO PFAM MATCH | - | - | - |
| WP_151770910.1 | 785556 | 783777 | - | 592 | TIGR02204<br>TIGR02203<br>TIGR00958<br>TIGR01846<br>TIGR03796 | TIGR02204<br>TIGR02203<br>TIGR00958<br>TIGR01846<br>TIGR03796 | MsbA_rel: ABC transporter, permease/ATP-binding protein<br>MsbA_lipidA: lipid A export permease/ATP-binding protein MsbA<br>3a01208: antigen peptide transporter 2<br>type_I_sec_HlyB: type I secretion system ATPase<br>NHLM_micro_ABC1: NHLM bacteriocin system ABC transporter, peptidase/ATP-binding protein | 5.40E-109<br>1.90E-98<br>1.20E-88<br>1.30E-83<br>7.90E-81 |
| WP_151770069.1 | 786398 | 785582 | - | 271 | PF13649<br>PF08241<br>PF13489<br>PF13847<br>PF08242 | Methyltransf_25<br>Methyltransf_11<br>Methyltransf_23<br>Methyltransf_31<br>Methyltransf_12 | Methyltransferase domain<br>Methyltransferase domain<br>Methyltransferase domain<br>Methyltransferase domain<br>Methyltransferase domain | 8.70E-16<br>2.10E-11<br>7.90E-09<br>2.10E-07<br>1.30E-06 |
| WP_151770070.1 | 787006 | 786394 | - | 203 | LinL | LinL | LinL | 3.20E-60 |
| WP_151770911.1 | 788817 | 787002 | - | 604 | LinH<br>LinG<br>LinE<br>PF12697<br>LinL | LinH<br>LinG<br>LinE<br>Abhydrolase_6<br>LinL | LinH (Fusion of LinE/LinG)<br>LinG Homolog<br>LinE Homolog<br>Alpha/beta hydrolase family<br>LinL | 1.70E-235<br>1.10E-112<br>1.60E-92<br>9.90E-15<br>8.80E-13 |
| WP_151770071.1 | 789284 | 789089 | - | 64 | NO PFAM MATCH | - | - | - |
| WP_151770072.1 | 789529 | 789340 | - | 62 | NO PFAM MATCH | - | - | - |
| WP_151770073.1 | 789763 | 789568 | - | 64 | LinA | LinA | LinA (linaridin precursor) | 6.20E-07 |
| WP_164540343.1 | 789986 | 789809 | - | 58 | LinA | LinA | LinA (linaridin precursor) | 5.00E-09 |
| WP_151770074.1 | 790293 | 790110 | - | 60 | LinA | LinA | LinA (linaridin precursor) | 2.80E-10 |
| WP_151770075.1 | 790632 | 791790 | + | 385 | PF01497<br>TIGR04281<br>TIGR03659<br>TIGR03868 | Peripla_BP_2<br>TIGR04281<br>TIGR03659<br>TIGR03868 | Periplasmic binding protein<br>peripla_PGF_1: putative ABC transporter PGF-CTERM-modified substrate-binding protein<br>IsdE: heme ABC transporter, heme-binding protein isdE<br>F420-O_ABCperi: proposed F420-0 ABC transporter, periplasmic F420-0 binding protein | 1.30E-22<br>6.30E-22<br>2.00E-14<br>2.10E-06 |
| WP_151770076.1 | 791770 | 792853 | + | 360 | PF01032<br>TIGR03869 | FecCD<br>TIGR03869 | FecCD transport family<br>F420-0_ABCperm: proposed F420-0 ABC transporter, permease protein | 9.60E-84<br>3.60E-51 |
| WP_151770077.1 | 792849 | 793725 | + | 291 | TIGR03873<br>TIGR03771<br>TIGR02315<br>TIGR01166<br>TIGR01189 | TIGR03873<br>TIGR03771<br>TIGR02315<br>TIGR01166<br>TIGR01189 | F420-0_ABC_ATP: proposed F420-0 ABC transporter, ATP-binding protein<br>anch_rpt_ABC: anchored repeat-type ABC transporter, ATP-binding subunit<br>ABC_phnC: phosphonate ABC transporter, ATP-binding protein<br>cbiO: cobalt ABC transporter, ATP-binding protein<br>ccmA: heme ABC exporter, ATP-binding protein CcmA | 8.80E-48<br>5.20E-42<br>2.30E-32<br>4.60E-30<br>4.60E-30 |

| index | leader | core | start | end | dir | score |
|---|---|---|---|---|---|---|
| 1 | MSTVPVPLPVPRALRARQELSANDPGHVPDTGAPVA | SAIVVTPSATGLPYGCYIGAAVGSKDPQ | 789284 | 789089 | - | 6 |
| 2 | MHDMLSPGAPRTDRGLREDDPGYVSKAIAPV | ASSVDVATSLVGLVAFGVAEVTQAIGSRDPR | 789529 | 789340 | - | 8 |
| 3 | MDNVSFLHRPGALTDIRADDPDGRVSPTHLPVA | STPAVVVGVAIGYALLNAFQAGRDGGPIDPR | 789763 | 789568 | - | 30 |
| 4 | MNNHRLPGTRHIDRELRESDRGHVAYATAPVT | ATPALFVAGIAVGFALAEAIGEKQPQ | 789986 | 789809 | - | 18 |
| 5 | MHQVGRPSWAGISGPLDEGAQGRVNASAAPVA | ATPVIVSVAAAVVVAYAAGALVGGHETQ | 790293 | 790110 | - | 30 |

### Results for WP_070012848.1 [Streptomyces abyssalis] back to top

previous - next

#### Architecture

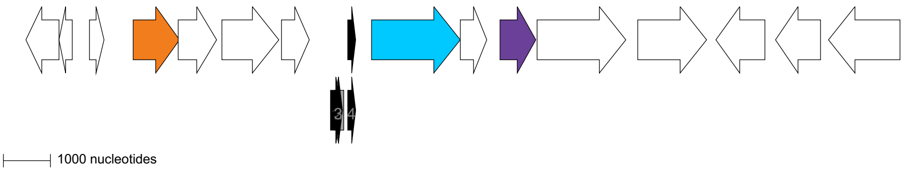

Link to nucleotide sequence

| Accession | start | end | direction | length (aa) | Pfam/HMM | name | description | E-value |
|---|---|---|---|---|---|---|---|---|
| WP_070012854.1 | 804740 | 804053 | - | 228 | TIGR01737<br>PF13507<br>TIGR01735<br>TIGR01857<br>PF07685 | TIGR01737<br>GATase_5<br>TIGR01735<br>TIGR01857<br>GATase_3 | FGAM_synth_I: phosphoribosylformylglycinamidine synthase I<br>CobB/CobQ-like glutamine amidotransferase domain<br>FGAM_synt: phosphoribosylformylglycinamidine synthase<br>FGAM-synthase: phosphoribosylformylglycinamidine synthase<br>CobB/CobQ-like glutamine amidotransferase domain | 9.70E-81<br>7.60E-34<br>9.70E-17<br>1.10E-15<br>2.00E-08 |
| WP_070012853.1 | 805021 | 804736 | - | 94 | PF02700<br>TIGR00302 | PurS<br>TIGR00302 | Phosphoribosylformylglycinamidine (FGAM) synthase<br>TIGR00302: phosphoribosylformylglycinamidine synthase, purS protein | 3.40E-28<br>1.30E-20 |
| WP_070013817.1 | 805403 | 805721 | + | 105 | PF11774 | Lsr2 | Lsr2 | 7.70E-36 |
| WP_070012852.1 | 806334 | 807303 | + | 322 | TIGR01188<br>TIGR03864<br>TIGR01288<br>TIGR04406<br>TIGR03522 | TIGR01188<br>TIGR03864<br>TIGR01288<br>TIGR04406<br>TIGR03522 | drrA: daunorubicin resistance ABC transporter, ATP-binding protein<br>PQQ_ABC_ATP: ABC transporter, ATP-binding subunit, PQQ-dependent alcohol dehydrogenase system<br>nodI: nodulation ABC transporter NodI<br>LPS_export_lptB: LPS export ABC transporter ATP-binding protein<br>GldA_ABC_ATP: gliding motility-associated ABC transporter ATP-binding subunit GldA | 1.10E-69<br>1.50E-49<br>4.60E-48<br>1.60E-39<br>9.60E-37 |
| WP_070012851.1 | 807299 | 808112 | + | 270 | PF01061<br>PF12698<br>TIGR03062<br>TIGR01247 | ABC2_membrane<br>ABC2_membrane_3<br>TIGR03062<br>TIGR01247 | ABC-2 type transporter<br>ABC-2 family transporter protein<br>pip_yhgE_Cterm: YhgE/Pip C-terminal domain<br>drrB: daunorubicin resistance ABC transporter membrane protein | 6.60E-20<br>2.40E-11<br>6.00E-08<br>6.80E-05 |
| WP_079123006.1 | 808227 | 809430 | + | 400 | PF07730 | HisKA_3 | Histidine kinase | 2.40E-15 |
| WP_107374361.1 | 809483 | 810086 | + | 200 | PF00072<br>PF00196<br>TIGR02875<br>TIGR02154<br>TIGR02956 | Response_reg<br>GerE<br>TIGR02875<br>TIGR02154<br>TIGR02956 | Response regulator receiver domain<br>Bacterial regulatory proteins, luxR family<br>spore_0_A: sporulation transcription factor Spo0A<br>PhoB: phosphate regulon transcriptional regulatory protein PhoB<br>TMAO_torS: TMAO reductase sytem sensor TorS | 3.40E-25<br>8.40E-14<br>9.90E-13<br>4.40E-12<br>2.70E-10 |
| WP_070012849.1 | 810887 | 811067 | + | 59 | LinA | LinA | LinA (linaridin precursor) | 2.80E-16 |
| WP_070012848.1 | 811387 | 813292 | + | 634 | LinH<br>LinG<br>LinE<br>PF12697<br>TIGR02427 | LinH<br>LinG<br>LinE<br>Abhydrolase_6<br>TIGR02427 | LinH (Fusion of LinE/LinG)<br>LinG Homolog<br>LinE Homolog<br>Alpha/beta hydrolase family<br>protocat_pcaD: 3-oxoadipate enol-lactonase | 2.40E-244<br>2.40E-121<br>3.70E-96<br>2.60E-13<br>5.10E-09 |
| WP_019435954.1 | 813288 | 813849 | + | 187 | INFERRED GENE | - | - | - |
| WP_070012846.1 | 814141 | 814900 | + | 252 | PF13649<br>PF08241<br>PF13847<br>PF13489<br>PF08242 | Methyltransf_25<br>Methyltransf_11<br>Methyltransf_31<br>Methyltransf_23<br>Methyltransf_12 | Methyltransferase domain<br>Methyltransferase domain<br>Methyltransferase domain<br>Methyltransferase domain<br>Methyltransferase domain | 2.00E-18<br>3.90E-13<br>5.20E-12<br>1.80E-11<br>3.00E-10 |
| WP_003991974.1 | 814907 | 816815 | + | 636 | INFERRED GENE | - | - | - |
| WP_070012844.1 | 817069 | 818536 | + | 488 | TIGR02734<br>TIGR03467<br>PF13450<br>TIGR00292<br>TIGR02733 | TIGR02734<br>TIGR03467<br>NAD_binding_8<br>TIGR00292<br>TIGR02733 | crtI_fam: phytoene desaturase<br>HpnE: squalene-associated FAD-dependent desaturase<br>NAD(P)-binding Rossmann-like domain<br>TIGR00292: thiazole biosynthesis enzyme<br>desat_CrtD: C-3',4' desaturase CrtD | 2.20E-09<br>3.00E-09<br>1.00E-07<br>1.80E-07<br>9.80E-07 |
| WP_079122885.1 | 819749 | 818723 | - | 341 | PF00196 | GerE | Bacterial regulatory proteins, luxR family | 4.10E-05 |
| WP_070012843.1 | 820947 | 819993 | - | 317 | PF01259<br>TIGR00081<br>TIGR02735 | SAICAR_synt<br>TIGR00081<br>TIGR02735 | SAICAR synthetase<br>purC: phosphoribosylaminoimidazolesuccinocarboxamide synthase<br>purC_vibrio: phosphoribosylaminoimidazole-succinocarboxamide synthase | 3.30E-99<br>1.60E-69<br>7.00E-21 |
| WP_070012842.1 | 822643 | 821116 | - | 508 | NO PFAM MATCH | - | - | - |

| index | leader | core | start | end | dir | score |
|---|---|---|---|---|---|---|

| | | | | | | |
|---|---|---|---|---|---|---|
| 1 | VTTLSFLASTTFARNSQSRLTSSEE...LNLQIQDQLHEEHAGEIAEEAAPVL | ATPALAGAFVGGAAVVTGAYAAGKAVG | 810532 | 810784 | + | 29 |
| 2 | MLEEKLNLQIQDQLHEEHAGEIAEEAAPVL | ATPALAGAFVGGAAVVTGAYAAGKAVG | 810610 | 810784 | + | 30 |
| 3 | MCQWSAHGLAGGVRARHDGRAADEGTRERGGGEDR | SRLFGDLAGVLLVELVLNLKVQLLFQHGFLRACET | 810796 | 810583 | - | 6 |
| 4 | MNDVTSHINASAELASVEITDAEGVVFTDYTPVM | GTPGAFLGGVAIASATVAAFEQGQG | 810887 | 811067 | + | 29 |

### Results for AOE46838.1 [Streptomyces antibioticus] back to top

previous - next

#### Architecture

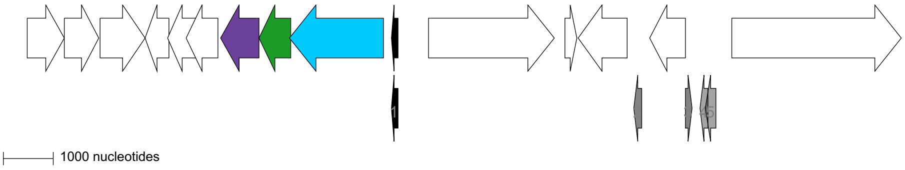

Link to nucleotide sequence

| Accession | start | end | direction | length (aa) | Pfam/HMM | name | description | E-value |
|---|---|---|---|---|---|---|---|---|
| AKA87338.1 | 22022 | 22766 | + | 247 | PF01259<br>TIGR00081 | SAICAR_synt<br>TIGR00081 | SAICAR synthetase<br>purC: phosphoribosylaminoimidazolesuccinocarboxamide synthase | 7.40E-12<br>6.60E-11 |
| AKA87339.1 | 22752 | 23457 | + | 234 | PF00106<br>PF13561<br>TIGR04316<br>TIGR01963<br>TIGR03971 | adh_short<br>adh_short_C2<br>TIGR04316<br>TIGR01963<br>TIGR03971 | short chain dehydrogenase<br>Enoyl-(Acyl carrier protein) reductase<br>dhbA_paeA: 2,3-dihydro-2,3-dihydroxybenzoate dehydrogenase<br>PHB_DH: 3-hydroxybutyrate dehydrogenase<br>SDR_subfam_1: SDR family mycofactocin-dependent oxidoreductase | 1.70E-32<br>1.20E-26<br>2.90E-25<br>2.60E-23<br>3.30E-23 |
| AKA87340.1 | 23466 | 24375 | + | 302 | TIGR00070<br>PF01634<br>TIGR03455 | TIGR00070<br>HisG<br>TIGR03455 | hisG: ATP phosphoribosyltransferase<br>ATP phosphoribosyltransferase<br>HisG_C-term: ATP phosphoribosyltransferase, C-terminal domain | 4.30E-49<br>4.10E-36<br>1.10E-05 |
| AOE46833.1 | 24864 | 24378 | - | 161 | NO PFAM MATCH | - | - | - |
| AOE46834.1 | 25402 | 24832 | - | 189 | NO PFAM MATCH | - | - | - |
| AOE46835.1 | 25850 | 25208 | - | 213 | TIGR00292<br>TIGR03467<br>PF13450<br>TIGR01317<br>TIGR01318 | TIGR00292<br>TIGR03467<br>NAD_binding_8<br>TIGR01317<br>TIGR01318 | TIGR00292: thiazole biosynthesis enzyme<br>HpnE: squalene-associated FAD-dependent desaturase<br>NAD(P)-binding Rossmann-like domain<br>GOGAT_sm_gam: glutamate synthase, NADH/NADPH, small subunit<br>gltD_gamma_fam: glutamate synthase, small subunit | 3.50E-07<br>3.80E-07<br>6.90E-07<br>9.60E-07<br>1.00E-05 |
| AOE46836.1 | 26655 | 25890 | - | 254 | PF13649<br>PF08241<br>PF13847<br>PF13489<br>PF08242 | Methyltransf_25<br>Methyltransf_11<br>Methyltransf_31<br>Methyltransf_23<br>Methyltransf_12 | Methyltransferase domain<br>Methyltransferase domain<br>Methyltransferase domain<br>Methyltransferase domain<br>Methyltransferase domain | 1.60E-16<br>1.90E-10<br>1.40E-09<br>6.00E-09<br>1.20E-08 |
| AOE46837.1 | 27287 | 26651 | - | 211 | LinL | LinL | LinL | 2.80E-69 |
| AOE46838.1 | 29164 | 27283 | - | 626 | LinH<br>LinG<br>LinE<br>PF12697<br>LinL | LinH<br>LinG<br>LinE<br>Abhydrolase_6<br>LinL | LinH (Fusion of LinE/LinG)<br>LinG Homolog<br>LinE Homolog<br>Alpha/beta hydrolase family<br>LinL | 1.40E-242<br>2.90E-113<br>3.00E-96<br>4.70E-17<br>7.60E-10 |
| AOE46839.1 | 29457 | 29304 | - | 50 | LinA | LinA | LinA (linaridin precursor) | 1.00E-08 |
| AOE46840.1 | 30060 | 32592 | + | 843 | PF07592<br>PF13613<br>PF13565 | DDE_Tnp_ISAZ013<br>HTH_Tnp_4<br>HTH_32 | Rhodopirellula transposase DDE domain<br>Helix-turn-helix of DDE superfamily endonuclease<br>Homeodomain-like domain | 3.60E-128<br>6.10E-05<br>4.60E-04 |
| AOE46841.1 | 32803 | 33028 | + | 74 | NO PFAM MATCH | - | - | - |
| AOE46842.1 | 34041 | 33054 | - | 328 | TIGR02779<br>TIGR02776<br>TIGR04120<br>PF01068<br>TIGR00574 | TIGR02779<br>TIGR02776<br>TIGR04120<br>DNA_ligase_A_M<br>TIGR00574 | NHEJ_ligase_lig: DNA ligase D, ligase domain<br>NHEJ_ligase_prk: DNA ligase D<br>DNA_lig_bact: DNA ligase, ATP-dependent, PP_1105 family<br>ATP dependent DNA ligase domain<br>dnl1: DNA ligase I, ATP-dependent (dnl1) | 8.70E-44<br>7.90E-37<br>3.10E-29<br>2.40E-23<br>6.90E-15 |
| AOE46843.1 | 35203 | 34489 | - | 237 | PF03009 | GDPD | Glycerophosphoryl diester phosphodiesterase family | 2.00E-33 |
| AOE46844.1 | 36131 | 39550 | + | 1139 | PF13385 | Laminin_G_3 | Concanavalin A-like lectin/glucanases superfamily | 2.20E-29 |

| index | leader | core | start | end | dir | score |
|---|---|---|---|---|---|---|
| 1 | MNDITAFINEDARLDEVPIERAEGTVLVEQGVTVL | ATPSANFCLTFAIGK | 29457 | 29304 | - | 27 |
| 2 | VRPPHRHTRPGYQRSGGGSCAWPRV | TTSVRSPPLAPGRNPFPRSTGGGRT | 34326 | 34173 | - | 7 |
| 3 | VHRDAAAPAVVSAAAAVSCAQGGEQAVIT | VTADATRVAIRTDLRFVTGQR | 35204 | 35357 | + | 7 |
| 4 | VSPPSRVTPPVSGPEGKGEFARRGR | APLPRRRAPCAGRGNSTGARPAPPAP | 35665 | 35509 | - | 6 |
| 5 | VPEASPEQAHAARRPDREASPRHRAHVAARSS | PDSELRQVGVRADAGVAGRGVATLARHTPCFWS | 35819 | 35621 | - | 6 |

### Results for AXK34881.1 [Streptomyces armeniacus] back to top

previous - next

#### Architecture

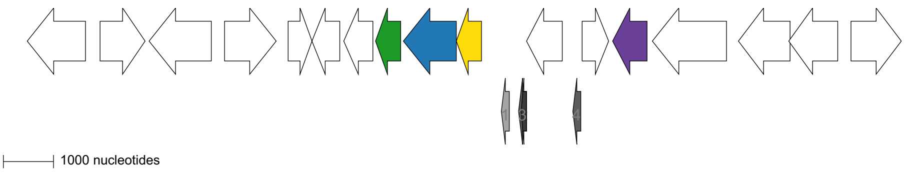

Link to nucleotide sequence

| Accession | start | end | direction | length (aa) | Pfam/HMM | name | description | E-value |
|---|---|---|---|---|---|---|---|---|
| AXK34873.1 | 4943598 | 4942437 | - | 386 | PF13556 | HTH_30 | PucR C-terminal helix-turn-helix domain | 2.80E-15 |
| AXK34874.1 | 4943877 | 4944780 | + | 300 | TIGR01426<br>PF00201<br>PF04101<br>TIGR00661 | TIGR01426<br>UDPGT<br>Glyco_tran_28_C<br>TIGR00661 | MGT: glycosyltransferase, MGT family<br>UDP-glucoronosyl and UDP-glucosyl transferase<br>Glycosyltransferase family 28 C-terminal domain<br>MJ1255: conserved hypothetical protein | 5.30E-40<br>2.80E-10<br>1.70E-04<br>5.20E-04 |
| AXK34875.1 | 4946105 | 4944863 | - | 413 | PF08924 | DUF1906 | Domain of unknown function (DUF1906) | 3.80E-44 |
| AXK34876.1 | 4946369 | 4947380 | + | 336 | PF12840<br>PF01022<br>PF12802<br>PF13412 | HTH_20<br>HTH_5<br>MarR_2<br>HTH_24 | Helix-turn-helix domain<br>Bacterial regulatory protein, arsR family<br>MarR family<br>Winged helix-turn-helix DNA-binding | 1.00E-10<br>9.80E-08<br>1.80E-07<br>6.90E-05 |
| AXK34877.1 | 4947630 | 4948101 | + | 156 | NO PFAM MATCH | - | - | - |
| AXK34878.1 | 4948666 | 4948111 | - | 184 | PF07853 | DUF1648 | Protein of unknown function (DUF1648) | 1.20E-10 |
| AXK34879.1 | 4949330 | 4948736 | - | 197 | NO PFAM MATCH | - | - | - |
| AXK34880.1 | 4949886 | 4949376 | - | 169 | LinL | LinL | LinL | 3.40E-55 |
| AXK34881.1 | 4950992 | 4949930 | - | 353 | LinE<br>LinH<br>PF12697<br>PF12146<br>TIGR03611 | LinE<br>LinH<br>Abhydrolase_6<br>Hydrolase_4<br>TIGR03611 | LinE Homolog<br>LinH (Fusion of LinE/LinG)<br>Alpha/beta hydrolase family<br>Serine aminopeptidase, S33<br>RutD: pyrimidine utilization protein D | 9.70E-93<br>6.20E-69<br>7.60E-14<br>1.70E-08<br>6.00E-08 |
| AXK34882.1 | 4951495 | 4950988 | - | 168 | LinG<br>LinH<br>PF05090 | LinG<br>LinH<br>VKG_Carbox | LinG Homolog<br>LinH (Fusion of LinE/LinG)<br>Vitamin K-dependent gamma-carboxylase | 4.20E-67<br>4.60E-64<br>1.90E-04 |
| AXK34883.1 | 4953110 | 4952393 | - | 238 | PF00392<br>TIGR02812<br>TIGR02325<br>PF07729<br>TIGR03338 | GntR<br>TIGR02812<br>TIGR02325<br>FCD<br>TIGR03338 | Bacterial regulatory proteins, gntR family<br>fadR_gamma: fatty acid metabolism transcriptional regulator FadR<br>C_P_lyase_phnF: phosphonate metabolism transcriptional regulator PhnF<br>FCD domain<br>phnR_burk: phosphonate utilization associated transcriptional regulator | 3.10E-17<br>1.90E-14<br>1.00E-12<br>6.20E-09<br>9.70E-08 |
| AXK37541.1 | 4953504 | 4954017 | + | 170 | PF00877<br>PF01471<br>TIGR02869<br>PF09374<br>PF05708 | NLPC_P60<br>PG_binding_1<br>TIGR02869<br>PG_binding_3<br>Peptidase_C92 | NlpC/P60 family<br>Putative peptidoglycan binding domain<br>spore_SleB: spore cortex-lytic enzyme<br>Predicted Peptidoglycan domain<br>Permuted papain-like amidase enzyme, YaeF/YiiX, C92 family | 7.30E-21<br>1.40E-14<br>1.40E-09<br>2.30E-06<br>5.40E-04 |
| AXK34884.1 | 4954787 | 4954109 | - | 225 | PF13649<br>PF08241<br>PF13489<br>TIGR02072<br>PF13847 | Methyltransf_25<br>Methyltransf_11<br>Methyltransf_23<br>TIGR02072<br>Methyltransf_31 | Methyltransferase domain<br>Methyltransferase domain<br>Methyltransferase domain<br>BioC: malonyl-acyl carrier protein O-methyltransferase BioC<br>Methyltransferase domain | 4.70E-17<br>1.50E-14<br>1.40E-11<br>5.20E-08<br>1.20E-07 |
| AXK34885.1 | 4956387 | 4954884 | - | 500 | PF01839<br>PF13517 | FG-GAP<br>VCBS | FG-GAP repeat<br>Repeat domain in Vibrio, Colwellia, Bradyrhizobium and Shewanella | 1.30E-24<br>7.80E-20 |
| AXK34886.1 | 4957647 | 4956612 | - | 344 | PF01471<br>TIGR02869<br>PF09374<br>TIGR03979 | PG_binding_1<br>TIGR02869<br>PG_binding_3<br>TIGR03979 | Putative peptidoglycan binding domain<br>spore_SleB: spore cortex-lytic enzyme<br>Predicted Peptidoglycan domain<br>His_Ser_Rich: His-Xaa-Ser repeat protein HxsA | 2.10E-30<br>1.80E-23<br>7.20E-10<br>3.70E-06 |
| AXK34887.1 | 4958599 | 4957627 | - | 323 | PF00288<br>TIGR00191<br>PF08544<br>TIGR01920<br>TIGR00549 | GHMP_kinases_N<br>TIGR00191<br>GHMP_kinases_C<br>TIGR01920<br>TIGR00549 | GHMP kinases N terminal domain<br>thrB: homoserine kinase<br>GHMP kinases C terminal<br>Shik_kin_archae: shikimate kinase<br>mevalon_kin: mevalonate kinase | 2.20E-08<br>2.70E-08<br>3.10E-08<br>1.10E-07<br>8.20E-05 |
| AXK34888.1 | 4958858 | 4959875 | + | 338 | TIGR03945<br>PF00291<br>TIGR01136<br>TIGR01139<br>TIGR01137 | TIGR03945<br>PALP<br>TIGR01136<br>TIGR01139<br>TIGR01137 | PLP_SbnA_fam: 2,3-diaminopropionate biosynthesis protein SbnA<br>Pyridoxal-phosphate dependent enzyme<br>cysKM: cysteine synthase<br>cysK: cysteine synthase A<br>cysta_beta: cystathionine beta-synthase | 5.10E-35<br>6.90E-28<br>8.40E-22<br>2.20E-21<br>5.70E-15 |

| index | leader | core | start | end | dir | score |
|---|---|---|---|---|---|---|
| 1 | VLLEKVNSDQEVSGYLEAAGPVS | GTTPVMDLTSVAYAIMEQYACMGSCLAAE | 4952056 | 4951897 | - | 6 |
| 2 | VRYPSSSRSCRRRRPP | VASRPMTTAVTCYVPR | 4952378 | 4952279 | - | 6 |
| 3 | VSAAVTCAIPVFVPLVPAE | TTPCRFPSHDDRGHVLRASVTGAPTAAGSQSVRLP | 4952397 | 4952232 | - | 8 |
| 4 | LSRSDSAQAADEVFADATEVPRVVRPPPSE | RAPQTTRAAALRVIFTGVPNARE | 4953476 | 4953314 | - | 8 |

### Results for WP_040898719.1 [Streptomyces auratus AGR0001] back to top

previous - next

#### Architecture

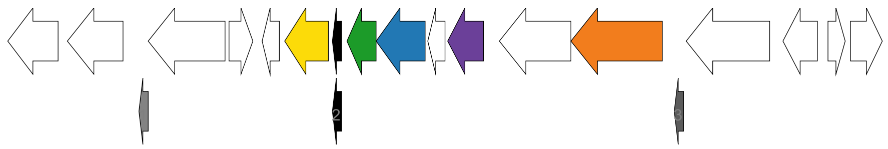

1000 nucleotides

Link to nucleotide sequence

| Accession | start | end | direction | length (aa) | Pfam/HMM | name | description | E-value |
|---|---|---|---|---|---|---|---|---|
| WP_006604045.1 | 2772650 | 2771591 | - | 352 | TIGR02432<br>PF01171<br>PF09179 | TIGR02432<br>ATP_bind_3<br>TilS | lysidine_TilS_N: tRNA(Ile)-lysidine synthetase<br>PP-loop family<br>TilS substrate binding domain | 4.30E-41<br>2.60E-36<br>4.80E-12 |
| WP_006604046.1 | 2773985 | 2772842 | - | 380 | TIGR03883<br>TIGR03624<br>PF10103 | TIGR03883<br>TIGR03624<br>Zincin_2 | DUF2342_F420: uncharacterized protein, coenzyme F420 biosynthesis associated<br>TIGR03624: putative hydrolase<br>Zincin-like metallopeptidase | 7.80E-142<br>1.10E-125<br>4.10E-121 |
| WP_006604047.1 | 2776109 | 2774501 | - | 535 | PF02113<br>TIGR00666 | Peptidase_S13<br>TIGR00666 | D-Ala-D-Ala carboxypeptidase 3 (S13) family<br>PBP4: D-alanyl-D-alanine carboxypeptidase/D-alanyl-D-alanine-endopeptidase | 2.90E-76<br>9.30E-57 |
| WP_005314187.1 | 2776195 | 2776690 | + | 164 | PF00719 | Pyrophosphatase | Inorganic pyrophosphatase | 2.50E-53 |
| WP_006604048.1 | 2777239 | 2776879 | - | 119 | NO PFAM MATCH | - | - | - |
| WP_040898716.1 | 2778240 | 2777328 | - | 303 | LinG<br>LinH<br>TIGR04033 | LinG<br>LinH<br>TIGR04033 | LinG Homolog<br>LinH (Fusion of LinE/LinG)<br>export_SdpB: antimicrobial peptide system protein, SdpB family | 1.40E-111<br>6.00E-79<br>2.50E-04 |
| WP_167551449.1 | 2778510 | 2778327 | - | 60 | LinA | LinA | LinA (linaridin precursor) | 1.80E-13 |
| WP_106430490.1 | 2779219 | 2778637 | - | 193 | LinL | LinL | LinL | 9.60E-56 |
| WP_040898719.1 | 2780241 | 2779221 | - | 339 | LinE<br>LinH<br>PF12697<br>TIGR02427<br>PF00561 | LinE<br>LinH<br>Abhydrolase_6<br>TIGR02427<br>Abhydrolase_1 | LinE Homolog<br>LinH (Fusion of LinE/LinG)<br>Alpha/beta hydrolase family<br>protocat_pcaD: 3-oxoadipate enol-lactonase<br>alpha/beta hydrolase fold | 2.90E-102<br>2.90E-51<br>4.10E-14<br>4.20E-10<br>8.40E-07 |
| WP_006604052.1 | 2780661 | 2780313 | - | 115 | PF13518<br>PF01527<br>PF13551<br>PF13011<br>PF13384 | HTH_28<br>HTH_Tnp_1<br>HTH_29<br>LZ_Tnp_IS481<br>HTH_23 | Helix-turn-helix domain<br>Transposase<br>Winged helix-turn helix<br>leucine-zipper of insertion element IS481<br>Homeodomain-like domain | 1.20E-09<br>1.70E-06<br>9.20E-06<br>9.60E-05<br>1.50E-04 |
| WP_040898722.1 | 2781462 | 2780703 | - | 252 | PF13649<br>PF08241<br>PF13489<br>PF08242<br>PF13847 | Methyltransf_25<br>Methyltransf_11<br>Methyltransf_23<br>Methyltransf_12<br>Methyltransf_31 | Methyltransferase domain<br>Methyltransferase domain<br>Methyltransferase domain<br>Methyltransferase domain<br>Methyltransferase domain | 8.30E-17<br>2.80E-12<br>2.10E-10<br>9.40E-10<br>6.50E-09 |
| WP_040898725.1 | 2783264 | 2781779 | - | 494 | TIGR02734<br>PF13450<br>TIGR00292<br>TIGR03467<br>PF01593 | TIGR02734<br>NAD_binding_8<br>TIGR00292<br>TIGR03467<br>Amino_oxidase | crtI_fam: phytoene desaturase<br>NAD(P)-binding Rossmann-like domain<br>TIGR00292: thiazole biosynthesis enzyme<br>HpnE: squalene-associated FAD-dependent desaturase<br>Flavin containing amine oxidoreductase | 1.80E-08<br>2.40E-08<br>4.70E-08<br>7.20E-08<br>1.40E-07 |
| WP_040900283.1 | 2785168 | 2783257 | - | 636 | TIGR02204<br>TIGR02203<br>TIGR00958<br>TIGR03796<br>TIGR03375 | TIGR02204<br>TIGR02203<br>TIGR00958<br>TIGR03796<br>TIGR03375 | MsbA_rel: ABC transporter, permease/ATP-binding protein<br>MsbA_lipidA: lipid A export permease/ATP-binding protein MsbA<br>3a01208: antigen peptide transporter 2<br>NHLM_micro_ABC1: NHLM bacteriocin system ABC transporter, peptidase/ATP-binding protein<br>type_I_sec_LssB: type I secretion system ATPase | 3.20E-101<br>2.80E-96<br>8.20E-90<br>1.70E-81<br>4.30E-80 |
| WP_006604056.1 | 2787391 | 2785654 | - | 578 | PF06738<br>PF12821 | ThrE<br>ThrE_2 | Putative threonine/serine exporter<br>Threonine/Serine exporter, ThrE | 5.10E-72<br>1.10E-18 |
| WP_006604057.1 | 2788382 | 2787653 | - | 242 | PF09335 | SNARE_assoc | SNARE associated Golgi protein | 2.60E-24 |
| WP_006604058.1 | 2788589 | 2788958 | + | 122 | PF01906 | YbjQ_1 | Putative heavy-metal-binding | 1.40E-33 |
| WP_006604059.1 | 2789048 | 2789711 | + | 220 | TIGR00949<br>PF01810<br>TIGR00948 | TIGR00949<br>LysE<br>TIGR00948 | 2A76: homoserine/Threonine efflux protein<br>LysE type translocator<br>2a75: L-lysine exporter | 2.70E-44<br>8.20E-44<br>1.80E-15 |

| index | leader | core | start | end | dir | score |
|---|---|---|---|---|---|---|
| 1 | VGVADFPRFRLSRRGGCSPLRFAAPPHVVGFP | AVVVLAVAPAAGVGGRVRCRSSGLRPAARPLP | 2774515 | 2774320 | - | 7 |
| 2 | MSVVANFANTELADVTPGRIGNDATPTM | LTPLAALATPEGVAVTSATAYALHEVTKDIQR | 2778510 | 2778327 | - | 33 |
| 3 | VLRRCGLVARACGKTKNAGDGESLAR | FAPAAEGSSPGSPMVGPTRSVRRSAAGFVTEQS | 2785594 | 2785414 | - | 8 |

### Results for SFX78248.1 [Streptomyces atratus] back to top

previous - next

#### Architecture

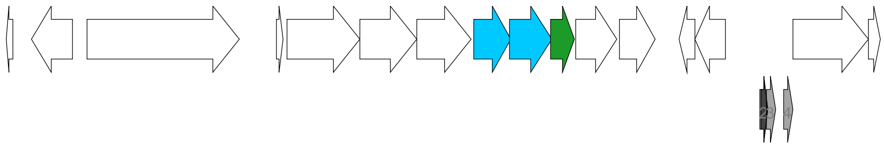

1000 nucleotides

Link to nucleotide sequence

| Accession | start | end | direction | length (aa) | Pfam/HMM | name | description | E-value |
|---|---|---|---|---|---|---|---|---|
| SFX78055.1 | 139971 | 139824 | - | 48 | NO PFAM MATCH | - | - | - |
| SFX78077.1 | 141363 | 140415 | - | 315 | TIGR03418<br>TIGR03339<br>TIGR02424<br>PF03466<br>PF00126 | TIGR03418<br>TIGR03339<br>TIGR02424<br>LysR_substrate<br>HTH_1 | chol_sulf_TF: putative choline sulfate-utilization transcription factor<br>phn_lysR: aminoethylphosphonate catabolism associated LysR family transcriptional regulator<br>TF_pcaQ: pca operon transcription factor PcaQ<br>LysR substrate binding domain<br>Bacterial regulatory helix-turn-helix protein, lysR family | 2.40E-18<br>7.40E-18<br>9.70E-17<br>9.90E-17<br>2.50E-16 |
| SFX78096.1 | 141716 | 145313 | + | 1198 | TIGR00594<br>PF07733<br>PF17657<br>TIGR01405<br>PF02811 | TIGR00594<br>DNA_pol3_alpha<br>DNA_pol3_finger<br>TIGR01405<br>PHP | polc: DNA polymerase III, alpha subunit<br>Bacterial DNA polymerase III alpha NTPase domain<br>Bacterial DNA polymerase III alpha subunit finger domain<br>polC_Gram_pos: DNA polymerase III, alpha subunit, Gram-positive type<br>PHP domain | 0.00E+00<br>1.50E-86<br>2.20E-56<br>3.40E-44<br>2.90E-41 |
| SFX78148.1 | 146191 | 146362 | + | 56 | NO PFAM MATCH | - | - | - |
| SFX78165.1 | 146458 | 148165 | + | 568 | PF04738 | Lant_dehydr_N | Lantibiotic dehydratase, N terminus | 4.60E-80 |
| SFX78185.1 | 148175 | 149519 | + | 447 | TIGR03891<br>PF14028<br>PF04738 | TIGR03891<br>Lant_dehydr_C<br>Lant_dehydr_N | thiopep_ocin: thiopeptide-type bacteriocin biosynthesis domain<br>Lantibiotic biosynthesis dehydratase C-term<br>Lantibiotic dehydratase, N terminus | 3.70E-50<br>1.20E-33<br>1.70E-25 |
| SFX78208.1 | 149515 | 150817 | + | 433 | PF05147<br>TIGR03897 | LANC_like<br>TIGR03897 | Lanthionine synthetase C-like protein<br>lanti_2_LanM: type 2 lantibiotic biosynthesis protein LanM | 1.80E-44<br>3.60E-15 |
| SFX78226.1 | 150858 | 151728 | + | 289 | LinH<br>LinG<br>PF05090<br>TIGR04033 | LinH<br>LinG<br>VKG_Carbox<br>TIGR04033 | LinH (Fusion of LinE/LinG)<br>LinG Homolog<br>Vitamin K-dependent gamma-carboxylase<br>export_SdpB: antimicrobial peptide system protein, SdpB family | 1.60E-112<br>1.00E-101<br>2.40E-04<br>7.70E-04 |
| SFX78248.1 | 151711 | 152701 | + | 329 | LinH<br>LinE<br>TIGR02427<br>PF12697<br>PF00561 | LinH<br>LinE<br>TIGR02427<br>Abhydrolase_6<br>Abhydrolase_1 | LinH (Fusion of LinE/LinG)<br>LinE Homolog<br>protocat_pcaD: 3-oxoadipate enol-lactonase<br>Alpha/beta hydrolase family<br>alpha/beta hydrolase fold | 1.50E-75<br>8.90E-75<br>2.30E-13<br>1.60E-12<br>1.60E-10 |
| SFX78265.1 | 152697 | 153231 | + | 177 | LinL | LinL | LinL | 6.20E-53 |
| SFX78287.1 | 153283 | 154261 | + | 325 | TIGR03564<br>TIGR03841<br>PF00296<br>TIGR04024<br>TIGR03617 | TIGR03564<br>TIGR03841<br>Bac_luciferase<br>TIGR04024<br>TIGR03617 | F420_MSMEG_4879: F420-dependent oxidoreductase, MSMEG_4879 family<br>F420_Rv3093c: probable F420-dependent oxidoreductase, Rv3093c family<br>Luciferase-like monooxygenase<br>F420_NP1902A: coenzyme F420-dependent oxidoreductase, NP1902A family<br>F420_MSMEG_2256: probable F420-dependent oxidoreductase, MSMEG_2256 family | 6.90E-38<br>1.50E-26<br>1.80E-22<br>5.80E-12<br>3.30E-11 |
| SFX78308.1 | 154300 | 155155 | + | 284 | PF00583<br>TIGR01575<br>PF08445<br>PF13673<br>PF13508 | Acetyltransf_1<br>TIGR01575<br>FR47<br>Acetyltransf_10<br>Acetyltransf_7 | Acetyltransferase (GNAT) family<br>riml: ribosomal-protein-alanine acetyltransferase<br>FR47-like protein<br>Acetyltransferase (GNAT) domain<br>Acetyltransferase (GNAT) domain | 5.30E-12<br>3.60E-09<br>3.70E-08<br>1.10E-07<br>7.70E-06 |
| SFX78328.1 | 156101 | 155711 | - | 129 | PF00392<br>TIGR02018<br>TIGR02325<br>TIGR02404<br>TIGR03337 | GntR<br>TIGR02018<br>TIGR02325<br>TIGR02404<br>TIGR03337 | Bacterial regulatory proteins, gntR family<br>his_ut_repres: histidine utilization repressor<br>C_P_lyase_phnF: phosphonate metabolism transcriptional regulator PhnF<br>trehalos_R_Bsub: trehalose operon repressor<br>phnR: phosphonate utilization transcriptional regulator PhnR | 1.30E-15<br>7.20E-11<br>1.10E-09<br>1.50E-08<br>9.20E-07 |
| SFX78346.1 | 156802 | 156097 | - | 234 | NO PFAM MATCH | - | - | - |
| SFX78370.1 | 158403 | 160200 | + | 598 | PF00082<br>TIGR03921 | Peptidase_S8<br>TIGR03921 | Subtilase family<br>T7SS_mycosin: type VII secretion-associated serine protease mycosin | 8.40E-29<br>2.90E-16 |
| SFX78397.1 | 160199 | 160496 | + | 98 | NO PFAM MATCH | - | - | - |

| index | leader | core | start | end | dir | score |
|---|---|---|---|---|---|---|
| 1 | MAPASAFTAGLSSARSPGCTASDVSASTGN | DATTSTKPSSASPPASSPTDTSNAFARTSY | 157633 | 157816 | + | 7 |
| 2 | LSSARSPGCTASDVSASTGNDATTS | TKPSSASPPASSPTDTSNAFARTSY | 157663 | 157816 | + | 7 |
| 3 | LPGPHIELEVTERTAFDTFSCSVGSRLPPTR | KTTRRSEPRTTRLSLRIVTHPSHSAASTSTAKGLPLRDS | 157796 | 158009 | + | 6 |
| 4 | VARRLGYGPGQDCGHVGESGLLATLIRAPSRDGE | GTAASDMRLRGRSCGAPPVGPVGVLIEGAERQQGQ | 158194 | 158404 | + | 6 |

### Results for WP_101256404.1 [Streptomyces barkulensis] back to top

previous - next

#### Architecture

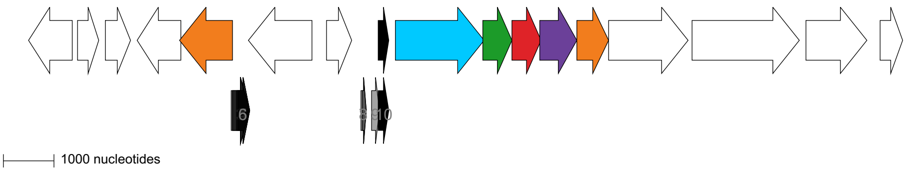

Link to nucleotide sequence

| Accession | start | end | direction | length (aa) | Pfam/HMM | name | description | E-value |
|---|---|---|---|---|---|---|---|---|

| | | | | | | | | |
|---|---|---|---|---|---|---|---|---|
| WP_101256398.1 | 30585 | 29712 | - | 290 | TIGR03446<br>TIGR03445<br>PF02585<br>TIGR04001<br>TIGR04000 | TIGR03446<br>TIGR03445<br>PIG-L<br>TIGR04001<br>TIGR04000 | mycothiol_Mca: mycothiol conjugate amidase Mca<br>mycothiol_MshB: N-acetyl-1-D-myo-inositol-2-amino-2-deoxy-alpha-D-glucopyranoside deacetylase<br>GlcNAc-PI de-N-acetylase<br>thiol_BshB1: bacillithiol biosynthesis deacetylase BshB1<br>thiol_BshB2: bacillithiol biosynthesis deacetylase BshB2 | 1.20E-150<br>9.70E-57<br>3.20E-27<br>1.70E-15<br>4.10E-10 |
| WP_101256426.1 | 30689 | 31109 | + | 139 | PF14155 | DUF4307 | Domain of unknown function (DUF4307) | 2.40E-29 |
| WP_101256399.1 | 31250 | 31748 | + | 165 | TIGR01462<br>PF03449<br>TIGR01461<br>PF01272 | TIGR01462<br>GreA_GreB_N<br>TIGR01461<br>GreA_GreB | greA: transcription elongation factor GreA<br>Transcription elongation factor, N-terminal<br>greB: transcription elongation factor GreB<br>Transcription elongation factor, GreA/GreB, C-term | 3.20E-45<br>5.30E-25<br>3.30E-17<br>5.40E-14 |
| WP_101256400.1 | 32724 | 31863 | - | 286 | PF01061<br>TIGR00025<br>TIGR01247<br>PF12698<br>TIGR01248 | ABC2_membrane<br>TIGR00025<br>TIGR01247<br>ABC2_membrane_3<br>TIGR01248 | ABC-2 type transporter<br>Mtu_efflux: ABC transporter efflux protein, DrrB family<br>drrB: daunorubicin resistance ABC transporter membrane protein<br>ABC-2 family transporter protein<br>drrC: daunorubicin resistance protein C | 3.90E-27<br>1.40E-20<br>2.10E-18<br>1.30E-16<br>1.00E-11 |
| WP_101256401.1 | 33746 | 32720 | - | 341 | TIGR01188<br>TIGR01288<br>TIGR03864<br>TIGR03522<br>TIGR01257 | TIGR01188<br>TIGR01288<br>TIGR03864<br>TIGR03522<br>TIGR01257 | drrA: daunorubicin resistance ABC transporter, ATP-binding protein<br>nodI: nodulation ABC transporter NodI<br>PQQ_ABC_ATP: ABC transporter, ATP-binding subunit, PQQ-dependent alcohol dehydrogenase system<br>GldA_ABC_ATP: gliding motility-associated ABC transporter ATP-binding subunit GldA<br>rim_protein: rim ABC transporter | 8.00E-111<br>1.70E-56<br>4.80E-53<br>6.90E-50<br>1.20E-46 |
| WP_101256402.1 | 35314 | 34069 | - | 414 | TIGR01127<br>TIGR02079<br>TIGR01124<br>PF00291<br>TIGR02991 | TIGR01127<br>TIGR02079<br>TIGR01124<br>PALP<br>TIGR02991 | ilvA_1Cterm: threonine ammonia-lyase<br>THD1: threonine dehydratase<br>ilvA_2Cterm: threonine ammonia-lyase, biosynthetic<br>Pyridoxal-phosphate dependent enzyme<br>ectoine_eutB: ectoine utilization protein EutB | 4.40E-130<br>9.50E-84<br>3.50E-83<br>5.50E-83<br>9.20E-80 |
| WP_101256427.1 | 35601 | 36105 | + | 167 | PF01047<br>PF12802<br>TIGR01884<br>TIGR02337<br>PF13463 | MarR<br>MarR_2<br>TIGR01884<br>TIGR02337<br>HTH_27 | MarR family<br>MarR family<br>cas_HTH: CRISPR locus-related DNA-binding protein<br>HpaR: homoprotocatechuate degradation operon regulator, HpaR<br>Winged helix DNA-binding domain | 3.00E-12<br>7.00E-11<br>5.50E-07<br>8.20E-07<br>2.10E-06 |
| WP_101256403.1 | 36647 | 36854 | + | 68 | LinA | LinA | LinA (linaridin precursor) | 1.80E-14 |
| WP_101256404.1 | 36967 | 38722 | + | 584 | LinH<br>LinG<br>LinE<br>PF12697<br>TIGR02427 | LinH<br>LinG<br>LinE<br>Abhydrolase_6<br>TIGR02427 | LinH (Fusion of LinE/LinG)<br>LinG Homolog<br>LinE Homolog<br>Alpha/beta hydrolase family<br>protocat_pcaD: 3-oxoadipate enol-lactonase | 5.00E-175<br>1.10E-72<br>9.40E-71<br>6.20E-11<br>3.10E-08 |
| WP_101256405.1 | 38711 | 39275 | + | 187 | LinL | LinL | LinL | 4.60E-52 |
| WP_101256406.1 | 39271 | 39844 | + | 190 | TIGR00521<br>PF02441<br>TIGR02113 | TIGR00521<br>Flavoprotein<br>TIGR02113 | coaBC_dfp: phosphopantothenoylcysteine decarboxylase / phosphopantothenate--cysteine ligase<br>Flavoprotein<br>coaC_strep: phosphopantothenoylcysteine decarboxylase | 1.40E-10<br>3.10E-09<br>2.30E-08 |
| WP_101256407.1 | 39836 | 40574 | + | 245 | PF13649<br>PF08241<br>TIGR02021<br>PF13847<br>TIGR01934 | Methyltransf_25<br>Methyltransf_11<br>TIGR02021<br>Methyltransf_31<br>TIGR01934 | Methyltransferase domain<br>Methyltransferase domain<br>BchM-ChlM: magnesium protoporphyrin O-methyltransferase<br>Methyltransferase domain<br>MenG_MenH_UbiE: ubiquinone/menaquinone biosynthesis methyltransferase | 1.60E-17<br>2.80E-15<br>2.40E-11<br>2.70E-11<br>9.10E-11 |
| WP_101256408.1 | 40570 | 41203 | + | 210 | TIGR02857<br>TIGR01189<br>TIGR02673<br>TIGR02982<br>TIGR03608 | TIGR02857<br>TIGR01189<br>TIGR02673<br>TIGR02982<br>TIGR03608 | CydD: thiol reductant ABC exporter, CydD subunit<br>ccmA: heme ABC exporter, ATP-binding protein CcmA<br>FtsE: cell division ATP-binding protein FtsE<br>heterocyst_DevA: ABC exporter ATP-binding subunit, DevA family<br>L_ocin_972_ABC: putative bacteriocin export ABC transporter, lactococcin 972 group | 3.20E-29<br>7.10E-27<br>1.30E-26<br>1.10E-23<br>3.10E-22 |
| WP_101256409.1 | 41199 | 42750 | + | 516 | NO PFAM MATCH | - | - | - |
| WP_018089273.1 | 42838 | 44950 | + | 704 | INFERRED GENE | - | - | - |
| WP_101256411.1 | 45091 | 46303 | + | 403 | TIGR01265<br>TIGR01264<br>PF00155<br>TIGR03540<br>TIGR01141 | TIGR01265<br>TIGR01264<br>Aminotran_1_2<br>TIGR03540<br>TIGR01141 | tyr_nico_aTase: tyrosine/nicotianamine family aminotransferase<br>tyr_amTase_E: tyrosine aminotransferase<br>Aminotransferase class I and II<br>DapC_direct: LL-diaminopimelate aminotransferase<br>hisC: histidinol-phosphate transaminase | 8.90E-72<br>5.20E-68<br>6.80E-54<br>6.50E-44<br>8.30E-32 |
| WP_101256412.1 | 46567 | 46993 | + | 141 | NO PFAM MATCH | - | - | - |

| index | leader | core | start | end | dir | score |
|---|---|---|---|---|---|---|
| 1 | VDGTWHGGLPWVWVVGKGRERWEPR...KRFTACPADVFQRRPTSVPTSVPTG | APPARRARGDPFRAAPCGRPRTPQPVTV | 33728 | 34088 | + | 7 |
| 2 | VWVVGKGRERWEPRQQWKPRRAAAP...KRFTACPADVFQRRPTSVPTSVPTG | APPARRARGDPFRAAPCGRPRTPQPVTV | 33761 | 34088 | + | 7 |
| 3 | VVGKGRERWEPRQQWKPRRAAAPHR...KRFTACPADVFQRRPTSVPTSVPTG | APPARRARGDPFRAAPCGRPRTPQPVTV | 33767 | 34088 | + | 7 |
| 4 | MRRRVRMPADTPRARCGSALCSNAI...KRFTACPADVFQRRPTSVPTSVPTG | APPARRARGDPFRAAPCGRPRTPQPVTV | 33842 | 34088 | + | 7 |
| 5 | VRMPADTPRARCGSALCSNAIYRVKRFTACPADVFQRRPTSVPTSVPTG | APPARRARGDPFRAAPCGRPRTPQPVTV | 33854 | 34088 | + | 7 |
| 6 | MPADTPRARCGSALCSNAIYRVKRFTACPADVFQRRPTSVPTSVPTG | APPARRARGDPFRAAPCGRPRTPQPVTV | 33860 | 34088 | + | 7 |
| 7 | VSKHEMNGRVQRSGPRSD | RRLSPSAAGTTPSSVTTV | 36301 | 36412 | + | 6 |
| 8 | MNGRVQRSGPRSDRR | LSPSAAGTTPSSVTTV | 36316 | 36412 | + | 7 |
| 9 | VFRGGCLQSLKRSSSVSCGASARSGR | REQCSTRRDHFARMIIDERLVKCDLN | 36504 | 36663 | + | 6 |
| 10 | MRSELTATRPDSADIRPEALAAQDFANTVLSGSAPGFHSDCETPAM | ATPATPTAAQFVIQGSTICLVC | 36647 | 36854 | + | 35 |

### Results for WP_101258824.1 [Streptomyces barkulensis] back to top

previous - next

#### Architecture

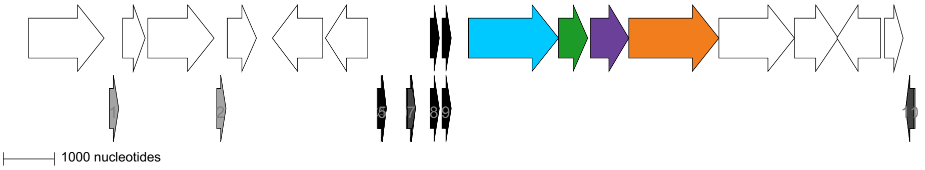

Link to nucleotide sequence

| Accession | start | end | direction | length (aa) | Pfam/HMM | name | description | E-value |
|---|---|---|---|---|---|---|---|---|
| WP_101258829.1 | 42343 | 43834 | + | 496 | PF00860 | Xan_ur_permease | Permease family | 1.70E-28 |
| WP_101258828.1 | 44192 | 44630 | + | 145 | PF12802<br>PF01047<br>TIGR02337<br>PF13412<br>PF09339 | MarR_2<br>MarR<br>TIGR02337<br>HTH_24<br>HTH_IclR | MarR family<br>MarR family<br>HpaR: homoprotocatechuate degradation operon regulator, HpaR<br>Winged helix-turn-helix DNA-binding<br>IclR helix-turn-helix domain | 3.30E-13<br>6.80E-13<br>2.60E-12<br>8.70E-05<br>1.20E-04 |
| WP_005315910.1 | 44680 | 45991 | + | 437 | INFERRED GENE | - | - | - |
| WP_101258827.1 | 46255 | 46825 | + | 189 | TIGR02258<br>PF02834<br>PF13563<br>PF10469 | TIGR02258<br>LigT_PEase<br>2_5_RNA_ligase2<br>AKAP7_NLS | 2_5_ligase: 2'-5' RNA ligase<br>LigT like Phosphoesterase<br>2'-5' RNA ligase superfamily<br>AKAP7 2'5' RNA ligase-like domain | 8.70E-34<br>2.50E-26<br>1.80E-18<br>4.20E-06 |
| WP_101258898.1 | 48108 | 47121 | - | 328 | PF00248<br>TIGR01293 | Aldo_ket_red<br>TIGR01293 | Aldo/keto reductase family<br>Kv_beta: voltage-dependent potassium channel beta subunit | 1.20E-72<br>2.20E-38 |
| WP_165869653.1 | 49006 | 48175 | - | 276 | PF13472<br>PF14606 | Lipase_GDSL_2<br>Lipase_GDSL_3 | GDSL-like Lipase/Acylhydrolase family<br>GDSL-like Lipase/Acylhydrolase family | 1.70E-17<br>9.60E-06 |
| WP_093846910.1 | 50222 | 50414 | + | 63 | LinA | LinA | LinA (linaridin precursor) | 2.90E-16 |
| WP_093846908.1 | 50451 | 50646 | + | 64 | LinA | LinA | LinA (linaridin precursor) | 3.00E-10 |
| WP_101258824.1 | 50991 | 52743 | + | 583 | LinH<br>LinE<br>LinG<br>PF12697<br>PF00561 | LinH<br>LinE<br>LinG<br>Abhydrolase_6<br>Abhydrolase_1 | LinH (Fusion of LinE/LinG)<br>LinE Homolog<br>LinG Homolog<br>Alpha/beta hydrolase family<br>alpha/beta hydrolase fold | 5.30E-224<br>3.50E-100<br>6.60E-96<br>1.70E-14<br>3.20E-13 |
| WP_101258823.1 | 52739 | 53318 | + | 192 | LinL | LinL | LinL | 2.30E-64 |
| WP_101258822.1 | 53367 | 54123 | + | 251 | PF13649<br>PF08241<br>PF13489<br>PF08242<br>PF13847 | Methyltransf_25<br>Methyltransf_11<br>Methyltransf_23<br>Methyltransf_12<br>Methyltransf_31 | Methyltransferase domain<br>Methyltransferase domain<br>Methyltransferase domain<br>Methyltransferase domain<br>Methyltransferase domain | 9.00E-14<br>2.70E-09<br>1.00E-08<br>1.60E-07<br>9.30E-07 |
| WP_101258821.1 | 54119 | 55898 | + | 592 | TIGR02204<br>TIGR02203<br>TIGR00958<br>TIGR03796<br>TIGR01846 | TIGR02204<br>TIGR02203<br>TIGR00958<br>TIGR03796<br>TIGR01846 | MsbA_rel: ABC transporter, permease/ATP-binding protein<br>MsbA_lipidA: lipid A export permease/ATP-binding protein MsbA<br>3a01208: antigen peptide transporter 2<br>NHLM_micro_ABC1: NHLM bacteriocin system ABC transporter, peptidase/ATP-binding protein<br>type_I_sec_HlyB: type I secretion system ATPase | 2.90E-99<br>4.90E-95<br>7.00E-86<br>1.40E-85<br>2.50E-78 |
| WP_101258820.1 | 55894 | 57376 | + | 493 | TIGR02734<br>TIGR01317<br>TIGR03467<br>PF13450<br>TIGR02733 | TIGR02734<br>TIGR01317<br>TIGR03467<br>NAD_binding_8<br>TIGR02733 | crtI_fam: phytoene desaturase<br>GOGAT_sm_gam: glutamate synthase, NADH/NADPH, small subunit<br>HpnE: squalene-associated FAD-dependent desaturase<br>NAD(P)-binding Rossmann-like domain<br>desat_CrtD: C-3',4' desaturase CrtD | 3.40E-11<br>1.10E-08<br>1.10E-08<br>1.30E-08<br>1.80E-08 |
| WP_101258819.1 | 57372 | 58224 | + | 283 | PF12680<br>PF14534 | SnoaL_2<br>DUF4440 | SnoaL-like domain<br>Domain of unknown function (DUF4440) | 3.90E-07<br>6.10E-04 |
| WP_101258818.1 | 59051 | 58208 | - | 280 | PF01551 | Peptidase_M23 | Peptidase family M23 | 8.90E-16 |
| WP_143688647.1 | 59137 | 59497 | + | 119 | PF03621 | MbtH | MbtH-like protein | 3.30E-27 |

| index | leader | core | start | end | dir | score |
|---|---|---|---|---|---|---|
| 1 | VGRTLAPRCAAWWWSRSSGAPSIRRLGTF | VSVDSMGTRSRGRRGRASAFLVLLLTHSYG | 43944 | 44124 | + | 6 |
| 2 | VASAEGPESIAGDGPAPQDRTPVSPRRTVPAAAPAA | APPSAPASAPGGERDHRGDVLA | 46051 | 46228 | + | 6 |
| 3 | MDGDSGNHLRCGVTEIDPRMAGVRRGCLDLFGQEEESNIPF | LPPGSSVARVYTPTGKHVAAPA | 49176 | 49368 | + | 9 |
| 4 | VTEIDPRMAGVRRGCLDLFGQEEESNIPF | LPPGSSVARVYTPTGKHVAAPA | 49212 | 49368 | + | 9 |
| 5 | MAGVRRGCLDLFGQEEESNIPF | LPPGSSVARVYTPTGKHVAAPA | 49233 | 49368 | + | 7 |
| 6 | VVGECCPGPRRVPARPFDRPRGPWGAARACGPRRGGPAPS | ACPVRTPQGVPFSVVM | 49769 | 49940 | + | 7 |
| 7 | VGECCPGPRRVPARPFDRPRGPWGAARACGPRRGGPAPS | ACPVRTPQGVPFSVVM | 49772 | 49940 | + | 7 |
| 8 | MEQKVAAFMNEQDFGPLEQGYQDDTPAPVL | ATPGAVATCWYVGGAVVGAGSVVGAYVTGRVVG | 50222 | 50414 | + | 29 |
| 9 | MSKIVESVVAVTPEELTDETHGEFGAMMPIQ | ATPGLVAFGVGFAGGAGAAWVTVQAYEAGRTAG | 50451 | 50646 | + | 31 |
| 10 | MELDGLGTTGQVYHHLRQLVSAGWLE | TAGRGRYRVPPTRLVPLLVVITAARH | 59716 | 59557 | - | 7 |
| 11 | VAELMELDGLGTTGQVYHHLRQLVSAGW | LETAGRGRYRVPPTRLVPLLVVITAARH | 59728 | 59557 | - | 7 |

### Results for WP_051827992.1 [Streptomyces bicolor] back to top

previous - next

#### Architecture

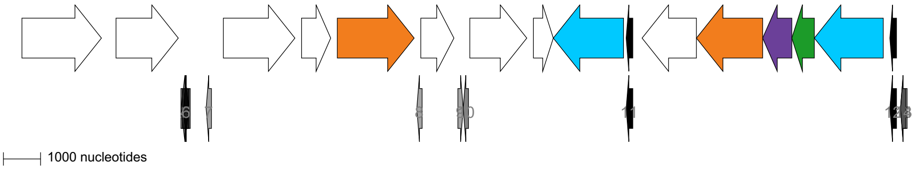

Link to nucleotide sequence

| Accession | start | end | direction | length (aa) | Pfam/HMM | name | description | E-value |
|---|---|---|---|---|---|---|---|---|
| WP_031486394.1 | 85952 | 88112 | + | 719 | TIGR01389<br>TIGR00614<br>PF00270<br>PF00271<br>TIGR04121 | TIGR01389<br>TIGR00614<br>DEAD<br>Helicase_C<br>TIGR04121 | recQ: ATP-dependent DNA helicase RecQ<br>recQ_fam: ATP-dependent DNA helicase, RecQ family<br>DEAD/DEAH box helicase<br>Helicase conserved C-terminal domain<br>DEXH_lig_assoc: DEXH box helicase, DNA ligase-associated | 4.90E-80<br>2.30E-66<br>2.50E-15<br>3.40E-15<br>5.10E-13 |
| WP_031486393.1 | 88499 | 90185 | + | 561 | PF13830 | DUF4192 | Domain of unknown function (DUF4192) | 1.00E-91 |
| WP_078662504.1 | 91378 | 93298 | + | 639 | PF14742 | GDE_N_bis | N-terminal domain of (some) glycogen debranching enzymes | 2.40E-16 |
| WP_031486391.1 | 93496 | 94255 | + | 252 | PF00293 | NUDIX | NUDIX domain | 3.30E-20 |
| WP_031486390.1 | 94464 | 96528 | + | 687 | TIGR01188<br>TIGR03740<br>TIGR01288<br>TIGR03864<br>TIGR03522 | TIGR01188<br>TIGR03740<br>TIGR01288<br>TIGR03864<br>TIGR03522 | drrA: daunorubicin resistance ABC transporter, ATP-binding protein<br>galliderm_ABC: lantibiotic protection ABC transporter, ATP-binding subunit<br>nodI: nodulation ABC transporter NodI<br>PQQ_ABC_ATP: ABC transporter, ATP-binding subunit, PQQ-dependent alcohol dehydrogenase system<br>GldA_ABC_ATP: gliding motility-associated ABC transporter ATP-binding subunit GldA | 6.10E-32<br>6.90E-28<br>5.70E-25<br>5.10E-23<br>5.70E-23 |
| WP_031486389.1 | 96691 | 97579 | + | 295 | PF07729<br>TIGR03338<br>PF00392<br>TIGR03337<br>TIGR02325 | FCD<br>TIGR03338<br>GntR<br>TIGR03337<br>TIGR02325 | FCD domain<br>phnR_burk: phosphonate utilization associated transcriptional regulator<br>Bacterial regulatory proteins, gntR family<br>phnR: phosphonate utilization transcriptional regulator PhnR<br>C_P_lyase_phnF: phosphonate metabolism transcriptional regulator PhnF | 4.10E-24<br>1.50E-22<br>1.20E-14<br>1.80E-04<br>1.90E-04 |
| WP_031486388.1 | 98007 | 99543 | + | 511 | TIGR02393<br>TIGR02997<br>TIGR02394<br>TIGR02392<br>TIGR02980 | TIGR02393<br>TIGR02997<br>TIGR02394<br>TIGR02392<br>TIGR02980 | RpoD_Cterm: RNA polymerase sigma factor RpoD<br>Sig70-cyanoRpoD: RNA polymerase sigma factor, cyanobacterial RpoD-like family<br>rpoS_proteo: RNA polymerase sigma factor RpoS<br>rpoH_proteo: alternative sigma factor RpoH<br>SigBFG: RNA polymerase sigma-70 factor, sigma-B/F/G subfamily | 7.50E-113<br>3.80E-105<br>3.30E-84<br>7.10E-42<br>2.10E-38 |
| WP_018824032.1 | 99720 | 100266 | + | 182 | INFERRED GENE | - | - | - |
| WP_051827992.1 | 102155 | 100262 | - | 630 | LinH<br>LinG<br>LinE<br>PF12697<br>PF12146 | LinH<br>LinG<br>LinE<br>Abhydrolase_6<br>Hydrolase_4 | LinH (Fusion of LinE/LinG)<br>LinG Homolog<br>LinE Homolog<br>Alpha/beta hydrolase family<br>Serine aminopeptidase, S33 | 1.10E-240<br>1.20E-120<br>4.30E-94<br>1.30E-16<br>1.50E-08 |
| WP_158751433.1 | 102412 | 102235 | - | 58 | LinA | LinA | LinA (linaridin precursor) | 3.90E-12 |
| WP_031486385.1 | 104118 | 102636 | - | 493 | TIGR02734<br>TIGR03467<br>PF13450<br>PF01593<br>TIGR02733 | TIGR02734<br>TIGR03467<br>NAD_binding_8<br>Amino_oxidase<br>TIGR02733 | crtI_fam: phytoene desaturase<br>HpnE: squalene-associated FAD-dependent desaturase<br>NAD(P)-binding Rossmann-like domain<br>Flavin containing amine oxidoreductase<br>desat_CrtD: C-3',4' desaturase CrtD | 4.70E-14<br>4.20E-10<br>6.70E-10<br>4.00E-09<br>6.50E-09 |
| WP_051827993.1 | 105902 | 104114 | - | 595 | TIGR02204<br>TIGR02203<br>TIGR03796<br>TIGR00958<br>TIGR03375 | TIGR02204<br>TIGR02203<br>TIGR03796<br>TIGR00958<br>TIGR03375 | MsbA_rel: ABC transporter, permease/ATP-binding protein<br>MsbA_lipidA: lipid A export permease/ATP-binding protein MsbA<br>NHLM_micro_ABC1: NHLM bacteriocin system ABC transporter, peptidase/ATP-binding protein<br>3a01208: antigen peptide transporter 2<br>type_I_sec_LssB: type I secretion system ATPase | 4.30E-104<br>7.30E-94<br>1.10E-84<br>1.30E-84<br>4.10E-82 |
| WP_051828000.1 | 106663 | 105898 | - | 254 | PF13649<br>PF08241<br>PF13847<br>PF08242<br>PF13489 | Methyltransf_25<br>Methyltransf_11<br>Methyltransf_31<br>Methyltransf_12<br>Methyltransf_23 | Methyltransferase domain<br>Methyltransferase domain<br>Methyltransferase domain<br>Methyltransferase domain<br>Methyltransferase domain | 5.30E-14<br>1.00E-11<br>1.10E-07<br>5.10E-07<br>9.40E-07 |
| WP_031498384.1 | 107271 | 106677 | - | 197 | LinL | LinL | LinL | 1.00E-62 |
| WP_051828002.1 | 109142 | 107267 | - | 624 | LinH<br>LinG<br>LinE<br>PF12697<br>PF00561 | LinH<br>LinG<br>LinE<br>Abhydrolase_6<br>Abhydrolase_1 | LinH (Fusion of LinE/LinG)<br>LinG Homolog<br>LinE Homolog<br>Alpha/beta hydrolase family<br>alpha/beta hydrolase fold | 2.90E-242<br>2.20E-121<br>4.40E-95<br>3.90E-12<br>1.20E-07 |
| WP_158751434.1 | 109470 | 109299 | - | 56 | LinA | LinA | LinA (linaridin precursor) | 1.50E-11 |

| index | leader | core | start | end | dir | score |
|---|---|---|---|---|---|---|
| 1 | LPDPVEMTAVAAGALGDAIKGRWLWDGARAPIGRGCP | CPGARRATPRSESARTAGAGSRAALRRPAPPQPAGRTR | 90245 | 90473 | + | 6 |
| 2 | MTAVAAGALGDAIKGRWLWDGARAPIGRGCPCPGARR | ATPRSESARTAGAGSRAALRRPAPPQPAGRTR | 90263 | 90473 | + | 8 |

| | | | | | | |
|---|---|---|---|---|---|---|
| 3 | VRGAEAQPCCRPRLSGPTRIGEL | LVAPRGMDTPGRSAPAPRPTATVP | 90444 | 90300 | - | 6 |
| 4 | LRRCGAPKRSPAAGPGCPGRLGSGSCSSRPGA | WTPPADRRPRPVPQPPSLDGIAECACRDGRHFDGIRQASGRG | 90452 | 90227 | - | 6 |
| 5 | VFARRVEAVRGAEAQPCCRPRLSGPTR | IGELLVAPRGMDTPGRSAPAPRPTATVP | 90468 | 90300 | - | 7 |
| 6 | MGQPRRRGAAFSVFARRVEAVRGAEAQPCCRPR | LSGPTRIGELLVAPRGMDTPGRSAPAPRPTATVP | 90504 | 90300 | - | 6 |
| 7 | MGSACGDVPPCPTGDEDAGLG | LRCVGLARRVGCTGSSDRPSHA | 91059 | 90927 | - | 6 |
| 8 | MVCARVLTDPLSAPS | STPLRSLVKSTRKGDRSPRRN | 96714 | 96603 | - | 6 |
| 9 | VRRGRGIAVQCMVEAMGAPFCR | YPPVFDRLRDVTRATQIGRNALGNSAMT | 97696 | 97849 | + | 6 |
| 10 | LENGFRPRPRRTGADHRLGDGPT | VRAGAGKMHSALSGVRIPSSAVTS | 97989 | 97845 | - | 7 |
| 11 | MQDIAEYMNNHPDLQGDITDADGAVRTEMLPVL | ATPVIVARTALATVAAYTVTRNHIG | 102412 | 102235 | - | 32 |
| 12 | MDVATQLNKQAELRSVAIDDEEGVVFSRQEPVL | ATPTLIMAARVAVATALVYTARR | 109470 | 109299 | - | 32 |
| 13 | VLNRYGLGAEGRRFQDELGRHLEAV | DTQANRFDEKRATVLARMADREKQGR | 109742 | 109586 | - | 7 |
| 14 | VLQPVLRHLNVLNRYGLGAEGRRFQDELGR | HLEAVDTQANRFDEKRATVLARMADREKQGR | 109772 | 109586 | - | 6 |

### Results for WP_051828002.1 [Streptomyces bicolor] back to top

previous - next

#### Architecture

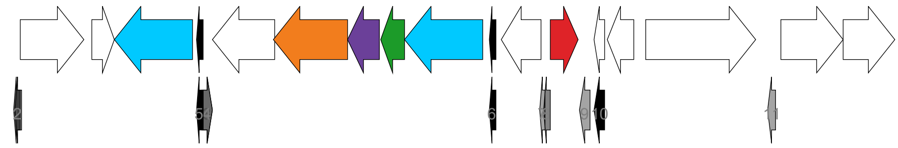

1000 nucleotides

Link to nucleotide sequence

| Accession | start | end | direction | length (aa) | Pfam/HMM | name | description | E-value |
|---|---|---|---|---|---|---|---|---|
| WP_031486388.1 | 98007 | 99543 | + | 511 | TIGR02393<br>TIGR02997<br>TIGR02394<br>TIGR02392<br>TIGR02980 | TIGR02393<br>TIGR02997<br>TIGR02394<br>TIGR02392<br>TIGR02980 | RpoD_Cterm: RNA polymerase sigma factor RpoD<br>Sig70-cyanoRpoD: RNA polymerase sigma factor, cyanobacterial RpoD-like family<br>rpoS_proteo: RNA polymerase sigma factor RpoS<br>rpoH_proteo: alternative sigma factor RpoH<br>SigBFG: RNA polymerase sigma-70 factor, sigma-B/F/G subfamily | 7.50E-113<br>3.80E-105<br>3.30E-84<br>7.10E-42<br>2.10E-38 |
| WP_018824032.1 | 99720 | 100266 | + | 182 | INFERRED GENE | - | - | - |
| WP_051827992.1 | 102155 | 100262 | - | 630 | LinH<br>LinG<br>LinE<br>PF12697<br>PF12146 | LinH<br>LinG<br>LinE<br>Abhydrolase_6<br>Hydrolase_4 | LinH (Fusion of LinE/LinG)<br>LinG Homolog<br>LinE Homolog<br>Alpha/beta hydrolase family<br>Serine aminopeptidase, S33 | 1.10E-240<br>1.20E-120<br>4.30E-94<br>1.30E-16<br>1.50E-08 |
| WP_158751433.1 | 102412 | 102235 | - | 58 | LinA | LinA | LinA (linaridin precursor) | 3.90E-12 |
| WP_031486385.1 | 104118 | 102636 | - | 493 | TIGR02734<br>TIGR03467<br>PF13450<br>PF01593<br>TIGR02733 | TIGR02734<br>TIGR03467<br>NAD_binding_8<br>Amino_oxidase<br>TIGR02733 | crtI_fam: phytoene desaturase<br>HpnE: squalene-associated FAD-dependent desaturase<br>NAD(P)-binding Rossmann-like domain<br>Flavin containing amine oxidoreductase<br>desat_CrtD: C-3',4' desaturase CrtD | 4.70E-14<br>4.20E-10<br>6.70E-10<br>4.00E-09<br>6.50E-09 |
| WP_051827993.1 | 105902 | 104114 | - | 595 | TIGR02204<br>TIGR02203<br>TIGR03796<br>TIGR00958<br>TIGR03375 | TIGR02204<br>TIGR02203<br>TIGR03796<br>TIGR00958<br>TIGR03375 | MsbA_rel: ABC transporter, permease/ATP-binding protein<br>MsbA_lipidA: lipid A export permease/ATP-binding protein MsbA<br>NHLM_micro_ABC1: NHLM bacteriocin system ABC transporter, peptidase/ATP-binding protein<br>3a01208: antigen peptide transporter 2<br>type_I_sec_LssB: type I secretion system ATPase | 4.30E-104<br>7.30E-94<br>1.10E-84<br>1.30E-84<br>4.10E-82 |
| WP_051828000.1 | 106663 | 105898 | - | 254 | PF13649<br>PF08241<br>PF13847<br>PF08242<br>PF13489 | Methyltransf_25<br>Methyltransf_11<br>Methyltransf_31<br>Methyltransf_12<br>Methyltransf_23 | Methyltransferase domain<br>Methyltransferase domain<br>Methyltransferase domain<br>Methyltransferase domain<br>Methyltransferase domain | 5.30E-14<br>1.00E-11<br>1.10E-07<br>5.10E-07<br>9.40E-07 |
| WP_031498384.1 | 107271 | 106677 | - | 197 | LinL | LinL | LinL | 1.00E-62 |
| WP_051828002.1 | 109142 | 107267 | - | 624 | LinH<br>LinG<br>LinE<br>PF12697<br>PF00561 | LinH<br>LinG<br>LinE<br>Abhydrolase_6<br>Abhydrolase_1 | LinH (Fusion of LinE/LinG)<br>LinG Homolog<br>LinE Homolog<br>Alpha/beta hydrolase family<br>alpha/beta hydrolase fold | 2.90E-242<br>2.20E-121<br>4.40E-95<br>3.90E-12<br>1.20E-07 |
| WP_158751434.1 | 109470 | 109299 | - | 56 | LinA | LinA | LinA (linaridin precursor) | 1.50E-11 |
| WP_031486380.1 | 110540 | 109586 | - | 317 | PF03405 | FA_desaturase_2 | Fatty acid desaturase | 2.60E-66 |
| WP_051828004.1 | 110781 | 111429 | + | 215 | TIGR00521<br>PF02441<br>TIGR02113<br>TIGR02699<br>TIGR02700 | TIGR00521<br>Flavoprotein<br>TIGR02113<br>TIGR02699<br>TIGR02700 | coaBC_dfp: phosphopantothenoylcysteine decarboxylase / phosphopantothenate--cysteine ligase<br>Flavoprotein<br>coaC_strep: phosphopantothenoylcysteine decarboxylase<br>archaeo_AfpA: archaeoflavoprotein AfpA<br>flavo_MJ0208: archaeoflavoprotein, MJ0208 family | 5.40E-25<br>9.90E-24<br>1.90E-13<br>3.80E-05<br>1.40E-04 |
| WP_031486378.1 | 112082 | 111815 | - | 88 | TIGR00517<br>PF00550<br>PF14573 | TIGR00517<br>PP-binding<br>PP-binding_2 | acyl_carrier: acyl carrier protein<br>Phosphopantetheine attachment site<br>Acyl-carrier | 2.80E-17<br>2.20E-16<br>1.10E-04 |

| | | | | | | | | |
|---|---|---|---|---|---|---|---|---|
| WP_031486377.1 | 112800 | 112137 | - | 220 | PF00196<br>TIGR03020<br>PF00072<br>PF08281<br>TIGR03541 | GerE<br>TIGR03020<br>Response_reg<br>Sigma70_r4_2<br>TIGR03541 | Bacterial regulatory proteins, luxR family<br>EpsA: transcriptional regulator EpsA<br>Response regulator receiver domain<br>Sigma-70, region 4<br>reg_near_HchA: LuxR family transcriptional regulatory, chaperone HchA-associated | 1.30E-19<br>3.40E-14<br>3.40E-10<br>4.70E-08<br>5.90E-08 |
| WP_031498388.1 | 113065 | 115717 | + | 883 | PF00069<br>PF05147<br>TIGR03897<br>TIGR03903<br>PF01636 | Pkinase<br>LANC_like<br>TIGR03897<br>TIGR03903<br>APH | Protein kinase domain<br>Lanthionine synthetase C-like protein<br>lanti_2_LanM: type 2 lantibiotic biosynthesis protein LanM<br>TOMM_kin_cyc: TOMM system kinase/cyclase fusion protein<br>Phosphotransferase enzyme family | 5.10E-15<br>1.10E-07<br>1.10E-06<br>1.60E-06<br>2.90E-06 |
| WP_051828007.1 | 116341 | 117835 | + | 497 | PF00501<br>TIGR01733<br>TIGR01923<br>TIGR03098<br>TIGR03208 | AMP-binding<br>TIGR01733<br>TIGR01923<br>TIGR03098<br>TIGR03208 | AMP-binding enzyme<br>AA-adenyl-dom: amino acid adenylation domain<br>menE: O-succinylbenzoate-CoA ligase<br>ligase_PEP_1: acyl-CoA ligase (AMP-forming), exosortase A-associated<br>cyc_hxne_CoA_lg: cyclohexanecarboxylate-CoA ligase | 8.00E-56<br>1.00E-42<br>3.00E-28<br>9.30E-27<br>6.10E-26 |
| WP_031486374.1 | 117839 | 119081 | + | 413 | TIGR03150<br>PF00109<br>PF02801<br>TIGR02813<br>TIGR01930 | TIGR03150<br>ketoacyl-synt<br>Ketoacyl-synt_C<br>TIGR02813<br>TIGR01930 | fabF: beta-ketoacyl-acyl-carrier-protein synthase II<br>Beta-ketoacyl synthase, N-terminal domain<br>Beta-ketoacyl synthase, C-terminal domain<br>omega_3_PfaA: polyketide-type polyunsaturated fatty acid synthase PfaA<br>AcCoA-C-Actrans: acetyl-CoA C-acyltransferase | 1.90E-152<br>1.20E-50<br>3.50E-38<br>3.10E-26<br>3.60E-08 |

| index | leader | core | start | end | dir | score |
|---|---|---|---|---|---|---|
| 1 | LENGFRPRPRRTGADHRLGDGPT | VRAGAGKMHSALSGVRIPSSAVTS | 97989 | 97845 | - | 8 |
| 2 | VLADTNNLSERWKTASGPVHGG | QEPTTGLGMGRRCGRGPGRCTAP | 98021 | 97883 | - | 6 |
| 3 | MYSAMSCMSSWVLS | RTVSYPVCDVRILHPLVHDREFLVL...IRFVIAASVLKRKIVKRGEFAWFES | 102389 | 102629 | + | 6 |
| 4 | MSCMSSWVLS | RTVSYPVCDVRILHPLVHDREFLVL...IRFVIAASVLKRKIVKRGEFAWFES | 102401 | 102629 | + | 6 |
| 5 | MQDIAEYMNNHPDLQGDITDADGAVRTEMLPVL | ATPVIVARTALATVAAYTVTRNHIG | 102412 | 102235 | - | 33 |
| 6 | MDVATQLNKQAELRSVAIDDEEGVVFSRQEPVL | ATPTLIMAARVAVATALVYTARR | 109470 | 109299 | - | 33 |
| 7 | LSPKTGRRLAVCAPTC | LTFECTGPDESVQHMSGMRPVTDEFR | 110655 | 110526 | - | 6 |
| 8 | MSLLRASCFLPPHSIESPIIPGSIAR | MTPASPLELPVANLSMICRRRPGDGLPFARRPA | 110783 | 110603 | - | 7 |
| 9 | MPRSADTPCRSPGGTSTSADHHRRRPAARGRRGAGVLDAGL | RADRPSTRSPASSARSTPVLVPVQGLSGGSCRRRGAWHAHLR | 111725 | 111473 | - | 6 |
| 10 | MTTQSADALDQELFGAVADLLVEIAEIPAED | ITPEARFRDDLDVDSLLMVEFGVSI...IPDEEFTQLHTVGALVAFVQRHKSG | 112082 | 111815 | - | 18 |
| 11 | VSCRSAVPAGRAGRHQGGLLLGGLQLLKVE | NRLGHVTHLLMRSRWSVAGRRGPAGLSTPSW | 116205 | 116019 | - | 6 |

### Results for WP_030214179.1 [Streptomyces bikiniensis] back to top

previous - next

#### Architecture

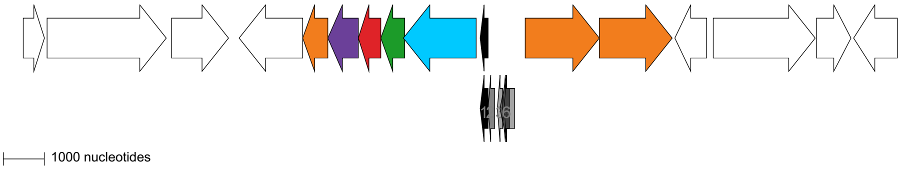

Link to nucleotide sequence

| Accession | start | end | direction | length (aa) | Pfam/HMM | name | description | E-value |
|---|---|---|---|---|---|---|---|---|
| WP_030214157.1 | 415232 | 415766 | + | 177 | PF12802<br>PF01047<br>TIGR02337<br>PF13463 | MarR_2<br>MarR<br>TIGR02337<br>HTH_27 | MarR family<br>MarR family<br>HpaR: homoprotocatechuate degradation operon regulator, HpaR<br>Winged helix DNA-binding domain | 6.60E-11<br>3.40E-10<br>1.10E-09<br>3.00E-08 |
| WP_030214160.1 | 415805 | 418730 | + | 974 | PF02913<br>TIGR00387<br>PF01565<br>PF13534<br>PF13183 | FAD-oxidase_C<br>TIGR00387<br>FAD_binding_4<br>Fer4_17<br>Fer4_8 | FAD linked oxidases, C-terminal domain<br>glcD: glycolate oxidase, subunit GlcD<br>FAD binding domain<br>4Fe-4S dicluster domain<br>4Fe-4S dicluster domain | 2.90E-51<br>2.90E-48<br>1.50E-29<br>2.20E-08<br>1.10E-07 |
| WP_030214163.1 | 418865 | 420254 | + | 462 | PF01408<br>TIGR04380 | GFO_IDH_MocA<br>TIGR04380 | Oxidoreductase family, NAD-binding Rossmann fold<br>myo_inos_iolG: inositol 2-dehydrogenase | 4.80E-15<br>1.80E-09 |
| WP_030214166.1 | 422063 | 420512 | - | 516 | NO PFAM MATCH | - | - | - |
| WP_030214168.1 | 422692 | 422059 | - | 210 | TIGR02857<br>TIGR01189<br>TIGR02673<br>TIGR02982<br>TIGR01166 | TIGR02857<br>TIGR01189<br>TIGR02673<br>TIGR02982<br>TIGR01166 | CydD: thiol reductant ABC exporter, CydD subunit<br>ccmA: heme ABC exporter, ATP-binding protein CcmA<br>FtsE: cell division ATP-binding protein FtsE<br>heterocyst_DevA: ABC exporter ATP-binding subunit, DevA family<br>cbiO: cobalt ABC transporter, ATP-binding protein | 6.20E-29<br>2.40E-27<br>5.70E-27<br>3.30E-23<br>1.90E-22 |
| WP_030214171.1 | 423429 | 422688 | - | 246 | PF13649<br>PF08241<br>TIGR02021<br>TIGR01934<br>PF13847 | Methyltransf_25<br>Methyltransf_11<br>TIGR02021<br>TIGR01934<br>Methyltransf_31 | Methyltransferase domain<br>Methyltransferase domain<br>BchM-ChlM: magnesium protoporphyrin O-methyltransferase<br>MenG_MenH_UbiE: ubiquinone/menaquinone biosynthesis methyltransferase<br>Methyltransferase domain | 1.90E-17<br>6.60E-15<br>9.20E-12<br>1.10E-10<br>4.80E-10 |
| WP_030214174.1 | 423994 | 423421 | - | 190 | TIGR00521<br>PF02441<br>TIGR02113 | TIGR00521<br>Flavoprotein<br>TIGR02113 | coaBC_dfp: phosphopantothenoylcysteine decarboxylase / phosphopantothenate--cysteine ligase<br>Flavoprotein<br>coaC_strep: phosphopantothenoylcysteine decarboxylase | 2.00E-10<br>2.40E-09<br>4.00E-08 |
| WP_030214176.1 | 424551 | 423990 | - | 186 | LinL | LinL | LinL | 4.20E-52 |

| | | | | | | | | |
|---|---|---|---|---|---|---|---|---|
| WP_030214179.1 | 426295 | 424540 | - | 584 | LinH<br>LinG<br>LinE<br>PF12697<br>TIGR02427 | LinH<br>LinG<br>LinE<br>Abhydrolase_6<br>TIGR02427 | LinH (Fusion of LinE/LinG)<br>LinG Homolog<br>LinE Homolog<br>Alpha/beta hydrolase family<br>protocat_pcaD: 3-oxoadipate enol-lactonase | 6.60E-174<br>5.00E-74<br>7.20E-69<br>4.00E-09<br>3.90E-08 |
| WP_030214182.1 | 426598 | 426409 | - | 62 | LinA | LinA | LinA (linaridin precursor) | 3.20E-14 |
| WP_030214185.1 | 427501 | 429325 | + | 607 | TIGR02203<br>TIGR03375<br>TIGR00958<br>TIGR01846<br>TIGR02204 | TIGR02203<br>TIGR03375<br>TIGR00958<br>TIGR01846<br>TIGR02204 | MsbA_lipidA: lipid A export permease/ATP-binding protein MsbA<br>type_I_sec_LssB: type I secretion system ATPase<br>3a01208: antigen peptide transporter 2<br>type_I_sec_HlyB: type I secretion system ATPase<br>MsbA_rel: ABC transporter, permease/ATP-binding protein | 3.10E-41<br>1.60E-38<br>2.30E-36<br>1.20E-34<br>1.30E-34 |
| WP_030214189.1 | 429321 | 431103 | + | 593 | TIGR02204<br>TIGR02203<br>TIGR01846<br>TIGR00958<br>TIGR03796 | TIGR02204<br>TIGR02203<br>TIGR01846<br>TIGR00958<br>TIGR03796 | MsbA_rel: ABC transporter, permease/ATP-binding protein<br>MsbA_lipidA: lipid A export permease/ATP-binding protein MsbA<br>type_I_sec_HlyB: type I secretion system ATPase<br>3a01208: antigen peptide transporter 2<br>NHLM_micro_ABC1: NHLM bacteriocin system ABC transporter, peptidase/ATP-binding protein | 9.80E-85<br>7.00E-76<br>3.90E-73<br>9.40E-72<br>2.00E-70 |
| WP_030214192.1 | 431962 | 431170 | - | 263 | PF04307<br>PF09988 | YdjM<br>DUF2227 | LexA-binding, inner membrane-associated putative hydrolase<br>Uncharacterized metal-binding protein (DUF2227) | 1.10E-34<br>6.30E-04 |
| WP_030214195.1 | 432093 | 434607 | + | 837 | PF12029<br>TIGR04121<br>PF00270<br>TIGR03817<br>TIGR00614 | DUF3516<br>TIGR04121<br>DEAD<br>TIGR03817<br>TIGR00614 | Domain of unknown function (DUF3516)<br>DEXH_lig_assoc: DEXH box helicase, DNA ligase-associated<br>DEAD/DEAH box helicase<br>DECH_helic: helicase/secretion neighborhood putative DEAH-box helicase<br>recQ_fam: ATP-dependent DNA helicase, RecQ family | 2.30E-215<br>4.50E-15<br>5.40E-12<br>2.00E-10<br>3.10E-08 |
| WP_030214197.1 | 434621 | 435485 | + | 287 | TIGR00189<br>PF13622<br>PF02551 | TIGR00189<br>4HBT_3<br>Acyl_CoA_thio | tesB: acyl-CoA thioesterase II<br>Thioesterase-like superfamily<br>Acyl-CoA thioesterase | 1.40E-86<br>3.80E-64<br>5.30E-43 |
| WP_030214200.1 | 436608 | 435537 | - | 356 | TIGR02306<br>PF09414 | TIGR02306<br>RNA_ligase | RNA_lig_DRB0094: RNA ligase<br>RNA ligase | 6.00E-137<br>5.10E-17 |

| index | leader | core | start | end | dir | score |
|---|---|---|---|---|---|---|
| 1 | MTRQDSTAVHPEVLAAQDFANTVLEGAAPGFHSNCETPAM | ATPATPTVAQFVIQGSTICLVC | 426598 | 426409 | - | 35 |
| 2 | VFRVGPRRRFLGAARHVHAKRVAGAAIF | GTPRPFARMTTDERLIKCDLYRP | 426750 | 426594 | - | 7 |
| 3 | MIRRPPVARRALPGAPC | CLPVLDEVRLRRTLASVSLDG | 426952 | 426835 | - | 6 |
| 4 | LRSASPGAAARSRCAGSEGAQSLRSVGP | WVRGSMAENMRHGATGRGECDALPSSTGR | 427122 | 426948 | - | 6 |
| 5 | VITPLAVRESGCRRPQQMRRVGRGSV | SPVRGSVGPWVHGREYAPRRDRARGM | 427135 | 426976 | - | 6 |
| 6 | VAVPEAAEEEGRREVSHGCVGPERG...DRGVRAAAGDHAACGPRVRVPPPAA | DAPGRKGLSLSGPWVRGSVGPWPRI...PADDSSPSGGPEGAPGRALLPPGTG | 427241 | 426881 | - | 6 |

### Results for WP_078489996.1 [Streptomyces bikiniensis] back to top

previous - next

#### Architecture

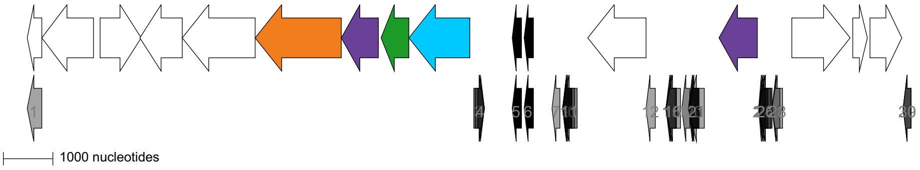

Link to nucleotide sequence

| Accession | start | end | direction | length (aa) | Pfam/HMM | name | description | E-value |
|---|---|---|---|---|---|---|---|---|
| WP_030221125.1 | 39708 | 39426 | - | 93 | PF00550<br>TIGR01688 | PP-binding<br>TIGR01688 | Phosphopantetheine attachment site<br>dltC: D-alanine--poly(phosphoribitol) ligase, subunit 2 | 2.70E-09<br>7.00E-06 |
| WP_051716317.1 | 40748 | 39704 | - | 347 | TIGR00747<br>PF08541 | TIGR00747<br>ACP_syn_III_C | fabH: 3-oxoacyl-[acyl-carrier-protein] synthase III<br>3-Oxoacyl-[acyl-carrier-protein (ACP)] synthase III C terminal | 1.10E-08<br>8.70E-05 |
| WP_033210586.1 | 40894 | 41722 | + | 275 | PF01551 | Peptidase_M23 | Peptidase family M23 | 9.80E-16 |
| WP_030221133.1 | 42558 | 41706 | - | 283 | PF12680<br>PF14534 | SnoaL_2<br>DUF4440 | SnoaL-like domain<br>Domain of unknown function (DUF4440) | 7.70E-07<br>1.80E-06 |
| WP_030221136.1 | 44039 | 42554 | - | 494 | TIGR02734<br>TIGR03467<br>TIGR01317<br>PF13450<br>TIGR00292 | TIGR02734<br>TIGR03467<br>TIGR01317<br>NAD_binding_8<br>TIGR00292 | crtI_fam: phytoene desaturase<br>HpnE: squalene-associated FAD-dependent desaturase<br>GOGAT_sm_gam: glutamate synthase, NADH/NADPH, small subunit<br>NAD(P)-binding Rossmann-like domain<br>TIGR00292: thiazole biosynthesis enzyme | 4.40E-10<br>4.90E-08<br>6.60E-08<br>1.30E-07<br>5.30E-07 |
| WP_030221139.1 | 45763 | 44035 | - | 575 | TIGR02204<br>TIGR02203<br>TIGR00958<br>TIGR01846<br>TIGR03375 | TIGR02204<br>TIGR02203<br>TIGR00958<br>TIGR01846<br>TIGR03375 | MsbA_rel: ABC transporter, permease/ATP-binding protein<br>MsbA_lipidA: lipid A export permease/ATP-binding protein MsbA<br>3a01208: antigen peptide transporter 2<br>type_I_sec_HlyB: type I secretion system ATPase<br>type_I_sec_LssB: type I secretion system ATPase | 3.50E-107<br>2.20E-101<br>1.00E-87<br>1.10E-85<br>7.40E-84 |
| WP_030221141.1 | 46515 | 45759 | - | 251 | PF13649<br>PF08241<br>PF13489<br>PF08242<br>PF13847 | Methyltransf_25<br>Methyltransf_11<br>Methyltransf_23<br>Methyltransf_12<br>Methyltransf_31 | Methyltransferase domain<br>Methyltransferase domain<br>Methyltransferase domain<br>Methyltransferase domain<br>Methyltransferase domain | 6.80E-13<br>7.40E-10<br>3.40E-07<br>6.10E-07<br>1.90E-06 |
| WP_030221144.1 | 47144 | 46565 | - | 192 | LinL | LinL | LinL | 5.00E-65 |

| | | | | | | | | |
|---|---|---|---|---|---|---|---|---|
| WP_078489996.1 | 48379 | 47140 | - | 412 | LinH<br>LinE<br>LinG<br>PF12697<br>PF00561 | LinH<br>LinE<br>LinG<br>Abhydrolase_6<br>Abhydrolase_1 | LinH (Fusion of LinE/LinG)<br>LinE Homolog<br>LinG Homolog<br>Alpha/beta hydrolase family<br>alpha/beta hydrolase fold | 1.00E-145<br>1.70E-103<br>5.40E-28<br>2.10E-15<br>1.40E-11 |
| WP_030221150.1 | 49420 | 49228 | - | 63 | LinA | LinA | LinA (linaridin precursor) | 4.80E-12 |
| WP_030221152.1 | 49655 | 49463 | - | 63 | LinA | LinA | LinA (linaridin precursor) | 9.60E-17 |
| WP_051716319.1 | 51941 | 50759 | - | 393 | NO PFAM MATCH | - | - | - |
| WP_030221156.1 | 54187 | 53398 | - | 262 | PF13649<br>PF08241<br>PF13847<br>TIGR01934<br>PF08242 | Methyltransf_25<br>Methyltransf_11<br>Methyltransf_31<br>TIGR01934<br>Methyltransf_12 | Methyltransferase domain<br>Methyltransferase domain<br>Methyltransferase domain<br>MenG_MenH_UbiE: ubiquinone/menaquinone biosynthesis methyltransferase<br>Methyltransferase domain | 8.10E-19<br>3.30E-15<br>2.20E-14<br>6.80E-11<br>1.50E-10 |
| WP_106969492.1 | 54863 | 56063 | + | 399 | PF01548<br>PF02371 | DEDD_Tnp_IS110<br>Transposase_20 | Transposase<br>Transposase IS116/IS110/IS902 family | 2.30E-46<br>3.50E-24 |
| WP_158708711.1 | 56105 | 56399 | + | 97 | NO PFAM MATCH | - | - | - |
| WP_010988951.1 | 56463 | 57102 | + | 213 | INFERRED GENE | - | - | - |

| index | leader | core | start | end | dir | score |
|---|---|---|---|---|---|---|
| 1 | MSRGSAETLDEITRFLTEALRREVGPDDDYFALGLVDSLFALELVT | HVEDRFELTVEVEDLDLDSFRTARRLTAFVHAKKADAETAPVHAGDS | 39708 | 39426 | - | 6 |
| 2 | VRVRRTPGRAEPSRHFVPASLARPD...MFAWIARYHIRASWTSGLRDSRARP | VTAWTWVATWSEPSVP | 48458 | 48671 | + | 7 |
| 3 | VRRTPGRAEPSRHFVPASLARPDSE...MFAWIARYHIRASWTSGLRDSRARP | VTAWTWVATWSEPSVP | 48464 | 48671 | + | 7 |
| 4 | LARPDSEYDMFAWIARYHIRASWTSGLRDSRARP | VTAWTWVATWSEPSVP | 48518 | 48671 | + | 7 |
| 5 | MSRIVDSVVAVTPQHLSDEAHGEFGAMMPVQ | ATPALIAFGVGFAGGAGAAWVTVQAYEAGAND | 49420 | 49228 | - | 30 |
| 6 | MEHKIAGFMNTQDFGTLEQGYQDDAAAPVL | ATPAAVATCWYVGGAVVGAGAVVGAYVTGRVNG | 49655 | 49463 | - | 30 |
| 7 | VLRRPTRPLGRRRVTSMIFRPELGAIW | TTFTGALTRASSGNGHVGMRPERQDL | 50200 | 50038 | - | 6 |
| 8 | VIVGHAGNRAADAVVRAGGGSERGEGGRC | VRVPAGARPTARRPYGHGSPAVRARSVRTS | 50428 | 50248 | - | 8 |
| 9 | VDVIVGHAGNRAADAVVRAGGGSERGEGGR | CVRVPAGARPTARRPYGHGSPAVRARSVRTS | 50434 | 50248 | - | 8 |
| 10 | LVSGSLAARERSGEGCVDVIVGHAGNRAADAVVRAGGG | SERGEGGRCVRVPAGARPTARRPYGHGSPAVRARSVRTS | 50482 | 50248 | - | 6 |
| 11 | VACVVLVKYCPKAGDVLIDPKRLVSGSLAARERSGEGCVDVIVGHAGNR | AADAVVRAGGGSERGEGGRCVRVPAGARPTARRPYGHGSPAVRARSVRTS | 50548 | 50248 | - | 7 |
| 12 | VGSGLGGVPAPDAGPGRWPRAVVEAADAEGIRAALPGPASAWWAAER | LTAALGEPLPFGRP | 52123 | 51937 | - | 6 |
| 13 | MPAVHRFPAARPLASCPPLYGRRRPHLGACPTP | GAPSETPRSCGGFHRSRGVLVDHPSE | 52520 | 52340 | - | 6 |
| 14 | VGARMPAVHRFPAARPLASCPPLYGRRRPHLGACPTP | GAPSETPRSCGGFHRSRGVLVDHPSE | 52532 | 52340 | - | 6 |
| 15 | VVVGARMPAVHRFPAARPLASCPPLYGRRRPHLGACPTP | GAPSETPRSCGGFHRSRGVLVDHPSE | 52538 | 52340 | - | 6 |
| 16 | MPGGAARSCCCRCAGRCPLFLRVAA...AARPLASCPPLYGRRRPHLGACPTP | GAPSETPRSCGGFHRSRGVLVDHPSE | 52634 | 52340 | - | 6 |
| 17 | VYRPERAEDCSATL | STACRCPVQPAGRGPVEGAMGMVTGQADPSMATRHPLL | 52804 | 52645 | - | 7 |
| 18 | VWNDSNTAGVKCTGGGAFIGVALCKNGKTAQGAKAAA | GTTSYAYCTSVNSSLKKPVQAWGVQA | 52982 | 52790 | - | 8 |
| 19 | VSCGTTPTPRVSSVRAAVPSSEWHSA | RTERLLRGQRRLPALPRMPTARPSTPP | 52987 | 52825 | - | 6 |
| 20 | MRCSVVWNDSNTAGVKCTGGGAFIGVALCKNGKTAQGAKAAA | GTTSYAYCTSVNSSLKKPVQAWGVQA | 52997 | 52790 | - | 8 |
| 21 | VKRKLGSVLAVAALAASAAVVNGAS...TGGGAFIGVALCKNGKTAQGAKAAA | GTTSYAYCTSVNSSLKKPVQAWGVQA | 53102 | 52790 | - | 6 |
| 22 | VPRFPVKSGGVSATGSGWVLLNAS | LTRPGTVRGAGTESQARSSPRVVE | 54348 | 54201 | - | 6 |
| 23 | MRRWWSRDLRRCGVPRFPVKSGGVSATGSG | WVLLNASLTRPGTVRGAGTESQARSSPRVVE | 54387 | 54201 | - | 8 |
| 24 | LVVAQDSPTRGQTPQLRPSWPMRRWWSRDLRRCGVPRFPVK | SGGVSATGSGWVLLNASLTRPGTVRGAGTESQARSSPRVVE | 54450 | 54201 | - | 6 |
| 25 | LVENPSTRSLVVAQDSPTRGQTPQLRPSWPMRRWWSRDLRRCGVP | RFPVKSGGVSATGSGWVLLNASLTRPGTVRGAGTESQARSSPRVVE | 54477 | 54201 | - | 6 |
| 26 | VILVENPSTRSLVVAQDSPTRGQTPQLRPSWPMRRWWSRDLRRCGV | PRFPVKSGGVSATGSGWVLLNASLTRPGTVRGAGTESQARSSPRVVE | 54483 | 54201 | - | 6 |
| 27 | LRSHEKGNGGGKAPPPHSDLLTPGCQLSG | QEPQGAAGRIDRGRGGNHTKLLADVGDLG | 54646 | 54469 | - | 6 |
| 28 | MNNKGGKSCRTRRSEAPLRSHEKGNGGGKAPPPHSDL | LTPGCQLSGQEPQGAAGRIDRGRGGNHTKLLADVGDLG | 54697 | 54469 | - | 6 |
| 29 | VCEGSGAWSGCRSAAGESVDHDADHG | PADHGLGVCRLLLVVAGEAAVCGDPRQ | 57277 | 57115 | - | 7 |
| 30 | MIDVCEGSGAWSGCRSAAGESVDHDADH | GPADHGLGVCRLLLVVAGEAAVCGDPRQ | 57286 | 57115 | - | 7 |

### Results for PJE97628.1 [Streptomyces carminius] back to top

previous - next

#### Architecture

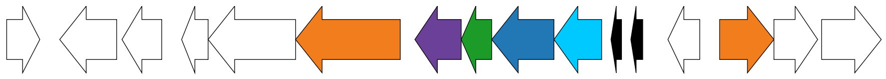

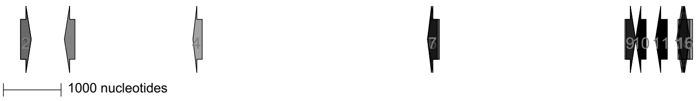

Link to nucleotide sequence

| Accession | start | end | direction | length (aa) | Pfam/HMM | name | description | E-value |
|---|---|---|---|---|---|---|---|---|
| PJE97621.1 | 65298 | 65865 | + | 188 | TIGR02258<br>PF02834<br>PF13563 | TIGR02258<br>LigT_PEase<br>2_5_RNA_ligase2 | 2_5_ligase: 2'-5' RNA ligase<br>LigT like Phosphoesterase<br>2'-5' RNA ligase superfamily | 5.40E-33<br>4.70E-24<br>5.90E-14 |
| PJE97622.1 | 67198 | 66211 | - | 328 | PF00248<br>TIGR01293 | Aldo_ket_red<br>TIGR01293 | Aldo/keto reductase family<br>Kv_beta: voltage-dependent potassium channel beta subunit | 1.10E-69<br>5.90E-39 |
| PJE97805.1 | 67985 | 67283 | - | 233 | PF13472 | Lipase_GDSL_2 | GDSL-like Lipase/Acylhydrolase family | 8.80E-16 |
| PJE97623.1 | 68779 | 68329 | - | 149 | NO PFAM MATCH | - | - | - |
| PJE97624.1 | 70296 | 68778 | - | 505 | TIGR02734<br>PF13450<br>TIGR01317<br>TIGR03467<br>TIGR00292 | TIGR02734<br>NAD_binding_8<br>TIGR01317<br>TIGR03467<br>TIGR00292 | crtI_fam: phytoene desaturase<br>NAD(P)-binding Rossmann-like domain<br>GOGAT_sm_gam: glutamate synthase, NADH/NADPH, small subunit<br>HpnE: squalene-associated FAD-dependent desaturase<br>TIGR00292: thiazole biosynthesis enzyme | 2.40E-13<br>1.70E-10<br>4.50E-08<br>4.90E-08<br>1.20E-07 |
| PJE97625.1 | 72107 | 70292 | - | 604 | TIGR02204<br>TIGR02203<br>TIGR00958<br>TIGR01846<br>TIGR03375 | TIGR02204<br>TIGR02203<br>TIGR00958<br>TIGR01846<br>TIGR03375 | MsbA_rel: ABC transporter, permease/ATP-binding protein<br>MsbA_lipidA: lipid A export permease/ATP-binding protein MsbA<br>3a01208: antigen peptide transporter 2<br>type_I_sec_HlyB: type I secretion system ATPase<br>type_I_sec_LssB: type I secretion system ATPase | 1.00E-110<br>2.40E-107<br>7.30E-88<br>4.70E-86<br>3.00E-82 |
| PJE97626.1 | 73169 | 72377 | - | 263 | PF13649<br>PF08241<br>PF13489<br>PF08242<br>TIGR01934 | Methyltransf_25<br>Methyltransf_11<br>Methyltransf_23<br>Methyltransf_12<br>TIGR01934 | Methyltransferase domain<br>Methyltransferase domain<br>Methyltransferase domain<br>Methyltransferase domain<br>MenG_MenH_UbiE: ubiquinone/menaquinone biosynthesis methyltransferase | 1.40E-11<br>6.20E-10<br>6.00E-08<br>3.00E-05<br>1.40E-04 |
| PJE97627.1 | 73705 | 73165 | - | 179 | LinL | LinL | LinL | 9.30E-62 |
| PJE97628.1 | 74778 | 73704 | - | 357 | LinE<br>LinH<br>PF12697<br>TIGR02427<br>TIGR03695 | LinE<br>LinH<br>Abhydrolase_6<br>TIGR02427<br>TIGR03695 | LinE Homolog<br>LinH (Fusion of LinE/LinG)<br>Alpha/beta hydrolase family<br>protocat_pcaD: 3-oxoadipate enol-lactonase<br>menH_SHCHC: 2-succinyl-6-hydroxy-2,4-cyclohexadiene-1-carboxylate synthase | 1.20E-123<br>2.70E-83<br>2.70E-17<br>1.30E-10<br>1.40E-09 |
| PJE97629.1 | 75611 | 74774 | - | 278 | LinH<br>LinG | LinH<br>LinG | LinH (Fusion of LinE/LinG)<br>LinG Homolog | 3.90E-105<br>4.80E-102 |
| PJE97630.1 | 75955 | 75772 | - | 60 | LinA | LinA | LinA (linaridin precursor) | 2.00E-11 |
| PJE97631.1 | 76328 | 76127 | - | 66 | LinA | LinA | LinA (linaridin precursor) | 4.90E-13 |
| PJE97632.1 | 77322 | 76758 | - | 187 | TIGR02983<br>TIGR02937<br>PF04542<br>PF08281<br>TIGR02985 | TIGR02983<br>TIGR02937<br>Sigma70_r2<br>Sigma70_r4_2<br>TIGR02985 | SigE-fam_strep: RNA polymerase sigma-70 factor, sigma-E family<br>sigma70-ECF: RNA polymerase sigma factor, sigma-70 family<br>Sigma-70 region 2<br>Sigma-70, region 4<br>Sig70_bacteroi1: RNA polymerase sigma-70 factor, Bacteroides expansion family 1 | 2.00E-16<br>7.10E-16<br>1.60E-11<br>7.00E-08<br>5.80E-07 |
| PJE97633.1 | 77647 | 78601 | + | 317 | TIGR01188<br>TIGR03864<br>TIGR01288<br>TIGR04406<br>TIGR02673 | TIGR01188<br>TIGR03864<br>TIGR01288<br>TIGR04406<br>TIGR02673 | drrA: daunorubicin resistance ABC transporter, ATP-binding protein<br>PQQ_ABC_ATP: ABC transporter, ATP-binding subunit, PQQ-dependent alcohol dehydrogenase system<br>nodI: nodulation ABC transporter NodI<br>LPS_export_lptB: LPS export ABC transporter ATP-binding protein<br>FtsE: cell division ATP-binding protein FtsE | 2.70E-59<br>2.00E-44<br>2.20E-42<br>6.90E-34<br>2.30E-33 |
| PJE97634.1 | 78597 | 79362 | + | 254 | PF01061<br>PF12698<br>TIGR01291 | ABC2_membrane<br>ABC2_membrane_3<br>TIGR01291 | ABC-2 type transporter<br>ABC-2 family transporter protein<br>nodJ: ABC-2 type transporter, NodJ family | 8.40E-15<br>5.50E-06<br>4.20E-05 |
| PJE97635.1 | 79428 | 80466 | + | 345 | PF07676<br>PF13449 | PD40<br>Phytase-like | WD40-like Beta Propeller Repeat<br>Esterase-like activity of phytase | 2.30E-04<br>2.30E-04 |

| index | leader | core | start | end | dir | score |
|---|---|---|---|---|---|---|
| 1 | VVAGPGRPGGGAAPEGKGEPGDIAG | GTAAPGPSPAGKENAPAGTGNVPAAVRAPERGRERRGKMPV | 65101 | 65302 | + | 6 |
| 2 | VAGPGRPGGGAAPEGKGEPGDIAG | GTAAPGPSPAGKENAPAGTGNVPAAVRAPERGRERRGKMPV | 65104 | 65302 | + | 6 |
| 3 | MRARAAAEATGRSPGGAAGCPPPGRPGQPSTAAR | ITTASSSASAPLTMRLRVFGSTPPP | 66054 | 65874 | - | 7 |
| 4 | MSGVNGVDRNELDETIAKAFKEVRTAI | DTHSEKSVQMYSNALRALVELRREVGAE | 68260 | 68092 | - | 6 |
| 5 | VACFTTALAI | ATPLTLVMFKDGPLTRETWAKGLGSAASLALLAAVLFGEDGP | 72330 | 72171 | - | 8 |
| 6 | VPTAHRVACFTTALAI | ATPLTLVMFKDGPLTRETWAKGLGSAASLALLAAVLFGEDGP | 72348 | 72171 | - | 7 |
| 7 | VTTQPHGKGPTV | PTAHRVACFTTALAIATPLTLVMFK...TWAKGLGSAASLALLAAVLFGEDGP | 72381 | 72171 | - | 7 |
| 8 | LRRRDSMICSWAGVPRVVVARIGLAGKRRAEGRRAETGGGETGSPGAVP | GLPAHPVAAGRVTS | 75591 | 75783 | + | 8 |
| 9 | MICSWAGVPRVVVARIGLAGKRRAEGRRAETGGGETGSPGAVP | GLPAHPVAAGRVTS | 75609 | 75783 | + | 8 |
| 10 | MSVLVEKVNSGAGRRTEDPVAPVM | ATPAAFVAGVVAGAKAAGALVAAAGVGAAVGAAVKK | 75955 | 75772 | - | 31 |
| 11 | MLTQKVNANTSLTGALEPGTPLGGAAPVM | ATPALIATPAIAKAAGAVVGAGAVTGAAYAAYRAVIN | 76328 | 76127 | - | 29 |

| | | | | | | |
|---|---|---|---|---|---|---|
| 12 | VAVGASPIPVLSGVFRGCRPMAGGT...APLCFRCGSGTGSGEGVDPAAAAGL | CTVPAPGSSVALLY | 76501 | 76711 | + | 6 |
| 13 | VLSGVFRGCRPMAGGTAGTGKAPLCFRCGSGTGSGEGVDPAAAAGL | CTVPAPGSSVALLY | 76528 | 76711 | + | 6 |
| 14 | VHSPAAAAGSTPSPLPVPDPHRKHKGALPVP | AVPPAIGRQPRKTPDSTGIGEAPTATSAPARR | 76671 | 76479 | - | 6 |
| 15 | LSVQESDGGSRSRNGAQSGCRGRVHAFAAPRA | GSAPETQGGFTGTGRPAGHRPTAPENTGQHRNR | 76714 | 76516 | - | 7 |
| 16 | MKIFPPSGHTPTGSPELSVQESDGGSRSRNGAQSGCRGRV | HAFAAPRAGSAPETQGGFTGTGRPAGHRPTAPENTGQHRNR | 76762 | 76516 | - | 6 |

### Results for WP_128380536.1 [Streptomyces cavernae] back to top

previous - next

#### Architecture

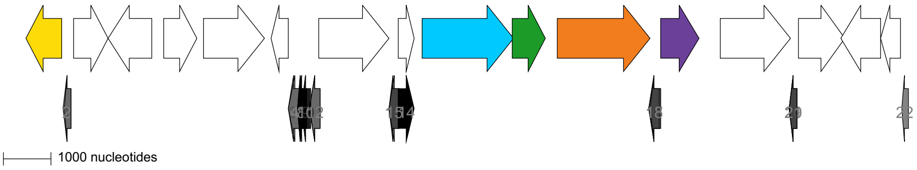

Link to nucleotide sequence

| Accession | start | end | direction | length (aa) | Pfam/HMM | name | description | E-value |
|---|---|---|---|---|---|---|---|---|
| WP_128380530.1 | 9809 | 9074 | - | 244 | PF12697<br>PF12146<br>LinH<br>TIGR01738<br>LinE | Abhydrolase_6<br>Hydrolase_4<br>LinH<br>TIGR01738<br>LinE | Alpha/beta hydrolase family<br>Serine aminopeptidase, S33<br>LinH (Fusion of LinE/LinG)<br>bioH: pimelyl-[acyl-carrier protein] methyl ester esterase<br>LinE Homolog | 1.60E-19<br>3.20E-13<br>5.50E-13<br>9.20E-12<br>2.00E-11 |
| WP_128380636.1 | 10075 | 10792 | + | 238 | PF03575<br>TIGR02069 | Peptidase_S51<br>TIGR02069 | Peptidase family S51<br>cyanophycinase: cyanophycinase | 2.40E-41<br>3.00E-07 |
| WP_128380531.1 | 11719 | 10780 | - | 312 | PF00196<br>TIGR03541<br>TIGR03020<br>PF08281 | GerE<br>TIGR03541<br>TIGR03020<br>Sigma70_r4_2 | Bacterial regulatory proteins, luxR family<br>reg_near_HchA: LuxR family transcriptional regulatory, chaperone HchA-associated<br>EpsA: transcriptional regulator EpsA<br>Sigma-70, region 4 | 1.00E-14<br>8.00E-09<br>5.70E-08<br>1.60E-04 |
| WP_128380532.1 | 11971 | 12655 | + | 227 | PF00072<br>PF00196<br>TIGR02875<br>TIGR03020<br>TIGR03541 | Response_reg<br>GerE<br>TIGR02875<br>TIGR03020<br>TIGR03541 | Response regulator receiver domain<br>Bacterial regulatory proteins, luxR family<br>spore_0_A: sporulation transcription factor Spo0A<br>EpsA: transcriptional regulator EpsA<br>reg_near_HchA: LuxR family transcriptional regulatory, chaperone HchA-associated | 1.50E-24<br>2.60E-18<br>1.10E-15<br>2.00E-12<br>2.90E-08 |
| WP_128380533.1 | 12803 | 14084 | + | 426 | PF07730<br>PF13581 | HisKA_3<br>HATPase_c_2 | Histidine kinase<br>Histidine kinase-like ATPase domain | 1.30E-13<br>4.40E-04 |
| WP_128380534.1 | 14574 | 14229 | - | 114 | PF00196<br>TIGR03541<br>TIGR03020<br>PF08281<br>PF13384 | GerE<br>TIGR03541<br>TIGR03020<br>Sigma70_r4_2<br>HTH_23 | Bacterial regulatory proteins, luxR family<br>reg_near_HchA: LuxR family transcriptional regulatory, chaperone HchA-associated<br>EpsA: transcriptional regulator EpsA<br>Sigma-70, region 4<br>Homeodomain-like domain | 3.40E-17<br>2.50E-12<br>6.20E-11<br>4.30E-08<br>2.30E-05 |
| WP_128380535.1 | 15226 | 16690 | + | 487 | TIGR02734<br>PF13450<br>TIGR03467<br>PF01593<br>TIGR01317 | TIGR02734<br>NAD_binding_8<br>TIGR03467<br>Amino_oxidase<br>TIGR01317 | crtI_fam: phytoene desaturase<br>NAD(P)-binding Rossmann-like domain<br>HpnE: squalene-associated FAD-dependent desaturase<br>Flavin containing amine oxidoreductase<br>GOGAT_sm_gam: glutamate synthase, NADH/NADPH, small subunit | 2.20E-12<br>2.70E-11<br>1.30E-08<br>1.80E-08<br>5.10E-07 |
| WP_155589799.1 | 16905 | 17229 | + | 107 | NO PFAM MATCH | - | - | - |
| WP_128380536.1 | 17395 | 19303 | + | 635 | LinH<br>LinG<br>LinE<br>PF12697<br>PF12146 | LinH<br>LinG<br>LinE<br>Abhydrolase_6<br>Hydrolase_4 | LinH (Fusion of LinE/LinG)<br>LinG Homolog<br>LinE Homolog<br>Alpha/beta hydrolase family<br>Serine aminopeptidase, S33 | 1.40E-238<br>3.80E-125<br>3.60E-89<br>4.40E-16<br>2.00E-10 |
| WP_128380537.1 | 19299 | 19986 | + | 228 | LinL | LinL | LinL | 3.80E-76 |
| WP_128380538.1 | 20229 | 22179 | + | 649 | TIGR02204<br>TIGR02203<br>TIGR00958<br>TIGR03796<br>TIGR01846 | TIGR02204<br>TIGR02203<br>TIGR00958<br>TIGR03796<br>TIGR01846 | MsbA_rel: ABC transporter, permease/ATP-binding protein<br>MsbA_lipidA: lipid A export permease/ATP-binding protein MsbA<br>3a01208: antigen peptide transporter 2<br>NHLM_micro_ABC1: NHLM bacteriocin system ABC transporter, peptidase/ATP-binding protein<br>type_I_sec_HlyB: type I secretion system ATPase | 8.20E-106<br>1.90E-102<br>2.10E-92<br>1.60E-86<br>2.70E-78 |
| WP_128380539.1 | 22422 | 23214 | + | 263 | PF13649<br>PF08241<br>PF13489<br>PF13847<br>TIGR01983 | Methyltransf_25<br>Methyltransf_11<br>Methyltransf_23<br>Methyltransf_31<br>TIGR01983 | Methyltransferase domain<br>Methyltransferase domain<br>Methyltransferase domain<br>Methyltransferase domain<br>UbiG: 3-demethylubiquinone-9 3-O-methyltransferase | 2.80E-15<br>2.00E-12<br>2.40E-10<br>5.50E-07<br>8.90E-07 |
| WP_162467260.1 | 23673 | 25134 | + | 486 | PF09339<br>TIGR02431<br>PF12802<br>PF13412 | HTH_IclR<br>TIGR02431<br>MarR_2<br>HTH_24 | IclR helix-turn-helix domain<br>pcaR_pcaU: beta-ketoadipate pathway transcriptional regulators, PcaR/PcaU/PobR family<br>MarR family<br>Winged helix-turn-helix DNA-binding | 4.90E-23<br>3.10E-18<br>1.20E-05<br>8.00E-05 |
| WP_128380541.1 | 25311 | 26292 | + | 326 | PF01636<br>TIGR02721 | APH<br>TIGR02721 | Phosphotransferase enzyme family<br>ycfN_thiK: thiamine kinase | 4.10E-17<br>1.50E-04 |
| WP_128380542.1 | 27023 | 26210 | - | 270 | PF02424 | ApbE | ApbE family | 1.30E-46 |
| WP_128380543.1 | 27465 | 27024 | - | 146 | PF04205<br>TIGR01938 | FMN_bind<br>TIGR01938 | FMN-binding domain<br>nqrC: NADH:ubiquinone oxidoreductase, Na(+)-translocating, C subunit | 1.30E-11<br>6.40E-04 |

| index | leader | core | start | end | dir | score |
|---|---|---|---|---|---|---|
| 1 | VPSHRLLPQSNSPHRDRAGRGTGMRGRCD | TRARRGNSAARPIGVMEAGVAKCRSRRTC | 10005 | 9828 | - | 6 |

| | | | | | | |
|---|---|---|---|---|---|---|
| 2 | MPEPCQAIGYFRNRIVRTAIAPAVAPVCAVG | VTPVHGEAIRLPVRLV | 10016 | 9872 | - | 6 |
| 3 | LSITDIACPLPSY | GAPEHGAGPRAVVDQTARGHGSTER...RESATTHRTGNRPHTERPGPRREAT | 14831 | 14570 | - | 6 |
| 4 | VNSASAVHVLRPPLSITDIACPLPSY | GAPEHGAGPRAVVDQTARGHGSTER...RESATTHRTGNRPHTERPGPRREAT | 14870 | 14570 | - | 6 |
| 5 | MPDGEFGECGPRS | ATPLVHHGHCLPFTVVWGP | 14880 | 14781 | - | 8 |
| 6 | VSLCQHGHSPAMPDGEFGECGPRS | ATPLVHHGHCLPFTVVWGP | 14913 | 14781 | - | 8 |
| 7 | MFQWAGHSHASAVSLCQHGHSPAMPDGEFGECGPRS | ATPLVHHGHCLPFTVVWGP | 14949 | 14781 | - | 10 |
| 8 | VSEVMFQWAGHSHASAVSLCQHGHSPAMPDGEFGECGPRS | ATPLVHHGHCLPFTVVWGP | 14961 | 14781 | - | 10 |
| 9 | MTRPNGETAASPLVETRETRGTGDSVG | GHVSVGGTFPCVSCEPVPTWAFPGDAGR | 15034 | 14866 | - | 8 |
| 10 | MGPMTRPNGETAASPLVETRETRGTGDSV | GGHVSVGGTFPCVSCEPVPTWAFPGDAGR | 15043 | 14866 | - | 8 |
| 11 | VLILAPLDADTPSSFTR | GTLDGAGAPRDQQNCLRAARSVSVPCPFRVHSVSFRTPLEFMTPHRSWAR | 15234 | 15030 | - | 6 |
| 12 | MAVLILAPLDADTPSSFTR | GTLDGAGAPRDQQNCLRAARSVSVPCPFRVHSVSFRTPLEFMTPHRSWAR | 15240 | 15030 | - | 6 |
| 13 | LFPFRRAPEFCFLRCASGDNNARSMSPLPLR | KIPISQVADFTSEFLAKRNGKGVIVVGPTNPPQ | 16903 | 16708 | - | 7 |
| 14 | MSHIAKAVDTLNLPVQEMVLTSVRPTL | GTPRIGRAGRGGGAGRGLAGLGAIA...AVANAAADVAAGAGAGAGAAGAGGA | 16905 | 17229 | + | 15 |
| 15 | LLFPFRRAPEFCFLRCASGDNNARSMSPLPLR | KIPISQVADFTSEFLAKRNGKGVIVVGPTNPPQ | 16906 | 16708 | - | 7 |
| 16 | VRPNGTRSAAPARAGRLTACVAIRNPMIPTA | TTTGQIRKSCIALFFPASGFLKKTLELRLVHGHS | 22392 | 22194 | - | 6 |
| 17 | LCGEAQRHQERGAGEGGETDGLRGDQESDDSDGHDDGAD | QEIVHRPFLSCEWILEKDARTPPGSWPFVELPIRFRRRCR | 22399 | 22159 | - | 6 |
| 18 | VLPLLPELCGEAQRHQERGAGEGGETDGLRGDQESDDSDGHDD | GADQEIVHRPFLSCEWILEKDARTPPGSWPFVELPIRFRRRCR | 22420 | 22159 | - | 6 |
| 19 | VDRAVRINLFFSGRPARVSPWGAVWTRWAVR | RAPRRPESSTRPTVSAVRGVR | 25269 | 25110 | - | 6 |
| 20 | VVDRAVRINLFFSGRPARVSPWGAVWTRWAVR | RAPRRPESSTRPTVSAVRGVR | 25272 | 25110 | - | 6 |
| 21 | VTTTVVDRAVRINLFFSGRPARVSPWGAVWTRWAVR | RAPRRPESSTRPTVSAVRGVR | 25284 | 25110 | - | 6 |
| 22 | VPDLTDRDVYVCGPPGM | TTAVLRTLRGLGVPKQQIHFERFSLAG | 27619 | 27484 | - | 7 |

### Results for GCD35929.1 [Streptomyces chrestomyceticus JCM 4735] back to top

previous - next

#### Architecture

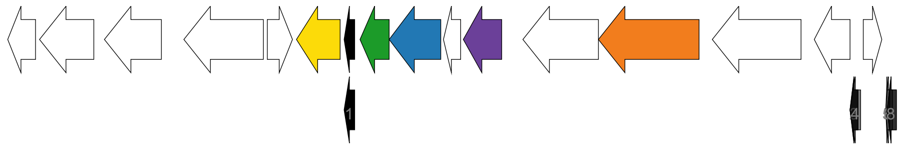

1000 nucleotides

Link to nucleotide sequence

| Accession | start | end | direction | length (aa) | Pfam/HMM | name | description | E-value |
|---|---|---|---|---|---|---|---|---|
| GCD35921.1 | 4290245 | 4289705 | - | 179 | TIGR01203<br>PF00156<br>TIGR01251<br>TIGR01367 | TIGR01203<br>Pribosyltran<br>TIGR01251<br>TIGR01367 | HGPRTase: hypoxanthine phosphoribosyltransferase<br>Phosphoribosyl transferase domain<br>ribP_PPkin: ribose-phosphate diphosphokinase<br>pyrE_Therm: orotate phosphoribosyltransferase | 3.20E-65<br>5.60E-26<br>1.90E-05<br>4.50E-04 |
| GCD35922.1 | 4291422 | 4290333 | - | 362 | TIGR02432<br>PF01171<br>PF09179 | TIGR02432<br>ATP_bind_3<br>TilS | lysidine_TilS_N: tRNA(Ile)-lysidine synthetase<br>PP-loop family<br>TilS substrate binding domain | 5.50E-43<br>3.80E-38<br>3.60E-11 |
| GCD35923.1 | 4292780 | 4291631 | - | 382 | TIGR03883<br>TIGR03624<br>PF10103 | TIGR03883<br>TIGR03624<br>Zincin_2 | DUF2342_F420: uncharacterized protein, coenzyme F420 biosynthesis associated<br>TIGR03624: putative hydrolase<br>Zincin-like metallopeptidase | 7.80E-141<br>1.30E-123<br>6.50E-121 |
| GCD35924.1 | 4294838 | 4293230 | - | 535 | PF02113<br>TIGR00666 | Peptidase_S13<br>TIGR00666 | D-Ala-D-Ala carboxypeptidase 3 (S13) family<br>PBP4: D-alanyl-D-alanine carboxypeptidase/D-alanyl-D-alanine-endopeptidase | 1.30E-78<br>1.80E-55 |
| GCD35925.1 | 4294924 | 4295422 | + | 165 | PF00719 | Pyrophosphatase | Inorganic pyrophosphatase | 2.50E-53 |
| GCD35926.1 | 4296373 | 4295515 | - | 285 | LinG<br>LinH<br>TIGR04033 | LinG<br>LinH<br>TIGR04033 | LinG Homolog<br>LinH (Fusion of LinE/LinG)<br>export_SdpB: antimicrobial peptide system protein, SdpB family | 4.60E-107<br>3.30E-77<br>7.20E-04 |
| GCD35927.1 | 4296663 | 4296471 | - | 63 | LinA | LinA | LinA (linaridin precursor) | 1.10E-10 |
| GCD35928.1 | 4297376 | 4296794 | - | 193 | LinL | LinL | LinL | 8.90E-57 |
| GCD35929.1 | 4298395 | 4297372 | - | 340 | LinE<br>LinH<br>PF12697<br>TIGR02427<br>PF00561 | LinE<br>LinH<br>Abhydrolase_6<br>TIGR02427<br>Abhydrolase_1 | LinE Homolog<br>LinH (Fusion of LinE/LinG)<br>Alpha/beta hydrolase family<br>protocat_pcaD: 3-oxoadipate enol-lactonase<br>alpha/beta hydrolase fold | 9.80E-101<br>4.10E-54<br>4.00E-11<br>4.00E-09<br>1.60E-07 |
| GCD35930.1 | 4298815 | 4298467 | - | 115 | PF01527<br>PF13518<br>PF13551<br>PF13384<br>PF13011 | HTH_Tnp_1<br>HTH_28<br>HTH_29<br>HTH_23<br>LZ_Tnp_IS481 | Transposase<br>Helix-turn-helix domain<br>Winged helix-turn helix<br>Homeodomain-like domain<br>leucine-zipper of insertion element IS481 | 7.40E-09<br>2.90E-07<br>3.40E-05<br>9.60E-05<br>9.10E-04 |

| | | | | | | | | |
|---|---|---|---|---|---|---|---|---|
| GCD35931.1 | 4299629 | 4298861 | - | 255 | PF13649<br>PF08241<br>PF08242<br>PF13489<br>PF13847 | Methyltransf_25<br>Methyltransf_11<br>Methyltransf_12<br>Methyltransf_23<br>Methyltransf_31 | Methyltransferase domain<br>Methyltransferase domain<br>Methyltransferase domain<br>Methyltransferase domain<br>Methyltransferase domain | 5.50E-16<br>6.10E-12<br>1.90E-10<br>2.70E-08<br>2.50E-07 |
| GCD35932.1 | 4301585 | 4300067 | - | 505 | TIGR02734<br>TIGR00292<br>TIGR03467<br>PF13450<br>PF01946 | TIGR02734<br>TIGR00292<br>TIGR03467<br>NAD_binding_8<br>Thi4 | crtI_fam: phytoene desaturase<br>TIGR00292: thiazole biosynthesis enzyme<br>HpnE: squalene-associated FAD-dependent desaturase<br>NAD(P)-binding Rossmann-like domain<br>Thi4 family | 1.60E-08<br>2.40E-08<br>2.90E-08<br>4.50E-08<br>2.20E-06 |
| GCD35933.1 | 4303606 | 4301581 | - | 674 | TIGR02204<br>TIGR02203<br>TIGR00958<br>TIGR03796<br>TIGR03375 | TIGR02204<br>TIGR02203<br>TIGR00958<br>TIGR03796<br>TIGR03375 | MsbA_rel: ABC transporter, permease/ATP-binding protein<br>MsbA_lipidA: lipid A export permease/ATP-binding protein MsbA<br>3a01208: antigen peptide transporter 2<br>NHLM_micro_ABC1: NHLM bacteriocin system ABC transporter, peptidase/ATP-binding protein<br>type_I_sec_LssB: type I secretion system ATPase | 2.70E-97<br>7.30E-96<br>1.30E-89<br>1.70E-82<br>8.30E-79 |
| GCD35934.1 | 4305648 | 4303860 | - | 595 | PF06738<br>PF12821 | ThrE<br>ThrE_2 | Putative threonine/serine exporter<br>Threonine/Serine exporter, ThrE | 2.00E-72<br>1.50E-18 |
| GCD35935.1 | 4306631 | 4305911 | - | 239 | PF09335 | SNARE_assoc | SNARE associated Golgi protein | 4.50E-25 |
| GCD35936.1 | 4306910 | 4307282 | + | 123 | PF01906 | YbjQ_1 | Putative heavy-metal-binding | 1.10E-33 |

| index | leader | core | start | end | dir | score |
|---|---|---|---|---|---|---|
| 1 | MSILADFANAELTDVKPGRIGADSTPTM | VTPLATIATPEATPVGFAATSATAVALDKVTHNIG | 4296663 | 4296471 | - | 33 |
| 2 | MSGRGGARRGVCGAVAGGWNHGSSSGVDSFGPHTAPASR | RTARSGALDPGAFTP | 4306792 | 4306627 | - | 8 |
| 3 | VMSGRGGARRGVCGAVAGGWNHGSSSGVDSFGPHTAPASR | RTARSGALDPGAFTP | 4306795 | 4306627 | - | 8 |
| 4 | MGEGRFGGSGGAGETGAVMSGRGGA...VAGGWNHGSSSGVDSFGPHTAPASR | RTARSGALDPGAFTP | 4306846 | 4306627 | - | 7 |
| 5 | MRPSGRGWRSICGTRC | WPTPYVTVESVRGRGV | 4307446 | 4307347 | - | 7 |
| 6 | MSRTSASPPLMRPSGRGWRSI | CGTRCWPTPYVTVESVRGRGV | 4307476 | 4307347 | - | 6 |
| 7 | VAEEHRRFISSGYYDCSYEMHTCLGEMYVA | DERFTATYEAIRPGLAVYMRDAMLANAVRNS | 4307558 | 4307372 | - | 7 |
| 8 | MDVAEEHRRFISSGYYDCSYEMHTCLGEMYV | ADERFTATYEAIRPGLAVYMRDAMLANAVRNS | 4307564 | 4307372 | - | 8 |

### Results for WP_150482280.1 [Streptomyces coeruleorubidus] back to top

previous - next

#### Architecture

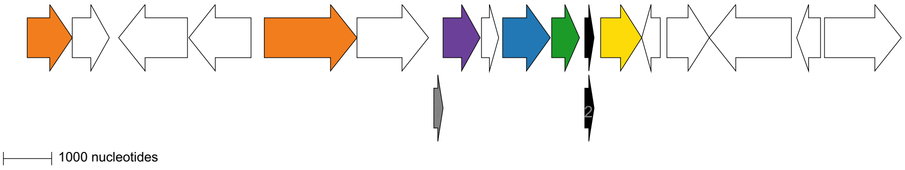

Link to nucleotide sequence

| Accession | start | end | direction | length (aa) | Pfam/HMM | name | description | E-value |
|---|---|---|---|---|---|---|---|---|
| WP_167537402.1 | 5250293 | 5251244 | + | 316 | TIGR01188<br>TIGR01288<br>TIGR03864<br>TIGR03522<br>TIGR03740 | TIGR01188<br>TIGR01288<br>TIGR03864<br>TIGR03522<br>TIGR03740 | drrA: daunorubicin resistance ABC transporter, ATP-binding protein<br>nodI: nodulation ABC transporter NodI<br>PQQ_ABC_ATP: ABC transporter, ATP-binding subunit, PQQ-dependent alcohol dehydrogenase system<br>GldA_ABC_ATP: gliding motility-associated ABC transporter ATP-binding subunit GldA<br>galliderm_ABC: lantibiotic protection ABC transporter, ATP-binding subunit | 1.10E-105<br>2.30E-61<br>7.60E-60<br>8.40E-51<br>2.00E-45 |
| WP_167537403.1 | 5251240 | 5252002 | + | 253 | PF01061<br>TIGR00025<br>PF12698<br>TIGR01247<br>TIGR01248 | ABC2_membrane<br>TIGR00025<br>ABC2_membrane_3<br>TIGR01247<br>TIGR01248 | ABC-2 type transporter<br>Mtu_efflux: ABC transporter efflux protein, DrrB family<br>ABC-2 family transporter protein<br>drrB: daunorubicin resistance ABC transporter membrane protein<br>drrC: daunorubicin resistance protein C | 1.80E-27<br>3.30E-24<br>2.40E-13<br>4.20E-12<br>6.10E-10 |
| WP_150482275.1 | 5253604 | 5252197 | - | 468 | TIGR01788<br>PF00282<br>TIGR03812<br>TIGR03799 | TIGR01788<br>Pyridoxal_deC<br>TIGR03812<br>TIGR03799 | Glu-decarb-GAD: glutamate decarboxylase<br>Pyridoxal-dependent decarboxylase conserved domain<br>tyr_de_CO2_Arch: tyrosine decarboxylase MnfA<br>NOD_PanD_pyr: putative pyridoxal-dependent aspartate 1-decarboxylase | 8.00E-187<br>1.70E-70<br>1.30E-38<br>1.30E-06 |
| WP_150482276.1 | 5254923 | 5253636 | - | 428 | PF00654 | Voltage_CLC | Voltage gated chloride channel | 1.50E-23 |
| WP_167537404.1 | 5255195 | 5257130 | + | 644 | TIGR02204<br>TIGR02203<br>TIGR00958<br>TIGR03796<br>TIGR03375 | TIGR02204<br>TIGR02203<br>TIGR00958<br>TIGR03796<br>TIGR03375 | MsbA_rel: ABC transporter, permease/ATP-binding protein<br>MsbA_lipidA: lipid A export permease/ATP-binding protein MsbA<br>3a01208: antigen peptide transporter 2<br>NHLM_micro_ABC1: NHLM bacteriocin system ABC transporter, peptidase/ATP-binding protein<br>type_I_sec_LssB: type I secretion system ATPase | 1.00E-98<br>4.80E-93<br>2.70E-86<br>3.60E-81<br>5.60E-78 |
| WP_150482277.1 | 5257126 | 5258611 | + | 494 | TIGR00292<br>TIGR02734<br>TIGR03467<br>PF01946<br>PF13450 | TIGR00292<br>TIGR02734<br>TIGR03467<br>Thi4<br>NAD_binding_8 | TIGR00292: thiazole biosynthesis enzyme<br>crtI_fam: phytoene desaturase<br>HpnE: squalene-associated FAD-dependent desaturase<br>Thi4 family<br>NAD(P)-binding Rossmann-like domain | 8.50E-09<br>2.00E-08<br>7.60E-08<br>7.60E-07<br>1.50E-06 |
| WP_150482278.1 | 5258908 | 5259667 | + | 252 | PF13649<br>PF08241<br>PF08242<br>PF13489<br>TIGR01983 | Methyltransf_25<br>Methyltransf_11<br>Methyltransf_12<br>Methyltransf_23<br>TIGR01983 | Methyltransferase domain<br>Methyltransferase domain<br>Methyltransferase domain<br>Methyltransferase domain<br>UbiG: 3-demethylubiquinone-9 3-O-methyltransferase | 5.10E-15<br>1.60E-10<br>1.20E-08<br>4.10E-08<br>2.20E-05 |

| | | | | | | | | |
|---|---|---|---|---|---|---|---|---|
| WP_150482279.1 | 5259710 | 5260058 | + | 115 | PF13518<br>PF01527<br>PF13551<br>PF13011<br>PF13384 | HTH_28<br>HTH_Tnp_1<br>HTH_29<br>LZ_Tnp_IS481<br>HTH_23 | Helix-turn-helix domain<br>Transposase<br>Winged helix-turn helix<br>leucine-zipper of insertion element IS481<br>Homeodomain-like domain | 6.90E-09<br>5.30E-08<br>2.10E-06<br>4.30E-06<br>3.00E-05 |
| WP_150482280.1 | 5260131 | 5261139 | + | 335 | LinE<br>LinH<br>PF12697<br>TIGR02427<br>PF00561 | LinE<br>LinH<br>Abhydrolase_6<br>TIGR02427<br>Abhydrolase_1 | LinE Homolog<br>LinH (Fusion of LinE/LinG)<br>Alpha/beta hydrolase family<br>protocat_pcaD: 3-oxoadipate enol-lactonase<br>alpha/beta hydrolase fold | 1.70E-101<br>3.40E-52<br>1.80E-14<br>2.80E-11<br>1.30E-10 |
| WP_150485486.1 | 5261162 | 5261723 | + | 186 | LinL | LinL | LinL | 1.90E-55 |
| WP_167537405.1 | 5261853 | 5262036 | + | 60 | LinA | LinA | LinA (linaridin precursor) | 5.20E-13 |
| WP_150482282.1 | 5262179 | 5263010 | + | 276 | LinG<br>LinH | LinG<br>LinH | LinG Homolog<br>LinH (Fusion of LinE/LinG) | 3.60E-109<br>6.50E-79 |
| WP_150482283.1 | 5263394 | 5263019 | - | 124 | PF01906<br>TIGR03884 | YbjQ_1<br>TIGR03884 | Putative heavy-metal-binding<br>sel_bind_Methan: selenium-binding protein | 2.60E-33<br>9.60E-05 |
| WP_150482284.1 | 5263529 | 5264405 | + | 291 | PF09335 | SNARE_assoc | SNARE associated Golgi protein | 1.60E-24 |
| WP_150482285.1 | 5266127 | 5264429 | - | 565 | PF06738<br>PF12821 | ThrE<br>ThrE_2 | Putative threonine/serine exporter<br>Threonine/Serine exporter, ThrE | 3.30E-69<br>2.40E-17 |
| WP_031487466.1 | 5266732 | 5266240 | - | 163 | PF00719 | Pyrophosphatase | Inorganic pyrophosphatase | 2.60E-53 |
| WP_150482286.1 | 5266811 | 5268386 | + | 524 | PF02113<br>TIGR00666<br>PF00768 | Peptidase_S13<br>TIGR00666<br>Peptidase_S11 | D-Ala-D-Ala carboxypeptidase 3 (S13) family<br>PBP4: D-alanyl-D-alanine carboxypeptidase/D-alanyl-D-alanine-endopeptidase<br>D-alanyl-D-alanine carboxypeptidase | 7.80E-74<br>2.90E-57<br>1.90E-08 |

| index | leader | core | start | end | dir | score |
|---|---|---|---|---|---|---|
| 1 | VEPTKLPTSSVEALPCCCLTNHAGGCGDF | RDRCSRHPTSRPAPTPRLHDAPIGMAVGTS | 5258715 | 5258895 | + | 7 |
| 2 | MSVVADFANTELADVTPGRIGNEATPTM | LTPTATLATPEGVAVTAATAYALNEVTNDLAG | 5261853 | 5262036 | + | 33 |

### Results for WP_046706697.1 [Streptomyces europaeiscabiei] back to top

previous - next

#### Architecture

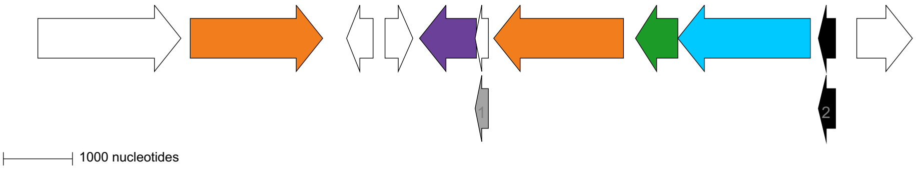

Link to nucleotide sequence

| Accession | start | end | direction | length (aa) | Pfam/HMM | name | description | E-value |
|---|---|---|---|---|---|---|---|---|
| WP_060904367.1 | 3375 | 5454 | + | 692 | PF07228<br>PF08448<br>TIGR00229<br>PF00989<br>PF01590 | SpoIIE<br>PAS_4<br>TIGR00229<br>PAS<br>GAF | Stage II sporulation protein E (SpoIIE)<br>PAS fold<br>sensory_box: PAS domain S-box protein<br>PAS fold<br>GAF domain | 1.60E-52<br>6.90E-31<br>8.70E-27<br>1.50E-18<br>2.80E-12 |
| WP_060904368.1 | 5590 | 7510 | + | 639 | TIGR02204<br>TIGR02203<br>TIGR01846<br>TIGR00958<br>TIGR03375 | TIGR02204<br>TIGR02203<br>TIGR01846<br>TIGR00958<br>TIGR03375 | MsbA_rel: ABC transporter, permease/ATP-binding protein<br>MsbA_lipidA: lipid A export permease/ATP-binding protein MsbA<br>type_I_sec_HlyB: type I secretion system ATPase<br>3a01208: antigen peptide transporter 2<br>type_I_sec_LssB: type I secretion system ATPase | 6.50E-101<br>1.40E-97<br>1.20E-88<br>6.00E-84<br>2.10E-77 |
| WP_046706703.1 | 8233 | 7861 | - | 123 | PF00717<br>TIGR02228 | Peptidase_S24<br>TIGR02228 | Peptidase S24-like<br>sigpep_I_arch: signal peptidase I | 7.50E-04<br>9.40E-04 |
| WP_046706702.1 | 8415 | 8811 | + | 131 | PF00583<br>PF13673<br>TIGR01575<br>PF13508<br>TIGR03448 | Acetyltransf_1<br>Acetyltransf_10<br>TIGR01575<br>Acetyltransf_7<br>TIGR03448 | Acetyltransferase (GNAT) family<br>Acetyltransferase (GNAT) domain<br>rimI: ribosomal-protein-alanine acetyltransferase<br>Acetyltransferase (GNAT) domain<br>mycothiol_MshD: mycothiol synthase | 5.50E-12<br>1.00E-05<br>6.60E-05<br>4.70E-04<br>5.90E-04 |
| WP_046706701.1 | 9722 | 8915 | - | 268 | PF13649<br>PF08241<br>PF13489<br>PF13847<br>PF08242 | Methyltransf_25<br>Methyltransf_11<br>Methyltransf_23<br>Methyltransf_31<br>Methyltransf_12 | Methyltransferase domain<br>Methyltransferase domain<br>Methyltransferase domain<br>Methyltransferase domain<br>Methyltransferase domain | 2.00E-14<br>2.50E-11<br>8.40E-08<br>1.40E-06<br>1.20E-05 |
| WP_046706700.1 | 9910 | 9718 | - | 63 | NO PFAM MATCH | - | - | - |
| WP_046706699.1 | 11867 | 9977 | - | 629 | TIGR02204<br>TIGR02203<br>TIGR00958<br>TIGR03796<br>TIGR01846 | TIGR02204<br>TIGR02203<br>TIGR00958<br>TIGR03796<br>TIGR01846 | MsbA_rel: ABC transporter, permease/ATP-binding protein<br>MsbA_lipidA: lipid A export permease/ATP-binding protein MsbA<br>3a01208: antigen peptide transporter 2<br>NHLM_micro_ABC1: NHLM bacteriocin system ABC transporter, peptidase/ATP-binding protein<br>type_I_sec_HlyB: type I secretion system ATPase | 1.50E-104<br>5.10E-102<br>2.40E-93<br>8.30E-85<br>1.70E-79 |
| WP_046706698.1 | 12655 | 12034 | - | 206 | LinL | LinL | LinL | 7.80E-71 |
| WP_046706697.1 | 14571 | 12651 | - | 639 | LinH<br>LinG<br>LinE<br>PF12697<br>PF12146 | LinH<br>LinG<br>LinE<br>Abhydrolase_6<br>Hydrolase_4 | LinH (Fusion of LinE/LinG)<br>LinG Homolog<br>LinE Homolog<br>Alpha/beta hydrolase family<br>Serine aminopeptidase, S33 | 6.10E-247<br>1.20E-160<br>5.00E-92<br>1.10E-17<br>2.80E-09 |

| | | | | | | | | |
|---|---|---|---|---|---|---|---|---|
| WP_046706714.1 | 14938 | 14686 | - | 83 | LinA | LinA | LinA (linaridin precursor) | 1.80E-04 |
| WP_013005807.1 | 15230 | 16033 | + | 267 | INFERRED GENE | - | - | - |

| index | leader | core | start | end | dir | score |
|---|---|---|---|---|---|---|
| 1 | MNDFLLIPSVVALGIVGFLIATRAVSTPAVI | GVALVVLWGACSQARSTIHTRNTPSKKRRQHP | 9910 | 9718 | - | 6 |
| 2 | MEIPVEGVVYVAARPTL | GTPRIARVGNRLGGLAQAAEGVGAI...NNLAAEANAQNAAALAAVGGAPPAA | 14938 | 14686 | - | 28 |

### Results for WP_060892339.1 [Streptomyces europaeiscabiei] back to top

previous - next

#### Architecture

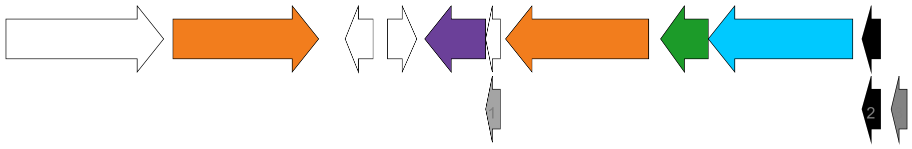

1000 nucleotides

Link to nucleotide sequence

| Accession | start | end | direction | length (aa) | Pfam/HMM | name | description | E-value |
|---|---|---|---|---|---|---|---|---|
| WP_060892334.1 | 3741 | 5820 | + | 692 | PF07228<br>PF08448<br>TIGR00229<br>PF00989<br>PF01590 | SpoIIE<br>PAS_4<br>TIGR00229<br>PAS<br>GAF | Stage II sporulation protein E (SpoIIE)<br>PAS fold<br>sensory_box: PAS domain S-box protein<br>PAS fold<br>GAF domain | 1.00E-51<br>6.50E-31<br>8.90E-27<br>1.50E-18<br>3.10E-12 |
| WP_060892335.1 | 5956 | 7876 | + | 639 | TIGR02204<br>TIGR02203<br>TIGR01846<br>TIGR00958<br>TIGR03375 | TIGR02204<br>TIGR02203<br>TIGR01846<br>TIGR00958<br>TIGR03375 | MsbA_rel: ABC transporter, permease/ATP-binding protein<br>MsbA_lipidA: lipid A export permease/ATP-binding protein MsbA<br>type_I_sec_HlyB: type I secretion system ATPase<br>3a01208: antigen peptide transporter 2<br>type_I_sec_LssB: type I secretion system ATPase | 4.20E-102<br>9.10E-98<br>7.10E-89<br>6.70E-85<br>1.00E-77 |
| WP_060892336.1 | 8599 | 8227 | - | 123 | PF00717<br>TIGR02228 | Peptidase_S24<br>TIGR02228 | Peptidase S24-like<br>sigpep_I_arch: signal peptidase I | 7.40E-04<br>9.10E-04 |
| WP_046706702.1 | 8781 | 9177 | + | 131 | PF00583<br>PF13673<br>TIGR01575<br>PF13508<br>TIGR03448 | Acetyltransf_1<br>Acetyltransf_10<br>TIGR01575<br>Acetyltransf_7<br>TIGR03448 | Acetyltransferase (GNAT) family<br>Acetyltransferase (GNAT) domain<br>rimI: ribosomal-protein-alanine acetyltransferase<br>Acetyltransferase (GNAT) domain<br>mycothiol_MshD: mycothiol synthase | 5.50E-12<br>1.00E-05<br>6.60E-05<br>4.70E-04<br>5.90E-04 |
| WP_046706701.1 | 10089 | 9282 | - | 268 | PF13649<br>PF08241<br>PF13489<br>PF13847<br>PF08242 | Methyltransf_25<br>Methyltransf_11<br>Methyltransf_23<br>Methyltransf_31<br>Methyltransf_12 | Methyltransferase domain<br>Methyltransferase domain<br>Methyltransferase domain<br>Methyltransferase domain<br>Methyltransferase domain | 2.00E-14<br>2.50E-11<br>8.40E-08<br>1.40E-06<br>1.20E-05 |
| WP_046706700.1 | 10277 | 10085 | - | 63 | NO PFAM MATCH | - | - | - |
| WP_060892337.1 | 12234 | 10344 | - | 629 | TIGR02204<br>TIGR02203<br>TIGR00958<br>TIGR03796<br>TIGR01846 | TIGR02204<br>TIGR02203<br>TIGR00958<br>TIGR03796<br>TIGR01846 | MsbA_rel: ABC transporter, permease/ATP-binding protein<br>MsbA_lipidA: lipid A export permease/ATP-binding protein MsbA<br>3a01208: antigen peptide transporter 2<br>NHLM_micro_ABC1: NHLM bacteriocin system ABC transporter, peptidase/ATP-binding protein<br>type_I_sec_HlyB: type I secretion system ATPase | 2.40E-104<br>3.20E-102<br>1.70E-92<br>1.00E-84<br>2.40E-79 |
| WP_060892338.1 | 13022 | 12401 | - | 206 | LinL | LinL | LinL | 7.60E-71 |
| WP_060892339.1 | 14938 | 13018 | - | 639 | LinH<br>LinG<br>LinE<br>PF12697<br>PF12146 | LinH<br>LinG<br>LinE<br>Abhydrolase_6<br>Hydrolase_4 | LinH (Fusion of LinE/LinG)<br>LinG Homolog<br>LinE Homolog<br>Alpha/beta hydrolase family<br>Serine aminopeptidase, S33 | 4.50E-245<br>3.40E-159<br>8.80E-92<br>3.10E-17<br>7.50E-09 |
| WP_046706714.1 | 15305 | 15053 | - | 83 | LinA | LinA | LinA (linaridin precursor) | 1.80E-04 |

| index | leader | core | start | end | dir | score |
|---|---|---|---|---|---|---|
| 1 | MNDFLLIPSVVALGIVGFLIATRAVSTPAVI | GVALVVLWGACSQARSTIHTRNTPSKKRRQHP | 10277 | 10085 | - | 6 |
| 2 | MEIPVEGVVYVAARPTL | GTPRIARVGNRLGGLAQAAEGVGAI...NNLAAEANAQNAAALAAVGGAPPAA | 15305 | 15053 | - | 28 |
| 3 | VARTPPRRPCVISGDGTGEDEPGGHGPHQSRKKAR | TDSVRAFFRAVVRAAACRPDSFPAKSRSGVKGGTEQ | 15657 | 15441 | - | 7 |

### Results for WP_119581921.1 [Streptomyces europaeiscabiei] back to top

previous - next

#### Architecture

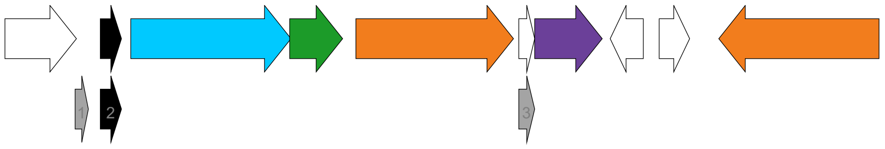

1000 nucleotides

Link to nucleotide sequence

| Accession | start | end | direction | length (aa) | Pfam/HMM | name | description | E-value |
|---|---|---|---|---|---|---|---|---|
| WP_162948329.1 | 0 | 851 | + | 282 | NO PFAM MATCH | - | - | - |
| WP_046706714.1 | 1142 | 1394 | + | 83 | LinA | LinA | LinA (linaridin precursor) | 1.80E-04 |
| WP_119581921.1 | 1509 | 3429 | + | 639 | LinH | LinH | LinH (Fusion of LinE/LinG) | 4.60E-246 |
| | | | | | LinG | LinG | LinG Homolog | 1.20E-160 |
| | | | | | LinE | LinE | LinE Homolog | 1.40E-91 |
| | | | | | PF12697 | Abhydrolase_6 | Alpha/beta hydrolase family | 8.20E-18 |
| | | | | | PF12146 | Hydrolase_4 | Serine aminopeptidase, S33 | 2.60E-09 |
| WP_060892338.1 | 3425 | 4046 | + | 206 | LinL | LinL | LinL | 7.60E-71 |
| WP_119581922.1 | 4213 | 6103 | + | 629 | TIGR02204 | TIGR02204 | MsbA_rel: ABC transporter, permease/ATP-binding protein | 3.30E-104 |
| | | | | | TIGR02203 | TIGR02203 | MsbA_lipidA: lipid A export permease/ATP-binding protein MsbA | 5.50E-102 |
| | | | | | TIGR00958 | TIGR00958 | 3a01208: antigen peptide transporter 2 | 2.30E-92 |
| | | | | | TIGR03796 | TIGR03796 | NHLM_micro_ABC1: NHLM bacteriocin system ABC transporter, peptidase/ATP-binding protein | 3.10E-85 |
| | | | | | TIGR01846 | TIGR01846 | type_I_sec_HlyB: type I secretion system ATPase | 5.20E-79 |
| WP_046706700.1 | 6170 | 6362 | + | 63 | NO PFAM MATCH | - | - | - |
| WP_119581923.1 | 6358 | 7165 | + | 268 | PF13649 | Methyltransf_25 | Methyltransferase domain | 1.70E-14 |
| | | | | | PF08241 | Methyltransf_11 | Methyltransferase domain | 2.50E-11 |
| | | | | | PF13489 | Methyltransf_23 | Methyltransferase domain | 7.40E-08 |
| | | | | | PF13847 | Methyltransf_31 | Methyltransferase domain | 1.60E-06 |
| | | | | | PF08242 | Methyltransf_12 | Methyltransferase domain | 1.20E-05 |
| WP_046706702.1 | 7665 | 7269 | - | 131 | PF00583 | Acetyltransf_1 | Acetyltransferase (GNAT) family | 5.50E-12 |
| | | | | | PF13673 | Acetyltransf_10 | Acetyltransferase (GNAT) domain | 1.00E-05 |
| | | | | | TIGR01575 | TIGR01575 | rimI: ribosomal-protein-alanine acetyltransferase | 6.60E-05 |
| | | | | | PF13508 | Acetyltransf_7 | Acetyltransferase (GNAT) domain | 4.70E-04 |
| | | | | | TIGR03448 | TIGR03448 | mycothiol_MshD: mycothiol synthase | 5.90E-04 |
| WP_046706703.1 | 7847 | 8219 | + | 123 | PF00717 | Peptidase_S24 | Peptidase S24-like | 7.50E-04 |
| | | | | | TIGR02228 | TIGR02228 | sigpep_I_arch: signal peptidase I | 9.40E-04 |
| WP_119581924.1 | 10490 | 8570 | - | 639 | TIGR02204 | TIGR02204 | MsbA_rel: ABC transporter, permease/ATP-binding protein | 3.30E-102 |
| | | | | | TIGR02203 | TIGR02203 | MsbA_lipidA: lipid A export permease/ATP-binding protein MsbA | 1.00E-98 |
| | | | | | TIGR01846 | TIGR01846 | type_I_sec_HlyB: type I secretion system ATPase | 8.90E-89 |
| | | | | | TIGR00958 | TIGR00958 | 3a01208: antigen peptide transporter 2 | 8.50E-85 |
| | | | | | TIGR03375 | TIGR03375 | type_I_sec_LssB: type I secretion system ATPase | 9.20E-78 |

| index | leader | core | start | end | dir | score |
|---|---|---|---|---|---|---|
| 1 | MNREAAGPHQSRKKARTDSVRAFFRA | VVRAAACRPDSFPAKSRSGVKGGTEQ | 847 | 1006 | + | 6 |
| 2 | MEIPVEGVVYVAARPTL | GTPRIARVGNRLGGLAQAAEGVGAI...NNLAAEANAQNAAALAAVGGAPPAA | 1142 | 1394 | + | 28 |
| 3 | MNDFLLIPSVVALGIVGFLIATRAVSTPAVI | GVALVVLWGACSQARSTIHTRNTPSKKRRQHP | 6170 | 6362 | + | 6 |

### Results for KND25157.1 [Streptomyces europaeiscabiei] back to top

previous - next

#### Architecture

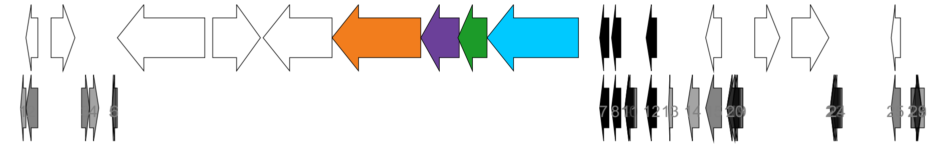

1000 nucleotides

Link to nucleotide sequence

| Accession | start | end | direction | length (aa) | Pfam/HMM | name | description | E-value |
|---|---|---|---|---|---|---|---|---|
| KND25149.1 | 1081220 | 1080971 | - | 82 | NO PFAM MATCH | - | - | - |
| KND25150.1 | 1081508 | 1082015 | + | 168 | PF12840 | HTH_20 | Helix-turn-helix domain | 3.10E-10 |
| | | | | | PF01022 | HTH_5 | Bacterial regulatory protein, arsR family | 2.90E-05 |
| KND25151.1 | 1084752 | 1082898 | - | 617 | NO PFAM MATCH | - | - | - |
| KND25152.1 | 1084941 | 1085955 | + | 337 | NO PFAM MATCH | - | - | - |
| KND25153.1 | 1087470 | 1086009 | - | 486 | TIGR02734 | TIGR02734 | crtI_fam: phytoene desaturase | 3.50E-11 |
| | | | | | PF13450 | NAD_binding_8 | NAD(P)-binding Rossmann-like domain | 6.40E-09 |
| | | | | | TIGR01317 | TIGR01317 | GOGAT_sm_gam: glutamate synthase, NADH/NADPH, small subunit | 1.20E-08 |
| | | | | | TIGR00292 | TIGR00292 | TIGR00292: thiazole biosynthesis enzyme | 1.40E-08 |
| | | | | | TIGR03467 | TIGR03467 | HpnE: squalene-associated FAD-dependent desaturase | 7.40E-08 |
| KND25154.1 | 1089347 | 1087475 | - | 623 | TIGR02204 | TIGR02204 | MsbA_rel: ABC transporter, permease/ATP-binding protein | 1.20E-103 |
| | | | | | TIGR02203 | TIGR02203 | MsbA_lipidA: lipid A export permease/ATP-binding protein MsbA | 9.90E-97 |
| | | | | | TIGR00958 | TIGR00958 | 3a01208: antigen peptide transporter 2 | 8.80E-85 |
| | | | | | TIGR03796 | TIGR03796 | NHLM_micro_ABC1: NHLM bacteriocin system ABC transporter, peptidase/ATP-binding protein | 8.50E-81 |
| | | | | | TIGR03375 | TIGR03375 | type_I_sec_LssB: type I secretion system ATPase | 2.20E-80 |
| KND25155.1 | 1090153 | 1089343 | - | 269 | PF13649 | Methyltransf_25 | Methyltransferase domain | 2.30E-16 |
| | | | | | PF08241 | Methyltransf_11 | Methyltransferase domain | 7.40E-12 |
| | | | | | PF08242 | Methyltransf_12 | Methyltransferase domain | 3.30E-08 |
| | | | | | PF13489 | Methyltransf_23 | Methyltransferase domain | 2.70E-07 |
| | | | | | PF13847 | Methyltransf_31 | Methyltransferase domain | 1.80E-06 |
| KND25156.1 | 1090758 | 1090149 | - | 202 | LinL | LinL | LinL | 1.90E-65 |

| | | | | | | | | |
|---|---|---|---|---|---|---|---|---|
| KND25157.1 | 1092710 | 1090754 | - | 651 | LinH<br>LinG<br>LinE<br>LinL<br>PF12697 | LinH<br>LinG<br>LinE<br>LinL<br>Abhydrolase_6 | LinH (Fusion of LinE/LinG)<br>LinG Homolog<br>LinE Homolog<br>LinL<br>Alpha/beta hydrolase family | 3.50E-257<br>5.40E-114<br>3.60E-95<br>1.30E-36<br>6.60E-15 |
| KND25158.1 | 1093350 | 1093149 | - | 66 | LinA | LinA | LinA (linaridin precursor) | 3.70E-14 |
| KND25159.1 | 1093602 | 1093407 | - | 64 | LinA | LinA | LinA (linaridin precursor) | 7.70E-11 |
| KND25160.1 | 1094344 | 1094140 | - | 67 | LinA | LinA | LinA (linaridin precursor) | 8.70E-12 |
| KND25161.1 | 1095724 | 1095403 | - | 106 | NO PFAM MATCH | - | - | - |
| KND25162.1 | 1096432 | 1096978 | + | 181 | NO PFAM MATCH | - | - | - |
| KND25163.1 | 1097230 | 1098007 | + | 258 | NO PFAM MATCH | - | - | - |
| KND25164.1 | 1099543 | 1099339 | - | 67 | NO PFAM MATCH | - | - | - |

| index | leader | core | start | end | dir | score |
|---|---|---|---|---|---|---|
| 1 | VDDVPLAPGGLVVRGPG | TEAAARWRRVPMSGLTSP | 1080956 | 1080848 | - | 6 |
| 2 | MIKNFVRGFSALSLALVPLL | SAPAAQAAPAAEVTTLTDAVGPVQV...SHGVGVIALTPYGVAHGVPTGSYRV | 1081220 | 1080971 | - | 7 |
| 3 | LTGCSRAGPQRVAAAASASRPRPPSRAG | ACSASSAVVPTRITDIVKAVVVPHHAST | 1082139 | 1082310 | + | 7 |
| 4 | MRPKRAPSSLAQHRFWELPYKQTPFAFRT | QTPEPSSASPAGASTSCSPCSATAPSTNPEPRTPTVTLAA | 1082306 | 1082516 | + | 6 |
| 5 | MTAAAVGGGPGASAVDL | ARLSPAGTRSCGTVTKV | 1082887 | 1082782 | - | 7 |
| 6 | MRGAGMTAAAVGGGPGASA | VDLARLSPAGTRSCGTVTKV | 1082902 | 1082782 | - | 6 |
| 7 | MSMSPTPAALRGTGGLSESDPGRALSSLAPVA | ATPGVVAGVAVGVALVNAFAAGYNHCGGNVELPM | 1093350 | 1093149 | - | 31 |
| 8 | MNKSSAPAVLTATGALTESAPGVTLPTLAPVA | ATPVAIAATMGVAFVAGYAAGRAATGNVELPM | 1093602 | 1093407 | - | 30 |
| 9 | LNTASIPAVLSETGALSESDHGRAPLD | TAPVASVTFTMTACVQVSVCLTGSVIQLPQ | 1093847 | 1093673 | - | 9 |
| 10 | MTSPHRPPTSEEKSLNTASIPAVLSETGALSESDHGRAPLD | TAPVASVTFTMTACVQVSVCLTGSVIQLPQ | 1093889 | 1093673 | - | 8 |
| 11 | VGPSRRRRRPPMTSPHRPPTSEEKS...TASIPAVLSETGALSESDHGRAPLD | TAPVASVTFTMTACVQVSVCLTGSVIQLPQ | 1093922 | 1093673 | - | 7 |
| 12 | MPEFREPGWTRGIAPLQEDAGGQVFGGASPVA | ATPAVVATAGAVVVAFAAGVAARHLANGGNVELPM | 1094344 | 1094140 | - | 30 |
| 13 | MATLKGCRLLRKLRCST | NRLTDLVKAVLALHHASA | 1094709 | 1094601 | - | 6 |
| 14 | MPDPAPDRWGGVDQPHELRDVVAVAACEDQGERSAVSVGDQVVLRPGS | ATICQTRSRPAPPLWRARGWSRPPHATSRDGPPR | 1095251 | 1095002 | - | 6 |
| 15 | MAGDPAEDVYLDPGVGEPGECGVPE...AQMLVAELGDHGAPMCGVAQDGGGD | TSASGAREQAGVRMGPGEDALGDEG...STTGTLRSRLPLVLLSVSPPGAGVV | 1095724 | 1095403 | - | 7 |
| 16 | VHDACLHAQLTPESSLLYPKIPLT | LTPGSQGVAGSNPVVPTGDSRRSRA...RPLIIFGDQLGTSWRFGVLDRVGGS | 1096094 | 1095848 | - | 7 |
| 17 | VVHDACLHAQLTPESSLLYPKIPLT | LTPGSQGVAGSNPVVPTGDSRRSRA...RPLIIFGDQLGTSWRFGVLDRVGGS | 1096097 | 1095848 | - | 7 |
| 18 | VGTSWGPHATRGGARCMPTRTTHARIKSAISE | NTPDPYSWESRGRRFKSCRPDWRQS | 1096131 | 1095957 | - | 6 |
| 19 | VLARGVGRTTGRGCVGTSWGPHATRGGARCMPTRTTHARIKSAISE | NTPDPYSWESRGRRFKSCRPDWRQS | 1096173 | 1095957 | - | 6 |
| 20 | LRCSREASAGQREGGAWGPAGDHTL...VVHDACLHAQLTPESSLLYPKIPLT | LTPGSQGVAGSNPVVPTGDSRRSRA...RPLIIFGDQLGTSWRFGVLDRVGGS | 1096178 | 1095848 | - | 6 |
| 21 | VARPGQEPHSRIWSCHAACDLQHADGSRID | RPGRQLCVTSPVVESCDEAATRRARSQEAA | 1098225 | 1098042 | - | 8 |
| 22 | VPEHRLRAVARPGQEPHSRIWSCHAACDLQHADG | SRIDRPGRQLCVTSPVVESCDEAATRRARSQEAA | 1098249 | 1098042 | - | 7 |
| 23 | MATPASSVPEHRLRAVARPGQEPHSRIWSCHAACDLQ | HADGSRIDRPGRQLCVTSPVVESCDEAATRRARSQEAA | 1098270 | 1098042 | - | 6 |
| 24 | MDTWPRPWLPPLRRCQSTGFEPWHAPARSRT | LAFGRVTRHAISNTPTGLGSTVRDGNSASRAR | 1098290 | 1098098 | - | 6 |
| 25 | MIADLLPTIADFDDEQTALICRRVDLQSDIGVW | GQIIADLEVFTEVRQVLEVHLSKTSPDGEPDALA | 1099543 | 1099339 | - | 7 |
| 26 | VPVIPLPLGGRAQSSGAVCGGRGWGVQPVTSRLMCGSLV | AGGHEAIAAGGINPAGAGACVSSTRRSLLARLVVEALPR | 1099747 | 1099984 | + | 6 |
| 27 | LGGRAQSSGAVCGGRGWGVQPVTSRLMCGSLVAGG | HEAIAAGGINPAGAGACVSSTRRSLLARLVVEALPR | 1099768 | 1099984 | + | 6 |
| 28 | VTSRLMCGSLVAGGHEAIAAGGINP | AGAGACVSSTRRSLLARLVVEALPR | 1099831 | 1099984 | + | 6 |
| 29 | VCPAALAAGPARSCGAYPRCVDLSR...CRRHARACPRRVDPARRDRLVPASD | QTAAHQAAGHRLDAPPPATAYSP | 1100050 | 1099789 | - | 6 |

### Results for WP_024756459.1 [Streptomyces exfoliatus DSM 41693] back to top

previous - next

#### Architecture

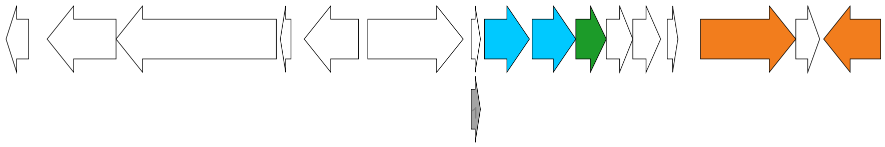

1000 nucleotides

Link to nucleotide sequence

| Accession | start | end | direction | length (aa) | Pfam/HMM | name | description | E-value |
|---|---|---|---|---|---|---|---|---|
| WP_024756454.1 | 16293 | 15855 | - | 145 | PF10901 | DUF2690 | Protein of unknown function (DUF2690) | 4.00E-14 |
| WP_024756455.1 | 18002 | 16670 | - | 443 | PF05147<br>TIGR03897 | LANC_like<br>TIGR03897 | Lanthionine synthetase C-like protein<br>lanti_2_LanM: type 2 lantibiotic biosynthesis protein LanM | 5.30E-54<br>6.20E-12 |
| WP_078626944.1 | 21163 | 18007 | - | 1051 | PF04738<br>TIGR03891<br>PF14028 | Lant_dehydr_N<br>TIGR03891<br>Lant_dehydr_C | Lantibiotic dehydratase, N terminus<br>thiopep_ocin: thiopeptide-type bacteriocin biosynthesis domain<br>Lantibiotic biosynthesis dehydratase C-term | 1.20E-158<br>1.10E-51<br>2.30E-33 |
| WP_157848182.1 | 21435 | 21228 | - | 68 | NO PFAM MATCH | - | - | - |
| WP_051494251.1 | 22758 | 21699 | - | 352 | TIGR03558<br>PF00296<br>TIGR03858<br>TIGR04036<br>TIGR04027 | TIGR03558<br>Bac_luciferase<br>TIGR03858<br>TIGR04036<br>TIGR04027 | oxido_grp_1: luciferase family oxidoreductase, group 1<br>Luciferase-like monooxygenase<br>LLM_2I7G: probable oxidoreductase, LLM family<br>LLM_CE1758_fam: putative luciferase-like monooxygenase, FMN-dependent, CE1758 family<br>LLM_KPN_01858: putative FMN-dependent luciferase-like monooxygenase, KPN_01858 family | 3.10E-116<br>2.50E-38<br>4.50E-20<br>7.20E-18<br>5.40E-17 |
| WP_024756457.1 | 22943 | 24803 | + | 619 | PF00326<br>PF02129<br>PF01738<br>PF00561<br>TIGR00976 | Peptidase_S9<br>Peptidase_S15<br>DLH<br>Abhydrolase_1<br>TIGR00976 | Prolyl oligopeptidase family<br>X-Pro dipeptidyl-peptidase (S15 family)<br>Dienelactone hydrolase family<br>alpha/beta hydrolase fold<br>/NonD: hydrolase CocE/NonD family protein | 1.40E-49<br>3.80E-16<br>1.10E-12<br>2.90E-11<br>1.20E-09 |
| WP_157848183.1 | 24971 | 25160 | + | 62 | NO PFAM MATCH | - | - | - |
| WP_024756458.1 | 25229 | 26105 | + | 291 | LinH<br>LinG | LinH<br>LinG | LinH (Fusion of LinE/LinG)<br>LinG Homolog | 1.60E-87<br>5.50E-81 |
| WP_024756459.1 | 26172 | 27030 | + | 285 | LinH<br>LinE<br>PF12697<br>PF12146<br>PF00561 | LinH<br>LinE<br>Abhydrolase_6<br>Hydrolase_4<br>Abhydrolase_1 | LinH (Fusion of LinE/LinG)<br>LinE Homolog<br>Alpha/beta hydrolase family<br>Serine aminopeptidase, S33<br>alpha/beta hydrolase fold | 4.40E-53<br>7.50E-49<br>1.80E-12<br>3.70E-09<br>5.00E-08 |
| WP_024756460.1 | 27026 | 27608 | + | 193 | LinL | LinL | LinL | 4.20E-52 |
| WP_024756461.1 | 27604 | 28132 | + | 175 | PF07291 | MauE | Methylamine utilisation protein MauE | 5.10E-08 |
| WP_024756462.1 | 28137 | 28677 | + | 179 | PF08534<br>TIGR02661<br>PF00578 | Redoxin<br>TIGR02661<br>AhpC-TSA | Redoxin<br>MauD: methylamine dehydrogenase accessory protein MauD<br>AhpC/TSA family | 2.60E-08<br>4.50E-07<br>4.40E-05 |
| WP_024756463.1 | 28806 | 29010 | + | 67 | NO PFAM MATCH | - | - | - |
| WP_024756464.1 | 29457 | 31320 | + | 620 | TIGR02203<br>TIGR02204<br>TIGR01846<br>TIGR03796<br>TIGR03797 | TIGR02203<br>TIGR02204<br>TIGR01846<br>TIGR03796<br>TIGR03797 | MsbA_lipidA: lipid A export permease/ATP-binding protein MsbA<br>MsbA_rel: ABC transporter, permease/ATP-binding protein<br>type_I_sec_HlyB: type I secretion system ATPase<br>NHLM_micro_ABC1: NHLM bacteriocin system ABC transporter, peptidase/ATP-binding protein<br>NHLM_micro_ABC2: NHLM bacteriocin system ABC transporter, ATP-binding protein | 5.00E-79<br>2.40E-71<br>1.30E-68<br>1.30E-61<br>3.20E-61 |
| WP_024756465.1 | 31316 | 31790 | + | 157 | TIGR02227<br>TIGR02754<br>PF00717<br>TIGR02771<br>PF10502 | TIGR02227<br>TIGR02754<br>Peptidase_S24<br>TIGR02771<br>Peptidase_S26 | sigpep_I_bact: signal peptidase I<br>sod_Ni_protease: nickel-type superoxide dismutase maturation protease<br>Peptidase S24-like<br>TraF_Ti: conjugative transfer signal peptidase TraF<br>Signal peptidase, peptidase S26 | 8.00E-16<br>2.60E-09<br>1.00E-07<br>5.60E-05<br>6.20E-05 |
| WP_024756466.1 | 32979 | 31869 | - | 369 | TIGR02769<br>TIGR01186<br>TIGR03265<br>TIGR02770<br>TIGR00968 | TIGR02769<br>TIGR01186<br>TIGR03265<br>TIGR02770<br>TIGR00968 | nickel_nikE: nickel import ATP-binding protein NikE<br>proV: glycine betaine/L-proline transport ATP binding subunit<br>PhnT2: putative 2-aminoethylphosphonate ABC transporter, ATP-binding protein<br>nickel_nikD: nickel import ATP-binding protein NikD<br>3a0106s01: sulfate ABC transporter, ATP-binding protein | 3.50E-80<br>6.10E-57<br>1.70E-52<br>3.40E-52<br>9.30E-51 |

| index | leader | core | start | end | dir | score |
|---|---|---|---|---|---|---|
| 1 | MDRSRLVAVATGILAVGLAASPAAAAGSSDESPSNSTYNATSAFK | GTLAEDTLGTNDFPCCV | 24971 | 25160 | + | 6 |

### Results for WP_108956003.1 [Streptomyces fragilis] back to top

previous - next

#### Architecture

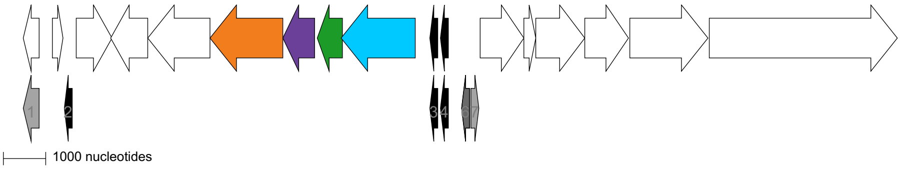

Link to nucleotide sequence

| Accession | start | end | direction | length (aa) | Pfam/HMM | name | description | E-value |
|---|---|---|---|---|---|---|---|---|
| WP_108955785.1 | 344780 | 344423 | - | 118 | PF00589<br>TIGR02249<br>TIGR02225<br>TIGR02224 | Phage_integrase<br>TIGR02249<br>TIGR02225<br>TIGR02224 | Phage integrase family<br>integrase_gron: integron integrase<br>recomb_XerD: tyrosine recombinase XerD<br>recomb_XerC: tyrosine recombinase XerC | 6.80E-12<br>3.70E-06<br>5.00E-05<br>2.90E-04 |
| WP_108955786.1 | 345111 | 345360 | + | 82 | NO PFAM MATCH | - | - | - |
| WP_108955787.1 | 345663 | 346503 | + | 279 | PF01551 | Peptidase_M23 | Peptidase family M23 | 7.10E-15 |
| WP_108955788.1 | 347339 | 346487 | - | 283 | PF12680<br>PF14534 | SnoaL_2<br>DUF4440 | SnoaL-like domain<br>Domain of unknown function (DUF4440) | 1.30E-04<br>3.10E-04 |

| | | | | | | | | |
|---|---|---|---|---|---|---|---|---|
| WP_108955789.1 | 348817 | 347335 | - | 493 | TIGR02734 | TIGR02734 | crtI_fam: phytoene desaturase | 3.40E-09 |
| | | | | | TIGR03467 | TIGR03467 | HpnE: squalene-associated FAD-dependent desaturase | 8.40E-09 |
| | | | | | TIGR00292 | TIGR00292 | TIGR00292: thiazole biosynthesis enzyme | 2.80E-08 |
| | | | | | TIGR01317 | TIGR01317 | GOGAT_sm_gam: glutamate synthase, NADH/NADPH, small subunit | 1.40E-07 |
| | | | | | TIGR02733 | TIGR02733 | desat_CrtD: C-3',4' desaturase CrtD | 1.90E-07 |
| WP_108955790.1 | 350550 | 348813 | - | 578 | TIGR02204 | TIGR02204 | MsbA_rel: ABC transporter, permease/ATP-binding protein | 2.20E-106 |
| | | | | | TIGR02203 | TIGR02203 | MsbA_lipidA: lipid A export permease/ATP-binding protein MsbA | 6.40E-100 |
| | | | | | TIGR00958 | TIGR00958 | 3a01208: antigen peptide transporter 2 | 1.40E-90 |
| | | | | | TIGR03796 | TIGR03796 | NHLM_micro_ABC1: NHLM bacteriocin system ABC transporter, peptidase/ATP-binding protein | 1.60E-84 |
| | | | | | TIGR01846 | TIGR01846 | type_I_sec_HlyB: type I secretion system ATPase | 1.40E-82 |
| WP_108955791.1 | 351302 | 350546 | - | 251 | PF13649 | Methyltransf_25 | Methyltransferase domain | 6.80E-15 |
| | | | | | PF08241 | Methyltransf_11 | Methyltransferase domain | 2.50E-10 |
| | | | | | PF13489 | Methyltransf_23 | Methyltransferase domain | 2.60E-09 |
| | | | | | PF08242 | Methyltransf_12 | Methyltransferase domain | 9.70E-09 |
| | | | | | TIGR01983 | TIGR01983 | UbiG: 3-demethylubiquinone-9 3-O-methyltransferase | 6.90E-07 |
| WP_108955792.1 | 351933 | 351354 | - | 192 | LinL | LinL | LinL | 4.00E-63 |
| WP_108956003.1 | 353663 | 351929 | - | 577 | LinH | LinH | LinH (Fusion of LinE/LinG) | 9.40E-226 |
| | | | | | LinE | LinE | LinE Homolog | 1.70E-99 |
| | | | | | LinG | LinG | LinG Homolog | 1.10E-94 |
| | | | | | PF12697 | Abhydrolase_6 | Alpha/beta hydrolase family | 4.10E-16 |
| | | | | | PF00561 | Abhydrolase_1 | alpha/beta hydrolase fold | 5.70E-14 |
| WP_108955793.1 | 354201 | 354009 | - | 63 | LinA | LinA | LinA (linaridin precursor) | 9.30E-10 |
| WP_108955794.1 | 354446 | 354251 | - | 64 | LinA | LinA | LinA (linaridin precursor) | 4.90E-17 |
| WP_108955795.1 | 355202 | 356243 | + | 346 | TIGR00747 | TIGR00747 | fabH: 3-oxoacyl-[acyl-carrier-protein] synthase III | 2.10E-12 |
| | | | | | PF08541 | ACP_syn_III_C | 3-Oxoacyl-[acyl-carrier-protein (ACP)] synthase III C terminal | 4.20E-06 |
| WP_108955796.1 | 356239 | 356521 | + | 93 | PF00550 | PP-binding | Phosphopantetheine attachment site | 8.00E-07 |
| | | | | | TIGR01688 | TIGR01688 | dltC: D-alanine--poly(phosphoribitol) ligase, subunit 2 | 8.80E-05 |
| WP_108955797.1 | 356504 | 357659 | + | 384 | PF00441 | Acyl-CoA_dh_1 | Acyl-CoA dehydrogenase, C-terminal domain | 6.30E-28 |
| | | | | | TIGR03203 | TIGR03203 | pimD_small: pimeloyl-CoA dehydrogenase, small subunit | 3.70E-22 |
| | | | | | TIGR03207 | TIGR03207 | cyc_hxne_CoA_dh: cyclohexanecarboxyl-CoA dehydrogenase | 1.10E-20 |
| | | | | | PF02770 | Acyl-CoA_dh_M | Acyl-CoA dehydrogenase, middle domain | 1.60E-17 |
| | | | | | PF02771 | Acyl-CoA_dh_N | Acyl-CoA dehydrogenase, N-terminal domain | 1.70E-17 |
| WP_108955798.1 | 357655 | 358714 | + | 352 | TIGR01686 | TIGR01686 | FkbH: FkbH domain | 2.50E-75 |
| | | | | | TIGR01681 | TIGR01681 | HAD-SF-IIIC: HAD phosphatase, family IIIC | 1.30E-13 |
| | | | | | PF12689 | Acid_PPase | Acid Phosphatase | 1.50E-04 |
| | | | | | PF00702 | Hydrolase | haloacid dehalogenase-like hydrolase | 3.60E-04 |
| WP_159105706.1 | 358724 | 360593 | + | 622 | TIGR01733 | TIGR01733 | AA-adenyl-dom: amino acid adenylation domain | 4.50E-112 |
| | | | | | PF00501 | AMP-binding | AMP-binding enzyme | 1.70E-81 |
| | | | | | TIGR01734 | TIGR01734 | D-ala-DACP-lig: D-alanine--poly(phosphoribitol) ligase, subunit 1 | 4.80E-55 |
| | | | | | TIGR03098 | TIGR03098 | ligase_PEP_1: acyl-CoA ligase (AMP-forming), exosortase A-associated | 2.80E-52 |
| | | | | | TIGR03443 | TIGR03443 | alpha_am_amid: L-aminoadipate-semialdehyde dehydrogenase | 4.90E-41 |
| WP_159105707.1 | 360609 | 365067 | + | 1485 | PF00668 | Condensation | Condensation domain | 4.30E-84 |
| | | | | | PF00109 | ketoacyl-synt | Beta-ketoacyl synthase, N-terminal domain | 2.20E-53 |
| | | | | | TIGR02813 | TIGR02813 | omega_3_PfaA: polyketide-type polyunsaturated fatty acid synthase PfaA | 8.70E-47 |
| | | | | | TIGR03150 | TIGR03150 | fabF: beta-ketoacyl-acyl-carrier-protein synthase II | 2.00E-33 |
| | | | | | PF02801 | Ketoacyl-synt_C | Beta-ketoacyl synthase, C-terminal domain | 6.50E-31 |

| index | leader | core | start | end | dir | score |
|---|---|---|---|---|---|---|
| 1 | MAEFGTWGEHGWLFESPKCKNQPFS...DAGLEHTGCTPKSLRHYFASVCIAG | GVPVFEVAKWLGHKGTRVTEATYVH...HLRSTDAIELAIVENAQHAQEARWS | 344780 | 344423 | - | 6 |
| 2 | MLVLGAFRLWPPGLGSPACSPNLYERAADEHGDL | SLRFRGTLAEAASGQDGAACADSSGVHGATELVV | 345572 | 345365 | - | 10 |
| 3 | MSKIVRSVAAVTPEVLTEDVHGDSGSMMPIQ | ATPALVAFGAGFAGGAGISWVCVQAYEAGAND | 354201 | 354009 | - | 29 |
| 4 | MEMKIAAFMNEQDLGSLERGYQDDAPAPVL | ATPLAAVATCWYVGGAVAGAAVVAGAYVTGRVVG | 354446 | 354251 | - | 30 |
| 5 | LAVLGAQSVRIGH | PTASASIPPRRLRRLNAQVRLNRRI...VVGCCTGNPARVAVQPVFLVGVRLA | 354949 | 354751 | - | 6 |
| 6 | MRPALAVLGAQSVRIGH | PTASASIPPRRLRRLNAQVRLNRRI...VVGCCTGNPARVAVQPVFLVGVRLA | 354961 | 354751 | - | 6 |
| 7 | MPADEHEEVGGGRTASDESLTKSEACG | PHGGRQGTWHSGVGLQTDDILRTCAIP | 354989 | 355154 | + | 6 |

### Results for WP_032775535.1 [Streptomyces fulvissimus] back to top

previous - next

#### Architecture

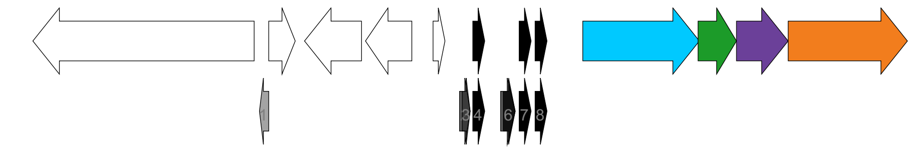

1000 nucleotides

Link to nucleotide sequence

| Accession | start | end | direction | length (aa) | Pfam/HMM | name | description | E-value |
|---|---|---|---|---|---|---|---|---|
| WP_164357055.1 | 13060 | 9559 | - | 1166 | PF06283 | ThuA | Trehalose utilisation | 3.10E-72 |
| | | | | | PF07995 | GSDH | Glucose / Sorbosone dehydrogenase | 2.20E-33 |
| | | | | | PF00801 | PKD | PKD domain | 2.20E-21 |
| | | | | | TIGR03606 | TIGR03606 | non_repeat_PQQ: dehydrogenase, PQQ-dependent, s-GDH family | 7.80E-21 |
| | | | | | PF03422 | CBM_6 | Carbohydrate binding module (family 6) | 9.50E-20 |

| | | | | | | | | |
|---|---|---|---|---|---|---|---|---|
| WP_164357056.1 | 13282 | 13708 | + | 141 | PF04237<br>PF04993 | YjbR<br>TfoX_N | YjbR<br>TfoX N-terminal domain | 3.90E-18<br>6.30E-04 |
| WP_164357057.1 | 14769 | 13866 | - | 300 | PF02779<br>TIGR00204<br>PF02780 | Transket_pyr<br>TIGR00204<br>Transketolase_C | Transketolase, pyrimidine binding domain<br>dxs: 1-deoxy-D-xylulose-5-phosphate synthase<br>Transketolase, C-terminal domain | 1.30E-19<br>1.30E-12<br>5.20E-06 |
| WP_164357058.1 | 15554 | 14834 | - | 239 | TIGR00232<br>PF00456<br>PF02775 | TIGR00232<br>Transketolase_N<br>TPP_enzyme_C | tktlase_bact: transketolase<br>Transketolase, thiamine diphosphate binding domain<br>Thiamine pyrophosphate enzyme, C-terminal TPP binding domain | 2.20E-24<br>4.20E-24<br>3.80E-04 |
| WP_011031194.1 | 15893 | 16076 | + | 61 | INFERRED GENE | - | - | - |
| WP_031123580.1 | 16518 | 16722 | + | 67 | LinA | LinA | LinA (linaridin precursor) | 1.10E-11 |
| WP_031123581.1 | 17260 | 17452 | + | 63 | LinA | LinA | LinA (linaridin precursor) | 2.30E-11 |
| WP_094211783.1 | 17509 | 17710 | + | 66 | LinA | LinA | LinA (linaridin precursor) | 4.50E-12 |
| WP_032775535.1 | 18260 | 20108 | + | 615 | LinH<br>LinG<br>LinE<br>LinL<br>PF12697 | LinH<br>LinG<br>LinE<br>LinL<br>Abhydrolase_6 | LinH (Fusion of LinE/LinG)<br>LinG Homolog<br>LinE Homolog<br>LinL<br>Alpha/beta hydrolase family | 1.70E-252<br>1.40E-111<br>5.20E-94<br>2.60E-37<br>2.00E-15 |
| WP_031123584.1 | 20104 | 20713 | + | 202 | LinL | LinL | LinL | 4.30E-64 |
| WP_031123585.1 | 20709 | 21522 | + | 270 | PF13649<br>PF08241<br>PF13847<br>PF13489<br>PF08242 | Methyltransf_25<br>Methyltransf_11<br>Methyltransf_31<br>Methyltransf_23<br>Methyltransf_12 | Methyltransferase domain<br>Methyltransferase domain<br>Methyltransferase domain<br>Methyltransferase domain<br>Methyltransferase domain | 2.30E-16<br>2.90E-12<br>1.70E-07<br>9.20E-07<br>1.00E-06 |
| WP_164357059.1 | 21518 | 23420 | + | 633 | TIGR02204<br>TIGR02203<br>TIGR00958<br>TIGR03796<br>TIGR02857 | TIGR02204<br>TIGR02203<br>TIGR00958<br>TIGR03796<br>TIGR02857 | MsbA_rel: ABC transporter, permease/ATP-binding protein<br>MsbA_lipidA: lipid A export permease/ATP-binding protein MsbA<br>3a01208: antigen peptide transporter 2<br>NHLM_micro_ABC1: NHLM bacteriocin system ABC transporter, peptidase/ATP-binding protein<br>CydD: thiol reductant ABC exporter, CydD subunit | 7.20E-99<br>1.50E-92<br>3.70E-83<br>2.50E-79<br>1.20E-75 |

| index | leader | core | start | end | dir | score |
|---|---|---|---|---|---|---|
| 1 | LRSSRFSDRVAPGGRCPGFDRNARDRHPPPA | GTPGTPGRTGPADPLTPSKR | 13292 | 13136 | - | 6 |
| 2 | VPEPAACCFGVRILMPAELSSLSAGL | SASCGAHHRTSVHAGVGSRVRSRAPG | 16322 | 16481 | + | 8 |
| 3 | MPAELSSLSAGLSASCGAH | HRTSVHAGVGSRVRSRAPG | 16364 | 16481 | + | 6 |
| 4 | MPEFRQPGWTRGIAPLDESAGGQVFGGASPVA | ATPAVVATAGAVVIAFAAGVAARQLANGGNVELPM | 16518 | 16722 | + | 30 |
| 5 | MTSPHRPPTSEEKSLSTASIPAVLSEAGALSESDRGRAPLD | TVPVASVTFTMTACVGISVCLTGSTIELPQ | 16971 | 17187 | + | 8 |
| 6 | LSTASIPAVLSEAGALSESDRGRAPLD | TVPVASVTFTMTACVGISVCLTGSTIELPQ | 17013 | 17187 | + | 9 |
| 7 | MSKSSAPAVLTATGALTESDPGVTLPALAPVA | ATPVAIAATLGAAFIAGYAAGRAAGNVELPM | 17260 | 17452 | + | 28 |
| 8 | MSTSSTPAALRGAGGLSESDPGRALSTLAPVA | ATPGIVAGAALGVALVNAFVAGYNHGGGNVELPM | 17509 | 17710 | + | 31 |

### Results for WP_033526318.1 [Streptomyces galbus] back to top

previous - next

#### Architecture

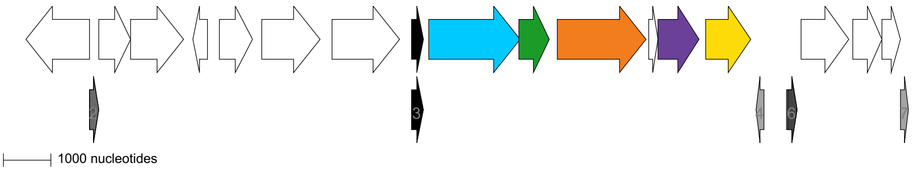

Link to nucleotide sequence

| Accession | start | end | direction | length (aa) | Pfam/HMM | name | description | E-value |
|---|---|---|---|---|---|---|---|---|
| WP_078937034.1 | 134675 | 133340 | - | 444 | PF04235<br>PF07786 | DUF418<br>DUF1624 | Protein of unknown function (DUF418)<br>Protein of unknown function (DUF1624) | 2.20E-09<br>2.00E-05 |
| WP_033526313.1 | 134867 | 135545 | + | 225 | PF00072<br>TIGR02875<br>PF00196<br>TIGR01818<br>TIGR03020 | Response_reg<br>TIGR02875<br>GerE<br>TIGR01818<br>TIGR03020 | Response regulator receiver domain<br>spore_0_A: sporulation transcription factor Spo0A<br>Bacterial regulatory proteins, luxR family<br>ntrC: nitrogen regulation protein NR(I)<br>EpsA: transcriptional regulator EpsA | 2.70E-24<br>1.10E-17<br>1.50E-17<br>5.30E-13<br>7.70E-11 |
| WP_052145933.1 | 135541 | 136663 | + | 373 | PF07730 | HisKA_3 | Histidine kinase | 2.30E-12 |
| WP_033526315.1 | 137145 | 136842 | - | 100 | PF00196<br>TIGR03541<br>TIGR03020<br>PF08281 | GerE<br>TIGR03541<br>TIGR03020<br>Sigma70_r4_2 | Bacterial regulatory proteins, luxR family<br>reg_near_HchA: LuxR family transcriptional regulatory, chaperone HchA-associated<br>EpsA: transcriptional regulator EpsA<br>Sigma-70, region 4 | 5.80E-15<br>1.60E-12<br>4.30E-09<br>5.10E-07 |
| WP_033526316.1 | 137401 | 138091 | + | 229 | PF00072<br>PF00196<br>TIGR02875<br>TIGR03020<br>TIGR03541 | Response_reg<br>GerE<br>TIGR02875<br>TIGR03020<br>TIGR03541 | Response regulator receiver domain<br>Bacterial regulatory proteins, luxR family<br>spore_0_A: sporulation transcription factor Spo0A<br>EpsA: transcriptional regulator EpsA<br>reg_near_HchA: LuxR family transcriptional regulatory, chaperone HchA-associated | 1.20E-24<br>1.40E-18<br>5.60E-16<br>2.50E-13<br>2.10E-08 |
| WP_052145934.1 | 138307 | 139537 | + | 409 | PF07730<br>PF13581 | HisKA_3<br>HATPase_c_2 | Histidine kinase<br>Histidine kinase-like ATPase domain | 1.90E-13<br>1.70E-04 |

| | | | | | | | | |
|---|---|---|---|---|---|---|---|---|
| WP_033526317.1 | 139777 | 141208 | + | 476 | PF13450 | NAD_binding_8 | NAD(P)-binding Rossmann-like domain | 1.10E-11 |
| | | | | | TIGR02734 | TIGR02734 | crtI_fam: phytoene desaturase | 1.60E-10 |
| | | | | | TIGR03467 | TIGR03467 | HpnE: squalene-associated FAD-dependent desaturase | 1.90E-08 |
| | | | | | TIGR03315 | TIGR03315 | Se_ygfK: putative selenate reductase, YgfK subunit | 4.50E-08 |
| | | | | | PF01266 | DAO | FAD dependent oxidoreductase | 1.80E-07 |
| WP_033526382.1 | 141451 | 141691 | + | 79 | LinA | LinA | LinA (linaridin precursor) | 1.00E-04 |
| WP_033526318.1 | 141806 | 143726 | + | 639 | LinH | LinH | LinH (Fusion of LinE/LinG) | 1.40E-248 |
| | | | | | LinG | LinG | LinG Homolog | 2.00E-157 |
| | | | | | LinE | LinE | LinE Homolog | 6.80E-92 |
| | | | | | PF12697 | Abhydrolase_6 | Alpha/beta hydrolase family | 1.20E-16 |
| | | | | | PF12146 | Hydrolase_4 | Serine aminopeptidase, S33 | 6.50E-09 |
| WP_033526319.1 | 143722 | 144343 | + | 206 | LinL | LinL | LinL | 2.30E-69 |
| WP_033526320.1 | 144509 | 146402 | + | 630 | TIGR02204 | TIGR02204 | MsbA_rel: ABC transporter, permease/ATP-binding protein | 1.70E-107 |
| | | | | | TIGR02203 | TIGR02203 | MsbA_lipidA: lipid A export permease/ATP-binding protein MsbA | 1.80E-103 |
| | | | | | TIGR00958 | TIGR00958 | 3a01208: antigen peptide transporter 2 | 1.10E-92 |
| | | | | | TIGR03796 | TIGR03796 | NHLM_micro_ABC1: NHLM bacteriocin system ABC transporter, peptidase/ATP-binding protein | 1.90E-86 |
| | | | | | TIGR03375 | TIGR03375 | type_I_sec_LssB: type I secretion system ATPase | 7.00E-81 |
| WP_033526321.1 | 146459 | 146651 | + | 63 | NO PFAM MATCH | - | - | - |
| WP_052145935.1 | 146647 | 147511 | + | 287 | PF13649 | Methyltransf_25 | Methyltransferase domain | 2.20E-15 |
| | | | | | PF08241 | Methyltransf_11 | Methyltransferase domain | 9.00E-13 |
| | | | | | PF13489 | Methyltransf_23 | Methyltransferase domain | 1.20E-07 |
| | | | | | PF08242 | Methyltransf_12 | Methyltransferase domain | 1.10E-06 |
| | | | | | TIGR01983 | TIGR01983 | UbiG: 3-demethylubiquinone-9 3-O-methyltransferase | 2.90E-05 |
| WP_033526322.1 | 147647 | 148604 | + | 318 | PF00561 | Abhydrolase_1 | alpha/beta hydrolase fold | 2.40E-25 |
| | | | | | PF12697 | Abhydrolase_6 | Alpha/beta hydrolase family | 6.40E-21 |
| | | | | | TIGR03056 | TIGR03056 | bchO_mg_che_rel: putative magnesium chelatase accessory protein | 7.40E-17 |
| | | | | | TIGR02427 | TIGR02427 | protocat_pcaD: 3-oxoadipate enol-lactonase | 2.60E-15 |
| | | | | | TIGR03343 | TIGR03343 | biphenyl_bphD: 2-hydroxy-6-oxo-6-phenylhexa-2,4-dienoate hydrolase | 2.60E-14 |
| WP_033526323.1 | 149661 | 150654 | + | 330 | TIGR02363 | TIGR02363 | dhaK1: dihydroxyacetone kinase, DhaK subunit | 1.40E-149 |
| | | | | | PF02733 | Dak1 | Dak1 domain | 4.90E-127 |
| | | | | | TIGR02361 | TIGR02361 | dak_ATP: dihydroxyacetone kinase | 2.90E-94 |
| | | | | | TIGR02362 | TIGR02362 | dhaK1b: probable dihydroxyacetone kinase DhaK1b subunit | 1.90E-74 |
| WP_033526324.1 | 150743 | 151358 | + | 204 | TIGR02365 | TIGR02365 | dha_L_ycgS: dihydroxyacetone kinase, L subunit | 1.70E-69 |
| | | | | | PF02734 | Dak2 | DAK2 domain | 2.40E-50 |
| | | | | | TIGR02361 | TIGR02361 | dak_ATP: dihydroxyacetone kinase | 9.90E-36 |
| | | | | | TIGR03599 | TIGR03599 | YloV: DAK2 domain fusion protein YloV | 5.30E-07 |
| WP_033526325.1 | 151350 | 151761 | + | 136 | TIGR02364 | TIGR02364 | dha_pts: dihydroxyacetone kinase, phosphotransfer subunit | 5.60E-29 |
| | | | | | PF03610 | EIIA-man | PTS system fructose IIA component | 2.00E-24 |

| index | leader | core | start | end | dir | score |
|---|---|---|---|---|---|---|
| 1 | VFGVMAFSVGGARGASPSGGWLDV | LPPTSRSHPRQVTGKGEEWEPDGDFGRRSGSSRSPVVVRRRAGHG | 134661 | 134871 | + | 6 |
| 2 | MAFSVGGARGASPSGGWLDV | LPPTSRSHPRQVTGKGEEWEPDGDFGRRSGSSRSPVVVRRRAGHG | 134673 | 134871 | + | 6 |
| 3 | MEIPVEGVVYVAARPTL | GTPRIARIGRIAQAAEGVGALAAAA...NNLAAEANAQNQAALAAVGGTPPAV | 141451 | 141691 | + | 28 |
| 4 | VAAGSAVHRELPGADLDPARHRRYGRGP | SRCMATRRSPGTAASNNLDAGARATALGA | 148874 | 148700 | - | 6 |
| 5 | VRAEAGHEKPLRYHRRLVGSPGLPDLSNRTRYRTQL | AQHYARFATDGHGREPAGRTHPAPTTPLAARHSKLVA | 149339 | 149561 | + | 7 |
| 6 | VGSPGLPDLSNRTRYRTQLAQHYARFAT | DGHGREPAGRTHPAPTTPLAARHSKLVA | 149390 | 149561 | + | 7 |
| 7 | VTLRSTATRETTDDSSRSSPTP | GTPFRGLPPHEAAGPERERPARTEATGSHGR | 151757 | 151919 | + | 6 |

### Results for WP_030581598.1 [Streptomyces globisporus subsp. globisporus]

back to top

previous - next

#### Architecture

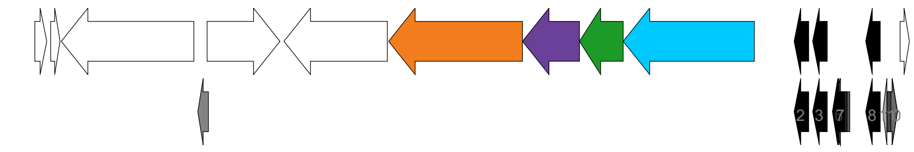

1000 nucleotides

Link to nucleotide sequence

| Accession | start | end | direction | length (aa) | Pfam/HMM | name | description | E-value |
|---|---|---|---|---|---|---|---|---|
| NP_624573.1 | 218182 | 218350 | + | 56 | INFERRED GENE | - | - | - |
| NP_630282.1 | 218392 | 218529 | + | 45 | INFERRED GENE | - | - | - |
| WP_030581580.1 | 220406 | 218552 | - | 617 | NO PFAM MATCH | - | - | - |
| WP_030581583.1 | 220595 | 221609 | + | 337 | NO PFAM MATCH | - | - | - |
| WP_078652008.1 | 223126 | 221665 | - | 486 | TIGR02734 | TIGR02734 | crtI_fam: phytoene desaturase | 2.80E-11 |
| | | | | | PF13450 | NAD_binding_8 | NAD(P)-binding Rossmann-like domain | 1.50E-09 |
| | | | | | TIGR01317 | TIGR01317 | GOGAT_sm_gam: glutamate synthase, NADH/NADPH, small subunit | 1.20E-08 |
| | | | | | TIGR00292 | TIGR00292 | TIGR00292: thiazole biosynthesis enzyme | 1.40E-08 |
| | | | | | TIGR03467 | TIGR03467 | HpnE: squalene-associated FAD-dependent desaturase | 1.60E-08 |

| | | | | | | | | |
|---|---|---|---|---|---|---|---|---|
| WP_030581589.1 | 225015 | 223131 | - | 627 | TIGR02204 | TIGR02204 | MsbA_rel: ABC transporter, permease/ATP-binding protein | 1.00E-103 |
| | | | | | TIGR02203 | TIGR02203 | MsbA_lipidA: lipid A export permease/ATP-binding protein MsbA | 3.50E-96 |
| | | | | | TIGR00958 | TIGR00958 | 3a01208: antigen peptide transporter 2 | 9.30E-87 |
| | | | | | TIGR03796 | TIGR03796 | NHLM_micro_ABC1: NHLM bacteriocin system ABC transporter, peptidase/ATP-binding protein | 1.90E-83 |
| | | | | | TIGR03375 | TIGR03375 | type_I_sec_LssB: type I secretion system ATPase | 1.60E-79 |
| WP_030581592.1 | 225824 | 225011 | - | 270 | PF13649 | Methyltransf_25 | Methyltransferase domain | 8.80E-17 |
| | | | | | PF08241 | Methyltransf_11 | Methyltransferase domain | 2.80E-12 |
| | | | | | PF08242 | Methyltransf_12 | Methyltransferase domain | 4.60E-08 |
| | | | | | PF13847 | Methyltransf_31 | Methyltransferase domain | 4.80E-07 |
| | | | | | PF13489 | Methyltransf_23 | Methyltransferase domain | 7.40E-07 |
| WP_030581595.1 | 226429 | 225820 | - | 202 | LinL | LinL | LinL | 1.20E-65 |
| WP_030581598.1 | 228276 | 226425 | - | 616 | LinH | LinH | LinH (Fusion of LinE/LinG) | 1.10E-250 |
| | | | | | LinG | LinG | LinG Homolog | 1.30E-111 |
| | | | | | LinE | LinE | LinE Homolog | 4.30E-94 |
| | | | | | LinL | LinL | LinL | 1.60E-34 |
| | | | | | PF12697 | Abhydrolase_6 | Alpha/beta hydrolase family | 6.80E-16 |
| WP_030581602.1 | 229038 | 228837 | - | 66 | LinA | LinA | LinA (linaridin precursor) | 3.10E-11 |
| WP_030581605.1 | 229292 | 229097 | - | 64 | LinA | LinA | LinA (linaridin precursor) | 3.30E-11 |
| WP_030581608.1 | 230033 | 229829 | - | 67 | LinA | LinA | LinA (linaridin precursor) | 4.10E-10 |
| WP_158709122.1 | 230314 | 230455 | + | 46 | NO PFAM MATCH | - | - | - |

| index | leader | core | start | end | dir | score |
|---|---|---|---|---|---|---|
| 1 | VGPHGPSLEHVRRDDGAQVCPGR | RVVRSQRSLTPRWYQGSSGCVTT | 220607 | 220466 | - | 7 |
| 2 | MNTSPIPAALRGVGGLSESASGRALSTFAPVA | ATPGIAAGAALGVALVNAFVAGYNHCGGNVELPM | 229038 | 228837 | - | 29 |
| 3 | MSKSSTPAVLTATGDLTESDPGVTLPALAPVT | ATPVAIAATLGAAFLAGYAAGRAAGSDVELPM | 229292 | 229097 | - | 28 |
| 4 | LIPAVLSETGALSESDPGRAPLD | TVPVASVTFTMSACVGISVCLTGSTIQLPQ | 229527 | 229365 | - | 9 |
| 5 | LSTALIPAVLSETGALSESDPGRAPLD | TVPVASVTFTMSACVGISVCLTGSTIQLPQ | 229539 | 229365 | - | 9 |
| 6 | MTSPQRPPTSEEKSLSTALIPAVLSETGALSESDPGRAPLD | TVPVASVTFTMSACVGISVCLTGSTIQLPQ | 229581 | 229365 | - | 8 |
| 7 | VGPSHRRRPPMTSPQRPPTSEEKSL...TALIPAVLSETGALSESDPGRAPLD | TVPVASVTFTMSACVGISVCLTGSTIQLPQ | 229611 | 229365 | - | 8 |
| 8 | MPEFRQPGWTRGIAPLDEGSGGQVFGAASPVA | ATPVVVASAGAVVIAFAAGVAARELANGGNVELPM | 230033 | 229829 | - | 31 |
| 9 | VMRSTRCREPGGKGRPLRWHEDADSK | ATSCGRVGAWGCAAPTVSLDPSWNAWR | 230127 | 230289 | + | 6 |
| 10 | MRSTRCREPGGKGRPLRWHEDADSKA | TSCGRVGAWGCAAPTVSLDPSWNAWR | 230130 | 230289 | + | 6 |
| 11 | MPAERSSLSAGLSASCGAH | HRTSVHAGAGSRVRLRAPG | 230187 | 230070 | - | 6 |

### Results for WP_105970077.1 [Streptomyces geranii] back to top

previous - next

#### Architecture

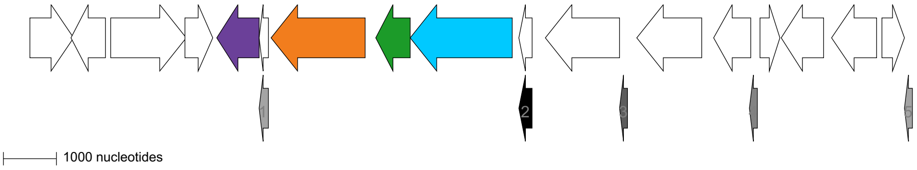

1000 nucleotides

Link to nucleotide sequence

| Accession | start | end | direction | length (aa) | Pfam/HMM | name | description | E-value |
|---|---|---|---|---|---|---|---|---|
| WP_105970071.1 | 205641 | 206487 | + | 281 | PF00459 | Inositol_P | Inositol monophosphatase family | 1.80E-58 |
| | | | | | TIGR02067 | TIGR02067 | his_9_HisN: histidinol-phosphatase | 7.80E-38 |
| | | | | | TIGR01331 | TIGR01331 | bisphos_cysQ: 3'(2'),5'-bisphosphate nucleotidase | 4.20E-26 |
| | | | | | TIGR01330 | TIGR01330 | bisphos_HAL2: 3'(2'),5'-bisphosphate nucleotidase | 2.50E-12 |
| WP_105970072.1 | 207085 | 206422 | - | 220 | NO PFAM MATCH | - | - | - |
| WP_105970191.1 | 207181 | 208591 | + | 469 | TIGR02734 | TIGR02734 | crtI_fam: phytoene desaturase | 5.40E-17 |
| | | | | | PF13450 | NAD_binding_8 | NAD(P)-binding Rossmann-like domain | 5.70E-13 |
| | | | | | TIGR03467 | TIGR03467 | HpnE: squalene-associated FAD-dependent desaturase | 1.40E-10 |
| | | | | | PF01593 | Amino_oxidase | Flavin containing amine oxidoreductase | 1.10E-09 |
| | | | | | PF01266 | DAO | FAD dependent oxidoreductase | 1.80E-09 |
| WP_105970073.1 | 208587 | 209100 | + | 170 | PF01661 | Macro | Macro domain | 2.00E-40 |
| WP_105970074.1 | 209973 | 209178 | - | 264 | PF13649 | Methyltransf_25 | Methyltransferase domain | 1.80E-15 |
| | | | | | PF08241 | Methyltransf_11 | Methyltransferase domain | 8.20E-12 |
| | | | | | PF13489 | Methyltransf_23 | Methyltransferase domain | 4.70E-08 |
| | | | | | PF08242 | Methyltransf_12 | Methyltransferase domain | 4.90E-07 |
| | | | | | TIGR01983 | TIGR01983 | UbiG: 3-demethylubiquinone-9 3-O-methyltransferase | 4.20E-05 |
| WP_105970075.1 | 210161 | 209969 | - | 63 | NO PFAM MATCH | - | - | - |
| WP_105970192.1 | 211966 | 210205 | - | 586 | TIGR02203 | TIGR02203 | MsbA_lipidA: lipid A export permease/ATP-binding protein MsbA | 7.50E-106 |
| | | | | | TIGR02204 | TIGR02204 | MsbA_rel: ABC transporter, permease/ATP-binding protein | 1.20E-105 |
| | | | | | TIGR00958 | TIGR00958 | 3a01208: antigen peptide transporter 2 | 9.90E-93 |
| | | | | | TIGR03796 | TIGR03796 | NHLM_micro_ABC1: NHLM bacteriocin system ABC transporter, peptidase/ATP-binding protein | 2.50E-88 |
| | | | | | TIGR01846 | TIGR01846 | type_I_sec_HlyB: type I secretion system ATPase | 1.10E-83 |
| WP_105970076.1 | 212834 | 212168 | - | 221 | LinL | LinL | LinL | 1.60E-74 |

| | | | | | | | | |
|---|---|---|---|---|---|---|---|---|
| WP_105970077.1 | 214741 | 212830 | - | 636 | LinH<br>LinG<br>LinE<br>PF12697<br>PF12146 | LinH<br>LinG<br>LinE<br>Abhydrolase_6<br>Hydrolase_4 | LinH (Fusion of LinE/LinG)<br>LinG Homolog<br>LinE Homolog<br>Alpha/beta hydrolase family<br>Serine aminopeptidase, S33 | 1.10E-233<br>2.10E-124<br>8.90E-88<br>1.30E-15<br>1.20E-09 |
| WP_105970193.1 | 215128 | 214867 | - | 86 | NO PFAM MATCH | - | - | - |
| WP_105970078.1 | 216786 | 215373 | - | 470 | TIGR02734<br>PF13450<br>TIGR03467<br>TIGR03315<br>TIGR00292 | TIGR02734<br>NAD_binding_8<br>TIGR03467<br>TIGR03315<br>TIGR00292 | crtI_fam: phytoene desaturase<br>NAD(P)-binding Rossmann-like domain<br>HpnE: squalene-associated FAD-dependent desaturase<br>Se_ygfK: putative selenate reductase, YgfK subunit<br>TIGR00292: thiazole biosynthesis enzyme | 2.80E-15<br>2.90E-11<br>5.90E-09<br>8.20E-08<br>1.70E-07 |
| WP_105970194.1 | 218314 | 217087 | - | 408 | PF07730 | HisKA_3 | Histidine kinase | 7.50E-14 |
| WP_105970195.1 | 219246 | 218556 | - | 229 | PF00072<br>PF00196<br>TIGR02875<br>TIGR03020<br>TIGR03541 | Response_reg<br>GerE<br>TIGR02875<br>TIGR03020<br>TIGR03541 | Response regulator receiver domain<br>Bacterial regulatory proteins, luxR family<br>spore_0_A: sporulation transcription factor Spo0A<br>EpsA: transcriptional regulator EpsA<br>reg_near_HchA: LuxR family transcriptional regulatory, chaperone HchA-associated | 1.30E-24<br>1.20E-17<br>7.50E-16<br>1.00E-13<br>2.30E-08 |
| WP_105970079.1 | 219432 | 219786 | + | 117 | PF00196<br>TIGR03541<br>TIGR03020<br>PF08281 | GerE<br>TIGR03541<br>TIGR03020<br>Sigma70_r4_2 | Bacterial regulatory proteins, luxR family<br>reg_near_HchA: LuxR family transcriptional regulatory, chaperone HchA-associated<br>EpsA: transcriptional regulator EpsA<br>Sigma-70, region 4 | 9.50E-16<br>1.90E-12<br>3.20E-09<br>5.10E-05 |
| WP_105970080.1 | 220633 | 219814 | - | 272 | PF00106<br>TIGR01963<br>TIGR01289<br>PF13561<br>TIGR04316 | adh_short<br>TIGR01963<br>TIGR01289<br>adh_short_C2<br>TIGR04316 | short chain dehydrogenase<br>PHB_DH: 3-hydroxybutyrate dehydrogenase<br>LPOR: light-dependent protochlorophyllide reductase<br>Enoyl-(Acyl carrier protein) reductase<br>dhbA_paeA: 2,3-dihydro-2,3-dihydroxybenzoate dehydrogenase | 7.20E-20<br>2.00E-13<br>5.20E-12<br>1.80E-11<br>2.10E-10 |
| WP_105970081.1 | 221629 | 220774 | - | 284 | TIGR03649<br>PF13460<br>PF05368<br>TIGR01746<br>TIGR03466 | TIGR03649<br>NAD_binding_10<br>NmrA<br>TIGR01746<br>TIGR03466 | ergot_EASG: ergot alkaloid biosynthesis protein, AFUA_2G17970 family<br>NAD(P)H-binding<br>NmrA-like family<br>Thioester-redct: thioester reductase domain<br>HpnA: hopanoid-associated sugar epimerase | 1.50E-17<br>3.80E-15<br>5.10E-07<br>1.10E-04<br>1.60E-04 |
| WP_105970082.1 | 221721 | 222135 | + | 137 | PF01638 | HxlR | HxlR-like helix-turn-helix | 1.30E-23 |

| index | leader | core | start | end | dir | score |
|---|---|---|---|---|---|---|
| 1 | MNDFLLVPFVVALGIVGFLIATRAVSTPAVI | GVALVVLLGACSQARRNVHPKNTPAKKRRQHP | 210161 | 209969 | - | 6 |
| 2 | MDLPVEGVILTSARPTL | GTPRIGRVANAANALNALGGLADLA...QAAADANQAVANAANAQAAAAGGTV | 215128 | 214867 | - | 28 |
| 3 | LTTDLAPQSGETARTGSFEGPGPSLSRCLPLA | RPPLLRTGAPRRRDAGVV | 216935 | 216782 | - | 8 |
| 4 | VTRAPRASTVRPIPGCGHRPIGLG | PSAPAETGIPRTAGPAACSMTSSAG | 219380 | 219230 | - | 7 |
| 5 | VGERRGRSVVYALHDHHVAGLLDQ | AVYHVEHLRLGLSDTDGDVETDGVE | 222294 | 222144 | - | 6 |

### Results for EMF28262.1 [Streptomyces gancidicus BKS 13-15] back to top

previous - next

#### Architecture

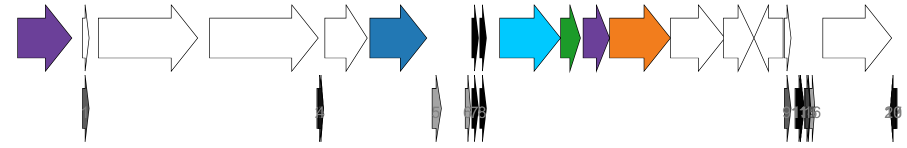

1000 nucleotides

Link to nucleotide sequence

| Accession | start | end | direction | length (aa) | Pfam/HMM | name | description | E-value |
|---|---|---|---|---|---|---|---|---|
| EMF28254.1 | 41723 | 43253 | + | 509 | TIGR03534<br>PF05175<br>TIGR03533<br>TIGR00537<br>TIGR00536 | TIGR03534<br>MTS<br>TIGR03533<br>TIGR00537<br>TIGR00536 | RF_mod_PrmC: protein-(glutamine-N5) methyltransferase, release factor-specific<br>Methyltransferase small domain<br>L3_gln_methyl: protein-(glutamine-N5) methyltransferase, ribosomal protein L3-specific<br>hemK_rel_arch: putative methylase<br>hemK_fam: methyltransferase, HemK family | 1.10E-14<br>8.90E-13<br>1.20E-10<br>6.40E-10<br>2.40E-09 |
| EMF28255.1 | 43547 | 43745 | + | 65 | NO PFAM MATCH | - | - | - |
| EMF28256.1 | 43990 | 46825 | + | 944 | TIGR01051<br>PF01131<br>TIGR01054<br>TIGR01057<br>PF13368 | TIGR01051<br>Topoisom_bac<br>TIGR01054<br>TIGR01057<br>Toprim_C_rpt | topA_bact: DNA topoisomerase I<br>DNA topoisomerase<br>rgy: reverse gyrase<br>topA_arch: DNA topoisomerase I<br>Topoisomerase C-terminal repeat | 3.40E-246<br>2.30E-131<br>2.00E-86<br>4.70E-72<br>9.20E-65 |
| EMF28257.1 | 47132 | 50195 | + | 1020 | TIGR00041<br>PF02223<br>TIGR00900<br>PF07690<br>TIGR00895 | TIGR00041<br>Thymidylate_kin<br>TIGR00900<br>MFS_1<br>TIGR00895 | DTMP_kinase: dTMP kinase<br>Thymidylate kinase<br>2A0121: H+ Antiporter protein<br>Major Facilitator Superfamily<br>2A0115: MFS transporter, aromatic acid:H+ symporter (AAHS) family | 1.20E-51<br>1.20E-46<br>8.40E-11<br>5.50E-08<br>4.00E-04 |
| EMF28258.1 | 50374 | 51580 | + | 401 | TIGR00678<br>PF13177<br>TIGR02397<br>PF00004 | TIGR00678<br>DNA_pol3_delta2<br>TIGR02397<br>AAA | holB: DNA polymerase III, delta' subunit<br>DNA polymerase III, delta subunit<br>dnaX_nterm: DNA polymerase III, subunit gamma and tau<br>ATPase family associated with various cellular activities (AAA) | 1.30E-54<br>9.20E-33<br>1.70E-29<br>3.60E-04 |
| EMF28259.1 | 51671 | 53261 | + | 529 | PF00561<br>PF08386<br>TIGR01249<br>LinE | Abhydrolase_1<br>Abhydrolase_4<br>TIGR01249<br>LinE | alpha/beta hydrolase fold<br>TAP-like protein<br>pro_imino_pep_1: prolyl aminopeptidase<br>LinE Homolog | 3.40E-36<br>1.30E-31<br>2.60E-06<br>9.80E-06 |

| | | | | | | | | |
|---|---|---|---|---|---|---|---|---|
| EMF28260.1 | 54543 | 54738 | + | 64 | LinA | LinA | LinA (linaridin precursor) | 1.10E-17 |
| EMF28261.1 | 54773 | 54965 | + | 63 | LinA | LinA | LinA (linaridin precursor) | 1.70E-11 |
| EMF28262.1 | 55310 | 57032 | + | 573 | LinH<br>LinE<br>LinG<br>PF12697<br>PF00561 | LinH<br>LinE<br>LinG<br>Abhydrolase_6<br>Abhydrolase_1 | LinH (Fusion of LinE/LinG)<br>LinE Homolog<br>LinG Homolog<br>Alpha/beta hydrolase family<br>alpha/beta hydrolase fold | 9.20E-224<br>1.90E-99<br>3.10E-96<br>5.30E-15<br>7.80E-12 |
| EMF28263.1 | 57028 | 57607 | + | 192 | LinL | LinL | LinL | 8.30E-66 |
| EMF28264.1 | 57661 | 58417 | + | 251 | PF13649<br>PF08241<br>PF13489<br>PF13847<br>PF08242 | Methyltransf_25<br>Methyltransf_11<br>Methyltransf_23<br>Methyltransf_31<br>Methyltransf_12 | Methyltransferase domain<br>Methyltransferase domain<br>Methyltransferase domain<br>Methyltransferase domain<br>Methyltransferase domain | 2.70E-13<br>1.50E-08<br>1.70E-08<br>4.80E-07<br>5.00E-07 |
| EMF28265.1 | 58413 | 60150 | + | 578 | TIGR02204<br>TIGR02203<br>TIGR03796<br>TIGR00958<br>TIGR01846 | TIGR02204<br>TIGR02203<br>TIGR03796<br>TIGR00958<br>TIGR01846 | MsbA_rel: ABC transporter, permease/ATP-binding protein<br>MsbA_lipidA: lipid A export permease/ATP-binding protein MsbA<br>NHLM_micro_ABC1: NHLM bacteriocin system ABC transporter, peptidase/ATP-binding protein<br>3a01208: antigen peptide transporter 2<br>type_I_sec_HlyB: type I secretion system ATPase | 8.80E-101<br>6.60E-98<br>2.80E-89<br>5.70E-85<br>9.40E-83 |
| EMF28266.1 | 60146 | 61652 | + | 501 | TIGR02734<br>TIGR00292<br>TIGR03467<br>TIGR01317<br>TIGR02733 | TIGR02734<br>TIGR00292<br>TIGR03467<br>TIGR01317<br>TIGR02733 | crtI_fam: phytoene desaturase<br>TIGR00292: thiazole biosynthesis enzyme<br>HpnE: squalene-associated FAD-dependent desaturase<br>GOGAT_sm_gam: glutamate synthase, NADH/NADPH, small subunit<br>desat_CrtD: C-3',4' desaturase CrtD | 1.30E-08<br>2.50E-08<br>9.20E-08<br>4.10E-07<br>6.80E-07 |
| EMF28267.1 | 61648 | 62500 | + | 283 | PF12680<br>PF14534 | SnoaL_2<br>DUF4440 | SnoaL-like domain<br>Domain of unknown function (DUF4440) | 5.30E-05<br>8.10E-04 |
| EMF28268.1 | 63324 | 62484 | - | 279 | PF01551 | Peptidase_M23 | Peptidase family M23 | 3.30E-17 |
| EMF28269.1 | 63354 | 63546 | + | 63 | NO PFAM MATCH | - | - | - |
| EMF28270.1 | 64449 | 66372 | + | 640 | PF03704<br>PF00486<br>PF13191<br>PF00931<br>TIGR02928 | BTAD<br>Trans_reg_C<br>AAA_16<br>NB-ARC<br>TIGR02928 | Bacterial transcriptional activator domain<br>Transcriptional regulatory protein, C terminal<br>AAA ATPase domain<br>NB-ARC domain<br>TIGR02928: orc1/cdc6 family replication initiation protein | 6.50E-41<br>1.30E-08<br>1.30E-07<br>3.00E-06<br>2.00E-04 |

| index | leader | core | start | end | dir | score |
|---|---|---|---|---|---|---|
| 1 | MESGPAIFAGTVFCLFGAALLAWTVTRARQHRPVALGVNRVASTTVA | GTVATMSLALGAWCFTRL | 43547 | 43745 | + | 8 |
| 2 | VAPALRRGCTVPGVPA | GTPGTAVFGVRGQACRCGPAQWIRRGTDATDTTDATDATDGYDGRAG | 50179 | 50371 | + | 8 |
| 3 | VPGVPAGTPGTAVFGVRGQACRCGPA | QWIRRGTDATDTTDATDATDGYDGRAG | 50209 | 50371 | + | 7 |
| 4 | VPAGTPGTAVFGVRGQACRCGPAQW | IRRGTDATDTTDATDATDGYDGRAG | 50218 | 50371 | + | 8 |
| 5 | MRRSRVRIPKAAPLEPQARSFCPWGFLLFEARR | DTPSEPPAAGGRLCLGPLPAPLGRR...GVVSARLPAGESGGIPVRESLVMLG | 53418 | 53673 | + | 6 |
| 6 | VTRVALDRTGRDAQARARAIPLTDL | SDVSRQAWSVRVFRAGSTTRRIGMR | 54361 | 54514 | + | 6 |
| 7 | MEQKIAAFMNEQDLGSLEQGYRDDAPMPVL | ATPAAAVATCWYVGGAAAGAAAVVGAYVTGRVVG | 54543 | 54738 | + | 30 |
| 8 | MSRIVDSVVAVTPTDLTDDSQGEFGAMAPVQ | ATPTLVAFGVGFAGGAGAAWVTVQAYEAGAND | 54773 | 54965 | + | 31 |
| 9 | MGAAFQISGPPSHPRFQREDDCVWEAALDGPLIGQVVREMAGPQSVVS | SAPRHTFSWPVVVGR | 63354 | 63546 | + | 8 |
| 10 | VPVFRSSPSMTRADRMPSCSVHRMPDAAAEPCAADHSTGAAVEG | AGPAAAECAADDKAATVKAIVARVRFGDAIRMRVTRLPGRRCGA | 63660 | 63927 | + | 8 |
| 11 | MTRADRMPSCSVHRMPDAAAEPCAADHSTGAAVEGAGPA | AAECAADDKAATVKAIVARVRFGDAIRMRVTRLPGRRCGA | 63687 | 63927 | + | 9 |
| 12 | MPSCSVHRMPDAAAEPCAADHSTGAAVEGAGPAAAE | CAADDKAATVKAIVARVRFGDAIRMRVTRLPGRRCGA | 63705 | 63927 | + | 9 |
| 13 | MPDAAAEPCAADHSTGAAVEGAGPAAAECAAD | DKAATVKAIVARVRFGDAIRMRVTRLPGRRCGA | 63729 | 63927 | + | 9 |
| 14 | VHDCVCLLRRWKKSEGQFYAGVGGMNKSE | SNERSGVGLFQVNSSRITSLIPSMSGVFRH | 63919 | 64099 | + | 7 |
| 15 | VCLLRRWKKSEGQFYAGVGGMNKSESN | ERSGVGLFQVNSSRITSLIPSMSGVFRH | 63931 | 64099 | + | 7 |
| 16 | VGFSGIDCLTSGRHTVEQDQIGRR | GIATGRTCLRLRGLATTEQPVRAG | 64079 | 64226 | + | 6 |
| 17 | VHCPEEETAAVTAAIREAAEQAGR | IAFGETPVRFPFTTAVVKCYADAK | 66500 | 66353 | - | 6 |
| 18 | VIVHCPEEETAAVTAAIREAAEQAG | RIAFGETPVRFPFTTAVVKCYADAK | 66506 | 66353 | - | 7 |
| 19 | VFFQHDEVIVHCPEE | ETAAVTAAIREAAEQAGRIAFGETPVRFPFTTAVVKCYADAK | 66527 | 66353 | - | 8 |
| 20 | MAAELVFFQHDEVIVHCPEE | ETAAVTAAIREAAEQAGRIAFGETPVRFPFTTAVVKCYADAK | 66542 | 66353 | - | 8 |

### Results for AXE28036.1 [Streptomyces globosus] back to top

previous - next

#### Architecture

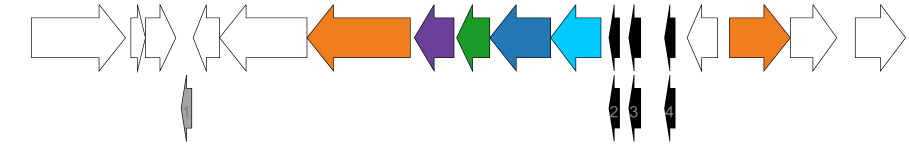

1000 nucleotides

Link to nucleotide sequence

| Accession | start | end | direction | length (aa) | Pfam/HMM | name | description | E-value |
|---|---|---|---|---|---|---|---|---|
| AXE28321.1 | 120011 | 121574 | + | 520 | PF13560<br>PF09339 | HTH_31<br>HTH_IclR | Helix-turn-helix domain<br>IclR helix-turn-helix domain | 1.40E-09<br>8.10E-04 |
| AXE28030.1 | 121649 | 121895 | + | 81 | PF01381<br>PF12844<br>PF13560<br>TIGR03070<br>TIGR02607 | HTH_3<br>HTH_19<br>HTH_31<br>TIGR03070<br>TIGR02607 | Helix-turn-helix<br>Helix-turn-helix domain<br>Helix-turn-helix domain<br>couple_hipB: transcriptional regulator, y4mF family<br>antidote_HigA: addiction module antidote protein, HigA family | 4.90E-13<br>1.80E-07<br>7.20E-07<br>3.00E-06<br>4.50E-05 |
| AXE28031.1 | 121891 | 122395 | + | 167 | NO PFAM MATCH | - | - | - |
| AXE28032.1 | 123124 | 122686 | - | 145 | NO PFAM MATCH | - | - | - |
| AXE28033.1 | 124581 | 123123 | - | 485 | PF13450<br>TIGR03467<br>TIGR00292<br>TIGR02734<br>TIGR01317 | NAD_binding_8<br>TIGR03467<br>TIGR00292<br>TIGR02734<br>TIGR01317 | NAD(P)-binding Rossmann-like domain<br>HpnE: squalene-associated FAD-dependent desaturase<br>TIGR00292: thiazole biosynthesis enzyme<br>crtI_fam: phytoene desaturase<br>GOGAT_sm_gam: glutamate synthase, NADH/NADPH, small subunit | 1.30E-10<br>5.70E-10<br>9.80E-09<br>1.90E-08<br>3.00E-08 |
| AXE28034.1 | 126296 | 124577 | - | 572 | TIGR02204<br>TIGR02203<br>TIGR00958<br>TIGR03796<br>TIGR01846 | TIGR02204<br>TIGR02203<br>TIGR00958<br>TIGR03796<br>TIGR01846 | MsbA_rel: ABC transporter, permease/ATP-binding protein<br>MsbA_lipidA: lipid A export permease/ATP-binding protein MsbA<br>3a01208: antigen peptide transporter 2<br>NHLM_micro_ABC1: NHLM bacteriocin system ABC transporter, peptidase/ATP-binding protein<br>type_I_sec_HlyB: type I secretion system ATPase | 7.20E-108<br>4.30E-104<br>5.60E-86<br>3.40E-83<br>9.60E-83 |
| AXE28322.1 | 127010 | 126356 | - | 217 | PF13649<br>PF08241<br>PF13489<br>TIGR01934<br>TIGR01983 | Methyltransf_25<br>Methyltransf_11<br>Methyltransf_23<br>TIGR01934<br>TIGR01983 | Methyltransferase domain<br>Methyltransferase domain<br>Methyltransferase domain<br>MenG_MenH_UbiE: ubiquinone/menaquinone biosynthesis methyltransferase<br>UbiG: 3-demethylubiquinone-9 3-O-methyltransferase | 1.50E-13<br>2.00E-09<br>9.50E-09<br>7.10E-06<br>8.10E-06 |
| AXE28035.1 | 127618 | 127072 | - | 181 | LinL | LinL | LinL | 7.40E-63 |
| AXE28036.1 | 128631 | 127614 | - | 338 | LinE<br>LinH<br>PF12697<br>TIGR02427<br>PF12146 | LinE<br>LinH<br>Abhydrolase_6<br>TIGR02427<br>Hydrolase_4 | LinE Homolog<br>LinH (Fusion of LinE/LinG)<br>Alpha/beta hydrolase family<br>protocat_pcaD: 3-oxoadipate enol-lactonase<br>Serine aminopeptidase, S33 | 9.70E-136<br>3.60E-81<br>3.00E-17<br>5.40E-12<br>7.80E-11 |
| AXE28037.1 | 129464 | 128627 | - | 278 | LinH<br>LinG | LinH<br>LinG | LinH (Fusion of LinE/LinG)<br>LinG Homolog | 1.40E-105<br>5.40E-103 |
| AXE28038.1 | 129771 | 129591 | - | 59 | LinA | LinA | LinA (linaridin precursor) | 3.60E-09 |
| AXE28039.1 | 130125 | 129924 | - | 66 | LinA | LinA | LinA (linaridin precursor) | 8.50E-10 |
| AXE28040.1 | 130692 | 130509 | - | 60 | LinA | LinA | LinA (linaridin precursor) | 1.40E-10 |
| AXE28041.1 | 131399 | 130886 | - | 170 | TIGR02983<br>TIGR02937<br>PF04542<br>PF08281<br>TIGR02985 | TIGR02983<br>TIGR02937<br>Sigma70_r2<br>Sigma70_r4_2<br>TIGR02985 | SigE-fam_strep: RNA polymerase sigma-70 factor, sigma-E family<br>sigma70-ECF: RNA polymerase sigma factor, sigma-70 family<br>Sigma-70 region 2<br>Sigma-70, region 4<br>Sig70_bacteroi1: RNA polymerase sigma-70 factor, Bacteroides expansion family 1 | 4.30E-19<br>2.50E-15<br>3.30E-10<br>3.50E-09<br>5.10E-08 |
| AXE28042.1 | 131593 | 132607 | + | 337 | TIGR01188<br>TIGR03864<br>TIGR01288<br>TIGR03873<br>TIGR03740 | TIGR01188<br>TIGR03864<br>TIGR01288<br>TIGR03873<br>TIGR03740 | drrA: daunorubicin resistance ABC transporter, ATP-binding protein<br>PQQ_ABC_ATP: ABC transporter, ATP-binding subunit, PQQ-dependent alcohol dehydrogenase system<br>nodI: nodulation ABC transporter NodI<br>F420-0_ABC_ATP: proposed F420-0 ABC transporter, ATP-binding protein<br>galliderm_ABC: lantibiotic protection ABC transporter, ATP-binding subunit | 7.30E-63<br>3.20E-43<br>8.80E-43<br>1.10E-32<br>1.70E-32 |
| AXE28043.1 | 132603 | 133368 | + | 254 | PF01061<br>PF12698<br>TIGR03062 | ABC2_membrane<br>ABC2_membrane_3<br>TIGR03062 | ABC-2 type transporter<br>ABC-2 family transporter protein<br>pip_yhgE_Cterm: YhgE/Pip C-terminal domain | 9.30E-14<br>4.80E-06<br>5.20E-04 |
| AXE28044.1 | 133672 | 134521 | + | 282 | NO PFAM MATCH | - | - | - |

| index | leader | core | start | end | dir | score |
|---|---|---|---|---|---|---|
| 1 | VNGIDKQDLDQLITHTYAEI | KTAITSHSEKSLLMYSQALSTLLELRRQLAENPPTT | 122660 | 122489 | - | 6 |
| 2 | MSVLIEKVNAGAARAAESPVAPVM | ATPAAFTVGFVGGAKATAIIVAAAGVGAAVKACQK | 129771 | 129591 | - | 30 |
| 3 | MLIQKVNLNPAVNGELHAGSPAGAAAPVM | ATPAIIATPVIAKTAGYVVGGAVVTGAAAVAYKTATG | 130125 | 129924 | - | 32 |
| 4 | MLLHKVNQNASLTGSLSQRTLEGAVSPVM | ATPSLVFTPCAGKAAVLVTAAAVGYATAKVK | 130692 | 130509 | - | 30 |

### Results for WP_148235529.1 [Streptomyces griseus] back to top

previous - next

#### Architecture

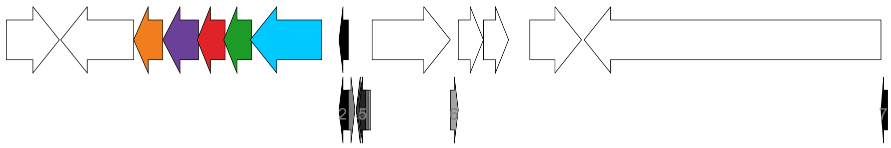

1000 nucleotides

Link to nucleotide sequence

| Accession | start | end | direction | length (aa) | Pfam/HMM | name | description | E-value |
|---|---|---|---|---|---|---|---|---|
| WP_164361063.1 | 0 | 1114 | + | 370 | NO PFAM MATCH | - | - | - |
| WP_012381884.1 | 2695 | 1150 | - | 514 | NO PFAM MATCH | - | - | - |
| WP_003970677.1 | 3324 | 2691 | - | 210 | TIGR02857<br>TIGR01189<br>TIGR02673<br>TIGR02982<br>TIGR03258 | TIGR02857<br>TIGR01189<br>TIGR02673<br>TIGR02982<br>TIGR03258 | CydD: thiol reductant ABC exporter, CydD subunit<br>ccmA: heme ABC exporter, ATP-binding protein CcmA<br>FtsE: cell division ATP-binding protein FtsE<br>heterocyst_DevA: ABC exporter ATP-binding subunit, DevA family<br>PhnT: 2-aminoethylphosphonate ABC transport system, ATP-binding component PhnT | 5.70E-28<br>2.70E-25<br>1.60E-24<br>3.30E-22<br>1.10E-21 |
| WP_012381886.1 | 4058 | 3320 | - | 245 | PF13649<br>PF08241<br>PF08242<br>TIGR02021<br>PF13489 | Methyltransf_25<br>Methyltransf_11<br>Methyltransf_12<br>TIGR02021<br>Methyltransf_23 | Methyltransferase domain<br>Methyltransferase domain<br>Methyltransferase domain<br>BchM-ChlM: magnesium protoporphyrin O-methyltransferase<br>Methyltransferase domain | 1.60E-16<br>2.20E-15<br>2.80E-11<br>1.20E-10<br>3.40E-10 |
| WP_003970679.1 | 4623 | 4050 | - | 190 | TIGR00521<br>TIGR02113<br>PF02441 | TIGR00521<br>TIGR02113<br>Flavoprotein | coaBC_dfp: phosphopantothenoylcysteine decarboxylase / phosphopantothenate--cysteine ligase<br>coaC_strep: phosphopantothenoylcysteine decarboxylase<br>Flavoprotein | 2.30E-14<br>3.70E-12<br>1.50E-11 |
| WP_003970680.1 | 5186 | 4619 | - | 188 | LinL | LinL | LinL | 1.10E-51 |
| WP_148235529.1 | 6684 | 5175 | - | 502 | LinH<br>LinG<br>LinE<br>PF12697<br>TIGR02427 | LinH<br>LinG<br>LinE<br>Abhydrolase_6<br>TIGR02427 | LinH (Fusion of LinE/LinG)<br>LinG Homolog<br>LinE Homolog<br>Alpha/beta hydrolase family<br>protocat_pcaD: 3-oxoadipate enol-lactonase | 2.70E-162<br>2.40E-67<br>3.20E-65<br>2.20E-06<br>2.60E-05 |
| WP_003970682.1 | 7244 | 7049 | - | 64 | LinA | LinA | LinA (linaridin precursor) | 6.80E-14 |
| WP_164361064.1 | 7766 | 9416 | + | 549 | PF03704<br>PF13191<br>TIGR03903<br>PF00486 | BTAD<br>AAA_16<br>TIGR03903<br>Trans_reg_C | Bacterial transcriptional activator domain<br>AAA ATPase domain<br>TOMM_kin_cyc: TOMM system kinase/cyclase fusion protein<br>Transcriptional regulatory protein, C terminal | 4.00E-43<br>3.40E-12<br>2.10E-07<br>2.70E-04 |
| WP_003970684.1 | 9596 | 10130 | + | 177 | PF01613<br>TIGR02296<br>TIGR03615 | Flavin_Reduct<br>TIGR02296<br>TIGR03615 | Flavin reductase like domain<br>HpaC: 4-hydroxyphenylacetate 3-monooxygenase, reductase component<br>RutF: pyrimidine utilization flavin reductase protein F | 8.90E-33<br>8.10E-24<br>1.80E-23 |
| WP_164361065.1 | 10133 | 10649 | + | 171 | NO PFAM MATCH | - | - | - |
| WP_012381890.1 | 11093 | 12191 | + | 365 | PF03704<br>PF13428<br>PF00486<br>PF13432<br>PF08279 | BTAD<br>TPR_14<br>Trans_reg_C<br>TPR_16<br>HTH_11 | Bacterial transcriptional activator domain<br>Tetratricopeptide repeat<br>Transcriptional regulatory protein, C terminal<br>Tetratricopeptide repeat<br>HTH domain | 7.20E-45<br>8.00E-06<br>3.30E-04<br>6.20E-04<br>6.70E-04 |
| WP_164361066.1 | 18569 | 12263 | - | 2101 | TIGR02813<br>PF00109<br>TIGR01746<br>PF08659<br>TIGR03443 | TIGR02813<br>ketoacyl-synt<br>TIGR01746<br>KR<br>TIGR03443 | omega_3_PfaA: polyketide-type polyunsaturated fatty acid synthase PfaA<br>Beta-ketoacyl synthase, N-terminal domain<br>Thioester-redct: thioester reductase domain<br>KR domain<br>alpha_am_amid: L-aminoadipate-semialdehyde dehydrogenase | 1.60E-117<br>5.30E-86<br>2.70E-85<br>5.30E-64<br>7.50E-58 |

| index | leader | core | start | end | dir | score |
|---|---|---|---|---|---|---|
| 1 | LQSNRVAFRCPSSEPMLSERDLAVP | SCPSPAASSELNALCTTSATRSAGIPQSW | 7222 | 7387 | + | 7 |
| 2 | MRLDSIATQETATALPESMATQDFANSVLAGAVPGFHSDAETPAM | ATPAVAQFVIQGSTICLVC | 7244 | 7049 | - | 34 |
| 3 | LLGGLGTCEASGAGTPTDTRRTGGTTGRSSKP | ASSRNRPTGAWQLVQIQRDLRLVKLILMLRVGC | 7626 | 7428 | - | 8 |
| 4 | VQGLGPSPHKTRTRNGPCSGASAHARR | PAPGHRPTREGPGERPGVRRNRHRHEIARQARGNLSKFSGT | 7676 | 7469 | - | 6 |
| 5 | VAAHDPGFRRSGLRIPAHRAGARSKPTQNSDKERTLLGGLGTCEASGAGT | PTDTRRTGGTTGRSSKPASSRNRPTGAWQLVQIQRDLRLVKLILMLRVGC | 7731 | 7428 | - | 6 |
| 6 | VRCRPRRRGGRSPAYGFRGRTRTVYGAVAR | RAPVWVTGYPGMRVREIPRSMFSV | 9412 | 9577 | + | 6 |
| 7 | VDRGPGREHRRRGRELRVRVGRGHLRLHRQPARTLHQ | LTPVATHEGKRDHRVR | 18711 | 18549 | - | 10 |

### Results for SEE39163.1 [Streptomyces griseus] back to top

previous - next

#### Architecture

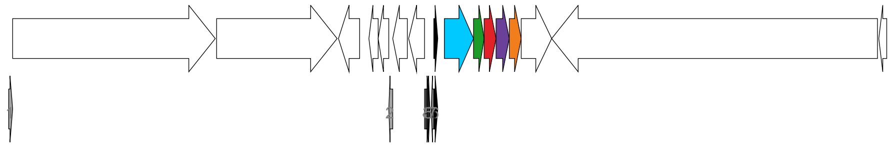

1000 nucleotides

Link to nucleotide sequence

| Accession | start | end | direction | length (aa) | Pfam/HMM | name | description | E-value |
|---|---|---|---|---|---|---|---|---|
| SEE39002.1 | 1100849 | 1111418 | + | 3522 | TIGR02813<br>PF00109<br>PF00698<br>TIGR00128<br>TIGR03150 | TIGR02813<br>ketoacyl-synt<br>Acyl_transf_1<br>TIGR00128<br>TIGR03150 | omega_3_PfaA: polyketide-type polyunsaturated fatty acid synthase PfaA<br>Beta-ketoacyl synthase, N-terminal domain<br>Acyl transferase domain<br>fabD: malonyl CoA-acyl carrier protein transacylase<br>fabF: beta-ketoacyl-acyl-carrier-protein synthase II | 1.50E-206<br>2.00E-173<br>2.00E-109<br>7.60E-105<br>4.00E-86 |

| | | | | | | | | |
|---|---|---|---|---|---|---|---|---|
| SEE39028.1 | 1111445 | 1117751 | + | 2101 | TIGR02813 | TIGR02813 | omega_3_PfaA: polyketide-type polyunsaturated fatty acid synthase PfaA | 1.30E-118 |
| | | | | | PF00109 | ketoacyl-synt | Beta-ketoacyl synthase, N-terminal domain | 3.10E-86 |
| | | | | | TIGR01746 | TIGR01746 | Thioester-redct: thioester reductase domain | 2.70E-85 |
| | | | | | PF08659 | KR | KR domain | 5.20E-64 |
| | | | | | TIGR03443 | TIGR03443 | alpha_am_amid: L-aminoadipate-semialdehyde dehydrogenase | 7.50E-58 |
| SEE39047.1 | 1118921 | 1117823 | - | 365 | PF03704 | BTAD | Bacterial transcriptional activator domain | 7.20E-45 |
| | | | | | PF13428 | TPR_14 | Tetratricopeptide repeat | 8.00E-06 |
| | | | | | PF00486 | Trans_reg_C | Transcriptional regulatory protein, C terminal | 3.30E-04 |
| | | | | | PF13432 | TPR_16 | Tetratricopeptide repeat | 6.20E-04 |
| | | | | | PF08279 | HTH_11 | HTH domain | 6.70E-04 |
| SEE39069.1 | 1119860 | 1119365 | - | 164 | NO PFAM MATCH | - | - | - |
| SEE39091.1 | 1120418 | 1119884 | - | 177 | PF01613 | Flavin_Reduct | Flavin reductase like domain | 8.90E-33 |
| | | | | | TIGR02296 | TIGR02296 | HpaC: 4-hydroxyphenylacetate 3-monooxygenase, reductase component | 8.10E-24 |
| | | | | | TIGR03615 | TIGR03615 | RutF: pyrimidine utilization flavin reductase protein F | 1.80E-23 |
| SEE39112.1 | 1121381 | 1120598 | - | 260 | PF13191 | AAA_16 | AAA ATPase domain | 1.70E-12 |
| | | | | | TIGR03903 | TIGR03903 | TOMM_kin_cyc: TOMM system kinase/cyclase fusion protein | 1.70E-08 |
| SEE39127.1 | 1122247 | 1121452 | - | 264 | PF03704 | BTAD | Bacterial transcriptional activator domain | 7.70E-44 |
| | | | | | PF00486 | Trans_reg_C | Transcriptional regulatory protein, C terminal | 2.50E-04 |
| SEE39148.1 | 1122769 | 1122964 | + | 64 | LinA | LinA | LinA (linaridin precursor) | 6.80E-14 |
| SEE39163.1 | 1123317 | 1124838 | + | 506 | LinH | LinH | LinH (Fusion of LinE/LinG) | 3.60E-163 |
| | | | | | LinG | LinG | LinG Homolog | 4.90E-68 |
| | | | | | LinE | LinE | LinE Homolog | 3.30E-65 |
| | | | | | PF12697 | Abhydrolase_6 | Alpha/beta hydrolase family | 5.30E-06 |
| | | | | | TIGR02427 | TIGR02427 | protocat_pcaD: 3-oxoadipate enol-lactonase | 2.60E-05 |
| SEE39183.1 | 1124827 | 1125394 | + | 188 | LinL | LinL | LinL | 1.10E-51 |
| SEE39196.1 | 1125390 | 1125963 | + | 190 | TIGR00521 | TIGR00521 | coaBC_dfp: phosphopantothenoylcysteine decarboxylase / phosphopantothenate--cysteine ligase | 2.30E-14 |
| | | | | | TIGR02113 | TIGR02113 | coaC_strep: phosphopantothenoylcysteine decarboxylase | 3.70E-12 |
| | | | | | PF02441 | Flavoprotein | Flavoprotein | 1.50E-11 |
| SEE39219.1 | 1125955 | 1126693 | + | 245 | PF13649 | Methyltransf_25 | Methyltransferase domain | 1.60E-16 |
| | | | | | PF08241 | Methyltransf_11 | Methyltransferase domain | 2.20E-15 |
| | | | | | PF08242 | Methyltransf_12 | Methyltransferase domain | 2.80E-11 |
| | | | | | TIGR02021 | TIGR02021 | BchM-ChlM: magnesium protoporphyrin O-methyltransferase | 1.20E-10 |
| | | | | | PF13489 | Methyltransf_23 | Methyltransferase domain | 3.40E-10 |
| SEE39233.1 | 1126689 | 1127322 | + | 210 | TIGR02857 | TIGR02857 | CydD: thiol reductant ABC exporter, CydD subunit | 1.20E-27 |
| | | | | | TIGR01189 | TIGR01189 | ccmA: heme ABC exporter, ATP-binding protein CcmA | 3.10E-25 |
| | | | | | TIGR02673 | TIGR02673 | FtsE: cell division ATP-binding protein FtsE | 1.80E-24 |
| | | | | | TIGR02982 | TIGR02982 | heterocyst_DevA: ABC exporter ATP-binding subunit, DevA family | 9.00E-22 |
| | | | | | TIGR03258 | TIGR03258 | PhnT: 2-aminoethylphosphonate ABC transport system, ATP-binding component PhnT | 1.70E-21 |
| SEE39254.1 | 1127318 | 1128863 | + | 514 | NO PFAM MATCH | - | - | - |
| SEE39266.1 | 1145792 | 1128899 | - | 5630 | NO PFAM MATCH | - | - | - |
| SEE39289.1 | 1146275 | 1145906 | - | 122 | NO PFAM MATCH | - | - | - |

| index | leader | core | start | end | dir | score |
|---|---|---|---|---|---|---|
| 1 | VEADDHSLITEHIEDLLAQWRGTGDGP | ASPADDSDLNAASDEELFSLVESNRKA | 1100678 | 1100843 | + | 6 |
| 2 | VRCRPRRRGGRSPAYGFRGRTRTVYGAVAR | RAPVWVTGYPGMRVREIPRSMFSV | 1120602 | 1120437 | - | 6 |
| 3 | VAAYDPGFRRSGLRIPAHRAGARSKPTQNSDKERALLGGLGTCEASG | AGTPTDTGRTGGTTGRSSKPASSRNRPTGAWQLVQIQRDLRLVKLILM | 1122282 | 1122570 | + | 7 |
| 4 | VQGLGPSPHKTRTRSGPCSGASAHARR | PAPGHRPTREGPGERPGVRRNRHRHEIARQARGNLSKFSGT | 1122337 | 1122544 | + | 6 |
| 5 | LLGGLGTCEASGAGTPTDTGRTGGTTGRSS | KPASSRNRPTGAWQLVQIQRDLRLVKLILM | 1122387 | 1122570 | + | 8 |
| 6 | MRLDSIATQETATALPESMATQDFANSVLAGAVPGFHSDAETPAM | ATPAVAQFVIQGSTICLVC | 1122769 | 1122964 | + | 34 |
| 7 | LQSNRVAFRCPSSEPMLSERDLAVP | SCPSPAASSELNALCTTSATRSAGIPQSW | 1122791 | 1122626 | - | 7 |

### Results for SQA25558.1 [Streptomyces griseus] back to top

previous - next

#### Architecture

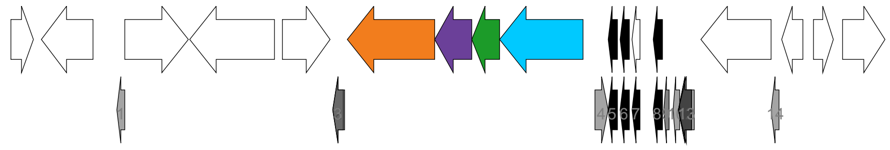

1000 nucleotides

Link to nucleotide sequence

| Accession | start | end | direction | length (aa) | Pfam/HMM | name | description | E-value |
|---|---|---|---|---|---|---|---|---|
| SQA25550.1 | 52254 | 52755 | + | 166 | PF00440 | TetR_N | Bacterial regulatory proteins, tetR family | 3.90E-12 |
| | | | | | PF09339 | HTH_IclR | IclR helix-turn-helix domain | 1.60E-04 |
| | | | | | PF17926 | TetR_C_21 | Tetracyclin repressor-like, C-terminal domain | 7.60E-04 |
| SQA25551.1 | 54051 | 52929 | - | 373 | NO PFAM MATCH | - | - | - |
| SQA25552.1 | 54758 | 56126 | + | 455 | PF13428 | TPR_14 | Tetratricopeptide repeat | 4.70E-09 |
| | | | | | PF14559 | TPR_19 | Tetratricopeptide repeat | 7.40E-09 |
| | | | | | PF13181 | TPR_8 | Tetratricopeptide repeat | 1.20E-05 |
| | | | | | PF12895 | ANAPC3 | Anaphase-promoting complex, cyclosome, subunit 3 | 1.80E-05 |
| | | | | | PF13432 | TPR_16 | Tetratricopeptide repeat | 9.80E-05 |

| | | | | | | | | |
|---|---|---|---|---|---|---|---|---|
| SQA25553.1 | 58012 | 56158 | - | 617 | NO PFAM MATCH | - | - | - |
| SQA25554.1 | 58201 | 59215 | + | 337 | NO PFAM MATCH | - | - | - |
| SQA25555.1 | 61507 | 59611 | - | 631 | TIGR02204<br>TIGR02203<br>TIGR00958<br>TIGR03796<br>TIGR03797 | TIGR02204<br>TIGR02203<br>TIGR00958<br>TIGR03796<br>TIGR03797 | MsbA_rel: ABC transporter, permease/ATP-binding protein<br>MsbA_lipidA: lipid A export permease/ATP-binding protein MsbA<br>3a01208: antigen peptide transporter 2<br>NHLM_micro_ABC1: NHLM bacteriocin system ABC transporter, peptidase/ATP-binding protein<br>NHLM_micro_ABC2: NHLM bacteriocin system ABC transporter, ATP-binding protein | 9.80E-102<br>1.70E-92<br>6.30E-85<br>1.40E-82<br>3.40E-77 |
| SQA25556.1 | 62316 | 61503 | - | 270 | PF13649<br>PF08241<br>PF08242<br>PF13489<br>PF13847 | Methyltransf_25<br>Methyltransf_11<br>Methyltransf_12<br>Methyltransf_23<br>Methyltransf_31 | Methyltransferase domain<br>Methyltransferase domain<br>Methyltransferase domain<br>Methyltransferase domain<br>Methyltransferase domain | 2.30E-16<br>7.20E-12<br>5.50E-08<br>1.20E-07<br>1.60E-06 |
| SQA25557.1 | 62921 | 62312 | - | 202 | LinL | LinL | LinL | 2.00E-65 |
| SQA25558.1 | 64759 | 62917 | - | 613 | LinH<br>LinG<br>LinE<br>LinL<br>PF12697 | LinH<br>LinG<br>LinE<br>LinL<br>Abhydrolase_6 | LinH (Fusion of LinE/LinG)<br>LinG Homolog<br>LinE Homolog<br>LinL<br>Alpha/beta hydrolase family | 1.50E-253<br>5.50E-112<br>3.40E-94<br>3.70E-35<br>2.00E-14 |
| SQA25559.1 | 65502 | 65301 | - | 66 | LinA | LinA | LinA (linaridin precursor) | 2.10E-13 |
| SQA25560.1 | 65754 | 65559 | - | 64 | LinA | LinA | LinA (linaridin precursor) | 1.20E-10 |
| SQA25561.1 | 65998 | 65824 | - | 57 | NO PFAM MATCH | - | - | - |
| SQA25562.1 | 66493 | 66289 | - | 67 | LinA | LinA | LinA (linaridin precursor) | 1.10E-11 |
| SQA25563.1 | 68860 | 67330 | - | 509 | TIGR01072<br>PF00275<br>TIGR01356<br>TIGR03070<br>PF01381 | TIGR01072<br>EPSP_synthase<br>TIGR01356<br>TIGR03070<br>HTH_3 | murA: UDP-N-acetylglucosamine 1-carboxyvinyltransferase<br>EPSP synthase (3-phosphoshikimate 1-carboxyvinyltransferase)<br>aroA: 3-phosphoshikimate 1-carboxyvinyltransferase<br>couple_hipB: transcriptional regulator, y4mF family<br>Helix-turn-helix | 4.00E-91<br>2.20E-39<br>1.10E-19<br>2.10E-14<br>1.70E-12 |
| SQA25564.1 | 69542 | 69089 | - | 150 | PF01042<br>TIGR00004<br>TIGR03610 | Ribonuc_L-PSP<br>TIGR00004<br>TIGR03610 | Endoribonuclease L-PSP<br>TIGR00004: reactive intermediate/imine deaminase<br>RutC: pyrimidine utilization protein C | 9.10E-15<br>4.40E-10<br>1.30E-08 |
| SQA25565.1 | 69788 | 70223 | + | 144 | PF18029 | Glyoxalase_6 | Glyoxalase-like domain | 1.60E-23 |
| SQA25566.1 | 70419 | 71337 | + | 305 | PF01636<br>TIGR02172<br>TIGR00938 | APH<br>TIGR02172<br>TIGR00938 | Phosphotransferase enzyme family<br>Fb_sc_TIGR02172: fibrobacter succinogenes paralogous family TIGR02172<br>thrB_alt: homoserine kinase | 1.50E-38<br>1.70E-06<br>7.30E-04 |

| index | leader | core | start | end | dir | score |
|---|---|---|---|---|---|---|
| 1 | VTSLRGDLRTSIARQRRPKRDVLAM | STEVRHVDDRLAVYASGEGARLACGR | 54734 | 54578 | - | 6 |
| 2 | VGGGANGPARPAGGPCRPARPGDCRGTGAARPRARRSSRSPAD | RPPGRTETGGPEAPRTETGRPLPALGPVGAGTPVGAGTPAA | 59528 | 59273 | - | 6 |
| 3 | VGSAASTAVVGGGANGPARPAGGPC...ARPGDCRGTGAARPRARRSSRSPAD | RPPGRTETGGPEAPRTETGRPLPALGPVGAGTPVGAGTPAA | 59555 | 59273 | - | 6 |
| 4 | VRDGLPPGRARAGRGRSCRGRAASP...VAWAGDRPGGARPHATRAPSLPAGP | GTVAAAPRAGAGATDLPPGARAAVAAHRPLGRHRDQRVPPPVTGTA | 64999 | 65296 | + | 6 |
| 5 | MSMSPTPAALRGAGGLSESDPGRALSSLAPVT | ATPGVVAGVALGVALVNAFAAGYNHCGGNVELPM | 65502 | 65301 | - | 31 |
| 6 | MNKSSAPAVLTATGALTESDAGVTLPTLAPVA | ATPVAIAATMGVAFVAGYAAGRAATGNVELPM | 65754 | 65559 | - | 30 |
| 7 | MNTASIPAVLSETGALSESDHGRALLDTVPVAS | VTFTMTACVEVSVCLTGSVIQLPQ | 65998 | 65824 | - | 20 |
| 8 | MPEFRQPGWTRGVAPLDESAGGQVFGGASPVA | ATPAVVATAGAVVVAFAAGVAARHLANGGNVELPM | 66493 | 66289 | - | 30 |
| 9 | MPAEWSSLCVGRSASCGAHHRSS | VHAGAGSRVRLRAPGWAPVTAHR | 66643 | 66502 | - | 7 |
| 10 | VRSKTLPPFDVTSGLAEYPSAVRAAAEMASRSVGE | APPIALNTTRSRAG | 66862 | 66712 | - | 6 |
| 11 | VAHSGSELPGEATSALERRVGVDDI...TGLPEAASHSVVLAGSPAGFEPPPV | IAPTALVRQGRSARSGRPEQVPVRQRAQQDAPAV | 67112 | 66833 | - | 6 |
| 12 | VDAVAHSGSELPGEATSALERRVGV...TGLPEAASHSVVLAGSPAGFEPPPV | IAPTALVRQGRSARSGRPEQVPVRQRAQQDAPAV | 67121 | 66833 | - | 6 |
| 13 | VDSWVGCRGVRTCPTVDAVAHSGSE...TGLPEAASHSVVLAGSPAGFEPPPV | IAPTALVRQGRSARSGRPEQVPVRQRAQQDAPAV | 67166 | 66833 | - | 6 |
| 14 | VRGGAVGCGRSLGRTRGPRPSLRRGARWGEACTRSRYTPVT | RTAHPHSSLISYLR | 69037 | 68869 | - | 6 |

### Results for BAG18344.1 [Streptomyces griseus subsp. griseus NBRC 13350]

back to top

previous - next

#### Architecture

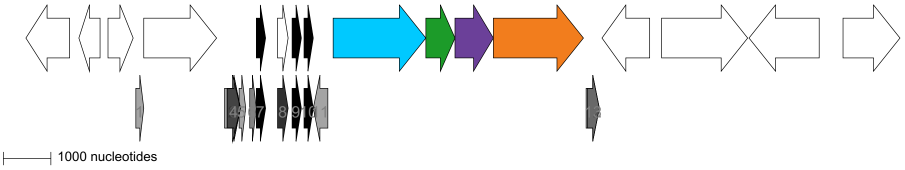

Link to nucleotide sequence

| Accession | start | end | direction | length (aa) | Pfam/HMM | name | description | E-value |
|---|---|---|---|---|---|---|---|---|

| | | | | | | | | |
|---|---|---|---|---|---|---|---|---|
| BAG18336.1 | 1769207 | 1768289 | - | 305 | PF01636<br>TIGR02172<br>TIGR00938 | APH<br>TIGR02172<br>TIGR00938 | Phosphotransferase enzyme family<br>Fb_sc_TIGR02172: fibrobacter succinogenes paralogous family TIGR02172<br>thrB_alt: homoserine kinase | 1.50E-38<br>1.70E-06<br>7.30E-04 |
| BAG18337.1 | 1769838 | 1769403 | - | 144 | PF18029 | Glyoxalase_6 | Glyoxalase-like domain | 1.60E-23 |
| BAG18338.1 | 1770006 | 1770537 | + | 176 | PF01042<br>TIGR00004<br>TIGR03610 | Ribonuc_L-PSP<br>TIGR00004<br>TIGR03610 | Endoribonuclease L-PSP<br>TIGR00004: reactive intermediate/imine deaminase<br>RutC: pyrimidine utilization protein C | 1.40E-14<br>6.90E-10<br>2.10E-08 |
| BAG18339.1 | 1770766 | 1772296 | + | 509 | TIGR01072<br>PF00275<br>TIGR01356<br>TIGR03070<br>PF01381 | TIGR01072<br>EPSP_synthase<br>TIGR01356<br>TIGR03070<br>HTH_3 | murA: UDP-N-acetylglucosamine 1-carboxyvinyltransferase<br>EPSP synthase (3-phosphoshikimate 1-carboxyvinyltransferase)<br>aroA: 3-phosphoshikimate 1-carboxyvinyltransferase<br>couple_hipB: transcriptional regulator, y4mF family<br>Helix-turn-helix | 4.00E-91<br>2.20E-39<br>1.10E-19<br>2.10E-14<br>1.70E-12 |
| BAG18340.1 | 1773133 | 1773337 | + | 67 | LinA | LinA | LinA (linaridin precursor) | 1.10E-11 |
| BAG18341.1 | 1773586 | 1773802 | + | 71 | NO PFAM MATCH | - | - | - |
| BAG18342.1 | 1773872 | 1774067 | + | 64 | LinA | LinA | LinA (linaridin precursor) | 1.20E-10 |
| BAG18343.1 | 1774124 | 1774325 | + | 66 | LinA | LinA | LinA (linaridin precursor) | 2.10E-13 |
| BAG18344.1 | 1774759 | 1776709 | + | 649 | LinH<br>LinG<br>LinE<br>LinL<br>PF12697 | LinH<br>LinG<br>LinE<br>LinL<br>Abhydrolase_6 | LinH (Fusion of LinE/LinG)<br>LinG Homolog<br>LinE Homolog<br>LinL<br>Alpha/beta hydrolase family | 2.40E-257<br>2.70E-114<br>3.80E-94<br>3.90E-35<br>2.30E-14 |
| BAG18345.1 | 1776705 | 1777314 | + | 202 | LinL | LinL | LinL | 2.00E-65 |
| BAG18346.1 | 1777310 | 1778123 | + | 270 | PF13649<br>PF08241<br>PF08242<br>PF13489<br>PF13847 | Methyltransf_25<br>Methyltransf_11<br>Methyltransf_12<br>Methyltransf_23<br>Methyltransf_31 | Methyltransferase domain<br>Methyltransferase domain<br>Methyltransferase domain<br>Methyltransferase domain<br>Methyltransferase domain | 2.30E-16<br>7.20E-12<br>5.50E-08<br>1.20E-07<br>1.60E-06 |
| BAG18347.1 | 1778119 | 1780015 | + | 631 | TIGR02204<br>TIGR02203<br>TIGR00958<br>TIGR03796<br>TIGR03797 | TIGR02204<br>TIGR02203<br>TIGR00958<br>TIGR03796<br>TIGR03797 | MsbA_rel: ABC transporter, permease/ATP-binding protein<br>MsbA_lipidA: lipid A export permease/ATP-binding protein MsbA<br>3a01208: antigen peptide transporter 2<br>NHLM_micro_ABC1: NHLM bacteriocin system ABC transporter, peptidase/ATP-binding protein<br>NHLM_micro_ABC2: NHLM bacteriocin system ABC transporter, ATP-binding protein | 9.80E-102<br>1.70E-92<br>6.30E-85<br>1.40E-82<br>3.40E-77 |
| BAG18348.1 | 1781410 | 1780411 | - | 332 | NO PFAM MATCH | - | - | - |
| BAG18349.1 | 1781674 | 1783468 | + | 597 | NO PFAM MATCH | - | - | - |
| BAG18350.1 | 1784985 | 1783500 | - | 494 | PF13428<br>PF14559<br>PF13181<br>PF12895<br>PF13432 | TPR_14<br>TPR_19<br>TPR_8<br>ANAPC3<br>TPR_16 | Tetratricopeptide repeat<br>Tetratricopeptide repeat<br>Tetratricopeptide repeat<br>Anaphase-promoting complex, cyclosome, subunit 3<br>Tetratricopeptide repeat | 1.00E-08<br>1.10E-08<br>1.90E-05<br>2.60E-05<br>1.70E-04 |
| BAG18351.1 | 1785497 | 1786697 | + | 399 | NO PFAM MATCH | - | - | - |

| index | leader | core | start | end | dir | score |
|---|---|---|---|---|---|---|
| 1 | VRGGAVGCGRSLGRTRGPRPSLRRGARWGEACTRSRYTPVT | RTAHPHSSLISYLR | 1770589 | 1770757 | + | 6 |
| 2 | VDSWVGCRGVRTCPTVDAVAHSGSE...TGLPEAASHSVVLAGSPAGFEPPPV | IAPTALVRQGRSARSGRPEQVPVRQRAQQDAPAV | 1772460 | 1772793 | + | 6 |
| 3 | VDAVAHSGSELPGEATSALERRVGV...TGLPEAASHSVVLAGSPAGFEPPPV | IAPTALVRQGRSARSGRPEQVPVRQRAQQDAPAV | 1772505 | 1772793 | + | 6 |
| 4 | VAHSGSELPGEATSALERRVGVDDI...TGLPEAASHSVVLAGSPAGFEPPPV | IAPTALVRQGRSARSGRPEQVPVRQRAQQDAPAV | 1772514 | 1772793 | + | 6 |
| 5 | VRSKTLPPFDVTSGLAEYPSAVRAAAEMASRSVGE | APPIALNTTRSRAG | 1772764 | 1772914 | + | 6 |
| 6 | MPAEWSSLCVGRSASCGAHHRSS | VHAGAGSRVRLRAPGWAPVTAHR | 1772983 | 1773124 | + | 7 |
| 7 | MPEFRQPGWTRGVAPLDESAGGQVFGGASPVA | ATPAVVATAGAVVVAFAAGVAARHLANGGNVELPM | 1773133 | 1773337 | + | 30 |
| 8 | MASPHRPPTSEEKSLNTASIPAVLSETGALSESDHGRALLDTVPVAS | VTFTMTACVEVSVCLTGSVIQLPQ | 1773586 | 1773802 | + | 9 |
| 9 | MNKSSAPAVLTATGALTESDAGVTLPTLAPVA | ATPVAIAATMGVAFVAGYAAGRAATGNVELPM | 1773872 | 1774067 | + | 30 |
| 10 | MSMSPTPAALRGAGGLSESDPGRALSSLAPVT | ATPGVVAGVALGVALVNAFAAGYNHCGGNVELPM | 1774124 | 1774325 | + | 31 |
| 11 | VRDGLPPGRARAGRGRSCRGRAASP...VAWAGDRPGGARPHATRAPSLPAGP | GTVAAAPRAGAGATDLPPGARAAVAAHRPLGRHRDQRVPPPVTGTA | 1774627 | 1774330 | - | 6 |
| 12 | VGSAASTAVVGGGANGPARPAGGPC...ARPGDCRGTGAARPRARRSSRSPAD | RPPGRTETGGPEAPRTETGRPLPALGPVGAGTPVGAGTPAA | 1780071 | 1780353 | + | 6 |
| 13 | VGGGANGPARPAGGPCRPARPGDCRGTGAARPRARRSSRSPAD | RPPGRTETGGPEAPRTETGRPLPALGPVGAGTPVGAGTPAA | 1780098 | 1780353 | + | 6 |

### Results for BAG23194.1 [Streptomyces griseus subsp. griseus NBRC 13350]

back to top

previous - next

#### Architecture

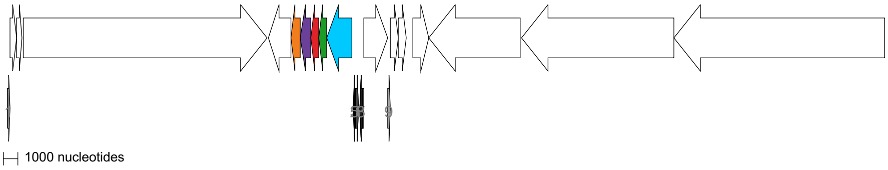

Link to nucleotide sequence

| Accession | start | end | direction | length (aa) | Pfam/HMM | name | description | E-value |
|---|---|---|---|---|---|---|---|---|
| BAG23186.1 | 7550833 | 7551253 | + | 139 | PF02575 | YbaB_DNA_bd | YbaB/EbfC DNA-binding family | 2.60E-09 |
| BAG23187.1 | 7551256 | 7551625 | + | 122 | NO PFAM MATCH | - | - | - |
| BAG23188.1 | 7551739 | 7568632 | + | 5630 | NO PFAM MATCH | - | - | - |
| BAG23189.1 | 7570213 | 7568668 | - | 514 | NO PFAM MATCH | - | - | - |
| BAG23190.1 | 7570842 | 7570209 | - | 210 | TIGR02857 | TIGR02857 | CydD: thiol reductant ABC exporter, CydD subunit | 1.20E-27 |
| | | | | | TIGR01189 | TIGR01189 | ccmA: heme ABC exporter, ATP-binding protein CcmA | 3.10E-25 |
| | | | | | TIGR02673 | TIGR02673 | FtsE: cell division ATP-binding protein FtsE | 1.80E-24 |
| | | | | | TIGR02982 | TIGR02982 | heterocyst_DevA: ABC exporter ATP-binding subunit, DevA family | 9.00E-22 |
| | | | | | TIGR03258 | TIGR03258 | PhnT: 2-aminoethylphosphonate ABC transport system, ATP-binding component PhnT | 1.70E-21 |
| BAG23191.1 | 7571576 | 7570838 | - | 245 | PF13649 | Methyltransf_25 | Methyltransferase domain | 1.60E-16 |
| | | | | | PF08241 | Methyltransf_11 | Methyltransferase domain | 2.20E-15 |
| | | | | | PF08242 | Methyltransf_12 | Methyltransferase domain | 2.80E-11 |
| | | | | | TIGR02021 | TIGR02021 | BchM-ChlM: magnesium protoporphyrin O-methyltransferase | 1.20E-10 |
| | | | | | PF13489 | Methyltransf_23 | Methyltransferase domain | 3.40E-10 |
| BAG23192.1 | 7572141 | 7571568 | - | 190 | TIGR00521 | TIGR00521 | coaBC_dfp: phosphopantothenoylcysteine decarboxylase / phosphopantothenate--cysteine ligase | 2.30E-14 |
| | | | | | TIGR02113 | TIGR02113 | coaC_strep: phosphopantothenoylcysteine decarboxylase | 3.70E-12 |
| | | | | | PF02441 | Flavoprotein | Flavoprotein | 1.50E-11 |
| BAG23193.1 | 7572704 | 7572137 | - | 188 | LinL | LinL | LinL | 1.10E-51 |
| BAG23194.1 | 7574451 | 7572693 | - | 585 | LinH | LinH | LinH (Fusion of LinE/LinG) | 1.90E-175 |
| | | | | | LinG | LinG | LinG Homolog | 1.40E-76 |
| | | | | | LinE | LinE | LinE Homolog | 4.50E-65 |
| | | | | | PF12697 | Abhydrolase_6 | Alpha/beta hydrolase family | 7.30E-06 |
| | | | | | TIGR02427 | TIGR02427 | protocat_pcaD: 3-oxoadipate enol-lactonase | 3.30E-05 |
| BAG23195.1 | 7575284 | 7576934 | + | 549 | PF03704 | BTAD | Bacterial transcriptional activator domain | 4.00E-43 |
| | | | | | PF13191 | AAA_16 | AAA ATPase domain | 7.00E-12 |
| | | | | | TIGR03903 | TIGR03903 | TOMM_kin_cyc: TOMM system kinase/cyclase fusion protein | 1.10E-07 |
| | | | | | PF00486 | Trans_reg_C | Transcriptional regulatory protein, C terminal | 2.70E-04 |
| BAG23196.1 | 7577114 | 7577648 | + | 177 | PF01613 | Flavin_Reduct | Flavin reductase like domain | 8.90E-33 |
| | | | | | TIGR02296 | TIGR02296 | HpaC: 4-hydroxyphenylacetate 3-monooxygenase, reductase component | 8.10E-24 |
| | | | | | TIGR03615 | TIGR03615 | RutF: pyrimidine utilization flavin reductase protein F | 1.80E-23 |
| BAG23197.1 | 7577651 | 7578167 | + | 171 | NO PFAM MATCH | - | - | - |
| BAG23198.1 | 7578611 | 7579709 | + | 365 | PF03704 | BTAD | Bacterial transcriptional activator domain | 7.20E-45 |
| | | | | | PF13428 | TPR_14 | Tetratricopeptide repeat | 8.00E-06 |
| | | | | | PF00486 | Trans_reg_C | Transcriptional regulatory protein, C terminal | 3.30E-04 |
| | | | | | PF13432 | TPR_16 | Tetratricopeptide repeat | 6.20E-04 |
| | | | | | PF08279 | HTH_11 | HTH domain | 6.70E-04 |
| BAG23199.1 | 7586087 | 7579781 | - | 2101 | TIGR02813 | TIGR02813 | omega_3_PfaA: polyketide-type polyunsaturated fatty acid synthase PfaA | 1.30E-118 |
| | | | | | PF00109 | ketoacyl-synt | Beta-ketoacyl synthase, N-terminal domain | 3.10E-86 |
| | | | | | TIGR01746 | TIGR01746 | Thioester-redct: thioester reductase domain | 2.70E-85 |
| | | | | | PF08659 | KR | KR domain | 5.20E-64 |
| | | | | | TIGR03443 | TIGR03443 | alpha_am_amid: L-aminoadipate-semialdehyde dehydrogenase | 7.50E-58 |
| BAG23200.1 | 7596683 | 7586114 | - | 3522 | TIGR02813 | TIGR02813 | omega_3_PfaA: polyketide-type polyunsaturated fatty acid synthase PfaA | 1.60E-206 |
| | | | | | PF00109 | ketoacyl-synt | Beta-ketoacyl synthase, N-terminal domain | 2.00E-173 |
| | | | | | PF00698 | Acyl_transf_1 | Acyl transferase domain | 2.00E-109 |
| | | | | | TIGR00128 | TIGR00128 | fabD: malonyl CoA-acyl carrier protein transacylase | 7.60E-105 |
| | | | | | TIGR03150 | TIGR03150 | fabF: beta-ketoacyl-acyl-carrier-protein synthase II | 4.00E-86 |
| BAG23201.1 | 7611221 | 7596689 | - | 4843 | TIGR02813 | TIGR02813 | omega_3_PfaA: polyketide-type polyunsaturated fatty acid synthase PfaA | 0.00E+00 |
| | | | | | PF00109 | ketoacyl-synt | Beta-ketoacyl synthase, N-terminal domain | 5.20E-266 |
| | | | | | TIGR00128 | TIGR00128 | fabD: malonyl CoA-acyl carrier protein transacylase | 5.40E-156 |
| | | | | | PF00698 | Acyl_transf_1 | Acyl transferase domain | 2.50E-152 |
| | | | | | PF08659 | KR | KR domain | 6.50E-149 |

| index | leader | core | start | end | dir | score |
|---|---|---|---|---|---|---|
| 1 | MGNGRGRHPGRHRQMTERQLHSPGPDVAPGAQD | RTVVRTEGKTEAEGGERA | 7550645 | 7550801 | + | 7 |
| 2 | MATQDFANSVLAGAVPGFHSDAETPAM | ATPAVAQFVIQGSTICLVC | 7574708 | 7574567 | - | 32 |
| 3 | VLWPSPALQSNRVAFRCPSSEPMLSERDLAVP | SCPSPAASSELNALCTTSATRSAGIPQSW | 7574719 | 7574905 | + | 7 |
| 4 | LQSNRVAFRCPSSEPMLSERDLAVP | SCPSPAASSELNALCTTSATRSAGIPQSW | 7574740 | 7574905 | + | 7 |
| 5 | MRLDSIATQETATALPESMATQDFANSVLAGAVPGFHSDAETPAM | ATPAVAQFVIQGSTICLVC | 7574762 | 7574567 | - | 34 |
| 6 | LLGGLGTCEASGAGTPTDTGRTGGTTGRSS | KPASSRNRPTGAWQLVQIQRDLRLVKLILM | 7575144 | 7574961 | - | 8 |
| 7 | VQGLGPSPHKTRTRSGPCSGASAHARR | PAPGHRPTREGPGERPGVRRNRHRHEIARQARGNLSKFSGT | 7575194 | 7574987 | - | 6 |
| 8 | VAAYDPGFRRSGLRIPAHRAGARSKPTQNSDKERALLGGLGTCEASG | AGTPTDTGRTGGTTGRSSKPASSRNRPTGAWQLVQIQRDLRLVKLILM | 7575249 | 7574961 | - | 7 |
| 9 | VRCRPRRRGGRSPAYGFRGRTRTVYGAVAR | RAPVWVTGYPGMRVREIPRSMFSV | 7576930 | 7577095 | + | 6 |

### Results for SFS58200.1 [Streptomyces harbinensis] back to top

previous - next

#### Architecture

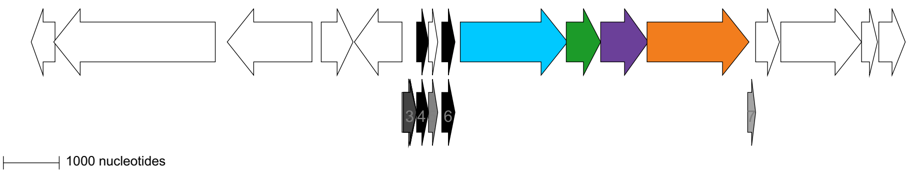

Link to nucleotide sequence

| Accession | start | end | direction | length (aa) | Pfam/HMM | name | description | E-value |
|---|---|---|---|---|---|---|---|---|
| SFS58172.1 | 660035 | 659609 | - | 141 | TIGR00941<br>PF00420 | TIGR00941<br>Oxidored_q2 | 2a6301s03: multicomponent Na+:H+ antiporter, MnhC subunit<br>NADH-ubiquinone/plastoquinone oxidoreductase chain 4L | 3.20E-24<br>2.60E-23 |
| SFS58176.1 | 662911 | 660031 | - | 959 | TIGR00940<br>TIGR01974<br>PF00361<br>TIGR01972<br>TIGR01770 | TIGR00940<br>TIGR01974<br>Proton_antipo_M<br>TIGR01972<br>TIGR01770 | 2a6301s01: monovalent cation:proton antiporter<br>NDH_I_L: proton-translocating NADH-quinone oxidoreductase, chain L<br>Proton-conducting membrane transporter<br>NDH_I_M: proton-translocating NADH-quinone oxidoreductase, chain M<br>NDH_I_N: proton-translocating NADH-quinone oxidoreductase, chain N | 1.30E-142<br>1.40E-73<br>3.40E-56<br>6.10E-41<br>7.50E-33 |
| SFS58179.1 | 664652 | 663131 | - | 506 | PF11066<br>PF13460<br>TIGR03466<br>PF05368<br>PF01370 | DUF2867<br>NAD_binding_10<br>TIGR03466<br>NmrA<br>Epimerase | Protein of unknown function (DUF2867)<br>NAD(P)H-binding<br>HpnA: hopanoid-associated sugar epimerase<br>NmrA-like family<br>NAD dependent epimerase/dehydratase family | 4.50E-42<br>5.10E-21<br>1.40E-15<br>8.80E-13<br>2.00E-09 |
| SFS58183.1 | 664816 | 665380 | + | 187 | PF01872 | RibD_C | RibD C-terminal domain | 4.50E-13 |
| SFS58186.1 | 666272 | 665399 | - | 290 | PF12146<br>PF12697<br>TIGR01738<br>TIGR03695<br>PF08386 | Hydrolase_4<br>Abhydrolase_6<br>TIGR01738<br>TIGR03695<br>Abhydrolase_4 | Serine aminopeptidase, S33<br>Alpha/beta hydrolase family<br>bioH: pimelyl-[acyl-carrier protein] methyl ester esterase<br>menH_SHCHC: 2-succinyl-6-hydroxy-2,4-cyclohexadiene-1-carboxylate synthase<br>TAP-like protein | 4.50E-10<br>1.20E-09<br>1.20E-09<br>6.70E-07<br>2.20E-05 |
| SFS58189.1 | 666529 | 666733 | + | 67 | LinA | LinA | LinA (linaridin precursor) | 1.70E-13 |
| SFS58193.1 | 666729 | 666906 | + | 58 | NO PFAM MATCH | - | - | - |
| SFS58196.1 | 666983 | 667202 | + | 72 | LinA | LinA | LinA (linaridin precursor) | 2.20E-13 |
| SFS58200.1 | 667307 | 669218 | + | 636 | LinH<br>LinG<br>LinE<br>PF12697<br>TIGR02427 | LinH<br>LinG<br>LinE<br>Abhydrolase_6<br>TIGR02427 | LinH (Fusion of LinE/LinG)<br>LinG Homolog<br>LinE Homolog<br>Alpha/beta hydrolase family<br>protocat_pcaD: 3-oxoadipate enol-lactonase | 3.00E-245<br>2.00E-112<br>1.70E-97<br>6.10E-16<br>1.60E-09 |
| SFS58203.1 | 669214 | 669814 | + | 199 | LinL | LinL | LinL | 1.20E-63 |
| SFS58205.1 | 669810 | 670665 | + | 284 | PF13649<br>PF08241<br>PF13489<br>PF08242<br>TIGR01983 | Methyltransf_25<br>Methyltransf_11<br>Methyltransf_23<br>Methyltransf_12<br>TIGR01983 | Methyltransferase domain<br>Methyltransferase domain<br>Methyltransferase domain<br>Methyltransferase domain<br>UbiG: 3-demethylubiquinone-9 3-O-methyltransferase | 1.20E-16<br>2.80E-11<br>8.00E-08<br>1.80E-07<br>1.40E-06 |
| SFS58209.1 | 670642 | 672469 | + | 608 | TIGR02204<br>TIGR02203<br>TIGR00958<br>TIGR03796<br>TIGR03375 | TIGR02204<br>TIGR02203<br>TIGR00958<br>TIGR03796<br>TIGR03375 | MsbA_rel: ABC transporter, permease/ATP-binding protein<br>MsbA_lipidA: lipid A export permease/ATP-binding protein MsbA<br>3a01208: antigen peptide transporter 2<br>NHLM_micro_ABC1: NHLM bacteriocin system ABC transporter, peptidase/ATP-binding protein<br>type_I_sec_LssB: type I secretion system ATPase | 8.90E-104<br>1.60E-99<br>2.00E-88<br>1.40E-86<br>5.20E-82 |
| SFS58218.1 | 672608 | 673022 | + | 137 | NO PFAM MATCH | - | - | - |
| SFS58229.1 | 673042 | 674503 | + | 486 | TIGR03467<br>TIGR01317<br>PF13450<br>TIGR02734<br>TIGR02733 | TIGR03467<br>TIGR01317<br>NAD_binding_8<br>TIGR02734<br>TIGR02733 | HpnE: squalene-associated FAD-dependent desaturase<br>GOGAT_sm_gam: glutamate synthase, NADH/NADPH, small subunit<br>NAD(P)-binding Rossmann-like domain<br>crtI_fam: phytoene desaturase<br>desat_CrtD: C-3',4' desaturase CrtD | 4.30E-10<br>1.10E-09<br>1.50E-09<br>2.00E-09<br>5.70E-09 |
| SFS58241.1 | 674499 | 674772 | + | 90 | NO PFAM MATCH | - | - | - |
| SFS58253.1 | 674814 | 675267 | + | 150 | PF13673<br>PF00583<br>PF13508<br>TIGR01575 | Acetyltransf_10<br>Acetyltransf_1<br>Acetyltransf_7<br>TIGR01575 | Acetyltransferase (GNAT) domain<br>Acetyltransferase (GNAT) family<br>Acetyltransferase (GNAT) domain<br>rimI: ribosomal-protein-alanine acetyltransferase | 2.40E-12<br>3.10E-08<br>1.70E-07<br>3.10E-06 |

| index | leader | core | start | end | dir | score |
|---|---|---|---|---|---|---|
| 1 | VLRRSDAWMPSPVSPWCPGRDSSTTVRSVFCPGR | TPPSHRRAARAPPERRPAVHDAAER...ATAAPSPPPAGSGGFCPAAERPPPG | 666250 | 666517 | + | 6 |
| 2 | MPSPVSPWCPGRDSSTTVRSVFCPGR | TPPSHRRAARAPPERRPAVHDAAER...ATAAPSPPPAGSGGFCPAAERPPPG | 666274 | 666517 | + | 6 |
| 3 | VSPWCPGRDSSTTVRSVFCPGR | TPPSHRRAARAPPERRPAVHDAAER...ATAAPSPPPAGSGGFCPAAERPPPG | 666286 | 666517 | + | 6 |
| 4 | MARSTPFAQRAVSGPLLEDSPGDIGTAASPVL | ATPAAAAAFAAGATIVVGAYVVGRMVGGETEPVIQ | 666529 | 666733 | + | 31 |
| 5 | MTTADRLPAALRHLPELTE | DAPGLASGTSEAVLCLVHSAVQVSVCVTSAGDQDSKLPL | 666729 | 666906 | + | 7 |
| 6 | MSRTTAYLNGCAALADLSVLDSAGRRSPAQPVL | ATPVTIVAFAGGVGAGAAIVAAFEAGRQAAGGSMEPIIT | 666983 | 667202 | + | 29 |
| 7 | MLRRGCSELPAHPLSPGRRSWQIRHVAPTG | TPPITRITHSDTAAEAGRG | 672446 | 672596 | + | 6 |

### Results for RKN35811.1 [Streptomyces hoynatensis] back to top

previous - next

#### Architecture

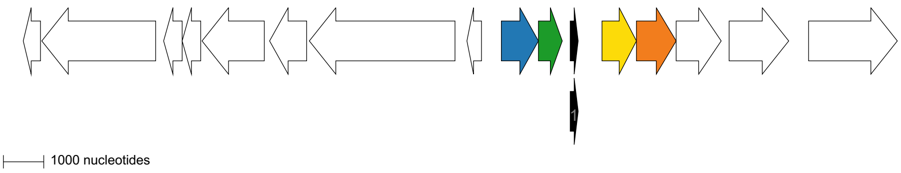

Link to nucleotide sequence

| Accession | start | end | direction | length (aa) | Pfam/HMM | name | description | E-value |
|---|---|---|---|---|---|---|---|---|
| RKN35803.1 | 4012 | 3580 | - | 143 | PF06841<br>TIGR02241 | Phage_T4_gp19<br>TIGR02241 | T4-like virus tail tube protein gp19<br>TIGR02241: conserved hypothetical phage tail region protein | 2.20E-27<br>3.60E-25 |
| RKN35804.1 | 6881 | 4049 | - | 943 | NO PFAM MATCH | - | - | - |
| RKN35805.1 | 7526 | 7058 | - | 155 | NO PFAM MATCH | - | - | - |
| RKN35806.1 | 7972 | 7525 | - | 148 | TIGR02241<br>PF06841 | TIGR02241<br>Phage_T4_gp19 | TIGR02241: conserved hypothetical phage tail region protein<br>T4-like virus tail tube protein gp19 | 2.20E-32<br>7.50E-30 |
| RKN35807.1 | 9559 | 8014 | - | 514 | PF04984<br>PF17482 | Phage_sheath_1<br>Phage_sheath_1C | Phage tail sheath protein subtilisin-like domain<br>Phage tail sheath C-terminal domain | 2.20E-33<br>3.50E-19 |
| RKN35808.1 | 10603 | 9697 | - | 301 | NO PFAM MATCH | - | - | - |
| RKN35809.1 | 14291 | 10673 | - | 1205 | PF07591<br>PF13403<br>PF14890<br>TIGR01443 | PT-HINT<br>Hint_2<br>Intein_splicing<br>TIGR01443 | Pretoxin HINT domain<br>Hint domain<br>Intein splicing domain<br>intein_Cterm: intein C-terminal splicing region | 1.20E-06<br>8.60E-06<br>9.40E-05<br>2.40E-04 |
| RKN35810.1 | 14945 | 14582 | - | 120 | NO PFAM MATCH | - | - | - |
| RKN35811.1 | 15434 | 16358 | + | 307 | LinE<br>LinH<br>PF12697<br>TIGR02427<br>PF00561 | LinE<br>LinH<br>Abhydrolase_6<br>TIGR02427<br>Abhydrolase_1 | LinE Homolog<br>LinH (Fusion of LinE/LinG)<br>Alpha/beta hydrolase family<br>protocat_pcaD: 3-oxoadipate enol-lactonase<br>alpha/beta hydrolase fold | 8.40E-86<br>5.00E-45<br>4.90E-18<br>4.50E-09<br>2.80E-07 |
| RKN35812.1 | 16354 | 16933 | + | 192 | LinL | LinL | LinL | 2.30E-51 |
| RKN35813.1 | 17119 | 17326 | + | 68 | LinA | LinA | LinA (linaridin precursor) | 3.50E-11 |
| RKN35822.1 | 17927 | 18761 | + | 277 | LinG<br>LinH | LinG<br>LinH | LinG Homolog<br>LinH (Fusion of LinE/LinG) | 3.40E-105<br>5.00E-79 |
| RKN35814.1 | 18757 | 19744 | + | 328 | TIGR03522<br>TIGR01188<br>TIGR03864<br>TIGR04406<br>TIGR01288 | TIGR03522<br>TIGR01188<br>TIGR03864<br>TIGR04406<br>TIGR01288 | GldA_ABC_ATP: gliding motility-associated ABC transporter ATP-binding subunit GldA<br>drrA: daunorubicin resistance ABC transporter, ATP-binding protein<br>PQQ_ABC_ATP: ABC transporter, ATP-binding subunit, PQQ-dependent alcohol dehydrogenase system<br>LPS_export_lptB: LPS export ABC transporter ATP-binding protein<br>nodI: nodulation ABC transporter NodI | 2.50E-53<br>2.70E-51<br>3.40E-46<br>5.40E-41<br>7.20E-38 |
| RKN35815.1 | 19754 | 20882 | + | 375 | PF12698<br>PF01061 | ABC2_membrane_3<br>ABC2_membrane | ABC-2 family transporter protein<br>ABC-2 type transporter | 5.70E-18<br>1.80E-04 |
| RKN35816.1 | 21077 | 22556 | + | 492 | PF13450<br>TIGR03467<br>TIGR00292<br>PF01593<br>TIGR02734 | NAD_binding_8<br>TIGR03467<br>TIGR00292<br>Amino_oxidase<br>TIGR02734 | NAD(P)-binding Rossmann-like domain<br>HpnE: squalene-associated FAD-dependent desaturase<br>TIGR00292: thiazole biosynthesis enzyme<br>Flavin containing amine oxidoreductase<br>crtI_fam: phytoene desaturase | 3.90E-08<br>6.70E-08<br>1.40E-07<br>2.10E-07<br>3.80E-07 |
| RKN35817.1 | 23023 | 25225 | + | 733 | PF03704<br>PF00931<br>PF00486<br>PF13191 | BTAD<br>NB-ARC<br>Trans_reg_C<br>AAA_16 | Bacterial transcriptional activator domain<br>NB-ARC domain<br>Transcriptional regulatory protein, C terminal<br>AAA ATPase domain | 5.70E-33<br>2.10E-07<br>2.80E-06<br>5.40E-05 |

| index | leader | core | start | end | dir | score |
|---|---|---|---|---|---|---|
| 1 | MSVVAEFANASLADVTPGPLMGTEAGLSSDAAPVL | CTPAATIACPEGWAILVAVADAVWNVTQEITQG | 17119 | 17326 | + | 32 |

### Results for WP_157852675.1 [Streptomyces iakyrus] back to top

previous - next

#### Architecture

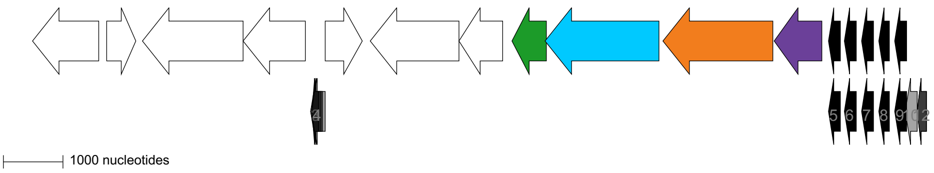

Link to nucleotide sequence

| Accession | start | end | direction | length (aa) | Pfam/HMM | name | description | E-value |
|---|---|---|---|---|---|---|---|---|
| WP_033306957.1 | 45870 | 44748 | - | 373 | TIGR03181<br>PF00676<br>TIGR03182<br>TIGR00204<br>PF13292 | TIGR03181<br>E1_dh<br>TIGR03182<br>TIGR00204<br>DXP_synthase_N | PDH_E1_alph_x: pyruvate dehydrogenase (acetyl-transferring) E1 component, alpha subunit<br>Dehydrogenase E1 component<br>PDH_E1_alph_y: pyruvate dehydrogenase (acetyl-transferring) E1 component, alpha subunit<br>dxs: 1-deoxy-D-xylulose-5-phosphate synthase<br>1-deoxy-D-xylulose-5-phosphate synthase | 3.90E-130<br>1.70E-70<br>6.20E-62<br>6.20E-09<br>3.20E-08 |
| WP_033306958.1 | 45999 | 46491 | + | 163 | PF01037<br>PF13412<br>PF13404<br>PF01978<br>PF12802 | AsnC_trans_reg<br>HTH_24<br>HTH_AsnC-type<br>TrmB<br>MarR_2 | Lrp/AsnC ligand binding domain<br>Winged helix-turn-helix DNA-binding<br>AsnC-type helix-turn-helix domain<br>Sugar-specific transcriptional regulator TrmB<br>MarR family | 7.20E-16<br>2.00E-13<br>1.70E-11<br>1.80E-04<br>3.30E-04 |

| | | | | | | | | |
|---|---|---|---|---|---|---|---|---|
| WP_051814460.1 | 48293 | 46589 | - | 567 | TIGR02288 | TIGR02288 | PaaN_2: phenylacetic acid degradation protein paaN | 9.50E-170 |
| | | | | | PF00171 | Aldedh | Aldehyde dehydrogenase family | 4.90E-19 |
| | | | | | TIGR01238 | TIGR01238 | D1pyr5carbox3: delta-1-pyrroline-5-carboxylate dehydrogenase | 1.60E-06 |
| | | | | | TIGR01237 | TIGR01237 | D1pyr5carbox2: putative delta-1-pyrroline-5-carboxylate dehydrogenase | 3.70E-05 |
| | | | | | TIGR04284 | TIGR04284 | aldehy_Rv0768: aldehyde dehydrogenase, Rv0768 family | 5.00E-05 |
| WP_033306959.1 | 49327 | 48280 | - | 348 | TIGR01141 | TIGR01141 | hisC: histidinol-phosphate transaminase | 4.60E-85 |
| | | | | | TIGR01140 | TIGR01140 | L_thr_O3P_dcar: threonine-phosphate decarboxylase | 2.80E-52 |
| | | | | | PF00155 | Aminotran_1_2 | Aminotransferase class I and II | 2.10E-43 |
| | | | | | TIGR03947 | TIGR03947 | viomycin_VioD: capreomycidine synthase | 1.40E-29 |
| | | | | | TIGR03540 | TIGR03540 | DapC_direct: LL-diaminopimelate aminotransferase | 1.50E-27 |
| WP_157852673.1 | 49655 | 50288 | + | 210 | NO PFAM MATCH | - | - | - |
| WP_033306961.1 | 51904 | 50410 | - | 497 | TIGR02734 | TIGR02734 | crtI_fam: phytoene desaturase | 5.80E-12 |
| | | | | | PF13450 | NAD_binding_8 | NAD(P)-binding Rossmann-like domain | 8.80E-10 |
| | | | | | TIGR02733 | TIGR02733 | desat_CrtD: C-3',4' desaturase CrtD | 1.20E-08 |
| | | | | | TIGR03467 | TIGR03467 | HpnE: squalene-associated FAD-dependent desaturase | 7.80E-08 |
| | | | | | TIGR00292 | TIGR00292 | TIGR00292: thiazole biosynthesis enzyme | 1.70E-07 |
| WP_157852674.1 | 52644 | 51900 | - | 247 | PF01551 | Peptidase_M23 | Peptidase family M23 | 7.30E-18 |
| WP_051814462.1 | 53369 | 52784 | - | 194 | LinL | LinL | LinL | 1.10E-65 |
| WP_157852675.1 | 55254 | 53349 | - | 634 | LinH | LinH | LinH (Fusion of LinE/LinG) | 6.60E-260 |
| | | | | | LinG | LinG | LinG Homolog | 5.70E-112 |
| | | | | | LinE | LinE | LinE Homolog | 4.30E-107 |
| | | | | | PF12697 | Abhydrolase_6 | Alpha/beta hydrolase family | 4.90E-14 |
| | | | | | PF00561 | Abhydrolase_1 | alpha/beta hydrolase fold | 1.80E-13 |
| WP_078625225.1 | 57183 | 55335 | - | 615 | TIGR02204 | TIGR02204 | MsbA_rel: ABC transporter, permease/ATP-binding protein | 1.70E-109 |
| | | | | | TIGR02203 | TIGR02203 | MsbA_lipidA: lipid A export permease/ATP-binding protein MsbA | 4.80E-99 |
| | | | | | TIGR00958 | TIGR00958 | 3a01208: antigen peptide transporter 2 | 6.10E-96 |
| | | | | | TIGR03796 | TIGR03796 | NHLM_micro_ABC1: NHLM bacteriocin system ABC transporter, peptidase/ATP-binding protein | 7.50E-89 |
| | | | | | TIGR01846 | TIGR01846 | type_I_sec_HlyB: type I secretion system ATPase | 4.10E-88 |
| WP_063739297.1 | 58003 | 57193 | - | 269 | PF13649 | Methyltransf_25 | Methyltransferase domain | 6.60E-17 |
| | | | | | PF08241 | Methyltransf_11 | Methyltransferase domain | 2.30E-13 |
| | | | | | PF13847 | Methyltransf_31 | Methyltransferase domain | 6.00E-11 |
| | | | | | PF08242 | Methyltransf_12 | Methyltransferase domain | 6.40E-09 |
| | | | | | TIGR01983 | TIGR01983 | UbiG: 3-demethylubiquinone-9 3-O-methyltransferase | 1.60E-08 |
| WP_033306962.1 | 58303 | 58117 | - | 61 | LinA | LinA | LinA (linaridin precursor) | 6.70E-15 |
| WP_033306963.1 | 58579 | 58381 | - | 65 | LinA | LinA | LinA (linaridin precursor) | 8.50E-16 |
| WP_033306964.1 | 58868 | 58655 | - | 70 | LinA | LinA | LinA (linaridin precursor) | 1.90E-17 |
| WP_033306965.1 | 59134 | 58945 | - | 62 | LinA | LinA | LinA (linaridin precursor) | 9.00E-13 |
| WP_033306966.1 | 59419 | 59215 | - | 67 | LinA | LinA | LinA (linaridin precursor) | 1.00E-16 |

| index | leader | core | start | end | dir | score |
|---|---|---|---|---|---|---|
| 1 | MWPTCDQHTLAGPGRPSCGSG | SDERSEGLLAGTDVADRQFVA | 49559 | 49430 | - | 6 |
| 2 | LRSTAAAGGPERGMWPTCDQHTLAGPG | RPSCGSGSDERSEGLLAGTDVADRQFVA | 49598 | 49430 | - | 8 |
| 3 | MAASRSEGSLCGARRRPGDRKGACGRLVINT | RLPGRAVRLAGAVLTSDRRDYLPVRMWLIGSLSPDFPIG | 49627 | 49414 | - | 6 |
| 4 | MVSLDRRGRGMAASRSEGSLCGARRRPGDRKGACGRLVINT | RLPGRAVRLAGAVLTSDRRDYLPVRMWLIGSLSPDFPIG | 49657 | 49414 | - | 6 |
| 5 | MSVLTERVNASSVLAELPLRSDRAQQMDELAPVL | ATPAIACMGFASVSGPSLAVAGSKQPS | 58303 | 58117 | - | 19 |
| 6 | MSVLTERVNASRVLAELPLRADRAQQMDELAPVL | ATPSAALVAAAAAVAVALADIYTVAGSKQPN | 58579 | 58381 | - | 30 |
| 7 | MSVLTERVNASSVLAELPLRADRAQQMDELAPVL | ATPATAVAFAAGVGVGVAVTNAFGAGLAAGGSKQPS | 58868 | 58655 | - | 30 |
| 8 | MLTEHVNASSVLAEFPLRADRARRMDELTPVL | ATPGAVVVAVGLYVVATPYVAGAIGSKQPS | 59134 | 58945 | - | 29 |
| 9 | MTSISDHVNGRSPMLFGDLRTTHGARHLSELTPVM | ATPAAFAAGVGVSVAAYAVGYAVGALGSKQPS | 59419 | 59215 | - | 30 |
| 10 | VRLTHMPARGSRTAAAFGVPEPAARPCPSAWPAVTAHAHEESALGTVPAD | FIPTQERNLHDDQYL | 59600 | 59402 | - | 6 |
| 11 | VEADVEATTLVLPGHVHGLGYSSRAFC | PTVAFLEAYLGPIPGTCTAP | 59748 | 59604 | - | 6 |
| 12 | MNAALVEADVEATTLVLPGHVHGLGYSSRAFC | PTVAFLEAYLGPIPGTCTAP | 59763 | 59604 | - | 8 |

### Results for TQE28105.1 [Streptomyces ipomoeae] back to top

previous - next

#### Architecture

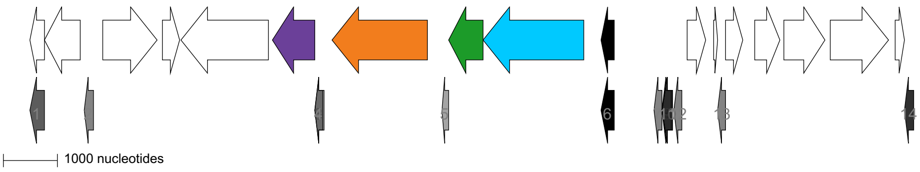

Link to nucleotide sequence

| Accession | start | end | direction | length (aa) | Pfam/HMM | name | description | E-value |
|---|---|---|---|---|---|---|---|---|

| | | | | | | | | |
|---|---|---|---|---|---|---|---|---|
| TQE28064.1 | 53247 | 52962 | - | 94 | PF05402<br>TIGR03859<br>TIGR04353<br>TIGR04248 | PqqD<br>TIGR03859<br>TIGR04353<br>TIGR04248 | Coenzyme PQQ synthesis protein D (PqqD)<br>PQQ_PqqD: coenzyme PQQ biosynthesis protein PqqD<br>PqqD_rel_X: PqqD family protein, HPr-rel-A system<br>SCM_PqqD_rel: SynChlorMet cassette protein ScmD | 2.90E-14<br>6.10E-06<br>3.00E-05<br>1.50E-04 |
| TQE28065.1 | 53909 | 53243 | - | 221 | NO PFAM MATCH | - | - | - |
| TQE28066.1 | 54318 | 55323 | + | 334 | TIGR01481<br>PF13377<br>TIGR02417<br>TIGR02405<br>PF00532 | TIGR01481<br>Peripla_BP_3<br>TIGR02417<br>TIGR02405<br>Peripla_BP_1 | ccpA: catabolite control protein A<br>Periplasmic binding protein-like domain<br>fruct_sucro_rep: D-fructose-responsive transcription factor<br>trehalos_R_Ecol: trehalose operon repressor<br>Periplasmic binding proteins and sugar binding domain of LacI family | 2.90E-49<br>2.90E-36<br>1.90E-29<br>1.30E-20<br>3.90E-19 |
| TQE28067.1 | 55426 | 55738 | + | 103 | NO PFAM MATCH | - | - | - |
| TQE28068.1 | 57372 | 55752 | - | 539 | PF09339<br>TIGR02431<br>PF12802<br>PF13412<br>PF12840 | HTH_IclR<br>TIGR02431<br>MarR_2<br>HTH_24<br>HTH_20 | IclR helix-turn-helix domain<br>pcaR_pcaU: beta-ketoadipate pathway transcriptional regulators, PcaR/PcaU/PobR family<br>MarR family<br>Winged helix-turn-helix DNA-binding<br>Helix-turn-helix domain | 6.80E-28<br>5.20E-19<br>4.80E-11<br>5.40E-07<br>1.00E-04 |
| TQE28069.1 | 58248 | 57444 | - | 267 | PF13649<br>PF08241<br>PF13489<br>TIGR01983<br>PF13847 | Methyltransf_25<br>Methyltransf_11<br>Methyltransf_23<br>TIGR01983<br>Methyltransf_31 | Methyltransferase domain<br>Methyltransferase domain<br>Methyltransferase domain<br>UbiG: 3-demethylubiquinone-9 3-O-methyltransferase<br>Methyltransferase domain | 1.70E-14<br>4.30E-12<br>2.40E-09<br>2.40E-07<br>1.50E-06 |
| TQE28104.1 | 60339 | 58569 | - | 589 | TIGR02203<br>TIGR02204<br>TIGR00958<br>TIGR03796<br>TIGR01846 | TIGR02203<br>TIGR02204<br>TIGR00958<br>TIGR03796<br>TIGR01846 | MsbA_lipidA: lipid A export permease/ATP-binding protein MsbA<br>MsbA_rel: ABC transporter, permease/ATP-binding protein<br>3a01208: antigen peptide transporter 2<br>NHLM_micro_ABC1: NHLM bacteriocin system ABC transporter, peptidase/ATP-binding protein<br>type_I_sec_HlyB: type I secretion system ATPase | 1.30E-104<br>1.20E-103<br>2.60E-92<br>1.40E-87<br>2.40E-78 |
| TQE28070.1 | 61362 | 60726 | - | 211 | LinL | LinL | LinL | 4.80E-76 |
| TQE28105.1 | 63227 | 61358 | - | 622 | LinH<br>LinG<br>LinE<br>PF12697<br>PF12146 | LinH<br>LinG<br>LinE<br>Abhydrolase_6<br>Hydrolase_4 | LinH (Fusion of LinE/LinG)<br>LinG Homolog<br>LinE Homolog<br>Alpha/beta hydrolase family<br>Serine aminopeptidase, S33 | 4.30E-234<br>5.10E-121<br>2.70E-88<br>9.10E-16<br>1.90E-10 |
| TQE28106.1 | 63796 | 63538 | - | 85 | LinA | LinA | LinA (linaridin precursor) | 8.60E-04 |
| TQE28071.1 | 65138 | 65480 | + | 113 | PF00196<br>TIGR03541<br>TIGR03020<br>PF08281<br>TIGR02937 | GerE<br>TIGR03541<br>TIGR03020<br>Sigma70_r4_2<br>TIGR02937 | Bacterial regulatory proteins, luxR family<br>reg_near_HchA: LuxR family transcriptional regulatory, chaperone HchA-associated<br>EpsA: transcriptional regulator EpsA<br>Sigma-70, region 4<br>sigma70-ECF: RNA polymerase sigma factor, sigma-70 family | 6.00E-19<br>2.20E-12<br>4.10E-11<br>7.70E-10<br>1.40E-06 |
| WP_009299312.1 | 65618 | 65696 | + | 26 | INFERRED GENE | - | - | - |
| TQE28072.1 | 65855 | 66164 | + | 102 | NO PFAM MATCH | - | - | - |
| TQE28073.1 | 66388 | 66862 | + | 157 | PF03779 | SPW | SPW repeat | 4.90E-21 |
| TQE28074.1 | 66917 | 67679 | + | 253 | PF09557<br>TIGR02271<br>PF05239 | DUF2382<br>TIGR02271<br>PRC | Domain of unknown function (DUF2382)<br>TIGR02271: conserved domain<br>PRC-barrel domain | 1.80E-28<br>9.80E-10<br>4.00E-07 |
| TQE28075.1 | 67784 | 68876 | + | 363 | PF03631<br>TIGR00765<br>TIGR00766 | Virul_fac_BrkB<br>TIGR00765<br>TIGR00766 | Virulence factor BrkB<br>yihY_not_rbn: YihY family inner membrane protein<br>TIGR00766: inner membrane protein YhjD | 2.40E-65<br>1.20E-36<br>1.20E-10 |
| WP_020117048.1 | 68987 | 69158 | + | 57 | INFERRED GENE | - | - | - |

| index | leader | core | start | end | dir | score |
|---|---|---|---|---|---|---|
| 1 | MSETKKPDTGTTRLQRYVSVSDVEDGMVLLDERKGRCFKLNSSGALI | LRALLDGLSAADVARTLVERYEVSQERAAADVAALTVELSGKGLVVV | 53247 | 52962 | - | 8 |
| 2 | MKQLLKLEYPLDTRLLDPAE | ITPCNHHFRHRNMYPEFRRSGRKPITGPLGHCA | 54139 | 53977 | - | 7 |
| 3 | MLCPFVVALGIVGFLIATQAVS | PPAVIGAALLLCGGVTVQLRKKR | 58391 | 58253 | - | 6 |
| 4 | MSDFMLCPFVVALGIVGFLIATQA | VSPPAVIGAALLLCGGVTVQLRKKR | 58403 | 58253 | - | 6 |
| 5 | MSLTNPFFLSGLGLVLIGLI | RAGALYVQVVRTRKQEGKDAE | 60724 | 60598 | - | 6 |
| 6 | MELPVEGIILTSARPTL | GTPRIGRVANANNALNALGGLADLA...NNAAAEANQAVANAANAQAAAAGGV | 63796 | 63538 | - | 28 |
| 7 | MKIGPSLISDPEASC | VPPAASVTCRHLPRQEGEPDADSPDLTATARGRNTRR | 64674 | 64515 | - | 7 |
| 8 | VEPRPWRRPESAGTAPDGGHLPCRHGMR | ATLTESPSRVAGPPMLAQFIGGAQERPA | 64844 | 64673 | - | 6 |
| 9 | MRAFLSVEPRPWRRPESAGTAPDGGHLPCRH | GMRATLTESPSRVAGPPMLAQFIGGAQERPA | 64862 | 64673 | - | 6 |
| 10 | MAMRAFLSVEPRPWRRPESAGTAPDGGHLPCR | HGMRATLTESPSRVAGPPMLAQFIGGAQERPA | 64868 | 64673 | - | 6 |
| 11 | MAMAMRAFLSVEPRPWRRPESAGTAPDGGHLPC | RHGMRATLTESPSRVAGPPMLAQFIGGAQERPA | 64874 | 64673 | - | 6 |
| 12 | VSCVRRGRFGTGSPASGVPTP | GTADRVVRRTSVDSLRVADDAEWAE | 65033 | 64892 | - | 7 |
| 13 | MAPSSYSPPHHPHRSLCAGEPTPIS | VPPHPPSTNHHPTSKTSQIAIKVV | 65857 | 65707 | - | 7 |
| 14 | LAAPETCFRSSAHFSPRSARGTGGAFG | SRPRHVRGGAVRRLFPDAAVLVVIATAR | 69330 | 69162 | - | 9 |

### Results for WP_100243889.1 [Streptomyces ipomoeae] back to top

previous - next

#### Architecture

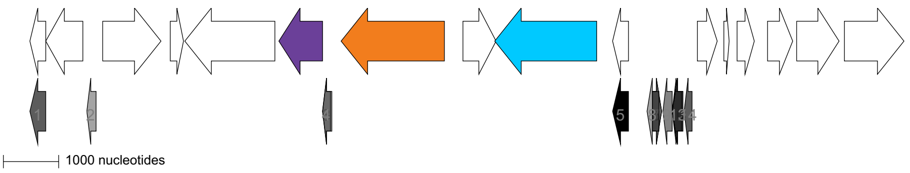

Link to nucleotide sequence

| Accession | start | end | direction | length (aa) | Pfam/HMM | name | description | E-value |
|---|---|---|---|---|---|---|---|---|
| WP_009308634.1 | 61460 | 61175 | - | 94 | PF05402<br>TIGR03859<br>TIGR04353<br>TIGR04248 | PqqD<br>TIGR03859<br>TIGR04353<br>TIGR04248 | Coenzyme PQQ synthesis protein D (PqqD)<br>PQQ_PqqD: coenzyme PQQ biosynthesis protein PqqD<br>PqqD_rel_X: PqqD family protein, HPr-rel-A system<br>SCM_PqqD_rel: SynChlorMet cassette protein ScmD | 2.90E-14<br>6.10E-06<br>3.00E-05<br>1.50E-04 |
| WP_141595397.1 | 62122 | 61456 | - | 221 | NO PFAM MATCH | - | - | - |
| WP_100243897.1 | 62483 | 63536 | + | 350 | TIGR01481<br>PF13377<br>TIGR02417<br>TIGR02405<br>PF00532 | TIGR01481<br>Peripla_BP_3<br>TIGR02417<br>TIGR02405<br>Peripla_BP_1 | ccpA: catabolite control protein A<br>Periplasmic binding protein-like domain<br>fruct_sucro_rep: D-fructose-responsive transcription factor<br>trehalos_R_Ecol: trehalose operon repressor<br>Periplasmic binding proteins and sugar binding domain of LacI family | 3.80E-49<br>3.20E-35<br>4.20E-29<br>1.60E-20<br>6.20E-19 |
| WP_048820115.1 | 63696 | 63951 | + | 84 | NO PFAM MATCH | - | - | - |
| WP_141595398.1 | 65585 | 63965 | - | 539 | PF09339<br>TIGR02431<br>PF12802<br>PF13412<br>PF12840 | HTH_IclR<br>TIGR02431<br>MarR_2<br>HTH_24<br>HTH_20 | IclR helix-turn-helix domain<br>pcaR_pcaU: beta-ketoadipate pathway transcriptional regulators, PcaR/PcaU/PobR family<br>MarR family<br>Winged helix-turn-helix DNA-binding<br>Helix-turn-helix domain | 6.90E-28<br>5.30E-19<br>4.90E-11<br>5.40E-07<br>1.10E-04 |
| WP_009307468.1 | 66461 | 65657 | - | 267 | PF13649<br>PF08241<br>PF13489<br>TIGR01983<br>PF13847 | Methyltransf_25<br>Methyltransf_11<br>Methyltransf_23<br>TIGR01983<br>Methyltransf_31 | Methyltransferase domain<br>Methyltransferase domain<br>Methyltransferase domain<br>UbiG: 3-demethylubiquinone-9 3-O-methyltransferase<br>Methyltransferase domain | 1.70E-14<br>4.30E-12<br>2.40E-09<br>2.40E-07<br>1.50E-06 |
| WP_009307479.1 | 68666 | 66782 | - | 627 | TIGR02204<br>TIGR02203<br>TIGR00958<br>TIGR03796<br>TIGR01846 | TIGR02204<br>TIGR02203<br>TIGR00958<br>TIGR03796<br>TIGR01846 | MsbA_rel: ABC transporter, permease/ATP-binding protein<br>MsbA_lipidA: lipid A export permease/ATP-binding protein MsbA<br>3a01208: antigen peptide transporter 2<br>NHLM_micro_ABC1: NHLM bacteriocin system ABC transporter, peptidase/ATP-binding protein<br>type_I_sec_HlyB: type I secretion system ATPase | 7.80E-108<br>9.50E-107<br>4.10E-93<br>3.90E-87<br>8.80E-80 |
| WP_005474565.1 | 68996 | 69584 | + | 196 | INFERRED GENE | - | - | - |
| WP_100243889.1 | 71410 | 69580 | - | 609 | LinH<br>LinG<br>LinE<br>PF12697<br>PF12146 | LinH<br>LinG<br>LinE<br>Abhydrolase_6<br>Hydrolase_4 | LinH (Fusion of LinE/LinG)<br>LinG Homolog<br>LinE Homolog<br>Alpha/beta hydrolase family<br>Serine aminopeptidase, S33 | 5.30E-231<br>2.30E-118<br>9.10E-89<br>1.00E-15<br>3.50E-11 |
| WP_048820058.1 | 71977 | 71698 | - | 92 | NO PFAM MATCH | - | - | - |
| WP_009307492.1 | 73241 | 73583 | + | 113 | PF00196<br>TIGR03541<br>TIGR03020<br>PF08281<br>TIGR02937 | GerE<br>TIGR03541<br>TIGR03020<br>Sigma70_r4_2<br>TIGR02937 | Bacterial regulatory proteins, luxR family<br>reg_near_HchA: LuxR family transcriptional regulatory, chaperone HchA-associated<br>EpsA: transcriptional regulator EpsA<br>Sigma-70, region 4<br>sigma70-ECF: RNA polymerase sigma factor, sigma-70 family | 5.80E-19<br>2.20E-12<br>4.00E-11<br>7.70E-10<br>1.40E-06 |
| WP_009299312.1 | 73721 | 73799 | + | 26 | INFERRED GENE | - | - | - |
| WP_009307484.1 | 73957 | 74266 | + | 102 | NO PFAM MATCH | - | - | - |
| WP_048820052.1 | 74490 | 74964 | + | 157 | PF03779 | SPW | SPW repeat | 4.90E-21 |
| WP_009307466.1 | 75019 | 75781 | + | 253 | PF09557<br>TIGR02271<br>PF05239 | DUF2382<br>TIGR02271<br>PRC | Domain of unknown function (DUF2382)<br>TIGR02271: conserved domain<br>PRC-barrel domain | 1.80E-28<br>9.80E-10<br>4.00E-07 |
| WP_009307496.1 | 75886 | 76975 | + | 362 | PF03631<br>TIGR00765<br>TIGR00766 | Virul_fac_BrkB<br>TIGR00765<br>TIGR00766 | Virulence factor BrkB<br>yihY_not_rbn: YihY family inner membrane protein<br>TIGR00766: inner membrane protein YhjD | 2.40E-65<br>1.20E-36<br>1.20E-10 |

| index | leader | core | start | end | dir | score |
|---|---|---|---|---|---|---|
| 1 | MSETKKPDTGTTRLQRYVSVSDVEDGMVLLDERKGRCFKLNSSGALI | LRALLDGLSAADVARTLVERYEVSQERAAADVAALTVELSGKGLVVV | 61460 | 61175 | - | 8 |
| 2 | MKQLLKLEYPLDTRLLEPAE | ITPCNHHFRHRNMYPEFRRSGRKPITGPLGHCA | 62352 | 62190 | - | 6 |
| 3 | MLCPFVVALGIVGFLIATQAVS | PPAVIGAALLLCGGVTVQLRKKR | 66604 | 66466 | - | 6 |
| 4 | MSDFMLCPFVVALGIVGFLIATQA | VSPPAVIGAALLLCGGVTVQLRKKR | 66616 | 66466 | - | 6 |
| 5 | MNLPVEGIILASARPTL | GTPRIGRVGRGAGGGGGRGLAALGA...ASVAADQAVAQAAADVAAGAGAPPA | 71977 | 71698 | - | 28 |
| 6 | VPCPVRGGCGAGSAGSGRSGGPGR | LPPDTWSAASSNTHAARCASLGVERAEAER | 72417 | 72582 | + | 6 |
| 7 | VRGGCGAGSAGSGRSGGPGR | LPPDTWSAASSNTHAARCASLGVERAEAER | 72429 | 72582 | + | 6 |
| 8 | VFDEAALHVSGGSRPGPPDRPLPAEPAP | HPPLTGHGTAAPRPDAATWLRSRNGPRPRSRNGTRP | 72527 | 72332 | - | 6 |
| 9 | MKIGPSLISDPGASC | VPPAASVTGRHLPRQEGEPDANSPDLTATARGRNTRR | 72777 | 72618 | - | 7 |
| 10 | VEPRPWRRPESAGTALDGRHLPCRHGMR | ATLTESPSRVAGPPMLAQFIGGAQERPA | 72947 | 72776 | - | 6 |
| 11 | MRAFLSVEPRPWRRPESAGTALDGRHLPCRH | GMRATLTESPSRVAGPPMLAQFIGGAQERPA | 72965 | 72776 | - | 6 |
| 12 | MAMRAFLSVEPRPWRRPESAGTALDGRHLPCR | HGMRATLTESPSRVAGPPMLAQFIGGAQERPA | 72971 | 72776 | - | 6 |
| 13 | MAMAMRAFLSVEPRPWRRPESAGTALDGRHLPC | RHGMRATLTESPSRVAGPPMLAQFIGGAQERPA | 72977 | 72776 | - | 6 |

| | | | | | | |
|---|---|---|---|---|---|---|
| 14 | VSCVRRGRFGTGSPAPGV | PTPGTADRVVRRTSVDSLRVADDAEWAE | 73136 | 72995 | - | 8 |

### Results for EKX67255.1 [Streptomyces ipomoeae 91-03] back to top

previous - next

#### Architecture

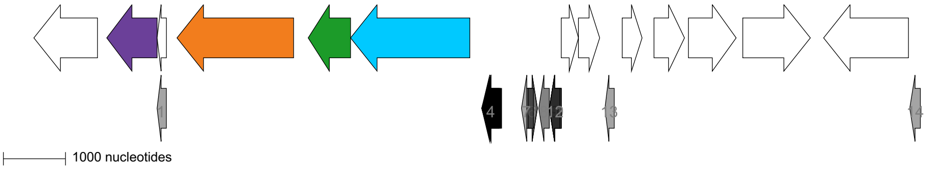

Link to nucleotide sequence

| Accession | start | end | direction | length (aa) | Pfam/HMM | name | description | E-value |
|---|---|---|---|---|---|---|---|---|
| EKX67249.1 | 1018 | 0 | - | 339 | PF09339<br>TIGR02431<br>PF12802<br>PF13412<br>PF12840 | HTH_IclR<br>TIGR02431<br>MarR_2<br>HTH_24<br>HTH_20 | IclR helix-turn-helix domain<br>pcaR_pcaU: beta-ketoadipate pathway transcriptional regulators, PcaR/PcaU/PobR family<br>MarR family<br>Winged helix-turn-helix DNA-binding<br>Helix-turn-helix domain | 4.70E-28<br>5.30E-18<br>7.30E-11<br>1.90E-07<br>3.90E-05 |
| EKX67235.1 | 1987 | 1183 | - | 267 | PF13649<br>PF08241<br>PF13489<br>TIGR01983<br>PF13847 | Methyltransf_25<br>Methyltransf_11<br>Methyltransf_23<br>TIGR01983<br>Methyltransf_31 | Methyltransferase domain<br>Methyltransferase domain<br>Methyltransferase domain<br>UbiG: 3-demethylubiquinone-9 3-O-methyltransferase<br>Methyltransferase domain | 1.70E-14<br>4.30E-12<br>2.40E-09<br>2.40E-07<br>1.50E-06 |
| EKX67252.1 | 2142 | 1992 | - | 49 | NO PFAM MATCH | - | - | - |
| EKX67242.1 | 4192 | 2308 | - | 627 | TIGR02204<br>TIGR02203<br>TIGR00958<br>TIGR03796<br>TIGR01846 | TIGR02204<br>TIGR02203<br>TIGR00958<br>TIGR03796<br>TIGR01846 | MsbA_rel: ABC transporter, permease/ATP-binding protein<br>MsbA_lipidA: lipid A export permease/ATP-binding protein MsbA<br>3a01208: antigen peptide transporter 2<br>NHLM_micro_ABC1: NHLM bacteriocin system ABC transporter, peptidase/ATP-binding protein<br>type_I_sec_HlyB: type I secretion system ATPase | 7.80E-108<br>9.50E-107<br>4.10E-93<br>3.90E-87<br>8.80E-80 |
| EKX67238.1 | 5110 | 4423 | - | 228 | LinL | LinL | LinL | 8.10E-77 |
| EKX67255.1 | 7032 | 5106 | - | 641 | LinH<br>LinG<br>LinE<br>PF12697<br>PF12146 | LinH<br>LinG<br>LinE<br>Abhydrolase_6<br>Hydrolase_4 | LinH (Fusion of LinE/LinG)<br>LinG Homolog<br>LinE Homolog<br>Alpha/beta hydrolase family<br>Serine aminopeptidase, S33 | 2.80E-236<br>2.20E-125<br>1.00E-88<br>1.20E-15<br>3.70E-11 |
| EKX67241.1 | 8489 | 8765 | + | 91 | NO PFAM MATCH | - | - | - |
| EKX67251.1 | 8767 | 9109 | + | 113 | PF00196<br>TIGR03541<br>TIGR03020<br>PF08281<br>TIGR02937 | GerE<br>TIGR03541<br>TIGR03020<br>Sigma70_r4_2<br>TIGR02937 | Bacterial regulatory proteins, luxR family<br>reg_near_HchA: LuxR family transcriptional regulatory, chaperone HchA-associated<br>EpsA: transcriptional regulator EpsA<br>Sigma-70, region 4<br>sigma70-ECF: RNA polymerase sigma factor, sigma-70 family | 5.80E-19<br>2.20E-12<br>4.00E-11<br>7.70E-10<br>1.40E-06 |
| EKX67245.1 | 9483 | 9792 | + | 102 | NO PFAM MATCH | - | - | - |
| EKX67257.1 | 9980 | 10490 | + | 169 | PF03779 | SPW | SPW repeat | 6.30E-21 |
| EKX67234.1 | 10545 | 11307 | + | 253 | PF09557<br>TIGR02271<br>PF05239 | DUF2382<br>TIGR02271<br>PRC | Domain of unknown function (DUF2382)<br>TIGR02271: conserved domain<br>PRC-barrel domain | 1.80E-28<br>9.80E-10<br>4.00E-07 |
| EKX67253.1 | 11412 | 12501 | + | 362 | PF03631<br>TIGR00765<br>TIGR00766 | Virul_fac_BrkB<br>TIGR00765<br>TIGR00766 | Virulence factor BrkB<br>yihY_not_rbn: YihY family inner membrane protein<br>TIGR00766: inner membrane protein YhjD | 2.40E-65<br>1.20E-36<br>1.20E-10 |
| EKX67258.1 | 14087 | 12728 | - | 452 | PF00781<br>TIGR00147<br>TIGR03702<br>PF01513<br>PF05567 | DAGK_cat<br>TIGR00147<br>TIGR03702<br>NAD_kinase<br>Neisseria_PilC | Diacylglycerol kinase catalytic domain<br>TIGR00147: lipid kinase, YegS/Rv2252/BmrU family<br>lip_kinase_YegS: lipid kinase YegS<br>ATP-NAD kinase<br>Neisseria PilC beta-propeller domain | 2.10E-20<br>1.10E-13<br>1.00E-08<br>2.60E-06<br>3.80E-06 |

| index | leader | core | start | end | dir | score |
|---|---|---|---|---|---|---|
| 1 | MSDFMLCPFVVALGIVGFLIATQA | VSPPAVIGAALLLCGGVTVQLRKKR | 2142 | 1992 | - | 6 |
| 2 | VEGIILASARPTL | GTPRIGRVGRGAGGGGGRGLAALGA...ASVAADQAVAQAAADVAAGAGAPPA | 7491 | 7224 | - | 18 |
| 3 | VDNLNLPVEGIILASARPTL | GTPRIGRVGRGAGGGGGRGLAALGA...ASVAADQAVAQAAADVAAGAGAPPA | 7512 | 7224 | - | 28 |
| 4 | MSHIAKAVDNLNLPVEGIILASARPTL | GTPRIGRVGRGAGGGGGRGLAALGA...ASVAADQAVAQAAADVAAGAGAPPA | 7533 | 7224 | - | 27 |
| 5 | VPCPVRGGCGAGSAGSGRSGGPGR | LPPDTWSAASSNTHAARCASLGVERAEAER | 7943 | 8108 | + | 6 |
| 6 | VRGGCGAGSAGSGRSGGPGR | LPPDTWSAASSNTHAARCASLGVERAEAER | 7955 | 8108 | + | 6 |
| 7 | VFDEAALHVSGGSRPGPPDRPLPAEPAP | HPPLTGHGTAAPRPDAATWLRSRNGPRPRSRNGTRP | 8053 | 7858 | - | 6 |
| 8 | MKIGPSLISDPGASC | VPPAASVTGRHLPRQEGEPDANSPDLTATARGRNTRR | 8303 | 8144 | - | 7 |
| 9 | VEPRPWRRPESAGTALDGRHLPCRHGMR | ATLTESPSRVAGPPMLAQFIGGAQERPA | 8473 | 8302 | - | 6 |
| 10 | MRAFLSVEPRPWRRPESAGTALDGRHLPCRH | GMRATLTESPSRVAGPPMLAQFIGGAQERPA | 8491 | 8302 | - | 6 |
| 11 | MAMRAFLSVEPRPWRRPESAGTALDGRHLPCR | HGMRATLTESPSRVAGPPMLAQFIGGAQERPA | 8497 | 8302 | - | 6 |
| 12 | MAMAMRAFLSVEPRPWRRPESAGTALDGRHLPC | RHGMRATLTESPSRVAGPPMLAQFIGGAQERPA | 8503 | 8302 | - | 6 |

| | | | | | | |
|---|---|---|---|---|---|---|
| 13 | LFKRQRRGRHRHRRGEGRSRNGR | AARPGQDRTVTGARKAPLTGWTR | 9344 | 9203 | - | 6 |
| 14 | VVGRDDHHLGLHHLGPVRGPAWQPLRPV | RGTSHVRGAPVGEDLVSARALLSTQRSGT | 14283 | 14109 | - | 6 |

### Results for WP_153524745.1 [Streptomyces jumonjinensis] back to top

previous - next

#### Architecture

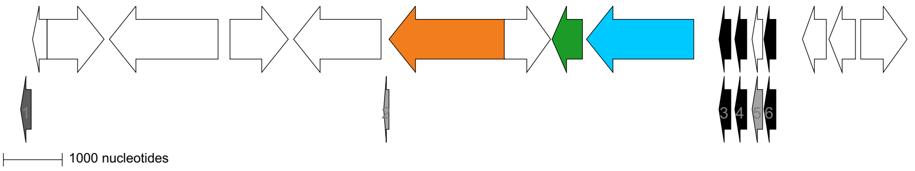

Link to nucleotide sequence

| Accession | start | end | direction | length (aa) | Pfam/HMM | name | description | E-value |
|---|---|---|---|---|---|---|---|---|
| WP_153483897.1 | 23024 | 22778 | - | 81 | PF10041 | DUF2277 | Uncharacterized conserved protein (DUF2277) | 1.80E-28 |
| WP_153524665.1 | 23022 | 24012 | + | 329 | PF08378 | NERD | Nuclease-related domain | 6.10E-15 |
| WP_153524666.1 | 25937 | 24098 | - | 612 | NO PFAM MATCH | - | - | - |
| WP_153524667.1 | 26133 | 27141 | + | 335 | NO PFAM MATCH | - | - | - |
| WP_153524668.1 | 28701 | 27213 | - | 495 | TIGR02734<br>TIGR03467<br>TIGR01317<br>TIGR00292<br>PF13450 | TIGR02734<br>TIGR03467<br>TIGR01317<br>TIGR00292<br>NAD_binding_8 | crtI_fam: phytoene desaturase<br>HpnE: squalene-associated FAD-dependent desaturase<br>GOGAT_sm_gam: glutamate synthase, NADH/NADPH, small subunit<br>TIGR00292: thiazole biosynthesis enzyme<br>NAD(P)-binding Rossmann-like domain | 7.20E-10<br>1.60E-08<br>3.10E-08<br>4.10E-08<br>7.30E-08 |
| WP_153524669.1 | 30815 | 28850 | - | 654 | TIGR02204<br>TIGR02203<br>TIGR00958<br>TIGR03796<br>TIGR01846 | TIGR02204<br>TIGR02203<br>TIGR00958<br>TIGR03796<br>TIGR01846 | MsbA_rel: ABC transporter, permease/ATP-binding protein<br>MsbA_lipidA: lipid A export permease/ATP-binding protein MsbA<br>3a01208: antigen peptide transporter 2<br>NHLM_micro_ABC1: NHLM bacteriocin system ABC transporter, peptidase/ATP-binding protein<br>type_I_sec_HlyB: type I secretion system ATPase | 1.50E-97<br>1.50E-91<br>1.10E-84<br>2.70E-84<br>7.60E-77 |
| WP_018487274.1 | 30811 | 31591 | + | 260 | INFERRED GENE | - | - | - |
| WP_153524744.1 | 32133 | 31602 | - | 176 | LinL | LinL | LinL | 1.70E-59 |
| WP_153524745.1 | 34025 | 32207 | - | 605 | LinH<br>LinG<br>LinE<br>PF12697<br>LinL | LinH<br>LinG<br>LinE<br>Abhydrolase_6<br>LinL | LinH (Fusion of LinE/LinG)<br>LinG Homolog<br>LinE Homolog<br>Alpha/beta hydrolase family<br>LinL | 8.90E-240<br>1.90E-114<br>3.20E-91<br>2.70E-15<br>5.20E-15 |
| WP_153524746.1 | 34646 | 34448 | - | 65 | LinA | LinA | LinA (linaridin precursor) | 2.10E-11 |
| WP_153524670.1 | 34929 | 34731 | - | 65 | LinA | LinA | LinA (linaridin precursor) | 8.50E-08 |
| WP_153524671.1 | 35196 | 35007 | - | 62 | NO PFAM MATCH | - | - | - |
| WP_153483909.1 | 35408 | 35219 | - | 62 | LinA | LinA | LinA (linaridin precursor) | 3.80E-09 |
| WP_153524672.1 | 36283 | 35860 | - | 140 | PF13581<br>TIGR01924 | HATPase_c_2<br>TIGR01924 | Histidine kinase-like ATPase domain<br>rsbW_low_gc: anti-sigma B factor | 6.10E-10<br>5.60E-04 |
| WP_153524673.1 | 36772 | 36328 | - | 147 | PF00293<br>TIGR00586<br>TIGR02705<br>PF14815 | NUDIX<br>TIGR00586<br>TIGR02705<br>NUDIX_4 | NUDIX domain<br>mutt: mutator mutT protein<br>nudix_YtkD: nucleoside triphosphatase YtkD<br>NUDIX domain | 2.00E-20<br>1.20E-09<br>8.00E-08<br>3.80E-06 |
| WP_153524674.1 | 36853 | 37642 | + | 262 | PF07702<br>TIGR02325<br>PF00392<br>TIGR03337<br>TIGR02018 | UTRA<br>TIGR02325<br>GntR<br>TIGR03337<br>TIGR02018 | UTRA domain<br>C_P_lyase_phnF: phosphonate metabolism transcriptional regulator PhnF<br>Bacterial regulatory proteins, gntR family<br>phnR: phosphonate utilization transcriptional regulator PhnR<br>his_ut_repres: histidine utilization repressor | 1.30E-27<br>3.70E-27<br>2.30E-23<br>2.40E-20<br>7.30E-19 |

| index | leader | core | start | end | dir | score |
|---|---|---|---|---|---|---|
| 1 | MDQETGAASGTGAAGPGA | GTAPARAAAAGRRITKVATAPAANSASTETAAPTQLAPSHRPPK | 22768 | 22579 | - | 8 |
| 2 | MTRALPDSDSDSDSDRAR | TRPARTPAGAAPAVNGWQ | 28854 | 28743 | - | 6 |
| 3 | MHRPGALTDIGIEDAKGRFVTAHAPVM | ATPAVAAGVIIGMALVNAFVAGRDGGPLEPSVPEIVLG | 34646 | 34448 | - | 29 |
| 4 | MSVSVCPVAPVTTCELSETDPGYAATAVAPVA | ATPVVVAVAAFGAGAAFGVAVTAAIGETQPVLE | 34929 | 34731 | - | 30 |
| 5 | MSVAGCPAAPVTVRELAETEPGFIAAAVAPIA | AAFVVVAEFGNGLGAGYASHVIAGSEERPL | 35196 | 35007 | - | 6 |
| 6 | MQQVRSRRWADAVGPLDEGAEGQVFDGVAPIA | ATQVVATVAAAVVVAYAAGAIIGESQPVLE | 35408 | 35219 | - | 18 |

### Results for WP_063778956.1 [Streptomyces katrae] back to top

previous - next

#### Architecture

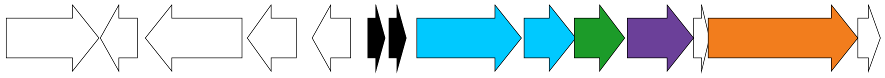

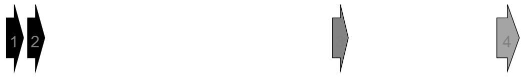

1000 nucleotides

Link to nucleotide sequence

| Accession | start | end | direction | length (aa) | Pfam/HMM | name | description | E-value |
|---|---|---|---|---|---|---|---|---|
| WP_045949462.1 | 1921 | 3022 | + | 366 | TIGR02050<br>PF04107<br>TIGR02048<br>TIGR03444 | TIGR02050<br>GCS2<br>TIGR02048<br>TIGR03444 | gshA_cyan_rel: carboxylate-amine ligase, YbdK family<br>Glutamate-cysteine ligase family 2(GCS2)<br>gshA_cyano: putative glutamate--cysteine ligase<br>EgtA_Cys_ligase: ergothioneine biosynthesis glutamate--cysteine ligase EgtA | 1.20E-79<br>9.00E-53<br>4.90E-11<br>1.90E-06 |
| WP_045949452.1 | 3470 | 3041 | - | 142 | PF13581 | HATPase_c_2 | Histidine kinase-like ATPase domain | 3.50E-07 |
| WP_045949453.1 | 4704 | 3561 | - | 380 | NO PFAM MATCH | - | - | - |
| WP_052713777.1 | 5347 | 4771 | - | 191 | PF06114 | Peptidase_M78 | IrrE N-terminal-like domain | 1.40E-04 |
| WP_045949454.1 | 5979 | 5535 | - | 147 | NO PFAM MATCH | - | - | - |
| WP_045949455.1 | 6195 | 6393 | + | 65 | LinA | LinA | LinA (linaridin precursor) | 1.40E-13 |
| WP_045949456.1 | 6433 | 6634 | + | 66 | LinA | LinA | LinA (linaridin precursor) | 7.70E-12 |
| WP_157880381.1 | 6764 | 7994 | + | 409 | LinH<br>LinG<br>LinE | LinH<br>LinG<br>LinE | LinH (Fusion of LinE/LinG)<br>LinG Homolog<br>LinE Homolog | 3.30E-163<br>1.10E-116<br>4.20E-27 |
| WP_063778956.1 | 8026 | 8635 | + | 202 | LinH<br>LinE | LinH<br>LinE | LinH (Fusion of LinE/LinG)<br>LinE Homolog | 5.00E-62<br>2.40E-50 |
| WP_045949457.1 | 8631 | 9219 | + | 195 | LinL | LinL | LinL | 2.10E-70 |
| WP_045949458.1 | 9247 | 10033 | + | 261 | PF13649<br>PF08241<br>TIGR01983<br>PF13489<br>TIGR02469 | Methyltransf_25<br>Methyltransf_11<br>TIGR01983<br>Methyltransf_23<br>TIGR02469 | Methyltransferase domain<br>Methyltransferase domain<br>UbiG: 3-demethylubiquinone-9 3-O-methyltransferase<br>Methyltransferase domain<br>CbiT: precorrin-6Y C5,15-methyltransferase (decarboxylating), CbiT subunit | 2.30E-10<br>4.80E-07<br>6.90E-06<br>2.30E-05<br>3.30E-05 |
| WP_157880382.1 | 10032 | 10209 | + | 58 | NO PFAM MATCH | - | - | - |
| WP_045949459.1 | 10205 | 11969 | + | 587 | TIGR02204<br>TIGR02203<br>TIGR00958<br>TIGR03796<br>TIGR01846 | TIGR02204<br>TIGR02203<br>TIGR00958<br>TIGR03796<br>TIGR01846 | MsbA_rel: ABC transporter, permease/ATP-binding protein<br>MsbA_lipidA: lipid A export permease/ATP-binding protein MsbA<br>3a01208: antigen peptide transporter 2<br>NHLM_micro_ABC1: NHLM bacteriocin system ABC transporter, peptidase/ATP-binding protein<br>type_I_sec_HlyB: type I secretion system ATPase | 2.60E-105<br>9.00E-101<br>2.70E-82<br>3.70E-82<br>5.00E-80 |
| WP_045949460.1 | 11965 | 12222 | + | 86 | PF13450<br>TIGR01317<br>TIGR03467<br>PF12831<br>TIGR01372 | NAD_binding_8<br>TIGR01317<br>TIGR03467<br>FAD_oxidored<br>TIGR01372 | NAD(P)-binding Rossmann-like domain<br>GOGAT_sm_gam: glutamate synthase, NADH/NADPH, small subunit<br>HpnE: squalene-associated FAD-dependent desaturase<br>FAD dependent oxidoreductase<br>soxA: sarcosine oxidase, alpha subunit family | 2.40E-11<br>1.00E-09<br>3.80E-09<br>7.10E-09<br>1.30E-08 |

| index | leader | core | start | end | dir | score |
|---|---|---|---|---|---|---|
| 1 | MAKIAAFMNEQEMGPITESVVNRDGTPVM | ATPAAVVAVPASVKITAAVVAFTVPVGAYVAGRVVN | 6195 | 6393 | + | 30 |
| 2 | MNRITESVRGIADAALSEDTGERTSRMMPVA | FTPAVLVVCPYAAGGLAAGAAAVGAYEAGRAAAKG | 6433 | 6634 | + | 29 |
| 3 | MNTTRTLWFLFSFAVAVPVM | AVMFREDGEFTSQSWTKGLIIGTAVAVVAAIAAGRARP | 10032 | 10209 | + | 7 |
| 4 | MSAGADAAVVGTGPNGLAAA | VTLARAGLRVVLYEQAGRIGGGLRT...HPMAAASPFFRAFDLPGRGVPLLQP | 11965 | 12222 | + | 6 |

### Results for WP_153483925.1 [Streptomyces katsurahamanus] back to top

previous - next

#### Architecture

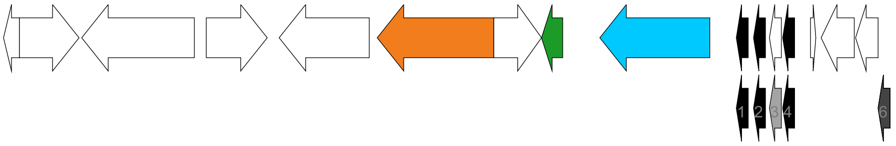

1000 nucleotides

Link to nucleotide sequence

| Accession | start | end | direction | length (aa) | Pfam/HMM | name | description | E-value |
|---|---|---|---|---|---|---|---|---|
| WP_153483897.1 | 299 | 53 | - | 81 | PF10041 | DUF2277 | Uncharacterized conserved protein (DUF2277) | 1.80E-28 |
| WP_153483898.1 | 297 | 1287 | + | 329 | PF08378 | NERD | Nuclease-related domain | 1.40E-14 |
| WP_153483899.1 | 3174 | 1335 | - | 612 | NO PFAM MATCH | - | - | - |
| WP_153483900.1 | 3371 | 4379 | + | 335 | NO PFAM MATCH | - | - | - |
| WP_153483901.1 | 6054 | 4584 | - | 489 | TIGR02734<br>TIGR03467<br>TIGR01317<br>TIGR00292<br>PF13450 | TIGR02734<br>TIGR03467<br>TIGR01317<br>TIGR00292<br>NAD_binding_8 | crtI_fam: phytoene desaturase<br>HpnE: squalene-associated FAD-dependent desaturase<br>GOGAT_sm_gam: glutamate synthase, NADH/NADPH, small subunit<br>TIGR00292: thiazole biosynthesis enzyme<br>NAD(P)-binding Rossmann-like domain | 1.30E-09<br>1.90E-08<br>3.10E-08<br>4.10E-08<br>7.10E-08 |

| | | | | | | | | |
|---|---|---|---|---|---|---|---|---|
| WP_153483903.1 | 8120 | 6191 | - | 642 | TIGR02204<br>TIGR02203<br>TIGR03796<br>TIGR00958<br>TIGR01846 | TIGR02204<br>TIGR02203<br>TIGR03796<br>TIGR00958<br>TIGR01846 | MsbA_rel: ABC transporter, permease/ATP-binding protein<br>MsbA_lipidA: lipid A export permease/ATP-binding protein MsbA<br>NHLM_micro_ABC1: NHLM bacteriocin system ABC transporter, peptidase/ATP-binding protein<br>3a01208: antigen peptide transporter 2<br>type_I_sec_HlyB: type I secretion system ATPase | 5.90E-99<br>1.60E-91<br>5.30E-87<br>2.60E-85<br>1.80E-77 |
| WP_018487274.1 | 8116 | 8896 | + | 260 | INFERRED GENE | - | - | - |
| WP_153483904.1 | 9240 | 8907 | - | 110 | LinL | LinL | LinL | 1.70E-31 |
| WP_153483925.1 | 11677 | 9847 | - | 609 | LinH<br>LinG<br>LinE<br>PF12697<br>PF00561 | LinH<br>LinG<br>LinE<br>Abhydrolase_6<br>Abhydrolase_1 | LinH (Fusion of LinE/LinG)<br>LinG Homolog<br>LinE Homolog<br>Alpha/beta hydrolase family<br>alpha/beta hydrolase fold | 3.70E-239<br>1.80E-114<br>3.90E-90<br>4.70E-15<br>8.00E-12 |
| WP_153483926.1 | 12299 | 12101 | - | 65 | LinA | LinA | LinA (linaridin precursor) | 3.60E-11 |
| WP_153483906.1 | 12579 | 12381 | - | 65 | LinA | LinA | LinA (linaridin precursor) | 1.50E-07 |
| WP_153483907.1 | 12846 | 12657 | - | 62 | NO PFAM MATCH | - | - | - |
| WP_153483909.1 | 13058 | 12869 | - | 62 | LinA | LinA | LinA (linaridin precursor) | 3.80E-09 |
| WP_003955481.1 | 13314 | 13419 | + | 35 | INFERRED GENE | - | - | - |
| WP_153483911.1 | 14054 | 13499 | - | 184 | PF13302<br>PF00583<br>TIGR03448<br>TIGR01575<br>TIGR03585 | Acetyltransf_3<br>Acetyltransf_1<br>TIGR03448<br>TIGR01575<br>TIGR03585 | Acetyltransferase (GNAT) domain<br>Acetyltransferase (GNAT) family<br>mycothiol_MshD: mycothiol synthase<br>rimI: ribosomal-protein-alanine acetyltransferase<br>PseH: pseudaminic acid biosynthesis N-acetyl transferase | 1.10E-14<br>7.10E-11<br>2.50E-07<br>6.50E-06<br>8.40E-06 |
| WP_153483912.1 | 14445 | 14061 | - | 127 | PF13471 | Transglut_core3 | Transglutaminase-like superfamily | 1.40E-15 |

| index | leader | core | start | end | dir | score |
|---|---|---|---|---|---|---|
| 1 | MRRPGALTDIGIEDAKGRFVTAHAPVM | ATPAVAAGVIIGVALVNAFVAGRDGGPLEPTVPEIGLG | 12299 | 12101 | - | 28 |
| 2 | MSVSVCPVAPVTTCALSETDPGYAATAVAPVA | ATPVVVAVAAFGAGAAFGVAVTAAIGETQPVLE | 12579 | 12381 | - | 30 |
| 3 | MSVAGCPAAPVTVRELAETEPGFIAAAVAPIA | AAFVVVAEFGHGLGAGYASHVIAGSEERPL | 12846 | 12657 | - | 6 |
| 4 | MQQVRSRRWADAVGPLDEGAEGQVFDGVAPIA | ATQVVATVAAAVVVAYAAGAIIGESQPVLE | 13058 | 12869 | - | 18 |
| 5 | VSVTDAGDSATIPRIASADAAAVHGRGLSMV | STLANHIAIHDTGEGHTVTAELTTARPGGHP | 14633 | 14444 | - | 7 |
| 6 | VLAVSVTDAGDSATIPRIASADAAAVHGRGLS | MVSTLANHIAIHDTGEGHTVTAELTTARPGGHP | 14642 | 14444 | - | 7 |

### Results for WP_069884696.1 [Streptomyces luteocolor] back to top

previous - next

#### Architecture

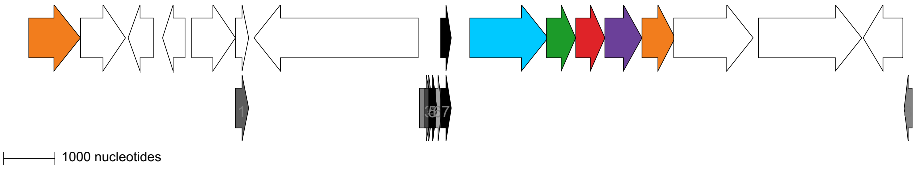

Link to nucleotide sequence

| Accession | start | end | direction | length (aa) | Pfam/HMM | name | description | E-value |
|---|---|---|---|---|---|---|---|---|
| WP_055697687.1 | 20906 | 21920 | + | 337 | TIGR01188<br>TIGR01288<br>TIGR03864<br>TIGR03522<br>TIGR01257 | TIGR01188<br>TIGR01288<br>TIGR03864<br>TIGR03522<br>TIGR01257 | drrA: daunorubicin resistance ABC transporter, ATP-binding protein<br>nodI: nodulation ABC transporter NodI<br>PQQ_ABC_ATP: ABC transporter, ATP-binding subunit, PQQ-dependent alcohol dehydrogenase system<br>GldA_ABC_ATP: gliding motility-associated ABC transporter ATP-binding subunit GldA<br>rim_protein: rim ABC transporter | 1.50E-107<br>1.40E-55<br>2.10E-51<br>5.20E-48<br>3.00E-43 |
| WP_069884672.1 | 21916 | 22786 | + | 289 | PF01061<br>TIGR01247<br>PF12698<br>TIGR00025<br>TIGR01248 | ABC2_membrane<br>TIGR01247<br>ABC2_membrane_3<br>TIGR00025<br>TIGR01248 | ABC-2 type transporter<br>drrB: daunorubicin resistance ABC transporter membrane protein<br>ABC-2 family transporter protein<br>Mtu_efflux: ABC transporter efflux protein, DrrB family<br>drrC: daunorubicin resistance protein C | 5.30E-30<br>1.30E-20<br>3.00E-18<br>4.70E-18<br>1.30E-10 |
| WP_055697685.1 | 23345 | 22847 | - | 165 | TIGR01462<br>PF03449<br>TIGR01461<br>PF01272 | TIGR01462<br>GreA_GreB_N<br>TIGR01461<br>GreA_GreB | greA: transcription elongation factor GreA<br>Transcription elongation factor, N-terminal<br>greB: transcription elongation factor GreB<br>Transcription elongation factor, GreA/GreB, C-term | 1.40E-44<br>6.30E-25<br>8.00E-15<br>2.00E-13 |
| WP_055697684.1 | 23948 | 23516 | - | 143 | PF14155 | DUF4307 | Domain of unknown function (DUF4307) | 4.80E-31 |
| WP_055697683.1 | 24084 | 24945 | + | 286 | TIGR03446<br>TIGR03445<br>PF02585<br>TIGR04001<br>TIGR04000 | TIGR03446<br>TIGR03445<br>PIG-L<br>TIGR04001<br>TIGR04000 | mycothiol_Mca: mycothiol conjugate amidase Mca<br>mycothiol_MshB: N-acetyl-1-D-myo-inositol-2-amino-2-deoxy-alpha-D-glucopyranoside deacetylase<br>GlcNAc-PI de-N-acetylase<br>thiol_BshB1: bacillithiol biosynthesis deacetylase BshB1<br>thiol_BshB2: bacillithiol biosynthesis deacetylase BshB2 | 2.70E-147<br>7.40E-52<br>1.80E-26<br>4.20E-13<br>4.40E-11 |
| WP_079036394.1 | 24946 | 25189 | + | 80 | NO PFAM MATCH | - | - | - |

| | | | | | | | | |
|---|---|---|---|---|---|---|---|---|
| WP_069884673.1 | 28522 | 25303 | - | 1072 | PF13424<br>PF17874<br>TIGR02917<br>PF00931<br>PF07719 | TPR_12<br>MalT<br>TIGR02917<br>NB-ARC<br>TPR_2 | Tetratricopeptide repeat<br>MalT-like TPR region<br>PEP_TPR_lipo: putative PEP-CTERM system TPR-repeat lipoprotein<br>NB-ARC domain<br>Tetratricopeptide repeat | 3.20E-18<br>1.30E-08<br>1.40E-08<br>1.40E-07<br>2.10E-06 |
| WP_158070518.1 | 28941 | 29145 | + | 67 | LinA | LinA | LinA (linaridin precursor) | 4.00E-14 |
| WP_069884696.1 | 29517 | 31035 | + | 505 | LinH<br>LinE<br>LinG<br>PF12697<br>TIGR02427 | LinH<br>LinE<br>LinG<br>Abhydrolase_6<br>TIGR02427 | LinH (Fusion of LinE/LinG)<br>LinE Homolog<br>LinG Homolog<br>Alpha/beta hydrolase family<br>protocat_pcaD: 3-oxoadipate enol-lactonase | 2.60E-<br>163<br>6.40E-70<br>1.20E-65<br>4.10E-10<br>1.00E-08 |
| WP_055697679.1 | 31024 | 31585 | + | 186 | LinL | LinL | LinL | 5.40E-53 |
| WP_069884674.1 | 31581 | 32154 | + | 190 | TIGR00521<br>PF02441<br>TIGR02113 | TIGR00521<br>Flavoprotein<br>TIGR02113 | coaBC_dfp: phosphopantothenoylcysteine decarboxylase / phosphopantothenate--cysteine ligase<br>Flavoprotein<br>coaC_strep: phosphopantothenoylcysteine decarboxylase | 2.20E-11<br>1.40E-09<br>4.00E-09 |
| WP_069884675.1 | 32146 | 32884 | + | 245 | PF13649<br>PF08241<br>PF13847<br>TIGR02021<br>PF08242 | Methyltransf_25<br>Methyltransf_11<br>Methyltransf_31<br>TIGR02021<br>Methyltransf_12 | Methyltransferase domain<br>Methyltransferase domain<br>Methyltransferase domain<br>BchM-ChlM: magnesium protoporphyrin O-methyltransferase<br>Methyltransferase domain | 6.60E-17<br>9.20E-15<br>5.20E-11<br>6.30E-11<br>6.60E-10 |
| WP_055699468.1 | 32880 | 33513 | + | 210 | TIGR02857<br>TIGR01189<br>TIGR02673<br>TIGR02982<br>TIGR01166 | TIGR02857<br>TIGR01189<br>TIGR02673<br>TIGR02982<br>TIGR01166 | CydD: thiol reductant ABC exporter, CydD subunit<br>ccmA: heme ABC exporter, ATP-binding protein CcmA<br>FtsE: cell division ATP-binding protein FtsE<br>heterocyst_DevA: ABC exporter ATP-binding subunit, DevA family<br>cbiO: cobalt ABC transporter, ATP-binding protein | 1.70E-29<br>2.60E-27<br>7.70E-27<br>9.10E-24<br>3.60E-23 |
| WP_055699467.1 | 33509 | 35060 | + | 516 | NO PFAM<br>MATCH | - | - | - |
| WP_069884676.1 | 35144 | 37166 | + | 673 | PF03190<br>PF13899<br>PF07221 | Thioredox_DsbH<br>Thioredoxin_7<br>GlcNAc_2-epim | Protein of unknown function, DUF255<br>Thioredoxin-like<br>N-acylglucosamine 2-epimerase (GlcNAc 2-epimerase) | 1.40E-74<br>8.90E-10<br>7.50E-05 |
| WP_055699465.1 | 37967 | 37199 | - | 255 | PF01927<br>PF14451 | Mut7-C<br>Ub-Mut7C | Mut7-C RNAse domain<br>Mut7-C ubiquitin | 6.00E-40<br>2.80E-23 |

| index | leader | core | start | end | dir | score |
|---|---|---|---|---|---|---|
| 1 | MNVHLAMTHLVPLAKEVDDDKVTPG...AVWGLMKSMNRHMGKVDFKEKEPVG | ATAGGDADRAADERTGEK | 24946 | 25189 | + | 8 |
| 2 | LPALHKCPAYGPGKRGRARIGDLPLAPVAGRGLAAAGR | GLLGGRAGPSRGRPSPLWLEIAVTSVTCQVPCCARCLKH | 28548 | 28782 | + | 7 |
| 3 | VAGRGLAAAGRGLLGGRAGPSRGRP | SPLWLEIAVTSVTCQVPCCARCLKH | 28629 | 28782 | + | 7 |
| 4 | MAGNSCHLCDVPSSLLRTVLEALTVLDVLLGPCLRSVRNVEL | QTLCSTFTQSGRRG | 28712 | 28883 | + | 10 |
| 5 | VPSSLLRTVLEALTVLDVLLGPCLRSVRNVEL | QTLCSTFTQSGRRG | 28742 | 28883 | + | 7 |
| 6 | VQHVHSERATGVATF | GTPRPFARMTTDERLVKCDLNRL | 28846 | 28963 | + | 6 |
| 7 | MRSESTVTRPDSAIRPEALAAQEFANTVLSGAAPGFHADCETPAM | ATPATPTAAQFVIQGSTICLVC | 28941 | 29145 | + | 35 |
| 8 | VRTFAMGLIGALTETTAYWAEHDHKDDLATLVD | RTLSTLTRGFAGAENPR | 38150 | 37997 | - | 7 |

### Results for WP_043371861.1 [Streptomyces luteus] back to top

previous - next

#### Architecture

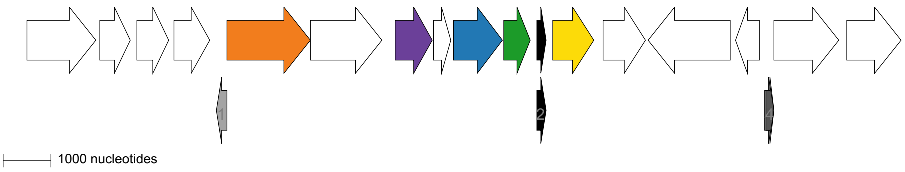

1000 nucleotides

Link to nucleotide sequence

| Accession | start | end | direction | length (aa) | Pfam/HMM | name | description | E-value |
|---|---|---|---|---|---|---|---|---|
| WP_043371850.1 | 334676 | 336137 | + | 486 | PF00171<br>TIGR02299<br>TIGR01780<br>TIGR01804<br>TIGR03216 | Aldedh<br>TIGR02299<br>TIGR01780<br>TIGR01804<br>TIGR03216 | Aldehyde dehydrogenase family<br>HpaE: 5-carboxymethyl-2-hydroxymuconate semialdehyde dehydrogenase<br>SSADH: succinate-semialdehyde dehydrogenase<br>BADH: betaine-aldehyde dehydrogenase<br>OH_muco_semi_DH: 2-hydroxymuconic semialdehyde dehydrogenase | 1.10E-148<br>3.90E-106<br>1.20E-101<br>7.00E-99<br>8.30E-91 |
| WP_043371852.1 | 336207 | 336849 | + | 213 | PF03551<br>TIGR03433<br>TIGR02719 | PadR<br>TIGR03433<br>TIGR02719 | Transcriptional regulator PadR-like family<br>padR_acidobact: transcriptional regulator, Acidobacterial, PadR-family<br>repress_PhaQ: poly-beta-hydroxybutyrate-responsive repressor | 1.10E-11<br>1.60E-06<br>2.00E-05 |
| WP_014674102.1 | 336994 | 337654 | + | 220 | INFERRED GENE | - | - | - |
| WP_043371856.1 | 337753 | 338515 | + | 253 | PF07739<br>PF13411<br>TIGR02043<br>TIGR02044<br>PF00376 | TipAS<br>MerR_1<br>TIGR02043<br>TIGR02044<br>MerR | TipAS antibiotic-recognition domain<br>MerR HTH family regulatory protein<br>ZntR: Zn(II)-responsive transcriptional regulator<br>CueR: Cu(I)-responsive transcriptional regulator<br>MerR family regulatory protein | 1.40E-38<br>9.20E-16<br>3.70E-14<br>4.00E-14<br>6.10E-13 |

| | | | | | | | | |
|---|---|---|---|---|---|---|---|---|
| WP_052412149.1 | 338866 | 340624 | + | 585 | TIGR02204<br>TIGR02203<br>TIGR00958<br>TIGR03375<br>TIGR03796 | TIGR02204<br>TIGR02203<br>TIGR00958<br>TIGR03375<br>TIGR03796 | MsbA_rel: ABC transporter, permease/ATP-binding protein<br>MsbA_lipidA: lipid A export permease/ATP-binding protein MsbA<br>3a01208: antigen peptide transporter 2<br>type_I_sec_LssB: type I secretion system ATPase<br>NHLM_micro_ABC1: NHLM bacteriocin system ABC transporter, peptidase/ATP-binding protein | 5.00E-98<br>3.20E-95<br>3.30E-86<br>2.50E-80<br>3.50E-78 |
| WP_043371858.1 | 340620 | 342102 | + | 493 | TIGR02734<br>TIGR03467<br>TIGR00292<br>TIGR02733<br>PF13450 | TIGR02734<br>TIGR03467<br>TIGR00292<br>TIGR02733<br>NAD_binding_8 | crtI_fam: phytoene desaturase<br>HpnE: squalene-associated FAD-dependent desaturase<br>TIGR00292: thiazole biosynthesis enzyme<br>desat_CrtD: C-3',4' desaturase CrtD<br>NAD(P)-binding Rossmann-like domain | 9.60E-11<br>8.20E-09<br>4.90E-08<br>3.40E-07<br>7.10E-07 |
| WP_043371860.1 | 342398 | 343157 | + | 252 | PF13649<br>PF08241<br>PF08242<br>PF13489<br>TIGR01983 | Methyltransf_25<br>Methyltransf_11<br>Methyltransf_12<br>Methyltransf_23<br>TIGR01983 | Methyltransferase domain<br>Methyltransferase domain<br>Methyltransferase domain<br>Methyltransferase domain<br>UbiG: 3-demethylubiquinone-9 3-O-methyltransferase | 1.20E-16<br>6.70E-12<br>2.10E-09<br>2.30E-08<br>2.40E-06 |
| WP_071528706.1 | 343198 | 343546 | + | 115 | PF13518<br>PF01527<br>PF13011<br>PF13551<br>PF13384 | HTH_28<br>HTH_Tnp_1<br>LZ_Tnp_IS481<br>HTH_29<br>HTH_23 | Helix-turn-helix domain<br>Transposase<br>leucine-zipper of insertion element IS481<br>Winged helix-turn helix<br>Homeodomain-like domain | 3.30E-09<br>9.90E-09<br>1.20E-06<br>5.00E-06<br>3.40E-05 |
| WP_043371861.1 | 343619 | 344645 | + | 341 | LinE<br>LinH<br>PF12697<br>TIGR02427<br>PF00975 | LinE<br>LinH<br>Abhydrolase_6<br>TIGR02427<br>Thioesterase | LinE Homolog<br>LinH (Fusion of LinE/LinG)<br>Alpha/beta hydrolase family<br>protocat_pcaD: 3-oxoadipate enol-lactonase<br>Thioesterase domain | 5.40E-98<br>6.70E-49<br>2.90E-13<br>9.10E-09<br>5.00E-07 |
| WP_043376870.1 | 344662 | 345220 | + | 185 | LinL | LinL | LinL | 2.30E-54 |
| WP_107063874.1 | 345361 | 345544 | + | 60 | LinA | LinA | LinA (linaridin precursor) | 3.00E-13 |
| WP_052412000.1 | 345687 | 346542 | + | 284 | LinG<br>LinH<br>TIGR04033 | LinG<br>LinH<br>TIGR04033 | LinG Homolog<br>LinH (Fusion of LinE/LinG)<br>export_SdpB: antimicrobial peptide system protein, SdpB family | 4.90E-109<br>3.80E-78<br>4.60E-04 |
| WP_043371863.1 | 346748 | 347627 | + | 292 | PF09335 | SNARE_assoc | SNARE associated Golgi protein | 2.40E-24 |
| WP_043371864.1 | 349393 | 347686 | - | 568 | PF06738<br>PF12821 | ThrE<br>ThrE_2 | Putative threonine/serine exporter<br>Threonine/Serine exporter, ThrE | 3.30E-67<br>1.10E-14 |
| WP_007452023.1 | 350012 | 349520 | - | 163 | PF00719 | Pyrophosphatase | Inorganic pyrophosphatase | 8.50E-53 |
| WP_043376876.1 | 350313 | 351687 | + | 457 | PF02113<br>TIGR00666<br>PF00768<br>PF13354 | Peptidase_S13<br>TIGR00666<br>Peptidase_S11<br>Beta-lactamase2 | D-Ala-D-Ala carboxypeptidase 3 (S13) family<br>PBP4: D-alanyl-D-alanine carboxypeptidase/D-alanyl-D-alanine-endopeptidase<br>D-alanyl-D-alanine carboxypeptidase<br>Beta-lactamase enzyme family | 1.30E-78<br>1.00E-57<br>6.60E-09<br>1.10E-05 |
| WP_043371866.1 | 351838 | 352966 | + | 375 | TIGR03883<br>TIGR03624<br>PF10103 | TIGR03883<br>TIGR03624<br>Zincin_2 | DUF2342_F420: uncharacterized protein, coenzyme F420 biosynthesis associated<br>TIGR03624: putative hydrolase<br>Zincin-like metallopeptidase | 7.50E-145<br>1.60E-127<br>8.70E-121 |

| index | leader | core | start | end | dir | score |
|---|---|---|---|---|---|---|
| 1 | VSSSRECCLLTRATDPAPFAAPDSGLAFSLWPSTSAPPAL | PTVTVNVPVPGGQPCRTSPSPRRPNVTRRSGLKLWVN | 338877 | 338643 | - | 6 |
| 2 | MSVVADFANAELTDVTPGRMGNDATPTM | LTPLATLATPEGVAVTSATAYALNEITNDLAG | 345361 | 345544 | + | 32 |
| 3 | MWCGPRVPSDRASRAPRGPRNRGSHGSHGPRRRSSPGSRGRRPGS | TPPVPPRPGWHWPPVW | 350134 | 350320 | + | 7 |
| 4 | VPSDRASRAPRGPRNRGSHGSHGPRRRSSPGSRGRRPGS | TPPVPPRPGWHWPPVW | 350152 | 350320 | + | 6 |

### Results for WP_149547799.1 [Streptomyces marokkonensis] back to top

previous - next

#### Architecture

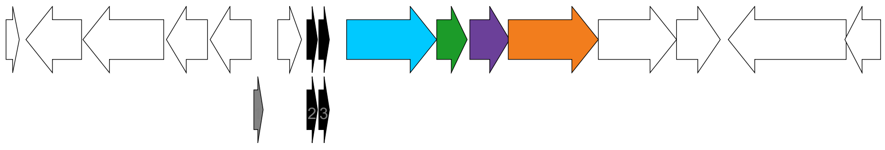

1000 nucleotides

Link to nucleotide sequence

| Accession | start | end | direction | length (aa) | Pfam/HMM | name | description | E-value |
|---|---|---|---|---|---|---|---|---|
| WP_149547791.1 | 33206 | 33455 | + | 82 | PF11273 | DUF3073 | Protein of unknown function (DUF3073) | 8.20E-22 |
| WP_149547792.1 | 34655 | 33587 | - | 355 | TIGR00878<br>PF02769<br>PF00586<br>TIGR01379<br>TIGR02124 | TIGR00878<br>AIRS_C<br>AIRS<br>TIGR01379<br>TIGR02124 | purM: phosphoribosylformylglycinamidine cyclo-ligase<br>AIR synthase related protein, C-terminal domain<br>AIR synthase related protein, N-terminal domain<br>thiL: thiamine-phosphate kinase<br>hypE: hydrogenase expression/formation protein HypE | 5.00E-124<br>1.20E-32<br>7.70E-16<br>3.80E-12<br>3.50E-11 |
| WP_149547793.1 | 36215 | 34688 | - | 508 | TIGR01134<br>TIGR01135<br>PF13522<br>PF13537<br>TIGR01536 | TIGR01134<br>TIGR01135<br>GATase_6<br>GATase_7<br>TIGR01536 | purF: amidophosphoribosyltransferase<br>glmS: glutamine-fructose-6-phosphate transaminase (isomerizing)<br>Glutamine amidotransferase domain<br>Glutamine amidotransferase domain<br>asn_synth_AEB: asparagine synthase (glutamine-hydrolyzing) | 3.10E-156<br>7.20E-27<br>1.70E-16<br>3.60E-14<br>1.10E-09 |
| WP_149547794.1 | 37063 | 36277 | - | 261 | PF03724 | META | META domain | 1.20E-51 |

| | | | | | | | | |
|---|---|---|---|---|---|---|---|---|
| WP_149547795.1 | 37910 | 37112 | - | 265 | PF17844<br>TIGR03083<br>PF11716<br>TIGR03086<br>TIGR03084 | SCP_3<br>TIGR03083<br>MDMPI_N<br>TIGR03086<br>TIGR03084 | Bacterial SCP ortholog<br>TIGR03083: uncharacterized Actinobacterial protein TIGR03083<br>Mycothiol maleylpyruvate isomerase N-terminal domain<br>TIGR03086: TIGR03086 family protein<br>TIGR03084: TIGR03084 family protein | 1.20E-39<br>4.80E-21<br>5.80E-15<br>8.50E-09<br>8.70E-06 |
| WP_149547796.1 | 38416 | 38860 | + | 147 | NO PFAM MATCH | - | - | - |
| WP_149547797.1 | 38979 | 39171 | + | 63 | LinA | LinA | LinA (linaridin precursor) | 2.40E-14 |
| WP_149547798.1 | 39200 | 39392 | + | 63 | LinA | LinA | LinA (linaridin precursor) | 1.70E-10 |
| WP_149547799.1 | 39725 | 41450 | + | 574 | LinH<br>LinG<br>LinE<br>PF12697<br>PF00561 | LinH<br>LinG<br>LinE<br>Abhydrolase_6<br>Abhydrolase_1 | LinH (Fusion of LinE/LinG)<br>LinG Homolog<br>LinE Homolog<br>Alpha/beta hydrolase family<br>alpha/beta hydrolase fold | 2.40E-219<br>6.60E-98<br>7.40E-97<br>5.10E-15<br>2.50E-11 |
| WP_149547800.1 | 41456 | 42035 | + | 192 | LinL | LinL | LinL | 1.50E-65 |
| WP_149547801.1 | 42094 | 42850 | + | 251 | PF13649<br>PF08241<br>PF13847<br>PF08242<br>PF13489 | Methyltransf_25<br>Methyltransf_11<br>Methyltransf_31<br>Methyltransf_12<br>Methyltransf_23 | Methyltransferase domain<br>Methyltransferase domain<br>Methyltransferase domain<br>Methyltransferase domain<br>Methyltransferase domain | 7.90E-15<br>1.50E-10<br>4.40E-08<br>1.50E-07<br>4.40E-07 |
| WP_149547802.1 | 42846 | 44574 | + | 575 | TIGR02204<br>TIGR02203<br>TIGR00958<br>TIGR03796<br>TIGR01846 | TIGR02204<br>TIGR02203<br>TIGR00958<br>TIGR03796<br>TIGR01846 | MsbA_rel: ABC transporter, permease/ATP-binding protein<br>MsbA_lipidA: lipid A export permease/ATP-binding protein MsbA<br>3a01208: antigen peptide transporter 2<br>NHLM_micro_ABC1: NHLM bacteriocin system ABC transporter, peptidase/ATP-binding protein<br>type_I_sec_HlyB: type I secretion system ATPase | 1.60E-103<br>6.40E-98<br>7.90E-87<br>2.40E-84<br>2.30E-82 |
| WP_149547803.1 | 44570 | 46055 | + | 494 | TIGR02734<br>TIGR03467<br>TIGR00292<br>PF12831<br>TIGR01317 | TIGR02734<br>TIGR03467<br>TIGR00292<br>FAD_oxidored<br>TIGR01317 | crtI_fam: phytoene desaturase<br>HpnE: squalene-associated FAD-dependent desaturase<br>TIGR00292: thiazole biosynthesis enzyme<br>FAD dependent oxidoreductase<br>GOGAT_sm_gam: glutamate synthase, NADH/NADPH, small subunit | 5.70E-10<br>1.40E-08<br>8.60E-08<br>1.50E-07<br>1.90E-07 |
| WP_149547804.1 | 46051 | 46897 | + | 281 | PF14534 | DUF4440 | Domain of unknown function (DUF4440) | 7.20E-04 |
| WP_149547805.1 | 49304 | 47045 | - | 752 | TIGR01736<br>PF02769<br>TIGR01735<br>TIGR01857<br>PF00586 | TIGR01736<br>AIRS_C<br>TIGR01735<br>TIGR01857<br>AIRS | FGAM_synth_II: phosphoribosylformylglycinamidine synthase II<br>AIR synthase related protein, C-terminal domain<br>FGAM_synt: phosphoribosylformylglycinamidine synthase<br>FGAM-synthase: phosphoribosylformylglycinamidine synthase<br>AIR synthase related protein, N-terminal domain | 1.80E-262<br>3.20E-54<br>1.30E-51<br>1.40E-49<br>1.10E-47 |
| WP_149547806.1 | 49981 | 49300 | - | 226 | TIGR01737<br>PF13507<br>TIGR01735<br>TIGR01857<br>PF07685 | TIGR01737<br>GATase_5<br>TIGR01735<br>TIGR01857<br>GATase_3 | FGAM_synth_I: phosphoribosylformylglycinamidine synthase I<br>CobB/CobQ-like glutamine amidotransferase domain<br>FGAM_synt: phosphoribosylformylglycinamidine synthase<br>FGAM-synthase: phosphoribosylformylglycinamidine synthase<br>CobB/CobQ-like glutamine amidotransferase domain | 6.40E-81<br>1.60E-34<br>2.30E-18<br>2.10E-17<br>1.40E-08 |

| index | leader | core | start | end | dir | score |
|---|---|---|---|---|---|---|
| 1 | LTSHSVGPTT | GTAHSGTAIAPLPATGSPTRDTEETARNDGFAPLRRLGFRVSEAARVR | 37952 | 38129 | + | 7 |
| 2 | MEQKISAFMNELDLGPLEGGYQEDSGLQVLA | TPAAVATCWYIAGGVAGAGAVVGAYVAGRNEG | 38979 | 39171 | + | 29 |
| 3 | MSKIIESVAAVTPAELTDEAHGEFGAMMPVQ | ATPGLVAFGVGFAGGAGVAWVTVQAYEAGAND | 39200 | 39392 | + | 31 |

### Results for WP_167503753.1 [Streptomyces malaysiensis] back to top

previous - next

#### Architecture

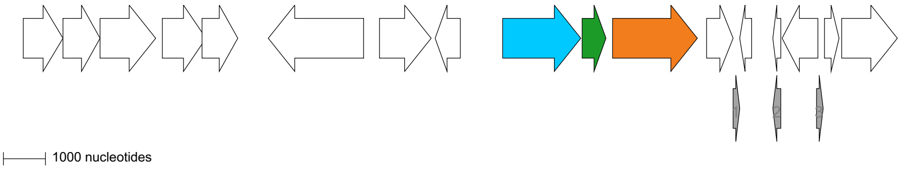

Link to nucleotide sequence

| Accession | start | end | direction | length (aa) | Pfam/HMM | name | description | E-value |
|---|---|---|---|---|---|---|---|---|
| WP_167503747.1 | 8462095 | 8463043 | + | 315 | PF00528<br>TIGR03262<br>TIGR00969<br>TIGR02141<br>TIGR01581 | BPD_transp_1<br>TIGR03262<br>TIGR00969<br>TIGR02141<br>TIGR01581 | Binding-protein-dependent transport system inner membrane component<br>PhnU2: putative 2-aminoethylphosphonate ABC transporter, permease protein<br>3a0106s02: sulfate ABC transporter, permease protein<br>modB_ABC: molybdate ABC transporter, permease protein<br>Mo_ABC_porter: NifC-like ABC-type porter | 3.20E-24<br>1.80E-16<br>1.60E-15<br>1.30E-14<br>2.80E-14 |
| WP_167503748.1 | 8463039 | 8463918 | + | 292 | PF00528<br>TIGR01581<br>TIGR03262<br>TIGR02141<br>TIGR02139 | BPD_transp_1<br>TIGR01581<br>TIGR03262<br>TIGR02141<br>TIGR02139 | Binding-protein-dependent transport system inner membrane component<br>Mo_ABC_porter: NifC-like ABC-type porter<br>PhnU2: putative 2-aminoethylphosphonate ABC transporter, permease protein<br>modB_ABC: molybdate ABC transporter, permease protein<br>permease_CysT: sulfate ABC transporter, permease protein CysT | 1.10E-21<br>8.70E-17<br>7.10E-12<br>4.70E-11<br>7.30E-10 |
| WP_167503749.1 | 8463917 | 8465231 | + | 437 | PF01547<br>PF13416 | SBP_bac_1<br>SBP_bac_8 | Bacterial extracellular solute-binding protein<br>Bacterial extracellular solute-binding protein | 1.00E-33<br>7.40E-22 |
| WP_167503750.1 | 8465383 | 8466433 | + | 349 | PF01066<br>TIGR00473 | CDP-OH_P_transf<br>TIGR00473 | CDP-alcohol phosphatidyltransferase<br>pssA: CDP-diacylglycerol-serine O-phosphatidyltransferase | 4.40E-10<br>4.70E-05 |
| WP_014055586.1 | 8466336 | 8467200 | + | 288 | INFERRED GENE | - | - | - |

| | | | | | | | | |
|---|---|---|---|---|---|---|---|---|
| WP_167503751.1 | 8470184 | 8467904 | - | 759 | PF03176<br>TIGR00833<br>TIGR00921<br>TIGR03480<br>TIGR00916 | MMPL<br>TIGR00833<br>TIGR00921<br>TIGR03480<br>TIGR00916 | MMPL family<br>actII: transport protein<br>2A067: efflux transporter, putative, hydrophobe/amphiphile efflux-3 (HAE3) family<br>HpnN: hopanoid biosynthesis associated RND transporter like protein HpnN<br>2A0604s01: protein-export membrane protein, SecD/SecF family | 5.70E-88<br>6.10E-53<br>2.10E-41<br>1.20E-14<br>5.90E-05 |
| WP_079259333.1 | 8470556 | 8471777 | + | 406 | PF00082<br>TIGR03921<br>TIGR03895<br>PF05922 | Peptidase_S8<br>TIGR03921<br>TIGR03895<br>Inhibitor_I9 | Subtilase family<br>T7SS_mycosin: type VII secretion-associated serine protease mycosin<br>protease_PatA: cyanobactin maturation protease, PatA/PatG family<br>Peptidase inhibitor I9 | 2.30E-43<br>4.20E-40<br>7.60E-12<br>3.70E-07 |
| WP_167503752.1 | 8472493 | 8471884 | - | 202 | PF05685 | Uma2 | Putative restriction endonuclease | 3.30E-19 |
| WP_167503753.1 | 8473498 | 8475346 | + | 615 | LinH<br>LinE<br>LinG<br>PF00561<br>TIGR02427 | LinH<br>LinE<br>LinG<br>Abhydrolase_1<br>TIGR02427 | LinH (Fusion of LinE/LinG)<br>LinE Homolog<br>LinG Homolog<br>alpha/beta hydrolase fold<br>protocat_pcaD: 3-oxoadipate enol-lactonase | 1.50E-169<br>1.50E-91<br>9.20E-90<br>4.60E-13<br>1.10E-11 |
| WP_167504604.1 | 8475390 | 8475954 | + | 187 | LinL | LinL | LinL | 3.60E-56 |
| WP_167504605.1 | 8476095 | 8478126 | + | 676 | TIGR02204<br>TIGR02203<br>TIGR00958<br>TIGR03796<br>TIGR03375 | TIGR02204<br>TIGR02203<br>TIGR00958<br>TIGR03796<br>TIGR03375 | MsbA_rel: ABC transporter, permease/ATP-binding protein<br>MsbA_lipidA: lipid A export permease/ATP-binding protein MsbA<br>3a01208: antigen peptide transporter 2<br>NHLM_micro_ABC1: NHLM bacteriocin system ABC transporter, peptidase/ATP-binding protein<br>type_I_sec_LssB: type I secretion system ATPase | 8.00E-107<br>1.00E-103<br>1.70E-89<br>2.40E-84<br>1.60E-78 |
| WP_167504606.1 | 8478325 | 8478979 | + | 217 | PF00196<br>PF08448<br>TIGR03020<br>TIGR00229 | GerE<br>PAS_4<br>TIGR03020<br>TIGR00229 | Bacterial regulatory proteins, luxR family<br>PAS fold<br>EpsA: transcriptional regulator EpsA<br>sensory_box: PAS domain S-box protein | 6.30E-09<br>4.20E-07<br>4.30E-06<br>3.80E-05 |
| WP_167503754.1 | 8479410 | 8479137 | - | 90 | NO PFAM MATCH | - | - | - |
| WP_167503755.1 | 8480119 | 8479915 | - | 67 | PF04149 | DUF397 | Domain of unknown function (DUF397) | 3.10E-13 |
| WP_167503756.1 | 8480982 | 8480130 | - | 283 | PF13560<br>PF12844<br>PF01381<br>PF13744<br>TIGR03070 | HTH_31<br>HTH_19<br>HTH_3<br>HTH_37<br>TIGR03070 | Helix-turn-helix domain<br>Helix-turn-helix domain<br>Helix-turn-helix<br>Helix-turn-helix domain<br>couple_hipB: transcriptional regulator, y4mF family | 6.20E-14<br>3.60E-07<br>2.20E-06<br>5.80E-05<br>3.00E-04 |
| WP_167503757.1 | 8481157 | 8481484 | + | 108 | NO PFAM MATCH | - | - | - |
| WP_167503758.1 | 8481538 | 8482888 | + | 449 | PF13304<br>PF02463<br>PF13175<br>TIGR04435<br>PF13476 | AAA_21<br>SMC_N<br>AAA_15<br>TIGR04435<br>AAA_23 | AAA domain, putative AbiEii toxin, Type IV TA system<br>RecF/RecN/SMC N terminal domain<br>AAA ATPase domain<br>restrict_AAA_1: restriction system-associated AAA family ATPase<br>AAA domain | 4.00E-23<br>2.10E-11<br>7.10E-11<br>7.10E-07<br>2.10E-06 |

| index | leader | core | start | end | dir | score |
|---|---|---|---|---|---|---|
| 1 | VTPRRRNAPAAGRGDLEDGPPKE | PTVRYEESRLRGLGGVVRSARAPHNPTQ | 8478975 | 8479131 | + | 6 |
| 2 | MEEPMTWRKSSFSGPGETNDCVELASVDGEIRLREGDTPADILT | TTPGELRTFIRAVKAGALDRFGR | 8480119 | 8479915 | - | 6 |
| 3 | LTSHRHARRLRLLDNPTGVRSVALSAPNARHKVVA | TTTLWRTLRDALITQR | 8480964 | 8481120 | + | 6 |

### Results for WP_078624126.1 [Streptomyces monomycini] back to top

previous - next

#### Architecture

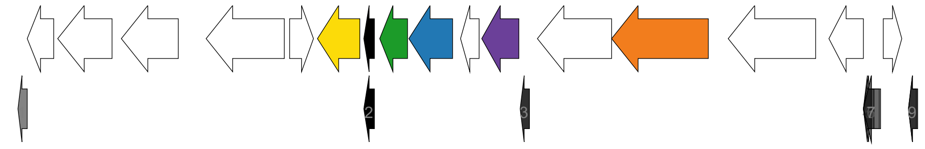

1000 nucleotides

Link to nucleotide sequence

| Accession | start | end | direction | length (aa) | Pfam/HMM | name | description | E-value |
|---|---|---|---|---|---|---|---|---|
| WP_030019223.1 | 33065 | 32525 | - | 179 | TIGR01203<br>PF00156<br>TIGR01251<br>TIGR01367 | TIGR01203<br>Pribosyltran<br>TIGR01251<br>TIGR01367 | HGPRTase: hypoxanthine phosphoribosyltransferase<br>Phosphoribosyl transferase domain<br>ribP_PPkin: ribose-phosphate diphosphokinase<br>pyrE_Therm: orotate phosphoribosyltransferase | 1.40E-64<br>3.70E-25<br>1.10E-05<br>4.30E-04 |
| WP_030019224.1 | 34242 | 33153 | - | 362 | TIGR02432<br>PF01171<br>PF09179 | TIGR02432<br>ATP_bind_3<br>TilS | lysidine_TilS_N: tRNA(Ile)-lysidine synthetase<br>PP-loop family<br>TilS substrate binding domain | 9.00E-43<br>5.00E-38<br>1.10E-11 |
| WP_030019225.1 | 35594 | 34451 | - | 380 | TIGR03883<br>TIGR03624<br>PF10103 | TIGR03883<br>TIGR03624<br>Zincin_2 | DUF2342_F420: uncharacterized protein, coenzyme F420 biosynthesis associated<br>TIGR03624: putative hydrolase<br>Zincin-like metallopeptidase | 1.40E-141<br>2.60E-124<br>9.00E-122 |
| WP_030019226.1 | 37773 | 36165 | - | 535 | PF02113<br>TIGR00666 | Peptidase_S13<br>TIGR00666 | D-Ala-D-Ala carboxypeptidase 3 (S13) family<br>PBP4: D-alanyl-D-alanine carboxypeptidase/D-alanyl-D-alanine-endopeptidase | 3.60E-78<br>1.60E-56 |
| WP_004571927.1 | 37859 | 38357 | + | 165 | PF00719 | Pyrophosphatase | Inorganic pyrophosphatase | 2.50E-53 |
| WP_030019227.1 | 39299 | 38438 | - | 286 | LinG<br>LinH<br>TIGR04033 | LinG<br>LinH<br>TIGR04033 | LinG Homolog<br>LinH (Fusion of LinE/LinG)<br>export_SdpB: antimicrobial peptide system protein, SdpB family | 4.80E-107<br>9.60E-77<br>5.50E-04 |
| WP_030019228.1 | 39589 | 39397 | - | 63 | LinA | LinA | LinA (linaridin precursor) | 1.20E-10 |

| | | | | | | | | |
|---|---|---|---|---|---|---|---|---|
| WP_050502306.1 | 40282 | 39721 | - | 186 | LinL | LinL | LinL | 3.50E-54 |
| WP_078624126.1 | 41205 | 40299 | - | 301 | LinE<br>LinH<br>PF12697<br>TIGR02427<br>PF00561 | LinE<br>LinH<br>Abhydrolase_6<br>TIGR02427<br>Abhydrolase_1 | LinE Homolog<br>LinH (Fusion of LinE/LinG)<br>Alpha/beta hydrolase family<br>protocat_pcaD: 3-oxoadipate enol-lactonase<br>alpha/beta hydrolase fold | 1.30E-95<br>5.50E-51<br>2.20E-11<br>1.30E-10<br>4.20E-07 |
| WP_078624119.1 | 41741 | 41363 | - | 125 | PF01527<br>PF13518<br>PF13551<br>PF13384<br>PF13011 | HTH_Tnp_1<br>HTH_28<br>HTH_29<br>HTH_23<br>LZ_Tnp_IS481 | Transposase<br>Helix-turn-helix domain<br>Winged helix-turn helix<br>Homeodomain-like domain<br>leucine-zipper of insertion element IS481 | 7.80E-09<br>6.00E-08<br>8.80E-06<br>4.70E-05<br>6.40E-04 |
| WP_030019231.1 | 42554 | 41786 | - | 255 | PF13649<br>PF08241<br>PF08242<br>PF13489<br>PF13847 | Methyltransf_25<br>Methyltransf_11<br>Methyltransf_12<br>Methyltransf_23<br>Methyltransf_31 | Methyltransferase domain<br>Methyltransferase domain<br>Methyltransferase domain<br>Methyltransferase domain<br>Methyltransferase domain | 1.50E-15<br>1.30E-11<br>3.40E-10<br>3.80E-08<br>2.50E-07 |
| WP_050502300.1 | 44440 | 42928 | - | 503 | TIGR03467<br>TIGR00292<br>TIGR02734<br>PF13450<br>TIGR03315 | TIGR03467<br>TIGR00292<br>TIGR02734<br>NAD_binding_8<br>TIGR03315 | HpnE: squalene-associated FAD-dependent desaturase<br>TIGR00292: thiazole biosynthesis enzyme<br>crtI_fam: phytoene desaturase<br>NAD(P)-binding Rossmann-like domain<br>Se_ygfK: putative selenate reductase, YgfK subunit | 7.30E-09<br>1.10E-08<br>1.60E-08<br>3.10E-08<br>5.90E-07 |
| WP_078624127.1 | 46398 | 44436 | - | 653 | TIGR02203<br>TIGR02204<br>TIGR00958<br>TIGR03796<br>TIGR03375 | TIGR02203<br>TIGR02204<br>TIGR00958<br>TIGR03796<br>TIGR03375 | MsbA_lipidA: lipid A export permease/ATP-binding protein MsbA<br>MsbA_rel: ABC transporter, permease/ATP-binding protein<br>3a01208: antigen peptide transporter 2<br>NHLM_micro_ABC1: NHLM bacteriocin system ABC transporter, peptidase/ATP-binding protein<br>type_I_sec_LssB: type I secretion system ATPase | 1.60E-95<br>5.50E-95<br>4.40E-90<br>6.60E-83<br>9.40E-77 |
| WP_030019234.1 | 48587 | 46799 | - | 595 | PF06738<br>PF12821 | ThrE<br>ThrE_2 | Putative threonine/serine exporter<br>Threonine/Serine exporter, ThrE | 1.50E-71<br>1.70E-18 |
| WP_030019235.1 | 49576 | 48856 | - | 239 | PF09335 | SNARE_assoc | SNARE associated Golgi protein | 5.70E-25 |
| WP_030019236.1 | 49977 | 50349 | + | 123 | PF01906<br>TIGR03884 | YbjQ_1<br>TIGR03884 | Putative heavy-metal-binding<br>sel_bind_Methan: selenium-binding protein | 4.20E-34<br>6.40E-04 |

| index | leader | core | start | end | dir | score |
|---|---|---|---|---|---|---|
| 1 | VDPSGRPRCGRGPGPTGARRREPLRGRRRWSR | ENGIRSCIVNRGRRHRVTMLGYRPKVSLFSGQ | 32520 | 32325 | - | 7 |
| 2 | MSILADFANAELADVKPGRIGADSTPTM | VTPLATIATPEATPVGFAATSATAVALDKVTHNVG | 39589 | 39397 | - | 34 |
| 3 | VTNEALSPHTCAVAALLLDVRLCRSDCRAVDRGFPAVGLADVFPSR | PVPAPLRHDAPIGMTVGT | 42764 | 42569 | - | 9 |
| 4 | VAGSGAGRAGAATGGWNHGPSSGVDSFGPHTAPAFR | RTARSGALDPGAFTP | 49728 | 49572 | - | 7 |
| 5 | VEGRGRTEAAEPAAGRPRPAVAGSG...ATGGWNHGPSSGVDSFGPHTAPAFR | RTARSGALDPGAFTP | 49788 | 49572 | - | 6 |
| 6 | VGGIGRVGEGGGGCVRPWRRRRGRG...ATGGWNHGPSSGVDSFGPHTAPAFR | RTARSGALDPGAFTP | 49899 | 49572 | - | 6 |
| 7 | MGERRVGGIGRVGEGGGGCVRPWRR...ATGGWNHGPSSGVDSFGPHTAPAFR | RTARSGALDPGAFTP | 49914 | 49572 | - | 6 |
| 8 | VAEEHRRFISSGYYDCSHEMHTCLGEMYVA | DERFTATYEAIRPGLAVYMRDAMLANAVRMS | 50658 | 50472 | - | 7 |
| 9 | MDVAEEHRRFISSGYYDCSHEMHTCLGEMYV | ADERFTATYEAIRPGLAVYMRDAMLANAVRMS | 50664 | 50472 | - | 8 |

### Results for OEV05017.1 [Streptomyces nanshensis] back to top

previous - next

#### Architecture

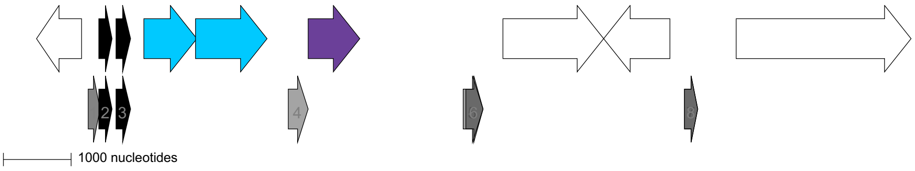

Link to nucleotide sequence

| Accession | start | end | direction | length (aa) | Pfam/HMM | name | description | E-value |
|---|---|---|---|---|---|---|---|---|
| OEV05013.1 | 777 | 114 | - | 220 | PF05719 | GPP34 | Golgi phosphoprotein 3 (GPP34) | 3.00E-42 |
| OEV05014.1 | 1033 | 1228 | + | 64 | LinA | LinA | LinA (linaridin precursor) | 6.20E-12 |
| OEV05015.1 | 1282 | 1495 | + | 70 | LinA | LinA | LinA (linaridin precursor) | 1.20E-15 |
| OEV05016.1 | 1700 | 2471 | + | 256 | LinH<br>LinG | LinH<br>LinG | LinH (Fusion of LinE/LinG)<br>LinG Homolog | 2.20E-114<br>7.60E-108 |
| OEV05017.1 | 2467 | 3508 | + | 346 | LinH<br>LinE<br>PF12697<br>PF12146<br>PF00561 | LinH<br>LinE<br>Abhydrolase_6<br>Hydrolase_4<br>Abhydrolase_1 | LinH (Fusion of LinE/LinG)<br>LinE Homolog<br>Alpha/beta hydrolase family<br>Serine aminopeptidase, S33<br>alpha/beta hydrolase fold | 2.20E-135<br>1.70E-122<br>2.10E-19<br>1.70E-09<br>3.50E-09 |
| OEV05018.1 | 4124 | 4895 | + | 256 | PF13649<br>PF08241<br>PF08242<br>PF13489<br>PF13847 | Methyltransf_25<br>Methyltransf_11<br>Methyltransf_12<br>Methyltransf_23<br>Methyltransf_31 | Methyltransferase domain<br>Methyltransferase domain<br>Methyltransferase domain<br>Methyltransferase domain<br>Methyltransferase domain | 3.00E-16<br>2.50E-10<br>1.40E-08<br>1.50E-08<br>1.70E-06 |

| | | | | | | | | |
|---|---|---|---|---|---|---|---|---|
| OEV05019.1 | 7005 | 8484 | + | 492 | TIGR02734<br>TIGR03467<br>PF13450<br>PF01593<br>TIGR01317 | TIGR02734<br>TIGR03467<br>NAD_binding_8<br>Amino_oxidase<br>TIGR01317 | crtI_fam: phytoene desaturase<br>HpnE: squalene-associated FAD-dependent desaturase<br>NAD(P)-binding Rossmann-like domain<br>Flavin containing amine oxidoreductase<br>GOGAT_sm_gam: glutamate synthase, NADH/NADPH, small subunit | 1.80E-12<br>9.60E-09<br>1.30E-08<br>1.50E-08<br>4.40E-08 |
| OEV05020.1 | 9466 | 8491 | - | 324 | PF01978<br>PF00196 | TrmB<br>GerE | Sugar-specific transcriptional regulator TrmB<br>Bacterial regulatory proteins, luxR family | 2.00E-05<br>3.60E-04 |
| OEV05021.1 | 10447 | 13027 | + | 859 | PF00196<br>PF13424<br>PF13176<br>PF13384<br>TIGR03020 | GerE<br>TPR_12<br>TPR_7<br>HTH_23<br>TIGR03020 | Bacterial regulatory proteins, luxR family<br>Tetratricopeptide repeat<br>Tetratricopeptide repeat<br>Homeodomain-like domain<br>EpsA: transcriptional regulator EpsA | 1.20E-13<br>4.30E-09<br>1.50E-07<br>2.90E-07<br>6.10E-07 |

| index | leader | core | start | end | dir | score |
|---|---|---|---|---|---|---|
| 1 | LTHAIVCDAGSELPRNPITILVTGFQVAISAHSAPESGAER | ETTKARERRRHTHVA | 873 | 1044 | + | 7 |
| 2 | MLLEKVNNSHLLEAVSMDAANVEDLSSAEPVL | STPAAAAGGALVTAGAFGAGVGLGVTQAQADG | 1033 | 1228 | + | 28 |
| 3 | MTAISIKVNASQELDGVSIHAGENEWNGRMSPVL | ATPAAVGAGLAGAAAVAGAGAAGIAVGNAAGQAAGG | 1282 | 1495 | + | 28 |
| 4 | VCGELGRLIVRKDVKVTDSAAYRLL...QFLVVLYSGHDDSEDPQYSFISPFI | RLPEPDPTDHADDVDHAGAADQLEPAA | 3831 | 4128 | + | 6 |
| 5 | MPQRPTSARPWRWPVSRMSSKVCRK...GSGCASPARCCRSPRSCFWTRPPPI | WTPTRSGISGKCSSGSPGTVRSSPSRTASPR | 6404 | 6701 | + | 6 |
| 6 | MSSKVCRKDSTPIWAIRAAGSPAVSGSGCASPARCCRSPRSCFWTRPPPI | WTPTRSGISGKCSSGSPGTVRSSPSRTASPR | 6455 | 6701 | + | 6 |
| 7 | LVSDPGGLWWKVRCSHCSWARAPMRRVSPD | ATGLVVIEVHRICGGVSGGLSSATGGPPPVR | 9685 | 9871 | + | 6 |
| 8 | VSDPGGLWWKVRCSHCSWARAPMRRVSPDA | TGLVVIEVHRICGGVSGGLSSATGGPPPVR | 9688 | 9871 | + | 6 |

### Results for SPE49561.1 [Streptomyces netropsis] back to top

previous - next

#### Architecture

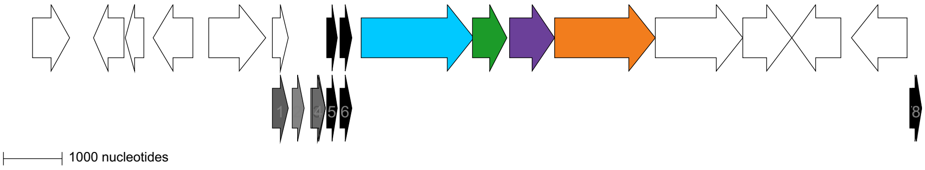

Link to nucleotide sequence

| Accession | start | end | direction | length (aa) | Pfam/HMM | name | description | E-value |
|---|---|---|---|---|---|---|---|---|
| SPE49553.1 | 590275 | 590917 | + | 213 | PF09335 | SNARE_assoc | SNARE associated Golgi protein | 2.30E-10 |
| SPE49554.1 | 591850 | 591325 | - | 174 | PF09348 | DUF1990 | Domain of unknown function (DUF1990) | 3.90E-51 |
| SPE49555.1 | 592179 | 591858 | - | 106 | NO PFAM MATCH | - | - | - |
| SPE49556.1 | 593005 | 592348 | - | 218 | PF02031<br>TIGR03296 | Peptidase_M7<br>TIGR03296 | Streptomyces extracellular neutral proteinase (M7) family<br>M6dom_TIGR03296: M6 family metalloprotease domain | 3.50E-79<br>2.50E-08 |
| SPE49557.1 | 593296 | 594253 | + | 318 | PF03466<br>TIGR03339<br>TIGR02424<br>PF00126 | LysR_substrate<br>TIGR03339<br>TIGR02424<br>HTH_1 | LysR substrate binding domain<br>phn_lysR: aminoethylphosphonate catabolism associated LysR family transcriptional regulator<br>TF_pcaQ: pca operon transcription factor PcaQ<br>Bacterial regulatory helix-turn-helix protein, lysR family | 2.90E-19<br>1.10E-11<br>9.90E-08<br>2.60E-04 |
| SPE49558.1 | 594381 | 594642 | + | 86 | NO PFAM MATCH | - | - | - |
| SPE49559.1 | 595294 | 595486 | + | 63 | LinA | LinA | LinA (linaridin precursor) | 3.40E-15 |
| SPE49560.1 | 595524 | 595716 | + | 63 | LinA | LinA | LinA (linaridin precursor) | 5.50E-11 |
| SPE49561.1 | 595886 | 597794 | + | 635 | LinH<br>LinG<br>LinE<br>PF12697<br>PF00561 | LinH<br>LinG<br>LinE<br>Abhydrolase_6<br>Abhydrolase_1 | LinH (Fusion of LinE/LinG)<br>LinG Homolog<br>LinE Homolog<br>Alpha/beta hydrolase family<br>alpha/beta hydrolase fold | 2.80E-243<br>3.30E-116<br>8.90E-103<br>1.70E-14<br>4.10E-13 |
| SPE49562.1 | 597790 | 598369 | + | 192 | LinL | LinL | LinL | 2.50E-63 |
| SPE49563.1 | 598419 | 599175 | + | 251 | PF13649<br>PF08241<br>PF13489<br>PF08242<br>PF13847 | Methyltransf_25<br>Methyltransf_11<br>Methyltransf_23<br>Methyltransf_12<br>Methyltransf_31 | Methyltransferase domain<br>Methyltransferase domain<br>Methyltransferase domain<br>Methyltransferase domain<br>Methyltransferase domain | 3.00E-14<br>6.80E-10<br>1.40E-09<br>5.40E-08<br>3.90E-07 |
| SPE49564.1 | 599171 | 600902 | + | 576 | TIGR02204<br>TIGR02203<br>TIGR00958<br>TIGR01846<br>TIGR03796 | TIGR02204<br>TIGR02203<br>TIGR00958<br>TIGR01846<br>TIGR03796 | MsbA_rel: ABC transporter, permease/ATP-binding protein<br>MsbA_lipidA: lipid A export permease/ATP-binding protein MsbA<br>3a01208: antigen peptide transporter 2<br>type_I_sec_HlyB: type I secretion system ATPase<br>NHLM_micro_ABC1: NHLM bacteriocin system ABC transporter, peptidase/ATP-binding protein | 7.40E-103<br>2.30E-97<br>1.20E-82<br>1.50E-82<br>3.60E-81 |
| SPE49565.1 | 600898 | 602383 | + | 494 | TIGR02734<br>TIGR03467<br>TIGR00292<br>TIGR02733<br>TIGR01317 | TIGR02734<br>TIGR03467<br>TIGR00292<br>TIGR02733<br>TIGR01317 | crtI_fam: phytoene desaturase<br>HpnE: squalene-associated FAD-dependent desaturase<br>TIGR00292: thiazole biosynthesis enzyme<br>desat_CrtD: C-3',4' desaturase CrtD<br>GOGAT_sm_gam: glutamate synthase, NADH/NADPH, small subunit | 3.90E-10<br>1.60E-08<br>2.60E-08<br>3.60E-08<br>4.10E-08 |
| SPE49566.1 | 602379 | 603228 | + | 282 | PF12680 | SnoaL_2 | SnoaL-like domain | 1.40E-06 |
| SPE49567.1 | 604055 | 603215 | - | 279 | PF01551 | Peptidase_M23 | Peptidase family M23 | 2.20E-17 |

| | | | | | | | | |
|---|---|---|---|---|---|---|---|---|
| SPE49568.1 | 605178 | 604233 | - | 314 | PF01408<br>TIGR04380 | GFO_IDH_MocA<br>TIGR04380 | Oxidoreductase family, NAD-binding Rossmann fold<br>myo_inos_iolG: inositol 2-dehydrogenase | 1.10E-19<br>1.40E-17 |

| index | leader | core | start | end | dir | score |
|---|---|---|---|---|---|---|
| 1 | MSGPRRQESRDAITVEISYAVASGC...LFLVVASPALFLDLGRGVESALFTC | ATLLAAAGFTMRAVTLLIRLRRAQDASADAR | 594381 | 594642 | + | 8 |
| 2 | MTDDARPPVQLLGVPNTPERYFRFLGKIGIPETFGRAES | ATPLPAHYAGRAGPRQQKPRSGPYRG | 594704 | 594902 | + | 7 |
| 3 | MDYRPTSGMIRSPNPDPKAAHVPSR...RTTNSILPVTHAASKGPISPSFPCG | ILPAAPVGKGRIRVPVRGNTTQYPVCSTSP | 595029 | 595275 | + | 6 |
| 4 | MIRSPNPDPKAAHVPSRWRTTNSILPVTHAASKGPISPSFPCG | ILPAAPVGKGRIRVPVRGNTTQYPVCSTSP | 595053 | 595275 | + | 6 |
| 5 | MEQKIAAFMNEQDLGALEQGYQDDAGPVL | ATPVAAVATCWYVGGAVAGAGAVVGAYVTGRVNG | 595294 | 595486 | + | 30 |
| 6 | MSKIIDSVVAVTPEKLTDETHGEFGAMMPIQ | ATPALVAFGVGFAGGAGAAWVTVQAYEAGAND | 595524 | 595716 | + | 31 |
| 7 | VVGERAQRVAGGVGWERRPKRFPRG...AGPAGAERGCCSTRAPSSRTAAAVA | ATSATRKVGMT | 605236 | 605440 | + | 10 |
| 8 | VGWERRPKRFPRGSRIRARAGPAGAERGCCSTRAPSSRTAAAVA | ATSATRKVGMT | 605272 | 605440 | + | 9 |

### Results for OEV08011.1 [Streptomyces nanshensis] back to top

previous - next

#### Architecture

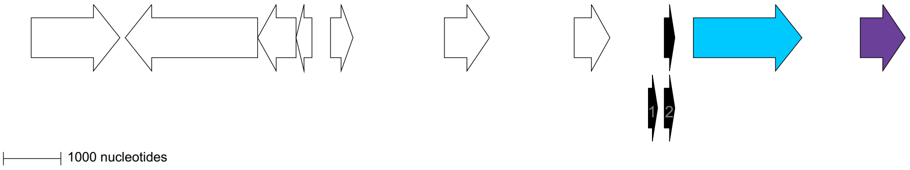

Link to nucleotide sequence

| Accession | start | end | direction | length (aa) | Pfam/HMM | name | description | E-value |
|---|---|---|---|---|---|---|---|---|
| OEV08000.1 | 5576 | 7106 | + | 509 | PF06964 | Alpha-L-AF_C | Alpha-L-arabinofuranosidase C-terminal domain | 6.70E-39 |
| OEV08001.1 | 9506 | 7217 | - | 762 | TIGR01736<br>PF02769<br>PF00586<br>TIGR01735<br>TIGR01857 | TIGR01736<br>AIRS_C<br>AIRS<br>TIGR01735<br>TIGR01857 | FGAM_synth_II: phosphoribosylformylglycinamidine synthase II<br>AIR synthase related protein, C-terminal domain<br>AIR synthase related protein, N-terminal domain<br>FGAM_synt: phosphoribosylformylglycinamidine synthase<br>FGAM-synthase: phosphoribosylformylglycinamidine synthase | 7.90E-262<br>4.70E-55<br>7.50E-49<br>7.20E-46<br>2.40E-45 |
| OEV08002.1 | 10192 | 9502 | - | 229 | TIGR01737<br>PF13507<br>TIGR01735<br>TIGR01857<br>PF07685 | TIGR01737<br>GATase_5<br>TIGR01735<br>TIGR01857<br>GATase_3 | FGAM_synth_I: phosphoribosylformylglycinamidine synthase I<br>CobB/CobQ-like glutamine amidotransferase domain<br>FGAM_synt: phosphoribosylformylglycinamidine synthase<br>FGAM-synthase: phosphoribosylformylglycinamidine synthase<br>CobB/CobQ-like glutamine amidotransferase domain | 3.50E-80<br>1.60E-33<br>6.00E-16<br>1.10E-14<br>1.50E-07 |
| OEV08003.1 | 10446 | 10188 | - | 85 | PF02700<br>TIGR00302 | PurS<br>TIGR00302 | Phosphoribosylformylglycinamidine (FGAM) synthase<br>TIGR00302: phosphoribosylformylglycinamidine synthase, purS protein | 1.40E-27<br>3.70E-20 |
| OEV08004.1 | 10790 | 11165 | + | 124 | PF11774 | Lsr2 | Lsr2 | 2.00E-34 |
| OEV08009.1 | 12756 | 13545 | + | 262 | PF01061<br>PF12698<br>TIGR03062<br>TIGR01247 | ABC2_membrane<br>ABC2_membrane_3<br>TIGR03062<br>TIGR01247 | ABC-2 type transporter<br>ABC-2 family transporter protein<br>pip_yhgE_Cterm: YhgE/Pip C-terminal domain<br>drrB: daunorubicin resistance ABC transporter membrane protein | 1.80E-19<br>2.10E-10<br>2.40E-06<br>1.90E-05 |
| OEV08010.1 | 15021 | 15654 | + | 210 | PF00072<br>TIGR02875<br>PF00196<br>TIGR02154<br>TIGR02956 | Response_reg<br>TIGR02875<br>GerE<br>TIGR02154<br>TIGR02956 | Response regulator receiver domain<br>spore_0_A: sporulation transcription factor Spo0A<br>Bacterial regulatory proteins, luxR family<br>PhoB: phosphate regulon transcriptional regulatory protein PhoB<br>TMAO_torS: TMAO reductase sytem sensor TorS | 2.50E-25<br>7.80E-14<br>8.50E-14<br>2.70E-11<br>9.50E-10 |
| OEV08005.1 | 16587 | 16767 | + | 59 | LinA | LinA | LinA (linaridin precursor) | 1.90E-15 |
| OEV08011.1 | 17088 | 18981 | + | 630 | LinH<br>LinG<br>LinE<br>PF12697<br>LinL | LinH<br>LinG<br>LinE<br>Abhydrolase_6<br>LinL | LinH (Fusion of LinE/LinG)<br>LinG Homolog<br>LinE Homolog<br>Alpha/beta hydrolase family<br>LinL | 5.70E-245<br>3.70E-118<br>9.20E-98<br>2.20E-14<br>2.70E-11 |
| OEV08006.1 | 20007 | 20787 | + | 259 | PF13649<br>PF08241<br>PF13489<br>PF13847<br>TIGR01983 | Methyltransf_25<br>Methyltransf_11<br>Methyltransf_23<br>Methyltransf_31<br>TIGR01983 | Methyltransferase domain<br>Methyltransferase domain<br>Methyltransferase domain<br>Methyltransferase domain<br>UbiG: 3-demethylubiquinone-9 3-O-methyltransferase | 5.30E-18<br>6.50E-13<br>1.20E-12<br>2.80E-11<br>7.90E-10 |

| index | leader | core | start | end | dir | score |
|---|---|---|---|---|---|---|
| 1 | MLEEKLNLQIQDELHEEHAGEIADKAAPVL | ATPALAGAFVGGAAVVTGAYAAGKAVG | 16306 | 16480 | + | 29 |
| 2 | MIDVTSHINESAELSSVEVTDAEGVVFTDYTPVL | ATPGAFLGGVAIASATVAAFEQGQG | 16587 | 16767 | + | 28 |

### Results for WP_150491658.1 [Streptomyces nitrosporeus] back to top

previous - next

#### Architecture

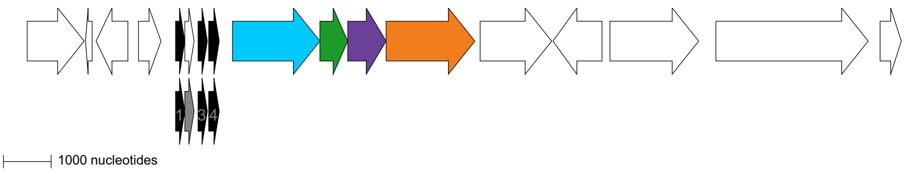

Link to nucleotide sequence

| Accession | start | end | direction | length (aa) | Pfam/HMM | name | description | E-value |
|---|---|---|---|---|---|---|---|---|
| WP_150486325.1 | 536666 | 537857 | + | 396 | PF03435<br>PF05368 | Sacchrp_dh_NADP<br>NmrA | Saccharopine dehydrogenase NADP binding domain<br>NmrA-like family | 1.30E-14<br>4.30E-04 |
| WP_167535313.1 | 538039 | 537892 | - | 48 | NO PFAM MATCH | - | - | - |
| WP_167535492.1 | 538787 | 538124 | - | 220 | PF04493 | Endonuclease_5 | Endonuclease V | 4.60E-64 |
| WP_150486327.1 | 539014 | 539488 | + | 157 | TIGR04431<br>PF13302<br>PF13523<br>PF00583<br>TIGR01575 | TIGR04431<br>Acetyltransf_3<br>Acetyltransf_8<br>Acetyltransf_1<br>TIGR01575 | N6_acetyl_AAC6: aminoglycoside N(6')-acetyltransferase, AacA4 family<br>Acetyltransferase (GNAT) domain<br>Acetyltransferase (GNAT) domain<br>Acetyltransferase (GNAT) family<br>rimI: ribosomal-protein-alanine acetyltransferase | 1.50E-12<br>7.00E-12<br>1.10E-10<br>1.10E-08<br>4.00E-06 |
| WP_150486328.1 | 539792 | 539978 | + | 61 | LinA | LinA | LinA (linaridin precursor) | 6.50E-08 |
| WP_150486329.1 | 539974 | 540172 | + | 65 | NO PFAM MATCH | - | - | - |
| WP_150486330.1 | 540244 | 540436 | + | 63 | LinA | LinA | LinA (linaridin precursor) | 3.20E-10 |
| WP_150486331.1 | 540475 | 540694 | + | 72 | LinA | LinA | LinA (linaridin precursor) | 1.90E-10 |
| WP_150491658.1 | 540981 | 542796 | + | 604 | LinH<br>LinG<br>LinE<br>LinL<br>PF12697 | LinH<br>LinG<br>LinE<br>LinL<br>Abhydrolase_6 | LinH (Fusion of LinE/LinG)<br>LinG Homolog<br>LinE Homolog<br>LinL<br>Alpha/beta hydrolase family | 5.50E-240<br>3.00E-114<br>3.20E-92<br>3.70E-18<br>1.10E-13 |
| WP_150486332.1 | 542792 | 543401 | + | 202 | LinL | LinL | LinL | 6.50E-63 |
| WP_150486333.1 | 543397 | 544204 | + | 268 | PF13649<br>PF08241<br>PF08242<br>PF13489<br>PF13847 | Methyltransf_25<br>Methyltransf_11<br>Methyltransf_12<br>Methyltransf_23<br>Methyltransf_31 | Methyltransferase domain<br>Methyltransferase domain<br>Methyltransferase domain<br>Methyltransferase domain<br>Methyltransferase domain | 1.40E-15<br>5.80E-12<br>1.90E-09<br>2.40E-08<br>7.10E-08 |
| WP_150486334.1 | 544200 | 546057 | + | 618 | TIGR02204<br>TIGR02203<br>TIGR00958<br>TIGR03796<br>TIGR01846 | TIGR02204<br>TIGR02203<br>TIGR00958<br>TIGR03796<br>TIGR01846 | MsbA_rel: ABC transporter, permease/ATP-binding protein<br>MsbA_lipidA: lipid A export permease/ATP-binding protein MsbA<br>3a01208: antigen peptide transporter 2<br>NHLM_micro_ABC1: NHLM bacteriocin system ABC transporter, peptidase/ATP-binding protein<br>type_I_sec_HlyB: type I secretion system ATPase | 1.40E-99<br>7.80E-93<br>2.40E-87<br>8.90E-86<br>2.70E-81 |
| WP_167535314.1 | 546163 | 547648 | + | 494 | TIGR02734<br>PF01593<br>TIGR03467<br>PF13450<br>TIGR00292 | TIGR02734<br>Amino_oxidase<br>TIGR03467<br>NAD_binding_8<br>TIGR00292 | crtI_fam: phytoene desaturase<br>Flavin containing amine oxidoreductase<br>HpnE: squalene-associated FAD-dependent desaturase<br>NAD(P)-binding Rossmann-like domain<br>TIGR00292: thiazole biosynthesis enzyme | 1.30E-10<br>2.30E-09<br>1.00E-08<br>1.40E-07<br>2.30E-07 |
| WP_150486335.1 | 548696 | 547691 | - | 334 | NO PFAM MATCH | - | - | - |
| WP_150486336.1 | 548890 | 550720 | + | 609 | NO PFAM MATCH | - | - | - |
| WP_150486337.1 | 551093 | 554285 | + | 1063 | PF04389<br>PF02868<br>PF01447<br>PF01546<br>PF07504 | Peptidase_M28<br>Peptidase_M4_C<br>Peptidase_M4<br>Peptidase_M20<br>FTP | Peptidase family M28<br>Thermolysin metallopeptidase, alpha-helical domain<br>Thermolysin metallopeptidase, catalytic domain<br>Peptidase family M20/M25/M40<br>Fungalysin/Thermolysin Propeptide Motif | 1.80E-42<br>1.00E-25<br>1.80E-24<br>8.00E-08<br>1.20E-06 |
| WP_150486338.1 | 554545 | 554962 | + | 138 | PF04686 | SsgA | Streptomyces sporulation and cell division protein, SsgA | 1.50E-31 |

| index | leader | core | start | end | dir | score |
|---|---|---|---|---|---|---|
| 1 | MQQARTPRWAAVGPLEEGAEGEVFGGAAPIA | ATQVVVTVAAAVVVAYAAGAIIGQTEPVMQ | 539792 | 539978 | + | 29 |
| 2 | MTGPRRAGAERLPFR | HTAPATTRELVPDDTGDPGAAAGPVASVVVAEFGAVVAVSFIGGSEYPHL | 539974 | 540172 | + | 7 |
| 3 | MNVALCPVTPATSRPLSETDTGFTRTEATPVA | ATPVTAAAFVAGIALGVAVCQAIGSQEPVIQ | 540244 | 540436 | + | 31 |
| 4 | MDMSDVSFLRRPGALTDIGIEDPEGRYTTAHAPVM | ATPAVASGVIIGYALVNAFVAGRDGGPIQPVPEIVLG | 540475 | 540694 | + | 29 |

### Results for WP_055540568.1 [Streptomyces neyagawaensis] back to top

previous - next

#### Architecture

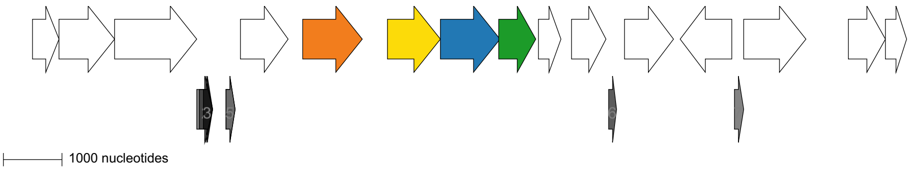

Link to nucleotide sequence

| Accession | start | end | direction | length (aa) | Pfam/HMM | name | description | E-value |
|---|---|---|---|---|---|---|---|---|
| WP_055540563.1 | 0 | 464 | + | 153 | PF00596<br>TIGR01086<br>TIGR03328<br>TIGR00760 | Aldolase_II<br>TIGR01086<br>TIGR03328<br>TIGR00760 | Class II Aldolase and Adducin N-terminal domain<br>fucA: L-fuculose phosphate aldolase<br>salvage_mtnB: methylthioribulose-1-phosphate dehydratase<br>araD: L-ribulose-5-phosphate 4-epimerase | 3.70E-26<br>1.30E-23<br>4.00E-10<br>4.70E-06 |
| WP_055540564.1 | 460 | 1402 | + | 313 | PF00294<br>TIGR02152<br>TIGR04382<br>TIGR03168<br>TIGR02198 | PfkB<br>TIGR02152<br>TIGR04382<br>TIGR03168<br>TIGR02198 | pfkB family carbohydrate kinase<br>D_ribokin_bact: ribokinase<br>myo_inos_iolC_N: 5-dehydro-2-deoxygluconokinase<br>1-PFK: hexose kinase, 1-phosphofructokinase family<br>rfaE_dom_I: bifunctional protein RfaE, domain I | 5.30E-44<br>6.40E-28<br>1.40E-27<br>2.50E-17<br>2.20E-16 |
| WP_055540565.1 | 1403 | 2807 | + | 467 | PF00171<br>TIGR01804<br>TIGR04284<br>TIGR01780<br>TIGR02299 | Aldedh<br>TIGR01804<br>TIGR04284<br>TIGR01780<br>TIGR02299 | Aldehyde dehydrogenase family<br>BADH: betaine-aldehyde dehydrogenase<br>aldehy_Rv0768: aldehyde dehydrogenase, Rv0768 family<br>SSADH: succinate-semialdehyde dehydrogenase<br>HpaE: 5-carboxymethyl-2-hydroxymuconate semialdehyde dehydrogenase | 1.50E-87<br>2.20E-55<br>9.20E-54<br>9.40E-54<br>1.20E-51 |
| WP_079055027.1 | 3557 | 4367 | + | 269 | PF00196<br>TIGR03020<br>TIGR03541<br>PF08281<br>TIGR02937 | GerE<br>TIGR03020<br>TIGR03541<br>Sigma70_r4_2<br>TIGR02937 | Bacterial regulatory proteins, luxR family<br>EpsA: transcriptional regulator EpsA<br>reg_near_HchA: LuxR family transcriptional regulatory, chaperone HchA-associated<br>Sigma-70, region 4<br>sigma70-ECF: RNA polymerase sigma factor, sigma-70 family | 1.30E-15<br>1.50E-07<br>3.40E-07<br>2.00E-06<br>1.10E-05 |
| WP_157873212.1 | 4618 | 5632 | + | 337 | TIGR01188<br>TIGR01288<br>TIGR03864<br>TIGR03522<br>TIGR00968 | TIGR01188<br>TIGR01288<br>TIGR03864<br>TIGR03522<br>TIGR00968 | drrA: daunorubicin resistance ABC transporter, ATP-binding protein<br>nodI: nodulation ABC transporter NodI<br>PQQ_ABC_ATP: ABC transporter, ATP-binding subunit, PQQ-dependent alcohol dehydrogenase system<br>GldA_ABC_ATP: gliding motility-associated ABC transporter ATP-binding subunit GldA<br>3a0106s01: sulfate ABC transporter, ATP-binding protein | 2.60E-86<br>5.40E-60<br>7.10E-59<br>5.20E-51<br>4.50E-41 |
| WP_055540567.1 | 6062 | 6959 | + | 298 | LinG<br>LinH | LinG<br>LinH | LinG Homolog<br>LinH (Fusion of LinE/LinG) | 8.10E-97<br>1.40E-91 |
| WP_055540568.1 | 6958 | 7975 | + | 338 | LinE<br>LinH<br>PF12697<br>PF00561<br>TIGR02427 | LinE<br>LinH<br>Abhydrolase_6<br>Abhydrolase_1<br>TIGR02427 | LinE Homolog<br>LinH (Fusion of LinE/LinG)<br>Alpha/beta hydrolase family<br>alpha/beta hydrolase fold<br>protocat_pcaD: 3-oxoadipate enol-lactonase | 1.80E-86<br>3.20E-74<br>4.60E-15<br>2.80E-11<br>6.80E-11 |
| WP_055540569.1 | 7971 | 8604 | + | 210 | LinL | LinL | LinL | 8.10E-50 |
| WP_063799259.1 | 8645 | 9029 | + | 127 | PF04134 | DUF393 | Protein of unknown function, DUF393 | 1.00E-17 |
| WP_157873213.1 | 9195 | 9801 | + | 201 | NO PFAM MATCH | - | - | - |
| WP_079055024.1 | 10115 | 10943 | + | 275 | PF01061<br>TIGR01247<br>TIGR03861 | ABC2_membrane<br>TIGR01247<br>TIGR03861 | ABC-2 type transporter<br>drrB: daunorubicin resistance ABC transporter membrane protein<br>phenyl_ABC_PedC: alcohol ABC transporter, permease protein | 4.70E-15<br>2.20E-07<br>8.00E-07 |
| WP_157873214.1 | 11942 | 11063 | - | 292 | TIGR03621<br>PF00296<br>TIGR03560<br>TIGR03856<br>TIGR03619 | TIGR03621<br>Bac_luciferase<br>TIGR03560<br>TIGR03856<br>TIGR03619 | F420_MSMEG_2516: probable F420-dependent oxidoreductase, MSMEG_2516 family<br>Luciferase-like monooxygenase<br>F420_Rv1855c: probable F420-dependent oxidoreductase, Rv1855c family<br>F420_MSMEG_2906: probable F420-dependent oxidoreductase, MSMEG_2906 family<br>F420_Rv2161c: probable F420-dependent oxidoreductase, Rv2161c family | 7.90E-61<br>7.60E-25<br>1.70E-23<br>1.10E-20<br>3.40E-19 |
| WP_055540574.1 | 12147 | 13197 | + | 349 | PF13469<br>PF00685 | Sulfotransfer_3<br>Sulfotransfer_1 | Sulfotransferase family<br>Sulfotransferase domain | 5.40E-20<br>2.90E-08 |

| | | | | | | | | |
|---|---|---|---|---|---|---|---|---|
| AOK14_RS44215 | 13935 | 14559 | + | 208 | INFERRED GENE | - | - | - |
| WP_055540575.1 | 14552 | 14915 | + | 120 | NO PFAM MATCH | - | - | - |

| index | leader | core | start | end | dir | score |
|---|---|---|---|---|---|---|
| 1 | MLRCWLRSGYGRCVMVLRARPGRDE...VIHRVGGEGRGAADARASQCPPPRA | RPPPFPGVQGTRATAALRRASADATRANSTAQAGLSY | 2814 | 3090 | + | 7 |
| 2 | MVLRARPGRDEGEQAVIHRVGGEGRGAADARASQCPPPRA | RPPPFPGVQGTRATAALRRASADATRANSTAQAGLSY | 2856 | 3090 | + | 7 |
| 3 | VGGEGRGAADARASQCPPPRA | RPPPFPGVQGTRATAALRRASADATRANSTAQAGLSY | 2913 | 3090 | + | 8 |
| 4 | LNMSGVGAVCKNTVIPSPWW | APPCWASVQEAAIQPDPAPEETTDKPKRLYTPR | 3304 | 3466 | + | 6 |
| 5 | MSGVGAVCKNTVIPSPWW | APPCWASVQEAAIQPDPAPEETTDKPKRLYTPR | 3310 | 3466 | + | 6 |
| 6 | VISGSPEDPRGYGFRTADASSHSSCLPLR | RPPPRASMTPTRSSLTDTSRT | 9829 | 9982 | + | 8 |
| 7 | MLLADRLRLPGGNQHVGADYSGTFDP | VISPAGRLASSGTWATCMRRGQRMVTT | 11989 | 12151 | + | 7 |

### Results for WP_078073573.1 [Streptomyces niveus] back to top

previous - next

#### Architecture

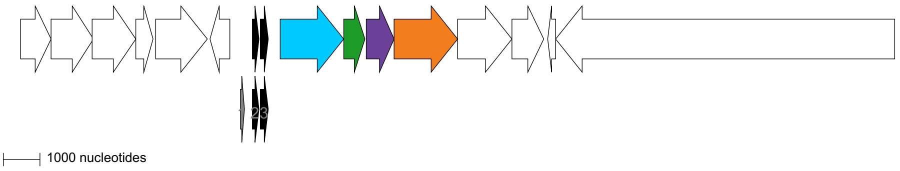

Link to nucleotide sequence

| Accession | start | end | direction | length (aa) | Pfam/HMM | name | description | E-value |
|---|---|---|---|---|---|---|---|---|
| WP_078073565.1 | 295932 | 296745 | + | 270 | NO PFAM MATCH | - | - | - |
| WP_078073566.1 | 296741 | 297914 | + | 390 | PF00441<br>PF02770<br>TIGR03207<br>TIGR03204<br>PF02771 | Acyl-CoA_dh_1<br>Acyl-CoA_dh_M<br>TIGR03207<br>TIGR03204<br>Acyl-CoA_dh_N | Acyl-CoA dehydrogenase, C-terminal domain<br>Acyl-CoA dehydrogenase, middle domain<br>cyc_hxne_CoA_dh: cyclohexanecarboxyl-CoA dehydrogenase<br>pimC_large: pimeloyl-CoA dehydrogenase, large subunit<br>Acyl-CoA dehydrogenase, N-terminal domain | 2.90E-21<br>2.50E-17<br>1.60E-15<br>2.20E-10<br>1.00E-06 |
| WP_078073567.1 | 297894 | 299067 | + | 390 | PF01648<br>TIGR00556 | ACPS<br>TIGR00556 | 4'-phosphopantetheinyl transferase superfamily<br>pantethn_trn: phosphopantetheine--protein transferase domain | 4.00E-06<br>9.80E-04 |
| WP_078073568.1 | 299063 | 299528 | + | 154 | NO PFAM MATCH | - | - | - |
| WP_078073569.1 | 299598 | 301011 | + | 470 | TIGR00900<br>PF07690<br>TIGR00881<br>PF05977<br>TIGR00880 | TIGR00900<br>MFS_1<br>TIGR00881<br>MFS_3<br>TIGR00880 | 2A0121: H+ Antiporter protein<br>Major Facilitator Superfamily<br>2A0104: phosphoglycerate transporter family protein<br>Transmembrane secretion effector<br>2_A_01_02: multidrug resistance protein | 5.00E-37<br>3.60E-29<br>3.10E-13<br>3.70E-13<br>4.40E-12 |
| WP_078073570.1 | 301649 | 301112 | - | 178 | PF12840<br>PF01022<br>PF13518<br>PF13338 | HTH_20<br>HTH_5<br>HTH_28<br>AbiEi_4 | Helix-turn-helix domain<br>Bacterial regulatory protein, arsR family<br>Helix-turn-helix domain<br>Transcriptional regulator, AbiEi antitoxin | 1.50E-11<br>2.20E-06<br>2.70E-04<br>6.80E-04 |
| WP_078073571.1 | 302265 | 302457 | + | 63 | LinA | LinA | LinA (linaridin precursor) | 6.40E-15 |
| WP_078073572.1 | 302495 | 302687 | + | 63 | LinA | LinA | LinA (linaridin precursor) | 6.10E-09 |
| WP_078073573.1 | 303032 | 304766 | + | 577 | LinH<br>LinE<br>LinG<br>PF12697<br>PF00561 | LinH<br>LinE<br>LinG<br>Abhydrolase_6<br>Abhydrolase_1 | LinH (Fusion of LinE/LinG)<br>LinE Homolog<br>LinG Homolog<br>Alpha/beta hydrolase family<br>alpha/beta hydrolase fold | 1.70E-217<br>4.20E-98<br>6.00E-92<br>2.30E-15<br>8.30E-12 |
| WP_078073574.1 | 304762 | 305341 | + | 192 | LinL | LinL | LinL | 2.30E-64 |
| WP_078073575.1 | 305386 | 306145 | + | 252 | PF13649<br>PF08241<br>PF08242<br>PF13489<br>PF13847 | Methyltransf_25<br>Methyltransf_11<br>Methyltransf_12<br>Methyltransf_23<br>Methyltransf_31 | Methyltransferase domain<br>Methyltransferase domain<br>Methyltransferase domain<br>Methyltransferase domain<br>Methyltransferase domain | 1.90E-14<br>2.80E-09<br>1.00E-08<br>3.90E-08<br>6.20E-08 |
| WP_078073576.1 | 306141 | 307884 | + | 580 | TIGR02204<br>TIGR02203<br>TIGR00958<br>TIGR03796<br>TIGR01846 | TIGR02204<br>TIGR02203<br>TIGR00958<br>TIGR03796<br>TIGR01846 | MsbA_rel: ABC transporter, permease/ATP-binding protein<br>MsbA_lipidA: lipid A export permease/ATP-binding protein MsbA<br>3a01208: antigen peptide transporter 2<br>NHLM_micro_ABC1: NHLM bacteriocin system ABC transporter, peptidase/ATP-binding protein<br>type_I_sec_HlyB: type I secretion system ATPase | 7.50E-105<br>3.50E-98<br>5.30E-89<br>5.40E-83<br>1.40E-81 |
| WP_078073577.1 | 307880 | 309365 | + | 494 | TIGR02734<br>TIGR01317<br>TIGR00292<br>TIGR03467<br>PF13450 | TIGR02734<br>TIGR01317<br>TIGR00292<br>TIGR03467<br>NAD_binding_8 | crtI_fam: phytoene desaturase<br>GOGAT_sm_gam: glutamate synthase, NADH/NADPH, small subunit<br>TIGR00292: thiazole biosynthesis enzyme<br>HpnE: squalene-associated FAD-dependent desaturase<br>NAD(P)-binding Rossmann-like domain | 2.40E-12<br>5.20E-09<br>2.00E-08<br>4.40E-08<br>5.70E-08 |
| WP_078073578.1 | 309361 | 310213 | + | 283 | PF12680 | SnoaL_2 | SnoaL-like domain | 2.40E-07 |
| WP_078073579.1 | 310542 | 310323 | - | 72 | PF03621 | MbtH | MbtH-like protein | 1.00E-28 |

| | | | | | | | | |
|---|---|---|---|---|---|---|---|---|
| WP_159392369.1 | 319838 | 310538 | - | 3099 | TIGR01733<br>PF00668<br>PF00501<br>TIGR01734<br>TIGR03443 | TIGR01733<br>Condensation<br>AMP-binding<br>TIGR01734<br>TIGR03443 | AA-adenyl-dom: amino acid adenylation domain<br>Condensation domain<br>AMP-binding enzyme<br>D-ala-DACP-lig: D-alanine--poly(phosphoribitol) ligase, subunit 1<br>alpha_am_amid: L-aminoadipate-semialdehyde dehydrogenase | 4.30E-246<br>8.40E-203<br>1.40E-178<br>2.40E-127<br>2.40E-111 |

| index | leader | core | start | end | dir | score |
|---|---|---|---|---|---|---|
| 1 | MAHPSIYFSRHGEMMGTER | MRRTPPPVSSVMCTTSVKA | 301939 | 302056 | + | 7 |
| 2 | MATIAEFMNDQKLGVLELGYKEDSQEPVL | ATPAAAVATCWYVGGAAAGVTVVAGAYVAGRVFG | 302265 | 302457 | + | 30 |
| 3 | MSRIIESVAAITPEHLTDGSHGDVVAMAPVQ | ATPGLVAFGIGFAGGAGVSWVGVQAYEAGAND | 302495 | 302687 | + | 31 |

### Results for AQU70555.1 [Streptomyces niveus] back to top

previous - next

#### Architecture

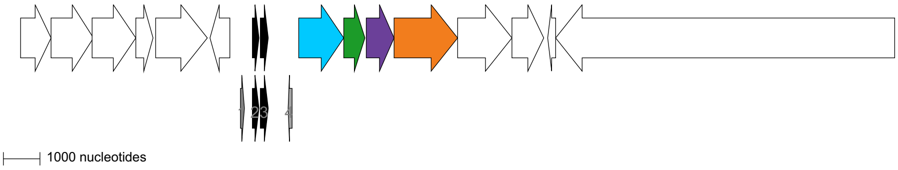

Link to nucleotide sequence

| Accession | start | end | direction | length (aa) | Pfam/HMM | name | description | E-value |
|---|---|---|---|---|---|---|---|---|
| AQU65083.1 | 295932 | 296745 | + | 270 | NO PFAM MATCH | - | - | - |
| AQU65084.1 | 296741 | 297914 | + | 390 | PF00441<br>PF02770<br>TIGR03207<br>TIGR03204<br>PF02771 | Acyl-CoA_dh_1<br>Acyl-CoA_dh_M<br>TIGR03207<br>TIGR03204<br>Acyl-CoA_dh_N | Acyl-CoA dehydrogenase, C-terminal domain<br>Acyl-CoA dehydrogenase, middle domain<br>cyc_hxne_CoA_dh: cyclohexanecarboxyl-CoA dehydrogenase<br>pimC_large: pimeloyl-CoA dehydrogenase, large subunit<br>Acyl-CoA dehydrogenase, N-terminal domain | 2.90E-21<br>2.50E-17<br>1.60E-15<br>2.20E-10<br>1.00E-06 |
| AQU65085.1 | 297894 | 299067 | + | 390 | PF01648<br>TIGR00556 | ACPS<br>TIGR00556 | 4'-phosphopantetheinyl transferase superfamily<br>pantethn_trn: phosphopantetheine--protein transferase domain | 4.00E-06<br>9.80E-04 |
| AQU65086.1 | 299063 | 299528 | + | 154 | NO PFAM MATCH | - | - | - |
| AQU65087.1 | 299598 | 301011 | + | 470 | TIGR00900<br>PF07690<br>TIGR00881<br>PF05977<br>TIGR00880 | TIGR00900<br>MFS_1<br>TIGR00881<br>MFS_3<br>TIGR00880 | 2A0121: H+ Antiporter protein<br>Major Facilitator Superfamily<br>2A0104: phosphoglycerate transporter family protein<br>Transmembrane secretion effector<br>2_A_01_02: multidrug resistance protein | 5.00E-37<br>3.60E-29<br>3.10E-13<br>3.70E-13<br>4.40E-12 |
| AQU65088.1 | 301649 | 301112 | - | 178 | PF12840<br>PF01022<br>PF13518<br>PF13338 | HTH_20<br>HTH_5<br>HTH_28<br>AbiEi_4 | Helix-turn-helix domain<br>Bacterial regulatory protein, arsR family<br>Helix-turn-helix domain<br>Transcriptional regulator, AbiEi antitoxin | 1.50E-11<br>2.20E-06<br>2.70E-04<br>6.80E-04 |
| AQU65089.1 | 302265 | 302457 | + | 63 | LinA | LinA | LinA (linaridin precursor) | 6.40E-15 |
| AQU65090.1 | 302495 | 302687 | + | 63 | LinA | LinA | LinA (linaridin precursor) | 6.10E-09 |
| AQU70555.1 | 303524 | 304766 | + | 413 | LinH<br>LinE<br>LinG<br>PF12697<br>PF00561 | LinH<br>LinE<br>LinG<br>Abhydrolase_6<br>Abhydrolase_1 | LinH (Fusion of LinE/LinG)<br>LinE Homolog<br>LinG Homolog<br>Alpha/beta hydrolase family<br>alpha/beta hydrolase fold | 6.40E-145<br>1.10E-98<br>1.10E-29<br>8.50E-16<br>4.80E-12 |
| AQU65091.1 | 304762 | 305341 | + | 192 | LinL | LinL | LinL | 2.30E-64 |
| AQU65092.1 | 305374 | 306145 | + | 256 | PF13649<br>PF08241<br>PF08242<br>PF13489<br>PF13847 | Methyltransf_25<br>Methyltransf_11<br>Methyltransf_12<br>Methyltransf_23<br>Methyltransf_31 | Methyltransferase domain<br>Methyltransferase domain<br>Methyltransferase domain<br>Methyltransferase domain<br>Methyltransferase domain | 2.00E-14<br>3.00E-09<br>1.00E-08<br>4.00E-08<br>6.40E-08 |
| AQU65093.1 | 306141 | 307884 | + | 580 | TIGR02204<br>TIGR02203<br>TIGR00958<br>TIGR03796<br>TIGR01846 | TIGR02204<br>TIGR02203<br>TIGR00958<br>TIGR03796<br>TIGR01846 | MsbA_rel: ABC transporter, permease/ATP-binding protein<br>MsbA_lipidA: lipid A export permease/ATP-binding protein MsbA<br>3a01208: antigen peptide transporter 2<br>NHLM_micro_ABC1: NHLM bacteriocin system ABC transporter, peptidase/ATP-binding protein<br>type_I_sec_HlyB: type I secretion system ATPase | 7.50E-105<br>3.50E-98<br>5.30E-89<br>5.40E-83<br>1.40E-81 |
| AQU65094.1 | 307880 | 309365 | + | 494 | TIGR02734<br>TIGR01317<br>TIGR00292<br>TIGR03467<br>PF13450 | TIGR02734<br>TIGR01317<br>TIGR00292<br>TIGR03467<br>NAD_binding_8 | crtI_fam: phytoene desaturase<br>GOGAT_sm_gam: glutamate synthase, NADH/NADPH, small subunit<br>TIGR00292: thiazole biosynthesis enzyme<br>HpnE: squalene-associated FAD-dependent desaturase<br>NAD(P)-binding Rossmann-like domain | 2.40E-12<br>5.20E-09<br>2.00E-08<br>4.40E-08<br>5.70E-08 |
| AQU65095.1 | 309361 | 310213 | + | 283 | PF12680 | SnoaL_2 | SnoaL-like domain | 2.40E-07 |
| AQU65096.1 | 310542 | 310323 | - | 72 | PF03621 | MbtH | MbtH-like protein | 1.00E-28 |
| AQU65097.1 | 319838 | 310538 | - | 3099 | TIGR01733<br>PF00668<br>PF00501<br>TIGR01734<br>TIGR03443 | TIGR01733<br>Condensation<br>AMP-binding<br>TIGR01734<br>TIGR03443 | AA-adenyl-dom: amino acid adenylation domain<br>Condensation domain<br>AMP-binding enzyme<br>D-ala-DACP-lig: D-alanine--poly(phosphoribitol) ligase, subunit 1<br>alpha_am_amid: L-aminoadipate-semialdehyde dehydrogenase | 4.30E-246<br>8.40E-203<br>1.40E-178<br>2.40E-127<br>2.40E-111 |

| index | leader | core | start | end | dir | score |
|---|---|---|---|---|---|---|
| 1 | MAHPSIYFSRHGEMMGTER | MRRTPPPVSSVMCTTSVKA | 301939 | 302056 | + | 7 |

| | | | | | | |
|---|---|---|---|---|---|---|
| 2 | MATIAEFMNDQKLGVLELGYKEDSQEPVL | ATPAAAVATCWYVGGAAAGVTVVAGAYVAGRVFG | 302265 | 302457 | + | 30 |
| 3 | MSRIIESVAAITPEHLTDGSHGDVVAMAPVQ | ATPGLVAFGIGFAGGAGVSWVGVQAYEAGAND | 302495 | 302687 | + | 31 |
| 4 | MLAWIARYHIRASLTCG | LTPSRASPVTAWTWVAT | 303358 | 303253 | - | 6 |

### Results for WP_130798439.1 [Streptomyces otsuchiensis] back to top

previous - next

#### Architecture

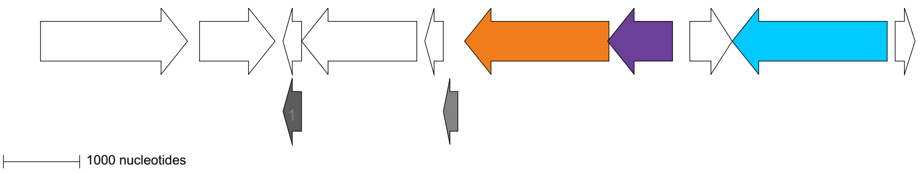

Link to nucleotide sequence

| Accession | start | end | direction | length (aa) | Pfam/HMM | name | description | E-value |
|---|---|---|---|---|---|---|---|---|
| WP_170214973.1 | 5584 | 7504 | + | 639 | PF13454 | NAD_binding_9 | FAD-NAD(P)-binding | 4.80E-41 |
| WP_130798443.1 | 7668 | 8649 | + | 326 | NO PFAM MATCH | - | - | - |
| WP_130798429.1 | 9004 | 8755 | - | 82 | NO PFAM MATCH | - | - | - |
| WP_130798431.1 | 10521 | 9006 | - | 504 | TIGR02734<br>TIGR03467<br>TIGR00292<br>PF13450<br>PF01946 | TIGR02734<br>TIGR03467<br>TIGR00292<br>NAD_binding_8<br>Thi4 | crtI_fam: phytoene desaturase<br>HpnE: squalene-associated FAD-dependent desaturase<br>TIGR00292: thiazole biosynthesis enzyme<br>NAD(P)-binding Rossmann-like domain<br>Thi4 family | 2.80E-12<br>1.10E-09<br>2.30E-09<br>4.50E-09<br>6.00E-08 |
| WP_130798433.1 | 10867 | 10618 | - | 82 | NO PFAM MATCH | - | - | - |
| WP_130798435.1 | 13031 | 11138 | - | 630 | TIGR02204<br>TIGR02203<br>TIGR00958<br>TIGR03796<br>TIGR01846 | TIGR02204<br>TIGR02203<br>TIGR00958<br>TIGR03796<br>TIGR01846 | MsbA_rel: ABC transporter, permease/ATP-binding protein<br>MsbA_lipidA: lipid A export permease/ATP-binding protein MsbA<br>3a01208: antigen peptide transporter 2<br>NHLM_micro_ABC1: NHLM bacteriocin system ABC transporter, peptidase/ATP-binding protein<br>type_I_sec_HlyB: type I secretion system ATPase | 6.20E-101<br>4.90E-94<br>6.60E-87<br>1.10E-81<br>9.50E-81 |
| WP_130798437.1 | 13866 | 13008 | - | 285 | PF13649<br>PF13489<br>PF08241<br>PF08242<br>PF13847 | Methyltransf_25<br>Methyltransf_23<br>Methyltransf_11<br>Methyltransf_12<br>Methyltransf_31 | Methyltransferase domain<br>Methyltransferase domain<br>Methyltransferase domain<br>Methyltransferase domain<br>Methyltransferase domain | 6.30E-14<br>5.60E-11<br>2.20E-08<br>3.10E-08<br>9.30E-07 |
| WP_019435954.1 | 14084 | 14651 | + | 189 | INFERRED GENE | - | - | - |
| WP_130798439.1 | 16675 | 14647 | - | 675 | LinH<br>LinG<br>LinE<br>PF12697<br>PF00561 | LinH<br>LinG<br>LinE<br>Abhydrolase_6<br>Abhydrolase_1 | LinH (Fusion of LinE/LinG)<br>LinG Homolog<br>LinE Homolog<br>Alpha/beta hydrolase family<br>alpha/beta hydrolase fold | 5.00E-254<br>2.00E-115<br>1.40E-97<br>1.20E-16<br>7.00E-11 |
| WP_170214974.1 | 16781 | 17049 | + | 89 | PF13340 | DUF4096 | Putative transposase of IS4/5 family (DUF4096) | 2.20E-24 |

| index | leader | core | start | end | dir | score |
|---|---|---|---|---|---|---|
| 1 | MSEITMELVYGGAGPPDGEVAVVRLLGRLPGGWHYEQQHIGPVLIQLRI | MAPLVGPADIEGAVVSALGDPALLNWRVLTADG | 9004 | 8755 | - | 8 |
| 2 | VYCQRFPFSRNPFSGIFMTPAKKPAE | ATPGIRNRTWGNRGIRIEFTRQRPEGLSCGTFSR | 11046 | 10863 | - | 7 |

### Results for PNE36149.1 [Streptomyces noursei] back to top

previous - next

#### Architecture

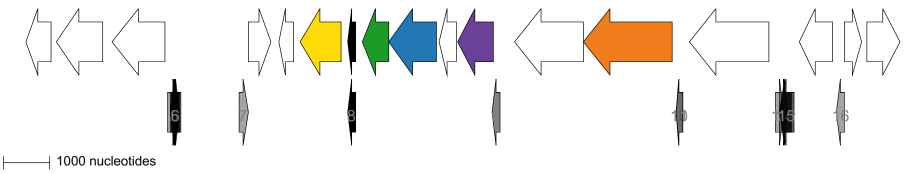

Link to nucleotide sequence

| Accession | start | end | direction | length (aa) | Pfam/HMM | name | description | E-value |
|---|---|---|---|---|---|---|---|---|
| PNE36821.1 | 435399 | 434859 | - | 179 | TIGR01203<br>PF00156<br>TIGR01251<br>TIGR01367 | TIGR01203<br>Pribosyltran<br>TIGR01251<br>TIGR01367 | HGPRTase: hypoxanthine phosphoribosyltransferase<br>Phosphoribosyl transferase domain<br>ribP_PPkin: ribose-phosphate diphosphokinase<br>pyrE_Therm: orotate phosphoribosyltransferase | 4.50E-65<br>1.40E-25<br>1.30E-05<br>5.40E-04 |
| PNE36142.1 | 436521 | 435513 | - | 335 | TIGR02432<br>PF01171<br>PF09179 | TIGR02432<br>ATP_bind_3<br>TilS | lysidine_TilS_N: tRNA(Ile)-lysidine synthetase<br>PP-loop family<br>TilS substrate binding domain | 3.10E-43<br>5.20E-38<br>3.00E-12 |

| | | | | | | | | |
|---|---|---|---|---|---|---|---|---|
| PNE36143.1 | 437860 | 436714 | - | 381 | TIGR03883 | TIGR03883 | DUF2342_F420: uncharacterized protein, coenzyme F420 biosynthesis associated | 8.30E-141 |
| | | | | | TIGR03624 | TIGR03624 | TIGR03624: putative hydrolase | 2.40E-125 |
| | | | | | PF10103 | Zincin_2 | Zincin-like metallopeptidase | 1.20E-120 |
| PNE36144.1 | 439670 | 440162 | + | 163 | PF00719 | Pyrophosphatase | Inorganic pyrophosphatase | 4.50E-53 |
| PNE36145.1 | 440637 | 440310 | - | 108 | NO PFAM MATCH | - | - | - |
| PNE36146.1 | 441651 | 440790 | - | 286 | LinG | LinG | LinG Homolog | 2.10E-108 |
| | | | | | LinH | LinH | LinH (Fusion of LinE/LinG) | 1.90E-76 |
| | | | | | TIGR04033 | TIGR04033 | export_SdpB: antimicrobial peptide system protein, SdpB family | 2.00E-04 |
| PNE36147.1 | 441984 | 441795 | - | 62 | LinA | LinA | LinA (linaridin precursor) | 2.10E-11 |
| PNE36148.1 | 442703 | 442121 | - | 193 | LinL | LinL | LinL | 1.50E-57 |
| PNE36149.1 | 443722 | 442699 | - | 340 | LinE | LinE | LinE Homolog | 1.00E-102 |
| | | | | | LinH | LinH | LinH (Fusion of LinE/LinG) | 8.20E-53 |
| | | | | | PF12697 | Abhydrolase_6 | Alpha/beta hydrolase family | 4.10E-15 |
| | | | | | TIGR02427 | TIGR02427 | protocat_pcaD: 3-oxoadipate enol-lactonase | 8.30E-11 |
| | | | | | PF00561 | Abhydrolase_1 | alpha/beta hydrolase fold | 8.90E-09 |
| PNE36150.1 | 444142 | 443794 | - | 115 | PF01527 | HTH_Tnp_1 | Transposase | 7.00E-08 |
| | | | | | PF13518 | HTH_28 | Helix-turn-helix domain | 2.10E-07 |
| | | | | | PF13384 | HTH_23 | Homeodomain-like domain | 4.80E-06 |
| | | | | | PF05344 | DUF746 | Domain of Unknown Function (DUF746) | 1.10E-05 |
| | | | | | PF13551 | HTH_29 | Winged helix-turn helix | 3.90E-05 |
| PNE36151.1 | 444939 | 444180 | - | 252 | PF13649 | Methyltransf_25 | Methyltransferase domain | 7.00E-16 |
| | | | | | PF08241 | Methyltransf_11 | Methyltransferase domain | 9.60E-13 |
| | | | | | PF13489 | Methyltransf_23 | Methyltransferase domain | 8.80E-10 |
| | | | | | PF08242 | Methyltransf_12 | Methyltransferase domain | 6.60E-09 |
| | | | | | PF13847 | Methyltransf_31 | Methyltransferase domain | 1.10E-08 |
| PNE36152.1 | 446888 | 445403 | - | 494 | TIGR03467 | TIGR03467 | HpnE: squalene-associated FAD-dependent desaturase | 1.60E-09 |
| | | | | | PF13450 | NAD_binding_8 | NAD(P)-binding Rossmann-like domain | 1.30E-08 |
| | | | | | TIGR00292 | TIGR00292 | TIGR00292: thiazole biosynthesis enzyme | 2.10E-08 |
| | | | | | TIGR02734 | TIGR02734 | crtI_fam: phytoene desaturase | 3.20E-08 |
| | | | | | TIGR03315 | TIGR03315 | Se_ygfK: putative selenate reductase, YgfK subunit | 1.10E-06 |
| PNE36153.1 | 448822 | 446884 | - | 645 | TIGR02204 | TIGR02204 | MsbA_rel: ABC transporter, permease/ATP-binding protein | 3.10E-97 |
| | | | | | TIGR02203 | TIGR02203 | MsbA_lipidA: lipid A export permease/ATP-binding protein MsbA | 1.20E-94 |
| | | | | | TIGR00958 | TIGR00958 | 3a01208: antigen peptide transporter 2 | 1.40E-88 |
| | | | | | TIGR03796 | TIGR03796 | NHLM_micro_ABC1: NHLM bacteriocin system ABC transporter, peptidase/ATP-binding protein | 7.50E-81 |
| | | | | | TIGR01846 | TIGR01846 | type_I_sec_HlyB: type I secretion system ATPase | 1.40E-77 |
| PNE36154.1 | 450908 | 449195 | - | 570 | PF06738 | ThrE | Putative threonine/serine exporter | 4.50E-73 |
| | | | | | PF12821 | ThrE_2 | Threonine/Serine exporter, ThrE | 2.70E-20 |
| PNE36155.1 | 452262 | 451551 | - | 236 | PF09335 | SNARE_assoc | SNARE associated Golgi protein | 2.00E-24 |
| PNE36156.1 | 452522 | 452891 | + | 122 | PF01906 | YbjQ_1 | Putative heavy-metal-binding | 1.30E-33 |
| | | | | | TIGR03884 | TIGR03884 | sel_bind_Methan: selenium-binding protein | 7.80E-04 |
| PNE36157.1 | 453016 | 453730 | + | 237 | PF01810 | LysE | LysE type translocator | 8.80E-43 |
| | | | | | TIGR00949 | TIGR00949 | 2A76: homoserine/Threonine efflux protein | 6.50E-40 |
| | | | | | TIGR00948 | TIGR00948 | 2a75: L-lysine exporter | 4.90E-14 |

| index | leader | core | start | end | dir | score |
|---|---|---|---|---|---|---|
| 1 | LAPGGRCAGGEDGGWAPGGWVVQRQ...RAFWAAWGSVVPVVMKAKARRRPSA | STTVPARLLTPVRVPVLARTVPAPAVSPT | 437909 | 438182 | + | 6 |
| 2 | LATVEAIRSRAFWAAWGSVVPVVMKAKARRRPSA | STTVPARLLTPVRVPVLARTVPAPAVSPT | 437990 | 438182 | + | 6 |
| 3 | VEAIRSRAFWAAWGSVVPVVMKAKARRRPSA | STTVPARLLTPVRVPVLARTVPAPAVSPT | 437999 | 438182 | + | 6 |
| 4 | VRAKTGTLTGVNSLAGTVVDADGRLLAFAF | MTTGTTDPQAAQKALDRMASTVANCGCR | 438151 | 437974 | - | 7 |
| 5 | VGDTAGAGTVRAKTGTLTGVNSLAGTVVDADGRLLAFAF | MTTGTTDPQAAQKALDRMASTVANCGCR | 438178 | 437974 | - | 9 |
| 6 | VAAFTGTLRGRYVGDTAGAGTVRAK...GTLTGVNSLAGTVVDADGRLLAFAF | MTTGTTDPQAAQKALDRMASTVANCGCR | 438214 | 437974 | - | 7 |
| 7 | MRGALCGPCRGACGTPRPACRHPRGTLRRLPCDLPCSGLGHFGPAPF | ATTHLRGGHLITRRMC | 439461 | 439653 | + | 6 |
| 8 | MAEFANTQLVDVKPGRIGSDAAPTM | MTPLATVATPEAVSAGIAVTSATAAALNMVTHNVQHG | 441984 | 441795 | - | 34 |
| 9 | LRRFHWIDRRCWSDQPCA | TTAVSPAVVLAYDVLPCSGSSASRRPPRDRMWGQSEGVR | 445109 | 444935 | - | 7 |
| 10 | VFLSGTAPSGQQTRLLTPAGPAGHSTG | QGVPANQLTRNLTPKGVLVSAILTTKL | 449051 | 448886 | - | 6 |
| 11 | VVFLSGTAPSGQQTRLLTPAGPAGHST | GQGVPANQLTRNLTPKGVLVSAILTTKL | 449054 | 448886 | - | 6 |
| 12 | LSAACPPDGGCCVVRCAGLLGRGDGPAGVVCRTAALYVLH | TTPSGPSPPVGGWERR | 451030 | 451201 | + | 6 |
| 13 | VQDCCALRPAHHPLRPVTSRGWVGKTVSGADASGPLPVI | ATPRRTRRGPPRRTPTA | 451119 | 451290 | + | 6 |
| 14 | VPGLWGGGCADCCALRP | HTPHPTDPSPPVDGWVSGRRGTPGW...GVGPTHRLSHPPTGGDGPEGVVCRT | 451389 | 451137 | - | 9 |
| 15 | MSAACPPDGGGASSAARGCSAAVPGLWGGGCADCCALRP | HTPHPTDPSPPVDGWVSGRRGTPGW...GVGPTHRLSHPPTGGDGPEGVVCRT | 451455 | 451137 | - | 8 |
| 16 | MPMVRTSCPCGQDHPGRPERPALGGGRVG | VGGRRRGGAGRPGRPWPGVFLEPRDVIVR | 452530 | 452353 | - | 6 |

### Results for PNE39469.1 [Streptomyces noursei] back to top

previous - next

#### Architecture

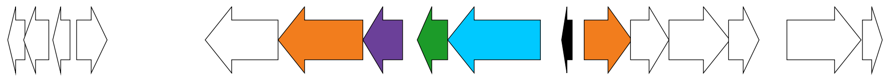

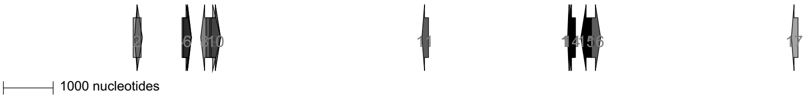

Link to nucleotide sequence

| Accession | start | end | direction | length (aa) | Pfam/HMM | name | description | E-value |
|---|---|---|---|---|---|---|---|---|
| PNE37321.1 | 686128 | 685801 | - | 108 | PF13601<br>PF12802<br>PF13412 | HTH_34<br>MarR_2<br>HTH_24 | Winged helix DNA-binding domain<br>MarR family<br>Winged helix-turn-helix DNA-binding | 5.00E-22<br>2.20E-04<br>4.50E-04 |
| PNE37322.1 | 686631 | 686124 | - | 168 | NO PFAM MATCH | - | - | - |
| PNE37323.1 | 687056 | 686687 | - | 122 | NO PFAM MATCH | - | - | - |
| PNE39468.1 | 687184 | 687787 | + | 200 | PF06889 | DUF1266 | Protein of unknown function (DUF1266) | 9.20E-27 |
| PNE37324.1 | 691221 | 689766 | - | 484 | TIGR02734<br>PF13450<br>TIGR00292<br>TIGR02733<br>TIGR03467 | TIGR02734<br>NAD_binding_8<br>TIGR00292<br>TIGR02733<br>TIGR03467 | crtI_fam: phytoene desaturase<br>NAD(P)-binding Rossmann-like domain<br>TIGR00292: thiazole biosynthesis enzyme<br>desat_CrtD: C-3',4' desaturase CrtD<br>HpnE: squalene-associated FAD-dependent desaturase | 1.00E-10<br>1.40E-08<br>7.20E-08<br>1.50E-07<br>2.50E-07 |
| PNE37325.1 | 692924 | 691217 | - | 568 | TIGR02204<br>TIGR02203<br>TIGR00958<br>TIGR01846<br>TIGR03375 | TIGR02204<br>TIGR02203<br>TIGR00958<br>TIGR01846<br>TIGR03375 | MsbA_rel: ABC transporter, permease/ATP-binding protein<br>MsbA_lipidA: lipid A export permease/ATP-binding protein MsbA<br>3a01208: antigen peptide transporter 2<br>type_I_sec_HlyB: type I secretion system ATPase<br>type_I_sec_LssB: type I secretion system ATPase | 6.10E-104<br>1.20E-98<br>4.20E-90<br>5.60E-87<br>1.40E-84 |
| PNE37326.1 | 693712 | 692920 | - | 263 | PF13649<br>PF08241<br>PF08242<br>PF13489<br>PF13847 | Methyltransf_25<br>Methyltransf_11<br>Methyltransf_12<br>Methyltransf_23<br>Methyltransf_31 | Methyltransferase domain<br>Methyltransferase domain<br>Methyltransferase domain<br>Methyltransferase domain<br>Methyltransferase domain | 3.30E-16<br>7.00E-12<br>1.00E-08<br>1.20E-07<br>3.00E-07 |
| PNE37327.1 | 694617 | 694005 | - | 203 | LinL | LinL | LinL | 9.60E-68 |
| PNE39469.1 | 696485 | 694613 | - | 623 | LinH<br>LinG<br>LinE<br>PF12697<br>PF12146 | LinH<br>LinG<br>LinE<br>Abhydrolase_6<br>Hydrolase_4 | LinH (Fusion of LinE/LinG)<br>LinG Homolog<br>LinE Homolog<br>Alpha/beta hydrolase family<br>Serine aminopeptidase, S33 | 2.10E-249<br>6.20E-127<br>7.40E-99<br>6.70E-16<br>1.60E-07 |
| PNE37328.1 | 697117 | 696916 | - | 66 | LinA | LinA | LinA (linaridin precursor) | 1.80E-13 |
| PNE37329.1 | 697356 | 698310 | + | 317 | TIGR01188<br>TIGR03864<br>TIGR01288<br>TIGR01257<br>TIGR03740 | TIGR01188<br>TIGR03864<br>TIGR01288<br>TIGR01257<br>TIGR03740 | drrA: daunorubicin resistance ABC transporter, ATP-binding protein<br>PQQ_ABC_ATP: ABC transporter, ATP-binding subunit, PQQ-dependent alcohol dehydrogenase system<br>nodI: nodulation ABC transporter NodI<br>rim_protein: rim ABC transporter<br>galliderm_ABC: lantibiotic protection ABC transporter, ATP-binding subunit | 9.90E-63<br>6.60E-44<br>1.10E-41<br>1.10E-32<br>1.60E-32 |
| PNE37330.1 | 698306 | 699074 | + | 255 | PF01061<br>PF12698<br>TIGR03062<br>TIGR01247<br>TIGR01291 | ABC2_membrane<br>ABC2_membrane_3<br>TIGR03062<br>TIGR01247<br>TIGR01291 | ABC-2 type transporter<br>ABC-2 family transporter protein<br>pip_yhgE_Cterm: YhgE/Pip C-terminal domain<br>drrB: daunorubicin resistance ABC transporter membrane protein<br>nodJ: ABC-2 type transporter, NodJ family | 6.80E-12<br>2.40E-09<br>2.90E-06<br>3.90E-05<br>1.90E-04 |
| PNE37331.1 | 699070 | 700282 | + | 403 | PF07730 | HisKA_3 | Histidine kinase | 7.50E-15 |
| PNE37332.1 | 700278 | 700884 | + | 201 | PF00072<br>TIGR02875<br>PF00196<br>TIGR02154<br>TIGR02956 | Response_reg<br>TIGR02875<br>GerE<br>TIGR02154<br>TIGR02956 | Response regulator receiver domain<br>spore_0_A: sporulation transcription factor Spo0A<br>Bacterial regulatory proteins, luxR family<br>PhoB: phosphate regulon transcriptional regulatory protein PhoB<br>TMAO_torS: TMAO reductase sytem sensor TorS | 6.50E-22<br>2.50E-16<br>7.20E-13<br>3.00E-08<br>4.70E-08 |
| PNE39470.1 | 701433 | 702927 | + | 497 | PF00135<br>PF07859 | COesterase<br>Abhydrolase_3 | Carboxylesterase family<br>alpha/beta hydrolase fold | 1.40E-85<br>2.80E-11 |
| PNE37333.1 | 702963 | 703359 | + | 131 | PF01042<br>TIGR00004<br>TIGR03610 | Ribonuc_L-PSP<br>TIGR00004<br>TIGR03610 | Endoribonuclease L-PSP<br>TIGR00004: reactive intermediate/imine deaminase<br>RutC: pyrimidine utilization protein C | 1.50E-22<br>2.20E-09<br>5.00E-06 |

| index | leader | core | start | end | dir | score |
|---|---|---|---|---|---|---|
| 1 | VRLPGPFSLDSRSRRSEVDSMDPCHCRRVQ | GAVAGWSSSVARWAHNPEVAGSNPVPATPM | 687934 | 688117 | + | 6 |
| 2 | LNLRPLGYEPSELPSC | STPRRHPEPYGNDTDPSSRPRIGVTASPG | 688091 | 687953 | - | 6 |
| 3 | LWVCCSLVAGLSASP | YLPSVLGAGLFLAASSATSLAACSANLFASALSLARASYWPTSRK | 688911 | 689094 | + | 6 |
| 4 | VCCSLVAGLSASP | YLPSVLGAGLFLAASSATSLAACSANLFASALSLARASYWPTSRK | 688917 | 689094 | + | 6 |
| 5 | VAGLSASPYLPSVLGAGLFLAASSAT | SLAACSANLFASALSLARASYWPTSRK | 688932 | 689094 | + | 6 |
| 6 | LSASPYLPSVLGAGLFLAASSATSL | AACSANLFASALSLARASYWPTSRK | 688941 | 689094 | + | 6 |
| 7 | LSAASAAFRTPGPEFWSERILRMAKRLFLSAFAFPCVSGLASGAVSG | VELAGDVAVSAATAGEANRTAGAIAAAATTVEIRDSFTGTLTFSGKW | 689394 | 689679 | + | 6 |
| 8 | LPFSRSFPTRTPGPASGRPPRRHSTALPRT | GFTSSASAASRLRTSTTGSGPRRSCTTAAPR | 689467 | 689281 | - | 7 |
| 9 | LFLSAFAFPCVSGLASGAVSGVELAGDVAVSAATAGEANRTAGAIAAA | ATTVEIRDSFTGTLTFSGKW | 689472 | 689679 | + | 6 |
| 10 | LSAFAFPCVSGLASGAVSGVELAGDVAVSAATAGEANRTAGAIAAA | ATTVEIRDSFTGTLTFSGKW | 689478 | 689679 | + | 6 |
| 11 | MNKSAVAGCLAVGLAVLGF | LTATGTVNIPIAMGAGVALFIAAGVEIKRKPTR | 693867 | 693708 | - | 8 |

| | | | | | | |
|---|---|---|---|---|---|---|
| 12 | MATPVLATPAAFMGGI | AISSAVVGAFEAGRNG | 696738 | 696639 | - | 15 |
| 13 | MEIPEEGAIFPMATPVL | ATPAAFMGGIAISSAVVGAFEAGRNG | 696771 | 696639 | - | 27 |
| 14 | MIDVAAAANEMEIPEEGAIFPMATPVL | ATPAAFMGGIAISSAVVGAFEAGRNG | 696801 | 696639 | - | 29 |
| 15 | MTEIVSFVNGLDLGAIETGEAHSDQAPVL | ATPAAALAAIGGAKAVGAFVGGATVVGGAYALGRAVG | 697117 | 696916 | - | 30 |
| 16 | MFWDPSHLTVRQECSAAGGYKHPCAEMGYGAGHSEMRSALPRS | LTVVMRGRDRCHTNP | 697134 | 697311 | + | 8 |
| 17 | VARNEGFHPHALAGMIIEVLEQYEPTGAVDTARF | PTLDDFAARHIAHRHP | 701283 | 701130 | - | 6 |

### Results for ANZ17123.1 [Streptomyces noursei ATCC 11455] back to top

previous - next

#### Architecture

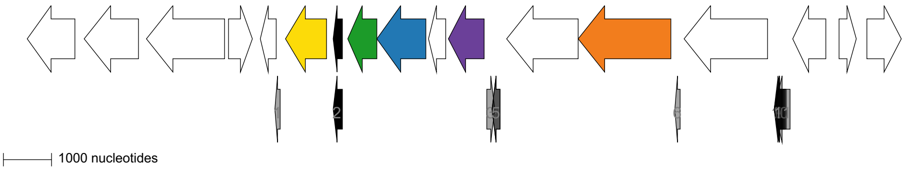

Link to nucleotide sequence

| Accession | start | end | direction | length (aa) | Pfam/HMM | name | description | E-value |
|---|---|---|---|---|---|---|---|---|
| ANZ17115.1 | 4365920 | 4364912 | - | 335 | TIGR02432<br>PF01171<br>PF09179 | TIGR02432<br>ATP_bind_3<br>TilS | lysidine_TilS_N: tRNA(Ile)-lysidine synthetase<br>PP-loop family<br>TilS substrate binding domain | 2.10E-43<br>2.00E-38<br>3.00E-12 |
| ANZ17116.1 | 4367235 | 4366113 | - | 373 | TIGR03883<br>TIGR03624<br>PF10103 | TIGR03883<br>TIGR03624<br>Zincin_2 | DUF2342_F420: uncharacterized protein, coenzyme F420 biosynthesis associated<br>TIGR03624: putative hydrolase<br>Zincin-like metallopeptidase | 8.80E-141<br>3.70E-126<br>1.90E-121 |
| ANZ17117.1 | 4369018 | 4367401 | - | 538 | PF02113<br>TIGR00666 | Peptidase_S13<br>TIGR00666 | D-Ala-D-Ala carboxypeptidase 3 (S13) family<br>PBP4: D-alanyl-D-alanine carboxypeptidase/D-alanyl-D-alanine-endopeptidase | 7.00E-77<br>1.70E-58 |
| ANZ17118.1 | 4369104 | 4369596 | + | 163 | PF00719 | Pyrophosphatase | Inorganic pyrophosphatase | 3.80E-53 |
| ANZ17119.1 | 4370080 | 4369753 | - | 108 | NO PFAM MATCH | - | - | - |
| ANZ17120.1 | 4371138 | 4370271 | - | 288 | LinG<br>LinH<br>TIGR04033 | LinG<br>LinH<br>TIGR04033 | LinG Homolog<br>LinH (Fusion of LinE/LinG)<br>export_SdpB: antimicrobial peptide system protein, SdpB family | 7.10E-110<br>2.20E-78<br>4.40E-04 |
| ANZ17121.1 | 4371465 | 4371282 | - | 60 | LinA | LinA | LinA (linaridin precursor) | 2.60E-14 |
| ANZ17122.1 | 4372173 | 4371588 | - | 194 | LinL | LinL | LinL | 5.10E-56 |
| ANZ17123.1 | 4373192 | 4372172 | - | 339 | LinE<br>LinH<br>PF12697<br>TIGR02427<br>PF00561 | LinE<br>LinH<br>Abhydrolase_6<br>TIGR02427<br>Abhydrolase_1 | LinE Homolog<br>LinH (Fusion of LinE/LinG)<br>Alpha/beta hydrolase family<br>protocat_pcaD: 3-oxoadipate enol-lactonase<br>alpha/beta hydrolase fold | 2.80E-102<br>1.40E-52<br>9.10E-16<br>4.10E-10<br>1.70E-07 |
| ANZ17124.1 | 4373612 | 4373264 | - | 115 | PF13518<br>PF01527<br>PF13551<br>PF13384<br>PF13011 | HTH_28<br>HTH_Tnp_1<br>HTH_29<br>HTH_23<br>LZ_Tnp_IS481 | Helix-turn-helix domain<br>Transposase<br>Winged helix-turn helix<br>Homeodomain-like domain<br>leucine-zipper of insertion element IS481 | 3.30E-10<br>2.50E-08<br>3.00E-07<br>3.40E-06<br>3.60E-05 |
| ANZ17125.1 | 4374413 | 4373654 | - | 252 | PF13649<br>PF08241<br>PF08242<br>PF13489<br>PF13847 | Methyltransf_25<br>Methyltransf_11<br>Methyltransf_12<br>Methyltransf_23<br>Methyltransf_31 | Methyltransferase domain<br>Methyltransferase domain<br>Methyltransferase domain<br>Methyltransferase domain<br>Methyltransferase domain | 2.20E-16<br>3.80E-11<br>4.30E-10<br>1.90E-09<br>1.80E-06 |
| ANZ17126.1 | 4376359 | 4374874 | - | 494 | TIGR03467<br>TIGR00292<br>TIGR02734<br>PF01946<br>PF13450 | TIGR03467<br>TIGR00292<br>TIGR02734<br>Thi4<br>NAD_binding_8 | HpnE: squalene-associated FAD-dependent desaturase<br>TIGR00292: thiazole biosynthesis enzyme<br>crtI_fam: phytoene desaturase<br>Thi4 family<br>NAD(P)-binding Rossmann-like domain | 7.70E-09<br>1.90E-08<br>1.70E-06<br>1.90E-06<br>1.20E-05 |
| ANZ17127.1 | 4378293 | 4376355 | - | 645 | TIGR02204<br>TIGR02203<br>TIGR00958<br>TIGR03796<br>TIGR03375 | TIGR02204<br>TIGR02203<br>TIGR00958<br>TIGR03796<br>TIGR03375 | MsbA_rel: ABC transporter, permease/ATP-binding protein<br>MsbA_lipidA: lipid A export permease/ATP-binding protein MsbA<br>3a01208: antigen peptide transporter 2<br>NHLM_micro_ABC1: NHLM bacteriocin system ABC transporter, peptidase/ATP-binding protein<br>type_I_sec_LssB: type I secretion system ATPase | 1.70E-97<br>1.20E-95<br>1.50E-88<br>7.90E-82<br>9.60E-79 |
| ANZ17128.1 | 4380285 | 4378572 | - | 570 | PF06738<br>PF12821 | ThrE<br>ThrE_2 | Putative threonine/serine exporter<br>Threonine/Serine exporter, ThrE | 5.50E-73<br>2.70E-20 |
| ANZ17129.1 | 4381516 | 4380811 | - | 234 | PF09335 | SNARE_assoc | SNARE associated Golgi protein | 5.80E-25 |
| ANZ17130.1 | 4381777 | 4382146 | + | 122 | PF01906<br>TIGR03884 | YbjQ_1<br>TIGR03884 | Putative heavy-metal-binding<br>sel_bind_Methan: selenium-binding protein | 1.30E-33<br>7.80E-04 |
| ANZ17131.1 | 4382348 | 4383062 | + | 237 | PF01810<br>TIGR00949<br>TIGR00948 | LysE<br>TIGR00949<br>TIGR00948 | LysE type translocator<br>2A76: homoserine/Threonine efflux protein<br>2a75: L-lysine exporter | 2.80E-42<br>6.40E-40<br>5.40E-14 |

| index | leader | core | start | end | dir | score |
|---|---|---|---|---|---|---|
| 1 | VAACHVPGRGNPRDD | GMNSRTGSRLSGETHS | 4370172 | 4370076 | - | 6 |
| 2 | MSIVADFANTELVDVEPGRIGNDATPTM | LTPLATLATPEGVAVTSATAYALHEVTKDIQR | 4371465 | 4371282 | - | 32 |

| | | | | | | |
|---|---|---|---|---|---|---|
| 3 | MKPKAGARKDVTGQHNCTVELQSLRMVGQ | TSTPVDPVVTPQRSVGNFRLCRVSPSYTPG | 4374458 | 4374638 | + | 6 |
| 4 | VTRANPDARRGGNRTCRKVHFLVVS | HIFTGAHPGVYDGLTRHNLKFPTLR | 4374730 | 4374577 | - | 6 |
| 5 | LACAEAVTRANPDARRGGNRTCRKVHFL | VVSHIFTGAHPGVYDGLTRHNLKFPTLR | 4374748 | 4374577 | - | 7 |
| 6 | LFTSAGRRRWPAGQ | GVPGNQVTRNFMPDAVLVSVTLTTNL | 4378481 | 4378358 | - | 6 |
| 7 | VGGGRRQGAGAGRPLVHDRYWPTKACPHPTA | PAPLPNSPAQRNRPAPQARDTNPAAG | 4380604 | 4380430 | - | 8 |
| 8 | VGHKDGGWGQASGGRCGSPLGPRPVLAHK | GVPPPHSPGTAAEQPRAAESPRAAGA | 4380620 | 4380452 | - | 8 |
| 9 | VPLRPVPSRGWVIKTVGGGRRQGAGAGRPLVHDRYWPTKACPHPTA | PAPLPNSPAQRNRPAPQARDTNPAAG | 4380649 | 4380430 | - | 7 |
| 10 | VSGLLRFTSGHLVPLRPVPSRGWVI...QGAGAGRPLVHDRYWPTKACPHPTA | PAPLPNSPAQRNRPAPQARDTNPAAG | 4380685 | 4380430 | - | 7 |
| 11 | VLRACGAGPLRGAVGCGDGSEGHEV...QGAGAGRPLVHDRYWPTKACPHPTA | PAPLPNSPAQRNRPAPQARDTNPAAG | 4380757 | 4380430 | - | 6 |

### Results for WP_100108206.1 [Streptomyces peucetius subsp. caesius ATCC 27952] back to top

previous - next

#### Architecture

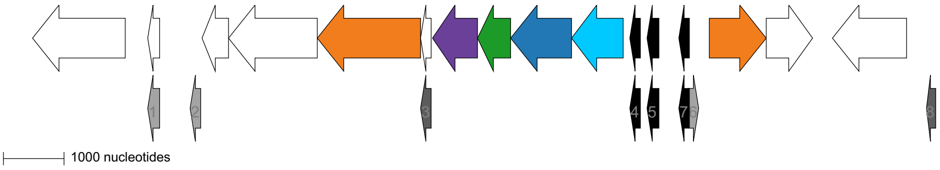

Link to nucleotide sequence

| Accession | start | end | direction | length (aa) | Pfam/HMM | name | description | E-value |
|---|---|---|---|---|---|---|---|---|
| WP_100108199.1 | 4093090 | 4091572 | - | 505 | PF00270<br>PF00271<br>TIGR04121<br>TIGR03817<br>TIGR01389 | DEAD<br>Helicase_C<br>TIGR04121<br>TIGR03817<br>TIGR01389 | DEAD/DEAH box helicase<br>Helicase conserved C-terminal domain<br>DEXH_lig_assoc: DEXH box helicase, DNA ligase-associated<br>DECH_helic: helicase/secretion neighborhood putative DEAH-box helicase<br>recQ: ATP-dependent DNA helicase RecQ | 3.10E-41<br>3.50E-26<br>5.80E-15<br>4.40E-13<br>1.70E-12 |
| WP_004986771.1 | 4093659 | 4093455 | - | 67 | PF00313<br>TIGR02381 | CSD<br>TIGR02381 | 'Cold-shock' DNA-binding domain<br>cspD: cold shock domain protein CspD | 1.30E-29<br>2.00E-28 |
| WP_100108200.1 | 4094788 | 4094362 | - | 141 | NO PFAM MATCH | - | - | - |
| WP_100108201.1 | 4096254 | 4094787 | - | 488 | TIGR02734<br>PF01593<br>PF13450<br>TIGR03467<br>TIGR01317 | TIGR02734<br>Amino_oxidase<br>NAD_binding_8<br>TIGR03467<br>TIGR01317 | crtI_fam: phytoene desaturase<br>Flavin containing amine oxidoreductase<br>NAD(P)-binding Rossmann-like domain<br>HpnE: squalene-associated FAD-dependent desaturase<br>GOGAT_sm_gam: glutamate synthase, NADH/NADPH, small subunit | 9.90E-12<br>1.00E-10<br>3.30E-10<br>3.30E-10<br>6.50E-08 |
| WP_100108202.1 | 4097969 | 4096250 | - | 572 | TIGR02204<br>TIGR02203<br>TIGR01846<br>TIGR03375<br>TIGR00958 | TIGR02204<br>TIGR02203<br>TIGR01846<br>TIGR03375<br>TIGR00958 | MsbA_rel: ABC transporter, permease/ATP-binding protein<br>MsbA_lipidA: lipid A export permease/ATP-binding protein MsbA<br>type_I_sec_HlyB: type I secretion system ATPase<br>type_I_sec_LssB: type I secretion system ATPase<br>3a01208: antigen peptide transporter 2 | 2.90E-106<br>3.70E-105<br>3.60E-87<br>5.20E-86<br>1.60E-82 |
| WP_100108203.1 | 4098145 | 4097965 | - | 59 | NO PFAM MATCH | - | - | - |
| WP_159026816.1 | 4098905 | 4098167 | - | 245 | PF13649<br>PF13489<br>PF08241<br>PF08242<br>TIGR01934 | Methyltransf_25<br>Methyltransf_23<br>Methyltransf_11<br>Methyltransf_12<br>TIGR01934 | Methyltransferase domain<br>Methyltransferase domain<br>Methyltransferase domain<br>Methyltransferase domain<br>MenG_MenH_UbiE: ubiquinone/menaquinone biosynthesis methyltransferase | 1.90E-14<br>4.20E-10<br>1.80E-09<br>8.90E-07<br>1.90E-05 |
| WP_100108205.1 | 4099447 | 4098901 | - | 181 | LinL | LinL | LinL | 7.90E-64 |
| WP_100108206.1 | 4100460 | 4099443 | - | 338 | LinE<br>LinH<br>PF12697<br>TIGR02427<br>PF00561 | LinE<br>LinH<br>Abhydrolase_6<br>TIGR02427<br>Abhydrolase_1 | LinE Homolog<br>LinH (Fusion of LinE/LinG)<br>Alpha/beta hydrolase family<br>protocat_pcaD: 3-oxoadipate enol-lactonase<br>alpha/beta hydrolase fold | 2.60E-126<br>6.00E-81<br>1.50E-14<br>2.10E-11<br>1.70E-10 |
| WP_100108207.1 | 4101293 | 4100456 | - | 278 | LinH<br>LinG | LinH<br>LinG | LinH (Fusion of LinE/LinG)<br>LinG Homolog | 1.00E-104<br>2.50E-103 |
| WP_100108208.1 | 4101586 | 4101406 | - | 59 | LinA | LinA | LinA (linaridin precursor) | 9.10E-10 |
| WP_100108209.1 | 4101890 | 4101689 | - | 66 | LinA | LinA | LinA (linaridin precursor) | 2.00E-10 |
| WP_100108210.1 | 4102392 | 4102212 | - | 59 | LinA | LinA | LinA (linaridin precursor) | 1.70E-09 |
| WP_100108211.1 | 4102728 | 4103661 | + | 310 | TIGR01188<br>TIGR03864<br>TIGR01288<br>TIGR01257<br>TIGR04406 | TIGR01188<br>TIGR03864<br>TIGR01288<br>TIGR01257<br>TIGR04406 | drrA: daunorubicin resistance ABC transporter, ATP-binding protein<br>PQQ_ABC_ATP: ABC transporter, ATP-binding subunit, PQQ-dependent alcohol dehydrogenase system<br>nodI: nodulation ABC transporter NodI<br>rim_protein: rim ABC transporter<br>LPS_export_lptB: LPS export ABC transporter ATP-binding protein | 2.20E-59<br>2.90E-43<br>6.90E-41<br>1.40E-32<br>1.50E-32 |

| | | | | | | | | |
|---|---|---|---|---|---|---|---|---|
| WP_159026817.1 | 4103672 | 4104422 | + | 249 | PF01061<br>PF12698 | ABC2_membrane<br>ABC2_membrane_3 | ABC-2 type transporter<br>ABC-2 family transporter protein | 3.50E-16<br>2.50E-05 |
| WP_100108213.1 | 4105985 | 4104743 | - | 413 | PF07228<br>TIGR02865<br>PF13185 | SpoIIE<br>TIGR02865<br>GAF_2 | Stage II sporulation protein E (SpoIIE)<br>spore_II_E: stage II sporulation protein E<br>GAF domain | 1.20E-40<br>4.90E-10<br>2.60E-04 |

| index | leader | core | start | end | dir | score |
|---|---|---|---|---|---|---|
| 1 | MATGTVKWFNAEKGFGFIEQDGGGADVFAHYSN | IAAQGFRELLEGQKVSFDIAQGGQKGPTAENIVPA | 4093659 | 4093455 | - | 6 |
| 2 | MNGIDRNDLDEVIA | KTFNEVKTAVNTHSEKSLQMYSGALRALVELRQQILSEERVAR | 4094337 | 4094163 | - | 6 |
| 3 | MLTANRLAWFAIAFAFAVPSMLVMFRDNGVITRDSWTKGFVFA | ATIAAVIAAVFGKGSR | 4098145 | 4097965 | - | 8 |
| 4 | MSVLVEKVNAGAGHAADGSIAPVM | ATPAAFVAGVVAGGKVSAVITAAAGVGAAVKACQK | 4101586 | 4101406 | - | 31 |
| 5 | MLTHKVNADASLKGALMAGAPARGTAPVM | ATPAIFATPVIAKAAGAVVGAGAVTGAAYAAYKTATG | 4101890 | 4101689 | - | 29 |
| 6 | LFRNMSFPVVCDHVDCADGREGPSGG | WAAQRRDVRGTRRRVVIPCAGERAASI | 4102378 | 4102540 | + | 6 |
| 7 | MFLNKVNQDVSLEDFVAERGPEARIAPIM | ATPAIFTPAAVKVGVSAGAAAAAYMATKVK | 4102392 | 4102212 | - | 31 |
| 8 | MPEQDLAAACTYLAGEGLITVDWARGNTPAM | VTLTHQGIRFMEAEEEEPR | 4106454 | 4106301 | - | 8 |

### Results for SFJ78423.1 [Streptomyces pini] back to top

previous - next

#### Architecture

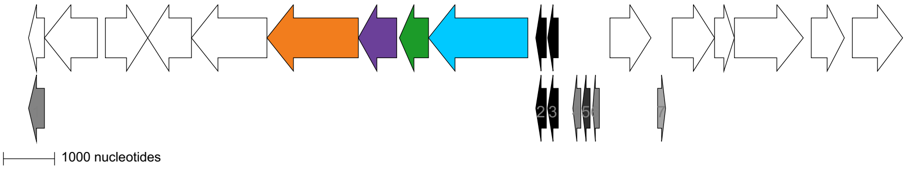

Link to nucleotide sequence

| Accession | start | end | direction | length (aa) | Pfam/HMM | name | description | E-value |
|---|---|---|---|---|---|---|---|---|
| SFJ78250.1 | 347917 | 347599 | - | 105 | PF00550<br>TIGR01688 | PP-binding<br>TIGR01688 | Phosphopantetheine attachment site<br>dltC: D-alanine--poly(phosphoribitol) ligase, subunit 2 | 7.30E-10<br>7.00E-07 |
| SFJ78269.1 | 348954 | 347913 | - | 346 | TIGR00747 | TIGR00747 | fabH: 3-oxoacyl-[acyl-carrier-protein] synthase III | 2.00E-11 |
| SFJ78292.1 | 349098 | 349941 | + | 280 | PF01551 | Peptidase_M23 | Peptidase family M23 | 4.80E-16 |
| SFJ78315.1 | 350777 | 349925 | - | 283 | PF12680 | SnoaL_2 | SnoaL-like domain | 5.80E-07 |
| SFJ78333.1 | 352255 | 350773 | - | 493 | TIGR02734<br>TIGR01317<br>TIGR02733<br>TIGR03467<br>PF13450 | TIGR02734<br>TIGR01317<br>TIGR02733<br>TIGR03467<br>NAD_binding_8 | crtI_fam: phytoene desaturase<br>GOGAT_sm_gam: glutamate synthase, NADH/NADPH, small subunit<br>desat_CrtD: C-3',4' desaturase CrtD<br>HpnE: squalene-associated FAD-dependent desaturase<br>NAD(P)-binding Rossmann-like domain | 2.70E-12<br>1.10E-08<br>1.10E-08<br>1.10E-08<br>5.80E-08 |
| SFJ78352.1 | 354039 | 352251 | - | 595 | TIGR02204<br>TIGR02203<br>TIGR03796<br>TIGR00958<br>TIGR01846 | TIGR02204<br>TIGR02203<br>TIGR03796<br>TIGR00958<br>TIGR01846 | MsbA_rel: ABC transporter, permease/ATP-binding protein<br>MsbA_lipidA: lipid A export permease/ATP-binding protein MsbA<br>NHLM_micro_ABC1: NHLM bacteriocin system ABC transporter, peptidase/ATP-binding protein<br>3a01208: antigen peptide transporter 2<br>type_I_sec_HlyB: type I secretion system ATPase | 2.50E-99<br>1.20E-94<br>9.20E-85<br>9.50E-85<br>7.80E-78 |
| SFJ78379.1 | 354791 | 354035 | - | 251 | PF13649<br>PF08241<br>PF13489<br>PF08242<br>PF13847 | Methyltransf_25<br>Methyltransf_11<br>Methyltransf_23<br>Methyltransf_12<br>Methyltransf_31 | Methyltransferase domain<br>Methyltransferase domain<br>Methyltransferase domain<br>Methyltransferase domain<br>Methyltransferase domain | 2.00E-13<br>4.00E-09<br>1.20E-08<br>5.60E-07<br>1.10E-06 |
| SFJ78400.1 | 355419 | 354840 | - | 192 | LinL | LinL | LinL | 1.40E-64 |
| SFJ78423.1 | 357350 | 355415 | - | 644 | LinH<br>LinG<br>LinE<br>PF12697<br>PF00561 | LinH<br>LinG<br>LinE<br>Abhydrolase_6<br>Abhydrolase_1 | LinH (Fusion of LinE/LinG)<br>LinG Homolog<br>LinE Homolog<br>Alpha/beta hydrolase family<br>alpha/beta hydrolase fold | 1.00E-243<br>2.40E-116<br>3.90E-100<br>3.90E-14<br>5.00E-13 |
| SFJ78431.1 | 357716 | 357521 | - | 64 | LinA | LinA | LinA (linaridin precursor) | 3.00E-10 |
| SFJ78454.1 | 357945 | 357753 | - | 63 | LinA | LinA | LinA (linaridin precursor) | 2.90E-16 |
| SFJ78480.1 | 358953 | 359769 | + | 271 | PF13472<br>PF14606 | Lipase_GDSL_2<br>Lipase_GDSL_3 | GDSL-like Lipase/Acylhydrolase family<br>GDSL-like Lipase/Acylhydrolase family | 2.40E-16<br>3.20E-05 |
| SFJ78504.1 | 360185 | 360998 | + | 270 | PF08892 | YqcI_YcgG | YqcI/YcgG family | 3.40E-66 |
| SFJ78524.1 | 361009 | 361396 | + | 128 | PF12973<br>PF07883 | Cupin_7<br>Cupin_2 | ChrR Cupin-like domain<br>Cupin domain | 5.40E-07<br>3.10E-04 |
| SFJ78547.1 | 361392 | 362733 | + | 446 | PF13520<br>TIGR00909<br>TIGR00911<br>TIGR00906<br>PF00324 | AA_permease_2<br>TIGR00909<br>TIGR00911<br>TIGR00906<br>AA_permease | Amino acid permease<br>2A0306: amino acid transporter<br>2A0308: L-type amino acid transporter<br>2A0303: cationic amino acid transport permease<br>Amino acid permease | 1.50E-31<br>2.80E-30<br>9.10E-29<br>1.60E-17<br>3.10E-16 |
| SFJ78568.1 | 362881 | 363550 | + | 222 | NO PFAM MATCH | - | - | - |
| SFJ78592.1 | 363685 | 364672 | + | 328 | PF00248<br>TIGR01293 | Aldo_ket_red<br>TIGR01293 | Aldo/keto reductase family<br>Kv_beta: voltage-dependent potassium channel beta subunit | 4.00E-72<br>1.50E-39 |

| index | leader | core | start | end | dir | score |
|---|---|---|---|---|---|---|
| 1 | MSHGTGETAGEITGETAEEITGEIT...YVEQRFDLTVEVEDLDLDSFRTARR | LTAFVRRKTGAETPDRTRAGDR | 347917 | 347599 | - | 7 |
| 2 | MSKIVESVVAVTPEELTDETHGEFGAMMPIQ | ATPGLVAFGVGFAGGAGAAWVTVQAYEAGRTAG | 357716 | 357521 | - | 31 |
| 3 | MEQKVAAFMNEQDFGPLEQGYQDDTPAPVL | ATPGAVATCWYVGGAVVGAGSVVGAYVTGRVVG | 357945 | 357753 | - | 29 |
| 4 | VAGKRCPGPRRVPACPFDRPHGPWDAARARGPCRGEPAPS | ACPVRTPRGIPFSMVM | 358396 | 358225 | - | 7 |
| 5 | METGQGETCEKQVPVAETSLCRCWFHR | TGPTTRPEVNENLFPSLSSVLPGGPALL | 358580 | 358412 | - | 9 |
| 6 | MAGVRRGCRDLSGQEEEPNIPF | LPSGSPVARVYTPTGKHVAAPA | 358760 | 358625 | - | 7 |
| 7 | MTTDIHRSPEMPGFTVAGKGSGSGHPSA | NTVLQQRCSARWTRSQPVVHTCRSGWK | 359879 | 360047 | + | 6 |

### Results for WP_069992530.1 [Streptomyces qinglanensis] back to top

previous - next

#### Architecture

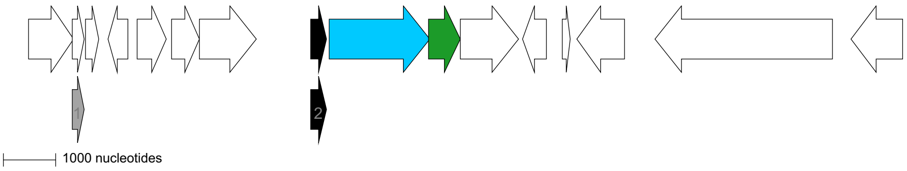

Link to nucleotide sequence

| Accession | start | end | direction | length (aa) | Pfam/HMM | name | description | E-value |
|---|---|---|---|---|---|---|---|---|
| WP_026004768.1 | 3913407 | 3914262 | + | 284 | PF13560 | HTH_31 | Helix-turn-helix domain | 5.40E-07 |
| WP_019357380.1 | 3914242 | 3914464 | + | 73 | PF04149 | DUF397 | Domain of unknown function (DUF397) | 2.10E-22 |
| WP_069992527.1 | 3914491 | 3914743 | + | 83 | PF04149 | DUF397 | Domain of unknown function (DUF397) | 2.90E-20 |
| WP_027759151.1 | 3915290 | 3914909 | - | 126 | NO PFAM MATCH | - | - | - |
| WP_069992528.1 | 3915472 | 3916039 | + | 188 | PF03795 | YCII | YCII-related domain | 1.10E-19 |
| WP_019357376.1 | 3916146 | 3916668 | + | 173 | PF01243<br>TIGR03618<br>TIGR03667<br>TIGR03666<br>TIGR03668 | Putative_PNPOx<br>TIGR03618<br>TIGR03667<br>TIGR03666<br>TIGR03668 | Pyridoxamine 5'-phosphate oxidase<br>Rv1155_F420: PPOX class probable F420-dependent enzyme<br>Rv3369: PPOX class probable F420-dependent enzyme, Rv3369 family<br>Rv2061_F420: PPOX class probable F420-dependent enzyme, Rv2061 family<br>Rv0121_F420: PPOX class probable F420-dependent enzyme, Rv0121 family | 2.70E-16<br>3.00E-16<br>1.10E-12<br>1.70E-07<br>4.60E-04 |
| WP_079135149.1 | 3916664 | 3917762 | + | 365 | PF00144 | Beta-lactamase | Beta-lactamase | 1.40E-56 |
| WP_019357374.1 | 3918790 | 3919087 | + | 98 | LinA | LinA | LinA (linaridin precursor) | 1.50E-08 |
| WP_069992530.1 | 3919143 | 3921063 | + | 639 | LinH<br>LinE<br>LinG<br>TIGR02427<br>PF00561 | LinH<br>LinE<br>LinG<br>TIGR02427<br>Abhydrolase_1 | LinH (Fusion of LinE/LinG)<br>LinE Homolog<br>LinG Homolog<br>protocat_pcaD: 3-oxoadipate enol-lactonase<br>alpha/beta hydrolase fold | 5.80E-197<br>1.00E-93<br>2.50E-87<br>6.60E-14<br>6.60E-09 |
| WP_069992531.1 | 3921049 | 3921652 | + | 200 | LinL | LinL | LinL | 3.90E-59 |
| WP_069992532.1 | 3921648 | 3922776 | + | 375 | PF00891<br>TIGR02716<br>PF16864<br>PF12802<br>TIGR03534 | Methyltransf_2<br>TIGR02716<br>Dimerisation2<br>MarR_2<br>TIGR03534 | O-methyltransferase domain<br>C20_methyl_CrtF: C-20 methyltransferase BchU<br>Dimerisation domain<br>MarR family<br>RF_mod_PrmC: protein-(glutamine-N5) methyltransferase, release factor-specific | 6.70E-44<br>3.50E-16<br>1.20E-08<br>2.40E-05<br>1.00E-04 |
| WP_019357370.1 | 3923301 | 3922839 | - | 153 | PF08327<br>PF10604<br>PF03364 | AHSA1<br>Polyketide_cyc2<br>Polyketide_cyc | Activator of Hsp90 ATPase homolog 1-like protein<br>Polyketide cyclase / dehydrase and lipid transport<br>Polyketide cyclase / dehydrase and lipid transport | 8.30E-07<br>6.60E-06<br>6.80E-04 |
| WP_006123527.1 | 3923600 | 3923745 | + | 48 | INFERRED GENE | - | - | - |
| WP_070032345.1 | 3924796 | 3923872 | - | 307 | PF14200<br>PF00652<br>PF03498 | RicinB_lectin_2<br>Ricin_B_lectin<br>CDtoxinA | Ricin-type beta-trefoil lectin domain-like<br>Ricin-type beta-trefoil lectin domain<br>Cytolethal distending toxin A/C domain | 4.20E-20<br>2.00E-17<br>8.20E-05 |
| WP_069992534.1 | 3928779 | 3925386 | - | 1130 | NO PFAM MATCH | - | - | - |
| WP_069992535.1 | 3930115 | 3929134 | - | 326 | PF07179 | SseB | SseB protein N-terminal domain | 2.70E-04 |

| index | leader | core | start | end | dir | score |
|---|---|---|---|---|---|---|
| 1 | MANTEIDLQQAVWRKSSYSNGSGGE...AVETVDASVPVRDTKTAPHGPVLTF | PTAQWAHFISTLHEGNVNA | 3914242 | 3914464 | + | 6 |
| 2 | MPITRESRAARWAQENAGEWCGHRVPEADEAVPVL | TTPIFATFAVAGAAGAGLGAARVVG...GSNEVSGYLEAGRPVKELLANHPAI | 3918790 | 3919087 | + | 26 |

### Results for SEQ27572.1 [Streptomyces radiopugnans] back to top

previous - next

#### Architecture

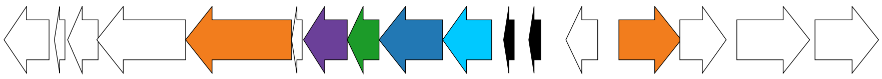

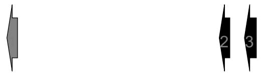

1000 nucleotides

Link to nucleotide sequence

| Accession | start | end | direction | length (aa) | Pfam/HMM | name | description | E-value |
|---|---|---|---|---|---|---|---|---|
| SEQ27380.1 | 231328 | 230572 | - | 251 | PF13472<br>PF14606 | Lipase_GDSL_2<br>Lipase_GDSL_3 | GDSL-like Lipase/Acylhydrolase family<br>GDSL-like Lipase/Acylhydrolase family | 1.70E-16<br>1.20E-04 |
| SEQ27405.1 | 231602 | 231428 | - | 57 | NO PFAM MATCH | - | - | - |
| SEQ27426.1 | 232155 | 231651 | - | 167 | NO PFAM MATCH | - | - | - |
| SEQ27450.1 | 233641 | 232147 | - | 497 | TIGR02734<br>TIGR03467<br>PF13450<br>TIGR00292<br>PF01593 | TIGR02734<br>TIGR03467<br>NAD_binding_8<br>TIGR00292<br>Amino_oxidase | crtI_fam: phytoene desaturase<br>HpnE: squalene-associated FAD-dependent desaturase<br>NAD(P)-binding Rossmann-like domain<br>TIGR00292: thiazole biosynthesis enzyme<br>Flavin containing amine oxidoreductase | 2.50E-13<br>4.90E-10<br>1.30E-09<br>1.50E-08<br>2.10E-08 |
| SEQ27467.1 | 235431 | 233637 | - | 597 | TIGR02204<br>TIGR02203<br>TIGR00958<br>TIGR03796<br>TIGR01846 | TIGR02204<br>TIGR02203<br>TIGR00958<br>TIGR03796<br>TIGR01846 | MsbA_rel: ABC transporter, permease/ATP-binding protein<br>MsbA_lipidA: lipid A export permease/ATP-binding protein MsbA<br>3a01208: antigen peptide transporter 2<br>NHLM_micro_ABC1: NHLM bacteriocin system ABC transporter, peptidase/ATP-binding protein<br>type_I_sec_HlyB: type I secretion system ATPase | 3.10E-103<br>3.40E-99<br>7.60E-86<br>4.00E-84<br>2.00E-81 |
| SEQ27491.1 | 235607 | 235427 | - | 59 | NO PFAM MATCH | - | - | - |
| SEQ27525.1 | 236379 | 235623 | - | 251 | PF13649<br>PF08241<br>PF13489<br>PF08242<br>TIGR01934 | Methyltransf_25<br>Methyltransf_11<br>Methyltransf_23<br>Methyltransf_12<br>TIGR01934 | Methyltransferase domain<br>Methyltransferase domain<br>Methyltransferase domain<br>Methyltransferase domain<br>MenG_MenH_UbiE: ubiquinone/menaquinone biosynthesis methyltransferase | 4.80E-13<br>1.40E-10<br>5.40E-08<br>1.70E-06<br>8.70E-06 |
| SEQ27555.1 | 236918 | 236375 | - | 180 | LinL | LinL | LinL | 4.00E-64 |
| SEQ27572.1 | 237976 | 236914 | - | 353 | LinE<br>LinH<br>PF12697<br>TIGR02427<br>TIGR03611 | LinE<br>LinH<br>Abhydrolase_6<br>TIGR02427<br>TIGR03611 | LinE Homolog<br>LinH (Fusion of LinE/LinG)<br>Alpha/beta hydrolase family<br>protocat_pcaD: 3-oxoadipate enol-lactonase<br>RutD: pyrimidine utilization protein D | 1.30E-129<br>1.10E-80<br>4.90E-19<br>4.30E-09<br>4.20E-08 |
| SEQ27597.1 | 238809 | 237972 | - | 278 | LinH<br>LinG | LinH<br>LinG | LinH (Fusion of LinE/LinG)<br>LinG Homolog | 3.50E-104<br>2.40E-102 |
| SEQ27616.1 | 239191 | 239011 | - | 59 | LinA | LinA | LinA (linaridin precursor) | 4.30E-11 |
| SEQ27644.1 | 239641 | 239440 | - | 66 | LinA | LinA | LinA (linaridin precursor) | 7.60E-13 |
| SEQ27682.1 | 240594 | 240069 | - | 174 | TIGR02983<br>TIGR02937<br>PF04542<br>PF08281<br>TIGR02984 | TIGR02983<br>TIGR02937<br>Sigma70_r2<br>Sigma70_r4_2<br>TIGR02984 | SigE-fam_strep: RNA polymerase sigma-70 factor, sigma-E family<br>sigma70-ECF: RNA polymerase sigma factor, sigma-70 family<br>Sigma-70 region 2<br>Sigma-70, region 4<br>Sig-70_plancto1: RNA polymerase sigma-70 factor, Planctomycetaceae-specific subfamily 1 | 2.70E-15<br>1.30E-11<br>2.20E-10<br>2.50E-06<br>2.80E-04 |
| SEQ27701.1 | 240961 | 242002 | + | 346 | TIGR01188<br>TIGR03864<br>TIGR01288<br>TIGR04406<br>TIGR02673 | TIGR01188<br>TIGR03864<br>TIGR01288<br>TIGR04406<br>TIGR02673 | drrA: daunorubicin resistance ABC transporter, ATP-binding protein<br>PQQ_ABC_ATP: ABC transporter, ATP-binding subunit, PQQ-dependent alcohol dehydrogenase system<br>nodI: nodulation ABC transporter NodI<br>LPS_export_lptB: LPS export ABC transporter ATP-binding protein<br>FtsE: cell division ATP-binding protein FtsE | 4.40E-61<br>2.00E-43<br>3.60E-41<br>3.00E-35<br>4.10E-33 |
| SEQ27723.1 | 241998 | 242763 | + | 254 | PF01061<br>PF12698<br>TIGR03062 | ABC2_membrane<br>ABC2_membrane_3<br>TIGR03062 | ABC-2 type transporter<br>ABC-2 family transporter protein<br>pip_yhgE_Cterm: YhgE/Pip C-terminal domain | 1.40E-15<br>4.60E-06<br>5.90E-04 |
| SEQ27750.1 | 242942 | 244190 | + | 415 | PF00144<br>PF13354 | Beta-lactamase<br>Beta-lactamase2 | Beta-lactamase<br>Beta-lactamase enzyme family | 4.00E-61<br>3.80E-07 |
| SEQ27772.1 | 244265 | 245342 | + | 358 | PF13449<br>PF07676 | Phytase-like<br>PD40 | Esterase-like activity of phytase<br>WD40-like Beta Propeller Repeat | 2.70E-04<br>3.20E-04 |

| index | leader | core | start | end | dir | score |
|---|---|---|---|---|---|---|
| 1 | MLTASRAAWFGAAFVFALPAMLLMFGDSGLTR | GTLWLQSALFAASIAAAVAVVFGRGPS | 235607 | 235427 | - | 7 |
| 2 | MSVLVEKVNAGVGRPVEDEAAPVM | ATPAAFAAGVVAGAKASAILVAAAGVGAAVANAQK | 239191 | 239011 | - | 31 |
| 3 | MLIRKVNADTSLRGTLEPGSPLGGAAPVM | ATPALIATPVIAKAAGAVVGAGAVTGAAYVAYKTATG | 239641 | 239440 | - | 30 |

### Results for GAQ68222.1 [Streptomyces scabiei] back to top

previous - next

#### Architecture

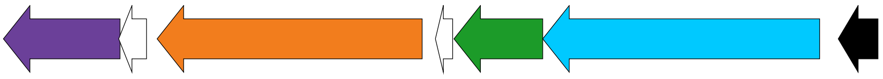

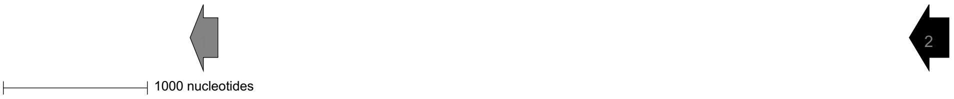

Link to nucleotide sequence

| Accession | start | end | direction | length (aa) | Pfam/HMM | name | description | E-value |
|---|---|---|---|---|---|---|---|---|
| GAQ68217.1 | 973 | 166 | - | 268 | PF13649<br>PF08241<br>PF13489<br>PF08242<br>PF13847 | Methyltransf_25<br>Methyltransf_11<br>Methyltransf_23<br>Methyltransf_12<br>Methyltransf_31 | Methyltransferase domain<br>Methyltransferase domain<br>Methyltransferase domain<br>Methyltransferase domain<br>Methyltransferase domain | 7.10E-15<br>4.90E-11<br>1.80E-08<br>5.20E-06<br>9.70E-06 |
| GAQ68218.1 | 1161 | 969 | - | 63 | NO PFAM MATCH | - | - | - |
| GAQ68219.1 | 3069 | 1230 | - | 612 | TIGR02204<br>TIGR02203<br>TIGR00958<br>TIGR03796<br>TIGR01846 | TIGR02204<br>TIGR02203<br>TIGR00958<br>TIGR03796<br>TIGR01846 | MsbA_rel: ABC transporter, permease/ATP-binding protein<br>MsbA_lipidA: lipid A export permease/ATP-binding protein MsbA<br>3a01208: antigen peptide transporter 2<br>NHLM_micro_ABC1: NHLM bacteriocin system ABC transporter, peptidase/ATP-binding protein<br>type_I_sec_HlyB: type I secretion system ATPase | 7.70E-105<br>1.80E-101<br>5.40E-91<br>1.00E-84<br>8.10E-78 |
| GAQ68220.1 | 3288 | 3162 | - | 41 | NO PFAM MATCH | - | - | - |
| GAQ68221.1 | 3911 | 3290 | - | 206 | LinL | LinL | LinL | 1.10E-70 |
| GAQ68222.1 | 5827 | 3907 | - | 639 | LinH<br>LinG<br>LinE<br>PF12697<br>PF12146 | LinH<br>LinG<br>LinE<br>Abhydrolase_6<br>Hydrolase_4 | LinH (Fusion of LinE/LinG)<br>LinG Homolog<br>LinE Homolog<br>Alpha/beta hydrolase family<br>Serine aminopeptidase, S33 | 3.90E-247<br>5.30E-163<br>2.90E-92<br>6.00E-16<br>2.60E-09 |
| GAQ68223.1 | 6235 | 5956 | - | 92 | LinA | LinA | LinA (linaridin precursor) | 2.50E-08 |

| index | leader | core | start | end | dir | score |
|---|---|---|---|---|---|---|
| 1 | MNDFLLIPSVVALGIVGFLIATRAVSTPAVI | GVALVVLWGACAQARSTVHPRHTPSKKRRQHP | 1161 | 969 | - | 7 |
| 2 | MSSIENALNSVEIPVEGVVYVAARPTL | GTPRIARVGNRLGGLAQAAEGVGAI...ANNLAAEANAQNAAALAAVGGAPAS | 6235 | 5956 | - | 28 |

### Results for GCD43950.1 [Streptomyces rimosus subsp. paromomycinus] back to top

previous - next

#### Architecture

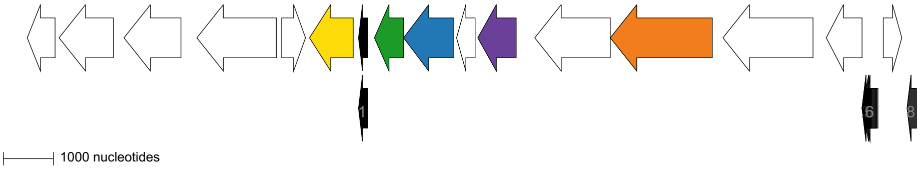

Link to nucleotide sequence

| Accession | start | end | direction | length (aa) | Pfam/HMM | name | description | E-value |
|---|---|---|---|---|---|---|---|---|
| GCD43942.1 | 4356193 | 4355653 | - | 179 | TIGR01203<br>PF00156<br>TIGR01251<br>TIGR01367 | TIGR01203<br>Pribosyltran<br>TIGR01251<br>TIGR01367 | HGPRTase: hypoxanthine phosphoribosyltransferase<br>Phosphoribosyl transferase domain<br>ribP_PPkin: ribose-phosphate diphosphokinase<br>pyrE_Therm: orotate phosphoribosyltransferase | 5.20E-65<br>1.30E-25<br>1.30E-05<br>5.70E-04 |
| GCD43943.1 | 4357370 | 4356281 | - | 362 | TIGR02432<br>PF01171<br>PF09179 | TIGR02432<br>ATP_bind_3<br>TilS | lysidine_TilS_N: tRNA(Ile)-lysidine synthetase<br>PP-loop family<br>TilS substrate binding domain | 5.50E-43<br>3.80E-38<br>2.30E-11 |
| GCD43944.1 | 4358728 | 4357579 | - | 382 | TIGR03883<br>TIGR03624<br>PF10103 | TIGR03883<br>TIGR03624<br>Zincin_2 | DUF2342_F420: uncharacterized protein, coenzyme F420 biosynthesis associated<br>TIGR03624: putative hydrolase<br>Zincin-like metallopeptidase | 7.70E-141<br>1.40E-123<br>6.40E-121 |
| GCD43945.1 | 4360632 | 4359024 | - | 535 | PF02113<br>TIGR00666 | Peptidase_S13<br>TIGR00666 | D-Ala-D-Ala carboxypeptidase 3 (S13) family<br>PBP4: D-alanyl-D-alanine carboxypeptidase/D-alanyl-D-alanine-endopeptidase | 7.10E-78<br>1.40E-55 |
| GCD43946.1 | 4360718 | 4361216 | + | 165 | PF00719 | Pyrophosphatase | Inorganic pyrophosphatase | 2.50E-53 |
| GCD43947.1 | 4362159 | 4361298 | - | 286 | LinG<br>LinH | LinG<br>LinH | LinG Homolog<br>LinH (Fusion of LinE/LinG) | 1.30E-106<br>1.50E-76 |
| GCD43948.1 | 4362449 | 4362257 | - | 63 | LinA | LinA | LinA (linaridin precursor) | 1.10E-10 |
| GCD43949.1 | 4363163 | 4362581 | - | 193 | LinL | LinL | LinL | 1.90E-55 |
| GCD43950.1 | 4364182 | 4363159 | - | 340 | LinE<br>LinH<br>PF12697<br>TIGR02427<br>PF00561 | LinE<br>LinH<br>Abhydrolase_6<br>TIGR02427<br>Abhydrolase_1 | LinE Homolog<br>LinH (Fusion of LinE/LinG)<br>Alpha/beta hydrolase family<br>protocat_pcaD: 3-oxoadipate enol-lactonase<br>alpha/beta hydrolase fold | 4.60E-100<br>1.40E-53<br>8.80E-12<br>3.30E-09<br>1.30E-07 |
| GCD43951.1 | 4364601 | 4364223 | - | 125 | PF01527<br>PF13518<br>PF13551<br>PF13384<br>PF13011 | HTH_Tnp_1<br>HTH_28<br>HTH_29<br>HTH_23<br>LZ_Tnp_IS481 | Transposase<br>Helix-turn-helix domain<br>Winged helix-turn helix<br>Homeodomain-like domain<br>leucine-zipper of insertion element IS481 | 9.10E-09<br>3.80E-07<br>4.10E-05<br>1.20E-04<br>9.80E-04 |
| GCD43952.1 | 4365415 | 4364647 | - | 255 | PF13649<br>PF08241<br>PF08242<br>PF13489<br>PF13847 | Methyltransf_25<br>Methyltransf_11<br>Methyltransf_12<br>Methyltransf_23<br>Methyltransf_31 | Methyltransferase domain<br>Methyltransferase domain<br>Methyltransferase domain<br>Methyltransferase domain<br>Methyltransferase domain | 1.30E-16<br>4.60E-12<br>1.40E-11<br>1.80E-08<br>1.00E-07 |

| | | | | | | | | |
|---|---|---|---|---|---|---|---|---|
| GCD43953.1 | 4367300 | 4365788 | - | 503 | TIGR03467 | TIGR03467 | HpnE: squalene-associated FAD-dependent desaturase | 2.00E-08 |
| | | | | | TIGR00292 | TIGR00292 | TIGR00292: thiazole biosynthesis enzyme | 2.40E-08 |
| | | | | | TIGR02734 | TIGR02734 | crtI_fam: phytoene desaturase | 2.60E-08 |
| | | | | | PF13450 | NAD_binding_8 | NAD(P)-binding Rossmann-like domain | 4.40E-08 |
| | | | | | PF01946 | Thi4 | Thi4 family | 2.10E-06 |
| GCD43954.1 | 4369342 | 4367296 | - | 681 | TIGR02204 | TIGR02204 | MsbA_rel: ABC transporter, permease/ATP-binding protein | 7.10E-96 |
| | | | | | TIGR02203 | TIGR02203 | MsbA_lipidA: lipid A export permease/ATP-binding protein MsbA | 4.60E-93 |
| | | | | | TIGR00958 | TIGR00958 | 3a01208: antigen peptide transporter 2 | 1.20E-87 |
| | | | | | TIGR03796 | TIGR03796 | NHLM_micro_ABC1: NHLM bacteriocin system ABC transporter, peptidase/ATP-binding protein | 3.90E-84 |
| | | | | | TIGR01846 | TIGR01846 | type_I_sec_HlyB: type I secretion system ATPase | 1.50E-76 |
| GCD43955.1 | 4371346 | 4369558 | - | 595 | PF06738 | ThrE | Putative threonine/serine exporter | 1.10E-71 |
| | | | | | PF12821 | ThrE_2 | Threonine/Serine exporter, ThrE | 1.70E-18 |
| GCD43956.1 | 4372327 | 4371607 | - | 239 | PF09335 | SNARE_assoc | SNARE associated Golgi protein | 5.10E-25 |
| GCD43957.1 | 4372765 | 4373137 | + | 123 | PF01906 | YbjQ_1 | Putative heavy-metal-binding | 2.70E-34 |
| | | | | | TIGR03884 | TIGR03884 | sel_bind_Methan: selenium-binding protein | 5.50E-04 |

| index | leader | core | start | end | dir | score |
|---|---|---|---|---|---|---|
| 1 | MSILADFANAELTDVKPGRIGADSTPTM | VTPLATIATPEATPVGFAATSATAVALDKVTHNIG | 4362449 | 4362257 | - | 33 |
| 2 | VEGSGAGCAGAAAGGWNHGSSSGVDSFGPHTAPASR | RTARSGALDPGAFTP | 4372479 | 4372323 | - | 9 |
| 3 | VAEPAAGAARTAVEGSGAGCAGAAAGGWNHGSSSGVDSFGPHTAPASR | RTARSGALDPGAFTP | 4372515 | 4372323 | - | 9 |
| 4 | MGGAGREEWGGKGWFCSSYGCLCGGGGACG | GGGPYGRGGVRRGVCGGRGGRLEPWLVIRR | 4372588 | 4372405 | - | 6 |
| 5 | MGGEGIGGDGRGWEGRAGRNGAERG...AAGGWNHGSSSGVDSFGPHTAPASR | RTARSGALDPGAFTP | 4372623 | 4372323 | - | 8 |
| 6 | MEGEGMGGEGIGGDGRGWEGRAGRN...AAGGWNHGSSSGVDSFGPHTAPASR | RTARSGALDPGAFTP | 4372638 | 4372323 | - | 7 |
| 7 | VAEEHRRFISSGYYDCSYEMHTCLGEMYVA | DERFTATYEAIRPGLAVYMRDAMLANAVRHS | 4373408 | 4373222 | - | 7 |
| 8 | MDVAEEHRRFISSGYYDCSYEMHTCLGEMYV | ADERFTATYEAIRPGLAVYMRDAMLANAVRHS | 4373414 | 4373222 | - | 8 |

### Results for KMO97782.1 [Streptomyces roseus] back to top

previous - next

#### Architecture

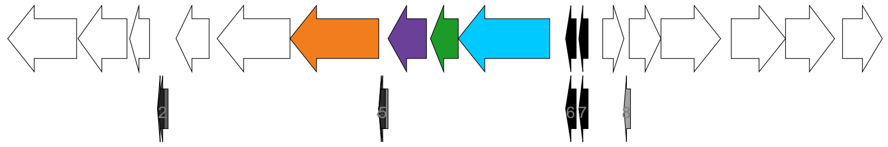

1000 nucleotides

Link to nucleotide sequence

| Accession | start | end | direction | length (aa) | Pfam/HMM | name | description | E-value |
|---|---|---|---|---|---|---|---|---|
| KMO97734.1 | 54042 | 52650 | - | 463 | TIGR01048 | TIGR01048 | lysA: diaminopimelate decarboxylase | 3.10E-144 |
| | | | | | PF02784 | Orn_Arg_deC_N | Pyridoxal-dependent decarboxylase, pyridoxal binding domain | 4.20E-67 |
| | | | | | TIGR03099 | TIGR03099 | dCO2ase_PEP1: pyridoxal-dependent decarboxylase, exosortase A system-associated | 1.40E-51 |
| | | | | | TIGR01273 | TIGR01273 | speA: arginine decarboxylase | 4.00E-22 |
| | | | | | PF00278 | Orn_DAP_Arg_deC | Pyridoxal-dependent decarboxylase, C-terminal sheet domain | 6.10E-14 |
| KMO97735.1 | 55057 | 54061 | - | 331 | TIGR00456 | TIGR00456 | argS: arginine--tRNA ligase | 6.70E-24 |
| | | | | | PF03485 | Arg_tRNA_synt_N | Arginyl tRNA synthetase N terminal domain | 2.00E-16 |
| | | | | | PF05746 | DALR_1 | DALR anticodon binding domain | 2.30E-16 |
| KMO97736.1 | 55529 | 55127 | - | 133 | PF00072 | Response_reg | Response regulator receiver domain | 7.90E-24 |
| | | | | | TIGR02154 | TIGR02154 | PhoB: phosphate regulon transcriptional regulatory protein PhoB | 1.30E-23 |
| | | | | | TIGR02875 | TIGR02875 | spore_0_A: sporulation transcription factor Spo0A | 5.90E-19 |
| | | | | | TIGR01818 | TIGR01818 | ntrC: nitrogen regulation protein NR(I) | 8.00E-18 |
| | | | | | TIGR02956 | TIGR02956 | TMAO_torS: TMAO reductase sytem sensor TorS | 2.60E-16 |
| KMO97781.1 | 56724 | 56043 | - | 226 | PF01551 | Peptidase_M23 | Peptidase family M23 | 2.90E-18 |
| KMO97737.1 | 58355 | 56891 | - | 487 | TIGR02734 | TIGR02734 | crtI_fam: phytoene desaturase | 4.60E-12 |
| | | | | | PF13450 | NAD_binding_8 | NAD(P)-binding Rossmann-like domain | 2.10E-10 |
| | | | | | TIGR03467 | TIGR03467 | HpnE: squalene-associated FAD-dependent desaturase | 2.60E-10 |
| | | | | | TIGR01372 | TIGR01372 | soxA: sarcosine oxidase, alpha subunit family | 1.70E-08 |
| | | | | | PF01593 | Amino_oxidase | Flavin containing amine oxidoreductase | 2.00E-08 |
| KMO97738.1 | 60172 | 58351 | - | 606 | TIGR02204 | TIGR02204 | MsbA_rel: ABC transporter, permease/ATP-binding protein | 4.10E-101 |
| | | | | | TIGR02203 | TIGR02203 | MsbA_lipidA: lipid A export permease/ATP-binding protein MsbA | 1.50E-98 |
| | | | | | TIGR03796 | TIGR03796 | NHLM_micro_ABC1: NHLM bacteriocin system ABC transporter, peptidase/ATP-binding protein | 1.30E-84 |
| | | | | | TIGR00958 | TIGR00958 | 3a01208: antigen peptide transporter 2 | 7.10E-83 |
| | | | | | TIGR01846 | TIGR01846 | type_I_sec_HlyB: type I secretion system ATPase | 3.00E-80 |
| KMO97739.1 | 61132 | 60346 | - | 261 | PF13649 | Methyltransf_25 | Methyltransferase domain | 3.90E-13 |
| | | | | | PF08241 | Methyltransf_11 | Methyltransferase domain | 6.60E-09 |
| | | | | | PF08242 | Methyltransf_12 | Methyltransferase domain | 6.20E-07 |
| | | | | | PF13489 | Methyltransf_23 | Methyltransferase domain | 5.20E-06 |
| | | | | | PF13847 | Methyltransf_31 | Methyltransferase domain | 1.00E-05 |
| KMO97740.1 | 61786 | 61198 | - | 195 | LinL | LinL | LinL | 3.10E-70 |
| KMO97782.1 | 63630 | 61782 | - | 615 | LinH | LinH | LinH (Fusion of LinE/LinG) | 5.90E-229 |
| | | | | | LinG | LinG | LinG Homolog | 3.50E-97 |
| | | | | | LinE | LinE | LinE Homolog | 5.40E-97 |
| | | | | | PF12697 | Abhydrolase_6 | Alpha/beta hydrolase family | 1.30E-15 |
| | | | | | PF12146 | Hydrolase_4 | Serine aminopeptidase, S33 | 1.60E-09 |

| | | | | | | | | |
|---|---|---|---|---|---|---|---|---|
| KMO97741.1 | 64153 | 63952 | - | 66 | LinA | LinA | LinA (linaridin precursor) | 5.90E-11 |
| KMO97742.1 | 64402 | 64207 | - | 64 | LinA | LinA | LinA (linaridin precursor) | 1.20E-12 |
| KMO97783.1 | 64690 | 65128 | + | 145 | PF13560 | HTH_31 | Helix-turn-helix domain | 2.20E-04 |
| KMO97743.1 | 65230 | 65878 | + | 215 | PF06114 | Peptidase_M78 | IrrE N-terminal-like domain | 2.90E-05 |
| KMO97744.1 | 65890 | 67096 | + | 401 | NO PFAM MATCH | - | - | - |
| KMO97745.1 | 67296 | 68406 | + | 369 | PF08028<br>TIGR04022<br>PF00441<br>PF02770 | Acyl-CoA_dh_2<br>TIGR04022<br>Acyl-CoA_dh_1<br>Acyl-CoA_dh_M | Acyl-CoA dehydrogenase, C-terminal domain<br>sulfur_SfnB: sulfur acquisition oxidoreductase, SfnB family<br>Acyl-CoA dehydrogenase, C-terminal domain<br>Acyl-CoA dehydrogenase, middle domain | 1.30E-25<br>5.90E-22<br>5.60E-05<br>2.70E-04 |
| KMO97746.1 | 68402 | 69389 | + | 328 | PF00248<br>TIGR01293 | Aldo_ket_red<br>TIGR01293 | Aldo/keto reductase family<br>Kv_beta: voltage-dependent potassium channel beta subunit | 3.20E-75<br>1.60E-26 |
| KMO97747.1 | 69557 | 70367 | + | 269 | PF03704<br>PF00486 | BTAD<br>Trans_reg_C | Bacterial transcriptional activator domain<br>Transcriptional regulatory protein, C terminal | 4.60E-42<br>8.30E-11 |

| index | leader | core | start | end | dir | score |
|---|---|---|---|---|---|---|
| 1 | MAGSLGTSCEVPPAGFEPAH | TAPEAVALSPELRGRVVRVAATG | 55806 | 55674 | - | 8 |
| 2 | MRRRVRNRRRTPAQGGSWPHIREDP...SANSLMAGSLGTSCEVPPAGFEPAH | TAPEAVALSPELRGRVVRVAATG | 55899 | 55674 | - | 8 |
| 3 | MAVIFREDGRFTSQSWTKGLIF | GTAVAVVAAIAAGRARQ | 60288 | 60168 | - | 6 |
| 4 | VAVPVMAVIFREDGRFTSQSWT | KGLIFGTAVAVVAAIAAGRARQ | 60303 | 60168 | - | 7 |
| 5 | MSTQRAVWFVTALAVAVPVM | AVIFREDGRFTSQSWTKGLIFGTAVAVVAAIAAGRARQ | 60345 | 60168 | - | 7 |
| 6 | MNSITESVRVIAGATLSQDTGERTTRMMPVA | FTPAVLVTTPYAAGGLAAGAAAVGAYEAGRAAAKG | 64153 | 63952 | - | 29 |
| 7 | MAKIAAFMNQQDMGPIAESELSVGGTPVM | ATPAVVAVPASVKITAGIVAFTIPVGAYVTGRVVN | 64402 | 64207 | - | 30 |
| 8 | LTCGLGNIGSGHLSVVRDGWISPVAC | HTFHRSMATRSRYR | 65253 | 65130 | - | 6 |

### Results for WP_078630323.1 [Streptomyces roseochromogenus subsp. oscitans DS 12.976] back to top

previous - next

#### Architecture

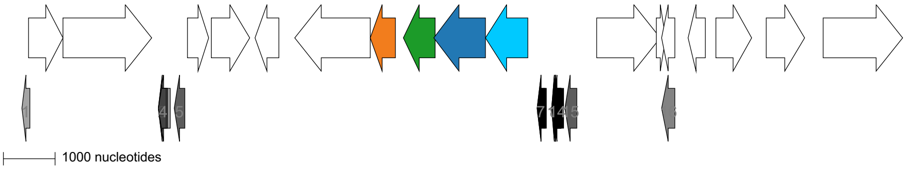

Link to nucleotide sequence

| Accession | start | end | direction | length (aa) | Pfam/HMM | name | description | E-value |
|---|---|---|---|---|---|---|---|---|
| WP_031225314.1 | 4545592 | 4546267 | + | 224 | PF05713 | MobC | Bacterial mobilisation protein (MobC) | 8.40E-09 |
| WP_023548214.1 | 4546263 | 4547970 | + | 568 | PF03432 | Relaxase | Relaxase/Mobilisation nuclease domain | 5.20E-06 |
| WP_023548216.1 | 4548673 | 4549066 | + | 130 | NO PFAM MATCH | - | - | - |
| WP_009307930.1 | 4549128 | 4549867 | + | 246 | INFERRED GENE | - | - | - |
| WP_158692712.1 | 4550428 | 4549981 | - | 148 | PF00072<br>TIGR02875<br>TIGR01818 | Response_reg<br>TIGR02875<br>TIGR01818 | Response regulator receiver domain<br>spore_0_A: sporulation transcription factor Spo0A<br>ntrC: nitrogen regulation protein NR(I) | 2.30E-13<br>3.20E-07<br>1.20E-04 |
| WP_023548221.1 | 4552207 | 4550737 | - | 489 | TIGR02734<br>PF13450<br>TIGR03467<br>TIGR01317<br>TIGR03315 | TIGR02734<br>NAD_binding_8<br>TIGR03467<br>TIGR01317<br>TIGR03315 | crtI_fam: phytoene desaturase<br>NAD(P)-binding Rossmann-like domain<br>HpnE: squalene-associated FAD-dependent desaturase<br>GOGAT_sm_gam: glutamate synthase, NADH/NADPH, small subunit<br>Se_ygfK: putative selenate reductase, YgfK subunit | 1.10E-13<br>1.40E-10<br>1.00E-09<br>3.30E-09<br>8.10E-08 |
| WP_023548222.1 | 4552679 | 4552199 | - | 159 | TIGR02204<br>TIGR02203<br>TIGR00958<br>TIGR01846<br>TIGR03796 | TIGR02204<br>TIGR02203<br>TIGR00958<br>TIGR01846<br>TIGR03796 | MsbA_rel: ABC transporter, permease/ATP-binding protein<br>MsbA_lipidA: lipid A export permease/ATP-binding protein MsbA<br>3a01208: antigen peptide transporter 2<br>type_I_sec_HlyB: type I secretion system ATPase<br>NHLM_micro_ABC1: NHLM bacteriocin system ABC transporter, peptidase/ATP-binding protein | 7.10E-45<br>4.90E-43<br>2.20E-40<br>4.40E-39<br>1.40E-35 |
| WP_158692713.1 | 4553446 | 4552846 | - | 199 | LinL | LinL | LinL | 1.20E-63 |
| WP_078630323.1 | 4554435 | 4553442 | - | 330 | LinE<br>LinH<br>PF12697<br>PF00561<br>TIGR02427 | LinE<br>LinH<br>Abhydrolase_6<br>Abhydrolase_1<br>TIGR02427 | LinE Homolog<br>LinH (Fusion of LinE/LinG)<br>Alpha/beta hydrolase family<br>alpha/beta hydrolase fold<br>protocat_pcaD: 3-oxoadipate enol-lactonase | 4.00E-111<br>2.30E-108<br>2.20E-21<br>1.60E-15<br>1.50E-12 |
| WP_158692714.1 | 4555238 | 4554431 | - | 268 | LinH<br>LinG | LinH<br>LinG | LinH (Fusion of LinE/LinG)<br>LinG Homolog | 3.30E-102<br>5.80E-92 |
| WP_158692715.1 | 4556567 | 4557806 | + | 412 | PF13814 | Replic_Relax | Replication-relaxation | 4.70E-11 |
| WP_009307930.1 | 4557729 | 4557903 | + | 58 | INFERRED GENE | - | - | - |
| WP_023548228.1 | 4558102 | 4557841 | - | 86 | NO PFAM MATCH | - | - | - |
| WP_078630635.1 | 4558673 | 4558337 | - | 111 | NO PFAM MATCH | - | - | - |

| | | | | | | | | |
|---|---|---|---|---|---|---|---|---|
| WP_009311981.1 | 4558887 | 4559573 | + | 228 | INFERRED GENE | - | - | - |
| WP_031225319.1 | 4559858 | 4560611 | + | 250 | NO PFAM MATCH | - | - | - |
| WP_078630636.1 | 4560967 | 4562497 | + | 509 | PF00665<br>PF09299 | rve<br>Mu-transpos_C | Integrase core domain<br>Mu transposase, C-terminal | 6.00E-11<br>3.10E-05 |

| index | leader | core | start | end | dir | score |
|---|---|---|---|---|---|---|
| 1 | VGCAWGLSCRGGRECWCCGDQRCE | SLASALSWALNARTTHPTSSLETV | 4545605 | 4545458 | - | 6 |
| 2 | VGAAAQAPDEDVADGGGSVARSAGA | GEWFGTVSAGGARGMGCALVRGCMG | 4548250 | 4548097 | - | 6 |
| 3 | VCRGGFVGRRAGGQVGAAA | QAPDEDVADGGGSVARSAGAGEWFGTVSAGGARGMGCALVRGCMG | 4548292 | 4548097 | - | 6 |
| 4 | VRCAPGRGACLGWVCRGGFVGRRAGGQVGAAA | QAPDEDVADGGGSVARSAGAGEWFGTVSAGGARGMGCALVRGCMG | 4548331 | 4548097 | - | 6 |
| 5 | MTCGCLGRDGVGCEGESPRVGVSVGRGPAVG | VTRVRLRWCDGPGSVAWGVWTPNAPENPRFR | 4548603 | 4548414 | - | 8 |
| 6 | VNNDERLDELQIVSAQSEILDETTVPVL | CTPAVAVGAVAAATALHSIW | 4555581 | 4555434 | - | 27 |
| 7 | MDSLTSIVNNDERLDELQIVSAQSEILDETTVPVL | CTPAVAVGAVAAATALHSIW | 4555602 | 4555434 | - | 29 |
| 8 | MFVHLSDVVNTALAVA | GVPTHAAVHCAVAALTH | 4555803 | 4555701 | - | 18 |
| 9 | MNITESVHGPLEPAMFVHLSDVVNTALAVA | GVPTHAAVHCAVAALTH | 4555845 | 4555701 | - | 18 |
| 10 | VNTSEEIGSLSMNITESVHGPLEPAMFVHLSDVVNTALAVA | GVPTHAAVHCAVAALTH | 4555878 | 4555701 | - | 20 |
| 11 | MVNTSEEIGSLSMNITESVHGPLEPAMFVHLSDVVNTALAVA | GVPTHAAVHCAVAALTH | 4555881 | 4555701 | - | 20 |
| 12 | VLEMVNTSEEIGSLSMNITESVHGPLEPAMFVHLSDVVNTALAVA | GVPTHAAVHCAVAALTH | 4555890 | 4555701 | - | 20 |
| 13 | MSPVLEMVNTSEEIGSLSMNITESVHGPLEPAMFVHLSDVVNTALAVA | GVPTHAAVHCAVAALTH | 4555899 | 4555701 | - | 20 |
| 14 | MESFRADATEKRSRMSPVLEMVNTS...SVHGPLEPAMFVHLSDVVNTALAVA | GVPTHAAVHCAVAALTH | 4555941 | 4555701 | - | 19 |
| 15 | LRDRQAPGSACHLQELSHGGVGCPYVTPIQSSRDL | RHRRTHNIRVRVFSINGGFHTGRVDNPFLTRRQST | 4556186 | 4555973 | - | 8 |
| 16 | MPAPLSSAALAKRLLDESDLGEGYTRTPQPPAQHDDMTVIGCP | ALEKRGSEVAVGCSLDFPHEAKSCARPTSIFRNTAAEPARHNR | 4558102 | 4557841 | - | 7 |

### Results for WP_086752480.1 [Streptomyces scabiei] back to top

previous - next

#### Architecture

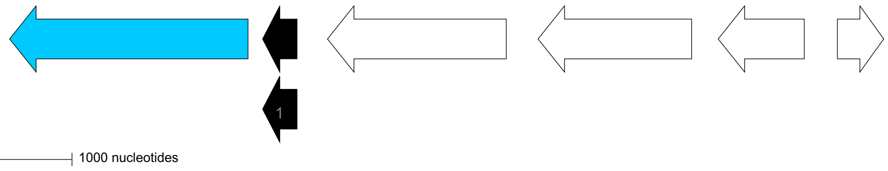

Link to nucleotide sequence

| Accession | start | end | direction | length (aa) | Pfam/HMM | name | description | E-value |
|---|---|---|---|---|---|---|---|---|
| WP_086752480.1 | 2001 | 81 | - | 639 | LinH<br>LinG<br>LinE<br>PF12697<br>PF12146 | LinH<br>LinG<br>LinE<br>Abhydrolase_6<br>Hydrolase_4 | LinH (Fusion of LinE/LinG)<br>LinG Homolog<br>LinE Homolog<br>Alpha/beta hydrolase family<br>Serine aminopeptidase, S33 | 1.40E-247<br>3.80E-164<br>2.30E-92<br>7.40E-17<br>5.50E-10 |
| WP_086752482.1 | 2393 | 2114 | - | 92 | LinA | LinA | LinA (linaridin precursor) | 9.90E-08 |
| WP_086752484.1 | 4071 | 2640 | - | 476 | TIGR02734<br>PF13450<br>TIGR03467<br>TIGR03315<br>TIGR00292 | TIGR02734<br>NAD_binding_8<br>TIGR03467<br>TIGR03315<br>TIGR00292 | crtI_fam: phytoene desaturase<br>NAD(P)-binding Rossmann-like domain<br>HpnE: squalene-associated FAD-dependent desaturase<br>Se_ygfK: putative selenate reductase, YgfK subunit<br>TIGR00292: thiazole biosynthesis enzyme | 6.20E-13<br>4.30E-11<br>1.50E-08<br>5.20E-08<br>1.90E-07 |
| WP_086752488.1 | 5562 | 4329 | - | 410 | PF07730 | HisKA_3 | Histidine kinase | 4.80E-14 |
| WP_086752490.1 | 6465 | 5775 | - | 229 | PF00072<br>PF00196<br>TIGR02875<br>TIGR03020<br>TIGR03541 | Response_reg<br>GerE<br>TIGR02875<br>TIGR03020<br>TIGR03541 | Response regulator receiver domain<br>Bacterial regulatory proteins, luxR family<br>spore_0_A: sporulation transcription factor Spo0A<br>EpsA: transcriptional regulator EpsA<br>reg_near_HchA: LuxR family transcriptional regulatory, chaperone HchA-associated | 1.50E-24<br>9.90E-19<br>1.80E-15<br>1.50E-13<br>1.90E-08 |
| WP_086752486.1 | 6732 | 7095 | + | 120 | PF00196<br>TIGR03541<br>TIGR03020<br>PF08281 | GerE<br>TIGR03541<br>TIGR03020<br>Sigma70_r4_2 | Bacterial regulatory proteins, luxR family<br>reg_near_HchA: LuxR family transcriptional regulatory, chaperone HchA-associated<br>EpsA: transcriptional regulator EpsA<br>Sigma-70, region 4 | 1.30E-15<br>2.90E-12<br>2.20E-09<br>1.10E-05 |

| index | leader | core | start | end | dir | score |
|---|---|---|---|---|---|---|
| 1 | MSSIENALNSVEIPVEGVVYVAARPTL | GTPRIARVGNRLGGLAQAAEGVGAI...ANNLAAEANAQNAAALAAVGGAPAA | 2393 | 2114 | - | 26 |

### Results for WP_086756723.1 [Streptomyces scabiei] back to top

previous - next

#### Architecture

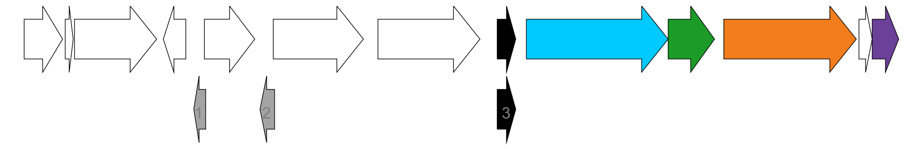

1000 nucleotides

Link to nucleotide sequence

| Accession | start | end | direction | length (aa) | Pfam/HMM | name | description | E-value |
|---|---|---|---|---|---|---|---|---|
| WP_004992705.1 | 0 | 541 | + | 180 | INFERRED GENE | - | - | - |
| WP_007166888.1 | 573 | 672 | + | 33 | INFERRED GENE | - | - | - |
| WP_086756716.1 | 702 | 1824 | + | 373 | PF07730 | HisKA_3 | Histidine kinase | 7.00E-12 |
| WP_086756718.1 | 2231 | 1922 | - | 102 | PF00196<br>TIGR03541<br>TIGR03020<br>PF08281<br>PF00486 | GerE<br>TIGR03541<br>TIGR03020<br>Sigma70_r4_2<br>Trans_reg_C | Bacterial regulatory proteins, luxR family<br>reg_near_HchA: LuxR family transcriptional regulatory, chaperone HchA-associated<br>EpsA: transcriptional regulator EpsA<br>Sigma-70, region 4<br>Transcriptional regulatory protein, C terminal | 3.00E-16<br>4.90E-12<br>3.60E-09<br>4.80E-06<br>3.80E-04 |
| WP_086756720.1 | 2484 | 3174 | + | 229 | PF00072<br>PF00196<br>TIGR02875<br>TIGR03020<br>TIGR03541 | Response_reg<br>GerE<br>TIGR02875<br>TIGR03020<br>TIGR03541 | Response regulator receiver domain<br>Bacterial regulatory proteins, luxR family<br>spore_0_A: sporulation transcription factor Spo0A<br>EpsA: transcriptional regulator EpsA<br>reg_near_HchA: LuxR family transcriptional regulatory, chaperone HchA-associated | 7.80E-25<br>2.90E-18<br>1.20E-15<br>6.10E-13<br>4.30E-08 |
| WP_086756729.1 | 3440 | 4673 | + | 410 | PF07730<br>PF13581 | HisKA_3<br>HATPase_c_2 | Histidine kinase<br>Histidine kinase-like ATPase domain | 7.50E-14<br>4.30E-04 |
| WP_086756722.1 | 4883 | 6299 | + | 471 | TIGR02734<br>PF13450<br>TIGR03467<br>TIGR03315<br>TIGR00292 | TIGR02734<br>NAD_binding_8<br>TIGR03467<br>TIGR03315<br>TIGR00292 | crtI_fam: phytoene desaturase<br>NAD(P)-binding Rossmann-like domain<br>HpnE: squalene-associated FAD-dependent desaturase<br>Se_ygfK: putative selenate reductase, YgfK subunit<br>TIGR00292: thiazole biosynthesis enzyme | 8.80E-14<br>3.60E-11<br>1.10E-08<br>6.70E-08<br>2.30E-07 |
| WP_086756731.1 | 6531 | 6789 | + | 85 | LinA | LinA | LinA (linaridin precursor) | 9.10E-04 |
| WP_086756723.1 | 6934 | 8890 | + | 651 | LinH<br>LinG<br>LinE<br>PF12697<br>PF12146 | LinH<br>LinG<br>LinE<br>Abhydrolase_6<br>Hydrolase_4 | LinH (Fusion of LinE/LinG)<br>LinG Homolog<br>LinE Homolog<br>Alpha/beta hydrolase family<br>Serine aminopeptidase, S33 | 1.80E-235<br>4.40E-124<br>4.40E-87<br>2.40E-16<br>5.20E-10 |
| WP_086756725.1 | 8886 | 9525 | + | 212 | LinL | LinL | LinL | 5.20E-74 |
| WP_086756726.1 | 9649 | 11488 | + | 612 | TIGR02204<br>TIGR02203<br>TIGR00958<br>TIGR03796<br>TIGR01846 | TIGR02204<br>TIGR02203<br>TIGR00958<br>TIGR03796<br>TIGR01846 | MsbA_rel: ABC transporter, permease/ATP-binding protein<br>MsbA_lipidA: lipid A export permease/ATP-binding protein MsbA<br>3a01208: antigen peptide transporter 2<br>NHLM_micro_ABC1: NHLM bacteriocin system ABC transporter, peptidase/ATP-binding protein<br>type_I_sec_HlyB: type I secretion system ATPase | 1.10E-105<br>4.40E-104<br>8.30E-92<br>2.40E-87<br>6.50E-83 |
| WP_086756728.1 | 11513 | 11705 | + | 63 | NO PFAM MATCH | - | - | - |
| WP_143673218.1 | 11701 | 12063 | + | 121 | PF13649<br>PF08241 | Methyltransf_25<br>Methyltransf_11 | Methyltransferase domain<br>Methyltransferase domain | 9.80E-07<br>9.50E-06 |

| index | leader | core | start | end | dir | score |
|---|---|---|---|---|---|---|
| 1 | VLLTRVSMPWITCPVRRSAEAGG | RTAVPNRAVGAFPAKDAPRADGPRSIGR | 2507 | 2351 | - | 6 |
| 2 | MSSSHSTPRAGYRAWWLSRRTGDRGRPWWCGWV | ARAESAASDEVVMEVQARHGHASPTGGASFCGE | 3460 | 3259 | - | 6 |
| 3 | MDLPVEGVILASARPTL | GTPRIGRVSGANALNALGGLADLAQ...NAAVDANQAVANAANAQAAAVGGAV | 6531 | 6789 | + | 29 |

### Results for KFG02471.1 [Streptomyces scabiei] back to top

previous - next

#### Architecture

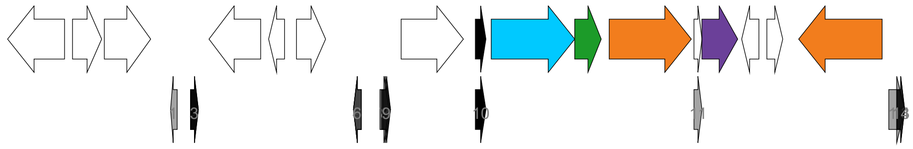

1000 nucleotides

Link to nucleotide sequence

| Accession | start | end | direction | length (aa) | Pfam/HMM | name | description | E-value |
|---|---|---|---|---|---|---|---|---|
| KFG02463.1 | 3637 | 2326 | - | 436 | PF04235<br>PF07786 | DUF418<br>DUF1624 | Protein of unknown function (DUF418)<br>Protein of unknown function (DUF1624) | 8.30E-11<br>4.80E-05 |
| KFG02464.1 | 3826 | 4504 | + | 225 | PF00072<br>PF00196<br>TIGR02875<br>TIGR01818<br>TIGR03020 | Response_reg<br>GerE<br>TIGR02875<br>TIGR01818<br>TIGR03020 | Response regulator receiver domain<br>Bacterial regulatory proteins, luxR family<br>spore_0_A: sporulation transcription factor Spo0A<br>ntrC: nitrogen regulation protein NR(I)<br>EpsA: transcriptional regulator EpsA | 2.00E-23<br>1.70E-17<br>1.90E-16<br>4.90E-13<br>1.50E-10 |
| KFG02465.1 | 4536 | 5622 | + | 361 | PF07730 | HisKA_3 | Histidine kinase | 5.90E-12 |

| | | | | | | | | |
|---|---|---|---|---|---|---|---|---|
| KFG02466.1 | 8132 | 6944 | - | 395 | PF00480 | ROK | ROK family | 2.00E-53 |
| | | | | | TIGR00744 | TIGR00744 | ROK_glcA_fam: ROK family protein (putative glucokinase) | 2.10E-40 |
| | | | | | PF12802 | MarR_2 | MarR family | 1.60E-06 |
| | | | | | PF13412 | HTH_24 | Winged helix-turn-helix DNA-binding | 1.50E-05 |
| | | | | | PF01047 | MarR | MarR family | 6.20E-04 |
| KFG02467.1 | 8702 | 8339 | - | 120 | PF00196 | GerE | Bacterial regulatory proteins, luxR family | 5.50E-16 |
| | | | | | TIGR03541 | TIGR03541 | reg_near_HchA: LuxR family transcriptional regulatory, chaperone HchA-associated | 1.50E-12 |
| | | | | | TIGR03020 | TIGR03020 | EpsA: transcriptional regulator EpsA | 1.60E-09 |
| | | | | | PF08281 | Sigma70_r4_2 | Sigma-70, region 4 | 9.20E-06 |
| | | | | | TIGR02937 | TIGR02937 | sigma70-ECF: RNA polymerase sigma factor, sigma-70 family | 9.50E-04 |
| KFG02468.1 | 8956 | 9646 | + | 229 | PF00072 | Response_reg | Response regulator receiver domain | 1.40E-24 |
| | | | | | PF00196 | GerE | Bacterial regulatory proteins, luxR family | 9.90E-19 |
| | | | | | TIGR02875 | TIGR02875 | spore_0_A: sporulation transcription factor Spo0A | 1.80E-15 |
| | | | | | TIGR03020 | TIGR03020 | EpsA: transcriptional regulator EpsA | 9.00E-14 |
| | | | | | TIGR03541 | TIGR03541 | reg_near_HchA: LuxR family transcriptional regulatory, chaperone HchA-associated | 2.10E-08 |
| KFG02469.1 | 11366 | 12797 | + | 476 | TIGR02734 | TIGR02734 | crtI_fam: phytoene desaturase | 6.00E-13 |
| | | | | | PF13450 | NAD_binding_8 | NAD(P)-binding Rossmann-like domain | 1.40E-11 |
| | | | | | TIGR03467 | TIGR03467 | HpnE: squalene-associated FAD-dependent desaturase | 1.40E-08 |
| | | | | | TIGR03315 | TIGR03315 | Se_ygfK: putative selenate reductase, YgfK subunit | 5.20E-08 |
| | | | | | PF01266 | DAO | FAD dependent oxidoreductase | 1.80E-07 |
| KFG02470.1 | 13086 | 13323 | + | 78 | LinA | LinA | LinA (linaridin precursor) | 1.20E-04 |
| KFG02471.1 | 13438 | 15358 | + | 639 | LinH | LinH | LinH (Fusion of LinE/LinG) | 2.10E-245 |
| | | | | | LinG | LinG | LinG Homolog | 7.50E-160 |
| | | | | | LinE | LinE | LinE Homolog | 1.30E-90 |
| | | | | | PF12697 | Abhydrolase_6 | Alpha/beta hydrolase family | 4.00E-16 |
| | | | | | PF12146 | Hydrolase_4 | Serine aminopeptidase, S33 | 1.90E-08 |
| KFG02472.1 | 15354 | 15975 | + | 206 | LinL | LinL | LinL | 2.20E-71 |
| KFG02473.1 | 16142 | 18032 | + | 629 | TIGR02204 | TIGR02204 | MsbA_rel: ABC transporter, permease/ATP-binding protein | 1.40E-104 |
| | | | | | TIGR02203 | TIGR02203 | MsbA_lipidA: lipid A export permease/ATP-binding protein MsbA | 7.70E-101 |
| | | | | | TIGR00958 | TIGR00958 | 3a01208: antigen peptide transporter 2 | 4.60E-92 |
| | | | | | TIGR03796 | TIGR03796 | NHLM_micro_ABC1: NHLM bacteriocin system ABC transporter, peptidase/ATP-binding protein | 4.30E-84 |
| | | | | | TIGR01846 | TIGR01846 | type_I_sec_HlyB: type I secretion system ATPase | 3.40E-79 |
| KFG02474.1 | 18100 | 18292 | + | 63 | NO PFAM MATCH | - | - | - |
| KFG02475.1 | 18288 | 19095 | + | 268 | PF13649 | Methyltransf_25 | Methyltransferase domain | 1.70E-15 |
| | | | | | PF08241 | Methyltransf_11 | Methyltransferase domain | 1.10E-12 |
| | | | | | PF13489 | Methyltransf_23 | Methyltransferase domain | 3.30E-08 |
| | | | | | PF13847 | Methyltransf_31 | Methyltransferase domain | 4.00E-07 |
| | | | | | PF08242 | Methyltransf_12 | Methyltransferase domain | 1.10E-06 |
| KFG02476.1 | 19595 | 19199 | - | 131 | PF00583 | Acetyltransf_1 | Acetyltransferase (GNAT) family | 6.00E-12 |
| | | | | | PF13673 | Acetyltransf_10 | Acetyltransferase (GNAT) domain | 1.40E-05 |
| | | | | | TIGR01575 | TIGR01575 | rimI: ribosomal-protein-alanine acetyltransferase | 1.30E-04 |
| | | | | | TIGR03448 | TIGR03448 | mycothiol_MshD: mycothiol synthase | 6.80E-04 |
| KFG02477.1 | 19776 | 20148 | + | 123 | PF00717 | Peptidase_S24 | Peptidase S24-like | 6.70E-04 |
| | | | | | TIGR02228 | TIGR02228 | sigpep_I_arch: signal peptidase I | 9.30E-04 |
| KFG02478.1 | 22419 | 20499 | - | 639 | TIGR02204 | TIGR02204 | MsbA_rel: ABC transporter, permease/ATP-binding protein | 1.50E-102 |
| | | | | | TIGR02203 | TIGR02203 | MsbA_lipidA: lipid A export permease/ATP-binding protein MsbA | 2.90E-98 |
| | | | | | TIGR01846 | TIGR01846 | type_I_sec_HlyB: type I secretion system ATPase | 4.80E-89 |
| | | | | | TIGR00958 | TIGR00958 | 3a01208: antigen peptide transporter 2 | 1.90E-85 |
| | | | | | TIGR03375 | TIGR03375 | type_I_sec_LssB: type I secretion system ATPase | 9.80E-78 |

| index | leader | core | start | end | dir | score |
|---|---|---|---|---|---|---|
| 1 | VSVPGTRSGRYTSFGAEFATDCGAGRPSP | RTAAGGPSPSTGGGSPSGTAEA | 6223 | 6067 | - | 6 |
| 2 | VTVNSREPCAGSACVTLSL | YTPRFSRFHRSRTFAGSRWIGRPVAAIAFCQAATSATLS | 6521 | 6698 | + | 9 |
| 3 | VNSREPCAGSACVTLSL | YTPRFSRFHRSRTFAGSRWIGRPVAAIAFCQAATSATLS | 6527 | 6698 | + | 9 |
| 4 | MGPTCWGELSW | ITPAACACAMMPVTWWATKSCRSRASCMRSASRTAATVRRRRASRVRR | 10452 | 10272 | - | 6 |
| 5 | LSRMGPTCWGELSW | ITPAACACAMMPVTWWATKSCRSRASCMRSASRTAATVRRRRASRVRR | 10461 | 10272 | - | 6 |
| 6 | MLSRMGPTCWGELSW | ITPAACACAMMPVTWWATKSCRSRASCMRSASRTAATVRRRRASRVRR | 10464 | 10272 | - | 6 |
| 7 | LTAGPADDGGWWLIAHFPVLP | EAPPEDMGKSEKSAASVSRRPAGRHAKPNAHTEPEPDMSWDAG | 10897 | 11092 | + | 6 |
| 8 | LVAHRPFPGTSGSPARGHGEVGEVRRFGVPPPG | RTPCQAECPYGTRTGHELGRGLVVRHRSVRGVN | 10926 | 11127 | + | 8 |
| 9 | VAHRPFPGTSGSPARGHGEVGEVRRFGVPPPG | RTPCQAECPYGTRTGHELGRGLVVRHRSVRGVN | 10929 | 11127 | + | 8 |
| 10 | MEIPVEGVVYVAARPTL | GTPRIARIGRIAQAAEGIGAVAAAA...ANNLAAEANAQNQAALAAVGGAPAA | 13086 | 13323 | + | 27 |
| 11 | MNDFLLIPSVVALGIVGFLIATRAVSAPAVI | GVALVVLWGACSQARSTIHARNTPSKKRRQHP | 18100 | 18292 | + | 6 |
| 12 | VIREHLRKPSSFANTAAERWFPVPA...GHRLRDHTSAPSAEPRPRAAGDPCR | SAPVHVRVRTRTSRSATSSSGCGRR...DLRKLVRVVRPVSMGSVRFSTSPSV | 22583 | 22943 | + | 6 |
| 13 | VRVRTRTSRSATSSSGCGRRALRRASLIRS | RLVLPGDLRKLVRVVRPVSMGSVRFSTSPSV | 22757 | 22943 | + | 8 |
| 14 | VRTRTSRSATSSSGCGRRALRRASLIRSR | LVLPGDLRKLVRVVRPVSMGSVRFSTSPSV | 22763 | 22943 | + | 8 |

### Results for WP_086801155.1 [Streptomyces scabiei] back to top

previous - next

#### Architecture

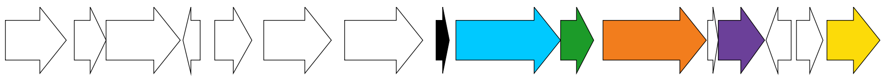

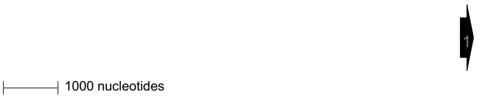

Link to nucleotide sequence

| Accession | start | end | direction | length (aa) | Pfam/HMM | name | description | E-value |
|---|---|---|---|---|---|---|---|---|
| WP_086801149.1 | 661 | 1783 | + | 373 | PF07730 | HisKA_3 | Histidine kinase | 1.60E-12 |
| WP_086801150.1 | 1915 | 2503 | + | 195 | TIGR02983<br>TIGR02937<br>PF08281<br>TIGR02954<br>PF04545 | TIGR02983<br>TIGR02937<br>Sigma70_r4_2<br>TIGR02954<br>Sigma70_r4 | SigE-fam_strep: RNA polymerase sigma-70 factor, sigma-E family<br>sigma70-ECF: RNA polymerase sigma factor, sigma-70 family<br>Sigma-70, region 4<br>Sig70_famx3: RNA polymerase sigma-70 factor, TIGR02954 family<br>Sigma-70, region 4 | 4.70E-54<br>4.30E-19<br>1.40E-16<br>2.00E-14<br>2.60E-12 |
| WP_086801151.1 | 2499 | 3855 | + | 451 | PF13360<br>PF01011<br>TIGR03300<br>TIGR03074<br>TIGR03075 | PQQ_2<br>PQQ<br>TIGR03300<br>TIGR03074<br>TIGR03075 | PQQ-like domain<br>PQQ enzyme repeat<br>assembly_YfgL: outer membrane assembly lipoprotein YfgL<br>PQQ_membr_DH: membrane-bound PQQ-dependent dehydrogenase, glucose/quinate/shikimate family<br>PQQ_enz_alc_DH: PQQ-dependent dehydrogenase, methanol/ethanol family | 1.00E-26<br>9.50E-14<br>6.70E-13<br>1.50E-07<br>7.00E-07 |
| WP_086801152.1 | 4230 | 3903 | - | 108 | PF00196<br>TIGR03541<br>TIGR03020<br>PF08281 | GerE<br>TIGR03541<br>TIGR03020<br>Sigma70_r4_2 | Bacterial regulatory proteins, luxR family<br>reg_near_HchA: LuxR family transcriptional regulatory, chaperone HchA-associated<br>EpsA: transcriptional regulator EpsA<br>Sigma-70, region 4 | 1.60E-15<br>1.70E-12<br>5.00E-09<br>2.20E-07 |
| WP_086801172.1 | 4485 | 5175 | + | 229 | PF00072<br>PF00196<br>TIGR02875<br>TIGR03020<br>TIGR03541 | Response_reg<br>GerE<br>TIGR02875<br>TIGR03020<br>TIGR03541 | Response regulator receiver domain<br>Bacterial regulatory proteins, luxR family<br>spore_0_A: sporulation transcription factor Spo0A<br>EpsA: transcriptional regulator EpsA<br>reg_near_HchA: LuxR family transcriptional regulatory, chaperone HchA-associated | 1.40E-24<br>9.70E-18<br>6.50E-16<br>9.50E-13<br>2.70E-08 |
| WP_086801153.1 | 5391 | 6621 | + | 409 | PF07730<br>PF13581 | HisKA_3<br>HATPase_c_2 | Histidine kinase<br>Histidine kinase-like ATPase domain | 1.30E-13<br>3.60E-04 |
| WP_086801154.1 | 6867 | 8298 | + | 476 | TIGR02734<br>PF13450<br>TIGR03467<br>TIGR00292<br>PF01266 | TIGR02734<br>NAD_binding_8<br>TIGR03467<br>TIGR00292<br>DAO | crtI_fam: phytoene desaturase<br>NAD(P)-binding Rossmann-like domain<br>HpnE: squalene-associated FAD-dependent desaturase<br>TIGR00292: thiazole biosynthesis enzyme<br>FAD dependent oxidoreductase | 1.00E-12<br>1.40E-11<br>2.70E-08<br>2.00E-07<br>2.80E-07 |
| WP_086801173.1 | 8546 | 8786 | + | 79 | LinA | LinA | LinA (linaridin precursor) | 2.10E-04 |
| WP_086801155.1 | 8900 | 10820 | + | 639 | LinH<br>LinG<br>LinE<br>PF12697<br>PF12146 | LinH<br>LinG<br>LinE<br>Abhydrolase_6<br>Hydrolase_4 | LinH (Fusion of LinE/LinG)<br>LinG Homolog<br>LinE Homolog<br>Alpha/beta hydrolase family<br>Serine aminopeptidase, S33 | 1.10E-<br>247<br>1.00E-<br>157<br>1.60E-92<br>4.60E-16<br>1.70E-08 |
| WP_086801156.1 | 10816 | 11437 | + | 206 | LinL | LinL | LinL | 5.00E-69 |
| WP_086801157.1 | 11600 | 13493 | + | 630 | TIGR02204<br>TIGR02203<br>TIGR00958<br>TIGR03796<br>TIGR03375 | TIGR02204<br>TIGR02203<br>TIGR00958<br>TIGR03796<br>TIGR03375 | MsbA_rel: ABC transporter, permease/ATP-binding protein<br>MsbA_lipidA: lipid A export permease/ATP-binding protein MsbA<br>3a01208: antigen peptide transporter 2<br>NHLM_micro_ABC1: NHLM bacteriocin system ABC transporter, peptidase/ATP-binding protein<br>type_I_sec_LssB: type I secretion system ATPase | 1.90E-<br>106<br>4.80E-<br>104<br>2.80E-92<br>8.50E-86<br>4.80E-81 |
| WP_086801158.1 | 13518 | 13710 | + | 63 | NO PFAM MATCH | - | - | - |
| WP_086801159.1 | 13706 | 14552 | + | 281 | PF13649<br>PF08241<br>PF08242<br>PF13489<br>TIGR01983 | Methyltransf_25<br>Methyltransf_11<br>Methyltransf_12<br>Methyltransf_23<br>TIGR01983 | Methyltransferase domain<br>Methyltransferase domain<br>Methyltransferase domain<br>Methyltransferase domain<br>UbiG: 3-demethylubiquinone-9 3-O-methyltransferase | 1.60E-15<br>1.90E-12<br>4.00E-07<br>5.00E-07<br>7.00E-05 |
| WP_086801160.1 | 15054 | 14577 | - | 158 | PF01638<br>PF03551 | HxlR<br>PadR | HxlR-like helix-turn-helix<br>Transcriptional regulator PadR-like family | 6.60E-20<br>5.20E-04 |
| WP_086801161.1 | 15143 | 15638 | + | 164 | TIGR00369<br>PF03061<br>TIGR02286<br>PF14539<br>PF13622 | TIGR00369<br>4HBT<br>TIGR02286<br>DUF4442<br>4HBT_3 | unchar_dom_1: uncharacterized domain 1<br>Thioesterase superfamily<br>PaaD: phenylacetic acid degradation protein PaaD<br>Domain of unknown function (DUF4442)<br>Thioesterase-like superfamily | 1.20E-26<br>7.20E-17<br>3.10E-09<br>2.50E-08<br>7.40E-05 |
| WP_086801162.1 | 15702 | 16674 | + | 323 | PF00561<br>PF12697<br>TIGR03056<br>TIGR02427<br>TIGR03343 | Abhydrolase_1<br>Abhydrolase_6<br>TIGR03056<br>TIGR02427<br>TIGR03343 | alpha/beta hydrolase fold<br>Alpha/beta hydrolase family<br>bchO_mg_che_rel: putative magnesium chelatase accessory protein<br>protocat_pcaD: 3-oxoadipate enol-lactonase<br>biphenyl_bphD: 2-hydroxy-6-oxo-6-phenylhexa-2,4-dienoate hydrolase | 1.70E-27<br>1.10E-22<br>4.10E-17<br>2.20E-16<br>9.20E-16 |

| index | leader | core | start | end | dir | score |
|---|---|---|---|---|---|---|
| 1 | MEIPVEGVVYVAARPTL | GTPRIARIGRIAQAAEGVGAIAAAA...NNLAAEANAQNAAALAAVGGTPPAA | 8546 | 8786 | + | 27 |

### Results for WP_086800803.1 [Streptomyces scabiei] back to top

previous - next

#### Architecture

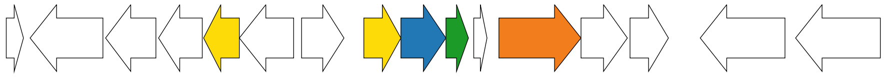

1000 nucleotides

Link to nucleotide sequence

| Accession | start | end | direction | length (aa) | Pfam/HMM | name | description | E-value |
|---|---|---|---|---|---|---|---|---|
| WP_006380350.1 | 4312 | 4714 | + | 134 | INFERRED GENE | - | - | - |
| WP_086800797.1 | 6611 | 4880 | - | 576 | TIGR00976<br>PF02129<br>PF08530<br>PF00326 | TIGR00976<br>Peptidase_S15<br>PepX_C<br>Peptidase_S9 | /NonD: hydrolase CocE/NonD family protein<br>X-Pro dipeptidyl-peptidase (S15 family)<br>X-Pro dipeptidyl-peptidase C-terminal non-catalytic domain<br>Prolyl oligopeptidase family | 1.00E-88<br>2.40E-52<br>1.10E-30<br>1.60E-06 |
| WP_086800798.1 | 7854 | 6669 | - | 394 | PF00144 | Beta-lactamase | Beta-lactamase | 2.50E-61 |
| WP_086800813.1 | 8951 | 7928 | - | 340 | PF00756<br>TIGR02821 | Esterase<br>TIGR02821 | Putative esterase<br>fghA_ester_D: S-formylglutathione hydrolase | 4.70E-30<br>2.40E-06 |
| WP_086800799.1 | 9831 | 9003 | - | 275 | TIGR02427<br>PF00561<br>TIGR03611<br>PF12146<br>TIGR03056 | TIGR02427<br>Abhydrolase_1<br>TIGR03611<br>Hydrolase_4<br>TIGR03056 | protocat_pcaD: 3-oxoadipate enol-lactonase<br>alpha/beta hydrolase fold<br>RutD: pyrimidine utilization protein D<br>Serine aminopeptidase, S33<br>bchO_mg_che_rel: putative magnesium chelatase accessory protein | 1.50E-27<br>1.70E-25<br>6.20E-23<br>6.40E-21<br>2.00E-16 |
| WP_086800800.1 | 11131 | 9847 | - | 427 | PF07690<br>TIGR00880<br>PF13347<br>TIGR00900<br>TIGR00792 | MFS_1<br>TIGR00880<br>MFS_2<br>TIGR00900<br>TIGR00792 | Major Facilitator Superfamily<br>2_A_01_02: multidrug resistance protein<br>MFS/sugar transport protein<br>2A0121: H+ Antiporter protein<br>gph: sugar (Glycoside-Pentoside-Hexuronide) transporter | 1.80E-27<br>9.00E-13<br>2.40E-11<br>1.60E-09<br>9.10E-07 |
| WP_086800801.1 | 11340 | 12336 | + | 331 | PF13377<br>TIGR01481<br>TIGR02417<br>TIGR02405<br>PF00356 | Peripla_BP_3<br>TIGR01481<br>TIGR02417<br>TIGR02405<br>LacI | Periplasmic binding protein-like domain<br>ccpA: catabolite control protein A<br>fruct_sucro_rep: D-fructose-responsive transcription factor<br>trehalos_R_Ecol: trehalose operon repressor<br>Bacterial regulatory proteins, lacI family | 1.10E-26<br>2.00E-21<br>2.90E-17<br>5.10E-15<br>2.20E-14 |
| WP_086800802.1 | 12821 | 13703 | + | 293 | LinG<br>LinH | LinG<br>LinH | LinG Homolog<br>LinH (Fusion of LinE/LinG) | 9.10E-99<br>4.70E-88 |
| WP_086800803.1 | 13699 | 14773 | + | 357 | LinE<br>LinH<br>PF00561<br>TIGR02427<br>PF12697 | LinE<br>LinH<br>Abhydrolase_1<br>TIGR02427<br>Abhydrolase_6 | LinE Homolog<br>LinH (Fusion of LinE/LinG)<br>alpha/beta hydrolase fold<br>protocat_pcaD: 3-oxoadipate enol-lactonase<br>Alpha/beta hydrolase family | 5.50E-80<br>1.50E-63<br>8.80E-17<br>7.50E-13<br>9.80E-13 |
| WP_086800804.1 | 14769 | 15309 | + | 179 | LinL | LinL | LinL | 3.30E-50 |
| WP_143674286.1 | 15411 | 15726 | + | 104 | NO PFAM MATCH | - | - | - |
| WP_086800806.1 | 16020 | 17961 | + | 646 | TIGR02203<br>TIGR02204<br>TIGR03375<br>TIGR01846<br>TIGR00958 | TIGR02203<br>TIGR02204<br>TIGR03375<br>TIGR01846<br>TIGR00958 | MsbA_lipidA: lipid A export permease/ATP-binding protein MsbA<br>MsbA_rel: ABC transporter, permease/ATP-binding protein<br>type_I_sec_LssB: type I secretion system ATPase<br>type_I_sec_HlyB: type I secretion system ATPase<br>3a01208: antigen peptide transporter 2 | 5.40E-72<br>1.60E-71<br>1.50E-64<br>2.30E-63<br>1.00E-62 |
| WP_086800807.1 | 17991 | 19083 | + | 363 | PF13469<br>PF00685 | Sulfotransfer_3<br>Sulfotransfer_1 | Sulfotransferase family<br>Sulfotransferase domain | 7.40E-21<br>4.60E-08 |
| WP_086800808.1 | 19151 | 20057 | + | 301 | TIGR03621<br>PF00296<br>TIGR03560<br>TIGR03619<br>TIGR03856 | TIGR03621<br>Bac_luciferase<br>TIGR03560<br>TIGR03619<br>TIGR03856 | F420_MSMEG_2516: probable F420-dependent oxidoreductase, MSMEG_2516 family<br>Luciferase-like monooxygenase<br>F420_Rv1855c: probable F420-dependent oxidoreductase, Rv1855c family<br>F420_Rv2161c: probable F420-dependent oxidoreductase, Rv2161c family<br>F420_MSMEG_2906: probable F420-dependent oxidoreductase, MSMEG_2906 family | 1.60E-67<br>2.70E-26<br>3.10E-23<br>9.80E-21<br>1.80E-16 |
| WP_086800810.1 | 22823 | 20798 | - | 674 | PF02836<br>PF02837<br>PF00703<br>PF13364<br>PF16355 | Glyco_hydro_2_C<br>Glyco_hydro_2_N<br>Glyco_hydro_2<br>BetaGal_dom4_5<br>DUF4982 | Glycosyl hydrolases family 2, TIM barrel domain<br>Glycosyl hydrolases family 2, sugar binding domain<br>Glycosyl hydrolases family 2<br>Beta-galactosidase jelly roll domain<br>Domain of unknown function (DUF4982) | 3.10E-29<br>9.20E-20<br>3.40E-12<br>7.10E-09<br>4.20E-07 |
| WP_086800811.1 | 25109 | 23066 | - | 680 | PF02449<br>PF08532<br>PF08533<br>PF01301 | Glyco_hydro_42<br>Glyco_hydro_42M<br>Glyco_hydro_42C<br>Glyco_hydro_35 | Beta-galactosidase<br>Beta-galactosidase trimerisation domain<br>Beta-galactosidase C-terminal domain<br>Glycosyl hydrolases family 35 | 6.40E-148<br>1.40E-58<br>6.70E-09<br>2.10E-05 |

| index | leader | core | start | end | dir | score |
|---|---|---|---|---|---|---|

### Results for WP_055697709.1 [Streptomyces silaceus] back to top

previous - next

#### Architecture

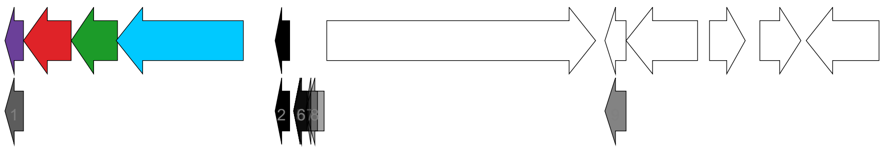

1000 nucleotides

Link to nucleotide sequence

| Accession | start | end | direction | length (aa) | Pfam/HMM | name | description | E-value |
|---|---|---|---|---|---|---|---|---|

| | | | | | | | | |
|---|---|---|---|---|---|---|---|---|
| WP_157872189.1 | 225 | 0 | - | 75 | TIGR02021<br>TIGR01983<br>PF13649<br>TIGR04345<br>TIGR04290 | TIGR02021<br>TIGR01983<br>Methyltransf_25<br>TIGR04345<br>TIGR04290 | BchM-ChlM: magnesium protoporphyrin O-methyltransferase<br>UbiG: 3-demethylubiquinone-9 3-O-methyltransferase<br>Methyltransferase domain<br>ovoA_Cterm: putative 4-mercaptohistidine N1-methyltranferase<br>meth_Rta_06860: methyltransferase, Rta_06860 family | 2.40E-10<br>9.10E-08<br>3.10E-07<br>6.10E-07<br>1.40E-06 |
| WP_055697678.1 | 790 | 217 | - | 190 | TIGR00521<br>PF02441<br>TIGR02113 | TIGR00521<br>Flavoprotein<br>TIGR02113 | coaBC_dfp: phosphopantothenoylcysteine decarboxylase / phosphopantothenate--cysteine ligase<br>Flavoprotein<br>coaC_strep: phosphopantothenoylcysteine decarboxylase | 5.10E-11<br>1.50E-09<br>1.50E-09 |
| WP_055697679.1 | 1347 | 786 | - | 186 | LinL | LinL | LinL | 5.40E-53 |
| WP_055697709.1 | 2854 | 1336 | - | 505 | LinH<br>LinE<br>LinG<br>PF12697<br>TIGR02427 | LinH<br>LinE<br>LinG<br>Abhydrolase_6<br>TIGR02427 | LinH (Fusion of LinE/LinG)<br>LinE Homolog<br>LinG Homolog<br>Alpha/beta hydrolase family<br>protocat_pcaD: 3-oxoadipate enol-lactonase | 5.20E-164<br>6.70E-70<br>1.70E-66<br>4.40E-10<br>9.10E-09 |
| WP_055697680.1 | 3412 | 3226 | - | 61 | LinA | LinA | LinA (linaridin precursor) | 2.00E-14 |
| WP_055697681.1 | 3849 | 7068 | + | 1072 | PF13424<br>PF17874<br>TIGR02917<br>PF00931<br>PF07719 | TPR_12<br>TPR_MalT<br>TIGR02917<br>NB-ARC<br>TPR_2 | Tetratricopeptide repeat<br>MalT-like TPR region<br>PEP_TPR_lipo: putative PEP-CTERM system TPR-repeat lipoprotein<br>NB-ARC domain<br>Tetratricopeptide repeat | 2.50E-19<br>2.50E-09<br>1.30E-08<br>1.40E-07<br>2.10E-07 |
| WP_079036394.1 | 7425 | 7182 | - | 80 | NO PFAM MATCH | - | - | - |
| WP_055697683.1 | 8287 | 7426 | - | 286 | TIGR03446<br>TIGR03445<br>PF02585<br>TIGR04001<br>TIGR04000 | TIGR03446<br>TIGR03445<br>PIG-L<br>TIGR04001<br>TIGR04000 | mycothiol_Mca: mycothiol conjugate amidase Mca<br>mycothiol_MshB: N-acetyl-1-D-myo-inositol-2-amino-2-deoxy-alpha-D-glucopyranoside deacetylase<br>GlcNAc-PI de-N-acetylase<br>thiol_BshB1: bacillithiol biosynthesis deacetylase BshB1<br>thiol_BshB2: bacillithiol biosynthesis deacetylase BshB2 | 2.70E-147<br>7.40E-52<br>1.80E-26<br>4.20E-13<br>4.40E-11 |
| WP_055697684.1 | 8423 | 8855 | + | 143 | PF14155 | DUF4307 | Domain of unknown function (DUF4307) | 4.80E-31 |
| WP_055697685.1 | 9026 | 9524 | + | 165 | TIGR01462<br>PF03449<br>TIGR01461<br>PF01272 | TIGR01462<br>GreA_GreB_N<br>TIGR01461<br>GreA_GreB | greA: transcription elongation factor GreA<br>Transcription elongation factor, N-terminal<br>greB: transcription elongation factor GreB<br>Transcription elongation factor, GreA/GreB, C-term | 1.40E-44<br>6.30E-25<br>8.00E-15<br>2.00E-13 |
| WP_055697686.1 | 10455 | 9585 | - | 289 | PF01061<br>TIGR01247<br>PF12698<br>TIGR00025<br>TIGR01248 | ABC2_membrane<br>TIGR01247<br>ABC2_membrane_3<br>TIGR00025<br>TIGR01248 | ABC-2 type transporter<br>drrB: daunorubicin resistance ABC transporter membrane protein<br>ABC-2 family transporter protein<br>Mtu_efflux: ABC transporter efflux protein, DrrB family<br>drrC: daunorubicin resistance protein C | 5.30E-30<br>1.30E-20<br>3.00E-18<br>4.70E-18<br>1.30E-10 |

| index | leader | core | start | end | dir | score |
|---|---|---|---|---|---|---|
| 1 | MSDPSVYDETAIEAYDLVSSMLSPGAGLAAWVSSHRP | LAGRSVLDLGCGTGVSSFALAEAGARVVAVDASRPSLD | 225 | 0 | - | 8 |
| 2 | MTRPDSAIRPEALAAQEFANTVLSGAAPGFHADCETPAM | ATPATPTAAQFVIQGSTICLVC | 3412 | 3226 | - | 34 |
| 3 | VLEALTVLDVLLGPCLRSVRNVEL | QTLCSTFTQSGRRGWQRSARRDHSLA | 3605 | 3452 | - | 7 |
| 4 | LRTVLEALTVLDVLLGPCLRSVRNVEL | QTLCSTFTQSGRRGWQRSARRDHSLA | 3614 | 3452 | - | 7 |
| 5 | VPSSLLRTVLEALTVLDVLLGPCLRSVRNVEL | QTLCSTFTQSGRRGWQRSARRDHSLA | 3629 | 3452 | - | 7 |
| 6 | MAGNSCHLCDVPSSLLRTVLEALTVLDVLLGPCLRSVRNVEL | QTLCSTFTQSGRRGWQRSARRDHSLA | 3659 | 3452 | - | 8 |
| 7 | VAGRGLAAAGRGLLGGRAGPSRGRP | SPLWLEIAVTSVTCQVPCCARCLKH | 3742 | 3589 | - | 6 |
| 8 | LPALHKCPAYGPGKRGRARIGDLPLAPVAGRGLAAAGR | GLLGGRAGPSRGRPSPLWLEIAVTSVTCQVPCCARCLKH | 3823 | 3589 | - | 6 |
| 9 | MNVHLAMTHLVPLAKEVDDDKVTPG...AVWGLMKSMNRHMGKVDFKEKEPVG | ATAGGDADRAADERTGEK | 7425 | 7182 | - | 7 |

### Results for CBG75367.1 [Streptomyces scabiei 87.22] back to top

previous - next

#### Architecture

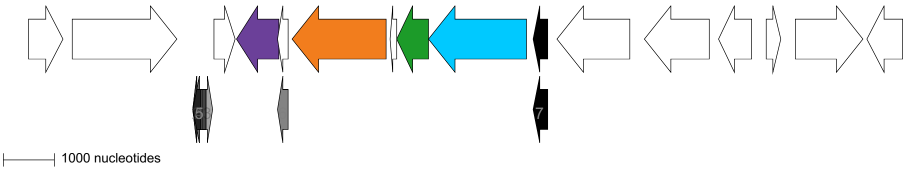

Link to nucleotide sequence

| Accession | start | end | direction | length (aa) | Pfam/HMM | name | description | E-value |
|---|---|---|---|---|---|---|---|---|
| CBG75359.1 | 9396048 | 9396729 | + | 226 | PF17837<br>PF01648 | 4PPT_N<br>ACPS | 4'-phosphopantetheinyl transferase N-terminal domain<br>4'-phosphopantetheinyl transferase superfamily | 1.50E-23<br>8.30E-09 |
| CBG75360.1 | 9396901 | 9398950 | + | 682 | PF03704<br>PF00931<br>PF00486<br>PF13191 | BTAD<br>NB-ARC<br>Trans_reg_C<br>AAA_16 | Bacterial transcriptional activator domain<br>NB-ARC domain<br>Transcriptional regulatory protein, C terminal<br>AAA ATPase domain | 4.20E-37<br>6.60E-15<br>1.10E-08<br>7.70E-05 |

| | | | | | | | | |
|---|---|---|---|---|---|---|---|---|
| CBG75361.1 | 9399700 | 9400096 | + | 131 | PF00583 | Acetyltransf_1 | Acetyltransferase (GNAT) family | 8.10E-13 |
| | | | | | PF13673 | Acetyltransf_10 | Acetyltransferase (GNAT) domain | 3.50E-06 |
| | | | | | PF13508 | Acetyltransf_7 | Acetyltransferase (GNAT) domain | 4.80E-05 |
| | | | | | TIGR01575 | TIGR01575 | rimI: ribosomal-protein-alanine acetyltransferase | 2.50E-04 |
| | | | | | PF14542 | Acetyltransf_CG | GCN5-related N-acetyl-transferase | 4.30E-04 |
| CBG75362.1 | 9400948 | 9400141 | - | 268 | PF13649 | Methyltransf_25 | Methyltransferase domain | 1.10E-14 |
| | | | | | PF08241 | Methyltransf_11 | Methyltransferase domain | 3.80E-11 |
| | | | | | PF13489 | Methyltransf_23 | Methyltransferase domain | 1.10E-08 |
| | | | | | PF08242 | Methyltransf_12 | Methyltransferase domain | 4.90E-06 |
| | | | | | TIGR01983 | TIGR01983 | UbiG: 3-demethylubiquinone-9 3-O-methyltransferase | 7.30E-06 |
| CBG75363.1 | 9401136 | 9400944 | - | 63 | NO PFAM MATCH | - | - | - |
| CBG75364.1 | 9403054 | 9401215 | - | 612 | TIGR02204 | TIGR02204 | MsbA_rel: ABC transporter, permease/ATP-binding protein | 9.80E-105 |
| | | | | | TIGR02203 | TIGR02203 | MsbA_lipidA: lipid A export permease/ATP-binding protein MsbA | 8.50E-102 |
| | | | | | TIGR00958 | TIGR00958 | 3a01208: antigen peptide transporter 2 | 4.20E-91 |
| | | | | | TIGR03796 | TIGR03796 | NHLM_micro_ABC1: NHLM bacteriocin system ABC transporter, peptidase/ATP-binding protein | 2.80E-85 |
| | | | | | TIGR01846 | TIGR01846 | type_I_sec_HlyB: type I secretion system ATPase | 1.40E-78 |
| CBG75365.1 | 9403273 | 9403147 | - | 41 | NO PFAM MATCH | - | - | - |
| CBG75366.1 | 9403896 | 9403275 | - | 206 | LinL | LinL | LinL | 7.60E-71 |
| CBG75367.1 | 9405812 | 9403892 | - | 639 | LinH | LinH | LinH (Fusion of LinE/LinG) | 1.40E-248 |
| | | | | | LinG | LinG | LinG Homolog | 9.60E-165 |
| | | | | | LinE | LinE | LinE Homolog | 3.10E-92 |
| | | | | | PF12697 | Abhydrolase_6 | Alpha/beta hydrolase family | 2.60E-16 |
| | | | | | PF12146 | Hydrolase_4 | Serine aminopeptidase, S33 | 3.90E-09 |
| CBG75368.1 | 9406220 | 9405953 | - | 88 | LinA | LinA | LinA (linaridin precursor) | 3.00E-08 |
| CBG75369.1 | 9407830 | 9406399 | - | 476 | TIGR02734 | TIGR02734 | crtI_fam: phytoene desaturase | 7.60E-12 |
| | | | | | PF13450 | NAD_binding_8 | NAD(P)-binding Rossmann-like domain | 1.40E-10 |
| | | | | | TIGR03467 | TIGR03467 | HpnE: squalene-associated FAD-dependent desaturase | 5.10E-08 |
| | | | | | TIGR00292 | TIGR00292 | TIGR00292: thiazole biosynthesis enzyme | 1.00E-07 |
| | | | | | TIGR03315 | TIGR03315 | Se_ygfK: putative selenate reductase, YgfK subunit | 4.00E-07 |
| CBG75370.1 | 9409395 | 9408135 | - | 419 | PF07730 | HisKA_3 | Histidine kinase | 1.30E-13 |
| CBG75371.1 | 9410217 | 9409566 | - | 216 | PF00072 | Response_reg | Response regulator receiver domain | 9.60E-24 |
| | | | | | PF00196 | GerE | Bacterial regulatory proteins, luxR family | 2.80E-18 |
| | | | | | TIGR02875 | TIGR02875 | spore_0_A: sporulation transcription factor Spo0A | 4.40E-14 |
| | | | | | TIGR03020 | TIGR03020 | EpsA: transcriptional regulator EpsA | 2.00E-13 |
| | | | | | TIGR03541 | TIGR03541 | reg_near_HchA: LuxR family transcriptional regulatory, chaperone HchA-associated | 1.40E-08 |
| CBG75372.1 | 9410501 | 9410801 | + | 99 | PF00196 | GerE | Bacterial regulatory proteins, luxR family | 7.60E-16 |
| | | | | | TIGR03541 | TIGR03541 | reg_near_HchA: LuxR family transcriptional regulatory, chaperone HchA-associated | 1.40E-11 |
| | | | | | TIGR03020 | TIGR03020 | EpsA: transcriptional regulator EpsA | 4.40E-09 |
| | | | | | PF08281 | Sigma70_r4_2 | Sigma-70, region 4 | 5.80E-06 |
| CBG75373.1 | 9411084 | 9412413 | + | 442 | PF02333 | Phytase | Phytase | 4.30E-52 |
| CBG75374.1 | 9413178 | 9412476 | - | 233 | PF07729 | FCD | FCD domain | 8.70E-23 |
| | | | | | PF00392 | GntR | Bacterial regulatory proteins, gntR family | 5.20E-18 |
| | | | | | TIGR03338 | TIGR03338 | phnR_burk: phosphonate utilization associated transcriptional regulator | 6.60E-18 |
| | | | | | TIGR02325 | TIGR02325 | C_P_lyase_phnF: phosphonate metabolism transcriptional regulator PhnF | 2.60E-11 |
| | | | | | TIGR02018 | TIGR02018 | his_ut_repres: histidine utilization repressor | 4.70E-11 |

| index | leader | core | start | end | dir | score |
|---|---|---|---|---|---|---|
| 1 | VDHARKRVQISNNRGHVNGWTGHG | SVFGICVTVDGAARPGTTGKTRPGG | 9399416 | 9399266 | - | 6 |
| 2 | VSSVDHARKRVQISNNRGHVNGWTGH | GSVFGICVTVDGAARPGTTGKTRPGG | 9399425 | 9399266 | - | 7 |
| 3 | MSPASTFVGSTGATVTCWRLRISGTIEEPLGRNVTV | APPATRSATMSKAPMRGAWQISAHSPTD | 9399465 | 9399660 | + | 6 |
| 4 | VPLIRSRQQVTVAPVDPTKVEAGDIVLARVAGTVYLHLVSSVDHA | RKRVQISNNRGHVNGWTGHGSVFGICVTVDGAARPGTTGKTRPGG | 9399539 | 9399266 | - | 6 |
| 5 | MVPLIRSRQQVTVAPVDPTKVEAGDIVLARVAGTVYLHLVSSVDH | ARKRVQISNNRGHVNGWTGHGSVFGICVTVDGAARPGTTGKTRPGG | 9399542 | 9399266 | - | 6 |
| 6 | MNDFLLIPSVVALGIVGFLIATRAVSTPAVI | GVALVVLWGACSQARSTVHPRHTSSKKRRQHP | 9401136 | 9400944 | - | 7 |
| 7 | MSSIENALNSVEIPVEGVVYVAARPTL | GTPRIARIGRIAQAAEGIGAIAAAA...ANNLAAEANAQNAAALAAVGGAPAS | 9406220 | 9405953 | - | 27 |

### Results for PVE04739.1 [Streptomyces scopuliridis RB72] back to top

previous - next

#### Architecture

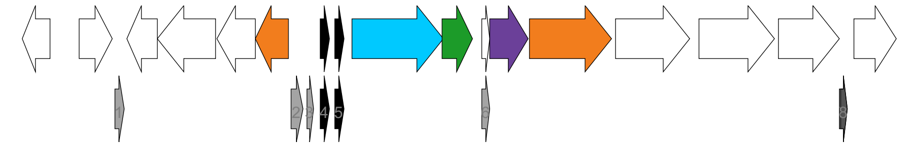

1000 nucleotides

Link to nucleotide sequence

| Accession | start | end | direction | length (aa) | Pfam/HMM | name | description | E-value |
|---|---|---|---|---|---|---|---|---|
| PVE04731.1 | 1848 | 1281 | - | 188 | PF01569 | PAP2 | PAP2 superfamily | 3.20E-04 |
| PVE04732.1 | 2431 | 3094 | + | 220 | PF05719 | GPP34 | Golgi phosphoprotein 3 (GPP34) | 7.90E-48 |

| | | | | | | | | |
|---|---|---|---|---|---|---|---|---|
| PVE04733.1 | 3999 | 3393 | - | 201 | PF00072<br>TIGR02875<br>PF00196<br>TIGR02154<br>TIGR02956 | Response_reg<br>TIGR02875<br>GerE<br>TIGR02154<br>TIGR02956 | Response regulator receiver domain<br>spore_0_A: sporulation transcription factor Spo0A<br>Bacterial regulatory proteins, luxR family<br>PhoB: phosphate regulon transcriptional regulatory protein PhoB<br>TMAO_torS: TMAO reductase sytem sensor TorS | 1.20E-24<br>1.90E-16<br>4.90E-13<br>4.20E-11<br>4.60E-10 |
| PVE04734.1 | 5183 | 3998 | - | 394 | PF07730 | HisKA_3 | Histidine kinase | 2.80E-14 |
| PVE04735.1 | 5981 | 5213 | - | 255 | PF01061<br>PF12698<br>TIGR03062 | ABC2_membrane<br>ABC2_membrane_3<br>TIGR03062 | ABC-2 type transporter<br>ABC-2 family transporter protein<br>pip_yhgE_Cterm: YhgE/Pip C-terminal domain | 6.70E-13<br>1.80E-08<br>8.10E-06 |
| PVE04736.1 | 6658 | 5977 | - | 226 | TIGR01188<br>TIGR03864<br>TIGR01288<br>TIGR03740<br>TIGR03522 | TIGR01188<br>TIGR03864<br>TIGR01288<br>TIGR03740<br>TIGR03522 | drrA: daunorubicin resistance ABC transporter, ATP-binding protein<br>PQQ_ABC_ATP: ABC transporter, ATP-binding subunit, PQQ-dependent alcohol dehydrogenase system<br>nodI: nodulation ABC transporter NodI<br>galliderm_ABC: lantibiotic protection ABC transporter, ATP-binding subunit<br>GldA_ABC_ATP: gliding motility-associated ABC transporter ATP-binding subunit GldA | 2.70E-41<br>1.10E-25<br>1.90E-25<br>1.00E-22<br>1.50E-20 |
| PVE04737.1 | 7285 | 7489 | + | 67 | LinA | LinA | LinA (linaridin precursor) | 1.40E-13 |
| PVE04738.1 | 7590 | 7776 | + | 61 | LinA | LinA | LinA (linaridin precursor) | 1.10E-15 |
| PVE04739.1 | 7930 | 9766 | + | 611 | LinH<br>LinG<br>LinE<br>PF12697<br>PF12146 | LinH<br>LinG<br>LinE<br>Abhydrolase_6<br>Hydrolase_4 | LinH (Fusion of LinE/LinG)<br>LinG Homolog<br>LinE Homolog<br>Alpha/beta hydrolase family<br>Serine aminopeptidase, S33 | 8.70E-253<br>2.60E-122<br>2.40E-100<br>4.40E-16<br>2.10E-10 |
| PVE04740.1 | 9762 | 10371 | + | 202 | LinL | LinL | LinL | 6.40E-69 |
| PVE04741.1 | 10549 | 10717 | + | 55 | NO PFAM MATCH | - | - | - |
| PVE04742.1 | 10713 | 11475 | + | 253 | PF13649<br>PF08241<br>PF13489<br>PF08242<br>PF13847 | Methyltransf_25<br>Methyltransf_11<br>Methyltransf_23<br>Methyltransf_12<br>Methyltransf_31 | Methyltransferase domain<br>Methyltransferase domain<br>Methyltransferase domain<br>Methyltransferase domain<br>Methyltransferase domain | 2.40E-15<br>8.40E-12<br>1.00E-08<br>6.00E-08<br>1.30E-07 |
| PVE04743.1 | 11504 | 13175 | + | 556 | TIGR02204<br>TIGR02203<br>TIGR00958<br>TIGR01846<br>TIGR03375 | TIGR02204<br>TIGR02203<br>TIGR00958<br>TIGR01846<br>TIGR03375 | MsbA_rel: ABC transporter, permease/ATP-binding protein<br>MsbA_lipidA: lipid A export permease/ATP-binding protein MsbA<br>3a01208: antigen peptide transporter 2<br>type_I_sec_HlyB: type I secretion system ATPase<br>type_I_sec_LssB: type I secretion system ATPase | 1.00E-108<br>1.60E-103<br>5.60E-96<br>9.10E-90<br>9.90E-88 |
| PVE04744.1 | 13239 | 14760 | + | 506 | TIGR02734<br>PF13450<br>PF01593<br>TIGR03467<br>TIGR00292 | TIGR02734<br>NAD_binding_8<br>Amino_oxidase<br>TIGR03467<br>TIGR00292 | crtI_fam: phytoene desaturase<br>NAD(P)-binding Rossmann-like domain<br>Flavin containing amine oxidoreductase<br>HpnE: squalene-associated FAD-dependent desaturase<br>TIGR00292: thiazole biosynthesis enzyme | 1.70E-11<br>4.30E-10<br>2.90E-09<br>8.90E-09<br>1.40E-07 |
| PVE04745.1 | 14941 | 16447 | + | 501 | TIGR01840<br>PF10503<br>PF00326 | TIGR01840<br>Esterase_phd<br>Peptidase_S9 | esterase_phb: esterase, PHB depolymerase family<br>Esterase PHB depolymerase<br>Prolyl oligopeptidase family | 2.10E-75<br>6.90E-41<br>4.70E-05 |
| PVE04746.1 | 16552 | 17764 | + | 403 | TIGR04458<br>PF00067 | TIGR04458<br>p450 | CYP450_TxtE: 4-nitrotryptophan synthase<br>Cytochrome P450 | 7.40E-40<br>5.20E-38 |
| PVE04747.1 | 18058 | 18925 | + | 288 | PF01370<br>TIGR03466<br>TIGR01777<br>PF13460<br>PF05368 | Epimerase<br>TIGR03466<br>TIGR01777<br>NAD_binding_10<br>NmrA | NAD dependent epimerase/dehydratase family<br>HpnA: hopanoid-associated sugar epimerase<br>yfcH: TIGR01777 family protein<br>NAD(P)H-binding<br>NmrA-like family | 1.80E-17<br>3.70E-13<br>5.60E-11<br>6.00E-09<br>1.30E-08 |

| index | leader | core | start | end | dir | score |
|---|---|---|---|---|---|---|
| 1 | VPERRATAPARASPARASPARASPARASPARASPVRTSPVRTTAPVP | TTAPVRPPEPTYGGG | 3158 | 3347 | + | 6 |
| 2 | VRAEKANAATGGAGEPFEEFDRRGLSCAICPEQSEYFALRD | REGDAPNGGEAPLVIGPVSPPKVCDLDHIFCSIGIHGRNRRR | 6703 | 6955 | + | 6 |
| 3 | MSVDGRSHSEYHCDLRHTGLVGA | GAFIQPHGRGIAASVAEVYPRITH | 7021 | 7165 | + | 6 |
| 4 | MVEIVSYFNEKLDLGAIDQGESNSEQSPVL | ATPAAALAAIGGAKVVGAFVGGATLVGGAYALGKAVG | 7285 | 7489 | + | 30 |
| 5 | MNNIIEAVNEMDIPAEGKVFPDATPVL | ATPGAFLGGVAVSSAVIGAFELGRGGGSAPATQL | 7590 | 7776 | + | 29 |
| 6 | MNKTPVILCALAAIAVLVFLAIGDVINYPFAAGI | GTALVLAAGLELHRNRKRSHQ | 10549 | 10717 | + | 6 |
| 7 | VTAGDRLKRPPVRPWTGSSRPGRRAADR | RQASGPRAITRSADPAANPLTWTDTRRA | 17770 | 17941 | + | 7 |
| 8 | VRPWTGSSRPGRRAADRRQASG | PRAITRSADPAANPLTWTDTRRA | 17803 | 17941 | + | 6 |

### Results for RXS62043.1 [Streptomyces sioyaensis] back to top

previous - next

#### Architecture

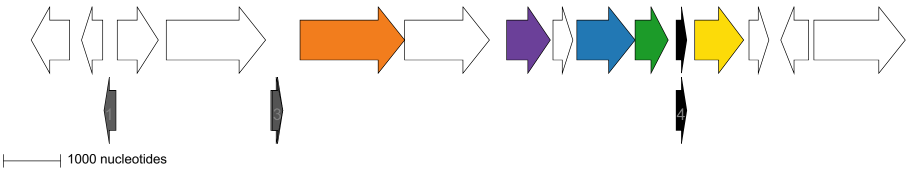

Link to nucleotide sequence

| Accession | start | end | direction | length (aa) | Pfam/HMM | name | description | E-value |
|---|---|---|---|---|---|---|---|---|
| RXS62036.1 | 12051 | 11388 | - | 220 | TIGR00949<br>PF01810<br>TIGR00948 | TIGR00949<br>LysE<br>TIGR00948 | 2A76: homoserine/Threonine efflux protein<br>LysE type translocator<br>2a75: L-lysine exporter | 8.70E-45<br>4.90E-44<br>4.90E-16 |
| RXS62037.1 | 12645 | 12276 | - | 122 | PF01906 | YbjQ_1 | Putative heavy-metal-binding | 1.40E-33 |
| RXS62038.1 | 12896 | 13604 | + | 235 | PF09335 | SNARE_assoc | SNARE associated Golgi protein | 3.10E-24 |
| RXS62039.1 | 13744 | 15478 | + | 577 | PF06738<br>PF12821 | ThrE<br>ThrE_2 | Putative threonine/serine exporter<br>Threonine/Serine exporter, ThrE | 9.80E-71<br>8.90E-19 |
| RXS62049.1 | 16085 | 17912 | + | 608 | TIGR02204<br>TIGR02203<br>TIGR00958<br>TIGR03375<br>TIGR03796 | TIGR02204<br>TIGR02203<br>TIGR00958<br>TIGR03375<br>TIGR03796 | MsbA_rel: ABC transporter, permease/ATP-binding protein<br>MsbA_lipidA: lipid A export permease/ATP-binding protein MsbA<br>3a01208: antigen peptide transporter 2<br>type_I_sec_LssB: type I secretion system ATPase<br>NHLM_micro_ABC1: NHLM bacteriocin system ABC transporter, peptidase/ATP-binding protein | 2.50E-99<br>8.60E-96<br>3.10E-89<br>1.50E-80<br>2.10E-80 |
| RXS62040.1 | 17908 | 19390 | + | 493 | TIGR02734<br>PF13450<br>TIGR03467<br>TIGR00292<br>PF01593 | TIGR02734<br>NAD_binding_8<br>TIGR03467<br>TIGR00292<br>Amino_oxidase | crtI_fam: phytoene desaturase<br>NAD(P)-binding Rossmann-like domain<br>HpnE: squalene-associated FAD-dependent desaturase<br>TIGR00292: thiazole biosynthesis enzyme<br>Flavin containing amine oxidoreductase | 1.20E-08<br>1.70E-08<br>4.30E-08<br>7.60E-08<br>2.00E-07 |
| RXS62041.1 | 19707 | 20466 | + | 252 | PF13649<br>PF08241<br>PF13489<br>PF08242<br>PF13847 | Methyltransf_25<br>Methyltransf_11<br>Methyltransf_23<br>Methyltransf_12<br>Methyltransf_31 | Methyltransferase domain<br>Methyltransferase domain<br>Methyltransferase domain<br>Methyltransferase domain<br>Methyltransferase domain | 5.00E-17<br>4.10E-12<br>1.80E-10<br>6.40E-10<br>9.90E-09 |
| RXS62042.1 | 20508 | 20856 | + | 115 | PF13518<br>PF01527<br>PF13551<br>PF13011<br>PF13384 | HTH_28<br>HTH_Tnp_1<br>HTH_29<br>LZ_Tnp_IS481<br>HTH_23 | Helix-turn-helix domain<br>Transposase<br>Winged helix-turn helix<br>leucine-zipper of insertion element IS481<br>Homeodomain-like domain | 4.20E-09<br>7.10E-06<br>7.80E-06<br>1.70E-04<br>4.80E-04 |
| RXS62043.1 | 20928 | 21948 | + | 339 | LinE<br>LinH<br>PF12697<br>TIGR02427<br>PF00561 | LinE<br>LinH<br>Abhydrolase_6<br>TIGR02427<br>Abhydrolase_1 | LinE Homolog<br>LinH (Fusion of LinE/LinG)<br>Alpha/beta hydrolase family<br>protocat_pcaD: 3-oxoadipate enol-lactonase<br>alpha/beta hydrolase fold | 4.00E-101<br>2.20E-51<br>2.70E-14<br>4.30E-09<br>7.70E-08 |
| RXS62044.1 | 21944 | 22532 | + | 195 | LinL | LinL | LinL | 1.90E-55 |
| RXS62045.1 | 22658 | 22841 | + | 60 | LinA | LinA | LinA (linaridin precursor) | 7.50E-14 |
| RXS62046.1 | 22986 | 23841 | + | 284 | LinG<br>LinH<br>TIGR04033 | LinG<br>LinH<br>TIGR04033 | LinG Homolog<br>LinH (Fusion of LinE/LinG)<br>export_SdpB: antimicrobial peptide system protein, SdpB family | 3.00E-111<br>9.10E-79<br>2.30E-04 |
| RXS62047.1 | 23932 | 24292 | + | 119 | NO PFAM MATCH | - | - | - |
| RXS62050.1 | 24974 | 24479 | - | 164 | PF00719 | Pyrophosphatase | Inorganic pyrophosphatase | 2.50E-53 |
| RXS62048.1 | 25060 | 26668 | + | 535 | PF02113<br>TIGR00666 | Peptidase_S13<br>TIGR00666 | D-Ala-D-Ala carboxypeptidase 3 (S13) family<br>PBP4: D-alanyl-D-alanine carboxypeptidase/D-alanyl-D-alanine-endopeptidase | 7.70E-76<br>6.70E-57 |

| index | leader | core | start | end | dir | score |
|---|---|---|---|---|---|---|
| 1 | VRCDERSDGGGGACAAQSYQRRTARRGSRRAHARPPDGPPWFQAGA | RTPTRRPPTAGRAPGAG | 12862 | 12670 | - | 8 |
| 2 | VWPTRSVRGPAAAFKGAVSNPQAPIRCEGSTSP | RAAPFSKAAYQRDPLRTLRATGQPWPAPGERAK | 15581 | 15782 | + | 7 |
| 3 | VRGPAAAFKGAVSNPQAPIRCEGSTSPRAA | PFSKAAYQRDPLRTLRATGQPWPAPGERAK | 15599 | 15782 | + | 6 |
| 4 | MSVVADFANTELADVTPGRIGNDATPTM | LTPLATLATPEGVAVTSATAYALHEVTKDIQR | 22658 | 22841 | + | 32 |

### Results for QBA31012.1 [Streptomyces sp.] back to top

previous - next

#### Architecture

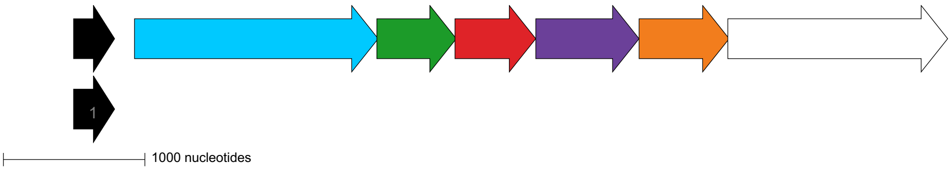

Link to nucleotide sequence

| Accession | start | end | direction | length (aa) | Pfam/HMM | name | description | E-value |
|---|---|---|---|---|---|---|---|---|
| QBA31011.1 | 0 | 288 | + | 95 | LinA | LinA | LinA (linaridin precursor) | 5.00E-14 |
| QBA31012.1 | 431 | 2153 | + | 573 | LinH<br>LinG<br>LinE<br>PF12697<br>TIGR02427 | LinH<br>LinG<br>LinE<br>Abhydrolase_6<br>TIGR02427 | LinH (Fusion of LinE/LinG)<br>LinG Homolog<br>LinE Homolog<br>Alpha/beta hydrolase family<br>protocat_pcaD: 3-oxoadipate enol-lactonase | 4.90E-169<br>4.90E-74<br>3.50E-70<br>1.50E-09<br>2.60E-08 |
| QBA31008.1 | 2142 | 2703 | + | 186 | LinL | LinL | LinL | 2.50E-52 |
| QBA31013.1 | 2699 | 3272 | + | 190 | TIGR00521<br>TIGR02113<br>PF02441 | TIGR00521<br>TIGR02113<br>Flavoprotein | coaBC_dfp: phosphopantothenoylcysteine decarboxylase / phosphopantothenate--cysteine ligase<br>coaC_strep: phosphopantothenoylcysteine decarboxylase<br>Flavoprotein | 9.20E-12<br>2.60E-10<br>1.80E-09 |

| | | | | | | | | |
|---|---|---|---|---|---|---|---|---|
| QBA31010.1 | 3264 | 4002 | + | 245 | PF13649<br>PF08241<br>PF13847<br>TIGR01934<br>PF08242 | Methyltransf_25<br>Methyltransf_11<br>Methyltransf_31<br>TIGR01934<br>Methyltransf_12 | Methyltransferase domain<br>Methyltransferase domain<br>Methyltransferase domain<br>MenG_MenH_UbiE: ubiquinone/menaquinone biosynthesis methyltransferase<br>Methyltransferase domain | 1.30E-16<br>2.40E-15<br>5.30E-11<br>9.20E-11<br>2.70E-10 |
| QBA31009.1 | 3998 | 4631 | + | 210 | TIGR02857<br>TIGR01189<br>TIGR02673<br>TIGR03608<br>TIGR02982 | TIGR02857<br>TIGR01189<br>TIGR02673<br>TIGR03608<br>TIGR02982 | CydD: thiol reductant ABC exporter, CydD subunit<br>ccmA: heme ABC exporter, ATP-binding protein CcmA<br>FtsE: cell division ATP-binding protein FtsE<br>L_ocin_972_ABC: putative bacteriocin export ABC transporter, lactococcin 972 group<br>heterocyst_DevA: ABC exporter ATP-binding subunit, DevA family | 1.80E-28<br>3.80E-27<br>7.00E-27<br>1.60E-23<br>6.30E-23 |
| QBA31007.1 | 4627 | 6178 | + | 516 | NO PFAM MATCH | - | - | - |

| index | leader | core | start | end | dir | score |
|---|---|---|---|---|---|---|
| 1 | MGAGDVLATFGTPRPFARIDHRRKV...AQEFANTALGAATPGFHADCETPAM | ATPATPTAAQFVIQGSTICLVC | 0 | 288 | + | 35 |

### Results for WP_161989239.1 [Streptomyces sioyaensis] back to top

previous - next

#### Architecture

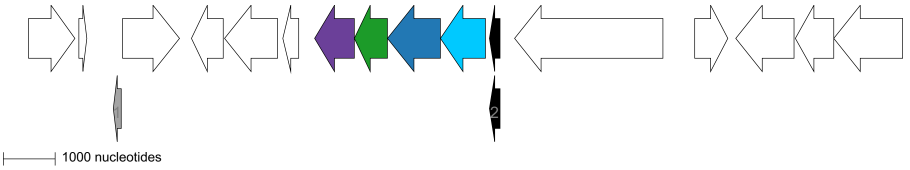

Link to nucleotide sequence

| Accession | start | end | direction | length (aa) | Pfam/HMM | name | description | E-value |
|---|---|---|---|---|---|---|---|---|
| WP_129244274.1 | 85662 | 86556 | + | 297 | PF00480<br>TIGR00744 | ROK<br>TIGR00744 | ROK family<br>ROK_glcA_fam: ROK family protein (putative glucokinase) | 2.50E-20<br>3.20E-19 |
| WP_154074006.1 | 86636 | 86789 | + | 50 | NO PFAM MATCH | - | - | - |
| WP_129244276.1 | 87482 | 88571 | + | 362 | TIGR00092<br>PF06071<br>PF01926<br>TIGR02729<br>TIGR03156 | TIGR00092<br>YchF-GTPase_C<br>MMR_HSR1<br>TIGR02729<br>TIGR03156 | TIGR00092: GTP-binding protein YchF<br>Protein of unknown function (DUF933)<br>50S ribosome-binding GTPase<br>Obg_CgtA: Obg family GTPase CgtA<br>GTP_HflX: GTP-binding protein HflX | 1.50E-131<br>3.80E-39<br>1.50E-22<br>2.90E-21<br>6.80E-11 |
| WP_129244278.1 | 89422 | 88804 | - | 205 | NO PFAM MATCH | - | - | - |
| WP_161989244.1 | 90465 | 89436 | - | 342 | TIGR03930<br>PF06013 | TIGR03930<br>WXG100 | WXG100_ESAT6: WXG100 family type VII secretion target<br>Proteins of 100 residues with WXG | 6.40E-09<br>8.10E-09 |
| WP_129244282.1 | 90884 | 90569 | - | 104 | PF10824<br>TIGR03930<br>PF06013<br>PF11932 | T7SS_ESX_EspC<br>TIGR03930<br>WXG100<br>DUF3450 | Excreted virulence factor EspC, type VII ESX diderm<br>WXG100_ESAT6: WXG100 family type VII secretion target<br>Proteins of 100 residues with WXG<br>Protein of unknown function (DUF3450) | 1.80E-09<br>5.30E-06<br>1.10E-04<br>1.30E-04 |
| WP_129244284.1 | 91954 | 91186 | - | 255 | PF13649<br>PF13489<br>PF08241<br>PF08242<br>TIGR03534 | Methyltransf_25<br>Methyltransf_23<br>Methyltransf_11<br>Methyltransf_12<br>TIGR03534 | Methyltransferase domain<br>Methyltransferase domain<br>Methyltransferase domain<br>Methyltransferase domain<br>RF_mod_PrmC: protein-(glutamine-N5) methyltransferase, release factor-specific | 1.50E-15<br>4.30E-11<br>5.40E-10<br>2.80E-08<br>9.80E-07 |
| WP_129244286.1 | 92595 | 91950 | - | 214 | LinL | LinL | LinL | 3.70E-72 |
| WP_161989239.1 | 93617 | 92594 | - | 340 | LinE<br>LinH<br>PF12697<br>TIGR02427<br>PF00561 | LinE<br>LinH<br>Abhydrolase_6<br>TIGR02427<br>Abhydrolase_1 | LinE Homolog<br>LinH (Fusion of LinE/LinG)<br>Alpha/beta hydrolase family<br>protocat_pcaD: 3-oxoadipate enol-lactonase<br>alpha/beta hydrolase fold | 1.80E-121<br>3.00E-118<br>1.00E-13<br>1.50E-08<br>2.00E-06 |
| WP_161989240.1 | 94477 | 93613 | - | 287 | LinH<br>LinG | LinH<br>LinG | LinH (Fusion of LinE/LinG)<br>LinG Homolog | 2.00E-125<br>2.00E-115 |
| WP_129244290.1 | 94771 | 94573 | - | 65 | LinA | LinA | LinA (linaridin precursor) | 2.60E-15 |
| WP_129244292.1 | 97908 | 95061 | - | 948 | PF13191<br>PF00196<br>TIGR03903<br>PF13401<br>TIGR03541 | AAA_16<br>GerE<br>TIGR03903<br>AAA_22<br>TIGR03541 | AAA ATPase domain<br>Bacterial regulatory proteins, luxR family<br>TOMM_kin_cyc: TOMM system kinase/cyclase fusion protein<br>AAA domain<br>reg_near_HchA: LuxR family transcriptional regulatory, chaperone HchA-associated | 2.10E-18<br>1.00E-11<br>4.30E-08<br>1.70E-06<br>5.50E-06 |
| WP_129244294.1 | 98536 | 99181 | + | 214 | PF00857<br>TIGR03614 | Isochorismatase<br>TIGR03614 | Isochorismatase family<br>RutB: pyrimidine utilization protein B | 8.20E-19<br>2.00E-05 |
| WP_129244377.1 | 100438 | 99334 | - | 367 | TIGR01392<br>PF00561 | TIGR01392<br>Abhydrolase_1 | homoserO_Ac_trn: homoserine O-acetyltransferase<br>alpha/beta hydrolase fold | 5.40E-115<br>9.60E-26 |

| | | | | | | | | |
|---|---|---|---|---|---|---|---|---|
| WP_129244296.1 | 101221 | 100486 | - | 244 | PF07722<br>PF00117<br>TIGR00888<br>TIGR01815<br>TIGR00566 | Peptidase_C26<br>GATase<br>TIGR00888<br>TIGR01815<br>TIGR00566 | Peptidase C26<br>Glutamine amidotransferase class-I<br>guaA_Nterm: GMP synthase (glutamine-hydrolyzing), N-terminal domain<br>TrpE-clade3: anthranilate synthase<br>trpG_papA: glutamine amidotransferase of anthranilate synthase or aminodeoxychorismate synthase | 6.90E-55<br>5.40E-12<br>1.70E-07<br>1.80E-07<br>5.20E-06 |
| WP_129244298.1 | 102549 | 101217 | - | 443 | TIGR01326<br>PF01053<br>TIGR01325<br>TIGR01328<br>TIGR02080 | TIGR01326<br>Cys_Met_Meta_PP<br>TIGR01325<br>TIGR01328<br>TIGR02080 | OAH_OAS_sulfhy: O-acetylhomoserine aminocarboxypropyltransferase/cysteine synthase<br>Cys/Met metabolism PLP-dependent enzyme<br>O_suc_HS_sulf: O-succinylhomoserine sulfhydrylase<br>met_gam_lyase: methionine gamma-lyase<br>O_succ_thio_ly: O-succinylhomoserine (thiol)-lyase | 2.70E-156<br>3.30E-119<br>4.90E-103<br>2.00E-99<br>2.50E-65 |

| index | leader | core | start | end | dir | score |
|---|---|---|---|---|---|---|
| 1 | VGSRSGTGTLDGRSTSLRGGGRRAAA | TGTGHAPRSAPAGSELGASSAKARVP | 87447 | 87288 | - | 6 |
| 2 | MNTMLHKVNTSEELRRLPIRSDETTGRDVSAPVL | ATPAIAATVGAVSAVADVVGATYAFAKAVGG | 94771 | 94573 | - | 29 |

### Results for WP_165274063.1 [Streptomyces sp. 196(2019)] back to top

previous - next

#### Architecture

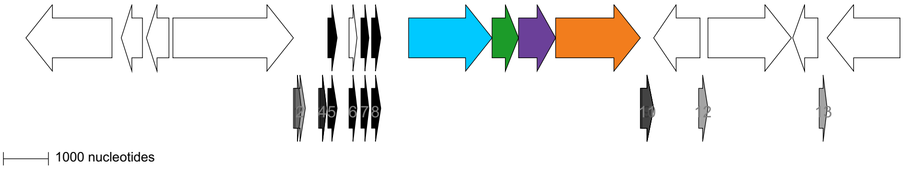

Link to nucleotide sequence

| Accession | start | end | direction | length (aa) | Pfam/HMM | name | description | E-value |
|---|---|---|---|---|---|---|---|---|
| WP_165274046.1 | 12602 | 10667 | - | 644 | PF10022 | DUF2264 | Uncharacterized protein conserved in bacteria (DUF2264) | 9.70E-97 |
| WP_165274048.1 | 13282 | 12799 | - | 160 | NO PFAM MATCH | - | - | - |
| WP_084990728.1 | 13846 | 13366 | - | 159 | PF03745 | DUF309 | Domain of unknown function (DUF309) | 4.90E-19 |
| WP_165274050.1 | 13956 | 16611 | + | 884 | TIGR03903<br>PF13191<br>PF13424<br>PF13401<br>PF07719 | TIGR03903<br>AAA_16<br>TPR_12<br>AAA_22<br>TPR_2 | TOMM_kin_cyc: TOMM system kinase/cyclase fusion protein<br>AAA ATPase domain<br>Tetratricopeptide repeat<br>AAA domain<br>Tetratricopeptide repeat | 2.00E-20<br>7.10E-19<br>2.40E-05<br>1.40E-04<br>2.80E-04 |
| WP_084990724.1 | 17374 | 17578 | + | 67 | LinA | LinA | LinA (linaridin precursor) | 3.10E-11 |
| WP_165274052.1 | 17866 | 18040 | + | 57 | NO PFAM MATCH | - | - | - |
| WP_084990722.1 | 18113 | 18305 | + | 63 | LinA | LinA | LinA (linaridin precursor) | 6.90E-09 |
| WP_084990720.1 | 18362 | 18563 | + | 66 | LinA | LinA | LinA (linaridin precursor) | 2.40E-12 |
| WP_165274063.1 | 19178 | 21023 | + | 614 | LinH<br>LinG<br>LinE<br>LinL<br>PF12697 | LinH<br>LinG<br>LinE<br>LinL<br>Abhydrolase_6 | LinH (Fusion of LinE/LinG)<br>LinG Homolog<br>LinE Homolog<br>LinL<br>Alpha/beta hydrolase family | 1.20E-251<br>2.10E-112<br>2.40E-95<br>3.90E-45<br>7.70E-14 |
| WP_084990715.1 | 21019 | 21628 | + | 202 | LinL | LinL | LinL | 7.10E-66 |
| WP_032756681.1 | 21624 | 22437 | + | 270 | PF13649<br>PF08241<br>PF13847<br>PF13489<br>PF08242 | Methyltransf_25<br>Methyltransf_11<br>Methyltransf_31<br>Methyltransf_23<br>Methyltransf_12 | Methyltransferase domain<br>Methyltransferase domain<br>Methyltransferase domain<br>Methyltransferase domain<br>Methyltransferase domain | 2.30E-16<br>2.80E-12<br>2.60E-07<br>8.20E-07<br>8.70E-07 |
| WP_165274054.1 | 22433 | 24335 | + | 633 | TIGR02204<br>TIGR02203<br>TIGR00958<br>TIGR03796<br>TIGR03797 | TIGR02204<br>TIGR02203<br>TIGR00958<br>TIGR03796<br>TIGR03797 | MsbA_rel: ABC transporter, permease/ATP-binding protein<br>MsbA_lipidA: lipid A export permease/ATP-binding protein MsbA<br>3a01208: antigen peptide transporter 2<br>NHLM_micro_ABC1: NHLM bacteriocin system ABC transporter, peptidase/ATP-binding protein<br>NHLM_micro_ABC2: NHLM bacteriocin system ABC transporter, ATP-binding protein | 8.40E-99<br>6.60E-92<br>1.40E-83<br>6.50E-82<br>2.40E-77 |
| WP_165274056.1 | 25647 | 24633 | - | 337 | NO PFAM MATCH | - | - | - |
| WP_084990709.1 | 25836 | 27690 | + | 617 | NO PFAM MATCH | - | - | - |
| WP_136016412.1 | 28252 | 27715 | - | 178 | PF12840<br>PF01022<br>PF13518 | HTH_20<br>HTH_5<br>HTH_28 | Helix-turn-helix domain<br>Bacterial regulatory protein, arsR family<br>Helix-turn-helix domain | 3.50E-11<br>1.30E-06<br>9.60E-05 |
| WP_165274058.1 | 30075 | 28482 | - | 530 | PF00501<br>TIGR03205<br>TIGR03098<br>TIGR02262<br>TIGR01923 | AMP-binding<br>TIGR03205<br>TIGR03098<br>TIGR02262<br>TIGR01923 | AMP-binding enzyme<br>pimA: dicarboxylate--CoA ligase PimA<br>ligase_PEP_1: acyl-CoA ligase (AMP-forming), exosortase A-associated<br>benz_CoA_lig: benzoate-CoA ligase family<br>menE: O-succinylbenzoate-CoA ligase | 3.80E-114<br>5.80E-79<br>1.60E-75<br>7.70E-75<br>8.00E-72 |

| index | leader | core | start | end | dir | score |
|---|---|---|---|---|---|---|
| 1 | MFDLRGRPDHRGNRSGTWRSLVAHLTGG | QGVAGSNPVVPTGDSRRSGPVSEQSETGP | 16615 | 16789 | + | 6 |
| 2 | VAQLGSALDWGSRGRRFKSCRPDWRQSQIRAGFG | AIRDRPLIIFGDQVGTGRDRLGPAGPGVPSRTPR | 16665 | 16872 | + | 6 |

| | | | | | | |
|---|---|---|---|---|---|---|
| 3 | VPEPAGCCCGVRILMPAELSSLSVGLSASC | GAHHRTSVHAGAGSRVRSRAPGRAPVTAHP | 17182 | 17365 | + | 8 |
| 4 | MPAELSSLSVGLSASCGAHHRTS | VHAGAGSRVRSRAPGRAPVTAHP | 17224 | 17365 | + | 8 |
| 5 | MPEFRQPGWTRGLAPLDEGAGGQVFGGASPVA | ATPAVVATAGAVVVAFAAGVAARHLANGGNVELPM | 17374 | 17578 | + | 31 |
| 6 | MSTASIPAVLSGTGALTESDRGRGPLD | TVPVTSVTVTMTACVGISVCLTGSTIELPQ | 17866 | 18040 | + | 10 |
| 7 | MNTSSTPAVLTATGALTESDPGATLPAIAPVA | ATPVVVVAALGAGFVAGYAAGRAAGNVELPM | 18113 | 18305 | + | 30 |
| 8 | MNMSPTPAALRGAGGLSESDPGRAHSTLVPVA | ATPGVLAGVAVGVALVNAFAAGYNHCGGNVELPM | 18362 | 18563 | + | 31 |
| 9 | VSTVDTTAVGSGPDKPAAA | VTPARAHHGRATGPGPAPYQRFTAR...PSFARAGCATRCPCSARSGAGTPAA | 24331 | 24613 | + | 6 |
| 10 | VDTTAVGSGPDKPAAA | VTPARAHHGRATGPGPAPYQRFTAR...PSFARAGCATRCPCSARSGAGTPAA | 24340 | 24613 | + | 6 |
| 11 | VGSGPDKPAAA | VTPARAHHGRATGPGPAPYQRFTAR...PSFARAGCATRCPCSARSGAGTPAA | 24355 | 24613 | + | 6 |
| 12 | VATSCRSPWPFPRTRPAGQRITSLPGEAGGEVS | AKSQASLAPGKFRMRDHLKERAPIAMTGGRAPK | 25622 | 25823 | + | 6 |
| 13 | VVSADGPSSAATADAWAERGRKHRRS | YRSGPVLTVVQVRCGPDGRPGPVRPG | 28308 | 28467 | + | 6 |

### Results for SOD81768.1 [Streptomyces sp. 1222.2] back to top

previous - next

#### Architecture

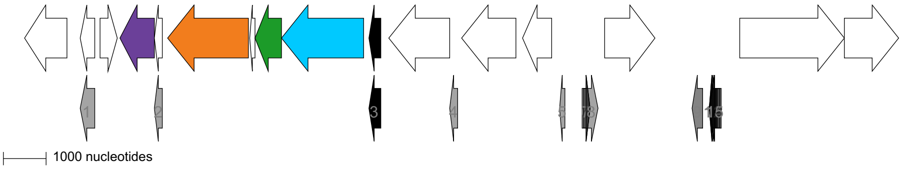

Link to nucleotide sequence

| Accession | start | end | direction | length (aa) | Pfam/HMM | name | description | E-value |
|---|---|---|---|---|---|---|---|---|
| SOD81744.1 | 8588065 | 8587072 | - | 330 | TIGR02363 | TIGR02363 | dhaK1: dihydroxyacetone kinase, DhaK subunit | 7.40E-148 |
| | | | | | PF02733 | Dak1 | Dak1 domain | 1.20E-125 |
| | | | | | TIGR02361 | TIGR02361 | dak_ATP: dihydroxyacetone kinase | 5.60E-93 |
| | | | | | TIGR02362 | TIGR02362 | dhaK1b: probable dihydroxyacetone kinase DhaK1b subunit | 1.50E-73 |
| SOD81747.1 | 8588720 | 8588384 | - | 111 | TIGR02228 | TIGR02228 | sigpep_I_arch: signal peptidase I | 5.40E-04 |
| | | | | | PF00717 | Peptidase_S24 | Peptidase S24-like | 9.40E-04 |
| SOD81750.1 | 8588855 | 8589251 | + | 131 | PF00583 | Acetyltransf_1 | Acetyltransferase (GNAT) family | 8.30E-11 |
| | | | | | PF13673 | Acetyltransf_10 | Acetyltransferase (GNAT) domain | 1.20E-05 |
| | | | | | TIGR01575 | TIGR01575 | rimI: ribosomal-protein-alanine acetyltransferase | 5.30E-05 |
| | | | | | PF13508 | Acetyltransf_7 | Acetyltransferase (GNAT) domain | 3.00E-04 |
| | | | | | PF14542 | Acetyltransf_CG | GCN5-related N-acetyl-transferase | 5.20E-04 |
| SOD81753.1 | 8590119 | 8589312 | - | 268 | PF13649 | Methyltransf_25 | Methyltransferase domain | 1.20E-13 |
| | | | | | PF08241 | Methyltransf_11 | Methyltransferase domain | 9.70E-11 |
| | | | | | PF13489 | Methyltransf_23 | Methyltransferase domain | 2.80E-06 |
| | | | | | TIGR01983 | TIGR01983 | UbiG: 3-demethylubiquinone-9 3-O-methyltransferase | 3.10E-05 |
| | | | | | PF08242 | Methyltransf_12 | Methyltransferase domain | 4.20E-05 |
| SOD81757.1 | 8590307 | 8590115 | - | 63 | NO PFAM MATCH | - | - | - |
| SOD81759.1 | 8592315 | 8590422 | - | 630 | TIGR02204 | TIGR02204 | MsbA_rel: ABC transporter, permease/ATP-binding protein | 5.60E-106 |
| | | | | | TIGR02203 | TIGR02203 | MsbA_lipidA: lipid A export permease/ATP-binding protein MsbA | 1.30E-101 |
| | | | | | TIGR00958 | TIGR00958 | 3a01208: antigen peptide transporter 2 | 3.20E-91 |
| | | | | | TIGR03796 | TIGR03796 | NHLM_micro_ABC1: NHLM bacteriocin system ABC transporter, peptidase/ATP-binding protein | 9.00E-85 |
| | | | | | TIGR01846 | TIGR01846 | type_I_sec_HlyB: type I secretion system ATPase | 2.20E-79 |
| SOD81762.1 | 8592480 | 8592354 | - | 41 | NO PFAM MATCH | - | - | - |
| SOD81765.1 | 8593104 | 8592483 | - | 206 | LinL | LinL | LinL | 4.80E-69 |
| SOD81768.1 | 8595020 | 8593100 | - | 639 | LinH | LinH | LinH (Fusion of LinE/LinG) | 2.10E-248 |
| | | | | | LinG | LinG | LinG Homolog | 8.40E-164 |
| | | | | | LinE | LinE | LinE Homolog | 9.00E-92 |
| | | | | | PF12697 | Abhydrolase_6 | Alpha/beta hydrolase family | 1.30E-16 |
| | | | | | PF12146 | Hydrolase_4 | Serine aminopeptidase, S33 | 1.50E-09 |
| SOD81772.1 | 8595415 | 8595136 | - | 92 | LinA | LinA | LinA (linaridin precursor) | 3.90E-08 |
| SOD81776.1 | 8597044 | 8595613 | - | 476 | TIGR02734 | TIGR02734 | crtI_fam: phytoene desaturase | 1.00E-12 |
| | | | | | PF13450 | NAD_binding_8 | NAD(P)-binding Rossmann-like domain | 2.50E-11 |
| | | | | | TIGR03467 | TIGR03467 | HpnE: squalene-associated FAD-dependent desaturase | 1.70E-08 |
| | | | | | TIGR00292 | TIGR00292 | TIGR00292: thiazole biosynthesis enzyme | 6.30E-08 |
| | | | | | PF01266 | DAO | FAD dependent oxidoreductase | 6.80E-08 |
| SOD81779.1 | 8598575 | 8597309 | - | 421 | PF07730 | HisKA_3 | Histidine kinase | 8.70E-14 |
| SOD81782.1 | 8599430 | 8598746 | - | 227 | PF00072 | Response_reg | Response regulator receiver domain | 1.40E-24 |
| | | | | | PF00196 | GerE | Bacterial regulatory proteins, luxR family | 2.50E-18 |
| | | | | | TIGR02875 | TIGR02875 | spore_0_A: sporulation transcription factor Spo0A | 2.30E-15 |
| | | | | | TIGR03020 | TIGR03020 | EpsA: transcriptional regulator EpsA | 1.60E-13 |
| | | | | | TIGR03541 | TIGR03541 | reg_near_HchA: LuxR family transcriptional regulatory, chaperone HchA-associated | 1.20E-08 |
| SOD81785.1 | 8600656 | 8601844 | + | 395 | PF00480 | ROK | ROK family | 2.70E-53 |
| | | | | | TIGR00744 | TIGR00744 | ROK_glcA_fam: ROK family protein (putative glucokinase) | 4.00E-40 |
| | | | | | PF12802 | MarR_2 | MarR family | 1.30E-07 |
| | | | | | PF13412 | HTH_24 | Winged helix-turn-helix DNA-binding | 4.70E-05 |
| | | | | | PF01978 | TrmB | Sugar-specific transcriptional regulator TrmB | 1.10E-04 |
| SOD81788.1 | 8603841 | 8606265 | + | 807 | PF14498 | Glyco_hyd_65N_2 | Glycosyl hydrolase family 65, N-terminal domain | 5.10E-36 |

| | | | | | | | | |
|---|---|---|---|---|---|---|---|---|
| SOD81791.1 | 8606268 | 8607537 | + | 422 | PF13472<br>PF00657 | Lipase_GDSL_2<br>Lipase_GDSL | GDSL-like Lipase/Acylhydrolase family<br>GDSL-like Lipase/Acylhydrolase | 1.60E-24<br>6.10E-08 |

| index | leader | core | start | end | dir | score |
|---|---|---|---|---|---|---|
| 1 | MVAERVVAGATVTFRPSGSSMVPLI...ISNNRGHVNGWTGHGRVFGICVAVD | GTARPRTAGKTRPGG | 8588720 | 8588384 | - | 6 |
| 2 | MNDFLLIPSVVALGIVGFLIATRAVSTPAVI | GVALVVLWGACSQARSTIHARHTPSKKRRQHP | 8590307 | 8590115 | - | 6 |
| 3 | MSSIENALNSVEIPVEGVVYVAARPTL | GTPRIARVGNRLGGLAQAAEGVGAI...ANNLAAEANAQNAAALAAVGGAPAA | 8595415 | 8595136 | - | 28 |
| 4 | MSESQRRRDGLISSRRVVNRPGPGPFEGPFR | LLSPAISRVGGPVSTVGKSQEKRNAEDAGDI | 8597229 | 8597040 | - | 6 |
| 5 | MGRQKEDLQFPGAEPAVAVHCHG | VPALVRRGSGTRGSSAWCAVTLS | 8599748 | 8599607 | - | 6 |
| 6 | VRVSRRRVATGDGFPAV | RTPSGRRSDAVRGCVSDPRPTSPADPGSTGTAVVPARFRSAATG | 8600145 | 8600331 | + | 6 |
| 7 | VATGDGFPAV | RTPSGRRSDAVRGCVSDPRPTSPADPGSTGTAVVPARFRSAATG | 8600166 | 8600331 | + | 7 |
| 8 | VCPTRGPPPRRTRAVRAPPSSRRGSARPPPDDLRGVGARGPRGRM | PRAARSVPGVRRPSSARCAVEPRQVVIGAICWTYGGMPVPAPRAAP | 8600236 | 8600512 | + | 6 |
| 9 | VAGPGGGVVPAVLDGVLPGDQGVAVALDVGV | DAPAVEYRFQSATWPRSRDQDSSRWIGRPAAAMAFCQAATSATLS | 8602945 | 8602714 | - | 7 |
| 10 | VKPPSGFFAAQTCWSSEGVTGLAGGVAPW | LTAMTSLLVSRERP | 8603232 | 8603100 | - | 6 |
| 11 | VTDAGASPRLNRATVKPPSGFFAAQTCWSSEGVTGLAGGVAPW | LTAMTSLLVSRERP | 8603274 | 8603100 | - | 7 |
| 12 | LPEKPRADQSAVTDAGASPRLNRAT...SGFFAAQTCWSSEGVTGLAGGVAPW | LTAMTSLLVSRERP | 8603307 | 8603100 | - | 7 |
| 13 | VPRFSRFHRSRTFALPEKPRADQSA...SGFFAAQTCWSSEGVTGLAGGVAPW | LTAMTSLLVSRERP | 8603349 | 8603100 | - | 6 |
| 14 | VILSRWVPRFSRFHRSRTFALPEKP...SGFFAAQTCWSSEGVTGLAGGVAPW | LTAMTSLLVSRERP | 8603367 | 8603100 | - | 6 |
| 15 | VNRREPCAGSACVILSRWVPRFSRF...SGFFAAQTCWSSEGVTGLAGGVAPW | LTAMTSLLVSRERP | 8603403 | 8603100 | - | 6 |

### Results for PIG13858.1 [Streptomyces sp. 1121.2] back to top

previous - next

#### Architecture

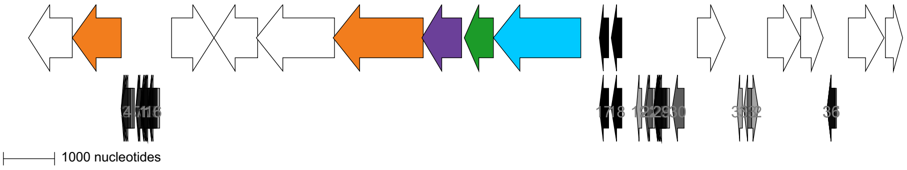

Link to nucleotide sequence

| Accession | start | end | direction | length (aa) | Pfam/HMM | name | description | E-value |
|---|---|---|---|---|---|---|---|---|
| PIG13850.1 | 4451144 | 4450292 | - | 283 | PF12698<br>PF01061<br>TIGR01247<br>TIGR00025<br>PF12679 | ABC2_membrane_3<br>ABC2_membrane<br>TIGR01247<br>TIGR00025<br>ABC2_membrane_2 | ABC-2 family transporter protein<br>ABC-2 type transporter<br>drrB: daunorubicin resistance ABC transporter membrane protein<br>Mtu_efflux: ABC transporter efflux protein, DrrB family<br>ABC-2 family transporter protein | 2.90E-18<br>7.40E-14<br>2.20E-10<br>1.10E-08<br>4.50E-05 |
| PIG13851.1 | 4452112 | 4451140 | - | 323 | TIGR01188<br>TIGR03864<br>TIGR03522<br>TIGR03873<br>TIGR01288 | TIGR01188<br>TIGR03864<br>TIGR03522<br>TIGR03873<br>TIGR01288 | drrA: daunorubicin resistance ABC transporter, ATP-binding protein<br>PQQ_ABC_ATP: ABC transporter, ATP-binding subunit, PQQ-dependent alcohol dehydrogenase system<br>GldA_ABC_ATP: gliding motility-associated ABC transporter ATP-binding subunit GldA<br>F420-0_ABC_ATP: proposed F420-0 ABC transporter, ATP-binding protein<br>nodI: nodulation ABC transporter NodI | 6.00E-58<br>7.00E-46<br>1.00E-40<br>3.40E-39<br>1.20E-38 |
| PIG13852.1 | 4453080 | 4453923 | + | 280 | PF01551 | Peptidase_M23 | Peptidase family M23 | 7.60E-16 |
| PIG13853.1 | 4454759 | 4453907 | - | 283 | PF12680<br>PF14534 | SnoaL_2<br>DUF4440 | SnoaL-like domain<br>Domain of unknown function (DUF4440) | 6.30E-05<br>2.50E-04 |
| PIG13854.1 | 4456261 | 4454755 | - | 501 | TIGR02734<br>TIGR00292<br>TIGR03467<br>TIGR01317<br>TIGR02733 | TIGR02734<br>TIGR00292<br>TIGR03467<br>TIGR01317<br>TIGR02733 | crtI_fam: phytoene desaturase<br>TIGR00292: thiazole biosynthesis enzyme<br>HpnE: squalene-associated FAD-dependent desaturase<br>GOGAT_sm_gam: glutamate synthase, NADH/NADPH, small subunit<br>desat_CrtD: C-3',4' desaturase CrtD | 6.30E-10<br>5.40E-08<br>7.60E-08<br>1.70E-07<br>1.90E-07 |
| PIG13855.1 | 4457991 | 4456257 | - | 577 | TIGR02204<br>TIGR02203<br>TIGR00958<br>TIGR03796<br>TIGR01846 | TIGR02204<br>TIGR02203<br>TIGR00958<br>TIGR03796<br>TIGR01846 | MsbA_rel: ABC transporter, permease/ATP-binding protein<br>MsbA_lipidA: lipid A export permease/ATP-binding protein MsbA<br>3a01208: antigen peptide transporter 2<br>NHLM_micro_ABC1: NHLM bacteriocin system ABC transporter, peptidase/ATP-binding protein<br>type_I_sec_HlyB: type I secretion system ATPase | 3.50E-102<br>6.10E-99<br>4.00E-86<br>5.20E-86<br>4.30E-83 |
| PIG13856.1 | 4458743 | 4457987 | - | 251 | PF13649<br>PF08241<br>PF13489<br>PF08242<br>TIGR01983 | Methyltransf_25<br>Methyltransf_11<br>Methyltransf_23<br>Methyltransf_12<br>TIGR01983 | Methyltransferase domain<br>Methyltransferase domain<br>Methyltransferase domain<br>Methyltransferase domain<br>UbiG: 3-demethylubiquinone-9 3-O-methyltransferase | 6.70E-13<br>1.80E-08<br>4.20E-08<br>2.60E-06<br>4.60E-05 |
| PIG13857.1 | 4459376 | 4458797 | - | 192 | LinL | LinL | LinL | 5.00E-66 |

| | | | | | | | | |
|---|---|---|---|---|---|---|---|---|
| PIG13858.1 | 4461094 | 4459372 | - | 573 | LinH<br>LinE<br>LinG<br>PF12697<br>PF00561 | LinH<br>LinE<br>LinG<br>Abhydrolase_6<br>Abhydrolase_1 | LinH (Fusion of LinE/LinG)<br>LinE Homolog<br>LinG Homolog<br>Alpha/beta hydrolase family<br>alpha/beta hydrolase fold | 7.00E-221<br>4.00E-99<br>4.00E-94<br>5.40E-15<br>4.00E-11 |
| PIG13859.1 | 4461633 | 4461441 | - | 63 | LinA | LinA | LinA (linaridin precursor) | 1.40E-12 |
| PIG13860.1 | 4461874 | 4461679 | - | 64 | LinA | LinA | LinA (linaridin precursor) | 1.10E-17 |
| BX282_4112 | 4463346 | 4463902 | + | 185 | INFERRED GENE | - | - | - |
| PIG13861.1 | 4464734 | 4465376 | + | 213 | PF04255<br>PF13411 | DUF433<br>MerR_1 | Protein of unknown function (DUF433)<br>MerR HTH family regulatory protein | 3.20E-16<br>8.80E-04 |
| PIG13862.1 | 4465375 | 4465816 | + | 146 | PF18478<br>PF18480 | PIN_10<br>DUF5615 | PIN like domain<br>Domain of unknown function (DUF5615) | 4.20E-16<br>1.00E-07 |
| PIG13863.1 | 4466317 | 4467013 | + | 231 | NO PFAM MATCH | - | - | - |
| PIG13864.1 | 4467032 | 4467362 | + | 109 | NO PFAM MATCH | - | - | - |

| index | leader | core | start | end | dir | score |
|---|---|---|---|---|---|---|
| 1 | LQREGGGADGGQTRSRDHEPRP | TGPLTRHTDTHRPRSMDRDGEHL | 4452246 | 4452108 | - | 6 |
| 2 | LCPVSALSTVGESRACLQREGGGADGGQTR | SRDHEPRPTGPLTRHTDTHRPRSMDRDGEHL | 4452294 | 4452108 | - | 8 |
| 3 | MCDCSPCGDCWKLSRDRLCPVSALSTVGESRACLQREGG | GADGGQTRSRDHEPRPTGPLTRHTDTHRPRSMDRDGEHL | 4452345 | 4452108 | - | 6 |
| 4 | MGGLMCDCSPCGDCWKLSRDRLCPVSALSTVGESRACLQRE | GGGADGGQTRSRDHEPRPTGPLTRHTDTHRPRSMDRDGEHL | 4452357 | 4452108 | - | 6 |
| 5 | LKDYALDGSLLMFRVSST | NPTLRSAPASALAGRSIL | 4452498 | 4452387 | - | 6 |
| 6 | LLSWCSSGITWVSLKDYALDGSLL | MFRVSSTNPTLRSAPASALAGRSIL | 4452537 | 4452387 | - | 6 |
| 7 | VFVLLSWCSSGITWVSLKDYALDGSL | LMFRVSSTNPTLRSAPASALAGRSIL | 4452546 | 4452387 | - | 8 |
| 8 | VAAEFAGLAAAEFAEEDKAAM | TTAMVESVRFAVLVFIWNHLGFLEGLCTGWFLADV | 4452628 | 4452457 | - | 7 |
| 9 | VPDALDEPWVADHSTGVAAEFAGLAAAEFAEEDKAAM | TTAMVESVRFAVLVFIWNHLGFLEGLCTGWFLADV | 4452676 | 4452457 | - | 6 |
| 10 | LTARTVLLPFCSVHRVPDALDEPWV...HSTGVAAEFAGLAAAEFAEEDKAAM | TTAMVESVRFAVLVFIWNHLGFLEGLCTGWFLADV | 4452721 | 4452457 | - | 7 |
| 11 | VFWISPLTARTVLLPFCSVHRVPDA...HSTGVAAEFAGLAAAEFAEEDKAAM | TTAMVESVRFAVLVFIWNHLGFLEGLCTGWFLADV | 4452739 | 4452457 | - | 7 |
| 12 | VPVFWISPLTARTVLLPFCSVHRVP...HSTGVAAEFAGLAAAEFAEEDKAAM | TTAMVESVRFAVLVFIWNHLGFLEGLCTGWFLADV | 4452745 | 4452457 | - | 7 |
| 13 | MKSRVHWTQTEKHSVPVFWISPLTA...HSTGVAAEFAGLAAAEFAEEDKAAM | TTAMVESVRFAVLVFIWNHLGFLEGLCTGWFLADV | 4452787 | 4452457 | - | 6 |
| 14 | MFGIGGRDEVTGPLDPDRKAFGACVLDFSVDRENCPAAVLF | GPPGAGRTRRTVGCRPLHRSGR | 4452809 | 4452617 | - | 7 |
| 15 | LSGMFGIGGRDEVTGPLDPDRKAFGACVLDFSVDRENCPAAVLF | GPPGAGRTRRTVGCRPLHRSGR | 4452818 | 4452617 | - | 7 |
| 16 | LIWAQPKVDGRDSGAALSGMFGIGG...DRKAFGACVLDFSVDRENCPAAVLF | GPPGAGRTRRTVGCRPLHRSGR | 4452866 | 4452617 | - | 6 |
| 17 | MSRIVDSVVAVTSADLTDDSQGEFGAMAPVQ | ATPTLVAFGVGFAGGAGAAWVTVQAYEAGAND | 4461633 | 4461441 | - | 31 |
| 18 | MEQKIAAFMNEQDLGSLEQGYRDDAPMPVL | ATPAAAVATCWYVGGAAAGAAAVVGAYVTGRVVG | 4461874 | 4461679 | - | 30 |
| 19 | MRLNLDTGIGAGGAAT | TFFMQGAIADLFSPAA | 4462274 | 4462175 | - | 6 |
| 20 | VALPSGSAATFLQGIAGRPLGRCIR | GVRAVVLLTSRQGLVTDHYRPIGCE | 4462479 | 4462326 | - | 7 |
| 21 | MVRIRPGRLHGLFSLISSRYCGVAFGKCSN | VLAGNCGSTARSLHPRRAGRRPAHQPTGASD | 4462540 | 4462354 | - | 6 |
| 22 | MSLSWEGTGHSITWTLLVGA | WVPFVKSQVDPRVEVRGGVRSAWFGSGQDGFMACSL | 4462665 | 4462494 | - | 6 |
| 23 | MSGIYDSMSLSWEGTGHSITWTLLVGA | WVPFVKSQVDPRVEVRGGVRSAWFGSGQDGFMACSL | 4462686 | 4462494 | - | 6 |
| 24 | MANAEIDQRACGRCGSMSGIYDSMSLSWEGTGHSITWTLLVGA | WVPFVKSQVDPRVEVRGGVRSAWFGSGQDGFMACSL | 4462734 | 4462494 | - | 6 |
| 25 | LMANAEIDQRACGRCGSMSGIYDSMSLSWEGTGHSITWTLLVGA | WVPFVKSQVDPRVEVRGGVRSAWFGSGQDGFMACSL | 4462737 | 4462494 | - | 6 |
| 26 | MGGLMANAEIDQRACGRCGSMSGIYDSMSLSWEGTGHSITWTLLVGA | WVPFVKSQVDPRVEVRGGVRSAWFGSGQDGFMACSL | 4462746 | 4462494 | - | 6 |
| 27 | VASPLMGGLMANAEIDQRACGRCGS...GIYDSMSLSWEGTGHSITWTLLVGA | WVPFVKSQVDPRVEVRGGVRSAWFGSGQDGFMACSL | 4462761 | 4462494 | - | 6 |
| 28 | LTPYVASPLMGGLMANAEIDQRACG...GIYDSMSLSWEGTGHSITWTLLVGA | WVPFVKSQVDPRVEVRGGVRSAWFGSGQDGFMACSL | 4462773 | 4462494 | - | 6 |
| 29 | VLELVLWWSTLTPYVASPLMGGLMA...AEIDQRACGRCGSMSGIYDSMSLSW | EGTGHSITWTLLVGAWVPFVKSQVD...RVEVRGGVRSAWFGSGQDGFMACSL | 4462803 | 4462494 | - | 6 |
| 30 | MDATSETTCREALLRQACRRAKVSA...EPGRKSAAGWQIRPGVVVDRTVPAS | VTPRSGRRTDHSVSCVR | 4463092 | 4462861 | - | 8 |
| 31 | MASPLSGHSWGQDGAGR | TVKRHERSTRPGRVHAC | 4464237 | 4464132 | - | 6 |
| 32 | MEGILDVADVLQGRLRAGPLAHFDR...GDNRAGHASRRRAEEGLLPCARAEP | MTVLAGQPRGSTVAPWR | 4464330 | 4464537 | + | 6 |
| 33 | MASPVIASSIKVGEWSRAKTALQDICN | VKDAFHSGAEPLFASPGRCVGRPFAARL | 4464429 | 4464261 | - | 6 |
| 34 | MLCHLHVVPVTVEVLGLLFGAISVFG | WVSCTVTLRRAEAGRTIRIPRSRWMW | 4466054 | 4465895 | - | 8 |
| 35 | VMLCHLHVVP | VTVEVLGLLFGAISVFGWVSCTVTLRRAEAGRTIRIPRSRWMW | 4466057 | 4465895 | - | 8 |
| 36 | LAVMLCHLHVVP | VTVEVLGLLFGAISVFGWVSCTVTLRRAEAGRTIRIPRSRWMW | 4466063 | 4465895 | - | 8 |

### Results for ROQ95786.1 [Streptomyces sp. 2132.2] back to top

previous - next

#### Architecture

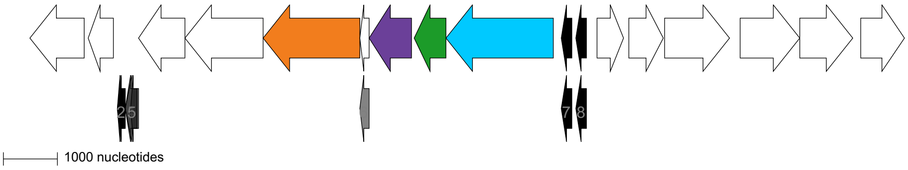

Link to nucleotide sequence

| Accession | start | end | direction | length (aa) | Pfam/HMM | name | description | E-value |
|---|---|---|---|---|---|---|---|---|
| ROQ95778.1 | 2519443 | 2518423 | - | 339 | TIGR00456<br>PF05746<br>PF03485 | TIGR00456<br>DALR_1<br>Arg_tRNA_synt_N | argS: arginine--tRNA ligase<br>DALR anticodon binding domain<br>Arginyl tRNA synthetase N terminal domain | 2.20E-22<br>1.90E-17<br>1.70E-16 |
| ROQ95779.1 | 2519983 | 2519512 | - | 156 | TIGR02154<br>PF00072<br>TIGR02875<br>TIGR01818<br>TIGR02956 | TIGR02154<br>Response_reg<br>TIGR02875<br>TIGR01818<br>TIGR02956 | PhoB: phosphate regulon transcriptional regulatory protein PhoB<br>Response regulator receiver domain<br>spore_0_A: sporulation transcription factor Spo0A<br>ntrC: nitrogen regulation protein NR(I)<br>TMAO_torS: TMAO reductase sytem sensor TorS | 4.00E-25<br>6.30E-24<br>8.80E-19<br>5.60E-17<br>5.30E-16 |
| ROQ95780.1 | 2521299 | 2520447 | - | 283 | PF01551 | Peptidase_M23 | Peptidase family M23 | 6.10E-18 |
| ROQ95781.1 | 2522759 | 2521295 | - | 487 | TIGR02734<br>TIGR03467<br>PF13450<br>TIGR01372<br>PF01593 | TIGR02734<br>TIGR03467<br>NAD_binding_8<br>TIGR01372<br>Amino_oxidase | crtI_fam: phytoene desaturase<br>HpnE: squalene-associated FAD-dependent desaturase<br>NAD(P)-binding Rossmann-like domain<br>soxA: sarcosine oxidase, alpha subunit family<br>Flavin containing amine oxidoreductase | 6.30E-12<br>2.40E-10<br>6.20E-10<br>1.70E-08<br>2.00E-08 |
| ROQ95782.1 | 2524540 | 2522755 | - | 594 | TIGR02204<br>TIGR02203<br>TIGR03796<br>TIGR00958<br>TIGR01846 | TIGR02204<br>TIGR02203<br>TIGR03796<br>TIGR00958<br>TIGR01846 | MsbA_rel: ABC transporter, permease/ATP-binding protein<br>MsbA_lipidA: lipid A export permease/ATP-binding protein MsbA<br>NHLM_micro_ABC1: NHLM bacteriocin system ABC transporter, peptidase/ATP-binding protein<br>3a01208: antigen peptide transporter 2<br>type_I_sec_HlyB: type I secretion system ATPase | 1.70E-100<br>2.80E-98<br>2.50E-85<br>6.30E-82<br>2.50E-80 |
| ROQ95783.1 | 2524713 | 2524536 | - | 58 | NO PFAM MATCH | - | - | - |
| ROQ95784.1 | 2525494 | 2524714 | - | 259 | PF13649<br>PF08241<br>PF08242<br>PF13489<br>PF13847 | Methyltransf_25<br>Methyltransf_11<br>Methyltransf_12<br>Methyltransf_23<br>Methyltransf_31 | Methyltransferase domain<br>Methyltransferase domain<br>Methyltransferase domain<br>Methyltransferase domain<br>Methyltransferase domain | 1.80E-13<br>4.40E-09<br>4.70E-08<br>5.20E-06<br>7.80E-06 |
| ROQ95785.1 | 2526130 | 2525542 | - | 195 | LinL | LinL | LinL | 2.20E-70 |
| ROQ95786.1 | 2528133 | 2526126 | - | 668 | LinH<br>LinG<br>LinE<br>PF12697<br>PF12146 | LinH<br>LinG<br>LinE<br>Abhydrolase_6<br>Hydrolase_4 | LinH (Fusion of LinE/LinG)<br>LinG Homolog<br>LinE Homolog<br>Alpha/beta hydrolase family<br>Serine aminopeptidase, S33 | 9.90E-252<br>1.10E-117<br>5.30E-100<br>2.60E-16<br>6.80E-10 |
| ROQ95787.1 | 2528481 | 2528280 | - | 66 | LinA | LinA | LinA (linaridin precursor) | 6.10E-11 |
| ROQ95788.1 | 2528730 | 2528535 | - | 64 | LinA | LinA | LinA (linaridin precursor) | 5.00E-12 |
| ROQ95789.1 | 2528934 | 2529420 | + | 161 | PF13560 | HTH_31 | Helix-turn-helix domain | 3.40E-04 |
| ROQ95790.1 | 2529522 | 2530170 | + | 215 | PF06114 | Peptidase_M78 | IrrE N-terminal-like domain | 6.60E-05 |
| ROQ95791.1 | 2530182 | 2531388 | + | 401 | NO PFAM MATCH | - | - | - |
| ROQ95792.1 | 2531592 | 2532702 | + | 369 | PF08028<br>TIGR04022<br>PF02770<br>PF00441 | Acyl-CoA_dh_2<br>TIGR04022<br>Acyl-CoA_dh_M<br>Acyl-CoA_dh_1 | Acyl-CoA dehydrogenase, C-terminal domain<br>sulfur_SfnB: sulfur acquisition oxidoreductase, SfnB family<br>Acyl-CoA dehydrogenase, middle domain<br>Acyl-CoA dehydrogenase, C-terminal domain | 4.30E-25<br>2.50E-23<br>1.10E-05<br>6.30E-05 |
| ROQ95793.1 | 2532698 | 2533685 | + | 328 | PF00248<br>TIGR01293 | Aldo_ket_red<br>TIGR01293 | Aldo/keto reductase family<br>Kv_beta: voltage-dependent potassium channel beta subunit | 2.90E-74<br>2.50E-27 |
| ROQ95794.1 | 2533819 | 2534629 | + | 269 | PF03704<br>PF00486 | BTAD<br>Trans_reg_C | Bacterial transcriptional activator domain<br>Transcriptional regulatory protein, C terminal | 5.20E-41<br>9.30E-11 |

| index | leader | core | start | end | dir | score |
|---|---|---|---|---|---|---|
| 1 | LGTSCEVPPAGFEPAH | TAPEAVALSPELRGRVVRVAATGGTLPASAG | 2520179 | 2520035 | - | 8 |
| 2 | MAGSLGTSCEVPPAGFEPAH | TAPEAVALSPELRGRVVRVAATGGTLPASAG | 2520191 | 2520035 | - | 10 |
| 3 | MGPARGAGSAALDSPRIDPEG | TAAAAGAVAALRGLLHRRLTAG | 2520420 | 2520288 | - | 6 |
| 4 | VHGACKGCGKRRARFTENRSGGNCRRSRCGRCSPRS | PTPAADRGLKNRSAEETRRAPLPRPPTHGCTYAKTPSQRTR | 2520424 | 2520190 | - | 7 |
| 5 | VTPVLPGPAVHGACKGCGKRRARFTENRSGGNCRRSRCGRCSPRS | PTPAADRGLKNRSAEETRRAPLPRPPTHGCTYAKTPSQRTR | 2520451 | 2520190 | - | 7 |
| 6 | MSTQRAVWFVTALAVAVPVM | AVMFREDGRFTSQSWTKGLIFGTAVAVVAAIAAGRARQ | 2524713 | 2524536 | - | 7 |
| 7 | MNSITESVRVIAGTTLSEDSGERTTRMMPVA | FTPAVLVTTPYAAGGLAAGAAAVGAYEAGRAAAKG | 2528481 | 2528280 | - | 29 |
| 8 | MAKIAAFMNQQDMGPIAESGLSVGGTPVM | ATPAVVAVPASVKITAGIVAFTIPVGAYVTGRVVN | 2528730 | 2528535 | - | 30 |

### Results for SED34674.1 [Streptomyces sp. 2314.4] back to top

previous - next

#### Architecture

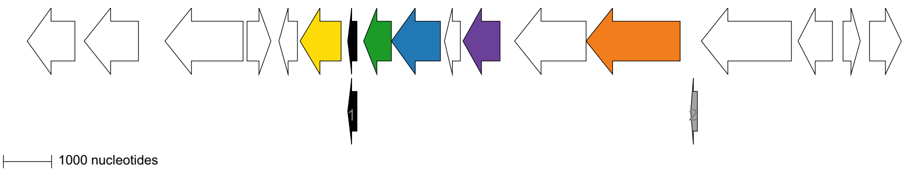

Link to nucleotide sequence

| Accession | start | end | direction | length (aa) | Pfam/HMM | name | description | E-value |
|---|---|---|---|---|---|---|---|---|
| SED34288.1 | 3939188 | 3938180 | - | 335 | TIGR02432<br>PF01171<br>PF09179 | TIGR02432<br>ATP_bind_3<br>TilS | lysidine_TilS_N: tRNA(Ile)-lysidine synthetase<br>PP-loop family<br>TilS substrate binding domain | 3.60E-43<br>6.60E-38<br>5.30E-12 |
| SED34339.1 | 3940499 | 3939380 | - | 372 | TIGR03883<br>TIGR03624<br>PF10103 | TIGR03883<br>TIGR03624<br>Zincin_2 | DUF2342_F420: uncharacterized protein, coenzyme F420 biosynthesis associated<br>TIGR03624: putative hydrolase<br>Zincin-like metallopeptidase | 7.20E-142<br>1.00E-125<br>3.80E-121 |
| SED34386.1 | 3942655 | 3941047 | - | 535 | PF02113<br>TIGR00666 | Peptidase_S13<br>TIGR00666 | D-Ala-D-Ala carboxypeptidase 3 (S13) family<br>PBP4: D-alanyl-D-alanine carboxypeptidase/D-alanyl-D-alanine-endopeptidase | 6.00E-76<br>6.70E-58 |
| SED34433.1 | 3942741 | 3943236 | + | 164 | PF00719 | Pyrophosphatase | Inorganic pyrophosphatase | 2.50E-53 |
| SED34481.1 | 3943777 | 3943417 | - | 119 | NO PFAM MATCH | - | - | - |
| SED34529.1 | 3944697 | 3943842 | - | 284 | LinG<br>LinH<br>TIGR04033 | LinG<br>LinH<br>TIGR04033 | LinG Homolog<br>LinH (Fusion of LinE/LinG)<br>export_SdpB: antimicrobial peptide system protein, SdpB family | 3.00E-111<br>3.70E-79<br>2.10E-04 |
| SED34577.1 | 3945025 | 3944842 | - | 60 | LinA | LinA | LinA (linaridin precursor) | 1.80E-13 |
| SED34630.1 | 3945740 | 3945152 | - | 195 | LinL | LinL | LinL | 2.50E-56 |
| SED34674.1 | 3946756 | 3945736 | - | 339 | LinE<br>LinH<br>PF12697<br>TIGR02427<br>PF00561 | LinE<br>LinH<br>Abhydrolase_6<br>TIGR02427<br>Abhydrolase_1 | LinE Homolog<br>LinH (Fusion of LinE/LinG)<br>Alpha/beta hydrolase family<br>protocat_pcaD: 3-oxoadipate enol-lactonase<br>alpha/beta hydrolase fold | 4.70E-104<br>4.10E-53<br>3.10E-15<br>8.80E-12<br>7.20E-10 |
| SED34720.1 | 3947176 | 3946828 | - | 115 | PF13518<br>PF01527<br>PF13551<br>PF13011<br>PF13384 | HTH_28<br>HTH_Tnp_1<br>HTH_29<br>LZ_Tnp_IS481<br>HTH_23 | Helix-turn-helix domain<br>Transposase<br>Winged helix-turn helix<br>leucine-zipper of insertion element IS481<br>Homeodomain-like domain | 4.20E-09<br>2.50E-06<br>1.30E-05<br>1.00E-04<br>4.20E-04 |
| SED34774.1 | 3947977 | 3947218 | - | 252 | PF13649<br>PF08241<br>PF13489<br>PF08242<br>PF13847 | Methyltransf_25<br>Methyltransf_11<br>Methyltransf_23<br>Methyltransf_12<br>Methyltransf_31 | Methyltransferase domain<br>Methyltransferase domain<br>Methyltransferase domain<br>Methyltransferase domain<br>Methyltransferase domain | 5.70E-17<br>1.90E-12<br>1.30E-10<br>5.60E-10<br>5.60E-09 |
| SED34822.1 | 3949779 | 3948294 | - | 494 | TIGR02734<br>TIGR00292<br>PF13450<br>TIGR03467<br>PF01593 | TIGR02734<br>TIGR00292<br>NAD_binding_8<br>TIGR03467<br>Amino_oxidase | crtI_fam: phytoene desaturase<br>TIGR00292: thiazole biosynthesis enzyme<br>NAD(P)-binding Rossmann-like domain<br>HpnE: squalene-associated FAD-dependent desaturase<br>Flavin containing amine oxidoreductase | 8.80E-09<br>4.70E-08<br>5.30E-08<br>7.40E-08<br>3.40E-07 |
| SED34867.1 | 3951710 | 3949772 | - | 645 | TIGR02204<br>TIGR02203<br>TIGR00958<br>TIGR03796<br>TIGR03375 | TIGR02204<br>TIGR02203<br>TIGR00958<br>TIGR03796<br>TIGR03375 | MsbA_rel: ABC transporter, permease/ATP-binding protein<br>MsbA_lipidA: lipid A export permease/ATP-binding protein MsbA<br>3a01208: antigen peptide transporter 2<br>NHLM_micro_ABC1: NHLM bacteriocin system ABC transporter, peptidase/ATP-binding protein<br>type_I_sec_LssB: type I secretion system ATPase | 5.70E-100<br>2.80E-96<br>6.80E-90<br>3.40E-81<br>2.20E-80 |
| SED34923.1 | 3954029 | 3952169 | - | 619 | PF06738<br>PF12821 | ThrE<br>ThrE_2 | Putative threonine/serine exporter<br>Threonine/Serine exporter, ThrE | 7.50E-73<br>6.00E-19 |
| SED34966.1 | 3954890 | 3954170 | - | 239 | PF09335 | SNARE_assoc | SNARE associated Golgi protein | 5.40E-24 |
| SED35008.1 | 3955097 | 3955466 | + | 122 | PF01906 | YbjQ_1 | Putative heavy-metal-binding | 1.40E-33 |
| SED35039.1 | 3955635 | 3956298 | + | 220 | TIGR00949<br>PF01810<br>TIGR00948 | TIGR00949<br>LysE<br>TIGR00948 | 2A76: homoserine/Threonine efflux protein<br>LysE type translocator<br>2a75: L-lysine exporter | 1.40E-45<br>7.60E-44<br>1.30E-16 |

| index | leader | core | start | end | dir | score |
|---|---|---|---|---|---|---|
| 1 | MSVVANFANTELADVNPGRIGNDATPTM | LTPLAALATPEGVAVTSATAYALHEVTKDIQR | 3945025 | 3944842 | - | 33 |
| 2 | LSRERAERQVTRWTENLWRVSSLPLR | VRPLGSPMVGPTRSVRRSAAGFVREQS | 3952091 | 3951929 | - | 6 |

### Results for PJJ03557.1 [Streptomyces sp. 2333.5] back to top

previous - next

#### Architecture

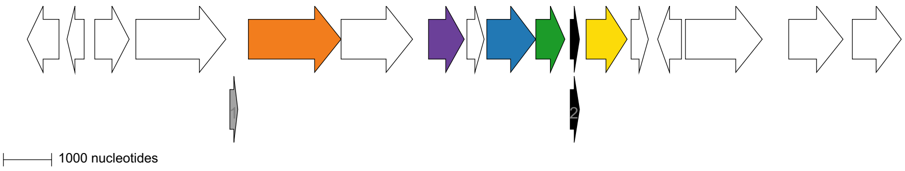

Link to nucleotide sequence

| Accession | start | end | direction | length (aa) | Pfam/HMM | name | description | E-value |
|---|---|---|---|---|---|---|---|---|
| PJJ03549.1 | 4463968 | 4463305 | - | 220 | TIGR00949<br>PF01810<br>TIGR00948 | TIGR00949<br>LysE<br>TIGR00948 | 2A76: homoserine/Threonine efflux protein<br>LysE type translocator<br>2a75: L-lysine exporter | 1.40E-45<br>7.60E-44<br>1.30E-16 |
| PJJ03550.1 | 4464506 | 4464137 | - | 122 | PF01906 | YbjQ_1 | Putative heavy-metal-binding | 1.40E-33 |
| PJJ03551.1 | 4464713 | 4465433 | + | 239 | PF09335 | SNARE_assoc | SNARE associated Golgi protein | 5.40E-24 |
| PJJ03552.1 | 4465574 | 4467434 | + | 619 | PF06738<br>PF12821 | ThrE<br>ThrE_2 | Putative threonine/serine exporter<br>Threonine/Serine exporter, ThrE | 7.50E-73<br>6.00E-19 |
| PJJ03553.1 | 4467893 | 4469831 | + | 645 | TIGR02204<br>TIGR02203<br>TIGR00958<br>TIGR03796<br>TIGR03375 | TIGR02204<br>TIGR02203<br>TIGR00958<br>TIGR03796<br>TIGR03375 | MsbA_rel: ABC transporter, permease/ATP-binding protein<br>MsbA_lipidA: lipid A export permease/ATP-binding protein MsbA<br>3a01208: antigen peptide transporter 2<br>NHLM_micro_ABC1: NHLM bacteriocin system ABC transporter, peptidase/ATP-binding protein<br>type_I_sec_LssB: type I secretion system ATPase | 5.70E-100<br>2.80E-96<br>6.80E-90<br>3.40E-81<br>2.20E-80 |
| PJJ03554.1 | 4469824 | 4471309 | + | 494 | TIGR02734<br>TIGR00292<br>PF13450<br>TIGR03467<br>PF01593 | TIGR02734<br>TIGR00292<br>NAD_binding_8<br>TIGR03467<br>Amino_oxidase | crtI_fam: phytoene desaturase<br>TIGR00292: thiazole biosynthesis enzyme<br>NAD(P)-binding Rossmann-like domain<br>HpnE: squalene-associated FAD-dependent desaturase<br>Flavin containing amine oxidoreductase | 8.80E-09<br>4.70E-08<br>5.30E-08<br>7.40E-08<br>3.40E-07 |
| PJJ03555.1 | 4471626 | 4472385 | + | 252 | PF13649<br>PF08241<br>PF13489<br>PF08242<br>PF13847 | Methyltransf_25<br>Methyltransf_11<br>Methyltransf_23<br>Methyltransf_12<br>Methyltransf_31 | Methyltransferase domain<br>Methyltransferase domain<br>Methyltransferase domain<br>Methyltransferase domain<br>Methyltransferase domain | 5.70E-17<br>1.90E-12<br>1.30E-10<br>5.60E-10<br>5.60E-09 |
| PJJ03556.1 | 4472427 | 4472775 | + | 115 | PF13518<br>PF01527<br>PF13551<br>PF13011<br>PF13384 | HTH_28<br>HTH_Tnp_1<br>HTH_29<br>LZ_Tnp_IS481<br>HTH_23 | Helix-turn-helix domain<br>Transposase<br>Winged helix-turn helix<br>leucine-zipper of insertion element IS481<br>Homeodomain-like domain | 4.20E-09<br>2.50E-06<br>1.30E-05<br>1.00E-04<br>4.20E-04 |
| PJJ03557.1 | 4472847 | 4473867 | + | 339 | LinE<br>LinH<br>PF12697<br>TIGR02427<br>PF00561 | LinE<br>LinH<br>Abhydrolase_6<br>TIGR02427<br>Abhydrolase_1 | LinE Homolog<br>LinH (Fusion of LinE/LinG)<br>Alpha/beta hydrolase family<br>protocat_pcaD: 3-oxoadipate enol-lactonase<br>alpha/beta hydrolase fold | 4.70E-104<br>4.10E-53<br>3.10E-15<br>8.80E-12<br>7.20E-10 |
| PJJ03558.1 | 4473863 | 4474451 | + | 195 | LinL | LinL | LinL | 2.50E-56 |
| PJJ03559.1 | 4474578 | 4474761 | + | 60 | LinA | LinA | LinA (linaridin precursor) | 1.80E-13 |
| PJJ03560.1 | 4474906 | 4475761 | + | 284 | LinG<br>LinH<br>TIGR04033 | LinG<br>LinH<br>TIGR04033 | LinG Homolog<br>LinH (Fusion of LinE/LinG)<br>export_SdpB: antimicrobial peptide system protein, SdpB family | 3.00E-111<br>3.70E-79<br>2.10E-04 |
| PJJ03561.1 | 4475826 | 4476186 | + | 119 | NO PFAM MATCH | - | - | - |
| PJJ03562.1 | 4476862 | 4476367 | - | 164 | PF00719 | Pyrophosphatase | Inorganic pyrophosphatase | 2.50E-53 |
| PJJ03563.1 | 4476948 | 4478556 | + | 535 | PF02113<br>TIGR00666 | Peptidase_S13<br>TIGR00666 | D-Ala-D-Ala carboxypeptidase 3 (S13) family<br>PBP4: D-alanyl-D-alanine carboxypeptidase/D-alanyl-D-alanine-endopeptidase | 6.00E-76<br>6.70E-58 |
| PJJ03564.1 | 4479104 | 4480223 | + | 372 | TIGR03883<br>TIGR03624<br>PF10103 | TIGR03883<br>TIGR03624<br>Zincin_2 | DUF2342_F420: uncharacterized protein, coenzyme F420 biosynthesis associated<br>TIGR03624: putative hydrolase<br>Zincin-like metallopeptidase | 7.20E-142<br>1.00E-125<br>3.80E-121 |
| PJJ03565.1 | 4480415 | 4481423 | + | 335 | TIGR02432<br>PF01171<br>PF09179 | TIGR02432<br>ATP_bind_3<br>TilS | lysidine_TilS_N: tRNA(Ile)-lysidine synthetase<br>PP-loop family<br>TilS substrate binding domain | 3.60E-43<br>6.60E-38<br>5.30E-12 |

| index | leader | core | start | end | dir | score |
|---|---|---|---|---|---|---|
| 1 | LSRERAERQVTRWTENLWRVSSLPLR | VRPLGSPMVGPTRSVRRSAAGFVREQS | 4467512 | 4467674 | + | 6 |
| 2 | MSVVANFANTELADVNPGRIGNDATPTM | LTPLAALATPEGVAVTSATAYALHEVTKDIQR | 4474578 | 4474761 | + | 33 |

### Results for SOE12061.1 [Streptomyces sp. 2323.1] back to top

previous - next

#### Architecture

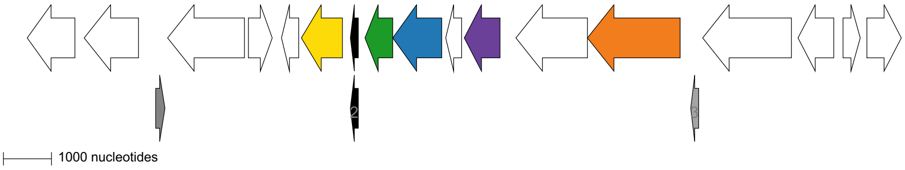

Link to nucleotide sequence

| Accession | start | end | direction | length (aa) | Pfam/HMM | name | description | E-value |
|---|---|---|---|---|---|---|---|---|
| SOE12052.1 | 3554055 | 3553047 | - | 335 | TIGR02432<br>PF01171<br>PF09179 | TIGR02432<br>ATP_bind_3<br>TilS | lysidine_TilS_N: tRNA(Ile)-lysidine synthetase<br>PP-loop family<br>TilS substrate binding domain | 3.60E-43<br>6.60E-38<br>5.30E-12 |
| SOE12053.1 | 3555366 | 3554247 | - | 372 | TIGR03883<br>TIGR03624<br>PF10103 | TIGR03883<br>TIGR03624<br>Zincin_2 | DUF2342_F420: uncharacterized protein, coenzyme F420 biosynthesis associated<br>TIGR03624: putative hydrolase<br>Zincin-like metallopeptidase | 7.20E-142<br>1.00E-125<br>3.80E-121 |
| SOE12055.1 | 3557581 | 3555973 | - | 535 | PF02113<br>TIGR00666 | Peptidase_S13<br>TIGR00666 | D-Ala-D-Ala carboxypeptidase 3 (S13) family<br>PBP4: D-alanyl-D-alanine carboxypeptidase/D-alanyl-D-alanine-endopeptidase | 9.20E-76<br>9.10E-58 |
| SOE12056.1 | 3557667 | 3558162 | + | 164 | PF00719 | Pyrophosphatase | Inorganic pyrophosphatase | 2.50E-53 |

| | | | | | | | | |
|---|---|---|---|---|---|---|---|---|
| SOE12057.1 | 3558703 | 3558343 | - | 119 | NO PFAM MATCH | - | - | - |
| SOE12058.1 | 3559621 | 3558766 | - | 284 | LinG<br>LinH<br>TIGR04033 | LinG<br>LinH<br>TIGR04033 | LinG Homolog<br>LinH (Fusion of LinE/LinG)<br>export_SdpB: antimicrobial peptide system protein, SdpB family | 3.00E-111<br>3.70E-79<br>2.10E-04 |
| SOE12059.1 | 3559949 | 3559766 | - | 60 | LinA | LinA | LinA (linaridin precursor) | 1.80E-13 |
| SOE12060.1 | 3560664 | 3560076 | - | 195 | LinL | LinL | LinL | 2.50E-56 |
| SOE12061.1 | 3561680 | 3560660 | - | 339 | LinE<br>LinH<br>PF12697<br>TIGR02427<br>PF00561 | LinE<br>LinH<br>Abhydrolase_6<br>TIGR02427<br>Abhydrolase_1 | LinE Homolog<br>LinH (Fusion of LinE/LinG)<br>Alpha/beta hydrolase family<br>protocat_pcaD: 3-oxoadipate enol-lactonase<br>alpha/beta hydrolase fold | 8.50E-104<br>5.00E-53<br>2.40E-15<br>8.80E-12<br>7.20E-10 |
| SOE12062.1 | 3562100 | 3561752 | - | 115 | PF13518<br>PF01527<br>PF13551<br>PF13011<br>PF13384 | HTH_28<br>HTH_Tnp_1<br>HTH_29<br>LZ_Tnp_IS481<br>HTH_23 | Helix-turn-helix domain<br>Transposase<br>Winged helix-turn helix<br>leucine-zipper of insertion element IS481<br>Homeodomain-like domain | 4.20E-09<br>2.50E-06<br>1.30E-05<br>1.00E-04<br>4.20E-04 |
| SOE12063.1 | 3562901 | 3562142 | - | 252 | PF13649<br>PF08241<br>PF13489<br>PF08242<br>PF13847 | Methyltransf_25<br>Methyltransf_11<br>Methyltransf_23<br>Methyltransf_12<br>Methyltransf_31 | Methyltransferase domain<br>Methyltransferase domain<br>Methyltransferase domain<br>Methyltransferase domain<br>Methyltransferase domain | 5.70E-17<br>1.90E-12<br>1.30E-10<br>5.60E-10<br>5.60E-09 |
| SOE12064.1 | 3564703 | 3563218 | - | 494 | TIGR02734<br>TIGR00292<br>PF13450<br>TIGR03467<br>PF01593 | TIGR02734<br>TIGR00292<br>NAD_binding_8<br>TIGR03467<br>Amino_oxidase | crtI_fam: phytoene desaturase<br>TIGR00292: thiazole biosynthesis enzyme<br>NAD(P)-binding Rossmann-like domain<br>HpnE: squalene-associated FAD-dependent desaturase<br>Flavin containing amine oxidoreductase | 1.60E-08<br>4.80E-08<br>5.40E-08<br>7.80E-08<br>5.10E-07 |
| SOE12065.1 | 3566634 | 3564696 | - | 645 | TIGR02204<br>TIGR02203<br>TIGR00958<br>TIGR03796<br>TIGR03375 | TIGR02204<br>TIGR02203<br>TIGR00958<br>TIGR03796<br>TIGR03375 | MsbA_rel: ABC transporter, permease/ATP-binding protein<br>MsbA_lipidA: lipid A export permease/ATP-binding protein MsbA<br>3a01208: antigen peptide transporter 2<br>NHLM_micro_ABC1: NHLM bacteriocin system ABC transporter, peptidase/ATP-binding protein<br>type_I_sec_LssB: type I secretion system ATPase | 2.60E-100<br>3.00E-96<br>2.30E-90<br>5.70E-81<br>3.40E-80 |
| SOE12066.1 | 3568953 | 3567093 | - | 619 | PF06738<br>PF12821 | ThrE<br>ThrE_2 | Putative threonine/serine exporter<br>Threonine/Serine exporter, ThrE | 7.50E-73<br>6.00E-19 |
| SOE12067.1 | 3569815 | 3569095 | - | 239 | PF09335 | SNARE_assoc | SNARE associated Golgi protein | 5.40E-24 |
| SOE12068.1 | 3570022 | 3570391 | + | 122 | PF01906 | YbjQ_1 | Putative heavy-metal-binding | 1.40E-33 |
| SOE12069.1 | 3570524 | 3571223 | + | 232 | TIGR00949<br>PF01810<br>TIGR00948 | TIGR00949<br>LysE<br>TIGR00948 | 2A76: homoserine/Threonine efflux protein<br>LysE type translocator<br>2a75: L-lysine exporter | 1.10E-45<br>6.80E-44<br>1.50E-16 |

| index | leader | core | start | end | dir | score |
|---|---|---|---|---|---|---|
| 1 | VGAGLIRRPRADEGRGPPPPACSHP...TGGAAPQDEVRRSGTAPNHPRPPPG | ATATNHHGGKADNVRGAA | 3555715 | 3555925 | + | 7 |
| 2 | MSVVANFANTELADVNPGRIGNDATPTM | LTPLAALATPEGVAVTSATAYALHEVTKDIQR | 3559949 | 3559766 | - | 33 |
| 3 | LSRERAERQVTRWTENLWRVSSLPL | RVRPLGSPMVGPTRSVRRSAAGFVRE | 3567015 | 3566859 | - | 6 |

### Results for OXY94523.1 [Streptomyces sp. 2R] back to top

previous - next

#### Architecture

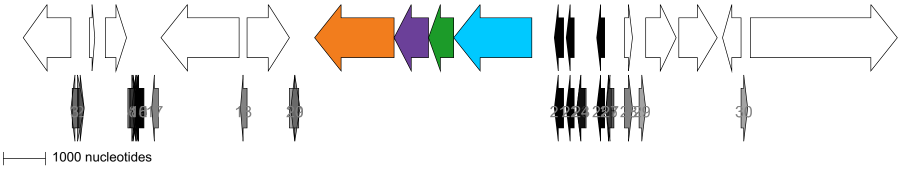

Link to nucleotide sequence

| Accession | start | end | direction | length (aa) | Pfam/HMM | name | description | E-value |
|---|---|---|---|---|---|---|---|---|
| OXY94516.1 | 9008 | 7883 | - | 374 | PF00724<br>TIGR03997<br>TIGR03996 | Oxidored_FMN<br>TIGR03997<br>TIGR03996 | NADH:flavin oxidoreductase / NADH oxidase family<br>mycofact_OYE_2: mycofactocin system FadH/OYE family oxidoreductase 2<br>mycofact_OYE_1: mycofactocin system FadH/OYE family oxidoreductase 1 | 8.90E-56<br>1.30E-43<br>1.90E-23 |
| WP_020123447.1 | 9441 | 9567 | + | 42 | INFERRED GENE | - | - | - |
| OXY94517.1 | 9826 | 10309 | + | 160 | NO PFAM MATCH | - | - | - |
| OXY94518.1 | 12981 | 11127 | - | 617 | NO PFAM MATCH | - | - | - |
| OXY94519.1 | 13170 | 14184 | + | 337 | NO PFAM MATCH | - | - | - |
| OXY94520.1 | 16685 | 14783 | - | 633 | TIGR02204<br>TIGR02203<br>TIGR00958<br>TIGR03796<br>TIGR02857 | TIGR02204<br>TIGR02203<br>TIGR00958<br>TIGR03796<br>TIGR02857 | MsbA_rel: ABC transporter, permease/ATP-binding protein<br>MsbA_lipidA: lipid A export permease/ATP-binding protein MsbA<br>3a01208: antigen peptide transporter 2<br>NHLM_micro_ABC1: NHLM bacteriocin system ABC transporter, peptidase/ATP-binding protein<br>CydD: thiol reductant ABC exporter, CydD subunit | 7.20E-99<br>1.50E-92<br>3.70E-83<br>2.50E-79<br>1.20E-75 |
| OXY94521.1 | 17494 | 16681 | - | 270 | PF13649<br>PF08241<br>PF13847<br>PF13489<br>PF08242 | Methyltransf_25<br>Methyltransf_11<br>Methyltransf_31<br>Methyltransf_23<br>Methyltransf_12 | Methyltransferase domain<br>Methyltransferase domain<br>Methyltransferase domain<br>Methyltransferase domain<br>Methyltransferase domain | 2.30E-16<br>2.90E-12<br>1.70E-07<br>9.20E-07<br>1.00E-06 |
| OXY94522.1 | 18099 | 17490 | - | 202 | LinL | LinL | LinL | 4.40E-64 |

| | | | | | | | | |
|---|---|---|---|---|---|---|---|---|
| OXY94523.1 | 19943 | 18095 | - | 615 | LinH<br>LinG<br>LinE<br>LinL<br>PF12697 | LinH<br>LinG<br>LinE<br>LinL<br>Abhydrolase_6 | LinH (Fusion of LinE/LinG)<br>LinG Homolog<br>LinE Homolog<br>LinL<br>Alpha/beta hydrolase family | 1.70E-252<br>1.40E-111<br>5.20E-94<br>2.60E-37<br>2.00E-15 |
| OXY94524.1 | 20691 | 20490 | - | 66 | LinA | LinA | LinA (linaridin precursor) | 4.50E-12 |
| OXY94525.1 | 20940 | 20748 | - | 63 | LinA | LinA | LinA (linaridin precursor) | 2.30E-11 |
| OXY94526.1 | 21682 | 21478 | - | 67 | LinA | LinA | LinA (linaridin precursor) | 1.10E-11 |
| OXY94527.1 | 22150 | 22333 | + | 60 | NO PFAM MATCH | - | - | - |
| OXY94528.1 | 22644 | 23364 | + | 239 | PF00456<br>TIGR00232<br>PF02775<br>TIGR03181 | Transketolase_N<br>TIGR00232<br>TPP_enzyme_C<br>TIGR03181 | Transketolase, thiamine diphosphate binding domain<br>tktlase_bact: transketolase<br>Thiamine pyrophosphate enzyme, C-terminal TPP binding domain<br>PDH_E1_alph_x: pyruvate dehydrogenase (acetyl-transferring) E1 component, alpha subunit | 7.10E-25<br>7.70E-25<br>7.60E-05<br>2.90E-04 |
| OXY94529.1 | 23429 | 24332 | + | 300 | PF02779<br>TIGR00204<br>PF02780 | Transket_pyr<br>TIGR00204<br>Transketolase_C | Transketolase, pyrimidine binding domain<br>dxs: 1-deoxy-D-xylulose-5-phosphate synthase<br>Transketolase, C-terminal domain | 2.50E-19<br>1.40E-12<br>3.00E-06 |
| OXY94530.1 | 24910 | 24481 | - | 142 | PF04237<br>PF04993 | YjbR<br>TfoX_N | YjbR<br>TfoX N-terminal domain | 4.00E-18<br>6.30E-04 |
| OXY94531.1 | 25132 | 28633 | + | 1166 | PF06283<br>PF07995<br>TIGR03606<br>PF00801<br>PF03422 | ThuA<br>GSDH<br>TIGR03606<br>PKD<br>CBM_6 | Trehalose utilisation<br>Glucose / Sorbosone dehydrogenase<br>non_repeat_PQQ: dehydrogenase, PQQ-dependent, s-GDH family<br>PKD domain<br>Carbohydrate binding module (family 6) | 3.10E-72<br>2.20E-33<br>7.80E-21<br>8.60E-21<br>9.50E-20 |

| index | leader | core | start | end | dir | score |
|---|---|---|---|---|---|---|
| 1 | VHHPSPTPRWTGSIPASRRYGTGPP...PCGFPLVPGRARGGGGGGAEKRPSR | PTVSATSYSVLEDDR | 9028 | 9319 | + | 6 |
| 2 | LLRTGPPVRCRRTPCGFPLVPGRARGGGGGGAEKRPSR | PTVSATSYSVLEDDR | 9157 | 9319 | + | 7 |
| 3 | VRRQRTGGPVRSNPNRPLIVSGGCSIRLPPS | GGPVPYRREAGMLPVHRGVGEGWCTPQPMNRN | 9195 | 9003 | - | 6 |
| 4 | LRLIQALAGGRSAVAELVELEGLGTTGQIYHHLR | QLVSAGWLETTGRGRYQIPPARLVPLLVVVTEARR | 10365 | 10575 | + | 6 |
| 5 | LAGGRSAVAELVELEGLGTTGQIYHHLRQLV | SAGWLETTGRGRYQIPPARLVPLLVVVTEARR | 10383 | 10575 | + | 6 |
| 6 | VELEGLGTTGQIYHHLRQLVSAGWLE | TTGRGRYQIPPARLVPLLVVVTEARR | 10416 | 10575 | + | 7 |
| 7 | MGEVCADRVAELPSQR | RASVTTTSSGTSRAGGI | 10616 | 10514 | - | 6 |
| 8 | VLCLDGRGMRRPSC | RTTQSAASLGDHDQQRNEPGRRNLISPTSGRLQPAGGNELS | 10629 | 10461 | - | 6 |
| 9 | MACSPADALFGAIRLARCFAWMGEVCA | DRVAELPSQRRASVTTTSSGTSRAGGI | 10679 | 10514 | - | 8 |
| 10 | VAMACSPADALFGAIRLARCFAWMGEVC | ADRVAELPSQRRASVTTTSSGTSRAGGI | 10685 | 10514 | - | 8 |
| 11 | LRVAMACSPADALFGAIRLARCFAWMGEV | CADRVAELPSQRRASVTTTSSGTSRAGGI | 10691 | 10514 | - | 6 |
| 12 | LRSGRQPLRVAMACSPADALFGAIRLARCFAW | MGEVCADRVAELPSQRRASVTTTSSGTSRAGGI | 10712 | 10514 | - | 7 |
| 13 | VLQRSLRSGRQPLRVAMACSPADALFGAIRLARCF | AWMGEVCADRVAELPSQRRASVTTTSSGTSRAGGI | 10727 | 10514 | - | 7 |
| 14 | LVLQRSLRSGRQPLRVAMACSPADALFGAIRLARC | FAWMGEVCADRVAELPSQRRASVTTTSSGTSRAGGI | 10730 | 10514 | - | 6 |
| 15 | VIRLVLQRSLRSGRQPLRVAMACSPADALFGAIRLAR | CFAWMGEVCADRVAELPSQRRASVTTTSSGTSRAGGI | 10739 | 10514 | - | 6 |
| 16 | MSVIRLVLQRSLRSGRQPLRVAMACSPADALFGAIRLA | RCFAWMGEVCADRVAELPSQRRASVTTTSSGTSRAGGI | 10745 | 10514 | - | 6 |
| 17 | VGGAGCPCGGPRPARPCRDQELRHRLE | GVAAGGEGVDEETGAQVIRDRPAGQMP | 11090 | 10925 | - | 7 |
| 18 | VGPHGPSLEHVRRDDGAQVCPGR | RGVRSQQSLTPRWYQGSSVCVTT | 13182 | 13041 | - | 7 |
| 19 | VTPGPRTGAGCHAAGVPAPTRPSAG...PHMPCTRVRAGWPPSRGGRQRGPTG | VPPGAGRNTVCRRPIAPVAVSPPA | 14180 | 14453 | + | 6 |
| 20 | VFRPAPGGTPVGPRC | RPPLLGGHPARTRVHGMCGLLAARSALRGEFGRPLPALGRVGAGTPAA | 14404 | 14212 | - | 6 |
| 21 | MSTSSTPAALRGAGGLSESDPGRALSTLAPVA | ATPGIVAGAALGVALVNAFVAGYNHGGGNVELPM | 20691 | 20490 | - | 31 |
| 22 | MSKSSAPAVLTATGALTESDPGVTLPALAPVA | ATPVAIAATLGAAFIAGYAAGRAAGNVELPM | 20940 | 20748 | - | 28 |
| 23 | LSTASIPAVLSEAGALSESDRGRAPLD | TVPVASVTFTMTACVGISVCLTGSTIELPQ | 21187 | 21013 | - | 9 |
| 24 | MTSPHRPPTSEEKSLSTASIPAVLSEAGALSESDRGRAPLD | TVPVASVTFTMTACVGISVCLTGSTIELPQ | 21229 | 21013 | - | 8 |
| 25 | MPEFRQPGWTRGIAPLDESAGGQVFGGASPVA | ATPAVVATAGAVVIAFAAGVAARQLANGGNVELPM | 21682 | 21478 | - | 30 |
| 26 | MPAELSSLSAGRSASCGAH | HRTSVHAGVGSRVRSRAPG | 21836 | 21719 | - | 6 |
| 27 | VPEPAACCFGVRILMPAELSSLSAGR | SASCGAHHRTSVHAGVGSRVRSRAPG | 21878 | 21719 | - | 8 |
| 28 | MRVAPAPGHLAVHMITNGAVDRTVQNWIADELSR | STVTNTIAVLVRVTEQSLTSPIVMTR | 22150 | 22333 | + | 7 |
| 29 | VRYQAAPRPVARLTWGFPWLNAHENN | TALGRVVTNRFLRLTSTLVDVERSVS | 22489 | 22648 | + | 6 |
| 30 | LRSSRFSDRVAPGGRCPGFDRNARDRHPPPA | GTPGTPGRTGPADPLTPSKR | 24900 | 25056 | + | 6 |

### Results for RIA63746.1 [Streptomyces sp. 3211.1] back to top

previous - next

#### Architecture

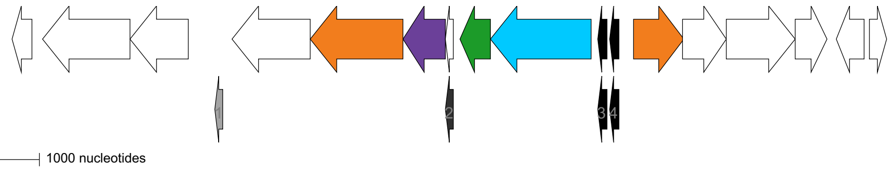

Link to nucleotide sequence

| Accession | start | end | direction | length (aa) | Pfam/HMM | name | description | E-value |
|---|---|---|---|---|---|---|---|---|
| RIA63738.1 | 478783 | 478405 | - | 125 | NO PFAM MATCH | - | - | - |
| RIA63739.1 | 480646 | 478966 | - | 559 | TIGR01613<br>PF08706 | TIGR01613<br>D5_N | primase_Cterm: phage/plasmid primase, P4 family, C-terminal domain<br>D5 N terminal like | 6.30E-76<br>1.50E-27 |
| RIA63740.1 | 481752 | 480642 | - | 369 | PF09250 | Prim-Pol | Bifunctional DNA primase/polymerase, N-terminal | 4.00E-36 |
| RIA63741.1 | 484083 | 482598 | - | 494 | TIGR02734<br>PF13450<br>TIGR03467<br>TIGR02733<br>TIGR01318 | TIGR02734<br>NAD_binding_8<br>TIGR03467<br>TIGR02733<br>TIGR01318 | crtI_fam: phytoene desaturase<br>NAD(P)-binding Rossmann-like domain<br>HpnE: squalene-associated FAD-dependent desaturase<br>desat_CrtD: C-3',4' desaturase CrtD<br>gltD_gamma_fam: glutamate synthase, small subunit | 1.20E-11<br>6.20E-09<br>3.20E-08<br>8.30E-08<br>3.00E-07 |
| RIA63742.1 | 485837 | 484079 | - | 585 | TIGR02204<br>TIGR02203<br>TIGR03796<br>TIGR00958<br>TIGR01846 | TIGR02204<br>TIGR02203<br>TIGR03796<br>TIGR00958<br>TIGR01846 | MsbA_rel: ABC transporter, permease/ATP-binding protein<br>MsbA_lipidA: lipid A export permease/ATP-binding protein MsbA<br>NHLM_micro_ABC1: NHLM bacteriocin system ABC transporter, peptidase/ATP-binding protein<br>3a01208: antigen peptide transporter 2<br>type_I_sec_HlyB: type I secretion system ATPase | 3.00E-102<br>2.00E-96<br>4.90E-87<br>1.50E-86<br>5.50E-86 |
| RIA63743.1 | 486652 | 485833 | - | 272 | PF13649<br>PF08241<br>PF13489<br>PF08242<br>PF13847 | Methyltransf_25<br>Methyltransf_11<br>Methyltransf_23<br>Methyltransf_12<br>Methyltransf_31 | Methyltransferase domain<br>Methyltransferase domain<br>Methyltransferase domain<br>Methyltransferase domain<br>Methyltransferase domain | 6.80E-16<br>3.20E-12<br>4.30E-08<br>3.50E-07<br>8.00E-07 |
| RIA63744.1 | 486810 | 486648 | - | 53 | NO PFAM MATCH | - | - | - |
| RIA63745.1 | 487503 | 486921 | - | 193 | LinL | LinL | LinL | 7.80E-68 |
| RIA63746.1 | 489434 | 487499 | - | 644 | LinH<br>LinG<br>LinE<br>PF12697<br>PF00561 | LinH<br>LinG<br>LinE<br>Abhydrolase_6<br>Abhydrolase_1 | LinH (Fusion of LinE/LinG)<br>LinG Homolog<br>LinE Homolog<br>Alpha/beta hydrolase family<br>alpha/beta hydrolase fold | 6.80E-252<br>2.40E-131<br>4.20E-100<br>1.20E-16<br>3.10E-09 |
| RIA63747.1 | 489729 | 489564 | - | 54 | LinA | LinA | LinA (linaridin precursor) | 2.10E-14 |
| RIA63748.1 | 489972 | 489771 | - | 66 | LinA | LinA | LinA (linaridin precursor) | 6.90E-14 |
| RIA63749.1 | 490229 | 491186 | + | 318 | TIGR01188<br>TIGR03864<br>TIGR01288<br>TIGR01166<br>TIGR03740 | TIGR01188<br>TIGR03864<br>TIGR01288<br>TIGR01166<br>TIGR03740 | drrA: daunorubicin resistance ABC transporter, ATP-binding protein<br>PQQ_ABC_ATP: ABC transporter, ATP-binding subunit, PQQ-dependent alcohol dehydrogenase system<br>nodI: nodulation ABC transporter NodI<br>cbiO: cobalt ABC transporter, ATP-binding protein<br>galliderm_ABC: lantibiotic protection ABC transporter, ATP-binding subunit | 1.50E-61<br>9.10E-44<br>6.10E-40<br>9.00E-33<br>3.20E-32 |
| RIA63750.1 | 491182 | 492007 | + | 274 | PF01061<br>PF12698<br>TIGR03062<br>TIGR01291<br>PF12679 | ABC2_membrane<br>ABC2_membrane_3<br>TIGR03062<br>TIGR01291<br>ABC2_membrane_2 | ABC-2 type transporter<br>ABC-2 family transporter protein<br>pip_yhgE_Cterm: YhgE/Pip C-terminal domain<br>nodJ: ABC-2 type transporter, NodJ family<br>ABC-2 family transporter protein | 3.50E-14<br>2.60E-11<br>5.60E-08<br>6.40E-05<br>8.70E-04 |
| RIA63751.1 | 492003 | 493314 | + | 436 | PF07730 | HisKA_3 | Histidine kinase | 3.70E-14 |
| RIA63752.1 | 493310 | 493919 | + | 202 | PF00072<br>TIGR02875<br>PF00196<br>TIGR02956<br>TIGR02154 | Response_reg<br>TIGR02875<br>GerE<br>TIGR02956<br>TIGR02154 | Response regulator receiver domain<br>spore_0_A: sporulation transcription factor Spo0A<br>Bacterial regulatory proteins, luxR family<br>TMAO_torS: TMAO reductase sytem sensor TorS<br>PhoB: phosphate regulon transcriptional regulatory protein PhoB | 1.30E-22<br>6.90E-18<br>2.00E-12<br>5.20E-10<br>8.90E-09 |
| RIA63753.1 | 494624 | 494096 | - | 175 | PF13302<br>PF00583 | Acetyltransf_3<br>Acetyltransf_1 | Acetyltransferase (GNAT) domain<br>Acetyltransferase (GNAT) family | 1.70E-31<br>1.80E-05 |
| RIA63754.1 | 494719 | 495067 | + | 115 | PF00392<br>TIGR02325<br>TIGR02018<br>TIGR03338<br>TIGR02404 | GntR<br>TIGR02325<br>TIGR02018<br>TIGR03338<br>TIGR02404 | Bacterial regulatory proteins, gntR family<br>C_P_lyase_phnF: phosphonate metabolism transcriptional regulator PhnF<br>his_ut_repres: histidine utilization repressor<br>phnR_burk: phosphonate utilization associated transcriptional regulator<br>trehalos_R_Bsub: trehalose operon repressor | 2.90E-17<br>7.70E-13<br>1.10E-11<br>4.00E-06<br>4.50E-06 |

| index | leader | core | start | end | dir | score |
|---|---|---|---|---|---|---|
| 1 | VGRLRSLSLQSLPKPAGQGVEAGGPGT | EGTQLSSYEPDMNYQASYTPARIHART | 482420 | 482255 | - | 6 |
| 2 | MHKPAVAGCLLIGLAVIGF | LTAADVMNTVSAIAVGVALFIAAGVELKRKKPNR | 486810 | 486648 | - | 9 |
| 3 | MNEVTALLDEMDIPEEGVVFPVAAPVL | ATPAAFMGGIAISCAVVSAFEAGRGNG | 489729 | 489564 | - | 30 |
| 4 | MTEIVTFVNNLDLGTIQAGETRTDQAPVL | ATPAAALAAIGGAKAVGAFVGGATVVGGAYALGRAIG | 489972 | 489771 | - | 29 |

### Results for SHI10722.1 [Streptomyces sp. 3214.6] back to top

### Architecture

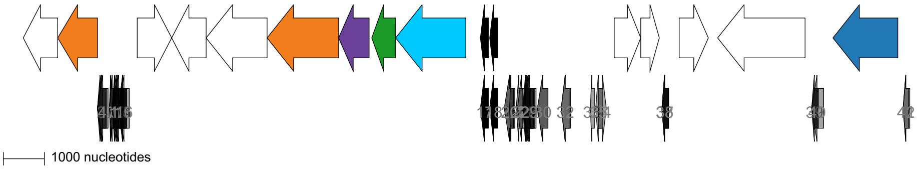

Link to nucleotide sequence

| Accession | start | end | direction | length (aa) | Pfam/HMM | name | description | E-value |
|---|---|---|---|---|---|---|---|---|
| SHI10635.1 | 4444158 | 4443306 | - | 283 | PF12698<br>PF01061<br>TIGR01247<br>TIGR00025<br>PF12679 | ABC2_membrane_3<br>ABC2_membrane<br>TIGR01247<br>TIGR00025<br>ABC2_membrane_2 | ABC-2 family transporter protein<br>ABC-2 type transporter<br>drrB: daunorubicin resistance ABC transporter membrane protein<br>Mtu_efflux: ABC transporter efflux protein, DrrB family<br>ABC-2 family transporter protein | 2.60E-18<br>1.40E-13<br>1.90E-10<br>1.10E-08<br>2.60E-05 |
| SHI10645.1 | 4445126 | 4444154 | - | 323 | TIGR01188<br>TIGR03864<br>TIGR03522<br>TIGR03873<br>TIGR01288 | TIGR01188<br>TIGR03864<br>TIGR03522<br>TIGR03873<br>TIGR01288 | drrA: daunorubicin resistance ABC transporter, ATP-binding protein<br>PQQ_ABC_ATP: ABC transporter, ATP-binding subunit, PQQ-dependent alcohol dehydrogenase system<br>GldA_ABC_ATP: gliding motility-associated ABC transporter ATP-binding subunit GldA<br>F420-0_ABC_ATP: proposed F420-0 ABC transporter, ATP-binding protein<br>nodI: nodulation ABC transporter NodI | 9.20E-58<br>7.00E-46<br>2.90E-40<br>7.10E-39<br>2.70E-38 |
| SHI10658.1 | 4446094 | 4446937 | + | 280 | PF01551 | Peptidase_M23 | Peptidase family M23 | 7.60E-16 |
| SHI10669.1 | 4447773 | 4446921 | - | 283 | PF12680<br>PF14534 | SnoaL_2<br>DUF4440 | SnoaL-like domain<br>Domain of unknown function (DUF4440) | 6.30E-05<br>2.50E-04 |
| SHI10682.1 | 4449275 | 4447769 | - | 501 | TIGR02734<br>TIGR00292<br>TIGR02733<br>TIGR03467<br>TIGR01317 | TIGR02734<br>TIGR00292<br>TIGR02733<br>TIGR03467<br>TIGR01317 | crtI_fam: phytoene desaturase<br>TIGR00292: thiazole biosynthesis enzyme<br>desat_CrtD: C-3',4' desaturase CrtD<br>HpnE: squalene-associated FAD-dependent desaturase<br>GOGAT_sm_gam: glutamate synthase, NADH/NADPH, small subunit | 2.70E-10<br>5.60E-08<br>7.10E-08<br>9.40E-08<br>1.70E-07 |
| SHI10690.1 | 4451005 | 4449271 | - | 577 | TIGR02204<br>TIGR02203<br>TIGR00958<br>TIGR03796<br>TIGR01846 | TIGR02204<br>TIGR02203<br>TIGR00958<br>TIGR03796<br>TIGR01846 | MsbA_rel: ABC transporter, permease/ATP-binding protein<br>MsbA_lipidA: lipid A export permease/ATP-binding protein MsbA<br>3a01208: antigen peptide transporter 2<br>NHLM_micro_ABC1: NHLM bacteriocin system ABC transporter, peptidase/ATP-binding protein<br>type_I_sec_HlyB: type I secretion system ATPase | 3.50E-102<br>6.10E-99<br>4.00E-86<br>5.20E-86<br>4.30E-83 |
| SHI10702.1 | 4451757 | 4451001 | - | 251 | PF13649<br>PF08241<br>PF13489<br>PF08242<br>TIGR01983 | Methyltransf_25<br>Methyltransf_11<br>Methyltransf_23<br>Methyltransf_12<br>TIGR01983 | Methyltransferase domain<br>Methyltransferase domain<br>Methyltransferase domain<br>Methyltransferase domain<br>UbiG: 3-demethylubiquinone-9 3-O-methyltransferase | 6.90E-13<br>1.80E-08<br>4.00E-08<br>2.60E-06<br>4.50E-05 |
| SHI10714.1 | 4452390 | 4451811 | - | 192 | LinL | LinL | LinL | 4.30E-66 |
| SHI10722.1 | 4454108 | 4452386 | - | 573 | LinH<br>LinE<br>LinG<br>PF12697<br>PF00561 | LinH<br>LinE<br>LinG<br>Abhydrolase_6<br>Abhydrolase_1 | LinH (Fusion of LinE/LinG)<br>LinE Homolog<br>LinG Homolog<br>Alpha/beta hydrolase family<br>alpha/beta hydrolase fold | 7.00E-221<br>6.50E-99<br>3.80E-94<br>5.50E-15<br>4.00E-11 |
| SHI10732.1 | 4454647 | 4454455 | - | 63 | LinA | LinA | LinA (linaridin precursor) | 1.40E-12 |
| SHI10744.1 | 4454888 | 4454693 | - | 64 | LinA | LinA | LinA (linaridin precursor) | 1.10E-17 |
| SHI10757.1 | 4457748 | 4458390 | + | 213 | PF04255<br>PF13411 | DUF433<br>MerR_1 | Protein of unknown function (DUF433)<br>MerR HTH family regulatory protein | 3.20E-16<br>8.80E-04 |
| SHI10768.1 | 4458389 | 4458830 | + | 146 | PF18478<br>PF18480 | PIN_10<br>DUF5615 | PIN like domain<br>Domain of unknown function (DUF5615) | 4.20E-16<br>1.00E-07 |
| SHI10778.1 | 4459331 | 4460027 | + | 231 | NO PFAM MATCH | - | - | - |
| SHI10788.1 | 4462388 | 4460261 | - | 708 | PF07591<br>TIGR03696<br>PF15542<br>PF05203<br>PF01079 | PT-HINT<br>TIGR03696<br>Ntox50<br>Hom_end_hint<br>Hint | Pretoxin HINT domain<br>Rhs_assc_core: RHS repeat-associated core domain<br>Bacterial toxin 50<br>Hom_end-associated Hint<br>Hint module | 1.80E-22<br>2.00E-19<br>4.50E-06<br>9.80E-06<br>2.50E-05 |
| SHI10801.1 | 4464658 | 4463080 | - | 525 | PF00561<br>PF08386<br>LinE<br>TIGR01249 | Abhydrolase_1<br>Abhydrolase_4<br>LinE<br>TIGR01249 | alpha/beta hydrolase fold<br>TAP-like protein<br>LinE Homolog<br>pro_imino_pep_1: prolyl aminopeptidase | 5.40E-38<br>2.00E-33<br>3.40E-06<br>4.20E-06 |

| index | leader | core | start | end | dir | score |
|---|---|---|---|---|---|---|
| 1 | LQREGGGADGGQTRSRDHEPRP | TGPLTRHTDTHRPRSMDRDGEHL | 4445260 | 4445122 | - | 6 |
| 2 | LCPVSALSTVGESRACLQREGGGADGGQTR | SRDHEPRPTGPLTRHTDTHRPRSMDRDGEHL | 4445308 | 4445122 | - | 8 |
| 3 | MCDCSPCGDCWKLSRDRLCPVSALSTVGESRACLQREGG | GADGGQTRSRDHEPRPTGPLTRHTDTHRPRSMDRDGEHL | 4445359 | 4445122 | - | 6 |
| 4 | MGGLMCDCSPCGDCWKLSRDRLCPVSALSTVGESRACLQRE | GGGADGGQTRSRDHEPRPTGPLTRHTDTHRPRSMDRDGEHL | 4445371 | 4445122 | - | 6 |
| 5 | LKDYALDGSLLMFRVSST | NPTLRSAPASALAGRSIL | 4445512 | 4445401 | - | 6 |
| 6 | LLSWCSSGITWVSLKDYALDGSLL | MFRVSSTNPTLRSAPASALAGRSIL | 4445551 | 4445401 | - | 6 |
| 7 | VFVLLSWCSSGITWVSLKDYALDGSL | LMFRVSSTNPTLRSAPASALAGRSIL | 4445560 | 4445401 | - | 8 |

| | | | | | | |
|---|---|---|---|---|---|---|
| 8 | VAAEFAGLAAAEFAEEDKAAM | TTAMVESVRFAVLVFIWNHLGFLEGLCTGWFLADV | 4445642 | 4445471 | - | 7 |
| 9 | VPDALDEPWVADHSTGVAAEFAGLAAAEFAEEDKAAM | TTAMVESVRFAVLVFIWNHLGFLEGLCTGWFLADV | 4445690 | 4445471 | - | 6 |
| 10 | LTARTVLLPFCSVHRVPDALDEPWV...HSTGVAAEFAGLAAAEFAEEDKAAM | TTAMVESVRFAVLVFIWNHLGFLEGLCTGWFLADV | 4445735 | 4445471 | - | 7 |
| 11 | VFWISPLTARTVLLPFCSVHRVPDA...HSTGVAAEFAGLAAAEFAEEDKAAM | TTAMVESVRFAVLVFIWNHLGFLEGLCTGWFLADV | 4445753 | 4445471 | - | 7 |
| 12 | VPVFWISPLTARTVLLPFCSVHRVP...HSTGVAAEFAGLAAAEFAEEDKAAM | TTAMVESVRFAVLVFIWNHLGFLEGLCTGWFLADV | 4445759 | 4445471 | - | 7 |
| 13 | MKSRVHWTQTEKHSVPVFWISPLTA...HSTGVAAEFAGLAAAEFAEEDKAAM | TTAMVESVRFAVLVFIWNHLGFLEGLCTGWFLADV | 4445801 | 4445471 | - | 6 |
| 14 | MFGIGGRDEVTGPLDPDRKAFGACVLDFSVDRENCPAAVLF | GPPGAGRTRRTVGCRPLHRSGR | 4445823 | 4445631 | - | 7 |
| 15 | LSGMFGIGGRDEVTGPLDPDRKAFGACVLDFSVDRENCPAAVLF | GPPGAGRTRRTVGCRPLHRSGR | 4445832 | 4445631 | - | 7 |
| 16 | LIWAQSKVDGRDSGAALSGMFGIGG...DRKAFGACVLDFSVDRENCPAAVLF | GPPGAGRTRRTVGCRPLHRSGR | 4445880 | 4445631 | - | 6 |
| 17 | MSRIVDSVVAVTSADLTDDSQGEFGAMAPVQ | ATPTLVAFGVGFAGGAGAAWVTVQAYEAGAND | 4454647 | 4454455 | - | 31 |
| 18 | MEQKIAAFMNEQDLGSLEQGYRDDAPMPVL | ATPAAAVATCWYVGGAAAGAAAVVGAYVTGRVVG | 4454888 | 4454693 | - | 30 |
| 19 | MRLNLDTGIGAGGAAT | TFFMQGAIADLFSPAA | 4455288 | 4455189 | - | 6 |
| 20 | MGEQVHEVEPGYGHWCRWCCDHVLHAGSDCGPVFPRCLTCLPAW | TLPGKDRAHTSQATPDVKTSYAFSRKFTLHSAFGVTRVASDR | 4455304 | 4455043 | - | 8 |
| 21 | VALPSGSAATFLQGIAGRPLGRCIR | GVRAVVLLTSRQGLVTDHYRPIGCE | 4455493 | 4455340 | - | 7 |
| 22 | MVRIRPGRLHGLFSLISSRYCGVAFGKCSN | VLAGNCGSTARSLHPRRAGRRPAHQPTGASD | 4455554 | 4455368 | - | 6 |
| 23 | MSLSWEGTGHSITWTLLVGA | WVPFVKSQVDPRVEVRGGVRSAWFGSGQDGFMACSL | 4455679 | 4455508 | - | 6 |
| 24 | MSGIYDSMSLSWEGTGHSITWTLLVGA | WVPFVKSQVDPRVEVRGGVRSAWFGSGQDGFMACSL | 4455700 | 4455508 | - | 6 |
| 25 | MANAEIDQGACGRCGSMSGIYDSMSLSWEGTGHSITWTLLVGA | WVPFVKSQVDPRVEVRGGVRSAWFGSGQDGFMACSL | 4455748 | 4455508 | - | 6 |
| 26 | MGGLMANAEIDQGACGRCGSMSGIYDSMSLSWEGTGHSITWTLLVGA | WVPFVKSQVDPRVEVRGGVRSAWFGSGQDGFMACSL | 4455760 | 4455508 | - | 6 |
| 27 | VASPLMGGLMANAEIDQGACGRCGS...GIYDSMSLSWEGTGHSITWTLLVGA | WVPFVKSQVDPRVEVRGGVRSAWFGSGQDGFMACSL | 4455775 | 4455508 | - | 6 |
| 28 | LTPYVASPLMGGLMANAEIDQGACG...GIYDSMSLSWEGTGHSITWTLLVGA | WVPFVKSQVDPRVEVRGGVRSAWFGSGQDGFMACSL | 4455787 | 4455508 | - | 6 |
| 29 | VLELALWWSTLTPYVASPLMGGLMA...AEIDQGACGRCGSMSGIYDSMSLSW | EGTGHSITWTLLVGAWVPFVKSQVD...RVEVRGGVRSAWFGSGQDGFMACSL | 4455817 | 4455508 | - | 6 |
| 30 | MDATSETTCREALLRQACRRAKVSA...EPGRKSAAGWQIRPGVVVDRTVPAS | VTPRSGRRTDHSVSCVR | 4456106 | 4455875 | - | 8 |
| 31 | VAAPCRRSCSRTGGRSEAATRRSKRSVTFPGWSQ | EPSSWVKTMPVSTQADLHSWRSASCCSRNRISVS | 4456647 | 4456440 | - | 6 |
| 32 | MSVAAPCRRSCSRTGGRSEAATRRSKRSVTFPGWS | QEPSSWVKTMPVSTQADLHSWRSASCCSRNRISVS | 4456653 | 4456440 | - | 6 |
| 33 | MASPLSGHSWGQDGAGR | TVKRHERSTRPGRVHAC | 4457251 | 4457146 | - | 6 |
| 34 | MEGILDVADVLQGRLRAGPLAHFDR...GDNRAGHASRRRAEEGLLPCARAEP | MTVLAGQPRGSTVAPWR | 4457344 | 4457551 | + | 6 |
| 35 | MASPVIASSIKVGEWSRAKTALQDICN | VKDAFHSGAEPLFASPGRCVGRPFAARL | 4457443 | 4457275 | - | 6 |
| 36 | MLCHLHVVPVTVEVLGLLFGAISVFG | WVSCTVTLRRAEAGRAIRIPRSRWMW | 4459068 | 4458909 | - | 7 |
| 37 | VMLCHLHVVP | VTVEVLGLLFGAISVFGWVSCTVTLRRAEAGRAIRIPRSRWMW | 4459071 | 4458909 | - | 8 |
| 38 | LAVMLCHLHVVP | VTVEVLGLLFGAISVFGWVSCTVTLRRAEAGRAIRIPRSRWMW | 4459077 | 4458909 | - | 8 |
| 39 | VRPSLHVPAGLADLGVPQDLHDRGHVD | PEFIEDRAGGVTASVVQTTVVDSGFGQ | 4462710 | 4462545 | - | 9 |
| 40 | LLLSDAATALSACRAAVCGLNSGLN...ARLRARHEGVSHAAHGVLREVRPSL | HVPAGLADLGVPQDLHDRGHVDPEFIEDRAGGVTASVVQTTVVDSGFGQ | 4462857 | 4462545 | - | 6 |
| 41 | LERLARDSRPETTLRRIEAIAAC | RDALDRNVAPLLAVEAMTMALRAG | 4464933 | 4464789 | - | 6 |
| 42 | VLERLARDSRPETTLRRIEAIAAC | RDALDRNVAPLLAVEAMTMALRAG | 4464936 | 4464789 | - | 6 |

### Results for PIG43643.1 [Streptomyces sp. 61] back to top

previous - next

#### Architecture

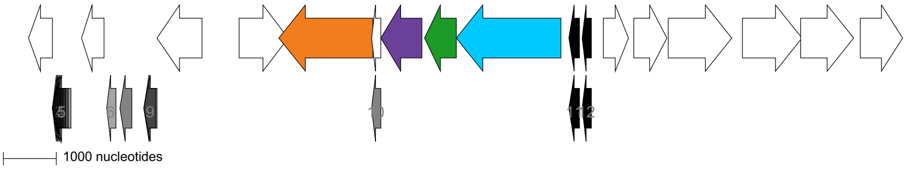

Link to nucleotide sequence

| Accession | start | end | direction | length (aa) | Pfam/HMM | name | description | E-value |
|---|---|---|---|---|---|---|---|---|
| PIG43635.1 | 2578511 | 2578076 | - | 144 | NO PFAM MATCH | - | - | - |
| PIG43636.1 | 2579499 | 2579055 | - | 147 | NO PFAM MATCH | - | - | - |
| PIG43637.1 | 2581341 | 2580489 | - | 283 | PF01551 | Peptidase_M23 | Peptidase family M23 | 6.00E-18 |
| PIG43638.1 | 2582051 | 2582969 | + | 305 | NO PFAM MATCH | - | - | - |

| | | | | | | | | |
|---|---|---|---|---|---|---|---|---|
| PIG43639.1 | 2584563 | 2582796 | - | 588 | TIGR02204<br>TIGR02203<br>TIGR03796<br>TIGR01846<br>TIGR00958 | TIGR02204<br>TIGR02203<br>TIGR03796<br>TIGR01846<br>TIGR00958 | MsbA_rel: ABC transporter, permease/ATP-binding protein<br>MsbA_lipidA: lipid A export permease/ATP-binding protein MsbA<br>NHLM_micro_ABC1: NHLM bacteriocin system ABC transporter, peptidase/ATP-binding protein<br>type_I_sec_HlyB: type I secretion system ATPase<br>3a01208: antigen peptide transporter 2 | 3.90E-104<br>1.10E-100<br>1.90E-85<br>1.50E-83<br>7.90E-83 |
| PIG43640.1 | 2584736 | 2584559 | - | 58 | NO PFAM MATCH | - | - | - |
| PIG43641.1 | 2585511 | 2584737 | - | 257 | PF13649<br>PF08241<br>PF08242<br>PF13847<br>PF13489 | Methyltransf_25<br>Methyltransf_11<br>Methyltransf_12<br>Methyltransf_31<br>Methyltransf_23 | Methyltransferase domain<br>Methyltransferase domain<br>Methyltransferase domain<br>Methyltransferase domain<br>Methyltransferase domain | 6.60E-14<br>2.30E-09<br>8.70E-09<br>3.30E-06<br>5.10E-06 |
| PIG43642.1 | 2586149 | 2585561 | - | 195 | LinL | LinL | LinL | 2.20E-70 |
| PIG43643.1 | 2588122 | 2586145 | - | 658 | LinH<br>LinG<br>LinE<br>PF12697<br>PF12146 | LinH<br>LinG<br>LinE<br>Abhydrolase_6<br>Hydrolase_4 | LinH (Fusion of LinE/LinG)<br>LinG Homolog<br>LinE Homolog<br>Alpha/beta hydrolase family<br>Serine aminopeptidase, S33 | 4.20E-251<br>5.70E-117<br>7.80E-101<br>1.60E-16<br>1.80E-10 |
| PIG43644.1 | 2588470 | 2588269 | - | 66 | LinA | LinA | LinA (linaridin precursor) | 2.70E-11 |
| PIG43645.1 | 2588719 | 2588524 | - | 64 | LinA | LinA | LinA (linaridin precursor) | 5.00E-12 |
| PIG43646.1 | 2588923 | 2589409 | + | 161 | PF13560 | HTH_31 | Helix-turn-helix domain | 3.50E-04 |
| PIG43647.1 | 2589511 | 2590141 | + | 209 | PF06114 | Peptidase_M78 | IrrE N-terminal-like domain | 3.10E-05 |
| PIG43648.1 | 2590153 | 2591359 | + | 401 | NO PFAM MATCH | - | - | - |
| PIG43649.1 | 2591566 | 2592676 | + | 369 | PF08028<br>TIGR04022<br>PF02770<br>PF00441 | Acyl-CoA_dh_2<br>TIGR04022<br>Acyl-CoA_dh_M<br>Acyl-CoA_dh_1 | Acyl-CoA dehydrogenase, C-terminal domain<br>sulfur_SfnB: sulfur acquisition oxidoreductase, SfnB family<br>Acyl-CoA dehydrogenase, middle domain<br>Acyl-CoA dehydrogenase, C-terminal domain | 1.30E-24<br>4.40E-23<br>1.10E-05<br>9.10E-05 |
| PIG43650.1 | 2592672 | 2593659 | + | 328 | PF00248<br>TIGR01293 | Aldo_ket_red<br>TIGR01293 | Aldo/keto reductase family<br>Kv_beta: voltage-dependent potassium channel beta subunit | 2.90E-74<br>2.50E-27 |
| PIG43651.1 | 2593793 | 2594603 | + | 269 | PF03704<br>PF00486 | BTAD<br>Trans_reg_C | Bacterial transcriptional activator domain<br>Transcriptional regulatory protein, C terminal | 5.20E-41<br>9.30E-11 |

| index | leader | core | start | end | dir | score |
|---|---|---|---|---|---|---|
| 1 | VLVSNSTATVPTATQQHTIQVSQTPSVSVLMSSPAI | LTPDSGPIILKEPR | 2578660 | 2578507 | - | 6 |
| 2 | LADVCDALEYELTGGLHPAGPRVLV...TATQQHTIQVSQTPSVSVLMSSPAI | LTPDSGPIILKEPR | 2578726 | 2578507 | - | 7 |
| 3 | MVQKDEAGGYETARLDRLADVCDAL...TATQQHTIQVSQTPSVSVLMSSPAI | LTPDSGPIILKEPR | 2578777 | 2578507 | - | 7 |
| 4 | LAARLEVSEQMVQKDEAGGYETARL...TATQQHTIQVSQTPSVSVLMSSPAI | LTPDSGPIILKEPR | 2578807 | 2578507 | - | 7 |
| 5 | LPNGLIKARIARKITQAELAARLEV...TATQQHTIQVSQTPSVSVLMSSPAI | LTPDSGPIILKEPR | 2578861 | 2578507 | - | 6 |
| 6 | LLQVREPLGSANRRRQGQGLSPSCHV | GTDLASDERSPNPIDWAIWLRACHYAE | 2579706 | 2579544 | - | 6 |
| 7 | VSHFGFGVLLLPPLGLVLRPLALMILASRAVHDRRPRT | RTVLGRRRATPGLSHSATTCLLPVDDGTRG | 2580006 | 2579799 | - | 7 |
| 8 | VHGAGKGCGKRRARFTENRSGGNCRRSRCGRCSPRS | PTPAADRGLKNGSAEETRRVPLPRPPTYGCTYAKTPSQLRR | 2580466 | 2580232 | - | 7 |
| 9 | VTPDLPGPAVHGAGKGCGKRRARFTENRSGGNCRRSRCGRCSPRS | PTPAADRGLKNGSAEETRRVPLPRPPTYGCTYAKTPSQLRR | 2580493 | 2580232 | - | 7 |
| 10 | MSTQRAVWFVTALAVAVPVM | AVMFREDGRFTSQSWTKGLIFGTAVAVVAAIAAGRARQ | 2584736 | 2584559 | - | 7 |
| 11 | MNSITESVRVIAGATLSEDSGERTTRMMPVA | FTPAVLVTTPYAAGGLAAGAAAVGAYEAGRAAAKG | 2588470 | 2588269 | - | 29 |
| 12 | MAKIAAFMNQQDMGPIAESGLSVGGTPVM | ATPAVVAVPASVKITAGIVAFTIPVGAYVTGRVVN | 2588719 | 2588524 | - | 30 |

### Results for WP_132931361.1 [Streptomyces sp. 8K308] back to top

previous - next

#### Architecture

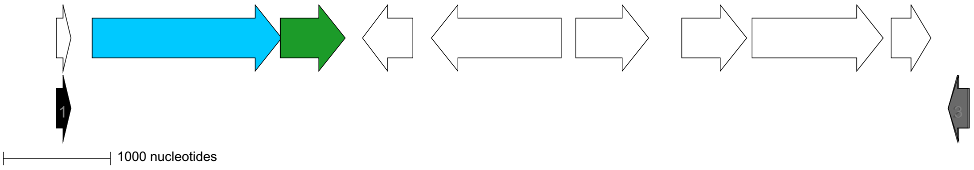

Link to nucleotide sequence

| Accession | start | end | direction | length (aa) | Pfam/HMM | name | description | E-value |
|---|---|---|---|---|---|---|---|---|
| WP_165953785.1 | 205 | 343 | + | 45 | NO PFAM MATCH | - | - | - |
| WP_132931361.1 | 535 | 2305 | + | 589 | LinH<br>LinG<br>LinE<br>PF12697<br>TIGR02427 | LinH<br>LinG<br>LinE<br>Abhydrolase_6<br>TIGR02427 | LinH (Fusion of LinE/LinG)<br>LinG Homolog<br>LinE Homolog<br>Alpha/beta hydrolase family<br>protocat_pcaD: 3-oxoadipate enol-lactonase | 2.40E-209<br>4.20E-96<br>3.80E-91<br>3.40E-14<br>2.00E-07 |
| WP_132931362.1 | 2301 | 2907 | + | 201 | LinL | LinL | LinL | 1.70E-52 |

| | | | | | | | | |
|---|---|---|---|---|---|---|---|---|
| WP_132931363.1 | 3531 | 3060 | - | 156 | PF01814<br>TIGR02481<br>TIGR03652 | Hemerythrin<br>TIGR02481<br>TIGR03652 | Hemerythrin HHE cation binding domain<br>hemeryth_dom: hemerythrin-like metal-binding domain<br>FeS_repair_RIC: iron-sulfur cluster repair di-iron protein | 3.20E-18<br>1.10E-05<br>4.90E-04 |
| WP_132931364.1 | 4913 | 3704 | - | 402 | PF12715<br>PF01738<br>PF12697<br>PF00326<br>PF02129 | Abhydrolase_7<br>DLH<br>Abhydrolase_6<br>Peptidase_S9<br>Peptidase_S15 | Abhydrolase family<br>Dienelactone hydrolase family<br>Alpha/beta hydrolase family<br>Prolyl oligopeptidase family<br>X-Pro dipeptidyl-peptidase (S15 family) | 1.00E-10<br>4.60E-09<br>1.70E-06<br>1.60E-05<br>7.20E-05 |
| WP_019993726.1 | 5057 | 5739 | + | 227 | INFERRED GENE | - | - | - |
| WP_005474110.1 | 6044 | 6654 | + | 203 | INFERRED GENE | - | - | - |
| WP_076611170.1 | 6693 | 7922 | + | 409 | INFERRED GENE | - | - | - |
| WP_008744193.1 | 7992 | 8367 | + | 125 | INFERRED GENE | - | - | - |

| index | leader | core | start | end | dir | score |
|---|---|---|---|---|---|---|
| 1 | MDDILSHFDIQDIDIDEGHVHVEQQPVA | CVVAVTVTVTTVAVTIG | 205 | 343 | + | 15 |
| 2 | MLNRPLGGGCSTGESGHPGRRVLRDDERHRR | YRGHFAFARTELSTFMRTPTGASTRAPCRAH | 8718 | 8529 | - | 6 |
| 3 | LMLNRPLGGGCSTGESGHPGRRVLRDDERHR | RYRGHFAFARTELSTFMRTPTGASTRAPCRAH | 8721 | 8529 | - | 6 |

### Results for WP_159062470.1 [Streptomyces sp. 96-12] back to top

previous - next

#### Architecture

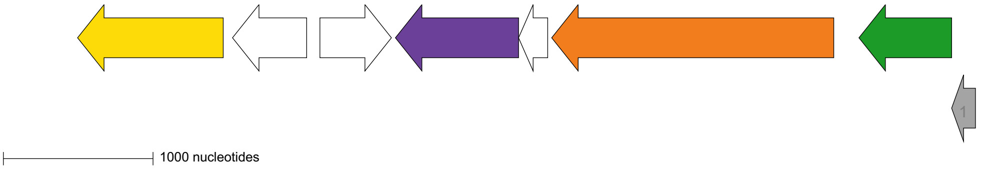

Link to nucleotide sequence

| Accession | start | end | direction | length (aa) | Pfam/HMM | name | description | E-value |
|---|---|---|---|---|---|---|---|---|
| WP_060885824.1 | 1023 | 51 | - | 323 | PF00561<br>PF12697<br>TIGR03056<br>TIGR02427<br>TIGR03343 | Abhydrolase_1<br>Abhydrolase_6<br>TIGR03056<br>TIGR02427<br>TIGR03343 | alpha/beta hydrolase fold<br>Alpha/beta hydrolase family<br>bchO_mg_che_rel: putative magnesium chelatase accessory protein<br>protocat_pcaD: 3-oxoadipate enol-lactonase<br>biphenyl_bphD: 2-hydroxy-6-oxo-6-phenylhexa-2,4-dienoate hydrolase | 4.10E-26<br>3.00E-19<br>2.00E-17<br>5.60E-16<br>8.80E-16 |
| WP_045555733.1 | 1582 | 1087 | - | 164 | TIGR00369<br>PF03061<br>TIGR02286<br>PF14539<br>PF13622 | TIGR00369<br>4HBT<br>TIGR02286<br>DUF4442<br>4HBT_3 | unchar_dom_1: uncharacterized domain 1<br>Thioesterase superfamily<br>PaaD: phenylacetic acid degradation protein PaaD<br>Domain of unknown function (DUF4442)<br>Thioesterase-like superfamily | 5.50E-27<br>2.50E-16<br>1.90E-09<br>2.20E-08<br>5.90E-05 |
| WP_060885825.1 | 1671 | 2148 | + | 158 | PF01638 | HxlR | HxlR-like helix-turn-helix | 9.70E-20 |
| WP_045555731.1 | 3002 | 2174 | - | 275 | PF13649<br>PF08241<br>PF08242<br>PF13489<br>TIGR01983 | Methyltransf_25<br>Methyltransf_11<br>Methyltransf_12<br>Methyltransf_23<br>TIGR01983 | Methyltransferase domain<br>Methyltransferase domain<br>Methyltransferase domain<br>Methyltransferase domain<br>UbiG: 3-demethylubiquinone-9 3-O-methyltransferase | 1.60E-15<br>1.30E-12<br>4.30E-07<br>5.20E-07<br>3.10E-05 |
| WP_045555730.1 | 3190 | 2998 | - | 63 | NO PFAM MATCH | - | - | - |
| WP_060885826.1 | 5108 | 3215 | - | 630 | TIGR02204<br>TIGR02203<br>TIGR00958<br>TIGR03796<br>TIGR03375 | TIGR02204<br>TIGR02203<br>TIGR00958<br>TIGR03796<br>TIGR03375 | MsbA_rel: ABC transporter, permease/ATP-binding protein<br>MsbA_lipidA: lipid A export permease/ATP-binding protein MsbA<br>3a01208: antigen peptide transporter 2<br>NHLM_micro_ABC1: NHLM bacteriocin system ABC transporter, peptidase/ATP-binding protein<br>type_I_sec_LssB: type I secretion system ATPase | 1.00E-106<br>1.60E-104<br>1.40E-92<br>1.20E-86<br>1.70E-81 |
| WP_060885827.1 | 5892 | 5271 | - | 206 | LinL | LinL | LinL | 4.50E-69 |

| index | leader | core | start | end | dir | score |
|---|---|---|---|---|---|---|
| 1 | VVVEGAGHDTLLINSRYAHQVTDRVL | AFFAGLDERPRRTRPGRSVTLEKEASR | 6050 | 5888 | - | 6 |

### Results for AJC56865.1 [Streptomyces sp. 769] back to top

previous - next

#### Architecture

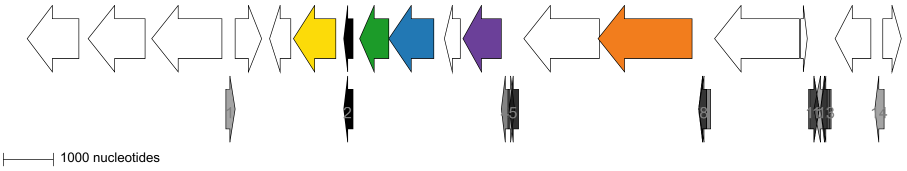

Link to nucleotide sequence

| Accession | start | end | direction | length (aa) | Pfam/HMM | name | description | E-value |
|---|---|---|---|---|---|---|---|---|
| AJC56857.1 | 4629729 | 4628709 | - | 339 | TIGR02432<br>PF01171<br>PF09179 | TIGR02432<br>ATP_bind_3<br>TilS | lysidine_TilS_N: tRNA(Ile)-lysidine synthetase<br>PP-loop family<br>TilS substrate binding domain | 2.20E-43<br>2.10E-38<br>3.10E-12 |
| AJC56858.1 | 4631057 | 4629911 | - | 381 | TIGR03883<br>TIGR03624<br>PF10103 | TIGR03883<br>TIGR03624<br>Zincin_2 | DUF2342_F420: uncharacterized protein, coenzyme F420 biosynthesis associated<br>TIGR03624: putative hydrolase<br>Zincin-like metallopeptidase | 9.50E-141<br>2.50E-125<br>8.50E-121 |
| AJC56859.1 | 4632590 | 4631171 | - | 472 | PF02113<br>TIGR00666 | Peptidase_S13<br>TIGR00666 | D-Ala-D-Ala carboxypeptidase 3 (S13) family<br>PBP4: D-alanyl-D-alanine carboxypeptidase/D-alanyl-D-alanine-endopeptidase | 2.70E-76<br>5.90E-58 |
| AJC56860.1 | 4632850 | 4633366 | + | 171 | PF00719 | Pyrophosphatase | Inorganic pyrophosphatase | 4.50E-53 |
| AJC56861.1 | 4633958 | 4633532 | - | 141 | NO PFAM MATCH | - | - | - |
| AJC56862.1 | 4634872 | 4634011 | - | 286 | LinG<br>LinH<br>TIGR04033 | LinG<br>LinH<br>TIGR04033 | LinG Homolog<br>LinH (Fusion of LinE/LinG)<br>export_SdpB: antimicrobial peptide system protein, SdpB family | 4.10E-108<br>1.80E-76<br>7.60E-05 |
| AJC56863.1 | 4635214 | 4635016 | - | 65 | LinA | LinA | LinA (linaridin precursor) | 1.20E-12 |
| AJC56864.1 | 4635924 | 4635342 | - | 193 | LinL | LinL | LinL | 1.50E-57 |
| AJC56865.1 | 4636826 | 4635920 | - | 301 | LinE<br>LinH<br>PF12697<br>TIGR02427<br>PF00561 | LinE<br>LinH<br>Abhydrolase_6<br>TIGR02427<br>Abhydrolase_1 | LinE Homolog<br>LinH (Fusion of LinE/LinG)<br>Alpha/beta hydrolase family<br>protocat_pcaD: 3-oxoadipate enol-lactonase<br>alpha/beta hydrolase fold | 2.10E-97<br>1.90E-51<br>3.30E-15<br>2.60E-11<br>3.40E-09 |
| AJC56866.1 | 4637345 | 4637015 | - | 109 | PF01527<br>PF13518<br>PF13384<br>PF05344<br>PF13551 | HTH_Tnp_1<br>HTH_28<br>HTH_23<br>DUF746<br>HTH_29 | Transposase<br>Helix-turn-helix domain<br>Homeodomain-like domain<br>Domain of Unknown Function (DUF746)<br>Winged helix-turn helix | 6.90E-08<br>1.90E-07<br>4.60E-06<br>9.60E-06<br>3.30E-05 |
| AJC56867.1 | 4638160 | 4637401 | - | 252 | PF13649<br>PF08241<br>PF13489<br>PF13847<br>PF08242 | Methyltransf_25<br>Methyltransf_11<br>Methyltransf_23<br>Methyltransf_31<br>Methyltransf_12 | Methyltransferase domain<br>Methyltransferase domain<br>Methyltransferase domain<br>Methyltransferase domain<br>Methyltransferase domain | 3.10E-16<br>4.70E-13<br>8.00E-10<br>6.90E-09<br>7.20E-09 |
| AJC56868.1 | 4640109 | 4638624 | - | 494 | TIGR03467<br>TIGR02734<br>TIGR00292<br>PF13450<br>TIGR03315 | TIGR03467<br>TIGR02734<br>TIGR00292<br>NAD_binding_8<br>TIGR03315 | HpnE: squalene-associated FAD-dependent desaturase<br>crtI_fam: phytoene desaturase<br>TIGR00292: thiazole biosynthesis enzyme<br>NAD(P)-binding Rossmann-like domain<br>Se_ygfK: putative selenate reductase, YgfK subunit | 1.90E-09<br>1.00E-08<br>2.20E-08<br>2.60E-08<br>1.10E-06 |
| AJC56869.1 | 4641983 | 4640105 | - | 625 | TIGR02204<br>TIGR02203<br>TIGR00958<br>TIGR03796<br>TIGR01846 | TIGR02204<br>TIGR02203<br>TIGR00958<br>TIGR03796<br>TIGR01846 | MsbA_rel: ABC transporter, permease/ATP-binding protein<br>MsbA_lipidA: lipid A export permease/ATP-binding protein MsbA<br>3a01208: antigen peptide transporter 2<br>NHLM_micro_ABC1: NHLM bacteriocin system ABC transporter, peptidase/ATP-binding protein<br>type_I_sec_HlyB: type I secretion system ATPase | 2.80E-98<br>3.90E-95<br>7.40E-89<br>1.60E-80<br>4.70E-78 |
| AJC56870.1 | 4644128 | 4642415 | - | 570 | PF06738<br>PF12821 | ThrE<br>ThrE_2 | Putative threonine/serine exporter<br>Threonine/Serine exporter, ThrE | 5.60E-73<br>1.60E-19 |
| AJC56871.1 | 4644143 | 4644278 | + | 44 | NO PFAM MATCH | - | - | - |
| AJC56872.1 | 4645533 | 4644822 | - | 236 | PF09335 | SNARE_assoc | SNARE associated Golgi protein | 2.00E-24 |
| AJC56873.1 | 4645784 | 4646162 | + | 125 | PF01906<br>TIGR03884 | YbjQ_1<br>TIGR03884 | Putative heavy-metal-binding<br>sel_bind_Methan: selenium-binding protein | 2.10E-33<br>7.50E-04 |

| index | leader | core | start | end | dir | score |
|---|---|---|---|---|---|---|
| 1 | MRGALCGPCRGACGTPRPACRHPRGTLRRLPCDLPCSGLGHFGPAPF | ATTHLRGGHLITRRMC | 4632665 | 4632857 | + | 6 |
| 2 | MSVVAEFANTQLVDVKPGRIGSDAAPTM | MTPLATVATPEAVSAGIAVTSATAAALNMVTHNVQHG | 4635214 | 4635016 | - | 34 |
| 3 | LVRPAAPVDPVETPQRSVGNFRLCR | VSPSYTSRWAPVNTCDTTRRQSFRAG | 4638285 | 4638441 | + | 6 |
| 4 | LRRFHWIDRRCWSDQPCA | TTAVSPAVVLAYDVLPCSGSSASRRPPRDRMWGQSEGVR | 4638330 | 4638156 | - | 7 |
| 5 | VVGVIRARGAGQSRLASDGNHPARKLCL | LVVSHVFTGAHRDVYDGLTRHNLKFPTLR | 4638500 | 4638326 | - | 9 |
| 6 | VFLSGTAPSGQQTRLLTPAGPAGHSTG | QGVPANQLTRNLTPEGVLVSAILTTKL | 4642272 | 4642107 | - | 6 |
| 7 | VVFLSGTAPSGQQTRLLTPAGPAGHST | GQGVPANQLTRNLTPEGVLVSAILTTKL | 4642275 | 4642107 | - | 6 |
| 8 | VSGESIAPGAPCVGGFVLLTREGTVVFLS | GTAPSGQQTRLLTPAGPAGHSTGQGVPANQLTRNLTPEGVLVSAILTTKL | 4642347 | 4642107 | - | 7 |
| 9 | LCARGAGPPIGGVFRCRSAARGCRVRWRVCGVGGVRTAALYV | RTPPTPQTHPLPWVGGKDGEWEPTQVVP | 4644287 | 4644500 | + | 7 |
| 10 | VFRCRSAARGCRVRWRVCGVGGVRTAALYV | RTPPTPQTHPLPWVGGKDGEWEPTQVVP | 4644323 | 4644500 | + | 7 |
| 11 | VPGLWGGGCADCCALRP | HIPHPTDPSPPVGGWVSGRRGTPGW...VGSHSPSFPPTHGRGWVCGVGGVRT | 4644661 | 4644406 | - | 6 |
| 12 | VPPDGGSFRFRCAGPIRCAGLVGRGAGSVG | WGMCGLLRFTSAHPPPHRPVPSRWWVGFRP | 4644734 | 4644551 | - | 6 |
| 13 | LCARAAVPPDGGSFRFRCAGPIRCAGLVGRGAG | SVGWGMCGLLRFTSAHPPPHRPVPSRWWVGFRP | 4644752 | 4644551 | - | 7 |
| 14 | MPMIRTSCPCDQDHPGRPERPALGGGRVG | VGGRRRGGAGRPGRPWPGVLLEPRDVIVR | 4645801 | 4645624 | - | 6 |

### Results for WP_106438501.1 [Streptomyces sp. AA0539] back to top

previous - next

#### Architecture

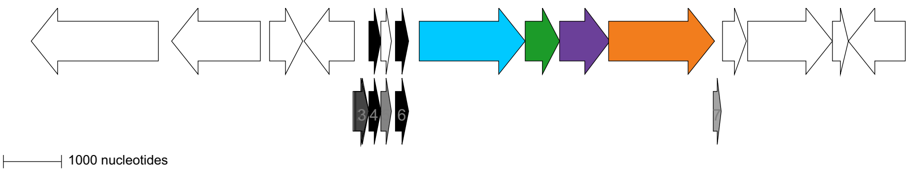

Link to nucleotide sequence

| Accession | start | end | direction | length (aa) | Pfam/HMM | name | description | E-value |
|---|---|---|---|---|---|---|---|---|
| WP_019435946.1 | 2189 | 0 | - | 729 | TIGR00940<br>TIGR01974<br>PF00361<br>TIGR01972<br>TIGR01770 | TIGR00940<br>TIGR01974<br>Proton_antipo_M<br>TIGR01972<br>TIGR01770 | 2a6301s01: monovalent cation:proton antiporter<br>NDH_I_L: proton-translocating NADH-quinone oxidoreductase, chain L<br>Proton-conducting membrane transporter<br>NDH_I_M: proton-translocating NADH-quinone oxidoreductase, chain M<br>NDH_I_N: proton-translocating NADH-quinone oxidoreductase, chain N | 1.30E-134<br>2.00E-73<br>2.30E-56<br>6.30E-41<br>5.00E-33 |
| WP_019435947.1 | 3930 | 2409 | - | 506 | PF11066<br>PF13460<br>TIGR03466<br>PF05368<br>PF01370 | DUF2867<br>NAD_binding_10<br>TIGR03466<br>NmrA<br>Epimerase | Protein of unknown function (DUF2867)<br>NAD(P)H-binding<br>HpnA: hopanoid-associated sugar epimerase<br>NmrA-like family<br>NAD dependent epimerase/dehydratase family | 3.90E-42<br>5.10E-21<br>1.40E-15<br>8.80E-13<br>2.00E-09 |
| WP_019435948.1 | 4094 | 4658 | + | 187 | PF01872 | RibD_C | RibD C-terminal domain | 4.20E-13 |
| WP_019435949.1 | 5553 | 4680 | - | 290 | PF12146<br>PF12697<br>TIGR01738<br>TIGR03695<br>PF08386 | Hydrolase_4<br>Abhydrolase_6<br>TIGR01738<br>TIGR03695<br>Abhydrolase_4 | Serine aminopeptidase, S33<br>Alpha/beta hydrolase family<br>bioH: pimelyl-[acyl-carrier protein] methyl ester esterase<br>menH_SHCHC: 2-succinyl-6-hydroxy-2,4-cyclohexadiene-1-carboxylate synthase<br>TAP-like protein | 5.10E-10<br>6.40E-10<br>7.40E-10<br>5.20E-07<br>1.40E-05 |
| WP_019435950.1 | 5810 | 6014 | + | 67 | LinA | LinA | LinA (linaridin precursor) | 1.70E-13 |
| WP_019435951.1 | 6010 | 6187 | + | 58 | NO PFAM MATCH | - | - | - |
| WP_019435952.1 | 6263 | 6482 | + | 72 | LinA | LinA | LinA (linaridin precursor) | 2.20E-13 |
| WP_106438501.1 | 6665 | 8498 | + | 610 | LinH<br>LinG<br>LinE<br>PF12697<br>TIGR02427 | LinH<br>LinG<br>LinE<br>Abhydrolase_6<br>TIGR02427 | LinH (Fusion of LinE/LinG)<br>LinG Homolog<br>LinE Homolog<br>Alpha/beta hydrolase family<br>protocat_pcaD: 3-oxoadipate enol-lactonase | 8.20E-242<br>2.80E-109<br>2.30E-97<br>2.60E-16<br>1.50E-09 |
| WP_019435954.1 | 8494 | 9094 | + | 199 | LinL | LinL | LinL | 2.00E-63 |
| WP_019435955.1 | 9090 | 9945 | + | 284 | PF13649<br>PF08241<br>PF13489<br>PF08242<br>TIGR01983 | Methyltransf_25<br>Methyltransf_11<br>Methyltransf_23<br>Methyltransf_12<br>TIGR01983 | Methyltransferase domain<br>Methyltransferase domain<br>Methyltransferase domain<br>Methyltransferase domain<br>UbiG: 3-demethylubiquinone-9 3-O-methyltransferase | 1.80E-16<br>2.40E-11<br>1.60E-07<br>1.90E-07<br>1.10E-06 |
| WP_019435956.1 | 9922 | 11746 | + | 607 | TIGR02204<br>TIGR02203<br>TIGR00958<br>TIGR03796<br>TIGR03375 | TIGR02204<br>TIGR02203<br>TIGR00958<br>TIGR03796<br>TIGR03375 | MsbA_rel: ABC transporter, permease/ATP-binding protein<br>MsbA_lipidA: lipid A export permease/ATP-binding protein MsbA<br>3a01208: antigen peptide transporter 2<br>NHLM_micro_ABC1: NHLM bacteriocin system ABC transporter, peptidase/ATP-binding protein<br>type_I_sec_LssB: type I secretion system ATPase | 5.40E-105<br>1.90E-100<br>5.90E-89<br>1.20E-87<br>1.40E-82 |
| WP_026047810.1 | 11885 | 12296 | + | 136 | NO PFAM MATCH | - | - | - |
| WP_026047811.1 | 12316 | 13777 | + | 486 | TIGR03467<br>TIGR01317<br>PF13450<br>TIGR02734<br>TIGR02733 | TIGR03467<br>TIGR01317<br>NAD_binding_8<br>TIGR02734<br>TIGR02733 | HpnE: squalene-associated FAD-dependent desaturase<br>GOGAT_sm_gam: glutamate synthase, NADH/NADPH, small subunit<br>NAD(P)-binding Rossmann-like domain<br>crtI_fam: phytoene desaturase<br>desat_CrtD: C-3',4' desaturase CrtD | 4.30E-10<br>1.10E-09<br>1.50E-09<br>2.00E-09<br>5.70E-09 |
| WP_019435958.1 | 13773 | 14046 | + | 90 | NO PFAM MATCH | - | - | - |
| WP_019435959.1 | 15041 | 14057 | - | 327 | NO PFAM MATCH | - | - | - |

| index | leader | core | start | end | dir | score |
|---|---|---|---|---|---|---|
| 1 | VPRRSDAWMPSPVSPWCPGRDSSTTVRSVFCPGR | TPPSHRRAARAPPERRPAVHDAAER...ATAAPSPPPAGSGGFCPAAERPPPG | 5531 | 5798 | + | 6 |
| 2 | MPSPVSPWCPGRDSSTTVRSVFCPGR | TPPSHRRAARAPPERRPAVHDAAER...ATAAPSPPPAGSGGFCPAAERPPPG | 5555 | 5798 | + | 6 |
| 3 | VSPWCPGRDSSTTVRSVFCPGR | TPPSHRRAARAPPERRPAVHDAAER...ATAAPSPPPAGSGGFCPAAERPPPG | 5567 | 5798 | + | 6 |
| 4 | MARSTPFAQRAVSGPLLEDSPGDIGTAASPVL | ATPAAAAAFAAGATIVVGAYVVGRMVGGETEPVIQ | 5810 | 6014 | + | 31 |
| 5 | MTTADRLPAALRHLPELTE | DAPGLASGTSEAVLCLVHSAVQVSVCVTSAGDQDSKLPL | 6010 | 6187 | + | 7 |
| 6 | MSRTTAYLNGCAALADLSVLDSAGRRSPAQPVL | ATPVTIVAFAGGVGAGAAIVAAFEAGRQAAGGSMEPIIT | 6263 | 6482 | + | 29 |
| 7 | MLRRGCSELPAHPLSPGRRSWQIRHVAPTG | TPPITRITHSDTAAEAGRG | 11723 | 11873 | + | 6 |

### Results for WP_111339449.1 [Streptomyces sp. AC1-42T] back to top

previous - next

#### Architecture

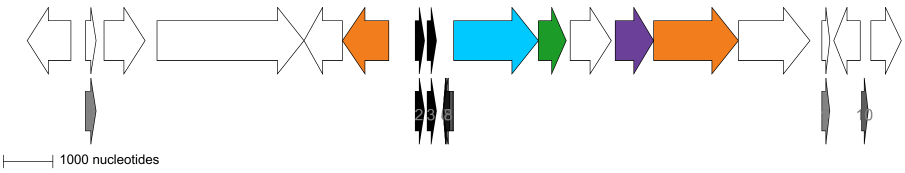

Link to nucleotide sequence

| Accession | start | end | direction | length (aa) | Pfam/HMM | name | description | E-value |
|---|---|---|---|---|---|---|---|---|
| WP_111337149.1 | 2567281 | 2566402 | - | 292 | PF02621 | VitK2_biosynth | Menaquinone biosynthesis | 6.50E-71 |
| WP_003967102.1 | 2567579 | 2567783 | + | 67 | PF00313<br>TIGR02381<br>PF08206 | CSD<br>TIGR02381<br>OB_RNB | 'Cold-shock' DNA-binding domain<br>cspD: cold shock domain protein CspD<br>Ribonuclease B OB domain | 2.00E-31<br>1.20E-26<br>3.90E-04 |
| WP_111339451.1 | 2567937 | 2568771 | + | 277 | PF13411<br>PF06445<br>TIGR02043<br>PF00376<br>TIGR02044 | MerR_1<br>GyrI-like<br>TIGR02043<br>MerR<br>TIGR02044 | MerR HTH family regulatory protein<br>GyrI-like small molecule binding domain<br>ZntR: Zn(II)-responsive transcriptional regulator<br>MerR family regulatory protein<br>CueR: Cu(I)-responsive transcriptional regulator | 2.00E-17<br>4.10E-16<br>1.30E-11<br>2.70E-11<br>5.40E-11 |
| WP_111337147.1 | 2569014 | 2571981 | + | 988 | PF03704<br>PF13424<br>PF00931<br>PF07719<br>PF13181 | BTAD<br>TPR_12<br>NB-ARC<br>TPR_2<br>TPR_8 | Bacterial transcriptional activator domain<br>Tetratricopeptide repeat<br>NB-ARC domain<br>Tetratricopeptide repeat<br>Tetratricopeptide repeat | 1.70E-40<br>1.80E-19<br>2.90E-15<br>4.30E-13<br>7.70E-11 |
| WP_111337145.1 | 2572774 | 2572000 | - | 257 | PF01061<br>PF12698<br>TIGR03062 | ABC2_membrane<br>ABC2_membrane_3<br>TIGR03062 | ABC-2 type transporter<br>ABC-2 family transporter protein<br>pip_yhgE_Cterm: YhgE/Pip C-terminal domain | 1.20E-13<br>4.60E-04<br>8.40E-04 |
| WP_111337143.1 | 2573718 | 2572770 | - | 315 | TIGR01188<br>TIGR03864<br>TIGR01288<br>TIGR03740<br>TIGR02673 | TIGR01188<br>TIGR03864<br>TIGR01288<br>TIGR03740<br>TIGR02673 | drrA: daunorubicin resistance ABC transporter, ATP-binding protein<br>PQQ_ABC_ATP: ABC transporter, ATP-binding subunit, PQQ-dependent alcohol dehydrogenase system<br>nodI: nodulation ABC transporter NodI<br>galliderm_ABC: lantibiotic protection ABC transporter, ATP-binding subunit<br>FtsE: cell division ATP-binding protein FtsE | 2.20E-58<br>4.40E-41<br>1.60E-40<br>3.60E-34<br>9.10E-32 |
| WP_111337141.1 | 2574234 | 2574426 | + | 63 | LinA | LinA | LinA (linaridin precursor) | 1.60E-14 |
| WP_111415608.1 | 2574472 | 2574676 | + | 67 | LinA | LinA | LinA (linaridin precursor) | 5.70E-10 |
| WP_111339449.1 | 2575013 | 2576723 | + | 569 | LinH<br>LinE<br>LinG<br>PF12697<br>PF12146 | LinH<br>LinE<br>LinG<br>Abhydrolase_6<br>Hydrolase_4 | LinH (Fusion of LinE/LinG)<br>LinE Homolog<br>LinG Homolog<br>Alpha/beta hydrolase family<br>Serine aminopeptidase, S33 | 2.60E-226<br>6.50E-99<br>2.10E-96<br>4.20E-17<br>1.40E-12 |
| WP_111337137.1 | 2576731 | 2577307 | + | 191 | LinL | LinL | LinL | 1.70E-64 |
| WP_111337135.1 | 2577385 | 2578210 | + | 274 | PF12680 | SnoaL_2 | SnoaL-like domain | 6.30E-08 |
| WP_111337133.1 | 2578292 | 2579054 | + | 253 | PF13649<br>PF08241<br>PF08242<br>PF13489<br>TIGR01983 | Methyltransf_25<br>Methyltransf_11<br>Methyltransf_12<br>Methyltransf_23<br>TIGR01983 | Methyltransferase domain<br>Methyltransferase domain<br>Methyltransferase domain<br>Methyltransferase domain<br>UbiG: 3-demethylubiquinone-9 3-O-methyltransferase | 1.10E-13<br>3.10E-11<br>7.50E-08<br>2.80E-07<br>1.40E-06 |
| WP_111337131.1 | 2579050 | 2580775 | + | 574 | TIGR02204<br>TIGR02203<br>TIGR00958<br>TIGR03375<br>TIGR03796 | TIGR02204<br>TIGR02203<br>TIGR00958<br>TIGR03375<br>TIGR03796 | MsbA_rel: ABC transporter, permease/ATP-binding protein<br>MsbA_lipidA: lipid A export permease/ATP-binding protein MsbA<br>3a01208: antigen peptide transporter 2<br>type_I_sec_LssB: type I secretion system ATPase<br>NHLM_micro_ABC1: NHLM bacteriocin system ABC transporter, peptidase/ATP-binding protein | 1.70E-108<br>1.00E-104<br>1.60E-93<br>5.20E-91<br>2.80E-86 |
| WP_111337129.1 | 2580767 | 2582210 | + | 480 | TIGR02734<br>PF13450<br>TIGR00292<br>TIGR03467<br>TIGR01317 | TIGR02734<br>NAD_binding_8<br>TIGR00292<br>TIGR03467<br>TIGR01317 | crtI_fam: phytoene desaturase<br>NAD(P)-binding Rossmann-like domain<br>TIGR00292: thiazole biosynthesis enzyme<br>HpnE: squalene-associated FAD-dependent desaturase<br>GOGAT_sm_gam: glutamate synthase, NADH/NADPH, small subunit | 2.00E-13<br>9.90E-10<br>5.60E-09<br>7.70E-09<br>3.40E-08 |
| WP_111337127.1 | 2582450 | 2582630 | + | 59 | NO PFAM MATCH | - | - | - |
| WP_111337125.1 | 2583244 | 2582710 | - | 177 | NO PFAM MATCH | - | - | - |
| WP_111337123.1 | 2583461 | 2584073 | + | 203 | NO PFAM MATCH | - | - | - |

| index | leader | core | start | end | dir | score |
|---|---|---|---|---|---|---|
| 1 | MATGTVKWFNAEKGFGFIAQDGGGPDVFVHYSA | INASGFRSLEENQVVNFDVTQGPKGPQAENVTPA | 2567579 | 2567783 | + | 7 |
| 2 | MAKIADFMNTQQLGPIEQGLTNTDGVPVM | ATPAVAVPVTVKITAGIVAFTIPVGAYVAGRVVN | 2574234 | 2574426 | + | 31 |
| 3 | MNRIIESVRTVTTPTLTGDAGISLKPMMPVT | FTPAILVTTPYAAGGLAAGAASVAAYEAGRAAGAGK | 2574472 | 2574676 | + | 27 |
| 4 | VSLSDIPMALLRLIGTGEN | TTEASFLNVLKIKPLDTTRV | 2574928 | 2574808 | - | 6 |
| 5 | VSAVSLSDIPMALLRLIGTGE | NTTEASFLNVLKIKPLDTTRV | 2574937 | 2574808 | - | 6 |
| 6 | VVSAVSLSDIPMALLRLIGTG | ENTTEASFLNVLKIKPLDTTRV | 2574940 | 2574808 | - | 6 |
| 7 | LRPPGSSCLRWCTYSREEESVVSAVSLSDIP | MALLRLIGTGENTTEASFLNVLKIKPLDTTRV | 2575000 | 2574808 | - | 7 |
| 8 | MISQSLRPPGSSCLRWCTYSREEESVVSAVSLSD | IPMALLRLIGTGENTTEASFLNVLKIKPLDTTRV | 2575015 | 2574808 | - | 6 |
| 9 | MGDKKKTDPIFKPADTHATGGDADLTTQD | THATVEPLDIHATGGEIKPLDTHATSEPAN | 2582450 | 2582630 | + | 7 |
| 10 | LSASLPFRCVAGPAAGRHGTVAHSM | ARGSGSPVMALSRSRLRLVAAVTRAA | 2583255 | 2583411 | + | 8 |

### Results for PZT77967.1 [Streptomyces sp. AC1-42W] back to top

previous - next

### Architecture

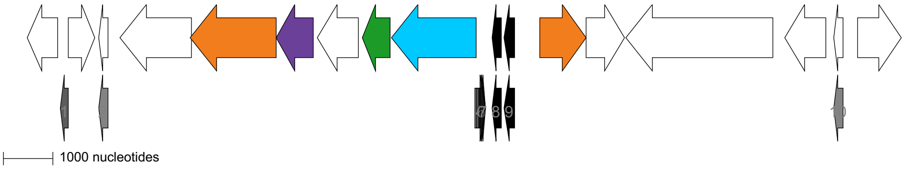

[Link to nucleotide sequence](#)

| Accession | start | end | direction | length (aa) | Pfam/HMM | name | description | E-value |
|---|---|---|---|---|---|---|---|---|
| PZT75929.1 | 750354 | 749742 | - | 203 | NO PFAM MATCH | - | - | - |
| PZT75930.1 | 750571 | 751105 | + | 177 | NO PFAM MATCH | - | - | - |
| PZT75931.1 | 751365 | 751185 | - | 59 | NO PFAM MATCH | - | - | - |
| PZT75932.1 | 753048 | 751605 | - | 480 | TIGR02734<br>PF13450<br>TIGR00292<br>TIGR03467<br>TIGR01317 | TIGR02734<br>NAD_binding_8<br>TIGR00292<br>TIGR03467<br>TIGR01317 | crtI_fam: phytoene desaturase<br>NAD(P)-binding Rossmann-like domain<br>TIGR00292: thiazole biosynthesis enzyme<br>HpnE: squalene-associated FAD-dependent desaturase<br>GOGAT_sm_gam: glutamate synthase, NADH/NADPH, small subunit | 2.00E-13<br>9.90E-10<br>5.60E-09<br>7.70E-09<br>3.40E-08 |
| PZT75933.1 | 754765 | 753040 | - | 574 | TIGR02204<br>TIGR02203<br>TIGR00958<br>TIGR03375<br>TIGR03796 | TIGR02204<br>TIGR02203<br>TIGR00958<br>TIGR03375<br>TIGR03796 | MsbA_rel: ABC transporter, permease/ATP-binding protein<br>MsbA_lipidA: lipid A export permease/ATP-binding protein MsbA<br>3a01208: antigen peptide transporter 2<br>type_I_sec_LssB: type I secretion system ATPase<br>NHLM_micro_ABC1: NHLM bacteriocin system ABC transporter, peptidase/ATP-binding protein | 1.70E-108<br>1.00E-104<br>1.60E-93<br>5.20E-91<br>2.80E-86 |
| PZT75934.1 | 755523 | 754761 | - | 253 | PF13649<br>PF08241<br>PF08242<br>PF13489<br>TIGR01983 | Methyltransf_25<br>Methyltransf_11<br>Methyltransf_12<br>Methyltransf_23<br>TIGR01983 | Methyltransferase domain<br>Methyltransferase domain<br>Methyltransferase domain<br>Methyltransferase domain<br>UbiG: 3-demethylubiquinone-9 3-O-methyltransferase | 1.10E-13<br>3.10E-11<br>7.50E-08<br>2.80E-07<br>1.40E-06 |
| PZT75935.1 | 756430 | 755605 | - | 274 | PF12680 | SnoaL_2 | SnoaL-like domain | 6.30E-08 |
| PZT75936.1 | 757084 | 756508 | - | 191 | LinL | LinL | LinL | 1.70E-64 |
| PZT77967.1 | 758802 | 757092 | - | 569 | LinH<br>LinE<br>LinG<br>PF12697<br>PF12146 | LinH<br>LinE<br>LinG<br>Abhydrolase_6<br>Hydrolase_4 | LinH (Fusion of LinE/LinG)<br>LinE Homolog<br>LinG Homolog<br>Alpha/beta hydrolase family<br>Serine aminopeptidase, S33 | 2.60E-226<br>6.50E-99<br>2.10E-96<br>4.20E-17<br>1.40E-12 |
| PZT75937.1 | 759322 | 759139 | - | 60 | LinA | LinA | LinA (linaridin precursor) | 7.50E-07 |
| PZT75938.1 | 759581 | 759389 | - | 63 | LinA | LinA | LinA (linaridin precursor) | 1.60E-14 |
| PZT75939.1 | 760097 | 761045 | + | 315 | TIGR01188<br>TIGR03864<br>TIGR01288<br>TIGR03740<br>TIGR02673 | TIGR01188<br>TIGR03864<br>TIGR01288<br>TIGR03740<br>TIGR02673 | drrA: daunorubicin resistance ABC transporter, ATP-binding protein<br>PQQ_ABC_ATP: ABC transporter, ATP-binding subunit, PQQ-dependent alcohol dehydrogenase system<br>nodI: nodulation ABC transporter NodI<br>galliderm_ABC: lantibiotic protection ABC transporter, ATP-binding subunit<br>FtsE: cell division ATP-binding protein FtsE | 2.20E-58<br>4.40E-41<br>1.60E-40<br>3.60E-34<br>9.10E-32 |
| PZT75940.1 | 761041 | 761815 | + | 257 | PF01061<br>PF12698<br>TIGR03062 | ABC2_membrane<br>ABC2_membrane_3<br>TIGR03062 | ABC-2 type transporter<br>ABC-2 family transporter protein<br>pip_yhgE_Cterm: YhgE/Pip C-terminal domain | 1.20E-13<br>4.60E-04<br>8.40E-04 |
| PZT75941.1 | 764801 | 761834 | - | 988 | PF03704<br>PF13424<br>PF00931<br>PF07719<br>PF13181 | BTAD<br>TPR_12<br>NB-ARC<br>TPR_2<br>TPR_8 | Bacterial transcriptional activator domain<br>Tetratricopeptide repeat<br>NB-ARC domain<br>Tetratricopeptide repeat<br>Tetratricopeptide repeat | 1.70E-40<br>1.80E-19<br>2.90E-15<br>4.30E-13<br>7.70E-11 |
| PZT75942.1 | 765878 | 765044 | - | 277 | PF13411<br>PF06445<br>TIGR02043<br>PF00376<br>TIGR02044 | MerR_1<br>GyrI-like<br>TIGR02043<br>MerR<br>TIGR02044 | MerR HTH family regulatory protein<br>GyrI-like small molecule binding domain<br>ZntR: Zn(II)-responsive transcriptional regulator<br>MerR family regulatory protein<br>CueR: Cu(I)-responsive transcriptional regulator | 2.00E-17<br>4.10E-16<br>1.30E-11<br>2.70E-11<br>5.40E-11 |
| PZT75943.1 | 766236 | 766032 | - | 67 | PF00313<br>TIGR02381<br>PF08206 | CSD<br>TIGR02381<br>OB_RNB | 'Cold-shock' DNA-binding domain<br>cspD: cold shock domain protein CspD<br>Ribonuclease B OB domain | 2.00E-31<br>1.20E-26<br>3.90E-04 |
| PZT75944.1 | 766534 | 767413 | + | 292 | PF02621 | VitK2_biosynth | Menaquinone biosynthesis | 6.50E-71 |

| index | leader | core | start | end | dir | score |
|---|---|---|---|---|---|---|
| 1 | LSASLPFRCVAGPAAGRHGTVAHSM | ARGSGSPVMALSRSRLRLVAAVTRAA | 750560 | 750404 | - | 8 |
| 2 | MGDKKKTDPIFKPADTHATGGDADLTTQD | THATVEPLDIHATGGEIKPLDTHATSEPAN | 751365 | 751185 | - | 7 |
| 3 | MISQSLRPPGSSCLRWCTYSREEESVVSAVSLSD | IPMALLRLIGTGENTTEASFLNVLKIKPLDTTRV | 758800 | 759007 | + | 6 |
| 4 | LRPPGSSCLRWCTYSREEESVVSAVSLSDIP | MALLRLIGTGENTTEASFLNVLKIKPLDTTRV | 758815 | 759007 | + | 7 |
| 5 | VVSAVSLSDIPMALLRLIGTG | ENTTEASFLNVLKIKPLDTTRV | 758875 | 759007 | + | 6 |
| 6 | VSAVSLSDIPMALLRLIGTGE | NTTEASFLNVLKIKPLDTTRV | 758878 | 759007 | + | 6 |
| 7 | VSLSDIPMALLRLIGTGEN | TTEASFLNVLKIKPLDTTRV | 758887 | 759007 | + | 6 |

| | | | | | | |
|---|---|---|---|---|---|---|
| 8 | MRTVTTPTLTGDAGISLKPMMPVT | FTPAILVTTPYAAGGLAAGAASVAAYEAGRAAGAGK | 759322 | 759139 | - | 27 |
| 9 | MAKIADFMNTQQLGPIEQGLTNTDGVPVM | ATPAVAVPVTVKITAGIVAFTIPVGAYVAGRVVN | 759581 | 759389 | - | 31 |
| 10 | MATGTVKWFNAEKGFGFIAQDGGGPDVFVHYSA | INASGFRSLEENQVVNFDVTQGPKGPQAENVTPA | 766236 | 766032 | - | 7 |

### Results for WP_107479021.1 [Streptomyces sp. alain-838] back to top

previous - next

#### Architecture

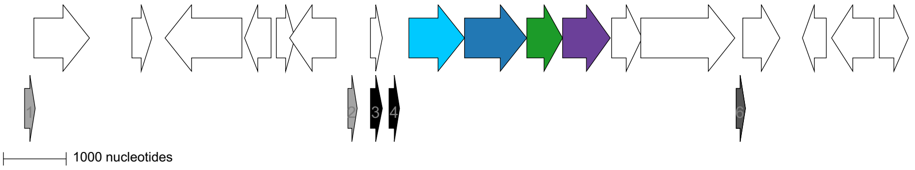

Link to nucleotide sequence

| Accession | start | end | direction | length (aa) | Pfam/HMM | name | description | E-value |
|---|---|---|---|---|---|---|---|---|
| WP_107479018.1 | 4014 | 4905 | + | 296 | PF00089<br>PF13365 | Trypsin<br>Trypsin_2 | Trypsin<br>Trypsin-like peptidase domain | 4.20E-32<br>6.20E-10 |
| WP_016827744.1 | 5580 | 5893 | + | 104 | INFERRED GENE | - | - | - |
| WP_107479019.1 | 7325 | 6101 | - | 407 | PF07690<br>TIGR00880<br>TIGR00895<br>TIGR00711<br>TIGR00710 | MFS_1<br>TIGR00880<br>TIGR00895<br>TIGR00711<br>TIGR00710 | Major Facilitator Superfamily<br>2_A_01_02: multidrug resistance protein<br>2A0115: MFS transporter, aromatic acid:H+ symporter (AAHS) family<br>efflux_EmrB: drug resistance MFS transporter, drug:H+ antiporter-2 (14 Spanner) (DHA2) family<br>efflux_Bcr_CflA: drug resistance transporter, Bcr/CflA subfamily | 7.50E-41<br>1.90E-22<br>2.40E-14<br>1.00E-13<br>1.40E-11 |
| WP_107479028.1 | 7787 | 7361 | - | 141 | PF13411<br>TIGR02043<br>TIGR02047<br>TIGR02044<br>TIGR02051 | MerR_1<br>TIGR02043<br>TIGR02047<br>TIGR02044<br>TIGR02051 | MerR HTH family regulatory protein<br>ZntR: Zn(II)-responsive transcriptional regulator<br>CadR-PbrR: Cd(II)/Pb(II)-responsive transcriptional regulator<br>CueR: Cu(I)-responsive transcriptional regulator<br>MerR: Hg(II)-responsive transcriptional regulator | 9.90E-18<br>3.20E-16<br>2.20E-13<br>9.30E-13<br>2.60E-12 |
| WP_008734492.1 | 7874 | 8168 | + | 98 | INFERRED GENE | - | - | - |
| WP_143657577.1 | 8821 | 8074 | - | 248 | NO PFAM MATCH | - | - | - |
| WP_143657578.1 | 9361 | 9559 | + | 65 | NO PFAM MATCH | - | - | - |
| WP_143657579.1 | 9990 | 10869 | + | 292 | LinH<br>LinG<br>PF05090 | LinH<br>LinG<br>VKG_Carbox | LinH (Fusion of LinE/LinG)<br>LinG Homolog<br>Vitamin K-dependent gamma-carboxylase | 3.80E-115<br>1.50E-102<br>8.10E-04 |
| WP_107479021.1 | 10865 | 11852 | + | 328 | LinE<br>LinH<br>PF12697<br>PF12146<br>PF00561 | LinE<br>LinH<br>Abhydrolase_6<br>Hydrolase_4<br>Abhydrolase_1 | LinE Homolog<br>LinH (Fusion of LinE/LinG)<br>Alpha/beta hydrolase family<br>Serine aminopeptidase, S33<br>alpha/beta hydrolase fold | 5.20E-116<br>5.50E-111<br>1.00E-18<br>1.30E-11<br>2.90E-11 |
| WP_143657580.1 | 11851 | 12430 | + | 192 | LinL | LinL | LinL | 7.60E-60 |
| WP_107479022.1 | 12433 | 13198 | + | 254 | PF13649<br>PF13489<br>PF08241<br>TIGR01983<br>PF08242 | Methyltransf_25<br>Methyltransf_23<br>Methyltransf_11<br>TIGR01983<br>Methyltransf_12 | Methyltransferase domain<br>Methyltransferase domain<br>Methyltransferase domain<br>UbiG: 3-demethylubiquinone-9 3-O-methyltransferase<br>Methyltransferase domain | 1.30E-14<br>2.90E-11<br>4.00E-10<br>1.20E-08<br>2.60E-07 |
| WP_107479023.1 | 13216 | 13702 | + | 161 | PF00072<br>PF18462 | Response_reg<br>DUF5612 | Response regulator receiver domain<br>Domain of unknown function (DUF5612) | 6.50E-06<br>9.40E-04 |
| WP_107479024.1 | 13664 | 15173 | + | 502 | TIGR02734<br>TIGR03467<br>TIGR00292<br>PF13450<br>TIGR01372 | TIGR02734<br>TIGR03467<br>TIGR00292<br>NAD_binding_8<br>TIGR01372 | crtI_fam: phytoene desaturase<br>HpnE: squalene-associated FAD-dependent desaturase<br>TIGR00292: thiazole biosynthesis enzyme<br>NAD(P)-binding Rossmann-like domain<br>soxA: sarcosine oxidase, alpha subunit family | 6.80E-09<br>1.10E-08<br>3.10E-08<br>4.80E-08<br>2.30E-07 |
| WP_005477319.1 | 15305 | 15891 | + | 195 | INFERRED GENE | - | - | - |
| WP_107479025.1 | 16622 | 16244 | - | 125 | PF14087 | DUF4267 | Domain of unknown function (DUF4267) | 4.30E-19 |
| WP_107479026.1 | 17377 | 16702 | - | 224 | PF00106<br>PF13561<br>TIGR04316<br>TIGR01963<br>TIGR02415 | adh_short<br>adh_short_C2<br>TIGR04316<br>TIGR01963<br>TIGR02415 | short chain dehydrogenase<br>Enoyl-(Acyl carrier protein) reductase<br>dhbA_paeA: 2,3-dihydro-2,3-dihydroxybenzoate dehydrogenase<br>PHB_DH: 3-hydroxybutyrate dehydrogenase<br>23BDH: acetoin reductases | 3.20E-25<br>7.90E-19<br>4.90E-15<br>8.50E-14<br>8.00E-13 |
| WP_018520665.1 | 17466 | 17940 | + | 158 | INFERRED GENE | - | - | - |

| index | leader | core | start | end | dir | score |
|---|---|---|---|---|---|---|
| 1 | VPGPPRGLDDALRVPA | SPPSGPGAGGSVRRFSPPRTRLVAVPREDPLTGWIGSP | 3868 | 4033 | + | 6 |
| 2 | VIATAPFQPRSSTSGQRSCSLGC | RTPRRGSLTCQNGSCPVVAPARGYPSQP | 9009 | 9165 | + | 6 |
| 3 | MSPVLEMVNSSQELASLSMHSHESIQGDRAPG | MWVADVAQVATNAISGNVGAVGGAVIATVGVIG | 9361 | 9559 | + | 18 |
| 4 | VHTLASIANEDERLEQVQLDADQPNLLGNRTAPVL | STPAALVTGVVTVNSVANAIW | 9658 | 9829 | + | 29 |
| 5 | VLAVTGGQVRGCVRVVCSGWCGLG | VVVGGARSTSAGCARWGGSTSTTR | 15183 | 15330 | + | 7 |
| 6 | VTGGQVRGCVRVVCSGWCGLGV | VVGGARSTSAGCARWGGSTSTTR | 15192 | 15330 | + | 6 |

### Results for RPK73392.1 [Streptomyces sp. ADI97-07] back to top

previous - next

#### Architecture

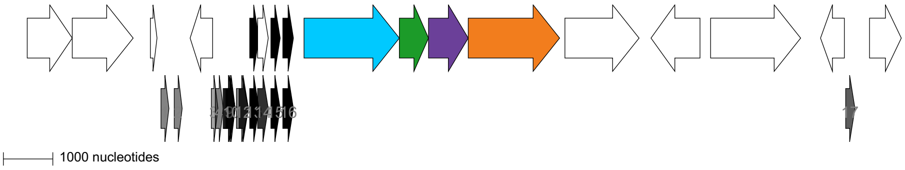

Link to nucleotide sequence

| Accession | start | end | direction | length (aa) | Pfam/HMM | name | description | E-value |
|---|---|---|---|---|---|---|---|---|
| RPK73384.1 | 760935 | 761841 | + | 301 | NO PFAM MATCH | - | - | - |
| RPK73385.1 | 761837 | 763076 | + | 412 | PF01547<br>PF13416<br>PF13531 | SBP_bac_1<br>SBP_bac_8<br>SBP_bac_11 | Bacterial extracellular solute-binding protein<br>Bacterial extracellular solute-binding protein<br>Bacterial extracellular solute-binding protein | 1.10E-14<br>1.60E-07<br>3.50E-05 |
| RPK73386.1 | 763433 | 763568 | + | 44 | NO PFAM MATCH | - | - | - |
| RPK73387.1 | 764678 | 764225 | - | 150 | PF12647 | RNHCP | RNHCP domain | 2.10E-34 |
| RPK73388.1 | 765420 | 765606 | + | 61 | LinA | LinA | LinA (linaridin precursor) | 1.40E-08 |
| RPK73389.1 | 765602 | 765797 | + | 64 | NO PFAM MATCH | - | - | - |
| RPK73390.1 | 765858 | 766053 | + | 64 | LinA | LinA | LinA (linaridin precursor) | 1.10E-09 |
| RPK73391.1 | 766096 | 766312 | + | 71 | LinA | LinA | LinA (linaridin precursor) | 1.30E-08 |
| RPK73392.1 | 766524 | 768450 | + | 641 | LinH<br>LinG<br>LinE<br>PF12697<br>PF00561 | LinH<br>LinG<br>LinE<br>Abhydrolase_6<br>Abhydrolase_1 | LinH (Fusion of LinE/LinG)<br>LinG Homolog<br>LinE Homolog<br>Alpha/beta hydrolase family<br>alpha/beta hydrolase fold | 2.20E-237<br>1.80E-114<br>8.60E-90<br>6.80E-14<br>3.50E-11 |
| RPK73393.1 | 768446 | 769055 | + | 202 | LinL | LinL | LinL | 3.80E-60 |
| RPK73394.1 | 769051 | 769858 | + | 268 | PF13649<br>PF08241<br>PF13847<br>PF13489<br>PF08242 | Methyltransf_25<br>Methyltransf_11<br>Methyltransf_31<br>Methyltransf_23<br>Methyltransf_12 | Methyltransferase domain<br>Methyltransferase domain<br>Methyltransferase domain<br>Methyltransferase domain<br>Methyltransferase domain | 9.40E-17<br>5.40E-11<br>9.10E-08<br>1.10E-07<br>1.30E-07 |
| RPK73395.1 | 769854 | 771699 | + | 614 | TIGR02204<br>TIGR02203<br>TIGR00958<br>TIGR03796<br>TIGR01846 | TIGR02204<br>TIGR02203<br>TIGR00958<br>TIGR03796<br>TIGR01846 | MsbA_rel: ABC transporter, permease/ATP-binding protein<br>MsbA_lipidA: lipid A export permease/ATP-binding protein MsbA<br>3a01208: antigen peptide transporter 2<br>NHLM_micro_ABC1: NHLM bacteriocin system ABC transporter, peptidase/ATP-binding protein<br>type_I_sec_HlyB: type I secretion system ATPase | 3.10E-101<br>7.90E-92<br>3.70E-85<br>8.40E-84<br>1.80E-80 |
| RPK73396.1 | 771802 | 773293 | + | 496 | TIGR02734<br>PF13450<br>TIGR03467<br>TIGR01317<br>PF01593 | TIGR02734<br>NAD_binding_8<br>TIGR03467<br>TIGR01317<br>Amino_oxidase | crtI_fam: phytoene desaturase<br>NAD(P)-binding Rossmann-like domain<br>HpnE: squalene-associated FAD-dependent desaturase<br>GOGAT_sm_gam: glutamate synthase, NADH/NADPH, small subunit<br>Flavin containing amine oxidoreductase | 6.50E-11<br>6.90E-10<br>7.20E-10<br>6.40E-09<br>3.50E-08 |
| RPK73034.1 | 774545 | 773537 | - | 335 | NO PFAM MATCH | - | - | - |
| RPK73035.1 | 774749 | 776585 | + | 611 | NO PFAM MATCH | - | - | - |
| RPK73036.1 | 777445 | 776977 | - | 155 | PF12802<br>PF13463<br>PF01047<br>TIGR01889<br>TIGR02337 | MarR_2<br>HTH_27<br>MarR<br>TIGR01889<br>TIGR02337 | MarR family<br>Winged helix DNA-binding domain<br>MarR family<br>Staph_reg_Sar: staphylococcal accessory regulator family<br>HpaR: homoprotocatechuate degradation operon regulator, HpaR | 2.40E-09<br>1.20E-08<br>1.50E-08<br>5.90E-06<br>4.00E-04 |
| RPK73037.1 | 777954 | 778614 | + | 219 | PF00857 | Isochorismatase | Isochorismatase family | 1.50E-27 |

| index | leader | core | start | end | dir | score |
|---|---|---|---|---|---|---|
| 1 | VWLECVLILFRMACGSGWLRCFHPLPN | GADVTPGGCGSPTGRLSRASSTCCGPV | 763644 | 763809 | + | 7 |
| 2 | VWPRLHAVLLTELRRAGLVDLDDCAVD | SSHVRALNGGSCRPLSCRPRPPRLVVVR | 763892 | 764060 | + | 7 |
| 3 | MRTERAPVGVVTGQTRQCMHRQ | GTVDAQCAGRARASSSARRTGTH | 764659 | 764797 | + | 6 |
| 4 | LTHNAQAGPGPVRLLGVLGLTDDTSQ | SCTGRSWPWAWKASPVIRPDWITGPC | 764732 | 764891 | + | 7 |
| 5 | LTLLCTGTGRAADVLTSRWRGQHLVQLVAGDGGRLQKPVQV | GVPSVRDLTGGLRVADVGGKQPGPCGVQVGDRGAACTGT | 764900 | 765143 | + | 7 |
| 6 | LLCTGTGRAADVLTSRWRGQHLVQLVAGDGGRLQKPVQV | GVPSVRDLTGGLRVADVGGKQPGPCGVQVGDRGAACTGT | 764906 | 765143 | + | 7 |
| 7 | LCTGTGRAADVLTSRWRGQHLVQLVAGDGGRLQKPVQV | GVPSVRDLTGGLRVADVGGKQPGPCGVQVGDRGAACTGT | 764909 | 765143 | + | 7 |
| 8 | VLTSRWRGQHLVQLVAGDGGRLQKPVQV | GVPSVRDLTGGLRVADVGGKQPGPCGVQVGDRGAACTGT | 764939 | 765143 | + | 7 |
| 9 | LTSRWRGQHLVQLVAGDGGRLQKPVQV | GVPSVRDLTGGLRVADVGGKQPGPCGVQVGDRGAACTGT | 764942 | 765143 | + | 7 |
| 10 | VQLVAGDGGRLQKPVQV | GVPSVRDLTGGLRVADVGGKQPGPCGVQVGDRGAACTGT | 764972 | 765143 | + | 7 |
| 11 | MRLSRPPRTAPAPVAPCLHHPAGGP...SCQPTVFPRAGGPAIVDGTDGTPPG | RTPRVRGAALPYGPVAGASGCRPG | 765176 | 765407 | + | 7 |
| 12 | VAPCLHHPAGGPAASCQPTVFPRAGGPAIVDGTDGTPPG | RTPRVRGAALPYGPVAGASGCRPG | 765215 | 765407 | + | 9 |
| 13 | MHQVRTPRWAAVGPLDERTEGQAFGGAAPIA | ATQVIVTVAAAVVVAYAAGAIIGSTEEPSL | 765420 | 765606 | + | 19 |

| | | | | | | |
|---|---|---|---|---|---|---|
| 14 | MTGTRWATASVCP | ATAVTTCALSENDAGHTGPAASPIA...VLVVVAEVCASPAISVIAGTHEPQL | 765602 | 765797 | + | 9 |
| 15 | MNASVFQTVPAAARPLTEEDPGYTATGTAPVA | ATPVLATAVFAAGFVAGVVVCDAIGDTSPVEQ | 765858 | 766053 | + | 30 |
| 16 | MNDTSFLTRPGALTDIRVDDPQGRVIINHPPVM | ATPAVVSGVIIGVALVNAFVAGRDGGPLEPPIEIDVPA | 766096 | 766312 | + | 29 |
| 17 | VRPSSARQGPPCPGNETWGGPHLAIVEADVTGRSVVRRIRRASTSASA | GTAIRVNCALRTVS | 777493 | 777682 | + | 8 |

### Results for RAJ66737.1 [Streptomyces sp. Amel2xB2] back to top

previous - next

#### Architecture

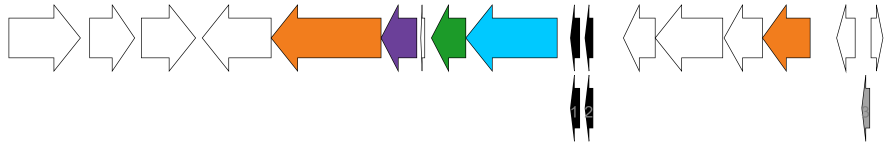

1000 nucleotides

Link to nucleotide sequence

| Accession | start | end | direction | length (aa) | Pfam/HMM | name | description | E-value |
|---|---|---|---|---|---|---|---|---|
| RAJ66729.1 | 369330 | 370827 | + | 498 | NO PFAM MATCH | - | - | - |
| RAJ66730.1 | 371015 | 371969 | + | 317 | PF01259<br>TIGR00081<br>TIGR02735 | SAICAR_synt<br>TIGR00081<br>TIGR02735 | SAICAR synthetase<br>purC: phosphoribosylaminoimidazolesuccinocarboxamide synthase<br>purC_vibrio: phosphoribosylaminoimidazole-succinocarboxamide synthase | 1.60E-98<br>2.10E-69<br>2.50E-20 |
| RAJ66731.1 | 372123 | 373254 | + | 376 | PF00196 | GerE | Bacterial regulatory proteins, luxR family | 2.90E-05 |
| RAJ66732.1 | 374859 | 373392 | - | 488 | TIGR02734<br>TIGR03467<br>PF13450<br>TIGR00292<br>PF01593 | TIGR02734<br>TIGR03467<br>NAD_binding_8<br>TIGR00292<br>Amino_oxidase | crtI_fam: phytoene desaturase<br>HpnE: squalene-associated FAD-dependent desaturase<br>NAD(P)-binding Rossmann-like domain<br>TIGR00292: thiazole biosynthesis enzyme<br>Flavin containing amine oxidoreductase | 1.10E-12<br>2.90E-09<br>5.50E-08<br>3.70E-07<br>1.60E-06 |
| RAJ66733.1 | 377153 | 374855 | - | 765 | TIGR02204<br>TIGR02203<br>TIGR00958<br>TIGR03796<br>TIGR01846 | TIGR02204<br>TIGR02203<br>TIGR00958<br>TIGR03796<br>TIGR01846 | MsbA_rel: ABC transporter, permease/ATP-binding protein<br>MsbA_lipidA: lipid A export permease/ATP-binding protein MsbA<br>3a01208: antigen peptide transporter 2<br>NHLM_micro_ABC1: NHLM bacteriocin system ABC transporter, peptidase/ATP-binding protein<br>type_I_sec_HlyB: type I secretion system ATPase | 4.60E-97<br>1.80E-93<br>7.20E-81<br>3.70E-79<br>3.00E-76 |
| RAJ66734.1 | 377919 | 377160 | - | 252 | PF13649<br>PF08241<br>PF13489<br>PF13847<br>TIGR01983 | Methyltransf_25<br>Methyltransf_11<br>Methyltransf_23<br>Methyltransf_31<br>TIGR01983 | Methyltransferase domain<br>Methyltransferase domain<br>Methyltransferase domain<br>Methyltransferase domain<br>UbiG: 3-demethylubiquinone-9 3-O-methyltransferase | 1.10E-18<br>2.10E-13<br>8.70E-13<br>4.80E-12<br>3.50E-10 |
| RAJ66735.1 | 378095 | 378002 | - | 30 | NO PFAM MATCH | - | - | - |
| RAJ66736.1 | 378957 | 378222 | - | 244 | LinL | LinL | LinL | 9.60E-62 |
| RAJ66737.1 | 380867 | 378953 | - | 637 | LinH<br>LinG<br>LinE<br>PF12697<br>LinL | LinH<br>LinG<br>LinE<br>Abhydrolase_6<br>LinL | LinH (Fusion of LinE/LinG)<br>LinG Homolog<br>LinE Homolog<br>Alpha/beta hydrolase family<br>LinL | 1.20E-245<br>8.10E-118<br>6.40E-98<br>1.70E-14<br>2.90E-11 |
| RAJ66738.1 | 381348 | 381168 | - | 59 | LinA | LinA | LinA (linaridin precursor) | 1.90E-15 |
| RAJ66739.1 | 381629 | 381455 | - | 57 | LinA | LinA | LinA (linaridin precursor) | 2.90E-14 |
| RAJ66740.1 | 382939 | 382282 | - | 218 | PF00072<br>PF00196<br>TIGR02875<br>TIGR02154<br>TIGR02956 | Response_reg<br>GerE<br>TIGR02875<br>TIGR02154<br>TIGR02956 | Response regulator receiver domain<br>Bacterial regulatory proteins, luxR family<br>spore_0_A: sporulation transcription factor Spo0A<br>PhoB: phosphate regulon transcriptional regulatory protein PhoB<br>TMAO_torS: TMAO reductase sytem sensor TorS | 2.70E-25<br>9.30E-14<br>9.30E-14<br>3.20E-11<br>1.10E-09 |
| RAJ66741.1 | 384375 | 382935 | - | 479 | PF07730 | HisKA_3 | Histidine kinase | 2.40E-14 |
| RAJ66742.1 | 385210 | 384397 | - | 270 | PF01061<br>PF12698<br>TIGR03062<br>TIGR01247 | ABC2_membrane<br>ABC2_membrane_3<br>TIGR03062<br>TIGR01247 | ABC-2 type transporter<br>ABC-2 family transporter protein<br>pip_yhgE_Cterm: YhgE/Pip C-terminal domain<br>drrB: daunorubicin resistance ABC transporter membrane protein | 2.00E-19<br>2.60E-11<br>2.80E-06<br>2.20E-05 |
| RAJ66743.1 | 386199 | 385206 | - | 330 | TIGR01188<br>TIGR03864<br>TIGR01288<br>TIGR04406<br>TIGR03522 | TIGR01188<br>TIGR03864<br>TIGR01288<br>TIGR04406<br>TIGR03522 | drrA: daunorubicin resistance ABC transporter, ATP-binding protein<br>PQQ_ABC_ATP: ABC transporter, ATP-binding subunit, PQQ-dependent alcohol dehydrogenase system<br>nodI: nodulation ABC transporter NodI<br>LPS_export_lptB: LPS export ABC transporter ATP-binding protein<br>GldA_ABC_ATP: gliding motility-associated ABC transporter ATP-binding subunit GldA | 2.00E-70<br>4.80E-50<br>3.80E-48<br>5.60E-39<br>6.20E-38 |
| RAJ66744.1 | 387152 | 386777 | - | 124 | PF11774 | Lsr2 | Lsr2 | 2.00E-34 |
| RAJ66745.1 | 387497 | 387755 | + | 85 | PF02700<br>TIGR00302 | PurS<br>TIGR00302 | Phosphoribosylformylglycinamidine (FGAM) synthase<br>TIGR00302: phosphoribosylformylglycinamidine synthase, purS protein | 1.40E-27<br>3.70E-20 |

| index | leader | core | start | end | dir | score |
|---|---|---|---|---|---|---|
| 1 | MIDVTSHINESAELSSVEVTDAEGVVFTDYTPVL | ATPGAFLGGVAIASATVAAFEQGQG | 381348 | 381168 | - | 29 |
| 2 | MLEEKLNLQIQDELHEEHAGEIADKAAPVL | ATPALAGAFVGGAAVVTGAYAAGKAVG | 381629 | 381455 | - | 30 |

| | | | | | | |
|---|---|---|---|---|---|---|
| 3 | VLSDGACCQRSDASNSLPGMEIYARGYSGRRPSP | RTLHALSIRIPHPPADLGSAGALGKSP | 387475 | 387289 | - | 6 |

### Results for RKN03588.1 [Streptomyces sp. AZ1-7] back to top

previous - next

#### Architecture

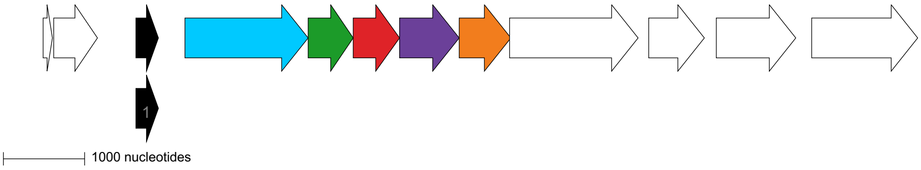

Link to nucleotide sequence

| Accession | start | end | direction | length (aa) | Pfam/HMM | name | description | E-value |
|---|---|---|---|---|---|---|---|---|
| RKN03563.1 | 0 | 111 | + | 36 | NO PFAM MATCH | - | - | - |
| WP_008745676.1 | 124 | 673 | + | 183 | INFERRED GENE | - | - | - |
| RKN03564.1 | 1140 | 1410 | + | 89 | LinA | LinA | LinA (linaridin precursor) | 1.90E-13 |
| RKN03588.1 | 1744 | 3265 | + | 506 | LinH<br>LinE<br>LinG<br>PF12697<br>TIGR02427 | LinH<br>LinE<br>LinG<br>Abhydrolase_6<br>TIGR02427 | LinH (Fusion of LinE/LinG)<br>LinE Homolog<br>LinG Homolog<br>Alpha/beta hydrolase family<br>protocat_pcaD: 3-oxoadipate enol-lactonase | 1.70E-160<br>8.60E-67<br>2.50E-66<br>4.50E-08<br>4.20E-07 |
| RKN03565.1 | 3254 | 3821 | + | 188 | LinL | LinL | LinL | 3.30E-53 |
| RKN03566.1 | 3817 | 4390 | + | 190 | TIGR00521<br>TIGR02113<br>PF02441 | TIGR00521<br>TIGR02113<br>Flavoprotein | coaBC_dfp: phosphopantothenoylcysteine decarboxylase / phosphopantothenate--cysteine ligase<br>coaC_strep: phosphopantothenoylcysteine decarboxylase<br>Flavoprotein | 5.50E-13<br>9.00E-11<br>1.40E-09 |
| RKN03567.1 | 4382 | 5120 | + | 245 | PF13649<br>PF08241<br>PF13847<br>TIGR02021<br>TIGR01934 | Methyltransf_25<br>Methyltransf_11<br>Methyltransf_31<br>TIGR02021<br>TIGR01934 | Methyltransferase domain<br>Methyltransferase domain<br>Methyltransferase domain<br>BchM-ChlM: magnesium protoporphyrin O-methyltransferase<br>MenG_MenH_UbiE: ubiquinone/menaquinone biosynthesis methyltransferase | 1.50E-15<br>2.90E-14<br>8.20E-11<br>1.30E-09<br>4.60E-09 |
| RKN03568.1 | 5116 | 5749 | + | 210 | TIGR02857<br>TIGR02673<br>TIGR01189<br>TIGR02982<br>TIGR03265 | TIGR02857<br>TIGR02673<br>TIGR01189<br>TIGR02982<br>TIGR03265 | CydD: thiol reductant ABC exporter, CydD subunit<br>FtsE: cell division ATP-binding protein FtsE<br>ccmA: heme ABC exporter, ATP-binding protein CcmA<br>heterocyst_DevA: ABC exporter ATP-binding subunit, DevA family<br>PhnT2: putative 2-aminoethylphosphonate ABC transporter, ATP-binding protein | 3.50E-25<br>6.80E-25<br>6.80E-24<br>1.20E-20<br>1.70E-20 |
| RKN03569.1 | 5745 | 7323 | + | 525 | NO PFAM MATCH | - | - | - |
| RKN03570.1 | 7443 | 8142 | + | 232 | PF02909<br>PF00440 | TetR_C_1<br>TetR_N | Tetracyclin repressor-like, C-terminal domain<br>Bacterial regulatory proteins, tetR family | 5.90E-14<br>2.10E-09 |
| RKN03571.1 | 8275 | 9265 | + | 329 | TIGR02824<br>TIGR01751<br>TIGR02817<br>TIGR02823<br>PF13602 | TIGR02824<br>TIGR01751<br>TIGR02817<br>TIGR02823<br>ADH_zinc_N_2 | quinone_pig3: putative NAD(P)H quinone oxidoreductase, PIG3 family<br>crot-CoA-red: crotonyl-CoA reductase<br>adh_fam_1: zinc-binding alcohol dehydrogenase family protein<br>oxido_YhdH: putative quinone oxidoreductase, YhdH/YhfP family<br>Zinc-binding dehydrogenase | 2.70E-48<br>1.30E-36<br>7.10E-28<br>1.10E-25<br>3.20E-25 |
| RKN03572.1 | 9455 | 10763 | + | 435 | TIGR00900<br>PF07690 | TIGR00900<br>MFS_1 | 2A0121: H+ Antiporter protein<br>Major Facilitator Superfamily | 7.30E-22<br>2.50E-14 |

| index | leader | core | start | end | dir | score |
|---|---|---|---|---|---|---|
| 1 | MGNFRSVETLSLSKETRKGSKMRLN...TQEFANGVLRDAAPGFHSDSEIPAM | ATPATPTAAQFVIQGSTICLVC | 1140 | 1410 | + | 34 |

### Results for SCK62389.1 [Streptomyces sp. AmelKG-D3] back to top

previous - next

#### Architecture

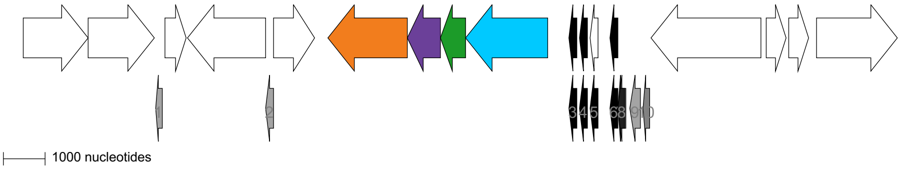

Link to nucleotide sequence

| Accession | start | end | direction | length (aa) | Pfam/HMM | name | description | E-value |
|---|---|---|---|---|---|---|---|---|
| SCK62373.1 | 5562 | 7107 | + | 514 | PF00501<br>TIGR03208<br>TIGR03098<br>TIGR02275<br>TIGR01733 | AMP-binding<br>TIGR03208<br>TIGR03098<br>TIGR02275<br>TIGR01733 | AMP-binding enzyme<br>cyc_hxne_CoA_lg: cyclohexanecarboxylate-CoA ligase<br>ligase_PEP_1: acyl-CoA ligase (AMP-forming), exosortase A-associated<br>DHB_AMP_lig: (2,3-dihydroxybenzoyl)adenylate synthase<br>AA-adenyl-dom: amino acid adenylation domain | 8.30E-68<br>4.40E-44<br>5.50E-43<br>1.00E-38<br>5.70E-38 |

| | | | | | | | | |
|---|---|---|---|---|---|---|---|---|
| SCK62375.1 | 7114 | 8707 | + | 530 | PF00501<br>TIGR03205<br>TIGR03098<br>TIGR02262<br>TIGR03208 | AMP-binding<br>TIGR03205<br>TIGR03098<br>TIGR02262<br>TIGR03208 | AMP-binding enzyme<br>pimA: dicarboxylate--CoA ligase PimA<br>ligase_PEP_1: acyl-CoA ligase (AMP-forming), exosortase A-associated<br>benz_CoA_lig: benzoate-CoA ligase family<br>cyc_hxne_CoA_lg: cyclohexanecarboxylate-CoA ligase | 8.00E-114<br>8.20E-79<br>4.00E-75<br>9.90E-75<br>9.00E-72 |
| SCK62378.1 | 8937 | 9474 | + | 178 | PF12840<br>PF01022<br>PF13518 | HTH_20<br>HTH_5<br>HTH_28 | Helix-turn-helix domain<br>Bacterial regulatory protein, arsR family<br>Helix-turn-helix domain | 3.50E-11<br>1.30E-06<br>9.50E-05 |
| SCK62380.1 | 11353 | 9499 | - | 617 | NO PFAM MATCH | - | - | - |
| SCK62382.1 | 11542 | 12556 | + | 337 | NO PFAM MATCH | - | - | - |
| SCK62384.1 | 14755 | 12853 | - | 633 | TIGR02204<br>TIGR02203<br>TIGR00958<br>TIGR03796<br>TIGR03797 | TIGR02204<br>TIGR02203<br>TIGR00958<br>TIGR03796<br>TIGR03797 | MsbA_rel: ABC transporter, permease/ATP-binding protein<br>MsbA_lipidA: lipid A export permease/ATP-binding protein MsbA<br>3a01208: antigen peptide transporter 2<br>NHLM_micro_ABC1: NHLM bacteriocin system ABC transporter, peptidase/ATP-binding protein<br>NHLM_micro_ABC2: NHLM bacteriocin system ABC transporter, ATP-binding protein | 6.70E-99<br>6.70E-92<br>9.70E-84<br>7.20E-82<br>2.60E-77 |
| SCK62385.1 | 15564 | 14751 | - | 270 | PF13649<br>PF08241<br>PF13847<br>PF13489<br>PF08242 | Methyltransf_25<br>Methyltransf_11<br>Methyltransf_31<br>Methyltransf_23<br>Methyltransf_12 | Methyltransferase domain<br>Methyltransferase domain<br>Methyltransferase domain<br>Methyltransferase domain<br>Methyltransferase domain | 4.70E-16<br>8.00E-12<br>6.80E-07<br>1.70E-06<br>1.80E-06 |
| SCK62387.1 | 16169 | 15560 | - | 202 | LinL | LinL | LinL | 7.10E-66 |
| SCK62389.1 | 18118 | 16165 | - | 650 | LinH<br>LinG<br>LinE<br>LinL<br>PF12697 | LinH<br>LinG<br>LinE<br>LinL<br>Abhydrolase_6 | LinH (Fusion of LinE/LinG)<br>LinG Homolog<br>LinE Homolog<br>LinL<br>Alpha/beta hydrolase family | 3.40E-255<br>3.30E-114<br>2.70E-95<br>1.20E-44<br>8.80E-14 |
| SCK62391.1 | 18826 | 18625 | - | 66 | LinA | LinA | LinA (linaridin precursor) | 2.40E-12 |
| SCK62393.1 | 19075 | 18883 | - | 63 | LinA | LinA | LinA (linaridin precursor) | 6.90E-09 |
| SCK62395.1 | 19322 | 19148 | - | 57 | NO PFAM MATCH | - | - | - |
| SCK62397.1 | 19814 | 19610 | - | 67 | LinA | LinA | LinA (linaridin precursor) | 3.10E-11 |
| SCK62400.1 | 23251 | 20596 | - | 884 | TIGR03903<br>PF13191<br>PF13424<br>PF07719<br>PF13401 | TIGR03903<br>AAA_16<br>TPR_12<br>TPR_2<br>AAA_22 | TOMM_kin_cyc: TOMM system kinase/cyclase fusion protein<br>AAA ATPase domain<br>Tetratricopeptide repeat<br>Tetratricopeptide repeat<br>AAA domain | 2.00E-20<br>7.10E-19<br>2.40E-05<br>9.30E-05<br>1.40E-04 |
| SCK62402.1 | 23361 | 23841 | + | 159 | PF03745 | DUF309 | Domain of unknown function (DUF309) | 4.90E-19 |
| SCK62404.1 | 23925 | 24390 | + | 154 | NO PFAM MATCH | - | - | - |
| SCK62406.1 | 24587 | 26522 | + | 644 | PF10022 | DUF2264 | Uncharacterized protein conserved in bacteria (DUF2264) | 9.70E-97 |

| index | leader | core | start | end | dir | score |
|---|---|---|---|---|---|---|
| 1 | VVSADGPSSAATADAWAERGRKHRRS | YRSGPVLTVVQVRCGPDGRPGPVRPG | 8881 | 8722 | - | 6 |
| 2 | VATSCRSPWPFPRTRPAGQRITSLPGEAGGEVS | AKSQASLAPGKFRMRDHLKERAPIAMTGGRAPK | 11567 | 11366 | - | 6 |
| 3 | MNMSPTPAALRGAGGLSESDPGRAHSTLVPVA | ATPGVLAGVAVGVALVNAFAAGYNHCGGNVELPM | 18826 | 18625 | - | 31 |
| 4 | MNTSSTPAVLTATGALTESDPGATLPAIAPVA | ATPVVVVAALGAGFVAGYAAGRAAGNVELPM | 19075 | 18883 | - | 30 |
| 5 | MSTASIPAVLSGTGALTESDRGRGPLD | TVPVTSVTVTMTACVGISVCLTGSTIELPQ | 19322 | 19148 | - | 10 |
| 6 | MPEFRQPGWTRGLAPLDEGAGGQVFGGASPVA | ATPAVVATAGAVVVAFAAGVAARHLANGGNVELPM | 19814 | 19610 | - | 31 |
| 7 | MPAELSSLSVGLSASCGAHHRTS | VHAGAGSRVRSRAPGRAPVTAHP | 19964 | 19823 | - | 8 |
| 8 | VPEPAGCCCGVRILMPAELSSLSVGLSASC | GAHHRTSVHAGAGSRVRSRAPGRAPVTAHP | 20006 | 19823 | - | 8 |
| 9 | VCRRERLVDRAGRHPSDLLGPRRPWCRSRRSAQRLGQTR | CAFGGEADDGAGAGTGGGLAGPWPGGDLGECGVLAPLRR | 20349 | 20112 | - | 6 |
| 10 | LTCAAARTTGGNRSGTWRSLVAHLTGGQ | GVAGSNPVVPTGDSRRSGPVSEQSETGP | 20588 | 20417 | - | 7 |

### Results for RKN04869.1 [Streptomyces sp. AZ1-7] back to top

previous - next

#### Architecture

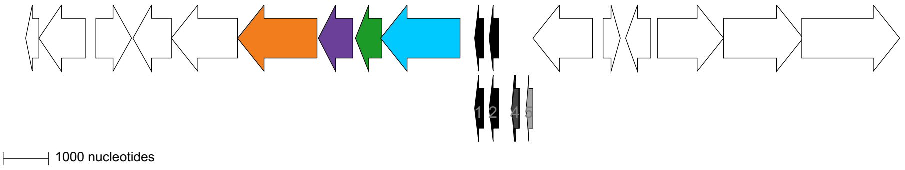

1000 nucleotides

Link to nucleotide sequence

| Accession | start | end | direction | length (aa) | Pfam/HMM | name | description | E-value |
|---|---|---|---|---|---|---|---|---|
| RKN04863.1 | 50794 | 50476 | - | 105 | PF00550<br>TIGR01688 | PP-binding<br>TIGR01688 | Phosphopantetheine attachment site<br>dltC: D-alanine--poly(phosphoribitol) ligase, subunit 2 | 2.20E-09<br>4.20E-06 |
| RKN04895.1 | 51807 | 50790 | - | 338 | TIGR00747 | TIGR00747 | fabH: 3-oxoacyl-[acyl-carrier-protein] synthase III | 4.80E-09 |
| RKN04896.1 | 52027 | 52819 | + | 263 | PF01551 | Peptidase_M23 | Peptidase family M23 | 3.30E-16 |

| | | | | | | | | |
|---|---|---|---|---|---|---|---|---|
| RKN04864.1 | 53704 | 52852 | - | 283 | NO PFAM MATCH | - | - | - |
| RKN04865.1 | 55185 | 53700 | - | 494 | TIGR02734<br>TIGR00292<br>TIGR02733<br>PF01946<br>TIGR01317 | TIGR02734<br>TIGR00292<br>TIGR02733<br>Thi4<br>TIGR01317 | crtI_fam: phytoene desaturase<br>TIGR00292: thiazole biosynthesis enzyme<br>desat_CrtD: C-3',4' desaturase CrtD<br>Thi4 family<br>GOGAT_sm_gam: glutamate synthase, NADH/NADPH, small subunit | 2.60E-09<br>4.50E-09<br>5.60E-08<br>2.30E-07<br>2.30E-07 |
| RKN04866.1 | 56933 | 55181 | - | 583 | TIGR02204<br>TIGR02203<br>TIGR00958<br>TIGR03796<br>TIGR01846 | TIGR02204<br>TIGR02203<br>TIGR00958<br>TIGR03796<br>TIGR01846 | MsbA_rel: ABC transporter, permease/ATP-binding protein<br>MsbA_lipidA: lipid A export permease/ATP-binding protein MsbA<br>3a01208: antigen peptide transporter 2<br>NHLM_micro_ABC1: NHLM bacteriocin system ABC transporter, peptidase/ATP-binding protein<br>type_I_sec_HlyB: type I secretion system ATPase | 7.50E-100<br>4.10E-96<br>5.60E-87<br>2.00E-84<br>7.30E-84 |
| RKN04867.1 | 57722 | 56951 | - | 256 | PF13649<br>PF13489<br>PF08241<br>PF08242<br>PF13847 | Methyltransf_25<br>Methyltransf_23<br>Methyltransf_11<br>Methyltransf_12<br>Methyltransf_31 | Methyltransferase domain<br>Methyltransferase domain<br>Methyltransferase domain<br>Methyltransferase domain<br>Methyltransferase domain | 3.30E-13<br>8.00E-09<br>1.70E-08<br>1.40E-07<br>6.50E-07 |
| RKN04868.1 | 58355 | 57776 | - | 192 | LinL | LinL | LinL | 4.00E-62 |
| RKN04869.1 | 60088 | 58351 | - | 578 | LinH<br>LinE<br>LinG<br>PF12697<br>PF00561 | LinH<br>LinE<br>LinG<br>Abhydrolase_6<br>Abhydrolase_1 | LinH (Fusion of LinE/LinG)<br>LinE Homolog<br>LinG Homolog<br>Alpha/beta hydrolase family<br>alpha/beta hydrolase fold | 1.10E-217<br>1.90E-96<br>2.10E-90<br>5.30E-15<br>1.30E-10 |
| RKN04870.1 | 60614 | 60422 | - | 63 | LinA | LinA | LinA (linaridin precursor) | 5.30E-09 |
| RKN04871.1 | 60937 | 60742 | - | 64 | LinA | LinA | LinA (linaridin precursor) | 9.60E-17 |
| RKN04897.1 | 63039 | 61719 | - | 439 | PF05139 | Erythro_esteras | Erythromycin esterase | 1.20E-107 |
| RKN04872.1 | 63264 | 63645 | + | 126 | PF11338 | DUF3140 | Protein of unknown function (DUF3140) | 5.60E-25 |
| RKN04873.1 | 64323 | 63771 | - | 183 | PF00754<br>PF06439 | F5_F8_type_C<br>DUF1080 | F5/8 type C domain<br>Domain of Unknown Function (DUF1080) | 7.20E-26<br>2.40E-06 |
| RKN04874.1 | 64452 | 65913 | + | 486 | TIGR02224<br>PF02899<br>PF00589 | TIGR02224<br>Phage_int_SAM_1<br>Phage_integrase | recomb_XerC: tyrosine recombinase XerC<br>Phage integrase, N-terminal SAM-like domain<br>Phage integrase family | 8.00E-06<br>4.00E-04<br>7.90E-04 |
| RKN04875.1 | 65912 | 67670 | + | 585 | NO PFAM MATCH | - | - | - |
| RKN04876.1 | 67666 | 69808 | + | 713 | NO PFAM MATCH | - | - | - |

| index | leader | core | start | end | dir | score |
|---|---|---|---|---|---|---|
| 1 | MSKIIESVVAVAPEELSDQTTGEFGAVMPVQ | ATPGLVAFGVGFAGGAGVAWLGVQAYEAGIND | 60614 | 60422 | - | 31 |
| 2 | MEQKIASFMNEQQLGPMEQGYQDDVKSPVL | ATPAAGVATCWYVGGAVAGAASVVGAYVCGAVFG | 60937 | 60742 | - | 30 |
| 3 | LLSDIDHIDRGGVEKGATCVPDSSP | EREEVIARHMVVSERGVSLRNLYDV | 61379 | 61226 | - | 7 |
| 4 | LEGCTKHSILLSDIDHIDRGGVEKGATCV | PDSSPEREEVIARHMVVSERGVSLRNLYDV | 61406 | 61226 | - | 7 |
| 5 | VTAQRACRERSVKRRHTACLRRVRGPVR | PVPGRDRGEGHVPGATRQFAENIA | 61713 | 61554 | - | 6 |

### Results for RKN10930.1 [Streptomyces sp. AZ1-7] back to top

previous - next

#### Architecture

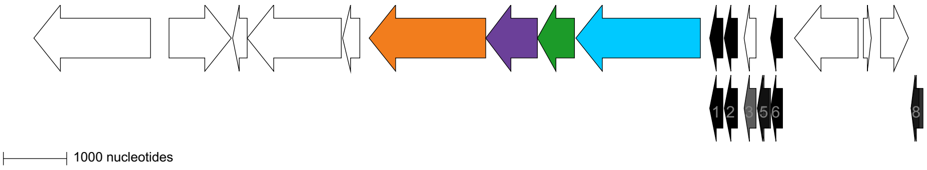

Link to nucleotide sequence

| Accession | start | end | direction | length (aa) | Pfam/HMM | name | description | E-value |
|---|---|---|---|---|---|---|---|---|
| RKN11093.1 | 102017 | 100178 | - | 612 | NO PFAM MATCH | - | - | - |
| RKN11094.1 | 102295 | 103276 | + | 326 | NO PFAM MATCH | - | - | - |
| RKN10925.1 | 103546 | 103303 | - | 80 | NO PFAM MATCH | - | - | - |
| RKN10926.1 | 105012 | 103545 | - | 488 | TIGR02734<br>TIGR03467<br>TIGR01317<br>PF13450<br>TIGR00292 | TIGR02734<br>TIGR03467<br>TIGR01317<br>NAD_binding_8<br>TIGR00292 | crtI_fam: phytoene desaturase<br>HpnE: squalene-associated FAD-dependent desaturase<br>GOGAT_sm_gam: glutamate synthase, NADH/NADPH, small subunit<br>NAD(P)-binding Rossmann-like domain<br>TIGR00292: thiazole biosynthesis enzyme | 6.40E-14<br>6.80E-09<br>5.20E-08<br>1.00E-07<br>1.70E-07 |
| RKN10927.1 | 105313 | 105034 | - | 92 | NO PFAM MATCH | - | - | - |
| RKN10928.1 | 107306 | 105458 | - | 615 | TIGR02204<br>TIGR02203<br>TIGR00958<br>TIGR01846<br>TIGR03796 | TIGR02204<br>TIGR02203<br>TIGR00958<br>TIGR01846<br>TIGR03796 | MsbA_rel: ABC transporter, permease/ATP-binding protein<br>MsbA_lipidA: lipid A export permease/ATP-binding protein MsbA<br>3a01208: antigen peptide transporter 2<br>type_I_sec_HlyB: type I secretion system ATPase<br>NHLM_micro_ABC1: NHLM bacteriocin system ABC transporter, peptidase/ATP-binding protein | 3.60E-102<br>2.20E-97<br>3.70E-82<br>5.00E-82<br>5.70E-82 |
| RKN10929.1 | 108120 | 107292 | - | 275 | PF13649<br>PF08241<br>PF13489<br>PF13847<br>PF08242 | Methyltransf_25<br>Methyltransf_11<br>Methyltransf_23<br>Methyltransf_31<br>Methyltransf_12 | Methyltransferase domain<br>Methyltransferase domain<br>Methyltransferase domain<br>Methyltransferase domain<br>Methyltransferase domain | 2.00E-16<br>2.10E-11<br>6.50E-11<br>2.00E-08<br>4.10E-08 |

| | | | | | | | | |
|---|---|---|---|---|---|---|---|---|
| RKN11095.1 | 108707 | 108116 | - | 196 | LinL | LinL | LinL | 1.20E-62 |
| RKN10930.1 | 110698 | 108727 | - | 656 | LinH<br>LinG<br>LinE<br>PF12697<br>PF00561 | LinH<br>LinG<br>LinE<br>Abhydrolase_6<br>Abhydrolase_1 | LinH (Fusion of LinE/LinG)<br>LinG Homolog<br>LinE Homolog<br>Alpha/beta hydrolase family<br>alpha/beta hydrolase fold | 5.00E-238<br>1.50E-114<br>1.80E-92<br>3.20E-16<br>2.30E-08 |
| RKN10931.1 | 111044 | 110843 | - | 66 | LinA | LinA | LinA (linaridin precursor) | 9.10E-09 |
| RKN10932.1 | 111268 | 111067 | - | 66 | LinA | LinA | LinA (linaridin precursor) | 1.60E-09 |
| RKN10933.1 | 111566 | 111374 | - | 63 | NO PFAM MATCH | - | - | - |
| RKN11096.1 | 111978 | 111792 | - | 61 | LinA | LinA | LinA (linaridin precursor) | 2.50E-10 |
| RKN10934.1 | 113172 | 112173 | - | 332 | TIGR01534<br>TIGR01532<br>PF02800<br>PF00044 | TIGR01534<br>TIGR01532<br>Gp_dh_C<br>Gp_dh_N | GAPDH-I: glyceraldehyde-3-phosphate dehydrogenase, type I<br>E4PD_g-proteo: erythrose-4-phosphate dehydrogenase<br>Glyceraldehyde 3-phosphate dehydrogenase, C-terminal domain<br>Glyceraldehyde 3-phosphate dehydrogenase, NAD binding domain | 8.50E-125<br>1.30E-94<br>4.50E-59<br>1.20E-35 |
| NP_631557.1 | 113272 | 113383 | + | 37 | INFERRED GENE | - | - | - |
| RKN10935.1 | 113527 | 113983 | + | 151 | PF05099<br>PF04391 | TerB<br>DUF533 | Tellurite resistance protein TerB<br>Protein of unknown function (DUF533) | 9.80E-31<br>3.50E-04 |

| index | leader | core | start | end | dir | score |
|---|---|---|---|---|---|---|
| 1 | MNDQSRRLYDSSALATVAVDDPRGWISARQKPVL | STPAVVAGLAIGYAMLNAFEAGRNGGPIRPAL | 111044 | 110843 | - | 28 |
| 2 | MSAMPVSQLPADLAAMSELKEGAAGRVPATAQPVM | ATPTIALAFAAGVGIGFAACQVVGDIRPVEN | 111268 | 111067 | - | 30 |
| 3 | MTVARLPADLAALPRLREDDAGQAPPT | LLPVTSTSCAAASAALGAARAVAWVADVIGPIRPSD | 111566 | 111374 | - | 8 |
| 4 | MSVARLPGDLAASPALREDDAGRPPVG | AVPVTSVLTSGQACVTVAIAITVGDTRPVLS | 111775 | 111598 | - | 9 |
| 5 | VTRKGRPMSVARLPGDLAASPALREDDAGRPPVG | AVPVTSVLTSGQACVTVAIAITVGDTRPVLS | 111796 | 111598 | - | 8 |
| 6 | MRRASVRIPQLDPGPLDEHAEGRVTRGALPVT | ATPVIVGAAGAIVIAYAAGRIVGEIRPTQ | 111978 | 111792 | - | 29 |
| 7 | MPTALAAWVAAGGAAVSTVSALNA | ATALVLILRAATPS | 114134 | 114017 | - | 7 |
| 8 | VPRRVAALLDEGPPARSWPPAHMPTALAAWVAAGGAAVSTVSALNA | ATALVLILRAATPS | 114200 | 114017 | - | 9 |

### Results for TCR48817.1 [Streptomyces sp. BK038] back to top

previous - next

#### Architecture

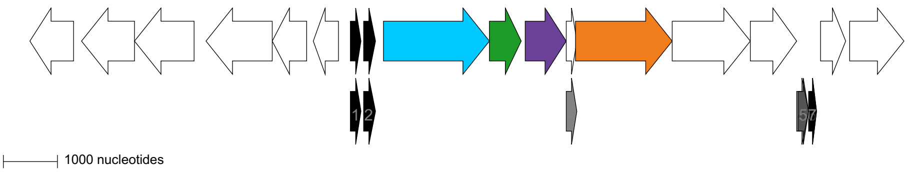

Link to nucleotide sequence

| Accession | start | end | direction | length (aa) | Pfam/HMM | name | description | E-value |
|---|---|---|---|---|---|---|---|---|
| TCR48809.1 | 527951 | 527141 | - | 269 | PF03704<br>PF00486 | BTAD<br>Trans_reg_C | Bacterial transcriptional activator domain<br>Transcriptional regulatory protein, C terminal | 2.10E-41<br>9.60E-11 |
| TCR48810.1 | 529074 | 528087 | - | 328 | PF00248<br>TIGR01293 | Aldo_ket_red<br>TIGR01293 | Aldo/keto reductase family<br>Kv_beta: voltage-dependent potassium channel beta subunit | 3.30E-75<br>3.10E-26 |
| TCR48811.1 | 530177 | 529070 | - | 368 | PF08028<br>TIGR04022<br>PF00441<br>PF02770 | Acyl-CoA_dh_2<br>TIGR04022<br>Acyl-CoA_dh_1<br>Acyl-CoA_dh_M | Acyl-CoA dehydrogenase, C-terminal domain<br>sulfur_SfnB: sulfur acquisition oxidoreductase, SfnB family<br>Acyl-CoA dehydrogenase, C-terminal domain<br>Acyl-CoA dehydrogenase, middle domain | 7.70E-27<br>3.50E-22<br>1.20E-05<br>2.60E-04 |
| TCR48812.1 | 531619 | 530413 | - | 401 | NO PFAM MATCH | - | - | - |
| TCR48813.1 | 532273 | 531631 | - | 213 | PF06114 | Peptidase_M78 | IrrE N-terminal-like domain | 6.40E-05 |
| TCR48814.1 | 532861 | 532375 | - | 161 | PF13560 | HTH_31 | Helix-turn-helix domain | 3.60E-04 |
| TCR48815.1 | 533080 | 533275 | + | 64 | LinA | LinA | LinA (linaridin precursor) | 1.20E-12 |
| TCR48816.1 | 533329 | 533530 | + | 66 | LinA | LinA | LinA (linaridin precursor) | 7.70E-11 |
| TCR48817.1 | 533677 | 535654 | + | 658 | LinH<br>LinG<br>LinE<br>PF12697<br>PF12146 | LinH<br>LinG<br>LinE<br>Abhydrolase_6<br>Hydrolase_4 | LinH (Fusion of LinE/LinG)<br>LinG Homolog<br>LinE Homolog<br>Alpha/beta hydrolase family<br>Serine aminopeptidase, S33 | 4.30E-252<br>1.80E-117<br>6.60E-101<br>2.50E-16<br>6.30E-10 |
| TCR48818.1 | 535650 | 536238 | + | 195 | LinL | LinL | LinL | 2.20E-70 |
| TCR48819.1 | 536300 | 537074 | + | 257 | PF13649<br>PF08241<br>PF08242<br>PF13847<br>PF13489 | Methyltransf_25<br>Methyltransf_11<br>Methyltransf_12<br>Methyltransf_31<br>Methyltransf_23 | Methyltransferase domain<br>Methyltransferase domain<br>Methyltransferase domain<br>Methyltransferase domain<br>Methyltransferase domain | 5.50E-14<br>3.40E-09<br>1.80E-08<br>4.00E-06<br>6.90E-06 |
| TCR48820.1 | 537075 | 537252 | + | 58 | NO PFAM MATCH | - | - | - |

| | | | | | | | | |
|---|---|---|---|---|---|---|---|---|
| TCR48821.1 | 537248 | 539021 | + | 590 | TIGR02204<br>TIGR02203<br>TIGR03796<br>TIGR00958<br>TIGR01846 | TIGR02204<br>TIGR02203<br>TIGR03796<br>TIGR00958<br>TIGR01846 | MsbA_rel: ABC transporter, permease/ATP-binding protein<br>MsbA_lipidA: lipid A export permease/ATP-binding protein MsbA<br>NHLM_micro_ABC1: NHLM bacteriocin system ABC transporter, peptidase/ATP-binding protein<br>3a01208: antigen peptide transporter 2<br>type_I_sec_HlyB: type I secretion system ATPase | 1.20E-103<br>1.10E-100<br>2.40E-85<br>6.80E-83<br>1.60E-82 |
| TCR48822.1 | 539017 | 540481 | + | 487 | TIGR02734<br>TIGR03467<br>PF13450<br>PF01593<br>TIGR01372 | TIGR02734<br>TIGR03467<br>NAD_binding_8<br>Amino_oxidase<br>TIGR01372 | crtI_fam: phytoene desaturase<br>HpnE: squalene-associated FAD-dependent desaturase<br>NAD(P)-binding Rossmann-like domain<br>Flavin containing amine oxidoreductase<br>soxA: sarcosine oxidase, alpha subunit family | 6.00E-12<br>2.30E-10<br>5.20E-10<br>1.00E-08<br>1.70E-08 |
| TCR48823.1 | 540477 | 541329 | + | 283 | PF01551 | Peptidase_M23 | Peptidase family M23 | 6.00E-18 |
| TCR48824.1 | 541757 | 542228 | + | 156 | TIGR02154<br>PF00072<br>TIGR02875<br>TIGR01818<br>TIGR01387 | TIGR02154<br>Response_reg<br>TIGR02875<br>TIGR01818<br>TIGR01387 | PhoB: phosphate regulon transcriptional regulatory protein PhoB<br>Response regulator receiver domain<br>spore_0_A: sporulation transcription factor Spo0A<br>ntrC: nitrogen regulation protein NR(I)<br>cztR_silR_copR: heavy metal response regulator | 2.90E-25<br>4.10E-24<br>9.30E-19<br>4.30E-17<br>2.00E-16 |
| TCR48825.1 | 542305 | 543310 | + | 334 | TIGR00456<br>PF05746<br>PF03485 | TIGR00456<br>DALR_1<br>Arg_tRNA_synt_N | argS: arginine--tRNA ligase<br>DALR anticodon binding domain<br>Arginyl tRNA synthetase N terminal domain | 4.00E-22<br>5.60E-18<br>9.20E-15 |

| index | leader | core | start | end | dir | score |
|---|---|---|---|---|---|---|
| 1 | MAKIAAFMNQQDMGPIAESELSVGGTPVM | ATPAVVAVPASVKITAGIVAFTIPVGAYVTGRVVN | 533080 | 533275 | + | 30 |
| 2 | MHSITESVRVIAGATLSEDSGEQTTRMMPVA | FTPAVLVTTPYAAGGLAAGAAAVGAYEAGRAAAKG | 533329 | 533530 | + | 29 |
| 3 | MSTQRAVWFVAALAVAVPVM | AVMFREDGRFTSQSWTKGLIFGTAVAVVAAIAAGRARQ | 537075 | 537252 | + | 7 |
| 4 | VTSVLPGPAVHGAENQSGGNCRRGR...RSPAPAADHGLKNRCAEETRRVPLP | RPPTYGCTYAKTPSQRRR | 541325 | 541550 | + | 6 |
| 5 | VHGAENQSGGNCRRGRRGRCSPRSPAPAADHGLKNRCAEETRRVPLP | RPPTYGCTYAKTPSQRRR | 541352 | 541550 | + | 7 |
| 6 | MAGSLGTPCEVPPAGFEPAH | TAPEAVALSPELRGRVVRVAATGGTLPASAG | 541549 | 541705 | + | 10 |
| 7 | LGTPCEVPPAGFEPAH | TAPEAVALSPELRGRVVRVAATGGTLPASAG | 541561 | 541705 | + | 8 |

### Results for WP_073790447.1 [Streptomyces sp. CB01580] back to top

previous - next

#### Architecture

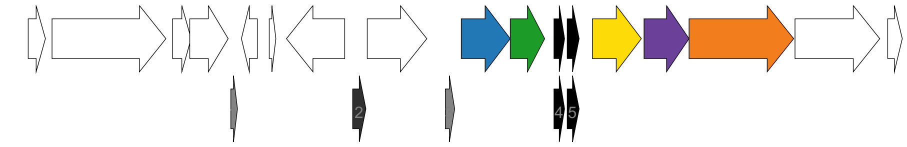

1000 nucleotides

Link to nucleotide sequence

| Accession | start | end | direction | length (aa) | Pfam/HMM | name | description | E-value |
|---|---|---|---|---|---|---|---|---|
| WP_010069772.1 | 90670 | 90952 | + | 94 | INFERRED GENE | - | - | - |
| WP_005477252.1 | 91074 | 93027 | + | 651 | INFERRED GENE | - | - | - |
| WP_010981551.1 | 93146 | 93452 | + | 102 | INFERRED GENE | - | - | - |
| WP_005477251.1 | 93440 | 94106 | + | 222 | INFERRED GENE | - | - | - |
| WP_107430674.1 | 94606 | 94318 | - | 95 | NO PFAM MATCH | - | - | - |
| WP_012948683.1 | 94807 | 94927 | + | 40 | INFERRED GENE | - | - | - |
| WP_073790445.1 | 96114 | 95091 | - | 340 | PF12840<br>PF12802<br>PF01022 | HTH_20<br>MarR_2<br>HTH_5 | Helix-turn-helix domain<br>MarR family<br>Bacterial regulatory protein, arsR family | 3.30E-10<br>2.70E-06<br>2.20E-04 |
| WP_073790491.1 | 96498 | 97521 | + | 340 | PF14525<br>PF12833<br>PF00165<br>TIGR04094<br>TIGR02297 | AraC_binding_2<br>HTH_18<br>HTH_AraC<br>TIGR04094<br>TIGR02297 | AraC-binding-like domain<br>Helix-turn-helix domain<br>Bacterial regulatory helix-turn-helix proteins, AraC family<br>adjacent_YSIRK: YSIRK-targeted surface antigen transcriptional regulator<br>HpaA: 4-hydroxyphenylacetate catabolism regulatory protein HpaA | 7.50E-25<br>2.40E-24<br>3.00E-13<br>1.30E-10<br>7.30E-06 |
| WP_073790447.1 | 98108 | 98954 | + | 281 | LinE<br>LinH<br>PF12697<br>PF00561<br>PF12146 | LinE<br>LinH<br>Abhydrolase_6<br>Abhydrolase_1<br>Hydrolase_4 | LinE Homolog<br>LinH (Fusion of LinE/LinG)<br>Alpha/beta hydrolase family<br>alpha/beta hydrolase fold<br>Serine aminopeptidase, S33 | 6.50E-84<br>1.20E-37<br>6.70E-15<br>2.90E-08<br>4.20E-07 |
| WP_073790449.1 | 98950 | 99538 | + | 195 | LinL | LinL | LinL | 3.50E-58 |
| WP_073790451.1 | 99696 | 99897 | + | 66 | LinA | LinA | LinA (linaridin precursor) | 2.80E-12 |
| WP_073790453.1 | 99931 | 100132 | + | 66 | LinA | LinA | LinA (linaridin precursor) | 2.50E-08 |
| WP_073790493.1 | 100360 | 101197 | + | 278 | LinG<br>LinH | LinG<br>LinH | LinG Homolog<br>LinH (Fusion of LinE/LinG) | 3.40E-100<br>3.90E-74 |
| WP_073790455.1 | 101250 | 102036 | + | 261 | PF13649<br>PF08241<br>PF13489<br>PF13847<br>PF08242 | Methyltransf_25<br>Methyltransf_11<br>Methyltransf_23<br>Methyltransf_31<br>Methyltransf_12 | Methyltransferase domain<br>Methyltransferase domain<br>Methyltransferase domain<br>Methyltransferase domain<br>Methyltransferase domain | 2.80E-12<br>1.50E-08<br>5.10E-08<br>2.10E-07<br>1.50E-05 |

| | | | | | | | | |
|---|---|---|---|---|---|---|---|---|
| WP_073790457.1 | 102032 | 103823 | + | 596 | TIGR02203<br>TIGR02204<br>TIGR00958<br>TIGR03796<br>TIGR01846 | TIGR02203<br>TIGR02204<br>TIGR00958<br>TIGR03796<br>TIGR01846 | MsbA_lipidA: lipid A export permease/ATP-binding protein MsbA<br>MsbA_rel: ABC transporter, permease/ATP-binding protein<br>3a01208: antigen peptide transporter 2<br>NHLM_micro_ABC1: NHLM bacteriocin system ABC transporter, peptidase/ATP-binding protein<br>type_I_sec_HlyB: type I secretion system ATPase | 3.20E-98<br>4.30E-96<br>4.80E-84<br>4.90E-84<br>2.10E-80 |
| WP_107430676.1 | 103855 | 105307 | + | 483 | TIGR02734<br>TIGR03467<br>TIGR02733<br>TIGR00292<br>TIGR01317 | TIGR02734<br>TIGR03467<br>TIGR02733<br>TIGR00292<br>TIGR01317 | crtI_fam: phytoene desaturase<br>HpnE: squalene-associated FAD-dependent desaturase<br>desat_CrtD: C-3',4' desaturase CrtD<br>TIGR00292: thiazole biosynthesis enzyme<br>GOGAT_sm_gam: glutamate synthase, NADH/NADPH, small subunit | 4.80E-12<br>1.20E-09<br>1.10E-08<br>4.50E-08<br>7.70E-08 |
| WP_005477326.1 | 105448 | 105706 | + | 86 | INFERRED GENE | - | - | - |

| index | leader | core | start | end | dir | score |
|---|---|---|---|---|---|---|
| 1 | MSRCPTAPFR | AAPHAHAAHPSHLTGAPTTPDTFHGF | 94155 | 94266 | + | 7 |
| 2 | MADRQGGLEDRLAGLARGPCHRHAAPDHSAHQQQTVRR | TAHQSGWTVTSLSGTALVPLALHAGRAFLAPVQRGRPTT | 96230 | 96464 | + | 9 |
| 3 | LHRPGSNQQARCRQRCAAHAGWAHL | SGLSPLTEPETGPTRPTRDQRGRSSR | 97845 | 98001 | + | 7 |
| 4 | MSLVSEFANETLTDVAPGMLSATDVATRLDTPADPVM | ATPLIVETVLLTTGWIANNIANYVDRVNY | 99696 | 99897 | + | 31 |
| 5 | MSLIAEFANETLTGVTPGLLSETDSIMCPDAPADPTM | STAFLTVNTCLDGLYTRTALTALIVVAAG | 99931 | 100132 | + | 31 |

### Results for OKJ59748.1 [Streptomyces sp. CB02009] back to top

previous - next

#### Architecture

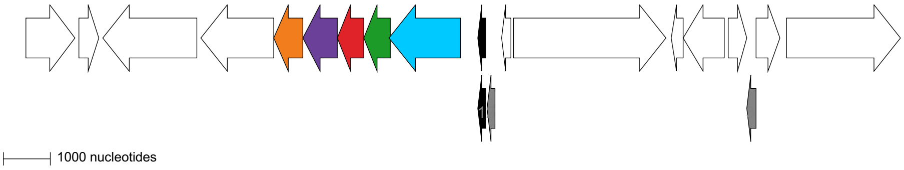

Link to nucleotide sequence

| Accession | start | end | direction | length (aa) | Pfam/HMM | name | description | E-value |
|---|---|---|---|---|---|---|---|---|
| OKJ59740.1 | 146554 | 147595 | + | 346 | TIGR00350<br>PF03816 | TIGR00350<br>LytR_cpsA_psr | lytR_cpsA_psr: cell envelope-related function transcriptional attenuator common domain<br>Cell envelope-related transcriptional attenuator domain | 6.30E-38<br>1.90E-37 |
| OKJ59741.1 | 147689 | 148097 | + | 135 | NO PFAM MATCH | - | - | - |
| OKJ59742.1 | 150195 | 148173 | - | 673 | PF03190<br>PF13899<br>PF07221 | Thioredox_DsbH<br>Thioredoxin_7<br>GlcNAc_2-epim | Protein of unknown function, DUF255<br>Thioredoxin-like<br>N-acylglucosamine 2-epimerase (GlcNAc 2-epimerase) | 8.20E-75<br>9.80E-11<br>4.40E-04 |
| OKJ59743.1 | 151825 | 150277 | - | 515 | NO PFAM MATCH | - | - | - |
| OKJ59744.1 | 152454 | 151821 | - | 210 | TIGR02857<br>TIGR02673<br>TIGR01189<br>TIGR03873<br>TIGR03265 | TIGR02857<br>TIGR02673<br>TIGR01189<br>TIGR03873<br>TIGR03265 | CydD: thiol reductant ABC exporter, CydD subunit<br>FtsE: cell division ATP-binding protein FtsE<br>ccmA: heme ABC exporter, ATP-binding protein CcmA<br>F420-0_ABC_ATP: proposed F420-0 ABC transporter, ATP-binding protein<br>PhnT2: putative 2-aminoethylphosphonate ABC transporter, ATP-binding protein | 1.40E-28<br>4.90E-27<br>1.40E-26<br>9.60E-25<br>4.80E-23 |
| OKJ59745.1 | 153188 | 152450 | - | 245 | PF13649<br>PF08241<br>TIGR02021<br>PF08242<br>TIGR01934 | Methyltransf_25<br>Methyltransf_11<br>TIGR02021<br>Methyltransf_12<br>TIGR01934 | Methyltransferase domain<br>Methyltransferase domain<br>BchM-ChlM: magnesium protoporphyrin O-methyltransferase<br>Methyltransferase domain<br>MenG_MenH_UbiE: ubiquinone/menaquinone biosynthesis methyltransferase | 5.70E-18<br>2.30E-16<br>2.00E-12<br>4.00E-12<br>8.30E-12 |
| OKJ59746.1 | 153753 | 153180 | - | 190 | TIGR00521<br>TIGR02113<br>PF02441 | TIGR00521<br>TIGR02113<br>Flavoprotein | coaBC_dfp: phosphopantothenoylcysteine decarboxylase / phosphopantothenate--cysteine ligase<br>coaC_strep: phosphopantothenoylcysteine decarboxylase<br>Flavoprotein | 9.50E-11<br>2.20E-09<br>1.60E-08 |
| OKJ59747.1 | 154310 | 153749 | - | 186 | LinL | LinL | LinL | 1.70E-53 |
| OKJ59748.1 | 155817 | 154299 | - | 505 | LinH<br>LinE<br>LinG<br>PF12697<br>TIGR02427 | LinH<br>LinE<br>LinG<br>Abhydrolase_6<br>TIGR02427 | LinH (Fusion of LinE/LinG)<br>LinE Homolog<br>LinG Homolog<br>Alpha/beta hydrolase family<br>protocat_pcaD: 3-oxoadipate enol-lactonase | 1.90E-160<br>1.00E-68<br>8.40E-67<br>3.30E-09<br>1.30E-07 |
| OKJ59749.1 | 156351 | 156168 | - | 60 | LinA | LinA | LinA (linaridin precursor) | 5.60E-14 |
| OKJ59750.1 | 156877 | 156691 | - | 61 | NO PFAM MATCH | - | - | - |
| OKJ59751.1 | 156939 | 160197 | + | 1085 | PF13424<br>PF17874<br>PF07719<br>PF13374<br>PF00931 | TPR_12<br>TPR_MalT<br>TPR_2<br>TPR_10<br>NB-ARC | Tetratricopeptide repeat<br>MalT-like TPR region<br>Tetratricopeptide repeat<br>Tetratricopeptide repeat<br>NB-ARC domain | 6.90E-21<br>1.40E-11<br>1.20E-09<br>9.30E-09<br>7.50E-08 |
| OKJ59752.1 | 160547 | 160310 | - | 78 | NO PFAM MATCH | - | - | - |
| OKJ59753.1 | 161430 | 160548 | - | 293 | TIGR03446<br>TIGR03445<br>PF02585<br>TIGR04001<br>TIGR04000 | TIGR03446<br>TIGR03445<br>PIG-L<br>TIGR04001<br>TIGR04000 | mycothiol_Mca: mycothiol conjugate amidase Mca<br>mycothiol_MshB: N-acetyl-1-D-myo-inositol-2-amino-2-deoxy-alpha-D-glucopyranoside deacetylase<br>GlcNAc-PI de-N-acetylase<br>thiol_BshB1: bacillithiol biosynthesis deacetylase BshB1<br>thiol_BshB2: bacillithiol biosynthesis deacetylase BshB2 | 2.10E-146<br>1.00E-51<br>3.30E-25<br>2.60E-13<br>4.60E-10 |
| OKJ59754.1 | 161528 | 161930 | + | 133 | PF14155 | DUF4307 | Domain of unknown function (DUF4307) | 1.60E-31 |

| | | | | | | | | |
|---|---|---|---|---|---|---|---|---|
| OKJ59755.1 | 162119 | 162617 | + | 165 | TIGR01462<br>PF03449<br>TIGR01461<br>PF01272 | TIGR01462<br>GreA_GreB_N<br>TIGR01461<br>GreA_GreB | greA: transcription elongation factor GreA<br>Transcription elongation factor, N-terminal<br>greB: transcription elongation factor GreB<br>Transcription elongation factor, GreA/GreB, C-term | 7.30E-45<br>7.70E-26<br>3.80E-16<br>9.30E-15 |
| OKJ59756.1 | 162760 | 165199 | + | 812 | PF00069<br>TIGR03903<br>PF07714<br>PF13360<br>TIGR03724 | Pkinase<br>TIGR03903<br>Pkinase_Tyr<br>PQQ_2<br>TIGR03724 | Protein kinase domain<br>TOMM_kin_cyc: TOMM system kinase/cyclase fusion protein<br>Protein tyrosine kinase<br>PQQ-like domain<br>arch_bud32: Kae1-associated kinase Bud32 | 1.00E-33<br>7.00E-26<br>1.40E-20<br>1.60E-19<br>8.10E-13 |

| index | leader | core | start | end | dir | score |
|---|---|---|---|---|---|---|
| 1 | MTLTSSNSAEALAAQDFANTVLSAAGPGFHADCETPAM | ATPATPTVAQFVVQGSTICLVC | 156351 | 156168 | - | 36 |
| 2 | VLYAEVAGSIALDEFLKVCLRSVRSFQL | QTLCSAFTQSGRRGWQRSIRLDHSLA | 156550 | 156385 | - | 7 |
| 3 | VLLVGTGNTKHRPTRESSAVPGGRNHEPNNEWRK | AGGHNHQDGVAPGHRGDAPGLDTRSLPTATEPYTP | 162117 | 161907 | - | 7 |

### Results for WP_148648000.1 [Streptomyces sp. CB01881] back to top

previous - next

#### Architecture

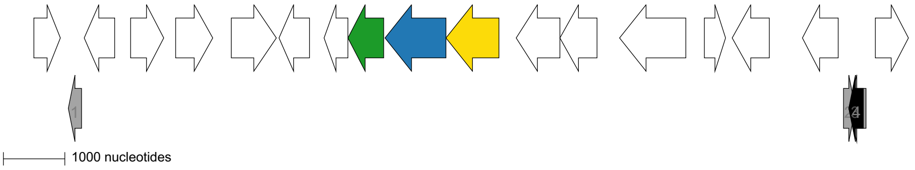

Link to nucleotide sequence

| Accession | start | end | direction | length (aa) | Pfam/HMM | name | description | E-value |
|---|---|---|---|---|---|---|---|---|
| WP_148647994.1 | 12543 | 12981 | + | 145 | PF00293<br>TIGR02705<br>TIGR00586<br>TIGR00052 | NUDIX<br>TIGR02705<br>TIGR00586<br>TIGR00052 | NUDIX domain<br>nudix_YtkD: nucleoside triphosphatase YtkD<br>mutt: mutator mutT protein<br>TIGR00052: nudix-type nucleoside diphosphatase, YffH/AdpP family | 1.50E-15<br>2.00E-08<br>1.80E-06<br>2.40E-06 |
| WP_148648068.1 | 13863 | 13380 | - | 160 | NO PFAM MATCH | - | - | - |
| WP_148647995.1 | 14128 | 14653 | + | 174 | PF08877 | MepB | MepB protein | 3.90E-50 |
| WP_020463643.1 | 14850 | 15469 | + | 206 | INFERRED GENE | - | - | - |
| WP_148647996.1 | 15758 | 16487 | + | 242 | PF00106<br>PF13561<br>TIGR03206<br>TIGR02632<br>TIGR01963 | adh_short<br>adh_short_C2<br>TIGR03206<br>TIGR02632<br>TIGR01963 | short chain dehydrogenase<br>Enoyl-(Acyl carrier protein) reductase<br>benzo_BadH: 2-hydroxycyclohexanecarboxyl-CoA dehydrogenase<br>RhaD_aldol-ADH: rhamnulose-1-phosphate aldolase/alcohol dehydrogenase<br>PHB_DH: 3-hydroxybutyrate dehydrogenase | 4.80E-33<br>4.10E-23<br>2.80E-17<br>7.20E-17<br>9.00E-17 |
| WP_148647997.1 | 17038 | 16546 | - | 163 | NO PFAM MATCH | - | - | - |
| WP_148647998.1 | 17654 | 17267 | - | 128 | NO PFAM MATCH | - | - | - |
| WP_148647999.1 | 18244 | 17659 | - | 194 | LinL | LinL | LinL | 5.60E-47 |
| WP_148648000.1 | 19257 | 18273 | - | 327 | LinE<br>LinH<br>TIGR02427<br>PF12697<br>TIGR03611 | LinE<br>LinH<br>TIGR02427<br>Abhydrolase_6<br>TIGR03611 | LinE Homolog<br>LinH (Fusion of LinE/LinG)<br>protocat_pcaD: 3-oxoadipate enol-lactonase<br>Alpha/beta hydrolase family<br>RutD: pyrimidine utilization protein D | 2.80E-81<br>1.70E-72<br>2.60E-13<br>5.50E-13<br>3.10E-08 |
| WP_148648001.1 | 20120 | 19253 | - | 288 | LinG<br>LinH | LinG<br>LinH | LinG Homolog<br>LinH (Fusion of LinE/LinG) | 4.40E-91<br>7.70E-87 |
| WP_162002685.1 | 21109 | 20401 | - | 235 | NO PFAM MATCH | - | - | - |
| WP_148648003.1 | 21726 | 21108 | - | 205 | TIGR02983<br>TIGR02937<br>TIGR02985<br>PF08281<br>TIGR02954 | TIGR02983<br>TIGR02937<br>TIGR02985<br>Sigma70_r4_2<br>TIGR02954 | SigE-fam_strep: RNA polymerase sigma-70 factor, sigma-E family<br>sigma70-ECF: RNA polymerase sigma factor, sigma-70 family<br>Sig70_bacteroi1: RNA polymerase sigma-70 factor, Bacteroides expansion family 1<br>Sigma-70, region 4<br>Sig70_famx3: RNA polymerase sigma-70 factor, TIGR02954 family | 2.30E-58<br>4.50E-23<br>1.30E-20<br>4.40E-18<br>2.00E-17 |
| WP_148648004.1 | 23169 | 22086 | - | 360 | PF03747 | ADP_ribosyl_GH | ADP-ribosylglycohydrolase | 8.40E-13 |
| WP_148648005.1 | 23471 | 23816 | + | 114 | NO PFAM MATCH | - | - | - |
| WP_148648006.1 | 24522 | 23925 | - | 198 | NO PFAM MATCH | - | - | - |
| WP_148648007.1 | 25643 | 25049 | - | 197 | PF01764 | Lipase_3 | Lipase (class 3) | 2.30E-24 |
| WP_009454356.1 | 26247 | 26778 | + | 177 | INFERRED GENE | - | - | - |

| index | leader | core | start | end | dir | score |
|---|---|---|---|---|---|---|
| 1 | LNDEARACSPRPGQRSGQAGD | RTTRASRQPPRTREVSQVPATGSSASAGAAARSLCTAPERAGLPPPHRR | 13326 | 13113 | - | 6 |
| 2 | LAAIACGTPERPPHPRQTHRIDGIQLEVVRIPH | AAMTSATTPSVVPSAPITLFVTLIRSALTRASA | 25723 | 25924 | + | 6 |
| 3 | VYAFQHTGHVAGTFEPGSRNDDWVIDSRSDQIADNWANI | AANPTAHAEARVNADLINVTNSVIGALGTTLGVVALVIAA | 26058 | 25818 | - | 16 |
| 4 | VWGVKDAQNKVYAFQHTGHVAGTFEPGSRNDDWVIDSRSDQIAD | NWANIAANPTAHAEARVNADLINVTNSVIGALGTTLGVVALVIAA | 26088 | 25818 | - | 6 |

### Results for OKI92952.1 [Streptomyces sp. CB02058] back to top

#### Architecture

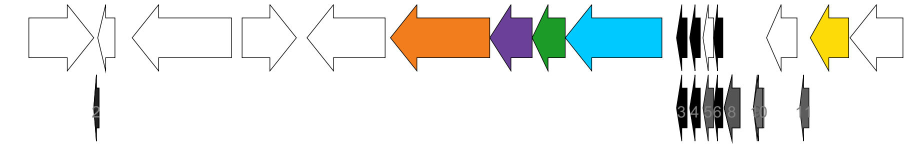

1000 nucleotides

Link to nucleotide sequence

| Accession | start | end | direction | length (aa) | Pfam/HMM | name | description | E-value |
|---|---|---|---|---|---|---|---|---|
| OKI92710.1 | 737054 | 738275 | + | 406 | PF00149<br>PF12850 | Metallophos<br>Metallophos_2 | Calcineurin-like phosphoesterase<br>Calcineurin-like phosphoesterase superfamily domain | 3.40E-08<br>1.60E-05 |
| OKI92711.1 | 738666 | 738357 | - | 102 | NO PFAM MATCH | - | - | - |
| OKI92712.1 | 740862 | 739005 | - | 618 | NO PFAM MATCH | - | - | - |
| OKI92713.1 | 741066 | 742071 | + | 334 | NO PFAM MATCH | - | - | - |
| OKI92714.1 | 743747 | 742262 | - | 494 | TIGR03467<br>TIGR02734<br>PF13450<br>TIGR02733<br>TIGR01317 | TIGR03467<br>TIGR02734<br>NAD_binding_8<br>TIGR02733<br>TIGR01317 | HpnE: squalene-associated FAD-dependent desaturase<br>crtI_fam: phytoene desaturase<br>NAD(P)-binding Rossmann-like domain<br>desat_CrtD: C-3',4' desaturase CrtD<br>GOGAT_sm_gam: glutamate synthase, NADH/NADPH, small subunit | 6.20E-09<br>8.20E-09<br>1.00E-08<br>1.50E-07<br>1.60E-07 |
| OKI92715.1 | 745698 | 743844 | - | 617 | TIGR02204<br>TIGR02203<br>TIGR00958<br>TIGR03796<br>TIGR01846 | TIGR02204<br>TIGR02203<br>TIGR00958<br>TIGR03796<br>TIGR01846 | MsbA_rel: ABC transporter, permease/ATP-binding protein<br>MsbA_lipidA: lipid A export permease/ATP-binding protein MsbA<br>3a01208: antigen peptide transporter 2<br>NHLM_micro_ABC1: NHLM bacteriocin system ABC transporter, peptidase/ATP-binding protein<br>type_I_sec_HlyB: type I secretion system ATPase | 4.40E-101<br>1.90E-92<br>6.80E-87<br>1.10E-86<br>1.00E-80 |
| OKI92716.1 | 746501 | 745694 | - | 268 | PF13649<br>PF08241<br>PF13489<br>PF13847<br>TIGR01983 | Methyltransf_25<br>Methyltransf_11<br>Methyltransf_23<br>Methyltransf_31<br>TIGR01983 | Methyltransferase domain<br>Methyltransferase domain<br>Methyltransferase domain<br>Methyltransferase domain<br>UbiG: 3-demethylubiquinone-9 3-O-methyltransferase | 9.60E-16<br>4.60E-11<br>1.30E-08<br>3.00E-06<br>3.80E-06 |
| OKI92717.1 | 747106 | 746497 | - | 202 | LinL | LinL | LinL | 1.10E-60 |
| OKI92952.1 | 748926 | 747102 | - | 607 | LinH<br>LinG<br>LinE<br>PF12697<br>PF00561 | LinH<br>LinG<br>LinE<br>Abhydrolase_6<br>Abhydrolase_1 | LinH (Fusion of LinE/LinG)<br>LinG Homolog<br>LinE Homolog<br>Alpha/beta hydrolase family<br>alpha/beta hydrolase fold | 1.70E-234<br>1.00E-112<br>1.10E-88<br>2.20E-14<br>5.40E-11 |
| OKI92718.1 | 749402 | 749186 | - | 71 | LinA | LinA | LinA (linaridin precursor) | 1.30E-09 |
| OKI92719.1 | 749640 | 749445 | - | 64 | LinA | LinA | LinA (linaridin precursor) | 2.10E-09 |
| OKI92720.1 | 749885 | 749690 | - | 64 | NO PFAM MATCH | - | - | - |
| OKI92721.1 | 750067 | 749881 | - | 61 | LinA | LinA | LinA (linaridin precursor) | 1.60E-08 |
| OKI92722.1 | 751459 | 750874 | - | 194 | PF00210 | Ferritin | Ferritin-like domain | 9.30E-29 |
| OKI92723.1 | 752414 | 751691 | - | 240 | PF12697<br>LinH<br>PF12146<br>LinE<br>TIGR03056 | Abhydrolase_6<br>LinH<br>Hydrolase_4<br>LinE<br>TIGR03056 | Alpha/beta hydrolase family<br>LinH (Fusion of LinE/LinG)<br>Serine aminopeptidase, S33<br>LinE Homolog<br>bchO_mg_che_rel: putative magnesium chelatase accessory protein | 1.90E-22<br>6.80E-08<br>7.10E-08<br>3.90E-07<br>1.20E-06 |
| OKI92724.1 | 753427 | 752455 | - | 323 | PF07859<br>PF00326 | Abhydrolase_3<br>Peptidase_S9 | alpha/beta hydrolase fold<br>Prolyl oligopeptidase family | 1.10E-69<br>3.50E-06 |

| index | leader | core | start | end | dir | score |
|---|---|---|---|---|---|---|
| 1 | MPAPLTGRGQAVQAGDAGH | HAPAHHVPAGTTSA | 738373 | 738271 | - | 8 |
| 2 | VGMPAPLTGRGQAVQAGDAGH | HAPAHHVPAGTTSA | 738379 | 738271 | - | 8 |
| 3 | MNDVSFLTRPGALTDIKVDDPQGRVIITQTPVM | ATPAVVAGVIIGAALVNAFVAGRDGGPLEPPIDVSVPS | 749402 | 749186 | - | 29 |
| 4 | MNAAVFQAVPAAARPLTEKDPGHTATGTAPVA | ATPVIVVTAFAAGVAAGVALCDAIGDTSPVEQ | 749640 | 749445 | - | 30 |
| 5 | MTGARWATASVCP | ATAVTTRALSESDAGYTGPAASPIA...VQIVVAEFVVGPAIDGTGGTHEPQL | 749885 | 749690 | - | 8 |
| 6 | MQQVRTPGWAAVGPLDERTEGQAFGGAAPIA | ATQVIVTVAAAVVIAYAAGAIIGSTTEPSL | 750067 | 749881 | - | 20 |
| 7 | MEPAGQCGIDLRPARDLLDDRSCTV...RQRRACTILRAALQPRASRRPSPRP | VAPPSLTSQTDRRPAGPLGVRGAALPYGPVAAASGCRPG | 750378 | 750081 | - | 7 |
| 8 | VGRPTPMEPAGQCGIDLRPARDLLD...RQRRACTILRAALQPRASRRPSPRP | VAPPSLTSQTDRRPAGPLGVRGAALPYGPVAAASGCRPG | 750396 | 750081 | - | 6 |
| 9 | VPWHALPHTPSG | GTPTPRCRRSSPSSRLRSSRGDPNRSASTPSCALRSHAPDPAP | 750805 | 750637 | - | 6 |
| 10 | LDLGGLACPGTHSPTLRPGVPPRRVVVGLPRALGFARAGETPTAPR | RLPPAPCDRTHQTPLPDPP | 750825 | 750627 | - | 6 |
| 11 | VSCRAADDPAARHPKPCARTRSGSVRTASACPET | AAPRCARTVISACAAGQRGTYAST | 751681 | 751504 | - | 8 |

### Results for WP_073756736.1 [Streptomyces sp. CB03234] back to top

#### Architecture

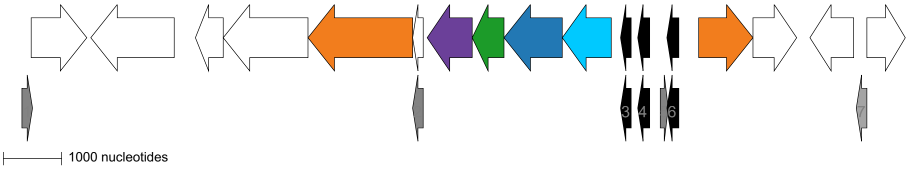

Link to nucleotide sequence

| Accession | start | end | direction | length (aa) | Pfam/HMM | name | description | E-value |
|---|---|---|---|---|---|---|---|---|
| WP_073756729.1 | 519447 | 520389 | + | 313 | PF01370<br>TIGR03466<br>PF16363<br>PF13460<br>TIGR02032 | Epimerase<br>TIGR03466<br>GDP_Man_Dehyd<br>NAD_binding_10<br>TIGR02032 | NAD dependent epimerase/dehydratase family<br>HpnA: hopanoid-associated sugar epimerase<br>GDP-mannose 4,6 dehydratase<br>NAD(P)H-binding<br>GG-red-SF: geranylgeranyl reductase family | 7.10E-14<br>5.30E-06<br>1.50E-05<br>4.10E-05<br>1.70E-04 |
| WP_073756730.1 | 521893 | 520465 | - | 475 | PF13360 | PQQ_2 | PQQ-like domain | 2.00E-04 |
| WP_073756731.1 | 522725 | 522245 | - | 159 | NO PFAM MATCH | - | - | - |
| WP_073756732.1 | 524191 | 522724 | - | 488 | PF01593<br>TIGR02734<br>PF13450<br>TIGR03467<br>TIGR01317 | Amino_oxidase<br>TIGR02734<br>NAD_binding_8<br>TIGR03467<br>TIGR01317 | Flavin containing amine oxidoreductase<br>crtI_fam: phytoene desaturase<br>NAD(P)-binding Rossmann-like domain<br>HpnE: squalene-associated FAD-dependent desaturase<br>GOGAT_sm_gam: glutamate synthase, NADH/NADPH, small subunit | 1.00E-10<br>1.30E-10<br>2.60E-10<br>2.60E-09<br>6.50E-08 |
| WP_079277773.1 | 525981 | 524187 | - | 597 | TIGR02204<br>TIGR02203<br>TIGR01846<br>TIGR03375<br>TIGR00958 | TIGR02204<br>TIGR02203<br>TIGR01846<br>TIGR03375<br>TIGR00958 | MsbA_rel: ABC transporter, permease/ATP-binding protein<br>MsbA_lipidA: lipid A export permease/ATP-binding protein MsbA<br>type_I_sec_HlyB: type I secretion system ATPase<br>type_I_sec_LssB: type I secretion system ATPase<br>3a01208: antigen peptide transporter 2 | 6.20E-105<br>7.30E-105<br>2.20E-86<br>1.70E-84<br>2.90E-81 |
| WP_073756733.1 | 526157 | 525977 | - | 59 | NO PFAM MATCH | - | - | - |
| WP_073756734.1 | 527010 | 526236 | - | 257 | PF13649<br>PF08241<br>PF13489<br>PF08242<br>PF13847 | Methyltransf_25<br>Methyltransf_11<br>Methyltransf_23<br>Methyltransf_12<br>Methyltransf_31 | Methyltransferase domain<br>Methyltransferase domain<br>Methyltransferase domain<br>Methyltransferase domain<br>Methyltransferase domain | 3.20E-14<br>2.50E-10<br>2.10E-09<br>5.10E-07<br>1.10E-05 |
| WP_158071144.1 | 527552 | 527006 | - | 181 | LinL | LinL | LinL | 5.70E-64 |
| WP_073756736.1 | 528565 | 527548 | - | 338 | LinE<br>LinH<br>PF12697<br>TIGR02427<br>PF00561 | LinE<br>LinH<br>Abhydrolase_6<br>TIGR02427<br>Abhydrolase_1 | LinE Homolog<br>LinH (Fusion of LinE/LinG)<br>Alpha/beta hydrolase family<br>protocat_pcaD: 3-oxoadipate enol-lactonase<br>alpha/beta hydrolase fold | 2.10E-125<br>5.30E-79<br>2.20E-14<br>1.80E-11<br>7.30E-10 |
| WP_073756737.1 | 529398 | 528561 | - | 278 | LinH<br>LinG | LinH<br>LinG | LinH (Fusion of LinE/LinG)<br>LinG Homolog | 1.90E-105<br>6.00E-105 |
| WP_073756738.1 | 529745 | 529565 | - | 59 | LinA | LinA | LinA (linaridin precursor) | 2.70E-09 |
| WP_073756739.1 | 530051 | 529850 | - | 66 | LinA | LinA | LinA (linaridin precursor) | 3.90E-11 |
| WP_073756740.1 | 530546 | 530363 | - | 60 | LinA | LinA | LinA (linaridin precursor) | 2.00E-10 |
| WP_073756741.1 | 530891 | 531824 | + | 310 | TIGR01188<br>TIGR03864<br>TIGR01288<br>TIGR04406<br>TIGR02673 | TIGR01188<br>TIGR03864<br>TIGR01288<br>TIGR04406<br>TIGR02673 | drrA: daunorubicin resistance ABC transporter, ATP-binding protein<br>PQQ_ABC_ATP: ABC transporter, ATP-binding subunit, PQQ-dependent alcohol dehydrogenase system<br>nodI: nodulation ABC transporter NodI<br>LPS_export_lptB: LPS export ABC transporter ATP-binding protein<br>FtsE: cell division ATP-binding protein FtsE | 3.70E-60<br>1.20E-43<br>2.70E-41<br>1.60E-33<br>1.20E-32 |
| WP_073757084.1 | 531835 | 532585 | + | 249 | PF01061<br>PF12698 | ABC2_membrane<br>ABC2_membrane_3 | ABC-2 type transporter<br>ABC-2 family transporter protein | 9.20E-16<br>1.10E-05 |
| WP_073756742.1 | 533562 | 532809 | - | 250 | PF13936<br>PF12802<br>PF01047<br>PF12840<br>PF13463 | HTH_38<br>MarR_2<br>MarR<br>HTH_20<br>HTH_27 | Helix-turn-helix domain<br>MarR family<br>MarR family<br>Helix-turn-helix domain<br>Winged helix DNA-binding domain | 1.90E-14<br>2.50E-13<br>8.90E-09<br>2.10E-07<br>3.10E-06 |
| WP_073756743.1 | 533788 | 534457 | + | 222 | PF13349 | DUF4097 | Putative adhesin | 3.30E-08 |

| index | leader | core | start | end | dir | score |
|---|---|---|---|---|---|---|
| 1 | VPALGGSTEAPTWCVVEG | LTPAWRWTAAPGSCCHVISASNCPSPPTVPTARASVRA | 519280 | 519451 | + | 7 |
| 2 | MLTASRLTWFVIAFAFALP | STLVMFRDNGVVTRDAWVKSFVFAAAVAAVIAVVFGKGSQ | 526157 | 525977 | - | 7 |
| 3 | MSVLVEKVNAGAGQAADGSVAPVM | ATPAAFVAGVVAGGKVSAVITAAAGVGAAVKAAQK | 529745 | 529565 | - | 31 |
| 4 | MLTQKVNADASLKGALDAGAPARGTAPVM | ATPAIFATPVIAKAAGAVVGAGAVTGAAYAAYKVATG | 530051 | 529850 | - | 30 |
| 5 | MAYCAGFTEVLHRGIRGPRR | TGRAYAGVVAAAVTGAARTT | 530248 | 530371 | + | 7 |
| 6 | MLLDKVNQDVSLAGFVAEHGPGARIAPVM | ATPAIFTPAAVKVGACVGAAVATYVATKAVK | 530546 | 530363 | - | 33 |
| 7 | VLVDVSGIAK | ATLRSRMWQRCRCTLIASAQVDAVNSLQRPWIQRNVRESVRCIG | 533777 | 533612 | - | 6 |

### Results for SFM59896.1 [Streptomyces sp. cf124] back to top

previous - next

#### Architecture

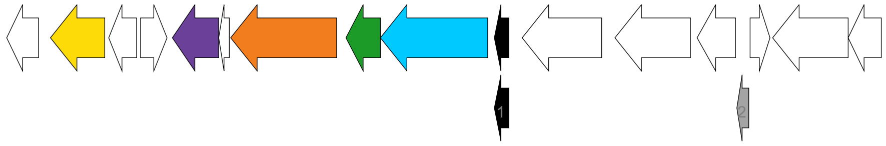

1000 nucleotides

Link to nucleotide sequence

| Accession | start | end | direction | length (aa) | Pfam/HMM | name | description | E-value |
|---|---|---|---|---|---|---|---|---|
| SFM59782.1 | 61193 | 60617 | - | 191 | PF13302<br>PF00583 | Acetyltransf_3<br>Acetyltransf_1 | Acetyltransferase (GNAT) domain<br>Acetyltransferase (GNAT) family | 2.30E-07<br>9.90E-04 |
| SFM59803.1 | 62373 | 61401 | - | 323 | PF00561<br>PF12697<br>TIGR03056<br>TIGR02427<br>TIGR03343 | Abhydrolase_1<br>Abhydrolase_6<br>TIGR03056<br>TIGR02427<br>TIGR03343 | alpha/beta hydrolase fold<br>Alpha/beta hydrolase family<br>bchO_mg_che_rel: putative magnesium chelatase accessory protein<br>protocat_pcaD: 3-oxoadipate enol-lactonase<br>biphenyl_bphD: 2-hydroxy-6-oxo-6-phenylhexa-2,4-dienoate hydrolase | 1.90E-27<br>1.10E-19<br>3.70E-17<br>2.30E-16<br>9.40E-16 |
| SFM59814.1 | 62932 | 62437 | - | 164 | TIGR00369<br>PF03061<br>TIGR02286<br>PF14539<br>PF13622 | TIGR00369<br>4HBT<br>TIGR02286<br>DUF4442<br>4HBT_3 | unchar_dom_1: uncharacterized domain 1<br>Thioesterase superfamily<br>PaaD: phenylacetic acid degradation protein PaaD<br>Domain of unknown function (DUF4442)<br>Thioesterase-like superfamily | 5.50E-27<br>2.50E-16<br>1.90E-09<br>2.20E-08<br>5.90E-05 |
| SFM59830.1 | 63021 | 63498 | + | 158 | PF01638 | HxlR | HxlR-like helix-turn-helix | 9.70E-20 |
| SFM59842.1 | 64405 | 63577 | - | 275 | PF13649<br>PF08241<br>PF08242<br>PF13489<br>TIGR01983 | Methyltransf_25<br>Methyltransf_11<br>Methyltransf_12<br>Methyltransf_23<br>TIGR01983 | Methyltransferase domain<br>Methyltransferase domain<br>Methyltransferase domain<br>Methyltransferase domain<br>UbiG: 3-demethylubiquinone-9 3-O-methyltransferase | 1.60E-15<br>1.30E-12<br>4.30E-07<br>5.20E-07<br>3.10E-05 |
| SFM59855.1 | 64593 | 64401 | - | 63 | NO PFAM MATCH | - | - | - |
| SFM59869.1 | 66511 | 64618 | - | 630 | TIGR02204<br>TIGR02203<br>TIGR00958<br>TIGR03796<br>TIGR03375 | TIGR02204<br>TIGR02203<br>TIGR00958<br>TIGR03796<br>TIGR03375 | MsbA_rel: ABC transporter, permease/ATP-binding protein<br>MsbA_lipidA: lipid A export permease/ATP-binding protein MsbA<br>3a01208: antigen peptide transporter 2<br>NHLM_micro_ABC1: NHLM bacteriocin system ABC transporter, peptidase/ATP-binding protein<br>type_I_sec_LssB: type I secretion system ATPase | 2.30E-106<br>2.10E-104<br>2.00E-92<br>4.70E-86<br>2.00E-81 |
| SFM59885.1 | 67295 | 66674 | - | 206 | LinL | LinL | LinL | 1.10E-68 |
| SFM59896.1 | 69211 | 67291 | - | 639 | LinH<br>LinG<br>LinE<br>PF12697<br>PF12146 | LinH<br>LinG<br>LinE<br>Abhydrolase_6<br>Hydrolase_4 | LinH (Fusion of LinE/LinG)<br>LinG Homolog<br>LinE Homolog<br>Alpha/beta hydrolase family<br>Serine aminopeptidase, S33 | 6.60E-247<br>3.40E-157<br>5.90E-92<br>9.20E-16<br>3.20E-08 |
| SFM59913.1 | 69595 | 69325 | - | 89 | LinA | LinA | LinA (linaridin precursor) | 2.30E-08 |
| SFM59926.1 | 71244 | 69813 | - | 476 | TIGR02734<br>PF13450<br>TIGR03467<br>TIGR00292<br>TIGR03315 | TIGR02734<br>NAD_binding_8<br>TIGR03467<br>TIGR00292<br>TIGR03315 | crtI_fam: phytoene desaturase<br>NAD(P)-binding Rossmann-like domain<br>HpnE: squalene-associated FAD-dependent desaturase<br>TIGR00292: thiazole biosynthesis enzyme<br>Se_ygfK: putative selenate reductase, YgfK subunit | 2.50E-12<br>3.80E-12<br>3.70E-08<br>2.00E-07<br>3.20E-07 |
| SFM59943.1 | 72837 | 71490 | - | 448 | PF07730<br>PF13581 | HisKA_3<br>HATPase_c_2 | Histidine kinase<br>Histidine kinase-like ATPase domain | 8.90E-14<br>4.80E-04 |
| SFM59953.1 | 73639 | 72937 | - | 233 | PF00072<br>PF00196<br>TIGR02875<br>TIGR03020<br>TIGR03541 | Response_reg<br>GerE<br>TIGR02875<br>TIGR03020<br>TIGR03541 | Response regulator receiver domain<br>Bacterial regulatory proteins, luxR family<br>spore_0_A: sporulation transcription factor Spo0A<br>EpsA: transcriptional regulator EpsA<br>reg_near_HchA: LuxR family transcriptional regulatory, chaperone HchA-associated | 1.50E-24<br>1.00E-17<br>7.40E-16<br>1.10E-12<br>2.90E-08 |
| SFM59970.1 | 73882 | 74245 | + | 120 | PF00196<br>TIGR03541<br>TIGR03020<br>PF08281 | GerE<br>TIGR03541<br>TIGR03020<br>Sigma70_r4_2 | Bacterial regulatory proteins, luxR family<br>reg_near_HchA: LuxR family transcriptional regulatory, chaperone HchA-associated<br>EpsA: transcriptional regulator EpsA<br>Sigma-70, region 4 | 2.00E-15<br>2.20E-12<br>6.40E-09<br>2.80E-07 |
| SFM59978.1 | 75649 | 74293 | - | 451 | PF13360<br>PF01011<br>TIGR03300<br>PF13570<br>TIGR03075 | PQQ_2<br>PQQ<br>TIGR03300<br>PQQ_3<br>TIGR03075 | PQQ-like domain<br>PQQ enzyme repeat<br>assembly_YfgL: outer membrane assembly lipoprotein YfgL<br>PQQ-like domain<br>PQQ_enz_alc_DH: PQQ-dependent dehydrogenase, methanol/ethanol family | 6.60E-27<br>3.90E-13<br>1.70E-12<br>5.20E-09<br>4.80E-07 |
| SFM59997.1 | 76224 | 75645 | - | 192 | TIGR02983<br>TIGR02937<br>PF08281<br>TIGR02954<br>PF04545 | TIGR02983<br>TIGR02937<br>Sigma70_r4_2<br>TIGR02954<br>Sigma70_r4 | SigE-fam_strep: RNA polymerase sigma-70 factor, sigma-E family<br>sigma70-ECF: RNA polymerase sigma factor, sigma-70 family<br>Sigma-70, region 4<br>Sig70_famx3: RNA polymerase sigma-70 factor, TIGR02954 family<br>Sigma-70, region 4 | 1.70E-54<br>2.30E-18<br>1.30E-16<br>3.70E-14<br>2.50E-12 |

| index | leader | core | start | end | dir | score |
|---|---|---|---|---|---|---|
| 1 | MSSIENALNSVEIPVEGVVYVAARPTL | GTPRIARIGRIAQAAEGVGAIAAAA...NNLAAEANAQNAAALAAVGGTPPAV | 69595 | 69325 | - | 28 |
| 2 | VGVEDPRAVRLLRCDSSVTSGRRASTIRPIPGSSHRPIGLGPPPRG | GTFSGLAPTARFGGGVRPAGFP | 73862 | 73655 | - | 6 |

### Results for WP_084898409.1 [Streptomyces sp. CB03238] back to top

previous - next

#### Architecture

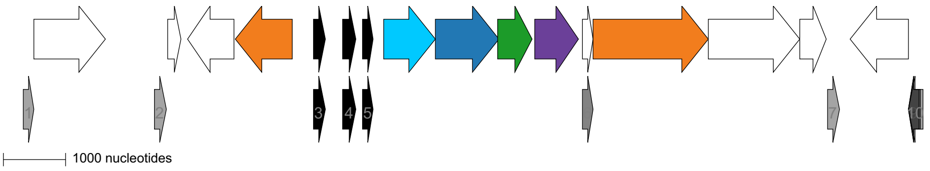

Link to nucleotide sequence

| Accession | start | end | direction | length (aa) | Pfam/HMM | name | description | E-value |
|---|---|---|---|---|---|---|---|---|
| WP_084898401.1 | 520055 | 521204 | + | 382 | PF05787 | DUF839 | Bacterial protein of unknown function (DUF839) | 3.10E-31 |
| WP_016033372.1 | 522211 | 522418 | + | 69 | INFERRED GENE | - | - | - |
| WP_143662655.1 | 523273 | 522523 | - | 249 | PF01061<br>PF12698 | ABC2_membrane<br>ABC2_membrane_3 | ABC-2 type transporter<br>ABC-2 family transporter protein | 1.40E-14<br>1.60E-04 |
| WP_084898404.1 | 524217 | 523284 | - | 310 | TIGR01188<br>TIGR03864<br>TIGR01288<br>TIGR04406<br>TIGR00968 | TIGR01188<br>TIGR03864<br>TIGR01288<br>TIGR04406<br>TIGR00968 | drrA: daunorubicin resistance ABC transporter, ATP-binding protein<br>PQQ_ABC_ATP: ABC transporter, ATP-binding subunit, PQQ-dependent alcohol dehydrogenase system<br>nodI: nodulation ABC transporter NodI<br>LPS_export_lptB: LPS export ABC transporter ATP-binding protein<br>3a0106s01: sulfate ABC transporter, ATP-binding protein | 5.80E-60<br>1.70E-43<br>8.30E-42<br>3.90E-33<br>9.70E-33 |
| WP_084898405.1 | 524554 | 524737 | + | 60 | LinA | LinA | LinA (linaridin precursor) | 1.90E-10 |
| WP_084898406.1 | 525024 | 525225 | + | 66 | LinA | LinA | LinA (linaridin precursor) | 3.20E-11 |
| WP_084898407.1 | 525330 | 525510 | + | 59 | LinA | LinA | LinA (linaridin precursor) | 5.60E-08 |
| WP_084898408.1 | 525672 | 526509 | + | 278 | LinH<br>LinG | LinH<br>LinG | LinH (Fusion of LinE/LinG)<br>LinG Homolog | 1.10E-104<br>3.30E-104 |
| WP_084898409.1 | 526505 | 527522 | + | 338 | LinE<br>LinH<br>PF12697<br>TIGR02427<br>PF00561 | LinE<br>LinH<br>Abhydrolase_6<br>TIGR02427<br>Abhydrolase_1 | LinE Homolog<br>LinH (Fusion of LinE/LinG)<br>Alpha/beta hydrolase family<br>protocat_pcaD: 3-oxoadipate enol-lactonase<br>alpha/beta hydrolase fold | 2.60E-125<br>2.20E-78<br>1.90E-14<br>1.70E-11<br>1.80E-10 |
| WP_143662656.1 | 527518 | 528064 | + | 181 | LinL | LinL | LinL | 6.40E-64 |
| WP_107417879.1 | 528108 | 528816 | + | 235 | PF13649<br>PF08241<br>PF13489<br>PF08242<br>TIGR02021 | Methyltransf_25<br>Methyltransf_11<br>Methyltransf_23<br>Methyltransf_12<br>TIGR02021 | Methyltransferase domain<br>Methyltransferase domain<br>Methyltransferase domain<br>Methyltransferase domain<br>BchM-ChlM: magnesium protoporphyrin O-methyltransferase | 5.30E-15<br>1.70E-10<br>1.80E-09<br>3.50E-07<br>9.20E-06 |
| WP_084898411.1 | 528866 | 529046 | + | 59 | NO PFAM MATCH | - | - | - |
| WP_084898412.1 | 529042 | 530896 | + | 617 | TIGR02203<br>TIGR02204<br>TIGR01846<br>TIGR03375<br>TIGR03797 | TIGR02203<br>TIGR02204<br>TIGR01846<br>TIGR03375<br>TIGR03797 | MsbA_lipidA: lipid A export permease/ATP-binding protein MsbA<br>MsbA_rel: ABC transporter, permease/ATP-binding protein<br>type_I_sec_HlyB: type I secretion system ATPase<br>type_I_sec_LssB: type I secretion system ATPase<br>NHLM_micro_ABC2: NHLM bacteriocin system ABC transporter, ATP-binding protein | 1.00E-104<br>1.60E-104<br>1.20E-86<br>3.60E-84<br>4.80E-81 |
| WP_084898413.1 | 530892 | 532359 | + | 488 | TIGR02734<br>TIGR03467<br>PF01593<br>PF13450<br>TIGR00292 | TIGR02734<br>TIGR03467<br>Amino_oxidase<br>NAD_binding_8<br>TIGR00292 | crtI_fam: phytoene desaturase<br>HpnE: squalene-associated FAD-dependent desaturase<br>Flavin containing amine oxidoreductase<br>NAD(P)-binding Rossmann-like domain<br>TIGR00292: thiazole biosynthesis enzyme | 3.70E-10<br>5.30E-10<br>9.60E-10<br>1.60E-09<br>1.00E-07 |
| WP_084898414.1 | 532358 | 532796 | + | 145 | NO PFAM MATCH | - | - | - |
| WP_084898415.1 | 534111 | 533169 | - | 313 | PF01370<br>PF16363<br>TIGR03466<br>PF13460<br>TIGR02032 | Epimerase<br>GDP_Man_Dehyd<br>TIGR03466<br>NAD_binding_10<br>TIGR02032 | NAD dependent epimerase/dehydratase family<br>GDP-mannose 4,6 dehydratase<br>HpnA: hopanoid-associated sugar epimerase<br>NAD(P)H-binding<br>GG-red-SF: geranylgeranyl reductase family | 1.30E-14<br>3.20E-06<br>4.00E-06<br>1.80E-05<br>1.50E-04 |

| index | leader | core | start | end | dir | score |
|---|---|---|---|---|---|---|
| 1 | MPDRFCAIPPLTGWPESVLLGEPYRPGAAALL | SLPGDQSVARNTKCSPFPG | 519889 | 520045 | + | 6 |
| 2 | MAAESAPRAPTRRTGRSRATGSRRSGPPPHPSWAAWWRRANH | STATSTATGDPRCPGRPSRCSP | 521992 | 522187 | + | 6 |
| 3 | MLLDKVNQDVSLAGFVAEHGPGARIAPVM | ATPAIFTPAAVKAGACVGVAVATYMATKAVK | 524554 | 524737 | + | 33 |
| 4 | MLTQKVNADASLKGALEAGAPARGTAPVM | ATPAIFATPVIAKAAGAVVGAGAVTGAAYAAYKVATG | 525024 | 525225 | + | 30 |
| 5 | MSVLVEKVNAGAGQAADGSVAPVM | ATPAAFVAGVVAGGKVSAVMTAAAGVGAAVKAAQK | 525330 | 525510 | + | 31 |
| 6 | MLTASRLTWFVIAFAFAVP | STLVMFRDNGVVTRDAWVKSFVFAAAVAAVIAVVFGKGSQ | 528866 | 529046 | + | 7 |
| 7 | MNGIDKNDLDEVIA | KTFNEVKTAVNSHSEKSIQMYSTALRALVELRQQILSEERVGS | 532822 | 532996 | + | 6 |
| 8 | LAEAPTWYFVGRLIARVAVDRRPRGAAATMTGRRARLCP | VAPPTTHRRRLIRPATLV | 534289 | 534115 | - | 6 |
| 9 | VPALRRVGGGAHLVLRRQAHRPRGRGPPPPGSCCHDDGTAGA | PVPGGASNYPSPPTDPTGHVSVSA | 534308 | 534107 | - | 6 |
| 10 | VTGREPDGAGPCRPSDGLAEAPTWY...ARVAVDRRPRGAAATMTGRRARLCP | VAPPTTHRRRLIRPATLV | 534340 | 534115 | - | 6 |

### Results for OKJ54325.1 [Streptomyces sp. CB02115] back to top

previous - next

#### Architecture

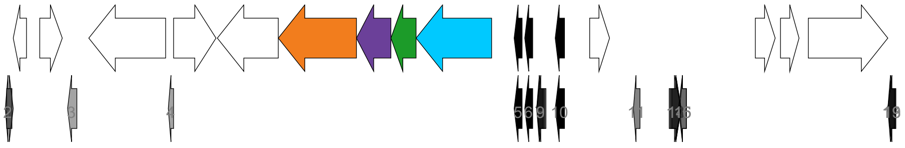

1000 nucleotides

Link to nucleotide sequence

| Accession | start | end | direction | length (aa) | Pfam/HMM | name | description | E-value |
|---|---|---|---|---|---|---|---|---|
| OKJ54317.1 | 578136 | 577818 | - | 105 | NO PFAM MATCH | - | - | - |
| OKJ54318.1 | 578454 | 578991 | + | 178 | PF12840<br>PF01022 | HTH_20<br>HTH_5 | Helix-turn-helix domain<br>Bacterial regulatory protein, arsR family | 4.90E-11<br>2.40E-05 |
| OKJ54319.1 | 581494 | 579640 | - | 617 | NO PFAM MATCH | - | - | - |
| OKJ54320.1 | 581683 | 582697 | + | 337 | NO PFAM MATCH | - | - | - |
| OKJ54321.1 | 584220 | 582750 | - | 489 | TIGR02734<br>PF13450<br>TIGR01317<br>TIGR00292<br>TIGR03467 | TIGR02734<br>NAD_binding_8<br>TIGR01317<br>TIGR00292<br>TIGR03467 | crtI_fam: phytoene desaturase<br>NAD(P)-binding Rossmann-like domain<br>GOGAT_sm_gam: glutamate synthase, NADH/NADPH, small subunit<br>TIGR00292: thiazole biosynthesis enzyme<br>HpnE: squalene-associated FAD-dependent desaturase | 3.80E-10<br>6.30E-09<br>1.00E-08<br>1.30E-08<br>6.30E-08 |
| OKJ54322.1 | 586118 | 584216 | - | 633 | TIGR02204<br>TIGR02203<br>TIGR00958<br>TIGR03796<br>TIGR03375 | TIGR02204<br>TIGR02203<br>TIGR00958<br>TIGR03796<br>TIGR03375 | MsbA_rel: ABC transporter, permease/ATP-binding protein<br>MsbA_lipidA: lipid A export permease/ATP-binding protein MsbA<br>3a01208: antigen peptide transporter 2<br>NHLM_micro_ABC1: NHLM bacteriocin system ABC transporter, peptidase/ATP-binding protein<br>type_I_sec_LssB: type I secretion system ATPase | 7.90E-98<br>1.30E-90<br>1.20E-81<br>4.80E-78<br>5.50E-75 |
| OKJ54323.1 | 586927 | 586114 | - | 270 | PF13649<br>PF08241<br>PF08242<br>PF13489<br>PF13847 | Methyltransf_25<br>Methyltransf_11<br>Methyltransf_12<br>Methyltransf_23<br>Methyltransf_31 | Methyltransferase domain<br>Methyltransferase domain<br>Methyltransferase domain<br>Methyltransferase domain<br>Methyltransferase domain | 3.60E-16<br>1.10E-11<br>2.10E-07<br>6.00E-07<br>9.10E-06 |
| OKJ54324.1 | 587532 | 586923 | - | 202 | LinL | LinL | LinL | 3.80E-65 |
| OKJ54325.1 | 589379 | 587528 | - | 616 | LinH<br>LinG<br>LinE<br>LinL<br>PF12697 | LinH<br>LinG<br>LinE<br>LinL<br>Abhydrolase_6 | LinH (Fusion of LinE/LinG)<br>LinG Homolog<br>LinE Homolog<br>LinL<br>Alpha/beta hydrolase family | 1.80E-249<br>6.10E-111<br>3.50E-94<br>2.70E-33<br>5.70E-16 |
| OKJ54326.1 | 590119 | 589921 | - | 65 | LinA | LinA | LinA (linaridin precursor) | 2.40E-11 |
| OKJ54327.1 | 590374 | 590179 | - | 64 | LinA | LinA | LinA (linaridin precursor) | 1.50E-11 |
| OKJ54328.1 | 591116 | 590912 | - | 67 | LinA | LinA | LinA (linaridin precursor) | 7.20E-11 |
| OKJ54329.1 | 591736 | 592207 | + | 156 | PF00583<br>TIGR03448<br>TIGR01575<br>PF13508<br>PF13302 | Acetyltransf_1<br>TIGR03448<br>TIGR01575<br>Acetyltransf_7<br>Acetyltransf_3 | Acetyltransferase (GNAT) family<br>mycothiol_MshD: mycothiol synthase<br>rimI: ribosomal-protein-alanine acetyltransferase<br>Acetyltransferase (GNAT) domain<br>Acetyltransferase (GNAT) domain | 8.30E-13<br>1.40E-08<br>8.20E-08<br>1.40E-06<br>2.70E-06 |
| OKJ54330.1 | 595737 | 596217 | + | 159 | PF03745 | DUF309 | Domain of unknown function (DUF309) | 6.30E-20 |
| OKJ54698.1 | 596355 | 596808 | + | 150 | NO PFAM MATCH | - | - | - |
| OKJ54331.1 | 597003 | 598941 | + | 645 | PF10022 | DUF2264 | Uncharacterized protein conserved in bacteria (DUF2264) | 5.10E-100 |

| index | leader | core | start | end | dir | score |
|---|---|---|---|---|---|---|
| 1 | VTARNSHSRPGADAEPDRFEESRPSS | PTPTGPMGRVRDTHVQGSPPGREAGSPRPVTA | 577652 | 577829 | + | 6 |
| 2 | LDVRVTDTPHGTSWGGRRGAALLEA | VRLGVGSRSAVAVPCRHPSDAPRPRT | 577778 | 577622 | - | 7 |
| 3 | VILGAPPRRSSRWCGRFWGGRSVSCCEGDGGGSGF | VEGAVAELGERDVDAVAGEAEKGLGVCVLNAKGVWL | 579342 | 579126 | - | 6 |
| 4 | MGPHGPSLEHVRRDDGAQVCPGR | RGVRSQRSLTPCRHQGSSGCVTT | 581695 | 581554 | - | 6 |
| 5 | MSTSPIPAALRGAGGLSESAPGQALSTFAPVA | ATPGIVAGAAIGVALVNAFVAGYNYTGNVELPM | 590119 | 589921 | - | 31 |
| 6 | MSKSSAPAALTATGALTESDPGVTLPGLAPVA | ATPVAIVASLGAAFVAGYAAGRAAGSDIELPM | 590374 | 590179 | - | 29 |
| 7 | LSTASIPVVLSETGALSESDPGRAPLD | TVPVASVTFTMSACVGISVCLIGSTIELPQ | 590621 | 590447 | - | 8 |
| 8 | MTSPHRPPTSEEKSLSTASIPVVLSETGALSESDPGRAPLD | TVPVASVTFTMSACVGISVCLIGSTIELPQ | 590663 | 590447 | - | 7 |
| 9 | VGPSRPRRSPMTSPHRPPTSEEKSL...TASIPVVLSETGALSESDPGRAPLD | TVPVASVTFTMSACVGISVCLIGSTIELPQ | 590693 | 590447 | - | 7 |
| 10 | MPEFRQPGWTRGIAPLQEGAGGQVFGGASPVA | ATPVVVAGAGAVVVAFAAGVAARTLANGGNIELPM | 591116 | 590912 | - | 31 |
| 11 | LSCAVTRGNRGKRDGTWRSLVAHLTGGQ | GVAGSNPVVPTGDSRRSGPVSEQSETGP | 592963 | 592792 | - | 7 |
| 12 | VDHAEQGVGERQVGPAPAGDLEGPDRFAEEALRAVQLVDEGVGDADH | GGEFEAPLGVGPAEATGRAAVDQGAFAQVHDELGVLTAAPWRRRPAA | 593670 | 593955 | + | 7 |
| 13 | VGERQVGPAPAGDLEGPDRFAEEALRAVQLVDEGVGDADHGGE | FEAPLGVGPAEATGRAAVDQGAFAQVHDELGVLTAAPWRRRPAA | 593691 | 593955 | + | 7 |
| 14 | VGPAPAGDLEGPDRFAEEALRAVQLVDEGVGDADHGGEFEA | PLGVGPAEATGRAAVDQGAFAQVHDELGVLTAAPWRRRPAA | 593706 | 593955 | + | 7 |

| | | | | | | |
|---|---|---|---|---|---|---|
| 15 | VGGHAGRRGRRTRPLPAAHPQR | RPPEGPRQRRAGAAHAAGLRRHGAAVRTPSSSWTCANAP | 594062 | 593876 | - | 6 |
| 16 | MRLQAAHFMWEGMQAAEDDGPARSRRLT | RNADHLKGRDNAERALLTLRAFDAMVRR | 594088 | 593917 | - | 6 |
| 17 | VPEAVARLAPER | PTTLISGPSATSDIELERVEGVHGPRTLVVVIRTDA | 599092 | 598945 | - | 6 |
| 18 | VDLSAVEAGVPEAVARLAPER | PTTLISGPSATSDIELERVEGVHGPRTLVVVIRTDA | 599119 | 598945 | - | 8 |
| 19 | VVDLSAVEAGVPEAVARLAPER | PTTLISGPSATSDIELERVEGVHGPRTLVVVIRTDA | 599122 | 598945 | - | 8 |

### Results for AUH40579.1 [Streptomyces sp. CMB-StM0423] back to top

previous - next

#### Architecture

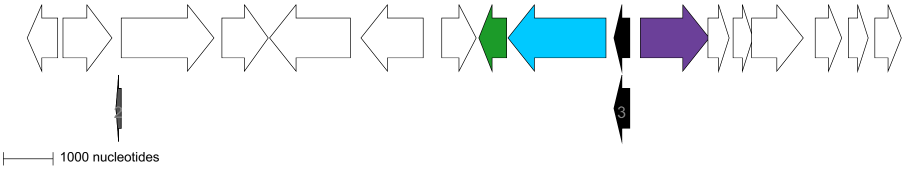

Link to nucleotide sequence

| Accession | start | end | direction | length (aa) | Pfam/HMM | name | description | E-value |
|---|---|---|---|---|---|---|---|---|
| AUH40572.1 | 2419971 | 2419356 | - | 204 | PF00440<br>PF13977<br>TIGR03384<br>TIGR03968<br>TIGR03613 | TetR_N<br>TetR_C_6<br>TIGR03384<br>TIGR03968<br>TIGR03613 | Bacterial regulatory proteins, tetR family<br>BetI-type transcriptional repressor, C-terminal<br>betaine_BetI: transcriptional repressor BetI<br>mycofact_TetR: mycofactocin system transcriptional regulator<br>RutR: pyrimidine utilization regulatory protein R | 3.60E-16<br>7.00E-11<br>8.90E-11<br>4.50E-09<br>5.10E-08 |
| AUH40573.1 | 2420077 | 2421052 | + | 324 | NO PFAM MATCH | - | - | - |
| AUH40574.1 | 2421238 | 2423110 | + | 623 | TIGR00976<br>PF02129<br>PF08530 | TIGR00976<br>Peptidase_S15<br>PepX_C | /NonD: hydrolase CocE/NonD family protein<br>X-Pro dipeptidyl-peptidase (S15 family)<br>X-Pro dipeptidyl-peptidase C-terminal non-catalytic domain | 7.90E-149<br>2.10E-55<br>7.70E-12 |
| AUH40575.1 | 2423274 | 2424207 | + | 310 | TIGR00212<br>PF01379<br>PF03900 | TIGR00212<br>Porphobil_deam<br>Porphobil_deamC | hemC: hydroxymethylbilane synthase<br>Porphobilinogen deaminase, dipyromethane cofactor binding domain<br>Porphobilinogen deaminase, C-terminal domain | 5.70E-97<br>9.40E-63<br>3.90E-12 |
| AUH40576.1 | 2425847 | 2424227 | - | 539 | PF01425<br>TIGR00132<br>TIGR02715<br>TIGR02713 | Amidase<br>TIGR00132<br>TIGR02715<br>TIGR02713 | Amidase<br>gatA: aspartyl/glutamyl-tRNA(Asn/Gln) amidotransferase, A subunit<br>amido_AtzE: amidohydrolase, AtzE family<br>allophanate_hyd: allophanate hydrolase | 6.60E-87<br>3.10E-60<br>8.90E-45<br>5.10E-41 |
| AUH40577.1 | 2427335 | 2426057 | - | 425 | PF13360<br>PF01011 | PQQ_2<br>PQQ | PQQ-like domain<br>PQQ enzyme repeat | 5.40E-10<br>2.10E-08 |
| AUH44821.1 | 2427703 | 2428381 | + | 225 | PF00196<br>TIGR03020<br>TIGR03541<br>TIGR02937<br>PF13936 | GerE<br>TIGR03020<br>TIGR03541<br>TIGR02937<br>HTH_38 | Bacterial regulatory proteins, luxR family<br>EpsA: transcriptional regulator EpsA<br>reg_near_HchA: LuxR family transcriptional regulatory, chaperone HchA-associated<br>sigma70-ECF: RNA polymerase sigma factor, sigma-70 family<br>Helix-turn-helix domain | 4.70E-19<br>6.20E-12<br>3.00E-10<br>2.10E-04<br>2.70E-04 |
| AUH40578.1 | 2429010 | 2428452 | - | 185 | LinL | LinL | LinL | 8.90E-59 |
| AUH40579.1 | 2431013 | 2429021 | - | 663 | LinH<br>LinE<br>LinG<br>PF00561<br>TIGR02427 | LinH<br>LinE<br>LinG<br>Abhydrolase_1<br>TIGR02427 | LinH (Fusion of LinE/LinG)<br>LinE Homolog<br>LinG Homolog<br>alpha/beta hydrolase fold<br>protocat_pcaD: 3-oxoadipate enol-lactonase | 1.00E-200<br>3.20E-94<br>2.80E-84<br>1.60E-11<br>1.60E-09 |
| AUH40580.1 | 2431477 | 2431174 | - | 100 | LinA | LinA | LinA (linaridin precursor) | 5.00E-09 |
| AUH44822.1 | 2431690 | 2433079 | + | 462 | PF00891<br>TIGR02716<br>PF16864<br>PF13649<br>PF08242 | Methyltransf_2<br>TIGR02716<br>Dimerisation2<br>Methyltransf_25<br>Methyltransf_12 | O-methyltransferase domain<br>C20_methyl_CrtF: C-20 methyltransferase BchU<br>Dimerisation domain<br>Methyltransferase domain<br>Methyltransferase domain | 1.50E-45<br>4.30E-17<br>3.40E-07<br>1.20E-06<br>1.10E-05 |
| AUH40581.1 | 2433056 | 2433470 | + | 137 | PF04134 | DUF393 | Protein of unknown function, DUF393 | 5.50E-11 |
| AUH40582.1 | 2433553 | 2433949 | + | 131 | PF12840<br>PF01022<br>PF12802<br>TIGR01884<br>PF01047 | HTH_20<br>HTH_5<br>MarR_2<br>TIGR01884<br>MarR | Helix-turn-helix domain<br>Bacterial regulatory protein, arsR family<br>MarR family<br>cas_HTH: CRISPR locus-related DNA-binding protein<br>MarR family | 3.40E-16<br>7.50E-10<br>6.30E-08<br>7.80E-07<br>1.50E-06 |
| AUH40583.1 | 2433945 | 2434986 | + | 346 | TIGR03086<br>PF08327<br>TIGR03083<br>PF11716<br>PF10604 | TIGR03086<br>AHSA1<br>TIGR03083<br>MDMPI_N<br>Polyketide_cyc2 | TIGR03086: TIGR03086 family protein<br>Activator of Hsp90 ATPase homolog 1-like protein<br>TIGR03083: uncharacterized Actinobacterial protein TIGR03083<br>Mycothiol maleylpyruvate isomerase N-terminal domain<br>Polyketide cyclase / dehydrase and lipid transport | 3.10E-32<br>7.00E-24<br>1.10E-16<br>1.60E-07<br>1.00E-05 |
| AUH40584.1 | 2435227 | 2435740 | + | 170 | PF01814<br>TIGR03652 | Hemerythrin<br>TIGR03652 | Hemerythrin HHE cation binding domain<br>FeS_repair_RIC: iron-sulfur cluster repair di-iron protein | 9.40E-20<br>2.30E-04 |
| WP_018835193.1 | 2435886 | 2436281 | + | 131 | INFERRED GENE | - | - | - |
| AUH40585.1 | 2436420 | 2436954 | + | 177 | PF13420<br>PF00583<br>TIGR01575<br>PF13508<br>PF13302 | Acetyltransf_4<br>Acetyltransf_1<br>TIGR01575<br>Acetyltransf_7<br>Acetyltransf_3 | Acetyltransferase (GNAT) domain<br>Acetyltransferase (GNAT) family<br>rimI: ribosomal-protein-alanine acetyltransferase<br>Acetyltransferase (GNAT) domain<br>Acetyltransferase (GNAT) domain | 6.20E-22<br>1.00E-11<br>1.40E-05<br>2.50E-05<br>1.10E-04 |

| index | leader | core | start | end | dir | score |
|---|---|---|---|---|---|---|

| | | | | | | |
|---|---|---|---|---|---|---|
| 1 | MLLRDDGTELWGDGGWA | HPGKPHVAGATGTDAWSV | 2421238 | 2421130 | - | 7 |
| 2 | VTSLHMLLRDDGTELWGDGG | WAHPGKPHVAGATGTDAWSV | 2421253 | 2421130 | - | 6 |
| 3 | MTTTLQPPTTADSRIAVWAQKNAAEHDEHFVAPTDRVVPVL | FTPLIALTVAGAVGCALGPVTSFVT...PDTPGGAALRGRSVDQLTGGHPGMV | 2431477 | 2431174 | - | 26 |

### Results for WP_018835201.1 [Streptomyces sp. CNQ766] back to top

previous - next

#### Architecture

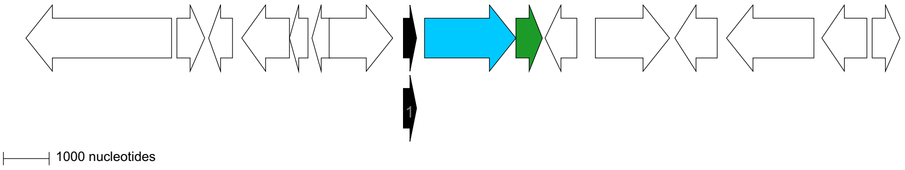

Link to nucleotide sequence

| Accession | start | end | direction | length (aa) | Pfam/HMM | name | description | E-value |
|---|---|---|---|---|---|---|---|---|
| WP_018835193.1 | 290474 | 287282 | - | 1063 | PF03704<br>PF13424<br>PF00486<br>PF13191<br>PF13401 | BTAD<br>TPR_12<br>Trans_reg_C<br>AAA_16<br>AAA_22 | Bacterial transcriptional activator domain<br>Tetratricopeptide repeat<br>Transcriptional regulatory protein, C terminal<br>AAA ATPase domain<br>AAA domain | 2.10E-32<br>4.90E-13<br>1.80E-08<br>7.10E-07<br>5.10E-04 |
| WP_018835194.1 | 290584 | 291190 | + | 201 | NO PFAM MATCH | - | - | - |
| WP_018835195.1 | 291803 | 291290 | - | 170 | PF01814 | Hemerythrin | Hemerythrin HHE cation binding domain | 2.30E-18 |
| WP_018835196.1 | 293049 | 292008 | - | 346 | TIGR03086<br>PF08327<br>TIGR03083<br>PF11716<br>PF10604 | TIGR03086<br>AHSA1<br>TIGR03083<br>MDMPI_N<br>Polyketide_cyc2 | TIGR03086: TIGR03086 family protein<br>Activator of Hsp90 ATPase homolog 1-like protein<br>TIGR03083: uncharacterized Actinobacterial protein TIGR03083<br>Mycothiol maleylpyruvate isomerase N-terminal domain<br>Polyketide cyclase / dehydrase and lipid transport | 2.40E-32<br>9.40E-24<br>2.60E-17<br>3.90E-07<br>1.80E-05 |
| WP_018835197.1 | 293450 | 293045 | - | 134 | PF12840<br>PF01022<br>PF12802<br>TIGR01884<br>PF01047 | HTH_20<br>HTH_5<br>MarR_2<br>TIGR01884<br>MarR | Helix-turn-helix domain<br>Bacterial regulatory protein, arsR family<br>MarR family<br>cas_HTH: CRISPR locus-related DNA-binding protein<br>MarR family | 1.20E-16<br>7.80E-10<br>1.00E-07<br>6.80E-07<br>2.10E-06 |
| WP_018835198.1 | 293945 | 293534 | - | 136 | PF04134 | DUF393 | Protein of unknown function, DUF393 | 2.10E-12 |
| WP_020610489.1 | 293922 | 295310 | + | 462 | INFERRED GENE | - | - | - |
| WP_018835200.1 | 295523 | 295826 | + | 100 | LinA | LinA | LinA (linaridin precursor) | 3.30E-09 |
| WP_018835201.1 | 295987 | 298006 | + | 672 | LinH<br>LinE<br>LinG<br>PF00561<br>TIGR02427 | LinH<br>LinE<br>LinG<br>Abhydrolase_1<br>TIGR02427 | LinH (Fusion of LinE/LinG)<br>LinE Homolog<br>LinG Homolog<br>alpha/beta hydrolase fold<br>protocat_pcaD: 3-oxoadipate enol-lactonase | 9.40E-202<br>5.50E-94<br>4.20E-84<br>1.60E-11<br>1.60E-08 |
| WP_018835202.1 | 298002 | 298560 | + | 185 | LinL | LinL | LinL | 7.60E-60 |
| WP_018835203.1 | 299309 | 298616 | - | 230 | PF00196<br>TIGR03020<br>TIGR03541<br>TIGR02937<br>PF13936 | GerE<br>TIGR03020<br>TIGR03541<br>TIGR02937<br>HTH_38 | Bacterial regulatory proteins, luxR family<br>EpsA: transcriptional regulator EpsA<br>reg_near_HchA: LuxR family transcriptional regulatory, chaperone HchA-associated<br>sigma70-ECF: RNA polymerase sigma factor, sigma-70 family<br>Helix-turn-helix domain | 4.70E-19<br>7.40E-12<br>3.00E-10<br>2.10E-04<br>2.50E-04 |
| WP_018835204.1 | 299731 | 301351 | + | 539 | PF01425<br>TIGR00132<br>TIGR02715<br>TIGR02713 | Amidase<br>TIGR00132<br>TIGR02715<br>TIGR02713 | Amidase<br>gatA: aspartyl/glutamyl-tRNA(Asn/Gln) amidotransferase, A subunit<br>amido_AtzE: amidohydrolase, AtzE family<br>allophanate_hyd: allophanate hydrolase | 3.40E-88<br>1.30E-59<br>3.00E-45<br>1.60E-42 |
| WP_026276125.1 | 302390 | 301457 | - | 310 | TIGR00212<br>PF01379<br>PF03900 | TIGR00212<br>Porphobil_deam<br>Porphobil_deamC | hemC: hydroxymethylbilane synthase<br>Porphobilinogen deaminase, dipyromethane cofactor binding domain<br>Porphobilinogen deaminase, C-terminal domain | 2.10E-98<br>2.50E-64<br>5.00E-11 |
| WP_018835206.1 | 304494 | 302604 | - | 629 | TIGR00976<br>PF02129<br>PF08530 | TIGR00976<br>Peptidase_S15<br>PepX_C | /NonD: hydrolase CocE/NonD family protein<br>X-Pro dipeptidyl-peptidase (S15 family)<br>X-Pro dipeptidyl-peptidase C-terminal non-catalytic domain | 3.00E-148<br>1.70E-53<br>7.10E-10 |
| WP_018835207.1 | 305655 | 304677 | - | 325 | PF12698 | ABC2_membrane_3 | ABC-2 family transporter protein | 5.60E-07 |
| WP_018835208.1 | 305761 | 306376 | + | 204 | PF00440<br>TIGR03384<br>PF13977<br>TIGR03968<br>TIGR03613 | TetR_N<br>TIGR03384<br>TetR_C_6<br>TIGR03968<br>TIGR03613 | Bacterial regulatory proteins, tetR family<br>betaine_BetI: transcriptional repressor BetI<br>BetI-type transcriptional repressor, C-terminal<br>mycofact_TetR: mycofactocin system transcriptional regulator<br>RutR: pyrimidine utilization regulatory protein R | 3.70E-16<br>2.60E-10<br>3.00E-09<br>7.60E-09<br>6.00E-08 |

| index | leader | core | start | end | dir | score |
|---|---|---|---|---|---|---|
| 1 | MTTTLQAPTTEDSRIAVWAQQHAAAHDEYFVAPTDRVVPVL | FTPLIALTVAGAVGCALGPVTSFVT...PDAPNGAALRGRSVDQLTGGHPGMV | 295523 | 295826 | + | 15 |

### Results for WP_078528867.1 [Streptomyces sp. CNS606] back to top

previous - next

#### Architecture

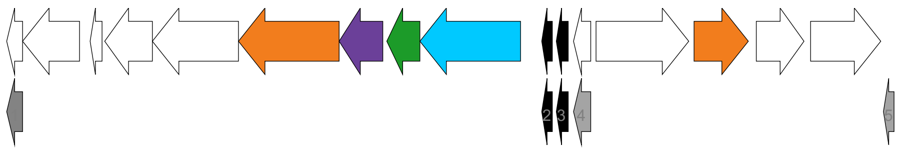

1000 nucleotides

Link to nucleotide sequence

| Accession | start | end | direction | length (aa) | Pfam/HMM | name | description | E-value |
|---|---|---|---|---|---|---|---|---|
| WP_027763595.1 | 34937 | 34679 | - | 85 | PF00550<br>TIGR01688 | PP-binding<br>TIGR01688 | Phosphopantetheine attachment site<br>dltC: D-alanine--poly(phosphoribitol) ligase, subunit 2 | 8.00E-07<br>1.10E-04 |
| WP_051308243.1 | 35914 | 34933 | - | 326 | TIGR00747<br>PF08541 | TIGR00747<br>ACP_syn_III_C | fabH: 3-oxoacyl-[acyl-carrier-protein] synthase III<br>3-Oxoacyl-[acyl-carrier-protein (ACP)] synthase III C terminal | 1.60E-10<br>1.00E-05 |
| WP_027763597.1 | 36315 | 36096 | - | 72 | PF03621 | MbtH | MbtH-like protein | 2.70E-29 |
| WP_027763598.1 | 37183 | 36361 | - | 273 | PF12680<br>PF14534<br>PF07366 | SnoaL_2<br>DUF4440<br>SnoaL | SnoaL-like domain<br>Domain of unknown function (DUF4440)<br>SnoaL-like polyketide cyclase | 9.40E-07<br>5.40E-06<br>2.90E-05 |
| WP_027763599.1 | 38661 | 37179 | - | 493 | TIGR02734<br>TIGR03467<br>TIGR00292<br>TIGR02733<br>PF13450 | TIGR02734<br>TIGR03467<br>TIGR00292<br>TIGR02733<br>NAD_binding_8 | crtI_fam: phytoene desaturase<br>HpnE: squalene-associated FAD-dependent desaturase<br>TIGR00292: thiazole biosynthesis enzyme<br>desat_CrtD: C-3',4' desaturase CrtD<br>NAD(P)-binding Rossmann-like domain | 3.20E-12<br>5.70E-10<br>1.20E-08<br>9.00E-08<br>2.80E-07 |
| WP_027763600.1 | 40382 | 38657 | - | 574 | TIGR02204<br>TIGR02203<br>TIGR00958<br>TIGR03796<br>TIGR01846 | TIGR02204<br>TIGR02203<br>TIGR00958<br>TIGR03796<br>TIGR01846 | MsbA_rel: ABC transporter, permease/ATP-binding protein<br>MsbA_lipidA: lipid A export permease/ATP-binding protein MsbA<br>3a01208: antigen peptide transporter 2<br>NHLM_micro_ABC1: NHLM bacteriocin system ABC transporter, peptidase/ATP-binding protein<br>type_I_sec_HlyB: type I secretion system ATPase | 5.10E-<br>100<br>2.30E-97<br>4.30E-85<br>1.10E-83<br>5.80E-82 |
| WP_051308203.1 | 41137 | 40378 | - | 252 | PF13649<br>PF08241<br>PF08242<br>PF13489<br>PF13847 | Methyltransf_25<br>Methyltransf_11<br>Methyltransf_12<br>Methyltransf_23<br>Methyltransf_31 | Methyltransferase domain<br>Methyltransferase domain<br>Methyltransferase domain<br>Methyltransferase domain<br>Methyltransferase domain | 7.20E-13<br>5.20E-09<br>1.40E-08<br>2.10E-08<br>5.70E-06 |
| WP_027763602.1 | 41786 | 41204 | - | 193 | LinL | LinL | LinL | 3.00E-62 |
| WP_078528867.1 | 43516 | 41782 | - | 577 | LinH<br>LinE<br>LinG<br>PF12697<br>PF00561 | LinH<br>LinE<br>LinG<br>Abhydrolase_6<br>Abhydrolase_1 | LinH (Fusion of LinE/LinG)<br>LinE Homolog<br>LinG Homolog<br>Alpha/beta hydrolase family<br>alpha/beta hydrolase fold | 8.50E-<br>217<br>1.10E-97<br>4.80E-94<br>2.00E-14<br>1.50E-10 |
| WP_027763603.1 | 44057 | 43865 | - | 63 | LinA | LinA | LinA (linaridin precursor) | 5.50E-09 |
| WP_027763604.1 | 44323 | 44128 | - | 64 | LinA | LinA | LinA (linaridin precursor) | 8.00E-15 |
| WP_158687324.1 | 44728 | 44425 | - | 100 | NO PFAM MATCH | - | - | - |
| WP_158687325.1 | 44815 | 46396 | + | 526 | NO PFAM MATCH | - | - | - |
| WP_037789827.1 | 46492 | 47431 | + | 312 | TIGR01188<br>TIGR03864<br>TIGR01288<br>TIGR03522<br>TIGR03265 | TIGR01188<br>TIGR03864<br>TIGR01288<br>TIGR03522<br>TIGR03265 | drrA: daunorubicin resistance ABC transporter, ATP-binding protein<br>PQQ_ABC_ATP: ABC transporter, ATP-binding subunit, PQQ-dependent alcohol dehydrogenase system<br>nodI: nodulation ABC transporter NodI<br>GldA_ABC_ATP: gliding motility-associated ABC transporter ATP-binding subunit GldA<br>PhnT2: putative 2-aminoethylphosphonate ABC transporter, ATP-binding protein | 1.70E-68<br>3.20E-48<br>1.80E-46<br>8.90E-36<br>1.20E-35 |
| WP_027763606.1 | 47572 | 48376 | + | 267 | PF01061<br>PF12698<br>TIGR03062 | ABC2_membrane<br>ABC2_membrane_3<br>TIGR03062 | ABC-2 type transporter<br>ABC-2 family transporter protein<br>pip_yhgE_Cterm: YhgE/Pip C-terminal domain | 1.40E-16<br>7.90E-11<br>4.70E-06 |
| WP_078528891.1 | 48502 | 49717 | + | 404 | PF07730 | HisKA_3 | Histidine kinase | 1.40E-14 |

| index | leader | core | start | end | dir | score |
|---|---|---|---|---|---|---|
| 1 | MNLITSEITAFLSDALRCEVGPEDDYFALGLADSLFALEL | VTFVEERFSLTVEVEDLDLDSFRTADRITGFVQRKSGAAAHPEAP | 34937 | 34679 | - | 7 |
| 2 | MSRITESVIAVTPEKLDDGATTELGAMAPVQ | ATPGFVAFGVGFAGGAGVAWLGVQAYEAGVND | 44057 | 43865 | - | 31 |
| 3 | MEQKIATFMNGQEFGSLENGFHETAPAPVM | GTPVAAVATCWYVAGGVAGAGAVVGAYTWGVANG | 44323 | 44128 | - | 30 |
| 4 | MIQITEGPLGRVFNCAYRQLRAIRNEFDFTVSLDQGTQGDDLSPIGWGIL | LMAYAQVKGCPGFNDEATPGDRGFMSLRGGKRSGLCVRLPLEERTTATRN | 44728 | 44425 | - | 6 |
| 5 | VTLRESRVRSRYGLTVVSVKTTDGEYSHAAP | DTVLNYSDVILVLGRIDDVERFAASN | 49934 | 49760 | - | 6 |

### Results for WP_027766445.1 [Streptomyces sp. CNQ865] back to top

previous - next

#### Architecture

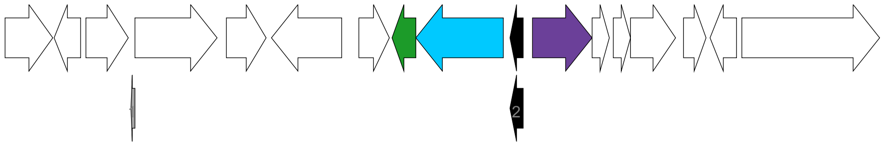

1000 nucleotides

Link to nucleotide sequence

| Accession | start | end | direction | length (aa) | Pfam/HMM | name | description | E-value |
|---|---|---|---|---|---|---|---|---|
| WP_027766440.1 | 62864 | 63968 | + | 367 | PF00144<br>PF13354 | Beta-lactamase<br>Beta-lactamase2 | Beta-lactamase<br>Beta-lactamase enzyme family | 4.90E-63<br>4.60E-04 |
| WP_027766441.1 | 64623 | 64008 | - | 204 | PF00440<br>TIGR03384<br>PF13977<br>TIGR03968<br>TIGR03613 | TetR_N<br>TIGR03384<br>TetR_C_6<br>TIGR03968<br>TIGR03613 | Bacterial regulatory proteins, tetR family<br>betaine_BetI: transcriptional repressor BetI<br>BetI-type transcriptional repressor, C-terminal<br>mycofact_TetR: mycofactocin system transcriptional regulator<br>RutR: pyrimidine utilization regulatory protein R | 3.70E-16<br>2.60E-10<br>3.00E-09<br>7.60E-09<br>6.00E-08 |
| WP_018835207.1 | 64729 | 65707 | + | 325 | PF12698 | ABC2_membrane_3 | ABC-2 family transporter protein | 5.60E-07 |
| WP_027766442.1 | 65890 | 67780 | + | 629 | TIGR00976<br>PF02129<br>PF08530 | TIGR00976<br>Peptidase_S15<br>PepX_C | /NonD: hydrolase CocE/NonD family protein<br>X-Pro dipeptidyl-peptidase (S15 family)<br>X-Pro dipeptidyl-peptidase C-terminal non-catalytic domain | 3.50E-148<br>1.70E-53<br>5.70E-10 |
| WP_026276125.1 | 67994 | 68927 | + | 310 | TIGR00212<br>PF01379<br>PF03900 | TIGR00212<br>Porphobil_deam<br>Porphobil_deamC | hemC: hydroxymethylbilane synthase<br>Porphobilinogen deaminase, dipyromethane cofactor binding domain<br>Porphobilinogen deaminase, C-terminal domain | 2.10E-98<br>2.50E-64<br>5.00E-11 |
| WP_027766443.1 | 70653 | 69033 | - | 539 | PF01425<br>TIGR00132<br>TIGR02715<br>TIGR02713 | Amidase<br>TIGR00132<br>TIGR02715<br>TIGR02713 | Amidase<br>gatA: aspartyl/glutamyl-tRNA(Asn/Gln) amidotransferase, A subunit<br>amido_AtzE: amidohydrolase, AtzE family<br>allophanate_hyd: allophanate hydrolase | 9.80E-87<br>3.60E-59<br>2.10E-45<br>2.40E-42 |
| WP_018842285.1 | 71074 | 71773 | + | 232 | PF00196<br>TIGR03020<br>TIGR03541<br>TIGR02937<br>PF13936 | GerE<br>TIGR03020<br>TIGR03541<br>TIGR02937<br>HTH_38 | Bacterial regulatory proteins, luxR family<br>EpsA: transcriptional regulator EpsA<br>reg_near_HchA: LuxR family transcriptional regulatory, chaperone HchA-associated<br>sigma70-ECF: RNA polymerase sigma factor, sigma-70 family<br>Helix-turn-helix domain | 4.80E-19<br>7.50E-12<br>3.00E-10<br>2.20E-04<br>2.50E-04 |
| WP_027766444.1 | 72387 | 71829 | - | 185 | LinL | LinL | LinL | 2.70E-60 |
| WP_027766445.1 | 74402 | 72383 | - | 672 | LinH<br>LinE<br>LinG<br>PF00561<br>TIGR02427 | LinH<br>LinE<br>LinG<br>Abhydrolase_1<br>TIGR02427 | LinH (Fusion of LinE/LinG)<br>LinE Homolog<br>LinG Homolog<br>alpha/beta hydrolase fold<br>protocat_pcaD: 3-oxoadipate enol-lactonase | 5.30E-202<br>8.70E-94<br>2.60E-84<br>1.60E-11<br>1.60E-08 |
| WP_018835200.1 | 74866 | 74563 | - | 100 | LinA | LinA | LinA (linaridin precursor) | 3.30E-09 |
| WP_027766446.1 | 75079 | 76468 | + | 462 | PF00891<br>TIGR02716<br>PF16864<br>PF13649<br>PF08242 | Methyltransf_2<br>TIGR02716<br>Dimerisation2<br>Methyltransf_25<br>Methyltransf_12 | O-methyltransferase domain<br>C20_methyl_CrtF: C-20 methyltransferase BchU<br>Dimerisation domain<br>Methyltransferase domain<br>Methyltransferase domain | 3.10E-45<br>7.60E-16<br>7.90E-07<br>2.00E-06<br>6.70E-05 |
| WP_018835198.1 | 76445 | 76856 | + | 136 | PF04134 | DUF393 | Protein of unknown function, DUF393 | 2.10E-12 |
| WP_018835197.1 | 76940 | 77345 | + | 134 | PF12840<br>PF01022<br>PF12802<br>TIGR01884<br>PF01047 | HTH_20<br>HTH_5<br>MarR_2<br>TIGR01884<br>MarR | Helix-turn-helix domain<br>Bacterial regulatory protein, arsR family<br>MarR family<br>cas_HTH: CRISPR locus-related DNA-binding protein<br>MarR family | 1.20E-16<br>7.80E-10<br>1.00E-07<br>6.80E-07<br>2.10E-06 |
| WP_018835196.1 | 77341 | 78382 | + | 346 | TIGR03086<br>PF08327<br>TIGR03083<br>PF11716<br>PF10604 | TIGR03086<br>AHSA1<br>TIGR03083<br>MDMPI_N<br>Polyketide_cyc2 | TIGR03086: TIGR03086 family protein<br>Activator of Hsp90 ATPase homolog 1-like protein<br>TIGR03083: uncharacterized Actinobacterial protein TIGR03083<br>Mycothiol maleylpyruvate isomerase N-terminal domain<br>Polyketide cyclase / dehydrase and lipid transport | 2.40E-32<br>9.40E-24<br>2.60E-17<br>3.90E-07<br>1.80E-05 |
| WP_018835195.1 | 78587 | 79100 | + | 170 | PF01814 | Hemerythrin | Hemerythrin HHE cation binding domain | 2.30E-18 |
| WP_018835194.1 | 79806 | 79200 | - | 201 | NO PFAM MATCH | - | - | - |
| WP_027766447.1 | 79914 | 83106 | + | 1063 | PF03704<br>PF13424<br>PF00486<br>PF13191<br>PF13401 | BTAD<br>TPR_12<br>Trans_reg_C<br>AAA_16<br>AAA_22 | Bacterial transcriptional activator domain<br>Tetratricopeptide repeat<br>Transcriptional regulatory protein, C terminal<br>AAA ATPase domain<br>AAA domain | 2.10E-32<br>6.50E-15<br>1.80E-08<br>4.90E-07<br>3.40E-04 |

| index | leader | core | start | end | dir | score |
|---|---|---|---|---|---|---|
| 1 | MLLRNDGTELWGDGGWA | HPGKPHVAGATGTDAWSV | 65890 | 65782 | - | 6 |
| 2 | MTTTLQAPTTEDSRIAVWAQQHAAAHDEYFVAPTDRVVPVL | FTPLIALTVAGAVGCALGPVTSFVT...PDAPNGAALRGRSVDQLTGGHPGMV | 74866 | 74563 | - | 26 |

### Results for WP_078528871.1 [Streptomyces sp. CNS606] back to top

previous - next

#### Architecture

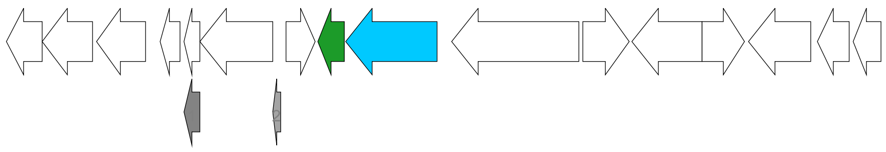

1000 nucleotides

Link to nucleotide sequence

| Accession | start | end | direction | length (aa) | Pfam/HMM | name | description | E-value |
|---|---|---|---|---|---|---|---|---|

| | | | | | | | | |
|---|---|---|---|---|---|---|---|---|
| WP_027763632.1 | 82387 | 81658 | - | 242 | TIGR03426<br>PF04093 | TIGR03426<br>MreD | shape_MreD: rod shape-determining protein MreD<br>rod shape-determining protein MreD | 1.80E-19<br>3.40E-04 |
| WP_051308209.1 | 83463 | 82392 | - | 356 | PF04085<br>TIGR00219 | MreC<br>TIGR00219 | rod shape-determining protein MreC<br>mreC: rod shape-determining protein MreC | 5.20E-33<br>9.30E-25 |
| WP_027763633.1 | 84579 | 83559 | - | 339 | PF06723<br>TIGR00904<br>TIGR01174<br>TIGR01991<br>PF00012 | MreB_Mbl<br>TIGR00904<br>TIGR01174<br>TIGR01991<br>HSP70 | MreB/Mbl protein<br>mreB: cell shape determining protein, MreB/Mrl family<br>ftsA: cell division protein FtsA<br>HscA: Fe-S protein assembly chaperone HscA<br>Hsp70 protein | 1.00E-155<br>2.90E-153<br>4.30E-18<br>6.00E-16<br>2.70E-14 |
| WP_027763634.1 | 85295 | 84881 | - | 137 | PF00334 | NDK | Nucleoside diphosphate kinase | 2.20E-50 |
| WP_027763635.1 | 85723 | 85375 | - | 115 | PF14017 | DUF4233 | Protein of unknown function (DUF4233) | 2.00E-32 |
| WP_027763636.1 | 87249 | 85728 | - | 506 | TIGR01499<br>TIGR01085<br>PF08245<br>TIGR02068<br>TIGR01082 | TIGR01499<br>TIGR01085<br>Mur_ligase_M<br>TIGR02068<br>TIGR01082 | folC: bifunctional protein FolC<br>murE: UDP-N-acetylmuramyl-tripeptide synthetase<br>Mur ligase middle domain<br>cya_phycin_syn: cyanophycin synthetase<br>murC: UDP-N-acetylmuramate--L-alanine ligase | 6.00E-98<br>1.20E-17<br>4.30E-14<br>2.70E-11<br>7.20E-11 |
| WP_037789893.1 | 87536 | 88154 | + | 205 | PF00196<br>TIGR03020<br>TIGR03541<br>TIGR02937 | GerE<br>TIGR03020<br>TIGR03541<br>TIGR02937 | Bacterial regulatory proteins, luxR family<br>EpsA: transcriptional regulator EpsA<br>reg_near_HchA: LuxR family transcriptional regulatory, chaperone HchA-associated<br>sigma70-ECF: RNA polymerase sigma factor, sigma-70 family | 1.80E-20<br>3.10E-11<br>2.00E-08<br>5.30E-04 |
| WP_051308211.1 | 88774 | 88201 | - | 190 | LinL | LinL | LinL | 1.40E-49 |
| WP_078528871.1 | 90710 | 88784 | - | 641 | LinH<br>LinE<br>LinG<br>TIGR02427<br>TIGR03611 | LinH<br>LinE<br>LinG<br>TIGR02427<br>TIGR03611 | LinH (Fusion of LinE/LinG)<br>LinE Homolog<br>LinG Homolog<br>protocat_pcaD: 3-oxoadipate enol-lactonase<br>RutD: pyrimidine utilization protein D | 5.80E-164<br>2.70E-91<br>8.30E-56<br>6.50E-11<br>3.50E-09 |
| WP_106965564.1 | 93698 | 91034 | - | 887 | TIGR00422<br>PF00133<br>TIGR00392<br>TIGR00395<br>TIGR00396 | TIGR00422<br>tRNA-synt_1<br>TIGR00392<br>TIGR00395<br>TIGR00396 | valS: valine--tRNA ligase<br>tRNA synthetases class I (I, L, M and V)<br>ileS: isoleucine--tRNA ligase<br>leuS_arch: leucine--tRNA ligase<br>leuS_bact: leucine--tRNA ligase | 0.00E+00<br>1.90E-196<br>2.50E-120<br>5.90E-57<br>1.10E-45 |
| WP_027763641.1 | 93803 | 94760 | + | 318 | NO PFAM MATCH | - | - | - |
| WP_027763642.1 | 96301 | 94837 | - | 487 | TIGR04461<br>PF00155<br>TIGR01140<br>TIGR01141<br>TIGR03947 | TIGR04461<br>Aminotran_1_2<br>TIGR01140<br>TIGR01141<br>TIGR03947 | endura_MppQ: enduracididine biosynthesis enzyme MppQ<br>Aminotransferase class I and II<br>L_thr_O3P_dcar: threonine-phosphate decarboxylase<br>hisC: histidinol-phosphate transaminase<br>viomycin_VioD: capreomycidine synthase | 1.00E-46<br>9.00E-27<br>8.00E-18<br>5.70E-17<br>9.70E-17 |
| WP_078528872.1 | 96299 | 97202 | + | 300 | PF00892<br>TIGR00950 | EamA<br>TIGR00950 | EamA-like transporter family<br>2A78: carboxylate/amino acid/amine transporter | 7.00E-21<br>1.20E-08 |
| WP_027763644.1 | 98584 | 97294 | - | 429 | TIGR00382<br>TIGR00390<br>PF07724<br>PF06689<br>PF00004 | TIGR00382<br>TIGR00390<br>AAA_2<br>zf-C4_ClpX<br>AAA | clpX: ATP-dependent Clp protease, ATP-binding subunit ClpX<br>hslU: ATP-dependent protease HslVU, ATPase subunit<br>AAA domain (Cdc48 subfamily)<br>ClpX C4-type zinc finger<br>ATPase family associated with various cellular activities (AAA) | 7.30E-197<br>3.70E-57<br>8.50E-51<br>1.10E-20<br>1.70E-16 |
| WP_027763645.1 | 99411 | 98721 | - | 229 | PF00574<br>TIGR00493<br>TIGR00706 | CLP_protease<br>TIGR00493<br>TIGR00706 | Clp protease<br>clpP: ATP-dependent Clp endopeptidase, proteolytic subunit ClpP<br>SppA_dom: signal peptide peptidase SppA, 36K type | 2.00E-75<br>3.10E-75<br>6.10E-04 |
| WP_106965565.1 | 100085 | 99479 | - | 201 | PF00574<br>TIGR00493<br>TIGR00705<br>TIGR00706 | CLP_protease<br>TIGR00493<br>TIGR00705<br>TIGR00706 | Clp protease<br>clpP: ATP-dependent Clp endopeptidase, proteolytic subunit ClpP<br>SppA_67K: signal peptide peptidase SppA, 67K type<br>SppA_dom: signal peptide peptidase SppA, 36K type | 1.30E-78<br>1.70E-76<br>6.90E-06<br>7.20E-06 |

| index | leader | core | start | end | dir | score |
|---|---|---|---|---|---|---|
| 1 | MRTLCASTLIGEFFVIGFAGLVAMQ...AVWAVSGTGMLLSLLLCGMLGRPGG | VQLGWALQIGLIASGFVVPAMFFLG...ASVHFGRKVDEAKAARAAAAEAETA | 85723 | 85375 | - | 7 |
| 2 | MNVRLGRDRDRVVPDFRPVRARTGRPAGAG | GPAGADRAAVAPAHRPDRAAVRAGSSVDWRT | 87431 | 87245 | - | 6 |

### Results for WP_018853313.1 [Streptomyces sp. CNY243] back to top

previous - next

#### Architecture

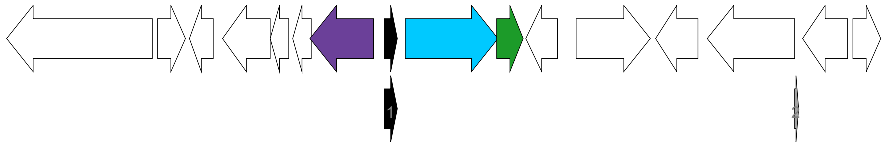

1000 nucleotides

Link to nucleotide sequence

| Accession | start | end | direction | length (aa) | Pfam/HMM | name | description | E-value |
|---|---|---|---|---|---|---|---|---|
| WP_018853310.1 | 144356 | 141167 | - | 1062 | PF03704<br>PF13424<br>PF00486<br>PF13191<br>PF13401 | BTAD<br>TPR_12<br>Trans_reg_C<br>AAA_16<br>AAA_22 | Bacterial transcriptional activator domain<br>Tetratricopeptide repeat<br>Transcriptional regulatory protein, C terminal<br>AAA ATPase domain<br>AAA domain | 1.30E-32<br>4.40E-14<br>1.90E-08<br>6.50E-07<br>5.00E-04 |
| WP_018835194.1 | 144464 | 145070 | + | 201 | NO PFAM MATCH | - | - | - |
| WP_018835195.1 | 145683 | 145170 | - | 170 | PF01814 | Hemerythrin | Hemerythrin HHE cation binding domain | 2.30E-18 |

| | | | | | | | | |
|---|---|---|---|---|---|---|---|---|
| WP_018853311.1 | 146929 | 145888 | - | 346 | TIGR03086<br>PF08327<br>TIGR03083<br>PF11716<br>PF10604 | TIGR03086<br>AHSA1<br>TIGR03083<br>MDMPI_N<br>Polyketide_cyc2 | TIGR03086: TIGR03086 family protein<br>Activator of Hsp90 ATPase homolog 1-like protein<br>TIGR03083: uncharacterized Actinobacterial protein TIGR03083<br>Mycothiol maleylpyruvate isomerase N-terminal domain<br>Polyketide cyclase / dehydrase and lipid transport | 4.90E-33<br>1.20E-23<br>3.80E-17<br>7.90E-08<br>3.10E-05 |
| WP_018835197.1 | 147330 | 146925 | - | 134 | PF12840<br>PF01022<br>PF12802<br>TIGR01884<br>PF01047 | HTH_20<br>HTH_5<br>MarR_2<br>TIGR01884<br>MarR | Helix-turn-helix domain<br>Bacterial regulatory protein, arsR family<br>MarR family<br>cas_HTH: CRISPR locus-related DNA-binding protein<br>MarR family | 1.20E-16<br>7.80E-10<br>1.00E-07<br>6.80E-07<br>2.10E-06 |
| WP_018835198.1 | 147825 | 147414 | - | 136 | PF04134 | DUF393 | Protein of unknown function, DUF393 | 2.10E-12 |
| WP_020610489.1 | 149203 | 147802 | - | 466 | PF00891<br>TIGR02716<br>PF16864<br>PF13649<br>PF08242 | Methyltransf_2<br>TIGR02716<br>Dimerisation2<br>Methyltransf_25<br>Methyltransf_12 | O-methyltransferase domain<br>C20_methyl_CrtF: C-20 methyltransferase BchU<br>Dimerisation domain<br>Methyltransferase domain<br>Methyltransferase domain | 5.10E-45<br>7.70E-16<br>8.10E-07<br>1.90E-06<br>6.30E-05 |
| WP_018835200.1 | 149416 | 149719 | + | 100 | LinA | LinA | LinA (linaridin precursor) | 3.30E-09 |
| WP_018853313.1 | 149880 | 151899 | + | 672 | LinH<br>LinE<br>LinG<br>PF00561<br>TIGR02427 | LinH<br>LinE<br>LinG<br>Abhydrolase_1<br>TIGR02427 | LinH (Fusion of LinE/LinG)<br>LinE Homolog<br>LinG Homolog<br>alpha/beta hydrolase fold<br>protocat_pcaD: 3-oxoadipate enol-lactonase | 6.80E-202<br>6.50E-94<br>2.60E-84<br>1.60E-11<br>1.60E-08 |
| WP_018835202.1 | 151895 | 152453 | + | 185 | LinL | LinL | LinL | 7.60E-60 |
| WP_018853315.1 | 153202 | 152509 | - | 230 | PF00196<br>TIGR03020<br>TIGR03541<br>TIGR02937<br>PF13936 | GerE<br>TIGR03020<br>TIGR03541<br>TIGR02937<br>HTH_38 | Bacterial regulatory proteins, luxR family<br>EpsA: transcriptional regulator EpsA<br>reg_near_HchA: LuxR family transcriptional regulatory, chaperone HchA-associated<br>sigma70-ECF: RNA polymerase sigma factor, sigma-70 family<br>Helix-turn-helix domain | 4.70E-19<br>7.40E-12<br>3.00E-10<br>2.10E-04<br>2.50E-04 |
| WP_018835204.1 | 153623 | 155243 | + | 539 | PF01425<br>TIGR00132<br>TIGR02715<br>TIGR02713 | Amidase<br>TIGR00132<br>TIGR02715<br>TIGR02713 | Amidase<br>gatA: aspartyl/glutamyl-tRNA(Asn/Gln) amidotransferase, A subunit<br>amido_AtzE: amidohydrolase, AtzE family<br>allophanate_hyd: allophanate hydrolase | 3.40E-88<br>1.30E-59<br>3.00E-45<br>1.60E-42 |
| WP_026278397.1 | 156282 | 155349 | - | 310 | TIGR00212<br>PF01379<br>PF03900 | TIGR00212<br>Porphobil_deam<br>Porphobil_deamC | hemC: hydroxymethylbilane synthase<br>Porphobilinogen deaminase, dipyromethane cofactor binding domain<br>Porphobilinogen deaminase, C-terminal domain | 6.90E-98<br>6.10E-64<br>4.60E-11 |
| WP_018853318.1 | 158386 | 156496 | - | 629 | TIGR00976<br>PF02129<br>PF08530 | TIGR00976<br>Peptidase_S15<br>PepX_C | /NonD: hydrolase CocE/NonD family protein<br>X-Pro dipeptidyl-peptidase (S15 family)<br>X-Pro dipeptidyl-peptidase C-terminal non-catalytic domain | 4.70E-148<br>1.70E-53<br>7.40E-10 |
| WP_018835207.1 | 159547 | 158569 | - | 325 | PF12698 | ABC2_membrane_3 | ABC-2 family transporter protein | 5.60E-07 |
| WP_018853319.1 | 159653 | 160268 | + | 204 | PF00440<br>TIGR03384<br>PF13977<br>TIGR03968<br>TIGR03613 | TetR_N<br>TIGR03384<br>TetR_C_6<br>TIGR03968<br>TIGR03613 | Bacterial regulatory proteins, tetR family<br>betaine_BetI: transcriptional repressor BetI<br>BetI-type transcriptional repressor, C-terminal<br>mycofact_TetR: mycofactocin system transcriptional regulator<br>RutR: pyrimidine utilization regulatory protein R | 3.70E-16<br>2.60E-10<br>3.00E-09<br>7.40E-09<br>6.00E-08 |

| index | leader | core | start | end | dir | score |
|---|---|---|---|---|---|---|
| 1 | MTTTLQAPTTEDSRIAVWAQQHAAAHDEYFVAPTDRVVPVL | FTPLIALTVAGAVGCALGPVTSFVT...PDAPNGAALRGRSVDQLTGGHPGMV | 149416 | 149719 | + | 26 |
| 2 | MLLRNDGTELWGDGGWA | HPGKPHVAGATGTDAWSV | 158386 | 158494 | + | 6 |

### Results for RZU87368.1 [Streptomyces sp. CNZ288] back to top

previous - next

#### Architecture

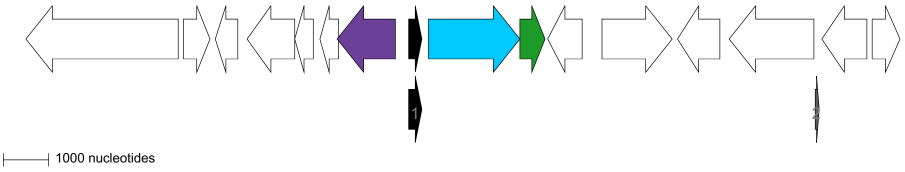

Link to nucleotide sequence

| Accession | start | end | direction | length (aa) | Pfam/HMM | name | description | E-value |
|---|---|---|---|---|---|---|---|---|
| RZU87360.1 | 1133579 | 1130207 | - | 1123 | PF03704<br>PF13424<br>PF00486<br>PF13191<br>PF13401 | BTAD<br>TPR_12<br>Trans_reg_C<br>AAA_16<br>AAA_22 | Bacterial transcriptional activator domain<br>Tetratricopeptide repeat<br>Transcriptional regulatory protein, C terminal<br>AAA ATPase domain<br>AAA domain | 6.80E-33<br>1.10E-13<br>4.20E-09<br>1.60E-06<br>2.20E-04 |
| RZU87361.1 | 1133687 | 1134293 | + | 201 | NO PFAM MATCH | - | - | - |
| RZU87362.1 | 1134906 | 1134393 | - | 170 | PF01814 | Hemerythrin | Hemerythrin HHE cation binding domain | 1.30E-18 |
| RZU87363.1 | 1136152 | 1135111 | - | 346 | TIGR03086<br>PF08327<br>TIGR03083<br>PF11716<br>PF10604 | TIGR03086<br>AHSA1<br>TIGR03083<br>MDMPI_N<br>Polyketide_cyc2 | TIGR03086: TIGR03086 family protein<br>Activator of Hsp90 ATPase homolog 1-like protein<br>TIGR03083: uncharacterized Actinobacterial protein TIGR03083<br>Mycothiol maleylpyruvate isomerase N-terminal domain<br>Polyketide cyclase / dehydrase and lipid transport | 2.70E-32<br>5.90E-24<br>1.70E-17<br>5.60E-07<br>1.40E-05 |
| RZU87364.1 | 1136553 | 1136148 | - | 134 | PF12840<br>PF01022<br>PF12802<br>TIGR01884<br>PF01047 | HTH_20<br>HTH_5<br>MarR_2<br>TIGR01884<br>MarR | Helix-turn-helix domain<br>Bacterial regulatory protein, arsR family<br>MarR family<br>cas_HTH: CRISPR locus-related DNA-binding protein<br>MarR family | 1.20E-16<br>8.00E-10<br>1.00E-07<br>8.40E-07<br>2.20E-06 |

| | | | | | | | | |
|---|---|---|---|---|---|---|---|---|
| RZU87365.1 | 1137104 | 1136693 | - | 136 | PF04134 | DUF393 | Protein of unknown function, DUF393 | 2.10E-12 |
| RZU87366.1 | 1138371 | 1137081 | - | 429 | PF00891 | Methyltransf_2 | O-methyltransferase domain | 1.20E-44 |
| | | | | | TIGR02716 | TIGR02716 | C20_methyl_CrtF: C-20 methyltransferase BchU | 2.30E-16 |
| | | | | | PF16864 | Dimerisation2 | Dimerisation domain | 5.40E-07 |
| | | | | | PF13649 | Methyltransf_25 | Methyltransferase domain | 2.20E-06 |
| | | | | | PF08242 | Methyltransf_12 | Methyltransferase domain | 4.20E-05 |
| RZU87367.1 | 1138650 | 1138953 | + | 100 | LinA | LinA | LinA (linaridin precursor) | 3.10E-09 |
| RZU87368.1 | 1139114 | 1141127 | + | 670 | LinH | LinH | LinH (Fusion of LinE/LinG) | 2.10E-201 |
| | | | | | LinE | LinE | LinE Homolog | 2.20E-94 |
| | | | | | LinG | LinG | LinG Homolog | 8.70E-84 |
| | | | | | PF12697 | Abhydrolase_6 | Alpha/beta hydrolase family | 4.80E-15 |
| | | | | | PF00561 | Abhydrolase_1 | alpha/beta hydrolase fold | 2.20E-11 |
| RZU87369.1 | 1141123 | 1141681 | + | 185 | LinL | LinL | LinL | 1.10E-59 |
| RZU87370.1 | 1142490 | 1141737 | - | 250 | PF00196 | GerE | Bacterial regulatory proteins, luxR family | 5.60E-19 |
| | | | | | TIGR03020 | TIGR03020 | EpsA: transcriptional regulator EpsA | 8.50E-12 |
| | | | | | TIGR03541 | TIGR03541 | reg_near_HchA: LuxR family transcriptional regulatory, chaperone HchA-associated | 3.50E-10 |
| | | | | | TIGR02937 | TIGR02937 | sigma70-ECF: RNA polymerase sigma factor, sigma-70 family | 2.40E-04 |
| | | | | | PF13936 | HTH_38 | Helix-turn-helix domain | 2.70E-04 |
| RZU87371.1 | 1142935 | 1144483 | + | 515 | PF01425 | Amidase | Amidase | 2.70E-88 |
| | | | | | TIGR00132 | TIGR00132 | gatA: aspartyl/glutamyl-tRNA(Asn/Gln) amidotransferase, A subunit | 9.10E-60 |
| | | | | | TIGR02715 | TIGR02715 | amido_AtzE: amidohydrolase, AtzE family | 6.80E-45 |
| | | | | | TIGR02713 | TIGR02713 | allophanate_hyd: allophanate hydrolase | 3.10E-42 |
| RZU87372.1 | 1145522 | 1144589 | - | 310 | TIGR00212 | TIGR00212 | hemC: hydroxymethylbilane synthase | 2.10E-98 |
| | | | | | PF01379 | Porphobil_deam | Porphobilinogen deaminase, dipyromethane cofactor binding domain | 2.50E-64 |
| | | | | | PF03900 | Porphobil_deamC | Porphobilinogen deaminase, C-terminal domain | 5.00E-11 |
| RZU87373.1 | 1147612 | 1145734 | - | 625 | TIGR00976 | TIGR00976 | /NonD: hydrolase CocE/NonD family protein | 1.80E-148 |
| | | | | | PF02129 | Peptidase_S15 | X-Pro dipeptidyl-peptidase (S15 family) | 1.60E-53 |
| | | | | | PF08530 | PepX_C | X-Pro dipeptidyl-peptidase C-terminal non-catalytic domain | 7.00E-10 |
| RZU87374.1 | 1148774 | 1147796 | - | 325 | PF12698 | ABC2_membrane_3 | ABC-2 family transporter protein | 2.50E-06 |
| RZU87375.1 | 1148880 | 1149495 | + | 204 | PF00440 | TetR_N | Bacterial regulatory proteins, tetR family | 3.70E-16 |
| | | | | | TIGR03384 | TIGR03384 | betaine_BetI: transcriptional repressor BetI | 3.00E-10 |
| | | | | | PF13977 | TetR_C_6 | BetI-type transcriptional repressor, C-terminal | 1.70E-09 |
| | | | | | TIGR03968 | TIGR03968 | mycofact_TetR: mycofactocin system transcriptional regulator | 8.30E-09 |
| | | | | | TIGR03613 | TIGR03613 | RutR: pyrimidine utilization regulatory protein R | 7.20E-08 |

| index | leader | core | start | end | dir | score |
|---|---|---|---|---|---|---|
| 1 | MTTTLQAPTTEDSRIAVWAQQHAADHDEHFVAPTDRVVPVL | FTPLIALTVAGAVGCALGPVTSFVT...PDAPNGAALRGRSVDQLTGGHPGMV | 1138650 | 1138953 | + | 26 |
| 2 | MTVRSCGGDGGWAHP | GKPHVAGATGTDAWSV | 1147625 | 1147721 | + | 8 |

### Results for WP_027759150.1 [Streptomyces sp. CNT318] back to top

previous - next

#### Architecture

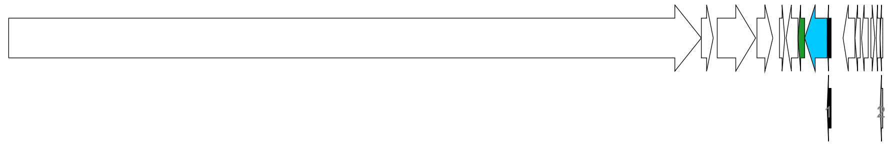

1000 nucleotides

Link to nucleotide sequence

| Accession | start | end | direction | length (aa) | Pfam/HMM | name | description | E-value |
|---|---|---|---|---|---|---|---|---|
| WP_158687847.1 | 0 | 61102 | + | 20366 | PF00691 | OmpA | OmpA family | 3.90E-04 |
| WP_063829181.1 | 61122 | 62103 | + | 326 | PF07179 | SseB | SseB protein N-terminal domain | 2.70E-04 |
| WP_051270988.1 | 62458 | 65914 | + | 1151 | NO PFAM MATCH | - | - | - |
| WP_027759148.1 | 65955 | 67428 | + | 490 | PF14200 | RicinB_lectin_2 | Ricin-type beta-trefoil lectin domain-like | 3.90E-19 |
| | | | | | PF00652 | Ricin_B_lectin | Ricin-type beta-trefoil lectin domain | 3.80E-16 |
| WP_019357370.1 | 68015 | 68477 | + | 153 | PF08327 | AHSA1 | Activator of Hsp90 ATPase homolog 1-like protein | 8.30E-07 |
| | | | | | PF10604 | Polyketide_cyc2 | Polyketide cyclase / dehydrase and lipid transport | 6.60E-06 |
| | | | | | PF03364 | Polyketide_cyc | Polyketide cyclase / dehydrase and lipid transport | 6.80E-04 |
| WP_027759149.1 | 69668 | 68540 | - | 375 | PF00891 | Methyltransf_2 | O-methyltransferase domain | 7.40E-44 |
| | | | | | TIGR02716 | TIGR02716 | C20_methyl_CrtF: C-20 methyltransferase BchU | 2.40E-16 |
| | | | | | PF16864 | Dimerisation2 | Dimerisation domain | 1.20E-08 |
| | | | | | PF12802 | MarR_2 | MarR family | 2.40E-05 |
| | | | | | TIGR03534 | TIGR03534 | RF_mod_PrmC: protein-(glutamine-N5) methyltransferase, release factor-specific | 1.10E-04 |
| WP_019357372.1 | 70267 | 69664 | - | 200 | LinL | LinL | LinL | 4.60E-59 |
| WP_027759150.1 | 72173 | 70253 | - | 639 | LinH | LinH | LinH (Fusion of LinE/LinG) | 4.00E-197 |
| | | | | | LinE | LinE | LinE Homolog | 8.50E-93 |
| | | | | | LinG | LinG | LinG Homolog | 2.20E-87 |
| | | | | | TIGR02427 | TIGR02427 | protocat_pcaD: 3-oxoadipate enol-lactonase | 4.50E-14 |
| | | | | | TIGR03056 | TIGR03056 | bchO_mg_che_rel: putative magnesium chelatase accessory protein | 4.00E-09 |
| WP_019357374.1 | 72526 | 72229 | - | 98 | LinA | LinA | LinA (linaridin precursor) | 1.50E-08 |

| | | | | | | | | |
|---|---|---|---|---|---|---|---|---|
| WP_078534059.1 | 74652 | 73554 | - | 365 | PF00144 | Beta-lactamase | Beta-lactamase | 2.50E-57 |
| WP_019357376.1 | 75170 | 74648 | - | 173 | PF01243<br>TIGR03618<br>TIGR03667<br>TIGR03666<br>TIGR03668 | Putative_PNPOx<br>TIGR03618<br>TIGR03667<br>TIGR03666<br>TIGR03668 | Pyridoxamine 5'-phosphate oxidase<br>Rv1155_F420: PPOX class probable F420-dependent enzyme<br>Rv3369: PPOX class probable F420-dependent enzyme, Rv3369 family<br>Rv2061_F420: PPOX class probable F420-dependent enzyme, Rv2061 family<br>Rv0121_F420: PPOX class probable F420-dependent enzyme, Rv0121 family | 2.70E-16<br>3.00E-16<br>1.10E-12<br>1.70E-07<br>4.60E-04 |
| WP_019357377.1 | 75844 | 75277 | - | 188 | PF03795 | YCII | YCII-related domain | 1.10E-20 |
| WP_027759151.1 | 76026 | 76407 | + | 126 | NO PFAM MATCH | - | - | - |
| WP_019357379.1 | 76825 | 76573 | - | 83 | PF04149 | DUF397 | Domain of unknown function (DUF397) | 2.80E-20 |
| WP_019357380.1 | 77074 | 76852 | - | 73 | PF04149 | DUF397 | Domain of unknown function (DUF397) | 2.10E-22 |

| index | leader | core | start | end | dir | score |
|---|---|---|---|---|---|---|
| 1 | MPITRESRAARWAQENAGEWCGHRVPEADEAVPVL | TTPIFATFAVAGAAGAGLGAARVVG...GSNEVSGYLEAGRPVKELLANHPAI | 72526 | 72229 | - | 26 |
| 2 | MANTEIDLQQAVWRKSSYSNGSGGE...AVETVDASVPVRDTKTAPHGPVLTF | PTAQWAHFISTLHEGNVNA | 77074 | 76852 | - | 6 |

### Results for RZU91185.1 [Streptomyces sp. CNZ301] back to top

previous - next

#### Architecture

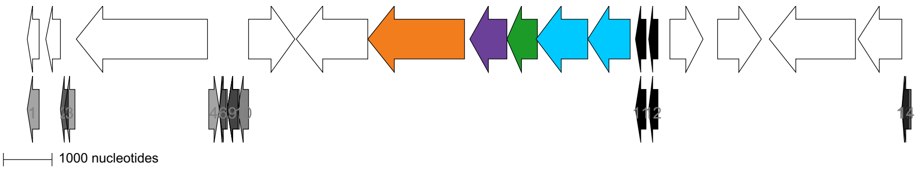

Link to nucleotide sequence

| Accession | start | end | direction | length (aa) | Pfam/HMM | name | description | E-value |
|---|---|---|---|---|---|---|---|---|
| RZU91177.1 | 4008688 | 4008427 | - | 86 | PF07362 | CcdA | Post-segregation antitoxin CcdA | 6.30E-05 |
| RZU91178.1 | 4009114 | 4008802 | - | 103 | PF11535 | Calci_bind_CcbP | Calcium binding | 2.10E-04 |
| RZU91179.1 | 4012143 | 4009452 | - | 896 | PF00196<br>PF13424<br>PF13384<br>PF13191<br>PF13176 | GerE<br>TPR_12<br>HTH_23<br>AAA_16<br>TPR_7 | Bacterial regulatory proteins, luxR family<br>Tetratricopeptide repeat<br>Homeodomain-like domain<br>AAA ATPase domain<br>Tetratricopeptide repeat | 1.20E-13<br>2.60E-09<br>9.20E-08<br>4.50E-07<br>6.80E-06 |
| RZU91180.1 | 4012975 | 4013947 | + | 323 | PF01978<br>TIGR03541<br>PF00196 | TrmB<br>TIGR03541<br>GerE | Sugar-specific transcriptional regulator TrmB<br>reg_near_HchA: LuxR family transcriptional regulatory, chaperone HchA-associated<br>Bacterial regulatory proteins, luxR family | 5.30E-07<br>4.10E-06<br>1.20E-05 |
| RZU91181.1 | 4015433 | 4013954 | - | 492 | TIGR02734<br>TIGR03467<br>PF13450<br>PF01593<br>TIGR01317 | TIGR02734<br>TIGR03467<br>NAD_binding_8<br>Amino_oxidase<br>TIGR01317 | crtI_fam: phytoene desaturase<br>HpnE: squalene-associated FAD-dependent desaturase<br>NAD(P)-binding Rossmann-like domain<br>Flavin containing amine oxidoreductase<br>GOGAT_sm_gam: glutamate synthase, NADH/NADPH, small subunit | 2.30E-12<br>5.10E-09<br>1.20E-08<br>1.50E-08<br>5.20E-08 |
| RZU91182.1 | 4017436 | 4015429 | - | 668 | TIGR02204<br>TIGR02203<br>TIGR00958<br>TIGR03796<br>TIGR03375 | TIGR02204<br>TIGR02203<br>TIGR00958<br>TIGR03796<br>TIGR03375 | MsbA_rel: ABC transporter, permease/ATP-binding protein<br>MsbA_lipidA: lipid A export permease/ATP-binding protein MsbA<br>3a01208: antigen peptide transporter 2<br>NHLM_micro_ABC1: NHLM bacteriocin system ABC transporter, peptidase/ATP-binding protein<br>type_I_sec_LssB: type I secretion system ATPase | 1.20E-101<br>2.30E-101<br>2.50E-90<br>8.70E-87<br>6.70E-82 |
| RZU91183.1 | 4018292 | 4017521 | - | 256 | PF13649<br>PF08241<br>PF13489<br>PF08242<br>PF13847 | Methyltransf_25<br>Methyltransf_11<br>Methyltransf_23<br>Methyltransf_12<br>Methyltransf_31 | Methyltransferase domain<br>Methyltransferase domain<br>Methyltransferase domain<br>Methyltransferase domain<br>Methyltransferase domain | 2.60E-17<br>8.30E-12<br>4.80E-10<br>2.70E-09<br>1.50E-07 |
| RZU91184.1 | 4018909 | 4018288 | - | 206 | LinL | LinL | LinL | 2.70E-71 |
| RZU91185.1 | 4019967 | 4018908 | - | 352 | LinH<br>LinE<br>PF12697<br>PF00561<br>PF12146 | LinH<br>LinE<br>Abhydrolase_6<br>Abhydrolase_1<br>Hydrolase_4 | LinH (Fusion of LinE/LinG)<br>LinE Homolog<br>Alpha/beta hydrolase family<br>alpha/beta hydrolase fold<br>Serine aminopeptidase, S33 | 8.10E-135<br>1.20E-121<br>1.00E-17<br>2.30E-10<br>3.00E-10 |
| RZU91186.1 | 4020836 | 4019963 | - | 290 | LinH<br>LinG | LinH<br>LinG | LinH (Fusion of LinE/LinG)<br>LinG Homolog | 1.30E-118<br>1.60E-113 |
| RZU91187.1 | 4021152 | 4020939 | - | 70 | LinA | LinA | LinA (linaridin precursor) | 3.20E-13 |
| RZU91188.1 | 4021397 | 4021205 | - | 63 | LinA | LinA | LinA (linaridin precursor) | 2.20E-11 |
| RZU91189.1 | 4021654 | 4022317 | + | 220 | PF05719 | GPP34 | Golgi phosphoprotein 3 (GPP34) | 2.00E-43 |
| RZU91190.1 | 4022639 | 4023539 | + | 299 | PF13391 | HNH_2 | HNH endonuclease | 1.50E-09 |
| RZU91191.1 | 4025460 | 4023699 | - | 586 | TIGR02403<br>PF00128<br>TIGR02456<br>TIGR02401<br>TIGR02402 | TIGR02403<br>Alpha-amylase<br>TIGR02456<br>TIGR02401<br>TIGR02402 | trehalose_treC: alpha,alpha-phosphotrehalase<br>Alpha amylase, catalytic domain<br>treS_nterm: trehalose synthase<br>trehalose_TreY: malto-oligosyltrehalose synthase<br>trehalose_TreZ: malto-oligosyltrehalose trehalohydrolase | 2.20E-203<br>2.60E-116<br>7.20E-98<br>2.30E-17<br>1.00E-16 |
| RZU91192.1 | 4026412 | 4025518 | - | 297 | PF00528<br>TIGR01581<br>TIGR02141<br>TIGR03262<br>TIGR00969 | BPD_transp_1<br>TIGR01581<br>TIGR02141<br>TIGR03262<br>TIGR00969 | Binding-protein-dependent transport system inner membrane component<br>Mo_ABC_porter: NifC-like ABC-type porter<br>modB_ABC: molybdate ABC transporter, permease protein<br>PhnU2: putative 2-aminoethylphosphonate ABC transporter, permease protein<br>3a0106s02: sulfate ABC transporter, permease protein | 4.60E-13<br>4.70E-13<br>1.80E-10<br>2.20E-08<br>5.70E-04 |

| index | leader | core | start | end | dir | score |
|---|---|---|---|---|---|---|
| 1 | MLAPMPDTTRITVTLPTEQVAELKKITDNISGYVAEAVARQIR | HQLLGADLRSHTDEHGAFSEEELAEARARILGTTSAAGGASAA | 4008688 | 4008427 | - | 6 |
| 2 | LPGPPLGRSLPNPTRQRHTPATASLGLTDGDGATT | DTPLLNRPVLNRTMRRIR | 4009280 | 4009118 | - | 8 |
| 3 | MAAHPAPNNSLTKADVSCRASGGAL...GSRTATLATHFRPVACRALPWAAHY | RTPPGNVTRRQPRALA | 4009414 | 4009198 | - | 6 |
| 4 | LYDKKREGMCVRHRVVFEPDRYAAG | RPPVTAGRPLSPPASGSGWSARAGADCLPRIAARSADAHSVSRR | 4012174 | 4012384 | + | 6 |
| 5 | LCVHYAVDLNGVAHRSHLWVLQSFS | SARSVWPPRRSDLPAPVTCLSILTGG | 4012526 | 4012370 | - | 7 |
| 6 | LPRLCVHYAVDLNGVAHRSHLWVLQSF | SSARSVWPPRRSDLPAPVTCLSILTGG | 4012535 | 4012370 | - | 6 |
| 7 | VSDPGGPGGRRTARTARTARTARAPARPCVGYC | PTRPARSISEIHRIRGSVSGGLSSVAGGLPPGR | 4012757 | 4012556 | - | 6 |
| 8 | VVSDPGGPGGRRTARTARTARTARAPARPCVGY | CPTRPARSISEIHRIRGSVSGGLSSVAGGLPPGR | 4012760 | 4012556 | - | 6 |
| 9 | VRELLWQVVSDPGGPGGRRTARTARTARTARAPARPC | VGYCPTRPARSISEIHRIRGSVSGGLSSVAGGLPPGR | 4012781 | 4012556 | - | 6 |
| 10 | LVSNPRPRCSWAGWAERPAPMRDGRRGGVGA | GTLRSTSRMGGLRAPHSGASSRGRATPLTGVGCA | 4012971 | 4012773 | - | 7 |
| 11 | MTAISTKVNASQELNGVSIHAGENECNGRMSPVL | ATPAAVGAGLAGAAAIAGAAAGGIAVGNAAGQAAGG | 4021152 | 4020939 | - | 29 |
| 12 | MLLEKVNNSHLLEGVLMDAANVEGLSPAEPVL | STPAAAVGGALVTAGAFGAGVGLGVTQAQAG | 4021397 | 4021205 | - | 31 |
| 13 | VPLTYMYKQAFEFLDFGYASALSFL | LAVAIAVVSFVQFHLFRDDGTGAPR | 4026561 | 4026408 | - | 8 |
| 14 | MTDGGPAGSTEVPLTYMYKQAFEFLDFGYA | SALSFLLAVAIAVVSFVQFHLFRDDGTGAPR | 4026594 | 4026408 | - | 8 |

### Results for PJJ50016.1 [Streptomyces sp. CNZ306] back to top

previous - next

#### Architecture

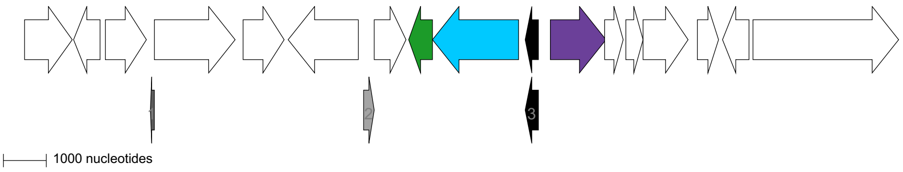

Link to nucleotide sequence

| Accession | start | end | direction | length (aa) | Pfam/HMM | name | description | E-value |
|---|---|---|---|---|---|---|---|---|
| PJJ50008.1 | 1398363 | 1399485 | + | 373 | PF00144<br>PF13354 | Beta-lactamase<br>Beta-lactamase2 | Beta-lactamase<br>Beta-lactamase enzyme family | 1.90E-61<br>5.80E-04 |
| PJJ50009.1 | 1400140 | 1399525 | - | 204 | PF00440<br>TIGR03384<br>PF13977<br>TIGR03968<br>TIGR03613 | TetR_N<br>TIGR03384<br>TetR_C_6<br>TIGR03968<br>TIGR03613 | Bacterial regulatory proteins, tetR family<br>betaine_BetI: transcriptional repressor BetI<br>BetI-type transcriptional repressor, C-terminal<br>mycofact_TetR: mycofactocin system transcriptional regulator<br>RutR: pyrimidine utilization regulatory protein R | 3.70E-16<br>2.60E-10<br>2.00E-09<br>7.60E-09<br>6.00E-08 |
| PJJ50010.1 | 1400246 | 1401224 | + | 325 | PF12698 | ABC2_membrane_3 | ABC-2 family transporter protein | 5.30E-07 |
| PJJ50011.1 | 1401408 | 1403286 | + | 625 | TIGR00976<br>PF02129<br>PF08530 | TIGR00976<br>Peptidase_S15<br>PepX_C | /NonD: hydrolase CocE/NonD family protein<br>X-Pro dipeptidyl-peptidase (S15 family)<br>X-Pro dipeptidyl-peptidase C-terminal non-catalytic domain | 1.80E-148<br>1.60E-53<br>7.20E-10 |
| PJJ50012.1 | 1403481 | 1404414 | + | 310 | TIGR00212<br>PF01379<br>PF03900 | TIGR00212<br>Porphobil_deam<br>Porphobil_deamC | hemC: hydroxymethylbilane synthase<br>Porphobilinogen deaminase, dipyromethane cofactor binding domain<br>Porphobilinogen deaminase, C-terminal domain | 2.90E-98<br>1.40E-64<br>5.00E-11 |
| PJJ50013.1 | 1406153 | 1404533 | - | 539 | PF01425<br>TIGR00132<br>TIGR02715<br>TIGR02713 | Amidase<br>TIGR00132<br>TIGR02715<br>TIGR02713 | Amidase<br>gatA: aspartyl/glutamyl-tRNA(Asn/Gln) amidotransferase, A subunit<br>amido_AtzE: amidohydrolase, AtzE family<br>allophanate_hyd: allophanate hydrolase | 5.20E-89<br>3.80E-60<br>3.50E-45<br>2.50E-42 |
| PJJ50014.1 | 1406526 | 1407261 | + | 244 | PF00196<br>TIGR03020<br>TIGR03541<br>TIGR02937<br>PF13936 | GerE<br>TIGR03020<br>TIGR03541<br>TIGR02937<br>HTH_38 | Bacterial regulatory proteins, luxR family<br>EpsA: transcriptional regulator EpsA<br>reg_near_HchA: LuxR family transcriptional regulatory, chaperone HchA-associated<br>sigma70-ECF: RNA polymerase sigma factor, sigma-70 family<br>Helix-turn-helix domain | 5.30E-19<br>8.10E-12<br>3.30E-10<br>2.30E-04<br>2.60E-04 |
| PJJ50015.1 | 1407875 | 1407317 | - | 185 | LinL | LinL | LinL | 1.20E-59 |
| PJJ50016.1 | 1409896 | 1407871 | - | 674 | LinH<br>LinE<br>LinG<br>PF00561<br>TIGR02427 | LinH<br>LinE<br>LinG<br>Abhydrolase_1<br>TIGR02427 | LinH (Fusion of LinE/LinG)<br>LinE Homolog<br>LinG Homolog<br>alpha/beta hydrolase fold<br>protocat_pcaD: 3-oxoadipate enol-lactonase | 2.60E-201<br>5.30E-94<br>9.90E-84<br>1.60E-11<br>1.60E-08 |
| PJJ50017.1 | 1410360 | 1410057 | - | 100 | LinA | LinA | LinA (linaridin precursor) | 3.10E-09 |
| PJJ50018.1 | 1410639 | 1411929 | + | 429 | PF00891<br>TIGR02716<br>PF16864<br>PF13649<br>PF08242 | Methyltransf_2<br>TIGR02716<br>Dimerisation2<br>Methyltransf_25<br>Methyltransf_12 | O-methyltransferase domain<br>C20_methyl_CrtF: C-20 methyltransferase BchU<br>Dimerisation domain<br>Methyltransferase domain<br>Methyltransferase domain | 5.00E-45<br>1.30E-16<br>2.70E-07<br>1.10E-06<br>3.20E-05 |
| PJJ50019.1 | 1411906 | 1412317 | + | 136 | PF04134 | DUF393 | Protein of unknown function, DUF393 | 2.20E-12 |

| | | | | | | | | |
|---|---|---|---|---|---|---|---|---|
| PJJ50020.1 | 1412402 | 1412804 | + | 133 | PF12840<br>PF01022<br>PF12802<br>TIGR01884<br>PF01047 | HTH_20<br>HTH_5<br>MarR_2<br>TIGR01884<br>MarR | Helix-turn-helix domain<br>Bacterial regulatory protein, arsR family<br>MarR family<br>cas_HTH: CRISPR locus-related DNA-binding protein<br>MarR family | 1.20E-16<br>7.80E-10<br>1.00E-07<br>6.60E-07<br>2.10E-06 |
| PJJ50021.1 | 1412800 | 1413841 | + | 346 | TIGR03086<br>PF08327<br>TIGR03083<br>PF11716<br>PF10604 | TIGR03086<br>AHSA1<br>TIGR03083<br>MDMPI_N<br>Polyketide_cyc2 | TIGR03086: TIGR03086 family protein<br>Activator of Hsp90 ATPase homolog 1-like protein<br>TIGR03083: uncharacterized Actinobacterial protein TIGR03083<br>Mycothiol maleylpyruvate isomerase N-terminal domain<br>Polyketide cyclase / dehydrase and lipid transport | 2.40E-32<br>8.50E-24<br>2.60E-17<br>4.70E-07<br>1.30E-05 |
| PJJ50022.1 | 1414046 | 1414559 | + | 170 | PF01814 | Hemerythrin | Hemerythrin HHE cation binding domain | 1.30E-18 |
| PJJ50023.1 | 1415265 | 1414659 | - | 201 | NO PFAM MATCH | - | - | - |
| PJJ50024.1 | 1415372 | 1418745 | + | 1123 | PF03704<br>PF13424<br>PF00486<br>PF13191<br>PF13401 | BTAD<br>TPR_12<br>Trans_reg_C<br>AAA_16<br>AAA_22 | Bacterial transcriptional activator domain<br>Tetratricopeptide repeat<br>Transcriptional regulatory protein, C terminal<br>AAA ATPase domain<br>AAA domain | 4.20E-32<br>1.50E-11<br>1.10E-08<br>9.80E-07<br>3.50E-04 |

| index | leader | core | start | end | dir | score |
|---|---|---|---|---|---|---|
| 1 | MTVRSCGGDGGWAHP | GKPHVAGATGTDAWSV | 1401395 | 1401299 | - | 8 |
| 2 | VGRKSATIQGCFRHKRAPRGGRAAKSRPFRARSAAGETGRTGQNPPGPTR | GTPTGTAVACELTSRVDCATVSVSVR | 1406289 | 1406520 | + | 6 |
| 3 | MTTTLQAPTTEDSRIAVWAQQHAADHDEHFVAPTDRVVPVL | FTPLIALTVAGAVGCALGPVTSFVT...PDAPNGAALRGRSVDQLTGGHPGMV | 1410360 | 1410057 | - | 26 |

### Results for RZU91753.1 [Streptomyces sp. CNZ301] back to top

previous - next

#### Architecture

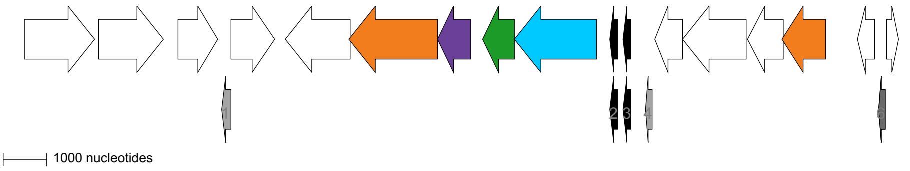

Link to nucleotide sequence

| Accession | start | end | direction | length (aa) | Pfam/HMM | name | description | E-value |
|---|---|---|---|---|---|---|---|---|
| RZU91745.1 | 4691247 | 4692879 | + | 543 | NO PFAM MATCH | - | - | - |
| RZU91746.1 | 4692962 | 4694459 | + | 498 | NO PFAM MATCH | - | - | - |
| RZU91747.1 | 4694795 | 4695740 | + | 314 | PF01259<br>TIGR00081<br>TIGR02735 | SAICAR_synt<br>TIGR00081<br>TIGR02735 | SAICAR synthetase<br>purC: phosphoribosylaminoimidazolesuccinocarboxamide synthase<br>purC_vibrio: phosphoribosylaminoimidazole-succinocarboxamide synthase | 3.50E-99<br>2.70E-69<br>3.20E-23 |
| RZU91748.1 | 4696028 | 4697051 | + | 340 | PF00196<br>PF13384 | GerE<br>HTH_23 | Bacterial regulatory proteins, luxR family<br>Homeodomain-like domain | 8.40E-05<br>4.20E-04 |
| RZU91749.1 | 4698776 | 4697297 | - | 492 | TIGR02734<br>TIGR03467<br>PF13450<br>TIGR00292<br>PF01593 | TIGR02734<br>TIGR03467<br>NAD_binding_8<br>TIGR00292<br>Amino_oxidase | crtI_fam: phytoene desaturase<br>HpnE: squalene-associated FAD-dependent desaturase<br>NAD(P)-binding Rossmann-like domain<br>TIGR00292: thiazole biosynthesis enzyme<br>Flavin containing amine oxidoreductase | 9.30E-11<br>5.60E-09<br>1.10E-07<br>6.60E-07<br>1.30E-06 |
| RZU91750.1 | 4700809 | 4698772 | - | 678 | TIGR02204<br>TIGR02203<br>TIGR00958<br>TIGR03796<br>TIGR03375 | TIGR02204<br>TIGR02203<br>TIGR00958<br>TIGR03796<br>TIGR03375 | MsbA_rel: ABC transporter, permease/ATP-binding protein<br>MsbA_lipidA: lipid A export permease/ATP-binding protein MsbA<br>3a01208: antigen peptide transporter 2<br>NHLM_micro_ABC1: NHLM bacteriocin system ABC transporter, peptidase/ATP-binding protein<br>type_I_sec_LssB: type I secretion system ATPase | 1.70E-99<br>4.80E-97<br>1.10E-83<br>4.40E-81<br>1.20E-80 |
| RZU91751.1 | 4701575 | 4700816 | - | 252 | PF13649<br>PF13489<br>PF08241<br>PF13847<br>TIGR01983 | Methyltransf_25<br>Methyltransf_23<br>Methyltransf_11<br>Methyltransf_31<br>TIGR01983 | Methyltransferase domain<br>Methyltransferase domain<br>Methyltransferase domain<br>Methyltransferase domain<br>UbiG: 3-demethylubiquinone-9 3-O-methyltransferase | 8.40E-18<br>1.70E-13<br>2.10E-12<br>1.10E-11<br>1.00E-10 |
| RZU91752.1 | 4702577 | 4701860 | - | 238 | LinL | LinL | LinL | 1.10E-62 |
| RZU91753.1 | 4704487 | 4702573 | - | 637 | LinH<br>LinG<br>LinE<br>PF12697<br>PF00561 | LinH<br>LinG<br>LinE<br>Abhydrolase_6<br>Abhydrolase_1 | LinH (Fusion of LinE/LinG)<br>LinG Homolog<br>LinE Homolog<br>Alpha/beta hydrolase family<br>alpha/beta hydrolase fold | 4.30E-247<br>2.60E-120<br>2.80E-97<br>6.30E-15<br>3.50E-09 |
| RZU91754.1 | 4704983 | 4704803 | - | 59 | LinA | LinA | LinA (linaridin precursor) | 2.60E-16 |
| RZU91755.1 | 4705270 | 4705096 | - | 57 | LinA | LinA | LinA (linaridin precursor) | 2.90E-15 |
| RZU91756.1 | 4706489 | 4705835 | - | 217 | PF00072<br>TIGR02875<br>PF00196<br>TIGR02154<br>TIGR02956 | Response_reg<br>TIGR02875<br>GerE<br>TIGR02154<br>TIGR02956 | Response regulator receiver domain<br>spore_0_A: sporulation transcription factor Spo0A<br>Bacterial regulatory proteins, luxR family<br>PhoB: phosphate regulon transcriptional regulatory protein PhoB<br>TMAO_torS: TMAO reductase sytem sensor TorS | 2.00E-25<br>2.40E-13<br>4.50E-13<br>3.90E-12<br>1.90E-10 |
| RZU91757.1 | 4707958 | 4706485 | - | 490 | PF07730 | HisKA_3 | Histidine kinase | 4.30E-15 |

| | | | | | | | | |
|---|---|---|---|---|---|---|---|---|
| RZU91758.1 | 4708792 | 4707979 | - | 270 | PF01061<br>PF12698<br>TIGR03062<br>TIGR01291 | ABC2_membrane<br>ABC2_membrane_3<br>TIGR03062<br>TIGR01291 | ABC-2 type transporter<br>ABC-2 family transporter protein<br>pip_yhgE_Cterm: YhgE/Pip C-terminal domain<br>nodJ: ABC-2 type transporter, NodJ family | 2.90E-19<br>1.80E-11<br>1.10E-07<br>3.10E-04 |
| RZU91759.1 | 4709775 | 4708788 | - | 328 | TIGR01188<br>TIGR03864<br>TIGR01288<br>TIGR04406<br>TIGR03740 | TIGR01188<br>TIGR03864<br>TIGR01288<br>TIGR04406<br>TIGR03740 | drrA: daunorubicin resistance ABC transporter, ATP-binding protein<br>PQQ_ABC_ATP: ABC transporter, ATP-binding subunit, PQQ-dependent alcohol dehydrogenase system<br>nodI: nodulation ABC transporter NodI<br>LPS_export_lptB: LPS export ABC transporter ATP-binding protein<br>galliderm_ABC: lantibiotic protection ABC transporter, ATP-binding subunit | 8.70E-70<br>3.30E-49<br>1.80E-47<br>3.90E-39<br>2.60E-36 |
| RZU91760.1 | 4710935 | 4710509 | - | 141 | PF11774 | Lsr2 | Lsr2 | 1.30E-34 |
| RZU91761.1 | 4711209 | 4711467 | + | 85 | PF02700<br>TIGR00302 | PurS<br>TIGR00302 | Phosphoribosylformylglycinamidine (FGAM) synthase<br>TIGR00302: phosphoribosylformylglycinamidine synthase, purS protein | 1.30E-27<br>6.30E-20 |

| index | leader | core | start | end | dir | score |
|---|---|---|---|---|---|---|
| 1 | VNSPCPAAPPGPVHVGGGQKLSTHMRNRTPSIEKTKAPNTPRR | DTFSKGPFINCASAPVCTLPDARFNSR | 4696027 | 4695814 | - | 6 |
| 2 | MKDVTSQINESAELASVDIADAEGVVFTDYTPVM | GTPGAFLGGVAIASATVAAFEAGQD | 4704983 | 4704803 | - | 29 |
| 3 | MLEEKLNLQIQDQLHEDHEGEIAEEAAPVL | ATPALAGAFVGGAAVVTGAYAAGKAVG | 4705270 | 4705096 | - | 30 |
| 4 | VLRDAPRCVNQAHTRTNSDHSAAVH | HDGTSIPEADIFGYPLLTLMPILKD | 4705774 | 4705621 | - | 6 |
| 5 | VRAASEATRPASLPATKIYARGYLPRT | LTAPAFHRRPTAPAPPGALGKMP | 4711173 | 4711020 | - | 6 |
| 6 | LVRAASEATRPASLPATKIYARGYLPRT | LTAPAFHRRPTAPAPPGALGKMP | 4711176 | 4711020 | - | 6 |

### Results for WP_078894000.1 [Streptomyces sp. CT34] back to top

previous - next

#### Architecture

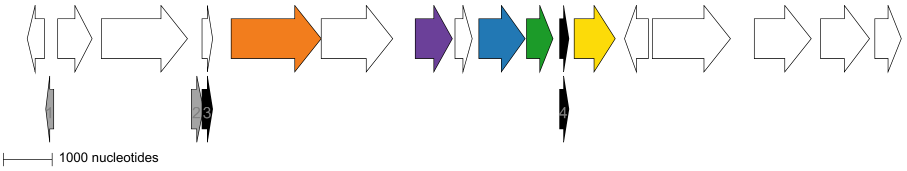

Link to nucleotide sequence

| Accession | start | end | direction | length (aa) | Pfam/HMM | name | description | E-value |
|---|---|---|---|---|---|---|---|---|
| WP_043265476.1 | 4618527 | 4618161 | - | 121 | PF01906<br>TIGR03884 | YbjQ_1<br>TIGR03884 | Putative heavy-metal-binding<br>sel_bind_Methan: selenium-binding protein | 9.30E-34<br>7.70E-04 |
| WP_043265477.1 | 4618782 | 4619502 | + | 239 | PF09335 | SNARE_assoc | SNARE associated Golgi protein | 3.20E-24 |
| WP_043265478.1 | 4619682 | 4621455 | + | 590 | PF06738<br>PF12821 | ThrE<br>ThrE_2 | Putative threonine/serine exporter<br>Threonine/Serine exporter, ThrE | 2.90E-73<br>1.40E-18 |
| WP_043265479.1 | 4621759 | 4621951 | + | 63 | NO PFAM MATCH | - | - | - |
| WP_052230046.1 | 4622332 | 4624183 | + | 616 | TIGR02204<br>TIGR02203<br>TIGR00958<br>TIGR03796<br>TIGR03375 | TIGR02204<br>TIGR02203<br>TIGR00958<br>TIGR03796<br>TIGR03375 | MsbA_rel: ABC transporter, permease/ATP-binding protein<br>MsbA_lipidA: lipid A export permease/ATP-binding protein MsbA<br>3a01208: antigen peptide transporter 2<br>NHLM_micro_ABC1: NHLM bacteriocin system ABC transporter, peptidase/ATP-binding protein<br>type_I_sec_LssB: type I secretion system ATPase | 6.30E-99<br>2.10E-98<br>8.30E-89<br>1.70E-80<br>6.50E-76 |
| WP_043270372.1 | 4624179 | 4625664 | + | 494 | TIGR02734<br>PF01593<br>PF13450<br>TIGR03467<br>TIGR00292 | TIGR02734<br>Amino_oxidase<br>NAD_binding_8<br>TIGR03467<br>TIGR00292 | crtI_fam: phytoene desaturase<br>Flavin containing amine oxidoreductase<br>NAD(P)-binding Rossmann-like domain<br>HpnE: squalene-associated FAD-dependent desaturase<br>TIGR00292: thiazole biosynthesis enzyme | 1.50E-09<br>3.00E-09<br>3.60E-09<br>2.10E-08<br>3.30E-08 |
| WP_107068144.1 | 4626132 | 4626891 | + | 252 | PF13649<br>PF08241<br>PF13489<br>PF08242<br>PF13847 | Methyltransf_25<br>Methyltransf_11<br>Methyltransf_23<br>Methyltransf_12<br>Methyltransf_31 | Methyltransferase domain<br>Methyltransferase domain<br>Methyltransferase domain<br>Methyltransferase domain<br>Methyltransferase domain | 5.30E-16<br>1.30E-12<br>1.90E-11<br>2.50E-09<br>1.70E-08 |
| WP_078893998.1 | 4626930 | 4627278 | + | 115 | PF01527<br>PF13518<br>PF13011<br>PF13384<br>PF05344 | HTH_Tnp_1<br>HTH_28<br>LZ_Tnp_IS481<br>HTH_23<br>DUF746 | Transposase<br>Helix-turn-helix domain<br>leucine-zipper of insertion element IS481<br>Homeodomain-like domain<br>Domain of Unknown Function (DUF746) | 2.90E-08<br>2.80E-07<br>1.80E-05<br>1.90E-05<br>1.50E-04 |
| WP_078894000.1 | 4627425 | 4628370 | + | 314 | LinE<br>LinH<br>PF12697<br>PF00561<br>TIGR02427 | LinE<br>LinH<br>Abhydrolase_6<br>Abhydrolase_1<br>TIGR02427 | LinE Homolog<br>LinH (Fusion of LinE/LinG)<br>Alpha/beta hydrolase family<br>alpha/beta hydrolase fold<br>protocat_pcaD: 3-oxoadipate enol-lactonase | 5.30E-96<br>2.30E-51<br>5.50E-14<br>3.40E-10<br>6.90E-09 |
| WP_078894692.1 | 4628393 | 4628954 | + | 186 | LinL | LinL | LinL | 4.40E-57 |
| WP_043265481.1 | 4629079 | 4629277 | + | 65 | LinA | LinA | LinA (linaridin precursor) | 6.10E-11 |
| WP_043265482.1 | 4629375 | 4630230 | + | 284 | LinG<br>LinH<br>TIGR04033 | LinG<br>LinH<br>TIGR04033 | LinG Homolog<br>LinH (Fusion of LinE/LinG)<br>export_SdpB: antimicrobial peptide system protein, SdpB family | 6.70E-105<br>1.20E-76<br>4.00E-05 |
| WP_019888410.1 | 4630905 | 4630410 | - | 164 | PF00719 | Pyrophosphatase | Inorganic pyrophosphatase | 2.20E-53 |

| | | | | | | | | |
|---|---|---|---|---|---|---|---|---|
| WP_043265483.1 | 4630991 | 4632581 | + | 529 | PF02113<br>TIGR00666 | Peptidase_S13<br>TIGR00666 | D-Ala-D-Ala carboxypeptidase 3 (S13) family<br>PBP4: D-alanyl-D-alanine carboxypeptidase/D-alanyl-D-alanine-endopeptidase | 1.70E-76<br>5.00E-58 |
| WP_043265484.1 | 4633076 | 4634225 | + | 382 | TIGR03883<br>TIGR03624<br>PF10103 | TIGR03883<br>TIGR03624<br>Zincin_2 | DUF2342_F420: uncharacterized protein, coenzyme F420 biosynthesis associated<br>TIGR03624: putative hydrolase<br>Zincin-like metallopeptidase | 9.40E-141<br>1.70E-123<br>1.20E-120 |
| WP_043265485.1 | 4634420 | 4635428 | + | 335 | TIGR02432<br>PF01171<br>PF09179 | TIGR02432<br>ATP_bind_3<br>TilS | lysidine_TilS_N: tRNA(Ile)-lysidine synthetase<br>PP-loop family<br>TilS substrate binding domain | 3.20E-43<br>5.50E-38<br>5.30E-12 |
| WP_043270376.1 | 4635541 | 4636081 | + | 179 | TIGR01203<br>PF00156<br>TIGR01251<br>TIGR01367<br>TIGR01091 | TIGR01203<br>Pribosyltran<br>TIGR01251<br>TIGR01367<br>TIGR01091 | HGPRTase: hypoxanthine phosphoribosyltransferase<br>Phosphoribosyl transferase domain<br>ribP_PPkin: ribose-phosphate diphosphokinase<br>pyrE_Therm: orotate phosphoribosyltransferase<br>upp: uracil phosphoribosyltransferase | 3.20E-65<br>1.50E-25<br>9.60E-06<br>4.80E-04<br>8.10E-04 |

| index | leader | core | start | end | dir | score |
|---|---|---|---|---|---|---|
| 1 | VCGPKLSTDYAASWFQPEDQEPAEPRAPAPRPRG | SAPATRPRPAAGRPRRPG | 4618711 | 4618552 | - | 6 |
| 2 | VAVAGILRSGKSTSDPGRAFRRAAD...LRAASNSSPEFSCRVRTTLSVEVVH | ATFMAFGRELPVEFAAPLNSEKKVAQL | 4621520 | 4621763 | + | 6 |
| 3 | MNSPVPTQISSYANDHLAKVTPGKVDNGQAVPALSGAKAQISA | ITPTFTDATVLIGIQQAVGH | 4621759 | 4621951 | + | 18 |
| 4 | MSVLAEFANTELVDVEPGRLGSEATPTM | ITPLATLATPEATPVGFAATSATAAAVNMITHDVTRH | 4629079 | 4629277 | + | 33 |

### Results for GFH39492.1 [Streptomyces sp. CWH03] back to top

previous - next

#### Architecture

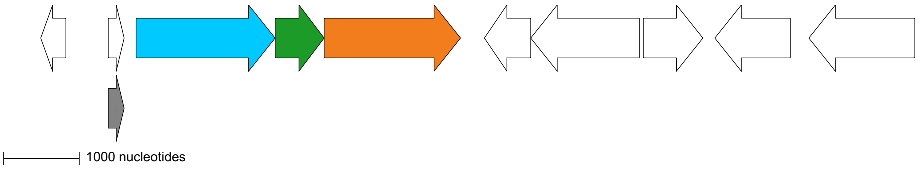

Link to nucleotide sequence

| Accession | start | end | direction | length (aa) | Pfam/HMM | name | description | E-value |
|---|---|---|---|---|---|---|---|---|
| GFH39490.1 | 1151 | 818 | - | 110 | NO PFAM MATCH | - | - | - |
| GFH39491.1 | 1718 | 1919 | + | 66 | NO PFAM MATCH | - | - | - |
| GFH39492.1 | 2079 | 3909 | + | 609 | LinH<br>LinE<br>LinG<br>PF12697<br>TIGR03695 | LinH<br>LinE<br>LinG<br>Abhydrolase_6<br>TIGR03695 | LinH (Fusion of LinE/LinG)<br>LinE Homolog<br>LinG Homolog<br>Alpha/beta hydrolase family<br>menH_SHCHC: 2-succinyl-6-hydroxy-2,4-cyclohexadiene-1-carboxylate synthase | 6.60E-227<br>2.80E-106<br>8.00E-106<br>1.70E-19<br>5.00E-14 |
| GFH39493.1 | 3908 | 4565 | + | 218 | LinL | LinL | LinL | 6.70E-64 |
| GFH39494.1 | 4561 | 6367 | + | 601 | TIGR02203<br>TIGR02204<br>TIGR01846<br>TIGR00958<br>TIGR03796 | TIGR02203<br>TIGR02204<br>TIGR01846<br>TIGR00958<br>TIGR03796 | MsbA_lipidA: lipid A export permease/ATP-binding protein MsbA<br>MsbA_rel: ABC transporter, permease/ATP-binding protein<br>type_I_sec_HlyB: type I secretion system ATPase<br>3a01208: antigen peptide transporter 2<br>NHLM_micro_ABC1: NHLM bacteriocin system ABC transporter, peptidase/ATP-binding protein | 3.10E-94<br>8.50E-94<br>4.20E-84<br>2.30E-81<br>7.50E-81 |
| GFH39495.1 | 7293 | 6678 | - | 204 | PF17940<br>TIGR03384<br>PF00440 | TetR_C_31<br>TIGR03384<br>TetR_N | Tetracyclin repressor-like, C-terminal domain<br>betaine_BetI: transcriptional repressor BetI<br>Bacterial regulatory proteins, tetR family | 1.80E-18<br>1.30E-12<br>7.70E-10 |
| GFH39496.1 | 8732 | 7298 | - | 477 | PF01425<br>TIGR02715<br>TIGR00132<br>TIGR02713 | Amidase<br>TIGR02715<br>TIGR00132<br>TIGR02713 | Amidase<br>amido_AtzE: amidohydrolase, AtzE family<br>gatA: aspartyl/glutamyl-tRNA(Asn/Gln) amidotransferase, A subunit<br>allophanate_hyd: allophanate hydrolase | 3.70E-108<br>8.60E-74<br>4.20E-60<br>3.60E-56 |
| GFH39497.1 | 8782 | 9577 | + | 264 | PF00106<br>PF13561<br>TIGR01963<br>TIGR01830<br>TIGR03971 | adh_short<br>adh_short_C2<br>TIGR01963<br>TIGR01830<br>TIGR03971 | short chain dehydrogenase<br>Enoyl-(Acyl carrier protein) reductase<br>PHB_DH: 3-hydroxybutyrate dehydrogenase<br>3oxo_ACP_reduc: 3-oxoacyl-[acyl-carrier-protein] reductase<br>SDR_subfam_1: SDR family mycofactocin-dependent oxidoreductase | 7.30E-38<br>1.80E-29<br>8.60E-29<br>5.90E-24<br>2.90E-20 |
| GFH39498.1 | 10720 | 9718 | - | 333 | TIGR03842<br>PF00296<br>TIGR03555<br>TIGR03857<br>TIGR04024 | TIGR03842<br>Bac_luciferase<br>TIGR03555<br>TIGR03857<br>TIGR04024 | F420_CPS_4043: F420-dependent oxidoreductase, CPS_4043 family<br>Luciferase-like monooxygenase<br>F420_mer: 5,10-methylenetetrahydromethanopterin reductase<br>F420_MSMEG_2249: probable F420-dependent oxidoreductase, MSMEG_2249 family<br>F420_NP1902A: coenzyme F420-dependent oxidoreductase, NP1902A family | 2.00E-176<br>6.20E-55<br>7.70E-32<br>9.70E-26<br>6.60E-17 |
| GFH39499.1 | 12371 | 10967 | - | 467 | TIGR02033<br>TIGR03178<br>TIGR00857<br>PF01979<br>PF07969 | TIGR02033<br>TIGR03178<br>TIGR00857<br>Amidohydro_1<br>Amidohydro_3 | D-hydantoinase: dihydropyrimidinase<br>allantoinase: allantoinase<br>pyrC_multi: dihydroorotase, multifunctional complex type<br>Amidohydrolase family<br>Amidohydrolase family | 1.40E-166<br>4.40E-82<br>6.50E-53<br>3.80E-22<br>1.50E-16 |

| index | leader | core | start | end | dir | score |
|---|---|---|---|---|---|---|
| 1 | MDLLHRVNCDSALA | EVPILAEQAIQVKGTEPILNIFNDM...FVNAVTQPAGEVTADTMEVTMATIG | 1718 | 1919 | + | 7 |

### Results for WP_045555727.1 [Streptomyces sp. FxanaA7] back to top

previous - next

#### Architecture

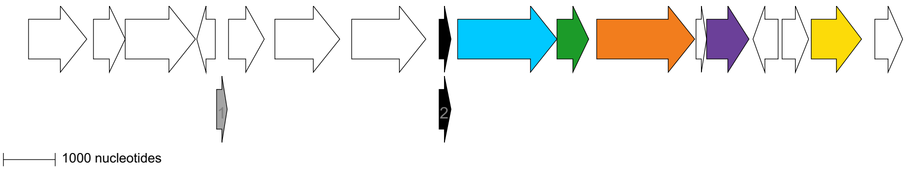

[Link to nucleotide sequence]

| Accession | start | end | direction | length (aa) | Pfam/HMM | name | description | E-value |
|---|---|---|---|---|---|---|---|---|
| WP_045555722.1 | 767391 | 768513 | + | 373 | PF07730 | HisKA_3 | Histidine kinase | 1.60E-12 |
| WP_078853284.1 | 768640 | 769246 | + | 201 | TIGR02983<br>TIGR02937<br>PF08281<br>TIGR02954<br>PF04545 | TIGR02983<br>TIGR02937<br>Sigma70_r4_2<br>TIGR02954<br>Sigma70_r4 | SigE-fam_strep: RNA polymerase sigma-70 factor, sigma-E family<br>sigma70-ECF: RNA polymerase sigma factor, sigma-70 family<br>Sigma-70, region 4<br>Sig70_famx3: RNA polymerase sigma-70 factor, TIGR02954 family<br>Sigma-70, region 4 | 1.90E-54<br>2.60E-18<br>1.50E-16<br>4.30E-14<br>2.70E-12 |
| WP_045555723.1 | 769242 | 770598 | + | 451 | PF13360<br>PF01011<br>TIGR03300<br>PF13570<br>TIGR03075 | PQQ_2<br>PQQ<br>TIGR03300<br>PQQ_3<br>TIGR03075 | PQQ-like domain<br>PQQ enzyme repeat<br>assembly_YfgL: outer membrane assembly lipoprotein YfgL<br>PQQ-like domain<br>PQQ_enz_alc_DH: PQQ-dependent dehydrogenase, methanol/ethanol family | 3.50E-21<br>3.90E-13<br>1.70E-12<br>5.20E-09<br>4.80E-07 |
| WP_045555724.1 | 770991 | 770646 | - | 114 | PF00196<br>TIGR03541<br>TIGR03020<br>PF08281 | GerE<br>TIGR03541<br>TIGR03020<br>Sigma70_r4_2 | Bacterial regulatory proteins, luxR family<br>reg_near_HchA: LuxR family transcriptional regulatory, chaperone HchA-associated<br>EpsA: transcriptional regulator EpsA<br>Sigma-70, region 4 | 1.80E-15<br>1.90E-12<br>5.80E-09<br>2.50E-07 |
| WP_045555880.1 | 771246 | 771936 | + | 229 | PF00072<br>PF00196<br>TIGR02875<br>TIGR03020<br>TIGR03541 | Response_reg<br>GerE<br>TIGR02875<br>TIGR03020<br>TIGR03541 | Response regulator receiver domain<br>Bacterial regulatory proteins, luxR family<br>spore_0_A: sporulation transcription factor Spo0A<br>EpsA: transcriptional regulator EpsA<br>reg_near_HchA: LuxR family transcriptional regulatory, chaperone HchA-associated | 1.40E-24<br>9.80E-18<br>7.00E-16<br>1.10E-12<br>2.80E-08 |
| WP_045555725.1 | 772152 | 773382 | + | 409 | PF07730<br>PF13581 | HisKA_3<br>HATPase_c_2 | Histidine kinase<br>Histidine kinase-like ATPase domain | 9.80E-14<br>3.80E-04 |
| WP_045555726.1 | 773628 | 775059 | + | 476 | TIGR02734<br>PF13450<br>TIGR03467<br>TIGR00292<br>TIGR03315 | TIGR02734<br>NAD_binding_8<br>TIGR03467<br>TIGR00292<br>TIGR03315 | crtI_fam: phytoene desaturase<br>NAD(P)-binding Rossmann-like domain<br>HpnE: squalene-associated FAD-dependent desaturase<br>TIGR00292: thiazole biosynthesis enzyme<br>Se_ygfK: putative selenate reductase, YgfK subunit | 2.50E-12<br>3.80E-12<br>3.70E-08<br>2.00E-07<br>3.20E-07 |
| WP_045555881.1 | 775307 | 775547 | + | 79 | LinA | LinA | LinA (linaridin precursor) | 1.30E-04 |
| WP_045555727.1 | 775661 | 777581 | + | 639 | LinH<br>LinG<br>LinE<br>PF12697<br>PF12146 | LinH<br>LinG<br>LinE<br>Abhydrolase_6<br>Hydrolase_4 | LinH (Fusion of LinE/LinG)<br>LinG Homolog<br>LinE Homolog<br>Alpha/beta hydrolase family<br>Serine aminopeptidase, S33 | 3.00E-247<br>3.40E-157<br>7.50E-92<br>1.20E-15<br>3.10E-08 |
| WP_045555728.1 | 777577 | 778198 | + | 206 | LinL | LinL | LinL | 1.10E-68 |
| WP_045555729.1 | 778361 | 780254 | + | 630 | TIGR02204<br>TIGR02203<br>TIGR00958<br>TIGR03796<br>TIGR03375 | TIGR02204<br>TIGR02203<br>TIGR00958<br>TIGR03796<br>TIGR03375 | MsbA_rel: ABC transporter, permease/ATP-binding protein<br>MsbA_lipidA: lipid A export permease/ATP-binding protein MsbA<br>3a01208: antigen peptide transporter 2<br>NHLM_micro_ABC1: NHLM bacteriocin system ABC transporter, peptidase/ATP-binding protein<br>type_I_sec_LssB: type I secretion system ATPase | 2.30E-106<br>2.10E-104<br>2.00E-92<br>4.70E-86<br>2.00E-81 |
| WP_045555730.1 | 780279 | 780471 | + | 63 | NO PFAM MATCH | - | - | - |
| WP_045555731.1 | 780467 | 781295 | + | 275 | PF13649<br>PF08241<br>PF08242<br>PF13489<br>TIGR01983 | Methyltransf_25<br>Methyltransf_11<br>Methyltransf_12<br>Methyltransf_23<br>TIGR01983 | Methyltransferase domain<br>Methyltransferase domain<br>Methyltransferase domain<br>Methyltransferase domain<br>UbiG: 3-demethylubiquinone-9 3-O-methyltransferase | 1.60E-15<br>1.30E-12<br>4.30E-07<br>5.20E-07<br>3.10E-05 |
| WP_045555732.1 | 781851 | 781374 | - | 158 | PF01638 | HxlR | HxlR-like helix-turn-helix | 9.70E-20 |
| WP_045555733.1 | 781940 | 782435 | + | 164 | TIGR00369<br>PF03061<br>TIGR02286<br>PF14539<br>PF13622 | TIGR00369<br>4HBT<br>TIGR02286<br>DUF4442<br>4HBT_3 | unchar_dom_1: uncharacterized domain 1<br>Thioesterase superfamily<br>PaaD: phenylacetic acid degradation protein PaaD<br>Domain of unknown function (DUF4442)<br>Thioesterase-like superfamily | 5.50E-27<br>2.50E-16<br>1.90E-09<br>2.20E-08<br>5.90E-05 |
| WP_045555734.1 | 782499 | 783471 | + | 323 | PF00561<br>PF12697<br>TIGR03056<br>TIGR02427<br>TIGR03343 | Abhydrolase_1<br>Abhydrolase_6<br>TIGR03056<br>TIGR02427<br>TIGR03343 | alpha/beta hydrolase fold<br>Alpha/beta hydrolase family<br>bchO_mg_che_rel: putative magnesium chelatase accessory protein<br>protocat_pcaD: 3-oxoadipate enol-lactonase<br>biphenyl_bphD: 2-hydroxy-6-oxo-6-phenylhexa-2,4-dienoate hydrolase | 1.90E-27<br>1.10E-19<br>3.70E-17<br>2.30E-16<br>9.40E-16 |
| WP_045555735.1 | 783718 | 784255 | + | 178 | PF13302<br>PF00583 | Acetyltransf_3<br>Acetyltransf_1 | Acetyltransferase (GNAT) domain<br>Acetyltransferase (GNAT) family | 9.60E-07<br>8.40E-04 |

| index | leader | core | start | end | dir | score |
|---|---|---|---|---|---|---|
| 1 | VGVEDPRAVRLLRCDSSVTSGRRASTIRPIPGSSHRPIGLGPPLRG | GTFSGLAPTARFGGGVRPAGFP | 771011 | 771218 | + | 6 |
| 2 | MEIPVEGVVYVAARPTL | GTPRIARIGRIAQAAEGVGAIAAAA...NNLAAEANAQNAAALAAVGGTPPAV | 775307 | 775547 | + | 28 |

### Results for WP_157878906.1 [Streptomyces sp. CT34] back to top

previous - next

#### Architecture

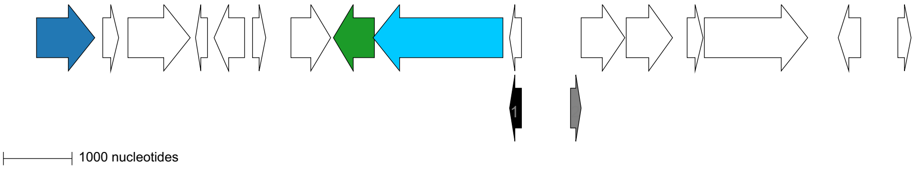

Link to nucleotide sequence

| Accession | start | end | direction | length (aa) | Pfam/HMM | name | description | E-value |
|---|---|---|---|---|---|---|---|---|
| WP_157878905.1 | 6362977 | 6363820 | + | 280 | LinE<br>LinH<br>PF12697<br>TIGR02427<br>PF12146 | LinE<br>LinH<br>Abhydrolase_6<br>TIGR02427<br>Hydrolase_4 | LinE Homolog<br>LinH (Fusion of LinE/LinG)<br>Alpha/beta hydrolase family<br>protocat_pcaD: 3-oxoadipate enol-lactonase<br>Serine aminopeptidase, S33 | 7.70E-32<br>7.30E-29<br>4.20E-20<br>5.00E-16<br>4.00E-12 |
| WP_020556386.1 | 6363939 | 6364158 | + | 73 | INFERRED GENE | - | - | - |
| WP_014182363.1 | 6364302 | 6365218 | + | 305 | INFERRED GENE | - | - | - |
| WP_052230180.1 | 6365469 | 6365289 | - | 59 | NO PFAM MATCH | - | - | - |
| WP_043266789.1 | 6366011 | 6365558 | - | 150 | NO PFAM MATCH | - | - | - |
| WP_043266791.1 | 6366108 | 6366315 | + | 68 | NO PFAM MATCH | - | - | - |
| WP_043266793.1 | 6366678 | 6367248 | + | 189 | NO PFAM MATCH | - | - | - |
| WP_063772705.1 | 6367888 | 6367291 | - | 198 | LinL | LinL | LinL | 1.60E-59 |
| WP_157878906.1 | 6369762 | 6367884 | - | 625 | LinH<br>LinG<br>LinE<br>PF12697<br>PF00561 | LinH<br>LinG<br>LinE<br>Abhydrolase_6<br>Abhydrolase_1 | LinH (Fusion of LinE/LinG)<br>LinG Homolog<br>LinE Homolog<br>Alpha/beta hydrolase family<br>alpha/beta hydrolase fold | 4.70E-239<br>2.60E-123<br>6.20E-94<br>3.60E-18<br>5.10E-12 |
| WP_157878907.1 | 6370030 | 6369868 | - | 53 | NO PFAM MATCH | - | - | - |
| WP_043266795.1 | 6370905 | 6371541 | + | 211 | NO PFAM MATCH | - | - | - |
| WP_043266798.1 | 6371565 | 6372243 | + | 225 | NO PFAM MATCH | - | - | - |
| WP_008746506.1 | 6372455 | 6372686 | + | 77 | INFERRED GENE | - | - | - |
| WP_003999825.1 | 6372699 | 6374216 | + | 505 | INFERRED GENE | - | - | - |
| WP_043266800.1 | 6374990 | 6374663 | - | 108 | PF11774 | Lsr2 | Lsr2 | 1.60E-33 |
| WP_043262638.1 | 6375514 | 6375724 | + | 69 | NO PFAM MATCH | - | - | - |

| index | leader | core | start | end | dir | score |
|---|---|---|---|---|---|---|
| 1 | MEDMKKIESFANSTLTLPMGEEADKFIQEVAPVIT | STLSSDMNINNHGCVVGW | 6370030 | 6369868 | - | 13 |
| 2 | VIAVEPPTARGSVSSGCGRSRGSIE | DLASAITGLGDTATETDLTTEEAWP | 6370762 | 6370915 | + | 7 |

### Results for WP_023423357.1 [Streptomyces sp. GBA 94-10] back to top

previous - next

#### Architecture

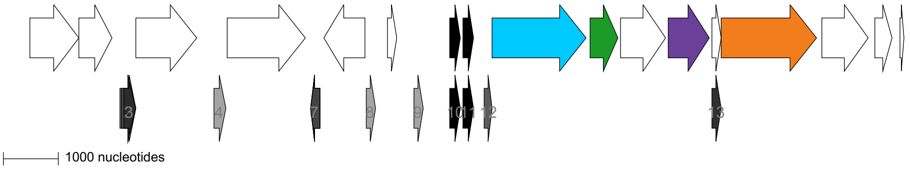

Link to nucleotide sequence

| Accession | start | end | direction | length (aa) | Pfam/HMM | name | description | E-value |
|---|---|---|---|---|---|---|---|---|
| WP_158698165.1 | 10490 | 11375 | + | 294 | PF07730<br>PF13581 | HisKA_3<br>HATPase_c_2 | Histidine kinase<br>Histidine kinase-like ATPase domain | 1.50E-09<br>1.60E-04 |
| WP_023423353.1 | 11380 | 11989 | + | 202 | PF00072<br>TIGR02875<br>TIGR02154<br>PF00196<br>TIGR01818 | Response_reg<br>TIGR02875<br>TIGR02154<br>GerE<br>TIGR01818 | Response regulator receiver domain<br>spore_0_A: sporulation transcription factor Spo0A<br>PhoB: phosphate regulon transcriptional regulatory protein PhoB<br>Bacterial regulatory proteins, luxR family<br>ntrC: nitrogen regulation protein NR(I) | 8.70E-24<br>7.70E-16<br>6.70E-12<br>3.10E-10<br>6.10E-09 |
| WP_076611826.1 | 12408 | 13519 | + | 370 | INFERRED GENE | - | - | - |
| WP_023423354.1 | 14068 | 15511 | + | 480 | TIGR02734<br>PF13450<br>TIGR03467<br>PF01593<br>TIGR01317 | TIGR02734<br>NAD_binding_8<br>TIGR03467<br>Amino_oxidase<br>TIGR01317 | crtI_fam: phytoene desaturase<br>NAD(P)-binding Rossmann-like domain<br>HpnE: squalene-associated FAD-dependent desaturase<br>Flavin containing amine oxidoreductase<br>GOGAT_sm_gam: glutamate synthase, NADH/NADPH, small subunit | 5.00E-11<br>1.50E-09<br>3.50E-08<br>7.00E-08<br>8.30E-07 |
| WP_023423355.1 | 16587 | 15831 | - | 251 | PF13359<br>PF13613<br>PF01609 | DDE_Tnp_4<br>HTH_Tnp_4<br>DDE_Tnp_1 | DDE superfamily endonuclease<br>Helix-turn-helix of DDE superfamily endonuclease<br>Transposase DDE domain | 3.70E-28<br>4.00E-14<br>1.10E-11 |
| WP_012377440.1 | 16996 | 17170 | + | 58 | INFERRED GENE | - | - | - |
| WP_037788156.1 | 18138 | 18330 | + | 63 | LinA | LinA | LinA (linaridin precursor) | 3.30E-14 |
| WP_023423356.1 | 18372 | 18573 | + | 66 | LinA | LinA | LinA (linaridin precursor) | 3.80E-11 |

| | | | | | | | | |
|---|---|---|---|---|---|---|---|---|
| WP_023423357.1 | 18908 | 20612 | + | 567 | LinH<br>LinE<br>LinG<br>PF12697<br>PF00561 | LinH<br>LinE<br>LinG<br>Abhydrolase_6<br>Abhydrolase_1 | LinH (Fusion of LinE/LinG)<br>LinE Homolog<br>LinG Homolog<br>Alpha/beta hydrolase family<br>alpha/beta hydrolase fold | 1.30E-229<br>1.80E-97<br>2.90E-96<br>3.50E-16<br>7.80E-10 |
| WP_078539199.1 | 20680 | 21190 | + | 169 | LinL | LinL | LinL | 2.90E-66 |
| WP_023423359.1 | 21241 | 22060 | + | 272 | PF12680 | SnoaL_2 | SnoaL-like domain | 3.00E-10 |
| WP_023423360.1 | 22115 | 22868 | + | 250 | PF13649<br>PF08241<br>PF08242<br>PF13847<br>PF13489 | Methyltransf_25<br>Methyltransf_11<br>Methyltransf_12<br>Methyltransf_31<br>Methyltransf_23 | Methyltransferase domain<br>Methyltransferase domain<br>Methyltransferase domain<br>Methyltransferase domain<br>Methyltransferase domain | 5.20E-13<br>1.20E-10<br>5.40E-09<br>1.00E-07<br>4.00E-06 |
| WP_157875472.1 | 22910 | 23087 | + | 58 | NO PFAM MATCH | - | - | - |
| WP_023423361.1 | 23083 | 24817 | + | 577 | TIGR02204<br>TIGR02203<br>TIGR01846<br>TIGR00958<br>TIGR03796 | TIGR02204<br>TIGR02203<br>TIGR01846<br>TIGR00958<br>TIGR03796 | MsbA_rel: ABC transporter, permease/ATP-binding protein<br>MsbA_lipidA: lipid A export permease/ATP-binding protein MsbA<br>type_I_sec_HlyB: type I secretion system ATPase<br>3a01208: antigen peptide transporter 2<br>NHLM_micro_ABC1: NHLM bacteriocin system ABC transporter, peptidase/ATP-binding protein | 8.80E-108<br>2.40E-107<br>2.20E-90<br>2.80E-90<br>1.90E-87 |
| WP_023423362.1 | 24899 | 25751 | + | 283 | PF01551 | Peptidase_M23 | Peptidase family M23 | 2.00E-15 |
| WP_023423363.1 | 25880 | 26186 | + | 101 | PF01527<br>PF13518 | HTH_Tnp_1<br>HTH_28 | Transposase<br>Helix-turn-helix domain | 4.90E-12<br>1.10E-05 |
| WP_078539201.1 | 26329 | 26410 | + | 26 | PF13276 | HTH_21 | HTH-like domain | 2.40E-07 |

| index | leader | core | start | end | dir | score |
|---|---|---|---|---|---|---|
| 1 | VPGAHRQTMVGRLVGPVDTCICVAG...HVPPAGRARRRVVREGRGAARQLRV | TTLADTRERLPRNGSTAGTWPSTAVPGEASMFFVK | 12134 | 12416 | + | 7 |
| 2 | MVGRLVGPVDTCICVAGGRRHVRRRHVPPAGRARRRVVREGRGAARQLRV | TTLADTRERLPRNGSTAGTWPSTAVPGEASMFFVK | 12158 | 12416 | + | 7 |
| 3 | VAGGRRHVRRRHVPPAGRARRRVVREGRGAARQLRV | TTLADTRERLPRNGSTAGTWPSTAVPGEASMFFVK | 12200 | 12416 | + | 6 |
| 4 | LAAEPRGRGGNRVTARGGTCHPAGRARQAVGVV | RSSCECDLRHNKDARVLVVSLHVHDGGSGAGPP | 13845 | 14046 | + | 6 |
| 5 | MQPAGAGADHSNYTGSGFTGMCVVR | SFARFVGGRRFARVGRAFRTSGRAS | 15761 | 15608 | - | 6 |
| 6 | MMQPAGAGADHSNYTGSGFTGMCVV | RSFARFVGGRRFARVGRAFRTSGRAS | 15764 | 15608 | - | 6 |
| 7 | VMMQPAGAGADHSNYTGSGFTGMCVV | RSFARFVGGRRFARVGRAFRTSGRAS | 15767 | 15608 | - | 6 |
| 8 | VDLGREPVYQELRRSADPSNTRPAPSGWKR | LTAAVNAYTSPAAAGESAAGRGDHRRAPPQ | 16606 | 16789 | + | 6 |
| 9 | VPCGPSRAVARARLAGLNELLQLEPRS | RTANRGWPARTLSTRSLIGGRVEEAR | 17483 | 17645 | + | 6 |
| 10 | MGKIAAFMNQQELGSLEDGLTKTDGTPVM | ATPAVAVPVTVKITAGIVAFTIPVGAYVAGRVVN | 18138 | 18330 | + | 32 |
| 11 | MTHIIESVRSITEPTLTEGQKDRLTRTMPVA | FTPAVLVTTPYAGGGLAAGAAVVGAYELGRAAGGK | 18372 | 18573 | + | 29 |
| 12 | LPRRFSLRCRYPGTRLWVCLSGFPRSLLS | PPPSNTCTSTSRWILVVSTTGKS | 18753 | 18912 | + | 8 |
| 13 | MTADRAVWFAVAFVFAAPSMSAFFREDGR | FTRRAVALGVVFATSVAAIVAIAFGQAPS | 22910 | 23087 | + | 9 |

### Results for WP_159508791.1 [Streptomyces sp. GS7] back to top

previous - next

#### Architecture

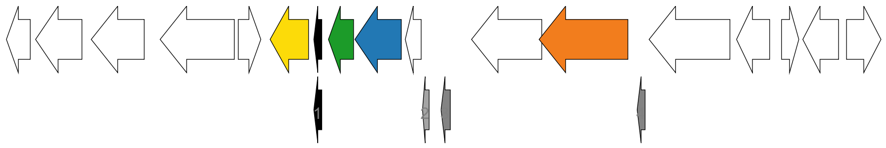

1000 nucleotides

Link to nucleotide sequence

| Accession | start | end | direction | length (aa) | Pfam/HMM | name | description | E-value |
|---|---|---|---|---|---|---|---|---|
| WP_159510475.1 | 8546000 | 8545460 | - | 179 | TIGR01203<br>PF00156<br>TIGR01251<br>TIGR01367<br>TIGR01091 | TIGR01203<br>Pribosyltran<br>TIGR01251<br>TIGR01367<br>TIGR01091 | HGPRTase: hypoxanthine phosphoribosyltransferase<br>Phosphoribosyl transferase domain<br>ribP_PPkin: ribose-phosphate diphosphokinase<br>pyrE_Therm: orotate phosphoribosyltransferase<br>upp: uracil phosphoribosyltransferase | 1.80E-65<br>5.80E-26<br>9.80E-06<br>3.70E-04<br>4.20E-04 |
| WP_159508782.1 | 8547121 | 8546113 | - | 335 | TIGR02432<br>PF01171<br>PF09179 | TIGR02432<br>ATP_bind_3<br>TilS | lysidine_TilS_N: tRNA(Ile)-lysidine synthetase<br>PP-loop family<br>TilS substrate binding domain | 4.10E-43<br>4.00E-38<br>2.20E-12 |
| WP_159508784.1 | 8548465 | 8547316 | - | 382 | TIGR03883<br>TIGR03624<br>PF10103 | TIGR03883<br>TIGR03624<br>Zincin_2 | DUF2342_F420: uncharacterized protein, coenzyme F420 biosynthesis associated<br>TIGR03624: putative hydrolase<br>Zincin-like metallopeptidase | 1.30E-139<br>5.90E-122<br>2.00E-119 |
| WP_159508786.1 | 8550440 | 8548826 | - | 537 | PF02113<br>TIGR00666 | Peptidase_S13<br>TIGR00666 | D-Ala-D-Ala carboxypeptidase 3 (S13) family<br>PBP4: D-alanyl-D-alanine carboxypeptidase/D-alanyl-D-alanine-endopeptidase | 2.70E-78<br>4.40E-59 |
| WP_019888410.1 | 8550526 | 8551021 | + | 164 | PF00719 | Pyrophosphatase | Inorganic pyrophosphatase | 2.20E-53 |
| WP_159508788.1 | 8552071 | 8551213 | - | 285 | LinG<br>LinH<br>TIGR04033 | LinG<br>LinH<br>TIGR04033 | LinG Homolog<br>LinH (Fusion of LinE/LinG)<br>export_SdpB: antimicrobial peptide system protein, SdpB family | 1.30E-107<br>1.80E-78<br>4.40E-05 |

| | | | | | | | | |
|---|---|---|---|---|---|---|---|---|
| WP_159508789.1 | 8552362 | 8552167 | - | 64 | LinA | LinA | LinA (linaridin precursor) | 1.40E-11 |
| WP_159510476.1 | 8553052 | 8552491 | - | 186 | LinL | LinL | LinL | 4.90E-56 |
| WP_159508791.1 | 8554095 | 8553075 | - | 339 | LinE | LinE | LinE Homolog | 4.90E-102 |
| | | | | | LinH | LinH | LinH (Fusion of LinE/LinG) | 1.50E-53 |
| | | | | | PF12697 | Abhydrolase_6 | Alpha/beta hydrolase family | 8.20E-14 |
| | | | | | TIGR02427 | TIGR02427 | protocat_pcaD: 3-oxoadipate enol-lactonase | 6.00E-11 |
| | | | | | PF00561 | Abhydrolase_1 | alpha/beta hydrolase fold | 8.20E-11 |
| WP_159508793.1 | 8554515 | 8554167 | - | 115 | PF01527 | HTH_Tnp_1 | Transposase | 4.90E-08 |
| | | | | | PF13518 | HTH_28 | Helix-turn-helix domain | 2.90E-07 |
| | | | | | PF13011 | LZ_Tnp_IS481 | leucine-zipper of insertion element IS481 | 1.40E-05 |
| | | | | | PF13384 | HTH_23 | Homeodomain-like domain | 2.00E-05 |
| | | | | | PF05344 | DUF746 | Domain of Unknown Function (DUF746) | 1.40E-04 |
| WP_159508795.1 | 8557159 | 8555632 | - | 508 | TIGR03467 | TIGR03467 | HpnE: squalene-associated FAD-dependent desaturase | 3.40E-09 |
| | | | | | TIGR00292 | TIGR00292 | TIGR00292: thiazole biosynthesis enzyme | 7.80E-09 |
| | | | | | TIGR02734 | TIGR02734 | crtI_fam: phytoene desaturase | 6.40E-08 |
| | | | | | PF13450 | NAD_binding_8 | NAD(P)-binding Rossmann-like domain | 4.10E-07 |
| | | | | | TIGR03315 | TIGR03315 | Se_ygfK: putative selenate reductase, YgfK subunit | 6.70E-07 |
| WP_159508796.1 | 8559030 | 8557095 | - | 644 | TIGR02203 | TIGR02203 | MsbA_lipidA: lipid A export permease/ATP-binding protein MsbA | 6.90E-97 |
| | | | | | TIGR02204 | TIGR02204 | MsbA_rel: ABC transporter, permease/ATP-binding protein | 8.70E-97 |
| | | | | | TIGR00958 | TIGR00958 | 3a01208: antigen peptide transporter 2 | 5.10E-90 |
| | | | | | TIGR03796 | TIGR03796 | NHLM_micro_ABC1: NHLM bacteriocin system ABC transporter, peptidase/ATP-binding protein | 3.10E-80 |
| | | | | | TIGR03375 | TIGR03375 | type_I_sec_LssB: type I secretion system ATPase | 1.50E-75 |
| WP_159508798.1 | 8561268 | 8559495 | - | 590 | PF06738 | ThrE | Putative threonine/serine exporter | 1.90E-71 |
| | | | | | PF12821 | ThrE_2 | Threonine/Serine exporter, ThrE | 1.40E-18 |
| WP_159508800.1 | 8562140 | 8561420 | - | 239 | PF09335 | SNARE_assoc | SNARE associated Golgi protein | 1.50E-24 |
| WP_159508802.1 | 8562397 | 8562763 | + | 121 | PF01906 | YbjQ_1 | Putative heavy-metal-binding | 1.20E-33 |
| | | | | | TIGR03884 | TIGR03884 | sel_bind_Methan: selenium-binding protein | 7.90E-04 |
| WP_159510477.1 | 8563627 | 8562868 | - | 252 | PF07739 | TipAS | TipAS antibiotic-recognition domain | 8.80E-37 |
| | | | | | PF13411 | MerR_1 | MerR HTH family regulatory protein | 5.50E-16 |
| | | | | | TIGR02043 | TIGR02043 | ZntR: Zn(II)-responsive transcriptional regulator | 2.50E-14 |
| | | | | | PF00376 | MerR | MerR family regulatory protein | 5.80E-14 |
| | | | | | TIGR02044 | TIGR02044 | CueR: Cu(I)-responsive transcriptional regulator | 2.30E-13 |
| WP_159508804.1 | 8563806 | 8564559 | + | 250 | PF01810 | LysE | LysE type translocator | 9.80E-43 |
| | | | | | TIGR00949 | TIGR00949 | 2A76: homoserine/Threonine efflux protein | 4.80E-42 |
| | | | | | TIGR00948 | TIGR00948 | 2a75: L-lysine exporter | 1.80E-14 |

| index | leader | core | start | end | dir | score |
|---|---|---|---|---|---|---|
| 1 | MSMLAEFANAELVDVKPGRIGSDATPTM | TTPLATVATPEATPVGFAATAATADALNMLTHNIAH | 8552362 | 8552167 | - | 31 |
| 2 | VGTAFTRPTTPDDAHDHARAAGLEC | EAHGPGRHSPQRKTVALFICCYVTP | 8554709 | 8554556 | - | 6 |
| 3 | MASFLVTASCLVPALLLHGVPVGFDSGDKTRE | HVWKTGEKTLAVGGSSKAGATASFLAADVPRRP | 8555166 | 8554968 | - | 7 |
| 4 | VFRLWRSGECEGPPAIPIRVFRTASERSCQPMRWALPSWRSAPGG | ATAALPGLPPGTCAK | 8559416 | 8559233 | - | 7 |

### Results for WP_165262038.1 [Streptomyces sp. HC44] back to top

previous - next

#### Architecture

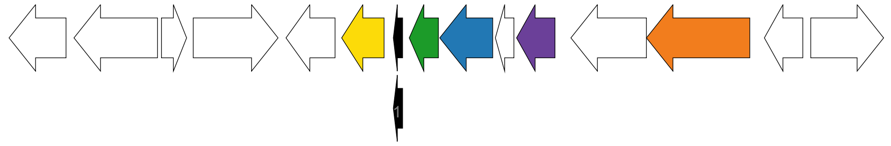

1000 nucleotides

Link to nucleotide sequence

| Accession | start | end | direction | length (aa) | Pfam/HMM | name | description | E-value |
|---|---|---|---|---|---|---|---|---|
| WP_165262026.1 | 30375 | 29247 | - | 375 | TIGR03883 | TIGR03883 | DUF2342_F420: uncharacterized protein, coenzyme F420 biosynthesis associated | 5.60E-145 |
| | | | | | TIGR03624 | TIGR03624 | TIGR03624: putative hydrolase | 1.80E-125 |
| | | | | | PF10103 | Zincin_2 | Zincin-like metallopeptidase | 2.00E-121 |
| WP_165262028.1 | 32155 | 30532 | - | 540 | PF02113 | Peptidase_S13 | D-Ala-D-Ala carboxypeptidase 3 (S13) family | 8.50E-74 |
| | | | | | TIGR00666 | TIGR00666 | PBP4: D-alanyl-D-alanine carboxypeptidase/D-alanyl-D-alanine-endopeptidase | 7.70E-55 |
| | | | | | PF00768 | Peptidase_S11 | D-alanyl-D-alanine carboxypeptidase | 3.10E-08 |
| WP_033321679.1 | 32234 | 32726 | + | 163 | PF00719 | Pyrophosphatase | Inorganic pyrophosphatase | 2.10E-53 |
| WP_165262030.1 | 32855 | 34502 | + | 548 | PF06738 | ThrE | Putative threonine/serine exporter | 4.20E-69 |
| | | | | | PF12821 | ThrE_2 | Threonine/Serine exporter, ThrE | 2.70E-14 |
| WP_165262032.1 | 35617 | 34663 | - | 317 | PF09335 | SNARE_assoc | SNARE associated Golgi protein | 3.00E-24 |
| WP_165262034.1 | 36577 | 35752 | - | 274 | LinG | LinG | LinG Homolog | 3.40E-108 |
| | | | | | LinH | LinH | LinH (Fusion of LinE/LinG) | 3.40E-78 |
| | | | | | TIGR04033 | TIGR04033 | export_SdpB: antimicrobial peptide system protein, SdpB family | 2.40E-04 |
| WP_165262036.1 | 36939 | 36750 | - | 62 | LinA | LinA | LinA (linaridin precursor) | 8.50E-13 |
| WP_165262058.1 | 37649 | 37067 | - | 193 | LinL | LinL | LinL | 3.90E-57 |
| WP_165262038.1 | 38686 | 37666 | - | 339 | LinE | LinE | LinE Homolog | 2.60E-99 |
| | | | | | LinH | LinH | LinH (Fusion of LinE/LinG) | 1.70E-51 |
| | | | | | PF12697 | Abhydrolase_6 | Alpha/beta hydrolase family | 1.90E-15 |
| | | | | | TIGR02427 | TIGR02427 | protocat_pcaD: 3-oxoadipate enol-lactonase | 2.70E-11 |
| | | | | | PF00561 | Abhydrolase_1 | alpha/beta hydrolase fold | 3.10E-08 |

| | | | | | | | | |
|---|---|---|---|---|---|---|---|---|
| WP_165262040.1 | 39107 | 38759 | - | 115 | PF13518 | HTH_28 | Helix-turn-helix domain | 4.90E-09 |
| | | | | | PF01527 | HTH_Tnp_1 | Transposase | 1.60E-08 |
| | | | | | PF13551 | HTH_29 | Winged helix-turn helix | 2.90E-06 |
| | | | | | PF13011 | LZ_Tnp_IS481 | leucine-zipper of insertion element IS481 | 4.60E-06 |
| | | | | | PF13384 | HTH_23 | Homeodomain-like domain | 3.00E-05 |
| WP_165262042.1 | 39920 | 39161 | - | 252 | PF13649 | Methyltransf_25 | Methyltransferase domain | 2.40E-15 |
| | | | | | PF08241 | Methyltransf_11 | Methyltransferase domain | 1.70E-11 |
| | | | | | PF08242 | Methyltransf_12 | Methyltransferase domain | 8.40E-09 |
| | | | | | PF13489 | Methyltransf_23 | Methyltransferase domain | 1.40E-08 |
| | | | | | TIGR01983 | TIGR01983 | UbiG: 3-demethylubiquinone-9 3-O-methyltransferase | 1.90E-06 |
| WP_165262044.1 | 41705 | 40220 | - | 494 | TIGR02734 | TIGR02734 | crtI_fam: phytoene desaturase | 1.10E-08 |
| | | | | | TIGR03467 | TIGR03467 | HpnE: squalene-associated FAD-dependent desaturase | 1.20E-08 |
| | | | | | TIGR00292 | TIGR00292 | TIGR00292: thiazole biosynthesis enzyme | 3.60E-08 |
| | | | | | PF13450 | NAD_binding_8 | NAD(P)-binding Rossmann-like domain | 3.50E-07 |
| | | | | | PF01946 | Thi4 | Thi4 family | 3.30E-06 |
| WP_165262046.1 | 43723 | 41701 | - | 673 | TIGR02204 | TIGR02204 | MsbA_rel: ABC transporter, permease/ATP-binding protein | 5.10E-98 |
| | | | | | TIGR02203 | TIGR02203 | MsbA_lipidA: lipid A export permease/ATP-binding protein MsbA | 1.60E-93 |
| | | | | | TIGR00958 | TIGR00958 | 3a01208: antigen peptide transporter 2 | 7.40E-86 |
| | | | | | TIGR03375 | TIGR03375 | type_I_sec_LssB: type I secretion system ATPase | 1.20E-79 |
| | | | | | TIGR03796 | TIGR03796 | NHLM_micro_ABC1: NHLM bacteriocin system ABC transporter, peptidase/ATP-binding protein | 1.10E-78 |
| WP_165262048.1 | 44764 | 44002 | - | 253 | PF07739 | TipAS | TipAS antibiotic-recognition domain | 1.00E-37 |
| | | | | | PF13411 | MerR_1 | MerR HTH family regulatory protein | 2.90E-16 |
| | | | | | TIGR02044 | TIGR02044 | CueR: Cu(I)-responsive transcriptional regulator | 2.90E-15 |
| | | | | | TIGR02043 | TIGR02043 | ZntR: Zn(II)-responsive transcriptional regulator | 1.60E-14 |
| | | | | | PF00376 | MerR | MerR family regulatory protein | 2.60E-13 |
| WP_165262050.1 | 44920 | 46327 | + | 468 | TIGR01788 | TIGR01788 | Glu-decarb-GAD: glutamate decarboxylase | 1.10E-186 |
| | | | | | PF00282 | Pyridoxal_deC | Pyridoxal-dependent decarboxylase conserved domain | 2.30E-70 |
| | | | | | TIGR03812 | TIGR03812 | tyr_de_CO2_Arch: tyrosine decarboxylase MnfA | 1.80E-40 |
| | | | | | TIGR03799 | TIGR03799 | NOD_PanD_pyr: putative pyridoxal-dependent aspartate 1-decarboxylase | 7.10E-07 |
| | | | | | TIGR03811 | TIGR03811 | tyr_de_CO2_Ent: tyrosine decarboxylase | 3.80E-04 |

| index | leader | core | start | end | dir | score |
|---|---|---|---|---|---|---|
| 1 | MSVVAEFANTELADVAPGRIGNEATPTM | LTPAATVGLAEGIAVTSATAYALNEVTNDLAAAG | 36939 | 36750 | - | 32 |

### Results for WP_152166802.1 [Streptomyces sp. GY16] back to top

previous - next

#### Architecture

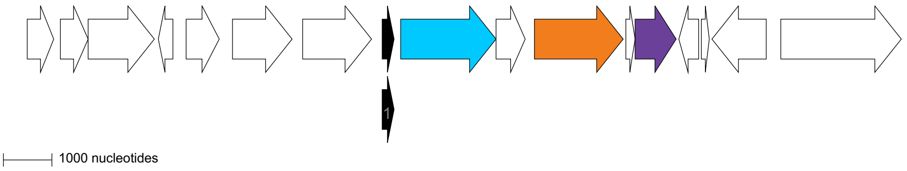

1000 nucleotides

Link to nucleotide sequence

| Accession | start | end | direction | length (aa) | Pfam/HMM | name | description | E-value |
|---|---|---|---|---|---|---|---|---|
| WP_006374913.1 | 476270 | 476825 | + | 185 | INFERRED GENE | - | - | - |
| WP_152166797.1 | 476969 | 477530 | + | 186 | TIGR02983 | TIGR02983 | SigE-fam_strep: RNA polymerase sigma-70 factor, sigma-E family | 8.60E-55 |
| | | | | | TIGR02937 | TIGR02937 | sigma70-ECF: RNA polymerase sigma factor, sigma-70 family | 2.50E-19 |
| | | | | | PF08281 | Sigma70_r4_2 | Sigma-70, region 4 | 2.10E-16 |
| | | | | | TIGR02954 | TIGR02954 | Sig70_famx3: RNA polymerase sigma-70 factor, TIGR02954 family | 3.50E-15 |
| | | | | | TIGR02960 | TIGR02960 | SigX5: RNA polymerase sigma-70 factor, TIGR02960 family | 9.10E-13 |
| WP_152166798.1 | 477526 | 478888 | + | 453 | PF13360 | PQQ_2 | PQQ-like domain | 1.40E-18 |
| | | | | | TIGR03300 | TIGR03300 | assembly_YfgL: outer membrane assembly lipoprotein YfgL | 9.30E-12 |
| | | | | | PF01011 | PQQ | PQQ enzyme repeat | 6.00E-07 |
| | | | | | TIGR03075 | TIGR03075 | PQQ_enz_alc_DH: PQQ-dependent dehydrogenase, methanol/ethanol family | 1.20E-06 |
| WP_152166799.1 | 479285 | 478985 | - | 99 | PF00196 | GerE | Bacterial regulatory proteins, luxR family | 2.00E-15 |
| | | | | | TIGR03541 | TIGR03541 | reg_near_HchA: LuxR family transcriptional regulatory, chaperone HchA-associated | 9.40E-12 |
| | | | | | TIGR03020 | TIGR03020 | EpsA: transcriptional regulator EpsA | 3.20E-08 |
| | | | | | PF08281 | Sigma70_r4_2 | Sigma-70, region 4 | 4.50E-06 |
| | | | | | PF13412 | HTH_24 | Winged helix-turn-helix DNA-binding | 7.40E-04 |
| WP_152166800.1 | 479537 | 480227 | + | 229 | PF00072 | Response_reg | Response regulator receiver domain | 3.50E-25 |
| | | | | | PF00196 | GerE | Bacterial regulatory proteins, luxR family | 4.30E-18 |
| | | | | | TIGR02875 | TIGR02875 | spore_0_A: sporulation transcription factor Spo0A | 5.90E-16 |
| | | | | | TIGR03020 | TIGR03020 | EpsA: transcriptional regulator EpsA | 1.80E-12 |
| | | | | | TIGR03541 | TIGR03541 | reg_near_HchA: LuxR family transcriptional regulatory, chaperone HchA-associated | 3.30E-08 |
| WP_152173649.1 | 480495 | 481725 | + | 409 | PF07730 | HisKA_3 | Histidine kinase | 7.20E-14 |
| WP_152166801.1 | 481945 | 483361 | + | 471 | TIGR02734 | TIGR02734 | crtI_fam: phytoene desaturase | 4.70E-15 |
| | | | | | PF13450 | NAD_binding_8 | NAD(P)-binding Rossmann-like domain | 3.00E-11 |
| | | | | | TIGR03467 | TIGR03467 | HpnE: squalene-associated FAD-dependent desaturase | 2.90E-09 |
| | | | | | TIGR03315 | TIGR03315 | Se_ygfK: putative selenate reductase, YgfK subunit | 3.60E-08 |
| | | | | | TIGR00292 | TIGR00292 | TIGR00292: thiazole biosynthesis enzyme | 6.80E-08 |
| WP_152173650.1 | 483578 | 483836 | + | 85 | LinA | LinA | LinA (linaridin precursor) | 2.00E-04 |
| WP_152166802.1 | 483983 | 485948 | + | 654 | LinH | LinH | LinH (Fusion of LinE/LinG) | 4.90E-230 |
| | | | | | LinG | LinG | LinG Homolog | 9.30E-124 |
| | | | | | LinE | LinE | LinE Homolog | 1.00E-86 |
| | | | | | PF12697 | Abhydrolase_6 | Alpha/beta hydrolase family | 6.90E-16 |
| | | | | | PF12146 | Hydrolase_4 | Serine aminopeptidase, S33 | 2.00E-09 |
| WP_005474565.1 | 485944 | 486535 | + | 197 | INFERRED GENE | - | - | - |

| | | | | | | | | |
|---|---|---|---|---|---|---|---|---|
| WP_152166803.1 | 486734 | 488570 | + | 611 | TIGR02204<br>TIGR02203<br>TIGR00958<br>TIGR03796<br>TIGR01846 | TIGR02204<br>TIGR02203<br>TIGR00958<br>TIGR03796<br>TIGR01846 | MsbA_rel: ABC transporter, permease/ATP-binding protein<br>MsbA_lipidA: lipid A export permease/ATP-binding protein MsbA<br>3a01208: antigen peptide transporter 2<br>NHLM_micro_ABC1: NHLM bacteriocin system ABC transporter, peptidase/ATP-binding protein<br>type_I_sec_HlyB: type I secretion system ATPase | 8.80E-107<br>1.40E-103<br>3.80E-92<br>6.40E-88<br>1.40E-82 |
| WP_152166804.1 | 488603 | 488795 | + | 63 | NO PFAM MATCH | - | - | - |
| WP_152166805.1 | 488791 | 489637 | + | 281 | PF13649<br>PF08241<br>PF13489<br>PF13847<br>PF08242 | Methyltransf_25<br>Methyltransf_11<br>Methyltransf_23<br>Methyltransf_31<br>Methyltransf_12 | Methyltransferase domain<br>Methyltransferase domain<br>Methyltransferase domain<br>Methyltransferase domain<br>Methyltransferase domain | 7.30E-13<br>3.70E-10<br>5.20E-08<br>2.50E-04<br>5.30E-04 |
| WP_152166806.1 | 490108 | 489712 | - | 131 | PF00583<br>PF13673<br>TIGR01575<br>TIGR03448<br>PF13508 | Acetyltransf_1<br>Acetyltransf_10<br>TIGR01575<br>TIGR03448<br>Acetyltransf_7 | Acetyltransferase (GNAT) family<br>Acetyltransferase (GNAT) domain<br>rimI: ribosomal-protein-alanine acetyltransferase<br>mycothiol_MshD: mycothiol synthase<br>Acetyltransferase (GNAT) domain | 1.50E-12<br>9.20E-06<br>1.50E-05<br>4.80E-05<br>3.10E-04 |
| WP_006607098.1 | 490167 | 490332 | + | 55 | INFERRED GENE | - | - | - |
| WP_152166807.1 | 491519 | 490376 | - | 380 | PF13407 | Peripla_BP_4 | Periplasmic binding protein domain | 8.50E-20 |
| WP_152166808.1 | 491795 | 494285 | + | 829 | PF02518<br>PF00672<br>TIGR02966 | HATPase_c<br>HAMP<br>TIGR02966 | Histidine kinase-, DNA gyrase B-, and HSP90-like ATPase<br>HAMP domain<br>phoR_proteo: phosphate regulon sensor kinase PhoR | 1.60E-07<br>2.80E-06<br>6.20E-04 |

| index | leader | core | start | end | dir | score |
|---|---|---|---|---|---|---|
| 1 | MDLPVEGVILASARPTL | GTPRIGRVNGANALNALAALGDLAQ...NAAVDANNAVANAANAQAAAVGGTV | 483578 | 483836 | + | 29 |

### Results for WP_165254437.1 [Streptomyces sp. HC44] back to top

previous - next

#### Architecture

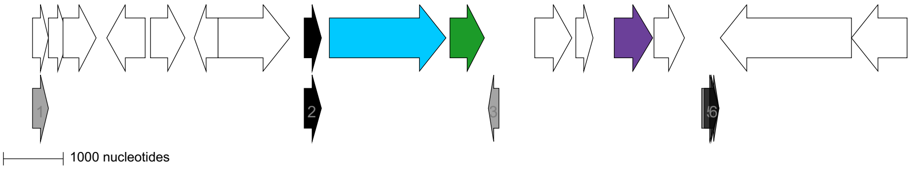

Link to nucleotide sequence

| Accession | start | end | direction | length (aa) | Pfam/HMM | name | description | E-value |
|---|---|---|---|---|---|---|---|---|
| WP_165254399.1 | 43808 | 44075 | + | 88 | NO PFAM MATCH | - | - | - |
| WP_165254432.1 | 44071 | 44413 | + | 113 | NO PFAM MATCH | - | - | - |
| WP_165254433.1 | 44312 | 44870 | + | 185 | TIGR03356<br>PF00232<br>TIGR01233 | TIGR03356<br>Glyco_hydro_1<br>TIGR01233 | BGL: beta-galactosidase<br>Glycosyl hydrolase family 1<br>lacG: 6-phospho-beta-galactosidase | 3.20E-24<br>1.70E-19<br>1.80E-12 |
| WP_165254434.1 | 45678 | 45054 | - | 207 | PF00400<br>PF16529<br>PF12894<br>TIGR02800<br>PF07676 | WD40<br>Ge1_WD40<br>ANAPC4_WD40<br>TIGR02800<br>PD40 | WD domain, G-beta repeat<br>WD40 region of Ge1, enhancer of mRNA-decapping protein<br>Anaphase-promoting complex subunit 4 WD40 domain<br>propeller_TolB: Tol-Pal system beta propeller repeat protein TolB<br>WD40-like Beta Propeller Repeat | 1.40E-36<br>6.70E-12<br>1.10E-11<br>2.50E-06<br>6.90E-05 |
| WP_017566699.1 | 45766 | 46362 | + | 198 | INFERRED GENE | - | - | - |
| WP_165254435.1 | 46900 | 46498 | - | 133 | PF04134 | DUF393 | Protein of unknown function, DUF393 | 2.70E-15 |
| WP_018835199.1 | 46896 | 48094 | + | 399 | INFERRED GENE | - | - | - |
| WP_165254436.1 | 48346 | 48637 | + | 96 | LinA | LinA | LinA (linaridin precursor) | 1.80E-05 |
| WP_165254437.1 | 48776 | 50720 | + | 647 | LinH<br>LinE<br>LinG<br>PF12697<br>PF00561 | LinH<br>LinE<br>LinG<br>Abhydrolase_6<br>Abhydrolase_1 | LinH (Fusion of LinE/LinG)<br>LinE Homolog<br>LinG Homolog<br>Alpha/beta hydrolase family<br>alpha/beta hydrolase fold | 3.30E-183<br>2.40E-89<br>1.60E-85<br>2.00E-15<br>1.40E-11 |
| WP_165254438.1 | 50786 | 51353 | + | 188 | LinL | LinL | LinL | 2.70E-54 |
| WP_165254501.1 | 52189 | 52822 | + | 210 | PF00196<br>TIGR03020<br>TIGR03541<br>PF13936<br>PF08281 | GerE<br>TIGR03020<br>TIGR03541<br>HTH_38<br>Sigma70_r4_2 | Bacterial regulatory proteins, luxR family<br>EpsA: transcriptional regulator EpsA<br>reg_near_HchA: LuxR family transcriptional regulatory, chaperone HchA-associated<br>Helix-turn-helix domain<br>Sigma-70, region 4 | 3.50E-17<br>1.00E-13<br>6.20E-08<br>9.70E-05<br>5.80E-04 |
| WP_165254439.1 | 52873 | 53179 | + | 101 | PF04472 | SepF | Cell division protein SepF | 1.80E-07 |
| WP_165254440.1 | 53534 | 54161 | + | 208 | PF13649<br>PF13489<br>PF08241<br>PF13847<br>TIGR00477 | Methyltransf_25<br>Methyltransf_23<br>Methyltransf_11<br>Methyltransf_31<br>TIGR00477 | Methyltransferase domain<br>Methyltransferase domain<br>Methyltransferase domain<br>Methyltransferase domain<br>tehB: tellurite resistance protein TehB | 5.20E-16<br>3.70E-15<br>1.00E-13<br>1.80E-13<br>6.70E-13 |
| WP_165254441.1 | 54189 | 54690 | + | 166 | PF04978<br>PF12867 | DUF664<br>DinB_2 | Protein of unknown function (DUF664)<br>DinB superfamily | 8.30E-38<br>3.30E-08 |
| WP_165254442.1 | 57489 | 55290 | - | 732 | NO PFAM MATCH | - | - | - |

| | | | | | | | | |
|---|---|---|---|---|---|---|---|---|
| WP_165254443.1 | 58421 | 57500 | - | 306 | PF00528<br>TIGR01581<br>TIGR03262<br>TIGR02141<br>TIGR02139 | BPD_transp_1<br>TIGR01581<br>TIGR03262<br>TIGR02141<br>TIGR02139 | Binding-protein-dependent transport system inner membrane component<br>Mo_ABC_porter: NifC-like ABC-type porter<br>PhnU2: putative 2-aminoethylphosphonate ABC transporter, permease protein<br>modB_ABC: molybdate ABC transporter, permease protein<br>permease_CysT: sulfate ABC transporter, permease protein CysT | 1.20E-20<br>1.10E-14<br>1.50E-13<br>2.20E-10<br>8.20E-10 |

| index | leader | core | start | end | dir | score |
|---|---|---|---|---|---|---|
| 1 | MRCLPSDPTTLATVHLREDRSAAYS...PDGGALPAGTALQLGSLGLLLEPLA | STLDNRSPLSISLERKSVT | 43808 | 44075 | + | 6 |
| 2 | MTLIQSVQESPIAQWAHENATAHDTHYIAPQDKTIPVL | FTPLVALTVAGAAGCAIGAGTGFAQ...ASMPTSGTAAGLSVDALTSRHPGMA | 48346 | 48637 | + | 27 |
| 3 | LLRGRPERARRPFLNDPRALSNGDGCDG | RSIPFTWHHRATRNPASVAAMTHAISEP | 51594 | 51423 | - | 6 |
| 4 | LLRQLGRRLGRRFVVGFVADGRENV...SSGGNGAWRQVTPPVAPPPRATPLT | AVPSEGAAVSRVWHAAWSGRTL | 54996 | 55275 | + | 7 |
| 5 | VVGFVADGRENVDRGEQALPFPTGS...SSGGNGAWRQVTPPVAPPPRATPLT | AVPSEGAAVSRVWHAAWSGRTL | 55035 | 55275 | + | 8 |
| 6 | VWTNSSGGNGAWRQVTPPVAPPPRATPLT | AVPSEGAAVSRVWHAAWSGRTL | 55119 | 55275 | + | 8 |

### Results for WP_147874565.1 [Streptomyces sp. IB2014 016-6] back to top

previous - next

#### Architecture

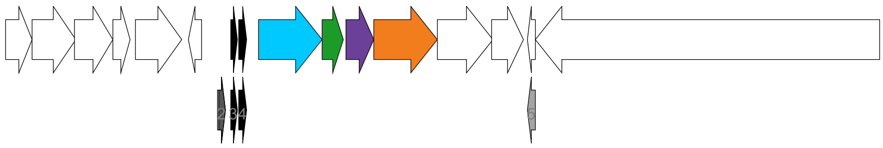

1000 nucleotides

Link to nucleotide sequence

| Accession | start | end | direction | length (aa) | Pfam/HMM | name | description | E-value |
|---|---|---|---|---|---|---|---|---|
| WP_147874559.1 | 30077 | 30797 | + | 239 | PF00441 | Acyl-CoA_dh_1 | Acyl-CoA dehydrogenase, C-terminal domain | 6.30E-04 |
| WP_147874560.1 | 30793 | 31966 | + | 390 | PF00441<br>PF02770<br>TIGR03207<br>TIGR03204<br>PF02771 | Acyl-CoA_dh_1<br>Acyl-CoA_dh_M<br>TIGR03207<br>TIGR03204<br>Acyl-CoA_dh_N | Acyl-CoA dehydrogenase, C-terminal domain<br>Acyl-CoA dehydrogenase, middle domain<br>cyc_hxne_CoA_dh: cyclohexanecarboxyl-CoA dehydrogenase<br>pimC_large: pimeloyl-CoA dehydrogenase, large subunit<br>Acyl-CoA dehydrogenase, N-terminal domain | 2.20E-20<br>9.80E-18<br>1.30E-14<br>1.20E-11<br>1.40E-06 |
| WP_147874561.1 | 31946 | 32987 | + | 346 | PF01648<br>TIGR00556 | ACPS<br>TIGR00556 | 4'-phosphopantetheinyl transferase superfamily<br>pantethn_trn: phosphopantetheine--protein transferase domain | 5.00E-06<br>6.00E-04 |
| WP_147874562.1 | 32983 | 33448 | + | 154 | NO PFAM MATCH | - | - | - |
| WP_147874851.1 | 33569 | 34838 | + | 422 | TIGR00900<br>PF07690<br>TIGR00881<br>PF05977<br>TIGR00880 | TIGR00900<br>MFS_1<br>TIGR00881<br>MFS_3<br>TIGR00880 | 2A0121: H+ Antiporter protein<br>Major Facilitator Superfamily<br>2A0104: phosphoglycerate transporter family protein<br>Transmembrane secretion effector<br>2_A_01_02: multidrug resistance protein | 2.20E-37<br>1.80E-30<br>2.50E-13<br>1.10E-12<br>3.60E-12 |
| WP_147874852.1 | 35368 | 35020 | - | 115 | PF12840<br>PF01022<br>PF13518<br>PF13338 | HTH_20<br>HTH_5<br>HTH_28<br>AbiEi_4 | Helix-turn-helix domain<br>Bacterial regulatory protein, arsR family<br>Helix-turn-helix domain<br>Transcriptional regulator, AbiEi antitoxin | 1.90E-11<br>1.00E-06<br>1.10E-04<br>5.10E-04 |
| WP_147874563.1 | 36156 | 36363 | + | 68 | LinA | LinA | LinA (linaridin precursor) | 1.00E-14 |
| WP_147874564.1 | 36392 | 36584 | + | 63 | LinA | LinA | LinA (linaridin precursor) | 1.60E-09 |
| WP_147874565.1 | 36928 | 38662 | + | 577 | LinH<br>LinE<br>LinG<br>PF12697<br>PF00561 | LinH<br>LinE<br>LinG<br>Abhydrolase_6<br>Abhydrolase_1 | LinH (Fusion of LinE/LinG)<br>LinE Homolog<br>LinG Homolog<br>Alpha/beta hydrolase family<br>alpha/beta hydrolase fold | 1.30E-216<br>1.10E-97<br>4.30E-92<br>1.70E-15<br>2.90E-13 |
| WP_147874566.1 | 38658 | 39237 | + | 192 | LinL | LinL | LinL | 3.70E-64 |
| WP_147874567.1 | 39286 | 40045 | + | 252 | PF13649<br>PF08241<br>PF13489<br>PF08242<br>PF13847 | Methyltransf_25<br>Methyltransf_11<br>Methyltransf_23<br>Methyltransf_12<br>Methyltransf_31 | Methyltransferase domain<br>Methyltransferase domain<br>Methyltransferase domain<br>Methyltransferase domain<br>Methyltransferase domain | 4.80E-14<br>3.70E-09<br>7.40E-08<br>8.00E-08<br>2.50E-07 |
| WP_147874568.1 | 40041 | 41784 | + | 580 | TIGR02204<br>TIGR02203<br>TIGR00958<br>TIGR03796<br>TIGR01846 | TIGR02204<br>TIGR02203<br>TIGR00958<br>TIGR03796<br>TIGR01846 | MsbA_rel: ABC transporter, permease/ATP-binding protein<br>MsbA_lipidA: lipid A export permease/ATP-binding protein MsbA<br>3a01208: antigen peptide transporter 2<br>NHLM_micro_ABC1: NHLM bacteriocin system ABC transporter, peptidase/ATP-binding protein<br>type_I_sec_HlyB: type I secretion system ATPase | 3.60E-103<br>1.00E-95<br>2.20E-89<br>1.30E-82<br>5.80E-79 |
| WP_147874569.1 | 41780 | 43265 | + | 494 | TIGR02734<br>TIGR01317<br>TIGR02733<br>TIGR03467<br>TIGR00292 | TIGR02734<br>TIGR01317<br>TIGR02733<br>TIGR03467<br>TIGR00292 | crtI_fam: phytoene desaturase<br>GOGAT_sm_gam: glutamate synthase, NADH/NADPH, small subunit<br>desat_CrtD: C-3',4' desaturase CrtD<br>HpnE: squalene-associated FAD-dependent desaturase<br>TIGR00292: thiazole biosynthesis enzyme | 2.50E-13<br>5.20E-09<br>6.70E-09<br>8.90E-09<br>2.00E-08 |
| WP_147874570.1 | 43261 | 44113 | + | 283 | PF12680 | SnoaL_2 | SnoaL-like domain | 1.10E-06 |
| WP_147874571.1 | 44425 | 44203 | - | 73 | PF03621 | MbtH | MbtH-like protein | 1.40E-28 |
| WP_147874572.1 | 53787 | 44421 | - | 3121 | TIGR01733<br>PF00668<br>PF00501<br>TIGR01734<br>TIGR03443 | TIGR01733<br>Condensation<br>AMP-binding<br>TIGR01734<br>TIGR03443 | AA-adenyl-dom: amino acid adenylation domain<br>Condensation domain<br>AMP-binding enzyme<br>D-ala-DACP-lig: D-alanine--poly(phosphoribitol) ligase, subunit 1<br>alpha_am_amid: L-aminoadipate-semialdehyde dehydrogenase | 2.20E-246<br>2.10E-212<br>3.70E-178<br>7.20E-127<br>5.00E-118 |

| index | leader | core | start | end | dir | score |
|---|---|---|---|---|---|---|
| 1 | LFQIPHHHRATSGFSLLGGLPIDPFFAPRGNDGQRKNAAC | VTPRLQCDVHHIDKGLIRQGPPVLLWLPQSQAFNQC | 35793 | 36024 | + | 6 |
| 2 | LGGLPIDPFFAPRGNDGQRKNAAC | VTPRLQCDVHHIDKGLIRQGPPVLLWLPQSQAFNQC | 35841 | 36024 | + | 7 |
| 3 | MKEQSVATIAEFMNDQKLGVLEQGYKEETQEPVL | ATPAAAVATCWYVGGAAAGVTVVAGAYVAGRVFG | 36156 | 36363 | + | 30 |
| 4 | MSRIIESVAAITPEHLTDGSHGEVTAMAPVQ | ATPGLVAFGIGFAGGAGVSWVGVQAYEAGAND | 36392 | 36584 | + | 31 |
| 5 | MTNPFDDESGIFLVLVNDEGQHSLWPRFADVPAG | WTPTFGPAGRAACLTHIESAWTDLRPKSLIAATDGSTTH | 44425 | 44203 | - | 6 |

### Results for RNL73410.1 [Streptomyces sp. I6] back to top

previous - next

#### Architecture

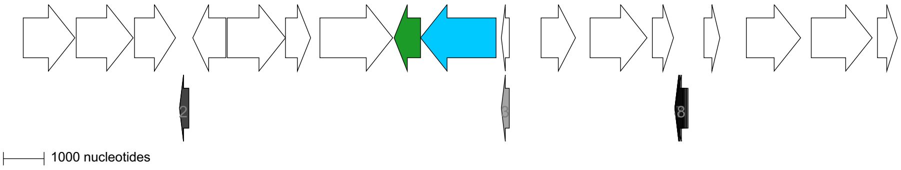

1000 nucleotides

Link to nucleotide sequence

| Accession | start | end | direction | length (aa) | Pfam/HMM | name | description | E-value |
|---|---|---|---|---|---|---|---|---|
| RNL73405.1 | 5892031 | 5893312 | + | 426 | PF00202<br>TIGR00707<br>TIGR00700<br>TIGR03246<br>TIGR01885 | Aminotran_3<br>TIGR00707<br>TIGR00700<br>TIGR03246<br>TIGR01885 | Aminotransferase class-III<br>argD: transaminase, acetylornithine/succinylornithine family<br>GABAtrnsam: 4-aminobutyrate transaminase<br>arg_catab_astC: succinylornithine transaminase family<br>Orn_aminotrans: ornithine--oxo-acid transaminase | 2.10E-85<br>1.40E-81<br>3.70E-69<br>1.90E-61<br>7.00E-60 |
| RNL73406.1 | 5893331 | 5894735 | + | 467 | TIGR02033<br>TIGR03178<br>TIGR00857<br>PF01979<br>PF07969 | TIGR02033<br>TIGR03178<br>TIGR00857<br>Amidohydro_1<br>Amidohydro_3 | D-hydantoinase: dihydropyrimidinase<br>allantoinase: allantoinase<br>pyrC_multi: dihydroorotase, multifunctional complex type<br>Amidohydrolase family<br>Amidohydrolase family | 1.60E-167<br>2.30E-81<br>1.20E-53<br>1.60E-24<br>1.30E-17 |
| WP_005309617.1 | 5894765 | 5895766 | + | 333 | INFERRED GENE | - | - | - |
| RNL73407.1 | 5897004 | 5896209 | - | 264 | PF00106<br>PF13561<br>TIGR01963<br>TIGR01830<br>TIGR03971 | adh_short<br>adh_short_C2<br>TIGR01963<br>TIGR01830<br>TIGR03971 | short chain dehydrogenase<br>Enoyl-(Acyl carrier protein) reductase<br>PHB_DH: 3-hydroxybutyrate dehydrogenase<br>3oxo_ACP_reduc: 3-oxoacyl-[acyl-carrier-protein] reductase<br>SDR_subfam_1: SDR family mycofactocin-dependent oxidoreductase | 1.90E-38<br>8.30E-30<br>7.30E-29<br>2.30E-24<br>1.70E-20 |
| RNL73408.1 | 5897060 | 5898488 | + | 475 | PF01425<br>TIGR02715<br>TIGR00132<br>TIGR02713 | Amidase<br>TIGR02715<br>TIGR00132<br>TIGR02713 | Amidase<br>amido_AtzE: amidohydrolase, AtzE family<br>gatA: aspartyl/glutamyl-tRNA(Asn/Gln) amidotransferase, A subunit<br>allophanate_hyd: allophanate hydrolase | 1.60E-108<br>8.20E-74<br>8.30E-61<br>9.00E-57 |
| RNL74348.1 | 5898493 | 5899108 | + | 204 | PF17940<br>TIGR03384<br>PF00440 | TetR_C_31<br>TIGR03384<br>TetR_N | Tetracyclin repressor-like, C-terminal domain<br>betaine_BetI: transcriptional repressor BetI<br>Bacterial regulatory proteins, tetR family | 2.80E-18<br>8.50E-13<br>2.20E-10 |
| WP_007452028.1 | 5899340 | 5901127 | + | 595 | INFERRED GENE | - | - | - |
| RNL73409.1 | 5901786 | 5901141 | - | 214 | LinL | LinL | LinL | 1.50E-65 |
| RNL73410.1 | 5903642 | 5901797 | - | 614 | LinH<br>LinE<br>LinG<br>PF12697<br>TIGR03695 | LinH<br>LinE<br>LinG<br>Abhydrolase_6<br>TIGR03695 | LinH (Fusion of LinE/LinG)<br>LinE Homolog<br>LinG Homolog<br>Alpha/beta hydrolase family<br>menH_SHCHC: 2-succinyl-6-hydroxy-2,4-cyclohexadiene-1-carboxylate synthase | 1.20E-227<br>2.10E-107<br>3.60E-107<br>1.90E-19<br>6.10E-14 |
| RNL73411.1 | 5903988 | 5903787 | - | 66 | NO PFAM MATCH | - | - | - |
| RNL73412.1 | 5904759 | 5905617 | + | 285 | PF00459<br>TIGR02067<br>TIGR01331<br>TIGR01330<br>PF03320 | Inositol_P<br>TIGR02067<br>TIGR01331<br>TIGR01330<br>FBPase_glpX | Inositol monophosphatase family<br>his_9_HisN: histidinol-phosphatase<br>bisphos_cysQ: 3'(2'),5'-bisphosphate nucleotidase<br>bisphos_HAL2: 3'(2'),5'-bisphosphate nucleotidase<br>Bacterial fructose-1,6-bisphosphatase, glpX-encoded | 2.70E-56<br>1.00E-40<br>8.20E-28<br>6.30E-11<br>8.30E-04 |
| RNL73413.1 | 5905962 | 5907381 | + | 472 | TIGR02734<br>PF13450<br>TIGR02733<br>TIGR03467<br>TIGR01316 | TIGR02734<br>NAD_binding_8<br>TIGR02733<br>TIGR03467<br>TIGR01316 | crtI_fam: phytoene desaturase<br>NAD(P)-binding Rossmann-like domain<br>desat_CrtD: C-3',4' desaturase CrtD<br>HpnE: squalene-associated FAD-dependent desaturase<br>gltA: glutamate synthase (NADPH), homotetrameric | 8.10E-16<br>5.80E-12<br>1.90E-09<br>6.50E-09<br>8.80E-09 |
| RNL74349.1 | 5907487 | 5908006 | + | 172 | PF01661 | Macro | Macro domain | 1.30E-39 |
| RNL73414.1 | 5908757 | 5909141 | + | 127 | NO PFAM MATCH | - | - | - |
| WP_018888836.1 | 5909812 | 5911143 | + | 443 | INFERRED GENE | - | - | - |
| RNL74350.1 | 5911401 | 5912883 | + | 493 | PF06029<br>PF02805<br>PF12833<br>PF00730<br>PF00165 | AlkA_N<br>Ada_Zn_binding<br>HTH_18<br>HhH-GPD<br>HTH_AraC | AlkA N-terminal domain<br>Metal binding domain of Ada<br>Helix-turn-helix domain<br>HhH-GPD superfamily base excision DNA repair protein<br>Bacterial regulatory helix-turn-helix proteins, AraC family | 3.30E-30<br>4.10E-26<br>8.30E-17<br>4.30E-05<br>5.80E-04 |
| RNL73415.1 | 5913017 | 5913512 | + | 164 | TIGR00589<br>PF01035<br>PF02870 | TIGR00589<br>DNA_binding_1<br>Methyltransf_1N | ogt: methylated-DNA--[protein]-cysteine S-methyltransferase<br>6-O-methylguanine DNA methyltransferase, DNA binding domain<br>6-O-methylguanine DNA methyltransferase, ribonuclease-like domain | 7.40E-34<br>1.90E-31<br>2.70E-05 |

| index | leader | core | start | end | dir | score |
|---|---|---|---|---|---|---|

| | | | | | | |
|---|---|---|---|---|---|---|
| 1 | VAAAPLRWGRPAGGAGRAPAASAPC...RAVPAGDRTGAGEGRPGAEGEPPTA | GTPPAVRAGA | 5896089 | 5895861 | - | 7 |
| 2 | VFHVAAAPLRWGRPAGGAGRAPAAS...RAVPAGDRTGAGEGRPGAEGEPPTA | GTPPAVRAGA | 5896098 | 5895861 | - | 7 |
| 3 | MELLHRVNSDSALA | AVPILTEEAVQVKGAEPILNIFNDM...FVNAATQPAGEVAADTMEVTMATVG | 5903988 | 5903787 | - | 6 |
| 4 | VDQAHHRLAPPVRHSPPARKPWV | TTPRSTAPPRASRTGPAASGQECARVPPAVLLAPEAL | 5908230 | 5908047 | - | 6 |
| 5 | VSVDQAHHRLAPPVRHSPPARKPWV | TTPRSTAPPRASRTGPAASGQECARVPPAVLLAPEAL | 5908236 | 5908047 | - | 6 |
| 6 | MTLGDPRPCGVSVDQAHHRLAPPVRHSPPARKPWV | TTPRSTAPPRASRTGPAASGQECARVPPAVLLAPEAL | 5908266 | 5908047 | - | 7 |
| 7 | LNSTPPRNRRRNTRRKPGGGPRVQR...VSVDQAHHRLAPPVRHSPPARKPWV | TTPRSTAPPRASRTGPAASGQECARVPPAVLLAPEAL | 5908347 | 5908047 | - | 8 |
| 8 | VGGASGTPSLNSTPPRNRRRNTRRK...VSVDQAHHRLAPPVRHSPPARKPWV | TTPRSTAPPRASRTGPAASGQECARVPPAVLLAPEAL | 5908374 | 5908047 | - | 7 |

### Results for WP_129773542.1 [Streptomyces sp. L-9-10] back to top

previous - next

#### Architecture

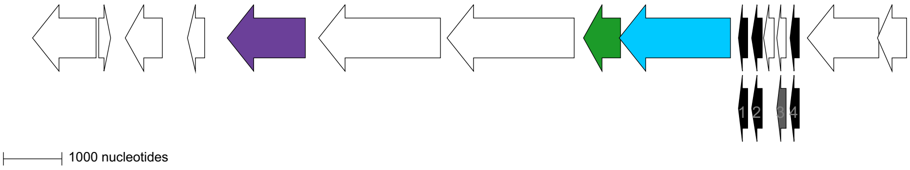

Link to nucleotide sequence

| Accession | start | end | direction | length (aa) | Pfam/HMM | name | description | E-value |
|---|---|---|---|---|---|---|---|---|
| WP_129773527.1 | 103641 | 102552 | - | 362 | PF00589<br>TIGR02224<br>TIGR02225<br>TIGR02249<br>PF14659 | Phage_integrase<br>TIGR02224<br>TIGR02225<br>TIGR02249<br>Phage_int_SAM_3 | Phage integrase family<br>recomb_XerC: tyrosine recombinase XerC<br>recomb_XerD: tyrosine recombinase XerD<br>integrase_gron: integron integrase<br>Phage integrase, N-terminal SAM-like domain | 3.80E-25<br>1.70E-23<br>7.10E-20<br>9.30E-13<br>1.60E-04 |
| WP_009324638.1 | 103700 | 103906 | + | 68 | INFERRED GENE | - | - | - |
| WP_129773529.1 | 104779 | 104149 | - | 209 | PF00589<br>TIGR02224 | Phage_integrase<br>TIGR02224 | Phage integrase family<br>recomb_XerC: tyrosine recombinase XerC | 9.20E-09<br>4.10E-05 |
| WP_129773531.1 | 105512 | 105212 | - | 99 | NO PFAM MATCH | - | - | - |
| WP_129773533.1 | 107221 | 105892 | - | 442 | TIGR00479<br>TIGR02085<br>TIGR02143<br>PF05958<br>PF01938 | TIGR00479<br>TIGR02085<br>TIGR02143<br>tRNA_U5-meth_tr<br>TRAM | rumA: 23S rRNA (uracil-5-)-methyltransferase RumA<br>meth_trns_rumB: 23S rRNA (uracil-5-)-methyltransferase RumB<br>trmA_only: tRNA (uracil(54)-C(5))-methyltransferase<br>tRNA (Uracil-5-)-methyltransferase<br>TRAM domain | 8.60E-62<br>4.00E-19<br>9.60E-16<br>2.10E-14<br>7.40E-13 |
| WP_129773535.1 | 109546 | 107464 | - | 693 | PF13520<br>PF02705<br>TIGR00908 | AA_permease_2<br>K_trans<br>TIGR00908 | Amino acid permease<br>K+ potassium transporter<br>2A0305: ethanolamine permease | 2.50E-32<br>3.20E-07<br>1.40E-04 |
| WP_129773538.1 | 111821 | 109643 | - | 725 | PF00933<br>PF01915<br>PF14310 | Glyco_hydro_3<br>Glyco_hydro_3_C<br>Fn3-like | Glycosyl hydrolase family 3 N terminal domain<br>Glycosyl hydrolase family 3 C-terminal domain<br>Fibronectin type III-like domain | 4.70E-77<br>4.20E-59<br>3.70E-20 |
| WP_129773540.1 | 112618 | 111982 | - | 211 | LinL | LinL | LinL | 1.00E-70 |
| WP_129773542.1 | 114494 | 112598 | - | 631 | LinH<br>LinG<br>LinE<br>PF12697<br>PF00975 | LinH<br>LinG<br>LinE<br>Abhydrolase_6<br>Thioesterase | LinH (Fusion of LinE/LinG)<br>LinG Homolog<br>LinE Homolog<br>Alpha/beta hydrolase family<br>Thioesterase domain | 2.20E-218<br>5.60E-102<br>4.00E-88<br>4.30E-14<br>7.30E-10 |
| WP_165372062.1 | 114796 | 114625 | - | 56 | LinA | LinA | LinA (linaridin precursor) | 2.10E-11 |
| WP_129773544.1 | 115034 | 114848 | - | 61 | LinA | LinA | LinA (linaridin precursor) | 4.90E-08 |
| WP_165372063.1 | 115234 | 115072 | - | 53 | NO PFAM MATCH | - | - | - |
| WP_165372064.1 | 115454 | 115286 | - | 55 | NO PFAM MATCH | - | - | - |
| WP_165372065.1 | 115680 | 115506 | - | 57 | LinA | LinA | LinA (linaridin precursor) | 1.70E-09 |
| WP_129773546.1 | 117026 | 115808 | - | 405 | NO PFAM MATCH | - | - | - |
| WP_129773673.1 | 117498 | 117018 | - | 159 | PF06114 | Peptidase_M78 | IrrE N-terminal-like domain | 5.60E-04 |

| index | leader | core | start | end | dir | score |
|---|---|---|---|---|---|---|
| 1 | MSLLANHVNESSALATVPL | HTTEVQRVDVPDSVICCPAAVQVGAALVAAFVAGYNL | 114796 | 114625 | - | 17 |
| 2 | MSSLSHQVNTSGALATVPLHTTQVHQMEPSDSAVCCPAVL | VTFAAGIAFGYALGGGFTEER | 115034 | 114848 | - | 16 |
| 3 | MSDLARSANDSGVLAAVPLH | ATAVQTPAEPNPVICIANTATNGASVDVAFAAHGG | 115454 | 115286 | - | 8 |
| 4 | MSVLARRVNDNNSLATIPLHTTEVRMVESPDSVISTPAT | AAAVVVAFAAGYVAGHSG | 115680 | 115506 | - | 26 |

### Results for WP_171052848.1 [Streptomyces sp. ICN19] back to top

previous - next

#### Architecture

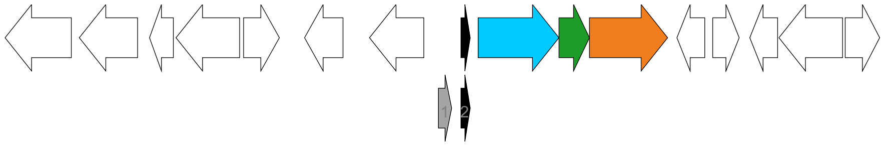

1000 nucleotides

Link to nucleotide sequence

| Accession | start | end | direction | length (aa) | Pfam/HMM | name | description | E-value |
|---|---|---|---|---|---|---|---|---|
| WP_138057321.1 | 1396210 | 1394728 | - | 493 | PF06029<br>PF02805<br>PF12833<br>PF00730<br>PF00165 | AlkA_N<br>Ada_Zn_binding<br>HTH_18<br>HhH-GPD<br>HTH_AraC | AlkA N-terminal domain<br>Metal binding domain of Ada<br>Helix-turn-helix domain<br>HhH-GPD superfamily base excision DNA repair protein<br>Bacterial regulatory helix-turn-helix proteins, AraC family | 6.80E-30<br>5.80E-27<br>6.10E-17<br>8.20E-06<br>1.70E-04 |
| WP_138052231.1 | 1397696 | 1396367 | - | 442 | TIGR00350<br>PF03816<br>PF13399 | TIGR00350<br>LytR_cpsA_psr<br>LytR_C | lytR_cpsA_psr: cell envelope-related function transcriptional attenuator common domain<br>Cell envelope-related transcriptional attenuator domain<br>LytR cell envelope-related transcriptional attenuator | 4.60E-41<br>4.80E-41<br>4.80E-04 |
| WP_138052233.1 | 1398469 | 1397947 | - | 173 | PF01661 | Macro | Macro domain | 4.60E-40 |
| WP_138052235.1 | 1399957 | 1398535 | - | 473 | TIGR02734<br>PF13450<br>TIGR03467<br>TIGR02733<br>PF01593 | TIGR02734<br>NAD_binding_8<br>TIGR03467<br>TIGR02733<br>Amino_oxidase | crtI_fam: phytoene desaturase<br>NAD(P)-binding Rossmann-like domain<br>HpnE: squalene-associated FAD-dependent desaturase<br>desat_CrtD: C-3',4' desaturase CrtD<br>Flavin containing amine oxidoreductase | 3.10E-16<br>5.90E-13<br>6.00E-10<br>9.80E-10<br>6.80E-09 |
| WP_138052237.1 | 1400044 | 1400845 | + | 266 | PF18765<br>PF01909 | Polbeta<br>NTP_transf_2 | Polymerase beta, Nucleotidyltransferase<br>Nucleotidyltransferase domain | 1.10E-06<br>5.10E-06 |
| WP_138052239.1 | 1402281 | 1401423 | - | 285 | PF00459<br>TIGR02067<br>TIGR01331<br>TIGR01330 | Inositol_P<br>TIGR02067<br>TIGR01331<br>TIGR01330 | Inositol monophosphatase family<br>his_9_HisN: histidinol-phosphatase<br>bisphos_cysQ: 3'(2'),5'-bisphosphate nucleotidase<br>bisphos_HAL2: 3'(2'),5'-bisphosphate nucleotidase | 5.40E-56<br>3.00E-39<br>2.60E-26<br>2.10E-10 |
| WP_138052241.1 | 1404074 | 1402874 | - | 399 | NO PFAM MATCH | - | - | - |
| WP_138052243.1 | 1404922 | 1405135 | + | 70 | LinA | LinA | LinA (linaridin precursor) | 5.80E-07 |
| WP_171052848.1 | 1405295 | 1407125 | + | 609 | LinH<br>LinE<br>LinG<br>PF12697<br>TIGR03695 | LinH<br>LinE<br>LinG<br>Abhydrolase_6<br>TIGR03695 | LinH (Fusion of LinE/LinG)<br>LinE Homolog<br>LinG Homolog<br>Alpha/beta hydrolase family<br>menH_SHCHC: 2-succinyl-6-hydroxy-2,4-cyclohexadiene-1-carboxylate synthase | 1.10E-226<br>1.40E-105<br>2.70E-105<br>3.00E-17<br>1.70E-13 |
| WP_138052246.1 | 1407124 | 1407784 | + | 219 | LinL | LinL | LinL | 5.10E-63 |
| WP_138052247.1 | 1407797 | 1409549 | + | 583 | TIGR02204<br>TIGR02203<br>TIGR01846<br>TIGR00958<br>TIGR03375 | TIGR02204<br>TIGR02203<br>TIGR01846<br>TIGR00958<br>TIGR03375 | MsbA_rel: ABC transporter, permease/ATP-binding protein<br>MsbA_lipidA: lipid A export permease/ATP-binding protein MsbA<br>type_I_sec_HlyB: type I secretion system ATPase<br>3a01208: antigen peptide transporter 2<br>type_I_sec_LssB: type I secretion system ATPase | 2.20E-98<br>1.20E-93<br>3.60E-88<br>1.70E-82<br>1.20E-79 |
| WP_138052248.1 | 1410362 | 1409762 | - | 199 | PF00440<br>TIGR03613<br>TIGR03384 | TetR_N<br>TIGR03613<br>TIGR03384 | Bacterial regulatory proteins, tetR family<br>RutR: pyrimidine utilization regulatory protein R<br>betaine_BetI: transcriptional repressor BetI | 3.80E-06<br>1.10E-04<br>1.30E-04 |
| WP_138052249.1 | 1410551 | 1411154 | + | 200 | PF13581 | HATPase_c_2 | Histidine kinase-like ATPase domain | 3.30E-07 |
| WP_138052250.1 | 1412013 | 1411392 | - | 206 | PF17940<br>TIGR03384<br>PF00440 | TetR_C_31<br>TIGR03384<br>TetR_N | Tetracyclin repressor-like, C-terminal domain<br>betaine_BetI: transcriptional repressor BetI<br>Bacterial regulatory proteins, tetR family | 1.90E-17<br>1.40E-13<br>1.20E-10 |
| WP_138052251.1 | 1413446 | 1412018 | - | 475 | PF01425<br>TIGR02715<br>TIGR00132<br>TIGR02713 | Amidase<br>TIGR02715<br>TIGR00132<br>TIGR02713 | Amidase<br>amido_AtzE: amidohydrolase, AtzE family<br>gatA: aspartyl/glutamyl-tRNA(Asn/Gln) amidotransferase, A subunit<br>allophanate_hyd: allophanate hydrolase | 4.10E-110<br>1.40E-73<br>2.90E-58<br>7.80E-56 |
| WP_138052252.1 | 1413502 | 1414297 | + | 264 | PF00106<br>TIGR01963<br>PF13561<br>TIGR01830<br>TIGR03206 | adh_short<br>TIGR01963<br>adh_short_C2<br>TIGR01830<br>TIGR03206 | short chain dehydrogenase<br>PHB_DH: 3-hydroxybutyrate dehydrogenase<br>Enoyl-(Acyl carrier protein) reductase<br>3oxo_ACP_reduc: 3-oxoacyl-[acyl-carrier-protein] reductase<br>benzo_BadH: 2-hydroxycyclohexanecarboxyl-CoA dehydrogenase | 2.00E-37<br>1.10E-27<br>3.20E-27<br>2.50E-23<br>4.40E-19 |

| index | leader | core | start | end | dir | score |
|---|---|---|---|---|---|---|
| 1 | VDPESLDSPEPAAGWCQWSSFRPEY...VPTTTSAATRQTTPRRTRTATTAGQ | RVPLRPRPGDFARRTSPAPGVTTGR | 1404426 | 1404714 | + | 6 |
| 2 | MELLHRVNGDSTLAEVPILAERAVQVKGTEPILNI | FNDMNNFVNAATQPAGEVAADALDVTLEACPATVA | 1404922 | 1405135 | + | 17 |

### Results for WP_147247722.1 [Streptomyces sp. ISID311] back to top

previous - next

#### Architecture

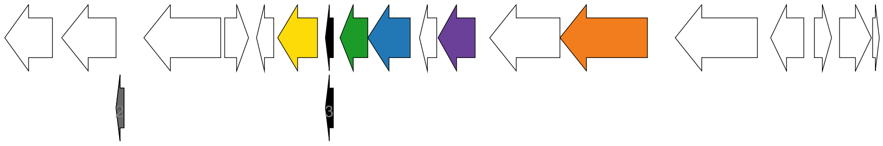

1000 nucleotides

Link to nucleotide sequence

| Accession | start | end | direction | length (aa) | Pfam/HMM | name | description | E-value |
|---|---|---|---|---|---|---|---|---|
| WP_147247718.1 | 83168 | 82160 | - | 335 | TIGR02432<br>PF01171<br>PF09179 | TIGR02432<br>ATP_bind_3<br>TilS | lysidine_TilS_N: tRNA(Ile)-lysidine synthetase<br>PP-loop family<br>TilS substrate binding domain | 3.60E-43<br>6.60E-38<br>2.20E-12 |
| WP_006604046.1 | 84503 | 83360 | - | 380 | TIGR03883<br>TIGR03624<br>PF10103 | TIGR03883<br>TIGR03624<br>Zincin_2 | DUF2342_F420: uncharacterized protein, coenzyme F420 biosynthesis associated<br>TIGR03624: putative hydrolase<br>Zincin-like metallopeptidase | 7.80E-142<br>1.10E-125<br>4.10E-121 |
| WP_147247719.1 | 86692 | 85084 | - | 535 | PF02113<br>TIGR00666<br>PF13354 | Peptidase_S13<br>TIGR00666<br>Beta-lactamase2 | D-Ala-D-Ala carboxypeptidase 3 (S13) family<br>PBP4: D-alanyl-D-alanine carboxypeptidase/D-alanyl-D-alanine-endopeptidase<br>Beta-lactamase enzyme family | 5.90E-76<br>7.20E-58<br>9.40E-04 |
| WP_005314187.1 | 86778 | 87273 | + | 164 | PF00719 | Pyrophosphatase | Inorganic pyrophosphatase | 2.50E-53 |
| WP_147247720.1 | 87815 | 87455 | - | 119 | NO PFAM MATCH | - | - | - |
| WP_093484461.1 | 88734 | 87879 | - | 284 | LinG<br>LinH<br>TIGR04033 | LinG<br>LinH<br>TIGR04033 | LinG Homolog<br>LinH (Fusion of LinE/LinG)<br>export_SdpB: antimicrobial peptide system protein, SdpB family | 3.00E-111<br>3.70E-79<br>2.10E-04 |
| WP_100315524.1 | 89062 | 88879 | - | 60 | LinA | LinA | LinA (linaridin precursor) | 1.80E-13 |
| WP_147247721.1 | 89777 | 89189 | - | 195 | LinL | LinL | LinL | 6.30E-56 |
| WP_147247722.1 | 90676 | 89773 | - | 300 | LinE<br>LinH<br>PF12697<br>TIGR02427<br>PF00561 | LinE<br>LinH<br>Abhydrolase_6<br>TIGR02427<br>Abhydrolase_1 | LinE Homolog<br>LinH (Fusion of LinE/LinG)<br>Alpha/beta hydrolase family<br>protocat_pcaD: 3-oxoadipate enol-lactonase<br>alpha/beta hydrolase fold | 5.30E-98<br>6.70E-52<br>9.90E-16<br>4.30E-12<br>5.80E-10 |
| WP_093484459.1 | 91213 | 90865 | - | 115 | PF13518<br>PF01527<br>PF13551<br>PF13011<br>PF13384 | HTH_28<br>HTH_Tnp_1<br>HTH_29<br>LZ_Tnp_IS481<br>HTH_23 | Helix-turn-helix domain<br>Transposase<br>Winged helix-turn helix<br>leucine-zipper of insertion element IS481<br>Homeodomain-like domain | 4.20E-09<br>2.50E-06<br>1.30E-05<br>1.00E-04<br>4.20E-04 |
| WP_147247723.1 | 92014 | 91255 | - | 252 | PF13649<br>PF08241<br>PF13489<br>PF08242<br>PF13847 | Methyltransf_25<br>Methyltransf_11<br>Methyltransf_23<br>Methyltransf_12<br>Methyltransf_31 | Methyltransferase domain<br>Methyltransferase domain<br>Methyltransferase domain<br>Methyltransferase domain<br>Methyltransferase domain | 5.70E-17<br>1.90E-12<br>1.60E-10<br>5.60E-10<br>7.50E-09 |
| WP_147247724.1 | 93813 | 92331 | - | 493 | TIGR03467<br>PF13450<br>TIGR00292<br>TIGR02734<br>PF01593 | TIGR03467<br>NAD_binding_8<br>TIGR00292<br>TIGR02734<br>Amino_oxidase | HpnE: squalene-associated FAD-dependent desaturase<br>NAD(P)-binding Rossmann-like domain<br>TIGR00292: thiazole biosynthesis enzyme<br>crtI_fam: phytoene desaturase<br>Flavin containing amine oxidoreductase | 2.50E-08<br>4.30E-08<br>5.40E-08<br>9.50E-08<br>1.30E-07 |
| WP_147247813.1 | 95636 | 93809 | - | 608 | TIGR02204<br>TIGR02203<br>TIGR00958<br>TIGR03796<br>TIGR03375 | TIGR02204<br>TIGR02203<br>TIGR00958<br>TIGR03796<br>TIGR03375 | MsbA_rel: ABC transporter, permease/ATP-binding protein<br>MsbA_lipidA: lipid A export permease/ATP-binding protein MsbA<br>3a01208: antigen peptide transporter 2<br>NHLM_micro_ABC1: NHLM bacteriocin system ABC transporter, peptidase/ATP-binding protein<br>type_I_sec_LssB: type I secretion system ATPase | 3.10E-100<br>2.70E-96<br>1.30E-89<br>2.50E-80<br>5.30E-80 |
| WP_147247725.1 | 97944 | 96207 | - | 578 | PF06738<br>PF12821 | ThrE<br>ThrE_2 | Putative threonine/serine exporter<br>Threonine/Serine exporter, ThrE | 4.10E-72<br>1.50E-18 |
| WP_147247726.1 | 98925 | 98205 | - | 239 | PF09335 | SNARE_assoc | SNARE associated Golgi protein | 4.90E-24 |
| WP_147247727.1 | 99132 | 99501 | + | 122 | PF01906<br>TIGR03884 | YbjQ_1<br>TIGR03884 | Putative heavy-metal-binding<br>sel_bind_Methan: selenium-binding protein | 2.00E-33<br>8.80E-04 |
| WP_147247728.1 | 99668 | 100334 | + | 221 | TIGR00949<br>PF01810<br>TIGR00948 | TIGR00949<br>LysE<br>TIGR00948 | 2A76: homoserine/Threonine efflux protein<br>LysE type translocator<br>2a75: L-lysine exporter | 7.70E-45<br>9.90E-44<br>8.60E-16 |
| WP_018103374.1 | 100360 | 100480 | + | 40 | INFERRED GENE | - | - | - |

| index | leader | core | start | end | dir | score |
|---|---|---|---|---|---|---|
| 1 | VRRKAQRREAPRRAGRQRAGAPQS | ATARSPAAGRPTTCGAAAKRNGYPPPRRPRTVMT | 84676 | 84499 | - | 6 |
| 2 | VRVRRKAQRREAPRRAGRQRAGAPQS | ATARSPAAGRPTTCGAAAKRNGYPPPRRPRTVMT | 84682 | 84499 | - | 6 |
| 3 | MSVVANFANTELADVNPGRIGNDATPTM | LTPLAALATPEGVAVTSATAYALHEVTKDIQR | 89062 | 88879 | - | 33 |

### Results for RYJ23219.1 [Streptomyces sp. L-9-10] back to top

previous - next

#### Architecture

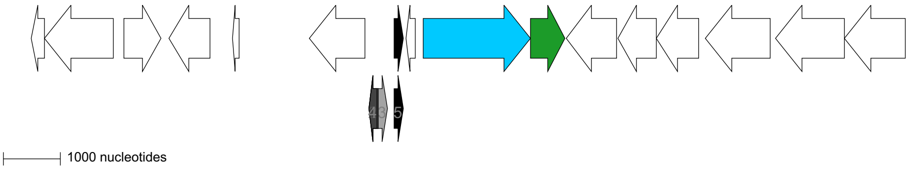

Link to nucleotide sequence

| Accession | start | end | direction | length (aa) | Pfam/HMM | name | description | E-value |
|---|---|---|---|---|---|---|---|---|
| RYJ23211.1 | 68928 | 68691 | - | 78 | PF13370<br>PF13459<br>PF06902 | Fer4_13<br>Fer4_15<br>Fer4_19 | 4Fe-4S single cluster domain of Ferredoxin I<br>4Fe-4S single cluster domain<br>Divergent 4Fe-4S mono-cluster | 7.30E-15<br>3.80E-11<br>3.70E-08 |

| | | | | | | | | |
|---|---|---|---|---|---|---|---|---|
| RYJ23212.1 | 70127 | 68924 | - | 400 | TIGR04458<br>PF00067 | TIGR04458<br>p450 | CYP450_TxtE: 4-nitrotryptophan synthase<br>Cytochrome P450 | 6.70E-41<br>5.00E-32 |
| RYJ23213.1 | 70316 | 70976 | + | 219 | PF13936<br>PF12802<br>PF01047<br>PF13412<br>PF09339 | HTH_38<br>MarR_2<br>MarR<br>HTH_24<br>HTH_IclR | Helix-turn-helix domain<br>MarR family<br>MarR family<br>Winged helix-turn-helix DNA-binding<br>IclR helix-turn-helix domain | 3.60E-16<br>5.80E-12<br>7.70E-09<br>1.60E-08<br>3.90E-08 |
| RYJ23214.1 | 71821 | 71104 | - | 238 | PF17765<br>PF13560 | MLTR_LBD<br>HTH_31 | MmyB-like transcription regulator ligand binding domain<br>Helix-turn-helix domain | 1.60E-46<br>5.10E-04 |
| RYJ23215.1 | 72336 | 72222 | - | 37 | NO PFAM MATCH | - | - | - |
| RYJ23216.1 | 74561 | 73577 | - | 327 | PF01636 | APH | Phosphotransferase enzyme family | 2.70E-13 |
| RYJ23217.1 | 75074 | 75227 | + | 50 | LinA | LinA | LinA (linaridin precursor) | 8.50E-09 |
| RYJ23218.1 | 75446 | 75281 | - | 54 | NO PFAM MATCH | - | - | - |
| RYJ23219.1 | 75584 | 77459 | + | 624 | LinH<br>LinG<br>LinE<br>PF12697<br>LinL | LinH<br>LinG<br>LinE<br>Abhydrolase_6<br>LinL | LinH (Fusion of LinE/LinG)<br>LinG Homolog<br>LinE Homolog<br>Alpha/beta hydrolase family<br>LinL | 1.10E-240<br>1.60E-112<br>5.00E-95<br>3.10E-16<br>4.90E-09 |
| RYJ23220.1 | 77467 | 78058 | + | 196 | LinL | LinL | LinL | 2.50E-69 |
| RYJ23221.1 | 78984 | 78081 | - | 300 | TIGR00070<br>PF01634<br>TIGR03455 | TIGR00070<br>HisG<br>TIGR03455 | hisG: ATP phosphoribosyltransferase<br>ATP phosphoribosyltransferase<br>HisG_C-term: ATP phosphoribosyltransferase, C-terminal domain | 7.90E-50<br>4.20E-38<br>4.40E-06 |
| RYJ23222.1 | 79677 | 78993 | - | 227 | PF00106<br>PF13561<br>TIGR04316<br>TIGR01963<br>TIGR03971 | adh_short<br>adh_short_C2<br>TIGR04316<br>TIGR01963<br>TIGR03971 | short chain dehydrogenase<br>Enoyl-(Acyl carrier protein) reductase<br>dhbA_paeA: 2,3-dihydro-2,3-dihydroxybenzoate dehydrogenase<br>PHB_DH: 3-hydroxybutyrate dehydrogenase<br>SDR_subfam_1: SDR family mycofactocin-dependent oxidoreductase | 1.00E-33<br>3.70E-26<br>2.50E-23<br>2.40E-22<br>3.60E-22 |
| RYJ23223.1 | 80428 | 79684 | - | 247 | PF01259<br>TIGR00081 | SAICAR_synt<br>TIGR00081 | SAICAR synthetase<br>purC: phosphoribosylaminoimidazolesuccinocarboxamide synthase | 9.10E-11<br>1.10E-10 |
| RYJ23224.1 | 81681 | 80523 | - | 385 | TIGR00936<br>PF00670<br>PF05221<br>PF02826<br>TIGR02853 | TIGR00936<br>AdoHcyase_NAD<br>AdoHcyase<br>2-Hacid_dh_C<br>TIGR02853 | ahcY: adenosylhomocysteinase<br>S-adenosyl-L-homocysteine hydrolase, NAD binding domain<br>S-adenosyl-L-homocysteine hydrolase<br>D-isomer specific 2-hydroxyacid dehydrogenase, NAD binding domain<br>spore_dpaA: dipicolinic acid synthetase, A subunit | 1.80E-71<br>3.30E-27<br>3.30E-10<br>2.90E-09<br>1.10E-04 |
| RYJ23225.1 | 82974 | 81768 | - | 401 | PF07690<br>TIGR00900<br>PF11872<br>TIGR00880 | MFS_1<br>TIGR00900<br>DUF3392<br>TIGR00880 | Major Facilitator Superfamily<br>2A0121: H+ Antiporter protein<br>Protein of unknown function (DUF3392)<br>2_A_01_02: multidrug resistance protein | 8.20E-21<br>1.20E-06<br>3.40E-04<br>5.50E-04 |
| RYJ23226.1 | 84047 | 82970 | - | 358 | NO PFAM MATCH | - | - | - |

| index | leader | core | start | end | dir | score |
|---|---|---|---|---|---|---|
| 1 | LWRGSGRAPRPVLADACG | FTPSTPASAAQAKASQHQGLPTRRGTARRGSG | 74773 | 74620 | - | 6 |
| 2 | VLWRGSGRAPRPVLADACG | FTPSTPASAAQAKASQHQGLPTRRGTARRGSG | 74776 | 74620 | - | 6 |
| 3 | LPPWGNFQSPRHRRRCGTVRVMTWTR | HRVRRTPKHPSIVNTPQQDEWVHPRTV | 74780 | 74942 | + | 6 |
| 4 | LPQGGKAVLWRGSGRAPRPVLADACG | FTPSTPASAAQAKASQHQGLPTRRGTARRGSG | 74797 | 74620 | - | 6 |
| 5 | MNDITAFINEDARLDEVPIERAEGTVLVEQGVTVL | ATPSANWCVTFAIGE | 75074 | 75227 | + | 26 |

### Results for PPS88678.1 [Streptomyces sp. MH60] back to top

previous - next

#### Architecture

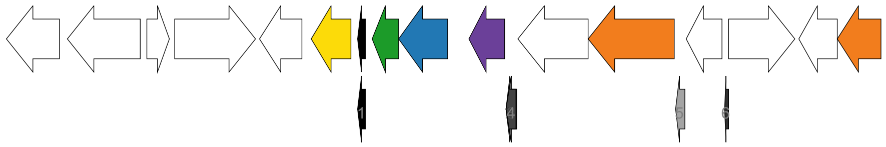

1000 nucleotides

Link to nucleotide sequence

| Accession | start | end | direction | length (aa) | Pfam/HMM | name | description | E-value |
|---|---|---|---|---|---|---|---|---|
| PPS88670.1 | 79287 | 78159 | - | 375 | TIGR03883<br>TIGR03624<br>PF10103 | TIGR03883<br>TIGR03624<br>Zincin_2 | DUF2342_F420: uncharacterized protein, coenzyme F420 biosynthesis associated<br>TIGR03624: putative hydrolase<br>Zincin-like metallopeptidase | 3.20E-144<br>4.80E-129<br>2.20E-120 |
| PPS88671.1 | 81010 | 79456 | - | 517 | PF02113<br>TIGR00666<br>PF00768 | Peptidase_S13<br>TIGR00666<br>Peptidase_S11 | D-Ala-D-Ala carboxypeptidase 3 (S13) family<br>PBP4: D-alanyl-D-alanine carboxypeptidase/D-alanyl-D-alanine-endopeptidase<br>D-alanyl-D-alanine carboxypeptidase | 1.40E-75<br>1.40E-57<br>3.00E-08 |
| PPS88672.1 | 81134 | 81626 | + | 163 | PF00719 | Pyrophosphatase | Inorganic pyrophosphatase | 8.50E-53 |
| PPS88673.1 | 81739 | 83446 | + | 568 | PF06738<br>PF12821 | ThrE<br>ThrE_2 | Putative threonine/serine exporter<br>Threonine/Serine exporter, ThrE | 1.90E-70<br>1.40E-15 |
| PPS88674.1 | 84449 | 83549 | - | 299 | PF09335 | SNARE_assoc | SNARE associated Golgi protein | 4.20E-23 |

| | | | | | | | | |
|---|---|---|---|---|---|---|---|---|
| PPS88675.1 | 85480 | 84634 | - | 281 | LinG<br>LinH<br>TIGR04033 | LinG<br>LinH<br>TIGR04033 | LinG Homolog<br>LinH (Fusion of LinE/LinG)<br>export_SdpB: antimicrobial peptide system protein, SdpB family | 6.40E-109<br>2.80E-78<br>8.50E-05 |
| PPS88676.1 | 85807 | 85624 | - | 60 | LinA | LinA | LinA (linaridin precursor) | 3.40E-13 |
| PPS88677.1 | 86517 | 85935 | - | 193 | LinL | LinL | LinL | 5.30E-57 |
| PPS88678.1 | 87539 | 86513 | - | 341 | LinE<br>LinH<br>PF12697<br>TIGR02427<br>PF00561 | LinE<br>LinH<br>Abhydrolase_6<br>TIGR02427<br>Abhydrolase_1 | LinE Homolog<br>LinH (Fusion of LinE/LinG)<br>Alpha/beta hydrolase family<br>protocat_pcaD: 3-oxoadipate enol-lactonase<br>alpha/beta hydrolase fold | 9.30E-99<br>5.80E-50<br>3.80E-13<br>3.90E-09<br>3.10E-07 |
| PPS88679.1 | 88759 | 88000 | - | 252 | PF13649<br>PF08241<br>PF13489<br>PF08242<br>PF13847 | Methyltransf_25<br>Methyltransf_11<br>Methyltransf_23<br>Methyltransf_12<br>Methyltransf_31 | Methyltransferase domain<br>Methyltransferase domain<br>Methyltransferase domain<br>Methyltransferase domain<br>Methyltransferase domain | 1.60E-15<br>6.30E-10<br>1.10E-07<br>1.30E-07<br>1.10E-06 |
| PPS88680.1 | 90540 | 89055 | - | 494 | TIGR02734<br>TIGR03467<br>TIGR00292<br>PF13450<br>PF01593 | TIGR02734<br>TIGR03467<br>TIGR00292<br>NAD_binding_8<br>Amino_oxidase | crtI_fam: phytoene desaturase<br>HpnE: squalene-associated FAD-dependent desaturase<br>TIGR00292: thiazole biosynthesis enzyme<br>NAD(P)-binding Rossmann-like domain<br>Flavin containing amine oxidoreductase | 6.60E-10<br>2.60E-08<br>3.50E-08<br>1.80E-07<br>9.40E-07 |
| PPS88681.1 | 92384 | 90536 | - | 615 | TIGR02204<br>TIGR02203<br>TIGR00958<br>TIGR03375<br>TIGR03796 | TIGR02204<br>TIGR02203<br>TIGR00958<br>TIGR03375<br>TIGR03796 | MsbA_rel: ABC transporter, permease/ATP-binding protein<br>MsbA_lipidA: lipid A export permease/ATP-binding protein MsbA<br>3a01208: antigen peptide transporter 2<br>type_I_sec_LssB: type I secretion system ATPase<br>NHLM_micro_ABC1: NHLM bacteriocin system ABC transporter, peptidase/ATP-binding protein | 4.00E-99<br>1.40E-96<br>5.50E-89<br>2.70E-79<br>6.60E-79 |
| PPS88682.1 | 93403 | 92641 | - | 253 | PF07739<br>PF13411<br>TIGR02043<br>TIGR02044<br>PF00376 | TipAS<br>MerR_1<br>TIGR02043<br>TIGR02044<br>MerR | TipAS antibiotic-recognition domain<br>MerR HTH family regulatory protein<br>ZntR: Zn(II)-responsive transcriptional regulator<br>CueR: Cu(I)-responsive transcriptional regulator<br>MerR family regulatory protein | 4.70E-38<br>6.40E-17<br>4.50E-15<br>3.10E-14<br>8.10E-14 |
| PPS88683.1 | 93522 | 94950 | + | 475 | TIGR01788<br>PF00282<br>TIGR03812<br>TIGR03799<br>TIGR03811 | TIGR01788<br>Pyridoxal_deC<br>TIGR03812<br>TIGR03799<br>TIGR03811 | Glu-decarb-GAD: glutamate decarboxylase<br>Pyridoxal-dependent decarboxylase conserved domain<br>tyr_de_CO2_Arch: tyrosine decarboxylase MnfA<br>NOD_PanD_pyr: putative pyridoxal-dependent aspartate 1-decarboxylase<br>tyr_de_CO2_Ent: tyrosine decarboxylase | 1.20E-188<br>1.90E-70<br>1.80E-40<br>1.70E-06<br>3.30E-04 |
| PPS88684.1 | 95845 | 95041 | - | 267 | PF01061<br>TIGR00025<br>TIGR01247<br>PF12698<br>TIGR01248 | ABC2_membrane<br>TIGR00025<br>TIGR01247<br>ABC2_membrane_3<br>TIGR01248 | ABC-2 type transporter<br>Mtu_efflux: ABC transporter efflux protein, DrrB family<br>drrB: daunorubicin resistance ABC transporter membrane protein<br>ABC-2 family transporter protein<br>drrC: daunorubicin resistance protein C | 8.70E-25<br>2.90E-21<br>1.30E-12<br>3.40E-12<br>9.20E-09 |
| PPS88685.1 | 96783 | 95841 | - | 313 | TIGR01188<br>TIGR01288<br>TIGR03864<br>TIGR03522<br>TIGR03740 | TIGR01188<br>TIGR01288<br>TIGR03864<br>TIGR03522<br>TIGR03740 | drrA: daunorubicin resistance ABC transporter, ATP-binding protein<br>nodI: nodulation ABC transporter NodI<br>PQQ_ABC_ATP: ABC transporter, ATP-binding subunit, PQQ-dependent alcohol dehydrogenase system<br>GldA_ABC_ATP: gliding motility-associated ABC transporter ATP-binding subunit GldA<br>galliderm_ABC: lantibiotic protection ABC transporter, ATP-binding subunit | 9.10E-103<br>6.50E-60<br>1.10E-59<br>1.70E-51<br>1.60E-43 |

| index | leader | core | start | end | dir | score |
|---|---|---|---|---|---|---|
| 1 | MSVVADFANTELTDVTPGRMGNDATPTM | LTPLATLATPEGVAVTSATAYALNEITNDLAG | 85807 | 85624 | - | 32 |
| 2 | VSESAADLHRSGHSAFGTVRQRGLPTDFRWGV | TVRRSTGRTAGCAFFAADPSAHHVLAGSRPPAS | 89001 | 88803 | - | 6 |
| 3 | VVSESAADLHRSGHSAFGTVRQRGLPTDFRWGV | TVRRSTGRTAGCAFFAADPSAHHVLAGSRPPAS | 89004 | 88803 | - | 6 |
| 4 | VARVVSESAADLHRSGHSAFGTVRQRGLPTDFRW | GVTVRRSTGRTAGCAFFAADPSAHHVLAGSRPPAS | 89013 | 88803 | - | 6 |
| 5 | VGRRGDPPAAGGAVTFPCAAPRLPAGTRSRPTG | QVTPNLRPGLRVTFGRLDDGLVRQGCPPDPARSP | 92614 | 92410 | - | 6 |
| 6 | LCSDMQPTVGRGPAP | RTRGRTGVHGLAPHVT | 93536 | 93440 | - | 9 |

### Results for SPF07382.1 [Streptomyces sp. MA5143a] back to top

previous - next

#### Architecture

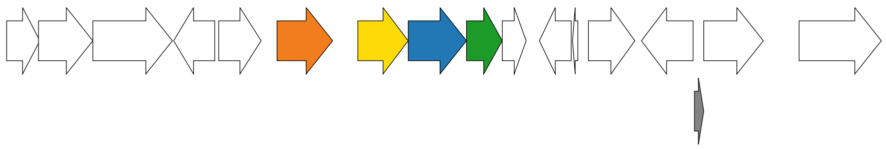

1000 nucleotides

Link to nucleotide sequence

| Accession | start | end | direction | length (aa) | Pfam/HMM | name | description | E-value |
|---|---|---|---|---|---|---|---|---|
| SPF07375.1 | 28 | 601 | + | 190 | PF00596<br>TIGR01086<br>TIGR03328<br>TIGR00760 | Aldolase_II<br>TIGR01086<br>TIGR03328<br>TIGR00760 | Class II Aldolase and Adducin N-terminal domain<br>fucA: L-fuculose phosphate aldolase<br>salvage_mtnB: methylthioribulose-1-phosphate dehydratase<br>araD: L-ribulose-5-phosphate 4-epimerase | 1.90E-31<br>1.00E-25<br>2.30E-14<br>9.20E-09 |
| SPF07376.1 | 597 | 1539 | + | 313 | PF00294<br>TIGR02152<br>TIGR04382<br>TIGR03168<br>TIGR02198 | PfkB<br>TIGR02152<br>TIGR04382<br>TIGR03168<br>TIGR02198 | pfkB family carbohydrate kinase<br>D_ribokin_bact: ribokinase<br>myo_inos_iolC_N: 5-dehydro-2-deoxygluconokinase<br>1-PFK: hexose kinase, 1-phosphofructokinase family<br>rfaE_dom_I: bifunctional protein RfaE, domain I | 1.50E-43<br>1.40E-27<br>4.60E-27<br>4.40E-17<br>2.90E-16 |

| | | | | | | | | |
|---|---|---|---|---|---|---|---|---|
| SPF07377.1 | 1540 | 2944 | + | 467 | PF00171<br>TIGR01804<br>TIGR04284<br>TIGR01780<br>TIGR02299 | Aldedh<br>TIGR01804<br>TIGR04284<br>TIGR01780<br>TIGR02299 | Aldehyde dehydrogenase family<br>BADH: betaine-aldehyde dehydrogenase<br>aldehy_Rv0768: aldehyde dehydrogenase, Rv0768 family<br>SSADH: succinate-semialdehyde dehydrogenase<br>HpaE: 5-carboxymethyl-2-hydroxymuconate semialdehyde dehydrogenase | 2.10E-83<br>1.10E-52<br>2.70E-52<br>1.50E-51<br>1.90E-49 |
| SPF07378.1 | 3697 | 2977 | - | 239 | NO PFAM MATCH | - | - | - |
| SPF07379.1 | 3770 | 4505 | + | 244 | PF00196<br>TIGR03541<br>TIGR03020<br>PF08281<br>TIGR02937 | GerE<br>TIGR03541<br>TIGR03020<br>Sigma70_r4_2<br>TIGR02937 | Bacterial regulatory proteins, luxR family<br>reg_near_HchA: LuxR family transcriptional regulatory, chaperone HchA-associated<br>EpsA: transcriptional regulator EpsA<br>Sigma-70, region 4<br>sigma70-ECF: RNA polymerase sigma factor, sigma-70 family | 3.00E-15<br>2.10E-07<br>7.50E-07<br>6.00E-06<br>6.20E-05 |
| SPF07380.1 | 4791 | 5769 | + | 325 | TIGR01188<br>TIGR03864<br>TIGR01288<br>TIGR03522<br>TIGR00968 | TIGR01188<br>TIGR03864<br>TIGR01288<br>TIGR03522<br>TIGR00968 | drrA: daunorubicin resistance ABC transporter, ATP-binding protein<br>PQQ_ABC_ATP: ABC transporter, ATP-binding subunit, PQQ-dependent alcohol dehydrogenase system<br>nodI: nodulation ABC transporter NodI<br>GldA_ABC_ATP: gliding motility-associated ABC transporter ATP-binding subunit GldA<br>3a0106s01: sulfate ABC transporter, ATP-binding protein | 2.20E-85<br>7.60E-59<br>4.20E-58<br>4.20E-50<br>5.40E-41 |
| SPF07381.1 | 6199 | 7096 | + | 298 | LinG<br>LinH | LinG<br>LinH | LinG Homolog<br>LinH (Fusion of LinE/LinG) | 9.10E-97<br>9.30E-92 |
| SPF07382.1 | 7095 | 8112 | + | 338 | LinE<br>LinH<br>PF12697<br>TIGR02427<br>PF00561 | LinE<br>LinH<br>Abhydrolase_6<br>TIGR02427<br>Abhydrolase_1 | LinE Homolog<br>LinH (Fusion of LinE/LinG)<br>Alpha/beta hydrolase family<br>protocat_pcaD: 3-oxoadipate enol-lactonase<br>alpha/beta hydrolase fold | 3.50E-86<br>3.50E-74<br>7.10E-15<br>1.10E-11<br>6.40E-11 |
| SPF07383.1 | 8108 | 8741 | + | 210 | LinL | LinL | LinL | 4.60E-50 |
| SPF07384.1 | 8737 | 9166 | + | 142 | PF04134 | DUF393 | Protein of unknown function, DUF393 | 2.70E-18 |
| SPF07385.1 | 9964 | 9403 | - | 186 | NO PFAM MATCH | - | - | - |
| SPF07386.1 | 10075 | 9985 | - | 29 | NO PFAM MATCH | - | - | - |
| SPF07387.1 | 10251 | 11079 | + | 275 | PF01061<br>TIGR03861<br>TIGR01247 | ABC2_membrane<br>TIGR03861<br>TIGR01247 | ABC-2 type transporter<br>phenyl_ABC_PedC: alcohol ABC transporter, permease protein<br>drrB: daunorubicin resistance ABC transporter membrane protein | 4.60E-15<br>9.80E-09<br>6.50E-08 |
| SPF07388.1 | 12099 | 11199 | - | 299 | TIGR03621<br>PF00296<br>TIGR03560<br>TIGR03856<br>TIGR03619 | TIGR03621<br>Bac_luciferase<br>TIGR03560<br>TIGR03856<br>TIGR03619 | F420_MSMEG_2516: probable F420-dependent oxidoreductase, MSMEG_2516 family<br>Luciferase-like monooxygenase<br>F420_Rv1855c: probable F420-dependent oxidoreductase, Rv1855c family<br>F420_MSMEG_2906: probable F420-dependent oxidoreductase, MSMEG_2906 family<br>F420_Rv2161c: probable F420-dependent oxidoreductase, Rv2161c family | 2.20E-60<br>8.70E-23<br>2.00E-22<br>2.10E-19<br>9.00E-18 |
| SPF07389.1 | 12283 | 13333 | + | 349 | PF13469<br>PF00685 | Sulfotransfer_3<br>Sulfotransfer_1 | Sulfotransferase family<br>Sulfotransferase domain | 6.60E-23<br>1.30E-09 |
| SPF07390.1 | 13976 | 15413 | + | 478 | PF13581<br>PF00668 | HATPase_c_2<br>Condensation | Histidine kinase-like ATPase domain<br>Condensation domain | 4.20E-07<br>1.10E-05 |

| index | leader | core | start | end | dir | score |
|---|---|---|---|---|---|---|
| 1 | MLLADRLRLPGGNQHVGADHSGTFDP | VISPAGRLASSGTWATCMRRGQRMVTT | 12125 | 12287 | + | 7 |

### Results for WP_063346941.1 [Streptomyces sp. MJM8645] back to top

previous - next

#### Architecture

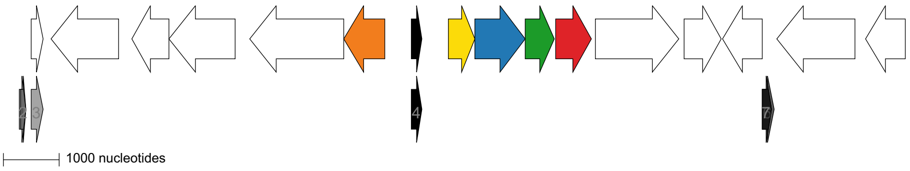

Link to nucleotide sequence

| Accession | start | end | direction | length (aa) | Pfam/HMM | name | description | E-value |
|---|---|---|---|---|---|---|---|---|
| WP_159398381.1 | 1615857 | 1616079 | + | 73 | PF06131 | DUF963 | Schizosaccharomyces pombe repeat of unknown function (DUF963) | 6.30E-05 |
| WP_063346936.1 | 1617441 | 1616232 | - | 402 | PF07859<br>PF00326<br>PF00135<br>PF01738 | Abhydrolase_3<br>Peptidase_S9<br>COesterase<br>DLH | alpha/beta hydrolase fold<br>Prolyl oligopeptidase family<br>Carboxylesterase family<br>Dienelactone hydrolase family | 1.10E-21<br>8.20E-10<br>2.10E-08<br>2.30E-06 |
| WP_063346937.1 | 1618347 | 1617672 | - | 224 | PF00072<br>PF00196<br>TIGR02875<br>TIGR03020<br>TIGR03541 | Response_reg<br>GerE<br>TIGR02875<br>TIGR03020<br>TIGR03541 | Response regulator receiver domain<br>Bacterial regulatory proteins, luxR family<br>spore_0_A: sporulation transcription factor Spo0A<br>EpsA: transcriptional regulator EpsA<br>reg_near_HchA: LuxR family transcriptional regulatory, chaperone HchA-associated | 3.50E-25<br>4.60E-20<br>8.80E-17<br>2.20E-14<br>4.10E-10 |
| WP_079136210.1 | 1619513 | 1618343 | - | 389 | PF07730 | HisKA_3 | Histidine kinase | 5.30E-19 |
| WP_159398382.1 | 1621472 | 1619795 | - | 558 | PF16962 | ABC_export | Putative ABC exporter | 3.60E-04 |
| WP_079135486.1 | 1622215 | 1621477 | - | 245 | TIGR01189<br>TIGR03265<br>TIGR02857<br>TIGR03864<br>TIGR00968 | TIGR01189<br>TIGR03265<br>TIGR02857<br>TIGR03864<br>TIGR00968 | ccmA: heme ABC exporter, ATP-binding protein CcmA<br>PhnT2: putative 2-aminoethylphosphonate ABC transporter, ATP-binding protein<br>CydD: thiol reductant ABC exporter, CydD subunit<br>PQQ_ABC_ATP: ABC transporter, ATP-binding subunit, PQQ-dependent alcohol dehydrogenase system<br>3a0106s01: sulfate ABC transporter, ATP-binding protein | 2.60E-34<br>1.10E-32<br>2.00E-31<br>2.10E-31<br>2.90E-31 |
| WP_159398383.1 | 1622693 | 1622867 | + | 57 | LinA | LinA | LinA (linaridin precursor) | 1.00E-10 |

| | | | | | | | | |
|---|---|---|---|---|---|---|---|---|
| WP_079136211.1 | 1623358 | 1623823 | + | 154 | LinG | LinG | LinG Homolog | 1.70E-57 |
| | | | | | LinH | LinH | LinH (Fusion of LinE/LinG) | 7.20E-53 |
| | | | | | TIGR04033 | TIGR04033 | export_SdpB: antimicrobial peptide system protein, SdpB family | 1.50E-06 |
| WP_063346941.1 | 1623819 | 1624731 | + | 303 | LinE | LinE | LinE Homolog | 2.40E-69 |
| | | | | | LinH | LinH | LinH (Fusion of LinE/LinG) | 3.70E-43 |
| | | | | | PF12697 | Abhydrolase_6 | Alpha/beta hydrolase family | 6.10E-09 |
| | | | | | TIGR02427 | TIGR02427 | protocat_pcaD: 3-oxoadipate enol-lactonase | 1.00E-08 |
| | | | | | TIGR03056 | TIGR03056 | bchO_mg_che_rel: putative magnesium chelatase accessory protein | 6.40E-07 |
| WP_063346942.1 | 1624727 | 1625258 | + | 176 | LinL | LinL | LinL | 5.00E-41 |
| WP_159398384.1 | 1625275 | 1625914 | + | 212 | TIGR00521 | TIGR00521 | coaBC_dfp: phosphopantothenoylcysteine decarboxylase / phosphopantothenate--cysteine ligase | 9.60E-14 |
| | | | | | PF02441 | Flavoprotein | Flavoprotein | 3.50E-13 |
| | | | | | TIGR02113 | TIGR02113 | coaC_strep: phosphopantothenoylcysteine decarboxylase | 1.70E-08 |
| WP_063346943.1 | 1625991 | 1627473 | + | 493 | TIGR02734 | TIGR02734 | crtI_fam: phytoene desaturase | 2.10E-12 |
| | | | | | TIGR03467 | TIGR03467 | HpnE: squalene-associated FAD-dependent desaturase | 9.00E-10 |
| | | | | | PF13450 | NAD_binding_8 | NAD(P)-binding Rossmann-like domain | 1.40E-09 |
| | | | | | TIGR01317 | TIGR01317 | GOGAT_sm_gam: glutamate synthase, NADH/NADPH, small subunit | 1.80E-09 |
| | | | | | TIGR01318 | TIGR01318 | gltD_gamma_fam: glutamate synthase, small subunit | 8.80E-08 |
| WP_063346944.1 | 1627563 | 1628235 | + | 223 | TIGR01840 | TIGR01840 | esterase_phb: esterase, PHB depolymerase family | 4.90E-15 |
| | | | | | PF00326 | Peptidase_S9 | Prolyl oligopeptidase family | 2.90E-14 |
| | | | | | PF10503 | Esterase_phd | Esterase PHB depolymerase | 5.40E-12 |
| | | | | | PF01738 | DLH | Dienelactone hydrolase family | 6.80E-12 |
| | | | | | PF12146 | Hydrolase_4 | Serine aminopeptidase, S33 | 1.30E-09 |
| WP_063346945.1 | 1628982 | 1628259 | - | 240 | PF10042 | DUF2278 | Uncharacterized conserved protein (DUF2278) | 4.30E-70 |
| WP_063346946.1 | 1630626 | 1629240 | - | 461 | PF00654 | Voltage_CLC | Voltage gated chloride channel | 1.70E-19 |
| WP_107453010.1 | 1631528 | 1630829 | - | 232 | TIGR03968 | TIGR03968 | mycofact_TetR: mycofactocin system transcriptional regulator | 2.00E-27 |
| | | | | | PF00440 | TetR_N | Bacterial regulatory proteins, tetR family | 2.20E-14 |
| | | | | | PF17754 | TetR_C_14 | MftR C-terminal domain | 2.30E-08 |
| | | | | | TIGR03613 | TIGR03613 | RutR: pyrimidine utilization regulatory protein R | 8.00E-08 |
| | | | | | TIGR03384 | TIGR03384 | betaine_BetI: transcriptional repressor BetI | 1.20E-04 |

| index | leader | core | start | end | dir | score |
|---|---|---|---|---|---|---|
| 1 | MRWLTTFDDDSLNRRSIM | DAYVGSCVLDERRSITVAT | 1615665 | 1615779 | + | 6 |
| 2 | LTTFDDDSLNRRSIMDA | YVGSCVLDERRSITVAT | 1615674 | 1615779 | + | 6 |
| 3 | MLKRVVMGVATAALFSLGTGVALANAGVDSTARPAPARYSTPVST | PTPVAESAPAGESGTDSTQAWWNHSTHY | 1615857 | 1616079 | + | 6 |
| 4 | MNATESRVEFVTDAVNASFAEAQEGFVNAHNAPAM | TTPTIIAVCEVVVTGSTICLGC | 1622693 | 1622867 | + | 31 |
| 5 | MGVVPRFPGCWSWCLAGRSAGVPPLS | RTPPTGGCGIRSCSSAVPRTGFRAAER | 1628980 | 1629142 | + | 7 |
| 6 | VWFLGSLAAGPGASPAGRPGFLRYRAHRRPGAV | ESAVVHLPSPERAFAPQNVDVGAPSSGDWTRAG | 1628984 | 1629185 | + | 8 |
| 7 | VVPRFPGCWSWCLAGRSAGVPPLS | RTPPTGGCGIRSCSSAVPRTGFRAAER | 1628986 | 1629142 | + | 7 |

### Results for SBU99212.1 [Streptomyces sp. MnatMP-M77] back to top

previous - next

#### Architecture

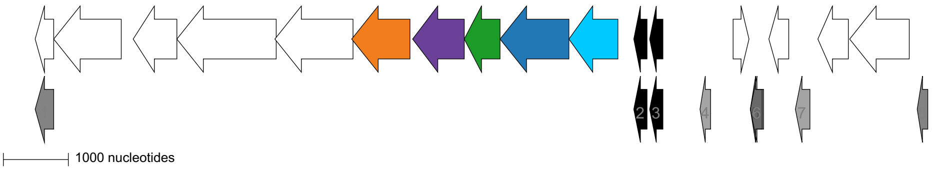

Link to nucleotide sequence

| Accession | start | end | direction | length (aa) | Pfam/HMM | name | description | E-value |
|---|---|---|---|---|---|---|---|---|
| SBU99185.1 | 46514 | 46232 | - | 93 | PF00550 | PP-binding | Phosphopantetheine attachment site | 7.70E-09 |
| | | | | | TIGR01688 | TIGR01688 | dltC: D-alanine--poly(phosphoribitol) ligase, subunit 2 | 1.90E-05 |
| SBU99187.1 | 47554 | 46510 | - | 347 | TIGR00747 | TIGR00747 | fabH: 3-oxoacyl-[acyl-carrier-protein] synthase III | 4.10E-08 |
| | | | | | PF08541 | ACP_syn_III_C | 3-Oxoacyl-[acyl-carrier-protein (ACP)] synthase III C terminal | 1.10E-04 |
| SBU99190.1 | 48406 | 47743 | - | 220 | PF05719 | GPP34 | Golgi phosphoprotein 3 (GPP34) | 6.90E-49 |
| SBU99192.1 | 49937 | 48422 | - | 504 | TIGR02734 | TIGR02734 | crtI_fam: phytoene desaturase | 1.50E-11 |
| | | | | | TIGR03467 | TIGR03467 | HpnE: squalene-associated FAD-dependent desaturase | 1.50E-08 |
| | | | | | PF13450 | NAD_binding_8 | NAD(P)-binding Rossmann-like domain | 4.10E-08 |
| | | | | | TIGR00292 | TIGR00292 | TIGR00292: thiazole biosynthesis enzyme | 4.70E-08 |
| | | | | | TIGR03997 | TIGR03997 | mycofact_OYE_2: mycofactocin system FadH/OYE family oxidoreductase 2 | 6.20E-07 |
| SBU99196.1 | 51120 | 49929 | - | 396 | PF12698 | ABC2_membrane_3 | ABC-2 family transporter protein | 5.80E-31 |
| SBU99200.1 | 52012 | 51100 | - | 303 | TIGR03522 | TIGR03522 | GldA_ABC_ATP: gliding motility-associated ABC transporter ATP-binding subunit GldA | 7.90E-53 |
| | | | | | TIGR01188 | TIGR01188 | drrA: daunorubicin resistance ABC transporter, ATP-binding protein | 1.00E-45 |
| | | | | | TIGR03864 | TIGR03864 | PQQ_ABC_ATP: ABC transporter, ATP-binding subunit, PQQ-dependent alcohol dehydrogenase system | 3.90E-44 |
| | | | | | TIGR04406 | TIGR04406 | LPS_export_lptB: LPS export ABC transporter ATP-binding protein | 4.40E-41 |
| | | | | | TIGR03740 | TIGR03740 | galliderm_ABC: lantibiotic protection ABC transporter, ATP-binding subunit | 1.00E-40 |
| SBU99204.1 | 52836 | 52047 | - | 262 | PF13649 | Methyltransf_25 | Methyltransferase domain | 5.10E-21 |
| | | | | | PF08241 | Methyltransf_11 | Methyltransferase domain | 2.70E-17 |
| | | | | | PF13847 | Methyltransf_31 | Methyltransferase domain | 1.00E-14 |
| | | | | | PF13489 | Methyltransf_23 | Methyltransferase domain | 2.20E-12 |
| | | | | | PF08242 | Methyltransf_12 | Methyltransferase domain | 6.20E-12 |

| | | | | | | | | |
|---|---|---|---|---|---|---|---|---|
| SBU99208.1 | 53399 | 52832 | - | 188 | LinL | LinL | LinL | 9.70E-71 |
| SBU99212.1 | 54460 | 53398 | - | 353 | LinE<br>LinH<br>PF12697<br>PF00561<br>TIGR03695 | LinE<br>LinH<br>Abhydrolase_6<br>Abhydrolase_1<br>TIGR03695 | LinE Homolog<br>LinH (Fusion of LinE/LinG)<br>Alpha/beta hydrolase family<br>alpha/beta hydrolase fold<br>menH_SHCHC: 2-succinyl-6-hydroxy-2,4-cyclohexadiene-1-carboxylate synthase | 9.00E-127<br>2.20E-113<br>8.60E-16<br>6.10E-09<br>4.40E-08 |
| SBU99216.1 | 55200 | 54456 | - | 247 | LinH<br>LinG | LinH<br>LinG | LinH (Fusion of LinE/LinG)<br>LinG Homolog | 6.00E-106<br>3.50E-101 |
| SBU99220.1 | 55661 | 55457 | - | 67 | LinA | LinA | LinA (linaridin precursor) | 5.60E-12 |
| SBU99224.1 | 55891 | 55693 | - | 65 | LinA | LinA | LinA (linaridin precursor) | 2.10E-08 |
| SBU99227.1 | 56976 | 57216 | + | 79 | NO PFAM MATCH | - | - | - |
| SBU99231.1 | 57840 | 57540 | - | 99 | NO PFAM MATCH | - | - | - |
| SBU99235.1 | 58757 | 58283 | - | 157 | NO PFAM MATCH | - | - | - |
| SBU99239.1 | 59692 | 58774 | - | 305 | PF13560<br>PF01381<br>PF12844<br>PF13744 | HTH_31<br>HTH_3<br>HTH_19<br>HTH_37 | Helix-turn-helix domain<br>Helix-turn-helix<br>Helix-turn-helix domain<br>Helix-turn-helix domain | 1.20E-09<br>1.50E-07<br>7.40E-04<br>9.10E-04 |

| index | leader | core | start | end | dir | score |
|---|---|---|---|---|---|---|
| 1 | MSRGPAETLDEITRFLTEALRREVGPDDDYFALGLVDSLFALELVT | YVEDRFELTVEVEDLDLDSFRTAGRLTAFVHAKKAGAETTPAHAGDN | 46514 | 46232 | - | 7 |
| 2 | MSVVLDKVNASVEFGAIPGHISEAEWGNATDVPVL | ATPVTLTPTIGKVTLGVGAFTGGIAVGDALFG | 55661 | 55457 | - | 30 |
| 3 | MSTLLQKVNTSQELHAVASDLADAKDLSLTLPVL | ATITTVCPATGYIVGGGLFATGIGITYTVNR | 55891 | 55693 | - | 29 |
| 4 | LSGRNAQSHPPLDEPDIRSSGLPC | RPPDSPRDHLEALGDNQTQSQCERIYCASL | 56631 | 56466 | - | 6 |
| 5 | MGEVIGVVGRRAHCLTCERPHEDAAMPEGVMCSNCART | AVPGPCRRCSARCPTARPFGTWTR | 57426 | 57237 | - | 6 |
| 6 | MSAYTPSMGEVIGVVGRRAHCLTCERPHEDAAMPEGVMCSNCART | AVPGPCRRCSARCPTARPFGTWTR | 57447 | 57237 | - | 6 |
| 7 | VCERDAAAGEAVAPSG | PAPRAPVATEVGIAPRTQHVHLTVL...APRSIPSFRPHRRRPPPAGGQPPSP | 58169 | 57944 | - | 6 |
| 8 | VECHRCHRAGKSSRCCRFPAAARLAELFR | PAGAARAGASADGLSWIVPGAPPDQPDLT | 59986 | 59809 | - | 7 |

### Results for REK89866.1 [Streptomyces sp. NEAU-D10] back to top

previous - next

#### Architecture

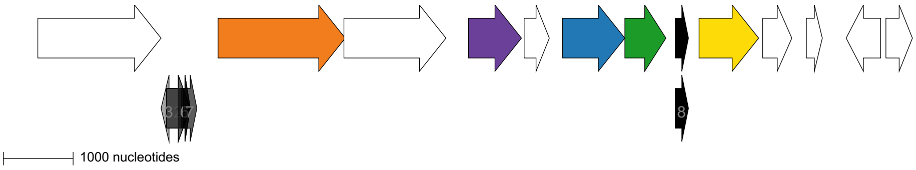

Link to nucleotide sequence

| Accession | start | end | direction | length (aa) | Pfam/HMM | name | description | E-value |
|---|---|---|---|---|---|---|---|---|
| REK89864.1 | 216 | 1989 | + | 590 | PF06738<br>PF12821 | ThrE<br>ThrE_2 | Putative threonine/serine exporter<br>Threonine/Serine exporter, ThrE | 1.00E-72<br>1.40E-18 |
| REK89873.1 | 2795 | 4622 | + | 608 | TIGR02204<br>TIGR02203<br>TIGR00958<br>TIGR03796<br>TIGR03375 | TIGR02204<br>TIGR02203<br>TIGR00958<br>TIGR03796<br>TIGR03375 | MsbA_rel: ABC transporter, permease/ATP-binding protein<br>MsbA_lipidA: lipid A export permease/ATP-binding protein MsbA<br>3a01208: antigen peptide transporter 2<br>NHLM_micro_ABC1: NHLM bacteriocin system ABC transporter, peptidase/ATP-binding protein<br>type_I_sec_LssB: type I secretion system ATPase | 9.50E-102<br>1.20E-97<br>3.60E-89<br>1.20E-82<br>1.60E-80 |
| REK89874.1 | 4618 | 6073 | + | 484 | TIGR00292<br>PF13450<br>TIGR03467<br>TIGR02734<br>PF01593 | TIGR00292<br>NAD_binding_8<br>TIGR03467<br>TIGR02734<br>Amino_oxidase | TIGR00292: thiazole biosynthesis enzyme<br>NAD(P)-binding Rossmann-like domain<br>HpnE: squalene-associated FAD-dependent desaturase<br>crtI_fam: phytoene desaturase<br>Flavin containing amine oxidoreductase | 1.40E-08<br>2.30E-08<br>2.70E-08<br>3.10E-08<br>5.90E-07 |
| REK89865.1 | 6401 | 7160 | + | 252 | PF13649<br>PF08241<br>PF08242<br>PF13489<br>PF13847 | Methyltransf_25<br>Methyltransf_11<br>Methyltransf_12<br>Methyltransf_23<br>Methyltransf_31 | Methyltransferase domain<br>Methyltransferase domain<br>Methyltransferase domain<br>Methyltransferase domain<br>Methyltransferase domain | 2.60E-15<br>1.70E-11<br>1.20E-09<br>1.50E-07<br>1.80E-07 |
| REK89875.1 | 7202 | 7550 | + | 115 | PF13518<br>PF01527<br>PF13551<br>PF13384<br>PF13011 | HTH_28<br>HTH_Tnp_1<br>HTH_29<br>HTH_23<br>LZ_Tnp_IS481 | Helix-turn-helix domain<br>Transposase<br>Winged helix-turn helix<br>Homeodomain-like domain<br>leucine-zipper of insertion element IS481 | 2.30E-09<br>3.70E-07<br>5.60E-07<br>1.10E-05<br>1.20E-05 |

| | | | | | | | | |
|---|---|---|---|---|---|---|---|---|
| REK89866.1 | 7739 | 8642 | + | 300 | LinE<br>LinH<br>PF12697<br>TIGR02427<br>PF00561 | LinE<br>LinH<br>Abhydrolase_6<br>TIGR02427<br>Abhydrolase_1 | LinE Homolog<br>LinH (Fusion of LinE/LinG)<br>Alpha/beta hydrolase family<br>protocat_pcaD: 3-oxoadipate enol-lactonase<br>alpha/beta hydrolase fold | 2.00E-96<br>1.50E-52<br>9.60E-17<br>4.00E-12<br>1.40E-10 |
| REK89867.1 | 8638 | 9226 | + | 195 | LinL | LinL | LinL | 1.80E-57 |
| REK89868.1 | 9355 | 9550 | + | 64 | LinA | LinA | LinA (linaridin precursor) | 7.00E-12 |
| REK89869.1 | 9696 | 10551 | + | 284 | LinG<br>LinH | LinG<br>LinH | LinG Homolog<br>LinH (Fusion of LinE/LinG) | 3.20E-107<br>1.30E-76 |
| REK89870.1 | 10617 | 11034 | + | 138 | NO PFAM MATCH | - | - | - |
| WP_014135373.1 | 11237 | 11460 | + | 74 | INFERRED GENE | - | - | - |
| REK89871.1 | 12296 | 11801 | - | 164 | PF00719 | Pyrophosphatase | Inorganic pyrophosphatase | 2.30E-53 |
| REK89872.1 | 12382 | 12757 | + | 125 | NO PFAM MATCH | - | - | - |

| index | leader | core | start | end | dir | score |
|---|---|---|---|---|---|---|
| 1 | LVQFAPVCEWRAGGMSMDFPVPTQVSAYANSHLGTVTPGA | VTAWQGVPVLRSAKAFHTAVSPSST...TVRSLPAGCSVAEASPVPQEFLKAL | 2060 | 2414 | + | 6 |
| 2 | VQFAPVCEWRAGGMSMDFPVPTQVSAYANSHLGTVTPGA | VTAWQGVPVLRSAKAFHTAVSPSST...TVRSLPAGCSVAEASPVPQEFLKAL | 2063 | 2414 | + | 6 |
| 3 | MESLCTPQYRDSLPRSDRPGSDRPQ...GHATRSPFTDRRELHQRRTEHTPTA | VTAREPPAVAPRAVP | 2231 | 1985 | - | 6 |
| 4 | VQMQVALQQEIGHSASQESCIRVSA | TTVRSLPAGCSVAEASPVPQEFLKAL | 2258 | 2414 | + | 8 |
| 5 | MQVALQQEIGHSASQESCIRVSA | TTVRSLPAGCSVAEASPVPQEFLKAL | 2264 | 2414 | + | 6 |
| 6 | VALQQEIGHSASQESCIRVSA | TTVRSLPAGCSVAEASPVPQEFLKAL | 2270 | 2414 | + | 6 |
| 7 | VYQSFRHYSPFASCRLFRSGSFASSSRVPQG | AVTARRAPVLLSHGATVVSPSGNASSKQRSH | 2314 | 2503 | + | 8 |
| 8 | MSVVAEFANTQLVDVQPGQVGAHAAPTM | ITPLATIATPEAVPIGVAVTSATAIAVNAVTQKVAG | 9355 | 9550 | + | 34 |

### Results for WP_043177785.1 [Streptomyces sp. NRRL F-2890] back to top

previous - next

#### Architecture

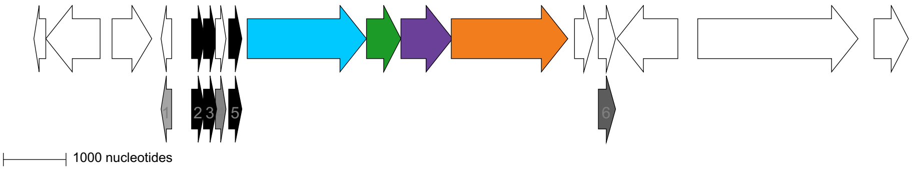

Link to nucleotide sequence

| Accession | start | end | direction | length (aa) | Pfam/HMM | name | description | E-value |
|---|---|---|---|---|---|---|---|---|
| WP_046724823.1 | 139023 | 138828 | - | 64 | PF04149 | DUF397 | Domain of unknown function (DUF397) | 2.30E-22 |
| WP_030733547.1 | 139889 | 139019 | - | 289 | PF13560<br>TIGR03830<br>PF01381<br>TIGR03070<br>PF12844 | HTH_31<br>TIGR03830<br>HTH_3<br>TIGR03070<br>HTH_19 | Helix-turn-helix domain<br>CxxCG_CxxCG_HTH: putative zinc finger/helix-turn-helix protein, YgiT family<br>Helix-turn-helix<br>couple_hipB: transcriptional regulator, y4mF family<br>Helix-turn-helix domain | 3.90E-15<br>4.10E-05<br>1.80E-04<br>2.50E-04<br>4.40E-04 |
| WP_030733548.1 | 140063 | 140714 | + | 216 | PF13581<br>PF02518 | HATPase_c_2<br>HATPase_c | Histidine kinase-like ATPase domain<br>Histidine kinase-, DNA gyrase B-, and HSP90-like ATPase | 3.70E-14<br>5.50E-04 |
| WP_158708039.1 | 141024 | 140850 | - | 57 | NO PFAM MATCH | - | - | - |
| WP_030733549.1 | 141351 | 141546 | + | 64 | LinA | LinA | LinA (linaridin precursor) | 2.00E-13 |
| WP_030733550.1 | 141542 | 141734 | + | 63 | LinA | LinA | LinA (linaridin precursor) | 5.10E-09 |
| WP_158708040.1 | 141730 | 141901 | + | 56 | NO PFAM MATCH | - | - | - |
| WP_030733552.1 | 141927 | 142143 | + | 71 | LinA | LinA | LinA (linaridin precursor) | 4.60E-15 |
| WP_043177785.1 | 142237 | 144145 | + | 635 | LinH<br>LinG<br>LinE<br>PF12697<br>PF00975 | LinH<br>LinG<br>LinE<br>Abhydrolase_6<br>Thioesterase | LinH (Fusion of LinE/LinG)<br>LinG Homolog<br>LinE Homolog<br>Alpha/beta hydrolase family<br>Thioesterase domain | 1.00E-238<br>3.50E-111<br>1.80E-97<br>4.60E-15<br>6.40E-09 |
| WP_043177786.1 | 144141 | 144687 | + | 181 | LinL | LinL | LinL | 3.30E-64 |
| WP_030733555.1 | 144683 | 145511 | + | 275 | PF13649<br>PF08241<br>PF13489<br>PF08242<br>PF13847 | Methyltransf_25<br>Methyltransf_11<br>Methyltransf_23<br>Methyltransf_12<br>Methyltransf_31 | Methyltransferase domain<br>Methyltransferase domain<br>Methyltransferase domain<br>Methyltransferase domain<br>Methyltransferase domain | 2.50E-16<br>1.40E-10<br>8.90E-08<br>1.20E-07<br>8.00E-07 |
| WP_078635559.1 | 145488 | 147354 | + | 621 | TIGR02204<br>TIGR02203<br>TIGR03797<br>TIGR00958<br>TIGR03796 | TIGR02204<br>TIGR02203<br>TIGR03797<br>TIGR00958<br>TIGR03796 | MsbA_rel: ABC transporter, permease/ATP-binding protein<br>MsbA_lipidA: lipid A export permease/ATP-binding protein MsbA<br>NHLM_micro_ABC2: NHLM bacteriocin system ABC transporter, ATP-binding protein<br>3a01208: antigen peptide transporter 2<br>NHLM_micro_ABC1: NHLM bacteriocin system ABC transporter, peptidase/ATP-binding protein | 7.40E-104<br>4.10E-101<br>6.50E-89<br>5.10E-88<br>3.30E-87 |
| WP_030733557.1 | 147454 | 147748 | + | 97 | NO PFAM MATCH | - | - | - |
| WP_030733558.1 | 147826 | 148108 | + | 93 | NO PFAM MATCH | - | - | - |

| | | | | | | | | |
|---|---|---|---|---|---|---|---|---|
| WP_030733560.1 | 149108 | 148124 | - | 327 | PF00196 | GerE | Bacterial regulatory proteins, luxR family | 3.20E-04 |
| WP_030733561.1 | 149429 | 151988 | + | 852 | PF03235<br>PF07510<br>PF01035 | DUF262<br>DUF1524<br>DNA_binding_1 | Protein of unknown function DUF262<br>Protein of unknown function (DUF1524)<br>6-O-methylguanine DNA methyltransferase, DNA binding domain | 5.50E-33<br>5.30E-29<br>8.80E-11 |
| WP_078635561.1 | 152228 | 152789 | + | 186 | PF00072<br>TIGR02154<br>TIGR01818<br>TIGR01387<br>TIGR02875 | Response_reg<br>TIGR02154<br>TIGR01818<br>TIGR01387<br>TIGR02875 | Response regulator receiver domain<br>PhoB: phosphate regulon transcriptional regulatory protein PhoB<br>ntrC: nitrogen regulation protein NR(I)<br>cztR_silR_copR: heavy metal response regulator<br>spore_0_A: sporulation transcription factor Spo0A | 1.80E-20<br>6.00E-18<br>5.20E-11<br>5.90E-10<br>1.10E-09 |

| index | leader | core | start | end | dir | score |
|---|---|---|---|---|---|---|
| 1 | MGDFNGSWAARLMEWSNGTIDEQRARTL | VTEIYEAAAERAVRLHEEEVAQREAEREE | 141024 | 140850 | - | 6 |
| 2 | MAGPSLARQWAHLGPLHENSPGEVGRAASPVL | ATPSAFAAGMGIVVGAYVVGRMIGSETEPVIQ | 141351 | 141546 | + | 19 |
| 3 | MTAADRVPALRDLSPLTENSPGQAPGSATAVL | ATPTPAAMIVIAYAAFEVGRMVCGTTEPVIQ | 141542 | 141734 | + | 31 |
| 4 | MTAADRIPAVLGEIGELNEEDGGRASAP | RAVVLCLHTSAYIAVSVVGDTEPSLPSL | 141730 | 141901 | + | 7 |
| 5 | MSRIAAYLDDSPVLATIPVLDRTGRVSPARPVL | ATPVTIVAFGAGVAAGSAIVAAFEAGRTAGSSTEPIMT | 141927 | 142143 | + | 29 |
| 6 | MTDGAGHGDGEGQGEGLDGEIALTL...RLLSLLPGHWTYRHRVTAARIMLWV | RTPAGAPAVHAALRAALADPALRGWLPDSD | 147826 | 148108 | + | 8 |

### Results for WP_158858022.1 [Streptomyces sp. NRRL B-1347] back to top

previous - next

#### Architecture

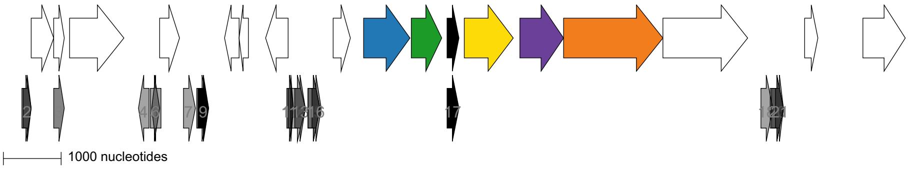

Link to nucleotide sequence

| Accession | start | end | direction | length (aa) | Pfam/HMM | name | description | E-value |
|---|---|---|---|---|---|---|---|---|
| WP_030688453.1 | 8884 | 9265 | + | 126 | NO PFAM MATCH | - | - | - |
| WP_030688455.1 | 9261 | 9456 | + | 64 | TIGR01292<br>TIGR02032<br>TIGR01350<br>TIGR03143<br>PF13738 | TIGR01292<br>TIGR02032<br>TIGR01350<br>TIGR03143<br>Pyr_redox_3 | TRX_reduct: thioredoxin-disulfide reductase<br>GG-red-SF: geranylgeranyl reductase family<br>lipoamide_DH: dihydrolipoyl dehydrogenase<br>AhpF_homolog: putative alkyl hydroperoxide reductase F subunit<br>Pyridine nucleotide-disulphide oxidoreductase | 1.50E-08<br>7.00E-08<br>1.50E-07<br>3.40E-07<br>5.80E-07 |
| WP_030688459.1 | 9540 | 10485 | + | 314 | TIGR02779<br>TIGR02776<br>TIGR00574<br>PF01068<br>TIGR04120 | TIGR02779<br>TIGR02776<br>TIGR00574<br>DNA_ligase_A_M<br>TIGR04120 | NHEJ_ligase_lig: DNA ligase D, ligase domain<br>NHEJ_ligase_prk: DNA ligase D<br>dnl1: DNA ligase I, ATP-dependent (dnl1)<br>ATP dependent DNA ligase domain<br>DNA_lig_bact: DNA ligase, ATP-dependent, PP_1105 family | 5.30E-60<br>7.10E-53<br>1.80E-24<br>8.70E-24<br>5.90E-23 |
| WP_005477366.1 | 11113 | 11441 | + | 109 | INFERRED GENE | - | - | - |
| WP_107059146.1 | 12479 | 12227 | - | 83 | NO PFAM MATCH | - | - | - |
| WP_107059147.1 | 12646 | 12475 | - | 56 | PF08681 | DUF1778 | Protein of unknown function (DUF1778) | 3.00E-04 |
| WP_107059150.1 | 13320 | 12945 | - | 124 | PF12840 | HTH_20 | Helix-turn-helix domain | 1.50E-07 |
| WP_158858020.1 | 14105 | 14408 | + | 100 | PF01527<br>PF13518<br>PF13384<br>PF13551 | HTH_Tnp_1<br>HTH_28<br>HTH_23<br>HTH_29 | Transposase<br>Helix-turn-helix domain<br>Homeodomain-like domain<br>Winged helix-turn helix | 3.30E-06<br>1.20E-04<br>1.50E-04<br>7.60E-04 |
| WP_158858022.1 | 14631 | 15447 | + | 271 | LinE<br>LinH<br>PF12697<br>TIGR02427<br>PF00561 | LinE<br>LinH<br>Abhydrolase_6<br>TIGR02427<br>Abhydrolase_1 | LinE Homolog<br>LinH (Fusion of LinE/LinG)<br>Alpha/beta hydrolase family<br>protocat_pcaD: 3-oxoadipate enol-lactonase<br>alpha/beta hydrolase fold | 6.70E-77<br>7.10E-43<br>7.70E-13<br>1.70E-10<br>4.60E-08 |
| WP_158858024.1 | 15452 | 15992 | + | 179 | LinL | LinL | LinL | 7.20E-51 |
| WP_158858026.1 | 16088 | 16280 | + | 63 | LinA | LinA | LinA (linaridin precursor) | 1.10E-10 |
| WP_078869533.1 | 16382 | 17219 | + | 278 | LinG<br>LinH<br>TIGR04033 | LinG<br>LinH<br>TIGR04033 | LinG Homolog<br>LinH (Fusion of LinE/LinG)<br>export_SdpB: antimicrobial peptide system protein, SdpB family | 2.10E-95<br>3.90E-70<br>9.00E-05 |
| WP_051856534.1 | 17340 | 18105 | + | 254 | PF13649<br>TIGR01983<br>PF13489<br>PF08241<br>PF08242 | Methyltransf_25<br>TIGR01983<br>Methyltransf_23<br>Methyltransf_11<br>Methyltransf_12 | Methyltransferase domain<br>UbiG: 3-demethylubiquinone-9 3-O-methyltransferase<br>Methyltransferase domain<br>Methyltransferase domain<br>Methyltransferase domain | 1.10E-13<br>2.30E-10<br>1.30E-09<br>1.40E-08<br>6.70E-07 |
| WP_030688470.1 | 18101 | 19820 | + | 572 | TIGR02203<br>TIGR02204<br>TIGR00958<br>TIGR03796<br>TIGR01846 | TIGR02203<br>TIGR02204<br>TIGR00958<br>TIGR03796<br>TIGR01846 | MsbA_lipidA: lipid A export permease/ATP-binding protein MsbA<br>MsbA_rel: ABC transporter, permease/ATP-binding protein<br>3a01208: antigen peptide transporter 2<br>NHLM_micro_ABC1: NHLM bacteriocin system ABC transporter, peptidase/ATP-binding protein<br>type_I_sec_HlyB: type I secretion system ATPase | 1.70E-104<br>9.80E-102<br>7.50E-90<br>1.60E-83<br>1.40E-81 |

| | | | | | | | | |
|---|---|---|---|---|---|---|---|---|
| WP_030688472.1 | 19816 | 21280 | + | 487 | TIGR02734<br>PF13450<br>TIGR03467<br>TIGR01317<br>TIGR00292 | TIGR02734<br>NAD_binding_8<br>TIGR03467<br>TIGR01317<br>TIGR00292 | crtI_fam: phytoene desaturase<br>NAD(P)-binding Rossmann-like domain<br>HpnE: squalene-associated FAD-dependent desaturase<br>GOGAT_sm_gam: glutamate synthase, NADH/NADPH, small subunit<br>TIGR00292: thiazole biosynthesis enzyme | 5.60E-12<br>2.80E-10<br>3.30E-09<br>8.90E-08<br>9.60E-08 |
| NP_639593.1 | 22266 | 22506 | + | 80 | INFERRED GENE | - | - | - |
| WP_014386339.1 | 23273 | 24021 | + | 249 | INFERRED GENE | - | - | - |

| index | leader | core | start | end | dir | score |
|---|---|---|---|---|---|---|
| 1 | MRAYLAQLAIEEENQVKLNK | ATLVFREVTGKPGIAEAFDAAFPEDAPGPRGTAGRAA | 8708 | 8882 | + | 7 |
| 2 | LAIEEENQVKLNK | ATLVFREVTGKPGIAEAFDAAFPEDAPGPRGTAGRAA | 8729 | 8882 | + | 7 |
| 3 | MTGFDLIVVGAGPAGVSAAVMASSLRM | RTVVVEAEQVGAKLRSIDALQKRLASTSPLGGQTHRR | 9261 | 9456 | + | 7 |
| 4 | MIREQDSRDADEGGRARDELQDHAGEGLA | FRRPTREPTRAARSQVHPLSRDRRGEPTR | 10908 | 10731 | - | 6 |
| 5 | LCGRAFMVPALMAEQARAALKAWNG | YAPTPAEEQGLDSELDRRLAQAAVR | 10952 | 11105 | + | 6 |
| 6 | MRTTSLTARPPGPGGGPARCPGPAP | QQGWARSRSTPSARPAPARPSARAP | 11121 | 10968 | - | 6 |
| 7 | MAGPRLERVGRWRQRAGPVAGHHRILRERFPG | SACVHRDQGPRVPRVGAHSFGGAPVNRGTSRL | 11520 | 11715 | + | 6 |
| 8 | MRRTASIVMTCTALALTVIG | ATPSIADPGPISEADCRAKNGVVDDIQGTYYCVAVHGVIEGAE | 11753 | 11945 | + | 7 |
| 9 | MTCTALALTVIG | ATPSIADPGPISEADCRAKNGVVDDIQGTYYCVAVHGVIEGAE | 11777 | 11945 | + | 18 |
| 10 | VQAGMLLSCCCSAERCCSS | SSTRWVNVGSGAASVTSSS | 13306 | 13423 | + | 7 |
| 11 | MLLSCCCSAERCCSSSS | TRWVNVGSGAASVTSSS | 13318 | 13423 | + | 7 |
| 12 | MSVLERLQSHPHRSAGVDHGFYSRHEGRRAVLLGPLRGEAGEPWQEPVE | ALPEWGGRDTVLGRYVSVGGGVRPVDHASVR | 13382 | 13625 | + | 6 |
| 13 | VDHGFYSRHEGRRAVLLGPLRGEAGEPWQEPVE | ALPEWGGRDTVLGRYVSVGGGVRPVDHASVR | 13430 | 13625 | + | 7 |
| 14 | VVVGELEQSGCRSRGAGLPRSTWPFGYRRRLA | AGSGMESALRSGARGKEVSLRALQAGLLAAIVA | 13670 | 13868 | + | 6 |
| 15 | VVGELEQSGCRSRGAGLPRSTWPFGYRRRLAA | GSGMESALRSGARGKEVSLRALQAGLLAAIVA | 13673 | 13868 | + | 6 |
| 16 | VAFRLSAAIGCWQWHGVGVAV | GCPRQRGESASASGRSSGCHRGLAHGGWAAALT | 13735 | 13900 | + | 7 |
| 17 | MSNVADYINSNDKITQGLQVALRPETNGPVM | ITPIAAVAAGAGVVATARVGQHVARYVAHRDH | 16088 | 16280 | + | 30 |
| 18 | LWTRFRFDRTPGPVRKHAEICGRSRVEVKMNQAY | QFITSGYVAVCEGTENPDNCGPWENITSWPVNHS | 21513 | 21720 | + | 6 |
| 19 | VGEHHLVAGQPQLSGRPSPASGPLDLCCRTVWCGRGARV | NTAQGDDDGPSAVSIPQTVRPASFAPVFSWAPGELR | 21680 | 21908 | + | 6 |
| 20 | VAGQPQLSGRPSPASGPLDLCCRTVWCGRGARV | NTAQGDDDGPSAVSIPQTVRPASFAPVFSWAPGELR | 21698 | 21908 | + | 7 |
| 21 | VWCGRGARVNTAQGDDDGPSAV | SIPQTVRPASFAPVFSWAPGELR | 21770 | 21908 | + | 6 |

### Results for WP_158716051.1 [Streptomyces sp. NRRL B-24720] back to top

previous - next

#### Architecture

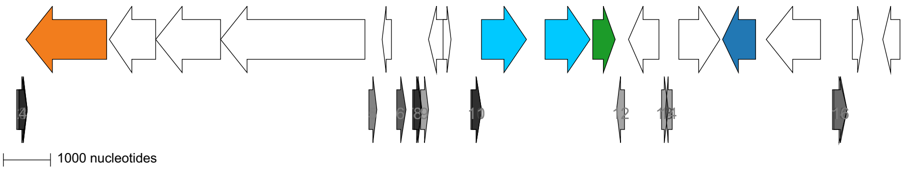

Link to nucleotide sequence

| Accession | start | end | direction | length (aa) | Pfam/HMM | name | description | E-value |
|---|---|---|---|---|---|---|---|---|
| WP_051879144.1 | 17666 | 15971 | - | 564 | TIGR02204<br>TIGR02203<br>TIGR00958<br>TIGR01846<br>TIGR03796 | TIGR02204<br>TIGR02203<br>TIGR00958<br>TIGR01846<br>TIGR03796 | MsbA_rel: ABC transporter, permease/ATP-binding protein<br>MsbA_lipidA: lipid A export permease/ATP-binding protein MsbA<br>3a01208: antigen peptide transporter 2<br>type_I_sec_HlyB: type I secretion system ATPase<br>NHLM_micro_ABC1: NHLM bacteriocin system ABC transporter, peptidase/ATP-binding protein | 4.10E-114<br>8.40E-113<br>6.30E-94<br>9.00E-93<br>7.40E-91 |
| WP_051879128.1 | 18709 | 17737 | - | 323 | TIGR03891<br>PF14028 | TIGR03891<br>Lant_dehydr_C | thiopep_ocin: thiopeptide-type bacteriocin biosynthesis domain<br>Lantibiotic biosynthesis dehydratase C-term | 1.60E-25<br>1.50E-13 |
| WP_078879182.1 | 20100 | 18732 | - | 455 | PF05147<br>TIGR03897 | LANC_like<br>TIGR03897 | Lanthionine synthetase C-like protein<br>lanti_2_LanM: type 2 lantibiotic biosynthesis protein LanM | 1.40E-53<br>2.10E-13 |
| WP_051879132.1 | 23147 | 20096 | - | 1016 | PF04738<br>TIGR03891<br>PF14028 | Lant_dehydr_N<br>TIGR03891<br>Lant_dehydr_C | Lantibiotic dehydratase, N terminus<br>thiopep_ocin: thiopeptide-type bacteriocin biosynthesis domain<br>Lantibiotic biosynthesis dehydratase C-term | 3.70E-151<br>1.00E-36<br>9.80E-22 |
| WP_030931945.1 | 23719 | 23521 | - | 65 | NO PFAM MATCH | - | - | - |
| WP_158716049.1 | 24863 | 24515 | - | 115 | PF13546<br>PF01609 | DDE_5<br>DDE_Tnp_1 | DDE superfamily endonuclease<br>Transposase DDE domain | 1.10E-16<br>1.40E-04 |
| IH49_RS49280 | 24810 | 24990 | + | 60 | INFERRED GENE | - | - | - |
| WP_158716050.1 | 25637 | 26585 | + | 315 | LinH<br>LinG | LinH<br>LinG | LinH (Fusion of LinE/LinG)<br>LinG Homolog | 4.30E-87<br>1.90E-81 |

| | | | | | | | | |
|---|---|---|---|---|---|---|---|---|
| WP_158716051.1 | 26971 | 27934 | + | 320 | LinH<br>LinE<br>PF12697<br>TIGR03611<br>PF12146 | LinH<br>LinE<br>Abhydrolase_6<br>TIGR03611<br>Hydrolase_4 | LinH (Fusion of LinE/LinG)<br>LinE Homolog<br>Alpha/beta hydrolase family<br>RutD: pyrimidine utilization protein D<br>Serine aminopeptidase, S33 | 1.90E-62<br>1.90E-61<br>2.60E-10<br>3.10E-09<br>3.00E-08 |
| WP_158716052.1 | 27999 | 28476 | + | 158 | LinL | LinL | LinL | 1.40E-47 |
| WP_051879136.1 | 29389 | 28762 | - | 208 | PF01548 | DEDD_Tnp_IS110 | Transposase | 5.60E-13 |
| WP_086558180.1 | 29818 | 30694 | + | 292 | INFERRED GENE | - | - | - |
| WP_030931961.1 | 31459 | 30745 | - | 237 | PF12697<br>LinE<br>LinH | Abhydrolase_6<br>LinE<br>LinH | Alpha/beta hydrolase family<br>LinE Homolog<br>LinH (Fusion of LinE/LinG) | 2.90E-14<br>2.80E-12<br>8.60E-11 |
| WP_030931964.1 | 32840 | 31667 | - | 390 | NO PFAM MATCH | - | - | - |
| WP_013002890.1 | 33503 | 33731 | + | 76 | INFERRED GENE | - | - | - |
| WP_158716053.1 | 34527 | 34161 | - | 121 | NO PFAM MATCH | - | - | - |

| index | leader | core | start | end | dir | score |
|---|---|---|---|---|---|---|
| 1 | MKFLFSRHPSTHMRLTKSAACCLVASS | GTAVHRRTSAFCQPAVMWSMSDAWVGRRINRSVVSTIGLMD | 15775 | 15982 | + | 6 |
| 2 | LFSRHPSTHMRLTKSAACCLVASS | GTAVHRRTSAFCQPAVMWSMSDAWVGRRINRSVVSTIGLMD | 15784 | 15982 | + | 6 |
| 3 | MRLTKSAACCLVASS | GTAVHRRTSAFCQPAVMWSMSDAWVGRRINRSVVSTIGLMD | 15811 | 15982 | + | 6 |
| 4 | LTKSAACCLVASS | GTAVHRRTSAFCQPAVMWSMSDAWVGRRINRSVVSTIGLMD | 15817 | 15982 | + | 6 |
| 5 | LPQEHRTGPSSTTQPTAVNACYFVLG | RTTVVGDYDRAACSGCWRRQSGARTRKL | 23252 | 23417 | + | 7 |
| 6 | VSLPARTTLVAVCSGWKVSGEDPLGLGRFRPS | HCAVSVEELLALALANYCPVLAFTAEIELVVP | 23831 | 24026 | + | 8 |
| 7 | VYRKCQVRKVRSTSTIQNARMVML | ETPRLVLRRWRADDVVPMADVNADPAVMRWILRATRPGA | 24164 | 24356 | + | 8 |
| 8 | VRSTSTIQNARMVML | ETPRLVLRRWRADDVVPMADVNADPAVMRWILRATRPGA | 24191 | 24356 | + | 7 |
| 9 | VTWAGHRPSCRTSRWRRSGPGGGPR | LRPTAPAAPGSASASVRGSLPGRSPR | 24352 | 24508 | + | 6 |
| 10 | LELFMDRSRIIAIASGVIAVG | LTVTPAVAATSNEESAPTTPYVSSAPTSNSAADEILGTNEYPCCV | 25423 | 25624 | + | 8 |
| 11 | MDRSRIIAIASGVIAVG | LTVTPAVAATSNEESAPTTPYVSSAPTSNSAADEILGTNEYPCCV | 25435 | 25624 | + | 7 |
| 12 | LGDTLPPLLIVGSLVRGDKVAVF | ATPYTPQMALLFALINRSVTACHAKVAASRTSLCLTSSF | 28681 | 28492 | - | 6 |
| 13 | LQEPLQFVLVGVSTCFGGLEAGGGSPA | CLRLAHPGESRFVSDTKPRRRKDDDHA | 29447 | 29612 | + | 6 |
| 14 | VSDAWGGRLVYGAGD | GTATPGICVSGVVVVLAAARFGVGDEPALAGMRQA | 29684 | 29531 | - | 6 |
| 15 | LPPAPQASPSVRISEAHSDARDETA...LKKAAVGAGSTPPKTSVNCVSRLGR | VTLRGRGGTVVALIQLASMPMASAEGSQVWGRR | 33097 | 33388 | + | 6 |
| 16 | VRISEAHSDARDETADFAAEPSANL...LKKAAVGAGSTPPKTSVNCVSRLGR | VTLRGRGGTVVALIQLASMPMASAEGSQVWGRR | 33127 | 33388 | + | 6 |

### Results for WP_158708080.1 [Streptomyces sp. NRRL F-2890] back to top

previous - next

#### Architecture

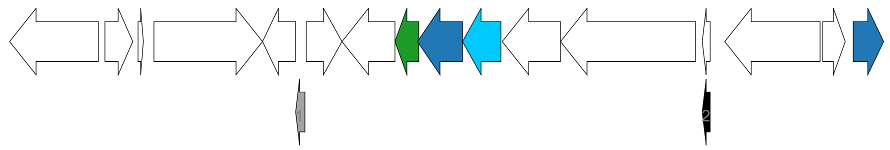

1000 nucleotides

Link to nucleotide sequence

| Accession | start | end | direction | length (aa) | Pfam/HMM | name | description | E-value |
|---|---|---|---|---|---|---|---|---|
| WP_052385436.1 | 283018 | 281008 | - | 669 | PF00496<br>TIGR02294<br>TIGR04028 | SBP_bac_5<br>TIGR02294<br>TIGR04028 | Bacterial extracellular solute-binding proteins, family 5 Middle<br>nickel_nikA: nickel ABC transporter, nickel/metallophore periplasmic binding protein<br>SBP_KPN_01854: ABC transporter substrate binding protein, KPN_01854 family | 3.00E-44<br>2.10E-17<br>3.10E-17 |
| WP_052385437.1 | 283180 | 283807 | + | 208 | PF04536 | TPM_phosphatase | TPM domain | 2.70E-12 |
| WP_014176736.1 | 283915 | 284038 | + | 41 | INFERRED GENE | - | - | - |
| WP_030737455.1 | 284277 | 286758 | + | 826 | PF07228<br>TIGR00229<br>PF08448<br>PF00989<br>PF13426 | SpoIIE<br>TIGR00229<br>PAS_4<br>PAS<br>PAS_9 | Stage II sporulation protein E (SpoIIE)<br>sensory_box: PAS domain S-box protein<br>PAS fold<br>PAS fold<br>PAS domain | 3.00E-51<br>8.10E-27<br>6.20E-25<br>8.50E-17<br>3.90E-14 |
| WP_030737458.1 | 287517 | 286764 | - | 250 | PF00392<br>PF07702<br>TIGR02018<br>TIGR02325<br>TIGR02404 | GntR<br>UTRA<br>TIGR02018<br>TIGR02325<br>TIGR02404 | Bacterial regulatory proteins, gntR family<br>UTRA domain<br>his_ut_repres: histidine utilization repressor<br>C_P_lyase_phnF: phosphonate metabolism transcriptional regulator PhnF<br>trehalos_R_Bsub: trehalose operon repressor | 3.00E-16<br>2.30E-13<br>2.60E-13<br>1.50E-12<br>1.00E-06 |
| WP_052385438.1 | 287742 | 288573 | + | 276 | PF00583<br>TIGR01575<br>PF13508<br>TIGR03448<br>PF13673 | Acetyltransf_1<br>TIGR01575<br>Acetyltransf_7<br>TIGR03448<br>Acetyltransf_10 | Acetyltransferase (GNAT) family<br>rimI: ribosomal-protein-alanine acetyltransferase<br>Acetyltransferase (GNAT) domain<br>mycothiol_MshD: mycothiol synthase<br>Acetyltransferase (GNAT) domain | 4.80E-14<br>7.30E-09<br>1.60E-06<br>9.60E-06<br>1.50E-05 |

| | | | | | | | | |
|---|---|---|---|---|---|---|---|---|
| WP_030737468.1 | 289774 | 288562 | - | 403 | TIGR04458<br>PF00067 | TIGR04458<br>p450 | CYP450_TxtE: 4-nitrotryptophan synthase<br>Cytochrome P450 | 2.10E-31<br>1.70E-27 |
| WP_030737472.1 | 290322 | 289770 | - | 183 | LinL | LinL | LinL | 1.10E-55 |
| WP_158708080.1 | 291316 | 290296 | - | 339 | LinE<br>LinH<br>PF12697<br>TIGR02427<br>PF12146 | LinE<br>LinH<br>Abhydrolase_6<br>TIGR02427<br>Hydrolase_4 | LinE Homolog<br>LinH (Fusion of LinE/LinG)<br>Alpha/beta hydrolase family<br>protocat_pcaD: 3-oxoadipate enol-lactonase<br>Serine aminopeptidase, S33 | 2.30E-90<br>1.70E-79<br>8.00E-17<br>7.00E-12<br>3.80E-11 |
| WP_030737478.1 | 292178 | 291299 | - | 292 | LinH<br>LinG | LinH<br>LinG | LinH (Fusion of LinE/LinG)<br>LinG Homolog | 6.10E-108<br>3.90E-101 |
| WP_052385439.1 | 293527 | 292219 | - | 435 | PF05147<br>TIGR03897 | LANC_like<br>TIGR03897 | Lanthionine synthetase C-like protein<br>lanti_2_LanM: type 2 lantibiotic biosynthesis protein LanM | 2.70E-45<br>5.10E-17 |
| WP_030737483.1 | 296613 | 293523 | - | 1029 | PF04738<br>TIGR03891<br>PF14028 | Lant_dehydr_N<br>TIGR03891<br>Lant_dehydr_C | Lantibiotic dehydratase, N terminus<br>thiopep_ocin: thiopeptide-type bacteriocin biosynthesis domain<br>Lantibiotic biosynthesis dehydratase C-term | 1.50E-142<br>4.10E-42<br>6.50E-25 |
| WP_030737486.1 | 296931 | 296748 | - | 60 | NO PFAM MATCH | - | - | - |
| WP_030737488.1 | 299437 | 297277 | - | 719 | TIGR00833<br>PF03176<br>TIGR00921<br>TIGR03480<br>PF00873 | TIGR00833<br>MMPL<br>TIGR00921<br>TIGR03480<br>ACR_tran | actII: transport protein<br>MMPL family<br>2A067: efflux transporter, putative, hydrophobe/amphiphile efflux-3 (HAE3) family<br>HpnN: hopanoid biosynthesis associated RND transporter like protein HpnN<br>AcrB/AcrD/AcrF family | 7.70E-123<br>5.10E-111<br>3.10E-32<br>5.20E-21<br>2.50E-04 |
| WP_030737490.1 | 299510 | 300017 | + | 168 | PF12802<br>PF01047<br>PF13463<br>TIGR02337<br>TIGR04472 | MarR_2<br>MarR<br>HTH_27<br>TIGR02337<br>TIGR04472 | MarR family<br>MarR family<br>Winged helix DNA-binding domain<br>HpaR: homoprotocatechuate degradation operon regulator, HpaR<br>reg_rSAM_mob: mobile rSAM pair MarR family regulator | 1.00E-11<br>9.40E-10<br>7.50E-07<br>5.50E-05<br>5.70E-05 |
| WP_052385440.1 | 300183 | 300891 | + | 235 | PF12146<br>PF12697<br>LinE<br>LinH<br>TIGR03695 | Hydrolase_4<br>Abhydrolase_6<br>LinE<br>LinH<br>TIGR03695 | Serine aminopeptidase, S33<br>Alpha/beta hydrolase family<br>LinE Homolog<br>LinH (Fusion of LinE/LinG)<br>menH_SHCHC: 2-succinyl-6-hydroxy-2,4-cyclohexadiene-1-carboxylate synthase | 1.70E-12<br>3.80E-10<br>1.20E-08<br>2.90E-07<br>6.10E-04 |

| index | leader | core | start | end | dir | score |
|---|---|---|---|---|---|---|
| 1 | VCHGARQGGMGVNGLRRNGTSAIRAGHGRVITSG | DSVASSCGRQPSVVKLLLFVKSADGSTPGRAYAVR | 287713 | 287503 | - | 6 |
| 2 | MGADLITAAANEELSDLDLDLTFAPNDENT | HVMLACAVDSMGNNVNTSRGIIPQAPVCCA | 296931 | 296748 | - | 15 |

### Results for WP_030999015.1 [Streptomyces sp. NRRL F-5630] back to top

previous - next

#### Architecture

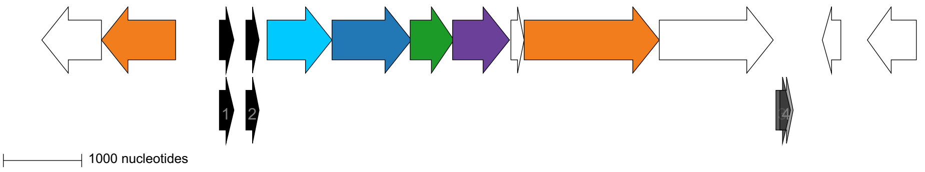

Link to nucleotide sequence

| Accession | start | end | direction | length (aa) | Pfam/HMM | name | description | E-value |
|---|---|---|---|---|---|---|---|---|
| WP_051703644.1 | 878 | 113 | - | 254 | PF01061<br>PF12698 | ABC2_membrane<br>ABC2_membrane_3 | ABC-2 type transporter<br>ABC-2 family transporter protein | 2.90E-15<br>2.50E-08 |
| WP_030999010.1 | 1822 | 874 | - | 315 | TIGR01188<br>TIGR03864<br>TIGR01288<br>TIGR00968<br>TIGR04406 | TIGR01188<br>TIGR03864<br>TIGR01288<br>TIGR00968<br>TIGR04406 | drrA: daunorubicin resistance ABC transporter, ATP-binding protein<br>PQQ_ABC_ATP: ABC transporter, ATP-binding subunit, PQQ-dependent alcohol dehydrogenase system<br>nodI: nodulation ABC transporter NodI<br>3a0106s01: sulfate ABC transporter, ATP-binding protein<br>LPS_export_lptB: LPS export ABC transporter ATP-binding protein | 5.30E-62<br>1.60E-43<br>4.20E-42<br>2.30E-34<br>2.50E-34 |
| WP_051703645.1 | 2372 | 2573 | + | 66 | LinA | LinA | LinA (linaridin precursor) | 1.00E-11 |
| WP_037827430.1 | 2714 | 2894 | + | 59 | LinA | LinA | LinA (linaridin precursor) | 4.20E-10 |
| WP_158716646.1 | 2983 | 3820 | + | 278 | LinH<br>LinG | LinH<br>LinG | LinH (Fusion of LinE/LinG)<br>LinG Homolog | 1.90E-104<br>6.30E-103 |
| WP_030999015.1 | 3816 | 4824 | + | 335 | LinE<br>LinH<br>PF12697<br>TIGR02427<br>TIGR03611 | LinE<br>LinH<br>Abhydrolase_6<br>TIGR02427<br>TIGR03611 | LinE Homolog<br>LinH (Fusion of LinE/LinG)<br>Alpha/beta hydrolase family<br>protocat_pcaD: 3-oxoadipate enol-lactonase<br>RutD: pyrimidine utilization protein D | 8.80E-122<br>7.90E-77<br>5.30E-12<br>2.10E-09<br>2.10E-06 |
| WP_158716647.1 | 4820 | 5366 | + | 181 | LinL | LinL | LinL | 8.10E-63 |
| WP_030999019.1 | 5362 | 6082 | + | 239 | PF13649<br>PF13489<br>PF08241<br>PF08242<br>TIGR01934 | Methyltransf_25<br>Methyltransf_23<br>Methyltransf_11<br>Methyltransf_12<br>TIGR01934 | Methyltransferase domain<br>Methyltransferase domain<br>Methyltransferase domain<br>Methyltransferase domain<br>MenG_MenH_UbiE: ubiquinone/menaquinone biosynthesis methyltransferase | 6.80E-12<br>1.50E-09<br>4.80E-09<br>7.20E-06<br>1.50E-05 |
| WP_030999021.1 | 6097 | 6277 | + | 59 | NO PFAM MATCH | - | - | - |

| | | | | | | | | |
|---|---|---|---|---|---|---|---|---|
| WP_030999023.1 | 6273 | 7992 | + | 572 | TIGR02204<br>TIGR02203<br>TIGR00958<br>TIGR01846<br>TIGR03796 | TIGR02204<br>TIGR02203<br>TIGR00958<br>TIGR01846<br>TIGR03796 | MsbA_rel: ABC transporter, permease/ATP-binding protein<br>MsbA_lipidA: lipid A export permease/ATP-binding protein MsbA<br>3a01208: antigen peptide transporter 2<br>type_I_sec_HlyB: type I secretion system ATPase<br>NHLM_micro_ABC1: NHLM bacteriocin system ABC transporter, peptidase/ATP-binding protein | 8.00E-111<br>1.30E-101<br>1.30E-88<br>2.10E-87<br>5.70E-86 |
| WP_030999025.1 | 7988 | 9446 | + | 485 | PF13450<br>TIGR03467<br>TIGR02734<br>TIGR01317<br>TIGR00292 | NAD_binding_8<br>TIGR03467<br>TIGR02734<br>TIGR01317<br>TIGR00292 | NAD(P)-binding Rossmann-like domain<br>HpnE: squalene-associated FAD-dependent desaturase<br>crtI_fam: phytoene desaturase<br>GOGAT_sm_gam: glutamate synthase, NADH/NADPH, small subunit<br>TIGR00292: thiazole biosynthesis enzyme | 9.50E-11<br>4.60E-10<br>5.20E-09<br>1.30E-07<br>6.30E-07 |
| WP_030999027.1 | 10306 | 10075 | - | 76 | NO PFAM MATCH | - | - | - |
| WP_078655237.1 | 11277 | 10647 | - | 209 | PF16859<br>PF00440 | TetR_C_11<br>TetR_N | Tetracyclin repressor-like, C-terminal domain<br>Bacterial regulatory proteins, tetR family | 1.50E-27<br>1.60E-10 |

| index | leader | core | start | end | dir | score |
|---|---|---|---|---|---|---|
| 1 | MLTQKVNANPSLQGALEAGAPMGGAAPVM | ATPALIATPVIAKAAGAVVGAGAVTGAAYAAYKVATG | 2372 | 2573 | + | 31 |
| 2 | MSVLLDKVNAGVGQASDVQIAPVM | ATPAAFVAGVVAGGKVCAVLTAAAGVGAAVKSAQK | 2714 | 2894 | + | 32 |
| 3 | VIRTATAPTADAEAGPECAEAGPPASAGSPAHHDPVG | RLPARAPERPRTARQGGATRRGA | 9484 | 9667 | + | 8 |
| 4 | VRGGGTARVRRITGTPRPGR | PTARKGTGTTADRAPRRSNPSWRVVRSQPTQQKE | 9534 | 9699 | + | 6 |

### Results for WP_037817104.1 [Streptomyces sp. NRRL F-5527] back to top

previous - next

#### Architecture

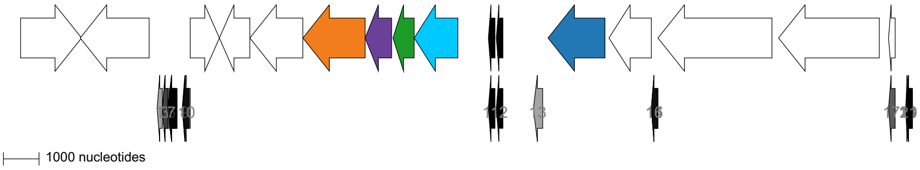

Link to nucleotide sequence

| Accession | start | end | direction | length (aa) | Pfam/HMM | name | description | E-value |
|---|---|---|---|---|---|---|---|---|
| WP_031020078.1 | 79834 | 81529 | + | 564 | PF00476<br>TIGR00593 | DNA_pol_A<br>TIGR00593 | DNA polymerase family A<br>pola: DNA polymerase I | 9.90E-30<br>7.20E-19 |
| WP_031062700.1 | 83448 | 81510 | - | 645 | PF03704<br>PF00486<br>PF13191<br>PF00931<br>TIGR02928 | BTAD<br>Trans_reg_C<br>AAA_16<br>NB-ARC<br>TIGR02928 | Bacterial transcriptional activator domain<br>Transcriptional regulatory protein, C terminal<br>AAA ATPase domain<br>NB-ARC domain<br>TIGR02928: orc1/cdc6 family replication initiation protein | 6.60E-41<br>1.30E-08<br>1.40E-08<br>1.10E-06<br>3.60E-04 |
| WP_006132587.1 | 84573 | 85413 | + | 279 | PF01551 | Peptidase_M23 | Peptidase family M23 | 3.30E-17 |
| WP_031062701.1 | 86249 | 85397 | - | 283 | PF12680 | SnoaL_2 | SnoaL-like domain | 4.70E-05 |
| WP_031020075.1 | 87751 | 86245 | - | 501 | TIGR02734<br>TIGR00292<br>TIGR03467<br>TIGR01317<br>TIGR02733 | TIGR02734<br>TIGR00292<br>TIGR03467<br>TIGR01317<br>TIGR02733 | crtI_fam: phytoene desaturase<br>TIGR00292: thiazole biosynthesis enzyme<br>HpnE: squalene-associated FAD-dependent desaturase<br>GOGAT_sm_gam: glutamate synthase, NADH/NADPH, small subunit<br>desat_CrtD: C-3',4' desaturase CrtD | 1.30E-08<br>2.50E-08<br>9.20E-08<br>4.10E-07<br>6.80E-07 |
| WP_031062704.1 | 89484 | 87747 | - | 578 | TIGR02204<br>TIGR02203<br>TIGR03796<br>TIGR00958<br>TIGR01846 | TIGR02204<br>TIGR02203<br>TIGR03796<br>TIGR00958<br>TIGR01846 | MsbA_rel: ABC transporter, permease/ATP-binding protein<br>MsbA_lipidA: lipid A export permease/ATP-binding protein MsbA<br>NHLM_micro_ABC1: NHLM bacteriocin system ABC transporter, peptidase/ATP-binding protein<br>3a01208: antigen peptide transporter 2<br>type_I_sec_HlyB: type I secretion system ATPase | 1.40E-100<br>2.00E-97<br>3.60E-89<br>9.80E-85<br>1.40E-82 |
| WP_006132583.1 | 90236 | 89480 | - | 251 | PF13649<br>PF08241<br>PF13489<br>PF13847<br>PF08242 | Methyltransf_25<br>Methyltransf_11<br>Methyltransf_23<br>Methyltransf_31<br>Methyltransf_12 | Methyltransferase domain<br>Methyltransferase domain<br>Methyltransferase domain<br>Methyltransferase domain<br>Methyltransferase domain | 2.70E-13<br>1.50E-08<br>1.70E-08<br>4.80E-07<br>5.00E-07 |
| WP_006132582.1 | 90869 | 90290 | - | 192 | LinL | LinL | LinL | 8.30E-66 |
| WP_037817104.1 | 92095 | 90865 | - | 409 | LinH<br>LinE<br>LinG<br>PF12697<br>PF00561 | LinH<br>LinE<br>LinG<br>Abhydrolase_6<br>Abhydrolase_1 | LinH (Fusion of LinE/LinG)<br>LinE Homolog<br>LinG Homolog<br>Alpha/beta hydrolase family<br>alpha/beta hydrolase fold | 1.10E-149<br>9.10E-100<br>3.90E-29<br>2.20E-15<br>4.50E-12 |
| WP_006132580.1 | 93124 | 92932 | - | 63 | LinA | LinA | LinA (linaridin precursor) | 1.70E-11 |
| WP_006132579.1 | 93354 | 93159 | - | 64 | LinA | LinA | LinA (linaridin precursor) | 1.10E-17 |
| WP_006132578.1 | 96226 | 94636 | - | 529 | PF00561<br>PF08386<br>TIGR01249<br>LinE | Abhydrolase_1<br>Abhydrolase_4<br>TIGR01249<br>LinE | alpha/beta hydrolase fold<br>TAP-like protein<br>pro_imino_pep_1: prolyl aminopeptidase<br>LinE Homolog | 3.40E-36<br>1.30E-31<br>2.60E-06<br>9.80E-06 |
| WP_006132577.1 | 97523 | 96317 | - | 401 | TIGR00678<br>PF13177<br>TIGR02397<br>PF00004 | TIGR00678<br>DNA_pol3_delta2<br>TIGR02397<br>AAA | holB: DNA polymerase III, delta' subunit<br>DNA polymerase III, delta subunit<br>dnaX_nterm: DNA polymerase III, subunit gamma and tau<br>ATPase family associated with various cellular activities (AAA) | 1.30E-54<br>9.20E-33<br>1.70E-29<br>3.60E-04 |

| | | | | | | | | |
|---|---|---|---|---|---|---|---|---|
| WP_031020071.1 | 100897 | 97702 | - | 1064 | TIGR00041<br>PF02223<br>TIGR00900<br>PF07690<br>TIGR00895 | TIGR00041<br>Thymidylate_kin<br>TIGR00900<br>MFS_1<br>TIGR00895 | DTMP_kinase: dTMP kinase<br>Thymidylate kinase<br>2A0121: H+ Antiporter protein<br>Major Facilitator Superfamily<br>2A0115: MFS transporter, aromatic acid:H+ symporter (AAHS) family | 1.30E-51<br>1.20E-46<br>2.70E-10<br>5.30E-08<br>4.20E-04 |
| WP_031062705.1 | 103907 | 101072 | - | 944 | TIGR01051<br>PF01131<br>TIGR01054<br>TIGR01057<br>PF13368 | TIGR01051<br>Topoisom_bac<br>TIGR01054<br>TIGR01057<br>Toprim_C_rpt | topA_bact: DNA topoisomerase I<br>DNA topoisomerase<br>rgy: reverse gyrase<br>topA_arch: DNA topoisomerase I<br>Topoisomerase C-terminal repeat | 3.90E-246<br>2.30E-131<br>2.00E-86<br>4.70E-72<br>9.20E-65 |
| WP_006132574.1 | 104350 | 104152 | - | 65 | NO PFAM MATCH | - | - | - |

| index | leader | core | start | end | dir | score |
|---|---|---|---|---|---|---|
| 1 | VGFSGIDCLTSGRHTVEQDQIGRR | GIATGRTCLRLRGLATTEQPVRAG | 83818 | 83671 | - | 6 |
| 2 | VCLLRRWKKSEGQFYAGVGGMNKSESN | ERSGVGLFQVNSSRITSLIPSMSGVFRH | 83966 | 83798 | - | 7 |
| 3 | VHDCVCLLRRWKKSEGQFYAGVGGMNKSE | SNERSGVGLFQVNSSRITSLIPSMSGVFRH | 83978 | 83798 | - | 7 |
| 4 | MPDAAAEPCAADHSTGAAVEGAGPAAAECAAD | DKAATVKAIVARVRFGDAIRMRVTRLPGRRCGA | 84168 | 83970 | - | 9 |
| 5 | MPSCSVHRMPDAAAEPCAADHSTGAAVEGAGPAAAE | CAADDKAATVKAIVARVRFGDAIRMRVTRLPGRRCGA | 84192 | 83970 | - | 9 |
| 6 | MTRADRMPSCSVHRMPDAAAEPCAADHSTGAAVEGAGPA | AAECAADDKAATVKAIVARVRFGDAIRMRVTRLPGRRCGA | 84210 | 83970 | - | 9 |
| 7 | VPVFRSSPSMTRADRMPSCSVHRMPDAAAEPCAADHSTGAAVEG | AGPAAAECAADDKAATVKAIVARVRFGDAIRMRVTRLPGRRCGA | 84237 | 83970 | - | 8 |
| 8 | MGAAFQISGPPSHPRFQREDDCVWEAALDGPLIGQVVREMAGPQSVVS | SAPRHTFSWPVVVGR | 84543 | 84351 | - | 8 |
| 9 | VGGSQMGAAFQISGPPSHPRFQRED...WEAALDGPLIGQVVREMAGPQSVVS | SAPRHTFSWPVVVGR | 84558 | 84351 | - | 8 |
| 10 | VGGLSVGGSQMGAAFQISGPPSHPR...WEAALDGPLIGQVVREMAGPQSVVS | SAPRHTFSWPVVVGR | 84573 | 84351 | - | 7 |
| 11 | MSRIVDSVVAVTPTDLTDDSQGEFGAMAPVQ | ATPTLVAFGVGFAGGAGAAWVTVQAYEAGAND | 93124 | 92932 | - | 31 |
| 12 | MEQKIAAFMNEQDLGSLEQGYRDDAPMPVL | ATPAAAVATCWYVGGAAAGAAAVVGAYVTGRVVG | 93354 | 93159 | - | 30 |
| 13 | MRRSRVRIPKAAPLEPQARSFCPWGFLLFEARR | DTPSEPPAAGGRLCLGPLPAPLGRR...GVVSARLPAGESGGIPVRESLVMLG | 94479 | 94224 | - | 6 |
| 14 | VPAGTPGTAVFGVRGQACRCGPAQW | IRRGTDATDTTDATDATDGYDGRAG | 97679 | 97526 | - | 8 |
| 15 | VPGVPAGTPGTAVFGVRGQACRCGPA | QWIRRGTDATDTTDATDATDGYDGRAG | 97688 | 97526 | - | 7 |
| 16 | VAPALRRGCTVPGVPA | GTPGTAVFGVRGQACRCGPAQWIRRGTDATDTTDATDATDGYDGRAG | 97718 | 97526 | - | 8 |
| 17 | MESGPAIFAGTVFCLFGAALLAWTVTRARQHRPVALGVNRVASTTVA | GTVATMSLALGAWCFTRL | 104350 | 104152 | - | 8 |
| 18 | MSAGRILDAIAQLLGEDPVLLRD | RTPAQIRLLVEQGFLNPVD | 104772 | 104643 | - | 6 |
| 19 | VCDGSMSAGRILDAIAQLLGEDPVLLRD | RTPAQIRLLVEQGFLNPVD | 104787 | 104643 | - | 7 |
| 20 | VDTIGAGFAGVCDGSMSAGRILDAIAQLLGEDPVLLRD | RTPAQIRLLVEQGFLNPVD | 104817 | 104643 | - | 10 |
| 21 | MRRATKVDTIGAGFAGVCDGSMSAGRILDAIAQLLGEDPVLLRD | RTPAQIRLLVEQGFLNPVD | 104835 | 104643 | - | 10 |

### Results for WP_030414971.1 [Streptomyces sp. NRRL S-1448] back to top

previous - next

#### Architecture

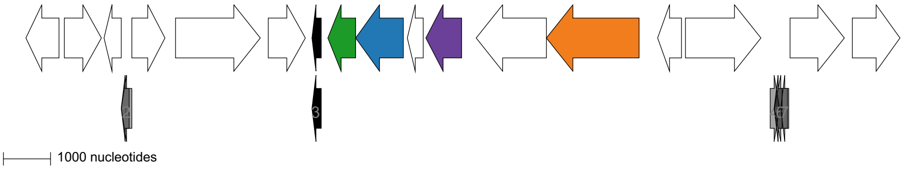

Link to nucleotide sequence

| Accession | start | end | direction | length (aa) | Pfam/HMM | name | description | E-value |
|---|---|---|---|---|---|---|---|---|
| WP_030414963.1 | 23707 | 23002 | - | 234 | PF01810<br>TIGR00949<br>TIGR00948 | LysE<br>TIGR00949<br>TIGR00948 | LysE type translocator<br>2A76: homoserine/Threonine efflux protein<br>2a75: L-lysine exporter | 1.10E-42<br>3.60E-41<br>1.60E-15 |
| WP_030414964.1 | 23821 | 24580 | + | 252 | PF07739<br>PF13411<br>TIGR02043<br>PF00376<br>TIGR02051 | TipAS<br>MerR_1<br>TIGR02043<br>MerR<br>TIGR02051 | TipAS antibiotic-recognition domain<br>MerR HTH family regulatory protein<br>ZntR: Zn(II)-responsive transcriptional regulator<br>MerR family regulatory protein<br>MerR: Hg(II)-responsive transcriptional regulator | 5.70E-37<br>1.50E-17<br>1.80E-14<br>2.70E-14<br>1.50E-13 |
| WP_030414965.1 | 25013 | 24644 | - | 122 | PF01906 | YbjQ_1 | Putative heavy-metal-binding | 1.90E-33 |
| WP_030414966.1 | 25234 | 25945 | + | 236 | PF09335 | SNARE_assoc | SNARE associated Golgi protein | 3.10E-24 |
| WP_078866335.1 | 26180 | 27959 | + | 592 | PF06738<br>PF12821 | ThrE<br>ThrE_2 | Putative threonine/serine exporter<br>Threonine/Serine exporter, ThrE | 1.40E-72<br>4.20E-19 |
| WP_006604049.1 | 28128 | 28920 | + | 264 | INFERRED GENE | - | - | - |
| WP_107052797.1 | 29262 | 29064 | - | 65 | LinA | LinA | LinA (linaridin precursor) | 3.10E-13 |

| | | | | | | | | |
|---|---|---|---|---|---|---|---|---|
| WP_030414970.1 | 29995 | 29392 | - | 200 | LinL | LinL | LinL | 5.20E-58 |
| WP_030414971.1 | 31008 | 29991 | - | 338 | LinE<br>LinH<br>PF12697<br>TIGR02427<br>PF00561 | LinE<br>LinH<br>Abhydrolase_6<br>TIGR02427<br>Abhydrolase_1 | LinE Homolog<br>LinH (Fusion of LinE/LinG)<br>Alpha/beta hydrolase family<br>protocat_pcaD: 3-oxoadipate enol-lactonase<br>alpha/beta hydrolase fold | 2.20E-102<br>5.10E-53<br>3.00E-14<br>8.00E-11<br>3.60E-10 |
| WP_078866336.1 | 31429 | 31081 | - | 115 | PF13518<br>PF01527<br>PF13384<br>PF13011<br>PF13551 | HTH_28<br>HTH_Tnp_1<br>HTH_23<br>LZ_Tnp_IS481<br>HTH_29 | Helix-turn-helix domain<br>Transposase<br>Homeodomain-like domain<br>leucine-zipper of insertion element IS481<br>Winged helix-turn helix | 1.40E-07<br>2.50E-07<br>2.10E-05<br>2.50E-05<br>7.80E-05 |
| WP_030414972.1 | 32231 | 31469 | - | 253 | PF13649<br>PF08241<br>PF08242<br>PF13489<br>PF13847 | Methyltransf_25<br>Methyltransf_11<br>Methyltransf_12<br>Methyltransf_23<br>Methyltransf_31 | Methyltransferase domain<br>Methyltransferase domain<br>Methyltransferase domain<br>Methyltransferase domain<br>Methyltransferase domain | 1.30E-16<br>2.10E-11<br>6.20E-11<br>1.30E-08<br>1.10E-07 |
| WP_030414973.1 | 34026 | 32541 | - | 494 | TIGR00292<br>PF13450<br>TIGR01317<br>TIGR02734<br>TIGR03315 | TIGR00292<br>NAD_binding_8<br>TIGR01317<br>TIGR02734<br>TIGR03315 | TIGR00292: thiazole biosynthesis enzyme<br>NAD(P)-binding Rossmann-like domain<br>GOGAT_sm_gam: glutamate synthase, NADH/NADPH, small subunit<br>crtI_fam: phytoene desaturase<br>Se_ygfK: putative selenate reductase, YgfK subunit | 3.20E-09<br>6.80E-09<br>2.70E-08<br>6.90E-08<br>2.00E-07 |
| WP_078866337.1 | 35984 | 34022 | - | 653 | TIGR02203<br>TIGR02204<br>TIGR00958<br>TIGR03796<br>TIGR03375 | TIGR02203<br>TIGR02204<br>TIGR00958<br>TIGR03796<br>TIGR03375 | MsbA_lipidA: lipid A export permease/ATP-binding protein MsbA<br>MsbA_rel: ABC transporter, permease/ATP-binding protein<br>3a01208: antigen peptide transporter 2<br>NHLM_micro_ABC1: NHLM bacteriocin system ABC transporter, peptidase/ATP-binding protein<br>type_I_sec_LssB: type I secretion system ATPase | 1.50E-100<br>1.70E-100<br>4.70E-91<br>1.50E-81<br>3.20E-80 |
| WP_030414975.1 | 36891 | 36399 | - | 163 | PF00719 | Pyrophosphatase | Inorganic pyrophosphatase | 5.00E-53 |
| WP_030414976.1 | 36977 | 38567 | + | 529 | PF02113<br>TIGR00666 | Peptidase_S13<br>TIGR00666 | D-Ala-D-Ala carboxypeptidase 3 (S13) family<br>PBP4: D-alanyl-D-alanine carboxypeptidase/D-alanyl-D-alanine-endopeptidase | 6.80E-73<br>3.90E-56 |
| WP_030414977.1 | 39190 | 40336 | + | 381 | TIGR03883<br>TIGR03624<br>PF10103 | TIGR03883<br>TIGR03624<br>Zincin_2 | DUF2342_F420: uncharacterized protein, coenzyme F420 biosynthesis associated<br>TIGR03624: putative hydrolase<br>Zincin-like metallopeptidase | 2.50E-142<br>2.50E-125<br>2.60E-122 |
| WP_030414978.1 | 40527 | 41535 | + | 335 | TIGR02432<br>PF01171<br>PF09179 | TIGR02432<br>ATP_bind_3<br>TilS | lysidine_TilS_N: tRNA(Ile)-lysidine synthetase<br>PP-loop family<br>TilS substrate binding domain | 7.90E-44<br>1.30E-38<br>3.20E-12 |

| index | leader | core | start | end | dir | score |
|---|---|---|---|---|---|---|
| 1 | VRRAGYERWTGPGAGCRPKAINAPRPP | SVPGARRGPHPTAPPPLAAPPDPGDPGH | 25195 | 25027 | - | 6 |
| 2 | VKAPGSRAPDRAVRRAGYERWTGPGAGCRPKAI | NAPRPPSVPGARRGPHPTAPPPLAAPPDPGDPGH | 25231 | 25027 | - | 6 |
| 3 | MSVVAEFANTNLVDVQPGRIGSDAAPTM | MTPLATVATPEAVSAGIAVTCATAIALNQVTHNVQHG | 29262 | 29064 | - | 34 |
| 4 | VDCCRFTSTQRPRDPYGICSNHPYGSQGPLAPPT | PTPTHLLPTPPTGGDGPEGPVFGT | 38788 | 38965 | + | 6 |
| 5 | VSGAVTGLRGRCLGLLSL | YVPNTGPSGPSPPVGGVGRRWVGVGVGGARGPWDP | 39018 | 38856 | - | 7 |
| 6 | LRFAAPPTYLAVPPRGSLAVALGGA...AVALCGWVSGAVTGLRGRCLGLLSL | YVPNTGPSGPSPPVGGVGRRWVGVGVGGARGPWDP | 39150 | 38856 | - | 6 |
| 7 | VGATAPGSRCALRRRPRTWLSRRGARSPLRL | AAPRTYLSVPPRGVSPLRFAGGCRVR | 39176 | 39002 | - | 6 |

### Results for WP_158744997.1 [Streptomyces sp. NRRL S-1448] back to top

previous - next

#### Architecture

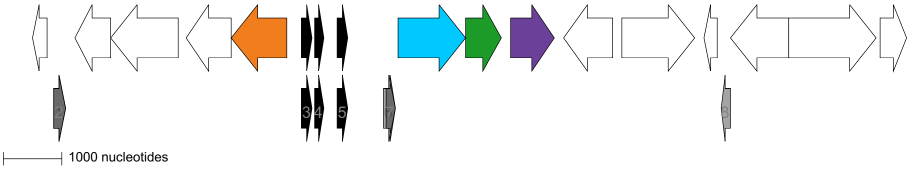

Link to nucleotide sequence

| Accession | start | end | direction | length (aa) | Pfam/HMM | name | description | E-value |
|---|---|---|---|---|---|---|---|---|
| WP_107052722.1 | 18085 | 17833 | - | 83 | NO PFAM MATCH | - | - | - |
| WP_030413056.1 | 19180 | 18574 | - | 201 | PF00072<br>TIGR02875<br>PF00196<br>TIGR02154<br>TIGR02956 | Response_reg<br>TIGR02875<br>GerE<br>TIGR02154<br>TIGR02956 | Response regulator receiver domain<br>spore_0_A: sporulation transcription factor Spo0A<br>Bacterial regulatory proteins, luxR family<br>PhoB: phosphate regulon transcriptional regulatory protein PhoB<br>TMAO_torS: TMAO reductase sytem sensor TorS | 1.70E-23<br>2.80E-14<br>1.50E-12<br>4.10E-10<br>5.40E-09 |
| WP_037808408.1 | 20346 | 19176 | - | 389 | PF07730 | HisKA_3 | Histidine kinase | 9.60E-15 |
| WP_030413058.1 | 21247 | 20482 | - | 254 | PF01061<br>PF12698<br>TIGR03062<br>TIGR01247 | ABC2_membrane<br>ABC2_membrane_3<br>TIGR03062<br>TIGR01247 | ABC-2 type transporter<br>ABC-2 family transporter protein<br>pip_yhgE_Cterm: YhgE/Pip C-terminal domain<br>drrB: daunorubicin resistance ABC transporter membrane protein | 2.00E-17<br>2.00E-13<br>4.40E-07<br>2.20E-04 |

| | | | | | | | | |
|---|---|---|---|---|---|---|---|---|
| WP_030413059.1 | 22203 | 21243 | - | 319 | TIGR01188<br>TIGR03864<br>TIGR01288<br>TIGR03522<br>TIGR03740 | TIGR01188<br>TIGR03864<br>TIGR01288<br>TIGR03522<br>TIGR03740 | drrA: daunorubicin resistance ABC transporter, ATP-binding protein<br>PQQ_ABC_ATP: ABC transporter, ATP-binding subunit, PQQ-dependent alcohol dehydrogenase system<br>nodI: nodulation ABC transporter NodI<br>GldA_ABC_ATP: gliding motility-associated ABC transporter ATP-binding subunit GldA<br>galliderm_ABC: lantibiotic protection ABC transporter, ATP-binding subunit | 4.50E-64<br>5.00E-46<br>8.00E-43<br>1.90E-36<br>2.60E-34 |
| WP_158744993.1 | 22446 | 22620 | + | 57 | LinA | LinA | LinA (linaridin precursor) | 6.80E-11 |
| WP_158744995.1 | 22665 | 22836 | + | 56 | LinA | LinA | LinA (linaridin precursor) | 3.80E-13 |
| WP_030413060.1 | 23062 | 23242 | + | 59 | LinA | LinA | LinA (linaridin precursor) | 6.90E-17 |
| WP_158744997.1 | 24103 | 25267 | + | 387 | LinH<br>LinE<br>LinG<br>PF12697<br>PF12146 | LinH<br>LinE<br>LinG<br>Abhydrolase_6<br>Hydrolase_4 | LinH (Fusion of LinE/LinG)<br>LinE Homolog<br>LinG Homolog<br>Alpha/beta hydrolase family<br>Serine aminopeptidase, S33 | 2.30E-142<br>1.50E-97<br>1.50E-19<br>1.10E-17<br>8.40E-09 |
| WP_037808412.1 | 25263 | 25875 | + | 203 | LinL | LinL | LinL | 1.50E-68 |
| WP_030413062.1 | 26017 | 26782 | + | 254 | PF13649<br>PF08241<br>PF13847<br>TIGR01983<br>PF08242 | Methyltransf_25<br>Methyltransf_11<br>Methyltransf_31<br>TIGR01983<br>Methyltransf_12 | Methyltransferase domain<br>Methyltransferase domain<br>Methyltransferase domain<br>UbiG: 3-demethylubiquinone-9 3-O-methyltransferase<br>Methyltransferase domain | 2.10E-17<br>6.00E-12<br>8.80E-10<br>3.10E-09<br>5.00E-09 |
| WP_030413063.1 | 27783 | 26925 | - | 285 | TIGR02279<br>PF02737<br>TIGR02440<br>TIGR02437<br>TIGR02441 | TIGR02279<br>3HCDH_N<br>TIGR02440<br>TIGR02437<br>TIGR02441 | PaaC-3OHAcCoADH: 3-hydroxyacyl-CoA dehydrogenase PaaC<br>3-hydroxyacyl-CoA dehydrogenase, NAD binding domain<br>FadJ: fatty oxidation complex, alpha subunit FadJ<br>FadB: fatty oxidation complex, alpha subunit FadB<br>fa_ox_alpha_mit: fatty acid oxidation complex, alpha subunit, mitochondrial | 2.30E-71<br>2.20E-59<br>2.80E-55<br>1.20E-51<br>1.80E-50 |
| WP_030413064.1 | 27942 | 29172 | + | 409 | PF02638 | GHL10 | Glycosyl hydrolase-like 10 | 4.40E-81 |
| WP_030413065.1 | 29555 | 29345 | - | 69 | PF08940 | DUF1918 | Domain of unknown function (DUF1918) | 3.00E-26 |
| WP_030413066.1 | 30789 | 29787 | - | 333 | PF00892<br>TIGR00950 | EamA<br>TIGR00950 | EamA-like transporter family<br>2A78: carboxylate/amino acid/amine transporter | 3.20E-23<br>3.60E-11 |
| WP_030413067.1 | 30787 | 32284 | + | 498 | TIGR04461<br>PF00155<br>TIGR03947<br>PF00392<br>TIGR01265 | TIGR04461<br>Aminotran_1_2<br>TIGR03947<br>GntR<br>TIGR01265 | endura_MppQ: enduracididine biosynthesis enzyme MppQ<br>Aminotransferase class I and II<br>viomycin_VioD: capreomycidine synthase<br>Bacterial regulatory proteins, gntR family<br>tyr_nico_aTase: tyrosine/nicotianamine family aminotransferase | 1.90E-34<br>6.00E-23<br>9.30E-14<br>4.20E-13<br>2.10E-12 |
| WP_030413068.1 | 32342 | 32795 | + | 150 | PF00583<br>PF13508<br>PF13673<br>PF08445<br>TIGR01575 | Acetyltransf_1<br>Acetyltransf_7<br>Acetyltransf_10<br>FR47<br>TIGR01575 | Acetyltransferase (GNAT) family<br>Acetyltransferase (GNAT) domain<br>Acetyltransferase (GNAT) domain<br>FR47-like protein<br>rimI: ribosomal-protein-alanine acetyltransferase | 1.70E-10<br>4.10E-09<br>3.30E-08<br>3.10E-06<br>5.70E-05 |

| index | leader | core | start | end | dir | score |
|---|---|---|---|---|---|---|
| 1 | LNSGLAWSSGSSKRRTPNAER | RTPNAERRAEAEPTPKPIGRHHSTLAAATVPLPLCRCAAVPLCQRTQGH | 18198 | 18411 | + | 6 |
| 2 | LAWSSGSSKRRTPNAER | RTPNAERRAEAEPTPKPIGRHHSTLAAATVPLPLCRCAAVPLCQRTQGH | 18210 | 18411 | + | 6 |
| 3 | MLDDTFASRLEGLGGLSEENAGRIEGLASPVL | ATPALAFIGGAAVVTGAYAAGQAVG | 22446 | 22620 | + | 31 |
| 4 | MQGIAEYLNTDVQVSAIDVEEMEGTIFSDHAPVL | ATPAALTVAAFASGFAVCRAVG | 22665 | 22836 | + | 30 |
| 5 | MTELITQVNESEVLTETAISADEGRVFPGATPVL | ATPAAFLGGVAVSCAVVGAFEAGRG | 23062 | 23242 | + | 30 |
| 6 | VVRGVAGQRLLPHIGLGEIAGQRVEGRDCTQR | RHADQDVRPRAHLQADAEVSEGREVPDARCARS | 23855 | 24053 | + | 6 |
| 7 | LGEIAGQRVEGRDCTQRRHADQDVR | PRAHLQADAEVSEGREVPDARCARS | 23900 | 24053 | + | 6 |
| 8 | VTGPGAAPPCPAAGSGGNRRPAPRV | RRAPADVNLPRWQIACRPGRARPVTP | 29791 | 29635 | - | 6 |

### Results for WP_111003021.1 [Streptomyces sp. NTH33] back to top

previous - next

#### Architecture

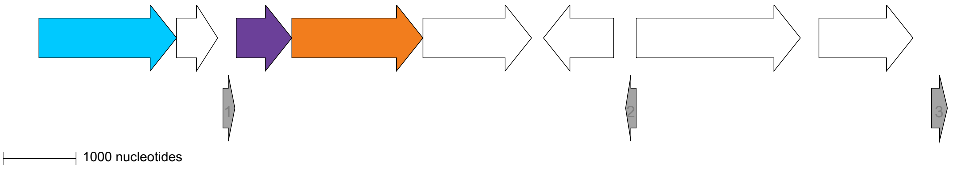

Link to nucleotide sequence

| Accession | start | end | direction | length (aa) | Pfam/HMM | name | description | E-value |
|---|---|---|---|---|---|---|---|---|
| WP_111003021.1 | 28 | 1924 | + | 631 | LinH<br>LinG<br>LinE<br>PF12697<br>LinL | LinH<br>LinG<br>LinE<br>Abhydrolase_6<br>LinL | LinH (Fusion of LinE/LinG)<br>LinG Homolog<br>LinE Homolog<br>Alpha/beta hydrolase family<br>LinL | 8.10E-249<br>1.30E-121<br>6.10E-116<br>2.30E-15<br>8.30E-10 |
| WP_019435954.1 | 1920 | 2481 | + | 187 | INFERRED GENE | - | - | - |

| | | | | | | | | |
|---|---|---|---|---|---|---|---|---|
| WP_111003012.1 | 2735 | 3497 | + | 253 | PF13649<br>PF08241<br>PF13489<br>PF08242<br>TIGR01983 | Methyltransf_25<br>Methyltransf_11<br>Methyltransf_23<br>Methyltransf_12<br>TIGR01983 | Methyltransferase domain<br>Methyltransferase domain<br>Methyltransferase domain<br>Methyltransferase domain<br>UbiG: 3-demethylubiquinone-9 3-O-methyltransferase | 7.40E-17<br>9.30E-11<br>2.30E-09<br>1.10E-07<br>1.20E-07 |
| WP_111003013.1 | 3500 | 5303 | + | 600 | TIGR02204<br>TIGR02203<br>TIGR00958<br>TIGR01846<br>TIGR03796 | TIGR02204<br>TIGR02203<br>TIGR00958<br>TIGR01846<br>TIGR03796 | MsbA_rel: ABC transporter, permease/ATP-binding protein<br>MsbA_lipidA: lipid A export permease/ATP-binding protein MsbA<br>3a01208: antigen peptide transporter 2<br>type_I_sec_HlyB: type I secretion system ATPase<br>NHLM_micro_ABC1: NHLM bacteriocin system ABC transporter, peptidase/ATP-binding protein | 5.10E-102<br>1.10E-101<br>2.00E-88<br>3.00E-84<br>1.50E-83 |
| WP_111003014.1 | 5299 | 6790 | + | 496 | TIGR03467<br>TIGR02734<br>PF13450<br>TIGR00292<br>PF01593 | TIGR03467<br>TIGR02734<br>NAD_binding_8<br>TIGR00292<br>Amino_oxidase | HpnE: squalene-associated FAD-dependent desaturase<br>crtI_fam: phytoene desaturase<br>NAD(P)-binding Rossmann-like domain<br>TIGR00292: thiazole biosynthesis enzyme<br>Flavin containing amine oxidoreductase | 6.60E-10<br>8.50E-10<br>3.70E-08<br>1.30E-07<br>2.20E-07 |
| WP_111003015.1 | 7917 | 6945 | - | 323 | TIGR03541 | TIGR03541 | reg_near_HchA: LuxR family transcriptional regulatory, chaperone HchA-associated | 4.20E-05 |
| WP_111003022.1 | 8218 | 10483 | + | 754 | TIGR01448<br>TIGR01447<br>PF13604<br>PF14490<br>TIGR02768 | TIGR01448<br>TIGR01447<br>AAA_30<br>HHH_4<br>TIGR02768 | recD_rel: helicase, RecD/TraA family<br>recD: exodeoxyribonuclease V, alpha subunit<br>AAA domain<br>Helix-hairpin-helix containing domain<br>TraA_Ti: Ti-type conjugative transfer relaxase TraA | 3.50E-201<br>7.40E-70<br>9.40E-46<br>2.90E-37<br>4.40E-35 |
| WP_111003016.1 | 10736 | 12026 | + | 429 | TIGR01798<br>PF00285<br>TIGR01800<br>TIGR01793 | TIGR01798<br>Citrate_synt<br>TIGR01800<br>TIGR01793 | cit_synth_I: citrate (Si)-synthase<br>Citrate synthase, C-terminal domain<br>cit_synth_II: 2-methylcitrate synthase/citrate synthase II<br>cit_synth_euk: citrate (Si)-synthase, eukaryotic | 2.50E-187<br>3.40E-135<br>6.30E-105<br>3.00E-31 |

| index | leader | core | start | end | dir | score |
|---|---|---|---|---|---|---|
| 1 | VICDEPSTKECSRNHVCSNDHHLR | HRHHHRRGDCLENHEVTETSVTKNP | 2562 | 2712 | + | 6 |
| 2 | MNVTVPPGCDSGEGRGAPAQEHA | VGSETESGCVRFVRNSTGKPRTGM | 8220 | 8076 | - | 6 |
| 3 | VSTTSRTRRALLVPQSYCRCDLPGQRTHGRAAQSFPLYLPRQRISE | EVPQDIPGCRSAPPKTPPGMTSPT | 12279 | 12492 | + | 6 |

### Results for ADR72963.1 [Streptomyces sp. OH-4156] back to top

previous - next

#### Architecture

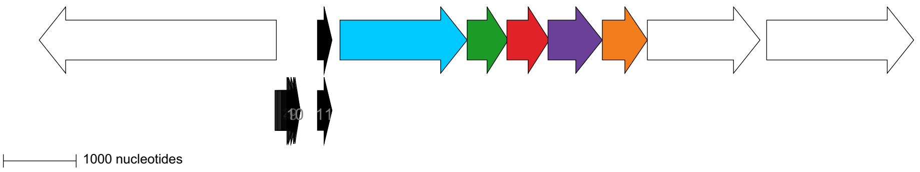

Link to nucleotide sequence

| Accession | start | end | direction | length (aa) | Pfam/HMM | name | description | E-value |
|---|---|---|---|---|---|---|---|---|
| ADR72961.1 | 3308 | 56 | - | 1083 | PF13424<br>PF17874<br>PF13374<br>PF07719<br>PF13191 | TPR_12<br>TPR_MalT<br>TPR_10<br>TPR_2<br>AAA_16 | Tetratricopeptide repeat<br>MalT-like TPR region<br>Tetratricopeptide repeat<br>Tetratricopeptide repeat<br>AAA ATPase domain | 1.30E-19<br>5.10E-12<br>1.30E-10<br>1.70E-09<br>1.90E-08 |
| ADR72962.1 | 3881 | 4076 | + | 64 | LinA | LinA | LinA (linaridin precursor) | 1.70E-13 |
| ADR72963.1 | 4190 | 5945 | + | 584 | LinH<br>LinG<br>LinE<br>PF12697<br>TIGR02427 | LinH<br>LinG<br>LinE<br>Abhydrolase_6<br>TIGR02427 | LinH (Fusion of LinE/LinG)<br>LinG Homolog<br>LinE Homolog<br>Alpha/beta hydrolase family<br>protocat_pcaD: 3-oxoadipate enol-lactonase | 3.90E-172<br>9.40E-75<br>6.40E-69<br>7.90E-08<br>3.50E-07 |
| ADR72964.1 | 5934 | 6495 | + | 186 | LinL | LinL | LinL | 1.40E-53 |
| ADR72965.1 | 6491 | 7064 | + | 190 | TIGR00521<br>TIGR02113<br>PF02441 | TIGR00521<br>TIGR02113<br>Flavoprotein | coaBC_dfp: phosphopantothenoylcysteine decarboxylase / phosphopantothenate--cysteine ligase<br>coaC_strep: phosphopantothenoylcysteine decarboxylase<br>Flavoprotein | 1.70E-11<br>1.10E-09<br>1.20E-08 |
| ADR72966.1 | 7056 | 7794 | + | 245 | PF13649<br>PF08241<br>TIGR02021<br>TIGR01934<br>PF08242 | Methyltransf_25<br>Methyltransf_11<br>TIGR02021<br>TIGR01934<br>Methyltransf_12 | Methyltransferase domain<br>Methyltransferase domain<br>BchM-ChlM: magnesium protoporphyrin O-methyltransferase<br>MenG_MenH_UbiE: ubiquinone/menaquinone biosynthesis methyltransferase<br>Methyltransferase domain | 2.00E-19<br>3.10E-17<br>9.30E-13<br>2.10E-12<br>2.60E-12 |
| ADR72967.1 | 7790 | 8423 | + | 210 | TIGR02857<br>TIGR01189<br>TIGR02673<br>TIGR03873<br>TIGR02982 | TIGR02857<br>TIGR01189<br>TIGR02673<br>TIGR03873<br>TIGR02982 | CydD: thiol reductant ABC exporter, CydD subunit<br>ccmA: heme ABC exporter, ATP-binding protein CcmA<br>FtsE: cell division ATP-binding protein FtsE<br>F420-0_ABC_ATP: proposed F420-0 ABC transporter, ATP-binding protein<br>heterocyst_DevA: ABC exporter ATP-binding subunit, DevA family | 2.20E-28<br>5.90E-26<br>7.70E-26<br>1.90E-24<br>7.00E-23 |
| ADR72968.1 | 8419 | 9970 | + | 516 | NO PFAM MATCH | - | - | - |
| ADR72969.1 | 10052 | 12074 | + | 673 | PF03190<br>PF13899<br>PF07221 | Thioredox_DsbH<br>Thioredoxin_7<br>GlcNAc_2-epim | Protein of unknown function, DUF255<br>Thioredoxin-like<br>N-acylglucosamine 2-epimerase (GlcNAc 2-epimerase) | 1.10E-74<br>2.20E-10<br>2.10E-05 |

| index | leader | core | start | end | dir | score |
|---|---|---|---|---|---|---|
| 1 | MAVPHVRPPPRVPAHPVPEALSDPG...TGRMPSGPGSVGPGTRGFRARPPGG | GAPPRGGPSNTVRGRRNHRPQSSESPRIKDP | 3293 | 3653 | + | 6 |

| | | | | | | |
|---|---|---|---|---|---|---|
| 2 | MCVPLRACPPIPSLRPCQTPDRARP...AGRAPWGPAPEASAPGPRAAGRPRE | AAPQTPFAAGEITALNLPSRPV | 3306 | 3642 | + | 7 |
| 3 | VRPPPRVPAHPVPEALSDPGPCASS...TGRMPSGPGSVGPGTRGFRARPPGG | GAPPRGGPSNTVRGRRNHRPQSSESPRIKDP | 3308 | 3653 | + | 6 |
| 4 | VPAHPVPEALSDPGPCASSVLRGAR...TGRMPSGPGSVGPGTRGFRARPPGG | GAPPRGGPSNTVRGRRNHRPQSSESPRIKDP | 3326 | 3653 | + | 6 |
| 5 | VCCAGREGARAGKGGPAAAGRSASR...AGRAPWGPAPEASAPGPRAAGRPRE | AAPQTPFAAGEITALNLPSRPV | 3381 | 3642 | + | 8 |
| 6 | VLRGARRRESGQRRAGRGGTVGIPN...TGRMPSGPGSVGPGTRGFRARPPGG | GAPPRGGPSNTVRGRRNHRPQSSESPRIKDP | 3383 | 3653 | + | 6 |
| 7 | VRGVVGRELAGCRAGRAPWGPAPEASAPGPRAAGRPRE | AAPQTPFAAGEITALNLPSRPV | 3459 | 3642 | + | 8 |
| 8 | VVGRELAGCRAGRAPWGPAPEASAPGPRAAGRPRE | AAPQTPFAAGEITALNLPSRPV | 3468 | 3642 | + | 8 |
| 9 | VGRELAGCRAGRAPWGPAPEASAPGPRAAGRPRE | AAPQTPFAAGEITALNLPSRPV | 3471 | 3642 | + | 8 |
| 10 | MPSGPGSVGPGTRGFRARPPGG | GAPPRGGPSNTVRGRRNHRPQSSESPRIKDP | 3491 | 3653 | + | 6 |
| 11 | MRSEMTLTSTNSAEALAAQDFANTVLSAAAPGFHADCETPAM | ATPATPTVAQFVIQGSTICLVC | 3881 | 4076 | + | 35 |

### Results for WP_030801616.1 [Streptomyces sp. NRRL S-337] back to top

previous - next

#### Architecture

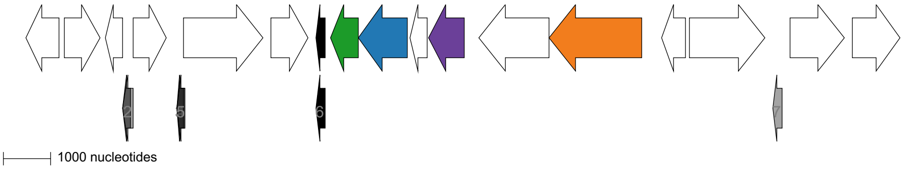

Link to nucleotide sequence

| Accession | start | end | direction | length (aa) | Pfam/HMM | name | description | E-value |
|---|---|---|---|---|---|---|---|---|
| WP_030801599.1 | 111349 | 110659 | - | 229 | PF01810<br>TIGR00949<br>TIGR00948 | LysE<br>TIGR00949<br>TIGR00948 | LysE type translocator<br>2A76: homoserine/Threonine efflux protein<br>2a75: L-lysine exporter | 2.30E-43<br>5.40E-42<br>2.10E-15 |
| WP_030801601.1 | 111463 | 112222 | + | 252 | PF07739<br>PF13411<br>TIGR02043<br>PF00376<br>TIGR02051 | TipAS<br>MerR_1<br>TIGR02043<br>MerR<br>TIGR02051 | TipAS antibiotic-recognition domain<br>MerR HTH family regulatory protein<br>ZntR: Zn(II)-responsive transcriptional regulator<br>MerR family regulatory protein<br>MerR: Hg(II)-responsive transcriptional regulator | 1.10E-35<br>2.90E-17<br>1.50E-14<br>2.70E-14<br>6.70E-14 |
| WP_030414965.1 | 112688 | 112319 | - | 122 | PF01906 | YbjQ_1 | Putative heavy-metal-binding | 1.90E-33 |
| WP_030801603.1 | 112909 | 113620 | + | 236 | PF09335 | SNARE_assoc | SNARE associated Golgi protein | 7.30E-25 |
| WP_030801606.1 | 113993 | 115676 | + | 560 | PF06738<br>PF12821 | ThrE<br>ThrE_2 | Putative threonine/serine exporter<br>Threonine/Serine exporter, ThrE | 4.00E-73<br>2.60E-19 |
| WP_006604049.1 | 115833 | 116625 | + | 264 | INFERRED GENE | - | - | - |
| WP_107052972.1 | 116981 | 116783 | - | 65 | LinA | LinA | LinA (linaridin precursor) | 5.30E-13 |
| WP_030801613.1 | 117697 | 117109 | - | 195 | LinL | LinL | LinL | 1.20E-57 |
| WP_030801616.1 | 118713 | 117693 | - | 339 | LinE<br>LinH<br>PF12697<br>TIGR02427<br>PF00561 | LinE<br>LinH<br>Abhydrolase_6<br>TIGR02427<br>Abhydrolase_1 | LinE Homolog<br>LinH (Fusion of LinE/LinG)<br>Alpha/beta hydrolase family<br>protocat_pcaD: 3-oxoadipate enol-lactonase<br>alpha/beta hydrolase fold | 7.60E-104<br>1.20E-54<br>2.00E-14<br>2.00E-12<br>6.80E-10 |
| WP_078871990.1 | 119134 | 118786 | - | 115 | PF13518<br>PF01527<br>PF13384<br>PF13011<br>PF13551 | HTH_28<br>HTH_Tnp_1<br>HTH_23<br>LZ_Tnp_IS481<br>HTH_29 | Helix-turn-helix domain<br>Transposase<br>Homeodomain-like domain<br>leucine-zipper of insertion element IS481<br>Winged helix-turn helix | 1.40E-07<br>2.50E-07<br>2.00E-05<br>2.30E-05<br>6.90E-05 |
| WP_030801619.1 | 119937 | 119175 | - | 253 | PF13649<br>PF08241<br>PF13489<br>PF08242<br>PF13847 | Methyltransf_25<br>Methyltransf_11<br>Methyltransf_23<br>Methyltransf_12<br>Methyltransf_31 | Methyltransferase domain<br>Methyltransferase domain<br>Methyltransferase domain<br>Methyltransferase domain<br>Methyltransferase domain | 8.40E-16<br>6.10E-11<br>6.40E-09<br>9.40E-09<br>8.70E-08 |
| WP_030801621.1 | 121732 | 120247 | - | 494 | TIGR00292<br>PF13450<br>TIGR01317<br>TIGR02734<br>TIGR01318 | TIGR00292<br>NAD_binding_8<br>TIGR01317<br>TIGR02734<br>TIGR01318 | TIGR00292: thiazole biosynthesis enzyme<br>NAD(P)-binding Rossmann-like domain<br>GOGAT_sm_gam: glutamate synthase, NADH/NADPH, small subunit<br>crtI_fam: phytoene desaturase<br>gltD_gamma_fam: glutamate synthase, small subunit | 3.20E-09<br>6.60E-09<br>1.10E-08<br>1.00E-07<br>2.50E-07 |
| WP_078871991.1 | 123687 | 121728 | - | 652 | TIGR02204<br>TIGR02203<br>TIGR00958<br>TIGR03796<br>TIGR03375 | TIGR02204<br>TIGR02203<br>TIGR00958<br>TIGR03796<br>TIGR03375 | MsbA_rel: ABC transporter, permease/ATP-binding protein<br>MsbA_lipidA: lipid A export permease/ATP-binding protein MsbA<br>3a01208: antigen peptide transporter 2<br>NHLM_micro_ABC1: NHLM bacteriocin system ABC transporter, peptidase/ATP-binding protein<br>type_I_sec_LssB: type I secretion system ATPase | 4.70E-104<br>7.60E-101<br>2.90E-92<br>4.10E-83<br>1.80E-82 |
| WP_030414975.1 | 124609 | 124117 | - | 163 | PF00719 | Pyrophosphatase | Inorganic pyrophosphatase | 5.00E-53 |
| WP_030801626.1 | 124695 | 126285 | + | 529 | PF02113<br>TIGR00666 | Peptidase_S13<br>TIGR00666 | D-Ala-D-Ala carboxypeptidase 3 (S13) family<br>PBP4: D-alanyl-D-alanine carboxypeptidase/D-alanyl-D-alanine-endopeptidase | 1.20E-72<br>1.20E-55 |
| WP_030801631.1 | 126813 | 127959 | + | 381 | TIGR03883<br>TIGR03624<br>PF10103 | TIGR03883<br>TIGR03624<br>Zincin_2 | DUF2342_F420: uncharacterized protein, coenzyme F420 biosynthesis associated<br>TIGR03624: putative hydrolase<br>Zincin-like metallopeptidase | 2.60E-142<br>2.50E-125<br>2.40E-122 |

| | | | | | | | | |
|---|---|---|---|---|---|---|---|---|
| WP_030801634.1 | 128150 | 129158 | + | 335 | TIGR02432<br>PF01171<br>PF09179 | TIGR02432<br>ATP_bind_3<br>TilS | lysidine_TilS_N: tRNA(Ile)-lysidine synthetase<br>PP-loop family<br>TilS substrate binding domain | 1.30E-43<br>1.90E-38<br>3.20E-12 |

| index | leader | core | start | end | dir | score |
|---|---|---|---|---|---|---|
| 1 | VRRAGYERWTGPGAGCRPKAINAPRPP | SVPGARPGPHPTAPPPLAVPPDPGDPGH | 112870 | 112702 | - | 7 |
| 2 | VKAPGSRAPDRAVRRAGYERWTGPGAGCRPKAI | NAPRPPSVPGARPGPHPTAPPPLAVPPDPGDPGH | 112906 | 112702 | - | 6 |
| 3 | MLLPVPLPLG | LVPLALVSLGLVSLGLVPLLSLSVLRWPPCGSDATVFAPPVPR | 113995 | 113833 | - | 7 |
| 4 | MGPMLLPVPLPLG | LVPLALVSLGLVSLGLVPLLSLSVLRWPPCGSDATVFAPPVPR | 114004 | 113833 | - | 7 |
| 5 | LRSSMGPMLLPVPLPLG | LVPLALVSLGLVSLGLVPLLSLSVLRWPPCGSDATVFAPPVPR | 114016 | 113833 | - | 6 |
| 6 | MSVVAEFANTNLVDVKPGRIGSDAAPTM | MTPLATVATPEAVSAGIAVTCATAIALNQVTHNVQHG | 116981 | 116783 | - | 34 |
| 7 | LRFAAPRTWLSFPSRGVSPLRLAAGVCLSGAVTA | LQTSSSGLSPPVSGGLEGRWGWVSSSWLYVVFGT | 126660 | 126453 | - | 6 |

### Results for WP_032773956.1 [Streptomyces sp. OspMP-M43] back to top

previous - next

#### Architecture

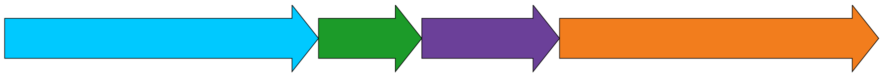

1000 nucleotides

Link to nucleotide sequence

| Accession | start | end | direction | length (aa) | Pfam/HMM | name | description | E-value |
|---|---|---|---|---|---|---|---|---|
| WP_032773956.1 | 284 | 2126 | + | 613 | LinH<br>LinG<br>LinE<br>LinL<br>PF12697 | LinH<br>LinG<br>LinE<br>LinL<br>Abhydrolase_6 | LinH (Fusion of LinE/LinG)<br>LinG Homolog<br>LinE Homolog<br>LinL<br>Alpha/beta hydrolase family | 3.40E-254<br>2.70E-112<br>2.60E-94<br>2.60E-35<br>2.00E-14 |
| WP_030703686.1 | 2122 | 2731 | + | 202 | LinL | LinL | LinL | 1.40E-65 |
| WP_003965565.1 | 2727 | 3540 | + | 270 | PF13649<br>PF08241<br>PF08242<br>PF13489<br>PF13847 | Methyltransf_25<br>Methyltransf_11<br>Methyltransf_12<br>Methyltransf_23<br>Methyltransf_31 | Methyltransferase domain<br>Methyltransferase domain<br>Methyltransferase domain<br>Methyltransferase domain<br>Methyltransferase domain | 2.30E-16<br>7.20E-12<br>5.40E-08<br>1.20E-07<br>1.60E-06 |
| WP_093564618.1 | 3536 | 5408 | + | 624 | TIGR02204<br>TIGR02203<br>TIGR00958<br>TIGR03796<br>TIGR03797 | TIGR02204<br>TIGR02203<br>TIGR00958<br>TIGR03796<br>TIGR03797 | MsbA_rel: ABC transporter, permease/ATP-binding protein<br>MsbA_lipidA: lipid A export permease/ATP-binding protein MsbA<br>3a01208: antigen peptide transporter 2<br>NHLM_micro_ABC1: NHLM bacteriocin system ABC transporter, peptidase/ATP-binding protein<br>NHLM_micro_ABC2: NHLM bacteriocin system ABC transporter, ATP-binding protein | 4.40E-102<br>3.40E-92<br>6.50E-85<br>1.30E-82<br>9.10E-77 |

| index | leader | core | start | end | dir | score |
|---|---|---|---|---|---|---|

### Results for WP_097967158.1 [Streptomyces sp. or20] back to top

previous - next

#### Architecture

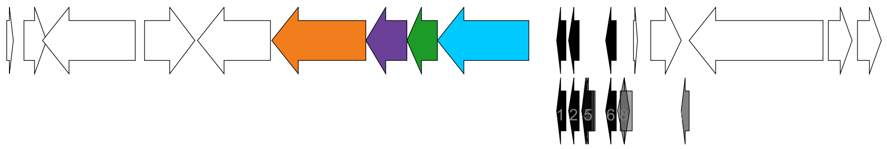

1000 nucleotides

Link to nucleotide sequence

| Accession | start | end | direction | length (aa) | Pfam/HMM | name | description | E-value |
|---|---|---|---|---|---|---|---|---|
| WP_009062325.1 | 4822 | 4948 | + | 42 | INFERRED GENE | - | - | - |
| WP_097967152.1 | 5144 | 5612 | + | 155 | PF13276 | HTH_21 | HTH-like domain | 4.80E-08 |
| WP_097967153.1 | 7380 | 5526 | - | 617 | NO PFAM MATCH | - | - | - |
| WP_097967154.1 | 7569 | 8583 | + | 337 | PF01978 | TrmB | Sugar-specific transcriptional regulator TrmB | 2.40E-04 |
| WP_097967172.1 | 10098 | 8637 | - | 486 | TIGR02734<br>PF13450<br>TIGR01317<br>TIGR00292<br>TIGR03467 | TIGR02734<br>NAD_binding_8<br>TIGR01317<br>TIGR00292<br>TIGR03467 | crtI_fam: phytoene desaturase<br>NAD(P)-binding Rossmann-like domain<br>GOGAT_sm_gam: glutamate synthase, NADH/NADPH, small subunit<br>TIGR00292: thiazole biosynthesis enzyme<br>HpnE: squalene-associated FAD-dependent desaturase | 6.20E-11<br>1.20E-08<br>1.20E-08<br>1.40E-08<br>1.60E-07 |

| | | | | | | | | |
|---|---|---|---|---|---|---|---|---|
| WP_097967155.1 | 12005 | 10103 | - | 633 | TIGR02204<br>TIGR02203<br>TIGR00958<br>TIGR03796<br>TIGR03375 | TIGR02204<br>TIGR02203<br>TIGR00958<br>TIGR03796<br>TIGR03375 | MsbA_rel: ABC transporter, permease/ATP-binding protein<br>MsbA_lipidA: lipid A export permease/ATP-binding protein MsbA<br>3a01208: antigen peptide transporter 2<br>NHLM_micro_ABC1: NHLM bacteriocin system ABC transporter, peptidase/ATP-binding protein<br>type_I_sec_LssB: type I secretion system ATPase | 3.80E-101<br>5.60E-93<br>1.70E-83<br>1.20E-81<br>6.90E-77 |
| WP_097967156.1 | 12814 | 12001 | - | 270 | PF13649<br>PF08241<br>PF08242<br>PF13489<br>PF13847 | Methyltransf_25<br>Methyltransf_11<br>Methyltransf_12<br>Methyltransf_23<br>Methyltransf_31 | Methyltransferase domain<br>Methyltransferase domain<br>Methyltransferase domain<br>Methyltransferase domain<br>Methyltransferase domain | 2.30E-16<br>7.50E-12<br>3.40E-08<br>2.70E-07<br>1.90E-06 |
| WP_097967157.1 | 13419 | 12810 | - | 202 | LinL | LinL | LinL | 3.60E-65 |
| WP_097967158.1 | 15263 | 13415 | - | 615 | LinH<br>LinG<br>LinE<br>LinL<br>PF12697 | LinH<br>LinG<br>LinE<br>LinL<br>Abhydrolase_6 | LinH (Fusion of LinE/LinG)<br>LinG Homolog<br>LinE Homolog<br>LinL<br>Alpha/beta hydrolase family | 1.40E-252<br>4.10E-111<br>2.10E-94<br>2.90E-36<br>1.80E-14 |
| WP_050362034.1 | 16003 | 15802 | - | 66 | LinA | LinA | LinA (linaridin precursor) | 3.70E-14 |
| WP_050362035.1 | 16255 | 16060 | - | 64 | LinA | LinA | LinA (linaridin precursor) | 7.70E-11 |
| WP_057667037.1 | 16996 | 16792 | - | 67 | LinA | LinA | LinA (linaridin precursor) | 3.40E-11 |
| WP_007460832.1 | 17353 | 17437 | + | 28 | INFERRED GENE | - | - | - |
| NP_630878.1 | 17686 | 18291 | + | 201 | INFERRED GENE | - | - | - |
| WP_097967159.1 | 21125 | 18470 | - | 884 | PF13191<br>TIGR03903<br>PF13424<br>PF07719<br>PF13401 | AAA_16<br>TIGR03903<br>TPR_12<br>TPR_2<br>AAA_22 | AAA ATPase domain<br>TOMM_kin_cyc: TOMM system kinase/cyclase fusion protein<br>Tetratricopeptide repeat<br>Tetratricopeptide repeat<br>AAA domain | 4.20E-20<br>4.70E-19<br>4.80E-05<br>5.50E-05<br>8.60E-05 |
| WP_073742616.1 | 21235 | 21715 | + | 159 | PF03745 | DUF309 | Domain of unknown function (DUF309) | 6.30E-20 |
| WP_097967160.1 | 21815 | 22313 | + | 165 | NO PFAM MATCH | - | - | - |

| index | leader | core | start | end | dir | score |
|---|---|---|---|---|---|---|
| 1 | MSMSPTPAALRGTGGLSESDPGRALSSLAPVA | ATPGVVAGVAVGVALVNAFAAGYNHCGGNVELPM | 16003 | 15802 | - | 31 |
| 2 | MNKSSAPAVLTATGALTESAPGVTLPTLAPVA | ATPVAIAATMGVAFVAGYAAGRAATGNVELPM | 16255 | 16060 | - | 30 |
| 3 | LNTASIPAVLSETGALSESDHGRAPLD | TVPVASVTFTMTACVEVSVCLTGSVIQLPQ | 16499 | 16325 | - | 9 |
| 4 | MTSPHRPPTSEEKSLNTASIPAVLSETGALSESDHGRAPLD | TVPVASVTFTMTACVEVSVCLTGSVIQLPQ | 16541 | 16325 | - | 8 |
| 5 | VGPSRRRRRPPMTSPHRPPTSEEKS...TASIPAVLSETGALSESDHGRAPLD | TVPVASVTFTMTACVEVSVCLTGSVIQLPQ | 16574 | 16325 | - | 7 |
| 6 | MPEFREPGWTRGIAPLQEGAGGQVFGGASPVA | ATPAVVATAGAVVVAFAAGVAARHLANGGNVELPM | 16996 | 16792 | - | 31 |
| 7 | MRSTRCREPDGKGRQLRWHEDADSEATC | CGFGHGGMRRVPIPIVGPGARAPFPTSR | 17091 | 17262 | + | 6 |
| 8 | LRPGCRNRRHRPGEGSPRASPRISVRWERAHALQDRLSESVRAAYPRAR | TRSMLLRSPHPHASGVVFPFRRALGIVWSASQNVCSRGAGSRVRLRAPG | 17329 | 17032 | - | 6 |
| 9 | LSCAVTRGNRGERDGTWRSLVAHLTGGQ | GVAGSNPVVPTGDSRRSGPVSEQSETGP | 18462 | 18291 | - | 7 |

### Results for WP_057667039.1 [Streptomyces sp. or3] back to top

previous - next

#### Architecture

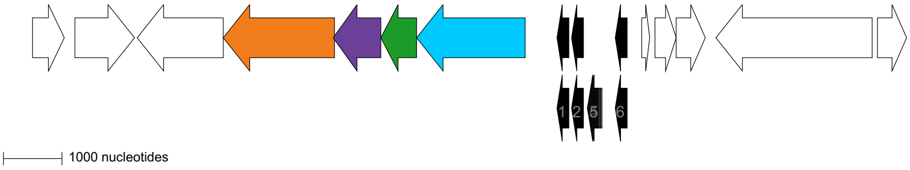

Link to nucleotide sequence

| Accession | start | end | direction | length (aa) | Pfam/HMM | name | description | E-value |
|---|---|---|---|---|---|---|---|---|
| WP_018487277.1 | 0 | 548 | + | 182 | INFERRED GENE | - | - | - |
| WP_056705451.1 | 737 | 1751 | + | 337 | NO PFAM MATCH | - | - | - |
| WP_098891769.1 | 3266 | 1805 | - | 486 | TIGR02734<br>PF13450<br>TIGR01317<br>TIGR00292<br>TIGR03467 | TIGR02734<br>NAD_binding_8<br>TIGR01317<br>TIGR00292<br>TIGR03467 | crtI_fam: phytoene desaturase<br>NAD(P)-binding Rossmann-like domain<br>GOGAT_sm_gam: glutamate synthase, NADH/NADPH, small subunit<br>TIGR00292: thiazole biosynthesis enzyme<br>HpnE: squalene-associated FAD-dependent desaturase | 3.50E-11<br>1.20E-08<br>1.20E-08<br>1.40E-08<br>7.40E-08 |
| WP_057667041.1 | 5155 | 3271 | - | 627 | TIGR02204<br>TIGR02203<br>TIGR00958<br>TIGR03796<br>TIGR03375 | TIGR02204<br>TIGR02203<br>TIGR00958<br>TIGR03796<br>TIGR03375 | MsbA_rel: ABC transporter, permease/ATP-binding protein<br>MsbA_lipidA: lipid A export permease/ATP-binding protein MsbA<br>3a01208: antigen peptide transporter 2<br>NHLM_micro_ABC1: NHLM bacteriocin system ABC transporter, peptidase/ATP-binding protein<br>type_I_sec_LssB: type I secretion system ATPase | 1.10E-102<br>1.00E-95<br>5.60E-84<br>8.10E-81<br>2.00E-79 |
| WP_056705456.1 | 5961 | 5151 | - | 269 | PF13649<br>PF08241<br>PF08242<br>PF13489<br>PF13847 | Methyltransf_25<br>Methyltransf_11<br>Methyltransf_12<br>Methyltransf_23<br>Methyltransf_31 | Methyltransferase domain<br>Methyltransferase domain<br>Methyltransferase domain<br>Methyltransferase domain<br>Methyltransferase domain | 2.30E-16<br>7.50E-12<br>3.30E-08<br>2.70E-07<br>1.80E-06 |
| WP_057667040.1 | 6566 | 5957 | - | 202 | LinL | LinL | LinL | 7.90E-65 |

| | | | | | | | | |
|---|---|---|---|---|---|---|---|---|
| WP_057667039.1 | 8410 | 6562 | - | 615 | LinH<br>LinG<br>LinE<br>LinL<br>PF12697 | LinH<br>LinG<br>LinE<br>LinL<br>Abhydrolase_6 | LinH (Fusion of LinE/LinG)<br>LinG Homolog<br>LinE Homolog<br>LinL<br>Alpha/beta hydrolase family | 9.40E-253<br>1.50E-111<br>1.90E-94<br>2.70E-36<br>1.80E-14 |
| WP_050362034.1 | 9158 | 8957 | - | 66 | LinA | LinA | LinA (linaridin precursor) | 3.70E-14 |
| WP_098891740.1 | 9410 | 9215 | - | 64 | LinA | LinA | LinA (linaridin precursor) | 5.10E-11 |
| WP_050362036.1 | 10152 | 9948 | - | 67 | LinA | LinA | LinA (linaridin precursor) | 8.70E-12 |
| WP_010988951.1 | 10409 | 10541 | + | 44 | INFERRED GENE | - | - | - |
| WP_007460832.1 | 10635 | 11004 | + | 123 | INFERRED GENE | - | - | - |
| NP_630878.1 | 11005 | 11495 | + | 163 | INFERRED GENE | - | - | - |
| WP_056705465.1 | 14331 | 11676 | - | 884 | PF13191<br>TIGR03903<br>PF13424<br>PF07719<br>PF13401 | AAA_16<br>TIGR03903<br>TPR_12<br>TPR_2<br>AAA_22 | AAA ATPase domain<br>TOMM_kin_cyc: TOMM system kinase/cyclase fusion protein<br>Tetratricopeptide repeat<br>Tetratricopeptide repeat<br>AAA domain | 4.20E-20<br>4.90E-19<br>4.80E-05<br>5.50E-05<br>8.70E-05 |
| WP_050362051.1 | 14441 | 14921 | + | 159 | PF03745 | DUF309 | Domain of unknown function (DUF309) | 6.30E-20 |

| index | leader | core | start | end | dir | score |
|---|---|---|---|---|---|---|
| 1 | MSMSPTPAALRGTGGLSESDPGRALSSLAPVA | ATPGVVAGVAVGVALVNAFAAGYNHCGGNVELPM | 9158 | 8957 | - | 31 |
| 2 | MNKSSAPAVLTTTGALTESAPGVTLPTLAPVA | ATPVAIAATMGVAFVAGYAAGRAATGNVELPM | 9410 | 9215 | - | 30 |
| 3 | LNTASIPAVLSETGALSESDHGRAPLN | TAPVASVTFTMTAGVQVSVCLTGSVIQLPQ | 9655 | 9481 | - | 9 |
| 4 | MTSPHRPPTSEEKSLNTASIPAVLSETGALSESDHGRAPLN | TAPVASVTFTMTAGVQVSVCLTGSVIQLPQ | 9697 | 9481 | - | 8 |
| 5 | VGPSRRRRRPPMTSPHRPPTSEEKS...TASIPAVLSETGALSESDHGRAPLN | TAPVASVTFTMTAGVQVSVCLTGSVIQLPQ | 9730 | 9481 | - | 7 |
| 6 | MPEFREPGWTRGIAPLQEDAGGQVFGGASPVA | ATPAVVATAGAVVVAFAAGVAARHLANGGNVELPM | 10152 | 9948 | - | 30 |

### Results for WP_093563556.1 [Streptomyces sp. OspMP-M43] back to top

previous - next

#### Architecture

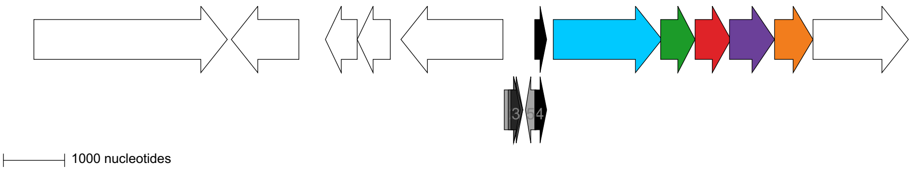

Link to nucleotide sequence

| Accession | start | end | direction | length (aa) | Pfam/HMM | name | description | E-value |
|---|---|---|---|---|---|---|---|---|
| WP_141716112.1 | 0 | 3163 | + | 1053 | TIGR01746<br>PF08659<br>TIGR03443<br>PF07993<br>PF01370 | TIGR01746<br>KR<br>TIGR03443<br>NAD_binding_4<br>Epimerase | Thioester-redct: thioester reductase domain<br>KR domain<br>alpha_am_amid: L-aminoadipate-semialdehyde dehydrogenase<br>Male sterility protein<br>NAD dependent epimerase/dehydratase family | 4.20E-86<br>1.50E-64<br>6.10E-59<br>5.90E-57<br>3.10E-21 |
| WP_012381890.1 | 4333 | 3235 | - | 365 | PF03704<br>PF13428<br>PF00486<br>PF13432<br>PF08279 | BTAD<br>TPR_14<br>Trans_reg_C<br>TPR_16<br>HTH_11 | Bacterial transcriptional activator domain<br>Tetratricopeptide repeat<br>Transcriptional regulatory protein, C terminal<br>Tetratricopeptide repeat<br>HTH domain | 7.20E-45<br>8.00E-06<br>3.30E-04<br>6.20E-04<br>6.70E-04 |
| WP_093563552.1 | 5293 | 4777 | - | 171 | NO PFAM MATCH | - | - | - |
| WP_003970684.1 | 5830 | 5296 | - | 177 | PF01613<br>TIGR02296<br>TIGR03615 | Flavin_Reduct<br>TIGR02296<br>TIGR03615 | Flavin reductase like domain<br>HpaC: 4-hydroxyphenylacetate 3-monooxygenase, reductase component<br>RutF: pyrimidine utilization flavin reductase protein F | 8.90E-33<br>8.10E-24<br>1.80E-23 |
| WP_093563553.1 | 7661 | 6011 | - | 549 | PF03704<br>PF13191<br>TIGR03903<br>PF00486 | BTAD<br>AAA_16<br>TIGR03903<br>Trans_reg_C | Bacterial transcriptional activator domain<br>AAA ATPase domain<br>TOMM_kin_cyc: TOMM system kinase/cyclase fusion protein<br>Transcriptional regulatory protein, C terminal | 4.00E-43<br>6.20E-12<br>3.90E-08<br>2.70E-04 |
| WP_003970682.1 | 8183 | 8378 | + | 64 | LinA | LinA | LinA (linaridin precursor) | 6.80E-14 |
| WP_093563556.1 | 8494 | 10252 | + | 585 | LinH<br>LinG<br>LinE<br>PF12697<br>TIGR02427 | LinH<br>LinG<br>LinE<br>Abhydrolase_6<br>TIGR02427 | LinH (Fusion of LinE/LinG)<br>LinG Homolog<br>LinE Homolog<br>Alpha/beta hydrolase family<br>protocat_pcaD: 3-oxoadipate enol-lactonase | 2.10E-175<br>1.50E-76<br>4.40E-65<br>7.30E-06<br>3.30E-05 |
| WP_003970680.1 | 10241 | 10808 | + | 188 | LinL | LinL | LinL | 1.10E-51 |
| WP_003970679.1 | 10804 | 11377 | + | 190 | TIGR00521<br>TIGR02113<br>PF02441 | TIGR00521<br>TIGR02113<br>Flavoprotein | coaBC_dfp: phosphopantothenoylcysteine decarboxylase / phosphopantothenate--cysteine ligase<br>coaC_strep: phosphopantothenoylcysteine decarboxylase<br>Flavoprotein | 2.30E-14<br>3.70E-12<br>1.50E-11 |
| WP_093563554.1 | 11369 | 12107 | + | 245 | PF13649<br>PF08241<br>PF08242<br>TIGR02021<br>TIGR01934 | Methyltransf_25<br>Methyltransf_11<br>Methyltransf_12<br>TIGR02021<br>TIGR01934 | Methyltransferase domain<br>Methyltransferase domain<br>Methyltransferase domain<br>BchM-ChlM: magnesium protoporphyrin O-methyltransferase<br>MenG_MenH_UbiE: ubiquinone/menaquinone biosynthesis methyltransferase | 9.20E-17<br>2.50E-15<br>1.70E-11<br>5.40E-11<br>2.40E-10 |

| | | | | | | | | |
|---|---|---|---|---|---|---|---|---|
| WP_003970677.1 | 12103 | 12736 | + | 210 | TIGR02857<br>TIGR01189<br>TIGR02673<br>TIGR02982<br>TIGR03258 | TIGR02857<br>TIGR01189<br>TIGR02673<br>TIGR02982<br>TIGR03258 | CydD: thiol reductant ABC exporter, CydD subunit<br>ccmA: heme ABC exporter, ATP-binding protein CcmA<br>FtsE: cell division ATP-binding protein FtsE<br>heterocyst_DevA: ABC exporter ATP-binding subunit, DevA family<br>PhnT: 2-aminoethylphosphonate ABC transport system, ATP-binding component PhnT | 5.70E-28<br>2.70E-25<br>1.60E-24<br>3.30E-22<br>1.10E-21 |
| WP_093563555.1 | 12732 | 14277 | + | 514 | NO PFAM MATCH | - | - | - |

| index | leader | core | start | end | dir | score |
|---|---|---|---|---|---|---|
| 1 | VAAYDPGFRRSGLRIPAHRAGARSKPTQNSDKERALLGGLGTCEASGAGT | PTDTGRTGGTTGRSSKPASSRNRPTGAWQLVQIQRDLRLVKLILMLRVGC | 7696 | 7999 | + | 6 |
| 2 | VQGLGPSPHKTRTRSGPCSGASAHARR | PAPGHRPTREGPGERPGVRRNRHRHEIARQARGNLSKFSGT | 7751 | 7958 | + | 6 |
| 3 | LLGGLGTCEASGAGTPTDTGRTGGTTGRSSKP | ASSRNRPTGAWQLVQIQRDLRLVKLILMLRVGC | 7801 | 7999 | + | 8 |
| 4 | MRLDSIATQETATALPESMATQDFANSVLAGAVPGFHSDAETPAM | ATPAVAQFVIQGSTICLVC | 8183 | 8378 | + | 34 |
| 5 | LQSNRVAFRCPSSEPMLSERDLAVP | SCPSPAASSELNALCTTSTTRSAGIPQSW | 8205 | 8040 | - | 6 |

### Results for SCE23008.1 [Streptomyces sp. OspMP-M43] back to top

previous - next

#### Architecture

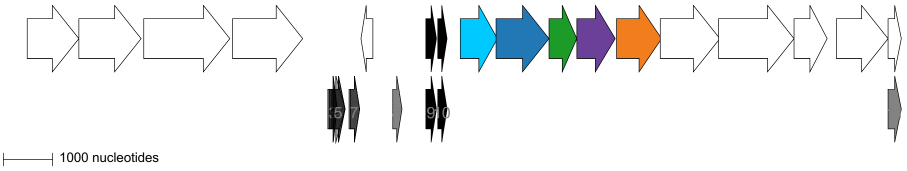

1000 nucleotides

Link to nucleotide sequence

| Accession | start | end | direction | length (aa) | Pfam/HMM | name | description | E-value |
|---|---|---|---|---|---|---|---|---|
| SCE22897.1 | 3082 | 4135 | + | 350 | TIGR03217<br>PF00682<br>PF07836<br>TIGR02660<br>TIGR02146 | TIGR03217<br>HMGL-like<br>DmpG_comm<br>TIGR02660<br>TIGR02146 | 4OH_2_O_val_ald: 4-hydroxy-2-oxovalerate aldolase<br>HMGL-like<br>DmpG-like communication domain<br>nifV_homocitr: homocitrate synthase<br>LysS_fung_arch: homocitrate synthase | 1.40E-166<br>9.00E-62<br>3.00E-29<br>9.90E-23<br>1.10E-18 |
| SCE22911.1 | 4128 | 5391 | + | 420 | TIGR01814<br>PF00266<br>TIGR01976<br>TIGR01979<br>TIGR01324 | TIGR01814<br>Aminotran_5<br>TIGR01976<br>TIGR01979<br>TIGR01324 | kynureninase: kynureninase<br>Aminotransferase class-V<br>am_tr_V_VC1184: cysteine desulfurase family protein<br>sufS: cysteine desulfurase, SufS family<br>cysta_beta_ly_B: cystathionine beta-lyase | 1.20E-76<br>2.10E-14<br>6.80E-10<br>1.40E-06<br>4.50E-06 |
| SCE22923.1 | 5436 | 7203 | + | 588 | PF08309 | LVIVD | LVIVD repeat | 1.70E-06 |
| SCE22933.1 | 7243 | 8692 | + | 482 | TIGR01773<br>PF00324<br>TIGR00913<br>PF13520<br>TIGR00909 | TIGR01773<br>AA_permease<br>TIGR00913<br>AA_permease_2<br>TIGR00909 | GABAperm: GABA permease<br>Amino acid permease<br>2A0310: amino acid permease (yeast)<br>Amino acid permease<br>2A0306: amino acid transporter | 9.90E-95<br>1.40E-88<br>1.20E-78<br>2.00E-42<br>1.00E-39 |
| SCE22948.1 | 10103 | 9863 | - | 79 | NO PFAM MATCH | - | - | - |
| SCE22961.1 | 11188 | 11386 | + | 65 | LinA | LinA | LinA (linaridin precursor) | 2.10E-08 |
| SCE22975.1 | 11418 | 11622 | + | 67 | LinA | LinA | LinA (linaridin precursor) | 5.60E-12 |
| SCE22992.1 | 11880 | 12624 | + | 247 | LinH<br>LinG | LinH<br>LinG | LinH (Fusion of LinE/LinG)<br>LinG Homolog | 1.10E-105<br>2.90E-101 |
| SCE23008.1 | 12620 | 13682 | + | 353 | LinE<br>LinH<br>PF12697<br>PF00561<br>TIGR03695 | LinE<br>LinH<br>Abhydrolase_6<br>Abhydrolase_1<br>TIGR03695 | LinE Homolog<br>LinH (Fusion of LinE/LinG)<br>Alpha/beta hydrolase family<br>alpha/beta hydrolase fold<br>menH_SHCHC: 2-succinyl-6-hydroxy-2,4-cyclohexadiene-1-carboxylate synthase | 2.80E-127<br>9.20E-114<br>6.40E-16<br>1.20E-08<br>4.40E-08 |
| SCE23021.1 | 13681 | 14248 | + | 188 | LinL | LinL | LinL | 9.20E-71 |
| SCE23035.1 | 14244 | 15033 | + | 262 | PF13649<br>PF08241<br>PF13847<br>PF13489<br>PF08242 | Methyltransf_25<br>Methyltransf_11<br>Methyltransf_31<br>Methyltransf_23<br>Methyltransf_12 | Methyltransferase domain<br>Methyltransferase domain<br>Methyltransferase domain<br>Methyltransferase domain<br>Methyltransferase domain | 5.10E-21<br>2.70E-17<br>1.00E-14<br>2.20E-12<br>6.20E-12 |
| SCE23052.1 | 15068 | 15980 | + | 303 | TIGR03522<br>TIGR01188<br>TIGR03864<br>TIGR03740<br>TIGR04406 | TIGR03522<br>TIGR01188<br>TIGR03864<br>TIGR03740<br>TIGR04406 | GldA_ABC_ATP: gliding motility-associated ABC transporter ATP-binding subunit GldA<br>drrA: daunorubicin resistance ABC transporter, ATP-binding protein<br>PQQ_ABC_ATP: ABC transporter, ATP-binding subunit, PQQ-dependent alcohol dehydrogenase system<br>galliderm_ABC: lantibiotic protection ABC transporter, ATP-binding subunit<br>LPS_export_lptB: LPS export ABC transporter ATP-binding protein | 2.00E-53<br>3.90E-46<br>2.90E-44<br>7.40E-42<br>6.70E-41 |
| SCE23068.1 | 15960 | 17151 | + | 396 | PF12698<br>PF12679 | ABC2_membrane_3<br>ABC2_membrane_2 | ABC-2 family transporter protein<br>ABC-2 family transporter protein | 9.30E-31<br>2.10E-04 |

| | | | | | | | | |
|---|---|---|---|---|---|---|---|---|
| SCE23082.1 | 17143 | 18658 | + | 504 | TIGR02734<br>TIGR03467<br>PF13450<br>TIGR00292<br>PF01593 | TIGR02734<br>TIGR03467<br>NAD_binding_8<br>TIGR00292<br>Amino_oxidase | crtI_fam: phytoene desaturase<br>HpnE: squalene-associated FAD-dependent desaturase<br>NAD(P)-binding Rossmann-like domain<br>TIGR00292: thiazole biosynthesis enzyme<br>Flavin containing amine oxidoreductase | 2.20E-11<br>1.40E-08<br>4.10E-08<br>4.60E-08<br>6.00E-07 |
| SCE23099.1 | 18674 | 19337 | + | 220 | PF05719 | GPP34 | Golgi phosphoprotein 3 (GPP34) | 5.20E-49 |
| SCE23116.1 | 19526 | 20570 | + | 347 | TIGR00747<br>PF08541 | TIGR00747<br>ACP_syn_III_C | fabH: 3-oxoacyl-[acyl-carrier-protein] synthase III<br>3-Oxoacyl-[acyl-carrier-protein (ACP)] synthase III C terminal | 5.30E-08<br>1.10E-04 |
| SCE23130.1 | 20566 | 20848 | + | 93 | PF00550<br>TIGR01688 | PP-binding<br>TIGR01688 | Phosphopantetheine attachment site<br>dltC: D-alanine--poly(phosphoribitol) ligase, subunit 2 | 7.60E-09<br>1.90E-05 |

| index | leader | core | start | end | dir | score |
|---|---|---|---|---|---|---|
| 1 | LPEPTDRVRLCGLRLLGGQACAAQE...LHAPSLLSMPRASSSPRTRVHCARR | YTPPPGLKATASMRSTRPGAARWFE...VAPWAAGSKLLPSDRRAHGVRAVPA | 9191 | 9539 | + | 6 |
| 2 | VFVSVGSGCSADRRAQRRKPLALWHFTP | PRCSVCRARAAAREPESTARGGTPRPQD | 9210 | 9381 | + | 7 |
| 3 | VGSGCSADRRAQRRKPLALWHFTPPR | CSVCRARAAAREPESTARGGTPRPQD | 9222 | 9381 | + | 7 |
| 4 | MALHAPSLLSMPRASSSPRTRVHCARR | YTPPPGLKATASMRSTRPGAARWFE...VAPWAAGSKLLPSDRRAHGVRAVPA | 9278 | 9539 | + | 7 |
| 5 | MPRASSSPRTRVHCARR | YTPPPGLKATASMRSTRPGAARWFE...VAPWAAGSKLLPSDRRAHGVRAVPA | 9308 | 9539 | + | 7 |
| 6 | MSAYTPSMGEVIGVVGRRAHCLTCERPHEDAAMPEGVMCSNCART | VVPGPCQRCSARCPTARSFGTWTR | 9632 | 9842 | + | 7 |
| 7 | MGEVIGVVGRRAHCLTCERPHEDAAMPEGVMCSNCART | VVPGPCQRCSARCPTARSFGTWTR | 9653 | 9842 | + | 7 |
| 8 | LPCRPPDSPRDHLEALGDNQTQSQCERIY | CASLRNGRARQVSIFQHRNFRTLSAARLF | 10511 | 10688 | + | 7 |
| 9 | MSTLLQKVNTSQELHAVASDLADAKDLSLTLPVL | ATITTVCPATGYIVGGGLFATGIGITYTVNR | 11188 | 11386 | + | 29 |
| 10 | MSVVLDKVNASVEFGAIPGHISEAEWGNATDVPVL | ATPVTLTPTIGKVTLGVGAFTGGIAVGDALFG | 11418 | 11622 | + | 30 |
| 11 | MSRGPAETLDEITRFLTEALRREVGPDDDYFALGLVDSLFALELVT | YVEDRFELTVEVEDLDLDSFRTAGRLTAFVHAKKAGAETTPAHASDN | 20566 | 20848 | + | 7 |

### Results for WP_113686667.1 [Streptomyces sp. PT12] back to top

previous - next

#### Architecture

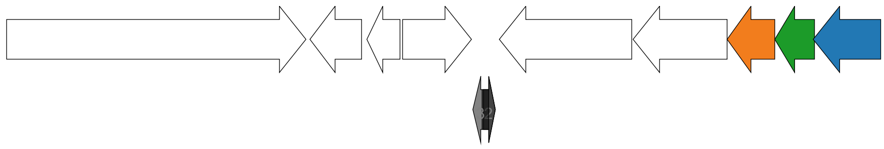

1000 nucleotides

Link to nucleotide sequence

| Accession | start | end | direction | length (aa) | Pfam/HMM | name | description | E-value |
|---|---|---|---|---|---|---|---|---|
| WP_113686660.1 | 27787 | 31885 | + | 1365 | PF13472<br>PF00657 | Lipase_GDSL_2<br>Lipase_GDSL | GDSL-like Lipase/Acylhydrolase family<br>GDSL-like Lipase/Acylhydrolase | 1.70E-12<br>1.30E-06 |
| WP_147255509.1 | 32649 | 31926 | - | 240 | NO PFAM MATCH | - | - | - |
| WP_113686672.1 | 33160 | 32707 | - | 150 | PF09424 | YqeY | Yqey-like protein | 3.60E-44 |
| WP_113686662.1 | 33202 | 34144 | + | 313 | PF12850<br>PF00149 | Metallophos_2<br>Metallophos | Calcineurin-like phosphoesterase superfamily domain<br>Calcineurin-like phosphoesterase | 5.30E-08<br>1.60E-06 |
| WP_113686663.1 | 36346 | 34531 | - | 604 | PF13738<br>TIGR04046<br>PF07992<br>PF00743<br>TIGR04018 | Pyr_redox_3<br>TIGR04046<br>Pyr_redox_2<br>FMO-like<br>TIGR04018 | Pyridine nucleotide-disulphide oxidoreductase<br>MSMEG_0569_nitr: flavin-dependent oxidoreductase, MSMEG_0569 family<br>Pyridine nucleotide-disulphide oxidoreductase<br>Flavin-binding monooxygenase-like<br>Bthiol_YpdA: putative bacillithiol system oxidoreductase, YpdA family | 1.80E-21<br>2.80E-21<br>8.40E-18<br>1.60E-15<br>3.70E-13 |
| WP_113686664.1 | 37640 | 36359 | - | 426 | NO PFAM MATCH | - | - | - |
| WP_113686665.1 | 38290 | 37636 | - | 217 | TIGR02857<br>TIGR01189<br>TIGR02868<br>TIGR03608<br>TIGR01166 | TIGR02857<br>TIGR01189<br>TIGR02868<br>TIGR03608<br>TIGR01166 | CydD: thiol reductant ABC exporter, CydD subunit<br>ccmA: heme ABC exporter, ATP-binding protein CcmA<br>CydC: thiol reductant ABC exporter, CydC subunit<br>L_ocin_972_ABC: putative bacteriocin export ABC transporter, lactococcin 972 group<br>cbiO: cobalt ABC transporter, ATP-binding protein | 3.40E-33<br>2.50E-30<br>7.90E-30<br>1.30E-27<br>6.30E-27 |
| WP_147255511.1 | 38832 | 38286 | - | 181 | LinL | LinL | LinL | 3.10E-44 |
| WP_113686667.1 | 39746 | 38828 | - | 305 | LinE<br>LinH<br>TIGR02427<br>TIGR03611 | LinE<br>LinH<br>TIGR02427<br>TIGR03611 | LinE Homolog<br>LinH (Fusion of LinE/LinG)<br>protocat_pcaD: 3-oxoadipate enol-lactonase<br>RutD: pyrimidine utilization protein D | 5.00E-74<br>6.60E-44<br>1.70E-06<br>2.40E-04 |

| index | leader | core | start | end | dir | score |
|---|---|---|---|---|---|---|
| 1 | VGRTLPVSPLIVWEKPGRRSWSGSPGPGTRPRAYANLAAAL | DTAEAVISGAVTTAESGT | 34297 | 34477 | + | 7 |
| 2 | VSPLIVWEKPGRRSWSGSPGPGTRPRAYANLAAAL | DTAEAVISGAVTTAESGT | 34315 | 34477 | + | 7 |
| 3 | VPGPGDPDQDLLPGFSQTISGDTGRVRPTLRIGPL | TCYLSVGVAGFEPTTSSSRTKRAAKLRHTPVQRSTL | 34383 | 34167 | - | 7 |

### Results for EXU61863.1 [Streptomyces sp. PRh5] back to top

### Architecture

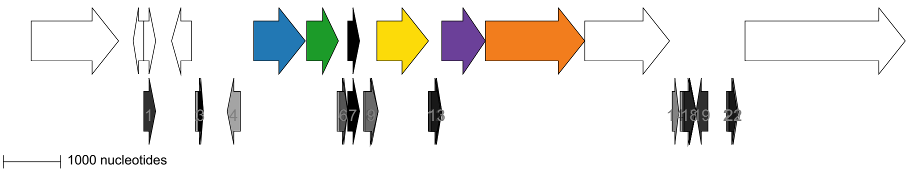

Link to nucleotide sequence

| Accession | start | end | direction | length (aa) | Pfam/HMM | name | description | E-value |
|---|---|---|---|---|---|---|---|---|
| EXU61861.1 | 635 | 2171 | + | 511 | PF00135 | COesterase | Carboxylesterase family | 3.70E-109 |
| | | | | | PF07859 | Abhydrolase_3 | alpha/beta hydrolase fold | 2.40E-05 |
| EXU61862.1 | 2599 | 2413 | - | 61 | NO PFAM MATCH | - | - | - |
| EXU61870.1 | 2611 | 2812 | + | 66 | NO PFAM MATCH | - | - | - |
| EXU61871.1 | 3441 | 3075 | - | 121 | PF01548 | DEDD_Tnp_IS110 | Transposase | 1.40E-27 |
| EXU61863.1 | 4509 | 5433 | + | 307 | LinE | LinE | LinE Homolog | 7.10E-92 |
| | | | | | LinH | LinH | LinH (Fusion of LinE/LinG) | 2.70E-41 |
| | | | | | PF12697 | Abhydrolase_6 | Alpha/beta hydrolase family | 1.80E-17 |
| | | | | | PF00561 | Abhydrolase_1 | alpha/beta hydrolase fold | 3.90E-09 |
| | | | | | TIGR02427 | TIGR02427 | protocat_pcaD: 3-oxoadipate enol-lactonase | 6.80E-08 |
| EXU61864.1 | 5447 | 5993 | + | 181 | LinL | LinL | LinL | 1.10E-51 |
| EXU61872.1 | 6153 | 6381 | + | 75 | LinA | LinA | LinA (linaridin precursor) | 2.00E-11 |
| EXU61865.1 | 6679 | 7573 | + | 297 | LinG | LinG | LinG Homolog | 2.20E-102 |
| | | | | | LinH | LinH | LinH (Fusion of LinE/LinG) | 7.70E-78 |
| EXU61866.1 | 7808 | 8564 | + | 251 | PF13649 | Methyltransf_25 | Methyltransferase domain | 2.10E-13 |
| | | | | | PF08241 | Methyltransf_11 | Methyltransferase domain | 6.30E-09 |
| | | | | | PF13489 | Methyltransf_23 | Methyltransferase domain | 1.40E-08 |
| | | | | | PF08242 | Methyltransf_12 | Methyltransferase domain | 1.60E-08 |
| | | | | | TIGR01983 | TIGR01983 | UbiG: 3-demethylubiquinone-9 3-O-methyltransferase | 3.10E-06 |
| EXU61867.1 | 8560 | 10303 | + | 580 | TIGR02204 | TIGR02204 | MsbA_rel: ABC transporter, permease/ATP-binding protein | 1.30E-100 |
| | | | | | TIGR02203 | TIGR02203 | MsbA_lipidA: lipid A export permease/ATP-binding protein MsbA | 6.50E-94 |
| | | | | | TIGR03796 | TIGR03796 | NHLM_micro_ABC1: NHLM bacteriocin system ABC transporter, peptidase/ATP-binding protein | 2.40E-84 |
| | | | | | TIGR00958 | TIGR00958 | 3a01208: antigen peptide transporter 2 | 2.30E-83 |
| | | | | | TIGR01846 | TIGR01846 | type_I_sec_HlyB: type I secretion system ATPase | 3.90E-78 |
| EXU61868.1 | 10299 | 11784 | + | 494 | TIGR02734 | TIGR02734 | crtI_fam: phytoene desaturase | 1.60E-09 |
| | | | | | TIGR00292 | TIGR00292 | TIGR00292: thiazole biosynthesis enzyme | 1.90E-08 |
| | | | | | TIGR02733 | TIGR02733 | desat_CrtD: C-3',4' desaturase CrtD | 3.90E-08 |
| | | | | | TIGR03467 | TIGR03467 | HpnE: squalene-associated FAD-dependent desaturase | 4.70E-08 |
| | | | | | TIGR01317 | TIGR01317 | GOGAT_sm_gam: glutamate synthase, NADH/NADPH, small subunit | 1.20E-07 |
| EXU61869.1 | 13109 | 15908 | + | 932 | PF03704 | BTAD | Bacterial transcriptional activator domain | 6.80E-39 |
| | | | | | PF07719 | TPR_2 | Tetratricopeptide repeat | 8.50E-20 |
| | | | | | PF13424 | TPR_12 | Tetratricopeptide repeat | 7.30E-18 |
| | | | | | PF00515 | TPR_1 | Tetratricopeptide repeat | 6.50E-12 |
| | | | | | PF00931 | NB-ARC | NB-ARC domain | 1.60E-10 |

| index | leader | core | start | end | dir | score |
|---|---|---|---|---|---|---|
| 1 | MPVSASGCVFAVDREVPEDFVVGARPAQEARCA | AVGKLVPSPTVTSKVAAVLTAIPGIEVRTLKRG | 2611 | 2812 | + | 9 |
| 2 | LLVEDFLAGRGEAVAVMGALADV | QAEEDARLVAVVHVVLPNGCRAC | 3510 | 3651 | + | 6 |
| 3 | VMGALADVQAEEDAR | LVAVVHVVLPNGCRAC | 3555 | 3651 | + | 14 |
| 4 | LKQQLKECFLRPGHRVMEQDPADGEVGRHWEEHGPWL | RPPILIDRRLLDLSVAVVVPMLRGARRTGVSIE | 4280 | 4067 | - | 6 |
| 5 | MVDRIGRESRPHAAVRMCEGGTPGRSGRH | ASLGSGHCSVGKTTKRSVTKIMMKGTRCLS | 5986 | 6166 | + | 6 |
| 6 | MCEGGTPGRSGRHASLGSGHC | SVGKTTKRSVTKIMMKGTRCLS | 6034 | 6166 | + | 7 |
| 7 | MSLVAQFANEALTDISPGMLSETDAASRLDGPADPVM | VTPFAALTVTCPEVAALAGAAFGALAANVANYVGRNAH | 6153 | 6381 | + | 31 |
| 8 | VTPSAPSPPPTMQHGCPEVTAEFCPVTPEPVGPTGRQLAQVPAL | SPPRRGADEGDRVFGARRNRTASSACPAPHSHQKEISCSVV | 6437 | 6695 | + | 6 |
| 9 | MQHGCPEVTAEFCPVTPEPVGPTGRQLAQVPAL | SPPRRGADEGDRVFGARRNRTASSACPAPHSHQKEISCSVV | 6470 | 6695 | + | 6 |
| 10 | LSRVDARCAHHRRRGLAMPAAAQHRPHLSRCWDDRDW | TCAGHSADRRAAAMPRCRGLWAAQDREALAARVLLRLT | 7569 | 7797 | + | 6 |
| 11 | VDARCAHHRRRGLAMPAAAQHRPHLSRCWDDRDWTC | AGHSADRRAAAMPRCRGLWAAQDREALAARVLLRLT | 7578 | 7797 | + | 6 |
| 12 | VRIIADEASQCLRRRSIARTSRGAGTIGTGPVP | GTARIAVRRPCRGAAACGPPKTARLSPRGCYFD | 7591 | 7792 | + | 6 |
| 13 | MPAAAQHRPHLSRCWDDRDWTCAGHSADR | RAAAMPRCRGLWAAQDREALAARVLLRLT | 7620 | 7797 | + | 8 |
| 14 | MPPLEPSHGADLGDGGPEA | GKKFSAGHASDVPARRRTT | 11843 | 11960 | + | 7 |
| 15 | VDRPANPASSPAER | MTLGQRPETSMTIPDGGSEASVSNI...LVAKVLGDVVRSPLRPLAALGSRGR | 11973 | 12240 | + | 6 |
| 16 | MTLGQRPETSMTIPDGGSEASVSN | IAPVLVGRGEVVQYGRVPSPSGGTGLVAKVLGDVVRSPLRPLAALGSRGR | 12015 | 12240 | + | 6 |
| 17 | LGQRPETSMTIPDGGSEASVSN | IAPVLVGRGEVVQYGRVPSPSGGTGLVAKVLGDVVRSPLRPLAALGSRGR | 12021 | 12240 | + | 6 |
| 18 | MTIPDGGSEASVSN | IAPVLVGRGEVVQYGRVPSPSGGTGLVAKVLGDVVRSPLRPLAALGSRGR | 12045 | 12240 | + | 7 |
| 19 | VAATSTPASPRHRLDGDGRNRLGGPLAEYMQLTDQ | VTTTGTIGRACVTRGLVQVRVLLPSHQTSEQFVQA | 12464 | 12251 | - | 9 |

| | | | | | | |
|---|---|---|---|---|---|---|
| 20 | VPLRIGQPLGLQCLVRLGSDLDLLGEPGVDR | DPDRMHTARHPSPPKATLSQGSVHPSGRTTA | 12773 | 12962 | + | 7 |
| 21 | LQCLVRLGSDLDLLGEPGVDRDPDRM | HTARHPSPPKATLSQGSVHPSGRTTA | 12803 | 12962 | + | 8 |
| 22 | LTCWASPASTAIQTACTPRDIRHRQRR | RYRKVRYTLAAARRLDGPAIEAMLVMR | 12834 | 12999 | + | 7 |

### Results for RBM10751.1 [Streptomyces sp. PT12] back to top

previous - next

#### Architecture

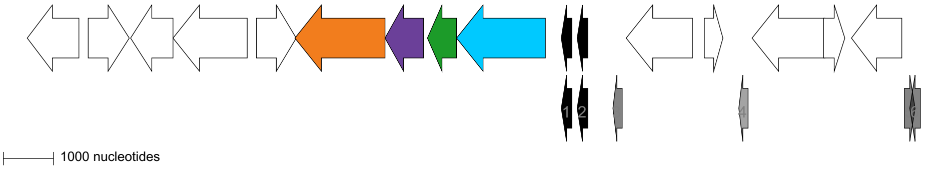

Link to nucleotide sequence

| Accession | start | end | direction | length (aa) | Pfam/HMM | name | description | E-value |
|---|---|---|---|---|---|---|---|---|
| RBM10765.1 | 28769 | 27752 | - | 338 | TIGR00747<br>PF08541 | TIGR00747<br>ACP_syn_III_C | fabH: 3-oxoacyl-[acyl-carrier-protein] synthase III<br>3-Oxoacyl-[acyl-carrier-protein (ACP)] synthase III C terminal | 7.40E-08<br>8.00E-04 |
| RBM10766.1 | 28954 | 29782 | + | 275 | PF01551 | Peptidase_M23 | Peptidase family M23 | 2.40E-16 |
| RBM10746.1 | 30656 | 29804 | - | 283 | PF12680 | SnoaL_2 | SnoaL-like domain | 6.10E-04 |
| RBM10747.1 | 32137 | 30652 | - | 494 | TIGR00292<br>TIGR02734<br>TIGR02733<br>TIGR01317<br>PF01946 | TIGR00292<br>TIGR02734<br>TIGR02733<br>TIGR01317<br>Thi4 | TIGR00292: thiazole biosynthesis enzyme<br>crtI_fam: phytoene desaturase<br>desat_CrtD: C-3',4' desaturase CrtD<br>GOGAT_sm_gam: glutamate synthase, NADH/NADPH, small subunit<br>Thi4 family | 8.60E-09<br>8.00E-08<br>1.10E-07<br>5.60E-07<br>5.70E-07 |
| RBM10748.1 | 32298 | 33093 | + | 264 | PF07282<br>PF01385<br>PF12323<br>TIGR01766 | OrfB_Zn_ribbon<br>OrfB_IS605<br>HTH_OrfB_IS605<br>TIGR01766 | Putative transposase DNA-binding domain<br>Probable transposase<br>Helix-turn-helix domain<br>tspaseT_teng_C: transposase, IS605 OrfB family | 8.10E-22<br>1.50E-20<br>1.30E-18<br>2.10E-17 |
| RBM10767.1 | 34861 | 33067 | - | 597 | TIGR02204<br>TIGR02203<br>TIGR00958<br>TIGR03796<br>TIGR01846 | TIGR02204<br>TIGR02203<br>TIGR00958<br>TIGR03796<br>TIGR01846 | MsbA_rel: ABC transporter, permease/ATP-binding protein<br>MsbA_lipidA: lipid A export permease/ATP-binding protein MsbA<br>3a01208: antigen peptide transporter 2<br>NHLM_micro_ABC1: NHLM bacteriocin system ABC transporter, peptidase/ATP-binding protein<br>type_I_sec_HlyB: type I secretion system ATPase | 3.00E-98<br>1.20E-93<br>1.50E-86<br>4.60E-83<br>5.20E-79 |
| RBM10749.1 | 35652 | 34881 | - | 256 | PF13649<br>PF13489<br>PF08241<br>PF08242<br>PF13847 | Methyltransf_25<br>Methyltransf_23<br>Methyltransf_11<br>Methyltransf_12<br>Methyltransf_31 | Methyltransferase domain<br>Methyltransferase domain<br>Methyltransferase domain<br>Methyltransferase domain<br>Methyltransferase domain | 2.00E-13<br>6.70E-09<br>1.20E-08<br>8.60E-08<br>1.30E-06 |
| RBM10750.1 | 36307 | 35728 | - | 192 | LinL | LinL | LinL | 5.40E-62 |
| RBM10751.1 | 38061 | 36303 | - | 585 | LinH<br>LinE<br>LinG<br>PF12697<br>PF00561 | LinH<br>LinE<br>LinG<br>Abhydrolase_6<br>Abhydrolase_1 | LinH (Fusion of LinE/LinG)<br>LinE Homolog<br>LinG Homolog<br>Alpha/beta hydrolase family<br>alpha/beta hydrolase fold | 4.80E-216<br>2.30E-95<br>3.70E-91<br>3.60E-15<br>4.80E-11 |
| RBM10752.1 | 38585 | 38393 | - | 63 | LinA | LinA | LinA (linaridin precursor) | 5.30E-09 |
| RBM10753.1 | 38909 | 38714 | - | 64 | LinA | LinA | LinA (linaridin precursor) | 9.60E-17 |
| RBM10768.1 | 40994 | 39692 | - | 433 | PF05139 | Erythro_esteras | Erythromycin esterase | 5.00E-106 |
| RBM10754.1 | 41230 | 41611 | + | 126 | PF11338 | DUF3140 | Protein of unknown function (DUF3140) | 6.00E-25 |
| RBM10755.1 | 43637 | 42179 | - | 485 | TIGR03150<br>PF00109<br>PF02801<br>TIGR02813<br>TIGR01930 | TIGR03150<br>ketoacyl-synt<br>Ketoacyl-synt_C<br>TIGR02813<br>TIGR01930 | fabF: beta-ketoacyl-acyl-carrier-protein synthase II<br>Beta-ketoacyl synthase, N-terminal domain<br>Beta-ketoacyl synthase, C-terminal domain<br>omega_3_PfaA: polyketide-type polyunsaturated fatty acid synthase PfaA<br>AcCoA-C-Actrans: acetyl-CoA C-acyltransferase | 6.70E-122<br>1.70E-48<br>5.70E-32<br>2.60E-30<br>9.70E-09 |
| RBM10769.1 | 43616 | 44048 | + | 143 | PF04075<br>TIGR00026 | F420H2_quin_red<br>TIGR00026 | F420H(2)-dependent quinone reductase<br>hi_GC_TIGR00026: deazaflavin-dependent oxidoreductase, nitroreductase family | 1.20E-50<br>1.10E-35 |
| RBM10756.1 | 45186 | 44166 | - | 339 | NO PFAM MATCH | - | - | - |

| index | leader | core | start | end | dir | score |
|---|---|---|---|---|---|---|
| 1 | MSKIIESVVAVAPEELSDQTTGEFGAVMPVQ | ATPGLVAFGVGFAGGAGVAWLGVQAYEAGIND | 38585 | 38393 | - | 31 |
| 2 | MEQKIASFMNEQQLGPMEQGYQDDVKSPVL | ATPAAGVATCWYVGGAVAGAASVVGAYVCGAVFG | 38909 | 38714 | - | 30 |
| 3 | LRDRLEDPDPGAYAGNLQRKSRESARKPFAG | QLPRAARPQSPACRATRATGEIRRAK | 39593 | 39419 | - | 7 |
| 4 | LWVVEGRPPARCAAGVRRTVGR | FTVVGLRPGEGCGRSGSLRAVAPGCRRSPNGGVARGR | 42104 | 41924 | - | 6 |
| 5 | MLNRHRSHQRHPHPPDHPNGTIGSVQRFPGAGPTPRPGVQAPWSCPCPSV | VTVNSAVRTTPSCWKSRKRR | 45229 | 45442 | + | 7 |
| 6 | VLRSGDLAYLHVHPGELAGGPDITFHT | RAPSAGTYRLFLDFQHEGVVRTAEFTVTTDGHGHDHGA | 45543 | 45345 | - | 7 |

### Results for RBM24328.1 [Streptomyces sp. PT12] back to top

previous - next

### Architecture

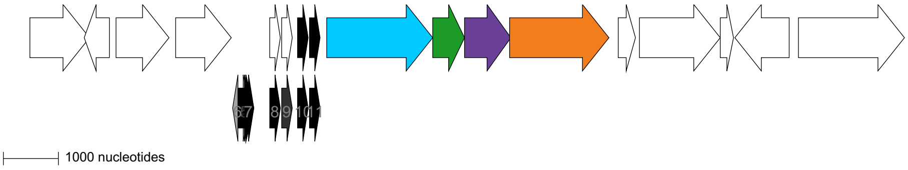

Link to nucleotide sequence

| Accession | start | end | direction | length (aa) | Pfam/HMM | name | description | E-value |
|---|---|---|---|---|---|---|---|---|
| RBM24320.1 | 26640 | 27717 | + | 358 | PF01435 | Peptidase_M48 | Peptidase family M48 | 7.20E-09 |
| RBM24321.1 | 28072 | 27616 | - | 151 | PF05099<br>PF04391 | TerB<br>DUF533 | Tellurite resistance protein TerB<br>Protein of unknown function (DUF533) | 6.00E-31<br>8.40E-05 |
| RBM24322.1 | 28209 | 29166 | + | 318 | PF12833<br>PF01965<br>PF00165<br>TIGR04094<br>TIGR01383 | HTH_18<br>DJ-1_PfpI<br>HTH_AraC<br>TIGR04094<br>TIGR01383 | Helix-turn-helix domain<br>DJ-1/PfpI family<br>Bacterial regulatory helix-turn-helix proteins, AraC family<br>adjacent_YSIRK: YSIRK-targeted surface antigen transcriptional regulator<br>not_thiJ: DJ-1 family protein | 3.10E-21<br>1.10E-18<br>1.10E-14<br>6.00E-08<br>9.80E-05 |
| RBM24323.1 | 29286 | 30285 | + | 332 | TIGR01534<br>TIGR01532<br>PF02800<br>PF00044 | TIGR01534<br>TIGR01532<br>Gp_dh_C<br>Gp_dh_N | GAPDH-I: glyceraldehyde-3-phosphate dehydrogenase, type I<br>E4PD_g-proteo: erythrose-4-phosphate dehydrogenase<br>Glyceraldehyde 3-phosphate dehydrogenase, C-terminal domain<br>Glyceraldehyde 3-phosphate dehydrogenase, NAD binding domain | 3.60E-122<br>1.30E-94<br>9.10E-60<br>1.70E-34 |
| RBM24324.1 | 31007 | 31187 | + | 59 | NO PFAM MATCH | - | - | - |
| RBM24325.1 | 31213 | 31405 | + | 63 | NO PFAM MATCH | - | - | - |
| RBM24326.1 | 31491 | 31683 | + | 63 | LinA | LinA | LinA (linaridin precursor) | 6.00E-09 |
| RBM24327.1 | 31718 | 31919 | + | 66 | LinA | LinA | LinA (linaridin precursor) | 4.90E-08 |
| RBM24328.1 | 32020 | 33964 | + | 647 | LinH<br>LinG<br>LinE<br>PF12697<br>PF00561 | LinH<br>LinG<br>LinE<br>Abhydrolase_6<br>Abhydrolase_1 | LinH (Fusion of LinE/LinG)<br>LinG Homolog<br>LinE Homolog<br>Alpha/beta hydrolase family<br>alpha/beta hydrolase fold | 2.10E-239<br>3.10E-114<br>2.90E-93<br>3.40E-16<br>1.00E-09 |
| RBM24329.1 | 33960 | 34545 | + | 194 | LinL | LinL | LinL | 5.40E-64 |
| RBM24330.1 | 34541 | 35372 | + | 276 | PF13649<br>PF08241<br>PF13489<br>PF08242<br>PF13847 | Methyltransf_25<br>Methyltransf_11<br>Methyltransf_23<br>Methyltransf_12<br>Methyltransf_31 | Methyltransferase domain<br>Methyltransferase domain<br>Methyltransferase domain<br>Methyltransferase domain<br>Methyltransferase domain | 7.50E-17<br>9.80E-12<br>6.70E-11<br>2.00E-08<br>7.80E-08 |
| RBM24331.1 | 35358 | 37161 | + | 600 | TIGR02204<br>TIGR02203<br>TIGR01846<br>TIGR00958<br>TIGR03796 | TIGR02204<br>TIGR02203<br>TIGR01846<br>TIGR00958<br>TIGR03796 | MsbA_rel: ABC transporter, permease/ATP-binding protein<br>MsbA_lipidA: lipid A export permease/ATP-binding protein MsbA<br>type_I_sec_HlyB: type I secretion system ATPase<br>3a01208: antigen peptide transporter 2<br>NHLM_micro_ABC1: NHLM bacteriocin system ABC transporter, peptidase/ATP-binding protein | 5.30E-102<br>7.00E-96<br>5.40E-82<br>5.60E-82<br>1.40E-80 |
| RBM24332.1 | 37331 | 37640 | + | 102 | NO PFAM MATCH | - | - | - |
| RBM24333.1 | 37722 | 39189 | + | 488 | TIGR02734<br>TIGR03467<br>TIGR01317<br>PF13450<br>TIGR00292 | TIGR02734<br>TIGR03467<br>TIGR01317<br>NAD_binding_8<br>TIGR00292 | crtI_fam: phytoene desaturase<br>HpnE: squalene-associated FAD-dependent desaturase<br>GOGAT_sm_gam: glutamate synthase, NADH/NADPH, small subunit<br>NAD(P)-binding Rossmann-like domain<br>TIGR00292: thiazole biosynthesis enzyme | 7.90E-12<br>7.50E-09<br>5.80E-08<br>6.70E-08<br>2.10E-07 |
| RBM24334.1 | 39188 | 39431 | + | 80 | NO PFAM MATCH | - | - | - |
| RBM24379.1 | 40438 | 39457 | - | 326 | NO PFAM MATCH | - | - | - |
| RBM24335.1 | 40621 | 42544 | + | 640 | NO PFAM MATCH | - | - | - |

| index | leader | core | start | end | dir | score |
|---|---|---|---|---|---|---|
| 1 | LLPGGCSLVPARRAPRRAAGRHRGG...IPQLDLGPLDEHTEGRVTRGALPVA | ATPVIVGAAGAIVVAYAAGRVVGEIRPT | 30407 | 30710 | + | 20 |
| 2 | LCQPGGPRGGPRAAIVVGTGRQERSAFPHPRNPQ | GGHPCAFPSWTSARSTSTPKAGSPEARCRSRPPR | 30429 | 30636 | + | 6 |
| 3 | VPARRAPRRAAGRHRGGHGTAGTFR...IPQLDLGPLDEHTEGRVTRGALPVA | ATPVIVGAAGAIVVAYAAGRVVGEIRPT | 30431 | 30710 | + | 19 |
| 4 | VVGTGRQERSAFPHPRNPQGGHPCAF | PSWTSARSTSTPKAGSPEARCRSRPPR | 30474 | 30636 | + | 7 |
| 5 | VGTGRQERSAFPHPRNPQGGHPCAFP | SWTSARSTSTPKAGSPEARCRSRPPR | 30477 | 30636 | + | 7 |
| 6 | LRGWGNAERSCRPVP | TTMAARGPPRGPPGWHKAATPRQQR...TRDATRGTTRSARVPSIRTRALRAW | 30524 | 30323 | - | 6 |
| 7 | VRIPQLDLGPLDEHTEGRVTRGALPVA | ATPVIVGAAGAIVVAYAAGRVVGEIRPT | 30542 | 30710 | + | 30 |
| 8 | MSVAQLPADLAAMPGLREDDAGRPPASAMPVTHVLTSGQAAC | VTVAVAIAVGDTRPVLK | 31007 | 31187 | + | 10 |
| 9 | MSFARLPADLAAMPGLREDDAGLPPST | LVPVTSTSCAAASAVSAVVRAAAWVADAIGPVRPSA | 31213 | 31405 | + | 9 |
| 10 | MSVSQLPADLAAVSELKVGAAGQVKAAAQPVM | ATPTLAVAFVAGIGVGLAACQANGTTRPVAE | 31491 | 31683 | + | 31 |
| 11 | MNDQSRRLYDSPALASVAVDDARGWISARQKPVL | ATPSVLAGIAIGYAMLNAFEAGRSGGPIRPTS | 31718 | 31919 | + | 17 |

### Results for WP_129844105.1 [Streptomyces sp. RFCAC02] back to top

previous - next

#### Architecture

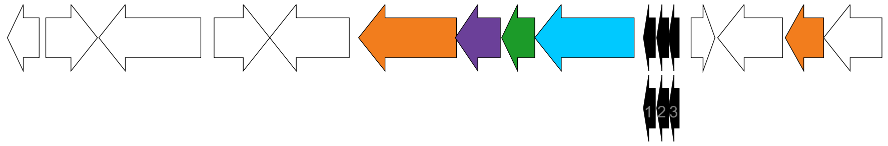

1000 nucleotides

Link to nucleotide sequence

| Accession | start | end | direction | length (aa) | Pfam/HMM | name | description | E-value |
|---|---|---|---|---|---|---|---|---|
| WP_129842020.1 | 3849151 | 3848599 | - | 183 | NO PFAM MATCH | - | - | - |
| WP_129842021.1 | 3849268 | 3850213 | + | 314 | PF02156 | Glyco_hydro_26 | Glycosyl hydrolase family 26 | 1.70E-54 |
| WP_129844103.1 | 3852059 | 3850229 | - | 609 | NO PFAM MATCH | - | - | - |
| WP_129842022.1 | 3852297 | 3853299 | + | 333 | NO PFAM MATCH | - | - | - |
| WP_129844104.1 | 3854744 | 3853304 | - | 479 | TIGR02734 | TIGR02734 | crtI_fam: phytoene desaturase | 4.90E-13 |
| | | | | | PF01593 | Amino_oxidase | Flavin containing amine oxidoreductase | 3.70E-09 |
| | | | | | TIGR01317 | TIGR01317 | GOGAT_sm_gam: glutamate synthase, NADH/NADPH, small subunit | 2.20E-08 |
| | | | | | TIGR03467 | TIGR03467 | HpnE: squalene-associated FAD-dependent desaturase | 2.80E-08 |
| | | | | | TIGR00292 | TIGR00292 | TIGR00292: thiazole biosynthesis enzyme | 1.40E-07 |
| WP_129842023.1 | 3856664 | 3854897 | - | 588 | TIGR02204 | TIGR02204 | MsbA_rel: ABC transporter, permease/ATP-binding protein | 1.10E-99 |
| | | | | | TIGR02203 | TIGR02203 | MsbA_lipidA: lipid A export permease/ATP-binding protein MsbA | 7.10E-94 |
| | | | | | TIGR03796 | TIGR03796 | NHLM_micro_ABC1: NHLM bacteriocin system ABC transporter, peptidase/ATP-binding protein | 4.80E-83 |
| | | | | | TIGR00958 | TIGR00958 | 3a01208: antigen peptide transporter 2 | 1.60E-82 |
| | | | | | TIGR03797 | TIGR03797 | NHLM_micro_ABC2: NHLM bacteriocin system ABC transporter, ATP-binding protein | 4.90E-80 |
| WP_129842024.1 | 3857469 | 3856650 | - | 272 | PF13649 | Methyltransf_25 | Methyltransferase domain | 8.70E-16 |
| | | | | | PF08241 | Methyltransf_11 | Methyltransferase domain | 2.90E-10 |
| | | | | | PF13489 | Methyltransf_23 | Methyltransferase domain | 1.90E-09 |
| | | | | | PF08242 | Methyltransf_12 | Methyltransferase domain | 3.90E-08 |
| | | | | | TIGR01983 | TIGR01983 | UbiG: 3-demethylubiquinone-9 3-O-methyltransferase | 5.60E-07 |
| WP_129842025.1 | 3858068 | 3857483 | - | 194 | LinL | LinL | LinL | 5.70E-65 |
| WP_129844105.1 | 3859879 | 3858073 | - | 601 | LinH | LinH | LinH (Fusion of LinE/LinG) | 7.10E-235 |
| | | | | | LinG | LinG | LinG Homolog | 5.10E-111 |
| | | | | | LinE | LinE | LinE Homolog | 1.40E-88 |
| | | | | | PF12697 | Abhydrolase_6 | Alpha/beta hydrolase family | 6.60E-15 |
| | | | | | PF00561 | Abhydrolase_1 | alpha/beta hydrolase fold | 1.40E-08 |
| WP_129842026.1 | 3860244 | 3860043 | - | 66 | LinA | LinA | LinA (linaridin precursor) | 1.90E-10 |
| WP_129842027.1 | 3860477 | 3860282 | - | 64 | LinA | LinA | LinA (linaridin precursor) | 2.40E-08 |
| WP_129842028.1 | 3860684 | 3860495 | - | 62 | LinA | LinA | LinA (linaridin precursor) | 2.00E-10 |
| WP_129842029.1 | 3860899 | 3861334 | + | 144 | NO PFAM MATCH | - | - | - |
| WP_129844106.1 | 3862544 | 3861368 | - | 391 | TIGR03434 | TIGR03434 | ADOP: acidobacterial duplicated orphan permease | 1.50E-21 |
| | | | | | PF12704 | MacB_PCD | MacB-like periplasmic core domain | 2.70E-17 |
| | | | | | TIGR02212 | TIGR02212 | lolCE: lipoprotein releasing system, transmembrane protein, LolC/E family | 4.70E-15 |
| | | | | | PF02687 | FtsX | FtsX-like permease family | 5.30E-11 |
| | | | | | TIGR02213 | TIGR02213 | lolE_release: lipoprotein releasing system, transmembrane protein LolE | 2.90E-08 |
| WP_129842030.1 | 3863272 | 3862594 | - | 225 | TIGR02211 | TIGR02211 | LolD_lipo_ex: lipoprotein releasing system, ATP-binding protein | 3.00E-64 |
| | | | | | TIGR02673 | TIGR02673 | FtsE: cell division ATP-binding protein FtsE | 8.70E-63 |
| | | | | | TIGR02982 | TIGR02982 | heterocyst_DevA: ABC exporter ATP-binding subunit, DevA family | 8.80E-54 |
| | | | | | TIGR03608 | TIGR03608 | L_ocin_972_ABC: putative bacteriocin export ABC transporter, lactococcin 972 group | 1.20E-53 |
| | | | | | TIGR02315 | TIGR02315 | ABC_phnC: phosphonate ABC transporter, ATP-binding protein | 4.50E-50 |
| WP_129842031.1 | 3864339 | 3863268 | - | 356 | PF01471 | PG_binding_1 | Putative peptidoglycan binding domain | 2.70E-09 |
| | | | | | TIGR02869 | TIGR02869 | spore_SleB: spore cortex-lytic enzyme | 2.20E-08 |
| | | | | | TIGR01730 | TIGR01730 | RND_mfp: efflux transporter, RND family, MFP subunit | 9.70E-04 |

| index | leader | core | start | end | dir | score |
|---|---|---|---|---|---|---|
| 1 | MSRETAHLNRSALLADIAVDDPDGRITMAAKPIM | STPAVLAGVAIGYAMLNAFEAGRSGGPLRPAS | 3860244 | 3860043 | - | 29 |
| 2 | MTAPTTVAAVLDSVADLREDSAGSAPGGARPVL | ATPTLAIVLVAGFAGGYAFATAIGDTRPVEN | 3860477 | 3860282 | - | 30 |
| 3 | MTEKILARNAGTGGPLDECAEGRVARGALPVA | ATPVIVGAAGAIVIAYAAGRIAGELRPVEN | 3860684 | 3860495 | - | 32 |

### Results for WP_097937323.1 [Streptomyces sp. rh206] back to top

previous - next

#### Architecture

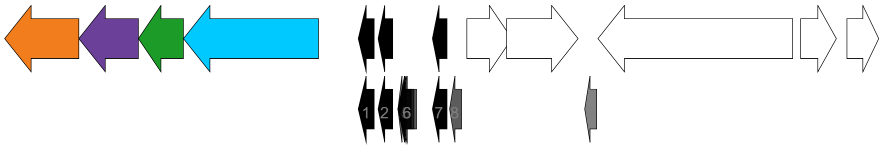

1000 nucleotides

Link to nucleotide sequence

| Accession | start | end | direction | length (aa) | Pfam/HMM | name | description | E-value |
|---|---|---|---|---|---|---|---|---|

| | | | | | | | | |
|---|---|---|---|---|---|---|---|---|
| WP_143672505.1 | 1004 | 0 | - | 335 | PF00664 | ABC_membrane | ABC transporter transmembrane region | 7.50E-35 |
| | | | | | TIGR02204 | TIGR02204 | MsbA_rel: ABC transporter, permease/ATP-binding protein | 4.90E-26 |
| | | | | | TIGR02203 | TIGR02203 | MsbA_lipidA: lipid A export permease/ATP-binding protein MsbA | 4.40E-22 |
| | | | | | TIGR00958 | TIGR00958 | 3a01208: antigen peptide transporter 2 | 7.30E-16 |
| | | | | | TIGR03796 | TIGR03796 | NHLM_micro_ABC1: NHLM bacteriocin system ABC transporter, peptidase/ATP-binding protein | 1.50E-15 |
| WP_097937321.1 | 1813 | 1000 | - | 270 | PF13649 | Methyltransf_25 | Methyltransferase domain | 7.20E-17 |
| | | | | | PF08241 | Methyltransf_11 | Methyltransferase domain | 2.50E-12 |
| | | | | | PF08242 | Methyltransf_12 | Methyltransferase domain | 2.50E-08 |
| | | | | | PF13847 | Methyltransf_31 | Methyltransferase domain | 3.30E-07 |
| | | | | | PF13489 | Methyltransf_23 | Methyltransferase domain | 3.70E-07 |
| WP_097937322.1 | 2418 | 1809 | - | 202 | LinL | LinL | LinL | 1.90E-65 |
| WP_097937323.1 | 4265 | 2414 | - | 616 | LinH | LinH | LinH (Fusion of LinE/LinG) | 1.30E-251 |
| | | | | | LinG | LinG | LinG Homolog | 1.40E-110 |
| | | | | | LinE | LinE | LinE Homolog | 1.90E-95 |
| | | | | | LinL | LinL | LinL | 4.50E-34 |
| | | | | | PF12697 | Abhydrolase_6 | Alpha/beta hydrolase family | 1.10E-15 |
| WP_097936642.1 | 5009 | 4805 | - | 67 | LinA | LinA | LinA (linaridin precursor) | 2.30E-11 |
| WP_097936643.1 | 5264 | 5069 | - | 64 | LinA | LinA | LinA (linaridin precursor) | 9.70E-12 |
| WP_097936644.1 | 6006 | 5802 | - | 67 | LinA | LinA | LinA (linaridin precursor) | 1.50E-11 |
| WP_097936645.1 | 6273 | 6879 | + | 201 | PF04471 | Mrr_cat | Restriction endonuclease | 1.80E-05 |
| WP_097936646.1 | 6805 | 7774 | + | 322 | PF00665 | rve | Integrase core domain | 6.00E-23 |
| | | | | | PF13276 | HTH_21 | HTH-like domain | 2.60E-20 |
| | | | | | PF13683 | rve_3 | Integrase core domain | 1.20E-10 |
| | | | | | PF01527 | HTH_Tnp_1 | Transposase | 9.80E-09 |
| | | | | | PF13384 | HTH_23 | Homeodomain-like domain | 2.70E-04 |
| WP_097936647.1 | 10698 | 8043 | - | 884 | PF13191 | AAA_16 | AAA ATPase domain | 4.20E-20 |
| | | | | | TIGR03903 | TIGR03903 | TOMM_kin_cyc: TOMM system kinase/cyclase fusion protein | 1.90E-19 |
| | | | | | PF13424 | TPR_12 | Tetratricopeptide repeat | 4.80E-05 |
| | | | | | PF07719 | TPR_2 | Tetratricopeptide repeat | 5.10E-05 |
| | | | | | PF13181 | TPR_8 | Tetratricopeptide repeat | 7.80E-05 |
| WP_097936648.1 | 10808 | 11288 | + | 159 | PF03745 | DUF309 | Domain of unknown function (DUF309) | 2.90E-20 |
| WP_097936650.1 | 11426 | 11870 | + | 147 | NO PFAM MATCH | - | - | - |

| index | leader | core | start | end | dir | score |
|---|---|---|---|---|---|---|
| 1 | MSTSPIPAALRGAGGLSESAPGQALSTFAPVA | ATPGIVAGAAVGVALVNAFVAGYNYTGNGNIELPM | 5009 | 4805 | - | 30 |
| 2 | MSKSSTPAALTATGALTESDPGVTLSGLAPVA | ATPVAIVACLGAAFVAGYAAGRAAGSDIELPM | 5264 | 5069 | - | 29 |
| 3 | MDTVPVASVTFTMSAC | VGISVCLTGSTIELPQ | 5436 | 5337 | - | 7 |
| 4 | LSTASIPAVLSETGALSESDPGRAPMD | TVPVASVTFTMSACVGISVCLTGSTIELPQ | 5511 | 5337 | - | 9 |
| 5 | MTSPHRPPTSEEKSLSTASIPAVLSETGALSESDPGRAPMD | TVPVASVTFTMSACVGISVCLTGSTIELPQ | 5553 | 5337 | - | 8 |
| 6 | VGPSRLRRPPMTSPHRPPTSEEKSL...TASIPAVLSETGALSESDPGRAPMD | TVPVASVTFTMSACVGISVCLTGSTIELPQ | 5583 | 5337 | - | 8 |
| 7 | MPEFRQPGWTRGIAPLDESAGGQVFGGASPVA | ATPVVVAGAGAVVIAFAAGVAARTLANGGNIELPM | 6006 | 5802 | - | 30 |
| 8 | VPEPAGCCFGVRILMPAEWSSLSAGL | SASCGAHHRTSVHAGSGSRVRLRAPG | 6202 | 6043 | - | 8 |
| 9 | LSCAVTRGNRGKRDGTWRSLVAHLTGGQ | GVAGSNPVVPTGDSRRSGPVSEQSETGP | 8035 | 7864 | - | 7 |

### Results for THA27824.1 [Streptomyces sp. RKND-216] back to top

previous - next

#### Architecture

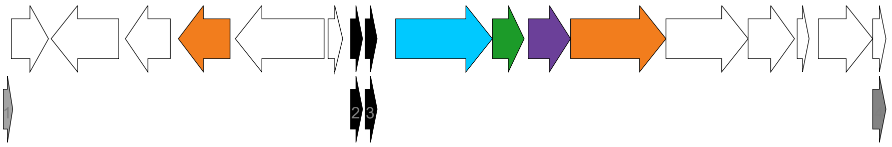

1000 nucleotides

Link to nucleotide sequence

| Accession | start | end | direction | length (aa) | Pfam/HMM | name | description | E-value |
|---|---|---|---|---|---|---|---|---|
| THA26442.1 | 4292202 | 4292865 | + | 220 | PF02254 | TrkA_N | TrkA-N domain | 1.20E-23 |
| | | | | | PF02080 | TrkA_C | TrkA-C domain | 1.50E-13 |
| | | | | | TIGR03026 | TIGR03026 | NDP-sugDHase: nucleotide sugar dehydrogenase | 8.80E-06 |
| | | | | | PF01494 | FAD_binding_3 | FAD binding domain | 1.20E-05 |
| | | | | | PF03446 | NAD_binding_2 | NAD binding domain of 6-phosphogluconate dehydrogenase | 1.50E-05 |
| THA26443.1 | 4294122 | 4292913 | - | 402 | PF07730 | HisKA_3 | Histidine kinase | 7.00E-15 |
| THA26444.1 | 4295049 | 4294245 | - | 267 | PF01061 | ABC2_membrane | ABC-2 type transporter | 2.40E-16 |
| | | | | | PF12698 | ABC2_membrane_3 | ABC-2 family transporter protein | 7.70E-11 |
| | | | | | TIGR03062 | TIGR03062 | pip_yhgE_Cterm: YhgE/Pip C-terminal domain | 2.30E-06 |

| | | | | | | | | |
|---|---|---|---|---|---|---|---|---|
| THA26445.1 | 4296129 | 4295190 | - | 312 | TIGR01188<br>TIGR03864<br>TIGR01288<br>TIGR03522<br>TIGR03265 | TIGR01188<br>TIGR03864<br>TIGR01288<br>TIGR03522<br>TIGR03265 | drrA: daunorubicin resistance ABC transporter, ATP-binding protein<br>PQQ_ABC_ATP: ABC transporter, ATP-binding subunit, PQQ-dependent alcohol dehydrogenase system<br>nodI: nodulation ABC transporter NodI<br>GldA_ABC_ATP: gliding motility-associated ABC transporter ATP-binding subunit GldA<br>PhnT2: putative 2-aminoethylphosphonate ABC transporter, ATP-binding protein | 4.20E-69<br>1.50E-48<br>2.00E-46<br>5.70E-37<br>2.50E-35 |
| THA26446.1 | 4297805 | 4296224 | - | 526 | NO PFAM MATCH | - | - | - |
| THA26447.1 | 4297892 | 4298138 | + | 81 | NO PFAM MATCH | - | - | - |
| THA26448.1 | 4298298 | 4298493 | + | 64 | LinA | LinA | LinA (linaridin precursor) | 8.00E-15 |
| THA26449.1 | 4298563 | 4298755 | + | 63 | LinA | LinA | LinA (linaridin precursor) | 5.50E-09 |
| THA27824.1 | 4299103 | 4300840 | + | 578 | LinH<br>LinE<br>LinG<br>PF12697<br>PF00561 | LinH<br>LinE<br>LinG<br>Abhydrolase_6<br>Abhydrolase_1 | LinH (Fusion of LinE/LinG)<br>LinE Homolog<br>LinG Homolog<br>Alpha/beta hydrolase family<br>alpha/beta hydrolase fold | 2.40E-<br>217<br>1.90E-98<br>7.30E-94<br>2.30E-15<br>1.10E-11 |
| THA26450.1 | 4300836 | 4301418 | + | 193 | LinL | LinL | LinL | 1.20E-62 |
| THA26451.1 | 4301485 | 4302244 | + | 252 | PF13649<br>PF08241<br>PF08242<br>PF13489<br>PF13847 | Methyltransf_25<br>Methyltransf_11<br>Methyltransf_12<br>Methyltransf_23<br>Methyltransf_31 | Methyltransferase domain<br>Methyltransferase domain<br>Methyltransferase domain<br>Methyltransferase domain<br>Methyltransferase domain | 5.90E-13<br>5.90E-09<br>8.60E-09<br>1.90E-08<br>2.40E-06 |
| THA26452.1 | 4302240 | 4303965 | + | 574 | TIGR02204<br>TIGR02203<br>TIGR00958<br>TIGR03796<br>TIGR01846 | TIGR02204<br>TIGR02203<br>TIGR00958<br>TIGR03796<br>TIGR01846 | MsbA_rel: ABC transporter, permease/ATP-binding protein<br>MsbA_lipidA: lipid A export permease/ATP-binding protein MsbA<br>3a01208: antigen peptide transporter 2<br>NHLM_micro_ABC1: NHLM bacteriocin system ABC transporter, peptidase/ATP-binding protein<br>type_I_sec_HlyB: type I secretion system ATPase | 7.70E-<br>100<br>3.80E-98<br>3.60E-87<br>4.10E-85<br>5.80E-83 |
| THA26453.1 | 4303961 | 4305443 | + | 493 | TIGR02734<br>TIGR03467<br>TIGR00292<br>TIGR02733<br>PF13450 | TIGR02734<br>TIGR03467<br>TIGR00292<br>TIGR02733<br>NAD_binding_8 | crtI_fam: phytoene desaturase<br>HpnE: squalene-associated FAD-dependent desaturase<br>TIGR00292: thiazole biosynthesis enzyme<br>desat_CrtD: C-3',4' desaturase CrtD<br>NAD(P)-binding Rossmann-like domain | 1.00E-11<br>4.40E-10<br>1.20E-08<br>1.10E-07<br>2.60E-07 |
| THA26454.1 | 4305439 | 4306261 | + | 273 | PF12680<br>PF14534<br>PF07366 | SnoaL_2<br>DUF4440<br>SnoaL | SnoaL-like domain<br>Domain of unknown function (DUF4440)<br>SnoaL-like polyketide cyclase | 6.50E-08<br>4.70E-05<br>2.80E-04 |
| THA26455.1 | 4306307 | 4306526 | + | 72 | PF03621 | MbtH | MbtH-like protein | 1.10E-29 |
| THA27825.1 | 4306691 | 4307672 | + | 326 | TIGR00747<br>PF08541 | TIGR00747<br>ACP_syn_III_C | fabH: 3-oxoacyl-[acyl-carrier-protein] synthase III<br>3-Oxoacyl-[acyl-carrier-protein (ACP)] synthase III C terminal | 2.50E-10<br>3.40E-05 |
| THA26456.1 | 4307668 | 4307926 | + | 85 | PF00550<br>TIGR01688 | PP-binding<br>TIGR01688 | Phosphopantetheine attachment site<br>dltC: D-alanine--poly(phosphoribitol) ligase, subunit 2 | 8.00E-07<br>1.10E-04 |

| index | leader | core | start | end | dir | score |
|---|---|---|---|---|---|---|
| 1 | VGLSTGITADLPRAGQCVLVALMFIGRIGP | LTLASALALRNRTRRYELPEERTIVG | 4292039 | 4292210 | + | 6 |
| 2 | MEQKIATFMNGQEFGSLENGFHETAPAPVM | GTPVAAVATCWYVAGGVAGAGAVVGAYTWGVANG | 4298298 | 4298493 | + | 30 |
| 3 | MSRITESVIAVTPEKLDDGATTELGAMAPVQ | ATPGFVAFGVSFAGGAGVAWLGVQAYEAGVND | 4298563 | 4298755 | + | 31 |
| 4 | MNLITSEITAFLSDALRCEVGPEDDYFALGLADSLFALEL | VTFVEERFSLTVEVEDLDLDSFRTADRITGFVQRKSGAAAHPEGP | 4307668 | 4307926 | + | 7 |

### Results for PPQ56339.1 [Streptomyces sp. QL37] back to top

previous - next

#### Architecture

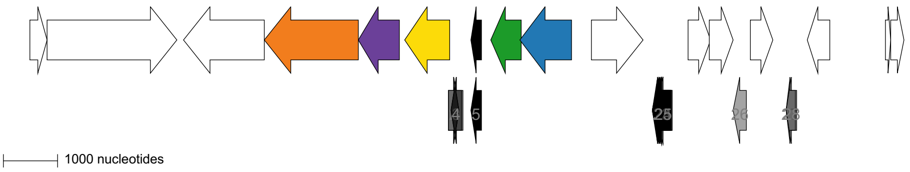

Link to nucleotide sequence

| Accession | start | end | direction | length (aa) | Pfam/HMM | name | description | E-value |
|---|---|---|---|---|---|---|---|---|
| PPQ56332.1 | 1435790 | 1436117 | + | 108 | PF13443 | HTH_26 | Cro/C1-type HTH DNA-binding domain | 1.20E-22 |
| PPQ56333.1 | 1436113 | 1438492 | + | 792 | NO PFAM MATCH | - | - | - |
| PPQ56334.1 | 1440114 | 1438629 | - | 494 | TIGR03467<br>TIGR00292<br>TIGR02734<br>TIGR02733<br>TIGR01317 | TIGR03467<br>TIGR00292<br>TIGR02734<br>TIGR02733<br>TIGR01317 | HpnE: squalene-associated FAD-dependent desaturase<br>TIGR00292: thiazole biosynthesis enzyme<br>crtI_fam: phytoene desaturase<br>desat_CrtD: C-3',4' desaturase CrtD<br>GOGAT_sm_gam: glutamate synthase, NADH/NADPH, small subunit | 2.10E-08<br>2.50E-08<br>3.90E-08<br>5.00E-08<br>1.60E-07 |
| PPQ56335.1 | 1441844 | 1440110 | - | 577 | TIGR02204<br>TIGR02203<br>TIGR03796<br>TIGR00958<br>TIGR01846 | TIGR02204<br>TIGR02203<br>TIGR03796<br>TIGR00958<br>TIGR01846 | MsbA_rel: ABC transporter, permease/ATP-binding protein<br>MsbA_lipidA: lipid A export permease/ATP-binding protein MsbA<br>NHLM_micro_ABC1: NHLM bacteriocin system ABC transporter, peptidase/ATP-binding protein<br>3a01208: antigen peptide transporter 2<br>type_I_sec_HlyB: type I secretion system ATPase | 1.10E-97<br>3.30E-95<br>1.30E-82<br>7.30E-80<br>1.90E-79 |

| | | | | | | | | |
|---|---|---|---|---|---|---|---|---|
| PPQ56336.1 | 1442596 | 1441840 | - | 251 | PF13649<br>PF08241<br>PF08242<br>PF13489<br>TIGR00477 | Methyltransf_25<br>Methyltransf_11<br>Methyltransf_12<br>Methyltransf_23<br>TIGR00477 | Methyltransferase domain<br>Methyltransferase domain<br>Methyltransferase domain<br>Methyltransferase domain<br>tehB: tellurite resistance protein TehB | 4.40E-14<br>4.30E-09<br>2.10E-08<br>2.40E-08<br>9.20E-06 |
| PPQ62208.1 | 1443533 | 1442699 | - | 277 | LinG<br>LinH<br>PF06762 | LinG<br>LinH<br>LMF1 | LinG Homolog<br>LinH (Fusion of LinE/LinG)<br>Lipase maturation factor | 5.30E-103<br>6.80E-73<br>9.50E-04 |
| PPQ56337.1 | 1444114 | 1443922 | - | 63 | LinA | LinA | LinA (linaridin precursor) | 8.20E-11 |
| PPQ56338.1 | 1444845 | 1444284 | - | 186 | LinL | LinL | LinL | 1.20E-54 |
| PPQ56339.1 | 1445792 | 1444841 | - | 316 | LinE<br>LinH<br>PF12697<br>PF00561<br>TIGR02427 | LinE<br>LinH<br>Abhydrolase_6<br>Abhydrolase_1<br>TIGR02427 | LinE Homolog<br>LinH (Fusion of LinE/LinG)<br>Alpha/beta hydrolase family<br>alpha/beta hydrolase fold<br>protocat_pcaD: 3-oxoadipate enol-lactonase | 1.70E-90<br>7.40E-43<br>6.00E-18<br>6.30E-09<br>2.00E-08 |
| PPQ62209.1 | 1446156 | 1447107 | + | 316 | PF14200<br>PF13560 | RicinB_lectin_2<br>HTH_31 | Ricin-type beta-trefoil lectin domain-like<br>Helix-turn-helix domain | 1.00E-06<br>1.10E-04 |
| PPQ62210.1 | 1447923 | 1448337 | + | 137 | NO PFAM MATCH | - | - | - |
| PPQ56340.1 | 1448333 | 1448774 | + | 146 | NO PFAM MATCH | - | - | - |
| PPQ56341.1 | 1449070 | 1449490 | + | 139 | NO PFAM MATCH | - | - | - |
| PPQ62211.1 | 1450548 | 1450131 | - | 138 | NO PFAM MATCH | - | - | - |
| WP_007268929.1 | 1451565 | 1451670 | + | 35 | INFERRED GENE | - | - | - |
| WP_010981454.1 | 1451671 | 1451914 | + | 81 | INFERRED GENE | - | - | - |

| index | leader | core | start | end | dir | score |
|---|---|---|---|---|---|---|
| 1 | VVGGCSEATDDAARFSRRNTGSPSRWEWVPGTGRCLSPTV | TVPDRADEAAGLATRRAPRRLLSGTG | 1443516 | 1443717 | + | 6 |
| 2 | VGGCSEATDDAARFSRRNTGSPSRWEWVPGTGRCLSPTV | TVPDRADEAAGLATRRAPRRLLSGTG | 1443519 | 1443717 | + | 6 |
| 3 | VGGGSTASRAVPATRCRSGGDGVPGAWLSPPPRRPGPAPSPSGTGTVLS | PAPTPTEKENPCSAD | 1443754 | 1443559 | - | 7 |
| 4 | VNDRVPVGGGSTASRAVPATRCRSG...GAWLSPPPRRPGPAPSPSGTGTVLS | PAPTPTEKENPCSAD | 1443772 | 1443559 | - | 7 |
| 5 | MSIVAQFANESLTDVTPGVLSGTATLGLNTPADPVM | ATPVLAFEVFLIATEMLYDNVQNYTNS | 1444114 | 1443922 | - | 30 |
| 6 | VGKSGSSFRLTDFTAGVPKYSVRESLVISRITGGRGLFRP | CTIPRIGASWNAAQS | 1447454 | 1447286 | - | 7 |
| 7 | VDGGQVGVEFSLDRLHRRGAEVLGQGVLG | DLADYWREGVIPTVHYSSHRGILERSPVLS | 1447461 | 1447281 | - | 7 |
| 8 | VIHQWTVGKSGSSFRLTDFTAGVPKYSVRESLVISRITGGRGLFRP | CTIPRIGASWNAAQS | 1447472 | 1447286 | - | 7 |
| 9 | MAWATGAVIHQWTVGKSGSSFRLTD...GVPKYSVRESLVISRITGGRGLFRP | CTIPRIGASWNAAQS | 1447493 | 1447286 | - | 7 |
| 10 | MAVSMAWATGAVIHQWTVGKSGSSF...GVPKYSVRESLVISRITGGRGLFRP | CTIPRIGASWNAAQS | 1447505 | 1447286 | - | 6 |
| 11 | VHGGLDGLGDRCGDPPVDGGQVGVEFSLDRLHRRGAE | VLGQGVLGDLADYWREGVIPTVHYSSHRGILERSPVLS | 1447509 | 1447281 | - | 9 |
| 12 | VHRQVHGGLDGLGDRCGDPPVDGGQVGVEFSLDRLHRRG | AEVLGQGVLGDLADYWREGVIPTVHYSSHRGILERSPVLS | 1447521 | 1447281 | - | 9 |
| 13 | VSCIDKCMAVSMAWATGAVIHQWTV...GVPKYSVRESLVISRITGGRGLFRP | CTIPRIGASWNAAQS | 1447526 | 1447286 | - | 7 |
| 14 | LKRGATAANARVSCIDKCMAVSMAW...GVPKYSVRESLVISRITGGRGLFRP | CTIPRIGASWNAAQS | 1447559 | 1447286 | - | 7 |
| 15 | VKPSPLKRGATAANARVSCIDKCMA...GVPKYSVRESLVISRITGGRGLFRP | CTIPRIGASWNAAQS | 1447574 | 1447286 | - | 7 |
| 16 | MAVWLVKPSPLKRGATAANARVSCI...GVPKYSVRESLVISRITGGRGLFRP | CTIPRIGASWNAAQS | 1447589 | 1447286 | - | 7 |
| 17 | LMAVWLVKPSPLKRGATAANARVSC...GVPKYSVRESLVISRITGGRGLFRP | CTIPRIGASWNAAQS | 1447592 | 1447286 | - | 6 |
| 18 | MKRLMAVWLVKPSPLKRGATAANAR...GVPKYSVRESLVISRITGGRGLFRP | CTIPRIGASWNAAQS | 1447601 | 1447286 | - | 6 |
| 19 | LMKRLMAVWLVKPSPLKRGATAANA...GVPKYSVRESLVISRITGGRGLFRP | CTIPRIGASWNAAQS | 1447604 | 1447286 | - | 6 |
| 20 | VLDEAVDGGLAGEAFTLEARGDCGE...VHRQVHGGLDGLGDRCGDPPVDGGQ | VGVEFSLDRLHRRGAEVLGQGVLGD...WREGVIPTVHYSSHRGILERSPVLS | 1447608 | 1447281 | - | 8 |
| 21 | LTCLMKRLMAVWLVKPSPLKRGATA...GVPKYSVRESLVISRITGGRGLFRP | CTIPRIGASWNAAQS | 1447613 | 1447286 | - | 6 |
| 22 | VPVDVLDEAVDGGLAGEAFTLEARG...GVVHRQVHGGLDGLGDRCGDPPVDG | GQVGVEFSLDRLHRRGAEVLGQGVL...WREGVIPTVHYSSHRGILERSPVLS | 1447620 | 1447281 | - | 8 |
| 23 | VPVPVDVLDEAVDGGLAGEAFTLEA...EGVVHRQVHGGLDGLGDRCGDPPVD | GGQVGVEFSLDRLHRRGAEVLGQGV...WREGVIPTVHYSSHRGILERSPVLS | 1447626 | 1447281 | - | 8 |
| 24 | VCRCRLTCLMKRLMAVWLVKPSPLK...GVPKYSVRESLVISRITGGRGLFRP | CTIPRIGASWNAAQS | 1447628 | 1447286 | - | 6 |
| 25 | VVCRCRLTCLMKRLMAVWLVKPSPL...GVPKYSVRESLVISRITGGRGLFRP | CTIPRIGASWNAAQS | 1447631 | 1447286 | - | 6 |
| 26 | LGDGAWKAESFAHL | FTTEHKGLLLVGLKCLENIAGRQVA...PDRRESSDQPTLFDASGGTGPSDNR | 1449019 | 1448770 | - | 6 |
| 27 | MRIAAAMYSQTSTSSRSISIRGCAFRA | QALNARVGRGRAANPTPFYGSPVSGNC | 1449909 | 1449744 | - | 6 |
| 28 | VFQCQQTMRIAAAMYSQTSTSSRSISIRGC | AFRAQALNARVGRGRAANPTPFYGSPVSGNC | 1449930 | 1449744 | - | 6 |

### Results for WP_057604735.1 [Streptomyces sp. Root1310] back to top

previous - next

#### Architecture

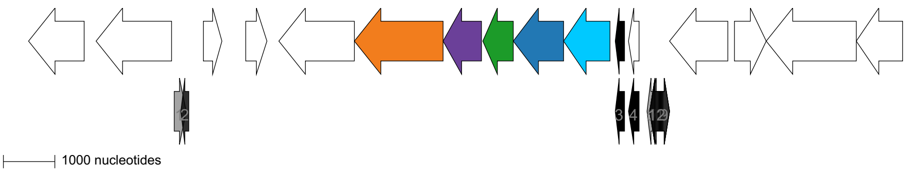

Link to nucleotide sequence

| Accession | start | end | direction | length (aa) | Pfam/HMM | name | description | E-value |
|---|---|---|---|---|---|---|---|---|
| WP_057604729.1 | 442658 | 441584 | - | 357 | TIGR00036<br>PF01113 | TIGR00036<br>DapB_N | dapB: 4-hydroxy-tetrahydrodipicolinate reductase<br>Dihydrodipicolinate reductase, N-terminus | 6.30E-07<br>2.60E-06 |
| WP_057604730.1 | 444376 | 442888 | - | 495 | NO PFAM MATCH | - | - | - |
| WP_079074079.1 | 444994 | 445336 | + | 113 | PF03795 | YCII | YCII-related domain | 5.70E-07 |
| WP_014669667.1 | 445800 | 446216 | + | 138 | INFERRED GENE | - | - | - |
| WP_057604731.1 | 447930 | 446460 | - | 489 | TIGR02734<br>PF13450<br>TIGR03467<br>PF01593<br>TIGR01317 | TIGR02734<br>NAD_binding_8<br>TIGR03467<br>Amino_oxidase<br>TIGR01317 | crtI_fam: phytoene desaturase<br>NAD(P)-binding Rossmann-like domain<br>HpnE: squalene-associated FAD-dependent desaturase<br>Flavin containing amine oxidoreductase<br>GOGAT_sm_gam: glutamate synthase, NADH/NADPH, small subunit | 3.50E-14<br>9.90E-11<br>5.10E-10<br>1.50E-09<br>1.50E-09 |
| WP_057604732.1 | 449642 | 447926 | - | 571 | TIGR02203<br>TIGR02204<br>TIGR00958<br>TIGR01846<br>TIGR03375 | TIGR02203<br>TIGR02204<br>TIGR00958<br>TIGR01846<br>TIGR03375 | MsbA_lipidA: lipid A export permease/ATP-binding protein MsbA<br>MsbA_rel: ABC transporter, permease/ATP-binding protein<br>3a01208: antigen peptide transporter 2<br>type_I_sec_HlyB: type I secretion system ATPase<br>type_I_sec_LssB: type I secretion system ATPase | 4.40E-104<br>2.30E-102<br>1.00E-89<br>4.50E-85<br>8.40E-84 |
| WP_057604733.1 | 450400 | 449638 | - | 253 | PF13649<br>PF13489<br>PF08241<br>TIGR01983<br>TIGR02469 | Methyltransf_25<br>Methyltransf_23<br>Methyltransf_11<br>TIGR01983<br>TIGR02469 | Methyltransferase domain<br>Methyltransferase domain<br>Methyltransferase domain<br>UbiG: 3-demethylubiquinone-9 3-O-methyltransferase<br>CbiT: precorrin-6Y C5,15-methyltransferase (decarboxylating), CbiT subunit | 2.00E-10<br>2.40E-06<br>6.10E-06<br>7.40E-06<br>3.60E-04 |
| WP_057604734.1 | 451006 | 450415 | - | 196 | LinL | LinL | LinL | 3.10E-63 |
| WP_057604735.1 | 451998 | 451002 | - | 331 | LinE<br>LinH<br>PF12697<br>PF00561<br>TIGR02427 | LinE<br>LinH<br>Abhydrolase_6<br>Abhydrolase_1<br>TIGR02427 | LinE Homolog<br>LinH (Fusion of LinE/LinG)<br>Alpha/beta hydrolase family<br>alpha/beta hydrolase fold<br>protocat_pcaD: 3-oxoadipate enol-lactonase | 1.10E-116<br>8.70E-109<br>7.90E-19<br>8.00E-13<br>9.50E-11 |
| WP_157880721.1 | 452879 | 451994 | - | 294 | LinH<br>LinG | LinH<br>LinG | LinH (Fusion of LinE/LinG)<br>LinG Homolog | 1.90E-111<br>1.20E-101 |
| WP_157880722.1 | 453164 | 452996 | - | 55 | LinA | LinA | LinA (linaridin precursor) | 7.50E-13 |
| WP_157880723.1 | 453455 | 453257 | - | 65 | NO PFAM MATCH | - | - | - |
| WP_057604738.1 | 455182 | 454042 | - | 379 | TIGR03207<br>PF00441<br>TIGR03203<br>TIGR03204<br>PF02770 | TIGR03207<br>Acyl-CoA_dh_1<br>TIGR03203<br>TIGR03204<br>Acyl-CoA_dh_M | cyc_hxne_CoA_dh: cyclohexanecarboxyl-CoA dehydrogenase<br>Acyl-CoA dehydrogenase, C-terminal domain<br>pimD_small: pimeloyl-CoA dehydrogenase, small subunit<br>pimC_large: pimeloyl-CoA dehydrogenase, large subunit<br>Acyl-CoA dehydrogenase, middle domain | 1.20E-50<br>2.50E-50<br>2.40E-40<br>1.90E-35<br>6.90E-23 |
| WP_057604739.1 | 455293 | 455920 | + | 208 | PF00440<br>TIGR03384<br>TIGR03968<br>TIGR03613<br>PF18556 | TetR_N<br>TIGR03384<br>TIGR03968<br>TIGR03613<br>TetR_C_35 | Bacterial regulatory proteins, tetR family<br>betaine_BetI: transcriptional repressor BetI<br>mycofact_TetR: mycofactocin system transcriptional regulator<br>RutR: pyrimidine utilization regulatory protein R<br>Bacterial Tetracyclin repressor, C-terminal domain | 1.60E-16<br>2.10E-09<br>3.00E-09<br>2.40E-08<br>1.50E-06 |
| WP_057604740.1 | 457683 | 455934 | - | 582 | PF00171<br>TIGR01780<br>TIGR01804<br>TIGR03216<br>TIGR02299 | Aldedh<br>TIGR01780<br>TIGR01804<br>TIGR03216<br>TIGR02299 | Aldehyde dehydrogenase family<br>SSADH: succinate-semialdehyde dehydrogenase<br>BADH: betaine-aldehyde dehydrogenase<br>OH_muco_semi_DH: 2-hydroxymuconic semialdehyde dehydrogenase<br>HpaE: 5-carboxymethyl-2-hydroxymuconate semialdehyde dehydrogenase | 5.20E-121<br>1.90E-78<br>1.80E-76<br>3.20E-72<br>4.20E-68 |
| WP_057604741.1 | 458573 | 457679 | - | 297 | NO PFAM MATCH | - | - | - |

| index | leader | core | start | end | dir | score |
|---|---|---|---|---|---|---|
| 1 | VRARARGRSRPDGCVGLGGHGCVRPGLLRGCGGSRQRQGAVVP | GTAAVGVRARADRGRRRFARDRRGVGA | 444429 | 444642 | + | 6 |
| 2 | MPVRSCFDHGHEVSVPGSHAPTPRR | SRANLRRPLSARALTPTAAVPGTTAP | 444695 | 444539 | - | 9 |
| 3 | MDSLTSIVNHDERLDALHIASTQTEILSETTVPVL | GTPAALAASVAVATAIHSIW | 453164 | 452996 | - | 29 |
| 4 | MSPVLEMVNASEEIGSLPMDITESIHGDLDPGMFISLSDAVNTALAVC | GLPAHAAVHCAVAALTH | 453455 | 453257 | - | 20 |
| 5 | LHNYRHARTACHLQELDEDYAERL | SRVFLSPHAKVAVIHTTDSSLCPR | 453764 | 453617 | - | 6 |
| 6 | MAGCIWSVTLEVAAMSDNRIAWWFACNVRLSRQGLRICR | ATPDGGSAEVRTRPGPGRAGAAVGLVASGAGPAQRP | 453827 | 454055 | + | 6 |
| 7 | VTLHIHPAIFDDQGQRFSPALKGFGHPVKSCTTTDTHGLHVTCKNLTR | ITPSAFRGSFSRPMPKSL | 453853 | 453652 | - | 7 |
| 8 | VAAMSDNRIAWWFACNVRLSRQGLRICR | ATPDGGSAEVRTRPGPGRAGAAVGLVASGAGPAQRP | 453860 | 454055 | + | 7 |
| 9 | MSDNRIAWWFACNVRLSRQGLRICR | ATPDGGSAEVRTRPGPGRAGAAVGLVASGAGPAQRP | 453869 | 454055 | + | 7 |
| 10 | MAATSRVTLHIHPAIFDDQGQRFSP...FGHPVKSCTTTDTHGLHVTCKNLTR | ITPSAFRGSFSRPMPKSL | 453871 | 453652 | - | 6 |
| 11 | LSDMAATSRVTLHIHPAIFDDQGQR...FGHPVKSCTTTDTHGLHVTCKNLTR | ITPSAFRGSFSRPMPKSL | 453880 | 453652 | - | 7 |
| 12 | LQANHQAIRLSDMAATSRVTLHIHP...FGHPVKSCTTTDTHGLHVTCKNLTR | ITPSAFRGSFSRPMPKSL | 453907 | 453652 | - | 7 |

### Results for KQX31486.1 [Streptomyces sp. Root1295] back to top

### Architecture

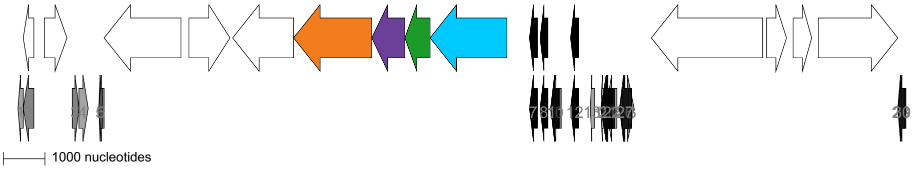

Link to nucleotide sequence

| Accession | start | end | direction | length (aa) | Pfam/HMM | name | description | E-value |
|---|---|---|---|---|---|---|---|---|
| KQX31039.1 | 934042 | 933793 | - | 82 | NO PFAM MATCH | - | - | - |
| KQX31040.1 | 934300 | 934837 | + | 178 | PF12840<br>PF01022 | HTH_20<br>HTH_5 | Helix-turn-helix domain<br>Bacterial regulatory protein, arsR family | 2.80E-10<br>3.30E-05 |
| KQX31041.1 | 937574 | 935720 | - | 617 | NO PFAM MATCH | - | - | - |
| KQX31042.1 | 937763 | 938777 | + | 337 | NO PFAM MATCH | - | - | - |
| KQX31043.1 | 940301 | 938831 | - | 489 | TIGR02734<br>PF13450<br>TIGR01317<br>TIGR00292<br>TIGR03467 | TIGR02734<br>NAD_binding_8<br>TIGR01317<br>TIGR00292<br>TIGR03467 | crtI_fam: phytoene desaturase<br>NAD(P)-binding Rossmann-like domain<br>GOGAT_sm_gam: glutamate synthase, NADH/NADPH, small subunit<br>TIGR00292: thiazole biosynthesis enzyme<br>HpnE: squalene-associated FAD-dependent desaturase | 3.60E-11<br>5.10E-09<br>1.00E-08<br>1.30E-08<br>7.50E-08 |
| KQX31044.1 | 942169 | 940297 | - | 623 | TIGR02204<br>TIGR02203<br>TIGR00958<br>TIGR03796<br>TIGR03375 | TIGR02204<br>TIGR02203<br>TIGR00958<br>TIGR03796<br>TIGR03375 | MsbA_rel: ABC transporter, permease/ATP-binding protein<br>MsbA_lipidA: lipid A export permease/ATP-binding protein MsbA<br>3a01208: antigen peptide transporter 2<br>NHLM_micro_ABC1: NHLM bacteriocin system ABC transporter, peptidase/ATP-binding protein<br>type_I_sec_LssB: type I secretion system ATPase | 1.20E-103<br>9.90E-97<br>8.80E-85<br>8.50E-81<br>2.20E-80 |
| KQX31045.1 | 942975 | 942165 | - | 269 | PF13649<br>PF08241<br>PF08242<br>PF13489<br>PF13847 | Methyltransf_25<br>Methyltransf_11<br>Methyltransf_12<br>Methyltransf_23<br>Methyltransf_31 | Methyltransferase domain<br>Methyltransferase domain<br>Methyltransferase domain<br>Methyltransferase domain<br>Methyltransferase domain | 2.30E-16<br>7.50E-12<br>3.30E-08<br>2.70E-07<br>1.80E-06 |
| KQX31046.1 | 943580 | 942971 | - | 202 | LinL | LinL | LinL | 2.80E-65 |
| KQX31486.1 | 945424 | 943576 | - | 615 | LinH<br>LinG<br>LinE<br>LinL<br>PF12697 | LinH<br>LinG<br>LinE<br>LinL<br>Abhydrolase_6 | LinH (Fusion of LinE/LinG)<br>LinG Homolog<br>LinE Homolog<br>LinL<br>Alpha/beta hydrolase family | 3.80E-253<br>1.40E-111<br>3.40E-95<br>1.20E-36<br>5.80E-15 |
| KQX31047.1 | 946173 | 945972 | - | 66 | LinA | LinA | LinA (linaridin precursor) | 3.70E-14 |
| KQX31048.1 | 946425 | 946230 | - | 64 | LinA | LinA | LinA (linaridin precursor) | 7.70E-11 |
| KQX31049.1 | 947166 | 946962 | - | 67 | LinA | LinA | LinA (linaridin precursor) | 8.70E-12 |
| KQX31050.1 | 951575 | 948920 | - | 884 | PF13191<br>TIGR03903<br>PF13424<br>PF07719<br>PF13401 | AAA_16<br>TIGR03903<br>TPR_12<br>TPR_2<br>AAA_22 | AAA ATPase domain<br>TOMM_kin_cyc: TOMM system kinase/cyclase fusion protein<br>Tetratricopeptide repeat<br>Tetratricopeptide repeat<br>AAA domain | 4.20E-20<br>4.90E-19<br>4.80E-05<br>5.50E-05<br>8.70E-05 |
| KQX31051.1 | 951685 | 952165 | + | 159 | PF03745 | DUF309 | Domain of unknown function (DUF309) | 6.20E-20 |
| KQX31487.1 | 952304 | 952769 | + | 154 | NO PFAM MATCH | - | - | - |
| KQX31052.1 | 952918 | 954844 | + | 641 | PF10022 | DUF2264 | Uncharacterized protein conserved in bacteria (DUF2264) | 4.90E-100 |

| index | leader | core | start | end | dir | score |
|---|---|---|---|---|---|---|
| 1 | VDDVPLAPGGLVVRGPG | TEAAARWRRVPMSGLTSP | 933778 | 933670 | - | 6 |
| 2 | MIKNFVRGFSALSLALVPLL | SAPAAQAAPAAEVTTLTDAVGPVQV...SHGVGVIALTPYGVAHGVPTGSYRV | 934042 | 933793 | - | 7 |
| 3 | LTGCSRAGPQRVAAAASASRPRPPSRAG | ACSASSAVVPTRITDIVKAVVVLHHAST | 934961 | 935132 | + | 7 |
| 4 | MRPKRAPSSLAQHRFWELPYKQTPFAFRT | QTPEPSSASPAGASTSCSPCSATAPSTNPEPRTPTVTLAA | 935128 | 935338 | + | 6 |
| 5 | MTAAAVGGGPGASAVDL | ARLSPAGTRSCGTVTKV | 935709 | 935604 | - | 7 |
| 6 | MRGAGMTAAAVGGGPGASA | VDLARLSPAGTRSCGTVTKV | 935724 | 935604 | - | 6 |
| 7 | MSMSPTPAALRGTGGLSESDPGRALSSLAPVA | ATPGVVAGVAVGVALVNAFAAGYNHCGGNVELPM | 946173 | 945972 | - | 31 |
| 8 | MNKSSAPAVLTATGALTESAPGVTLPTLAPVA | ATPVAIAATMGVAFVAGYAAGRAATGNVELPM | 946425 | 946230 | - | 30 |
| 9 | LNTASIPAVLSETGALSESDHGRAPLD | TVPVASVTFTMTACVEVSVCLTGSVIQLPQ | 946669 | 946495 | - | 9 |
| 10 | MTSPHRPPTSEEKSLNTASIPAVLSETGALSESDHGRAPLD | TVPVASVTFTMTACVEVSVCLTGSVIQLPQ | 946711 | 946495 | - | 8 |
| 11 | VGPSRRRRRLPMTSPHRPPTSEEKS...TASIPAVLSETGALSESDHGRAPLD | TVPVASVTFTMTACVEVSVCLTGSVIQLPQ | 946744 | 946495 | - | 7 |
| 12 | MPEFREPGWTRGIAPLQEDAGGQVFGGASPVA | ATPAVVATAGAVVVAFAAGVAARHLANGGNVELPM | 947166 | 946962 | - | 30 |
| 13 | MATLKGCRLLRKLRCST | NRLTDLVKAVLALHHASA | 947531 | 947423 | - | 6 |
| 14 | MQLLPDAGVVPVPQ | PTPARHARPEAQLLWQVLPLDAGVRHE | 947805 | 947679 | - | 6 |
| 15 | VIRWCFDPVLPRSAKLGPVRLPLYG...GRRVELGRQLLMQLLPDAGVVPVPQ | PTPARHARPEAQLLWQVLPLDAGVRHE | 947958 | 947679 | - | 7 |
| 16 | VPCPSVIRWCFDPVLPRSAKLGPVR...GRRVELGRQLLMQLLPDAGVVPVPQ | PTPARHARPEAQLLWQVLPLDAGVRHE | 947973 | 947679 | - | 7 |

| | | | | | | |
|---|---|---|---|---|---|---|
| 17 | VRIRASGVPCPSVIRWCFDPVLPRS...GRRVELGRQLLMQLLPDAGVVPVPQ | PTPARHARPEAQLLWQVLPLDAGVRHE | 947994 | 947679 | - | 6 |
| 18 | LPPVRIRASGVPCPSVIRWCFDPVL...GRRVELGRQLLMQLLPDAGVVPVPQ | PTPARHARPEAQLLWQVLPLDAGVRHE | 948003 | 947679 | - | 6 |
| 19 | LRLPPVRIRASGVPCPSVIRWCFDP...GRRVELGRQLLMQLLPDAGVVPVPQ | PTPARHARPEAQLLWQVLPLDAGVRHE | 948009 | 947679 | - | 6 |
| 20 | VTSLRLPPVRIRASGVPCPSVIRWC...GRRVELGRQLLMQLLPDAGVVPVPQ | PTPARHARPEAQLLWQVLPLDAGVRHE | 948018 | 947679 | - | 6 |
| 21 | MSCVTSLRLPPVRIRASGVPCPSVI...GRRVELGRQLLMQLLPDAGVVPVPQ | PTPARHARPEAQLLWQVLPLDAGVRHE | 948027 | 947679 | - | 6 |
| 22 | MPDPAPDRWGGVDQPHELRDVVAVAACEDQGERSAVSVGDQVVLRPGS | ATICQTRSRPAPPLWRARGWSRPPHATSRDGPPR | 948073 | 947824 | - | 6 |
| 23 | VDLDRAPSDHTRPGWADRQEHQGQA...LSPGPIRTPACSRAPEAEVSPPPSC | ATPHTGAPWSPSSATSICAATTSGTPR | 948204 | 948492 | + | 7 |
| 24 | VNRRASGWGRGTGCAGRRGADLLDDRNLAGPLALGALVG | QPTRGGCGLTAHGPDPLGGVDVSDAAAGDLPDPCGGTRG | 948370 | 948133 | - | 6 |
| 25 | VEIPPPRGPVNRRASGWGRGTGCAGRRGADLLDDRNLAGPLAL | GALVGQPTRGGCGLTAHGPDPLGGVDVSDAAAGDLPDPCGGTRG | 948397 | 948133 | - | 6 |
| 26 | VASRRTAVEIPPPRGPVNRRASGWGRGTGCAGRRGADLLDDRNLAGP | LALGALVGQPTRGGCGLTAHGPDPLGGVDVSDAAAGDLPDPCGGTRG | 948418 | 948133 | - | 6 |
| 27 | VPQCVASRRTAVEIPPPRGPVNRRASGWGRGTGCAGRRGADLLDDRNLA | GPLALGALVGQPTRGGCGLTAHGPDPLGGVDVSDAAAGDLPDPCGGTRG | 948430 | 948133 | - | 6 |
| 28 | VPEAVARLAPER | PTTLISGPSATSDIELERVEGVHGPRTLVVVIRTDA | 954995 | 954848 | - | 6 |
| 29 | VDLSAVEAGVPEAVARLAPER | PTTLISGPSATSDIELERVEGVHGPRTLVVVIRTDA | 955022 | 954848 | - | 8 |
| 30 | VVDLSAVEAGVPEAVARLAPER | PTTLISGPSATSDIELERVEGVHGPRTLVVVIRTDA | 955025 | 954848 | - | 8 |

### Results for WP_056796007.1 [Streptomyces sp. Root1319] back to top

previous - next

#### Architecture

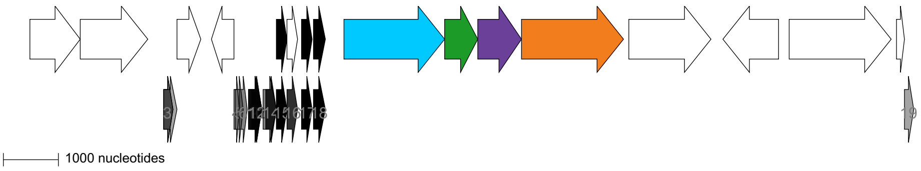

Link to nucleotide sequence

| Accession | start | end | direction | length (aa) | Pfam/HMM | name | description | E-value |
|---|---|---|---|---|---|---|---|---|
| WP_056795368.1 | 236292 | 237198 | + | 301 | NO PFAM MATCH | - | - | - |
| WP_056795366.1 | 237194 | 238433 | + | 412 | PF01547<br>PF13416<br>PF13531 | SBP_bac_1<br>SBP_bac_8<br>SBP_bac_11 | Bacterial extracellular solute-binding protein<br>Bacterial extracellular solute-binding protein<br>Bacterial extracellular solute-binding protein | 1.10E-14<br>1.60E-07<br>3.50E-05 |
| WP_015036771.1 | 238967 | 239400 | + | 144 | INFERRED GENE | - | - | - |
| WP_056796012.1 | 240005 | 239582 | - | 140 | PF12647 | RNHCP | RNHCP domain | 1.70E-34 |
| WP_056795364.1 | 240777 | 240963 | + | 61 | LinA | LinA | LinA (linaridin precursor) | 1.40E-08 |
| WP_056795361.1 | 240959 | 241154 | + | 64 | NO PFAM MATCH | - | - | - |
| WP_056795357.1 | 241215 | 241410 | + | 64 | LinA | LinA | LinA (linaridin precursor) | 1.10E-09 |
| WP_056795354.1 | 241453 | 241669 | + | 71 | LinA | LinA | LinA (linaridin precursor) | 1.30E-08 |
| WP_056796007.1 | 241999 | 243826 | + | 608 | LinH<br>LinG<br>LinE<br>PF12697<br>PF00561 | LinH<br>LinG<br>LinE<br>Abhydrolase_6<br>Abhydrolase_1 | LinH (Fusion of LinE/LinG)<br>LinG Homolog<br>LinE Homolog<br>Alpha/beta hydrolase family<br>alpha/beta hydrolase fold | 1.50E-234<br>4.60E-112<br>7.60E-90<br>6.10E-14<br>3.30E-11 |
| WP_056795351.1 | 243822 | 244431 | + | 202 | LinL | LinL | LinL | 3.80E-60 |
| WP_056795348.1 | 244427 | 245234 | + | 268 | PF13649<br>PF08241<br>PF13489<br>PF13847<br>PF08242 | Methyltransf_25<br>Methyltransf_11<br>Methyltransf_23<br>Methyltransf_31<br>Methyltransf_12 | Methyltransferase domain<br>Methyltransferase domain<br>Methyltransferase domain<br>Methyltransferase domain<br>Methyltransferase domain | 9.40E-17<br>5.50E-11<br>9.00E-08<br>9.20E-08<br>1.30E-07 |
| WP_056795344.1 | 245230 | 247075 | + | 614 | TIGR02204<br>TIGR02203<br>TIGR00958<br>TIGR03796<br>TIGR01846 | TIGR02204<br>TIGR02203<br>TIGR00958<br>TIGR03796<br>TIGR01846 | MsbA_rel: ABC transporter, permease/ATP-binding protein<br>MsbA_lipidA: lipid A export permease/ATP-binding protein MsbA<br>3a01208: antigen peptide transporter 2<br>NHLM_micro_ABC1: NHLM bacteriocin system ABC transporter, peptidase/ATP-binding protein<br>type_I_sec_HlyB: type I secretion system ATPase | 1.20E-100<br>3.50E-92<br>2.40E-85<br>8.30E-84<br>1.60E-80 |
| WP_056796004.1 | 247178 | 248669 | + | 496 | TIGR02734<br>PF13450<br>TIGR03467<br>TIGR01317<br>PF01593 | TIGR02734<br>NAD_binding_8<br>TIGR03467<br>TIGR01317<br>Amino_oxidase | crtI_fam: phytoene desaturase<br>NAD(P)-binding Rossmann-like domain<br>HpnE: squalene-associated FAD-dependent desaturase<br>GOGAT_sm_gam: glutamate synthase, NADH/NADPH, small subunit<br>Flavin containing amine oxidoreductase | 6.60E-11<br>2.00E-10<br>6.90E-10<br>6.40E-09<br>1.10E-08 |
| WP_056795340.1 | 249895 | 248887 | - | 335 | NO PFAM MATCH | - | - | - |
| WP_056796001.1 | 250099 | 251935 | + | 611 | NO PFAM MATCH | - | - | - |
| WP_076611639.1 | 252051 | 252192 | + | 47 | INFERRED GENE | - | - | - |

| index | leader | core | start | end | dir | score |
|---|---|---|---|---|---|---|
| 1 | VWCGTGSRCRSWSSSAVR | STPRPHAREGPHPGRFGFAARFEEP...RRPASCDRVRRLMERTRITDRYRMT | 238720 | 238966 | + | 6 |

| | | | | | | |
|---|---|---|---|---|---|---|
| 2 | VVRHRESVPQLEFVGCQVDS | ATACTRRSSSWTVRFCGTVRRTCLRTSGRRT | 238722 | 238878 | + | 6 |
| 3 | VRHRESVPQLEFVGCQVDS | ATACTRRSSSWTVRFCGTVRRTCLRTSGRRT | 238725 | 238878 | + | 6 |
| 4 | VLWANPMGYGDAYRTGT | RRRRHWADTAMHASAGDC | 239985 | 240093 | + | 6 |
| 5 | MRTERAPVGVVTGQTRQCMHRQ | GTVDAQCAGRARASSSARRTGTH | 240016 | 240154 | + | 6 |
| 6 | LTHNAQAGPGPVRLLGVLGLTDDTSQ | SCTGRSWPWAWKASPVIRPDWITGPC | 240089 | 240248 | + | 7 |
| 7 | LTLLCTGTGRAADVLTSRWRGQHLVQLVAGDGGRLQKPVQV | GVPSVRDLTGGLRVADVGGKQPGPCGVQVGDRGAACTGT | 240257 | 240500 | + | 7 |
| 8 | LLCTGTGRAADVLTSRWRGQHLVQLVAGDGGRLQKPVQV | GVPSVRDLTGGLRVADVGGKQPGPCGVQVGDRGAACTGT | 240263 | 240500 | + | 7 |
| 9 | LCTGTGRAADVLTSRWRGQHLVQLVAGDGGRLQKPVQV | GVPSVRDLTGGLRVADVGGKQPGPCGVQVGDRGAACTGT | 240266 | 240500 | + | 7 |
| 10 | VLTSRWRGQHLVQLVAGDGGRLQKPVQV | GVPSVRDLTGGLRVADVGGKQPGPCGVQVGDRGAACTGT | 240296 | 240500 | + | 7 |
| 11 | LTSRWRGQHLVQLVAGDGGRLQKPVQV | GVPSVRDLTGGLRVADVGGKQPGPCGVQVGDRGAACTGT | 240299 | 240500 | + | 7 |
| 12 | VQLVAGDGGRLQKPVQV | GVPSVRDLTGGLRVADVGGKQPGPCGVQVGDRGAACTGT | 240329 | 240500 | + | 7 |
| 13 | MRLSRPPRTAPAPVAPCLHHPAGGP...SCQPTVFPRAGGPAIVDGTDGTPPG | RTPRVRGAALPYGPVAGASGCRPG | 240533 | 240764 | + | 7 |
| 14 | VAPCLHHPAGGPAASCQPTVFPRAGGPAIVDGTDGTPPG | RTPRVRGAALPYGPVAGASGCRPG | 240572 | 240764 | + | 9 |
| 15 | MHQVRTPRWAAVGPLDERTEGQAFGGAAPIA | ATQVIVTVAAAVVVAYAAGAIIGSTEEPSL | 240777 | 240963 | + | 19 |
| 16 | MTGTRWATASVCP | ATAVTTCALSENDAGHTGPAASPIA...VLVVVAEVCASPAISVIAGTHEPQL | 240959 | 241154 | + | 9 |
| 17 | MNASVFQTVPAAARPLTEEDPGYTATGTAPVA | ATPVLATAVFAAGFVAGVVVCDAIGDTSPVEQ | 241215 | 241410 | + | 30 |
| 18 | MNDTSFLTRPGALTDIRVDDPQGRVIINHPPVM | ATPAVVSGVIIGVALVNAFVAGRDGGPLEPPIEIDVPA | 241453 | 241669 | + | 29 |
| 19 | VGPGRLRCRSAARAWSVPRPRRARAASPRPVSR | AAPPCAPGRARRYGRVP | 252196 | 252349 | + | 6 |

### Results for KRA40986.1 [Streptomyces sp. Root63] back to top

previous - next

#### Architecture

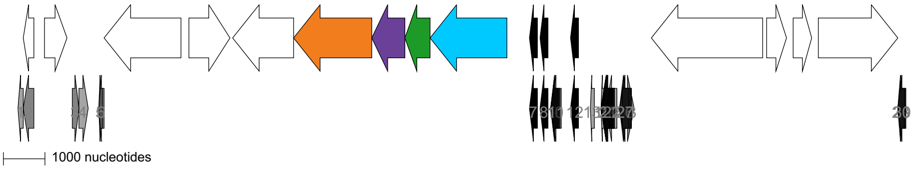

Link to nucleotide sequence

| Accession | start | end | direction | length (aa) | Pfam/HMM | name | description | E-value |
|---|---|---|---|---|---|---|---|---|
| KRA40978.1 | 935622 | 935373 | - | 82 | NO PFAM MATCH | - | - | - |
| KRA40979.1 | 935880 | 936417 | + | 178 | PF12840<br>PF01022 | HTH_20<br>HTH_5 | Helix-turn-helix domain<br>Bacterial regulatory protein, arsR family | 2.80E-10<br>3.30E-05 |
| KRA40980.1 | 939154 | 937300 | - | 617 | NO PFAM MATCH | - | - | - |
| KRA40981.1 | 939343 | 940357 | + | 337 | NO PFAM MATCH | - | - | - |
| KRA40982.1 | 941881 | 940411 | - | 489 | TIGR02734<br>PF13450<br>TIGR01317<br>TIGR00292<br>TIGR03467 | TIGR02734<br>NAD_binding_8<br>TIGR01317<br>TIGR00292<br>TIGR03467 | crtI_fam: phytoene desaturase<br>NAD(P)-binding Rossmann-like domain<br>GOGAT_sm_gam: glutamate synthase, NADH/NADPH, small subunit<br>TIGR00292: thiazole biosynthesis enzyme<br>HpnE: squalene-associated FAD-dependent desaturase | 3.60E-11<br>5.10E-09<br>1.00E-08<br>1.30E-08<br>7.50E-08 |
| KRA40983.1 | 943749 | 941877 | - | 623 | TIGR02204<br>TIGR02203<br>TIGR00958<br>TIGR03796<br>TIGR03375 | TIGR02204<br>TIGR02203<br>TIGR00958<br>TIGR03796<br>TIGR03375 | MsbA_rel: ABC transporter, permease/ATP-binding protein<br>MsbA_lipidA: lipid A export permease/ATP-binding protein MsbA<br>3a01208: antigen peptide transporter 2<br>NHLM_micro_ABC1: NHLM bacteriocin system ABC transporter, peptidase/ATP-binding protein<br>type_I_sec_LssB: type I secretion system ATPase | 1.20E-103<br>9.90E-97<br>8.80E-85<br>8.50E-81<br>2.20E-80 |
| KRA40984.1 | 944555 | 943745 | - | 269 | PF13649<br>PF08241<br>PF08242<br>PF13489<br>PF13847 | Methyltransf_25<br>Methyltransf_11<br>Methyltransf_12<br>Methyltransf_23<br>Methyltransf_31 | Methyltransferase domain<br>Methyltransferase domain<br>Methyltransferase domain<br>Methyltransferase domain<br>Methyltransferase domain | 2.30E-16<br>7.50E-12<br>3.30E-08<br>2.70E-07<br>1.80E-06 |
| KRA40985.1 | 945160 | 944551 | - | 202 | LinL | LinL | LinL | 2.80E-65 |
| KRA40986.1 | 947004 | 945156 | - | 615 | LinH<br>LinG<br>LinE<br>LinL<br>PF12697 | LinH<br>LinG<br>LinE<br>LinL<br>Abhydrolase_6 | LinH (Fusion of LinE/LinG)<br>LinG Homolog<br>LinE Homolog<br>LinL<br>Alpha/beta hydrolase family | 3.80E-253<br>1.40E-111<br>3.40E-95<br>1.20E-36<br>5.80E-15 |
| KRA40987.1 | 947753 | 947552 | - | 66 | LinA | LinA | LinA (linaridin precursor) | 3.70E-14 |
| KRA40988.1 | 948005 | 947810 | - | 64 | LinA | LinA | LinA (linaridin precursor) | 7.70E-11 |
| KRA40989.1 | 948746 | 948542 | - | 67 | LinA | LinA | LinA (linaridin precursor) | 8.70E-12 |

| | | | | | | | | |
|---|---|---|---|---|---|---|---|---|
| KRA40990.1 | 953155 | 950500 | - | 884 | PF13191<br>TIGR03903<br>PF13424<br>PF07719<br>PF13401 | AAA_16<br>TIGR03903<br>TPR_12<br>TPR_2<br>AAA_22 | AAA ATPase domain<br>TOMM_kin_cyc: TOMM system kinase/cyclase fusion protein<br>Tetratricopeptide repeat<br>Tetratricopeptide repeat<br>AAA domain | 4.20E-20<br>4.90E-19<br>4.80E-05<br>5.50E-05<br>8.70E-05 |
| KRA40991.1 | 953265 | 953745 | + | 159 | PF03745 | DUF309 | Domain of unknown function (DUF309) | 6.20E-20 |
| KRA41431.1 | 953884 | 954349 | + | 154 | NO PFAM MATCH | - | - | - |
| KRA40992.1 | 954498 | 956424 | + | 641 | PF10022 | DUF2264 | Uncharacterized protein conserved in bacteria (DUF2264) | 4.90E-100 |

| index | leader | core | start | end | dir | score |
|---|---|---|---|---|---|---|
| 1 | VDDVPLAPGGLVVRGPG | TEAAARWRRVPMSGLTSP | 935358 | 935250 | - | 6 |
| 2 | MIKNFVRGFSALSLALVPLL | SAPAAQAAPAAEVTTLTDAVGPVQV...SHGVGVIALTPYGVAHGVPTGSYRV | 935622 | 935373 | - | 7 |
| 3 | LTGCSRAGPQRVAAAASASRPRPPSRAG | ACSASSAVVPTRITDIVKAVVVLHHAST | 936541 | 936712 | + | 7 |
| 4 | MRPKRAPSSLAQHRFWELPYKQTPFAFRT | QTPEPSSASPAGASTSCSPCSATAPSTNPEPRTPTVTLAA | 936708 | 936918 | + | 6 |
| 5 | MTAAAVGGGPGASAVDL | ARLSPAGTRSCGTVTKV | 937289 | 937184 | - | 7 |
| 6 | MRGAGMTAAAVGGGPGASA | VDLARLSPAGTRSCGTVTKV | 937304 | 937184 | - | 6 |
| 7 | MSMSPTPAALRGTGGLSESDPGRALSSLAPVA | ATPGVVAGVAVGVALVNAFAAGYNHCGGNVELPM | 947753 | 947552 | - | 31 |
| 8 | MNKSSAPAVLTATGALTESAPGVTLPTLAPVA | ATPVAIAATMGVAFVAGYAAGRAATGNVELPM | 948005 | 947810 | - | 30 |
| 9 | LNTASIPAVLSETGALSESDHGRAPLD | TVPVASVTFTMTACVEVSVCLTGSVIQLPQ | 948249 | 948075 | - | 9 |
| 10 | MTSPHRPPTSEEKSLNTASIPAVLSETGALSESDHGRAPLD | TVPVASVTFTMTACVEVSVCLTGSVIQLPQ | 948291 | 948075 | - | 8 |
| 11 | VGPSRRRRRLPMTSPHRPPTSEEKS...TASIPAVLSETGALSESDHGRAPLD | TVPVASVTFTMTACVEVSVCLTGSVIQLPQ | 948324 | 948075 | - | 7 |
| 12 | MPEFREPGWTRGIAPLQEDAGGQVFGGASPVA | ATPAVVATAGAVVVAFAAGVAARHLANGGNVELPM | 948746 | 948542 | - | 30 |
| 13 | MATLKGCRLLRKLRCST | NRLTDLVKAVLALHHASA | 949111 | 949003 | - | 6 |
| 14 | MQLLPDAGVVPVPQ | PTPARHARPEAQLLWQVLPLDAGVRHE | 949385 | 949259 | - | 6 |
| 15 | VIRWCFDPVLPRSAKLGPVRLPLYG...GRRVELGRQLLMQLLPDAGVVPVPQ | PTPARHARPEAQLLWQVLPLDAGVRHE | 949538 | 949259 | - | 7 |
| 16 | VPCPSVIRWCFDPVLPRSAKLGPVR...GRRVELGRQLLMQLLPDAGVVPVPQ | PTPARHARPEAQLLWQVLPLDAGVRHE | 949553 | 949259 | - | 7 |
| 17 | VRIRASGVPCPSVIRWCFDPVLPRS...GRRVELGRQLLMQLLPDAGVVPVPQ | PTPARHARPEAQLLWQVLPLDAGVRHE | 949574 | 949259 | - | 6 |
| 18 | LPPVRIRASGVPCPSVIRWCFDPVL...GRRVELGRQLLMQLLPDAGVVPVPQ | PTPARHARPEAQLLWQVLPLDAGVRHE | 949583 | 949259 | - | 6 |
| 19 | LRLPPVRIRASGVPCPSVIRWCFDP...GRRVELGRQLLMQLLPDAGVVPVPQ | PTPARHARPEAQLLWQVLPLDAGVRHE | 949589 | 949259 | - | 6 |
| 20 | VTSLRLPPVRIRASGVPCPSVIRWC...GRRVELGRQLLMQLLPDAGVVPVPQ | PTPARHARPEAQLLWQVLPLDAGVRHE | 949598 | 949259 | - | 6 |
| 21 | MSCVTSLRLPPVRIRASGVPCPSVI...GRRVELGRQLLMQLLPDAGVVPVPQ | PTPARHARPEAQLLWQVLPLDAGVRHE | 949607 | 949259 | - | 6 |
| 22 | MPDPAPDRWGGVDQPHELRDVVAVAACEDQGERSAVSVGDQVVLRPGS | ATICQTRSRPAPPLWRARGWSRPPHATSRDGPPR | 949653 | 949404 | - | 6 |
| 23 | VDLDRAPSDHTRPGWADRQEHQGQA...LSPGPIRTPACSRAPEAEVSPPPSC | ATPHTGAPWSPSSATSICAATTSGTPR | 949784 | 950072 | + | 7 |
| 24 | VNRRASGWGRGTGCAGRRGADLLDDRNLAGPLALGALVG | QPTRGGCGLTAHGPDPLGGVDVSDAAAGDLPDPCGGTRG | 949950 | 949713 | - | 6 |
| 25 | VEIPPPRGPVNRRASGWGRGTGCAGRRGADLLDDRNLAGPLAL | GALVGQPTRGGCGLTAHGPDPLGGVDVSDAAAGDLPDPCGGTRG | 949977 | 949713 | - | 6 |
| 26 | VASRRTAVEIPPPRGPVNRRASGWGRGTGCAGRRGADLLDDRNLAGP | LALGALVGQPTRGGCGLTAHGPDPLGGVDVSDAAAGDLPDPCGGTRG | 949998 | 949713 | - | 6 |
| 27 | VPQCVASRRTAVEIPPPRGPVNRRASGWGRGTGCAGRRGADLLDDRNLA | GPLALGALVGQPTRGGCGLTAHGPDPLGGVDVSDAAAGDLPDPCGGTRG | 950010 | 949713 | - | 6 |
| 28 | VPEAVARLAPER | PTTLISGPSATSDIELERVEGVHGPRTLVVVIRTDA | 956575 | 956428 | - | 6 |
| 29 | VDLSAVEAGVPEAVARLAPER | PTTLISGPSATSDIELERVEGVHGPRTLVVVIRTDA | 956602 | 956428 | - | 8 |
| 30 | VVDLSAVEAGVPEAVARLAPER | PTTLISGPSATSDIELERVEGVHGPRTLVVVIRTDA | 956605 | 956428 | - | 8 |

### Results for WP_161264539.1 [Streptomyces sp. SID2119] back to top

previous - next

#### Architecture

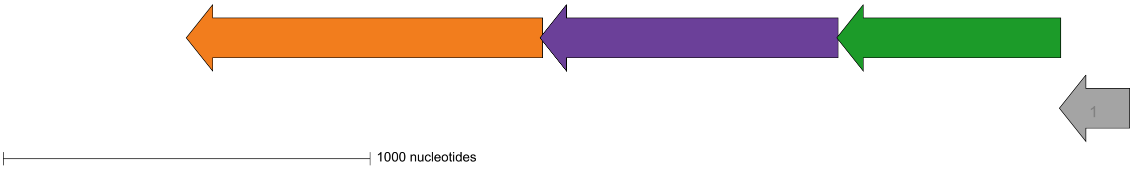

Link to nucleotide sequence

| Accession | start | end | direction | length (aa) | Pfam/HMM | name | description | E-value |
|---|---|---|---|---|---|---|---|---|

| | | | | | | | | |
|---|---|---|---|---|---|---|---|---|
| WP_161264536.1 | 971 | 0 | - | 324 | PF00664<br>TIGR02204<br>TIGR02203<br>TIGR00958<br>TIGR03796 | ABC_membrane<br>TIGR02204<br>TIGR02203<br>TIGR00958<br>TIGR03796 | ABC transporter transmembrane region<br>MsbA_rel: ABC transporter, permease/ATP-binding protein<br>MsbA_lipidA: lipid A export permease/ATP-binding protein MsbA<br>3a01208: antigen peptide transporter 2<br>NHLM_micro_ABC1: NHLM bacteriocin system ABC transporter, peptidase/ATP-binding protein | 2.10E-34<br>7.70E-25<br>3.70E-20<br>5.30E-16<br>6.60E-14 |
| WP_161264537.1 | 1780 | 967 | - | 270 | PF13649<br>PF08241<br>PF13847<br>PF08242<br>PF13489 | Methyltransf_25<br>Methyltransf_11<br>Methyltransf_31<br>Methyltransf_12<br>Methyltransf_23 | Methyltransferase domain<br>Methyltransferase domain<br>Methyltransferase domain<br>Methyltransferase domain<br>Methyltransferase domain | 3.30E-16<br>4.70E-12<br>1.90E-07<br>2.60E-07<br>1.90E-06 |
| WP_161264538.1 | 2385 | 1776 | - | 202 | LinL | LinL | LinL | 7.10E-65 |

| index | leader | core | start | end | dir | score |
|---|---|---|---|---|---|---|
| 1 | VRSVVIEGADHDGILTDPQYAAETLRHLLAFLA | DTAGRGPGGPTGLLTGTEPSGLTASSEENR | 2573 | 2381 | - | 6 |

### Results for WP_161245905.1 [Streptomyces sp. SID1034] back to top

previous - next

#### Architecture

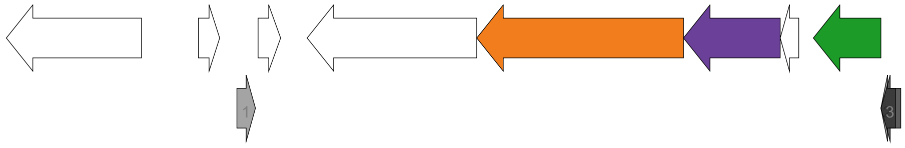

1000 nucleotides

Link to nucleotide sequence

| Accession | start | end | direction | length (aa) | Pfam/HMM | name | description | E-value |
|---|---|---|---|---|---|---|---|---|
| WP_161245899.1 | 6024 | 4845 | - | 392 | TIGR04022<br>PF08028<br>PF00441<br>PF02771 | TIGR04022<br>Acyl-CoA_dh_2<br>Acyl-CoA_dh_1<br>Acyl-CoA_dh_N | sulfur_SfnB: sulfur acquisition oxidoreductase, SfnB family<br>Acyl-CoA dehydrogenase, C-terminal domain<br>Acyl-CoA dehydrogenase, C-terminal domain<br>Acyl-CoA dehydrogenase, N-terminal domain | 1.80E-25<br>8.80E-21<br>5.40E-07<br>5.80E-04 |
| WP_005477046.1 | 6517 | 6703 | + | 62 | INFERRED GENE | - | - | - |
| WP_018560249.1 | 7039 | 7234 | + | 65 | INFERRED GENE | - | - | - |
| WP_161245900.1 | 8947 | 7465 | - | 493 | TIGR02734<br>PF13450<br>TIGR03467<br>TIGR02733<br>PF01593 | TIGR02734<br>NAD_binding_8<br>TIGR03467<br>TIGR02733<br>Amino_oxidase | crtI_fam: phytoene desaturase<br>NAD(P)-binding Rossmann-like domain<br>HpnE: squalene-associated FAD-dependent desaturase<br>desat_CrtD: C-3',4' desaturase CrtD<br>Flavin containing amine oxidoreductase | 3.60E-10<br>1.70E-09<br>4.10E-09<br>1.30E-07<br>9.30E-07 |
| WP_161245901.1 | 10746 | 8943 | - | 600 | TIGR02204<br>TIGR02203<br>TIGR00958<br>TIGR01846<br>TIGR03796 | TIGR02204<br>TIGR02203<br>TIGR00958<br>TIGR01846<br>TIGR03796 | MsbA_rel: ABC transporter, permease/ATP-binding protein<br>MsbA_lipidA: lipid A export permease/ATP-binding protein MsbA<br>3a01208: antigen peptide transporter 2<br>type_I_sec_HlyB: type I secretion system ATPase<br>NHLM_micro_ABC1: NHLM bacteriocin system ABC transporter, peptidase/ATP-binding protein | 2.00E-103<br>2.70E-95<br>2.30E-88<br>6.40E-85<br>1.10E-82 |
| WP_161245902.1 | 11582 | 10742 | - | 279 | PF13649<br>PF08241<br>PF08242<br>PF13847<br>PF13489 | Methyltransf_25<br>Methyltransf_11<br>Methyltransf_12<br>Methyltransf_31<br>Methyltransf_23 | Methyltransferase domain<br>Methyltransferase domain<br>Methyltransferase domain<br>Methyltransferase domain<br>Methyltransferase domain | 4.10E-15<br>5.90E-11<br>7.20E-09<br>3.00E-07<br>3.40E-07 |
| WP_161245903.1 | 11740 | 11578 | - | 53 | NO PFAM MATCH | - | - | - |
| WP_161245904.1 | 12458 | 11867 | - | 196 | LinL | LinL | LinL | 1.40E-67 |

| index | leader | core | start | end | dir | score |
|---|---|---|---|---|---|---|
| 1 | VARAWSASLSKRGEVKSKTGPHPGMG | SGPVLVTRQRPSTFTRATATARSPTAR | 6854 | 7016 | + | 6 |
| 2 | MSRYAHQVADAIKEFFAQGPGG | GDATADADDARTAGTATDKETAR | 12592 | 12454 | - | 6 |
| 3 | VVVEGAGHDTLLTMSRYAHQVADAIKEFF | AQGPGGGDATADADDARTAGTATDKETAR | 12631 | 12454 | - | 8 |

### Results for QHY99994.1 [Streptomyces sp. S4.7] back to top

previous - next

#### Architecture

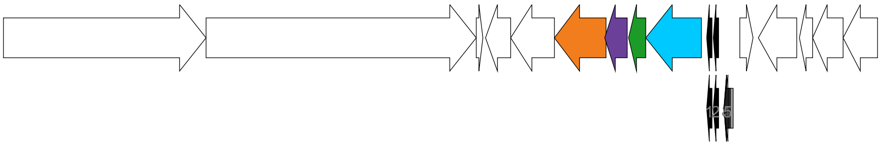

1000 nucleotides

Link to nucleotide sequence

| Accession | start | end | direction | length (aa) | Pfam/HMM | name | description | E-value |
|---|---|---|---|---|---|---|---|---|

| | | | | | | | | |
|---|---|---|---|---|---|---|---|---|
| QHY99986.1 | 7468167 | 7475088 | + | 2306 | PF00668<br>TIGR01733<br>PF00501<br>TIGR01734<br>TIGR03098 | Condensation<br>TIGR01733<br>AMP-binding<br>TIGR01734<br>TIGR03098 | Condensation domain<br>AA-adenyl-dom: amino acid adenylation domain<br>AMP-binding enzyme<br>D-ala-DACP-lig: D-alanine--poly(phosphoribitol) ligase, subunit 1<br>ligase_PEP_1: acyl-CoA ligase (AMP-forming), exosortase A-associated | 2.10E-131<br>7.70E-127<br>1.40E-92<br>3.20E-73<br>1.10E-60 |
| QHY99987.1 | 7475084 | 7484300 | + | 3071 | TIGR01733<br>PF00668<br>PF00501<br>TIGR01734<br>TIGR03443 | TIGR01733<br>Condensation<br>AMP-binding<br>TIGR01734<br>TIGR03443 | AA-adenyl-dom: amino acid adenylation domain<br>Condensation domain<br>AMP-binding enzyme<br>D-ala-DACP-lig: D-alanine--poly(phosphoribitol) ligase, subunit 1<br>alpha_am_amid: L-aminoadipate-semialdehyde dehydrogenase | 1.30E-246<br>2.00E-214<br>1.70E-178<br>2.90E-127<br>1.30E-115 |
| QHY99988.1 | 7484296 | 7484518 | + | 73 | PF03621 | MbtH | MbtH-like protein | 1.40E-28 |
| QHY99989.1 | 7485461 | 7484609 | - | 283 | PF12680 | SnoaL_2 | SnoaL-like domain | 2.30E-06 |
| QHY99990.1 | 7486942 | 7485457 | - | 494 | TIGR02734<br>TIGR01317<br>TIGR02733<br>TIGR03467<br>TIGR00292 | TIGR02734<br>TIGR01317<br>TIGR02733<br>TIGR03467<br>TIGR00292 | crtI_fam: phytoene desaturase<br>GOGAT_sm_gam: glutamate synthase, NADH/NADPH, small subunit<br>desat_CrtD: C-3',4' desaturase CrtD<br>HpnE: squalene-associated FAD-dependent desaturase<br>TIGR00292: thiazole biosynthesis enzyme | 4.70E-13<br>5.20E-09<br>7.60E-09<br>1.30E-08<br>2.00E-08 |
| QHY99991.1 | 7488681 | 7486938 | - | 580 | TIGR02204<br>TIGR02203<br>TIGR00958<br>TIGR03796<br>TIGR01846 | TIGR02204<br>TIGR02203<br>TIGR00958<br>TIGR03796<br>TIGR01846 | MsbA_rel: ABC transporter, permease/ATP-binding protein<br>MsbA_lipidA: lipid A export permease/ATP-binding protein MsbA<br>3a01208: antigen peptide transporter 2<br>NHLM_micro_ABC1: NHLM bacteriocin system ABC transporter, peptidase/ATP-binding protein<br>type_I_sec_HlyB: type I secretion system ATPase | 3.60E-103<br>1.00E-95<br>2.20E-89<br>1.30E-82<br>5.80E-79 |
| QHY99992.1 | 7489436 | 7488677 | - | 252 | PF13649<br>PF08241<br>PF13489<br>PF08242<br>PF13847 | Methyltransf_25<br>Methyltransf_11<br>Methyltransf_23<br>Methyltransf_12<br>Methyltransf_31 | Methyltransferase domain<br>Methyltransferase domain<br>Methyltransferase domain<br>Methyltransferase domain<br>Methyltransferase domain | 4.80E-14<br>3.70E-09<br>7.40E-08<br>8.00E-08<br>2.50E-07 |
| QHY99993.1 | 7490064 | 7489485 | - | 192 | LinL | LinL | LinL | 3.70E-64 |
| QHY99994.1 | 7491968 | 7490060 | - | 635 | LinH<br>LinG<br>LinE<br>PF12697<br>PF00561 | LinH<br>LinG<br>LinE<br>Abhydrolase_6<br>Abhydrolase_1 | LinH (Fusion of LinE/LinG)<br>LinG Homolog<br>LinE Homolog<br>Alpha/beta hydrolase family<br>alpha/beta hydrolase fold | 1.00E-236<br>7.00E-112<br>5.50E-98<br>1.10E-15<br>2.30E-13 |
| QHY99995.1 | 7492330 | 7492138 | - | 63 | LinA | LinA | LinA (linaridin precursor) | 1.60E-09 |
| QHY99996.1 | 7492551 | 7492359 | - | 63 | LinA | LinA | LinA (linaridin precursor) | 4.80E-15 |
| QHY99997.1 | 7493249 | 7493702 | + | 150 | PF12840<br>PF01022<br>PF13518<br>PF13338 | HTH_20<br>HTH_5<br>HTH_28<br>AbiEi_4 | Helix-turn-helix domain<br>Bacterial regulatory protein, arsR family<br>Helix-turn-helix domain<br>Transcriptional regulator, AbiEi antitoxin | 3.20E-11<br>1.80E-06<br>1.90E-04<br>8.30E-04 |
| QHY99998.1 | 7495202 | 7493882 | - | 439 | TIGR00900<br>PF07690<br>TIGR00881<br>PF05977<br>TIGR00880 | TIGR00900<br>MFS_1<br>TIGR00881<br>MFS_3<br>TIGR00880 | 2A0121: H+ Antiporter protein<br>Major Facilitator Superfamily<br>2A0104: phosphoglycerate transporter family protein<br>Transmembrane secretion effector<br>2_A_01_02: multidrug resistance protein | 3.20E-37<br>1.80E-30<br>1.20E-13<br>2.60E-13<br>3.90E-12 |
| QHY99999.1 | 7495737 | 7495272 | - | 154 | NO PFAM MATCH | - | - | - |
| QHZ00001.1 | 7496774 | 7495733 | - | 346 | PF01648<br>TIGR00556 | ACPS<br>TIGR00556 | 4'-phosphopantetheinyl transferase superfamily<br>pantethn_trn: phosphopantetheine--protein transferase domain | 4.20E-06<br>6.20E-04 |
| QHZ00002.1 | 7497927 | 7496754 | - | 390 | PF00441<br>PF02770<br>TIGR03207<br>TIGR03204<br>PF02771 | Acyl-CoA_dh_1<br>Acyl-CoA_dh_M<br>TIGR03207<br>TIGR03204<br>Acyl-CoA_dh_N | Acyl-CoA dehydrogenase, C-terminal domain<br>Acyl-CoA dehydrogenase, middle domain<br>cyc_hxne_CoA_dh: cyclohexanecarboxyl-CoA dehydrogenase<br>pimC_large: pimeloyl-CoA dehydrogenase, large subunit<br>Acyl-CoA dehydrogenase, N-terminal domain | 2.30E-20<br>8.10E-18<br>4.00E-15<br>7.30E-12<br>1.00E-06 |

| index | leader | core | start | end | dir | score |
|---|---|---|---|---|---|---|
| 1 | MSRIIESVAAITPEHLTDGSHGEVTAMAPVQ | ATPGLVAFGIGFAGGAGVSWVGVQAYEAGAND | 7492330 | 7492138 | - | 31 |
| 2 | MATIAEFMNDQKLGVLEQGYKEETQEPVL | ATPAAAVATCWYVGGAAAGVTVVAGAYVAGRVFG | 7492551 | 7492359 | - | 30 |
| 3 | LGGLSIDPFFAPRGNGGQRMNAAC | VTPRLQCDVHHIGKALIRQGPPVLLWLPQSQAFNQC | 7492881 | 7492698 | - | 8 |
| 4 | LFQIPHHHRATSGFSPLGGLSIDPFFAPRGNGGQRMNAAC | VTPRLQCDVHHIGKALIRQGPPVLLWLPQSQAFNQC | 7492929 | 7492698 | - | 7 |
| 5 | VTDSGQPGISRAWILTQNAIRFRPR...PLGGLSIDPFFAPRGNGGQRMNAAC | VTPRLQCDVHHIGKALIRQGPPVLLWLPQSQAFNQC | 7493019 | 7492698 | - | 6 |

### Results for ARI51159.1 [Streptomyces sp. S8] back to top

previous - next

#### Architecture

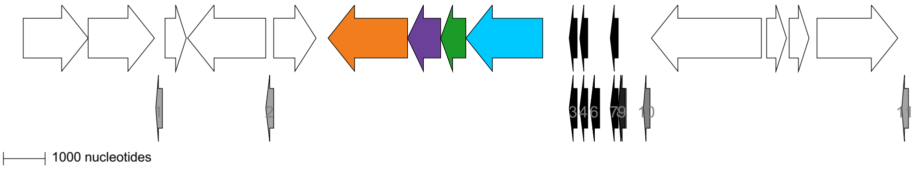

Link to nucleotide sequence

| Accession | start | end | direction | length (aa) | Pfam/HMM | name | description | E-value |
|---|---|---|---|---|---|---|---|---|

| | | | | | | | | |
|---|---|---|---|---|---|---|---|---|
| ARI51151.1 | 705695 | 707240 | + | 514 | PF00501<br>TIGR03208<br>TIGR03098<br>TIGR02275<br>TIGR01733 | AMP-binding<br>TIGR03208<br>TIGR03098<br>TIGR02275<br>TIGR01733 | AMP-binding enzyme<br>cyc_hxne_CoA_lg: cyclohexanecarboxylate-CoA ligase<br>ligase_PEP_1: acyl-CoA ligase (AMP-forming), exosortase A-associated<br>DHB_AMP_lig: (2,3-dihydroxybenzoyl)adenylate synthase<br>AA-adenyl-dom: amino acid adenylation domain | 4.40E-68<br>4.40E-43<br>1.10E-42<br>1.10E-38<br>4.00E-38 |
| ARI51152.1 | 707247 | 708840 | + | 530 | PF00501<br>TIGR03205<br>TIGR03098<br>TIGR02262<br>TIGR03208 | AMP-binding<br>TIGR03205<br>TIGR03098<br>TIGR02262<br>TIGR03208 | AMP-binding enzyme<br>pimA: dicarboxylate--CoA ligase PimA<br>ligase_PEP_1: acyl-CoA ligase (AMP-forming), exosortase A-associated<br>benz_CoA_lig: benzoate-CoA ligase family<br>cyc_hxne_CoA_lg: cyclohexanecarboxylate-CoA ligase | 8.00E-114<br>8.20E-79<br>4.00E-75<br>9.90E-75<br>9.00E-72 |
| ARI51153.1 | 709070 | 709607 | + | 178 | PF12840<br>PF01022<br>PF13518 | HTH_20<br>HTH_5<br>HTH_28 | Helix-turn-helix domain<br>Bacterial regulatory protein, arsR family<br>Helix-turn-helix domain | 3.60E-11<br>1.30E-06<br>1.00E-04 |
| ARI51154.1 | 711486 | 709632 | - | 617 | NO PFAM MATCH | - | - | - |
| ARI51155.1 | 711675 | 712689 | + | 337 | NO PFAM MATCH | - | - | - |
| ARI51156.1 | 714889 | 712987 | - | 633 | TIGR02204<br>TIGR02203<br>TIGR00958<br>TIGR03796<br>TIGR03797 | TIGR02204<br>TIGR02203<br>TIGR00958<br>TIGR03796<br>TIGR03797 | MsbA_rel: ABC transporter, permease/ATP-binding protein<br>MsbA_lipidA: lipid A export permease/ATP-binding protein MsbA<br>3a01208: antigen peptide transporter 2<br>NHLM_micro_ABC1: NHLM bacteriocin system ABC transporter, peptidase/ATP-binding protein<br>NHLM_micro_ABC2: NHLM bacteriocin system ABC transporter, ATP-binding protein | 5.10E-99<br>2.20E-91<br>2.10E-83<br>6.60E-82<br>1.70E-77 |
| ARI51157.1 | 715698 | 714885 | - | 270 | PF13649<br>PF08241<br>PF13847<br>PF13489<br>PF08242 | Methyltransf_25<br>Methyltransf_11<br>Methyltransf_31<br>Methyltransf_23<br>Methyltransf_12 | Methyltransferase domain<br>Methyltransferase domain<br>Methyltransferase domain<br>Methyltransferase domain<br>Methyltransferase domain | 4.70E-16<br>8.00E-12<br>6.80E-07<br>1.70E-06<br>1.80E-06 |
| ARI51158.1 | 716303 | 715694 | - | 202 | LinL | LinL | LinL | 7.10E-66 |
| ARI51159.1 | 718144 | 716299 | - | 614 | LinH<br>LinG<br>LinE<br>LinL<br>PF12697 | LinH<br>LinG<br>LinE<br>LinL<br>Abhydrolase_6 | LinH (Fusion of LinE/LinG)<br>LinG Homolog<br>LinE Homolog<br>LinL<br>Alpha/beta hydrolase family | 1.50E-251<br>2.00E-112<br>2.10E-95<br>3.90E-45<br>7.30E-14 |
| ARI51160.1 | 718960 | 718759 | - | 66 | LinA | LinA | LinA (linaridin precursor) | 2.40E-12 |
| ARI51161.1 | 719209 | 719017 | - | 63 | LinA | LinA | LinA (linaridin precursor) | 6.90E-09 |
| ARI51162.1 | 719948 | 719744 | - | 67 | LinA | LinA | LinA (linaridin precursor) | 3.10E-11 |
| ARI51163.1 | 723378 | 720723 | - | 884 | TIGR03903<br>PF13191<br>PF13424<br>PF07719<br>PF13401 | TIGR03903<br>AAA_16<br>TPR_12<br>TPR_2<br>AAA_22 | TOMM_kin_cyc: TOMM system kinase/cyclase fusion protein<br>AAA ATPase domain<br>Tetratricopeptide repeat<br>Tetratricopeptide repeat<br>AAA domain | 3.30E-20<br>7.10E-19<br>2.40E-05<br>9.30E-05<br>1.40E-04 |
| ARI51164.1 | 723488 | 723968 | + | 159 | PF03745 | DUF309 | Domain of unknown function (DUF309) | 4.90E-19 |
| ARI51165.1 | 724052 | 724517 | + | 154 | NO PFAM MATCH | - | - | - |
| ARI51166.1 | 724714 | 726649 | + | 644 | PF10022 | DUF2264 | Uncharacterized protein conserved in bacteria (DUF2264) | 9.70E-97 |

| index | leader | core | start | end | dir | score |
|---|---|---|---|---|---|---|
| 1 | VVSADGPSSAATADAWAERGRKHRRS | YRSGPVLTVVQVRCGPDGRPGPVRPG | 709014 | 708855 | - | 6 |
| 2 | VATSCRSPWPFPRTRPAGQRITSLPGEAGGEVS | AKSQASLAPGKFRMRDHLKERAPIAMTGGRAPK | 711700 | 711499 | - | 6 |
| 3 | MNMSPTPAALRGAGGLSESDPGRAHSTLVPVA | ATPGVLAGVAVGVALVNAFAAGYNHCGGNVELPM | 718960 | 718759 | - | 31 |
| 4 | MNTSSTPAVLTATGALTESDPGATLPAIAPVA | ATPVVVVAALGAGFVAGYAAGRAAGNVELPM | 719209 | 719017 | - | 30 |
| 5 | LSTASIPAVLSGTGALTESDRGRGPLD | TVPVTSVTVTMTACVGISVCLTGSTIELPQ | 719456 | 719282 | - | 10 |
| 6 | VTSTLPPTSEEKSLSTASIPAVLSGTGALTESDRGRGPLD | TVPVTSVTVTMTACVGISVCLTGSTIELPQ | 719495 | 719282 | - | 10 |
| 7 | MPEFRQPGWTRGLAPLDEGAGGQVFGGASPVA | ATPAVVATAGAVVVAFAAGVAARHLANGGNVELPM | 719948 | 719744 | - | 31 |
| 8 | MPAELSSLSVGLSASCGAHHRTS | VHAGAGSRVRSRAPGRAPVTAHP | 720098 | 719957 | - | 8 |
| 9 | VPEPAGCCCGVRILMPAELSSLSVGLSASC | GAHHRTSVHAGAGSRVRSRAPGRAPVTAHP | 720140 | 719957 | - | 8 |
| 10 | LTCAAARTTGGNRSGTWRSLVAHLTGGQ | GVAGSNPVVPTRESRRSGPVSEQSETGP | 720715 | 720544 | - | 7 |
| 11 | VPEAVARLVPER | PTTLISGPSATSDIELERVEGVHGPRRLVVVIRTDR | 726887 | 726740 | - | 6 |

### Results for WP_161184067.1 [Streptomyces sp. SID724] back to top

previous - next

#### Architecture

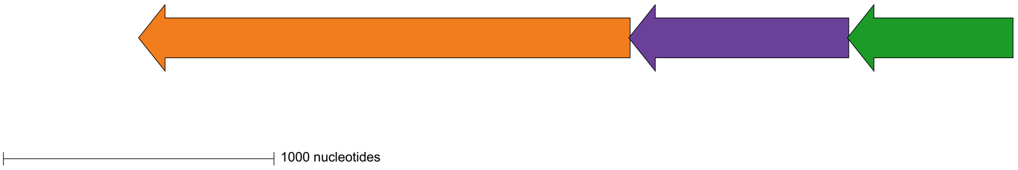

Link to nucleotide sequence

| Accession | start | end | direction | length (aa) | Pfam/HMM | name | description | E-value |
|---|---|---|---|---|---|---|---|---|

| | | | | | | | | |
|---|---|---|---|---|---|---|---|---|
| WP_161184066.1 | 1818 | 0 | - | 606 | TIGR02204<br>TIGR02203<br>TIGR00958<br>TIGR03796<br>TIGR03797 | TIGR02204<br>TIGR02203<br>TIGR00958<br>TIGR03796<br>TIGR03797 | MsbA_rel: ABC transporter, permease/ATP-binding protein<br>MsbA_lipidA: lipid A export permease/ATP-binding protein MsbA<br>3a01208: antigen peptide transporter 2<br>NHLM_micro_ABC1: NHLM bacteriocin system ABC transporter, peptidase/ATP-binding protein<br>NHLM_micro_ABC2: NHLM bacteriocin system ABC transporter, ATP-binding protein | 4.80E-102<br>2.90E-92<br>4.60E-85<br>1.20E-82<br>7.90E-77 |
| WP_003965565.1 | 2627 | 1814 | - | 270 | PF13649<br>PF08241<br>PF08242<br>PF13489<br>PF13847 | Methyltransf_25<br>Methyltransf_11<br>Methyltransf_12<br>Methyltransf_23<br>Methyltransf_31 | Methyltransferase domain<br>Methyltransferase domain<br>Methyltransferase domain<br>Methyltransferase domain<br>Methyltransferase domain | 2.30E-16<br>7.20E-12<br>5.40E-08<br>1.20E-07<br>1.60E-06 |
| WP_003965564.1 | 3232 | 2623 | - | 202 | LinL | LinL | LinL | 2.00E-65 |

| index | leader | core | start | end | dir | score |
|---|---|---|---|---|---|---|

### Results for WP_161184048.1 [Streptomyces sp. SID724] back to top

previous - next

#### Architecture

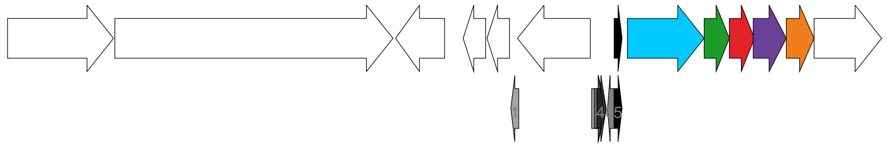

1000 nucleotides

Link to nucleotide sequence

| Accession | start | end | direction | length (aa) | Pfam/HMM | name | description | E-value |
|---|---|---|---|---|---|---|---|---|
| WP_161184045.1 | 0 | 2418 | + | 805 | PF08659<br>PF00550 | KR<br>PP-binding | KR domain<br>Phosphopantetheine attachment site | 5.30E-17<br>2.20E-13 |
| WP_161184046.1 | 2445 | 8751 | + | 2101 | TIGR02813<br>PF00109<br>TIGR01746<br>PF08659<br>TIGR03443 | TIGR02813<br>ketoacyl-synt<br>TIGR01746<br>KR<br>TIGR03443 | omega_3_PfaA: polyketide-type polyunsaturated fatty acid synthase PfaA<br>Beta-ketoacyl synthase, N-terminal domain<br>Thioester-redct: thioester reductase domain<br>KR domain<br>alpha_am_amid: L-aminoadipate-semialdehyde dehydrogenase | 1.90E-118<br>3.70E-86<br>2.70E-85<br>5.30E-64<br>7.50E-58 |
| WP_012381890.1 | 9921 | 8823 | - | 365 | PF03704<br>PF13428<br>PF00486<br>PF13432<br>PF08279 | BTAD<br>TPR_14<br>Trans_reg_C<br>TPR_16<br>HTH_11 | Bacterial transcriptional activator domain<br>Tetratricopeptide repeat<br>Transcriptional regulatory protein, C terminal<br>Tetratricopeptide repeat<br>HTH domain | 7.20E-45<br>8.00E-06<br>3.30E-04<br>6.20E-04<br>6.70E-04 |
| WP_003970685.1 | 10860 | 10365 | - | 164 | NO PFAM MATCH | - | - | - |
| WP_003970684.1 | 11418 | 10884 | - | 177 | PF01613<br>TIGR02296<br>TIGR03615 | Flavin_Reduct<br>TIGR02296<br>TIGR03615 | Flavin reductase like domain<br>HpaC: 4-hydroxyphenylacetate 3-monooxygenase, reductase component<br>RutF: pyrimidine utilization flavin reductase protein F | 8.90E-33<br>8.10E-24<br>1.80E-23 |
| WP_161184047.1 | 13248 | 11598 | - | 549 | PF03704<br>PF13191<br>TIGR03903<br>PF00486 | BTAD<br>AAA_16<br>TIGR03903<br>Trans_reg_C | Bacterial transcriptional activator domain<br>AAA ATPase domain<br>TOMM_kin_cyc: TOMM system kinase/cyclase fusion protein<br>Transcriptional regulatory protein, C terminal | 4.00E-43<br>5.20E-12<br>1.20E-07<br>2.70E-04 |
| WP_003970682.1 | 13770 | 13965 | + | 64 | LinA | LinA | LinA (linaridin precursor) | 6.80E-14 |
| WP_161184048.1 | 14081 | 15839 | + | 585 | LinH<br>LinG<br>LinE<br>PF12697<br>TIGR02427 | LinH<br>LinG<br>LinE<br>Abhydrolase_6<br>TIGR02427 | LinH (Fusion of LinE/LinG)<br>LinG Homolog<br>LinE Homolog<br>Alpha/beta hydrolase family<br>protocat_pcaD: 3-oxoadipate enol-lactonase | 4.40E-175<br>1.50E-76<br>9.90E-65<br>7.30E-06<br>3.30E-05 |
| WP_003970680.1 | 15828 | 16395 | + | 188 | LinL | LinL | LinL | 1.10E-51 |
| WP_003970679.1 | 16391 | 16964 | + | 190 | TIGR00521<br>TIGR02113<br>PF02441 | TIGR00521<br>TIGR02113<br>Flavoprotein | coaBC_dfp: phosphopantothenoylcysteine decarboxylase / phosphopantothenate--cysteine ligase<br>coaC_strep: phosphopantothenoylcysteine decarboxylase<br>Flavoprotein | 2.30E-14<br>3.70E-12<br>1.50E-11 |
| WP_012381886.1 | 16956 | 17694 | + | 245 | PF13649<br>PF08241<br>PF08242<br>TIGR02021<br>PF13489 | Methyltransf_25<br>Methyltransf_11<br>Methyltransf_12<br>TIGR02021<br>Methyltransf_23 | Methyltransferase domain<br>Methyltransferase domain<br>Methyltransferase domain<br>BchM-ChlM: magnesium protoporphyrin O-methyltransferase<br>Methyltransferase domain | 1.60E-16<br>2.20E-15<br>2.80E-11<br>1.20E-10<br>3.40E-10 |
| WP_003970677.1 | 17690 | 18323 | + | 210 | TIGR02857<br>TIGR01189<br>TIGR02673<br>TIGR02982<br>TIGR03258 | TIGR02857<br>TIGR01189<br>TIGR02673<br>TIGR02982<br>TIGR03258 | CydD: thiol reductant ABC exporter, CydD subunit<br>ccmA: heme ABC exporter, ATP-binding protein CcmA<br>FtsE: cell division ATP-binding protein FtsE<br>heterocyst_DevA: ABC exporter ATP-binding subunit, DevA family<br>PhnT: 2-aminoethylphosphonate ABC transport system, ATP-binding component PhnT | 5.70E-28<br>2.70E-25<br>1.60E-24<br>3.30E-22<br>1.10E-21 |
| WP_012381884.1 | 18319 | 19864 | + | 514 | NO PFAM MATCH | - | - | - |

| index | leader | core | start | end | dir | score |
|---|---|---|---|---|---|---|
| 1 | VRCRPRRRGGRSPAYGFRGRTRTVYGAVAR | RAPVWVTGYPGMRVREIPRSMFSV | 11602 | 11437 | - | 6 |
| 2 | VAAHDPGFRRSGLRIPAHRAGARSKPTQNSDKERTLLGGLGTCEASGAGT | PTDTRRTGGTTGRSSKPASSRNRPTGAWQLVQIQRDLRLVKLILMLRVGC | 13283 | 13586 | + | 6 |
| 3 | VQGLGPSPHKTRTRNGPCSGASAHARR | PAPGHRPTREGPGERPGVRRNRHRHEIARQARGNLSKFSGT | 13338 | 13545 | + | 6 |
| 4 | LLGGLGTCEASGAGTPTDTRRTGGTTGRSSKP | ASSRNRPTGAWQLVQIQRDLRLVKLILMLRVGC | 13388 | 13586 | + | 8 |
| 5 | MRLDSIATQETATALPESMATQDFANSVLAGAVPGFHSDAETPAM | ATPAVAQFVIQGSTICLVC | 13770 | 13965 | + | 34 |

| | | | | | | |
|---|---|---|---|---|---|---|
| 6 | LQSNRVAFRCPSSEPMLSERDLAVP | SCPSPAASSELNALCTTSATRSAGIPQSW | 13792 | 13627 | - | 7 |

### Results for WP_063743169.1 [Streptomyces sp. SID5476] back to top

previous - next

#### Architecture

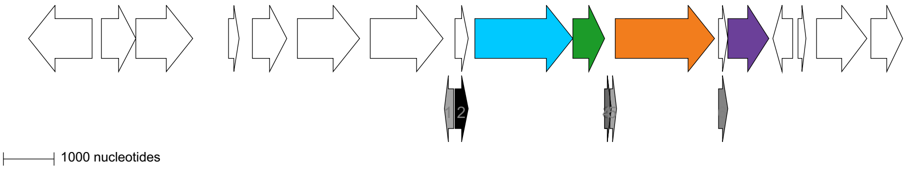

Link to nucleotide sequence

| Accession | start | end | direction | length (aa) | Pfam/HMM | name | description | E-value |
|---|---|---|---|---|---|---|---|---|
| WP_005474549.1 | 652826 | 651566 | - | 419 | PF04235<br>PF07786 | DUF418<br>DUF1624 | Protein of unknown function (DUF418)<br>Protein of unknown function (DUF1624) | 8.90E-10<br>7.20E-06 |
| WP_005474551.1 | 653016 | 653694 | + | 225 | PF00072<br>PF00196<br>TIGR02875<br>TIGR01818<br>TIGR03020 | Response_reg<br>GerE<br>TIGR02875<br>TIGR01818<br>TIGR03020 | Response regulator receiver domain<br>Bacterial regulatory proteins, luxR family<br>spore_0_A: sporulation transcription factor Spo0A<br>ntrC: nitrogen regulation protein NR(I)<br>EpsA: transcriptional regulator EpsA | 6.80E-24<br>1.50E-17<br>9.20E-17<br>4.40E-14<br>3.10E-11 |
| WP_005474553.1 | 653690 | 654812 | + | 373 | PF07730 | HisKA_3 | Histidine kinase | 6.80E-13 |
| WP_013005810.1 | 655521 | 655734 | + | 71 | INFERRED GENE | - | - | - |
| WP_005474556.1 | 655985 | 656669 | + | 227 | PF00072<br>PF00196<br>TIGR02875<br>TIGR03020<br>TIGR03541 | Response_reg<br>GerE<br>TIGR02875<br>TIGR03020<br>TIGR03541 | Response regulator receiver domain<br>Bacterial regulatory proteins, luxR family<br>spore_0_A: sporulation transcription factor Spo0A<br>EpsA: transcriptional regulator EpsA<br>reg_near_HchA: LuxR family transcriptional regulatory, chaperone HchA-associated | 2.00E-24<br>3.80E-18<br>8.40E-16<br>8.70E-13<br>1.50E-08 |
| WP_005474557.1 | 656882 | 658103 | + | 406 | PF07730 | HisKA_3 | Histidine kinase | 7.10E-14 |
| WP_005474559.1 | 658333 | 659746 | + | 470 | TIGR02734<br>PF13450<br>TIGR03467<br>TIGR03315<br>TIGR00292 | TIGR02734<br>NAD_binding_8<br>TIGR03467<br>TIGR03315<br>TIGR00292 | crtI_fam: phytoene desaturase<br>NAD(P)-binding Rossmann-like domain<br>HpnE: squalene-associated FAD-dependent desaturase<br>Se_ygfK: putative selenate reductase, YgfK subunit<br>TIGR00292: thiazole biosynthesis enzyme | 7.50E-14<br>2.80E-11<br>2.30E-09<br>7.00E-08<br>2.80E-07 |
| WP_005474562.1 | 659995 | 660247 | + | 83 | NO PFAM MATCH | - | - | - |
| WP_063743169.1 | 660392 | 662318 | + | 641 | LinH<br>LinG<br>LinE<br>PF12697<br>PF12146 | LinH<br>LinG<br>LinE<br>Abhydrolase_6<br>Hydrolase_4 | LinH (Fusion of LinE/LinG)<br>LinG Homolog<br>LinE Homolog<br>Alpha/beta hydrolase family<br>Serine aminopeptidase, S33 | 1.50E-238<br>1.10E-126<br>1.70E-90<br>4.70E-16<br>1.90E-11 |
| WP_005474565.1 | 662314 | 662938 | + | 207 | LinL | LinL | LinL | 1.50E-74 |
| WP_005474570.1 | 663169 | 665113 | + | 647 | TIGR02204<br>TIGR02203<br>TIGR00958<br>TIGR03796<br>TIGR03375 | TIGR02204<br>TIGR02203<br>TIGR00958<br>TIGR03796<br>TIGR03375 | MsbA_rel: ABC transporter, permease/ATP-binding protein<br>MsbA_lipidA: lipid A export permease/ATP-binding protein MsbA<br>3a01208: antigen peptide transporter 2<br>NHLM_micro_ABC1: NHLM bacteriocin system ABC transporter, peptidase/ATP-binding protein<br>type_I_sec_LssB: type I secretion system ATPase | 1.60E-102<br>4.90E-100<br>6.00E-93<br>1.10E-85<br>3.70E-78 |
| WP_005474572.1 | 665198 | 665390 | + | 63 | NO PFAM MATCH | - | - | - |
| WP_005474574.1 | 665386 | 666193 | + | 268 | PF13649<br>PF08241<br>PF13489<br>PF08242<br>TIGR01983 | Methyltransf_25<br>Methyltransf_11<br>Methyltransf_23<br>Methyltransf_12<br>TIGR01983 | Methyltransferase domain<br>Methyltransferase domain<br>Methyltransferase domain<br>Methyltransferase domain<br>UbiG: 3-demethylubiquinone-9 3-O-methyltransferase | 6.90E-14<br>8.20E-11<br>8.70E-07<br>1.00E-05<br>3.90E-05 |
| WP_005474576.1 | 666650 | 666254 | - | 131 | PF00583<br>PF13673<br>PF13508<br>TIGR01575<br>PF14542 | Acetyltransf_1<br>Acetyltransf_10<br>Acetyltransf_7<br>TIGR01575<br>Acetyltransf_CG | Acetyltransferase (GNAT) family<br>Acetyltransferase (GNAT) domain<br>Acetyltransferase (GNAT) domain<br>rimI: ribosomal-protein-alanine acetyltransferase<br>GCN5-related N-acetyl-transferase | 1.10E-12<br>6.20E-08<br>5.40E-06<br>1.60E-05<br>9.80E-05 |
| WP_017241969.1 | 666755 | 666926 | + | 57 | INFERRED GENE | - | - | - |
| WP_005474591.1 | 667122 | 668115 | + | 330 | TIGR02363<br>PF02733<br>TIGR02361<br>TIGR02362 | TIGR02363<br>Dak1<br>TIGR02361<br>TIGR02362 | dhaK1: dihydroxyacetone kinase, DhaK subunit<br>Dak1 domain<br>dak_ATP: dihydroxyacetone kinase<br>dhaK1b: probable dihydroxyacetone kinase DhaK1b subunit | 4.50E-148<br>1.40E-126<br>8.80E-94<br>3.60E-74 |
| WP_005474593.1 | 668201 | 668819 | + | 205 | TIGR02365<br>PF02734<br>TIGR02361<br>TIGR03599 | TIGR02365<br>Dak2<br>TIGR02361<br>TIGR03599 | dha_L_ycgS: dihydroxyacetone kinase, L subunit<br>DAK2 domain<br>dak_ATP: dihydroxyacetone kinase<br>YloV: DAK2 domain fusion protein YloV | 2.40E-68<br>9.30E-50<br>1.40E-35<br>3.10E-07 |

| index | leader | core | start | end | dir | score |
|---|---|---|---|---|---|---|
| 1 | VRFPFDFEGFGNLCFVPFV | GTTVRLHPEAPHRQKYLFPSVILFREQVERRKARTVRSGPSW | 659968 | 659782 | - | 6 |
| 2 | MDLPLEEIILTSARPTL | GTPRIGRGARAVQAAQVAGGLGEAA...QAAVEANQAVEAQANAANAQAAGGV | 659995 | 660247 | + | 18 |
| 3 | MSFSNPVFLLVCGFALLLPGL | ITAALMYAQARRQMKEEEGKTT | 662934 | 663066 | + | 6 |
| 4 | VFLLVCGFALLLPGL | ITAALMYAQARRQMKEEEGKTT | 662952 | 663066 | + | 6 |
| 5 | MTPVCTGAGAAVRPAGIR | PDRAWPPHRVSVHVKGTR | 663062 | 663173 | + | 6 |

| | | | | | | |
|---|---|---|---|---|---|---|
| 6 | MNDFLLIPSVVALGIVGFLIATRAVSTPAVI | GVALVVLWGACSQARAHTHARHTPSKKRRQHP | 665198 | 665390 | + | 7 |

### Results for WP_003965563.1 [Streptomyces sp. SID4928] back to top

previous - next

#### Architecture

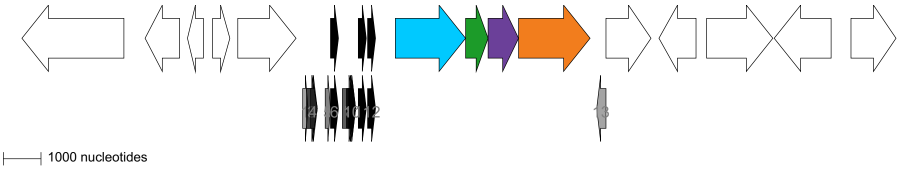

Link to nucleotide sequence

| Accession | start | end | direction | length (aa) | Pfam/HMM | name | description | E-value |
|---|---|---|---|---|---|---|---|---|
| WP_003965554.1 | 1971070 | 1968334 | - | 911 | TIGR01341<br>TIGR02333<br>PF00330<br>TIGR01340<br>TIGR01342 | TIGR01341<br>TIGR02333<br>Aconitase<br>TIGR01340<br>TIGR01342 | aconitase_1: aconitate hydratase 1<br>2met_isocit_dHY: 2-methylisocitrate dehydratase, Fe/S-dependent<br>Aconitase family (aconitate hydratase)<br>aconitase_mito: aconitate hydratase, mitochondrial<br>acon_putative: putative aconitate hydratase | 0.00E+00<br>9.40E-256<br>1.50E-181<br>3.90E-78<br>4.00E-75 |
| WP_003965555.1 | 1972539 | 1971621 | - | 305 | PF01636<br>TIGR02172<br>TIGR00938 | APH<br>TIGR02172<br>TIGR00938 | Phosphotransferase enzyme family<br>Fb_sc_TIGR02172: fibrobacter succinogenes paralogous family TIGR02172<br>thrB_alt: homoserine kinase | 2.60E-38<br>1.70E-06<br>6.30E-04 |
| WP_003965556.1 | 1973169 | 1972734 | - | 144 | PF18029 | Glyoxalase_6 | Glyoxalase-like domain | 1.00E-23 |
| WP_003965557.1 | 1973415 | 1973868 | + | 150 | PF01042<br>TIGR00004<br>TIGR03610 | Ribonuc_L-PSP<br>TIGR00004<br>TIGR03610 | Endoribonuclease L-PSP<br>TIGR00004: reactive intermediate/imine deaminase<br>RutC: pyrimidine utilization protein C | 9.20E-15<br>4.40E-10<br>1.30E-08 |
| WP_003965558.1 | 1974087 | 1975617 | + | 509 | TIGR01072<br>PF00275<br>TIGR01356<br>TIGR03070<br>PF01381 | TIGR01072<br>EPSP_synthase<br>TIGR01356<br>TIGR03070<br>HTH_3 | murA: UDP-N-acetylglucosamine 1-carboxyvinyltransferase<br>EPSP synthase (3-phosphoshikimate 1-carboxyvinyltransferase)<br>aroA: 3-phosphoshikimate 1-carboxyvinyltransferase<br>couple_hipB: transcriptional regulator, y4mF family<br>Helix-turn-helix | 4.00E-91<br>2.20E-39<br>1.10E-19<br>2.10E-14<br>1.70E-12 |
| WP_003965559.1 | 1976550 | 1976754 | + | 67 | LinA | LinA | LinA (linaridin precursor) | 1.10E-11 |
| WP_003965561.1 | 1977290 | 1977485 | + | 64 | LinA | LinA | LinA (linaridin precursor) | 1.20E-10 |
| WP_003965562.1 | 1977542 | 1977743 | + | 66 | LinA | LinA | LinA (linaridin precursor) | 2.10E-13 |
| WP_003965563.1 | 1978288 | 1980130 | + | 613 | LinH<br>LinG<br>LinE<br>LinL<br>PF12697 | LinH<br>LinG<br>LinE<br>LinL<br>Abhydrolase_6 | LinH (Fusion of LinE/LinG)<br>LinG Homolog<br>LinE Homolog<br>LinL<br>Alpha/beta hydrolase family | 3.90E-253<br>1.80E-111<br>3.10E-94<br>2.50E-35<br>1.80E-14 |
| WP_003965564.1 | 1980126 | 1980735 | + | 202 | LinL | LinL | LinL | 2.00E-65 |
| WP_003965565.1 | 1980731 | 1981544 | + | 270 | PF13649<br>PF08241<br>PF08242<br>PF13489<br>PF13847 | Methyltransf_25<br>Methyltransf_11<br>Methyltransf_12<br>Methyltransf_23<br>Methyltransf_31 | Methyltransferase domain<br>Methyltransferase domain<br>Methyltransferase domain<br>Methyltransferase domain<br>Methyltransferase domain | 2.30E-16<br>7.20E-12<br>5.40E-08<br>1.20E-07<br>1.60E-06 |
| WP_003965566.1 | 1981540 | 1983436 | + | 631 | TIGR02204<br>TIGR02203<br>TIGR00958<br>TIGR03796<br>TIGR03797 | TIGR02204<br>TIGR02203<br>TIGR00958<br>TIGR03796<br>TIGR03797 | MsbA_rel: ABC transporter, permease/ATP-binding protein<br>MsbA_lipidA: lipid A export permease/ATP-binding protein MsbA<br>3a01208: antigen peptide transporter 2<br>NHLM_micro_ABC1: NHLM bacteriocin system ABC transporter, peptidase/ATP-binding protein<br>NHLM_micro_ABC2: NHLM bacteriocin system ABC transporter, ATP-binding protein | 6.00E-102<br>3.50E-92<br>6.90E-85<br>1.40E-82<br>9.50E-77 |
| WP_106731625.1 | 1983860 | 1985069 | + | 402 | PF01548<br>PF02371 | DEDD_Tnp_IS110<br>Transposase_20 | Transposase<br>Transposase IS116/IS110/IS902 family | 1.20E-46<br>2.30E-23 |
| WP_003965568.1 | 1986285 | 1985271 | - | 337 | NO PFAM MATCH | - | - | - |
| WP_106731626.1 | 1986534 | 1988328 | + | 597 | NO PFAM MATCH | - | - | - |
| WP_106731553.1 | 1989845 | 1988360 | - | 494 | PF14559<br>PF13428<br>PF12895<br>PF13181<br>PF13432 | TPR_19<br>TPR_14<br>ANAPC3<br>TPR_8<br>TPR_16 | Tetratricopeptide repeat<br>Tetratricopeptide repeat<br>Anaphase-promoting complex, cyclosome, subunit 3<br>Tetratricopeptide repeat<br>Tetratricopeptide repeat | 1.30E-08<br>2.30E-08<br>6.40E-05<br>1.30E-04<br>7.20E-04 |
| WP_106731555.1 | 1990375 | 1991575 | + | 399 | NO PFAM MATCH | - | - | - |

| index | leader | core | start | end | dir | score |
|---|---|---|---|---|---|---|
| 1 | VTSAPREHGTGGRSGRLSMAPPIR | WTVGRVPGSQDVPNRGRRGPQWLRTSR | 1975807 | 1975963 | + | 6 |
| 2 | VDAVAHSGSELPGEATSALERRVGV...TCLPEAASYSVVLAGSPVGFEPPPV | IAPTALVRQGRSARSGRPEQVPVRQRAQQDAPAV | 1975922 | 1976210 | + | 7 |
| 3 | VAHSGSELPGEATSALERRVGVDDI...TCLPEAASYSVVLAGSPVGFEPPPV | IAPTALVRQGRSARSGRPEQVPVRQRAQQDAPAV | 1975931 | 1976210 | + | 7 |
| 4 | LPEAASYSVVLAGSPVGFEPPPV | IAPTALVRQGRSARSGRPEQVPVRQRAQQDAPAV | 1976036 | 1976210 | + | 7 |

| | | | | | | |
|---|---|---|---|---|---|---|
| 5 | MPAEWSSLCVGRSASCGAHHRSS | VHAGAGSRVRLRAPGWAPVTAHR | 1976400 | 1976541 | + | 7 |
| 6 | MPEFRQPGWTRGVAPLDESAGGQVFGGASPVA | ATPAVVATAGAVVVAFAAGVAARHLANGGNVELPM | 1976550 | 1976754 | + | 30 |
| 7 | VSPSGAARVGCRADSGSTRRRPSAD...VLSETGALSESDHGRALLDTVPVAS | VTFTMTACVEVSVCLTGSVIQLPQ | 1976862 | 1977219 | + | 8 |
| 8 | VGPSRRRRPPMASPHRPPTSEEKSL...VLSETGALSESDHGRALLDTVPVAS | VTFTMTACVEVSVCLTGSVIQLPQ | 1976973 | 1977219 | + | 8 |
| 9 | MASPHRPPTSEEKSLNTASIPAVLSETGALSESDHGRALLDTVPVAS | VTFTMTACVEVSVCLTGSVIQLPQ | 1977003 | 1977219 | + | 9 |
| 10 | LNTASIPAVLSETGALSESDHGRALLDTVPVAS | VTFTMTACVEVSVCLTGSVIQLPQ | 1977045 | 1977219 | + | 10 |
| 11 | MNKSSAPAVLTATGALTESDAGVTLPTLAPVA | ATPVAIAATMGVAFVAGYAAGRAATGNVELPM | 1977290 | 1977485 | + | 30 |
| 12 | MSMSPTPAALRGAGGLSESDPGRALSSLAPVT | ATPGVVAGVALGVALVNAFAAGYNHCGGNVELPM | 1977542 | 1977743 | + | 31 |
| 13 | VHPSVALLPGLVHGTDRHIHITKILPTCEQAGGHAPNQRSVSASGPGD | TTPRIMVNKGGEVMPGPEAGSPVAGPPKR | 1983866 | 1983632 | - | 6 |

### Results for WP_099217667.1 [Streptomyces sp. SID8350] back to top

previous - next

#### Architecture

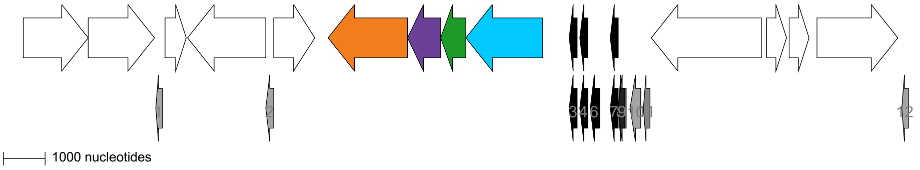

Link to nucleotide sequence

| Accession | start | end | direction | length (aa) | Pfam/HMM | name | description | E-value |
|---|---|---|---|---|---|---|---|---|
| WP_099217664.1 | 5562 | 7107 | + | 514 | PF00501<br>TIGR03208<br>TIGR03098<br>TIGR02275<br>TIGR01733 | AMP-binding<br>TIGR03208<br>TIGR03098<br>TIGR02275<br>TIGR01733 | AMP-binding enzyme<br>cyc_hxne_CoA_lg: cyclohexanecarboxylate-CoA ligase<br>ligase_PEP_1: acyl-CoA ligase (AMP-forming), exosortase A-associated<br>DHB_AMP_lig: (2,3-dihydroxybenzoyl)adenylate synthase<br>AA-adenyl-dom: amino acid adenylation domain | 8.30E-68<br>4.40E-44<br>5.50E-43<br>1.00E-38<br>5.70E-38 |
| WP_084990705.1 | 7114 | 8707 | + | 530 | PF00501<br>TIGR03205<br>TIGR03098<br>TIGR02262<br>TIGR03208 | AMP-binding<br>TIGR03205<br>TIGR03098<br>TIGR02262<br>TIGR03208 | AMP-binding enzyme<br>pimA: dicarboxylate--CoA ligase PimA<br>ligase_PEP_1: acyl-CoA ligase (AMP-forming), exosortase A-associated<br>benz_CoA_lig: benzoate-CoA ligase family<br>cyc_hxne_CoA_lg: cyclohexanecarboxylate-CoA ligase | 8.00E-114<br>8.20E-79<br>4.00E-75<br>9.90E-75<br>9.00E-72 |
| WP_099217665.1 | 8937 | 9474 | + | 178 | PF12840<br>PF01022<br>PF13518 | HTH_20<br>HTH_5<br>HTH_28 | Helix-turn-helix domain<br>Bacterial regulatory protein, arsR family<br>Helix-turn-helix domain | 3.50E-11<br>1.30E-06<br>9.50E-05 |
| WP_084990709.1 | 11353 | 9499 | - | 617 | NO PFAM MATCH | - | - | - |
| WP_028419792.1 | 11542 | 12556 | + | 337 | NO PFAM MATCH | - | - | - |
| WP_099217666.1 | 14755 | 12853 | - | 633 | TIGR02204<br>TIGR02203<br>TIGR00958<br>TIGR03796<br>TIGR03797 | TIGR02204<br>TIGR02203<br>TIGR00958<br>TIGR03796<br>TIGR03797 | MsbA_rel: ABC transporter, permease/ATP-binding protein<br>MsbA_lipidA: lipid A export permease/ATP-binding protein MsbA<br>3a01208: antigen peptide transporter 2<br>NHLM_micro_ABC1: NHLM bacteriocin system ABC transporter, peptidase/ATP-binding protein<br>NHLM_micro_ABC2: NHLM bacteriocin system ABC transporter, ATP-binding protein | 6.70E-99<br>6.70E-92<br>9.70E-84<br>7.20E-82<br>2.60E-77 |
| WP_084990713.1 | 15564 | 14751 | - | 270 | PF13649<br>PF08241<br>PF13847<br>PF13489<br>PF08242 | Methyltransf_25<br>Methyltransf_11<br>Methyltransf_31<br>Methyltransf_23<br>Methyltransf_12 | Methyltransferase domain<br>Methyltransferase domain<br>Methyltransferase domain<br>Methyltransferase domain<br>Methyltransferase domain | 4.70E-16<br>8.00E-12<br>6.80E-07<br>1.70E-06<br>1.80E-06 |
| WP_084990715.1 | 16169 | 15560 | - | 202 | LinL | LinL | LinL | 7.10E-66 |
| WP_099217667.1 | 18010 | 16165 | - | 614 | LinH<br>LinG<br>LinE<br>LinL<br>PF12697 | LinH<br>LinG<br>LinE<br>LinL<br>Abhydrolase_6 | LinH (Fusion of LinE/LinG)<br>LinG Homolog<br>LinE Homolog<br>LinL<br>Alpha/beta hydrolase family | 2.10E-251<br>1.00E-111<br>2.50E-95<br>3.90E-45<br>7.70E-14 |
| WP_084990720.1 | 18826 | 18625 | - | 66 | LinA | LinA | LinA (linaridin precursor) | 2.40E-12 |
| WP_084990722.1 | 19075 | 18883 | - | 63 | LinA | LinA | LinA (linaridin precursor) | 6.90E-09 |
| WP_084990724.1 | 19814 | 19610 | - | 67 | LinA | LinA | LinA (linaridin precursor) | 3.10E-11 |
| WP_099217668.1 | 23251 | 20596 | - | 884 | TIGR03903<br>PF13191<br>PF13424<br>PF07719<br>PF13401 | TIGR03903<br>AAA_16<br>TPR_12<br>TPR_2<br>AAA_22 | TOMM_kin_cyc: TOMM system kinase/cyclase fusion protein<br>AAA ATPase domain<br>Tetratricopeptide repeat<br>Tetratricopeptide repeat<br>AAA domain | 2.00E-20<br>7.10E-19<br>2.40E-05<br>9.30E-05<br>1.40E-04 |
| WP_099217669.1 | 23361 | 23841 | + | 159 | PF03745 | DUF309 | Domain of unknown function (DUF309) | 4.90E-19 |
| WP_099217670.1 | 23925 | 24390 | + | 154 | NO PFAM MATCH | - | - | - |
| WP_099217671.1 | 24587 | 26522 | + | 644 | PF10022 | DUF2264 | Uncharacterized protein conserved in bacteria (DUF2264) | 9.70E-97 |

| index | leader | core | start | end | dir | score |
|---|---|---|---|---|---|---|
| 1 | VVSADGPSSAATADAWAERGRKHRRS | YRSGPVLTVVQVRCGPDGRPGPVRPG | 8881 | 8722 | - | 6 |

| | | | | | | |
|---|---|---|---|---|---|---|
| 2 | VATSCRSPWPFPRTRPAGQRITSLPGEAGGEVS | AKSQASLAPGKFRMRDHLKERAPIAMTGGRAPK | 11567 | 11366 | - | 6 |
| 3 | MNMSPTPAALRGAGGLSESDPGRAHSTLVPVA | ATPGVLAGVAVGVALVNAFAAGYNHCGGNVELPM | 18826 | 18625 | - | 31 |
| 4 | MNTSSTPAVLTATGALTESDPGATLPAIAPVA | ATPVVVVAALGAGFVAGYAAGRAAGNVELPM | 19075 | 18883 | - | 30 |
| 5 | LSTASIPAVLSGTGALTESDRGRGPLD | TVPVTSVTVTMTACVGISVCLTGSTIELPQ | 19322 | 19148 | - | 10 |
| 6 | VTSTLPPTSEEKSLSTASIPAVLSGTGALTESDRGRGPLD | TVPVTSVTVTMTACVGISVCLTGSTIELPQ | 19361 | 19148 | - | 10 |
| 7 | MPEFRQPGWTRGLAPLDEGAGGQVFGGASPVA | ATPAVVATAGAVVVAFAAGVAARHLANGGNVELPM | 19814 | 19610 | - | 31 |
| 8 | MPAELSSLSVGLSASCGAHHRTS | VHAGAGSRVRSRAPGRAPVTAHP | 19964 | 19823 | - | 8 |
| 9 | VPEPAGCCCGVRILMPAELSSLSVGLSASC | GAHHRTSVHAGAGSRVRSRAPGRAPVTAHP | 20006 | 19823 | - | 8 |
| 10 | VCRRERLVDRAGRHPSDLLGPRRPWCRSRRSAQRLGQTR | CAFGGEADDGAGAGTGGGLAGPWPGGDLGECGVLAPLRR | 20349 | 20112 | - | 6 |
| 11 | LTCAAARTTGGNRSGTWRSLVAHLTGGQ | GVAGSNPVVPTGDSRRSGPVSEQSETGP | 20588 | 20417 | - | 7 |
| 12 | VPEAVARLVPER | PTTLISGPSATSDIELERVEGVHGPRRLVVVIRTDR | 26760 | 26613 | - | 6 |

### Results for WP_042495855.1 [Streptomyces sp. SID7816] back to top

previous - next

#### Architecture

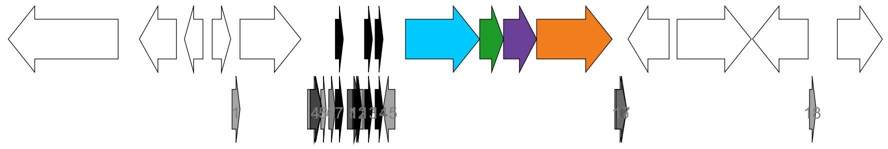

1000 nucleotides

Link to nucleotide sequence

| Accession | start | end | direction | length (aa) | Pfam/HMM | name | description | E-value |
|---|---|---|---|---|---|---|---|---|
| WP_003965554.1 | 1767747 | 1765011 | - | 911 | TIGR01341<br>TIGR02333<br>PF00330<br>TIGR01340<br>TIGR01342 | TIGR01341<br>TIGR02333<br>Aconitase<br>TIGR01340<br>TIGR01342 | aconitase_1: aconitate hydratase 1<br>2met_isocit_dHY: 2-methylisocitrate dehydratase, Fe/S-dependent<br>Aconitase family (aconitate hydratase)<br>aconitase_mito: aconitate hydratase, mitochondrial<br>acon_putative: putative aconitate hydratase | 0.00E+00<br>9.40E-256<br>1.50E-181<br>3.90E-78<br>4.00E-75 |
| WP_012378583.1 | 1769207 | 1768289 | - | 305 | PF01636<br>TIGR02172<br>TIGR00938 | APH<br>TIGR02172<br>TIGR00938 | Phosphotransferase enzyme family<br>Fb_sc_TIGR02172: fibrobacter succinogenes paralogous family TIGR02172<br>thrB_alt: homoserine kinase | 1.50E-38<br>1.70E-06<br>7.30E-04 |
| WP_012378584.1 | 1769838 | 1769403 | - | 144 | PF18029 | Glyoxalase_6 | Glyoxalase-like domain | 1.60E-23 |
| WP_042498244.1 | 1770084 | 1770537 | + | 150 | PF01042<br>TIGR00004<br>TIGR03610 | Ribonuc_L-PSP<br>TIGR00004<br>TIGR03610 | Endoribonuclease L-PSP<br>TIGR00004: reactive intermediate/imine deaminase<br>RutC: pyrimidine utilization protein C | 9.10E-15<br>4.40E-10<br>1.30E-08 |
| WP_003965558.1 | 1770766 | 1772296 | + | 509 | TIGR01072<br>PF00275<br>TIGR01356<br>TIGR03070<br>PF01381 | TIGR01072<br>EPSP_synthase<br>TIGR01356<br>TIGR03070<br>HTH_3 | murA: UDP-N-acetylglucosamine 1-carboxyvinyltransferase<br>EPSP synthase (3-phosphoshikimate 1-carboxyvinyltransferase)<br>aroA: 3-phosphoshikimate 1-carboxyvinyltransferase<br>couple_hipB: transcriptional regulator, y4mF family<br>Helix-turn-helix | 4.00E-91<br>2.20E-39<br>1.10E-19<br>2.10E-14<br>1.70E-12 |
| WP_003965559.1 | 1773133 | 1773337 | + | 67 | LinA | LinA | LinA (linaridin precursor) | 1.10E-11 |
| WP_003965561.1 | 1773872 | 1774067 | + | 64 | LinA | LinA | LinA (linaridin precursor) | 1.20E-10 |
| WP_003965562.1 | 1774124 | 1774325 | + | 66 | LinA | LinA | LinA (linaridin precursor) | 2.10E-13 |
| WP_042495855.1 | 1774867 | 1776709 | + | 613 | LinH<br>LinG<br>LinE<br>LinL<br>PF12697 | LinH<br>LinG<br>LinE<br>LinL<br>Abhydrolase_6 | LinH (Fusion of LinE/LinG)<br>LinG Homolog<br>LinE Homolog<br>LinL<br>Alpha/beta hydrolase family | 1.50E-253<br>5.50E-112<br>3.40E-94<br>3.70E-35<br>2.00E-14 |
| WP_003965564.1 | 1776705 | 1777314 | + | 202 | LinL | LinL | LinL | 2.00E-65 |
| WP_012378588.1 | 1777310 | 1778123 | + | 270 | PF13649<br>PF08241<br>PF08242<br>PF13489<br>PF13847 | Methyltransf_25<br>Methyltransf_11<br>Methyltransf_12<br>Methyltransf_23<br>Methyltransf_31 | Methyltransferase domain<br>Methyltransferase domain<br>Methyltransferase domain<br>Methyltransferase domain<br>Methyltransferase domain | 2.30E-16<br>7.20E-12<br>5.50E-08<br>1.20E-07<br>1.60E-06 |
| WP_012378589.1 | 1778119 | 1780015 | + | 631 | TIGR02204<br>TIGR02203<br>TIGR00958<br>TIGR03796<br>TIGR03797 | TIGR02204<br>TIGR02203<br>TIGR00958<br>TIGR03796<br>TIGR03797 | MsbA_rel: ABC transporter, permease/ATP-binding protein<br>MsbA_lipidA: lipid A export permease/ATP-binding protein MsbA<br>3a01208: antigen peptide transporter 2<br>NHLM_micro_ABC1: NHLM bacteriocin system ABC transporter, peptidase/ATP-binding protein<br>NHLM_micro_ABC2: NHLM bacteriocin system ABC transporter, ATP-binding protein | 9.80E-102<br>1.70E-92<br>6.30E-85<br>1.40E-82<br>3.40E-77 |
| WP_030703682.1 | 1781425 | 1780411 | - | 337 | NO PFAM MATCH | - | - | - |
| WP_042495859.1 | 1781614 | 1783468 | + | 617 | NO PFAM MATCH | - | - | - |
| WP_042495862.1 | 1784868 | 1783500 | - | 455 | PF13428<br>PF14559<br>PF13181<br>PF12895<br>PF13432 | TPR_14<br>TPR_19<br>TPR_8<br>ANAPC3<br>TPR_16 | Tetratricopeptide repeat<br>Tetratricopeptide repeat<br>Tetratricopeptide repeat<br>Anaphase-promoting complex, cyclosome, subunit 3<br>Tetratricopeptide repeat | 4.70E-09<br>7.40E-09<br>1.20E-05<br>1.80E-05<br>9.80E-05 |
| WP_042495864.1 | 1785575 | 1786697 | + | 373 | NO PFAM MATCH | - | - | - |

| index | leader | core | start | end | dir | score |
|---|---|---|---|---|---|---|
| 1 | VRGGAVGCGRSLGRTRGPRPSLRRGARWGEACTRSRYTPVT | RTAHPHSSLISYLR | 1770589 | 1770757 | + | 6 |
| 2 | VDSWVGCRGVRTCPTVDAVAHSGSE...TGLPEAASHSVVLAGSPAGFEPPPV | IAPTALVRQGRSARSGRPEQVPVRQRAQQDAPAV | 1772460 | 1772793 | + | 6 |
| 3 | VDAVAHSGSELPGEATSALERRVGV...TGLPEAASHSVVLAGSPAGFEPPPV | IAPTALVRQGRSARSGRPEQVPVRQRAQQDAPAV | 1772505 | 1772793 | + | 6 |
| 4 | VAHSGSELPGEATSALERRVGVDDI...TGLPEAASHSVVLAGSPAGFEPPPV | IAPTALVRQGRSARSGRPEQVPVRQRAQQDAPAV | 1772514 | 1772793 | + | 6 |
| 5 | VRSKTLPPFDVTSGLAEYPSAVRAAAEMASRSVGE | APPIALNTTRSRAG | 1772764 | 1772914 | + | 6 |
| 6 | MPAEWSSLCVGRSASCGAHHRSS | VHAGAGSRVRLRAPGWAPVTAHR | 1772983 | 1773124 | + | 7 |
| 7 | MPEFRQPGWTRGVAPLDESAGGQVFGGASPVA | ATPAVVATAGAVVVAFAAGVAARHLANGGNVELPM | 1773133 | 1773337 | + | 30 |
| 8 | VSPSGAARVGCRAESGSTRRRPSAD...VLSETGALSESDHGRALLDTVPVAS | VTFTMTACVEVSVCLTGSVIQLPQ | 1773445 | 1773802 | + | 8 |
| 9 | VGPSRRRRPPMASPHRPPTSEEKSL...VLSETGALSESDHGRALLDTVPVAS | VTFTMTACVEVSVCLTGSVIQLPQ | 1773556 | 1773802 | + | 8 |
| 10 | MASPHRPPTSEEKSLNTASIPAVLSETGALSESDHGRALLDTVPVAS | VTFTMTACVEVSVCLTGSVIQLPQ | 1773586 | 1773802 | + | 9 |
| 11 | LNTASIPAVLSETGALSESDHGRALLDTVPVAS | VTFTMTACVEVSVCLTGSVIQLPQ | 1773628 | 1773802 | + | 10 |
| 12 | MRGHPRGRPVGLLRELNDASGEADRDLDACRHGERDGGDR | NGVEQRPAVVGLGQGTGLGENGRYRSGVQGLLLAGGRPVG | 1773835 | 1773592 | - | 6 |
| 13 | MNKSSAPAVLTATGALTESDAGVTLPTLAPVA | ATPVAIAATMGVAFVAGYAAGRAATGNVELPM | 1773872 | 1774067 | + | 30 |
| 14 | MSMSPTPAALRGAGGLSESDPGRALSSLAPVT | ATPGVVAGVALGVALVNAFAAGYNHCGGNVELPM | 1774124 | 1774325 | + | 31 |
| 15 | VRDGLPPGRARAGRGRSCRGRAASP...VAWAGDRPGGARPHATRAPSLPAGP | GTVAAAPRAGAGATDLPPGARAAVAAHRPLGRHRDQRVPPPVTGTA | 1774627 | 1774330 | - | 6 |
| 16 | VGSAASTAVVGGGANGPARPAGGPC...ARPGDCRGTGAARPRARRSSRSPAD | RPPGRTETGGPEAPRTETGRPLPALGPVGAGTPVGAGTPAA | 1780071 | 1780353 | + | 6 |
| 17 | VGGGANGPARPAGGPCRPARPGDCRGTGAARPRARRSSRSPAD | RPPGRTETGGPEAPRTETGRPLPALGPVGAGTPVGAGTPAA | 1780098 | 1780353 | + | 6 |
| 18 | VTSLRGDLRTSIARQRRPKRDVLAM | STEVRHVDDRLAVYASGEGARLACGR | 1784892 | 1785048 | + | 6 |

### Results for WP_161229806.1 [Streptomyces sp. SID8352] back to top

previous - next

#### Architecture

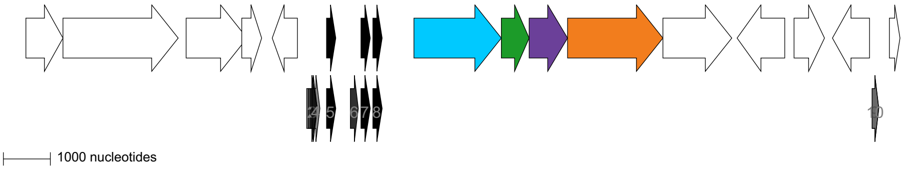

Link to nucleotide sequence

| Accession | start | end | direction | length (aa) | Pfam/HMM | name | description | E-value |
|---|---|---|---|---|---|---|---|---|
| WP_161229608.1 | 86308 | 87085 | + | 258 | TIGR03193<br>TIGR03198<br>TIGR03311<br>TIGR02963<br>PF01799 | TIGR03193<br>TIGR03198<br>TIGR03311<br>TIGR02963<br>Fer2_2 | 4hydroxCoAred: 4-hydroxybenzoyl-CoA reductase, gamma subunit<br>pucE: xanthine dehydrogenase E subunit<br>Se_dep_XDH: selenium-dependent xanthine dehydrogenase<br>xanthine_xdhA: xanthine dehydrogenase, small subunit<br>[2Fe-2S] binding domain | 8.20E-45<br>5.20E-42<br>9.00E-40<br>9.40E-34<br>1.10E-28 |
| WP_161229609.1 | 87081 | 89547 | + | 821 | TIGR03196<br>PF02738<br>TIGR03194<br>TIGR03311<br>TIGR02965 | TIGR03196<br>Ald_Xan_dh_C2<br>TIGR03194<br>TIGR03311<br>TIGR02965 | pucD: xanthine dehydrogenase D subunit<br>Molybdopterin-binding domain of aldehyde dehydrogenase<br>4hydrxCoA_A: 4-hydroxybenzoyl-CoA reductase, alpha subunit<br>Se_dep_XDH: selenium-dependent xanthine dehydrogenase<br>xanthine_xdhB: xanthine dehydrogenase, molybdopterin binding subunit | 7.80E-232<br>8.50E-153<br>1.20E-127<br>1.50E-125<br>3.80E-83 |
| WP_161229610.1 | 89714 | 91004 | + | 429 | PF13478<br>TIGR02964<br>PF02625 | XdhC_C<br>TIGR02964<br>XdhC_CoxI | XdhC Rossmann domain<br>xanthine_xdhC: xanthine dehydrogenase accessory protein XdhC<br>XdhC and CoxI family | 8.50E-41<br>4.30E-27<br>7.60E-27 |
| WP_161229611.1 | 90891 | 91314 | + | 140 | PF14525 | AraC_binding_2 | AraC-binding-like domain | 1.60E-09 |
| WP_161229612.1 | 92077 | 91558 | - | 172 | NO PFAM MATCH | - | - | - |
| WP_161229613.1 | 92702 | 92906 | + | 67 | LinA | LinA | LinA (linaridin precursor) | 1.20E-11 |
| WP_161229614.1 | 93433 | 93637 | + | 67 | LinA | LinA | LinA (linaridin precursor) | 1.00E-10 |
| WP_161229615.1 | 93694 | 93895 | + | 66 | LinA | LinA | LinA (linaridin precursor) | 4.70E-12 |
| WP_161229806.1 | 94574 | 96425 | + | 616 | LinH<br>LinG<br>LinE<br>LinL<br>PF12697 | LinH<br>LinG<br>LinE<br>LinL<br>Abhydrolase_6 | LinH (Fusion of LinE/LinG)<br>LinG Homolog<br>LinE Homolog<br>LinL<br>Alpha/beta hydrolase family | 1.20E-248<br>2.40E-111<br>1.30E-92<br>9.10E-35<br>1.20E-15 |
| WP_161229616.1 | 96421 | 97030 | + | 202 | LinL | LinL | LinL | 1.60E-64 |
| WP_161229617.1 | 97026 | 97830 | + | 267 | PF13649<br>PF08241<br>PF08242<br>PF13847<br>PF13489 | Methyltransf_25<br>Methyltransf_11<br>Methyltransf_12<br>Methyltransf_31<br>Methyltransf_23 | Methyltransferase domain<br>Methyltransferase domain<br>Methyltransferase domain<br>Methyltransferase domain<br>Methyltransferase domain | 1.20E-17<br>4.90E-13<br>3.90E-09<br>5.70E-09<br>1.90E-07 |

| | | | | | | | | |
|---|---|---|---|---|---|---|---|---|
| WP_161229618.1 | 97826 | 99863 | + | 678 | TIGR02204<br>TIGR02203<br>TIGR00958<br>TIGR03796<br>TIGR01846 | TIGR02204<br>TIGR02203<br>TIGR00958<br>TIGR03796<br>TIGR01846 | MsbA_rel: ABC transporter, permease/ATP-binding protein<br>MsbA_lipidA: lipid A export permease/ATP-binding protein MsbA<br>3a01208: antigen peptide transporter 2<br>NHLM_micro_ABC1: NHLM bacteriocin system ABC transporter, peptidase/ATP-binding protein<br>type_I_sec_HlyB: type I secretion system ATPase | 1.90E-98<br>2.30E-95<br>1.20E-85<br>3.10E-82<br>5.70E-77 |
| WP_161229807.1 | 99868 | 101329 | + | 486 | TIGR02734<br>TIGR03467<br>PF13450<br>PF01593<br>TIGR01317 | TIGR02734<br>TIGR03467<br>NAD_binding_8<br>Amino_oxidase<br>TIGR01317 | crtI_fam: phytoene desaturase<br>HpnE: squalene-associated FAD-dependent desaturase<br>NAD(P)-binding Rossmann-like domain<br>Flavin containing amine oxidoreductase<br>GOGAT_sm_gam: glutamate synthase, NADH/NADPH, small subunit | 2.00E-13<br>3.20E-09<br>3.70E-09<br>1.10E-08<br>1.20E-08 |
| WP_161229619.1 | 102471 | 101457 | - | 337 | NO PFAM MATCH | - | - | - |
| WP_018487277.1 | 102660 | 103321 | + | 220 | INFERRED GENE | - | - | - |
| WP_161229620.1 | 104264 | 103478 | - | 261 | TIGR02764<br>TIGR02884<br>PF01522<br>TIGR04243<br>TIGR02873 | TIGR02764<br>TIGR02884<br>Polysacc_deac_1<br>TIGR04243<br>TIGR02873 | spore_ybaN_pdaB: polysaccharide deacetylase family sporulation protein PdaB<br>spore_pdaA: delta-lactam-biosynthetic de-N-acetylase<br>Polysaccharide deacetylase<br>nodulat_NodB: chitooligosaccharide deacetylase NodB<br>spore_ylxY: probable sporulation protein, polysaccharide deacetylase family | 1.40E-42<br>5.60E-39<br>9.10E-38<br>2.70E-37<br>3.30E-26 |
| WP_161229621.1 | 104695 | 104926 | + | 76 | NO PFAM MATCH | - | - | - |

| index | leader | core | start | end | dir | score |
|---|---|---|---|---|---|---|
| 1 | VPPGRRNVPPGRRNTAGRPSPPLRRPPLAGEPAECR | HPRLRVPFTAAFRNVLPTRRVSGWVTGTAPLRIPPCP | 92294 | 92516 | + | 6 |
| 2 | VPPGRRNTAGRPSPPLRRPPLAGEPAECRHPRL | RVPFTAAFRNVLPTRRVSGWVTGTAPLRIPPCP | 92315 | 92516 | + | 8 |
| 3 | LAGEPAECRHPRLRVPFTAAFRN | VLPTRRVSGWVTGTAPLRIPPCP | 92375 | 92516 | + | 6 |
| 4 | MPTSPPPRPVHRRIPQCPSNAPRFRLGYRNRSAPDTPVP | VTAGCCLGVRILMPVA | 92395 | 92563 | + | 7 |
| 5 | MPELRQPGWTRGIAPLDESAGGQMFGDASPVA | ATPAVVTVAGAVVIAFAAGVAAREFACGSNTRLPM | 92702 | 92906 | + | 30 |
| 6 | LSTPSVPAALSETGALTESDPGRTPLD | SVPVASVSFTMTAGVSFAVCLTGSTTRLPQ | 93203 | 93377 | + | 9 |
| 7 | MSMSLAPAVLTETGVLTESDPGVALPTLTPVA | ATPVVAITALGVAGAGFVAGYMAGRAAGSDTRLPM | 93433 | 93637 | + | 29 |
| 8 | MNMSSTPAALRAAGSLSESDAGRALSAFAPVA | ATPAVVAGAAIGVALVNAFVAGYNYSGSNTKLPM | 93694 | 93895 | + | 30 |
| 9 | VKGMPGGRPVGGRRGSVRGAHPAS | VRTGPGTTVGGDADVARRSGTGCRV | 104323 | 104473 | + | 6 |
| 10 | MPGGRPVGGRRGSVRGAHPASVR | TGPGTTVGGDADVARRSGTGCRV | 104332 | 104473 | + | 6 |

### Results for WP_018487272.1 [Streptomyces sp. SID8356] back to top

previous - next

#### Architecture

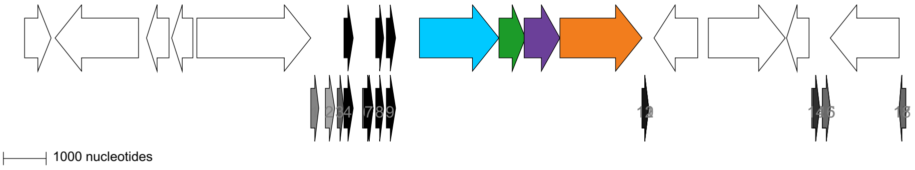

Link to nucleotide sequence

| Accession | start | end | direction | length (aa) | Pfam/HMM | name | description | E-value |
|---|---|---|---|---|---|---|---|---|
| WP_018487263.1 | 11534 | 12167 | + | 210 | PF02589 | LUD_dom | LUD domain | 2.10E-19 |
| WP_018487264.1 | 14194 | 12259 | - | 644 | PF10022 | DUF2264 | Uncharacterized protein conserved in bacteria (DUF2264) | 1.60E-96 |
| WP_018487265.1 | 14906 | 14399 | - | 168 | NO PFAM MATCH | - | - | - |
| WP_018487266.1 | 15470 | 14990 | - | 159 | PF03745 | DUF309 | Domain of unknown function (DUF309) | 7.30E-19 |
| WP_018487267.1 | 15580 | 18235 | + | 884 | TIGR03903<br>PF13191<br>PF13401<br>PF13424<br>PF07719 | TIGR03903<br>AAA_16<br>AAA_22<br>TPR_12<br>TPR_2 | TOMM_kin_cyc: TOMM system kinase/cyclase fusion protein<br>AAA ATPase domain<br>AAA domain<br>Tetratricopeptide repeat<br>Tetratricopeptide repeat | 2.10E-20<br>7.10E-19<br>1.40E-04<br>2.20E-04<br>3.00E-04 |
| WP_018487268.1 | 19019 | 19223 | + | 67 | LinA | LinA | LinA (linaridin precursor) | 3.40E-11 |
| WP_018487270.1 | 19757 | 19949 | + | 63 | LinA | LinA | LinA (linaridin precursor) | 2.60E-09 |
| WP_018487271.1 | 20007 | 20208 | + | 66 | LinA | LinA | LinA (linaridin precursor) | 7.60E-13 |
| WP_018487272.1 | 20788 | 22633 | + | 614 | LinH<br>LinG<br>LinE<br>LinL<br>PF12697 | LinH<br>LinG<br>LinE<br>LinL<br>Abhydrolase_6 | LinH (Fusion of LinE/LinG)<br>LinG Homolog<br>LinE Homolog<br>LinL<br>Alpha/beta hydrolase family | 2.50E-252<br>5.50E-112<br>1.80E-96<br>2.10E-43<br>1.40E-14 |
| WP_018487273.1 | 22629 | 23238 | + | 202 | LinL | LinL | LinL | 4.70E-66 |
| WP_018487274.1 | 23234 | 24047 | + | 270 | PF13649<br>PF08241<br>PF13847<br>PF13489<br>PF08242 | Methyltransf_25<br>Methyltransf_11<br>Methyltransf_31<br>Methyltransf_23<br>Methyltransf_12 | Methyltransferase domain<br>Methyltransferase domain<br>Methyltransferase domain<br>Methyltransferase domain<br>Methyltransferase domain | 2.30E-16<br>2.80E-12<br>2.50E-07<br>8.20E-07<br>8.60E-07 |

| | | | | | | | | |
|---|---|---|---|---|---|---|---|---|
| WP_018487275.1 | 24043 | 25969 | + | 641 | TIGR02204<br>TIGR02203<br>TIGR00958<br>TIGR03796<br>TIGR03797 | TIGR02204<br>TIGR02203<br>TIGR00958<br>TIGR03796<br>TIGR03797 | MsbA_rel: ABC transporter, permease/ATP-binding protein<br>MsbA_lipidA: lipid A export permease/ATP-binding protein MsbA<br>3a01208: antigen peptide transporter 2<br>NHLM_micro_ABC1: NHLM bacteriocin system ABC transporter, peptidase/ATP-binding protein<br>NHLM_micro_ABC2: NHLM bacteriocin system ABC transporter, ATP-binding protein | 5.90E-100<br>8.90E-93<br>4.20E-83<br>8.10E-81<br>8.50E-76 |
| WP_018487276.1 | 27279 | 26265 | - | 337 | NO PFAM MATCH | - | - | - |
| WP_106669312.1 | 27528 | 29322 | + | 597 | NO PFAM MATCH | - | - | - |
| WP_050565940.1 | 29884 | 29347 | - | 178 | PF12840<br>PF01022<br>PF13518 | HTH_20<br>HTH_5<br>HTH_28 | Helix-turn-helix domain<br>Bacterial regulatory protein, arsR family<br>Helix-turn-helix domain | 9.90E-12<br>9.90E-07<br>1.30E-04 |
| WP_018487279.1 | 31976 | 30383 | - | 530 | PF00501<br>TIGR03205<br>TIGR02262<br>TIGR03098<br>TIGR01923 | AMP-binding<br>TIGR03205<br>TIGR02262<br>TIGR03098<br>TIGR01923 | AMP-binding enzyme<br>pimA: dicarboxylate--CoA ligase PimA<br>benz_CoA_lig: benzoate-CoA ligase family<br>ligase_PEP_1: acyl-CoA ligase (AMP-forming), exosortase A-associated<br>menE: O-succinylbenzoate-CoA ligase | 6.80E-114<br>1.00E-79<br>1.40E-75<br>4.80E-75<br>1.30E-72 |

| index | leader | core | start | end | dir | score |
|---|---|---|---|---|---|---|
| 1 | LSCAAARTTGRNRSGTWRSLVAHLTGGQ | GVAGSNPVVPTGDSRRSGPVSEQSETGP | 18243 | 18414 | + | 7 |
| 2 | VRSAFGGPGARSAER | PTTAPVPARAAGSLVHGPAAISESA...AVARRDGRSPRSRSASLNASRSRPD | 18575 | 18800 | + | 6 |
| 3 | MPAELSSLSVGLSASCGAHHRTS | VHAGAGSRVRLRAPGRAPVTAHP | 18869 | 19010 | + | 8 |
| 4 | MPEFRQPGWTRGIAPLDEGAGGQVFGGASPVA | ATPAVVATAGAVVVAFAAGVAARHLANGGNVELPM | 19019 | 19223 | + | 31 |
| 5 | VGLFFRCHPPVT | STLPPTSEEKSLSTASIPAVLSGTG...SVTVTMTACVGISVCLTGSTIELPQ | 19442 | 19685 | + | 7 |
| 6 | VTSTLPPTSEEKSLSTASIPAVLSGTGALTESDRGRAPLD | SVPVTSVTVTMTACVGISVCLTGSTIELPQ | 19472 | 19685 | + | 10 |
| 7 | LSTASIPAVLSGTGALTESDRGRAPLD | SVPVTSVTVTMTACVGISVCLTGSTIELPQ | 19511 | 19685 | + | 10 |
| 8 | MNTSSTPAVLTATGALTESDPGAALPSLAPVT | ATPVVIVAALGAGFVAGYAAGRAAGNVELPM | 19757 | 19949 | + | 30 |
| 9 | MNMSPTPAALRGAGGLSESDPGRALSTLAPVA | ATPGVVAGVALGVALVNAFVAGYNHCGGNVELPM | 20007 | 20208 | + | 30 |
| 10 | VSTVDTTVVGSGPDGSAAA | VTPARAHHGCAPGPGPAPYRRSTARPGPRSWCDRPGA | 25965 | 26136 | + | 8 |
| 11 | VDTTVVGSGPDGSAAA | VTPARAHHGCAPGPGPAPYRRSTARPGPRSWCDRPGA | 25974 | 26136 | + | 8 |
| 12 | VGSGPDGSAAA | VTPARAHHGCAPGPGPAPYRRSTARPGPRSWCDRPGA | 25989 | 26136 | + | 6 |
| 13 | VASADGPSSSGSPGAAVDG | STPPPAASRGGVTTAESALGCGGFTAPRASGVPWSGPWCGTSSVAQP | 29940 | 30141 | + | 7 |
| 14 | VDGSTPPPAASRGGVTTAESALGCG | GFTAPRASGVPWSGPWCGTSSVAQP | 29988 | 30141 | + | 8 |
| 15 | VRSPVPDLVLAGDQPEVRGGADA | ATADLWADCGRKPDGRPSPVRRPSGHPVIRSHGCPVRPG | 30179 | 30368 | + | 6 |
| 16 | VLAGDQPEVRGGADA | ATADLWADCGRKPDGRPSPVRRPSGHPVIRSHGCPVRPG | 30203 | 30368 | + | 6 |
| 17 | VLREAATATPDGLAEHIRATISPKHVP | ARIHVRSALPLTGVGKPDKARLAAEAGG | 32151 | 31983 | - | 6 |
| 18 | VVLREAATATPDGLAEHIRATISPKHVP | ARIHVRSALPLTGVGKPDKARLAAEAGG | 32154 | 31983 | - | 6 |

### Results for WP_093683568.1 [Streptomyces sp. SID8364] back to top

previous - next

#### Architecture

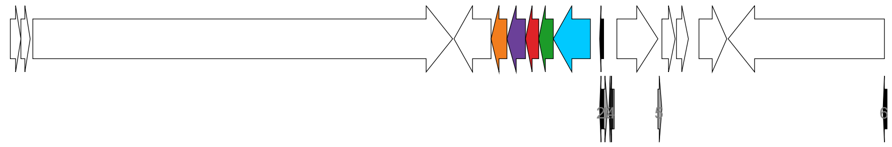

1000 nucleotides

Link to nucleotide sequence

| Accession | start | end | direction | length (aa) | Pfam/HMM | name | description | E-value |
|---|---|---|---|---|---|---|---|---|
| WP_003970673.1 | 234051 | 234507 | + | 151 | PF02575 | YbaB_DNA_bd | YbaB/EbfC DNA-binding family | 3.20E-09 |
| WP_030704389.1 | 234510 | 234879 | + | 122 | NO PFAM MATCH | - | - | - |
| WP_093683579.1 | 234993 | 251937 | + | 5647 | NO PFAM MATCH | - | - | - |
| WP_093683564.1 | 253518 | 251973 | - | 514 | NO PFAM MATCH | - | - | - |
| WP_093683565.1 | 254147 | 253514 | - | 210 | TIGR02857<br>TIGR01189<br>TIGR02673<br>TIGR02982<br>TIGR03258 | TIGR02857<br>TIGR01189<br>TIGR02673<br>TIGR02982<br>TIGR03258 | CydD: thiol reductant ABC exporter, CydD subunit<br>ccmA: heme ABC exporter, ATP-binding protein CcmA<br>FtsE: cell division ATP-binding protein FtsE<br>heterocyst_DevA: ABC exporter ATP-binding subunit, DevA family<br>PhnT: 2-aminoethylphosphonate ABC transport system, ATP-binding component PhnT | 1.30E-27<br>1.60E-25<br>3.70E-24<br>4.40E-22<br>1.40E-21 |
| WP_093683566.1 | 254881 | 254143 | - | 245 | PF13649<br>PF08241<br>TIGR02021<br>PF08242<br>TIGR01934 | Methyltransf_25<br>Methyltransf_11<br>TIGR02021<br>Methyltransf_12<br>TIGR01934 | Methyltransferase domain<br>Methyltransferase domain<br>BchM-ChlM: magnesium protoporphyrin O-methyltransferase<br>Methyltransferase domain<br>MenG_MenH_UbiE: ubiquinone/menaquinone biosynthesis methyltransferase | 1.30E-15<br>2.10E-14<br>1.10E-10<br>1.50E-10<br>8.50E-10 |

| | | | | | | | | |
|---|---|---|---|---|---|---|---|---|
| WP_093683567.1 | 255446 | 254873 | - | 190 | TIGR00521<br>TIGR02113<br>PF02441 | TIGR00521<br>TIGR02113<br>Flavoprotein | coaBC_dfp: phosphopantothenoylcysteine decarboxylase / phosphopantothenate--cysteine ligase<br>coaC_strep: phosphopantothenoylcysteine decarboxylase<br>Flavoprotein | 1.20E-14<br>7.60E-12<br>1.50E-11 |
| WP_003970680.1 | 256009 | 255442 | - | 188 | LinL | LinL | LinL | 1.10E-51 |
| WP_093683568.1 | 257519 | 255998 | - | 506 | LinH<br>LinG<br>LinE<br>PF12697<br>TIGR02427 | LinH<br>LinG<br>LinE<br>Abhydrolase_6<br>TIGR02427 | LinH (Fusion of LinE/LinG)<br>LinG Homolog<br>LinE Homolog<br>Alpha/beta hydrolase family<br>protocat_pcaD: 3-oxoadipate enol-lactonase | 6.70E-163<br>1.40E-67<br>4.40E-65<br>2.20E-06<br>2.60E-05 |
| WP_003970682.1 | 258067 | 257872 | - | 64 | LinA | LinA | LinA (linaridin precursor) | 6.80E-14 |
| WP_093683569.1 | 258588 | 260238 | + | 549 | PF03704<br>PF13191<br>TIGR03903<br>PF00486 | BTAD<br>AAA_16<br>TIGR03903<br>Trans_reg_C | Bacterial transcriptional activator domain<br>AAA ATPase domain<br>TOMM_kin_cyc: TOMM system kinase/cyclase fusion protein<br>Transcriptional regulatory protein, C terminal | 3.20E-43<br>4.80E-12<br>9.60E-08<br>2.70E-04 |
| WP_003970684.1 | 260415 | 260949 | + | 177 | PF01613<br>TIGR02296<br>TIGR03615 | Flavin_Reduct<br>TIGR02296<br>TIGR03615 | Flavin reductase like domain<br>HpaC: 4-hydroxyphenylacetate 3-monooxygenase, reductase component<br>RutF: pyrimidine utilization flavin reductase protein F | 8.90E-33<br>8.10E-24<br>1.80E-23 |
| WP_003970685.1 | 260973 | 261468 | + | 164 | NO PFAM MATCH | - | - | - |
| WP_093683570.1 | 261912 | 263016 | + | 367 | PF03704<br>PF13428<br>PF00486<br>PF13432<br>PF08279 | BTAD<br>TPR_14<br>Trans_reg_C<br>TPR_16<br>HTH_11 | Bacterial transcriptional activator domain<br>Tetratricopeptide repeat<br>Transcriptional regulatory protein, C terminal<br>Tetratricopeptide repeat<br>HTH domain | 7.20E-45<br>7.90E-06<br>4.40E-04<br>6.00E-04<br>6.90E-04 |
| WP_093683571.1 | 269394 | 263088 | - | 2101 | TIGR02813<br>PF00109<br>TIGR01746<br>PF08659<br>TIGR03443 | TIGR02813<br>ketoacyl-synt<br>TIGR01746<br>KR<br>TIGR03443 | omega_3_PfaA: polyketide-type polyunsaturated fatty acid synthase PfaA<br>Beta-ketoacyl synthase, N-terminal domain<br>Thioester-redct: thioester reductase domain<br>KR domain<br>alpha_am_amid: L-aminoadipate-semialdehyde dehydrogenase | 1.80E-118<br>3.10E-86<br>4.40E-85<br>5.50E-64<br>8.30E-58 |

| index | leader | core | start | end | dir | score |
|---|---|---|---|---|---|---|
| 1 | LQSNRVAFRCPSSEPMLSERDLAVP | SCPSPAASSELNALCTTSTTRSAGIPQSW | 258045 | 258210 | + | 6 |
| 2 | MRLDSIATQETATALPESMATQDFANSVLAGAVPGFHSDAETPAM | ATPAVAQFVIQGSTICLVC | 258067 | 257872 | - | 34 |
| 3 | LLGGLGTCEASGAGTPTDTGRTEGTTGRSSKP | ASSRNRPTGAWQLVQIQRDLRLVKLILMLRVGC | 258449 | 258251 | - | 9 |
| 4 | VQGLGPSPHKTRTRSGPCSGASAHARR | PAPGHRPTREGPRERPGVRRNRHRHEIARQARGNLSKFSGT | 258499 | 258292 | - | 6 |
| 5 | VRCRPRRRGGRSPAYGFRGRTRTIRR | RGAPRPVWVTGYPGMRVREIPRSMFSV | 260234 | 260396 | + | 6 |
| 6 | VDRGPGREHRRRGRELRVRVGRGHLRLHRQPARTLHQ | LTPVATHEGKRDHRVR | 269536 | 269374 | - | 10 |

### Results for WP_032756679.1 [Streptomyces sp. SID8359] back to top

previous - next

#### Architecture

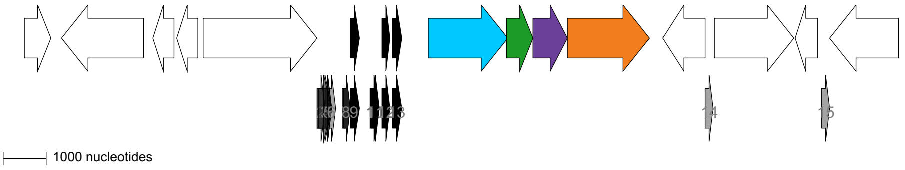

Link to nucleotide sequence

| Accession | start | end | direction | length (aa) | Pfam/HMM | name | description | E-value |
|---|---|---|---|---|---|---|---|---|
| WP_028419781.1 | 11281 | 11914 | + | 210 | PF02589 | LUD_dom | LUD domain | 1.70E-19 |
| WP_028419782.1 | 14073 | 12138 | - | 644 | PF10022 | DUF2264 | Uncharacterized protein conserved in bacteria (DUF2264) | 1.20E-96 |
| WP_028419783.1 | 14761 | 14278 | - | 160 | NO PFAM MATCH | - | - | - |
| WP_028419784.1 | 15325 | 14845 | - | 159 | PF03745 | DUF309 | Domain of unknown function (DUF309) | 5.00E-19 |
| WP_028419785.1 | 15435 | 18090 | + | 884 | TIGR03903<br>PF13191<br>PF13401<br>PF13424<br>PF07719 | TIGR03903<br>AAA_16<br>AAA_22<br>TPR_12<br>TPR_2 | TOMM_kin_cyc: TOMM system kinase/cyclase fusion protein<br>AAA ATPase domain<br>AAA domain<br>Tetratricopeptide repeat<br>Tetratricopeptide repeat | 1.20E-20<br>6.30E-19<br>1.70E-04<br>2.20E-04<br>3.20E-04 |
| WP_018487268.1 | 18873 | 19077 | + | 67 | LinA | LinA | LinA (linaridin precursor) | 3.40E-11 |
| WP_028419786.1 | 19612 | 19804 | + | 63 | LinA | LinA | LinA (linaridin precursor) | 1.30E-08 |
| WP_028419787.1 | 19861 | 20062 | + | 66 | LinA | LinA | LinA (linaridin precursor) | 1.40E-13 |
| WP_032756679.1 | 20676 | 22521 | + | 614 | LinH<br>LinG<br>LinE<br>LinL<br>PF12697 | LinH<br>LinG<br>LinE<br>LinL<br>Abhydrolase_6 | LinH (Fusion of LinE/LinG)<br>LinG Homolog<br>LinE Homolog<br>LinL<br>Alpha/beta hydrolase family | 5.80E-251<br>4.20E-112<br>1.00E-94<br>2.40E-45<br>9.60E-14 |
| WP_028419789.1 | 22517 | 23126 | + | 202 | LinL | LinL | LinL | 1.40E-65 |
| WP_032756681.1 | 23122 | 23935 | + | 270 | PF13649<br>PF08241<br>PF13847<br>PF13489<br>PF08242 | Methyltransf_25<br>Methyltransf_11<br>Methyltransf_31<br>Methyltransf_23<br>Methyltransf_12 | Methyltransferase domain<br>Methyltransferase domain<br>Methyltransferase domain<br>Methyltransferase domain<br>Methyltransferase domain | 2.30E-16<br>2.80E-12<br>2.60E-07<br>8.20E-07<br>8.70E-07 |

| | | | | | | | | |
|---|---|---|---|---|---|---|---|---|
| WP_028419791.1 | 23931 | 25833 | + | 633 | TIGR02204<br>TIGR02203<br>TIGR00958<br>TIGR03796<br>TIGR03797 | TIGR02204<br>TIGR02203<br>TIGR00958<br>TIGR03796<br>TIGR03797 | MsbA_rel: ABC transporter, permease/ATP-binding protein<br>MsbA_lipidA: lipid A export permease/ATP-binding protein MsbA<br>3a01208: antigen peptide transporter 2<br>NHLM_micro_ABC1: NHLM bacteriocin system ABC transporter, peptidase/ATP-binding protein<br>NHLM_micro_ABC2: NHLM bacteriocin system ABC transporter, ATP-binding protein | 2.70E-99<br>9.00E-92<br>2.80E-83<br>2.30E-81<br>1.70E-77 |
| WP_028419792.1 | 27142 | 26128 | - | 337 | NO PFAM MATCH | - | - | - |
| WP_028419793.1 | 27331 | 29185 | + | 617 | NO PFAM MATCH | - | - | - |
| WP_050451768.1 | 29747 | 29210 | - | 178 | PF12840<br>PF01022<br>PF13518 | HTH_20<br>HTH_5<br>HTH_28 | Helix-turn-helix domain<br>Bacterial regulatory protein, arsR family<br>Helix-turn-helix domain | 8.60E-11<br>1.90E-06<br>1.60E-04 |
| WP_028419795.1 | 31617 | 30024 | - | 530 | PF00501<br>TIGR03205<br>TIGR03098<br>TIGR02262<br>TIGR01923 | AMP-binding<br>TIGR03205<br>TIGR03098<br>TIGR02262<br>TIGR01923 | AMP-binding enzyme<br>pimA: dicarboxylate--CoA ligase PimA<br>ligase_PEP_1: acyl-CoA ligase (AMP-forming), exosortase A-associated<br>benz_CoA_lig: benzoate-CoA ligase family<br>menE: O-succinylbenzoate-CoA ligase | 1.30E-113<br>7.10E-79<br>5.50E-76<br>5.50E-75<br>2.20E-71 |

| index | leader | core | start | end | dir | score |
|---|---|---|---|---|---|---|
| 1 | MFVLRGHPDHREEPQRDVAQLGSAL...FKSCRPDWRQSQIRAGFGAIRDRPL | IIFGDQVGTGRDRSGPVGAPGVRAC...AGRHPSDLLGPPQAVVPVAPECAAP | 18094 | 18445 | + | 6 |
| 2 | LSCAATRTTGRNRSGTWRSLVAHLTGGQ | GVAGSNPVVPTGDSRRSGPVSEQSETGP | 18098 | 18269 | + | 7 |
| 3 | VAQLGSALDWGSRGRRFKSCRPDWRQSQIRAGFGAIRDRPLIIFGDQVG | TGRDRSGPVGAPGVRACRRERLVDRAGRHPSDLLGPPQAVVPVAPECAAP | 18145 | 18445 | + | 6 |
| 4 | LIIFGDQVGTGRDRSGPVGAPGVRACRRE | RLVDRAGRHPSDLLGPPQAVVPVAPECAAP | 18265 | 18445 | + | 7 |
| 5 | VGTGRDRSGPVGAPGVRACRRERLVD | RAGRHPSDLLGPPQAVVPVAPECAAP | 18286 | 18445 | + | 7 |
| 6 | LTGRGDIRRTSSGPRRPWCRSRRSAQR | LNQTRCAFGGEADDGAGAGTGGGLPDP | 18359 | 18524 | + | 6 |
| 7 | VPEPAGCCCGVRILMPAELSSLSVGLSASC | GAHHRTSVHAGAGSRVRSRAPGRAPVTAHP | 18681 | 18864 | + | 8 |
| 8 | MPAELSSLSVGLSASCGAHHRTS | VHAGAGSRVRSRAPGRAPVTAHP | 18723 | 18864 | + | 8 |
| 9 | MPEFRQPGWTRGIAPLDEGAGGQVFGGASPVA | ATPAVVATAGAVVVAFAAGVAARHLANGGNVELPM | 18873 | 19077 | + | 31 |
| 10 | VTSTLPPTSEEKSLSTASIPAVLSGTGALTESDRGRGPLD | TVPVTSVTVTMTACVGISVCLTGSTIELPQ | 19326 | 19539 | + | 10 |
| 11 | LSTASIPAVLSGTGALTESDRGRGPLD | TVPVTSVTVTMTACVGISVCLTGSTIELPQ | 19365 | 19539 | + | 10 |
| 12 | MNTSSTPAVLTATGALTESDPGATLPAIAPVT | ATPVVIVAALGAGFVAGYAAGRAAGNVELPM | 19612 | 19804 | + | 30 |
| 13 | MNMSPTPAALRGAGGLSESDPGRALPTLAPVA | ATPAVVAGVAVGVALVNAFVAGYNHCGGNVELPM | 19861 | 20062 | + | 31 |
| 14 | VATSCRSPWPFPRTRPAGQRITSLPGEAGGEVS | AKSQASLAPGKFRMRDHLKERAPIAMTGGRAPK | 27117 | 27318 | + | 6 |
| 15 | VGGTRQEAPTVVQVRSRRPSGSGPVP | TAVRVRSDPGRQSRSGPDCRPGPVRPG | 29847 | 30009 | + | 6 |

### Results for WP_093684647.1 [Streptomyces sp. SID8364] back to top

previous - next

#### Architecture

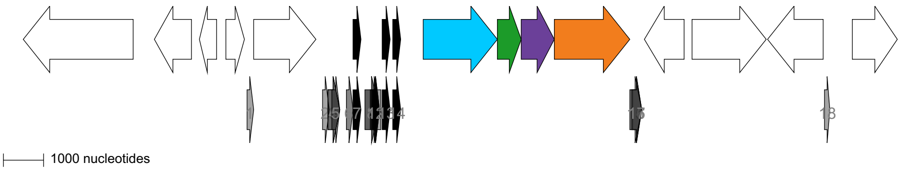

Link to nucleotide sequence

| Accession | start | end | direction | length (aa) | Pfam/HMM | name | description | E-value |
|---|---|---|---|---|---|---|---|---|
| WP_003965554.1 | 341821 | 339085 | - | 911 | TIGR01341<br>TIGR02333<br>PF00330<br>TIGR01340<br>TIGR01342 | TIGR01341<br>TIGR02333<br>Aconitase<br>TIGR01340<br>TIGR01342 | aconitase_1: aconitate hydratase 1<br>2met_isocit_dHY: 2-methylisocitrate dehydratase, Fe/S-dependent<br>Aconitase family (aconitate hydratase)<br>aconitase_mito: aconitate hydratase, mitochondrial<br>acon_putative: putative aconitate hydratase | 0.00E+00<br>9.40E-256<br>1.50E-181<br>3.90E-78<br>4.00E-75 |
| WP_093684645.1 | 343260 | 342342 | - | 305 | PF01636<br>TIGR02172<br>TIGR00938 | APH<br>TIGR02172<br>TIGR00938 | Phosphotransferase enzyme family<br>Fb_sc_TIGR02172: fibrobacter succinogenes paralogous family TIGR02172<br>thrB_alt: homoserine kinase | 9.20E-39<br>1.60E-06<br>4.80E-04 |
| WP_012378584.1 | 343899 | 343464 | - | 144 | PF18029 | Glyoxalase_6 | Glyoxalase-like domain | 1.60E-23 |
| WP_093562706.1 | 344145 | 344598 | + | 150 | PF01042<br>TIGR00004<br>TIGR03610 | Ribonuc_L-PSP<br>TIGR00004<br>TIGR03610 | Endoribonuclease L-PSP<br>TIGR00004: reactive intermediate/imine deaminase<br>RutC: pyrimidine utilization protein C | 9.20E-15<br>4.40E-10<br>1.30E-08 |
| WP_093684646.1 | 344827 | 346357 | + | 509 | TIGR01072<br>PF00275<br>TIGR01356<br>TIGR03070<br>PF01381 | TIGR01072<br>EPSP_synthase<br>TIGR01356<br>TIGR03070<br>HTH_3 | murA: UDP-N-acetylglucosamine 1-carboxyvinyltransferase<br>EPSP synthase (3-phosphoshikimate 1-carboxyvinyltransferase)<br>aroA: 3-phosphoshikimate 1-carboxyvinyltransferase<br>couple_hipB: transcriptional regulator, y4mF family<br>Helix-turn-helix | 3.60E-91<br>1.80E-39<br>8.90E-20<br>2.10E-14<br>1.70E-12 |
| WP_003965559.1 | 347290 | 347494 | + | 67 | LinA | LinA | LinA (linaridin precursor) | 1.10E-11 |
| WP_003965561.1 | 348030 | 348225 | + | 64 | LinA | LinA | LinA (linaridin precursor) | 1.20E-10 |
| WP_093562707.1 | 348288 | 348489 | + | 66 | LinA | LinA | LinA (linaridin precursor) | 1.50E-13 |

| | | | | | | | | |
|---|---|---|---|---|---|---|---|---|
| WP_093684647.1 | 349033 | 350875 | + | 613 | LinH<br>LinG<br>LinE<br>LinL<br>PF12697 | LinH<br>LinG<br>LinE<br>LinL<br>Abhydrolase_6 | LinH (Fusion of LinE/LinG)<br>LinG Homolog<br>LinE Homolog<br>LinL<br>Alpha/beta hydrolase family | 4.80E-254<br>3.10E-112<br>2.60E-94<br>2.40E-35<br>2.20E-14 |
| WP_030703686.1 | 350871 | 351480 | + | 202 | LinL | LinL | LinL | 1.40E-65 |
| WP_093684648.1 | 351476 | 352289 | + | 270 | PF13649<br>PF08241<br>PF08242<br>PF13489<br>PF13847 | Methyltransf_25<br>Methyltransf_11<br>Methyltransf_12<br>Methyltransf_23<br>Methyltransf_31 | Methyltransferase domain<br>Methyltransferase domain<br>Methyltransferase domain<br>Methyltransferase domain<br>Methyltransferase domain | 2.20E-16<br>7.00E-12<br>5.40E-08<br>1.20E-07<br>1.60E-06 |
| WP_003965566.1 | 352285 | 354181 | + | 631 | TIGR02204<br>TIGR02203<br>TIGR00958<br>TIGR03796<br>TIGR03797 | TIGR02204<br>TIGR02203<br>TIGR00958<br>TIGR03796<br>TIGR03797 | MsbA_rel: ABC transporter, permease/ATP-binding protein<br>MsbA_lipidA: lipid A export permease/ATP-binding protein MsbA<br>3a01208: antigen peptide transporter 2<br>NHLM_micro_ABC1: NHLM bacteriocin system ABC transporter, peptidase/ATP-binding protein<br>NHLM_micro_ABC2: NHLM bacteriocin system ABC transporter, ATP-binding protein | 6.00E-102<br>3.50E-92<br>6.90E-85<br>1.40E-82<br>9.50E-77 |
| WP_030703682.1 | 355534 | 354520 | - | 337 | NO PFAM MATCH | - | - | - |
| WP_042495859.1 | 355723 | 357577 | + | 617 | NO PFAM MATCH | - | - | - |
| WP_093684649.1 | 358977 | 357609 | - | 455 | PF14559<br>PF13428<br>PF13181<br>PF12895<br>PF13432 | TPR_19<br>TPR_14<br>TPR_8<br>ANAPC3<br>TPR_16 | Tetratricopeptide repeat<br>Tetratricopeptide repeat<br>Tetratricopeptide repeat<br>Anaphase-promoting complex, cyclosome, subunit 3<br>Tetratricopeptide repeat | 1.40E-09<br>1.90E-07<br>1.30E-05<br>2.20E-05<br>7.70E-05 |
| WP_093684650.1 | 359690 | 360812 | + | 373 | NO PFAM MATCH | - | - | - |

| index | leader | core | start | end | dir | score |
|---|---|---|---|---|---|---|
| 1 | VRGGAVGCGRSPGRTRGPRPSLRRGARWGEACTRSRYTPVT | RTAHPHSSLISYLR | 344650 | 344818 | + | 6 |
| 2 | VTSAPREHGTGGRSGRPSM | APPMRWTVGRVPGSQDVPNRGRRGPQWLRTSR | 346547 | 346703 | + | 6 |
| 3 | VDAVAHSGSELPGEATSALERRVGV...TCLPEAASYSVVLAGSPVGFEPPPV | IAPTALVRQGRSARSGRPEQVPVRQRAQQDTPAV | 346662 | 346950 | + | 6 |
| 4 | VAHSGSELPGEATSALERRVGVDDI...TCLPEAASYSVVLAGSPVGFEPPPV | IAPTALVRQGRSARSGRPEQVPVRQRAQQDTPAV | 346671 | 346950 | + | 6 |
| 5 | LPEAASYSVVLAGSPVGFEPPPV | IAPTALVRQGRSARSGRPEQVPVRQRAQQDTPAV | 346776 | 346950 | + | 6 |
| 6 | MPAEWSSLCVGRSASCGAHHRPS | VHAGAGSRVRLRAPGWAPVTAHR | 347140 | 347281 | + | 7 |
| 7 | MPEFRQPGWTRGVAPLDESAGGQVFGGASPVA | ATPAVVATAGAVVVAFAAGVAARHLANGGNVELPM | 347290 | 347494 | + | 30 |
| 8 | VSPSGAARAGCRADSGSTRRRPSAD...VLSETGALSESDHGRALLDTVPVAS | VTFTMTACVEVSVCLTGSVIQLPQ | 347602 | 347959 | + | 8 |
| 9 | VGPSRRRRPPMASPHRPPTSEEKSL...VLSETGALSESDHGRALLDTVPVAS | VTFTMTACVEVSVCLTGSVIQLPQ | 347713 | 347959 | + | 8 |
| 10 | MASPHRPPTSEEKSLNTASIPAVLSETGALSESDHGRALLDTVPVAS | VTFTMTACVEVSVCLTGSVIQLPQ | 347743 | 347959 | + | 9 |
| 11 | LNTASIPAVLSETGALSESDHGRALLDTVPVAS | VTFTMTACVEVSVCLTGSVIQLPQ | 347785 | 347959 | + | 10 |
| 12 | VRCGATRDGRPVGLLRELNDASGEADRDLDAGRHGERDGGD | RNGVEQRPAVVGLGQGTGLGENGRYRSGVQGLLLAGGRPVG | 347998 | 347749 | - | 6 |
| 13 | MNKSSAPAVLTATGALTESDAGVTLPTLAPVA | ATPVAIAATMGVAFVAGYAAGRAATGNVELPM | 348030 | 348225 | + | 30 |
| 14 | MSPTPTPAALRGAGGLSESDPGRALSSLAPVT | ATPGVAAGVALGVALVNAFAAGYNHCGGNVELPM | 348288 | 348489 | + | 31 |
| 15 | VRTAGAAGPPVVGSAASTAVVGGGA...ARPGDCRGTGAARPRARRSSCSPAD | RPPGRSETGRPLPALGPVGAGTPVGAGTPAA | 354177 | 354462 | + | 6 |
| 16 | VGSAASTAVVGGGANGPARPAGGPR...ARPGDCRGTGAARPRARRSSCSPAD | RPPGRSETGRPLPALGPVGAGTPVGAGTPAA | 354210 | 354462 | + | 6 |
| 17 | VGGGANGPARPAGGPRRPARPGDCRGTGAARPRARRSSCSPAD | RPPGRSETGRPLPALGPVGAGTPVGAGTPAA | 354237 | 354462 | + | 6 |
| 18 | VTSLRGGLRTSIARQRRPKRDVLAM | STEVRHVDDRLAVYASGEGARLACGR | 359001 | 359157 | + | 6 |

### Results for WP_161223844.1 [Streptomyces sp. SID8364] back to top

previous - next

#### Architecture

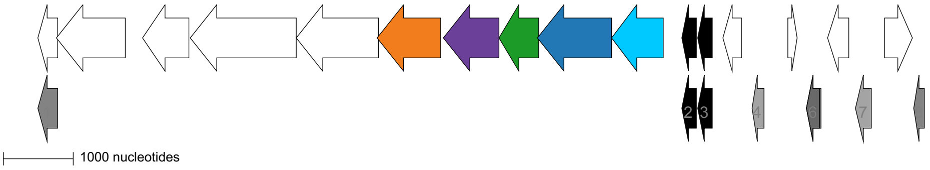

Link to nucleotide sequence

| Accession | start | end | direction | length (aa) | Pfam/HMM | name | description | E-value |
|---|---|---|---|---|---|---|---|---|
| WP_093683971.1 | 46514 | 46232 | - | 93 | PF00550<br>TIGR01688 | PP-binding<br>TIGR01688 | Phosphopantetheine attachment site<br>dltC: D-alanine--poly(phosphoribitol) ligase, subunit 2 | 7.70E-09<br>1.90E-05 |
| WP_161223848.1 | 47488 | 46510 | - | 325 | TIGR00747<br>PF08541 | TIGR00747<br>ACP_syn_III_C | fabH: 3-oxoacyl-[acyl-carrier-protein] synthase III<br>3-Oxoacyl-[acyl-carrier-protein (ACP)] synthase III C terminal | 2.70E-07<br>9.90E-05 |
| WP_093683973.1 | 48406 | 47743 | - | 220 | PF05719 | GPP34 | Golgi phosphoprotein 3 (GPP34) | 6.90E-49 |

| | | | | | | | | |
|---|---|---|---|---|---|---|---|---|
| WP_093683974.1 | 49937 | 48422 | - | 504 | TIGR02734<br>TIGR03467<br>PF13450<br>TIGR00292<br>TIGR03997 | TIGR02734<br>TIGR03467<br>NAD_binding_8<br>TIGR00292<br>TIGR03997 | crtI_fam: phytoene desaturase<br>HpnE: squalene-associated FAD-dependent desaturase<br>NAD(P)-binding Rossmann-like domain<br>TIGR00292: thiazole biosynthesis enzyme<br>mycofact_OYE_2: mycofactocin system FadH/OYE family oxidoreductase 2 | 1.50E-11<br>1.50E-08<br>4.10E-08<br>4.70E-08<br>6.20E-07 |
| WP_141697899.1 | 51120 | 49929 | - | 396 | PF12698 | ABC2_membrane_3 | ABC-2 family transporter protein | 5.80E-31 |
| WP_093683976.1 | 52012 | 51100 | - | 303 | TIGR03522<br>TIGR01188<br>TIGR03864<br>TIGR04406<br>TIGR03740 | TIGR03522<br>TIGR01188<br>TIGR03864<br>TIGR04406<br>TIGR03740 | GldA_ABC_ATP: gliding motility-associated ABC transporter ATP-binding subunit GldA<br>drrA: daunorubicin resistance ABC transporter, ATP-binding protein<br>PQQ_ABC_ATP: ABC transporter, ATP-binding subunit, PQQ-dependent alcohol dehydrogenase system<br>LPS_export_lptB: LPS export ABC transporter ATP-binding protein<br>galliderm_ABC: lantibiotic protection ABC transporter, ATP-binding subunit | 7.90E-53<br>1.00E-45<br>3.90E-44<br>4.40E-41<br>1.00E-40 |
| WP_093563034.1 | 52836 | 52047 | - | 262 | PF13649<br>PF08241<br>PF13847<br>PF13489<br>PF08242 | Methyltransf_25<br>Methyltransf_11<br>Methyltransf_31<br>Methyltransf_23<br>Methyltransf_12 | Methyltransferase domain<br>Methyltransferase domain<br>Methyltransferase domain<br>Methyltransferase domain<br>Methyltransferase domain | 5.10E-21<br>2.70E-17<br>1.00E-14<br>2.20E-12<br>6.20E-12 |
| WP_093683977.1 | 53399 | 52832 | - | 188 | LinL | LinL | LinL | 9.70E-71 |
| WP_161223844.1 | 54460 | 53398 | - | 353 | LinE<br>LinH<br>PF12697<br>PF00561<br>TIGR03695 | LinE<br>LinH<br>Abhydrolase_6<br>Abhydrolase_1<br>TIGR03695 | LinE Homolog<br>LinH (Fusion of LinE/LinG)<br>Alpha/beta hydrolase family<br>alpha/beta hydrolase fold<br>menH_SHCHC: 2-succinyl-6-hydroxy-2,4-cyclohexadiene-1-carboxylate synthase | 9.00E-127<br>2.20E-113<br>8.60E-16<br>6.10E-09<br>4.40E-08 |
| WP_161223845.1 | 55200 | 54456 | - | 247 | LinH<br>LinG | LinH<br>LinG | LinH (Fusion of LinE/LinG)<br>LinG Homolog | 6.00E-106<br>3.50E-101 |
| WP_093563031.1 | 55661 | 55457 | - | 67 | LinA | LinA | LinA (linaridin precursor) | 5.60E-12 |
| WP_093563030.1 | 55891 | 55693 | - | 65 | LinA | LinA | LinA (linaridin precursor) | 2.10E-08 |
| WP_093563029.1 | 56318 | 56039 | - | 92 | PF00196<br>TIGR03541<br>TIGR03020<br>TIGR01884<br>PF08279 | GerE<br>TIGR03541<br>TIGR03020<br>TIGR01884<br>HTH_11 | Bacterial regulatory proteins, luxR family<br>reg_near_HchA: LuxR family transcriptional regulatory, chaperone HchA-associated<br>EpsA: transcriptional regulator EpsA<br>cas_HTH: CRISPR locus-related DNA-binding protein<br>HTH domain | 3.60E-15<br>7.60E-10<br>2.40E-08<br>2.90E-04<br>5.10E-04 |
| WP_014043973.1 | 56976 | 57108 | + | 44 | INFERRED GENE | - | - | - |
| WP_093683979.1 | 57840 | 57540 | - | 99 | NO PFAM MATCH | - | - | - |
| WP_018523445.1 | 58361 | 58757 | + | 132 | INFERRED GENE | - | - | - |

| index | leader | core | start | end | dir | score |
|---|---|---|---|---|---|---|
| 1 | MSRGPAETLDEITRFLTEALRREVGPDDDYFALGLVDSLFALELVT | YVEDRFELTVEVEDLDLDSFRTAGRLTAFVHAKKAGAETTPAHAGDN | 46514 | 46232 | - | 7 |
| 2 | MSVVLDKVNASVEFGAIPGHISEAEWGNATDVPVL | ATPVTLTPTIGKVTLGVGAFTGGIAVGDALFG | 55661 | 55457 | - | 30 |
| 3 | MSTLLQKVNTSQELHAVASDLADAKDLSLTLPVL | ATITTVCPATGYIVGGGLFATGIGITYTVNR | 55891 | 55693 | - | 29 |
| 4 | LSGRNAQSHPPLDEPDIRSSGLPC | RPPDSPRDHLEALGDNQTQSQCERIYCASL | 56631 | 56466 | - | 6 |
| 5 | MGEVIGVVGRRAHCLTCERPHEDAAMPEGVMCSNCART | AVPGPCRRCSARCPTARPFGTWTR | 57426 | 57237 | - | 6 |
| 6 | MSAYTPSMGEVIGVVGRRAHCLTCERPHEDAAMPEGVMCSNCART | AVPGPCRRCSARCPTARPFGTWTR | 57447 | 57237 | - | 6 |
| 7 | VCERDAAAGEAVAPSG | PAPRAPVATEVGIAPRTQHVHLTVL...APRSIPSFRPHRRRPPPAGGQPPSP | 58169 | 57944 | - | 6 |
| 8 | MEELRQALDGPGVPSWPVVNHVTTALL | GLPEITRPLWRSAQNSTVSTIRAEAFG | 58939 | 58774 | - | 7 |

### Results for TXS69815.1 [Streptomyces sp. sk2.1] back to top

previous - next

#### Architecture

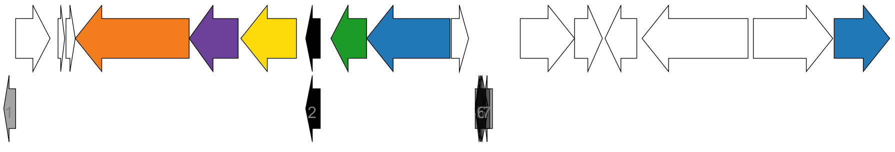

1000 nucleotides

Link to nucleotide sequence

| Accession | start | end | direction | length (aa) | Pfam/HMM | name | description | E-value |
|---|---|---|---|---|---|---|---|---|
| WP_019075229.1 | 11025 | 11544 | + | 173 | INFERRED GENE | - | - | - |
| WP_006374571.1 | 11676 | 11778 | + | 34 | INFERRED GENE | - | - | - |
| EAO76_24650 | 11796 | 11946 | + | 50 | INFERRED GENE | - | - | - |

| | | | | | | | | |
|---|---|---|---|---|---|---|---|---|
| TXS69811.1 | 13682 | 11942 | - | 579 | TIGR02204<br>TIGR02203<br>TIGR00958<br>TIGR03796<br>TIGR01846 | TIGR02204<br>TIGR02203<br>TIGR00958<br>TIGR03796<br>TIGR01846 | MsbA_rel: ABC transporter, permease/ATP-binding protein<br>MsbA_lipidA: lipid A export permease/ATP-binding protein MsbA<br>3a01208: antigen peptide transporter 2<br>NHLM_micro_ABC1: NHLM bacteriocin system ABC transporter, peptidase/ATP-binding protein<br>type_I_sec_HlyB: type I secretion system ATPase | 5.50E-99<br>9.20E-96<br>8.90E-84<br>1.30E-83<br>3.40E-79 |
| TXS69812.1 | 14434 | 13678 | - | 251 | PF13649<br>PF08241<br>PF13489<br>PF08242<br>PF13847 | Methyltransf_25<br>Methyltransf_11<br>Methyltransf_23<br>Methyltransf_12<br>Methyltransf_31 | Methyltransferase domain<br>Methyltransferase domain<br>Methyltransferase domain<br>Methyltransferase domain<br>Methyltransferase domain | 6.40E-15<br>3.50E-09<br>6.30E-09<br>5.30E-08<br>2.20E-07 |
| TXS69833.1 | 15341 | 14486 | - | 284 | LinG<br>LinH | LinG<br>LinH | LinG Homolog<br>LinH (Fusion of LinE/LinG) | 9.80E-103<br>4.80E-75 |
| TXS69813.1 | 15691 | 15484 | - | 68 | LinA | LinA | LinA (linaridin precursor) | 6.70E-14 |
| TXS69814.1 | 16418 | 15854 | - | 187 | LinL | LinL | LinL | 1.40E-55 |
| TXS69815.1 | 17689 | 16414 | - | 424 | LinE<br>LinH<br>PF12697<br>PF00561<br>TIGR02427 | LinE<br>LinH<br>Abhydrolase_6<br>Abhydrolase_1<br>TIGR02427 | LinE Homolog<br>LinH (Fusion of LinE/LinG)<br>Alpha/beta hydrolase family<br>alpha/beta hydrolase fold<br>protocat_pcaD: 3-oxoadipate enol-lactonase | 8.00E-93<br>1.30E-44<br>2.40E-16<br>2.30E-11<br>2.40E-08 |
| WP_019524132.1 | 17703 | 17976 | + | 91 | INFERRED GENE | - | - | - |
| TXS69816.1 | 18773 | 19604 | + | 276 | PF13340<br>PF01609<br>PF13586 | DUF4096<br>DDE_Tnp_1<br>DDE_Tnp_1_2 | Putative transposase of IS4/5 family (DUF4096)<br>Transposase DDE domain<br>Transposase DDE domain | 3.60E-23<br>3.90E-07<br>8.20E-07 |
| TXS69817.1 | 19600 | 20023 | + | 140 | PF12126 | DUF3583 | Protein of unknown function (DUF3583) | 4.60E-04 |
| TXS69818.1 | 20564 | 20063 | - | 166 | NO PFAM MATCH | - | - | - |
| TXS69819.1 | 22261 | 20635 | - | 541 | PF13738<br>PF00743<br>PF13450<br>TIGR04046<br>PF07992 | Pyr_redox_3<br>FMO-like<br>NAD_binding_8<br>TIGR04046<br>Pyr_redox_2 | Pyridine nucleotide-disulphide oxidoreductase<br>Flavin-binding monooxygenase-like<br>NAD(P)-binding Rossmann-like domain<br>MSMEG_0569_nitr: flavin-dependent oxidoreductase, MSMEG_0569 family<br>Pyridine nucleotide-disulphide oxidoreductase | 1.40E-14<br>2.30E-14<br>3.30E-13<br>8.80E-13<br>2.40E-11 |
| TXS69820.1 | 22346 | 23576 | + | 409 | PF03435<br>PF16653 | Sacchrp_dh_NADP<br>Sacchrp_dh_C | Saccharopine dehydrogenase NADP binding domain<br>Saccharopine dehydrogenase C-terminal domain | 4.60E-17<br>5.00E-04 |
| TXS69821.1 | 23596 | 24445 | + | 282 | PF00561<br>TIGR02427<br>TIGR03343<br>PF12697<br>TIGR01738 | Abhydrolase_1<br>TIGR02427<br>TIGR03343<br>Abhydrolase_6<br>TIGR01738 | alpha/beta hydrolase fold<br>protocat_pcaD: 3-oxoadipate enol-lactonase<br>biphenyl_bphD: 2-hydroxy-6-oxo-6-phenylhexa-2,4-dienoate hydrolase<br>Alpha/beta hydrolase family<br>bioH: pimelyl-[acyl-carrier protein] methyl ester esterase | 1.20E-30<br>5.00E-28<br>5.30E-24<br>5.10E-21<br>5.70E-21 |

| index | leader | core | start | end | dir | score |
|---|---|---|---|---|---|---|
| 1 | LTSIGGSTPSPSRSSRASSASCSTDAAGPRRH | RPGSSYSAGSRTTTGAVAIPNSATSHQPSSNNN | 11029 | 10831 | - | 6 |
| 2 | MSLVAQFANQSLTDIAPGMLSETDAARLDTPADPVM | ATPTTVLAVEVFAVATGMLAANIANYVDRNNY | 15691 | 15484 | - | 33 |
| 3 | LGHRSVNDSPSCRAVVGSAGEPILDARAHEIFE | TAVDEVCVHEELSYTAVVTAVGVLHFVVAAGVG | 18071 | 18272 | + | 7 |
| 4 | VDADLIYSGLKYLMSSCI | QNGFTGTPYNCATRWGVVD | 18197 | 18083 | - | 7 |
| 5 | VQDANGSNNGCVRELFVDADLIYSGL | KYLMSSCIQNGFTGTPYNCATRWGVVD | 18245 | 18083 | - | 8 |
| 6 | MGPGDSRRAVTTISTNAGGYYEVQDANGSNNGCVREL | FVDADLIYSGLKYLMSSCIQNGFTGTPYNCATRWGVVD | 18311 | 18083 | - | 7 |
| 7 | LAVPPPSSKHATWDRATAVARS | PPSQPTPAATTKCRTPTAVTTAV | 18346 | 18208 | - | 6 |

### Results for WP_148838111.1 [Streptomyces sp. sk2.1] back to top

previous - next

#### Architecture

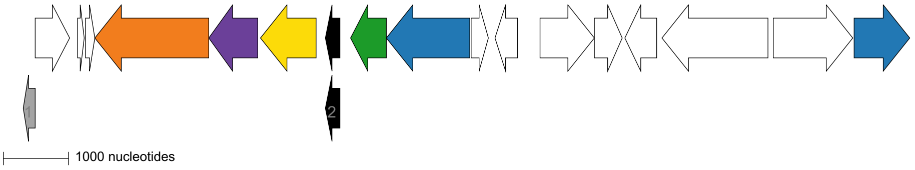

Link to nucleotide sequence

| Accession | start | end | direction | length (aa) | Pfam/HMM | name | description | E-value |
|---|---|---|---|---|---|---|---|---|
| WP_019075229.1 | 11025 | 11544 | + | 173 | INFERRED GENE | - | - | - |
| WP_006374571.1 | 11676 | 11778 | + | 34 | INFERRED GENE | - | - | - |
| EAO76_RS24570 | 11796 | 11946 | + | 50 | INFERRED GENE | - | - | - |
| WP_148838107.1 | 13682 | 11942 | - | 579 | TIGR02204<br>TIGR02203<br>TIGR00958<br>TIGR03796<br>TIGR01846 | TIGR02204<br>TIGR02203<br>TIGR00958<br>TIGR03796<br>TIGR01846 | MsbA_rel: ABC transporter, permease/ATP-binding protein<br>MsbA_lipidA: lipid A export permease/ATP-binding protein MsbA<br>3a01208: antigen peptide transporter 2<br>NHLM_micro_ABC1: NHLM bacteriocin system ABC transporter, peptidase/ATP-binding protein<br>type_I_sec_HlyB: type I secretion system ATPase | 5.50E-99<br>9.20E-96<br>8.90E-84<br>1.30E-83<br>3.40E-79 |
| WP_148838108.1 | 14434 | 13678 | - | 251 | PF13649<br>PF08241<br>PF13489<br>PF08242<br>PF13847 | Methyltransf_25<br>Methyltransf_11<br>Methyltransf_23<br>Methyltransf_12<br>Methyltransf_31 | Methyltransferase domain<br>Methyltransferase domain<br>Methyltransferase domain<br>Methyltransferase domain<br>Methyltransferase domain | 6.40E-15<br>3.50E-09<br>6.30E-09<br>5.30E-08<br>2.20E-07 |

| | | | | | | | | |
|---|---|---|---|---|---|---|---|---|
| WP_148838131.1 | 15341 | 14486 | - | 284 | LinG<br>LinH | LinG<br>LinH | LinG Homolog<br>LinH (Fusion of LinE/LinG) | 9.80E-103<br>4.80E-75 |
| WP_148838109.1 | 15691 | 15484 | - | 68 | LinA | LinA | LinA (linaridin precursor) | 6.70E-14 |
| WP_148838110.1 | 16418 | 15854 | - | 187 | LinL | LinL | LinL | 1.40E-55 |
| WP_148838111.1 | 17683 | 16414 | - | 422 | LinE<br>LinH<br>PF12697<br>PF00561<br>TIGR02427 | LinE<br>LinH<br>Abhydrolase_6<br>Abhydrolase_1<br>TIGR02427 | LinE Homolog<br>LinH (Fusion of LinE/LinG)<br>Alpha/beta hydrolase family<br>alpha/beta hydrolase fold<br>protocat_pcaD: 3-oxoadipate enol-lactonase | 7.90E-93<br>1.30E-44<br>2.40E-16<br>2.30E-11<br>2.40E-08 |
| WP_019524132.1 | 17703 | 17976 | + | 91 | INFERRED GENE | - | - | - |
| WP_148838112.1 | 18431 | 18083 | - | 115 | NO PFAM MATCH | - | - | - |
| WP_148838113.1 | 18773 | 19604 | + | 276 | PF13340<br>PF01609<br>PF13586 | DUF4096<br>DDE_Tnp_1<br>DDE_Tnp_1_2 | Putative transposase of IS4/5 family (DUF4096)<br>Transposase DDE domain<br>Transposase DDE domain | 3.60E-23<br>3.90E-07<br>8.20E-07 |
| WP_148838114.1 | 19600 | 20023 | + | 140 | PF12126 | DUF3583 | Protein of unknown function (DUF3583) | 4.60E-04 |
| WP_148838115.1 | 20564 | 20063 | - | 166 | NO PFAM MATCH | - | - | - |
| WP_148838116.1 | 22261 | 20635 | - | 541 | PF13738<br>PF00743<br>PF13450<br>TIGR04046<br>PF07992 | Pyr_redox_3<br>FMO-like<br>NAD_binding_8<br>TIGR04046<br>Pyr_redox_2 | Pyridine nucleotide-disulphide oxidoreductase<br>Flavin-binding monooxygenase-like<br>NAD(P)-binding Rossmann-like domain<br>MSMEG_0569_nitr: flavin-dependent oxidoreductase, MSMEG_0569 family<br>Pyridine nucleotide-disulphide oxidoreductase | 1.40E-14<br>2.30E-14<br>3.30E-13<br>8.80E-13<br>2.40E-11 |
| WP_148838117.1 | 22346 | 23576 | + | 409 | PF03435<br>PF16653 | Sacchrp_dh_NADP<br>Sacchrp_dh_C | Saccharopine dehydrogenase NADP binding domain<br>Saccharopine dehydrogenase C-terminal domain | 4.60E-17<br>5.00E-04 |
| WP_148838118.1 | 23596 | 24445 | + | 282 | PF00561<br>TIGR02427<br>TIGR03343<br>PF12697<br>TIGR01738 | Abhydrolase_1<br>TIGR02427<br>TIGR03343<br>Abhydrolase_6<br>TIGR01738 | alpha/beta hydrolase fold<br>protocat_pcaD: 3-oxoadipate enol-lactonase<br>biphenyl_bphD: 2-hydroxy-6-oxo-6-phenylhexa-2,4-dienoate hydrolase<br>Alpha/beta hydrolase family<br>bioH: pimelyl-[acyl-carrier protein] methyl ester esterase | 1.20E-30<br>5.00E-28<br>5.30E-24<br>5.10E-21<br>5.70E-21 |

| index | leader | core | start | end | dir | score |
|---|---|---|---|---|---|---|
| 1 | LTSIGGSTPSPSRSSRASSASCSTDAAGPRRH | RPGSSYSAGSRTTTGAVAIPNSATSHQPSSNNN | 11029 | 10831 | - | 6 |
| 2 | MSLVAQFANQSLTDIAPGMLSETDAARLDTPADPVM | ATPTTVLAVEVFAVATGMLAANIANYVDRNNY | 15691 | 15484 | - | 33 |

### Results for WP_103502974.1 [Streptomyces sp. SM14] back to top

previous - next

#### Architecture

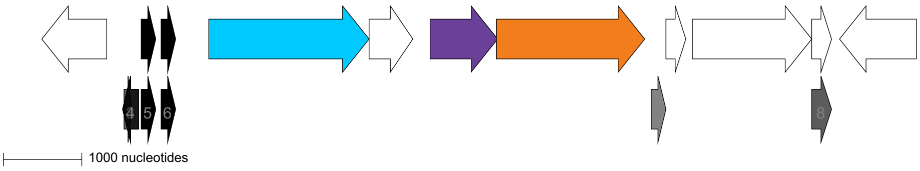

Link to nucleotide sequence

| Accession | start | end | direction | length (aa) | Pfam/HMM | name | description | E-value |
|---|---|---|---|---|---|---|---|---|
| WP_103501345.1 | 838 | 10 | - | 275 | PF13359<br>PF13613<br>PF01609 | DDE_Tnp_4<br>HTH_Tnp_4<br>DDE_Tnp_1 | DDE superfamily endonuclease<br>Helix-turn-helix of DDE superfamily endonuclease<br>Transposase DDE domain | 8.10E-24<br>3.30E-05<br>6.60E-04 |
| WP_103502972.1 | 1278 | 1467 | + | 62 | LinA | LinA | LinA (linaridin precursor) | 7.70E-10 |
| WP_103502973.1 | 1532 | 1721 | + | 62 | LinA | LinA | LinA (linaridin precursor) | 1.90E-11 |
| WP_103502974.1 | 2145 | 4185 | + | 679 | LinH<br>LinG<br>LinE<br>PF12697<br>PF00561 | LinH<br>LinG<br>LinE<br>Abhydrolase_6<br>Abhydrolase_1 | LinH (Fusion of LinE/LinG)<br>LinG Homolog<br>LinE Homolog<br>Alpha/beta hydrolase family<br>alpha/beta hydrolase fold | 1.90E-254<br>4.00E-115<br>3.20E-97<br>2.40E-17<br>1.20E-10 |
| WP_019435954.1 | 4181 | 4748 | + | 189 | INFERRED GENE | - | - | - |
| WP_103502975.1 | 4964 | 5822 | + | 285 | PF13649<br>PF13489<br>PF08241<br>PF08242<br>PF13847 | Methyltransf_25<br>Methyltransf_23<br>Methyltransf_11<br>Methyltransf_12<br>Methyltransf_31 | Methyltransferase domain<br>Methyltransferase domain<br>Methyltransferase domain<br>Methyltransferase domain<br>Methyltransferase domain | 8.50E-14<br>8.00E-11<br>1.80E-08<br>2.10E-08<br>1.70E-06 |
| WP_103502976.1 | 5799 | 7692 | + | 630 | TIGR02204<br>TIGR02203<br>TIGR00958<br>TIGR03796<br>TIGR01846 | TIGR02204<br>TIGR02203<br>TIGR00958<br>TIGR03796<br>TIGR01846 | MsbA_rel: ABC transporter, permease/ATP-binding protein<br>MsbA_lipidA: lipid A export permease/ATP-binding protein MsbA<br>3a01208: antigen peptide transporter 2<br>NHLM_micro_ABC1: NHLM bacteriocin system ABC transporter, peptidase/ATP-binding protein<br>type_I_sec_HlyB: type I secretion system ATPase | 1.50E-101<br>1.30E-94<br>1.60E-87<br>2.40E-82<br>4.10E-81 |
| WP_146057161.1 | 7963 | 8212 | + | 82 | NO PFAM MATCH | - | - | - |
| WP_103502978.1 | 8309 | 9824 | + | 504 | TIGR02734<br>TIGR03467<br>PF13450<br>TIGR00292<br>PF01946 | TIGR02734<br>TIGR03467<br>NAD_binding_8<br>TIGR00292<br>Thi4 | crtI_fam: phytoene desaturase<br>HpnE: squalene-associated FAD-dependent desaturase<br>NAD(P)-binding Rossmann-like domain<br>TIGR00292: thiazole biosynthesis enzyme<br>Thi4 family | 2.60E-12<br>3.90E-10<br>2.90E-09<br>3.00E-09<br>6.00E-08 |
| WP_103502979.1 | 9826 | 10075 | + | 82 | NO PFAM MATCH | - | - | - |
| WP_103502980.1 | 11162 | 10181 | - | 326 | NO PFAM MATCH | - | - | - |

| index | leader | core | start | end | dir | score |
|---|---|---|---|---|---|---|
| 1 | MARETGLPVL | STPVTASVTIITCGQRVTVAGSREPEIL | 1048 | 1165 | + | 8 |
| 2 | VRWCGGAPPGRPGAAAGGGAWSERRQRISGSRL | PATVTRWPQVMIVTDAVTGVDSTGRPVSRAMPW | 1239 | 1038 | - | 8 |
| 3 | VVRWCGGAPPGRPGAAAGGGAWSERRQRISGSR | LPATVTRWPQVMIVTDAVTGVDSTGRPVSRAMPW | 1242 | 1038 | - | 7 |
| 4 | LTVVRWCGGAPPGRPGAAAGGGAWSERRQRISGS | RLPATVTRWPQVMIVTDAVTGVDSTGRPVSRAMPW | 1248 | 1038 | - | 7 |
| 5 | MRENKLVQNINGADLGGLAEDTAGETRGQGAAVL | ATPAVFTAGVVIGMTVTLAAGSREPVIS | 1278 | 1467 | + | 31 |
| 6 | MRETKIAQHLDSSDLGDLRENSAGENRGKGSAVL | ATPAMFTAGVVIGMTVTLAAGSREPVIS | 1532 | 1721 | + | 32 |
| 7 | VYCQRFPFSRNPFSGIFMTPAKKPAE | ATPGIRNRTWGNRGIRIEFTRQRPEGLSCGTFSR | 7784 | 7967 | + | 7 |
| 8 | MSEITMELVYGGAGPPDGEVAVVRLLGRLPGGWHYEQQHIGPVLIQLRI | MAPLAGPADIEGAVVSALGDPALLNWRVLTADG | 9826 | 10075 | + | 8 |

### Results for WP_103529472.1 [Streptomyces sp. SM12] back to top

previous - next

#### Architecture

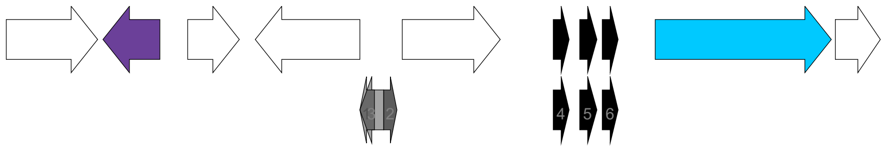

1000 nucleotides

Link to nucleotide sequence

| Accession | start | end | direction | length (aa) | Pfam/HMM | name | description | E-value |
|---|---|---|---|---|---|---|---|---|
| WP_103529474.1 | 5940 | 6993 | + | 350 | TIGR01686<br>TIGR01681 | TIGR01686<br>TIGR01681 | FkbH: FkbH domain<br>HAD-SF-IIIC: HAD phosphatase, family IIIC | 1.40E-113<br>1.00E-18 |
| WP_103529473.1 | 7714 | 7051 | - | 220 | PF01596<br>PF13578<br>PF01135<br>PF13847<br>PF13649 | Methyltransf_3<br>Methyltransf_24<br>PCMT<br>Methyltransf_31<br>Methyltransf_25 | O-methyltransferase<br>Methyltransferase domain<br>Protein-L-isoaspartate(D-aspartate) O-methyltransferase (PCMT)<br>Methyltransferase domain<br>Methyltransferase domain | 7.10E-68<br>3.00E-18<br>1.20E-04<br>1.40E-04<br>1.80E-04 |
| WP_103529469.1 | 8027 | 8633 | + | 201 | TIGR01426<br>PF00201<br>PF04101<br>TIGR01133<br>TIGR00661 | TIGR01426<br>UDPGT<br>Glyco_tran_28_C<br>TIGR01133<br>TIGR00661 | MGT: glycosyltransferase, MGT family<br>UDP-glucoronosyl and UDP-glucosyl transferase<br>Glycosyltransferase family 28 C-terminal domain<br>murG: undecaprenyldiphospho-muramoylpentapeptide beta-N-acetylglucosaminyltransferase<br>MJ1255: conserved hypothetical protein | 4.30E-70<br>3.10E-19<br>5.00E-12<br>3.80E-09<br>5.30E-07 |
| WP_103529470.1 | 10018 | 8815 | - | 400 | PF00149<br>PF12850 | Metallophos<br>Metallophos_2 | Calcineurin-like phosphoesterase<br>Calcineurin-like phosphoesterase superfamily domain | 5.80E-09<br>3.70E-06 |
| WP_103502987.1 | 10511 | 11633 | + | 373 | PF01794 | Ferric_reduct | Ferric reductase like transmembrane component | 2.20E-12 |
| WP_103529471.1 | 12255 | 12435 | + | 59 | LinA | LinA | LinA (linaridin precursor) | 1.10E-05 |
| WP_103502972.1 | 12556 | 12745 | + | 62 | LinA | LinA | LinA (linaridin precursor) | 7.70E-10 |
| WP_103502973.1 | 12810 | 12999 | + | 62 | LinA | LinA | LinA (linaridin precursor) | 1.90E-11 |
| WP_103529472.1 | 13425 | 15450 | + | 674 | LinH<br>LinG<br>LinE<br>PF12697<br>TIGR03695 | LinH<br>LinG<br>LinE<br>Abhydrolase_6<br>TIGR03695 | LinH (Fusion of LinE/LinG)<br>LinG Homolog<br>LinE Homolog<br>Alpha/beta hydrolase family<br>menH_SHCHC: 2-succinyl-6-hydroxy-2,4-cyclohexadiene-1-carboxylate synthase | 8.50E-254<br>1.20E-114<br>4.10E-97<br>1.10E-16<br>1.60E-10 |
| WP_007452032.1 | 15497 | 16025 | + | 176 | INFERRED GENE | - | - | - |

| index | leader | core | start | end | dir | score |
|---|---|---|---|---|---|---|
| 1 | LSRRLGRLSPERESGHGRRLTPGPLADGRYEDPAGPPPG | STPGTIRPTEPQEEA | 10179 | 10014 | - | 6 |
| 2 | MSACCPESRSGGRPEEGRGHPTDPP | CAQGAPGVSTRVAIPKRPLPPGGSLA | 10285 | 10441 | + | 8 |
| 3 | MSLRAGRTLRGTHRVAVGGGPTGAC...GHGRRLTPGPLADGRYEDPAGPPPG | STPGTIRPTEPQEEA | 10287 | 10014 | - | 6 |
| 4 | MLNGIEGKLEGIEPLHEDYQGMARETGLPVL | STPVTASVTIITCGQRVTVAGSREPEIL | 12255 | 12435 | + | 29 |
| 5 | MRENKLVQNINGADLGGLAEDTAGETRGQGAAVL | ATPAVFTAGVVIGMTVTLAAGSREPVIS | 12556 | 12745 | + | 30 |
| 6 | MRETKIAQHLDSSDLGDLRENSAGENRGKGSAVL | ATPAMFTAGVVIGMTVTLAAGSREPVIS | 12810 | 12999 | + | 31 |

### Results for WP_051266157.1 [Streptomyces sp. TAA204] back to top

previous - next

#### Architecture

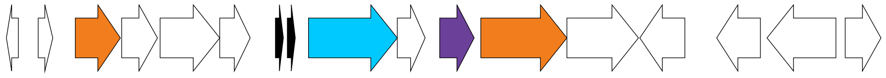

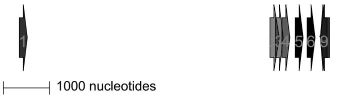

Link to nucleotide sequence

| Accession | start | end | direction | length (aa) | Pfam/HMM | name | description | E-value |
|---|---|---|---|---|---|---|---|---|
| WP_028433649.1 | 125439 | 125166 | - | 90 | PF02700<br>TIGR00302 | PurS<br>TIGR00302 | Phosphoribosylformylglycinamidine (FGAM) synthase<br>TIGR00302: phosphoribosylformylglycinamidine synthase, purS protein | 1.00E-26<br>4.00E-19 |
| WP_028433650.1 | 125849 | 126167 | + | 105 | PF11774 | Lsr2 | Lsr2 | 5.80E-36 |
| WP_051266144.1 | 126654 | 127638 | + | 327 | TIGR01188<br>TIGR03864<br>TIGR01288<br>TIGR04406<br>TIGR03522 | TIGR01188<br>TIGR03864<br>TIGR01288<br>TIGR04406<br>TIGR03522 | drrA: daunorubicin resistance ABC transporter, ATP-binding protein<br>PQQ_ABC_ATP: ABC transporter, ATP-binding subunit, PQQ-dependent alcohol dehydrogenase system<br>nodI: nodulation ABC transporter NodI<br>LPS_export_lptB: LPS export ABC transporter ATP-binding protein<br>GldA_ABC_ATP: gliding motility-associated ABC transporter ATP-binding subunit GldA | 1.10E-70<br>1.40E-49<br>5.00E-49<br>2.90E-39<br>8.60E-38 |
| WP_028433651.1 | 127658 | 128447 | + | 262 | PF01061<br>PF12698<br>TIGR03062 | ABC2_membrane<br>ABC2_membrane_3<br>TIGR03062 | ABC-2 type transporter<br>ABC-2 family transporter protein<br>pip_yhgE_Cterm: YhgE/Pip C-terminal domain | 9.30E-19<br>1.50E-12<br>5.80E-08 |
| WP_037905468.1 | 128512 | 129787 | + | 424 | PF07730 | HisKA_3 | Histidine kinase | 2.70E-14 |
| WP_051266154.1 | 129783 | 130452 | + | 222 | PF00072<br>TIGR02875<br>PF00196<br>TIGR02956<br>TIGR02154 | Response_reg<br>TIGR02875<br>GerE<br>TIGR02956<br>TIGR02154 | Response regulator receiver domain<br>spore_0_A: sporulation transcription factor Spo0A<br>Bacterial regulatory proteins, luxR family<br>TMAO_torS: TMAO reductase sytem sensor TorS<br>PhoB: phosphate regulon transcriptional regulatory protein PhoB | 3.90E-23<br>8.30E-13<br>3.30E-12<br>7.60E-12<br>3.20E-11 |
| WP_158679626.1 | 130999 | 131173 | + | 57 | LinA | LinA | LinA (linaridin precursor) | 1.10E-13 |
| WP_028433653.1 | 131255 | 131435 | + | 59 | LinA | LinA | LinA (linaridin precursor) | 4.50E-16 |
| WP_051266157.1 | 131718 | 133656 | + | 645 | LinH<br>LinG<br>LinE<br>PF12697<br>LinL | LinH<br>LinG<br>LinE<br>Abhydrolase_6<br>LinL | LinH (Fusion of LinE/LinG)<br>LinG Homolog<br>LinE Homolog<br>Alpha/beta hydrolase family<br>LinL | 6.60E-246<br>1.10E-120<br>2.30E-99<br>1.30E-13<br>1.60E-09 |
| WP_003965564.1 | 133652 | 134252 | + | 200 | INFERRED GENE | - | - | - |
| WP_028433654.1 | 134556 | 135315 | + | 252 | PF13649<br>PF13489<br>PF13847<br>PF08241<br>PF08242 | Methyltransf_25<br>Methyltransf_23<br>Methyltransf_31<br>Methyltransf_11<br>Methyltransf_12 | Methyltransferase domain<br>Methyltransferase domain<br>Methyltransferase domain<br>Methyltransferase domain<br>Methyltransferase domain | 1.20E-16<br>2.60E-13<br>1.30E-11<br>1.00E-10<br>2.60E-10 |
| WP_078506149.1 | 135443 | 137324 | + | 626 | TIGR02204<br>TIGR02203<br>TIGR00958<br>TIGR03796<br>TIGR01846 | TIGR02204<br>TIGR02203<br>TIGR00958<br>TIGR03796<br>TIGR01846 | MsbA_rel: ABC transporter, permease/ATP-binding protein<br>MsbA_lipidA: lipid A export permease/ATP-binding protein MsbA<br>3a01208: antigen peptide transporter 2<br>NHLM_micro_ABC1: NHLM bacteriocin system ABC transporter, peptidase/ATP-binding protein<br>type_I_sec_HlyB: type I secretion system ATPase | 2.50E-106<br>8.40E-100<br>5.90E-90<br>7.20E-86<br>3.20E-85 |
| WP_037905356.1 | 137320 | 138874 | + | 517 | TIGR02734<br>TIGR03467<br>TIGR02733<br>TIGR00292<br>TIGR01372 | TIGR02734<br>TIGR03467<br>TIGR02733<br>TIGR00292<br>TIGR01372 | crtI_fam: phytoene desaturase<br>HpnE: squalene-associated FAD-dependent desaturase<br>desat_CrtD: C-3',4' desaturase CrtD<br>TIGR00292: thiazole biosynthesis enzyme<br>soxA: sarcosine oxidase, alpha subunit family | 1.10E-09<br>1.30E-08<br>2.70E-08<br>4.00E-08<br>6.70E-07 |
| WP_037905359.1 | 139871 | 138893 | - | 325 | PF00196 | GerE | Bacterial regulatory proteins, luxR family | 1.70E-05 |
| WP_028433656.1 | 141499 | 140557 | - | 313 | PF01259<br>TIGR00081<br>TIGR02735 | SAICAR_synt<br>TIGR00081<br>TIGR02735 | SAICAR synthetase<br>purC: phosphoribosylaminoimidazolesuccinocarboxamide synthase<br>purC_vibrio: phosphoribosylaminoimidazole-succinocarboxamide synthase | 4.80E-99<br>3.20E-68<br>4.60E-21 |
| WP_028433657.1 | 143161 | 141646 | - | 504 | NO PFAM MATCH | - | - | - |
| WP_028433658.1 | 143362 | 144124 | + | 253 | PF05721 | PhyH | Phytanoyl-CoA dioxygenase (PhyH) | 3.20E-07 |

| index | leader | core | start | end | dir | score |
|---|---|---|---|---|---|---|
| 1 | VQILVAVPEGDGFAARERDGAPCV | AVVERSGEGDDPDTADTAAHAGVP | 125038 | 125185 | + | 9 |
| 2 | MAVHGGTGRDTAVQGPCQPPGRGQSGLSRPHGRSRWKQSGPGYPSPSR | FTLAEPRANSSGLSFT | 130458 | 130653 | + | 7 |
| 3 | VEAERTRISFTIQIYPCRTPCEFLRIE | LYLSLVSGGLETQHWPVTSWFRPSPSVA | 130562 | 130730 | + | 7 |
| 4 | MKLSCCPAANCHEFLFNGSADTGIF | IPAEANPDIDAGVSVRTDTRFARVFR | 130726 | 130882 | + | 8 |
| 5 | MLEEKLNLRIQDELHEGHTGEIAEEATPVL | ATPALAGAFVGGAAVVTGAYAAGQAVG | 130999 | 131173 | + | 30 |
| 6 | MNDVTSHINKSAELSAVEITDADGVVFTDYTPVM | GTPGAFLGGVAIASATVAAFEAGQG | 131255 | 131435 | + | 29 |
| 7 | VSLGGGRLECGGIRQEAWCEEYGE | LSGGTGVQKTVEIYTETTAERSARE | 131699 | 131549 | - | 6 |
| 8 | VPVSLGGGRLECGGIRQEAWCEEYG | ELSGGTGVQKTVEIYTETTAERSARE | 131705 | 131549 | - | 8 |
| 9 | LASDFTSVSSVPVSLGGGRLECGGIRQEAW | CEEYGELSGGTGVQKTVEIYTETTAERSARE | 131735 | 131549 | - | 7 |

### Results for WP_037978036.1 [Streptomyces sp. TAA486] back to top

#### Architecture

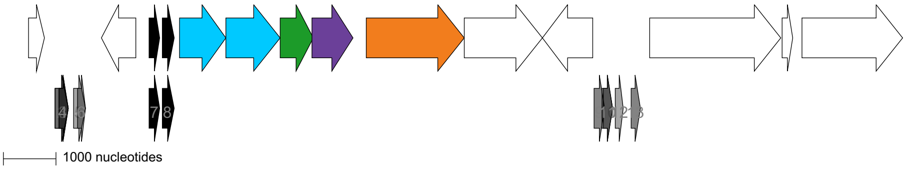

Link to nucleotide sequence

| Accession | start | end | direction | length (aa) | Pfam/HMM | name | description | E-value |
|---|---|---|---|---|---|---|---|---|
| WP_019354801.1 | 0 | 287 | + | 95 | INFERRED GENE | - | - | - |
| WP_158677113.1 | 2033 | 1376 | - | 218 | PF05719 | GPP34 | Golgi phosphoprotein 3 (GPP34) | 3.80E-42 |
| WP_028438060.1 | 2290 | 2482 | + | 63 | LinA | LinA | LinA (linaridin precursor) | 2.70E-12 |
| WP_028438061.1 | 2536 | 2749 | + | 70 | LinA | LinA | LinA (linaridin precursor) | 1.70E-13 |
| WP_037978033.1 | 2852 | 3740 | + | 295 | LinH<br>LinG | LinH<br>LinG | LinH (Fusion of LinE/LinG)<br>LinG Homolog | 8.60E-120<br>1.40E-114 |
| WP_037978036.1 | 3736 | 4774 | + | 345 | LinH<br>LinE<br>PF12697<br>PF12146<br>PF00561 | LinH<br>LinE<br>Abhydrolase_6<br>Hydrolase_4<br>Abhydrolase_1 | LinH (Fusion of LinE/LinG)<br>LinE Homolog<br>Alpha/beta hydrolase family<br>Serine aminopeptidase, S33<br>alpha/beta hydrolase fold | 7.60E-134<br>3.40E-120<br>1.40E-18<br>3.00E-09<br>1.10E-08 |
| WP_051345488.1 | 4773 | 5394 | + | 206 | LinL | LinL | LinL | 3.30E-70 |
| WP_028438062.1 | 5390 | 6161 | + | 256 | PF13649<br>PF08241<br>PF13489<br>PF08242<br>TIGR00537 | Methyltransf_25<br>Methyltransf_11<br>Methyltransf_23<br>Methyltransf_12<br>TIGR00537 | Methyltransferase domain<br>Methyltransferase domain<br>Methyltransferase domain<br>Methyltransferase domain<br>hemK_rel_arch: putative methylase | 1.10E-15<br>5.50E-10<br>3.70E-08<br>4.50E-08<br>5.00E-06 |
| WP_078511858.1 | 6414 | 8274 | + | 619 | TIGR02204<br>TIGR02203<br>TIGR00958<br>TIGR03796<br>TIGR03375 | TIGR02204<br>TIGR02203<br>TIGR00958<br>TIGR03796<br>TIGR03375 | MsbA_rel: ABC transporter, permease/ATP-binding protein<br>MsbA_lipidA: lipid A export permease/ATP-binding protein MsbA<br>3a01208: antigen peptide transporter 2<br>NHLM_micro_ABC1: NHLM bacteriocin system ABC transporter, peptidase/ATP-binding protein<br>type_I_sec_LssB: type I secretion system ATPase | 1.40E-101<br>3.00E-99<br>4.70E-90<br>1.60E-84<br>1.20E-81 |
| WP_028438063.1 | 8270 | 9749 | + | 492 | TIGR02734<br>PF01593<br>TIGR03467<br>PF13450<br>TIGR02733 | TIGR02734<br>Amino_oxidase<br>TIGR03467<br>NAD_binding_8<br>TIGR02733 | crtI_fam: phytoene desaturase<br>Flavin containing amine oxidoreductase<br>HpnE: squalene-associated FAD-dependent desaturase<br>NAD(P)-binding Rossmann-like domain<br>desat_CrtD: C-3',4' desaturase CrtD | 1.30E-12<br>2.20E-10<br>6.40E-09<br>9.30E-09<br>2.20E-08 |
| WP_028438064.1 | 10731 | 9756 | - | 324 | PF01978<br>PF00196<br>TIGR03541 | TrmB<br>GerE<br>TIGR03541 | Sugar-specific transcriptional regulator TrmB<br>Bacterial regulatory proteins, luxR family<br>reg_near_HchA: LuxR family transcriptional regulatory, chaperone HchA-associated | 1.30E-06<br>1.20E-04<br>1.60E-04 |
| WP_158677114.1 | 11800 | 14287 | + | 828 | PF00196<br>PF13424<br>TIGR03020<br>PF13191<br>TIGR03541 | GerE<br>TPR_12<br>TIGR03020<br>AAA_16<br>TIGR03541 | Bacterial regulatory proteins, luxR family<br>Tetratricopeptide repeat<br>EpsA: transcriptional regulator EpsA<br>AAA ATPase domain<br>reg_near_HchA: LuxR family transcriptional regulatory, chaperone HchA-associated | 7.50E-14<br>1.10E-09<br>5.70E-07<br>7.60E-07<br>1.30E-06 |
| WP_018844060.1 | 14311 | 14518 | + | 69 | INFERRED GENE | - | - | - |
| WP_078511851.1 | 14697 | 16620 | + | 640 | PF13556<br>PF09862 | HTH_30<br>DUF2089 | PucR C-terminal helix-turn-helix domain<br>Protein of unknown function (DUF2089) | 1.70E-08<br>6.50E-04 |

| index | leader | core | start | end | dir | score |
|---|---|---|---|---|---|---|
| 1 | VATEMERTRRRERLRRCCAARPLDA...APVRGRSGTAEPPPGGWGREGLIPR | RTPGSPPDGAALQGVYEYSHPLR | 486 | 741 | + | 6 |
| 2 | MERTRRRERLRRCCAARPLDAHGVP...APVRGRSGTAEPPPGGWGREGLIPR | RTPGSPPDGAALQGVYEYSHPLR | 498 | 741 | + | 6 |
| 3 | VMDRRRAPVRGRSGTAEPPPGGWGREGLIPR | RTPGSPPDGAALQGVYEYSHPLR | 576 | 741 | + | 6 |
| 4 | MDRRRAPVRGRSGTAEPPPGGWGREGLIPR | RTPGSPPDGAALQGVYEYSHPLR | 579 | 741 | + | 6 |
| 5 | MNRCHTPRARVHTSAQHRKIVTMTHGRAHSWNESVA | AAAELAETPSEGPRRRRALDTKSEAADSKNGAGPSGA | 851 | 1073 | + | 6 |
| 6 | MTHGRAHSWNESVAAAAELAETPSE | GPRRRRALDTKSEAADSKNGAGPSGA | 917 | 1073 | + | 6 |
| 7 | MLLEKVNNSHLLDTVLTDAANAEDLSPAEPVL | STPAAVAGGALVTAGAFGAGVGLGVTQAQAG | 2290 | 2482 | + | 31 |
| 8 | MTAVSTKVNASQELNGVSIHAGENEWNGRMSPVL | ATPAAVGAGLAGAAAIAGAAAGGIAVGNAAGQAAGG | 2536 | 2749 | + | 28 |
| 9 | LVSNPRPRCSRVGWGLRPALMRDGRGGGVGA | GTLRSTSRMGVGLRAPHSGAGSCGRATPLTGVGRT | 10735 | 10936 | + | 7 |
| 10 | VGGPHRSQVSAVRELCWQLVSDPGGLGGRRAAR | GPARPCVGYCPTRPVRSISRFTESVAAFQVGCLR | 10895 | 11099 | + | 6 |
| 11 | VSAVRELCWQLVSDPGGLGGRRAARGPAR | PCVGYCPTRPVRSISRFTESVAAFQVGCLR | 10919 | 11099 | + | 7 |
| 12 | LPRLCVLSAVDLNGLAHRSHLWVLES | FSSARSVGLRAKVICPITCLSILTGC | 11146 | 11305 | + | 6 |
| 13 | MTDSTPTQVERNARDRNQTPLDFSRT | SQSRPGPCHTSTMNSAHFAATCTMRW | 11456 | 11615 | + | 7 |

### Results for SNX62950.1 [Streptomyces sp. TLI_55] back to top

previous - next

### Architecture

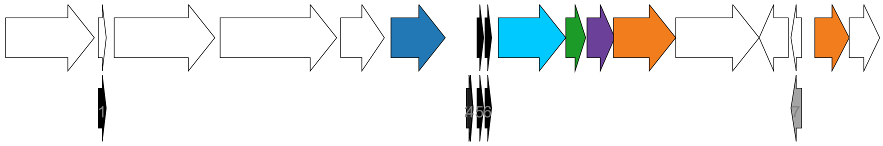

1000 nucleotides

Link to nucleotide sequence

| Accession | start | end | direction | length (aa) | Pfam/HMM | name | description | E-value |
|---|---|---|---|---|---|---|---|---|
| SNX62942.1 | 5158431 | 5160915 | + | 827 | PF00069<br>TIGR03903<br>PF07714<br>PF13845<br>TIGR03724 | Pkinase<br>TIGR03903<br>Pkinase_Tyr<br>Septum_form<br>TIGR03724 | Protein kinase domain<br>TOMM_kin_cyc: TOMM system kinase/cyclase fusion protein<br>Protein tyrosine kinase<br>Septum formation<br>arch_bud32: Kae1-associated kinase Bud32 | 1.30E-33<br>4.40E-33<br>4.20E-13<br>3.90E-08<br>1.70E-04 |
| SNX62943.1 | 5161057 | 5161255 | + | 65 | NO PFAM MATCH | - | - | - |
| SNX62944.1 | 5161499 | 5164325 | + | 941 | TIGR01051<br>PF01131<br>TIGR01054<br>TIGR01057<br>PF13368 | TIGR01051<br>Topoisom_bac<br>TIGR01054<br>TIGR01057<br>Toprim_C_rpt | topA_bact: DNA topoisomerase I<br>DNA topoisomerase<br>rgy: reverse gyrase<br>topA_arch: DNA topoisomerase I<br>Topoisomerase C-terminal repeat | 9.10E-245<br>1.30E-131<br>3.40E-85<br>7.80E-73<br>3.10E-61 |
| SNX62945.1 | 5164471 | 5167744 | + | 1090 | TIGR00041<br>PF02223<br>TIGR00900<br>PF07690<br>PF17380 | TIGR00041<br>Thymidylate_kin<br>TIGR00900<br>MFS_1<br>DUF5401 | DTMP_kinase: dTMP kinase<br>Thymidylate kinase<br>2A0121: H+ Antiporter protein<br>Major Facilitator Superfamily<br>Family of unknown function (DUF5401) | 2.00E-52<br>2.60E-47<br>1.40E-12<br>2.20E-08<br>2.20E-05 |
| SNX62946.1 | 5167875 | 5169081 | + | 401 | TIGR00678<br>PF13177<br>TIGR02397<br>PF00004<br>PF09115 | TIGR00678<br>DNA_pol3_delta2<br>TIGR02397<br>AAA<br>DNApol3-delta_C | holB: DNA polymerase III, delta' subunit<br>DNA polymerase III, delta subunit<br>dnaX_nterm: DNA polymerase III, subunit gamma and tau<br>ATPase family associated with various cellular activities (AAA)<br>DNA polymerase III, delta subunit, C terminal | 1.20E-55<br>7.40E-34<br>3.20E-31<br>6.90E-05<br>6.60E-04 |
| SNX62947.1 | 5169286 | 5170810 | + | 507 | PF00561<br>PF08386<br>LinE<br>LinH<br>TIGR01249 | Abhydrolase_1<br>Abhydrolase_4<br>LinE<br>LinH<br>TIGR01249 | alpha/beta hydrolase fold<br>TAP-like protein<br>LinE Homolog<br>LinH (Fusion of LinE/LinG)<br>pro_imino_pep_1: prolyl aminopeptidase | 1.60E-38<br>3.90E-32<br>1.30E-06<br>1.40E-06<br>3.80E-06 |
| SNX62948.1 | 5171691 | 5171886 | + | 64 | LinA | LinA | LinA (linaridin precursor) | 1.10E-17 |
| SNX62949.1 | 5171930 | 5172122 | + | 63 | LinA | LinA | LinA (linaridin precursor) | 2.00E-11 |
| SNX62950.1 | 5172295 | 5174191 | + | 631 | LinH<br>LinG<br>LinE<br>PF12697<br>PF00561 | LinH<br>LinG<br>LinE<br>Abhydrolase_6<br>Abhydrolase_1 | LinH (Fusion of LinE/LinG)<br>LinG Homolog<br>LinE Homolog<br>Alpha/beta hydrolase family<br>alpha/beta hydrolase fold | 3.50E-239<br>1.40E-114<br>7.50E-97<br>1.70E-13<br>6.50E-11 |
| SNX62951.1 | 5174187 | 5174766 | + | 192 | LinL | LinL | LinL | 3.50E-65 |
| SNX62952.1 | 5174796 | 5175552 | + | 251 | PF13649<br>PF08241<br>PF13489<br>PF08242<br>PF13847 | Methyltransf_25<br>Methyltransf_11<br>Methyltransf_23<br>Methyltransf_12<br>Methyltransf_31 | Methyltransferase domain<br>Methyltransferase domain<br>Methyltransferase domain<br>Methyltransferase domain<br>Methyltransferase domain | 3.80E-13<br>2.70E-09<br>3.80E-08<br>2.40E-06<br>1.60E-05 |
| SNX62953.1 | 5175548 | 5177282 | + | 577 | TIGR02204<br>TIGR02203<br>TIGR03796<br>TIGR00958<br>TIGR01846 | TIGR02204<br>TIGR02203<br>TIGR03796<br>TIGR00958<br>TIGR01846 | MsbA_rel: ABC transporter, permease/ATP-binding protein<br>MsbA_lipidA: lipid A export permease/ATP-binding protein MsbA<br>NHLM_micro_ABC1: NHLM bacteriocin system ABC transporter, peptidase/ATP-binding protein<br>3a01208: antigen peptide transporter 2<br>type_I_sec_HlyB: type I secretion system ATPase | 1.90E-100<br>1.90E-97<br>7.50E-86<br>2.70E-83<br>7.70E-81 |
| SNX62954.1 | 5177278 | 5179633 | + | 784 | TIGR02734<br>TIGR00292<br>TIGR03467<br>TIGR01317<br>TIGR02733 | TIGR02734<br>TIGR00292<br>TIGR03467<br>TIGR01317<br>TIGR02733 | crtI_fam: phytoene desaturase<br>TIGR00292: thiazole biosynthesis enzyme<br>HpnE: squalene-associated FAD-dependent desaturase<br>GOGAT_sm_gam: glutamate synthase, NADH/NADPH, small subunit<br>desat_CrtD: C-3',4' desaturase CrtD | 2.60E-09<br>9.80E-08<br>1.60E-07<br>2.90E-07<br>5.60E-07 |
| SNX62955.1 | 5180460 | 5179617 | - | 280 | PF01551 | Peptidase_M23 | Peptidase family M23 | 1.80E-16 |
| SNX62956.1 | 5180829 | 5180538 | - | 96 | NO PFAM MATCH | - | - | - |
| SNX62957.1 | 5181194 | 5182169 | + | 324 | TIGR01188<br>TIGR03864<br>TIGR03522<br>TIGR01288<br>TIGR03873 | TIGR01188<br>TIGR03864<br>TIGR03522<br>TIGR01288<br>TIGR03873 | drrA: daunorubicin resistance ABC transporter, ATP-binding protein<br>PQQ_ABC_ATP: ABC transporter, ATP-binding subunit, PQQ-dependent alcohol dehydrogenase system<br>GldA_ABC_ATP: gliding motility-associated ABC transporter ATP-binding subunit GldA<br>nodI: nodulation ABC transporter NodI<br>F420-0_ABC_ATP: proposed F420-0 ABC transporter, ATP-binding protein | 7.30E-58<br>1.30E-46<br>2.10E-41<br>1.30E-39<br>1.40E-39 |
| SNX62958.1 | 5182165 | 5183017 | + | 283 | PF12698<br>PF01061<br>TIGR01247<br>TIGR00025<br>PF12679 | ABC2_membrane_3<br>ABC2_membrane<br>TIGR01247<br>TIGR00025<br>ABC2_membrane_2 | ABC-2 family transporter protein<br>ABC-2 type transporter<br>drrB: daunorubicin resistance ABC transporter membrane protein<br>Mtu_efflux: ABC transporter efflux protein, DrrB family<br>ABC-2 family transporter protein | 2.40E-17<br>3.70E-13<br>3.30E-10<br>2.80E-08<br>5.10E-05 |

| index | leader | core | start | end | dir | score |
|---|---|---|---|---|---|---|
| 1 | MESGPAIFAGSVFALFGAGLLA | WTAVRVRHREPVAHGVSPVASVMIASSAAVIALVAGAWCFTQL | 5161057 | 5161255 | + | 10 |

| | | | | | | |
|---|---|---|---|---|---|---|
| 2 | LTLRGMRLNGGIVLSDMPVIA | PTPFRGSSLCIIAFGVTRVVSGRRSRCRSDAISCNAPHRLA | 5171400 | 5171589 | + | 6 |
| 3 | MRLNGGIVLSDMPVIA | PTPFRGSSLCIIAFGVTRVVSGRRSRCRSDAISCNAPHRLA | 5171415 | 5171589 | + | 7 |
| 4 | VLSDMPVIAPTPFRGSSLCIIAFGV | TRVVSGRRSRCRSDAISCNAPHRLA | 5171436 | 5171589 | + | 8 |
| 5 | MEQKIAAFMNEQDLGSLEQGYRDDAPMPVL | ATPAAAVATCWYVGGAAAGAAAVVGAYVTGRVVG | 5171691 | 5171886 | + | 30 |
| 6 | MSRIVDSVVAVTPAHLTDDSQGEFGAMAPVQ | ATPTLVAFGVGFAGGAGAAWVTVQAYEAGAND | 5171930 | 5172122 | + | 31 |
| 7 | MVNGEIQNTGTECFSVWVQWTRDFI...SPYTKHATQCGAGVSPVNLRLSPYQ | LTTTGRLKVCQGAADTTDCGSAISLTTWPINNSKKTASTNQSPPR | 5180829 | 5180538 | - | 6 |

### Results for WP_078511723.1 [Streptomyces sp. TAA486] back to top

previous - next

#### Architecture

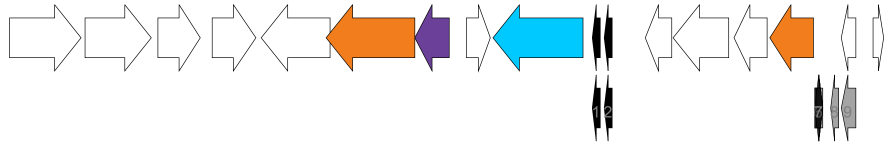

1000 nucleotides

Link to nucleotide sequence

| Accession | start | end | direction | length (aa) | Pfam/HMM | name | description | E-value |
|---|---|---|---|---|---|---|---|---|
| WP_028437548.1 | 77297 | 78926 | + | 542 | NO PFAM MATCH | - | - | - |
| WP_028437549.1 | 79008 | 80505 | + | 498 | NO PFAM MATCH | - | - | - |
| WP_028437550.1 | 80664 | 81615 | + | 316 | PF01259<br>TIGR00081<br>TIGR02735 | SAICAR_synt<br>TIGR00081<br>TIGR02735 | SAICAR synthetase<br>purC: phosphoribosylaminoimidazolesuccinocarboxamide synthase<br>purC_vibrio: phosphoribosylaminoimidazole-succinocarboxamide synthase | 1.80E-98<br>7.60E-70<br>4.50E-23 |
| WP_028437551.1 | 81887 | 82889 | + | 333 | PF00196 | GerE | Bacterial regulatory proteins, luxR family | 1.20E-04 |
| WP_078511721.1 | 84579 | 83010 | - | 522 | TIGR02734<br>TIGR03467<br>PF13450<br>TIGR00292<br>PF01946 | TIGR02734<br>TIGR03467<br>NAD_binding_8<br>TIGR00292<br>Thi4 | crtI_fam: phytoene desaturase<br>HpnE: squalene-associated FAD-dependent desaturase<br>NAD(P)-binding Rossmann-like domain<br>TIGR00292: thiazole biosynthesis enzyme<br>Thi4 family | 2.40E-12<br>1.40E-09<br>6.10E-08<br>7.50E-08<br>2.70E-06 |
| WP_078511722.1 | 86495 | 84473 | - | 673 | TIGR02204<br>TIGR02203<br>TIGR00958<br>TIGR01846<br>TIGR03796 | TIGR02204<br>TIGR02203<br>TIGR00958<br>TIGR01846<br>TIGR03796 | MsbA_rel: ABC transporter, permease/ATP-binding protein<br>MsbA_lipidA: lipid A export permease/ATP-binding protein MsbA<br>3a01208: antigen peptide transporter 2<br>type_I_sec_HlyB: type I secretion system ATPase<br>NHLM_micro_ABC1: NHLM bacteriocin system ABC transporter, peptidase/ATP-binding protein | 5.80E-101<br>1.10E-94<br>2.10E-85<br>6.90E-81<br>1.10E-80 |
| WP_037977524.1 | 87261 | 86502 | - | 252 | PF13649<br>PF13489<br>PF08241<br>PF13847<br>TIGR01983 | Methyltransf_25<br>Methyltransf_23<br>Methyltransf_11<br>Methyltransf_31<br>TIGR01983 | Methyltransferase domain<br>Methyltransferase domain<br>Methyltransferase domain<br>Methyltransferase domain<br>UbiG: 3-demethylubiquinone-9 3-O-methyltransferase | 1.40E-18<br>1.20E-13<br>5.60E-13<br>4.10E-12<br>6.00E-11 |
| WP_006132582.1 | 87666 | 88218 | + | 184 | INFERRED GENE | - | - | - |
| WP_078511723.1 | 90305 | 88262 | - | 680 | LinH<br>LinG<br>LinE<br>PF12697<br>PF00561 | LinH<br>LinG<br>LinE<br>Abhydrolase_6<br>Abhydrolase_1 | LinH (Fusion of LinE/LinG)<br>LinG Homolog<br>LinE Homolog<br>Alpha/beta hydrolase family<br>alpha/beta hydrolase fold | 8.80E-247<br>6.40E-122<br>5.70E-98<br>1.80E-15<br>2.30E-09 |
| WP_028437555.1 | 90692 | 90512 | - | 59 | LinA | LinA | LinA (linaridin precursor) | 4.50E-16 |
| WP_158677079.1 | 90972 | 90798 | - | 57 | LinA | LinA | LinA (linaridin precursor) | 5.60E-15 |
| WP_028437556.1 | 92315 | 91712 | - | 200 | PF00072<br>TIGR02875<br>TIGR02154<br>PF00196<br>TIGR02956 | Response_reg<br>TIGR02875<br>TIGR02154<br>GerE<br>TIGR02956 | Response regulator receiver domain<br>spore_0_A: sporulation transcription factor Spo0A<br>PhoB: phosphate regulon transcriptional regulatory protein PhoB<br>Bacterial regulatory proteins, luxR family<br>TMAO_torS: TMAO reductase sytem sensor TorS | 2.80E-25<br>2.80E-13<br>3.20E-12<br>4.90E-12<br>9.60E-11 |
| WP_078511744.1 | 93628 | 92362 | - | 421 | PF07730 | HisKA_3 | Histidine kinase | 3.60E-15 |
| WP_106965432.1 | 94492 | 93736 | - | 251 | PF01061<br>PF12698<br>TIGR03062 | ABC2_membrane<br>ABC2_membrane_3<br>TIGR03062 | ABC-2 type transporter<br>ABC-2 family transporter protein<br>pip_yhgE_Cterm: YhgE/Pip C-terminal domain | 8.20E-20<br>8.20E-13<br>5.50E-08 |
| WP_051345363.1 | 95529 | 94545 | - | 327 | TIGR01188<br>TIGR03864<br>TIGR01288<br>TIGR04406<br>TIGR03740 | TIGR01188<br>TIGR03864<br>TIGR01288<br>TIGR04406<br>TIGR03740 | drrA: daunorubicin resistance ABC transporter, ATP-binding protein<br>PQQ_ABC_ATP: ABC transporter, ATP-binding subunit, PQQ-dependent alcohol dehydrogenase system<br>nodI: nodulation ABC transporter NodI<br>LPS_export_lptB: LPS export ABC transporter ATP-binding protein<br>galliderm_ABC: lantibiotic protection ABC transporter, ATP-binding subunit | 2.20E-69<br>5.10E-48<br>1.20E-46<br>1.20E-37<br>2.00E-36 |
| WP_028437558.1 | 96500 | 96182 | - | 105 | PF11774 | Lsr2 | Lsr2 | 4.20E-36 |
| WP_028437559.1 | 96882 | 97140 | + | 85 | PF02700<br>TIGR00302 | PurS<br>TIGR00302 | Phosphoribosylformylglycinamidine (FGAM) synthase<br>TIGR00302: phosphoribosylformylglycinamidine synthase, purS protein | 1.30E-27<br>6.30E-20 |

| index | leader | core | start | end | dir | score |
|---|---|---|---|---|---|---|
| 1 | MKDVTSQINESAELASVAITDAEGVVFTDYTPVM | GTPGAFLGGVAIASATVAAFEAGQD | 90692 | 90512 | - | 29 |
| 2 | MLEEKLNLQIQDQLHEAHEGEIAEEAAPVL | ATPALAGAFVGGAAVVTGAYAAGKAVG | 90972 | 90798 | - | 30 |
| 3 | VLSRVRMTNLMRGQGGLHTEAGAPVGERCAW | RTVFRTVDTGRPDSGPETEGTPVRGPGFSHC | 95568 | 95757 | + | 7 |
| 4 | VRMTNLMRGQGGLHTEAGAPVGERCAWRT | VFRTVDTGRPDSGPETEGTPVRGPGFSHC | 95580 | 95757 | + | 6 |
| 5 | MTNLMRGQGGLHTEAGAPVGERCAWRTV | FRTVDTGRPDSGPETEGTPVRGPGFSHC | 95586 | 95757 | + | 6 |
| 6 | MRGQGGLHTEAGAPVGERCAWRTVFR | TVDTGRPDSGPETEGTPVRGPGFSHC | 95598 | 95757 | + | 7 |
| 7 | VREPRSPDRGSFRFRTGIRPPRIHGPKDR | SPGAPLTHRRAGLRVQAALPAHEVRHAHS | 95751 | 95574 | - | 6 |
| 8 | LPRTPGDPLESGPSPRGEPHGARGCSSVVERPLSKGEAQC | AIPVTRSRSHMDRHR | 96109 | 95941 | - | 6 |
| 9 | MAQRVVVTLSDDLDGGEASETVSFG...KKLREDLARFVEAGRKRAKSGKAYH | RTAVAPDPRAVRAWAESNGFEVPARGRIPKKVYDAFNASR | 96500 | 96182 | - | 6 |

### Results for RXS68531.1 [Streptomyces sp. TM32] back to top

previous - next

#### Architecture

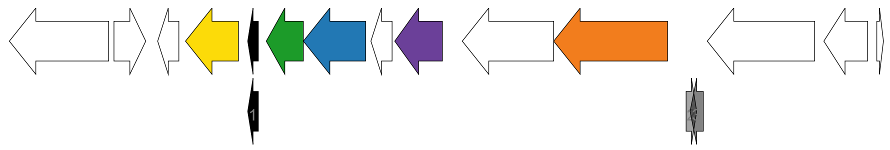

1000 nucleotides

Link to nucleotide sequence

| Accession | start | end | direction | length (aa) | Pfam/HMM | name | description | E-value |
|---|---|---|---|---|---|---|---|---|
| RXS68526.1 | 1779 | 171 | - | 535 | PF02113<br>TIGR00666 | Peptidase_S13<br>TIGR00666 | D-Ala-D-Ala carboxypeptidase 3 (S13) family<br>PBP4: D-alanyl-D-alanine carboxypeptidase/D-alanyl-D-alanine-endopeptidase | 3.50E-75<br>1.50E-56 |
| RXS68537.1 | 1865 | 2360 | + | 164 | PF00719 | Pyrophosphatase | Inorganic pyrophosphatase | 2.50E-53 |
| RXS68527.1 | 2909 | 2549 | - | 119 | NO PFAM MATCH | - | - | - |
| RXS68528.1 | 3854 | 2999 | - | 284 | LinG<br>LinH<br>TIGR04033 | LinG<br>LinH<br>TIGR04033 | LinG Homolog<br>LinH (Fusion of LinE/LinG)<br>export_SdpB: antimicrobial peptide system protein, SdpB family | 2.90E-111<br>1.60E-79<br>2.30E-04 |
| RXS68529.1 | 4182 | 3999 | - | 60 | LinA | LinA | LinA (linaridin precursor) | 7.50E-14 |
| RXS68530.1 | 4896 | 4308 | - | 195 | LinL | LinL | LinL | 1.90E-55 |
| RXS68531.1 | 5912 | 4892 | - | 339 | LinE<br>LinH<br>PF12697<br>TIGR02427<br>PF00561 | LinE<br>LinH<br>Abhydrolase_6<br>TIGR02427<br>Abhydrolase_1 | LinE Homolog<br>LinH (Fusion of LinE/LinG)<br>Alpha/beta hydrolase family<br>protocat_pcaD: 3-oxoadipate enol-lactonase<br>alpha/beta hydrolase fold | 7.30E-102<br>6.00E-52<br>1.80E-14<br>2.80E-09<br>5.20E-08 |
| RXS68532.1 | 6332 | 5984 | - | 115 | PF13518<br>PF01527<br>PF13551<br>PF13011<br>PF13384 | HTH_28<br>HTH_Tnp_1<br>HTH_29<br>LZ_Tnp_IS481<br>HTH_23 | Helix-turn-helix domain<br>Transposase<br>Winged helix-turn helix<br>leucine-zipper of insertion element IS481<br>Homeodomain-like domain | 4.30E-09<br>6.20E-06<br>8.40E-06<br>1.40E-04<br>2.60E-04 |
| RXS68533.1 | 7133 | 6374 | - | 252 | PF13649<br>PF08241<br>PF13489<br>PF08242<br>PF13847 | Methyltransf_25<br>Methyltransf_11<br>Methyltransf_23<br>Methyltransf_12<br>Methyltransf_31 | Methyltransferase domain<br>Methyltransferase domain<br>Methyltransferase domain<br>Methyltransferase domain<br>Methyltransferase domain | 5.00E-17<br>4.10E-12<br>1.80E-10<br>6.40E-10<br>9.90E-09 |
| RXS68534.1 | 8932 | 7450 | - | 493 | PF13450<br>TIGR02734<br>TIGR00292<br>PF01593<br>TIGR03467 | NAD_binding_8<br>TIGR02734<br>TIGR00292<br>Amino_oxidase<br>TIGR03467 | NAD(P)-binding Rossmann-like domain<br>crtI_fam: phytoene desaturase<br>TIGR00292: thiazole biosynthesis enzyme<br>Flavin containing amine oxidoreductase<br>HpnE: squalene-associated FAD-dependent desaturase | 1.80E-08<br>5.60E-08<br>9.70E-08<br>1.50E-07<br>2.80E-07 |
| RXS68538.1 | 10758 | 8928 | - | 609 | TIGR02204<br>TIGR02203<br>TIGR00958<br>TIGR03375<br>TIGR03796 | TIGR02204<br>TIGR02203<br>TIGR00958<br>TIGR03375<br>TIGR03796 | MsbA_rel: ABC transporter, permease/ATP-binding protein<br>MsbA_lipidA: lipid A export permease/ATP-binding protein MsbA<br>3a01208: antigen peptide transporter 2<br>type_I_sec_LssB: type I secretion system ATPase<br>NHLM_micro_ABC1: NHLM bacteriocin system ABC transporter, peptidase/ATP-binding protein | 2.10E-99<br>1.70E-95<br>5.30E-89<br>2.90E-80<br>4.30E-80 |
| RXS68535.1 | 13134 | 11397 | - | 578 | PF06738<br>PF12821 | ThrE<br>ThrE_2 | Putative threonine/serine exporter<br>Threonine/Serine exporter, ThrE | 5.70E-72<br>2.90E-19 |
| RXS68536.1 | 13989 | 13275 | - | 237 | PF09335 | SNARE_assoc | SNARE associated Golgi protein | 3.20E-24 |
| WP_015104329.1 | 14136 | 14235 | + | 33 | INFERRED GENE | - | - | - |

| index | leader | core | start | end | dir | score |
|---|---|---|---|---|---|---|
| 1 | MSVVADFANTELADVTPGRIGNDATPTM | LTPLATLATPEGVAVTSATAYALHEVTKDIQR | 4182 | 3999 | - | 32 |
| 2 | LPRSAGAGQGCPVAQSVRSGSRCYG | ALENGAARGEVCALAADRRPGIATAP | 11065 | 11221 | + | 6 |
| 3 | VSPRGRPLAVPSPSPRRTLGRSAGVADAVGAG | TGRSLQGAVAIPGRRSAARAHTSPRAAPFSKAP | 11331 | 11133 | - | 7 |

### Results for WP_079127392.1 [Streptomyces sp. TP-A0874] back to top

previous - next

#### Architecture

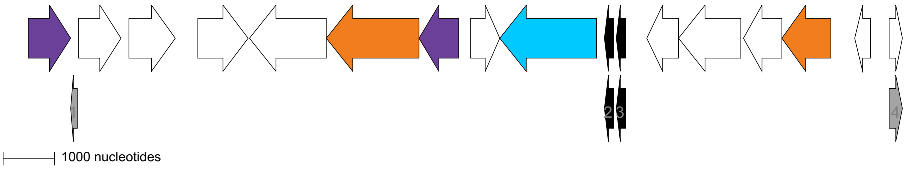

Link to nucleotide sequence

| Accession | start | end | direction | length (aa) | Pfam/HMM | name | description | E-value |
|---|---|---|---|---|---|---|---|---|
| WP_069813889.1 | 585182 | 586013 | + | 276 | TIGR01444<br>PF05050<br>TIGR04364<br>PF01135<br>TIGR00080 | TIGR01444<br>Methyltransf_21<br>TIGR04364<br>PCMT<br>TIGR00080 | fkbM_fam: methyltransferase, FkbM family<br>Methyltransferase FkbM domain<br>methyltran_FxLD: methyltransferase, FxLD system<br>Protein-L-isoaspartate(D-aspartate) O-methyltransferase (PCMT)<br>pimt: protein-L-isoaspartate O-methyltransferase | 2.10E-30<br>9.90E-19<br>3.10E-08<br>2.90E-07<br>3.50E-06 |
| WP_069813891.1 | 586159 | 586996 | + | 278 | PF12706<br>PF13483 | Lactamase_B_2<br>Lactamase_B_3 | Beta-lactamase superfamily domain<br>Beta-lactamase superfamily domain | 9.40E-17<br>2.30E-09 |
| WP_069813893.1 | 587130 | 588036 | + | 301 | PF00355<br>PF13806<br>TIGR02378<br>PF09990<br>TIGR02377 | Rieske<br>Rieske_2<br>TIGR02378<br>DUF2231<br>TIGR02377 | Rieske [2Fe-2S] domain<br>Rieske-like [2Fe-2S] domain<br>nirD_assim_sml: nitrite reductase [NAD(P)H], small subunit<br>Predicted membrane protein (DUF2231)<br>MocE_fam_FeS: Rieske [2Fe-2S] domain protein, MocE subfamily | 1.00E-16<br>1.20E-07<br>1.90E-05<br>2.20E-05<br>2.70E-04 |
| WP_069813895.1 | 588478 | 589453 | + | 324 | PF13412<br>TIGR03541<br>PF01978 | HTH_24<br>TIGR03541<br>TrmB | Winged helix-turn-helix DNA-binding<br>reg_near_HchA: LuxR family transcriptional regulatory, chaperone HchA-associated<br>Sugar-specific transcriptional regulator TrmB | 9.40E-06<br>6.70E-05<br>7.80E-05 |
| WP_069813897.1 | 590985 | 589494 | - | 496 | TIGR02734<br>TIGR03467<br>PF13450<br>TIGR00292<br>PF01593 | TIGR02734<br>TIGR03467<br>NAD_binding_8<br>TIGR00292<br>Amino_oxidase | crtI_fam: phytoene desaturase<br>HpnE: squalene-associated FAD-dependent desaturase<br>NAD(P)-binding Rossmann-like domain<br>TIGR00292: thiazole biosynthesis enzyme<br>Flavin containing amine oxidoreductase | 2.60E-11<br>2.20E-09<br>1.20E-07<br>1.90E-07<br>3.20E-06 |
| WP_069813900.1 | 592784 | 590981 | - | 600 | TIGR02204<br>TIGR02203<br>TIGR00958<br>TIGR03796<br>TIGR01846 | TIGR02204<br>TIGR02203<br>TIGR00958<br>TIGR03796<br>TIGR01846 | MsbA_rel: ABC transporter, permease/ATP-binding protein<br>MsbA_lipidA: lipid A export permease/ATP-binding protein MsbA<br>3a01208: antigen peptide transporter 2<br>NHLM_micro_ABC1: NHLM bacteriocin system ABC transporter, peptidase/ATP-binding protein<br>type_I_sec_HlyB: type I secretion system ATPase | 4.10E-107<br>1.50E-105<br>1.80E-91<br>2.00E-88<br>2.60E-88 |
| WP_069814016.1 | 593549 | 592787 | - | 253 | PF13649<br>PF08241<br>PF13489<br>PF08242<br>TIGR01983 | Methyltransf_25<br>Methyltransf_11<br>Methyltransf_23<br>Methyltransf_12<br>TIGR01983 | Methyltransferase domain<br>Methyltransferase domain<br>Methyltransferase domain<br>Methyltransferase domain<br>UbiG: 3-demethylubiquinone-9 3-O-methyltransferase | 4.30E-17<br>3.20E-11<br>7.00E-09<br>9.60E-09<br>3.40E-07 |
| WP_019435954.1 | 593796 | 594357 | + | 187 | INFERRED GENE | - | - | - |
| WP_079127392.1 | 596249 | 594353 | - | 631 | LinH<br>LinG<br>LinE<br>PF12697<br>TIGR02427 | LinH<br>LinG<br>LinE<br>Abhydrolase_6<br>TIGR02427 | LinH (Fusion of LinE/LinG)<br>LinG Homolog<br>LinE Homolog<br>Alpha/beta hydrolase family<br>protocat_pcaD: 3-oxoadipate enol-lactonase | 4.40E-247<br>3.50E-120<br>5.10E-108<br>4.60E-14<br>1.20E-08 |
| WP_069813904.1 | 596575 | 596395 | - | 59 | LinA | LinA | LinA (linaridin precursor) | 7.90E-15 |
| WP_170837784.1 | 596802 | 596628 | - | 57 | LinA | LinA | LinA (linaridin precursor) | 2.20E-12 |
| WP_069813906.1 | 597830 | 597206 | - | 207 | PF00072<br>TIGR02875<br>PF00196<br>TIGR02154<br>TIGR02956 | Response_reg<br>TIGR02875<br>GerE<br>TIGR02154<br>TIGR02956 | Response regulator receiver domain<br>spore_0_A: sporulation transcription factor Spo0A<br>Bacterial regulatory proteins, luxR family<br>PhoB: phosphate regulon transcriptional regulatory protein PhoB<br>TMAO_torS: TMAO reductase sytem sensor TorS | 5.80E-24<br>3.10E-13<br>5.10E-13<br>2.70E-10<br>1.20E-09 |
| WP_069813908.1 | 599044 | 597826 | - | 405 | PF07730 | HisKA_3 | Histidine kinase | 9.50E-15 |
| WP_069813909.1 | 599856 | 599088 | - | 255 | PF01061<br>PF12698<br>TIGR03062<br>TIGR01247 | ABC2_membrane<br>ABC2_membrane_3<br>TIGR03062<br>TIGR01247 | ABC-2 type transporter<br>ABC-2 family transporter protein<br>pip_yhgE_Cterm: YhgE/Pip C-terminal domain<br>drrB: daunorubicin resistance ABC transporter membrane protein | 4.10E-15<br>1.50E-12<br>1.70E-07<br>1.00E-05 |
| WP_069813910.1 | 600788 | 599852 | - | 311 | TIGR01188<br>TIGR03864<br>TIGR01288<br>TIGR03265<br>TIGR04406 | TIGR01188<br>TIGR03864<br>TIGR01288<br>TIGR03265<br>TIGR04406 | drrA: daunorubicin resistance ABC transporter, ATP-binding protein<br>PQQ_ABC_ATP: ABC transporter, ATP-binding subunit, PQQ-dependent alcohol dehydrogenase system<br>nodI: nodulation ABC transporter NodI<br>PhnT2: putative 2-aminoethylphosphonate ABC transporter, ATP-binding protein<br>LPS_export_lptB: LPS export ABC transporter ATP-binding protein | 5.70E-68<br>2.70E-49<br>5.00E-48<br>1.80E-37<br>2.50E-36 |
| WP_069813911.1 | 601578 | 601260 | - | 105 | PF11774 | Lsr2 | Lsr2 | 2.30E-37 |
| WP_069813912.1 | 601919 | 602186 | + | 88 | PF02700<br>TIGR00302 | PurS<br>TIGR00302 | Phosphoribosylformylglycinamidine (FGAM) synthase<br>TIGR00302: phosphoribosylformylglycinamidine synthase, purS protein | 2.20E-28<br>4.70E-21 |

| index | leader | core | start | end | dir | score |
|---|---|---|---|---|---|---|
| 1 | LTRATGADRSGPMR | STPRRSSHARPSTARRHAATARPG | 586126 | 586009 | - | 6 |

| | | | | | | |
|---|---|---|---|---|---|---|
| 2 | MIEMDSYINRSAALADIDVEDTAGTLFSGAHPVL | VTPAAFLGGVAVASATVAAFEAGRG | 596575 | 596395 | - | 31 |
| 3 | MLEEKLNTTDLGLLDETTEGAVTESAMVVL | ATPAAAAAFVAGAAIVTGAYAAGKAVG | 596802 | 596628 | - | 30 |
| 4 | MARVVVDVMLKPEILDPQGQAVQRA...PVDDAALARVRELAETFLANTVIED | FTVRVEPETEKAAS | 601919 | 602186 | + | 6 |

### Results for WP_149180181.1 [Streptomyces sp. TRM49041] back to top

previous - next

#### Architecture

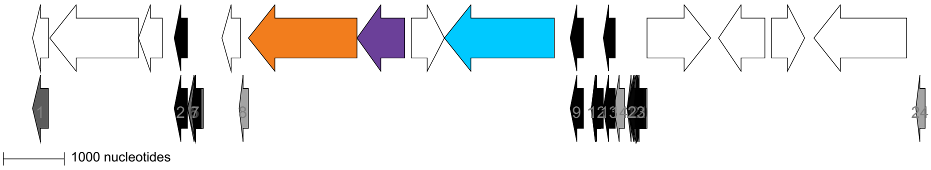

Link to nucleotide sequence

| Accession | start | end | direction | length (aa) | Pfam/HMM | name | description | E-value |
|---|---|---|---|---|---|---|---|---|
| WP_149180163.1 | 64075 | 63823 | - | 83 | NO PFAM MATCH | - | - | - |
| WP_149180164.1 | 65555 | 64097 | - | 485 | TIGR03467<br>TIGR02734<br>PF13450<br>TIGR00292<br>PF01593 | TIGR03467<br>TIGR02734<br>NAD_binding_8<br>TIGR00292<br>Amino_oxidase | HpnE: squalene-associated FAD-dependent desaturase<br>crtI_fam: phytoene desaturase<br>NAD(P)-binding Rossmann-like domain<br>TIGR00292: thiazole biosynthesis enzyme<br>Flavin containing amine oxidoreductase | 1.10E-10<br>6.00E-10<br>2.70E-09<br>3.70E-09<br>3.50E-08 |
| WP_149180165.1 | 65944 | 65545 | - | 132 | NO PFAM MATCH | - | - | - |
| WP_149180166.1 | 66365 | 66143 | - | 73 | LinA | LinA | LinA (linaridin precursor) | 2.00E-11 |
| WP_149180167.1 | 67219 | 66925 | - | 97 | NO PFAM MATCH | - | - | - |
| WP_149180168.1 | 69131 | 67355 | - | 591 | TIGR02204<br>TIGR02203<br>TIGR00958<br>TIGR03375<br>TIGR03796 | TIGR02204<br>TIGR02203<br>TIGR00958<br>TIGR03375<br>TIGR03796 | MsbA_rel: ABC transporter, permease/ATP-binding protein<br>MsbA_lipidA: lipid A export permease/ATP-binding protein MsbA<br>3a01208: antigen peptide transporter 2<br>type_I_sec_LssB: type I secretion system ATPase<br>NHLM_micro_ABC1: NHLM bacteriocin system ABC transporter, peptidase/ATP-binding protein | 8.80E-108<br>6.80E-101<br>1.20E-89<br>1.00E-86<br>3.40E-81 |
| WP_149180169.1 | 69929 | 69149 | - | 259 | PF13649<br>PF08241<br>PF13489<br>PF08242<br>PF13847 | Methyltransf_25<br>Methyltransf_11<br>Methyltransf_23<br>Methyltransf_12<br>Methyltransf_31 | Methyltransferase domain<br>Methyltransferase domain<br>Methyltransferase domain<br>Methyltransferase domain<br>Methyltransferase domain | 1.40E-16<br>2.80E-11<br>1.80E-09<br>5.50E-09<br>2.30E-08 |
| WP_019435954.1 | 70027 | 70582 | + | 185 | INFERRED GENE | - | - | - |
| WP_149180181.1 | 72393 | 70578 | - | 604 | LinH<br>LinG<br>LinE<br>PF12697<br>PF12146 | LinH<br>LinG<br>LinE<br>Abhydrolase_6<br>Hydrolase_4 | LinH (Fusion of LinE/LinG)<br>LinG Homolog<br>LinE Homolog<br>Alpha/beta hydrolase family<br>Serine aminopeptidase, S33 | 6.40E-247<br>2.60E-115<br>1.30E-99<br>5.40E-13<br>1.40E-08 |
| WP_149180170.1 | 72854 | 72653 | - | 66 | LinA | LinA | LinA (linaridin precursor) | 8.20E-17 |
| WP_149180171.1 | 73375 | 73180 | - | 64 | LinA | LinA | LinA (linaridin precursor) | 3.40E-13 |
| WP_149180172.1 | 73907 | 74939 | + | 343 | PF14027 | Questin_oxidase | Questin oxidase-like | 7.00E-31 |
| WP_149180182.1 | 75832 | 75070 | - | 253 | PF02900 | LigB | Catalytic LigB subunit of aromatic ring-opening dioxygenase | 4.50E-43 |
| WP_149180173.1 | 75955 | 76501 | + | 181 | PF12802<br>PF01047<br>TIGR02337<br>PF13463<br>PF09339 | MarR_2<br>MarR<br>TIGR02337<br>HTH_27<br>HTH_IclR | MarR family<br>MarR family<br>HpaR: homoprotocatechuate degradation operon regulator, HpaR<br>Winged helix DNA-binding domain<br>IclR helix-turn-helix domain | 1.30E-13<br>6.00E-11<br>2.30E-08<br>1.40E-06<br>1.30E-05 |
| WP_149180174.1 | 78179 | 76643 | - | 511 | TIGR00711<br>PF07690<br>TIGR00710<br>TIGR00880<br>TIGR00881 | TIGR00711<br>MFS_1<br>TIGR00710<br>TIGR00880<br>TIGR00881 | efflux_EmrB: drug resistance MFS transporter, drug:H+ antiporter-2 (14 Spanner) (DHA2) family<br>Major Facilitator Superfamily<br>efflux_Bcr_CflA: drug resistance transporter, Bcr/CflA subfamily<br>2_A_01_02: multidrug resistance protein<br>2A0104: phosphoglycerate transporter family protein | 4.60E-77<br>2.70E-53<br>4.20E-22<br>2.40E-17<br>4.90E-14 |

| index | leader | core | start | end | dir | score |
|---|---|---|---|---|---|---|
| 1 | MTVYIEYVELSHRGPDTSDGEVAVL...LPRHWRCRHAATADRIRIWITVPGD | ATPERVRAAVAESLNDPALAAWRLEEAG | 64075 | 63823 | - | 8 |
| 2 | MAAAPHVVTWSANGAPKLAEISIRDARGHVVPGSLPVM | ATPGAFTAGVAIASAVLGANELGKAAAGDSGIQPA | 66365 | 66143 | - | 16 |
| 3 | LVGGAALSGALRETDQGKISDRAVPVM | ATPTLAFAAPA | 66546 | 66429 | - | 18 |
| 4 | LAGPGRTHHQLLVGGAALSGALRETDQGKISDRAVPVM | ATPTLAFAAPA | 66579 | 66429 | - | 18 |
| 5 | VPLAGPGRTHHQLLVGGAALSGALRETDQGKISDRAVPVM | ATPTLAFAAPA | 66585 | 66429 | - | 20 |
| 6 | LSGSRAVGRARA | HTPSAVGWRRRPVRRAARDRPGEDQ...RAGVTVVGAAYAVGYAVGGDAEEGR | 66598 | 66364 | - | 6 |
| 7 | VAAASDQRPSLSGSRAVGRARA | HTPSAVGWRRRPVRRAARDRPGEDQ...RAGVTVVGAAYAVGYAVGGDAEEGR | 66628 | 66364 | - | 6 |
| 8 | VSLAGCCTRSRRHLPRPPGGRRP | RSASVGCRKRTTPCRTDVVAEDEK | 67359 | 67215 | - | 6 |
| 9 | MFTVTEFANTSADLTGMEMRDTRGHIPAGAAPVM | ATPGAFVAGVAVASAVVAAYEAGNADDDDPGC | 72854 | 72653 | - | 31 |
| 10 | MDFANGSAELAEVDVRDEGGYAAAASVPVL | ATTQAVVAVARAVAVAYG | 73139 | 72992 | - | 29 |

| | | | | | | |
|---|---|---|---|---|---|---|
| 11 | MSTVMDFANGSAELAEVDVRDEGGYAAAASVPVL | ATTQAVVAVARAVAVAYG | 73151 | 72992 | - | 31 |
| 12 | VLTTERTEGRKHMSTVMDFANGSAELAEVDVRDEGGYAAAASVPVL | ATTQAVVAVARAVAVAYG | 73187 | 72992 | - | 31 |
| 13 | MLTKPLISDAAPSGALCETDQGQITDRALPVM | ATPAAFAAGAAVVVVAYAAGDGLGDDEEDPWC | 73375 | 73180 | - | 31 |
| 14 | LLNGACSMRTATFLKW | ILPRWQGEERPSAPTDHGRRRARRTHHQITCPRRETC | 73530 | 73368 | - | 6 |
| 15 | VGGGREGDVMVPSAPGVGDE | AVTVPRALVAGGPAARRCASA | 73708 | 73582 | - | 8 |
| 16 | MRRSSTAAGPAWPGGLGRRSGRRWGFWVESAKREEGVG | GGREGDVMVPSAPGVGDEAVTVPRALVAGGPAARRCASA | 73816 | 73582 | - | 8 |
| 17 | VAGGACAGPAQQQVLHGLEDLVEG | RADAGAFGSSLRSVRKAWAAVARVT | 73830 | 73680 | - | 8 |
| 18 | VGGGGRRGMRRSSTAAGPAWPGGLGRRSGRRWGFWVESAKRE | EGVGGGREGDVMVPSAPGVGDEAVTVPRALVAGGPAARRCASA | 73840 | 73582 | - | 7 |
| 19 | LGRWAAVAGGACAGPAQQQVLHGLEDL | VEGRADAGAFGSSLRSVRKAWAAVARVT | 73848 | 73680 | - | 11 |
| 20 | VESRGYLGRWAAVAGGACAGPAQQQVLHGL | EDLVEGRADAGAFGSSLRSVRKAWAAVARVT | 73866 | 73680 | - | 9 |
| 21 | LVLRVESRGYLGRWAAVAGGACAGPAQQQVLH | GLEDLVEGRADAGAFGSSLRSVRKAWAAVARVT | 73878 | 73680 | - | 9 |
| 22 | VHRQVGPASGKSGVFGPVGGGGRRG...RRSSTAAGPAWPGGLGRRSGRRWGF | WVESAKREEGVGGGREGDVMVPSAP...VGDEAVTVPRALVAGGPAARRCASA | 73891 | 73582 | - | 6 |
| 23 | MTRTLVVHRQVGPASGKSGVFGPVG...RGMRRSSTAAGPAWPGGLGRRSGRR | WGFWVESAKREEGVGGGREGDVMVP...VGDEAVTVPRALVAGGPAARRCASA | 73909 | 73582 | - | 6 |
| 24 | VPGQRPVRPPPSAAAARRPGGPRALR | AARNRRHPGGAAAARPLPMSDGRRPTD | 78472 | 78310 | - | 6 |

### Results for WP_149823165.1 [Streptomyces sp. TRM68348] back to top

previous - next

#### Architecture

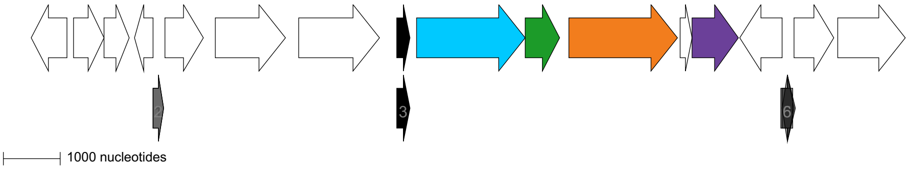

Link to nucleotide sequence

| Accession | start | end | direction | length (aa) | Pfam/HMM | name | description | E-value |
|---|---|---|---|---|---|---|---|---|
| WP_149823161.1 | 376101 | 375462 | - | 212 | PF00857 | Isochorismatase | Isochorismatase family | 1.20E-28 |
| WP_015662222.1 | 376209 | 376735 | + | 175 | INFERRED GENE | - | - | - |
| NP_631450.1 | 376753 | 377187 | + | 144 | INFERRED GENE | - | - | - |
| WP_149823162.1 | 377603 | 377279 | - | 107 | PF00196<br>TIGR03541<br>TIGR03020<br>PF08281 | GerE<br>TIGR03541<br>TIGR03020<br>Sigma70_r4_2 | Bacterial regulatory proteins, luxR family<br>reg_near_HchA: LuxR family transcriptional regulatory, chaperone HchA-associated<br>EpsA: transcriptional regulator EpsA<br>Sigma-70, region 4 | 1.30E-15<br>1.90E-13<br>7.10E-10<br>2.30E-05 |
| WP_149823163.1 | 377813 | 378503 | + | 229 | PF00072<br>PF00196<br>TIGR02875<br>TIGR03020<br>TIGR03541 | Response_reg<br>GerE<br>TIGR02875<br>TIGR03020<br>TIGR03541 | Response regulator receiver domain<br>Bacterial regulatory proteins, luxR family<br>spore_0_A: sporulation transcription factor Spo0A<br>EpsA: transcriptional regulator EpsA<br>reg_near_HchA: LuxR family transcriptional regulatory, chaperone HchA-associated | 8.90E-25<br>8.70E-19<br>2.80E-16<br>7.20E-14<br>2.20E-08 |
| WP_005474557.1 | 378715 | 379944 | + | 409 | INFERRED GENE | - | - | - |
| WP_149823164.1 | 380165 | 381596 | + | 476 | PF13450<br>TIGR02734<br>TIGR03467<br>TIGR00292<br>PF01266 | NAD_binding_8<br>TIGR02734<br>TIGR03467<br>TIGR00292<br>DAO | NAD(P)-binding Rossmann-like domain<br>crtI_fam: phytoene desaturase<br>HpnE: squalene-associated FAD-dependent desaturase<br>TIGR00292: thiazole biosynthesis enzyme<br>FAD dependent oxidoreductase | 6.80E-12<br>8.60E-12<br>8.60E-09<br>3.80E-08<br>7.40E-08 |
| WP_149823215.1 | 381887 | 382124 | + | 78 | LinA | LinA | LinA (linaridin precursor) | 3.30E-04 |
| WP_149823165.1 | 382237 | 384157 | + | 639 | LinH<br>LinG<br>LinE<br>PF12697<br>PF12146 | LinH<br>LinG<br>LinE<br>Abhydrolase_6<br>Hydrolase_4 | LinH (Fusion of LinE/LinG)<br>LinG Homolog<br>LinE Homolog<br>Alpha/beta hydrolase family<br>Serine aminopeptidase, S33 | 2.30E-245<br>1.40E-159<br>8.00E-93<br>1.50E-15<br>1.20E-09 |
| WP_149823166.1 | 384153 | 384774 | + | 206 | LinL | LinL | LinL | 2.70E-68 |
| WP_149823167.1 | 384935 | 386828 | + | 630 | TIGR02204<br>TIGR02203<br>TIGR00958<br>TIGR03796<br>TIGR01846 | TIGR02204<br>TIGR02203<br>TIGR00958<br>TIGR03796<br>TIGR01846 | MsbA_rel: ABC transporter, permease/ATP-binding protein<br>MsbA_lipidA: lipid A export permease/ATP-binding protein MsbA<br>3a01208: antigen peptide transporter 2<br>NHLM_micro_ABC1: NHLM bacteriocin system ABC transporter, peptidase/ATP-binding protein<br>type_I_sec_HlyB: type I secretion system ATPase | 6.90E-106<br>2.20E-101<br>4.50E-92<br>4.90E-83<br>9.40E-80 |
| WP_149823168.1 | 386897 | 387089 | + | 63 | NO PFAM MATCH | - | - | - |
| WP_149823169.1 | 387085 | 387904 | + | 272 | PF13649<br>PF08241<br>PF13489<br>PF08242<br>TIGR02469 | Methyltransf_25<br>Methyltransf_11<br>Methyltransf_23<br>Methyltransf_12<br>TIGR02469 | Methyltransferase domain<br>Methyltransferase domain<br>Methyltransferase domain<br>Methyltransferase domain<br>CbiT: precorrin-6Y C5,15-methyltransferase (decarboxylating), CbiT subunit | 2.20E-15<br>1.60E-12<br>3.90E-08<br>3.40E-06<br>1.20E-05 |
| WP_149823170.1 | 388670 | 387929 | - | 246 | PF07987 | DUF1775 | Domain of unkown function (DUF1775) | 8.60E-15 |

| | | | | | | | | |
|---|---|---|---|---|---|---|---|---|
| WP_149823171.1 | 388893 | 389586 | + | 230 | TIGR02154 | TIGR02154 | PhoB: phosphate regulon transcriptional regulatory protein PhoB | 1.90E-52 |
| | | | | | TIGR01387 | TIGR01387 | cztR_silR_copR: heavy metal response regulator | 7.40E-48 |
| | | | | | TIGR03787 | TIGR03787 | marine_sort_RR: proteobacterial dedicated sortase system response regulator | 1.70E-23 |
| | | | | | PF00072 | Response_reg | Response regulator receiver domain | 9.60E-23 |
| | | | | | PF00486 | Trans_reg_C | Transcriptional regulatory protein, C terminal | 1.30E-22 |
| WP_149823216.1 | 389659 | 390847 | + | 395 | TIGR02966 | TIGR02966 | phoR_proteo: phosphate regulon sensor kinase PhoR | 3.60E-15 |
| | | | | | PF00512 | HisKA | His Kinase A (phospho-acceptor) domain | 1.80E-13 |
| | | | | | TIGR01386 | TIGR01386 | cztS_silS_copS: heavy metal sensor kinase | 3.90E-13 |
| | | | | | PF00672 | HAMP | HAMP domain | 4.40E-07 |

| index | leader | core | start | end | dir | score |
|---|---|---|---|---|---|---|
| 1 | MVPALFERGSRTRDVVRLVRCDSAVTPGPPA | STIRPIPGRDHRPTGLGPSARLGPTIQNAPTAVPAA | 377601 | 377805 | + | 6 |
| 2 | LFERGSRTRDVVRLVRCDSAVTPGPPA | STIRPIPGRDHRPTGLGPSARLGPTIQNAPTAVPAA | 377613 | 377805 | + | 6 |
| 3 | MEIPVEGVFRLVARPTL | GTPRIARIARVADVAQAAGAVVAAG...ANALAAEANAQNAQLAAAAGGAPVA | 381887 | 382124 | + | 29 |
| 4 | MGAGHYGTPWCSATGRPSGSGRQSGGSLQESDRPGGKFPADQVIHITQ | PTPRLSTDTAQPAPTGSEHASRFRQSRRPDQCVDRR | 388655 | 388910 | + | 7 |
| 5 | VFSDGAAFGKRTPKRWFAPGVGSPRR | EVPGRSGDPHHTAHPEAQHRHRPTCSDRVGARLALPTVTKT | 388684 | 388888 | + | 7 |
| 6 | VLRPCRSRLGGVCAEPRGGLCDVDHLIGRELPAGAIRLLE | RTTALASASRRPPRR | 388856 | 388688 | - | 6 |

### Results for WP_155070885.1 [Streptomyces sp. TRM43335] back to top

previous - next

#### Architecture

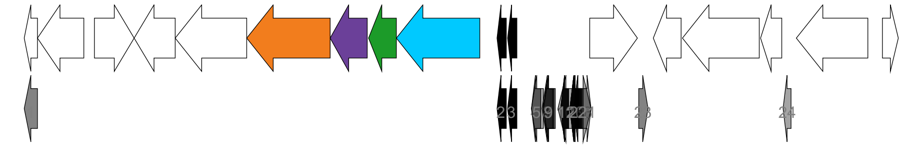

1000 nucleotides

Link to nucleotide sequence

| Accession | start | end | direction | length (aa) | Pfam/HMM | name | description | E-value |
|---|---|---|---|---|---|---|---|---|
| WP_155070879.1 | 2459884 | 2459602 | - | 93 | PF00550 | PP-binding | Phosphopantetheine attachment site | 9.10E-10 |
| | | | | | TIGR01688 | TIGR01688 | dltC: D-alanine--poly(phosphoribitol) ligase, subunit 2 | 5.00E-06 |
| WP_155073458.1 | 2460858 | 2459880 | - | 325 | TIGR00747 | TIGR00747 | fabH: 3-oxoacyl-[acyl-carrier-protein] synthase III | 5.40E-07 |
| WP_155070880.1 | 2461062 | 2461905 | + | 280 | PF01551 | Peptidase_M23 | Peptidase family M23 | 9.10E-16 |
| WP_155070881.1 | 2462741 | 2461889 | - | 283 | PF12680 | SnoaL_2 | SnoaL-like domain | 5.30E-05 |
| WP_155070882.1 | 2464222 | 2462737 | - | 494 | TIGR02734 | TIGR02734 | crtI_fam: phytoene desaturase | 9.70E-10 |
| | | | | | TIGR02733 | TIGR02733 | desat_CrtD: C-3',4' desaturase CrtD | 8.90E-09 |
| | | | | | TIGR00292 | TIGR00292 | TIGR00292: thiazole biosynthesis enzyme | 1.40E-08 |
| | | | | | TIGR03467 | TIGR03467 | HpnE: squalene-associated FAD-dependent desaturase | 2.50E-08 |
| | | | | | TIGR01317 | TIGR01317 | GOGAT_sm_gam: glutamate synthase, NADH/NADPH, small subunit | 1.00E-07 |
| WP_155070883.1 | 2465967 | 2464218 | - | 582 | TIGR02204 | TIGR02204 | MsbA_rel: ABC transporter, permease/ATP-binding protein | 8.50E-105 |
| | | | | | TIGR02203 | TIGR02203 | MsbA_lipidA: lipid A export permease/ATP-binding protein MsbA | 5.30E-98 |
| | | | | | TIGR03796 | TIGR03796 | NHLM_micro_ABC1: NHLM bacteriocin system ABC transporter, peptidase/ATP-binding protein | 5.50E-87 |
| | | | | | TIGR00958 | TIGR00958 | 3a01208: antigen peptide transporter 2 | 1.50E-83 |
| | | | | | TIGR01846 | TIGR01846 | type_I_sec_HlyB: type I secretion system ATPase | 2.60E-82 |
| WP_155070884.1 | 2466719 | 2465963 | - | 251 | PF13649 | Methyltransf_25 | Methyltransferase domain | 4.60E-13 |
| | | | | | PF08241 | Methyltransf_11 | Methyltransferase domain | 4.10E-09 |
| | | | | | PF13489 | Methyltransf_23 | Methyltransferase domain | 7.50E-08 |
| | | | | | PF08242 | Methyltransf_12 | Methyltransferase domain | 4.20E-07 |
| | | | | | PF13847 | Methyltransf_31 | Methyltransferase domain | 4.20E-06 |
| WP_162466045.1 | 2467348 | 2466769 | - | 192 | LinL | LinL | LinL | 6.10E-64 |
| WP_155070885.1 | 2469078 | 2467344 | - | 577 | LinH | LinH | LinH (Fusion of LinE/LinG) | 2.40E-221 |
| | | | | | LinE | LinE | LinE Homolog | 9.50E-101 |
| | | | | | LinG | LinG | LinG Homolog | 1.00E-95 |
| | | | | | PF12697 | Abhydrolase_6 | Alpha/beta hydrolase family | 9.40E-15 |
| | | | | | PF00561 | Abhydrolase_1 | alpha/beta hydrolase fold | 2.00E-12 |
| WP_155070886.1 | 2469618 | 2469423 | - | 64 | LinA | LinA | LinA (linaridin precursor) | 4.80E-10 |
| WP_155070887.1 | 2469849 | 2469657 | - | 63 | LinA | LinA | LinA (linaridin precursor) | 3.20E-15 |
| WP_155070888.1 | 2471359 | 2472346 | + | 328 | PF00248 | Aldo_ket_red | Aldo/keto reductase family | 1.10E-73 |
| | | | | | TIGR01293 | TIGR01293 | Kv_beta: voltage-dependent potassium channel beta subunit | 4.00E-39 |
| WP_155070889.1 | 2473249 | 2472667 | - | 193 | TIGR02258 | TIGR02258 | 2_5_ligase: 2'-5' RNA ligase | 2.40E-36 |
| | | | | | PF02834 | LigT_PEase | LigT like Phosphoesterase | 9.20E-24 |
| | | | | | PF13563 | 2_5_RNA_ligase2 | 2'-5' RNA ligase superfamily | 6.60E-17 |
| | | | | | PF10469 | AKAP7_NLS | AKAP7 2'5' RNA ligase-like domain | 5.30E-04 |
| WP_155070890.1 | 2474875 | 2473276 | - | 532 | TIGR00900 | TIGR00900 | 2A0121: H+ Antiporter protein | 5.00E-60 |
| | | | | | PF05977 | MFS_3 | Transmembrane secretion effector | 1.10E-56 |
| | | | | | PF07690 | MFS_1 | Major Facilitator Superfamily | 3.70E-33 |
| | | | | | TIGR00880 | TIGR00880 | 2_A_01_02: multidrug resistance protein | 1.20E-11 |
| | | | | | TIGR00893 | TIGR00893 | 2A0114: D-galactonate transporter | 7.30E-09 |
| WP_155070891.1 | 2475345 | 2474907 | - | 145 | PF12802 | MarR_2 | MarR family | 8.90E-15 |
| | | | | | PF01047 | MarR | MarR family | 1.80E-14 |
| | | | | | TIGR02337 | TIGR02337 | HpaR: homoprotocatechuate degradation operon regulator, HpaR | 2.50E-12 |
| | | | | | PF09339 | HTH_IclR | IclR helix-turn-helix domain | 5.80E-05 |
| | | | | | PF13412 | HTH_24 | Winged helix-turn-helix DNA-binding | 8.40E-05 |

| | | | | | | | | |
|---|---|---|---|---|---|---|---|---|
| WP_155070892.1 | 2477121 | 2475639 | - | 493 | PF00860<br>TIGR00815 | Xan_ur_permease<br>TIGR00815 | Permease family<br>sulP: sulfate permease | 4.40E-29<br>4.60E-04 |
| WP_155070893.1 | 2477414 | 2477744 | + | 109 | PF10745 | DUF2530 | Protein of unknown function (DUF2530) | 6.40E-07 |

| index | leader | core | start | end | dir | score |
|---|---|---|---|---|---|---|
| 1 | MNHGTEKTTEEITRFLTEALRRPIGPDDDYFALGLVDSLFALELVT | HVEQRFTLTVEVEDLDLDSFRTARRIAEFVHRKTGADTPTRTRAGDR | 2459884 | 2459602 | - | 7 |
| 2 | MNRIVESVAALTPEELTDETGGEFGAMMPVQ | ATPGLVAFGVGFAGGAGAAWVTVQAYEAGRTAG | 2469618 | 2469423 | - | 31 |
| 3 | MEQKISVFMNDQDLGPLEQGYQDDAPAPVL | ATPGAVATCWYVGGAVVGAGSVVGAYVTGRVVG | 2469849 | 2469657 | - | 29 |
| 4 | LSAVSQGIPLHGSLFRVESGRGWILL | VLPRSSPNWYFCRVVGVRSSGVHRFGCSQTTVPV | 2470326 | 2470143 | - | 7 |
| 5 | VSQFWTNGSRSAGRSRTDLSAVSQGIPLHGSLFRVESGRGWILL | VLPRSSPNWYFCRVVGVRSSGVHRFGCSQTTVPV | 2470380 | 2470143 | - | 6 |
| 6 | MGRRRPAAAPRSRVDSYAEPVRP | SVPVFPFPRATRATCGFLVVYSSGAHCLRSAPSRVTA | 2470569 | 2470386 | - | 6 |
| 7 | MPGCPACCRMGRRRPAAAPRSRVDSYAEPVRP | SVPVFPFPRATRATCGFLVVYSSGAHCLRSAPSRVTA | 2470596 | 2470386 | - | 7 |
| 8 | VRCSGLPDARVPRLLSYGPPKACGCAAESCRLVRRARPPICSRV | PVSAGHQGDLRFPCGVQLRRALFTLGPVPRYRLKRSVPILDQRQ | 2470618 | 2470351 | - | 6 |
| 9 | VIRRECGVPGFRMPGCPACCRMGRRRPAAAPRSRVDSYAEPVRP | SVPVFPFPRATRATCGFLVVYSSGAHCLRSAPSRVTA | 2470632 | 2470386 | - | 6 |
| 10 | VWCASMLLSLPDQQGQWKSSIPEAW | TYGNLPQGETCTRSNRTRGDPPRRQ | 2470903 | 2470750 | - | 9 |
| 11 | MCIVWCASMLLSLPDQQGQWKSSIPE | AWTYGNLPQGETCTRSNRTRGDPPRRQ | 2470912 | 2470750 | - | 8 |
| 12 | VFAWNNDRKESSVGWCLDRPCGPFKVEW | ITPLFFPGAGGPDVYSVVREHVAEP...GSSASTVKFPQRCDRLPEEEREILP | 2471033 | 2470694 | - | 6 |
| 13 | VADGLVFDLADGKRPAVPISDDPWRYKPTHAHTCPLRAGRRTAPPPP | DTPGPVTGLVPPTGAAGRGPVPSDI...ERSSVRRVTWPVLALSALQRVGWWP | 2471048 | 2471363 | + | 6 |
| 14 | MGTAGRFPSARSKTNPSATGGLDRCLPG | ITTGKNLRSAGVLIVRVVHSKWSG | 2471104 | 2470945 | - | 7 |
| 15 | LYRHGSSEMGTAGRFPSARSKTNPSATGGLDRCLPG | ITTGKNLRSAGVLIVRVVHSKWSG | 2471128 | 2470945 | - | 7 |
| 16 | VGLYRHGSSEMGTAGRFPSARSKTNPSATGGLDRCLPG | ITTGKNLRSAGVLIVRVVHSKWSG | 2471134 | 2470945 | - | 7 |
| 17 | VWACVGLYRHGSSEMGTAGRFPSARSKTNPSATGGLDRCLPG | ITTGKNLRSAGVLIVRVVHSKWSG | 2471146 | 2470945 | - | 7 |
| 18 | VRRPARSGHVWACVGLYRHGSSEMG...AGRFPSARSKTNPSATGGLDRCLPG | ITTGKNLRSAGVLIVRVVHSKWSG | 2471173 | 2470945 | - | 6 |
| 19 | VRCGARPGAGMCGRVWACIAMGRLKW | VQQDAFRPPGQKLTHRPLGDSIGVCLE | 2471178 | 2471016 | - | 7 |
| 20 | VSGGGGAVRRPARSGHVWACVGLYR...AGRFPSARSKTNPSATGGLDRCLPG | ITTGKNLRSAGVLIVRVVHSKWSG | 2471194 | 2470945 | - | 7 |
| 21 | VTGLVPPTGAAGRGPVPSDIRTETGG | AERSSVRRVTWPVLALSALQRVGWWP | 2471204 | 2471363 | + | 6 |
| 22 | VARGPSPAPGCRAEEVRCGARPGAGMCGRVWACI | AMGRLKWVQQDAFRPPGQKLTHRPLGDSIGVCLE | 2471223 | 2471016 | - | 6 |
| 23 | VRRTRARPGTGPARPDGTGRPRCRRCVRRLPQPTRAAT | STTASISTRAPLTMRVRVFGSTPQR | 2472363 | 2472555 | + | 7 |
| 24 | VPAVRRLSAFDSVGFVRTRSRGRRGQ | CVVRRTGAPVRVMKDTPPPPASHWSLD | 2475529 | 2475367 | - | 6 |

### Results for EHM28125.1 [Streptomyces sp. W007] back to top

previous - next

#### Architecture

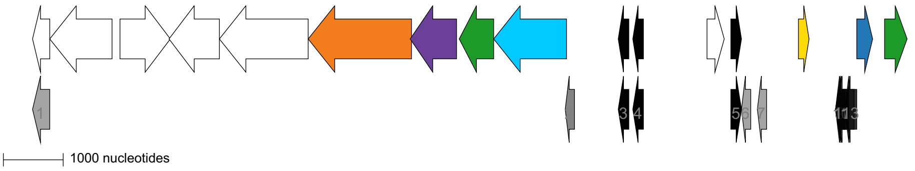

Link to nucleotide sequence

| Accession | start | end | direction | length (aa) | Pfam/HMM | name | description | E-value |
|---|---|---|---|---|---|---|---|---|
| EHM28117.1 | 12900 | 12618 | - | 93 | PF00550<br>TIGR01688 | PP-binding<br>TIGR01688 | Phosphopantetheine attachment site<br>dltC: D-alanine--poly(phosphoribitol) ligase, subunit 2 | 1.20E-08<br>4.20E-05 |
| EHM28118.1 | 13940 | 12896 | - | 347 | TIGR00747<br>PF08541 | TIGR00747<br>ACP_syn_III_C | fabH: 3-oxoacyl-[acyl-carrier-protein] synthase III<br>3-Oxoacyl-[acyl-carrier-protein (ACP)] synthase III C terminal | 7.40E-11<br>5.00E-04 |
| EHM28119.1 | 14086 | 14914 | + | 275 | PF01551 | Peptidase_M23 | Peptidase family M23 | 2.00E-16 |
| EHM28120.1 | 15750 | 14898 | - | 283 | PF12680<br>PF14534 | SnoaL_2<br>DUF4440 | SnoaL-like domain<br>Domain of unknown function (DUF4440) | 5.40E-06<br>3.90E-05 |
| EHM28121.1 | 17231 | 15746 | - | 494 | TIGR02734<br>TIGR00292<br>TIGR02733<br>TIGR03467<br>TIGR01317 | TIGR02734<br>TIGR00292<br>TIGR02733<br>TIGR03467<br>TIGR01317 | crtI_fam: phytoene desaturase<br>TIGR00292: thiazole biosynthesis enzyme<br>desat_CrtD: C-3',4' desaturase CrtD<br>HpnE: squalene-associated FAD-dependent desaturase<br>GOGAT_sm_gam: glutamate synthase, NADH/NADPH, small subunit | 4.40E-09<br>4.40E-08<br>8.90E-08<br>1.70E-07<br>1.80E-07 |
| EHM28122.1 | 18946 | 17227 | - | 572 | TIGR02204<br>TIGR02203<br>TIGR00958<br>TIGR01846<br>TIGR03796 | TIGR02204<br>TIGR02203<br>TIGR00958<br>TIGR01846<br>TIGR03796 | MsbA_rel: ABC transporter, permease/ATP-binding protein<br>MsbA_lipidA: lipid A export permease/ATP-binding protein MsbA<br>3a01208: antigen peptide transporter 2<br>type_I_sec_HlyB: type I secretion system ATPase<br>NHLM_micro_ABC1: NHLM bacteriocin system ABC transporter, peptidase/ATP-binding protein | 6.70E-109<br>9.50E-105<br>1.10E-91<br>2.50E-87<br>7.00E-85 |

| | | | | | | | | |
|---|---|---|---|---|---|---|---|---|
| EHM28123.1 | 19698 | 18942 | - | 251 | PF13649<br>PF08241<br>PF08242<br>PF13489<br>PF13847 | Methyltransf_25<br>Methyltransf_11<br>Methyltransf_12<br>Methyltransf_23<br>Methyltransf_31 | Methyltransferase domain<br>Methyltransferase domain<br>Methyltransferase domain<br>Methyltransferase domain<br>Methyltransferase domain | 2.50E-12<br>1.50E-09<br>3.30E-07<br>1.70E-06<br>3.10E-06 |
| EHM28124.1 | 20328 | 19749 | - | 192 | LinL | LinL | LinL | 4.10E-65 |
| EHM28125.1 | 21542 | 20324 | - | 405 | LinH<br>LinE<br>LinG<br>PF12697<br>PF00561 | LinH<br>LinE<br>LinG<br>Abhydrolase_6<br>Abhydrolase_1 | LinH (Fusion of LinE/LinG)<br>LinE Homolog<br>LinG Homolog<br>Alpha/beta hydrolase family<br>alpha/beta hydrolase fold | 6.60E-140<br>6.70E-105<br>3.60E-22<br>6.70E-14<br>2.20E-13 |
| EHM28126.1 | 22591 | 22411 | - | 59 | LinA | LinA | LinA (linaridin precursor) | 3.10E-10 |
| EHM28127.1 | 22839 | 22647 | - | 63 | LinA | LinA | LinA (linaridin precursor) | 2.20E-18 |
| EHM28128.1 | 23903 | 24182 | + | 92 | TIGR01288<br>TIGR01188 | TIGR01288<br>TIGR01188 | nodI: nodulation ABC transporter NodI<br>drrA: daunorubicin resistance ABC transporter, ATP-binding protein | 5.30E-05<br>2.60E-04 |
| EHM28129.1 | 24284 | 24461 | + | 58 | LinA | LinA | LinA (linaridin precursor) | 1.50E-08 |
| EHM28130.1 | 25419 | 25611 | + | 63 | LinG<br>LinH | LinG<br>LinH | LinG Homolog<br>LinH (Fusion of LinE/LinG) | 4.90E-21<br>1.60E-18 |
| EHM28131.1 | 26405 | 26663 | + | 85 | LinE<br>LinH | LinE<br>LinH | LinE Homolog<br>LinH (Fusion of LinE/LinG) | 1.00E-18<br>7.30E-11 |
| EHM28132.1 | 26869 | 27241 | + | 123 | LinL | LinL | LinL | 9.00E-32 |

| index | leader | core | start | end | dir | score |
|---|---|---|---|---|---|---|
| 1 | MSRGPAETNDEITRFLTDSLRREVGPDDDYFALGLVDSLFALELVT | YVEERFKLTVEVDDLDLDSFRTARRLTAFVHAKEAAAETASFHVGDS | 12900 | 12618 | - | 6 |
| 2 | VTAPPFPASCGPAPTDLSARTGLRS | ATPEPAKSRSTRSWLWNASSPSCTSLAGG | 21682 | 21517 | - | 7 |
| 3 | MDSVVAVTSQHLSDEAHGEFGVMMPVQ | ATPTLIAFGVGFAGGAGAAWVTVQAYEAGAND | 22591 | 22411 | - | 30 |
| 4 | MEHKIAGFMNTQDLGTLEQGYQDDDAAPVL | ATPAAVATCWYVGGAVAGAGAVVGAYVTGRVNG | 22839 | 22647 | - | 30 |
| 5 | MMLEKINQDTKMAKELAAGEAEGASAPIM | ATFTIVAAPVGVTIVGVGAAYLTGKIAGK | 24284 | 24461 | + | 28 |
| 6 | VATGAPGTAAFSAVP | GTLKDGRTAPSLGGGGPQVTRGRTTSRWRALTRGNPAR | 24624 | 24462 | - | 6 |
| 7 | LRPPGPRSALRARYSRLEISCVAAPS | RSAVPSGSAKADAEDLRIFDHIRIKAT | 24890 | 24728 | - | 6 |
| 8 | MWTPLEPSRCRRDRISSPGWVGS | MSTTPSIAGADTRDSVVRLAVVAT | 26194 | 26050 | - | 8 |
| 9 | MSRPPPAREVSGPSFSAMWTPLEPSRCRRDRI | SSPGWVGSMSTTPSIAGADTRDSVVRLAVVAT | 26245 | 26050 | - | 8 |
| 10 | LSMSRPPPAREVSGPSFSAMWTPLEPSRCRRDR | ISSPGWVGSMSTTPSIAGADTRDSVVRLAVVAT | 26251 | 26050 | - | 8 |
| 11 | MFFLSMSRPPPAREVSGPSFSAMWTPLEPSRCRR | DRISSPGWVGSMSTTPSIAGADTRDSVVRLAVVAT | 26260 | 26050 | - | 8 |
| 12 | LPPLSTGGQPGADGRALQAPKFLVRA | TRTERVHVLLVHEQTATGQGGKRPQF | 26357 | 26198 | - | 7 |
| 13 | VQLAPTVQHEYVCLPPLSTGGQPGADGRALQA | PKFLVRATRTERVHVLLVHEQTATGQGGKRPQF | 26396 | 26198 | - | 9 |

### Results for EHM28131.1 [Streptomyces sp. W007] back to top

previous - next

#### Architecture

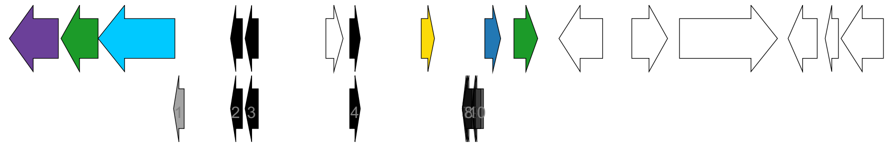

1000 nucleotides

Link to nucleotide sequence

| Accession | start | end | direction | length (aa) | Pfam/HMM | name | description | E-value |
|---|---|---|---|---|---|---|---|---|
| EHM28123.1 | 19698 | 18942 | - | 251 | PF13649<br>PF08241<br>PF08242<br>PF13489<br>PF13847 | Methyltransf_25<br>Methyltransf_11<br>Methyltransf_12<br>Methyltransf_23<br>Methyltransf_31 | Methyltransferase domain<br>Methyltransferase domain<br>Methyltransferase domain<br>Methyltransferase domain<br>Methyltransferase domain | 2.50E-12<br>1.50E-09<br>3.30E-07<br>1.70E-06<br>3.10E-06 |
| EHM28124.1 | 20328 | 19749 | - | 192 | LinL | LinL | LinL | 4.10E-65 |
| EHM28125.1 | 21542 | 20324 | - | 405 | LinH<br>LinE<br>LinG<br>PF12697<br>PF00561 | LinH<br>LinE<br>LinG<br>Abhydrolase_6<br>Abhydrolase_1 | LinH (Fusion of LinE/LinG)<br>LinE Homolog<br>LinG Homolog<br>Alpha/beta hydrolase family<br>alpha/beta hydrolase fold | 6.60E-140<br>6.70E-105<br>3.60E-22<br>6.70E-14<br>2.20E-13 |
| EHM28126.1 | 22591 | 22411 | - | 59 | LinA | LinA | LinA (linaridin precursor) | 3.10E-10 |
| EHM28127.1 | 22839 | 22647 | - | 63 | LinA | LinA | LinA (linaridin precursor) | 2.20E-18 |
| EHM28128.1 | 23903 | 24182 | + | 92 | TIGR01288<br>TIGR01188 | TIGR01288<br>TIGR01188 | nodI: nodulation ABC transporter NodI<br>drrA: daunorubicin resistance ABC transporter, ATP-binding protein | 5.30E-05<br>2.60E-04 |
| EHM28129.1 | 24284 | 24461 | + | 58 | LinA | LinA | LinA (linaridin precursor) | 1.50E-08 |

| | | | | | | | | |
|---|---|---|---|---|---|---|---|---|
| EHM28130.1 | 25419 | 25611 | + | 63 | LinG<br>LinH | LinG<br>LinH | LinG Homolog<br>LinH (Fusion of LinE/LinG) | 4.90E-21<br>1.60E-18 |
| EHM28131.1 | 26405 | 26663 | + | 85 | LinE<br>LinH | LinE<br>LinH | LinE Homolog<br>LinH (Fusion of LinE/LinG) | 1.00E-18<br>7.30E-11 |
| EHM28132.1 | 26869 | 27241 | + | 123 | LinL | LinL | LinL | 9.00E-32 |
| EHM28133.1 | 28267 | 27586 | - | 226 | PF13359<br>PF01609<br>PF13613 | DDE_Tnp_4<br>DDE_Tnp_1<br>HTH_Tnp_4 | DDE superfamily endonuclease<br>Transposase DDE domain<br>Helix-turn-helix of DDE superfamily endonuclease | 5.70E-22<br>5.50E-10<br>1.20E-09 |
| EHM28134.1 | 28732 | 29287 | + | 184 | PF05719 | GPP34 | Golgi phosphoprotein 3 (GPP34) | 2.60E-40 |
| EHM28135.1 | 29469 | 31023 | + | 517 | PF13701<br>PF01609 | DDE_Tnp_1_4<br>DDE_Tnp_1 | Transposase DDE domain group 1<br>Transposase DDE domain | 1.20E-32<br>1.40E-04 |
| EHM28136.1 | 31648 | 31177 | - | 156 | NO PFAM MATCH | - | - | - |
| EHM28137.1 | 31982 | 31763 | - | 72 | NO PFAM MATCH | - | - | - |
| EHM28138.1 | 32694 | 32013 | - | 226 | TIGR01723 | TIGR01723 | hmd_TIGR: 5,10-methenyltetrahydromethanopterin hydrogenase | 9.00E-04 |

| index | leader | core | start | end | dir | score |
|---|---|---|---|---|---|---|
| 1 | VTAPPFPASCGPAPTDLSARTGLRS | ATPEPAKSRSTRSWLWNASSPSCTSLAGG | 21682 | 21517 | - | 6 |
| 2 | MDSVVAVTSQHLSDEAHGEFGVMMPVQ | ATPTLIAFGVGFAGGAGAAWVTVQAYEAGAND | 22591 | 22411 | - | 29 |
| 3 | MEHKIAGFMNTQDLGTLEQGYQDDDAAPVL | ATPAAVATCWYVGGAVAGAGAVVGAYVTGRVNG | 22839 | 22647 | - | 29 |
| 4 | MMLEKINQDTKMAKELAAGEAEGASAPIM | ATFTIVAAPVGVTIVGVGAAYLTGKIAGK | 24284 | 24461 | + | 27 |
| 5 | MWTPLEPSRCRRDRISSPGWVGS | MSTTPSIAGADTRDSVVRLAVVAT | 26194 | 26050 | - | 7 |
| 6 | MSRPPPAREVSGPSFSAMWTPLEPSRCRRDRI | SSPGWVGSMSTTPSIAGADTRDSVVRLAVVAT | 26245 | 26050 | - | 7 |
| 7 | LSMSRPPPAREVSGPSFSAMWTPLEPSRCRRDR | ISSPGWVGSMSTTPSIAGADTRDSVVRLAVVAT | 26251 | 26050 | - | 7 |
| 8 | MFFLSMSRPPPAREVSGPSFSAMWTPLEPSRCRR | DRISSPGWVGSMSTTPSIAGADTRDSVVRLAVVAT | 26260 | 26050 | - | 7 |
| 9 | LPPLSTGGQPGADGRALQAPKFLVRA | TRTERVHVLLVHEQTATGQGGKRPQF | 26357 | 26198 | - | 6 |
| 10 | VQLAPTVQHEYVCLPPLSTGGQPGADGRALQA | PKFLVRATRTERVHVLLVHEQTATGQGGKRPQF | 26396 | 26198 | - | 8 |

### Results for WP_050987754.1 [Streptomyces sp. W007] back to top

previous - next

#### Architecture

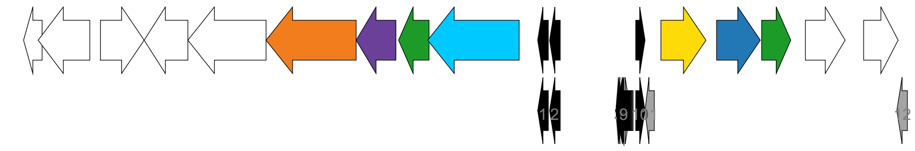

1000 nucleotides

Link to nucleotide sequence

| Accession | start | end | direction | length (aa) | Pfam/HMM | name | description | E-value |
|---|---|---|---|---|---|---|---|---|
| WP_106431539.1 | 12954 | 12618 | - | 111 | PF00550<br>TIGR01688 | PP-binding<br>TIGR01688 | Phosphopantetheine attachment site<br>dltC: D-alanine--poly(phosphoribitol) ligase, subunit 2 | 1.90E-08<br>6.40E-05 |
| WP_032792111.1 | 13874 | 12896 | - | 325 | TIGR00747<br>PF08541 | TIGR00747<br>ACP_syn_III_C | fabH: 3-oxoacyl-[acyl-carrier-protein] synthase III<br>3-Oxoacyl-[acyl-carrier-protein (ACP)] synthase III C terminal | 9.50E-09<br>4.40E-04 |
| WP_007452022.1 | 14086 | 14914 | + | 275 | PF01551 | Peptidase_M23 | Peptidase family M23 | 2.00E-16 |
| WP_007452024.1 | 15750 | 14898 | - | 283 | PF12680<br>PF14534 | SnoaL_2<br>DUF4440 | SnoaL-like domain<br>Domain of unknown function (DUF4440) | 5.40E-06<br>3.90E-05 |
| WP_007452027.1 | 17231 | 15746 | - | 494 | TIGR02734<br>TIGR00292<br>TIGR02733<br>TIGR03467<br>TIGR01317 | TIGR02734<br>TIGR00292<br>TIGR02733<br>TIGR03467<br>TIGR01317 | crtI_fam: phytoene desaturase<br>TIGR00292: thiazole biosynthesis enzyme<br>desat_CrtD: C-3',4' desaturase CrtD<br>HpnE: squalene-associated FAD-dependent desaturase<br>GOGAT_sm_gam: glutamate synthase, NADH/NADPH, small subunit | 4.40E-09<br>4.40E-08<br>8.90E-08<br>1.70E-07<br>1.80E-07 |
| WP_007452028.1 | 18946 | 17227 | - | 572 | TIGR02204<br>TIGR02203<br>TIGR00958<br>TIGR01846<br>TIGR03796 | TIGR02204<br>TIGR02203<br>TIGR00958<br>TIGR01846<br>TIGR03796 | MsbA_rel: ABC transporter, permease/ATP-binding protein<br>MsbA_lipidA: lipid A export permease/ATP-binding protein MsbA<br>3a01208: antigen peptide transporter 2<br>type_I_sec_HlyB: type I secretion system ATPase<br>NHLM_micro_ABC1: NHLM bacteriocin system ABC transporter, peptidase/ATP-binding protein | 6.70E-109<br>9.50E-105<br>1.10E-91<br>2.50E-87<br>7.00E-85 |
| WP_007452030.1 | 19698 | 18942 | - | 251 | PF13649<br>PF08241<br>PF08242<br>PF13489<br>PF13847 | Methyltransf_25<br>Methyltransf_11<br>Methyltransf_12<br>Methyltransf_23<br>Methyltransf_31 | Methyltransferase domain<br>Methyltransferase domain<br>Methyltransferase domain<br>Methyltransferase domain<br>Methyltransferase domain | 2.50E-12<br>1.50E-09<br>3.30E-07<br>1.70E-06<br>3.10E-06 |
| WP_007452032.1 | 20328 | 19749 | - | 192 | LinL | LinL | LinL | 4.10E-65 |
| WP_050987754.1 | 22055 | 20324 | - | 576 | LinH<br>LinE<br>LinG<br>PF12697<br>PF00561 | LinH<br>LinE<br>LinG<br>Abhydrolase_6<br>Abhydrolase_1 | LinH (Fusion of LinE/LinG)<br>LinE Homolog<br>LinG Homolog<br>Alpha/beta hydrolase family<br>alpha/beta hydrolase fold | 1.70E-218<br>1.60E-104<br>3.80E-90<br>1.70E-13<br>5.40E-13 |
| WP_032792089.1 | 22603 | 22411 | - | 63 | LinA | LinA | LinA (linaridin precursor) | 6.30E-12 |

| | | | | | | | | |
|---|---|---|---|---|---|---|---|---|
| WP_007452038.1 | 22839 | 22647 | - | 63 | LinA | LinA | LinA (linaridin precursor) | 2.20E-18 |
| WP_007452042.1 | 24284 | 24461 | + | 58 | LinA | LinA | LinA (linaridin precursor) | 1.50E-08 |
| WP_032792091.1 | 24747 | 25611 | + | 287 | LinG<br>LinH | LinG<br>LinH | LinG Homolog<br>LinH (Fusion of LinE/LinG) | 2.70E-100<br>4.00E-99 |
| WP_158000644.1 | 25826 | 26663 | + | 278 | LinE<br>LinH<br>PF12697<br>PF00975<br>TIGR03611 | LinE<br>LinH<br>Abhydrolase_6<br>Thioesterase<br>TIGR03611 | LinE Homolog<br>LinH (Fusion of LinE/LinG)<br>Alpha/beta hydrolase family<br>Thioesterase domain<br>RutD: pyrimidine utilization protein D | 2.00E-85<br>2.40E-58<br>1.10E-14<br>9.80E-08<br>3.00E-07 |
| WP_050987756.1 | 26683 | 27241 | + | 185 | LinL | LinL | LinL | 3.20E-55 |
| WP_008747277.1 | 27499 | 28267 | + | 256 | INFERRED GENE | - | - | - |
| WP_032792094.1 | 28624 | 29287 | + | 220 | PF05719 | GPP34 | Golgi phosphoprotein 3 (GPP34) | 8.60E-49 |

| index | leader | core | start | end | dir | score |
|---|---|---|---|---|---|---|
| 1 | MSRIVDSVVAVTSQHLSDEAHGEFGVMMPVQ | ATPTLIAFGVGFAGGAGAAWVTVQAYEAGAND | 22603 | 22411 | - | 30 |
| 2 | MEHKIAGFMNTQDLGTLEQGYQDDDAAPVL | ATPAAVATCWYVGGAVAGAGAVVGAYVTGRVNG | 22839 | 22647 | - | 30 |
| 3 | MTSARCVSLASL | RTPTTGRGRVVVATSIGTPSSIVK | 24004 | 23893 | - | 8 |
| 4 | VQNLSGNEKPLGLAIAPLSRLEILFLDEP | AAGQGARRHADSSARNVDTHPIAEPALTT | 24005 | 24182 | + | 6 |
| 5 | MARPRGFSFPDRFCTMTSARCVSLASL | RTPTTGRGRVVVATSIGTPSSIVK | 24049 | 23893 | - | 10 |
| 6 | VSTFRAEESACRRAPWPAAGSSRKR...RPRGFSFPDRFCTMTSARCVSLASL | RTPTTGRGRVVVATSIGTPSSIVK | 24148 | 23893 | - | 9 |
| 7 | VSAGSAIGCVSTFRAEESACRRAPW...RPRGFSFPDRFCTMTSARCVSLASL | RTPTTGRGRVVVATSIGTPSSIVK | 24175 | 23893 | - | 9 |
| 8 | VNNCIRTRAEPQVVSAGSAIGCVST...RPRGFSFPDRFCTMTSARCVSLASL | RTPTTGRGRVVVATSIGTPSSIVK | 24214 | 23893 | - | 8 |
| 9 | LFTAIAVNNCIRTRAEPQVVSAGSA...RPRGFSFPDRFCTMTSARCVSLASL | RTPTTGRGRVVVATSIGTPSSIVK | 24232 | 23893 | - | 8 |
| 10 | MMLEKINQDTKMAKELAAGEAEGASAPIM | ATFTIVAAPVGVTIVGVGAAYLTGKIAGK | 24284 | 24461 | + | 28 |
| 11 | VATGAPGTAAFSAVP | GTLKDGRTAPSLGGGGPQVTRGRTTSRWRALTRGNPAR | 24624 | 24462 | - | 6 |
| 12 | VCAGGWRVTRSGEAPDPLGSRSFATTRLLMPVRMQPALIGPAAPGR | VAPARSAPGAGAVSCRRWLM | 29462 | 29261 | - | 6 |

### Results for WP_158000644.1 [Streptomyces sp. W007] back to top

previous - next

#### Architecture

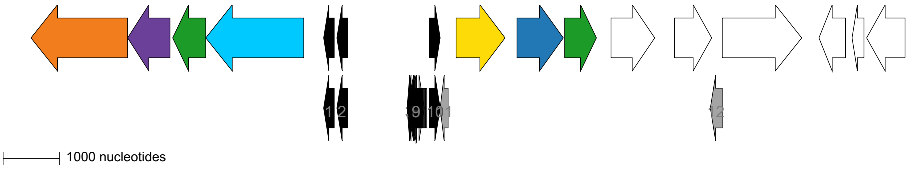

Link to nucleotide sequence

| Accession | start | end | direction | length (aa) | Pfam/HMM | name | description | E-value |
|---|---|---|---|---|---|---|---|---|
| WP_007452028.1 | 18946 | 17227 | - | 572 | TIGR02204<br>TIGR02203<br>TIGR00958<br>TIGR01846<br>TIGR03796 | TIGR02204<br>TIGR02203<br>TIGR00958<br>TIGR01846<br>TIGR03796 | MsbA_rel: ABC transporter, permease/ATP-binding protein<br>MsbA_lipidA: lipid A export permease/ATP-binding protein MsbA<br>3a01208: antigen peptide transporter 2<br>type_I_sec_HlyB: type I secretion system ATPase<br>NHLM_micro_ABC1: NHLM bacteriocin system ABC transporter, peptidase/ATP-binding protein | 6.70E-109<br>9.50E-105<br>1.10E-91<br>2.50E-87<br>7.00E-85 |
| WP_007452030.1 | 19698 | 18942 | - | 251 | PF13649<br>PF08241<br>PF08242<br>PF13489<br>PF13847 | Methyltransf_25<br>Methyltransf_11<br>Methyltransf_12<br>Methyltransf_23<br>Methyltransf_31 | Methyltransferase domain<br>Methyltransferase domain<br>Methyltransferase domain<br>Methyltransferase domain<br>Methyltransferase domain | 2.50E-12<br>1.50E-09<br>3.30E-07<br>1.70E-06<br>3.10E-06 |
| WP_007452032.1 | 20328 | 19749 | - | 192 | LinL | LinL | LinL | 4.10E-65 |
| WP_050987754.1 | 22055 | 20324 | - | 576 | LinH<br>LinE<br>LinG<br>PF12697<br>PF00561 | LinH<br>LinE<br>LinG<br>Abhydrolase_6<br>Abhydrolase_1 | LinH (Fusion of LinE/LinG)<br>LinE Homolog<br>LinG Homolog<br>Alpha/beta hydrolase family<br>alpha/beta hydrolase fold | 1.70E-218<br>1.60E-104<br>3.80E-90<br>1.70E-13<br>5.40E-13 |
| WP_032792089.1 | 22603 | 22411 | - | 63 | LinA | LinA | LinA (linaridin precursor) | 6.30E-12 |
| WP_007452038.1 | 22839 | 22647 | - | 63 | LinA | LinA | LinA (linaridin precursor) | 2.20E-18 |
| WP_007452042.1 | 24284 | 24461 | + | 58 | LinA | LinA | LinA (linaridin precursor) | 1.50E-08 |
| WP_032792091.1 | 24747 | 25611 | + | 287 | LinG<br>LinH | LinG<br>LinH | LinG Homolog<br>LinH (Fusion of LinE/LinG) | 2.70E-100<br>4.00E-99 |
| WP_158000644.1 | 25826 | 26663 | + | 278 | LinE<br>LinH<br>PF12697<br>PF00975<br>TIGR03611 | LinE<br>LinH<br>Abhydrolase_6<br>Thioesterase<br>TIGR03611 | LinE Homolog<br>LinH (Fusion of LinE/LinG)<br>Alpha/beta hydrolase family<br>Thioesterase domain<br>RutD: pyrimidine utilization protein D | 2.00E-85<br>2.40E-58<br>1.10E-14<br>9.80E-08<br>3.00E-07 |
| WP_050987756.1 | 26683 | 27241 | + | 185 | LinL | LinL | LinL | 3.20E-55 |

| | | | | | | | | |
|---|---|---|---|---|---|---|---|---|
| WP_008747277.1 | 27499 | 28267 | + | 256 | INFERRED GENE | - | - | - |
| WP_032792094.1 | 28624 | 29287 | + | 220 | PF05719 | GPP34 | Golgi phosphoprotein 3 (GPP34) | 8.60E-49 |
| WP_019031128.1 | 29469 | 30869 | + | 466 | INFERRED GENE | - | - | - |
| WP_032792098.1 | 31639 | 31177 | - | 153 | NO PFAM MATCH | - | - | - |
| WP_007452053.1 | 31982 | 31763 | - | 72 | NO PFAM MATCH | - | - | - |
| WP_007452054.1 | 32694 | 32013 | - | 226 | TIGR01723 | TIGR01723 | hmd_TIGR: 5,10-methenyltetrahydromethanopterin hydrogenase | 9.00E-04 |

| index | leader | core | start | end | dir | score |
|---|---|---|---|---|---|---|
| 1 | MSRIVDSVVAVTSQHLSDEAHGEFGVMMPVQ | ATPTLIAFGVGFAGGAGAAWVTVQAYEAGAND | 22603 | 22411 | - | 30 |
| 2 | MEHKIAGFMNTQDLGTLEQGYQDDDAAPVL | ATPAAVATCWYVGGAVAGAGAVVGAYVTGRVNG | 22839 | 22647 | - | 30 |
| 3 | MTSARCVSLASL | RTPTTGRGRVVVATSIGTPSSIVK | 24004 | 23893 | - | 8 |
| 4 | VQNLSGNEKPLGLAIAPLSRLEILFLDEP | AAGQGARRHADSSARNVDTHPIAEPALTT | 24005 | 24182 | + | 6 |
| 5 | MARPRGFSFPDRFCTMTSARCVSLASL | RTPTTGRGRVVVATSIGTPSSIVK | 24049 | 23893 | - | 10 |
| 6 | VSTFRAEESACRRAPWPAAGSSRKR...RPRGFSFPDRFCTMTSARCVSLASL | RTPTTGRGRVVVATSIGTPSSIVK | 24148 | 23893 | - | 9 |
| 7 | VSAGSAIGCVSTFRAEESACRRAPW...RPRGFSFPDRFCTMTSARCVSLASL | RTPTTGRGRVVVATSIGTPSSIVK | 24175 | 23893 | - | 9 |
| 8 | VNNCIRTRAEPQVVSAGSAIGCVST...RPRGFSFPDRFCTMTSARCVSLASL | RTPTTGRGRVVVATSIGTPSSIVK | 24214 | 23893 | - | 8 |
| 9 | LFTAIAVNNCIRTRAEPQVVSAGSA...RPRGFSFPDRFCTMTSARCVSLASL | RTPTTGRGRVVVATSIGTPSSIVK | 24232 | 23893 | - | 8 |
| 10 | MMLEKINQDTKMAKELAAGEAEGASAPIM | ATFTIVAAPVGVTIVGVGAAYLTGKIAGK | 24284 | 24461 | + | 28 |
| 11 | VATGAPGTAAFSAVP | GTLKDGRTAPSLGGGGPQVTRGRTTSRWRALTRGNPAR | 24624 | 24462 | - | 6 |
| 12 | VCAGGWRVTRSGEAPDPLGSRSFATTRLLMPVRMQPALIGPAAPGR | VAPARSAPGAGAVSCRRWLM | 29462 | 29261 | - | 6 |

### Results for WP_164721115.1 [Streptomyces sp. W1SF4] back to top

previous - next

#### Architecture

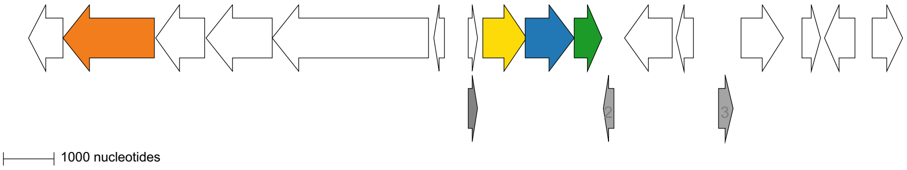

Link to nucleotide sequence

| Accession | start | end | direction | length (aa) | Pfam/HMM | name | description | E-value |
|---|---|---|---|---|---|---|---|---|
| WP_125824642.1 | 386071 | 385381 | - | 229 | PF13302<br>PF00583<br>PF13523 | Acetyltransf_3<br>Acetyltransf_1<br>Acetyltransf_8 | Acetyltransferase (GNAT) domain<br>Acetyltransferase (GNAT) family<br>Acetyltransferase (GNAT) domain | 4.70E-22<br>2.20E-06<br>1.90E-04 |
| WP_125824643.1 | 387873 | 386058 | - | 604 | TIGR02204<br>TIGR02203<br>TIGR01846<br>TIGR03796<br>TIGR00958 | TIGR02204<br>TIGR02203<br>TIGR01846<br>TIGR03796<br>TIGR00958 | MsbA_rel: ABC transporter, permease/ATP-binding protein<br>MsbA_lipidA: lipid A export permease/ATP-binding protein MsbA<br>type_I_sec_HlyB: type I secretion system ATPase<br>NHLM_micro_ABC1: NHLM bacteriocin system ABC transporter, peptidase/ATP-binding protein<br>3a01208: antigen peptide transporter 2 | 2.40E-112<br>1.60E-111<br>6.40E-103<br>2.00E-97<br>1.00E-96 |
| WP_125824644.1 | 388874 | 387893 | - | 326 | TIGR03891<br>PF14028 | TIGR03891<br>Lant_dehydr_C | thiopep_ocin: thiopeptide-type bacteriocin biosynthesis domain<br>Lantibiotic biosynthesis dehydratase C-term | 8.10E-25<br>1.20E-12 |
| WP_125824645.1 | 390210 | 388896 | - | 437 | PF05147<br>TIGR03897 | LANC_like<br>TIGR03897 | Lanthionine synthetase C-like protein<br>lanti_2_LanM: type 2 lantibiotic biosynthesis protein LanM | 6.20E-42<br>1.10E-05 |
| WP_125824646.1 | 393290 | 390206 | - | 1027 | PF04738<br>TIGR03891<br>PF14028 | Lant_dehydr_N<br>TIGR03891<br>Lant_dehydr_C | Lantibiotic dehydratase, N terminus<br>thiopep_ocin: thiopeptide-type bacteriocin biosynthesis domain<br>Lantibiotic biosynthesis dehydratase C-term | 4.70E-155<br>1.90E-38<br>1.30E-16 |
| WP_125594456.1 | 393594 | 393393 | - | 66 | NO PFAM MATCH | - | - | - |
| WP_125594457.1 | 394069 | 394258 | + | 62 | NO PFAM MATCH | - | - | - |
| WP_164721114.1 | 394349 | 395216 | + | 288 | LinG<br>LinH | LinG<br>LinH | LinG Homolog<br>LinH (Fusion of LinE/LinG) | 1.80E-86<br>4.10E-83 |
| WP_164721115.1 | 395205 | 396165 | + | 319 | LinE<br>LinH<br>PF12146<br>PF00561 | LinE<br>LinH<br>Hydrolase_4<br>Abhydrolase_1 | LinE Homolog<br>LinH (Fusion of LinE/LinG)<br>Serine aminopeptidase, S33<br>alpha/beta hydrolase fold | 2.80E-65<br>4.20E-62<br>4.00E-11<br>2.10E-05 |
| WP_125824648.1 | 396161 | 396719 | + | 185 | LinL | LinL | LinL | 3.30E-56 |
| WP_125824753.1 | 398102 | 397160 | - | 313 | TIGR04187<br>-<br>-<br>-<br>TIGR04185 | TIGR04187<br>Subclass_4<br>Subclass_5<br>Subclass_6<br>TIGR04185 | GRASP_SAV_5884: ATP-grasp ribosomal peptide maturase, SAV_5884 family<br>-<br>-<br>-<br>ATPgraspMvdC: ATP-grasp ribosomal peptide maturase, MvdC family | 3.10E-45<br>2.50E-41<br>3.30E-36<br>3.20E-35<br>4.00E-32 |
| WP_125824649.1 | 398512 | 398185 | - | 108 | NO PFAM MATCH | - | - | - |
| WP_125824754.1 | 399451 | 400306 | + | 284 | NO PFAM MATCH | - | - | - |

| | | | | | | | | |
|---|---|---|---|---|---|---|---|---|
| WP_125824755.1 | 400661 | 401027 | + | 121 | PF01471<br>TIGR02869 | PG_binding_1<br>TIGR02869 | Putative peptidoglycan binding domain<br>spore_SleB: spore cortex-lytic enzyme | 2.90E-15<br>2.90E-12 |
| WP_125824650.1 | 401721 | 401112 | - | 202 | PF07719 | TPR_2 | Tetratricopeptide repeat | 5.00E-05 |
| WP_164721116.1 | 402048 | 402642 | + | 197 | PF01548 | DEDD_Tnp_IS110 | Transposase | 1.30E-37 |

| index | leader | core | start | end | dir | score |
|---|---|---|---|---|---|---|
| 1 | MDRSRILAIAGGVLTVGVLAAGPAAAAERTE | QTEGASYAVQASGDRSAAGGDLVVNKFPCCV | 394069 | 394258 | + | 7 |
| 2 | LCDGQRQCAHAHPCQPGHDDDPGGRQDDDPPPH | GTPHGSTMTPRTAAPRRNRRNRAARRARGTEGVVETSPA | 396956 | 396737 | - | 6 |
| 3 | VIDTDAADESGTTRNAAALGCTPVS...RATVFNCLHEWHTDNTWRQQAGPET | KTLKPGGGSVAPHGATMWTSM | 399015 | 399315 | + | 6 |

### Results for AZM48683.1 [Streptomyces sp. WAC 06738] back to top

previous - next

#### Architecture

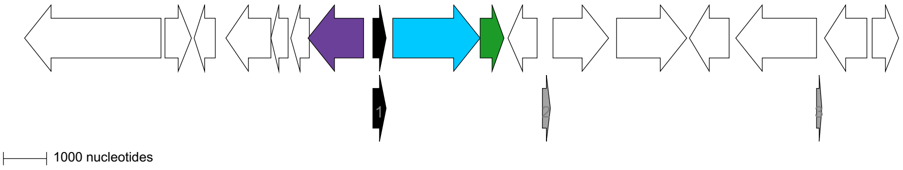

Link to nucleotide sequence

| Accession | start | end | direction | length (aa) | Pfam/HMM | name | description | E-value |
|---|---|---|---|---|---|---|---|---|
| AZM48676.1 | 5723413 | 5720254 | - | 1052 | PF03704<br>PF13424<br>PF00486<br>PF13191<br>PF13401 | BTAD<br>TPR_12<br>Trans_reg_C<br>AAA_16<br>AAA_22 | Bacterial transcriptional activator domain<br>Tetratricopeptide repeat<br>Transcriptional regulatory protein, C terminal<br>AAA ATPase domain<br>AAA domain | 2.60E-33<br>1.40E-13<br>6.70E-08<br>1.70E-05<br>3.00E-04 |
| AZM48677.1 | 5723520 | 5724126 | + | 201 | NO PFAM MATCH | - | - | - |
| AZM48678.1 | 5724682 | 5724169 | - | 170 | PF01814<br>TIGR03652 | Hemerythrin<br>TIGR03652 | Hemerythrin HHE cation binding domain<br>FeS_repair_RIC: iron-sulfur cluster repair di-iron protein | 8.70E-20<br>4.70E-04 |
| AZM48679.1 | 5725961 | 5724920 | - | 346 | TIGR03086<br>PF08327<br>TIGR03083<br>PF11716<br>PF10604 | TIGR03086<br>AHSA1<br>TIGR03083<br>MDMPI_N<br>Polyketide_cyc2 | TIGR03086: TIGR03086 family protein<br>Activator of Hsp90 ATPase homolog 1-like protein<br>TIGR03083: uncharacterized Actinobacterial protein TIGR03083<br>Mycothiol maleylpyruvate isomerase N-terminal domain<br>Polyketide cyclase / dehydrase and lipid transport | 8.80E-32<br>1.60E-24<br>1.60E-15<br>1.70E-07<br>2.10E-06 |
| AZM48680.1 | 5726347 | 5725957 | - | 129 | PF12840<br>PF01022<br>PF12802<br>TIGR01884<br>TIGR02702 | HTH_20<br>HTH_5<br>MarR_2<br>TIGR01884<br>TIGR02702 | Helix-turn-helix domain<br>Bacterial regulatory protein, arsR family<br>MarR family<br>cas_HTH: CRISPR locus-related DNA-binding protein<br>SufR_cyano: iron-sulfur cluster biosynthesis transcriptional regulator SufR | 1.50E-16<br>5.60E-10<br>4.20E-08<br>6.60E-07<br>1.40E-06 |
| AZM48681.1 | 5726841 | 5726430 | - | 136 | PF04134 | DUF393 | Protein of unknown function, DUF393 | 2.00E-11 |
| AZM50978.1 | 5728111 | 5726818 | - | 430 | PF00891<br>TIGR02716<br>PF13649<br>PF16864<br>PF08242 | Methyltransf_2<br>TIGR02716<br>Methyltransf_25<br>Dimerisation2<br>Methyltransf_12 | O-methyltransferase domain<br>C20_methyl_CrtF: C-20 methyltransferase BchU<br>Methyltransferase domain<br>Dimerisation domain<br>Methyltransferase domain | 5.30E-45<br>2.10E-18<br>1.20E-07<br>5.10E-07<br>1.90E-05 |
| AZM48682.1 | 5728323 | 5728626 | + | 100 | LinA | LinA | LinA (linaridin precursor) | 1.60E-08 |
| AZM48683.1 | 5728787 | 5730788 | + | 666 | LinH<br>LinE<br>LinG<br>PF00561<br>TIGR02427 | LinH<br>LinE<br>LinG<br>Abhydrolase_1<br>TIGR02427 | LinH (Fusion of LinE/LinG)<br>LinE Homolog<br>LinG Homolog<br>alpha/beta hydrolase fold<br>protocat_pcaD: 3-oxoadipate enol-lactonase | 1.40E-200<br>1.00E-93<br>5.40E-85<br>2.60E-11<br>2.30E-09 |
| AZM48684.1 | 5730799 | 5731357 | + | 185 | LinL | LinL | LinL | 1.60E-59 |
| AZM50979.1 | 5732103 | 5731428 | - | 224 | PF00196<br>TIGR03020<br>TIGR03541<br>TIGR02937<br>PF13936 | GerE<br>TIGR03020<br>TIGR03541<br>TIGR02937<br>HTH_38 | Bacterial regulatory proteins, luxR family<br>EpsA: transcriptional regulator EpsA<br>reg_near_HchA: LuxR family transcriptional regulatory, chaperone HchA-associated<br>sigma70-ECF: RNA polymerase sigma factor, sigma-70 family<br>Helix-turn-helix domain | 5.30E-19<br>7.10E-12<br>2.90E-10<br>2.10E-04<br>2.40E-04 |
| AZM48685.1 | 5732472 | 5733753 | + | 426 | PF13360<br>PF01011<br>TIGR03300<br>TIGR03075 | PQQ_2<br>PQQ<br>TIGR03300<br>TIGR03075 | PQQ-like domain<br>PQQ enzyme repeat<br>assembly_YfgL: outer membrane assembly lipoprotein YfgL<br>PQQ_enz_alc_DH: PQQ-dependent dehydrogenase, methanol/ethanol family | 9.50E-12<br>1.80E-11<br>1.70E-05<br>4.20E-04 |
| AZM48686.1 | 5733958 | 5735578 | + | 539 | PF01425<br>TIGR00132<br>TIGR02715<br>TIGR02713 | Amidase<br>TIGR00132<br>TIGR02715<br>TIGR02713 | Amidase<br>gatA: aspartyl/glutamyl-tRNA(Asn/Gln) amidotransferase, A subunit<br>amido_AtzE: amidohydrolase, AtzE family<br>allophanate_hyd: allophanate hydrolase | 1.90E-86<br>4.10E-60<br>3.90E-45<br>1.40E-40 |
| AZM50980.1 | 5736556 | 5735623 | - | 310 | TIGR00212<br>PF01379<br>PF03900 | TIGR00212<br>Porphobil_deam<br>Porphobil_deamC | hemC: hydroxymethylbilane synthase<br>Porphobilinogen deaminase, dipyromethane cofactor binding domain<br>Porphobilinogen deaminase, C-terminal domain | 7.00E-98<br>2.60E-64<br>5.80E-12 |
| AZM48687.1 | 5738589 | 5736717 | - | 623 | TIGR00976<br>PF02129<br>PF08530<br>PF00326 | TIGR00976<br>Peptidase_S15<br>PepX_C<br>Peptidase_S9 | /NonD: hydrolase CocE/NonD family protein<br>X-Pro dipeptidyl-peptidase (S15 family)<br>X-Pro dipeptidyl-peptidase C-terminal non-catalytic domain<br>Prolyl oligopeptidase family | 6.40E-151<br>7.20E-55<br>2.50E-10<br>8.90E-04 |
| AZM48688.1 | 5739750 | 5738775 | - | 324 | PF12698 | ABC2_membrane_3 | ABC-2 family transporter protein | 2.50E-06 |

| | | | | | | | | |
|---|---|---|---|---|---|---|---|---|
| AZM48689.1 | 5739856 | 5740471 | + | 204 | PF00440<br>TIGR03384<br>PF13977<br>TIGR03968<br>TIGR03613 | TetR_N<br>TIGR03384<br>TetR_C_6<br>TIGR03968<br>TIGR03613 | Bacterial regulatory proteins, tetR family<br>betaine_BetI: transcriptional repressor BetI<br>BetI-type transcriptional repressor, C-terminal<br>mycofact_TetR: mycofactocin system transcriptional regulator<br>RutR: pyrimidine utilization regulatory protein R | 5.00E-16<br>1.00E-10<br>3.20E-10<br>9.80E-09<br>3.60E-08 |

| index | leader | core | start | end | dir | score |
|---|---|---|---|---|---|---|
| 1 | MTTTLQAPTTADSRIAVWAQKNAAEHDEHFVAPTDRVVPVL | FTPLIALTVAGAVGCALGPATAFAT...PDAPDGAALRGRSVDQLTGGHPGMV | 5728323 | 5728626 | + | 26 |
| 2 | VGDPRVGPGG | SCPVRPVSAAAPRALNGRDGTARPA...WGKQPCIVADFRPTRSGTILSAVAS | 5732228 | 5732426 | + | 6 |
| 3 | MLLRNDGTELWGDGGWA | HPGKPHVAGATGTDAWSV | 5738589 | 5738697 | + | 6 |

### Results for RSS64107.1 [Streptomyces sp. WAC07061] back to top

previous - next

#### Architecture

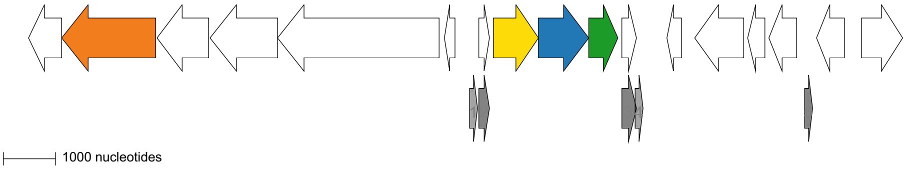

Link to nucleotide sequence

| Accession | start | end | direction | length (aa) | Pfam/HMM | name | description | E-value |
|---|---|---|---|---|---|---|---|---|
| RSS64099.1 | 2321 | 1679 | - | 213 | PF13302<br>PF00583<br>PF13523 | Acetyltransf_3<br>Acetyltransf_1<br>Acetyltransf_8 | Acetyltransferase (GNAT) domain<br>Acetyltransferase (GNAT) family<br>Acetyltransferase (GNAT) domain | 3.90E-22<br>1.80E-06<br>1.60E-04 |
| RSS64100.1 | 4123 | 2308 | - | 604 | TIGR02204<br>TIGR02203<br>TIGR01846<br>TIGR03796<br>TIGR00958 | TIGR02204<br>TIGR02203<br>TIGR01846<br>TIGR03796<br>TIGR00958 | MsbA_rel: ABC transporter, permease/ATP-binding protein<br>MsbA_lipidA: lipid A export permease/ATP-binding protein MsbA<br>type_I_sec_HlyB: type I secretion system ATPase<br>NHLM_micro_ABC1: NHLM bacteriocin system ABC transporter, peptidase/ATP-binding protein<br>3a01208: antigen peptide transporter 2 | 1.50E-110<br>2.20E-110<br>3.60E-100<br>5.10E-97<br>4.70E-96 |
| RSS64101.1 | 5124 | 4143 | - | 326 | TIGR03891<br>PF14028 | TIGR03891<br>Lant_dehydr_C | thiopep_ocin: thiopeptide-type bacteriocin biosynthesis domain<br>Lantibiotic biosynthesis dehydratase C-term | 2.00E-24<br>5.10E-13 |
| RSS64102.1 | 6460 | 5146 | - | 437 | PF05147<br>TIGR03897 | LANC_like<br>TIGR03897 | Lanthionine synthetase C-like protein<br>lanti_2_LanM: type 2 lantibiotic biosynthesis protein LanM | 4.10E-41<br>3.00E-05 |
| RSS64103.1 | 9540 | 6456 | - | 1027 | PF04738<br>TIGR03891<br>PF14028 | Lant_dehydr_N<br>TIGR03891<br>Lant_dehydr_C | Lantibiotic dehydratase, N terminus<br>thiopep_ocin: thiopeptide-type bacteriocin biosynthesis domain<br>Lantibiotic biosynthesis dehydratase C-term | 5.40E-156<br>1.30E-39<br>4.00E-18 |
| RSS64104.1 | 9844 | 9643 | - | 66 | NO PFAM MATCH | - | - | - |
| RSS64105.1 | 10320 | 10509 | + | 62 | NO PFAM MATCH | - | - | - |
| RSS64106.1 | 10600 | 11467 | + | 288 | LinG<br>LinH | LinG<br>LinH | LinG Homolog<br>LinH (Fusion of LinE/LinG) | 2.10E-87<br>2.80E-84 |
| RSS64107.1 | 11456 | 12416 | + | 319 | LinE<br>LinH<br>PF12146<br>PF00561<br>TIGR03611 | LinE<br>LinH<br>Hydrolase_4<br>Abhydrolase_1<br>TIGR03611 | LinE Homolog<br>LinH (Fusion of LinE/LinG)<br>Serine aminopeptidase, S33<br>alpha/beta hydrolase fold<br>RutD: pyrimidine utilization protein D | 3.20E-66<br>3.70E-63<br>1.60E-11<br>1.20E-06<br>5.80E-04 |
| RSS64108.1 | 12412 | 12970 | + | 185 | LinL | LinL | LinL | 1.40E-56 |
| RSS64109.1 | 13052 | 13328 | + | 91 | NO PFAM MATCH | - | - | - |
| RSS64110.1 | 14208 | 13920 | - | 95 | NO PFAM MATCH | - | - | - |
| RSS64134.1 | 15395 | 14453 | - | 313 | TIGR04187<br>-<br>-<br>-<br>TIGR04185 | TIGR04187<br>Subclass_4<br>Subclass_5<br>Subclass_6<br>TIGR04185 | GRASP_SAV_5884: ATP-grasp ribosomal peptide maturase, SAV_5884 family<br>-<br>-<br>-<br>ATPgraspMvdC: ATP-grasp ribosomal peptide maturase, MvdC family | 6.00E-45<br>2.30E-40<br>1.60E-36<br>1.60E-35<br>6.80E-32 |
| RSS64111.1 | 15805 | 15478 | - | 108 | NO PFAM MATCH | - | - | - |
| RSS64112.1 | 16408 | 15874 | - | 177 | NO PFAM MATCH | - | - | - |
| RSS64113.1 | 17314 | 16795 | - | 172 | NO PFAM MATCH | - | - | - |
| WP_015031706.1 | 17666 | 18447 | + | 260 | INFERRED GENE | - | - | - |

| index | leader | core | start | end | dir | score |
|---|---|---|---|---|---|---|
| 1 | VSAAYCGCRSGPEF | LTVTRSVPSGHRVRIHGGFLFFCECSRHCVRPPNLRFVSR | 10130 | 10295 | + | 6 |
| 2 | MDRSRILAIAGGVLTVGVLAAGPAAAAERTE | QTEGASYAVQASGDRSAAGGDLVVNKFPCCV | 10320 | 10509 | + | 7 |
| 3 | MTSRERLLHHDFPSRVLALGELPPS...RRRLERLSPRDRSGLPTGVRAAFDC | PTAGCRRTGPGAGAAA | 13052 | 13328 | + | 7 |
| 4 | VLGLLPEHPGVHC | TTAVTVALVHTGDDEPPQWSAGRSPPCGNCSPAWPTPVC | 13308 | 13467 | + | 6 |
| 5 | LSDRFRSGSGGLRVTPARQAGLRGTA | AEGPAGGWATPPPAPSTARWSRTRRS | 16555 | 16714 | + | 7 |

### Results for KOU14214.1 [Streptomyces sp. WM6372] back to top

previous - next

### Architecture

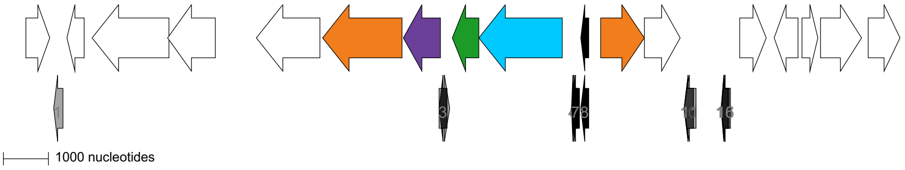

Link to nucleotide sequence

| Accession | start | end | direction | length (aa) | Pfam/HMM | name | description | E-value |
|---|---|---|---|---|---|---|---|---|
| KOU14192.1 | 1045 | 1600 | + | 184 | PF13384<br>TIGR01764<br>TIGR03830<br>PF12728<br>PF13542 | HTH_23<br>TIGR01764<br>TIGR03830<br>HTH_17<br>HTH_Tnp_ISL3 | Homeodomain-like domain<br>excise: DNA binding domain, excisionase family<br>CxxCG_CxxCG_HTH: putative zinc finger/helix-turn-helix protein, YgiT family<br>Helix-turn-helix domain<br>Helix-turn-helix domain of transposase family ISL3 | 1.90E-06<br>1.80E-04<br>2.30E-04<br>2.90E-04<br>4.20E-04 |
| KOU14193.1 | 2360 | 1982 | - | 125 | NO PFAM MATCH | - | - | - |
| KOU14194.1 | 4223 | 2543 | - | 559 | TIGR01613<br>PF08706 | TIGR01613<br>D5_N | primase_Cterm: phage/plasmid primase, P4 family, C-terminal domain<br>D5 N terminal like | 6.30E-76<br>1.50E-27 |
| KOU14211.1 | 5278 | 4219 | - | 352 | PF09250 | Prim-Pol | Bifunctional DNA primase/polymerase, N-terminal | 3.60E-36 |
| KOU14212.1 | 7585 | 6175 | - | 469 | TIGR02734 | TIGR02734 | crtI_fam: phytoene desaturase | 2.00E-05 |
| KOU14213.1 | 9405 | 7656 | - | 582 | TIGR02204<br>TIGR02203<br>TIGR00958<br>TIGR03796<br>TIGR01846 | TIGR02204<br>TIGR02203<br>TIGR00958<br>TIGR03796<br>TIGR01846 | MsbA_rel: ABC transporter, permease/ATP-binding protein<br>MsbA_lipidA: lipid A export permease/ATP-binding protein MsbA<br>3a01208: antigen peptide transporter 2<br>NHLM_micro_ABC1: NHLM bacteriocin system ABC transporter, peptidase/ATP-binding protein<br>type_I_sec_HlyB: type I secretion system ATPase | 6.20E-99<br>1.10E-92<br>3.80E-84<br>7.90E-84<br>4.20E-82 |
| KOU14195.1 | 10250 | 9437 | - | 270 | PF13649<br>PF08241<br>PF13489<br>PF08242<br>PF13847 | Methyltransf_25<br>Methyltransf_11<br>Methyltransf_23<br>Methyltransf_12<br>Methyltransf_31 | Methyltransferase domain<br>Methyltransferase domain<br>Methyltransferase domain<br>Methyltransferase domain<br>Methyltransferase domain | 4.00E-16<br>9.90E-13<br>1.90E-09<br>4.70E-07<br>6.30E-07 |
| KOU14196.1 | 11108 | 10526 | - | 193 | LinL | LinL | LinL | 7.80E-68 |
| KOU14214.1 | 12961 | 11104 | - | 618 | LinH<br>LinG<br>LinE<br>PF12697<br>PF00561 | LinH<br>LinG<br>LinE<br>Abhydrolase_6<br>Abhydrolase_1 | LinH (Fusion of LinE/LinG)<br>LinG Homolog<br>LinE Homolog<br>Alpha/beta hydrolase family<br>alpha/beta hydrolase fold | 1.20E-246<br>1.70E-125<br>4.70E-100<br>1.20E-16<br>3.00E-09 |
| KOU14197.1 | 13571 | 13370 | - | 66 | LinA | LinA | LinA (linaridin precursor) | 6.90E-14 |
| KOU14198.1 | 13828 | 14785 | + | 318 | TIGR01188<br>TIGR03864<br>TIGR01288<br>TIGR01166<br>TIGR03740 | TIGR01188<br>TIGR03864<br>TIGR01288<br>TIGR01166<br>TIGR03740 | drrA: daunorubicin resistance ABC transporter, ATP-binding protein<br>PQQ_ABC_ATP: ABC transporter, ATP-binding subunit, PQQ-dependent alcohol dehydrogenase system<br>nodI: nodulation ABC transporter NodI<br>cbiO: cobalt ABC transporter, ATP-binding protein<br>galliderm_ABC: lantibiotic protection ABC transporter, ATP-binding subunit | 4.10E-61<br>1.00E-43<br>5.00E-40<br>2.50E-32<br>4.50E-32 |
| KOU14199.1 | 14781 | 15594 | + | 270 | PF01061<br>PF12698<br>TIGR03062<br>TIGR01291<br>PF12679 | ABC2_membrane<br>ABC2_membrane_3<br>TIGR03062<br>TIGR01291<br>ABC2_membrane_2 | ABC-2 type transporter<br>ABC-2 family transporter protein<br>pip_yhgE_Cterm: YhgE/Pip C-terminal domain<br>nodJ: ABC-2 type transporter, NodJ family<br>ABC-2 family transporter protein | 3.30E-14<br>1.20E-11<br>6.00E-08<br>5.80E-05<br>4.40E-04 |
| KOU14200.1 | 16897 | 17506 | + | 202 | PF00072<br>TIGR02875<br>PF00196<br>TIGR02956<br>TIGR02154 | Response_reg<br>TIGR02875<br>GerE<br>TIGR02956<br>TIGR02154 | Response regulator receiver domain<br>spore_0_A: sporulation transcription factor Spo0A<br>Bacterial regulatory proteins, luxR family<br>TMAO_torS: TMAO reductase sytem sensor TorS<br>PhoB: phosphate regulon transcriptional regulatory protein PhoB | 1.30E-22<br>6.90E-18<br>2.00E-12<br>5.20E-10<br>8.90E-09 |
| KOU14201.1 | 18211 | 17683 | - | 175 | PF13302<br>PF00583 | Acetyltransf_3<br>Acetyltransf_1 | Acetyltransferase (GNAT) domain<br>Acetyltransferase (GNAT) family | 3.20E-31<br>1.80E-05 |
| KOU14202.1 | 18306 | 18654 | + | 115 | PF00392<br>TIGR02325<br>TIGR02018<br>TIGR03338<br>TIGR02404 | GntR<br>TIGR02325<br>TIGR02018<br>TIGR03338<br>TIGR02404 | Bacterial regulatory proteins, gntR family<br>C_P_lyase_phnF: phosphonate metabolism transcriptional regulator PhnF<br>his_ut_repres: histidine utilization repressor<br>phnR_burk: phosphonate utilization associated transcriptional regulator<br>trehalos_R_Bsub: trehalose operon repressor | 2.90E-17<br>7.70E-13<br>1.10E-11<br>4.00E-06<br>4.50E-06 |
| KOU14203.1 | 18692 | 19610 | + | 305 | PF07853<br>TIGR03074 | DUF1648<br>TIGR03074 | Protein of unknown function (DUF1648)<br>PQQ_membr_DH: membrane-bound PQQ-dependent dehydrogenase, glucose/quinate/shikimate family | 6.40E-05<br>2.10E-04 |
| KOU14204.1 | 19762 | 20455 | + | 230 | NO PFAM MATCH | - | - | - |

| index | leader | core | start | end | dir | score |
|---|---|---|---|---|---|---|
| 1 | LSPSGPCSVPLLSPAPASGRGEDADLVALRIGQ | DPPRDAALGHVGGRRTEGLKACDQLRSVGRGGGRERRQEP | 1889 | 1667 | - | 6 |
| 2 | VGRCHRFGFFLFS | STPAAMNRATPTAIALTVFMTSAAV...HPATAGLCMVFSLGYAVARGPYVRT | 10235 | 10460 | + | 6 |
| 3 | MHKPAVAGCLLIGLAVIGF | LTAADVMNTVSAIAVGVALFIAAGVELKRKKPNR | 10408 | 10246 | - | 9 |
| 4 | VFPVAAPVLATPAAFMGG | IAISCAVVSAFEAGRGNG | 13274 | 13163 | - | 14 |
| 5 | MDIPEEGVVFPVAAPVL | ATPAAFMGGIAISCAVVSAFEAGRGNG | 13298 | 13163 | - | 27 |

| | | | | | | |
|---|---|---|---|---|---|---|
| 6 | MNEVTALLDEMDIPEEGVVFPVAAPVL | ATPAAFMGGIAISCAVVSAFEAGRGNG | 13328 | 13163 | - | 30 |
| 7 | VRKDAVMNEVTALLDEMDIPEEGVVFPVAAPVL | ATPAAFMGGIAISCAVVSAFEAGRGNG | 13346 | 13163 | - | 30 |
| 8 | MTEIVTFVNNLDLGTIQAGETRTDQAPVL | ATPAAALAAIGGAKAVGAFVGGATVVGGAYALGRAIG | 13571 | 13370 | - | 29 |
| 9 | VTPATAQPIPAGSRLASAASAASAASRWAS | SGQSSEASSSHGIRQSIQRVCVVARRTSVW | 15868 | 15685 | - | 6 |
| 10 | VVAPRPWRAVTP | ATAQPIPAGSRLASAASAASAASRW...EASSSHGIRQSIQRVCVVARRTSVW | 15895 | 15685 | - | 7 |
| 11 | MRPAERFARPSPYAIAVVAPRPWRAVTP | ATAQPIPAGSRLASAASAASAASRW...EASSSHGIRQSIQRVCVVARRTSVW | 15943 | 15685 | - | 6 |
| 12 | VTPSRTTQPRAVCASWDRSDPVVRR | IRQAMPAASRMAWAPWSSVARSVER | 16639 | 16486 | - | 6 |
| 13 | LVTPSRTTQPRAVCASWDRSDPVVR | RIRQAMPAASRMAWAPWSSVARSVER | 16642 | 16486 | - | 6 |
| 14 | VHVPAWEWRSTLVTPSRTTQPRAVCASWDRS | DPVVRRIRQAMPAASRMAWAPWSSVARSVER | 16675 | 16486 | - | 6 |
| 15 | MVHVPAWEWRSTLVTPSRTTQPRAVCASWDR | SDPVVRRIRQAMPAASRMAWAPWSSVARSVER | 16678 | 16486 | - | 6 |
| 16 | VSESEMVHVPAWEWRSTLVTPSRTTQPRAVCASW | DRSDPVVRRIRQAMPAASRMAWAPWSSVARSVER | 16693 | 16486 | - | 6 |

### Results for SCK38574.1 [Streptomyces sp. WMMB 322] back to top

previous - next

#### Architecture

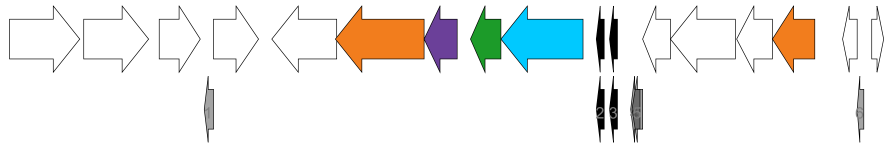

1000 nucleotides

Link to nucleotide sequence

| Accession | start | end | direction | length (aa) | Pfam/HMM | name | description | E-value |
|---|---|---|---|---|---|---|---|---|
| SCK38496.1 | 127939 | 129571 | + | 543 | NO PFAM MATCH | - | - | - |
| SCK38506.1 | 129654 | 131151 | + | 498 | NO PFAM MATCH | - | - | - |
| SCK38514.1 | 131391 | 132345 | + | 317 | PF01259<br>TIGR00081<br>TIGR02735 | SAICAR_synt<br>TIGR00081<br>TIGR02735 | SAICAR synthetase<br>purC: phosphoribosylaminoimidazolesuccinocarboxamide synthase<br>purC_vibrio: phosphoribosylaminoimidazole-succinocarboxamide synthase | 4.30E-99<br>1.10E-68<br>4.20E-23 |
| SCK38525.1 | 132656 | 133679 | + | 340 | PF00196<br>PF13384 | GerE<br>HTH_23 | Bacterial regulatory proteins, luxR family<br>Homeodomain-like domain | 8.50E-05<br>4.20E-04 |
| SCK38534.1 | 135475 | 133996 | - | 492 | TIGR02734<br>TIGR03467<br>PF13450<br>TIGR00292<br>TIGR01372 | TIGR02734<br>TIGR03467<br>NAD_binding_8<br>TIGR00292<br>TIGR01372 | crtI_fam: phytoene desaturase<br>HpnE: squalene-associated FAD-dependent desaturase<br>NAD(P)-binding Rossmann-like domain<br>TIGR00292: thiazole biosynthesis enzyme<br>soxA: sarcosine oxidase, alpha subunit family | 1.00E-10<br>7.10E-10<br>3.00E-08<br>3.30E-07<br>8.80E-06 |
| SCK38543.1 | 137511 | 135471 | - | 679 | TIGR02204<br>TIGR02203<br>TIGR00958<br>TIGR03796<br>TIGR03375 | TIGR02204<br>TIGR02203<br>TIGR00958<br>TIGR03796<br>TIGR03375 | MsbA_rel: ABC transporter, permease/ATP-binding protein<br>MsbA_lipidA: lipid A export permease/ATP-binding protein MsbA<br>3a01208: antigen peptide transporter 2<br>NHLM_micro_ABC1: NHLM bacteriocin system ABC transporter, peptidase/ATP-binding protein<br>type_I_sec_LssB: type I secretion system ATPase | 6.50E-101<br>1.50E-98<br>7.00E-85<br>5.00E-82<br>4.20E-81 |
| SCK38555.1 | 138277 | 137518 | - | 252 | PF13649<br>PF13489<br>PF08241<br>PF13847<br>TIGR01983 | Methyltransf_25<br>Methyltransf_23<br>Methyltransf_11<br>Methyltransf_31<br>TIGR01983 | Methyltransferase domain<br>Methyltransferase domain<br>Methyltransferase domain<br>Methyltransferase domain<br>UbiG: 3-demethylubiquinone-9 3-O-methyltransferase | 3.00E-18<br>4.80E-14<br>5.40E-13<br>6.70E-13<br>5.60E-11 |
| SCK38564.1 | 139270 | 138562 | - | 235 | LinL | LinL | LinL | 8.40E-63 |
| SCK38574.1 | 141180 | 139266 | - | 637 | LinH<br>LinG<br>LinE<br>PF12697<br>TIGR02427 | LinH<br>LinG<br>LinE<br>Abhydrolase_6<br>TIGR02427 | LinH (Fusion of LinE/LinG)<br>LinG Homolog<br>LinE Homolog<br>Alpha/beta hydrolase family<br>protocat_pcaD: 3-oxoadipate enol-lactonase | 1.50E-246<br>6.70E-121<br>2.90E-98<br>3.50E-14<br>1.10E-08 |
| SCK38582.1 | 141672 | 141492 | - | 59 | LinA | LinA | LinA (linaridin precursor) | 3.00E-16 |
| SCK38591.1 | 141958 | 141784 | - | 57 | LinA | LinA | LinA (linaridin precursor) | 1.20E-14 |
| SCK38601.1 | 143199 | 142533 | - | 221 | PF00072<br>TIGR02875<br>PF00196<br>TIGR02154<br>TIGR02956 | Response_reg<br>TIGR02875<br>GerE<br>TIGR02154<br>TIGR02956 | Response regulator receiver domain<br>spore_0_A: sporulation transcription factor Spo0A<br>Bacterial regulatory proteins, luxR family<br>PhoB: phosphate regulon transcriptional regulatory protein PhoB<br>TMAO_torS: TMAO reductase sytem sensor TorS | 2.40E-25<br>6.00E-13<br>2.60E-12<br>8.20E-12<br>6.80E-10 |
| SCK38610.1 | 144695 | 143195 | - | 499 | PF07730 | HisKA_3 | Histidine kinase | 1.30E-14 |
| SCK38620.1 | 145532 | 144719 | - | 270 | PF01061<br>PF12698<br>TIGR03062<br>TIGR01247 | ABC2_membrane<br>ABC2_membrane_3<br>TIGR03062<br>TIGR01247 | ABC-2 type transporter<br>ABC-2 family transporter protein<br>pip_yhgE_Cterm: YhgE/Pip C-terminal domain<br>drrB: daunorubicin resistance ABC transporter membrane protein | 1.30E-19<br>6.10E-12<br>7.70E-08<br>7.60E-04 |

| | | | | | | | | |
|---|---|---|---|---|---|---|---|---|
| SCK38628.1 | 146515 | 145528 | - | 328 | TIGR01188<br>TIGR03864<br>TIGR01288<br>TIGR04406<br>TIGR03740 | TIGR01188<br>TIGR03864<br>TIGR01288<br>TIGR04406<br>TIGR03740 | drrA: daunorubicin resistance ABC transporter, ATP-binding protein<br>PQQ_ABC_ATP: ABC transporter, ATP-binding subunit, PQQ-dependent alcohol dehydrogenase system<br>nodI: nodulation ABC transporter NodI<br>LPS_export_lptB: LPS export ABC transporter ATP-binding protein<br>galliderm_ABC: lantibiotic protection ABC transporter, ATP-binding subunit | 3.30E-69<br>5.00E-49<br>4.10E-47<br>4.20E-39<br>7.70E-36 |
| SCK38641.1 | 147504 | 147147 | - | 118 | PF11774 | Lsr2 | Lsr2 | 7.50E-35 |
| SCK38650.1 | 147847 | 148105 | + | 85 | PF02700<br>TIGR00302 | PurS<br>TIGR00302 | Phosphoribosylformylglycinamidine (FGAM) synthase<br>TIGR00302: phosphoribosylformylglycinamidine synthase, purS protein | 6.50E-28<br>4.50E-20 |

| index | leader | core | start | end | dir | score |
|---|---|---|---|---|---|---|
| 1 | VNSPCPVAPPGPVHVGGGQKLSTHMRNRTPSIEKTKAPNTPRR | DTFSKGPFIYCASAPVCTLPDGRFNSG | 132655 | 132442 | - | 6 |
| 2 | MKDVTSQINESAELASVDIADTEGVVFTDYTPVM | GTPGAFLGGVAIASATVAAFEAGQG | 141672 | 141492 | - | 29 |
| 3 | MLEEKLNLQIQDQLHEGHEGEIAEEAAPVL | ATPALAGAFVGGAAVVTGAYAAGKAVG | 141958 | 141784 | - | 30 |
| 4 | VCGAGRRCAGMRPDTSTGLTRGRIRTIQQWCTMT | GHPFPRRISLAIPCLPSCQFSTIEVYLSLPSGKP | 142482 | 142275 | - | 6 |
| 5 | VTRGPGPARAPAFLARVHGVRGRAAVRRDAPRYVNGA | HTRTNSDHSAVVHHDGTSIPEADIFGNPLLTLVPILNN | 142537 | 142309 | - | 6 |
| 6 | VKKVWSRLRSDTRVEILSLHIAKSAA | VQKKCPTSIMRITRTSPWSSHVSAGIP | 147662 | 147500 | - | 6 |

### Results for WP_079100284.1 [Streptomyces sp. WMMB 322] back to top

previous - next

#### Architecture

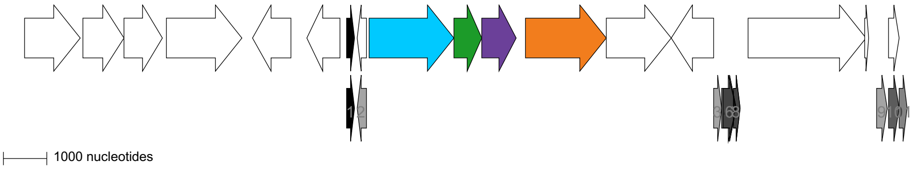

Link to nucleotide sequence

| Accession | start | end | direction | length (aa) | Pfam/HMM | name | description | E-value |
|---|---|---|---|---|---|---|---|---|
| WP_055487588.1 | 8205 | 9501 | + | 431 | PF13416<br>PF01547<br>TIGR03850<br>TIGR03851<br>PF13343 | SBP_bac_8<br>SBP_bac_1<br>TIGR03850<br>TIGR03851<br>SBP_bac_6 | Bacterial extracellular solute-binding protein<br>Bacterial extracellular solute-binding protein<br>bind_CPR_0540: carbohydrate ABC transporter substrate-binding protein, CPR_0540 family<br>chitin_NgcE: carbohydrate ABC transporter, N-acetylglucosamine/diacetylchitobiose-binding protein<br>Bacterial extracellular solute-binding protein | 5.50E-45<br>2.00E-36<br>6.90E-16<br>1.60E-15<br>1.10E-09 |
| WP_055487704.1 | 9565 | 10501 | + | 311 | PF00528<br>TIGR03262<br>TIGR02140<br>TIGR00969<br>TIGR02141 | BPD_transp_1<br>TIGR03262<br>TIGR02140<br>TIGR00969<br>TIGR02141 | Binding-protein-dependent transport system inner membrane component<br>PhnU2: putative 2-aminoethylphosphonate ABC transporter, permease protein<br>permease_CysW: sulfate ABC transporter, permease protein CysW<br>3a0106s02: sulfate ABC transporter, permease protein<br>modB_ABC: molybdate ABC transporter, permease protein | 5.10E-18<br>1.30E-10<br>3.00E-09<br>1.40E-05<br>5.80E-05 |
| WP_079100272.1 | 10497 | 11391 | + | 297 | PF00528<br>TIGR01581<br>TIGR02141<br>TIGR03262<br>TIGR00969 | BPD_transp_1<br>TIGR01581<br>TIGR02141<br>TIGR03262<br>TIGR00969 | Binding-protein-dependent transport system inner membrane component<br>Mo_ABC_porter: NifC-like ABC-type porter<br>modB_ABC: molybdate ABC transporter, permease protein<br>PhnU2: putative 2-aminoethylphosphonate ABC transporter, permease protein<br>3a0106s02: sulfate ABC transporter, permease protein | 8.60E-14<br>1.80E-13<br>5.00E-11<br>2.10E-08<br>4.40E-04 |
| WP_055487706.1 | 11472 | 13218 | + | 581 | TIGR02403<br>PF00128<br>TIGR02456<br>TIGR02402<br>TIGR02401 | TIGR02403<br>Alpha-amylase<br>TIGR02456<br>TIGR02402<br>TIGR02401 | trehalose_treC: alpha,alpha-phosphotrehalase<br>Alpha amylase, catalytic domain<br>treS_nterm: trehalose synthase<br>trehalose_TreZ: malto-oligosyltrehalose trehalohydrolase<br>trehalose_TreY: malto-oligosyltrehalose synthase | 2.50E-203<br>1.10E-117<br>1.40E-97<br>1.80E-16<br>3.30E-16 |
| WP_055487589.1 | 14351 | 13451 | - | 299 | PF13391 | HNH_2 | HNH endonuclease | 1.50E-09 |
| WP_079100273.1 | 15468 | 14700 | - | 255 | PF05719 | GPP34 | Golgi phosphoprotein 3 (GPP34) | 7.10E-42 |
| WP_055487591.1 | 15618 | 15813 | + | 64 | LinA | LinA | LinA (linaridin precursor) | 4.30E-12 |
| WP_107097714.1 | 16086 | 15855 | - | 76 | NO PFAM MATCH | - | - | - |
| WP_079100284.1 | 16153 | 18109 | + | 651 | LinH<br>LinE<br>PF12697<br>PF12146<br>PF00561 | LinH<br>LinE<br>Abhydrolase_6<br>Hydrolase_4<br>Abhydrolase_1 | LinH (Fusion of LinE/LinG)<br>LinE Homolog<br>Alpha/beta hydrolase family<br>Serine aminopeptidase, S33<br>alpha/beta hydrolase fold | 3.80E-174<br>6.70E-122<br>1.00E-18<br>1.40E-09<br>2.60E-09 |
| WP_139151414.1 | 18108 | 18753 | + | 214 | LinL | LinL | LinL | 7.30E-73 |
| WP_055487596.1 | 18749 | 19520 | + | 256 | PF13649<br>PF08241<br>PF08242<br>PF13489<br>PF13847 | Methyltransf_25<br>Methyltransf_11<br>Methyltransf_12<br>Methyltransf_23<br>Methyltransf_31 | Methyltransferase domain<br>Methyltransferase domain<br>Methyltransferase domain<br>Methyltransferase domain<br>Methyltransferase domain | 2.10E-15<br>7.40E-10<br>8.30E-09<br>6.80E-08<br>9.00E-06 |
| WP_079100285.1 | 19752 | 21615 | + | 620 | TIGR02203<br>TIGR02204<br>TIGR00958<br>TIGR03796<br>TIGR03375 | TIGR02203<br>TIGR02204<br>TIGR00958<br>TIGR03796<br>TIGR03375 | MsbA_lipidA: lipid A export permease/ATP-binding protein MsbA<br>MsbA_rel: ABC transporter, permease/ATP-binding protein<br>3a01208: antigen peptide transporter 2<br>NHLM_micro_ABC1: NHLM bacteriocin system ABC transporter, peptidase/ATP-binding protein<br>type_I_sec_LssB: type I secretion system ATPase | 2.50E-101<br>9.40E-101<br>1.20E-89<br>9.10E-86<br>1.30E-81 |

| | | | | | | | | |
|---|---|---|---|---|---|---|---|---|
| WP_055487597.1 | 21611 | 23090 | + | 492 | TIGR02734<br>TIGR03467<br>PF13450<br>PF01593<br>TIGR00292 | TIGR02734<br>TIGR03467<br>NAD_binding_8<br>Amino_oxidase<br>TIGR00292 | crtI_fam: phytoene desaturase<br>HpnE: squalene-associated FAD-dependent desaturase<br>NAD(P)-binding Rossmann-like domain<br>Flavin containing amine oxidoreductase<br>TIGR00292: thiazole biosynthesis enzyme | 3.00E-12<br>5.30E-09<br>1.00E-08<br>1.80E-08<br>4.80E-08 |
| WP_055487598.1 | 24072 | 23097 | - | 324 | PF01978<br>PF00196<br>TIGR03541 | TrmB<br>GerE<br>TIGR03541 | Sugar-specific transcriptional regulator TrmB<br>Bacterial regulatory proteins, luxR family<br>reg_near_HchA: LuxR family transcriptional regulatory, chaperone HchA-associated | 2.60E-06<br>1.80E-05<br>6.80E-05 |
| WP_139151415.1 | 24884 | 27575 | + | 896 | PF00196<br>PF13424<br>PF13384<br>PF13191<br>TIGR03020 | GerE<br>TPR_12<br>HTH_23<br>AAA_16<br>TIGR03020 | Bacterial regulatory proteins, luxR family<br>Tetratricopeptide repeat<br>Homeodomain-like domain<br>AAA ATPase domain<br>EpsA: transcriptional regulator EpsA | 2.20E-14<br>9.80E-08<br>1.20E-07<br>1.60E-07<br>1.40E-06 |
| WP_010355355.1 | 27571 | 27655 | + | 28 | INFERRED GENE | - | - | - |
| WP_107097715.1 | 28115 | 28349 | + | 77 | NO PFAM MATCH | - | - | - |

| index | leader | core | start | end | dir | score |
|---|---|---|---|---|---|---|
| 1 | MLLEKVNNSHLLEAVPMDAANVEDLSSAEPVL | STPAAAAGGALVTAGAFGAGVGLGVTQAQADG | 15618 | 15813 | + | 29 |
| 2 | MGQPPAACPAAFPTAMPAAPAP | ATAAAPASPAPTAAGVARTGDIRPF...PAWIETPSSSWLALTLVDIAVTFTP | 16086 | 15855 | - | 6 |
| 3 | LVSGPRPRCSWAGWGRRPALMRDGRRGGVGA | GTLRSRSRMGVGLRAPHSGASSRGRAAPLTGVG | 24076 | 24271 | + | 6 |
| 4 | VSAERELLWQSVSDPGGLGGWRTAR...IGGSQNPWQRFRRAVFGIWRSAPSP | LTPTAFDPLAALVCPLRRGYEWWSGSSQSSLGSTVIFFGPKCVDSAPK | 24260 | 24611 | + | 6 |
| 5 | LWQSVSDPGGLGGWRTARTACAPAR...IGGSQNPWQRFRRAVFGIWRSAPSP | LTPTAFDPLAALVCPLRRGYEWWSGSSQSSLGSTVIFFGPKCVDSAPK | 24281 | 24611 | + | 6 |
| 6 | VSDPGGLGGWRTARTACAPARRVLP...IGGSQNPWQRFRRAVFGIWRSAPSP | LTPTAFDPLAALVCPLRRGYEWWSGSSQSSLGSTVIFFGPKCVDSAPK | 24293 | 24611 | + | 6 |
| 7 | MVVWLIAVISGFYSHLLRPEVCGLRAK | VICLPQSPVLSILTGCRLSVHRQIVQR | 24526 | 24691 | + | 7 |
| 8 | VVWLIAVISGFYSHLLRPEVCGLRAK | VICLPQSPVLSILTGCRLSVHRQIVQR | 24529 | 24691 | + | 7 |
| 9 | VALGGKSTAPRRRWVGPGGHPLAEHGGCSSPPSPVSCSPL | LRACSTSWSISLACAAGYGGAARGQGIPRRSRPGGAEISA | 27833 | 28076 | + | 6 |
| 10 | MDDEPNQQTSDGGTDEARDWNWGPGAALGISLGVALGA | AFGNVAVGVALGVAFFPVFAMAFRLNGGDRRRTGQEPGR | 28115 | 28349 | + | 8 |
| 11 | VPARIRPSELQVALTRWRLDRGRLRCKG | GLPGSASADTELVNADAHGHRAGDAADH | 28365 | 28536 | + | 7 |

### Results for SCK46711.1 [Streptomyces sp. WMMB 322] back to top

previous - next

#### Architecture

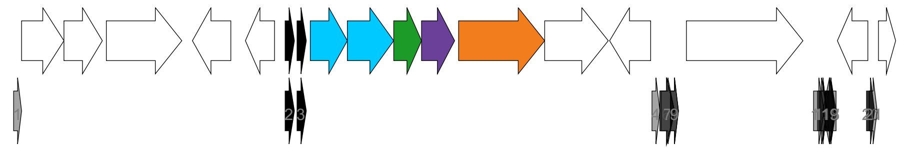

Link to nucleotide sequence

| Accession | start | end | direction | length (aa) | Pfam/HMM | name | description | E-value |
|---|---|---|---|---|---|---|---|---|
| SCK46640.1 | 9505 | 10501 | + | 331 | PF00528<br>TIGR03262<br>TIGR02140<br>TIGR00969<br>TIGR02141 | BPD_transp_1<br>TIGR03262<br>TIGR02140<br>TIGR00969<br>TIGR02141 | Binding-protein-dependent transport system inner membrane component<br>PhnU2: putative 2-aminoethylphosphonate ABC transporter, permease protein<br>permease_CysW: sulfate ABC transporter, permease protein CysW<br>3a0106s02: sulfate ABC transporter, permease protein<br>modB_ABC: molybdate ABC transporter, permease protein | 6.10E-18<br>1.60E-10<br>3.00E-09<br>2.00E-05<br>8.20E-05 |
| SCK46650.1 | 10497 | 11391 | + | 297 | PF00528<br>TIGR01581<br>TIGR02141<br>TIGR03262<br>TIGR00969 | BPD_transp_1<br>TIGR01581<br>TIGR02141<br>TIGR03262<br>TIGR00969 | Binding-protein-dependent transport system inner membrane component<br>Mo_ABC_porter: NifC-like ABC-type porter<br>modB_ABC: molybdate ABC transporter, permease protein<br>PhnU2: putative 2-aminoethylphosphonate ABC transporter, permease protein<br>3a0106s02: sulfate ABC transporter, permease protein | 8.60E-14<br>1.80E-13<br>5.00E-11<br>2.10E-08<br>4.40E-04 |
| SCK46659.1 | 11457 | 13218 | + | 586 | TIGR02403<br>PF00128<br>TIGR02456<br>TIGR02402<br>TIGR02401 | TIGR02403<br>Alpha-amylase<br>TIGR02456<br>TIGR02402<br>TIGR02401 | trehalose_treC: alpha,alpha-phosphotrehalase<br>Alpha amylase, catalytic domain<br>treS_nterm: trehalose synthase<br>trehalose_TreZ: malto-oligosyltrehalose trehalohydrolase<br>trehalose_TreY: malto-oligosyltrehalose synthase | 2.60E-203<br>1.20E-117<br>1.50E-97<br>1.30E-16<br>3.40E-16 |
| SCK46670.1 | 14351 | 13451 | - | 299 | PF13391 | HNH_2 | HNH endonuclease | 1.50E-09 |
| SCK46677.1 | 15363 | 14700 | - | 220 | PF05719 | GPP34 | Golgi phosphoprotein 3 (GPP34) | 4.30E-42 |
| SCK46687.1 | 15618 | 15813 | + | 64 | LinA | LinA | LinA (linaridin precursor) | 4.30E-12 |
| SCK46695.1 | 15868 | 16081 | + | 70 | LinA | LinA | LinA (linaridin precursor) | 2.20E-15 |
| SCK46704.1 | 16184 | 17057 | + | 290 | LinH<br>LinG | LinH<br>LinG | LinH (Fusion of LinE/LinG)<br>LinG Homolog | 1.80E-121<br>3.60E-115 |
| SCK46711.1 | 17053 | 18109 | + | 351 | LinH<br>LinE<br>PF12697<br>PF12146<br>PF00561 | LinH<br>LinE<br>Abhydrolase_6<br>Hydrolase_4<br>Abhydrolase_1 | LinH (Fusion of LinE/LinG)<br>LinE Homolog<br>Alpha/beta hydrolase family<br>Serine aminopeptidase, S33<br>alpha/beta hydrolase fold | 6.10E-137<br>1.10E-122<br>1.50E-19<br>3.90E-10<br>8.60E-10 |
| SCK46720.1 | 18108 | 18753 | + | 214 | LinL | LinL | LinL | 7.30E-73 |

| | | | | | | | | |
|---|---|---|---|---|---|---|---|---|
| SCK46728.1 | 18749 | 19520 | + | 256 | PF13649<br>PF08241<br>PF08242<br>PF13489<br>PF13847 | Methyltransf_25<br>Methyltransf_11<br>Methyltransf_12<br>Methyltransf_23<br>Methyltransf_31 | Methyltransferase domain<br>Methyltransferase domain<br>Methyltransferase domain<br>Methyltransferase domain<br>Methyltransferase domain | 2.10E-15<br>7.40E-10<br>8.30E-09<br>6.80E-08<br>9.00E-06 |
| SCK46737.1 | 19605 | 21615 | + | 669 | TIGR02204<br>TIGR02203<br>TIGR00958<br>TIGR03796<br>TIGR03375 | TIGR02204<br>TIGR02203<br>TIGR00958<br>TIGR03796<br>TIGR03375 | MsbA_rel: ABC transporter, permease/ATP-binding protein<br>MsbA_lipidA: lipid A export permease/ATP-binding protein MsbA<br>3a01208: antigen peptide transporter 2<br>NHLM_micro_ABC1: NHLM bacteriocin system ABC transporter, peptidase/ATP-binding protein<br>type_I_sec_LssB: type I secretion system ATPase | 4.50E-102<br>3.40E-101<br>1.10E-90<br>1.50E-86<br>4.90E-81 |
| SCK46745.1 | 21611 | 23090 | + | 492 | TIGR02734<br>TIGR03467<br>PF13450<br>PF01593<br>TIGR00292 | TIGR02734<br>TIGR03467<br>NAD_binding_8<br>Amino_oxidase<br>TIGR00292 | crtI_fam: phytoene desaturase<br>HpnE: squalene-associated FAD-dependent desaturase<br>NAD(P)-binding Rossmann-like domain<br>Flavin containing amine oxidoreductase<br>TIGR00292: thiazole biosynthesis enzyme | 3.00E-12<br>5.30E-09<br>1.00E-08<br>1.80E-08<br>4.80E-08 |
| SCK46752.1 | 24072 | 23097 | - | 324 | PF01978<br>PF00196<br>TIGR03541 | TrmB<br>GerE<br>TIGR03541 | Sugar-specific transcriptional regulator TrmB<br>Bacterial regulatory proteins, luxR family<br>reg_near_HchA: LuxR family transcriptional regulatory, chaperone HchA-associated | 2.60E-06<br>1.80E-05<br>6.80E-05 |
| SCK46758.1 | 24884 | 27575 | + | 896 | PF00196<br>PF13424<br>PF13384<br>PF13191<br>TIGR03020 | GerE<br>TPR_12<br>HTH_23<br>AAA_16<br>TIGR03020 | Bacterial regulatory proteins, luxR family<br>Tetratricopeptide repeat<br>Homeodomain-like domain<br>AAA ATPase domain<br>EpsA: transcriptional regulator EpsA | 2.20E-14<br>9.80E-08<br>1.20E-07<br>1.60E-07<br>1.40E-06 |
| SCK46764.1 | 29082 | 28392 | - | 229 | NO PFAM MATCH | - | - | - |
| SCK46771.1 | 29329 | 29719 | + | 129 | PF00903<br>TIGR03211<br>PF12681<br>PF18029<br>TIGR02295 | Glyoxalase<br>TIGR03211<br>Glyoxalase_2<br>Glyoxalase_6<br>TIGR02295 | Glyoxalase/Bleomycin resistance protein/Dioxygenase superfamily<br>catechol_2_3: catechol 2,3 dioxygenase<br>Glyoxalase-like domain<br>Glyoxalase-like domain<br>HpaD: 3,4-dihydroxyphenylacetate 2,3-dioxygenase | 7.10E-11<br>7.20E-07<br>1.50E-05<br>2.80E-05<br>6.00E-05 |

| index | leader | core | start | end | dir | score |
|---|---|---|---|---|---|---|
| 1 | MKQLKTARPRP | ATPSWTKIDDILRKQVQLAVRGDVPVQKALDDAVEKIDPLLARD | 9333 | 9501 | + | 6 |
| 2 | MLLEKVNNSHLLEAVPMDAANVEDLSSAEPVL | STPAAAAGGALVTAGAFGAGVGLGVTQAQADG | 15618 | 15813 | + | 29 |
| 3 | MTAISTKVNASQELDGVSIHAGENEWNGRMSPVL | ATPAAVGAGLAGAAAVAGAGAAGIAVGNAAGQAAGG | 15868 | 16081 | + | 29 |
| 4 | LVSGPRPRCSWAGWGRRPALMRDGRRGGVGA | GTLRSRSRMGVGLRAPHSGASSRGRAAPLTGVG | 24076 | 24271 | + | 6 |
| 5 | VSAERELLWQSVSDPGGLGGWRTAR...IGGSQNPWQRFRRAVFGIWRSAPSP | LTPTAFDPLAALVCPLRRGYEWWSGSSQSSLGSTVIFFGPKCVDSAPK | 24260 | 24611 | + | 6 |
| 6 | LWQSVSDPGGLGGWRTARTACAPAR...IGGSQNPWQRFRRAVFGIWRSAPSP | LTPTAFDPLAALVCPLRRGYEWWSGSSQSSLGSTVIFFGPKCVDSAPK | 24281 | 24611 | + | 6 |
| 7 | VSDPGGLGGWRTARTACAPARRVLP...IGGSQNPWQRFRRAVFGIWRSAPSP | LTPTAFDPLAALVCPLRRGYEWWSGSSQSSLGSTVIFFGPKCVDSAPK | 24293 | 24611 | + | 6 |
| 8 | MVVWLIAVISGFYSHLLRPEVCGLRAK | VICLPQSPVLSILTGCRLSVHRQIVQR | 24526 | 24691 | + | 7 |
| 9 | VVWLIAVISGFYSHLLRPEVCGLRAK | VICLPQSPVLSILTGCRLSVHRQIVQR | 24529 | 24691 | + | 7 |
| 10 | VALGGKSTAPRRRWVGPGGHPLAEHGGCSSPPSPVSCSPL | LRACSTSWSISLACAAGYGGAARGQGIPRRSRPGGAEISA | 27833 | 28076 | + | 6 |
| 11 | VQQPAFAGQLQSLATGLLDELVDQSRVRGRLRRRSSGSGHPTA | VTPGWRGDLRLRPGCLAGRLLWWLDGRRTESADE | 27913 | 28147 | + | 6 |
| 12 | LQSLATGLLDELVDQSRVRGRLRRRSSGSGHPTA | VTPGWRGDLRLRPGCLAGRLLWWLDGRRTESADE | 27940 | 28147 | + | 7 |
| 13 | VRGRLRRRSSGSGHPTA | VTPGWRGDLRLRPGCLAGRLLWWLDGRRTESADE | 27991 | 28147 | + | 6 |
| 14 | MVAGWTTNRISRRVTVAPTRLATGTGAQEPLSE | SLSAWRWAPRSATSPWESRLALRSSPYSPWPSA | 28104 | 28305 | + | 6 |
| 15 | VAGWTTNRISRRVTVAPTRLATGTGAQEPLSE | SLSAWRWAPRSATSPWESRLALRSSPYSPWPSA | 28107 | 28305 | + | 6 |
| 16 | MDDEPNQQTSDGGTDEARDWNWGPGAALGISLGVALGA | AFGNVAVGVALGVAFFPVFAMAFRLNGGDRRRTGQEPGR | 28115 | 28349 | + | 8 |
| 17 | VTVAPTRLATGTGAQEPLSESLSAWR | WAPRSATSPWESRLALRSSPYSPWPSA | 28143 | 28305 | + | 6 |
| 18 | VAPTRLATGTGAQEPLSESLSAWR | WAPRSATSPWESRLALRSSPYSPWPSA | 28149 | 28305 | + | 6 |
| 19 | LSSSPTVATVQAEGHGEYGEERNAKRDSHGDVAERGAQRHAERDSESG | SWAPVPVASLVGATVTRLLIRFVVHPATIEAGLRDNLASGGDLRATRA | 28334 | 28043 | - | 6 |
| 20 | VIAYCTYPPADSPVVG | IVPGAKAINGRGSATTWAFRARFSPAFVGIAAALVNYRRLTS | 29065 | 29242 | + | 9 |
| 21 | LSASYPGQRPSTGGDQPLHGHSVPGFPPPSSASPQPSS | TTADSLHDHTRISGDIACRSLAG | 29105 | 29291 | + | 6 |

### Results for SCK25365.1 [Streptomyces sp. WMMB 714] back to top

previous - next

#### Architecture

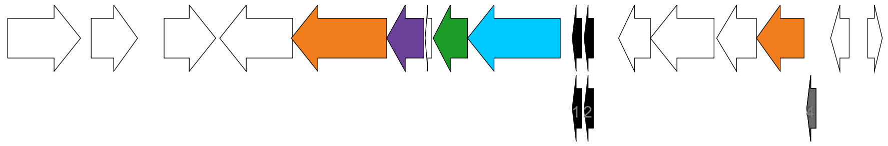

1000 nucleotides

Link to nucleotide sequence

| Accession | start | end | direction | length (aa) | Pfam/HMM | name | description | E-value |
|---|---|---|---|---|---|---|---|---|
| SCK25291.1 | 414898 | 416392 | + | 497 | NO PFAM MATCH | - | - | - |
| SCK25298.1 | 416625 | 417564 | + | 312 | PF01259<br>TIGR00081<br>TIGR02735 | SAICAR_synt<br>TIGR00081<br>TIGR02735 | SAICAR synthetase<br>purC: phosphoribosylaminoimidazolesuccinocarboxamide synthase<br>purC_vibrio: phosphoribosylaminoimidazole-succinocarboxamide synthase | 4.40E-97<br>1.00E-68<br>9.40E-20 |
| SCK25310.1 | 418119 | 419169 | + | 349 | PF00196 | GerE | Bacterial regulatory proteins, luxR family | 6.70E-05 |
| SCK25320.1 | 420729 | 419241 | - | 495 | TIGR02734<br>TIGR03467<br>TIGR02733<br>TIGR00292<br>PF13450 | TIGR02734<br>TIGR03467<br>TIGR02733<br>TIGR00292<br>NAD_binding_8 | crtI_fam: phytoene desaturase<br>HpnE: squalene-associated FAD-dependent desaturase<br>desat_CrtD: C-3',4' desaturase CrtD<br>TIGR00292: thiazole biosynthesis enzyme<br>NAD(P)-binding Rossmann-like domain | 2.90E-11<br>1.00E-08<br>7.00E-08<br>1.40E-07<br>2.10E-07 |
| SCK25330.1 | 422663 | 420725 | - | 645 | TIGR02204<br>TIGR02203<br>TIGR00958<br>TIGR01846<br>TIGR03796 | TIGR02204<br>TIGR02203<br>TIGR00958<br>TIGR01846<br>TIGR03796 | MsbA_rel: ABC transporter, permease/ATP-binding protein<br>MsbA_lipidA: lipid A export permease/ATP-binding protein MsbA<br>3a01208: antigen peptide transporter 2<br>type_I_sec_HlyB: type I secretion system ATPase<br>NHLM_micro_ABC1: NHLM bacteriocin system ABC transporter, peptidase/ATP-binding protein | 3.00E-105<br>3.30E-102<br>3.60E-90<br>3.30E-87<br>1.30E-86 |
| SCK25339.1 | 423429 | 422673 | - | 251 | PF13649<br>PF08241<br>PF13489<br>PF13847<br>TIGR01983 | Methyltransf_25<br>Methyltransf_11<br>Methyltransf_23<br>Methyltransf_31<br>TIGR01983 | Methyltransferase domain<br>Methyltransferase domain<br>Methyltransferase domain<br>Methyltransferase domain<br>UbiG: 3-demethylubiquinone-9 3-O-methyltransferase | 2.50E-18<br>3.90E-13<br>1.30E-12<br>5.70E-11<br>2.30E-10 |
| SCK25348.1 | 423584 | 423464 | - | 39 | NO PFAM MATCH | - | - | - |
| SCK25358.1 | 424329 | 423624 | - | 234 | LinL | LinL | LinL | 9.10E-66 |
| SCK25365.1 | 426233 | 424325 | - | 635 | LinH<br>LinG<br>LinE<br>PF12697<br>TIGR02427 | LinH<br>LinG<br>LinE<br>Abhydrolase_6<br>TIGR02427 | LinH (Fusion of LinE/LinG)<br>LinG Homolog<br>LinE Homolog<br>Alpha/beta hydrolase family<br>protocat_pcaD: 3-oxoadipate enol-lactonase | 8.00E-248<br>4.90E-119<br>1.80E-99<br>3.80E-15<br>7.00E-09 |
| SCK25375.1 | 426651 | 426480 | - | 56 | LinA | LinA | LinA (linaridin precursor) | 1.90E-15 |
| SCK25385.1 | 426905 | 426731 | - | 57 | LinA | LinA | LinA (linaridin precursor) | 1.10E-13 |
| SCK25394.1 | 428086 | 427414 | - | 223 | PF00072<br>TIGR02875<br>TIGR02154<br>PF00196<br>TIGR02956 | Response_reg<br>TIGR02875<br>TIGR02154<br>GerE<br>TIGR02956 | Response regulator receiver domain<br>spore_0_A: sporulation transcription factor Spo0A<br>PhoB: phosphate regulon transcriptional regulatory protein PhoB<br>Bacterial regulatory proteins, luxR family<br>TMAO_torS: TMAO reductase sytem sensor TorS | 3.10E-25<br>2.80E-14<br>1.50E-12<br>3.30E-12<br>8.30E-12 |
| SCK25402.1 | 429378 | 428082 | - | 431 | PF07730 | HisKA_3 | Histidine kinase | 5.50E-15 |
| SCK25413.1 | 430244 | 429428 | - | 271 | PF01061<br>PF12698<br>TIGR03062 | ABC2_membrane<br>ABC2_membrane_3<br>TIGR03062 | ABC-2 type transporter<br>ABC-2 family transporter protein<br>pip_yhgE_Cterm: YhgE/Pip C-terminal domain | 3.40E-18<br>8.50E-12<br>1.30E-04 |
| SCK25422.1 | 431227 | 430240 | - | 328 | TIGR01188<br>TIGR03864<br>TIGR01288<br>TIGR04406<br>TIGR03522 | TIGR01188<br>TIGR03864<br>TIGR01288<br>TIGR04406<br>TIGR03522 | drrA: daunorubicin resistance ABC transporter, ATP-binding protein<br>PQQ_ABC_ATP: ABC transporter, ATP-binding subunit, PQQ-dependent alcohol dehydrogenase system<br>nodI: nodulation ABC transporter NodI<br>LPS_export_lptB: LPS export ABC transporter ATP-binding protein<br>GldA_ABC_ATP: gliding motility-associated ABC transporter ATP-binding subunit GldA | 1.50E-69<br>2.70E-49<br>7.80E-48<br>7.50E-38<br>4.60E-37 |
| SCK25435.1 | 432142 | 431761 | - | 126 | PF11774 | Lsr2 | Lsr2 | 1.30E-34 |
| SCK25445.1 | 432534 | 432828 | + | 97 | PF02700<br>TIGR00302 | PurS<br>TIGR00302 | Phosphoribosylformylglycinamidine (FGAM) synthase<br>TIGR00302: phosphoribosylformylglycinamidine synthase, purS protein | 2.10E-27<br>6.10E-20 |

| index | leader | core | start | end | dir | score |
|---|---|---|---|---|---|---|
| 1 | MNDITSHINESAELAAVDAEGAIFADNTPVL | ATPGAFLGGVAIASATVAAFEAGQG | 426651 | 426480 | - | 29 |
| 2 | MLEAKLNLQIQDELHEGHDGEIAEEATPVL | ATPALAGAFVGGAAVVTGAYAAGKAVG | 426905 | 426731 | - | 30 |
| 3 | LRSRVRTSSSAPFRNRPRFPPGPIRLPGPCG | PVPLRSASTGFAVRPALPLCAKRMVLP | 431471 | 431294 | - | 6 |
| 4 | LPRLRSRVRTSSSAPFRNRPRFPPGPIRLPGPCG | PVPLRSASTGFAVRPALPLCAKRMVLP | 431480 | 431294 | - | 6 |

### Results for SCK35447.1 [Streptomyces sp. WMMB 714] back to top

previous - next

#### Architecture

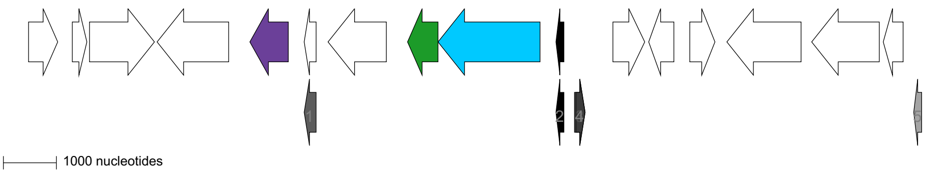

1000 nucleotides

Link to nucleotide sequence

| Accession | start | end | direction | length (aa) | Pfam/HMM | name | description | E-value |
|---|---|---|---|---|---|---|---|---|

| | | | | | | | | |
|---|---|---|---|---|---|---|---|---|
| SCK35373.1 | 67491 | 68028 | + | 178 | NO PFAM MATCH | - | - | - |
| SCK35383.1 | 68306 | 68567 | + | 86 | NO PFAM MATCH | - | - | - |
| SCK35392.1 | 68625 | 69867 | + | 413 | PF13378<br>PF02746<br>TIGR02534<br>TIGR01928 | MR_MLE_C<br>MR_MLE_N<br>TIGR02534<br>TIGR01928 | Enolase C-terminal domain-like<br>Mandelate racemase / muconate lactonizing enzyme, N-terminal domain<br>mucon_cyclo: muconate and chloromuconate cycloisomerases<br>menC_lowGC/arch: o-succinylbenzoate synthase | 1.50E-44<br>8.50E-33<br>9.40E-24<br>1.20E-10 |
| SCK35401.1 | 71257 | 69904 | - | 450 | PF01547<br>PF13416<br>TIGR03850 | SBP_bac_1<br>SBP_bac_8<br>TIGR03850 | Bacterial extracellular solute-binding protein<br>Bacterial extracellular solute-binding protein<br>bind_CPR_0540: carbohydrate ABC transporter substrate-binding protein, CPR_0540 family | 3.30E-42<br>1.60E-21<br>1.20E-06 |
| SCK35409.1 | 72371 | 71657 | - | 237 | PF13649<br>TIGR00755<br>PF08242<br>PF08241<br>PF13489 | Methyltransf_25<br>TIGR00755<br>Methyltransf_12<br>Methyltransf_11<br>Methyltransf_23 | Methyltransferase domain<br>ksgA: ribosomal RNA small subunit methyltransferase A<br>Methyltransferase domain<br>Methyltransferase domain<br>Methyltransferase domain | 9.20E-08<br>1.50E-06<br>1.80E-05<br>3.20E-05<br>5.90E-05 |
| SCK35417.1 | 72903 | 72678 | - | 74 | NO PFAM MATCH | - | - | - |
| SCK35428.1 | 74239 | 73132 | - | 368 | PF07228 | SpoIIE | Stage II sporulation protein E (SpoIIE) | 1.10E-40 |
| SCK35439.1 | 75205 | 74626 | - | 192 | LinL | LinL | LinL | 6.00E-64 |
| SCK35447.1 | 77124 | 75201 | - | 640 | LinH<br>LinG<br>LinE<br>PF12697<br>LinL | LinH<br>LinG<br>LinE<br>Abhydrolase_6<br>LinL | LinH (Fusion of LinE/LinG)<br>LinG Homolog<br>LinE Homolog<br>Alpha/beta hydrolase family<br>LinL | 8.30E-238<br>8.20E-120<br>3.30E-90<br>3.40E-11<br>1.10E-08 |
| SCK35457.1 | 77592 | 77439 | - | 50 | LinA | LinA | LinA (linaridin precursor) | 1.10E-09 |
| SCK35466.1 | 78507 | 79116 | + | 202 | PF00072<br>TIGR02875<br>PF00196<br>TIGR01818<br>TIGR02154 | Response_reg<br>TIGR02875<br>GerE<br>TIGR01818<br>TIGR02154 | Response regulator receiver domain<br>spore_0_A: sporulation transcription factor Spo0A<br>Bacterial regulatory proteins, luxR family<br>ntrC: nitrogen regulation protein NR(I)<br>PhoB: phosphate regulon transcriptional regulatory protein PhoB | 6.60E-26<br>1.10E-17<br>2.30E-13<br>2.50E-10<br>4.00E-10 |
| SCK35475.1 | 79658 | 79190 | - | 155 | NO PFAM MATCH | - | - | - |
| SCK35484.1 | 79964 | 80435 | + | 156 | NO PFAM MATCH | - | - | - |
| SCK35494.1 | 82064 | 80666 | - | 465 | TIGR00720<br>PF03313<br>PF03315<br>TIGR00718<br>TIGR00719 | TIGR00720<br>SDH_alpha<br>SDH_beta<br>TIGR00718<br>TIGR00719 | sda_mono: L-serine ammonia-lyase<br>Serine dehydratase alpha chain<br>Serine dehydratase beta chain<br>sda_alpha: L-serine dehydratase, iron-sulfur-dependent, alpha subunit<br>sda_beta: L-serine dehydratase, iron-sulfur-dependent, beta subunit | 4.10E-204<br>3.50E-102<br>1.20E-57<br>3.40E-56<br>2.80E-09 |
| SCK35502.1 | 83534 | 82259 | - | 424 | PF00464<br>TIGR04506<br>PF00155<br>TIGR00858<br>PF01212 | SHMT<br>TIGR04506<br>Aminotran_1_2<br>TIGR00858<br>Beta_elim_lyase | Serine hydroxymethyltransferase<br>F_threo_transal: fluorothreonine transaldolase<br>Aminotransferase class I and II<br>bioF: 8-amino-7-oxononanoate synthase<br>Beta-eliminating lyase | 2.10E-161<br>1.20E-30<br>2.40E-10<br>7.00E-08<br>1.60E-07 |
| SCK35512.1 | 84001 | 83620 | - | 126 | TIGR00527<br>PF01597<br>TIGR03077<br>PF00364<br>PF13375 | TIGR00527<br>GCV_H<br>TIGR03077<br>Biotin_lipoyl<br>RnfC_N | gcvH: glycine cleavage system H protein<br>Glycine cleavage H-protein<br>not_gcvH: glycine cleavage protein H-like protein<br>Biotin-requiring enzyme<br>RnfC Barrel sandwich hybrid domain | 5.70E-47<br>2.30E-42<br>2.20E-21<br>5.40E-05<br>8.80E-04 |

| index | leader | core | start | end | dir | score |
|---|---|---|---|---|---|---|
| 1 | MLGLGAGEAAGTAVPPGRDELIAYGPLLSDDGGTWVGTAALVGAPDPAA | APAVLTADAYAEAGVHSWQFGGRPS | 72903 | 72678 | - | 8 |
| 2 | MREIIVKAEPKLSESDKGQVVVPQGPVL | ATPVAAATVAFATGVAVTKAVG | 77592 | 77439 | - | 31 |
| 3 | LSDVGHGPRMTGVRACHESTHALSMGFEVVETGPERFAERNVTD | ATPSGVPPVSICGGAFSA | 77788 | 77977 | + | 8 |
| 4 | MTGVRACHESTHALSMGFEVVETGPERFAERNVTD | ATPSGVPPVSICGGAFSA | 77815 | 77977 | + | 6 |
| 5 | VTSGAPSPTLGKPIAIAYVDAAHAEPG | TEGVAVDIRGKHEPYEVVALPFYKRQR | 84344 | 84179 | - | 6 |

### Results for SCK51308.1 [Streptomyces sp. WMMB 714] back to top

previous - next

#### Architecture

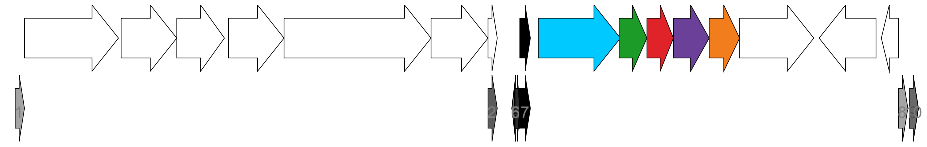

1000 nucleotides

Link to nucleotide sequence

| Accession | start | end | direction | length (aa) | Pfam/HMM | name | description | E-value |
|---|---|---|---|---|---|---|---|---|
| SCK51234.1 | 34753 | 36715 | + | 653 | PF00326<br>PF00135<br>PF12146<br>PF01738 | Peptidase_S9<br>COesterase<br>Hydrolase_4<br>DLH | Prolyl oligopeptidase family<br>Carboxylesterase family<br>Serine aminopeptidase, S33<br>Dienelactone hydrolase family | 3.20E-39<br>3.50E-05<br>5.30E-04<br>8.90E-04 |
| SCK51243.1 | 36773 | 37940 | + | 388 | PF00487 | FA_desaturase | Fatty acid desaturase | 1.90E-25 |
| SCK51252.1 | 37936 | 38914 | + | 325 | PF01619 | Pro_dh | Proline dehydrogenase | 5.10E-09 |

| | | | | | | | | |
|---|---|---|---|---|---|---|---|---|
| SCK51262.1 | 39002 | 40157 | + | 384 | PF00266<br>TIGR04343<br>TIGR01979<br>TIGR01977<br>TIGR01976 | Aminotran_5<br>TIGR04343<br>TIGR01979<br>TIGR01977<br>TIGR01976 | Aminotransferase class-V<br>egtE_PLP_lyase: ergothioneine biosynthesis PLP-dependent enzyme EgtE<br>sufS: cysteine desulfurase, SufS family<br>am_tr_V_EF2568: cysteine desulfurase family protein<br>am_tr_V_VC1184: cysteine desulfurase family protein | 5.60E-43<br>5.40E-30<br>1.10E-27<br>7.40E-24<br>1.80E-20 |
| SCK51270.1 | 40153 | 43213 | + | 1019 | TIGR01733<br>PF00501<br>TIGR01734<br>PF00668<br>TIGR03443 | TIGR01733<br>AMP-binding<br>TIGR01734<br>Condensation<br>TIGR03443 | AA-adenyl-dom: amino acid adenylation domain<br>AMP-binding enzyme<br>D-ala-DACP-lig: D-alanine--poly(phosphoribitol) ligase, subunit 1<br>Condensation domain<br>alpha_am_amid: L-aminoadipate-semialdehyde dehydrogenase | 5.50E-90<br>1.00E-70<br>5.80E-47<br>1.20E-40<br>4.10E-37 |
| SCK51282.1 | 43209 | 44412 | + | 400 | TIGR04458<br>PF00067 | TIGR04458<br>p450 | CYP450_TxtE: 4-nitrotryptophan synthase<br>Cytochrome P450 | 2.50E-34<br>1.70E-23 |
| SCK51291.1 | 44408 | 44597 | + | 62 | NO PFAM MATCH | - | - | - |
| SCK51299.1 | 45076 | 45274 | + | 65 | LinA | LinA | LinA (linaridin precursor) | 1.30E-13 |
| SCK51308.1 | 45453 | 47145 | + | 563 | LinH<br>LinG<br>LinE<br>PF12697<br>TIGR02427 | LinH<br>LinG<br>LinE<br>Abhydrolase_6<br>TIGR02427 | LinH (Fusion of LinE/LinG)<br>LinG Homolog<br>LinE Homolog<br>Alpha/beta hydrolase family<br>protocat_pcaD: 3-oxoadipate enol-lactonase | 1.80E-171<br>2.50E-75<br>4.20E-65<br>5.40E-07<br>9.50E-07 |
| SCK51315.1 | 47134 | 47698 | + | 187 | LinL | LinL | LinL | 7.70E-55 |
| SCK51324.1 | 47694 | 48267 | + | 190 | TIGR00521<br>TIGR02113<br>PF02441 | TIGR00521<br>TIGR02113<br>Flavoprotein | coaBC_dfp: phosphopantothenoylcysteine decarboxylase / phosphopantothenate--cysteine ligase<br>coaC_strep: phosphopantothenoylcysteine decarboxylase<br>Flavoprotein | 1.30E-13<br>1.10E-10<br>7.60E-10 |
| SCK51331.1 | 48259 | 48997 | + | 245 | PF13649<br>PF08241<br>TIGR02072<br>TIGR02021<br>PF08242 | Methyltransf_25<br>Methyltransf_11<br>TIGR02072<br>TIGR02021<br>Methyltransf_12 | Methyltransferase domain<br>Methyltransferase domain<br>BioC: malonyl-acyl carrier protein O-methyltransferase BioC<br>BchM-ChlM: magnesium protoporphyrin O-methyltransferase<br>Methyltransferase domain | 7.20E-16<br>1.00E-14<br>7.30E-10<br>8.00E-10<br>1.00E-09 |
| SCK51340.1 | 48993 | 49626 | + | 210 | TIGR02857<br>TIGR01189<br>TIGR02673<br>TIGR02982<br>TIGR03608 | TIGR02857<br>TIGR01189<br>TIGR02673<br>TIGR02982<br>TIGR03608 | CydD: thiol reductant ABC exporter, CydD subunit<br>ccmA: heme ABC exporter, ATP-binding protein CcmA<br>FtsE: cell division ATP-binding protein FtsE<br>heterocyst_DevA: ABC exporter ATP-binding subunit, DevA family<br>L_ocin_972_ABC: putative bacteriocin export ABC transporter, lactococcin 972 group | 1.80E-27<br>6.10E-26<br>3.80E-24<br>1.20E-22<br>2.30E-21 |
| SCK51349.1 | 49622 | 51167 | + | 514 | NO PFAM MATCH | - | - | - |
| SCK51357.1 | 52485 | 51282 | - | 400 | PF06889 | DUF1266 | Protein of unknown function (DUF1266) | 1.20E-43 |
| SCK51363.1 | 52930 | 52588 | - | 113 | PF03795 | YCII | YCII-related domain | 2.40E-07 |

| index | leader | core | start | end | dir | score |
|---|---|---|---|---|---|---|
| 1 | MLGCFLEPPRAIAYAAQLAGLADWSDLDGHF | WVSDDPAVPGYALDSSEPGIPRISYAKDGPA | 34568 | 34757 | + | 6 |
| 2 | MNLYALMRLAAPPAQAARPARKRTGAATARTKAWSFGRRAGP | GTAAEPAPGTGAGTESSDAA | 44408 | 44597 | + | 8 |
| 3 | LPTLQGTFNSERATGEG | QHEVSTETLSLGMSNEKG | 44964 | 45072 | + | 6 |
| 4 | LQGTFNSERATGEGQH | EVSTETLSLGMSNEKG | 44973 | 45072 | + | 6 |
| 5 | MPSERVSVDTSCCPSPVARSELNVPCN | VGNSHCGSRVTSGARERQRSTEPFSGA | 45053 | 44888 | - | 6 |
| 6 | LLMPSERVSVDTSCCPSPVARSELNVPC | NVGNSHCGSRVTSGARERQRSTEPFSGA | 45059 | 44888 | - | 6 |
| 7 | MRVNVMTDPDTVSPESVATQEFANSVLAGAAPGFYSDSDVPAM | ATPATPTAAQFVVQGSTICLVC | 45076 | 45274 | + | 35 |
| 8 | LSRLLPRRPHPACATEAGSAPGRLSASLAAGAKGALNDAAEAAS | VLPADPRSPGIQGVREIRGSS | 52941 | 53139 | + | 6 |
| 9 | VLHRYRPDDASLTRQPRSRGLKSGGPFRNGSTVTAVGPAH | VTPRAERRPSRRYSFRSRRRRTSRK | 53158 | 53356 | + | 6 |
| 10 | LHRYRPDDASLTRQPRSRGLKSGGPFRNGSTVTAVGPAH | VTPRAERRPSRRYSFRSRRRRTSRK | 53161 | 53356 | + | 6 |

### Results for WP_045865779.1 [Streptomyces sp. WMMB 714] back to top

previous - next

#### Architecture

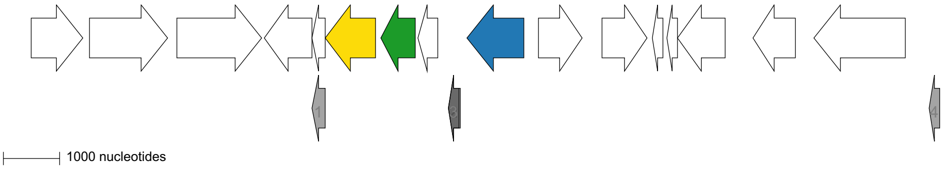

Link to nucleotide sequence

| Accession | start | end | direction | length (aa) | Pfam/HMM | name | description | E-value |
|---|---|---|---|---|---|---|---|---|
| WP_045865774.1 | 147255 | 148179 | + | 307 | TIGR03383<br>PF01014 | TIGR03383<br>Uricase | urate_oxi: urate oxidase<br>Uricase | 5.60E-106<br>5.90E-52 |
| WP_045865775.1 | 148286 | 149681 | + | 464 | PF01979<br>TIGR02967<br>TIGR03314<br>TIGR02022<br>PF07969 | Amidohydro_1<br>TIGR02967<br>TIGR03314<br>TIGR02022<br>Amidohydro_3 | Amidohydrolase family<br>guan_deamin: guanine deaminase<br>Se_ssnA: putative selenium metabolism protein SsnA<br>hutF: formiminoglutamate deiminase<br>Amidohydrolase family | 2.70E-62<br>3.40E-46<br>4.60E-42<br>3.60E-36<br>2.40E-14 |

| | | | | | | | | |
|---|---|---|---|---|---|---|---|---|
| WP_078895834.1 | 149841 | 151350 | + | 502 | TIGR03173<br>TIGR00801<br>PF00860<br>TIGR03616 | TIGR03173<br>TIGR00801<br>Xan_ur_permease<br>TIGR03616 | pbuX: xanthine permease<br>ncs2: uracil-xanthine permease<br>Permease family<br>RutG: pyrimidine utilization transport protein G | 4.20E-156<br>8.40E-108<br>3.60E-93<br>5.90E-38 |
| WP_045865777.1 | 152250 | 151404 | - | 281 | PF00144 | Beta-lactamase | Beta-lactamase | 7.80E-37 |
| WP_139131725.1 | 152498 | 152246 | - | 83 | NO PFAM MATCH | - | - | - |
| WP_107068805.1 | 153388 | 152494 | - | 297 | TIGR04033<br>LinG<br>LinH<br>PF05090 | TIGR04033<br>LinG<br>LinH<br>VKG_Carbox | export_SdpB: antimicrobial peptide system protein, SdpB family<br>LinG Homolog<br>LinH (Fusion of LinE/LinG)<br>Vitamin K-dependent gamma-carboxylase | 6.50E-22<br>1.10E-20<br>3.30E-15<br>9.10E-14 |
| WP_139131726.1 | 154098 | 153486 | - | 203 | LinL | LinL | LinL | 9.90E-09 |
| WP_139131727.1 | 154478 | 154124 | - | 117 | NO PFAM MATCH | - | - | - |
| WP_045865779.1 | 156011 | 155003 | - | 335 | PF12697<br>LinE<br>LinH<br>PF12146<br>TIGR03695 | Abhydrolase_6<br>LinE<br>LinH<br>Hydrolase_4<br>TIGR03695 | Alpha/beta hydrolase family<br>LinE Homolog<br>LinH (Fusion of LinE/LinG)<br>Serine aminopeptidase, S33<br>menH_SHCHC: 2-succinyl-6-hydroxy-2,4-cyclohexadiene-1-carboxylate synthase | 1.90E-16<br>1.20E-10<br>4.60E-10<br>4.50E-07<br>1.50E-05 |
| WP_078895836.1 | 156279 | 157044 | + | 254 | PF06475 | Glycolipid_bind | Putative glycolipid-binding | 5.10E-47 |
| WP_139131728.1 | 157402 | 158203 | + | 266 | PF14016 | DUF4232 | Protein of unknown function (DUF4232) | 1.00E-10 |
| WP_045865781.1 | 158494 | 158302 | - | 63 | PF04149 | DUF397 | Domain of unknown function (DUF397) | 2.20E-21 |
| WP_045865782.1 | 158754 | 158562 | - | 63 | PF04149 | DUF397 | Domain of unknown function (DUF397) | 7.30E-21 |
| WP_045865783.1 | 159596 | 158750 | - | 281 | PF13560<br>TIGR03070<br>PF01381<br>PF12844 | HTH_31<br>TIGR03070<br>HTH_3<br>HTH_19 | Helix-turn-helix domain<br>couple_hipB: transcriptional regulator, y4mF family<br>Helix-turn-helix<br>Helix-turn-helix domain | 5.20E-14<br>1.80E-06<br>2.40E-06<br>5.30E-05 |
| WP_045865784.1 | 160835 | 160088 | - | 248 | NO PFAM MATCH | - | - | - |
| WP_045865785.1 | 162793 | 161167 | - | 541 | TIGR01344<br>PF01274<br>TIGR01345 | TIGR01344<br>Malate_synthase<br>TIGR01345 | malate_syn_A: malate synthase A<br>Malate synthase<br>malate_syn_G: malate synthase G | 2.90E-251<br>9.80E-201<br>9.60E-04 |

| index | leader | core | start | end | dir | score |
|---|---|---|---|---|---|---|
| 1 | MMVLAEFLTNFFPRRPGSFREYVIS...WVAVARFYGAGPDTVVHVSWVLAAF | ETAAYAVSWFVFRHRAGGEGEA | 152498 | 152246 | - | 6 |
| 2 | MTPPGLDLSCNERHRITDPRHIEPAHGATGIGTRSSPSECCRLSRPLQ | VTAGKRDFPRPGTCVKIFS | 154871 | 154667 | - | 6 |
| 3 | VHEEDPEMTPPGLDLSCNERHRITD...PAHGATGIGTRSSPSECCRLSRPLQ | VTAGKRDFPRPGTCVKIFS | 154892 | 154667 | - | 6 |
| 4 | VLFGSGHWQGVTDRAEGDQGARGYLRERAGEIELVECGDIADPADI | DTPADLERLRAYGS | 163403 | 163220 | - | 6 |

### Results for PWS41421.1 [Streptomyces sp. ZEA17I] back to top

previous - next

#### Architecture

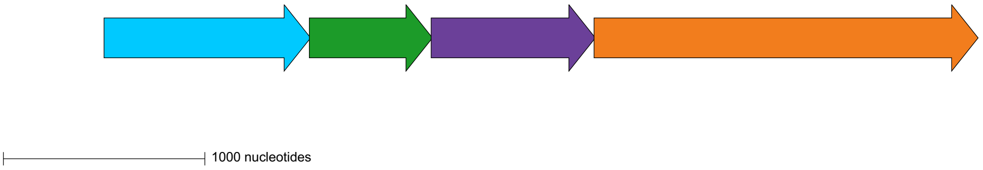

Link to nucleotide sequence

| Accession | start | end | direction | length (aa) | Pfam/HMM | name | description | E-value |
|---|---|---|---|---|---|---|---|---|
| PWS41421.1 | 0 | 1027 | + | 341 | LinH<br>LinE<br>LinL<br>PF12697<br>PF00561 | LinH<br>LinE<br>LinL<br>Abhydrolase_6<br>Abhydrolase_1 | LinH (Fusion of LinE/LinG)<br>LinE Homolog<br>LinL<br>Alpha/beta hydrolase family<br>alpha/beta hydrolase fold | 2.80E-120<br>1.10E-94<br>6.20E-46<br>1.10E-14<br>7.00E-12 |
| PWS41422.1 | 1023 | 1632 | + | 202 | LinL | LinL | LinL | 4.10E-66 |
| PWS41423.1 | 1628 | 2441 | + | 270 | PF13649<br>PF08241<br>PF13847<br>PF13489<br>PF08242 | Methyltransf_25<br>Methyltransf_11<br>Methyltransf_31<br>Methyltransf_23<br>Methyltransf_12 | Methyltransferase domain<br>Methyltransferase domain<br>Methyltransferase domain<br>Methyltransferase domain<br>Methyltransferase domain | 2.30E-16<br>2.80E-12<br>2.60E-07<br>8.20E-07<br>8.70E-07 |
| PWS41424.1 | 2437 | 4339 | + | 633 | TIGR02204<br>TIGR02203<br>TIGR00958<br>TIGR03796<br>TIGR03797 | TIGR02204<br>TIGR02203<br>TIGR00958<br>TIGR03796<br>TIGR03797 | MsbA_rel: ABC transporter, permease/ATP-binding protein<br>MsbA_lipidA: lipid A export permease/ATP-binding protein MsbA<br>3a01208: antigen peptide transporter 2<br>NHLM_micro_ABC1: NHLM bacteriocin system ABC transporter, peptidase/ATP-binding protein<br>NHLM_micro_ABC2: NHLM bacteriocin system ABC transporter, ATP-binding protein | 2.70E-99<br>9.00E-92<br>2.80E-83<br>2.30E-81<br>1.70E-77 |

| index | leader | core | start | end | dir | score |
|---|---|---|---|---|---|---|

### Results for WP_165339856.1 [Streptomyces sp. YC419] back to top

previous - next

#### Architecture

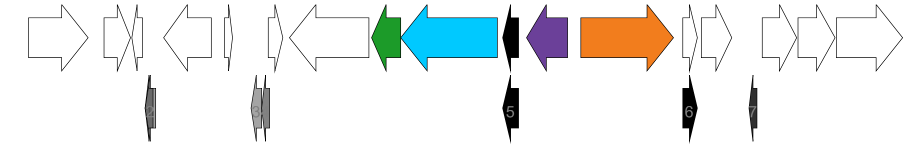

1000 nucleotides

Link to nucleotide sequence

| Accession | start | end | direction | length (aa) | Pfam/HMM | name | description | E-value |
|---|---|---|---|---|---|---|---|---|
| WP_165339851.1 | 30433 | 31573 | + | 379 | PF13191<br>TIGR03903 | AAA_16<br>TIGR03903 | AAA ATPase domain<br>TOMM_kin_cyc: TOMM system kinase/cyclase fusion protein | 7.30E-18<br>8.90E-16 |
| WP_165339852.1 | 31877 | 32405 | + | 175 | NO PFAM MATCH | - | - | - |
| WP_165339853.1 | 32623 | 32422 | - | 66 | PF13683<br>PF13333 | rve_3<br>rve_2 | Integrase core domain<br>Integrase core domain | 4.20E-07<br>5.30E-05 |
| WP_165339854.1 | 33950 | 33026 | - | 307 | PF07730 | HisKA_3 | Histidine kinase | 5.20E-09 |
| WP_014676472.1 | 34201 | 34351 | + | 50 | INFERRED GENE | - | - | - |
| WP_013226009.1 | 35033 | 35328 | + | 98 | INFERRED GENE | - | - | - |
| WP_165339855.1 | 36970 | 35461 | - | 502 | TIGR02734<br>TIGR03467<br>PF01593<br>TIGR02733<br>PF13450 | TIGR02734<br>TIGR03467<br>Amino_oxidase<br>TIGR02733<br>NAD_binding_8 | crtI_fam: phytoene desaturase<br>HpnE: squalene-associated FAD-dependent desaturase<br>Flavin containing amine oxidoreductase<br>desat_CrtD: C-3',4' desaturase CrtD<br>NAD(P)-binding Rossmann-like domain | 3.80E-18<br>3.60E-11<br>1.60E-10<br>1.20E-09<br>2.00E-09 |
| WP_165339872.1 | 37583 | 37031 | - | 183 | LinL | LinL | LinL | 3.00E-54 |
| WP_165339856.1 | 39459 | 37605 | - | 617 | LinH<br>LinE<br>LinG<br>PF00561<br>TIGR02427 | LinH<br>LinE<br>LinG<br>Abhydrolase_1<br>TIGR02427 | LinH (Fusion of LinE/LinG)<br>LinE Homolog<br>LinG Homolog<br>alpha/beta hydrolase fold<br>protocat_pcaD: 3-oxoadipate enol-lactonase | 5.40E-170<br>5.30E-90<br>4.60E-89<br>4.40E-14<br>3.20E-09 |
| WP_165339857.1 | 39863 | 39563 | - | 99 | LinA | LinA | LinA (linaridin precursor) | 1.30E-04 |
| WP_165339873.1 | 40806 | 40020 | - | 261 | PF13649<br>PF08241<br>PF08242<br>PF13489<br>PF13847 | Methyltransf_25<br>Methyltransf_11<br>Methyltransf_12<br>Methyltransf_23<br>Methyltransf_31 | Methyltransferase domain<br>Methyltransferase domain<br>Methyltransferase domain<br>Methyltransferase domain<br>Methyltransferase domain | 4.40E-13<br>1.10E-10<br>2.10E-10<br>1.10E-09<br>1.70E-08 |
| WP_165339874.1 | 41066 | 42845 | + | 592 | TIGR02204<br>TIGR02203<br>TIGR00958<br>TIGR01846<br>TIGR03375 | TIGR02204<br>TIGR02203<br>TIGR00958<br>TIGR01846<br>TIGR03375 | MsbA_rel: ABC transporter, permease/ATP-binding protein<br>MsbA_lipidA: lipid A export permease/ATP-binding protein MsbA<br>3a01208: antigen peptide transporter 2<br>type_I_sec_HlyB: type I secretion system ATPase<br>type_I_sec_LssB: type I secretion system ATPase | 3.50E-110<br>2.00E-101<br>6.50E-97<br>7.00E-87<br>8.90E-83 |
| WP_165339858.1 | 43025 | 43310 | + | 94 | NO PFAM MATCH | - | - | - |
| WP_014150745.1 | 43382 | 43975 | + | 197 | INFERRED GENE | - | - | - |
| WP_165339875.1 | 44541 | 45219 | + | 225 | TIGR02154<br>TIGR01387<br>PF00072<br>TIGR03787<br>TIGR01818 | TIGR02154<br>TIGR01387<br>Response_reg<br>TIGR03787<br>TIGR01818 | PhoB: phosphate regulon transcriptional regulatory protein PhoB<br>cztR_silR_copR: heavy metal response regulator<br>Response regulator receiver domain<br>marine_sort_RR: proteobacterial dedicated sortase system response regulator<br>ntrC: nitrogen regulation protein NR(I) | 3.20E-35<br>6.30E-33<br>7.40E-23<br>1.90E-16<br>2.70E-11 |
| WP_165339833.1 | 45249 | 45951 | + | 233 | PF03358<br>TIGR03567 | FMN_red<br>TIGR03567 | NADPH-dependent FMN reductase<br>FMN_reduc_SsuE: FMN reductase | 1.00E-12<br>5.70E-04 |
| WP_165339859.1 | 45975 | 47256 | + | 426 | PF00174 | Oxidored_molyb | Oxidoreductase molybdopterin binding domain | 5.00E-24 |

| index | leader | core | start | end | dir | score |
|---|---|---|---|---|---|---|
| 1 | VQAARRWGRCWRTEVTRSGAAVGRRC | NRGASCGHCRGPRSATVTVAPSSWSF | 32826 | 32667 | - | 6 |
| 2 | LKVPRPSMTLETKVQAARRWGRCWRTEVTRSG | AAVGRRCNRGASCGHCRGPRSATVTVAPSSWSF | 32865 | 32667 | - | 6 |
| 3 | MRRCSRPSSTFWAAAVHRRRRGALSA | RTTSWSPRTSCTPNWPPGNSWQPQGEGVSRGPGAGGCVRLV | 34920 | 34716 | - | 6 |
| 4 | VRRTLTQHGAGNVELDCSARAV | GDREAADPAVEGAVAGRRNTGHA | 35057 | 34919 | - | 7 |
| 5 | MSLMTSTHVAEWANRNAAAGEQSSVWAGGREVTPVL | ATPAAAAAAQAARVLVATQAAGTLA...APEAALTGRTAPEGASFSALMDAVG | 39863 | 39563 | - | 29 |
| 6 | MSMITTSRAAEWANHNATGEEHSTWHSAVAAPVL | GTPNVITPNVIDLEATNRAIIDEAF...PEASTVPGRVAPQGSSFSALMATYA | 43025 | 43310 | + | 16 |
| 7 | LSSGHPGESPGFTALRKPNGCKSLLDE | YGKVQEYFKVRDSLSRIVEYFHCVTNG | 44459 | 44294 | - | 9 |

### Results for RII13759.1 [Streptomyces sp. YIM 130001] back to top

previous - next

#### Architecture

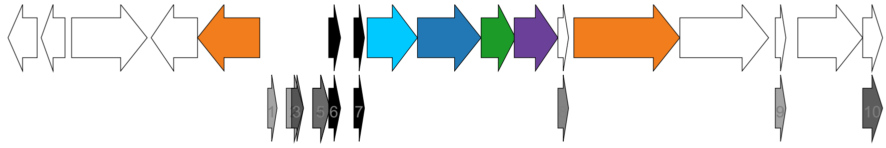

1000 nucleotides

Link to nucleotide sequence

| Accession | start | end | direction | length (aa) | Pfam/HMM | name | description | E-value |
|---|---|---|---|---|---|---|---|---|
| RII13751.1 | 62661 | 62181 | - | 159 | PF00583<br>PF13508<br>PF13673<br>TIGR01575<br>PF08445 | Acetyltransf_1<br>Acetyltransf_7<br>Acetyltransf_10<br>TIGR01575<br>FR47 | Acetyltransferase (GNAT) family<br>Acetyltransferase (GNAT) domain<br>Acetyltransferase (GNAT) domain<br>rimI: ribosomal-protein-alanine acetyltransferase<br>FR47-like protein | 8.00E-15<br>1.00E-08<br>5.70E-06<br>1.20E-05<br>2.00E-04 |
| RII13752.1 | 63117 | 62724 | - | 130 | NO PFAM MATCH | - | - | - |
| RII13753.1 | 63218 | 64472 | + | 417 | PF13560<br>PF01381<br>TIGR03070 | HTH_31<br>HTH_3<br>TIGR03070 | Helix-turn-helix domain<br>Helix-turn-helix<br>couple_hipB: transcriptional regulator, y4mF family | 7.80E-09<br>7.80E-07<br>3.70E-05 |
| RII13754.1 | 65307 | 64539 | - | 255 | PF01061<br>PF12698<br>TIGR03062<br>TIGR01247 | ABC2_membrane<br>ABC2_membrane_3<br>TIGR03062<br>TIGR01247 | ABC-2 type transporter<br>ABC-2 family transporter protein<br>pip_yhgE_Cterm: YhgE/Pip C-terminal domain<br>drrB: daunorubicin resistance ABC transporter membrane protein | 5.80E-20<br>3.10E-10<br>7.80E-07<br>8.10E-06 |
| RII13755.1 | 66326 | 65303 | - | 340 | TIGR01188<br>TIGR03864<br>TIGR01288<br>TIGR04406<br>TIGR02673 | TIGR01188<br>TIGR03864<br>TIGR01288<br>TIGR04406<br>TIGR02673 | drrA: daunorubicin resistance ABC transporter, ATP-binding protein<br>PQQ_ABC_ATP: ABC transporter, ATP-binding subunit, PQQ-dependent alcohol dehydrogenase system<br>nodI: nodulation ABC transporter NodI<br>LPS_export_lptB: LPS export ABC transporter ATP-binding protein<br>FtsE: cell division ATP-binding protein FtsE | 2.60E-61<br>2.30E-46<br>4.30E-42<br>9.20E-36<br>3.40E-33 |
| RII13756.1 | 67465 | 67666 | + | 66 | LinA | LinA | LinA (linaridin precursor) | 7.30E-09 |
| RII13757.1 | 67874 | 68057 | + | 60 | LinA | LinA | LinA (linaridin precursor) | 7.30E-11 |
| RII13758.1 | 68096 | 68927 | + | 276 | LinH<br>LinG | LinH<br>LinG | LinH (Fusion of LinE/LinG)<br>LinG Homolog | 2.50E-101<br>1.60E-100 |
| RII13759.1 | 68923 | 69979 | + | 351 | LinE<br>LinH<br>PF12697<br>TIGR03611<br>PF00561 | LinE<br>LinH<br>Abhydrolase_6<br>TIGR03611<br>Abhydrolase_1 | LinE Homolog<br>LinH (Fusion of LinE/LinG)<br>Alpha/beta hydrolase family<br>RutD: pyrimidine utilization protein D<br>alpha/beta hydrolase fold | 3.00E-99<br>2.00E-69<br>1.40E-12<br>7.30E-08<br>8.40E-08 |
| RII13760.1 | 69975 | 70536 | + | 186 | LinL | LinL | LinL | 2.60E-58 |
| RII13761.1 | 70532 | 71252 | + | 239 | PF13649<br>PF08241<br>PF13847<br>PF08242<br>PF13489 | Methyltransf_25<br>Methyltransf_11<br>Methyltransf_31<br>Methyltransf_12<br>Methyltransf_23 | Methyltransferase domain<br>Methyltransferase domain<br>Methyltransferase domain<br>Methyltransferase domain<br>Methyltransferase domain | 2.60E-15<br>3.40E-11<br>3.20E-09<br>2.40E-08<br>5.60E-08 |
| RII13762.1 | 71254 | 71428 | + | 57 | NO PFAM MATCH | - | - | - |
| RII13763.1 | 71501 | 73256 | + | 584 | TIGR02204<br>TIGR02203<br>TIGR03796<br>TIGR01846<br>TIGR00958 | TIGR02204<br>TIGR02203<br>TIGR03796<br>TIGR01846<br>TIGR00958 | MsbA_rel: ABC transporter, permease/ATP-binding protein<br>MsbA_lipidA: lipid A export permease/ATP-binding protein MsbA<br>NHLM_micro_ABC1: NHLM bacteriocin system ABC transporter, peptidase/ATP-binding protein<br>type_I_sec_HlyB: type I secretion system ATPase<br>3a01208: antigen peptide transporter 2 | 3.20E-107<br>1.10E-102<br>1.80E-88<br>4.60E-88<br>6.00E-88 |
| RII13764.1 | 73252 | 74719 | + | 488 | TIGR02734<br>TIGR03467<br>PF13450<br>TIGR01317<br>TIGR00292 | TIGR02734<br>TIGR03467<br>NAD_binding_8<br>TIGR01317<br>TIGR00292 | crtI_fam: phytoene desaturase<br>HpnE: squalene-associated FAD-dependent desaturase<br>NAD(P)-binding Rossmann-like domain<br>GOGAT_sm_gam: glutamate synthase, NADH/NADPH, small subunit<br>TIGR00292: thiazole biosynthesis enzyme | 5.30E-11<br>9.50E-09<br>1.60E-08<br>5.60E-07<br>1.30E-06 |
| RII13765.1 | 74825 | 74999 | + | 57 | NO PFAM MATCH | - | - | - |
| RII13766.1 | 75213 | 76284 | + | 356 | TIGR00747<br>PF08541 | TIGR00747<br>ACP_syn_III_C | fabH: 3-oxoacyl-[acyl-carrier-protein] synthase III<br>3-Oxoacyl-[acyl-carrier-protein (ACP)] synthase III C terminal | 9.00E-20<br>1.20E-05 |
| RII13767.1 | 76280 | 76610 | + | 109 | PF00550 | PP-binding | Phosphopantetheine attachment site | 1.00E-04 |

| index | leader | core | start | end | dir | score |
|---|---|---|---|---|---|---|
| 1 | VRYRGRPAPVTAALASRPRPLDGQLSR | ATQSSVRVPDKWPRYRSSASTRLATPQG | 66447 | 66615 | + | 6 |
| 2 | MNQTHARWVLHVNFPRGALTQFTQK...AGHHSSAPPTGRHTPLTCIYVPPPA | RVPPHPGTTSSGMTTP | 66756 | 67050 | + | 6 |
| 3 | MRARAREKRTDSPIHRKLSTQPRVAGHHSSAPPTGRHTPLTCIYVPPPA | RVPPHPGTTSSGMTTP | 66852 | 67050 | + | 7 |
| 4 | LKLPILGDYVLNPARQHRNTSATTG...AVRRFSAAQESSHTLVFPLSLSLHS | CVPLTRVHICRIERDRCDRVSIGTRQSRNGEVHAS | 67191 | 67476 | + | 6 |
| 5 | LPILGDYVLNPARQHRNTSATTGVG...AVRRFSAAQESSHTLVFPLSLSLHS | CVPLTRVHICRIERDRCDRVSIGTRQSRNGEVHAS | 67197 | 67476 | + | 6 |
| 6 | MLLEKVNENEALAGELTADGPVAYNVPVM | ATPALIATPVIAKAAGAVVGAGAVGGAAYGAYRAVVN | 67465 | 67666 | + | 30 |
| 7 | MSIMLERINAGEGEMTEQSIAPVM | ATPAAFVAGAVAGAKVSGALVAAAAGGAAVGAAVKK | 67874 | 68057 | + | 31 |
| 8 | MMTDSRIAWFLGVFLL | SLPLLLAAFKGEGRITRHVLARSLGTAAAIGVANALLWGHA | 71254 | 71428 | + | 7 |
| 9 | MASIDSNELDEVLA | TTFKELKTAVNNYNKDSIALYSQALSALVGLRQQVDAQDRAKA | 74825 | 74999 | + | 6 |
| 10 | MNGTTPDLGSTGTGTAGALPDPELI...LRVSVGPDDDYFAQGLANSLFSLEL | VTFVEDTFGLTVEVEDLNLDSFRTARRVTDFVLSKAGPGTGEAHADAV | 76280 | 76610 | + | 8 |

### Results for WP_019544895.1 [Streptomyces sulphureus DSM 40104] back to top

previous - next

#### Architecture

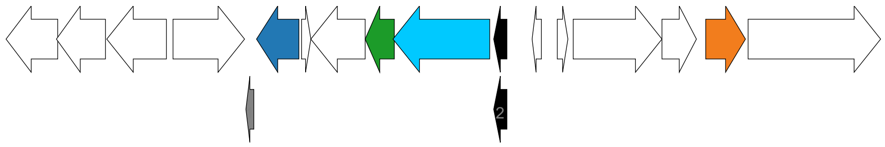

1000 nucleotides

Link to nucleotide sequence

| Accession | start | end | direction | length (aa) | Pfam/HMM | name | description | E-value |
|---|---|---|---|---|---|---|---|---|
| WP_019544888.1 | 126304 | 125305 | - | 332 | PF02737 | 3HCDH_N | 3-hydroxyacyl-CoA dehydrogenase, NAD binding domain | 4.40E-40 |
| | | | | | TIGR02279 | TIGR02279 | PaaC-3OHAcCoADH: 3-hydroxyacyl-CoA dehydrogenase PaaC | 6.70E-24 |
| | | | | | PF00725 | 3HCDH | 3-hydroxyacyl-CoA dehydrogenase, C-terminal domain | 5.20E-16 |
| | | | | | TIGR02437 | TIGR02437 | FadB: fatty oxidation complex, alpha subunit FadB | 3.30E-15 |
| | | | | | TIGR02441 | TIGR02441 | fa_ox_alpha_mit: fatty acid oxidation complex, alpha subunit, mitochondrial | 1.60E-12 |
| WP_019544889.1 | 127220 | 126290 | - | 309 | PF05853 | BKACE | beta-keto acid cleavage enzyme | 5.10E-101 |
| WP_037955413.1 | 128413 | 127258 | - | 384 | TIGR03228 | TIGR03228 | anthran_1_2_A: anthranilate 1,2-dioxygenase, large subunit | 5.10E-30 |
| | | | | | PF00848 | Ring_hydroxyl_A | Ring hydroxylating alpha subunit (catalytic domain) | 5.90E-28 |
| | | | | | TIGR03229 | TIGR03229 | benzo_1_2_benA: benzoate 1,2-dioxygenase, large subunit | 2.10E-20 |
| | | | | | PF00355 | Rieske | Rieske [2Fe-2S] domain | 2.90E-18 |
| | | | | | TIGR02378 | TIGR02378 | nirD_assim_sml: nitrite reductase [NAD(P)H], small subunit | 2.10E-05 |
| WP_019544891.1 | 128553 | 129915 | + | 453 | PF00171 | Aldedh | Aldehyde dehydrogenase family | 5.00E-128 |
| | | | | | TIGR01780 | TIGR01780 | SSADH: succinate-semialdehyde dehydrogenase | 2.70E-100 |
| | | | | | TIGR01804 | TIGR01804 | BADH: betaine-aldehyde dehydrogenase | 1.10E-92 |
| | | | | | TIGR02299 | TIGR02299 | HpaE: 5-carboxymethyl-2-hydroxymuconate semialdehyde dehydrogenase | 1.20E-81 |
| | | | | | TIGR04284 | TIGR04284 | aldehy_Rv0768: aldehyde dehydrogenase, Rv0768 family | 1.10E-76 |
| WP_019544892.1 | 130973 | 130151 | - | 273 | TIGR02427 | TIGR02427 | protocat_pcaD: 3-oxoadipate enol-lactonase | 2.10E-29 |
| | | | | | PF12697 | Abhydrolase_6 | Alpha/beta hydrolase family | 5.20E-17 |
| | | | | | TIGR03056 | TIGR03056 | bchO_mg_che_rel: putative magnesium chelatase accessory protein | 6.30E-16 |
| | | | | | TIGR03611 | TIGR03611 | RutD: pyrimidine utilization protein D | 2.30E-15 |
| | | | | | TIGR03695 | TIGR03695 | menH_SHCHC: 2-succinyl-6-hydroxy-2,4-cyclohexadiene-1-carboxylate synthase | 1.90E-14 |
| F603_RS34985 | 131033 | 131201 | + | 56 | INFERRED GENE | - | - | - |
| WP_019544893.1 | 132270 | 131208 | - | 353 | PF00891 | Methyltransf_2 | O-methyltransferase domain | 2.70E-37 |
| | | | | | TIGR02716 | TIGR02716 | C20_methyl_CrtF: C-20 methyltransferase BchU | 1.20E-08 |
| | | | | | PF16864 | Dimerisation2 | Dimerisation domain | 1.30E-05 |
| | | | | | TIGR03534 | TIGR03534 | RF_mod_PrmC: protein-(glutamine-N5) methyltransferase, release factor-specific | 6.40E-04 |
| WP_019544894.1 | 132818 | 132266 | - | 183 | LinL | LinL | LinL | 9.40E-58 |
| WP_019544895.1 | 134671 | 132814 | - | 618 | LinH | LinH | LinH (Fusion of LinE/LinG) | 3.70E-199 |
| | | | | | LinE | LinE | LinE Homolog | 8.80E-94 |
| | | | | | LinG | LinG | LinG Homolog | 5.70E-93 |
| | | | | | TIGR02427 | TIGR02427 | protocat_pcaD: 3-oxoadipate enol-lactonase | 6.40E-11 |
| | | | | | PF00561 | Abhydrolase_1 | alpha/beta hydrolase fold | 9.20E-11 |
| WP_106431256.1 | 135022 | 134746 | - | 91 | LinA | LinA | LinA (linaridin precursor) | 2.50E-13 |
| WP_019544897.1 | 135666 | 135495 | - | 56 | NO PFAM MATCH | - | - | - |
| WP_019544898.1 | 135991 | 136183 | + | 63 | NO PFAM MATCH | - | - | - |
| WP_019544899.1 | 136283 | 138005 | + | 573 | PF07730 | HisKA_3 | Histidine kinase | 5.50E-14 |
| WP_019544900.1 | 138001 | 138688 | + | 228 | PF00072 | Response_reg | Response regulator receiver domain | 3.20E-27 |
| | | | | | TIGR02875 | TIGR02875 | spore_0_A: sporulation transcription factor Spo0A | 2.60E-20 |
| | | | | | PF00196 | GerE | Bacterial regulatory proteins, luxR family | 1.30E-12 |
| | | | | | TIGR01818 | TIGR01818 | ntrC: nitrogen regulation protein NR(I) | 2.50E-12 |
| | | | | | TIGR02154 | TIGR02154 | PhoB: phosphate regulon transcriptional regulatory protein PhoB | 4.60E-12 |
| WP_019544901.1 | 138854 | 139640 | + | 261 | TIGR02982 | TIGR02982 | heterocyst_DevA: ABC exporter ATP-binding subunit, DevA family | 3.40E-58 |
| | | | | | TIGR02211 | TIGR02211 | LolD_lipo_ex: lipoprotein releasing system, ATP-binding protein | 3.50E-57 |
| | | | | | TIGR02673 | TIGR02673 | FtsE: cell division ATP-binding protein FtsE | 3.80E-55 |
| | | | | | TIGR03608 | TIGR03608 | L_ocin_972_ABC: putative bacteriocin export ABC transporter, lactococcin 972 group | 2.00E-53 |
| | | | | | TIGR02315 | TIGR02315 | ABC_phnC: phosphonate ABC transporter, ATP-binding protein | 2.00E-46 |
| WP_019544902.1 | 139690 | 142249 | + | 852 | TIGR03434 | TIGR03434 | ADOP: acidobacterial duplicated orphan permease | 4.10E-31 |
| | | | | | TIGR02212 | TIGR02212 | lolCE: lipoprotein releasing system, transmembrane protein, LolC/E family | 8.40E-30 |
| | | | | | PF12704 | MacB_PCD | MacB-like periplasmic core domain | 7.00E-24 |
| | | | | | PF02687 | FtsX | FtsX-like permease family | 8.20E-21 |
| | | | | | TIGR02213 | TIGR02213 | lolE_release: lipoprotein releasing system, transmembrane protein LolE | 1.70E-09 |

| index | leader | core | start | end | dir | score |
|---|---|---|---|---|---|---|
| 1 | VRGFALRSRASLRGVGSRWPGL | LLPATGELRPATAPGRTTRSSVTSVEVPA | 130104 | 129948 | - | 7 |
| 2 | MSRTTRIADWAQHHAEFGERHVPRSDEPLPVL | LTPAAVFAGVTAAQTAAQVGLTAAA...STVPSTALDTGRSVGDTLAHHPQGF | 135022 | 134746 | - | 18 |

### Results for KND26783.1 [Streptomyces stelliscabiei] back to top

previous - next

#### Architecture

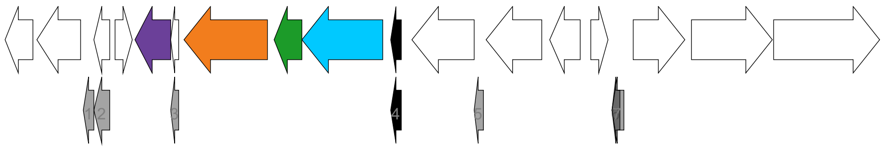

1000 nucleotides

Link to nucleotide sequence

| Accession | start | end | direction | length (aa) | Pfam/HMM | name | description | E-value |
|---|---|---|---|---|---|---|---|---|
| KND26775.1 | 1459495 | 1458877 | - | 205 | TIGR02365<br>PF02734<br>TIGR02361<br>TIGR03599 | TIGR02365<br>Dak2<br>TIGR02361<br>TIGR03599 | dha_L_ycgS: dihydroxyacetone kinase, L subunit<br>DAK2 domain<br>dak_ATP: dihydroxyacetone kinase<br>YloV: DAK2 domain fusion protein YloV | 2.40E-68<br>9.20E-50<br>1.40E-35<br>3.10E-07 |
| KND26776.1 | 1460578 | 1459585 | - | 330 | TIGR02363<br>PF02733<br>TIGR02361<br>TIGR02362 | TIGR02363<br>Dak1<br>TIGR02361<br>TIGR02362 | dhaK1: dihydroxyacetone kinase, DhaK subunit<br>Dak1 domain<br>dak_ATP: dihydroxyacetone kinase<br>dhaK1b: probable dihydroxyacetone kinase DhaK1b subunit | 4.30E-148<br>1.20E-125<br>5.50E-93<br>2.70E-73 |
| KND26777.1 | 1461249 | 1460898 | - | 116 | TIGR02228 | TIGR02228 | sigpep_I_arch: signal peptidase I | 5.40E-04 |
| KND26778.1 | 1461369 | 1461765 | + | 131 | PF00583<br>PF13673<br>TIGR01575<br>PF13508<br>PF14542 | Acetyltransf_1<br>Acetyltransf_10<br>TIGR01575<br>Acetyltransf_7<br>Acetyltransf_CG | Acetyltransferase (GNAT) family<br>Acetyltransferase (GNAT) domain<br>rimI: ribosomal-protein-alanine acetyltransferase<br>Acetyltransferase (GNAT) domain<br>GCN5-related N-acetyl-transferase | 8.30E-11<br>1.20E-05<br>5.30E-05<br>3.00E-04<br>5.20E-04 |
| KND26779.1 | 1462633 | 1461826 | - | 268 | PF13649<br>PF08241<br>PF13489<br>TIGR01983<br>PF08242 | Methyltransf_25<br>Methyltransf_11<br>Methyltransf_23<br>TIGR01983<br>Methyltransf_12 | Methyltransferase domain<br>Methyltransferase domain<br>Methyltransferase domain<br>UbiG: 3-demethylubiquinone-9 3-O-methyltransferase<br>Methyltransferase domain | 1.20E-13<br>9.70E-11<br>2.80E-06<br>3.10E-05<br>4.20E-05 |
| KND26780.1 | 1462821 | 1462629 | - | 63 | NO PFAM MATCH | - | - | - |
| KND26781.1 | 1464829 | 1462936 | - | 630 | TIGR02204<br>TIGR02203<br>TIGR00958<br>TIGR03796<br>TIGR01846 | TIGR02204<br>TIGR02203<br>TIGR00958<br>TIGR03796<br>TIGR01846 | MsbA_rel: ABC transporter, permease/ATP-binding protein<br>MsbA_lipidA: lipid A export permease/ATP-binding protein MsbA<br>3a01208: antigen peptide transporter 2<br>NHLM_micro_ABC1: NHLM bacteriocin system ABC transporter, peptidase/ATP-binding protein<br>type_I_sec_HlyB: type I secretion system ATPase | 2.50E-106<br>1.40E-101<br>1.90E-91<br>1.10E-84<br>1.30E-79 |
| KND26782.1 | 1465618 | 1464997 | - | 206 | LinL | LinL | LinL | 2.40E-69 |
| KND26783.1 | 1467462 | 1465614 | - | 615 | LinH<br>LinG<br>LinE<br>PF12697<br>PF12146 | LinH<br>LinG<br>LinE<br>Abhydrolase_6<br>Hydrolase_4 | LinH (Fusion of LinE/LinG)<br>LinG Homolog<br>LinE Homolog<br>Alpha/beta hydrolase family<br>Serine aminopeptidase, S33 | 9.70E-246<br>1.30E-154<br>6.40E-92<br>1.10E-16<br>5.10E-09 |
| KND26784.1 | 1467899 | 1467650 | - | 82 | LinA | LinA | LinA (linaridin precursor) | 3.30E-04 |
| KND26785.1 | 1469558 | 1468127 | - | 476 | TIGR02734<br>PF13450<br>TIGR03467<br>TIGR00292<br>PF01266 | TIGR02734<br>NAD_binding_8<br>TIGR03467<br>TIGR00292<br>DAO | crtI_fam: phytoene desaturase<br>NAD(P)-binding Rossmann-like domain<br>HpnE: squalene-associated FAD-dependent desaturase<br>TIGR00292: thiazole biosynthesis enzyme<br>FAD dependent oxidoreductase | 1.00E-12<br>2.50E-11<br>1.70E-08<br>6.30E-08<br>6.80E-08 |
| KND26786.1 | 1471089 | 1469823 | - | 421 | PF07730 | HisKA_3 | Histidine kinase | 8.70E-14 |
| KND26787.1 | 1471944 | 1471260 | - | 227 | PF00072<br>PF00196<br>TIGR02875<br>TIGR03020<br>TIGR03541 | Response_reg<br>GerE<br>TIGR02875<br>TIGR03020<br>TIGR03541 | Response regulator receiver domain<br>Bacterial regulatory proteins, luxR family<br>spore_0_A: sporulation transcription factor Spo0A<br>EpsA: transcriptional regulator EpsA<br>reg_near_HchA: LuxR family transcriptional regulatory, chaperone HchA-associated | 1.40E-24<br>2.50E-18<br>2.30E-15<br>1.60E-13<br>1.20E-08 |
| KND26788.1 | 1472196 | 1472595 | + | 132 | PF00196<br>TIGR03541<br>TIGR03020<br>PF08281<br>PF13518 | GerE<br>TIGR03541<br>TIGR03020<br>Sigma70_r4_2<br>HTH_28 | Bacterial regulatory proteins, luxR family<br>reg_near_HchA: LuxR family transcriptional regulatory, chaperone HchA-associated<br>EpsA: transcriptional regulator EpsA<br>Sigma-70, region 4<br>Helix-turn-helix domain | 1.40E-15<br>6.10E-12<br>1.00E-08<br>1.00E-05<br>8.90E-04 |
| KND26789.1 | 1473159 | 1474347 | + | 395 | PF00480<br>TIGR00744<br>PF12802<br>PF13412<br>PF01978 | ROK<br>TIGR00744<br>MarR_2<br>HTH_24<br>TrmB | ROK family<br>ROK_glcA_fam: ROK family protein (putative glucokinase)<br>MarR family<br>Winged helix-turn-helix DNA-binding<br>Sugar-specific transcriptional regulator TrmB | 6.80E-53<br>6.40E-40<br>1.30E-07<br>4.70E-05<br>1.10E-04 |
| KND26790.1 | 1474477 | 1476319 | + | 613 | PF16499<br>PF17801<br>PF16990<br>PF03422 | Melibiase_2<br>Melibiase_C<br>CBM_35<br>CBM_6 | Alpha galactosidase A<br>Alpha galactosidase C-terminal beta sandwich domain<br>Carbohydrate binding module (family 35)<br>Carbohydrate binding module (family 6) | 4.50E-37<br>3.70E-16<br>1.90E-07<br>6.80E-07 |
| KND26791.1 | 1476345 | 1478769 | + | 807 | PF14498 | Glyco_hyd_65N_2 | Glycosyl hydrolase family 65, N-terminal domain | 5.10E-36 |

| index | leader | core | start | end | dir | score |
|---|---|---|---|---|---|---|
| 1 | VTRPPPTHGVPDDGPPPPADRADHLCGRPSRSPSPTS | ALPSPLPPSRQVGRAHRRVDARDLRLLKVDTFTRRASTDPCPTTPRI | 1460902 | 1460647 | - | 6 |
| 2 | MGALDMVAERVVAGATVTFRPSGSS...ISNNRGHVNGWTGHGRVFGICVAVD | GTARPRTAGKTRPGG | 1461249 | 1460898 | - | 6 |
| 3 | MNDFLLIPSVVALGIVGFLIATRAVSTPAVI | GVALVVLWGACSQARSTIHARHTPSKKRRQHP | 1462821 | 1462629 | - | 6 |
| 4 | MEIPVEGVVYVAARPTL | GTPRIARVGNRLGGLAQAAEGVGAI...ANNLAAEANAQNAAALAAVGGAPAA | 1467899 | 1467650 | - | 28 |
| 5 | MSESQRRRDGLISSRRVVNRPGPGPFEGPFR | LLSPAISRVGGPVSTVGKSQEKRNAEDAGDI | 1469743 | 1469554 | - | 6 |
| 6 | MRPRGPLAPTRRRSSGGGRAEPRRDGGGARTARVRRGGGPRVGH | TAPDGGRTAAERRENRPLLPRAGG | 1472873 | 1472666 | - | 6 |
| 7 | MTTCRGSAAHRADDGLRTPGTERAA...AEPRRDGGGARTARVRRGGGPRVGH | TAPDGGRTAAERRENRPLLPRAGG | 1472954 | 1472666 | - | 6 |

### Results for WP_055713858.1 [Streptomyces torulosus] back to top

previous - next

#### Architecture

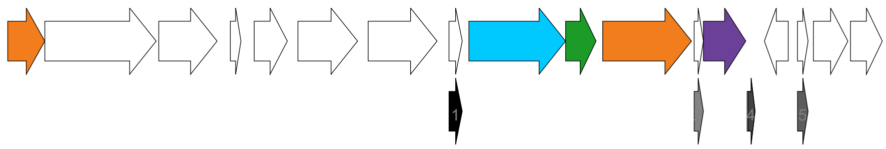

1000 nucleotides

Link to nucleotide sequence

| Accession | start | end | direction | length (aa) | Pfam/HMM | name | description | E-value |
|---|---|---|---|---|---|---|---|---|
| WP_055713854.1 | 22511 | 23261 | + | 249 | TIGR02673<br>TIGR02211<br>TIGR03608<br>TIGR02982<br>TIGR02315 | TIGR02673<br>TIGR02211<br>TIGR03608<br>TIGR02982<br>TIGR02315 | FtsE: cell division ATP-binding protein FtsE<br>LolD_lipo_ex: lipoprotein releasing system, ATP-binding protein<br>L_ocin_972_ABC: putative bacteriocin export ABC transporter, lactococcin 972 group<br>heterocyst_DevA: ABC exporter ATP-binding subunit, DevA family<br>ABC_phnC: phosphonate ABC transporter, ATP-binding protein | 5.60E-61<br>1.70E-60<br>6.20E-55<br>1.40E-54<br>5.00E-46 |
| WP_055713855.1 | 23264 | 25556 | + | 763 | PF12704<br>TIGR03434<br>PF02687<br>TIGR02212<br>TIGR00439 | MacB_PCD<br>TIGR03434<br>FtsX<br>TIGR02212<br>TIGR00439 | MacB-like periplasmic core domain<br>ADOP: acidobacterial duplicated orphan permease<br>FtsX-like permease family<br>lolCE: lipoprotein releasing system, transmembrane protein, LolC/E family<br>ftsX: putative protein insertion permease FtsX | 3.50E-20<br>2.70E-13<br>2.40E-09<br>9.00E-05<br>1.10E-04 |
| WP_020135148.1 | 25596 | 26796 | + | 400 | INFERRED GENE | - | - | - |
| WP_013005810.1 | 27076 | 27289 | + | 71 | INFERRED GENE | - | - | - |
| WP_055713856.1 | 27543 | 28233 | + | 229 | PF00072<br>PF00196<br>TIGR02875<br>TIGR03020<br>TIGR03541 | Response_reg<br>GerE<br>TIGR02875<br>TIGR03020<br>TIGR03541 | Response regulator receiver domain<br>Bacterial regulatory proteins, luxR family<br>spore_0_A: sporulation transcription factor Spo0A<br>EpsA: transcriptional regulator EpsA<br>reg_near_HchA: LuxR family transcriptional regulatory, chaperone HchA-associated | 1.80E-24<br>8.70E-18<br>1.70E-16<br>5.80E-13<br>3.70E-08 |
| WP_079049558.1 | 28445 | 29675 | + | 409 | PF07730<br>PF13581 | HisKA_3<br>HATPase_c_2 | Histidine kinase<br>Histidine kinase-like ATPase domain | 2.60E-14<br>6.70E-04 |
| WP_055713857.1 | 29882 | 31298 | + | 471 | TIGR02734<br>PF13450<br>TIGR03467<br>TIGR03315<br>PF01266 | TIGR02734<br>NAD_binding_8<br>TIGR03467<br>TIGR03315<br>DAO | crtI_fam: phytoene desaturase<br>NAD(P)-binding Rossmann-like domain<br>HpnE: squalene-associated FAD-dependent desaturase<br>Se_ygfK: putative selenate reductase, YgfK subunit<br>FAD dependent oxidoreductase | 4.10E-13<br>3.60E-11<br>8.60E-09<br>6.30E-08<br>2.00E-07 |
| WP_055713870.1 | 31540 | 31801 | + | 86 | NO PFAM MATCH | - | - | - |
| WP_055713858.1 | 31948 | 33919 | + | 656 | LinH<br>LinG<br>LinE<br>PF12697<br>PF12146 | LinH<br>LinG<br>LinE<br>Abhydrolase_6<br>Hydrolase_4 | LinH (Fusion of LinE/LinG)<br>LinG Homolog<br>LinE Homolog<br>Alpha/beta hydrolase family<br>Serine aminopeptidase, S33 | 9.40E-234<br>1.90E-124<br>5.80E-85<br>2.90E-15<br>5.30E-08 |
| WP_055713859.1 | 33915 | 34548 | + | 210 | LinL | LinL | LinL | 9.20E-75 |
| WP_079049555.1 | 34672 | 36508 | + | 611 | TIGR02204<br>TIGR02203<br>TIGR00958<br>TIGR03796<br>TIGR01846 | TIGR02204<br>TIGR02203<br>TIGR00958<br>TIGR03796<br>TIGR01846 | MsbA_rel: ABC transporter, permease/ATP-binding protein<br>MsbA_lipidA: lipid A export permease/ATP-binding protein MsbA<br>3a01208: antigen peptide transporter 2<br>NHLM_micro_ABC1: NHLM bacteriocin system ABC transporter, peptidase/ATP-binding protein<br>type_I_sec_HlyB: type I secretion system ATPase | 4.30E-106<br>1.20E-104<br>3.30E-93<br>9.30E-87<br>1.60E-84 |
| WP_055713860.1 | 36548 | 36740 | + | 63 | NO PFAM MATCH | - | - | - |
| WP_079049556.1 | 36736 | 37615 | + | 292 | PF13649<br>PF08241<br>PF13489<br>PF08242<br>TIGR01983 | Methyltransf_25<br>Methyltransf_11<br>Methyltransf_23<br>Methyltransf_12<br>TIGR01983 | Methyltransferase domain<br>Methyltransferase domain<br>Methyltransferase domain<br>Methyltransferase domain<br>UbiG: 3-demethylubiquinone-9 3-O-methyltransferase | 1.70E-15<br>5.20E-13<br>6.30E-10<br>5.50E-07<br>2.00E-06 |
| WP_055713862.1 | 38472 | 37992 | - | 159 | NO PFAM MATCH | - | - | - |
| WP_055518518.1 | 38662 | 38896 | + | 77 | NO PFAM MATCH | - | - | - |
| WP_169800310.1 | 38989 | 39709 | + | 239 | PF09844<br>PF06314 | DUF2071<br>ADC | Uncharacterized conserved protein (COG2071)<br>Acetoacetate decarboxylase (ADC) | 3.30E-60<br>3.30E-06 |
| WP_079049560.1 | 39762 | 40404 | + | 213 | NO PFAM MATCH | - | - | - |

| index | leader | core | start | end | dir | score |
|---|---|---|---|---|---|---|
| 1 | MDIPVEAVILASARPTL | GTPRIGRVSGANALNALAGLADLAQ...AAAEANQAVANAANAQAAAAGGVPA | 31540 | 31801 | + | 18 |
| 2 | MNDFLLIPSVVALGIVGFLIATRAVSAPAVI | GVALVVLWGACSQARGNAHARNTPSKKRRHHP | 36548 | 36740 | + | 7 |
| 3 | VRALRAPVRHASPVPGPAPVP | LPPCVTHVFRTSSAGTPATRPSSASRCGSTTAPRAAGTW | 37630 | 37813 | + | 7 |
| 4 | VPGPAPVPLPPCVTHVFRTSSAG | TPATRPSSASRCGSTTAPRAAGTW | 37669 | 37813 | + | 7 |
| 5 | MTSPDPEPNRTTGLEPGGGVPPG | ETPPAEGGTSGVARPDPPAVRRKGW...LVLIILLAAVIAIGLIGMAVVLIVQ | 38662 | 38896 | + | 8 |

### Results for AWI33039.1 [Streptomyces tirandamycinicus] back to top

previous - next

#### Architecture

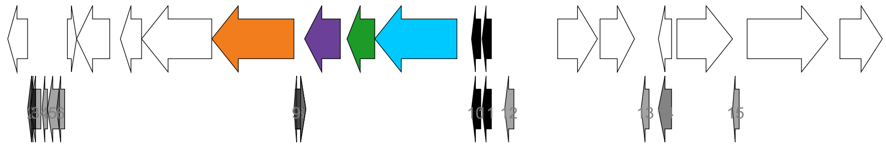

1000 nucleotides

Link to nucleotide sequence

| Accession | start | end | direction | length (aa) | Pfam/HMM | name | description | E-value |
|---|---|---|---|---|---|---|---|---|
| AWI29761.1 | 3153004 | 3152569 | - | 144 | PF09346 | SMI1_KNR4 | SMI1 / KNR4 family (SUKH-1) | 3.60E-05 |
| WP_010038104.1 | 3153829 | 3154017 | + | 62 | INFERRED GENE | - | - | - |
| AWI33038.1 | 3154718 | 3154022 | - | 231 | PF01551 | Peptidase_M23 | Peptidase family M23 | 1.20E-18 |
| AWI29762.1 | 3155391 | 3154950 | - | 146 | NO PFAM MATCH | - | - | - |
| AWI29763.1 | 3156851 | 3155390 | - | 486 | TIGR02734<br>TIGR03467<br>TIGR00292<br>PF13450<br>PF01946 | TIGR02734<br>TIGR03467<br>TIGR00292<br>NAD_binding_8<br>Thi4 | crtI_fam: phytoene desaturase<br>HpnE: squalene-associated FAD-dependent desaturase<br>TIGR00292: thiazole biosynthesis enzyme<br>NAD(P)-binding Rossmann-like domain<br>Thi4 family | 1.40E-10<br>9.70E-09<br>1.40E-08<br>3.00E-08<br>3.20E-07 |
| AWI29764.1 | 3158569 | 3156847 | - | 573 | TIGR02204<br>TIGR02203<br>TIGR00958<br>TIGR01846<br>TIGR03796 | TIGR02204<br>TIGR02203<br>TIGR00958<br>TIGR01846<br>TIGR03796 | MsbA_rel: ABC transporter, permease/ATP-binding protein<br>MsbA_lipidA: lipid A export permease/ATP-binding protein MsbA<br>3a01208: antigen peptide transporter 2<br>type_I_sec_HlyB: type I secretion system ATPase<br>NHLM_micro_ABC1: NHLM bacteriocin system ABC transporter, peptidase/ATP-binding protein | 8.20E-107<br>1.60E-104<br>5.40E-88<br>1.90E-85<br>1.70E-80 |
| AWI29765.1 | 3159554 | 3158792 | - | 253 | PF13649<br>PF08241<br>PF08242<br>PF13489<br>PF13847 | Methyltransf_25<br>Methyltransf_11<br>Methyltransf_12<br>Methyltransf_23<br>Methyltransf_31 | Methyltransferase domain<br>Methyltransferase domain<br>Methyltransferase domain<br>Methyltransferase domain<br>Methyltransferase domain | 3.60E-13<br>1.30E-09<br>5.90E-09<br>9.70E-08<br>3.00E-07 |
| AWI29766.1 | 3160267 | 3159682 | - | 194 | LinL | LinL | LinL | 8.00E-69 |
| AWI33039.1 | 3161976 | 3160263 | - | 570 | LinH<br>LinE<br>LinG<br>PF12697<br>PF12146 | LinH<br>LinE<br>LinG<br>Abhydrolase_6<br>Hydrolase_4 | LinH (Fusion of LinE/LinG)<br>LinE Homolog<br>LinG Homolog<br>Alpha/beta hydrolase family<br>Serine aminopeptidase, S33 | 4.20E-226<br>2.00E-100<br>3.00E-97<br>3.50E-16<br>7.70E-10 |
| AWI29767.1 | 3162498 | 3162303 | - | 64 | LinA | LinA | LinA (linaridin precursor) | 1.10E-09 |
| AWI29768.1 | 3162708 | 3162513 | - | 64 | LinA | LinA | LinA (linaridin precursor) | 6.40E-13 |
| WP_005482622.1 | 3164087 | 3164920 | + | 277 | INFERRED GENE | - | - | - |
| AWI33040.1 | 3164979 | 3165714 | + | 244 | PF00589<br>TIGR02225<br>TIGR02224<br>TIGR02249 | Phage_integrase<br>TIGR02225<br>TIGR02224<br>TIGR02249 | Phage integrase family<br>recomb_XerD: tyrosine recombinase XerD<br>recomb_XerC: tyrosine recombinase XerC<br>integrase_gron: integron integrase | 9.10E-20<br>1.30E-12<br>2.10E-12<br>2.10E-08 |
| AWI29769.1 | 3166476 | 3166212 | - | 87 | NO PFAM MATCH | - | - | - |
| AWI29770.1 | 3166609 | 3167770 | + | 386 | PF14072<br>TIGR03187<br>TIGR03233 | DndB<br>TIGR03187<br>TIGR03233 | DNA-sulfur modification-associated<br>DGQHR: DGQHR domain<br>DNA_S_dndB: DNA sulfur modification protein DndB | 1.90E-29<br>4.90E-22<br>8.90E-15 |
| AWI29771.1 | 3168078 | 3169755 | + | 558 | PF01844 | HNH | HNH endonuclease | 2.40E-07 |
| AWI29772.1 | 3170001 | 3170883 | + | 293 | PF13365 | Trypsin_2 | Trypsin-like peptidase domain | 5.70E-09 |

| index | leader | core | start | end | dir | score |
|---|---|---|---|---|---|---|
| 1 | MAAAGTNQALGRRRAVCC | DSCAQSVRGRAVCNAAGT | 3153152 | 3153041 | - | 6 |
| 2 | VGRRRHGGRRYQPGVGAATRCLLRFMR | TVGAGPRCVQRGGNVTAVGAPRCPQRK | 3153168 | 3153003 | - | 8 |
| 3 | MPPARRRAPPSATSSKGWTALLPDRPSGWADPSWGDA | DMAAAGTNQALGRRRAVCCDSCAQSVRGRAVCNAAGT | 3153266 | 3153041 | - | 6 |
| 4 | MTWGFCCPGCDGARRGATG | PTVGTVRNGLRERRG | 3153414 | 3153309 | - | 6 |
| 5 | LTGAAAGAVRPTDLRERCVSGWCSTGPHRMDRPSGQTSLTR | QNQTAEHHPEPPVVIAQALVMRRSSVRIRQGAPRNPRSHVP | 3153659 | 3153410 | - | 6 |
| 6 | LPDVSMAART | STAPEPQAGRVAEVYQTRSARRRASSLRTAAMLDRGRRRCRQTHGPA | 3153786 | 3153612 | - | 6 |
| 7 | MLPPKIRLLAQLALVISPLSR | KTVSMTGTATMKATTNDTTRLVVTMIPFLVVARRQAVQLRLKCPSDR | 3158608 | 3158815 | + | 6 |
| 8 | LAQLALVISPLSR | KTVSMTGTATMKATTNDTTRLVVTMIPFLVVARRQAVQLRLKCPSDR | 3158632 | 3158815 | + | 6 |
| 9 | VSFVVAFIVAVPVM | LTVFRESGEITRASWAKSLIFGGSIAAIAAIAFGRSRQ | 3158724 | 3158565 | - | 6 |
| 10 | MNRITESARKAASPTLSEVQSDKVVRMMPVA | FTPSVVALCPYAAGGLAAGAAVVGAYEAGRAAG | 3162498 | 3162303 | - | 29 |
| 11 | MAEIAAFMNRQEMSPLEEGMAKTDGTPIM | ATPAAVAIPAGAKIAAAAVAFTIPVGAYVTGRVFG | 3162708 | 3162513 | - | 30 |
| 12 | MQFLASRKRGCSRCAIGRIPWSGQLWGVGILWKISATQNFWRLNRSEILQ | SAPPVSCDVHHSRSP | 3163196 | 3162998 | - | 6 |
| 13 | LRLAPSESVVVRLSPASLLTCTGASMM | DADGRTWSKIGQLVMRRSSVRIRQGALW | 3166003 | 3165835 | - | 6 |
| 14 | MTLHTLGGSSSGSEAGGNGGTDGNGDLFRFRRPSGGLVG | GTALGLAVAVHMTGVIPEGLPSYAAGGLVGVSVMGVFLTKCLRPGKAC | 3166476 | 3166212 | - | 7 |
| 15 | VPSRLSLAQLRPNSTDNDAPVLLLL | DRACAFRRWVGSGPSHQPTGVIRMSS | 3167909 | 3167753 | - | 6 |

### Results for WP_159744032.1 [Streptomyces tubercidicus] back to top

previous - next

#### Architecture

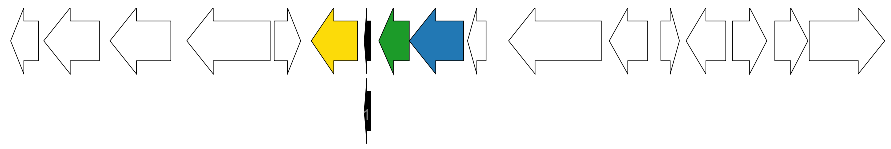

1000 nucleotides

Link to nucleotide sequence

| Accession | start | end | direction | length (aa) | Pfam/HMM | name | description | E-value |
|---|---|---|---|---|---|---|---|---|
| WP_030077134.1 | 3190394 | 3189854 | - | 179 | TIGR01203<br>PF00156<br>TIGR01251<br>TIGR01367 | TIGR01203<br>Pribosyltran<br>TIGR01251<br>TIGR01367 | HGPRTase: hypoxanthine phosphoribosyltransferase<br>Phosphoribosyl transferase domain<br>ribP_PPkin: ribose-phosphate diphosphokinase<br>pyrE_Therm: orotate phosphoribosyltransferase | 4.60E-65<br>1.50E-25<br>9.60E-06<br>5.60E-04 |
| WP_159744027.1 | 3191539 | 3190480 | - | 352 | TIGR02432<br>PF01171<br>PF09179 | TIGR02432<br>ATP_bind_3<br>TilS | lysidine_TilS_N: tRNA(Ile)-lysidine synthetase<br>PP-loop family<br>TilS substrate binding domain | 2.20E-41<br>2.90E-36<br>1.10E-12 |
| WP_159744028.1 | 3192876 | 3191730 | - | 381 | TIGR03883<br>TIGR03624<br>PF10103 | TIGR03883<br>TIGR03624<br>Zincin_2 | DUF2342_F420: uncharacterized protein, coenzyme F420 biosynthesis associated<br>TIGR03624: putative hydrolase<br>Zincin-like metallopeptidase | 1.50E-141<br>3.50E-125<br>6.90E-121 |
| WP_159744029.1 | 3194753 | 3193178 | - | 524 | PF02113<br>TIGR00666 | Peptidase_S13<br>TIGR00666 | D-Ala-D-Ala carboxypeptidase 3 (S13) family<br>PBP4: D-alanyl-D-alanine carboxypeptidase/D-alanyl-D-alanine-endopeptidase | 7.60E-75<br>1.70E-56 |
| WP_018090094.1 | 3194839 | 3195334 | + | 164 | PF00719 | Pyrophosphatase | Inorganic pyrophosphatase | 2.10E-53 |
| WP_159744030.1 | 3196393 | 3195538 | - | 284 | LinG<br>LinH | LinG<br>LinH | LinG Homolog<br>LinH (Fusion of LinE/LinG) | 3.20E-100<br>2.00E-75 |
| WP_159744031.1 | 3196652 | 3196529 | - | 40 | LinA | LinA | LinA (linaridin precursor) | 8.20E-09 |
| WP_167539154.1 | 3197378 | 3196790 | - | 195 | LinL | LinL | LinL | 4.80E-54 |
| WP_159744032.1 | 3198393 | 3197379 | - | 337 | LinE<br>LinH<br>PF12697<br>TIGR02427<br>PF00561 | LinE<br>LinH<br>Abhydrolase_6<br>TIGR02427<br>Abhydrolase_1 | LinE Homolog<br>LinH (Fusion of LinE/LinG)<br>Alpha/beta hydrolase family<br>protocat_pcaD: 3-oxoadipate enol-lactonase<br>alpha/beta hydrolase fold | 1.70E-94<br>4.30E-50<br>1.60E-12<br>1.20E-08<br>2.10E-08 |
| WP_159744033.1 | 3198813 | 3198465 | - | 115 | PF13518<br>PF01527<br>PF13551<br>PF13011<br>PF13384 | HTH_28<br>HTH_Tnp_1<br>HTH_29<br>LZ_Tnp_IS481<br>HTH_23 | Helix-turn-helix domain<br>Transposase<br>Winged helix-turn helix<br>leucine-zipper of insertion element IS481<br>Homeodomain-like domain | 5.50E-09<br>5.70E-07<br>2.70E-06<br>8.60E-05<br>1.20E-04 |
| WP_159744034.1 | 3200988 | 3199242 | - | 581 | PF06738<br>PF12821 | ThrE<br>ThrE_2 | Putative threonine/serine exporter<br>Threonine/Serine exporter, ThrE | 5.90E-71<br>9.30E-19 |
| WP_159744035.1 | 3201849 | 3201147 | - | 233 | PF09335 | SNARE_assoc | SNARE associated Golgi protein | 1.90E-24 |
| WP_159744036.1 | 3202103 | 3202472 | + | 122 | PF01906 | YbjQ_1 | Putative heavy-metal-binding | 2.30E-33 |
| WP_159744037.1 | 3203334 | 3202575 | - | 252 | PF07739<br>PF13411<br>TIGR02043<br>TIGR02051<br>PF00376 | TipAS<br>MerR_1<br>TIGR02043<br>TIGR02051<br>MerR | TipAS antibiotic-recognition domain<br>MerR HTH family regulatory protein<br>ZntR: Zn(II)-responsive transcriptional regulator<br>MerR: Hg(II)-responsive transcriptional regulator<br>MerR family regulatory protein | 1.60E-34<br>5.40E-17<br>4.50E-15<br>1.60E-14<br>3.10E-14 |
| WP_159744038.1 | 3203449 | 3204118 | + | 222 | TIGR00949<br>PF01810<br>TIGR00948 | TIGR00949<br>LysE<br>TIGR00948 | 2A76: homoserine/Threonine efflux protein<br>LysE type translocator<br>2a75: L-lysine exporter | 2.30E-44<br>6.40E-44<br>3.90E-16 |
| WP_159744039.1 | 3204258 | 3204882 | + | 207 | TIGR01755<br>PF03358<br>PF02525<br>TIGR01753<br>PF12724 | TIGR01755<br>FMN_red<br>Flavodoxin_2<br>TIGR01753<br>Flavodoxin_5 | flav_wrbA: NAD(P)H:quinone oxidoreductase, type IV<br>NADPH-dependent FMN reductase<br>Flavodoxin-like fold<br>flav_short: flavodoxin<br>Flavodoxin domain | 1.70E-42<br>2.20E-15<br>2.50E-09<br>3.20E-09<br>7.70E-07 |
| WP_159744040.1 | 3204898 | 3206314 | + | 471 | TIGR01788<br>PF00282<br>TIGR03812<br>TIGR03799 | TIGR01788<br>Pyridoxal_deC<br>TIGR03812<br>TIGR03799 | Glu-decarb-GAD: glutamate decarboxylase<br>Pyridoxal-dependent decarboxylase conserved domain<br>tyr_de_CO2_Arch: tyrosine decarboxylase MnfA<br>NOD_PanD_pyr: putative pyridoxal-dependent aspartate 1-decarboxylase | 8.30E-188<br>2.40E-74<br>4.60E-40<br>3.80E-05 |

| index | leader | core | start | end | dir | score |
|---|---|---|---|---|---|---|
| 1 | MSELADFANAELVDVEPGRI | GADATPTMVLVHNATTDIIG | 3196652 | 3196529 | - | 26 |

### Results for WP_106434293.1 [Streptomyces viridochromogenes DSM 40736]

back to top

previous - next

#### Architecture

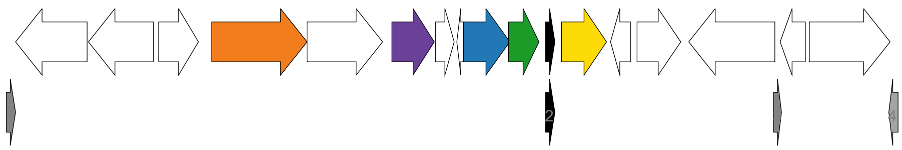

1000 nucleotides

Link to nucleotide sequence

| Accession | start | end | direction | length (aa) | Pfam/HMM | name | description | E-value |
|---|---|---|---|---|---|---|---|---|

| | | | | | | | | |
|---|---|---|---|---|---|---|---|---|
| WP_039933341.1 | 4919164 | 4917757 | - | 468 | TIGR01788<br>PF00282<br>TIGR03812<br>TIGR03799<br>TIGR03811 | TIGR01788<br>Pyridoxal_deC<br>TIGR03812<br>TIGR03799<br>TIGR03811 | Glu-decarb-GAD: glutamate decarboxylase<br>Pyridoxal-dependent decarboxylase conserved domain<br>tyr_de_CO2_Arch: tyrosine decarboxylase MnfA<br>NOD_PanD_pyr: putative pyridoxal-dependent aspartate 1-decarboxylase<br>tyr_de_CO2_Ent: tyrosine decarboxylase | 3.00E-187<br>1.60E-71<br>2.20E-38<br>2.50E-06<br>9.90E-04 |
| WP_003991972.1 | 4920489 | 4919202 | - | 428 | PF00654 | Voltage_CLC | Voltage gated chloride channel | 1.60E-21 |
| WP_003991973.1 | 4920591 | 4921365 | + | 257 | PF07739<br>PF13411<br>TIGR02044<br>TIGR02043<br>TIGR02051 | TipAS<br>MerR_1<br>TIGR02044<br>TIGR02043<br>TIGR02051 | TipAS antibiotic-recognition domain<br>MerR HTH family regulatory protein<br>CueR: Cu(I)-responsive transcriptional regulator<br>ZntR: Zn(II)-responsive transcriptional regulator<br>MerR: Hg(II)-responsive transcriptional regulator | 1.70E-39<br>5.20E-17<br>3.70E-14<br>8.50E-14<br>1.90E-13 |
| WP_003991974.1 | 4921640 | 4923515 | + | 624 | TIGR02204<br>TIGR02203<br>TIGR00958<br>TIGR03796<br>TIGR03375 | TIGR02204<br>TIGR02203<br>TIGR00958<br>TIGR03796<br>TIGR03375 | MsbA_rel: ABC transporter, permease/ATP-binding protein<br>MsbA_lipidA: lipid A export permease/ATP-binding protein MsbA<br>3a01208: antigen peptide transporter 2<br>NHLM_micro_ABC1: NHLM bacteriocin system ABC transporter, peptidase/ATP-binding protein<br>type_I_sec_LssB: type I secretion system ATPase | 2.70E-97<br>4.40E-94<br>2.60E-85<br>9.30E-81<br>4.90E-78 |
| WP_017883627.1 | 4923511 | 4925007 | + | 498 | INFERRED GENE | - | - | - |
| WP_078506889.1 | 4925194 | 4926004 | + | 269 | PF13649<br>PF08241<br>PF13489<br>PF08242<br>TIGR01983 | Methyltransf_25<br>Methyltransf_11<br>Methyltransf_23<br>Methyltransf_12<br>TIGR01983 | Methyltransferase domain<br>Methyltransferase domain<br>Methyltransferase domain<br>Methyltransferase domain<br>UbiG: 3-demethylubiquinone-9 3-O-methyltransferase | 5.90E-15<br>6.70E-11<br>6.60E-09<br>9.50E-09<br>1.20E-06 |
| WP_003991977.1 | 4926049 | 4926397 | + | 115 | PF13518<br>PF01527<br>PF13011<br>PF13551<br>PF13384 | HTH_28<br>HTH_Tnp_1<br>LZ_Tnp_IS481<br>HTH_29<br>HTH_23 | Helix-turn-helix domain<br>Transposase<br>leucine-zipper of insertion element IS481<br>Winged helix-turn helix<br>Homeodomain-like domain | 6.40E-09<br>1.20E-08<br>5.80E-06<br>1.50E-05<br>2.40E-05 |
| WP_167545764.1 | 4926632 | 4926449 | - | 60 | NO PFAM MATCH | - | - | - |
| WP_106434293.1 | 4926587 | 4927493 | + | 301 | LinE<br>LinH<br>PF12697<br>TIGR02427<br>PF00561 | LinE<br>LinH<br>Abhydrolase_6<br>TIGR02427<br>Abhydrolase_1 | LinE Homolog<br>LinH (Fusion of LinE/LinG)<br>Alpha/beta hydrolase family<br>protocat_pcaD: 3-oxoadipate enol-lactonase<br>alpha/beta hydrolase fold | 4.20E-94<br>1.30E-49<br>4.20E-14<br>2.50E-11<br>9.60E-11 |
| WP_063749169.1 | 4927489 | 4928077 | + | 195 | LinL | LinL | LinL | 1.20E-56 |
| WP_158681081.1 | 4928220 | 4928394 | + | 57 | LinA | LinA | LinA (linaridin precursor) | 7.40E-12 |
| WP_052310327.1 | 4928535 | 4929414 | + | 292 | LinG<br>LinH | LinG<br>LinH | LinG Homolog<br>LinH (Fusion of LinE/LinG) | 5.00E-107<br>1.10E-78 |
| WP_003991982.1 | 4929871 | 4929496 | - | 124 | PF01906<br>TIGR03884 | YbjQ_1<br>TIGR03884 | Putative heavy-metal-binding<br>sel_bind_Methan: selenium-binding protein | 3.40E-33<br>2.10E-04 |
| WP_003991983.1 | 4930007 | 4930880 | + | 290 | PF09335 | SNARE_assoc | SNARE associated Golgi protein | 1.90E-25 |
| WP_003991984.1 | 4932715 | 4931017 | - | 565 | PF06738<br>PF12821 | ThrE<br>ThrE_2 | Putative threonine/serine exporter<br>Threonine/Serine exporter, ThrE | 1.80E-68<br>2.90E-16 |
| WP_003991985.1 | 4933329 | 4932837 | - | 163 | PF00719 | Pyrophosphatase | Inorganic pyrophosphatase | 1.60E-53 |
| WP_039933344.1 | 4933408 | 4934989 | + | 526 | PF02113<br>TIGR00666<br>PF00768 | Peptidase_S13<br>TIGR00666<br>Peptidase_S11 | D-Ala-D-Ala carboxypeptidase 3 (S13) family<br>PBP4: D-alanyl-D-alanine carboxypeptidase/D-alanyl-D-alanine-endopeptidase<br>D-alanyl-D-alanine carboxypeptidase | 6.20E-76<br>1.10E-56<br>5.30E-07 |

| index | leader | core | start | end | dir | score |
|---|---|---|---|---|---|---|
| 1 | MPGWLGTVVEWNPMS | HTATAVRDLFGVPGAESGHVWPAVIWPLALLAVFFPLAVRRFGRLSR | 4917572 | 4917761 | + | 7 |
| 2 | MADFANTELADVTPGRIGNDATPTM | LTPLAALATPEGVAVTAATAYALNEVTNDLAG | 4928220 | 4928394 | + | 33 |
| 3 | LRSSASVTAWCSLYDTSASYPPLPQYA | HTRTNGPHAHACGPSASGEGEVSGRPAP | 4932692 | 4932860 | + | 7 |
| 4 | VGQGDEGGIRRGRLRVLGVVCRAHPCRPGG | TTAHGGARRPAHGGAGAASGRGAPGR | 4935156 | 4934985 | - | 6 |

### Results for WP_132756788.1 [Streptomyces vinaceus] back to top

previous - next

#### Architecture

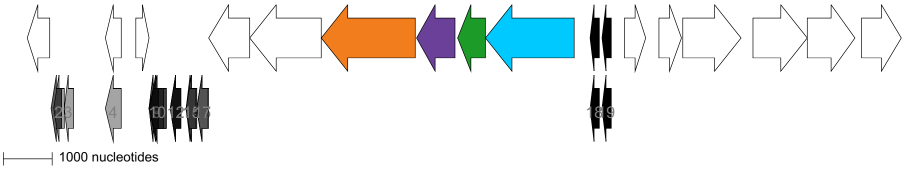

Link to nucleotide sequence

| Accession | start | end | direction | length (aa) | Pfam/HMM | name | description | E-value |
|---|---|---|---|---|---|---|---|---|
| WP_150528057.1 | 2475047 | 2474591 | - | 151 | PF13274 | DUF4065 | Protein of unknown function (DUF4065) | 1.20E-21 |
| WP_150528059.1 | 2476531 | 2476195 | - | 111 | NO PFAM MATCH | - | - | - |
| WP_150528060.1 | 2476807 | 2477077 | + | 89 | NO PFAM MATCH | - | - | - |
| WP_132756794.1 | 2479152 | 2478300 | - | 283 | PF01551 | Peptidase_M23 | Peptidase family M23 | 6.00E-18 |

| | | | | | | | | |
|---|---|---|---|---|---|---|---|---|
| WP_150528062.1 | 2480612 | 2479148 | - | 487 | TIGR02734 | TIGR02734 | crtI_fam: phytoene desaturase | 5.80E-12 |
| | | | | | TIGR03467 | TIGR03467 | HpnE: squalene-associated FAD-dependent desaturase | 2.40E-10 |
| | | | | | PF13450 | NAD_binding_8 | NAD(P)-binding Rossmann-like domain | 6.30E-10 |
| | | | | | PF01593 | Amino_oxidase | Flavin containing amine oxidoreductase | 1.10E-08 |
| | | | | | TIGR01372 | TIGR01372 | soxA: sarcosine oxidase, alpha subunit family | 1.70E-08 |
| WP_150528063.1 | 2482558 | 2480608 | - | 649 | TIGR02204 | TIGR02204 | MsbA_rel: ABC transporter, permease/ATP-binding protein | 1.50E-100 |
| | | | | | TIGR02203 | TIGR02203 | MsbA_lipidA: lipid A export permease/ATP-binding protein MsbA | 8.90E-99 |
| | | | | | TIGR03796 | TIGR03796 | NHLM_micro_ABC1: NHLM bacteriocin system ABC transporter, peptidase/ATP-binding protein | 1.20E-85 |
| | | | | | TIGR00958 | TIGR00958 | 3a01208: antigen peptide transporter 2 | 1.20E-82 |
| | | | | | TIGR01846 | TIGR01846 | type_I_sec_HlyB: type I secretion system ATPase | 2.20E-80 |
| WP_150528064.1 | 2483359 | 2482579 | - | 259 | PF13649 | Methyltransf_25 | Methyltransferase domain | 7.50E-14 |
| | | | | | PF08241 | Methyltransf_11 | Methyltransferase domain | 2.60E-09 |
| | | | | | PF08242 | Methyltransf_12 | Methyltransferase domain | 1.50E-08 |
| | | | | | PF13489 | Methyltransf_23 | Methyltransferase domain | 5.60E-06 |
| | | | | | PF13847 | Methyltransf_31 | Methyltransferase domain | 5.90E-06 |
| WP_099889448.1 | 2483996 | 2483408 | - | 195 | LinL | LinL | LinL | 2.20E-70 |
| WP_132756788.1 | 2485795 | 2483992 | - | 600 | LinH | LinH | LinH (Fusion of LinE/LinG) | 6.00E-231 |
| | | | | | LinE | LinE | LinE Homolog | 5.10E-101 |
| | | | | | LinG | LinG | LinG Homolog | 1.50E-96 |
| | | | | | PF12697 | Abhydrolase_6 | Alpha/beta hydrolase family | 2.00E-16 |
| | | | | | PF12146 | Hydrolase_4 | Serine aminopeptidase, S33 | 5.20E-10 |
| WP_132756786.1 | 2486317 | 2486116 | - | 66 | LinA | LinA | LinA (linaridin precursor) | 7.70E-11 |
| WP_048476417.1 | 2486566 | 2486371 | - | 64 | LinA | LinA | LinA (linaridin precursor) | 1.20E-12 |
| WP_132757254.1 | 2486833 | 2487271 | + | 145 | PF13560 | HTH_31 | Helix-turn-helix domain | 2.80E-04 |
| WP_165937276.1 | 2487544 | 2488015 | + | 156 | PF06114 | Peptidase_M78 | IrrE N-terminal-like domain | 2.60E-05 |
| WP_150528066.1 | 2488027 | 2489233 | + | 401 | NO PFAM MATCH | - | - | - |
| WP_150528067.1 | 2489470 | 2490577 | + | 368 | PF08028 | Acyl-CoA_dh_2 | Acyl-CoA dehydrogenase, C-terminal domain | 1.40E-26 |
| | | | | | TIGR04022 | TIGR04022 | sulfur_SfnB: sulfur acquisition oxidoreductase, SfnB family | 3.20E-22 |
| | | | | | PF00441 | Acyl-CoA_dh_1 | Acyl-CoA dehydrogenase, C-terminal domain | 1.70E-05 |
| | | | | | PF02770 | Acyl-CoA_dh_M | Acyl-CoA dehydrogenase, middle domain | 2.60E-04 |
| WP_150528068.1 | 2490573 | 2491560 | + | 328 | PF00248 | Aldo_ket_red | Aldo/keto reductase family | 1.50E-75 |
| | | | | | TIGR01293 | TIGR01293 | Kv_beta: voltage-dependent potassium channel beta subunit | 5.30E-27 |
| WP_132756779.1 | 2491696 | 2492506 | + | 269 | PF03704 | BTAD | Bacterial transcriptional activator domain | 2.10E-41 |
| | | | | | PF00486 | Trans_reg_C | Transcriptional regulatory protein, C terminal | 9.60E-11 |

| index | leader | core | start | end | dir | score |
|---|---|---|---|---|---|---|
| 1 | LVSLPRKPHVRGLACHYTGNPSDPTFEGSHCVHIF | TRPLTSVSAGGYAETVWGRASSATPPANTTGRRLV | 2475291 | 2475078 | - | 8 |
| 2 | VPETWNRLSPRSIVWIFGQPAAKAPCQRPRVS | LHRQSVRSYVRRVALRPHLHPPTYECQCRWIR | 2475340 | 2475145 | - | 6 |
| 3 | MVSPRRAAGKAQTRRTGAPPAHGFGVSVGLGACA | GHGQRASRPQAVLAKPVRYRGGRSTAPPAAQRWL | 2475547 | 2475340 | - | 6 |
| 4 | MATELVGIVGAGRESQRSPDRAVAF...NRDVDRVEAVKHCRAEPVDLEAPLR | HTLVVHKDLDGHLWGQFAVEAALLDPSYSIGGGSSGLCCAVS | 2476531 | 2476195 | - | 6 |
| 5 | MGAETLGWERAEELLRERPQPIRIGQITASDRLWV | YTFFLTADGRELVACAGVRRADRIPDQ | 2477283 | 2477094 | - | 6 |
| 6 | MQRMGAETLGWERAEELLRERPQPIRIGQITASDRLWV | YTFFLTADGRELVACAGVRRADRIPDQ | 2477292 | 2477094 | - | 6 |
| 7 | LAALADGAEYDVWDAGTRLSDGHAG...AEELLRERPQPIRIGQITASDRLWV | YTFFLTADGRELVACAGVRRADRIPDQ | 2477391 | 2477094 | - | 7 |
| 8 | MRLLAGDDAASTAALAALADGAEYD...AEELLRERPQPIRIGQITASDRLWV | YTFFLTADGRELVACAGVRRADRIPDQ | 2477433 | 2477094 | - | 6 |
| 9 | LMRLLAGDDAASTAALAALADGAEY...AEELLRERPQPIRIGQITASDRLWV | YTFFLTADGRELVACAGVRRADRIPDQ | 2477436 | 2477094 | - | 6 |
| 10 | MERHELMRLLAGDDAASTAALAALA...AEELLRERPQPIRIGQITASDRLWV | YTFFLTADGRELVACAGVRRADRIPDQ | 2477451 | 2477094 | - | 6 |
| 11 | MRCIVGHKVPLLSQGPRNSNHPGAWGPSLEDRARAHPVTSQPPDLHVPTR | STTFPAKPRPLGHAGDTAL | 2477744 | 2477534 | - | 9 |
| 12 | VSMRCIVGHKVPLLSQGPRNSNHPG...GPSLEDRARAHPVTSQPPDLHVPTR | STTFPAKPRPLGHAGDTAL | 2477750 | 2477534 | - | 8 |
| 13 | VPPAGFEPAHTAPEAPCVSRVPAGQATTSALPDHCRPRIGRKPRR | QLPQLINDVARRVIPTGNCRSEGE | 2478050 | 2477840 | - | 7 |
| 14 | LGTSCEVPPAGFEPAHTAPEAPCVS...VPAGQATTSALPDHCRPRIGRKPRR | QLPQLINDVARRVIPTGNCRSEGE | 2478068 | 2477840 | - | 6 |
| 15 | MAGSLGTSCEVPPAGFEPAHTAPEA...VPAGQATTSALPDHCRPRIGRKPRR | QLPQLINDVARRVIPTGNCRSEGE | 2478080 | 2477840 | - | 6 |
| 16 | VHGAENQSGGNCRRGRRGRCSPRSPAPAADHGLKNRCAEEARRVPLP | RPPTYGCTYAKTPSQRTR | 2478277 | 2478079 | - | 7 |
| 17 | VTSVLPGPAVHGAENQSGGNCRRGR...RSPAPAADHGLKNRCAEEARRVPLP | RPPTYGCTYAKTPSQRTR | 2478304 | 2478079 | - | 6 |
| 18 | MHSITESVRVIAGATLSEDSGEQTTRMMPVA | FTPAVLVTTPYAAGGLAAGAAAVGAYEAGRAAAKG | 2486317 | 2486116 | - | 29 |
| 19 | MAKIAAFMNQQDMGPIAESELSVGGTPVM | ATPAVVAVPASVKITAGIVAFTIPVGAYVTGRVVN | 2486566 | 2486371 | - | 30 |

### Results for AKG45062.1 [Streptomyces xiamenensis] back to top

previous - next

#### Architecture

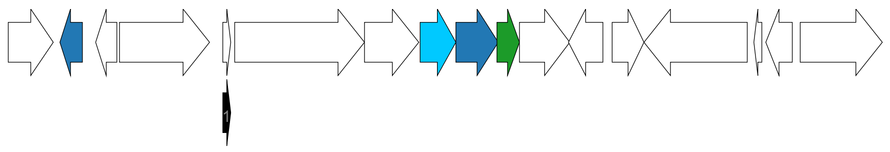

1000 nucleotides

Link to nucleotide sequence

| Accession | start | end | direction | length (aa) | Pfam/HMM | name | description | E-value |
|---|---|---|---|---|---|---|---|---|
| AKG45054.1 | 3917698 | 3918787 | + | 362 | TIGR00092<br>PF06071<br>PF01926<br>TIGR02729<br>TIGR03156 | TIGR00092<br>YchF-GTPase_C<br>MMR_HSR1<br>TIGR02729<br>TIGR03156 | TIGR00092: GTP-binding protein YchF<br>Protein of unknown function (DUF933)<br>50S ribosome-binding GTPase<br>Obg_CgtA: Obg family GTPase CgtA<br>GTP_HflX: GTP-binding protein HflX | 7.60E-130<br>1.20E-38<br>7.60E-23<br>8.70E-21<br>4.20E-11 |
| AKG45055.1 | 3919480 | 3918940 | - | 179 | PF12146<br>LinE<br>LinH<br>PF12697<br>TIGR03695 | Hydrolase_4<br>LinE<br>LinH<br>Abhydrolase_6<br>TIGR03695 | Serine aminopeptidase, S33<br>LinE Homolog<br>LinH (Fusion of LinE/LinG)<br>Alpha/beta hydrolase family<br>menH_SHCHC: 2-succinyl-6-hydroxy-2,4-cyclohexadiene-1-carboxylate synthase | 3.10E-12<br>7.00E-09<br>2.00E-07<br>4.50E-07<br>4.50E-04 |
| AKG45056.1 | 3920312 | 3919805 | - | 168 | PF12802<br>PF01047<br>PF13463<br>TIGR02337<br>TIGR04472 | MarR_2<br>MarR<br>HTH_27<br>TIGR02337<br>TIGR04472 | MarR family<br>MarR family<br>Winged helix DNA-binding domain<br>HpaR: homoprotocatechuate degradation operon regulator, HpaR<br>reg_rSAM_mob: mobile rSAM pair MarR family regulator | 1.00E-11<br>9.40E-10<br>7.50E-07<br>5.50E-05<br>5.70E-05 |
| AKG45057.1 | 3920385 | 3922545 | + | 719 | TIGR00833<br>PF03176<br>TIGR00921<br>TIGR03480<br>PF00873 | TIGR00833<br>MMPL<br>TIGR00921<br>TIGR03480<br>ACR_tran | actII: transport protein<br>MMPL family<br>2A067: efflux transporter, putative, hydrophobe/amphiphile efflux-3 (HAE3) family<br>HpnN: hopanoid biosynthesis associated RND transporter like protein HpnN<br>AcrB/AcrD/AcrF family | 7.70E-123<br>5.10E-111<br>3.10E-32<br>5.20E-21<br>2.50E-04 |
| AKG45058.1 | 3922891 | 3923074 | + | 60 | NO PFAM MATCH | - | - | - |
| AKG45059.1 | 3923176 | 3926299 | + | 1040 | PF04738<br>TIGR03891<br>PF14028 | Lant_dehydr_N<br>TIGR03891<br>Lant_dehydr_C | Lantibiotic dehydratase, N terminus<br>thiopep_ocin: thiopeptide-type bacteriocin biosynthesis domain<br>Lantibiotic biosynthesis dehydratase C-term | 1.20E-142<br>4.40E-42<br>5.80E-25 |
| AKG45060.1 | 3926295 | 3927603 | + | 435 | PF05147<br>TIGR03897 | LANC_like<br>TIGR03897 | Lanthionine synthetase C-like protein<br>lanti_2_LanM: type 2 lantibiotic biosynthesis protein LanM | 2.60E-45<br>5.10E-17 |
| AKG45061.1 | 3927644 | 3928523 | + | 292 | LinH<br>LinG | LinH<br>LinG | LinH (Fusion of LinE/LinG)<br>LinG Homolog | 6.00E-108<br>3.50E-101 |
| AKG45062.1 | 3928506 | 3929526 | + | 339 | LinE<br>LinH<br>PF12697<br>TIGR02427<br>PF12146 | LinE<br>LinH<br>Abhydrolase_6<br>TIGR02427<br>Hydrolase_4 | LinE Homolog<br>LinH (Fusion of LinE/LinG)<br>Alpha/beta hydrolase family<br>protocat_pcaD: 3-oxoadipate enol-lactonase<br>Serine aminopeptidase, S33 | 4.00E-90<br>1.90E-79<br>7.90E-17<br>7.00E-12<br>3.80E-11 |
| AKG45063.1 | 3929500 | 3930052 | + | 183 | LinL | LinL | LinL | 1.10E-55 |
| AKG45064.1 | 3930048 | 3931260 | + | 403 | TIGR04458<br>PF00067 | TIGR04458<br>p450 | CYP450_TxtE: 4-nitrotryptophan synthase<br>Cytochrome P450 | 4.90E-30<br>2.20E-27 |
| AKG45065.1 | 3932080 | 3931249 | - | 276 | PF00583<br>TIGR01575<br>PF13508<br>TIGR03448<br>PF13673 | Acetyltransf_1<br>TIGR01575<br>Acetyltransf_7<br>TIGR03448<br>Acetyltransf_10 | Acetyltransferase (GNAT) family<br>rimI: ribosomal-protein-alanine acetyltransferase<br>Acetyltransferase (GNAT) domain<br>mycothiol_MshD: mycothiol synthase<br>Acetyltransferase (GNAT) domain | 4.80E-14<br>7.30E-09<br>1.60E-06<br>9.60E-06<br>1.50E-05 |
| AKG45066.1 | 3932305 | 3933058 | + | 250 | PF00392<br>PF07702<br>TIGR02018<br>TIGR02325<br>TIGR02404 | GntR<br>UTRA<br>TIGR02018<br>TIGR02325<br>TIGR02404 | Bacterial regulatory proteins, gntR family<br>UTRA domain<br>his_ut_repres: histidine utilization repressor<br>C_P_lyase_phnF: phosphonate metabolism transcriptional regulator PhnF<br>trehalos_R_Bsub: trehalose operon repressor | 3.00E-16<br>2.30E-13<br>2.60E-13<br>1.50E-12<br>1.00E-06 |
| AKG45067.1 | 3935545 | 3933064 | - | 826 | PF07228<br>TIGR00229<br>PF08448<br>PF00989<br>PF13426 | SpoIIE<br>TIGR00229<br>PAS_4<br>PAS<br>PAS_9 | Stage II sporulation protein E (SpoIIE)<br>sensory_box: PAS domain S-box protein<br>PAS fold<br>PAS fold<br>PAS domain | 2.30E-51<br>8.10E-27<br>6.20E-25<br>8.50E-17<br>3.90E-14 |
| AKG45068.1 | 3935907 | 3935703 | - | 67 | PF05036 | SPOR | Sporulation related domain | 7.50E-04 |
| AKG45069.1 | 3936642 | 3936015 | - | 208 | PF04536 | TPM_phosphatase | TPM domain | 1.90E-12 |
| AKG45070.1 | 3936849 | 3938826 | + | 658 | PF00496<br>TIGR02294<br>TIGR04028 | SBP_bac_5<br>TIGR02294<br>TIGR04028 | Bacterial extracellular solute-binding proteins, family 5 Middle<br>nickel_nikA: nickel ABC transporter, nickel/metallophore periplasmic binding protein<br>SBP_KPN_01854: ABC transporter substrate binding protein, KPN_01854 family | 8.90E-44<br>2.00E-17<br>3.40E-17 |

| index | leader | core | start | end | dir | score |
|---|---|---|---|---|---|---|
| 1 | MGADLITAAANEELSDLDLDLTFAPNDENT | HVMLACAVDSMGNNVNTSRGIIPQAPVCCA | 3922891 | 3923074 | + | 13 |

### Results for AKG45482.1 [Streptomyces xiamenensis] back to top

previous - next

#### Architecture

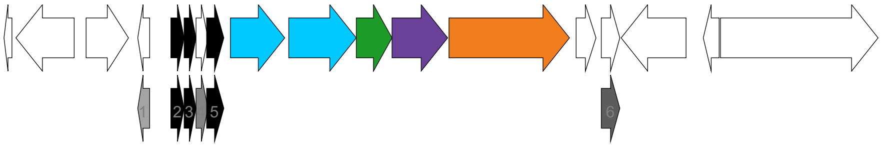

1000 nucleotides

Link to nucleotide sequence

| Accession | start | end | direction | length (aa) | Pfam/HMM | name | description | E-value |
|---|---|---|---|---|---|---|---|---|

| | | | | | | | | |
|---|---|---|---|---|---|---|---|---|
| AKG45473.1 | 4371646 | 4371526 | - | 39 | PF04149 | DUF397 | Domain of unknown function (DUF397) | 3.90E-12 |
| AKG45474.1 | 4372587 | 4371717 | - | 289 | PF13560<br>TIGR03830<br>PF01381<br>TIGR03070<br>PF12844 | HTH_31<br>TIGR03830<br>HTH_3<br>TIGR03070<br>HTH_19 | Helix-turn-helix domain<br>CxxCG_CxxCG_HTH: putative zinc finger/helix-turn-helix protein, YgiT family<br>Helix-turn-helix<br>couple_hipB: transcriptional regulator, y4mF family<br>Helix-turn-helix domain | 3.90E-15<br>4.10E-05<br>1.80E-04<br>2.50E-04<br>4.40E-04 |
| AKG45475.1 | 4372761 | 4373412 | + | 216 | PF13581<br>PF02518 | HATPase_c_2<br>HATPase_c | Histidine kinase-like ATPase domain<br>Histidine kinase-, DNA gyrase B-, and HSP90-like ATPase | 3.70E-14<br>5.50E-04 |
| AKG45476.1 | 4373722 | 4373548 | - | 57 | NO PFAM MATCH | - | - | - |
| AKG45477.1 | 4374049 | 4374244 | + | 64 | LinA | LinA | LinA (linaridin precursor) | 2.00E-13 |
| AKG45478.1 | 4374240 | 4374432 | + | 63 | LinA | LinA | LinA (linaridin precursor) | 5.10E-09 |
| AKG45479.1 | 4374428 | 4374599 | + | 56 | NO PFAM MATCH | - | - | - |
| AKG45480.1 | 4374595 | 4374841 | + | 81 | LinA | LinA | LinA (linaridin precursor) | 7.30E-15 |
| AKG45481.1 | 4374935 | 4375769 | + | 277 | LinH<br>LinG | LinH<br>LinG | LinH (Fusion of LinE/LinG)<br>LinG Homolog | 9.80E-118<br>6.70E-107 |
| AKG45482.1 | 4375817 | 4376843 | + | 341 | LinH<br>LinE<br>PF12697<br>PF00975<br>TIGR03695 | LinH<br>LinE<br>Abhydrolase_6<br>Thioesterase<br>TIGR03695 | LinH (Fusion of LinE/LinG)<br>LinE Homolog<br>Alpha/beta hydrolase family<br>Thioesterase domain<br>menH_SHCHC: 2-succinyl-6-hydroxy-2,4-cyclohexadiene-1-carboxylate synthase | 3.20E-113<br>1.90E-97<br>5.40E-16<br>2.00E-09<br>3.20E-09 |
| AKG45483.1 | 4376839 | 4377385 | + | 181 | LinL | LinL | LinL | 3.30E-64 |
| AKG45484.1 | 4377381 | 4378209 | + | 275 | PF13649<br>PF08241<br>PF13489<br>PF08242<br>PF13847 | Methyltransf_25<br>Methyltransf_11<br>Methyltransf_23<br>Methyltransf_12<br>Methyltransf_31 | Methyltransferase domain<br>Methyltransferase domain<br>Methyltransferase domain<br>Methyltransferase domain<br>Methyltransferase domain | 2.50E-16<br>1.40E-10<br>8.90E-08<br>1.20E-07<br>8.00E-07 |
| AKG45485.1 | 4378237 | 4380052 | + | 604 | TIGR02204<br>TIGR02203<br>TIGR03797<br>TIGR00958<br>TIGR03796 | TIGR02204<br>TIGR02203<br>TIGR03797<br>TIGR00958<br>TIGR03796 | MsbA_rel: ABC transporter, permease/ATP-binding protein<br>MsbA_lipidA: lipid A export permease/ATP-binding protein MsbA<br>NHLM_micro_ABC2: NHLM bacteriocin system ABC transporter, ATP-binding protein<br>3a01208: antigen peptide transporter 2<br>NHLM_micro_ABC1: NHLM bacteriocin system ABC transporter, peptidase/ATP-binding protein | 6.70E-104<br>3.70E-101<br>5.80E-89<br>4.70E-88<br>3.20E-87 |
| AKG45486.1 | 4380152 | 4380446 | + | 97 | NO PFAM MATCH | - | - | - |
| AKG45487.1 | 4380523 | 4380805 | + | 93 | NO PFAM MATCH | - | - | - |
| AKG45488.1 | 4381805 | 4380821 | - | 327 | PF00196 | GerE | Bacterial regulatory proteins, luxR family | 3.20E-04 |
| AKG45489.1 | 4382304 | 4382061 | - | 80 | NO PFAM MATCH | - | - | - |
| AKG45490.1 | 4382319 | 4384686 | + | 788 | PF07510<br>PF03235<br>PF01035 | DUF1524<br>DUF262<br>DNA_binding_1 | Protein of unknown function (DUF1524)<br>Protein of unknown function DUF262<br>6-O-methylguanine DNA methyltransferase, DNA binding domain | 4.80E-29<br>9.30E-17<br>7.80E-11 |

| index | leader | core | start | end | dir | score |
|---|---|---|---|---|---|---|
| 1 | MGDFNGSWAARLMEWSNGTIDEQRARTL | VTEIYEAAAERAVRLHEEEVAQREAEREE | 4373722 | 4373548 | - | 6 |
| 2 | MAGPSLARQWAHLGPLHENSPGEVGRAASPVL | ATPSAFAAGMGIVVGAYVVGRMIGSETEPVIQ | 4374049 | 4374244 | + | 19 |
| 3 | MTAADRVPALRDLSPLTENSPGQAPGSATAVL | ATPTPAAMIVIAYAAFEVGRMVCGTTEPVIQ | 4374240 | 4374432 | + | 31 |
| 4 | MTAVDRIPAVLGEIGELNEEDGGRASAP | RAVVLCLHTSAYIAVSVVGDTEPSLPSL | 4374428 | 4374599 | + | 7 |
| 5 | MTGQEGHRIPMSRIAAYLDDSPVLATIPVLDRTGRVSPARPVL | ATPVTIVAFGAGVAAGSAIVAAFEAGRTAGSSTEPIMT | 4374595 | 4374841 | + | 29 |
| 6 | MTDGAGHGDGEGQGEGLDGEIALTL...RLLSLLPGHWTYRHRVTAARIMLWV | RTPAGAPAVHAALRAALADPALRGWLPDSD | 4380523 | 4380805 | + | 8 |

### Results for SEG54329.1 [Streptomyces yanglinensis] back to top

previous - next

#### Architecture

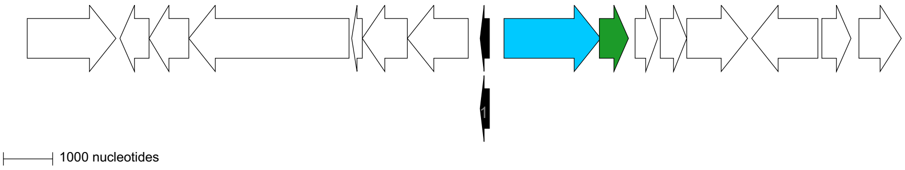

Link to nucleotide sequence

| Accession | start | end | direction | length (aa) | Pfam/HMM | name | description | E-value |
|---|---|---|---|---|---|---|---|---|
| SEG54186.1 | 129468 | 131271 | + | 600 | PF03704<br>PF00486<br>PF00931<br>PF13191 | BTAD<br>Trans_reg_C<br>NB-ARC<br>AAA_16 | Bacterial transcriptional activator domain<br>Transcriptional regulatory protein, C terminal<br>NB-ARC domain<br>AAA ATPase domain | 5.20E-30<br>2.20E-09<br>3.10E-07<br>4.80E-05 |
| SEG54213.1 | 131932 | 131350 | - | 193 | PF13185<br>PF01590<br>PF13492<br>TIGR02916 | GAF_2<br>GAF<br>GAF_3<br>TIGR02916 | GAF domain<br>GAF domain<br>GAF domain<br>PEP_his_kin: putative PEP-CTERM system histidine kinase | 1.20E-19<br>4.70E-18<br>3.50E-13<br>7.10E-04 |
| SEG54229.1 | 132753 | 131940 | - | 270 | TIGR03921<br>PF00082<br>TIGR03895 | TIGR03921<br>Peptidase_S8<br>TIGR03895 | T7SS_mycosin: type VII secretion-associated serine protease mycosin<br>Subtilase family<br>protease_PatA: cyanobactin maturation protease, PatA/PatG family | 3.80E-33<br>1.80E-23<br>1.30E-07 |

| | | | | | | | | |
|---|---|---|---|---|---|---|---|---|
| SEG54247.1 | 136001 | 132737 | - | 1087 | TIGR03903<br>PF00211<br>PF13191<br>PF13424<br>PF13401 | TIGR03903<br>Guanylate_cyc<br>AAA_16<br>TPR_12<br>AAA_22 | TOMM_kin_cyc: TOMM system kinase/cyclase fusion protein<br>Adenylate and Guanylate cyclase catalytic domain<br>AAA ATPase domain<br>Tetratricopeptide repeat<br>AAA domain | 2.30E-46<br>5.10E-30<br>1.10E-22<br>3.80E-09<br>1.40E-05 |
| SEG54263.1 | 136267 | 136051 | - | 71 | NO PFAM MATCH | - | - | - |
| SEG54278.1 | 137175 | 136263 | - | 303 | PF14518 | Haem_oxygenas_2 | Iron-containing redox enzyme | 3.50E-07 |
| SEG54296.1 | 138407 | 137171 | - | 411 | PF04055<br>TIGR02109<br>TIGR02666<br>TIGR04051<br>TIGR04055 | Radical_SAM<br>TIGR02109<br>TIGR02666<br>TIGR04051<br>TIGR04055 | Radical SAM superfamily<br>PQQ_syn_pqqE: coenzyme PQQ biosynthesis enzyme PqqE<br>moaA: molybdenum cofactor biosynthesis protein A<br>rSAM_NirJ: heme d1 biosynthesis radical SAM protein NirJ<br>rSAM_NirJ2: putative heme d1 biosynthesis radical SAM protein NirJ2 | 7.30E-11<br>3.50E-07<br>6.20E-07<br>2.80E-06<br>4.90E-06 |
| SEG54314.1 | 138830 | 138662 | - | 55 | LinA | LinA | LinA (linaridin precursor) | 2.50E-05 |
| SEG54329.1 | 139121 | 141095 | + | 657 | LinH<br>LinG<br>LinE<br>PF12697<br>PF12146 | LinH<br>LinG<br>LinE<br>Abhydrolase_6<br>Hydrolase_4 | LinH (Fusion of LinE/LinG)<br>LinG Homolog<br>LinE Homolog<br>Alpha/beta hydrolase family<br>Serine aminopeptidase, S33 | 5.30E-194<br>8.30E-102<br>7.10E-71<br>1.10E-08<br>4.30E-08 |
| SEG54348.1 | 141075 | 141669 | + | 197 | LinL | LinL | LinL | 2.80E-63 |
| SEG54365.1 | 141804 | 142242 | + | 145 | PF13560 | HTH_31 | Helix-turn-helix domain | 5.40E-06 |
| SEG54382.1 | 142292 | 142847 | + | 184 | NO PFAM MATCH | - | - | - |
| SEG54398.1 | 142839 | 144084 | + | 414 | NO PFAM MATCH | - | - | - |
| SEG54413.1 | 145497 | 144144 | - | 450 | PF07690<br>TIGR00711<br>TIGR00880<br>TIGR00710<br>TIGR00895 | MFS_1<br>TIGR00711<br>TIGR00880<br>TIGR00710<br>TIGR00895 | Major Facilitator Superfamily<br>efflux_EmrB: drug resistance MFS transporter, drug:H+ antiporter-2 (14 Spanner) (DHA2) family<br>2_A_01_02: multidrug resistance protein<br>efflux_Bcr_CflA: drug resistance transporter, Bcr/CflA subfamily<br>2A0115: MFS transporter, aromatic acid:H+ symporter (AAHS) family | 9.90E-41<br>2.40E-22<br>1.20E-15<br>4.80E-15<br>5.20E-13 |
| SEG54430.1 | 145585 | 146164 | + | 192 | PF01047<br>PF12802<br>PF13463<br>TIGR02337<br>PF09339 | MarR<br>MarR_2<br>HTH_27<br>TIGR02337<br>HTH_IclR | MarR family<br>MarR family<br>Winged helix DNA-binding domain<br>HpaR: homoprotocatechuate degradation operon regulator, HpaR<br>IclR helix-turn-helix domain | 3.60E-15<br>2.00E-11<br>8.80E-10<br>1.50E-09<br>1.60E-04 |
| SEG54447.1 | 146342 | 147194 | + | 283 | PF04954<br>PF08021 | SIP<br>FAD_binding_9 | Siderophore-interacting protein<br>Siderophore-interacting FAD-binding domain | 4.10E-39<br>4.00E-34 |

| index | leader | core | start | end | dir | score |
|---|---|---|---|---|---|---|
| 1 | MSDLLERANANGALA | AIPLHAPEVQLMSHPDLAPILCVHLVTSSVCISVCVNTGQ | 138830 | 138662 | - | 12 |

### Results for WP_016907298.1 [Streptomyces xiaopingdaonensis] back to top

previous - next

#### Architecture

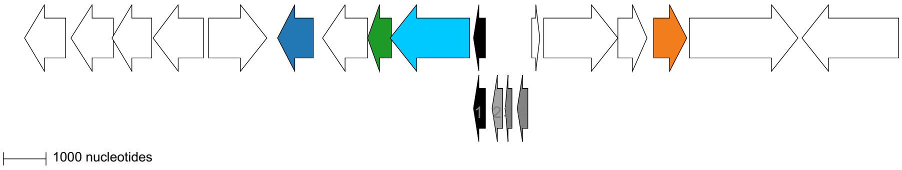

Link to nucleotide sequence

| Accession | start | end | direction | length (aa) | Pfam/HMM | name | description | E-value |
|---|---|---|---|---|---|---|---|---|
| WP_016907290.1 | 8889 | 7920 | - | 322 | TIGR02160<br>PF00111<br>PF00175<br>TIGR02008<br>TIGR01941 | TIGR02160<br>Fer2<br>NAD_binding_1<br>TIGR02008<br>TIGR01941 | PA_CoA_Oxy5: phenylacetate-CoA oxygenase/reductase, PaaK subunit<br>2Fe-2S iron-sulfur cluster binding domain<br>Oxidoreductase NAD-binding domain<br>fdx_plant: ferredoxin [2Fe-2S]<br>nqrF: NADH:ubiquinone oxidoreductase, Na(+)-translocating, F subunit | 1.10E-25<br>3.20E-11<br>4.10E-07<br>6.60E-05<br>7.00E-05 |
| WP_016907291.1 | 10004 | 9005 | - | 332 | PF02737<br>TIGR02279<br>TIGR02437<br>PF00725<br>TIGR02441 | 3HCDH_N<br>TIGR02279<br>TIGR02437<br>3HCDH<br>TIGR02441 | 3-hydroxyacyl-CoA dehydrogenase, NAD binding domain<br>PaaC-3OHAcCoADH: 3-hydroxyacyl-CoA dehydrogenase PaaC<br>FadB: fatty oxidation complex, alpha subunit FadB<br>3-hydroxyacyl-CoA dehydrogenase, C-terminal domain<br>fa_ox_alpha_mit: fatty acid oxidation complex, alpha subunit, mitochondrial | 6.60E-40<br>4.90E-24<br>7.30E-17<br>4.20E-16<br>1.90E-14 |
| WP_016907292.1 | 10920 | 9990 | - | 309 | PF05853 | BKACE | beta-keto acid cleavage enzyme | 5.70E-101 |
| WP_029391353.1 | 12125 | 10958 | - | 388 | TIGR03228<br>PF00848<br>TIGR03229<br>PF00355<br>TIGR02378 | TIGR03228<br>Ring_hydroxyl_A<br>TIGR03229<br>Rieske<br>TIGR02378 | anthran_1_2_A: anthranilate 1,2-dioxygenase, large subunit<br>Ring hydroxylating alpha subunit (catalytic domain)<br>benzo_1_2_benA: benzoate 1,2-dioxygenase, large subunit<br>Rieske [2Fe-2S] domain<br>nirD_assim_sml: nitrite reductase [NAD(P)H], small subunit | 1.20E-28<br>3.60E-27<br>2.00E-19<br>5.50E-19<br>7.10E-07 |
| WP_016907294.1 | 12257 | 13619 | + | 453 | PF00171<br>TIGR01780<br>TIGR01804<br>TIGR02299<br>TIGR04284 | Aldedh<br>TIGR01780<br>TIGR01804<br>TIGR02299<br>TIGR04284 | Aldehyde dehydrogenase family<br>SSADH: succinate-semialdehyde dehydrogenase<br>BADH: betaine-aldehyde dehydrogenase<br>HpaE: 5-carboxymethyl-2-hydroxymuconate semialdehyde dehydrogenase<br>aldehy_Rv0768: aldehyde dehydrogenase, Rv0768 family | 2.50E-128<br>5.90E-101<br>9.60E-93<br>1.90E-81<br>4.60E-77 |
| WP_016907295.1 | 14695 | 13873 | - | 273 | TIGR02427<br>PF12697<br>TIGR03695<br>TIGR03056<br>TIGR03611 | TIGR02427<br>Abhydrolase_6<br>TIGR03695<br>TIGR03056<br>TIGR03611 | protocat_pcaD: 3-oxoadipate enol-lactonase<br>Alpha/beta hydrolase family<br>menH_SHCHC: 2-succinyl-6-hydroxy-2,4-cyclohexadiene-1-carboxylate synthase<br>bchO_mg_che_rel: putative magnesium chelatase accessory protein<br>RutD: pyrimidine utilization protein D | 4.90E-28<br>4.20E-15<br>5.80E-14<br>6.00E-14<br>2.70E-13 |

| | | | | | | | | |
|---|---|---|---|---|---|---|---|---|
| WP_016907296.1 | 15975 | 14913 | - | 353 | PF00891<br>TIGR02716<br>PF16864 | Methyltransf_2<br>TIGR02716<br>Dimerisation2 | O-methyltransferase domain<br>C20_methyl_CrtF: C-20 methyltransferase BchU<br>Dimerisation domain | 4.90E-38<br>9.00E-11<br>5.20E-05 |
| WP_016907297.1 | 16523 | 15971 | - | 183 | LinL | LinL | LinL | 1.80E-58 |
| WP_016907298.1 | 18376 | 16519 | - | 618 | LinH<br>LinE<br>LinG<br>TIGR02427<br>PF00561 | LinH<br>LinE<br>LinG<br>TIGR02427<br>Abhydrolase_1 | LinH (Fusion of LinE/LinG)<br>LinE Homolog<br>LinG Homolog<br>protocat_pcaD: 3-oxoadipate enol-lactonase<br>alpha/beta hydrolase fold | 1.80E-200<br>3.50E-94<br>9.80E-93<br>1.70E-11<br>2.20E-10 |
| WP_033196079.1 | 18733 | 18451 | - | 93 | LinA | LinA | LinA (linaridin precursor) | 1.90E-11 |
| WP_016907302.1 | 19827 | 20019 | + | 63 | NO PFAM MATCH | - | - | - |
| WP_016907303.1 | 20119 | 21841 | + | 573 | PF07730 | HisKA_3 | Histidine kinase | 5.50E-14 |
| WP_016907304.1 | 21837 | 22524 | + | 228 | PF00072<br>TIGR02875<br>PF00196<br>TIGR01818<br>TIGR02154 | Response_reg<br>TIGR02875<br>GerE<br>TIGR01818<br>TIGR02154 | Response regulator receiver domain<br>spore_0_A: sporulation transcription factor Spo0A<br>Bacterial regulatory proteins, luxR family<br>ntrC: nitrogen regulation protein NR(I)<br>PhoB: phosphate regulon transcriptional regulatory protein PhoB | 5.00E-27<br>7.60E-20<br>1.60E-12<br>3.50E-12<br>5.80E-12 |
| WP_016907305.1 | 22690 | 23476 | + | 261 | TIGR02211<br>TIGR02982<br>TIGR02673<br>TIGR03608<br>TIGR02315 | TIGR02211<br>TIGR02982<br>TIGR02673<br>TIGR03608<br>TIGR02315 | LolD_lipo_ex: lipoprotein releasing system, ATP-binding protein<br>heterocyst_DevA: ABC exporter ATP-binding subunit, DevA family<br>FtsE: cell division ATP-binding protein FtsE<br>L_ocin_972_ABC: putative bacteriocin export ABC transporter, lactococcin 972 group<br>ABC_phnC: phosphonate ABC transporter, ATP-binding protein | 1.10E-58<br>1.70E-57<br>8.70E-56<br>3.70E-53<br>5.10E-48 |
| WP_016907306.1 | 23526 | 26085 | + | 852 | TIGR03434<br>TIGR02212<br>PF12704<br>PF02687<br>TIGR02213 | TIGR03434<br>TIGR02212<br>MacB_PCD<br>FtsX<br>TIGR02213 | ADOP: acidobacterial duplicated orphan permease<br>lolCE: lipoprotein releasing system, transmembrane protein, LolC/E family<br>MacB-like periplasmic core domain<br>FtsX-like permease family<br>lolE_release: lipoprotein releasing system, transmembrane protein LolE | 1.30E-30<br>1.50E-29<br>1.60E-22<br>4.40E-21<br>4.00E-09 |
| WP_016907307.1 | 28419 | 26154 | - | 754 | PF00199<br>PF18011<br>PF06628 | Catalase<br>Catalase_C<br>Catalase-rel | Catalase<br>C-terminal domain found in long catalases<br>Catalase-related immune-responsive | 1.00E-163<br>2.90E-31<br>8.00E-22 |

| index | leader | core | start | end | dir | score |
|---|---|---|---|---|---|---|
| 1 | MAVSRTTRIADWAQHHAEFGERHVPRSEEPLPVL | LTPAIVFAGVTAAQTAAQVGLTAAA...STVPSTALDTGRSVGDTLAHHPQRF | 18733 | 18451 | - | 18 |
| 2 | VNAARGDPLTRSCRARGAGGGGFPSRGLLQHPFDVRKLP | PGRRAGPLRGHATTVATTPKRVAGASSRARLRAPQRYAR | 19141 | 18904 | - | 6 |
| 3 | MSTGRDREPRLDAAEANDAIRRFVAG | RTVWTAADLVELDRLRTEWRAARRGCTAAA | 19374 | 19203 | - | 7 |
| 4 | VIGAGATVLPFLGAAARIDVLAHDE...RLGADPSGRHRRSRCRDRRGRGPGA | PTARATAPTCSRRSSR | 19735 | 19498 | - | 7 |

### Results for SHM24347.1 [Streptomyces yunnanensis] back to top

previous - next

#### Architecture

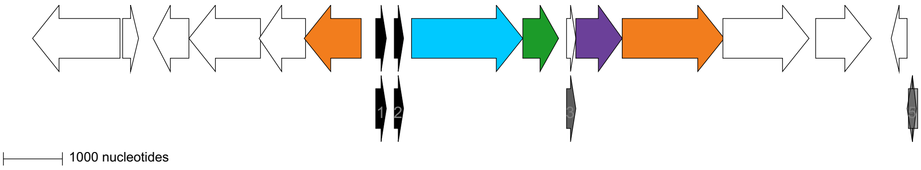

Link to nucleotide sequence

| Accession | start | end | direction | length (aa) | Pfam/HMM | name | description | E-value |
|---|---|---|---|---|---|---|---|---|
| SHM24093.1 | 232338 | 230844 | - | 497 | PF00135<br>PF07859 | COesterase<br>Abhydrolase_3 | Carboxylesterase family<br>alpha/beta hydrolase fold | 3.10E-84<br>2.80E-11 |
| SHM24116.1 | 232380 | 232641 | + | 86 | NO PFAM MATCH | - | - | - |
| SHM24146.1 | 233493 | 232887 | - | 201 | PF00072<br>TIGR02875<br>PF00196<br>TIGR02154<br>TIGR02956 | Response_reg<br>TIGR02875<br>GerE<br>TIGR02154<br>TIGR02956 | Response regulator receiver domain<br>spore_0_A: sporulation transcription factor Spo0A<br>Bacterial regulatory proteins, luxR family<br>PhoB: phosphate regulon transcriptional regulatory protein PhoB<br>TMAO_torS: TMAO reductase sytem sensor TorS | 6.50E-22<br>2.50E-16<br>7.20E-13<br>3.00E-08<br>4.70E-08 |
| SHM24177.1 | 234701 | 233489 | - | 403 | PF07730 | HisKA_3 | Histidine kinase | 7.50E-15 |
| SHM24208.1 | 235465 | 234697 | - | 255 | PF01061<br>PF12698<br>TIGR03062<br>TIGR01291<br>TIGR01247 | ABC2_membrane<br>ABC2_membrane_3<br>TIGR03062<br>TIGR01291<br>TIGR01247 | ABC-2 type transporter<br>ABC-2 family transporter protein<br>pip_yhgE_Cterm: YhgE/Pip C-terminal domain<br>nodJ: ABC-2 type transporter, NodJ family<br>drrB: daunorubicin resistance ABC transporter membrane protein | 1.30E-11<br>1.90E-09<br>2.60E-06<br>6.00E-05<br>7.40E-05 |
| SHM24233.1 | 236415 | 235461 | - | 317 | TIGR01188<br>TIGR03864<br>TIGR01288<br>TIGR04406<br>TIGR03740 | TIGR01188<br>TIGR03864<br>TIGR01288<br>TIGR04406<br>TIGR03740 | drrA: daunorubicin resistance ABC transporter, ATP-binding protein<br>PQQ_ABC_ATP: ABC transporter, ATP-binding subunit, PQQ-dependent alcohol dehydrogenase system<br>nodI: nodulation ABC transporter NodI<br>LPS_export_lptB: LPS export ABC transporter ATP-binding protein<br>galliderm_ABC: lantibiotic protection ABC transporter, ATP-binding subunit | 4.40E-62<br>9.20E-44<br>1.90E-41<br>1.60E-31<br>1.90E-31 |
| SHM24264.1 | 236654 | 236855 | + | 66 | LinA | LinA | LinA (linaridin precursor) | 1.80E-13 |
| SHM24314.1 | 236970 | 237132 | + | 53 | LinA | LinA | LinA (linaridin precursor) | 1.30E-14 |

| | | | | | | | | |
|---|---|---|---|---|---|---|---|---|
| SHM24347.1 | 237268 | 239158 | + | 629 | LinH<br>LinG<br>LinE<br>PF12697<br>PF12146 | LinH<br>LinG<br>LinE<br>Abhydrolase_6<br>Hydrolase_4 | LinH (Fusion of LinE/LinG)<br>LinG Homolog<br>LinE Homolog<br>Alpha/beta hydrolase family<br>Serine aminopeptidase, S33 | 8.30E-251<br>9.80E-129<br>8.00E-99<br>5.20E-16<br>1.40E-07 |
| SHM24381.1 | 239154 | 239766 | + | 203 | LinL | LinL | LinL | 9.60E-68 |
| SHM24408.1 | 239904 | 240063 | + | 52 | NO PFAM MATCH | - | - | - |
| SHM24437.1 | 240059 | 240851 | + | 263 | PF13649<br>PF08241<br>PF08242<br>PF13489<br>PF13847 | Methyltransf_25<br>Methyltransf_11<br>Methyltransf_12<br>Methyltransf_23<br>Methyltransf_31 | Methyltransferase domain<br>Methyltransferase domain<br>Methyltransferase domain<br>Methyltransferase domain<br>Methyltransferase domain | 5.10E-16<br>7.60E-12<br>5.30E-09<br>2.30E-07<br>5.40E-07 |
| SHM24471.1 | 240847 | 242560 | + | 570 | TIGR02204<br>TIGR02203<br>TIGR00958<br>TIGR01846<br>TIGR03796 | TIGR02204<br>TIGR02203<br>TIGR00958<br>TIGR01846<br>TIGR03796 | MsbA_rel: ABC transporter, permease/ATP-binding protein<br>MsbA_lipidA: lipid A export permease/ATP-binding protein MsbA<br>3a01208: antigen peptide transporter 2<br>type_I_sec_HlyB: type I secretion system ATPase<br>NHLM_micro_ABC1: NHLM bacteriocin system ABC transporter, peptidase/ATP-binding protein | 1.90E-104<br>2.30E-99<br>9.40E-92<br>6.20E-87<br>2.80E-85 |
| SHM24501.1 | 242556 | 244011 | + | 484 | TIGR02734<br>PF13450<br>TIGR00292<br>TIGR02733<br>TIGR03467 | TIGR02734<br>NAD_binding_8<br>TIGR00292<br>TIGR02733<br>TIGR03467 | crtI_fam: phytoene desaturase<br>NAD(P)-binding Rossmann-like domain<br>TIGR00292: thiazole biosynthesis enzyme<br>desat_CrtD: C-3',4' desaturase CrtD<br>HpnE: squalene-associated FAD-dependent desaturase | 1.80E-11<br>1.70E-08<br>7.10E-08<br>2.80E-07<br>2.90E-07 |
| SHM24530.1 | 244129 | 245071 | + | 313 | PF03752 | ALF | Short repeats of unknown function | 2.70E-43 |
| SHM24556.1 | 245664 | 245406 | - | 85 | PF07510 | DUF1524 | Protein of unknown function (DUF1524) | 1.10E-04 |

| index | leader | core | start | end | dir | score |
|---|---|---|---|---|---|---|
| 1 | MTEIVSFVNGLDLGAIETGEAHSDQAPVL | ATPAAALAAIGGAKAVGAFVGGATVVGGAYALGRAVG | 236654 | 236855 | + | 30 |
| 2 | MIDVAAAANEMEIPEEGAIFAVATPVL | ATPAAFMGGIAISSAVVGAFEAGRNG | 236970 | 237132 | + | 29 |
| 3 | MNKSAVAGCLAVGLAVLGF | LTATGTVNIPIAMGAGVALFIAAGVEIKRKPTR | 239904 | 240063 | + | 8 |
| 4 | LNLRPLGYEPSELPSC | STPRRHPEPYGNDTDPSSRPRIGVTASPG | 245686 | 245824 | + | 6 |
| 5 | VRLPGPFSLDSRSRRSEVDSMDPCHCRRVQ | GAVAGWSSSVARWAHNPEVAGSNPVPATPM | 245843 | 245660 | - | 6 |

### Results for SHL65197.1 [Streptomyces yunnanensis] back to top

previous - next

#### Architecture

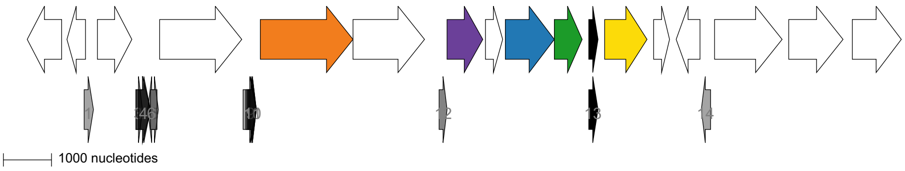

Link to nucleotide sequence

| Accession | start | end | direction | length (aa) | Pfam/HMM | name | description | E-value |
|---|---|---|---|---|---|---|---|---|
| SHL64916.1 | 384142 | 383428 | - | 237 | PF01810<br>TIGR00949<br>TIGR00948 | LysE<br>TIGR00949<br>TIGR00948 | LysE type translocator<br>2A76: homoserine/Threonine efflux protein<br>2a75: L-lysine exporter | 1.50E-42<br>2.10E-39<br>4.70E-14 |
| SHL64941.1 | 384636 | 384267 | - | 122 | PF01906<br>TIGR03884 | YbjQ_1<br>TIGR03884 | Putative heavy-metal-binding<br>sel_bind_Methan: selenium-binding protein | 1.30E-33<br>7.80E-04 |
| SHL64978.1 | 384896 | 385607 | + | 236 | PF09335 | SNARE_assoc | SNARE associated Golgi protein | 2.00E-24 |
| SHL65011.1 | 386189 | 387902 | + | 570 | PF06738<br>PF12821 | ThrE<br>ThrE_2 | Putative threonine/serine exporter<br>Threonine/Serine exporter, ThrE | 4.50E-73<br>2.70E-20 |
| SHL65045.1 | 388276 | 390214 | + | 645 | TIGR02204<br>TIGR02203<br>TIGR00958<br>TIGR03796<br>TIGR01846 | TIGR02204<br>TIGR02203<br>TIGR00958<br>TIGR03796<br>TIGR01846 | MsbA_rel: ABC transporter, permease/ATP-binding protein<br>MsbA_lipidA: lipid A export permease/ATP-binding protein MsbA<br>3a01208: antigen peptide transporter 2<br>NHLM_micro_ABC1: NHLM bacteriocin system ABC transporter, peptidase/ATP-binding protein<br>type_I_sec_HlyB: type I secretion system ATPase | 6.60E-97<br>1.00E-94<br>1.70E-88<br>2.10E-80<br>1.60E-77 |
| SHL65083.1 | 390210 | 391695 | + | 494 | TIGR03467<br>PF13450<br>TIGR00292<br>TIGR02734<br>TIGR03315 | TIGR03467<br>NAD_binding_8<br>TIGR00292<br>TIGR02734<br>TIGR03315 | HpnE: squalene-associated FAD-dependent desaturase<br>NAD(P)-binding Rossmann-like domain<br>TIGR00292: thiazole biosynthesis enzyme<br>crtI_fam: phytoene desaturase<br>Se_ygfK: putative selenate reductase, YgfK subunit | 1.60E-09<br>1.30E-08<br>2.10E-08<br>9.80E-08<br>1.10E-06 |
| SHL65127.1 | 392159 | 392918 | + | 252 | PF13649<br>PF08241<br>PF13489<br>PF08242<br>PF13847 | Methyltransf_25<br>Methyltransf_11<br>Methyltransf_23<br>Methyltransf_12<br>Methyltransf_31 | Methyltransferase domain<br>Methyltransferase domain<br>Methyltransferase domain<br>Methyltransferase domain<br>Methyltransferase domain | 7.10E-16<br>9.60E-13<br>5.80E-10<br>6.70E-09<br>9.60E-09 |
| SHL65160.1 | 392956 | 393304 | + | 115 | PF01527<br>PF13518<br>PF13384<br>PF05344<br>PF13551 | HTH_Tnp_1<br>HTH_28<br>HTH_23<br>DUF746<br>HTH_29 | Transposase<br>Helix-turn-helix domain<br>Homeodomain-like domain<br>Domain of Unknown Function (DUF746)<br>Winged helix-turn helix | 1.30E-07<br>1.90E-07<br>4.80E-06<br>1.10E-05<br>3.70E-05 |

| | | | | | | | | |
|---|---|---|---|---|---|---|---|---|
| SHL65197.1 | 393376 | 394399 | + | 340 | LinE<br>LinH<br>PF12697<br>TIGR02427<br>PF00561 | LinE<br>LinH<br>Abhydrolase_6<br>TIGR02427<br>Abhydrolase_1 | LinE Homolog<br>LinH (Fusion of LinE/LinG)<br>Alpha/beta hydrolase family<br>protocat_pcaD: 3-oxoadipate enol-lactonase<br>alpha/beta hydrolase fold | 1.00E-102<br>8.20E-53<br>4.10E-15<br>8.30E-11<br>8.90E-09 |
| SHL65234.1 | 394395 | 394977 | + | 193 | LinL | LinL | LinL | 1.90E-57 |
| SHL65271.1 | 395105 | 395303 | + | 65 | LinA | LinA | LinA (linaridin precursor) | 1.20E-12 |
| SHL65302.1 | 395447 | 396308 | + | 286 | LinG<br>LinH<br>TIGR04033 | LinG<br>LinH<br>TIGR04033 | LinG Homolog<br>LinH (Fusion of LinE/LinG)<br>export_SdpB: antimicrobial peptide system protein, SdpB family | 1.60E-108<br>1.10E-76<br>7.60E-05 |
| SHL65338.1 | 396461 | 396788 | + | 108 | NO PFAM MATCH | - | - | - |
| SHL65370.1 | 397428 | 396936 | - | 163 | PF00719 | Pyrophosphatase | Inorganic pyrophosphatase | 4.50E-53 |
| SHL65408.1 | 397712 | 399131 | + | 472 | PF02113<br>TIGR00666 | Peptidase_S13<br>TIGR00666 | D-Ala-D-Ala carboxypeptidase 3 (S13) family<br>PBP4: D-alanyl-D-alanine carboxypeptidase/D-alanyl-D-alanine-endopeptidase | 2.60E-75<br>3.60E-57 |
| SHL65439.1 | 399269 | 400391 | + | 373 | TIGR03883<br>TIGR03624<br>PF10103 | TIGR03883<br>TIGR03624<br>Zincin_2 | DUF2342_F420: uncharacterized protein, coenzyme F420 biosynthesis associated<br>TIGR03624: putative hydrolase<br>Zincin-like metallopeptidase | 7.60E-141<br>2.20E-125<br>1.10E-120 |
| SHL65471.1 | 400584 | 401592 | + | 335 | TIGR02432<br>PF01171<br>PF09179 | TIGR02432<br>ATP_bind_3<br>TilS | lysidine_TilS_N: tRNA(Ile)-lysidine synthetase<br>PP-loop family<br>TilS substrate binding domain | 3.10E-43<br>5.20E-38<br>3.00E-12 |

| index | leader | core | start | end | dir | score |
|---|---|---|---|---|---|---|
| 1 | MPMVRTSCPCGQDHPGRPERPALGGGRVG | VGGRRRGGAGRPGRPWPGVFLEPRDVIVR | 384628 | 384805 | + | 6 |
| 2 | VFRCPLRGSAARGWSAAVPGLWGGGCAYCCALRP | HTPHPTDPSPPVDGWVSGRRGTPGW...GVGPTHRLSHPPTGGDGAEGVVCRT | 385695 | 385998 | + | 8 |
| 3 | MSAARVRCAGLVGRGAGSVGW | GVCVLLRFTSAHPPPHRPVPSR | 385703 | 385835 | + | 6 |
| 4 | VPGLWGGGCAYCCALRP | HTPHPTDPSPPVDGWVSGRRGTPGW...GVGPTHRLSHPPTGGDGAEGVVCRT | 385746 | 385998 | + | 8 |
| 5 | VQDVKRSSPAHHPLRP | VTAPDSPAQRTRHPRRQADTQRTGPRRSAVRSHAASCA | 385985 | 386150 | + | 6 |
| 6 | MAADGAPSGARPLRVRLTAGVPCPLRGAVGCGDGPEGVVCRTAALYVLH | TTPSAPSPPVGGWERR | 386132 | 385934 | - | 6 |
| 7 | VHRTDPQGPSDAPGPRPRCWVSRSPPGVPCVGGFVLLAREGTVVFLS | GTAPSGQQTRLLTPAGPAGHSTGQGVPANQLTRNLTPKGVLVSAILTTKL | 387918 | 388212 | + | 6 |
| 8 | VSRSPPGVPCVGGFVLLAREGTVVFLS | GTAPSGQQTRLLTPAGPAGHSTGQGVPANQLTRNLTPKGVLVSAILTTKL | 387978 | 388212 | + | 6 |
| 9 | VPCVGGFVLLAREGTVVFLS | GTAPSGQQTRLLTPAGPAGHSTGQGVPANQLTRNLTPKGVLVSAILTTKL | 387999 | 388212 | + | 8 |
| 10 | VVFLSGTAPSGQQTRLLTPAGPAGHST | GQGVPANQLTRNLTPKGVLVSAILTTKL | 388044 | 388212 | + | 6 |
| 11 | VFLSGTAPSGQQTRLLTPAGPAGHSTG | QGVPANQLTRNLTPKGVLVSAILTTKL | 388047 | 388212 | + | 6 |
| 12 | LRRFHWIDRRCWSDQPCA | TTAVSPAVVLAYDVLPCSGSSASRRPPRDRMWGQSEGVR | 391989 | 392163 | + | 7 |
| 13 | MSVVAEFANTQLVDVKPGRIGSDAAPTM | MTPLATVATPEAVSAGIAVTSATAAALNMVTHNVQHG | 395105 | 395303 | + | 34 |
| 14 | MRGALCGPCRGACGTPRPACRHPRGTLRRLPCDLPCSGLGHFGPAPF | ATTHLRGGHLITRRMC | 397637 | 397445 | - | 6 |

### Results for WP_148281456.1 [Tsukamurella sp. 1534] back to top

previous - next

#### Architecture

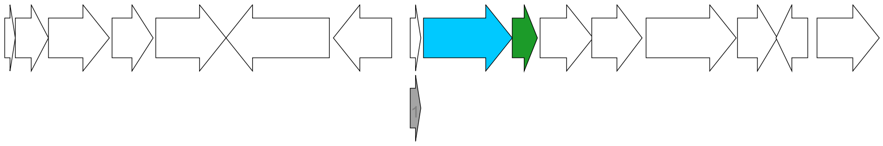

1000 nucleotides

Link to nucleotide sequence

| Accession | start | end | direction | length (aa) | Pfam/HMM | name | description | E-value |
|---|---|---|---|---|---|---|---|---|
| WP_040766109.1 | 1983890 | 1984130 | + | 79 | PF02700<br>TIGR00302 | PurS<br>TIGR00302 | Phosphoribosylformylglycinamidine (FGAM) synthase<br>TIGR00302: phosphoribosylformylglycinamidine synthase, purS protein | 6.10E-24<br>2.90E-14 |
| WP_019202174.1 | 1984126 | 1984804 | + | 225 | TIGR01737<br>PF13507<br>TIGR01857<br>TIGR01735<br>PF07685 | TIGR01737<br>GATase_5<br>TIGR01857<br>TIGR01735<br>GATase_3 | FGAM_synth_I: phosphoribosylformylglycinamidine synthase I<br>CobB/CobQ-like glutamine amidotransferase domain<br>FGAM-synthase: phosphoribosylformylglycinamidine synthase<br>FGAM_synt: phosphoribosylformylglycinamidine synthase<br>CobB/CobQ-like glutamine amidotransferase domain | 9.80E-81<br>3.20E-38<br>5.30E-21<br>1.30E-20<br>2.70E-09 |
| WP_019202175.1 | 1984803 | 1986084 | + | 426 | PF02127 | Peptidase_M18 | Aminopeptidase I zinc metalloprotease (M18) | 2.90E-128 |
| WP_040767233.1 | 1986145 | 1987000 | + | 284 | TIGR01505<br>TIGR01692<br>PF03446<br>PF14833 | TIGR01505<br>TIGR01692<br>NAD_binding_2<br>NAD_binding_11 | tartro_sem_red: 2-hydroxy-3-oxopropionate reductase<br>HIBADH: 3-hydroxyisobutyrate dehydrogenase<br>NAD binding domain of 6-phosphogluconate dehydrogenase<br>NAD-binding of NADP-dependent 3-hydroxyisobutyrate dehydrogenase | 1.60E-44<br>2.60E-34<br>3.30E-24<br>1.10E-21 |
| WP_019202177.1 | 1987036 | 1988509 | + | 490 | PF00324<br>TIGR01773<br>TIGR00913<br>PF13520<br>TIGR00909 | AA_permease<br>TIGR01773<br>TIGR00913<br>AA_permease_2<br>TIGR00909 | Amino acid permease<br>GABAperm: GABA permease<br>2A0310: amino acid permease (yeast)<br>Amino acid permease<br>2A0306: amino acid transporter | 1.40E-106<br>3.80E-93<br>1.90E-81<br>2.70E-50<br>4.50E-41 |

| | | | | | | | | |
|---|---|---|---|---|---|---|---|---|
| WP_019202178.1 | 1990678 | 1988521 | - | 718 | TIGR01591 | TIGR01591 | Fdh-alpha: formate dehydrogenase, alpha subunit | 5.10E-70 |
| | | | | | PF00384 | Molybdopterin | Molybdopterin oxidoreductase | 7.90E-37 |
| | | | | | TIGR00509 | TIGR00509 | bisC_fam: molybdopterin guanine dinucleotide-containing S/N-oxide reductases | 5.50E-34 |
| | | | | | TIGR02166 | TIGR02166 | dmsA_ynfE: anaerobic dimethyl sulfoxide reductase, A subunit, DmsA/YnfE family | 2.20E-32 |
| | | | | | TIGR03479 | TIGR03479 | DMSO_red_II_alp: DMSO reductase family type II enzyme, molybdopterin subunit | 8.80E-25 |
| WP_019202179.1 | 1991971 | 1990753 | - | 405 | PF00817 | IMS | impB/mucB/samB family | 6.20E-44 |
| | | | | | PF11799 | IMS_C | impB/mucB/samB family C-terminal domain | 2.10E-18 |
| | | | | | PF11798 | IMS_HHH | IMS family HHH motif | 4.20E-09 |
| WP_019202180.1 | 1992366 | 1992561 | + | 64 | NO PFAM MATCH | - | - | - |
| WP_148281456.1 | 1992629 | 1994486 | + | 618 | LinH | LinH | LinH (Fusion of LinE/LinG) | 8.80E-273 |
| | | | | | LinG | LinG | LinG Homolog | 4.80E-92 |
| | | | | | LinE | LinE | LinE Homolog | 3.70E-79 |
| | | | | | PF12697 | Abhydrolase_6 | Alpha/beta hydrolase family | 1.80E-15 |
| | | | | | PF00975 | Thioesterase | Thioesterase domain | 2.60E-07 |
| WP_019202182.1 | 1994472 | 1995009 | + | 178 | LinL | LinL | LinL | 1.50E-60 |
| WP_019202183.1 | 1995060 | 1996155 | + | 364 | TIGR02824 | TIGR02824 | quinone_pig3: putative NAD(P)H quinone oxidoreductase, PIG3 family | 1.80E-43 |
| | | | | | TIGR01751 | TIGR01751 | crot-CoA-red: crotonyl-CoA reductase | 2.30E-36 |
| | | | | | TIGR02822 | TIGR02822 | adh_fam_2: zinc-binding alcohol dehydrogenase family protein | 2.20E-23 |
| | | | | | TIGR00692 | TIGR00692 | tdh: L-threonine 3-dehydrogenase | 6.00E-21 |
| | | | | | PF00107 | ADH_zinc_N | Zinc-binding dehydrogenase | 1.60E-19 |
| WP_019202184.1 | 1996141 | 1997179 | + | 345 | PF05544 | Pro_racemase | Proline racemase | 5.90E-80 |
| WP_040767251.1 | 1997256 | 1999149 | + | 630 | PF00106 | adh_short | short chain dehydrogenase | 5.90E-50 |
| | | | | | PF07993 | NAD_binding_4 | Male sterility protein | 1.30E-39 |
| | | | | | TIGR01746 | TIGR01746 | Thioester-redct: thioester reductase domain | 1.60E-39 |
| | | | | | TIGR01963 | TIGR01963 | PHB_DH: 3-hydroxybutyrate dehydrogenase | 3.90E-34 |
| | | | | | PF13561 | adh_short_C2 | Enoyl-(Acyl carrier protein) reductase | 4.20E-34 |
| WP_040766112.1 | 1999175 | 2000015 | + | 279 | PF00106 | adh_short | short chain dehydrogenase | 5.30E-45 |
| | | | | | TIGR03206 | TIGR03206 | benzo_BadH: 2-hydroxycyclohexanecarboxyl-CoA dehydrogenase | 4.40E-39 |
| | | | | | TIGR01963 | TIGR01963 | PHB_DH: 3-hydroxybutyrate dehydrogenase | 2.40E-35 |
| | | | | | PF13561 | adh_short_C2 | Enoyl-(Acyl carrier protein) reductase | 1.00E-31 |
| | | | | | TIGR02415 | TIGR02415 | 23BDH: acetoin reductases | 5.40E-31 |
| WP_019202187.1 | 2000653 | 1999978 | - | 224 | PF13302 | Acetyltransf_3 | Acetyltransferase (GNAT) domain | 6.40E-13 |
| WP_019202188.1 | 2000824 | 2002120 | + | 431 | PF06965 | Na_H_antiport_1 | Na+/H+ antiporter 1 | 7.10E-124 |
| | | | | | TIGR00773 | TIGR00773 | NhaA: Na+/H+ antiporter NhaA | 2.70E-94 |

| index | leader | core | start | end | dir | score |
|---|---|---|---|---|---|---|
| 1 | MAEMISKVNDSAVLAGLPVELDRPEYVGI | TAPVTSVGFAAGGGGGVDVAGMGGGADFAAAAGAF | 1992366 | 1992561 | + | 6 |

### Results for SED38833.1 [Tsukamurella tyrosinosolvens] back to top

previous - next

#### Architecture

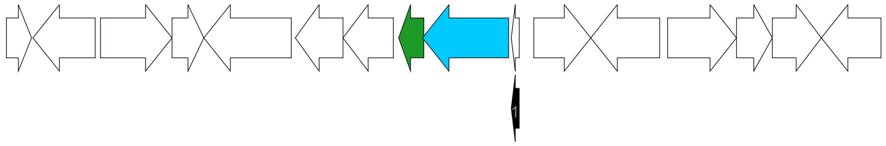

1000 nucleotides

Link to nucleotide sequence

| Accession | start | end | direction | length (aa) | Pfam/HMM | name | description | E-value |
|---|---|---|---|---|---|---|---|---|
| SED38549.1 | 4353084 | 4353624 | + | 179 | TIGR01129 | TIGR01129 | secD: protein-export membrane protein SecD | 1.20E-18 |
| | | | | | TIGR00916 | TIGR00916 | 2A0604s01: protein-export membrane protein, SecD/SecF family | 3.80E-08 |
| SED38588.1 | 4355009 | 4353662 | - | 448 | PF00375 | SDF | Sodium:dicarboxylate symporter family | 1.10E-96 |
| SED38622.1 | 4355108 | 4356662 | + | 517 | PF17203 | sCache_3_2 | Single cache domain 3 | 1.20E-21 |
| | | | | | PF02518 | HATPase_c | Histidine kinase-, DNA gyrase B-, and HSP90-like ATPase | 2.30E-15 |
| | | | | | TIGR02966 | TIGR02966 | phoR_proteo: phosphate regulon sensor kinase PhoR | 5.10E-12 |
| | | | | | TIGR02916 | TIGR02916 | PEP_his_kin: putative PEP-CTERM system histidine kinase | 2.80E-11 |
| | | | | | TIGR01386 | TIGR01386 | cztS_silS_copS: heavy metal sensor kinase | 7.50E-10 |
| SED38660.1 | 4356658 | 4357345 | + | 228 | PF00072 | Response_reg | Response regulator receiver domain | 4.30E-18 |
| | | | | | TIGR02875 | TIGR02875 | spore_0_A: sporulation transcription factor Spo0A | 4.00E-12 |
| | | | | | TIGR01818 | TIGR01818 | ntrC: nitrogen regulation protein NR(I) | 1.80E-11 |
| | | | | | TIGR02154 | TIGR02154 | PhoB: phosphate regulon transcriptional regulatory protein PhoB | 3.50E-09 |
| | | | | | TIGR01387 | TIGR01387 | cztR_silR_copR: heavy metal response regulator | 6.20E-08 |
| SED38692.1 | 4359227 | 4357349 | - | 625 | PF00106 | adh_short | short chain dehydrogenase | 3.90E-52 |
| | | | | | TIGR01746 | TIGR01746 | Thioester-redct: thioester reductase domain | 3.10E-40 |
| | | | | | PF07993 | NAD_binding_4 | Male sterility protein | 4.40E-39 |
| | | | | | PF13561 | adh_short_C2 | Enoyl-(Acyl carrier protein) reductase | 4.10E-37 |
| | | | | | TIGR01963 | TIGR01963 | PHB_DH: 3-hydroxybutyrate dehydrogenase | 5.60E-36 |
| SED38731.1 | 4360344 | 4359306 | - | 345 | PF05544 | Pro_racemase | Proline racemase | 3.70E-80 |
| SED38765.1 | 4361425 | 4360330 | - | 364 | TIGR02824 | TIGR02824 | quinone_pig3: putative NAD(P)H quinone oxidoreductase, PIG3 family | 6.10E-43 |
| | | | | | TIGR01751 | TIGR01751 | crot-CoA-red: crotonyl-CoA reductase | 2.80E-31 |
| | | | | | TIGR02822 | TIGR02822 | adh_fam_2: zinc-binding alcohol dehydrogenase family protein | 6.70E-21 |
| | | | | | TIGR00692 | TIGR00692 | tdh: L-threonine 3-dehydrogenase | 2.00E-19 |
| | | | | | TIGR02823 | TIGR02823 | oxido_YhdH: putative quinone oxidoreductase, YhdH/YhfP family | 5.50E-19 |
| SED38800.1 | 4362070 | 4361521 | - | 182 | LinL | LinL | LinL | 1.70E-65 |

| | | | | | | | | |
|---|---|---|---|---|---|---|---|---|
| SED38833.1 | 4363895 | 4362056 | - | 612 | LinH<br>LinG<br>LinE<br>PF12697<br>PF00561 | LinH<br>LinG<br>LinE<br>Abhydrolase_6<br>Abhydrolase_1 | LinH (Fusion of LinE/LinG)<br>LinG Homolog<br>LinE Homolog<br>Alpha/beta hydrolase family<br>alpha/beta hydrolase fold | 8.40E-248<br>1.50E-91<br>3.70E-79<br>1.40E-14<br>1.20E-09 |
| SED38869.1 | 4364145 | 4363950 | - | 64 | NO PFAM MATCH | - | - | - |
| SED38919.1 | 4364450 | 4365662 | + | 403 | PF00817<br>PF11799<br>PF11798 | IMS<br>IMS_C<br>IMS_HHH | impB/mucB/samB family<br>impB/mucB/samB family C-terminal domain<br>IMS family HHH motif | 1.50E-41<br>1.70E-18<br>1.50E-08 |
| SED38952.1 | 4367152 | 4365679 | - | 490 | PF00324<br>TIGR01773<br>TIGR00913<br>PF13520<br>TIGR00909 | AA_permease<br>TIGR01773<br>TIGR00913<br>AA_permease_2<br>TIGR00909 | Amino acid permease<br>GABAperm: GABA permease<br>2A0310: amino acid permease (yeast)<br>Amino acid permease<br>2A0306: amino acid transporter | 1.60E-106<br>1.70E-95<br>2.40E-81<br>3.90E-49<br>5.70E-44 |
| SED38992.1 | 4367344 | 4368805 | + | 486 | TIGR00883<br>PF00083<br>PF07690<br>TIGR00891<br>TIGR00895 | TIGR00883<br>Sugar_tr<br>MFS_1<br>TIGR00891<br>TIGR00895 | 2A0106: MFS transporter, metabolite:H+ symporter (MHS) family protein<br>Sugar (and other) transporter<br>Major Facilitator Superfamily<br>2A0112: MFS transporter, sialate:H+ symporter (SHS) family<br>2A0115: MFS transporter, aromatic acid:H+ symporter (AAHS) family | 1.20E-114<br>1.30E-32<br>7.20E-27<br>3.80E-24<br>2.10E-21 |
| SED39027.1 | 4368814 | 4369585 | + | 256 | TIGR01832<br>TIGR01830<br>TIGR01963<br>PF13561<br>PF00106 | TIGR01832<br>TIGR01830<br>TIGR01963<br>adh_short_C2<br>adh_short | kduD: 2-deoxy-D-gluconate 3-dehydrogenase<br>3oxo_ACP_reduc: 3-oxoacyl-[acyl-carrier-protein] reductase<br>PHB_DH: 3-hydroxybutyrate dehydrogenase<br>Enoyl-(Acyl carrier protein) reductase<br>short chain dehydrogenase | 3.30E-61<br>1.00E-59<br>1.10E-57<br>1.60E-57<br>2.30E-54 |
| SED39069.1 | 4369596 | 4370628 | + | 343 | TIGR00692<br>TIGR03366<br>TIGR01202<br>PF00107<br>TIGR03451 | TIGR00692<br>TIGR03366<br>TIGR01202<br>ADH_zinc_N<br>TIGR03451 | tdh: L-threonine 3-dehydrogenase<br>HpnZ_proposed: putative phosphonate catabolism associated alcohol dehydrogenase<br>bchC: chlorophyll synthesis pathway protein BchC<br>Zinc-binding dehydrogenase<br>mycoS_dep_FDH: S-(hydroxymethyl)mycothiol dehydrogenase | 2.40E-37<br>1.80E-23<br>2.40E-19<br>8.80E-19<br>2.20E-17 |
| SED39104.1 | 4371927 | 4370649 | - | 425 | PF02127 | Peptidase_M18 | Aminopeptidase I zinc metalloprotease (M18) | 2.60E-129 |

| index | leader | core | start | end | dir | score |
|---|---|---|---|---|---|---|
| 1 | MTEMISKVNRSAVLA | GTPVDLDRPAYVGATAPVTSVGFAAGGGGGVDVAGMGGDANFAAAAGAF | 4364145 | 4363950 | - | 17 |

### Results for WP_147276785.1 [Tsukamurella tyrosinosolvens] back to top

previous - next

#### Architecture

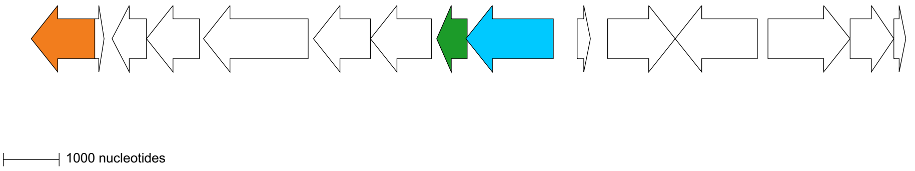

Link to nucleotide sequence

| Accession | start | end | direction | length (aa) | Pfam/HMM | name | description | E-value |
|---|---|---|---|---|---|---|---|---|
| WP_147276788.1 | 24768 | 23611 | - | 386 | TIGR02770<br>TIGR02769<br>TIGR03005<br>TIGR03269<br>TIGR00972 | TIGR02770<br>TIGR02769<br>TIGR03005<br>TIGR03269<br>TIGR00972 | nickel_nikD: nickel import ATP-binding protein NikD<br>nickel_nikE: nickel import ATP-binding protein NikE<br>ectoine_ehuA: ectoine/hydroxyectoine ABC transporter, ATP-binding protein EhuA<br>met_CoM_red_A2: methyl coenzyme M reductase system, component A2<br>3a0107s01c2: phosphate ABC transporter, ATP-binding protein | 2.70E-79<br>4.00E-60<br>5.20E-48<br>2.60E-44<br>9.50E-44 |
| WP_013128178.1 | 24764 | 24920 | + | 52 | INFERRED GENE | - | - | - |
| WP_147276789.1 | 25699 | 25071 | - | 209 | TIGR02790<br>PF00528<br>TIGR01097<br>TIGR03416<br>PF12911 | TIGR02790<br>BPD_transp_1<br>TIGR01097<br>TIGR03416<br>OppC_N | nickel_nikC: nickel ABC transporter, permease subunit NikC<br>Binding-protein-dependent transport system inner membrane component<br>PhnE: phosphonate ABC transporter, permease protein PhnE<br>ABC_choXWV_perm: choline ABC transporter, permease protein<br>N-terminal TM domain of oligopeptide transport permease C | 4.60E-42<br>3.10E-16<br>7.10E-11<br>2.80E-07<br>7.40E-06 |
| WP_114515649.1 | 26649 | 25695 | - | 317 | TIGR02789<br>PF00528 | TIGR02789<br>BPD_transp_1 | nickel_nikB: nickel ABC transporter, permease subunit NikB<br>Binding-protein-dependent transport system inner membrane component | 4.20E-56<br>2.50E-26 |
| WP_114515650.1 | 28600 | 26722 | - | 625 | PF00106<br>TIGR01746<br>PF07993<br>PF13561<br>TIGR01963 | adh_short<br>TIGR01746<br>NAD_binding_4<br>adh_short_C2<br>TIGR01963 | short chain dehydrogenase<br>Thioester-redct: thioester reductase domain<br>Male sterility protein<br>Enoyl-(Acyl carrier protein) reductase<br>PHB_DH: 3-hydroxybutyrate dehydrogenase | 3.90E-52<br>8.50E-40<br>1.60E-39<br>4.10E-37<br>5.60E-36 |
| WP_068523287.1 | 29714 | 28679 | - | 344 | PF05544 | Pro_racemase | Proline racemase | 3.60E-80 |
| WP_114515651.1 | 30798 | 29703 | - | 364 | TIGR02824<br>TIGR01751<br>TIGR02822<br>TIGR00692<br>TIGR02823 | TIGR02824<br>TIGR01751<br>TIGR02822<br>TIGR00692<br>TIGR02823 | quinone_pig3: putative NAD(P)H quinone oxidoreductase, PIG3 family<br>crot-CoA-red: crotonyl-CoA reductase<br>adh_fam_2: zinc-binding alcohol dehydrogenase family protein<br>tdh: L-threonine 3-dehydrogenase<br>oxido_YhdH: putative quinone oxidoreductase, YhdH/YhfP family | 8.90E-44<br>8.00E-32<br>7.10E-21<br>1.90E-19<br>5.70E-19 |
| WP_147276784.1 | 31443 | 30894 | - | 182 | LinL | LinL | LinL | 2.10E-65 |
| WP_147276785.1 | 32986 | 31429 | - | 518 | LinH<br>LinE<br>LinG<br>PF12697<br>PF00561 | LinH<br>LinE<br>LinG<br>Abhydrolase_6<br>Abhydrolase_1 | LinH (Fusion of LinE/LinG)<br>LinE Homolog<br>LinG Homolog<br>Alpha/beta hydrolase family<br>alpha/beta hydrolase fold | 5.40E-215<br>1.40E-78<br>1.30E-70<br>2.90E-15<br>2.90E-10 |
| WP_114515654.1 | 33404 | 33653 | + | 82 | NO PFAM MATCH | - | - | - |

| | | | | | | | | |
|---|---|---|---|---|---|---|---|---|
| WP_068522805.1 | 33956 | 35168 | + | 403 | PF00817<br>PF11799<br>PF11798 | IMS<br>IMS_C<br>IMS_HHH | impB/mucB/samB family<br>impB/mucB/samB family C-terminal domain<br>IMS family HHH motif | 1.50E-41<br>1.70E-18<br>1.50E-08 |
| WP_114515655.1 | 36658 | 35185 | - | 490 | PF00324<br>TIGR01773<br>TIGR00913<br>PF13520<br>TIGR00909 | AA_permease<br>TIGR01773<br>TIGR00913<br>AA_permease_2<br>TIGR00909 | Amino acid permease<br>GABAperm: GABA permease<br>2A0310: amino acid permease (yeast)<br>Amino acid permease<br>2A0306: amino acid transporter | 1.30E-106<br>1.60E-95<br>3.60E-81<br>1.10E-49<br>4.20E-44 |
| WP_114515656.1 | 36850 | 38311 | + | 486 | TIGR00883<br>PF00083<br>PF07690<br>TIGR00891<br>TIGR00895 | TIGR00883<br>Sugar_tr<br>MFS_1<br>TIGR00891<br>TIGR00895 | 2A0106: MFS transporter, metabolite:H+ symporter (MHS) family protein<br>Sugar (and other) transporter<br>Major Facilitator Superfamily<br>2A0112: MFS transporter, sialate:H+ symporter (SHS) family<br>2A0115: MFS transporter, aromatic acid:H+ symporter (AAHS) family | 9.60E-115<br>1.10E-32<br>5.70E-27<br>3.80E-24<br>1.50E-21 |
| WP_114515657.1 | 38320 | 39091 | + | 256 | TIGR01832<br>TIGR01830<br>TIGR01963<br>PF13561<br>PF00106 | TIGR01832<br>TIGR01830<br>TIGR01963<br>adh_short_C2<br>adh_short | kduD: 2-deoxy-D-gluconate 3-dehydrogenase<br>3oxo_ACP_reduc: 3-oxoacyl-[acyl-carrier-protein] reductase<br>PHB_DH: 3-hydroxybutyrate dehydrogenase<br>Enoyl-(Acyl carrier protein) reductase<br>short chain dehydrogenase | 9.10E-61<br>2.50E-59<br>3.20E-57<br>3.30E-57<br>8.00E-54 |
| WP_013350118.1 | 39087 | 39300 | + | 71 | INFERRED GENE | - | - | - |

| index | leader | core | start | end | dir | score |
|---|---|---|---|---|---|---|

### Results for AUN42108.1 [Tsukamurella tyrosinosolvens] back to top

previous - next

#### Architecture

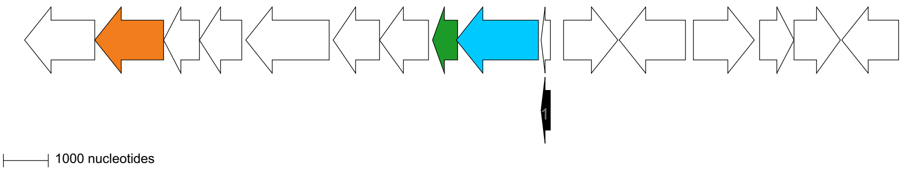

Link to nucleotide sequence

| Accession | start | end | direction | length (aa) | Pfam/HMM | name | description | E-value |
|---|---|---|---|---|---|---|---|---|
| AUN42101.1 | 4188870 | 4187328 | - | 513 | PF00496<br>TIGR02294<br>TIGR04028 | SBP_bac_5<br>TIGR02294<br>TIGR04028 | Bacterial extracellular solute-binding proteins, family 5 Middle<br>nickel_nikA: nickel ABC transporter, nickel/metallophore periplasmic binding protein<br>SBP_KPN_01854: ABC transporter substrate binding protein, KPN_01854 family | 5.80E-63<br>5.30E-33<br>8.50E-22 |
| AUN42102.1 | 4190420 | 4188866 | - | 517 | TIGR02770<br>TIGR02769<br>TIGR03005<br>TIGR00968<br>TIGR00972 | TIGR02770<br>TIGR02769<br>TIGR03005<br>TIGR00968<br>TIGR00972 | nickel_nikD: nickel import ATP-binding protein NikD<br>nickel_nikE: nickel import ATP-binding protein NikE<br>ectoine_ehuA: ectoine/hydroxyectoine ABC transporter, ATP-binding protein EhuA<br>3a0106s01: sulfate ABC transporter, ATP-binding protein<br>3a0107s01c2: phosphate ABC transporter, ATP-binding protein | 5.50E-118<br>4.70E-100<br>7.70E-78<br>1.30E-73<br>2.90E-72 |
| AUN42103.1 | 4191208 | 4190416 | - | 263 | TIGR02790<br>PF00528<br>TIGR01097<br>TIGR02789<br>TIGR03416 | TIGR02790<br>BPD_transp_1<br>TIGR01097<br>TIGR02789<br>TIGR03416 | nickel_nikC: nickel ABC transporter, permease subunit NikC<br>Binding-protein-dependent transport system inner membrane component<br>PhnE: phosphonate ABC transporter, permease protein PhnE<br>nickel_nikB: nickel ABC transporter, permease subunit NikB<br>ABC_choXWV_perm: choline ABC transporter, permease protein | 2.60E-48<br>1.20E-19<br>2.90E-10<br>4.80E-07<br>7.50E-07 |
| AUN42104.1 | 4192149 | 4191204 | - | 314 | TIGR02789<br>PF00528 | TIGR02789<br>BPD_transp_1 | nickel_nikB: nickel ABC transporter, permease subunit NikB<br>Binding-protein-dependent transport system inner membrane component | 5.20E-56<br>1.20E-25 |
| AUN42105.1 | 4194109 | 4192231 | - | 625 | PF00106<br>TIGR01746<br>PF07993<br>PF13561<br>TIGR01963 | adh_short<br>TIGR01746<br>NAD_binding_4<br>adh_short_C2<br>TIGR01963 | short chain dehydrogenase<br>Thioester-redct: thioester reductase domain<br>Male sterility protein<br>Enoyl-(Acyl carrier protein) reductase<br>PHB_DH: 3-hydroxybutyrate dehydrogenase | 6.00E-52<br>7.30E-40<br>1.00E-39<br>4.20E-37<br>5.40E-36 |
| AUN43200.1 | 4195223 | 4194188 | - | 344 | PF05544 | Pro_racemase | Proline racemase | 3.60E-80 |
| AUN42106.1 | 4196307 | 4195212 | - | 364 | TIGR02824<br>TIGR01751<br>TIGR02822<br>TIGR00692<br>TIGR02823 | TIGR02824<br>TIGR01751<br>TIGR02822<br>TIGR00692<br>TIGR02823 | quinone_pig3: putative NAD(P)H quinone oxidoreductase, PIG3 family<br>crot-CoA-red: crotonyl-CoA reductase<br>adh_fam_2: zinc-binding alcohol dehydrogenase family protein<br>tdh: L-threonine 3-dehydrogenase<br>oxido_YhdH: putative quinone oxidoreductase, YhdH/YhfP family | 8.70E-44<br>6.30E-32<br>9.50E-21<br>1.30E-19<br>4.90E-19 |
| AUN42107.1 | 4196961 | 4196403 | - | 185 | LinL | LinL | LinL | 1.60E-65 |
| AUN42108.1 | 4198777 | 4196938 | - | 612 | LinH<br>LinG<br>LinE<br>PF12697<br>PF00561 | LinH<br>LinG<br>LinE<br>Abhydrolase_6<br>Abhydrolase_1 | LinH (Fusion of LinE/LinG)<br>LinG Homolog<br>LinE Homolog<br>Alpha/beta hydrolase family<br>alpha/beta hydrolase fold | 4.70E-248<br>2.00E-91<br>3.60E-79<br>1.40E-14<br>1.20E-09 |
| AUN42109.1 | 4199018 | 4198832 | - | 61 | NO PFAM MATCH | - | - | - |
| AUN42110.1 | 4199331 | 4200543 | + | 403 | PF00817<br>PF11799<br>PF11798 | IMS<br>IMS_C<br>IMS_HHH | impB/mucB/samB family<br>impB/mucB/samB family C-terminal domain<br>IMS family HHH motif | 1.50E-41<br>1.70E-18<br>1.50E-08 |
| AUN42111.1 | 4202033 | 4200560 | - | 490 | PF00324<br>TIGR01773<br>TIGR00913<br>PF13520<br>TIGR00909 | AA_permease<br>TIGR01773<br>TIGR00913<br>AA_permease_2<br>TIGR00909 | Amino acid permease<br>GABAperm: GABA permease<br>2A0310: amino acid permease (yeast)<br>Amino acid permease<br>2A0306: amino acid transporter | 1.30E-106<br>2.10E-95<br>3.60E-81<br>1.10E-49<br>4.20E-44 |
| WP_017828447.1 | 4202225 | 4203581 | + | 452 | INFERRED GENE | - | - | - |

| | | | | | | | | |
|---|---|---|---|---|---|---|---|---|
| AUN42112.1 | 4203695 | 4204466 | + | 256 | TIGR01832<br>TIGR01830<br>TIGR01963<br>PF13561<br>PF00106 | TIGR01832<br>TIGR01830<br>TIGR01963<br>adh_short_C2<br>adh_short | kduD: 2-deoxy-D-gluconate 3-dehydrogenase<br>3oxo_ACP_reduc: 3-oxoacyl-[acyl-carrier-protein] reductase<br>PHB_DH: 3-hydroxybutyrate dehydrogenase<br>Enoyl-(Acyl carrier protein) reductase<br>short chain dehydrogenase | 3.30E-61<br>1.00E-59<br>1.10E-57<br>1.60E-57<br>2.30E-54 |
| AUN42113.1 | 4204462 | 4205509 | + | 348 | TIGR00692<br>TIGR03366<br>TIGR01202<br>PF00107<br>TIGR03451 | TIGR00692<br>TIGR03366<br>TIGR01202<br>ADH_zinc_N<br>TIGR03451 | tdh: L-threonine 3-dehydrogenase<br>HpnZ_proposed: putative phosphonate catabolism associated alcohol dehydrogenase<br>bchC: chlorophyll synthesis pathway protein BchC<br>Zinc-binding dehydrogenase<br>mycoS_dep_FDH: S-(hydroxymethyl)mycothiol dehydrogenase | 4.80E-36<br>2.30E-22<br>1.60E-17<br>4.50E-17<br>2.70E-16 |
| AUN42114.1 | 4206808 | 4205530 | - | 425 | PF02127 | Peptidase_M18 | Aminopeptidase I zinc metalloprotease (M18) | 5.70E-129 |

| index | leader | core | start | end | dir | score |
|---|---|---|---|---|---|---|
| 1 | MISKVNRSAVLA | GTPVDLDRPAYVGATAPVTSVGFAAGGGGGVDVAGMGGDANFAAAAGAF | 4199018 | 4198832 | - | 18 |

### Results for RDB46095.1 [Tsukamurella tyrosinosolvens] back to top

previous - next

#### Architecture

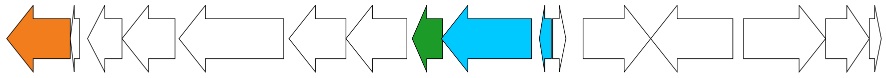

1000 nucleotides

Link to nucleotide sequence

| Accession | start | end | direction | length (aa) | Pfam/HMM | name | description | E-value |
|---|---|---|---|---|---|---|---|---|
| RDB46103.1 | 24768 | 23611 | - | 386 | TIGR02770<br>TIGR02769<br>TIGR03005<br>TIGR03269<br>TIGR00972 | TIGR02770<br>TIGR02769<br>TIGR03005<br>TIGR03269<br>TIGR00972 | nickel_nikD: nickel import ATP-binding protein NikD<br>nickel_nikE: nickel import ATP-binding protein NikE<br>ectoine_ehuA: ectoine/hydroxyectoine ABC transporter, ATP-binding protein EhuA<br>met_CoM_red_A2: methyl coenzyme M reductase system, component A2<br>3a0107s01c2: phosphate ABC transporter, ATP-binding protein | 2.70E-79<br>4.00E-60<br>5.20E-48<br>2.60E-44<br>9.50E-44 |
| RDB46089.1 | 24920 | 24764 | - | 51 | NO PFAM MATCH | - | - | - |
| RDB46090.1 | 25699 | 25071 | - | 209 | TIGR02790<br>PF00528<br>TIGR01097<br>TIGR03416<br>PF12911 | TIGR02790<br>BPD_transp_1<br>TIGR01097<br>TIGR03416<br>OppC_N | nickel_nikC: nickel ABC transporter, permease subunit NikC<br>Binding-protein-dependent transport system inner membrane component<br>PhnE: phosphonate ABC transporter, permease protein PhnE<br>ABC_choXWV_perm: choline ABC transporter, permease protein<br>N-terminal TM domain of oligopeptide transport permease C | 4.60E-42<br>3.10E-16<br>7.10E-11<br>2.80E-07<br>7.40E-06 |
| RDB46091.1 | 26649 | 25695 | - | 317 | TIGR02789<br>PF00528 | TIGR02789<br>BPD_transp_1 | nickel_nikB: nickel ABC transporter, permease subunit NikB<br>Binding-protein-dependent transport system inner membrane component | 4.20E-56<br>2.50E-26 |
| RDB46092.1 | 28600 | 26722 | - | 625 | PF00106<br>TIGR01746<br>PF07993<br>PF13561<br>TIGR01963 | adh_short<br>TIGR01746<br>NAD_binding_4<br>adh_short_C2<br>TIGR01963 | short chain dehydrogenase<br>Thioester-redct: thioester reductase domain<br>Male sterility protein<br>Enoyl-(Acyl carrier protein) reductase<br>PHB_DH: 3-hydroxybutyrate dehydrogenase | 3.90E-52<br>8.50E-40<br>1.60E-39<br>4.10E-37<br>5.60E-36 |
| RDB46104.1 | 29714 | 28679 | - | 344 | PF05544 | Pro_racemase | Proline racemase | 3.60E-80 |
| RDB46093.1 | 30798 | 29703 | - | 364 | TIGR02824<br>TIGR01751<br>TIGR02822<br>TIGR00692<br>TIGR02823 | TIGR02824<br>TIGR01751<br>TIGR02822<br>TIGR00692<br>TIGR02823 | quinone_pig3: putative NAD(P)H quinone oxidoreductase, PIG3 family<br>crot-CoA-red: crotonyl-CoA reductase<br>adh_fam_2: zinc-binding alcohol dehydrogenase family protein<br>tdh: L-threonine 3-dehydrogenase<br>oxido_YhdH: putative quinone oxidoreductase, YhdH/YhfP family | 8.90E-44<br>8.00E-32<br>7.10E-21<br>1.90E-19<br>5.70E-19 |
| RDB46094.1 | 31452 | 30894 | - | 185 | LinL | LinL | LinL | 1.90E-65 |
| RDB46095.1 | 33037 | 31429 | - | 535 | LinH<br>LinE<br>LinG<br>PF12697<br>PF00561 | LinH<br>LinE<br>LinG<br>Abhydrolase_6<br>Abhydrolase_1 | LinH (Fusion of LinE/LinG)<br>LinE Homolog<br>LinG Homolog<br>Alpha/beta hydrolase family<br>alpha/beta hydrolase fold | 5.10E-221<br>1.60E-78<br>1.00E-75<br>3.20E-15<br>3.00E-10 |
| RDB46096.1 | 33401 | 33178 | - | 74 | LinH<br>LinG | LinH<br>LinG | LinH (Fusion of LinE/LinG)<br>LinG Homolog | 2.60E-20<br>1.30E-11 |
| RDB46097.1 | 33404 | 33653 | + | 82 | NO PFAM MATCH | - | - | - |
| RDB46098.1 | 33956 | 35168 | + | 403 | PF00817<br>PF11799<br>PF11798 | IMS<br>IMS_C<br>IMS_HHH | impB/mucB/samB family<br>impB/mucB/samB family C-terminal domain<br>IMS family HHH motif | 1.50E-41<br>1.70E-18<br>1.50E-08 |
| RDB46099.1 | 36658 | 35185 | - | 490 | PF00324<br>TIGR01773<br>TIGR00913<br>PF13520<br>TIGR00909 | AA_permease<br>TIGR01773<br>TIGR00913<br>AA_permease_2<br>TIGR00909 | Amino acid permease<br>GABAperm: GABA permease<br>2A0310: amino acid permease (yeast)<br>Amino acid permease<br>2A0306: amino acid transporter | 1.30E-106<br>1.60E-95<br>3.60E-81<br>1.10E-49<br>4.20E-44 |
| RDB46100.1 | 36850 | 38311 | + | 486 | TIGR00883<br>PF00083<br>PF07690<br>TIGR00891<br>TIGR00895 | TIGR00883<br>Sugar_tr<br>MFS_1<br>TIGR00891<br>TIGR00895 | 2A0106: MFS transporter, metabolite:H+ symporter (MHS) family protein<br>Sugar (and other) transporter<br>Major Facilitator Superfamily<br>2A0112: MFS transporter, sialate:H+ symporter (SHS) family<br>2A0115: MFS transporter, aromatic acid:H+ symporter (AAHS) family | 9.60E-115<br>1.10E-32<br>5.70E-27<br>3.80E-24<br>1.50E-21 |
| RDB46101.1 | 38320 | 39091 | + | 256 | TIGR01832<br>TIGR01830<br>TIGR01963<br>PF13561<br>PF00106 | TIGR01832<br>TIGR01830<br>TIGR01963<br>adh_short_C2<br>adh_short | kduD: 2-deoxy-D-gluconate 3-dehydrogenase<br>3oxo_ACP_reduc: 3-oxoacyl-[acyl-carrier-protein] reductase<br>PHB_DH: 3-hydroxybutyrate dehydrogenase<br>Enoyl-(Acyl carrier protein) reductase<br>short chain dehydrogenase | 9.10E-61<br>2.50E-59<br>3.20E-57<br>3.30E-57<br>8.00E-54 |

| | | | | | | | | |
|---|---|---|---|---|---|---|---|---|
| RDB46102.1 | 39087 | 39300 | + | 71 | PF08240<br>TIGR00692<br>TIGR02822<br>TIGR03201<br>TIGR03451 | ADH_N<br>TIGR00692<br>TIGR02822<br>TIGR03201<br>TIGR03451 | Alcohol dehydrogenase GroES-like domain<br>tdh: L-threonine 3-dehydrogenase<br>adh_fam_2: zinc-binding alcohol dehydrogenase family protein<br>dearomat_had: 6-hydroxycyclohex-1-ene-1-carbonyl-CoA dehydrogenase<br>mycoS_dep_FDH: S-(hydroxymethyl)mycothiol dehydrogenase | 1.00E-07<br>9.20E-06<br>1.90E-05<br>5.50E-05<br>6.10E-05 |

| index | leader | core | start | end | dir | score |
|---|---|---|---|---|---|---|

### Results for SFQ42880.1 [Yuhushiella deserti] back to top

previous - next

#### Architecture

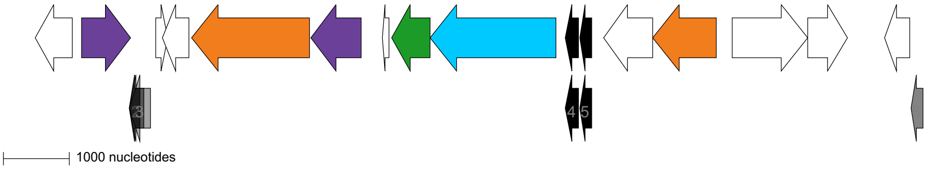

Link to nucleotide sequence

| Accession | start | end | direction | length (aa) | Pfam/HMM | name | description | E-value |
|---|---|---|---|---|---|---|---|---|
| SFQ42749.1 | 108865 | 108310 | - | 184 | NO PFAM MATCH | - | - | - |
| SFQ42763.1 | 109011 | 109749 | + | 245 | PF13649<br>PF13847<br>PF08241<br>TIGR02021<br>TIGR01983 | Methyltransf_25<br>Methyltransf_31<br>Methyltransf_11<br>TIGR02021<br>TIGR01983 | Methyltransferase domain<br>Methyltransferase domain<br>Methyltransferase domain<br>BchM-ChlM: magnesium protoporphyrin O-methyltransferase<br>UbiG: 3-demethylubiquinone-9 3-O-methyltransferase | 4.00E-22<br>3.60E-15<br>4.60E-15<br>2.60E-13<br>1.50E-11 |
| SFQ42778.1 | 110139 | 110310 | + | 56 | NO PFAM MATCH | - | - | - |
| SFQ42794.1 | 110639 | 110228 | - | 136 | NO PFAM MATCH | - | - | - |
| SFQ42811.1 | 112474 | 110683 | - | 596 | TIGR02204<br>TIGR02203<br>TIGR00958<br>TIGR03796<br>TIGR03375 | TIGR02204<br>TIGR02203<br>TIGR00958<br>TIGR03796<br>TIGR03375 | MsbA_rel: ABC transporter, permease/ATP-binding protein<br>MsbA_lipidA: lipid A export permease/ATP-binding protein MsbA<br>3a01208: antigen peptide transporter 2<br>NHLM_micro_ABC1: NHLM bacteriocin system ABC transporter, peptidase/ATP-binding protein<br>type_I_sec_LssB: type I secretion system ATPase | 3.20E-105<br>2.20E-102<br>7.20E-90<br>1.70E-87<br>1.90E-87 |
| SFQ42826.1 | 113241 | 112479 | - | 253 | PF13649<br>PF08241<br>PF13489<br>PF08242<br>PF13847 | Methyltransf_25<br>Methyltransf_11<br>Methyltransf_23<br>Methyltransf_12<br>Methyltransf_31 | Methyltransferase domain<br>Methyltransferase domain<br>Methyltransferase domain<br>Methyltransferase domain<br>Methyltransferase domain | 8.20E-18<br>2.30E-12<br>2.70E-10<br>5.60E-10<br>4.60E-09 |
| SFQ42842.1 | 113674 | 113581 | - | 30 | NO PFAM MATCH | - | - | - |
| SFQ42856.1 | 114304 | 113722 | - | 193 | LinL | LinL | LinL | 4.40E-69 |
| SFQ42880.1 | 116204 | 114305 | - | 632 | LinH<br>LinG<br>LinE<br>PF12697<br>LinL | LinH<br>LinG<br>LinE<br>Abhydrolase_6<br>LinL | LinH (Fusion of LinE/LinG)<br>LinG Homolog<br>LinE Homolog<br>Alpha/beta hydrolase family<br>LinL | 3.70E-252<br>1.80E-119<br>2.10E-100<br>7.80E-14<br>6.80E-10 |
| SFQ42900.1 | 116541 | 116349 | - | 63 | LinA | LinA | LinA (linaridin precursor) | 8.00E-16 |
| SFQ42919.1 | 116746 | 116566 | - | 59 | LinA | LinA | LinA (linaridin precursor) | 5.30E-14 |
| SFQ42935.1 | 117679 | 116929 | - | 249 | PF01061<br>PF12698<br>TIGR00025 | ABC2_membrane<br>ABC2_membrane_3<br>TIGR00025 | ABC-2 type transporter<br>ABC-2 family transporter protein<br>Mtu_efflux: ABC transporter efflux protein, DrrB family | 1.20E-09<br>3.10E-08<br>4.00E-07 |
| SFQ42951.1 | 118635 | 117678 | - | 318 | TIGR01188<br>TIGR03864<br>TIGR01288<br>TIGR03522<br>TIGR03740 | TIGR01188<br>TIGR03864<br>TIGR01288<br>TIGR03522<br>TIGR03740 | drrA: daunorubicin resistance ABC transporter, ATP-binding protein<br>PQQ_ABC_ATP: ABC transporter, ATP-binding subunit, PQQ-dependent alcohol dehydrogenase system<br>nodI: nodulation ABC transporter NodI<br>GldA_ABC_ATP: gliding motility-associated ABC transporter ATP-binding subunit GldA<br>galliderm_ABC: lantibiotic protection ABC transporter, ATP-binding subunit | 9.10E-67<br>2.50E-51<br>6.60E-47<br>1.20E-37<br>6.30E-36 |
| SFQ42965.1 | 118885 | 120025 | + | 379 | PF07730 | HisKA_3 | Histidine kinase | 8.10E-14 |
| SFQ42978.1 | 120021 | 120624 | + | 200 | PF00072<br>TIGR02875<br>PF00196<br>TIGR02154<br>TIGR03020 | Response_reg<br>TIGR02875<br>GerE<br>TIGR02154<br>TIGR03020 | Response regulator receiver domain<br>spore_0_A: sporulation transcription factor Spo0A<br>Bacterial regulatory proteins, luxR family<br>PhoB: phosphate regulon transcriptional regulatory protein PhoB<br>EpsA: transcriptional regulator EpsA | 1.30E-23<br>2.80E-14<br>6.10E-14<br>1.00E-07<br>1.10E-06 |
| SFQ42994.1 | 121582 | 121198 | - | 127 | PF04134 | DUF393 | Protein of unknown function, DUF393 | 1.40E-20 |

| index | leader | core | start | end | dir | score |
|---|---|---|---|---|---|---|
| 1 | LEGCAQRAVAVDNPARARSGQLGAGTGVGRLPGR | LTTLTRAPREPGVTRSRG | 109888 | 109729 | - | 8 |
| 2 | VLPWSSLSGRSRQAGGKRFPLEGCA...AVDNPARARSGQLGAGTGVGRLPGR | LTTLTRAPREPGVTRSRG | 109948 | 109729 | - | 7 |

| | | | | | | |
|---|---|---|---|---|---|---|
| 3 | MAAGGGVRPRVSTNGAFAPREAAPQ...AVDNPARARSGQLGAGTGVGRLPGR | LTTLTRAPREPGVTRSRG | 110044 | 109729 | - | 6 |
| 4 | MESTMDQLVARLNTSPDLASVEVSSTEGAVFPGASPVL | GTPGAFLGGVAIASATVAAFEAGRG | 116541 | 116349 | - | 29 |
| 5 | MLEEKLNIQAEEFGKLDENHAGRIAEAATPVL | ATPALAGAFVAGAGVVTGAYAAGKAVG | 116746 | 116566 | - | 30 |
| 6 | VLRILRKLDVANRTEAAAIAHRVGLTGLTQVNRVPLR | PTPMDGVAVSQAMNRVPMNDIAG | 121769 | 121586 | - | 7 |

### Results for WP_114385254.1 [Brevibacterium aurantiacum] back to top

previous - next

#### Architecture

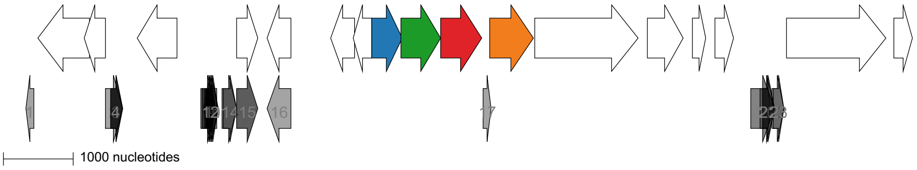

Link to nucleotide sequence

| Accession | start | end | direction | length (aa) | Pfam/HMM | name | description | E-value |
|---|---|---|---|---|---|---|---|---|
| WP_127362988.1 | 2055474 | 2054739 | - | 244 | PF13276<br>PF00665 | HTH_21<br>rve | HTH-like domain<br>Integrase core domain | 4.00E-17<br>2.10E-12 |
| WP_009883477.1 | 2055710 | 2055398 | - | 103 | PF01527 | HTH_Tnp_1 | Transposase | 5.70E-14 |
| WP_125178125.1 | 2056726 | 2056168 | - | 185 | NO PFAM MATCH | - | - | - |
| WP_127362989.1 | 2057582 | 2057885 | + | 100 | PF02861<br>TIGR02639 | Clp_N<br>TIGR02639 | Clp amino terminal domain, pathogenicity island component<br>ClpA: ATP-dependent Clp protease ATP-binding subunit ClpA | 2.20E-04<br>4.80E-04 |
| WP_125178127.1 | 2058370 | 2058022 | - | 115 | NO PFAM MATCH | - | - | - |
| WP_009885501.1 | 2059271 | 2058938 | - | 110 | NO PFAM MATCH | - | - | - |
| WP_081448679.1 | 2059528 | 2059267 | - | 86 | PF13936<br>PF13384<br>PF01381<br>PF13542<br>TIGR04111 | HTH_38<br>HTH_23<br>HTH_3<br>HTH_Tnp_ISL3<br>TIGR04111 | Helix-turn-helix domain<br>Homeodomain-like domain<br>Helix-turn-helix<br>Helix-turn-helix domain of transposase family ISL3<br>BcepMu_gp16: phage-associated protein, BcepMu gp16 family | 1.70E-12<br>1.80E-10<br>2.00E-08<br>8.20E-08<br>2.20E-07 |
| WP_127362990.1 | 2059520 | 2059961 | + | 146 | LinE<br>LinH | LinE<br>LinH | LinE Homolog<br>LinH (Fusion of LinE/LinG) | 1.00E-30<br>5.30E-21 |
| WP_114385254.1 | 2059950 | 2060508 | + | 185 | LinL | LinL | LinL | 1.10E-45 |
| WP_114385253.1 | 2060504 | 2061098 | + | 197 | PF02441<br>TIGR00521<br>TIGR02113 | Flavoprotein<br>TIGR00521<br>TIGR02113 | Flavoprotein<br>coaBC_dfp: phosphopantothenoylcysteine decarboxylase / phosphopantothenate--cysteine ligase<br>coaC_strep: phosphopantothenoylcysteine decarboxylase | 8.00E-10<br>8.60E-10<br>4.80E-04 |
| WP_101621248.1 | 2061218 | 2061848 | + | 209 | TIGR02673<br>TIGR01189<br>TIGR03873<br>TIGR02857<br>TIGR01166 | TIGR02673<br>TIGR01189<br>TIGR03873<br>TIGR02857<br>TIGR01166 | FtsE: cell division ATP-binding protein FtsE<br>ccmA: heme ABC exporter, ATP-binding protein CcmA<br>F420-0_ABC_ATP: proposed F420-0 ABC transporter, ATP-binding protein<br>CydD: thiol reductant ABC exporter, CydD subunit<br>cbiO: cobalt ABC transporter, ATP-binding protein | 1.30E-30<br>1.20E-26<br>1.40E-26<br>1.40E-25<br>7.10E-25 |
| WP_127362991.1 | 2061868 | 2063344 | + | 491 | NO PFAM MATCH | - | - | - |
| WP_127362992.1 | 2063477 | 2063978 | + | 166 | PF03372 | Exo_endo_phos | Endonuclease/Exonuclease/phosphatase family | 9.20E-06 |
| WP_009212852.1 | 2064121 | 2064313 | + | 64 | INFERRED GENE | - | - | - |
| WP_005961783.1 | 2064435 | 2064714 | + | 93 | INFERRED GENE | - | - | - |
| WP_009883329.1 | 2065466 | 2066900 | + | 477 | TIGR04416<br>PF00078<br>PF08388 | TIGR04416<br>RVT_1<br>GIIM | group_II_RT_mat: group II intron reverse transcriptase/maturase<br>Reverse transcriptase (RNA-dependent DNA polymerase)<br>Group II intron, maturase-specific domain | 2.80E-113<br>1.30E-38<br>1.70E-23 |
| WP_076611758.1 | 2067007 | 2067268 | + | 87 | INFERRED GENE | - | - | - |

| index | leader | core | start | end | dir | score |
|---|---|---|---|---|---|---|
| 1 | LRLYKASSCESVSCGAVGVGV | ATVMAWRIMSIVANA | 2054689 | 2054578 | - | 6 |
| 2 | MRESPSSNLPASIIPGAIHSAQYRLGDMLMRCEADCRQVRV | PPACTQGLLDSSQVKIFKYVLVSSIFSLGIYVASTSCKFPWP | 2055708 | 2055960 | + | 6 |
| 3 | MLMRCEADCRQVRVPPACTQGLLDSSQV | KIFKYVLVSSIFSLGIYVASTSCKFPWP | 2055789 | 2055960 | + | 8 |
| 4 | MRCEADCRQVRVPPACTQGLLDSSQVK | IFKYVLVSSIFSLGIYVASTSCKFPWP | 2055795 | 2055960 | + | 7 |
| 5 | VSQLGCHLSVYQSLTSAQYRLGDML...DGGVEQTVDKLGECRSKNFCGVPHV | KIPGRWNLVTVGEQEYAQSPN | 2057068 | 2057287 | + | 7 |
| 6 | VYQSLTSAQYRLGDMLVDGGVEQTVDKLGECRSKNFCGVPHV | KIPGRWNLVTVGEQEYAQSPN | 2057095 | 2057287 | + | 8 |
| 7 | MLVDGGVEQTVDKLGECRSKNFCGVPHV | KIPGRWNLVTVGEQEYAQSPN | 2057137 | 2057287 | + | 6 |
| 8 | MVVWSRLSISLESADPKTSAECLMSRSPAD | GTLSRSESRSMLSRPISLAFWIWPGTRIGE | 2057147 | 2057330 | + | 6 |
| 9 | VVWSRLSISLESADPKTSAECLMSRSPAD | GTLSRSESRSMLSRPISLAFWIWPGTRIGE | 2057150 | 2057330 | + | 6 |

| | | | | | | |
|---|---|---|---|---|---|---|
| 10 | VWSRLSISLESADPKTSAECLMSRSPAD | GTLSRSESRSMLSRPISLAFWIWPGTRIGE | 2057153 | 2057330 | + | 6 |
| 11 | LESADPKTSAECLMSRSPAD | GTLSRSESRSMLSRPISLAFWIWPGTRIGE | 2057177 | 2057330 | + | 6 |
| 12 | VPGQIQKARLIGRLSILLLSDRDKV...GDLDMRHSAEVFGSALSKLIDSLLH | TTIDEHVAEAVLRGG | 2057314 | 2057107 | - | 6 |
| 13 | VAVVALGVLGV | AAPASASASPSHEQQEVVALDPYVA...CRNGGGYPLNGICFGGRWDGAIIIV | 2057385 | 2057586 | + | 7 |
| 14 | VALGVLGVAAPASASASPSHEQQEVVALDPY | VAVTEATCRNGGGYPLNGICFGGRWDGAIIIV | 2057394 | 2057586 | + | 6 |
| 15 | MSLPSGELQVGRLCIDGWVTPRLEA...GVEHVMLALIAQGDSPTLVDRVVPV | ETVREWRDELSKVIASTKPEIASDSINVSVERE | 2057582 | 2057885 | + | 8 |
| 16 | MDEDQMVQFYERRRIYPECQSGWYP...ERVDKLVDAAEAKAYTTCEVCGVSD | STVTTSGPSWVKTLCTTHRGKIR | 2058370 | 2058022 | - | 6 |
| 17 | LHIFIVMVCF | STLVRTQWDTFCHNETFRMVEQSLYDRTQ | 2061115 | 2061235 | + | 6 |
| 18 | MGARAGSSPDNAETIPVLPDGSGSANTTERYTHRNQCPNAPQV | RPPTRNLVDRGRVASQTEAGVPVEG...AGVKLGTDPVNRLNSERGNPAAGPC | 2064946 | 2065300 | + | 7 |
| 19 | VDRGRVASQTEAGVPVEGTSGSENVAGPGGHSE | SLRRSYGEDAGVKLGTDPVNRLNSERGNPAAGPC | 2065096 | 2065300 | + | 6 |
| 20 | VCQSRELPDRRMSPVR | EAIVKACGVAMARMQE | 2065133 | 2065232 | + | 6 |
| 21 | VPVEGTSGSENVAGPGGHSESLRRSYG | EDAGVKLGTDPVNRLNSERGNPAAGPC | 2065135 | 2065300 | + | 7 |
| 22 | VNVGTPPLVPADHGFHAWSAGKVRRRLI | PVVWGRRVRSSPRSGKPTTWRRDSVPIPQ | 2065265 | 2065439 | + | 6 |
| 23 | VGTPPLVPADHGFHAWSAGKVRRRLIP | VVWGRRVRSSPRSGKPTTWRRDSVPIPQ | 2065271 | 2065439 | + | 6 |

### Results for KMS87535.1 [Streptomyces regensis] back to top

previous - next

#### Architecture

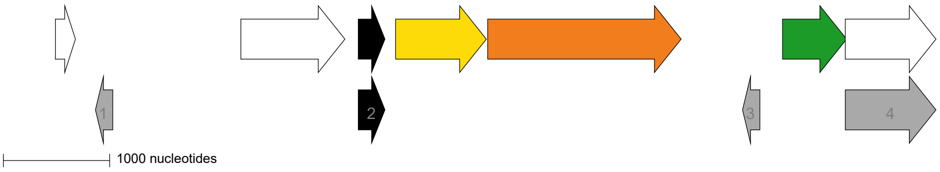

Link to nucleotide sequence

| Accession | start | end | direction | length (aa) | Pfam/HMM | name | description | E-value |
|---|---|---|---|---|---|---|---|---|
| KMS87529.1 | 0 | 183 | + | 60 | NO PFAM MATCH | - | - | - |
| KMS87530.1 | 1734 | 2718 | + | 327 | PF08028<br>TIGR04022<br>PF00441 | Acyl-CoA_dh_2<br>TIGR04022<br>Acyl-CoA_dh_1 | Acyl-CoA dehydrogenase, C-terminal domain<br>sulfur_SfnB: sulfur acquisition oxidoreductase, SfnB family<br>Acyl-CoA dehydrogenase, C-terminal domain | 1.90E-18<br>1.20E-06<br>6.30E-05 |
| KMS87531.1 | 2838 | 3099 | + | 86 | LinA | LinA | LinA (linaridin precursor) | 8.30E-15 |
| KMS87535.1 | 3194 | 4052 | + | 285 | LinG<br>LinH | LinG<br>LinH | LinG Homolog<br>LinH (Fusion of LinE/LinG) | 1.10E-105<br>1.90E-89 |
| KMS87532.1 | 4056 | 5874 | + | 605 | TIGR02204<br>TIGR02203<br>TIGR00958<br>TIGR03796<br>TIGR01846 | TIGR02204<br>TIGR02203<br>TIGR00958<br>TIGR03796<br>TIGR01846 | MsbA_rel: ABC transporter, permease/ATP-binding protein<br>MsbA_lipidA: lipid A export permease/ATP-binding protein MsbA<br>3a01208: antigen peptide transporter 2<br>NHLM_micro_ABC1: NHLM bacteriocin system ABC transporter, peptidase/ATP-binding protein<br>type_I_sec_HlyB: type I secretion system ATPase | 4.20E-103<br>4.10E-94<br>6.40E-85<br>1.50E-81<br>1.60E-78 |
| KMS87533.1 | 6837 | 7431 | + | 197 | LinL | LinL | LinL | 1.30E-62 |
| KMS87534.1 | 7423 | 8281 | + | 286 | TIGR02032<br>PF01494<br>PF01266<br>PF04820<br>TIGR01988 | TIGR02032<br>FAD_binding_3<br>DAO<br>Trp_halogenase<br>TIGR01988 | GG-red-SF: geranylgeranyl reductase family<br>FAD binding domain<br>FAD dependent oxidoreductase<br>Tryptophan halogenase<br>Ubi-OHases: ubiquinone biosynthesis hydroxylase, UbiH/UbiF/VisC/COQ6 family | 5.70E-15<br>4.90E-11<br>7.70E-11<br>7.50E-10<br>1.80E-07 |

| index | leader | core | start | end | dir | score |
|---|---|---|---|---|---|---|
| 1 | VTCQRRMPVPSSASSTRQRSAGASASSGVSVATRSRASV | ATVTSALASGVVSR | 540 | 378 | - | 7 |
| 2 | MSNVEAFANQHADNVLSPLEVTSAVGPVL | ATPAYAGYTAAAAAVFTAGLISDAV...VTPTGPVPNYGSVGDLLDMRVSSIN | 2838 | 3099 | + | 25 |
| 3 | VSSSSCSLRCVMSLRRERPDT | TTPTGTSGGKASVATCAGSAFFQATNSAAAAR | 6622 | 6460 | - | 7 |
| 4 | MRDDRPVVVAGAGIAGATVAAGLAS...LVGAGWLSVGDAAGRQDPMFLDPAT | ITLLAAQAAVPVVADLAFG | 7423 | 8281 | + | 7 |

### Results for RKN13450.1 [Streptomyces sp. DS1-2] back to top

previous - next

#### Architecture

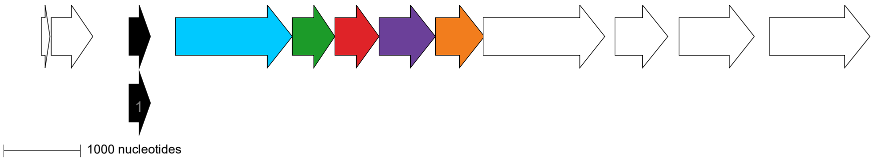

Link to nucleotide sequence

| Accession | start | end | direction | length (aa) | Pfam/HMM | name | description | E-value |
|---|---|---|---|---|---|---|---|---|
| RKN13424.1 | 0 | 111 | + | 36 | NO PFAM MATCH | - | - | - |
| WP_008745676.1 | 124 | 673 | + | 183 | INFERRED GENE | - | - | - |
| RKN13425.1 | 1140 | 1410 | + | 89 | LinA | LinA | LinA (linaridin precursor) | 1.90E-13 |
| RKN13450.1 | 1744 | 3265 | + | 506 | LinH<br>LinE<br>LinG<br>PF12697<br>TIGR02427 | LinH<br>LinE<br>LinG<br>Abhydrolase_6<br>TIGR02427 | LinH (Fusion of LinE/LinG)<br>LinE Homolog<br>LinG Homolog<br>Alpha/beta hydrolase family<br>protocat_pcaD: 3-oxoadipate enol-lactonase | 1.70E-160<br>8.60E-67<br>2.50E-66<br>4.50E-08<br>4.20E-07 |
| RKN13426.1 | 3254 | 3821 | + | 188 | LinL | LinL | LinL | 3.30E-53 |
| RKN13427.1 | 3817 | 4390 | + | 190 | TIGR00521<br>TIGR02113<br>PF02441 | TIGR00521<br>TIGR02113<br>Flavoprotein | coaBC_dfp: phosphopantothenoylcysteine decarboxylase / phosphopantothenate--cysteine ligase<br>coaC_strep: phosphopantothenoylcysteine decarboxylase<br>Flavoprotein | 5.60E-13<br>9.10E-11<br>1.40E-09 |
| RKN13428.1 | 4382 | 5120 | + | 245 | PF13649<br>PF08241<br>PF13847<br>TIGR02021<br>TIGR01934 | Methyltransf_25<br>Methyltransf_11<br>Methyltransf_31<br>TIGR02021<br>TIGR01934 | Methyltransferase domain<br>Methyltransferase domain<br>Methyltransferase domain<br>BchM-ChlM: magnesium protoporphyrin O-methyltransferase<br>MenG_MenH_UbiE: ubiquinone/menaquinone biosynthesis methyltransferase | 1.50E-15<br>2.90E-14<br>8.20E-11<br>1.30E-09<br>4.70E-09 |
| RKN13429.1 | 5116 | 5749 | + | 210 | TIGR02857<br>TIGR02673<br>TIGR01189<br>TIGR02982<br>TIGR03265 | TIGR02857<br>TIGR02673<br>TIGR01189<br>TIGR02982<br>TIGR03265 | CydD: thiol reductant ABC exporter, CydD subunit<br>FtsE: cell division ATP-binding protein FtsE<br>ccmA: heme ABC exporter, ATP-binding protein CcmA<br>heterocyst_DevA: ABC exporter ATP-binding subunit, DevA family<br>PhnT2: putative 2-aminoethylphosphonate ABC transporter, ATP-binding protein | 3.60E-25<br>6.90E-25<br>6.90E-24<br>1.20E-20<br>1.70E-20 |
| RKN13430.1 | 5745 | 7323 | + | 525 | NO PFAM MATCH | - | - | - |
| RKN13431.1 | 7443 | 8142 | + | 232 | PF02909<br>PF00440 | TetR_C_1<br>TetR_N | Tetracyclin repressor-like, C-terminal domain<br>Bacterial regulatory proteins, tetR family | 5.90E-14<br>2.10E-09 |
| RKN13432.1 | 8275 | 9265 | + | 329 | TIGR02824<br>TIGR01751<br>TIGR02817<br>TIGR02823<br>PF13602 | TIGR02824<br>TIGR01751<br>TIGR02817<br>TIGR02823<br>ADH_zinc_N_2 | quinone_pig3: putative NAD(P)H quinone oxidoreductase, PIG3 family<br>crot-CoA-red: crotonyl-CoA carboxylase/reductase<br>adh_fam_1: zinc-binding alcohol dehydrogenase family protein<br>oxido_YhdH: putative quinone oxidoreductase, YhdH/YhfP family<br>Zinc-binding dehydrogenase | 2.70E-48<br>1.30E-36<br>7.20E-28<br>1.10E-25<br>3.20E-25 |
| RKN13433.1 | 9455 | 10763 | + | 435 | TIGR00900<br>PF07690 | TIGR00900<br>MFS_1 | 2A0121: H+ Antiporter protein<br>Major Facilitator Superfamily | 7.40E-22<br>2.50E-14 |

| index | leader | core | start | end | dir | score |
|---|---|---|---|---|---|---|
| 1 | MGNFRSVETLSLSKETRKGSKMRLN...TQEFANGVLRDAAPGFHSDSEIPAM | ATPATPTAAQFVIQGSTICLVC | 1140 | 1410 | + | 34 |

### Results for SEE50185.1 [Streptomyces sp. 2112.2] back to top

previous - next

#### Architecture

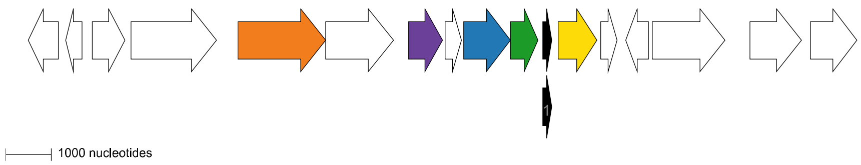

Link to nucleotide sequence

| Accession | start | end | direction | length (aa) | Pfam/HMM | name | description | E-value |
|---|---|---|---|---|---|---|---|---|
| SEE49873.1 | 4546529 | 4545866 | - | 220 | TIGR00949<br>PF01810<br>TIGR00948 | TIGR00949<br>LysE<br>TIGR00948 | 2A76: homoserine/Threonine efflux protein<br>LysE type translocator<br>2a75: L-lysine exporter | 1.40E-45<br>7.60E-44<br>1.40E-16 |
| SEE49919.1 | 4547067 | 4546698 | - | 122 | PF01906 | YbjQ_1 | Putative heavy-metal-binding | 1.40E-33 |
| SEE49956.1 | 4547274 | 4547994 | + | 239 | PF09335 | SNARE_assoc | SNARE associated Golgi protein | 5.40E-24 |
| SEE50001.1 | 4548135 | 4549995 | + | 619 | PF06738<br>PF12821 | ThrE<br>ThrE_2 | Putative threonine/serine exporter<br>Threonine/Serine exporter, ThrE | 7.50E-73<br>6.00E-19 |
| SEE50027.1 | 4550454 | 4552392 | + | 645 | TIGR02204<br>TIGR02203<br>TIGR00958<br>TIGR03796<br>TIGR03375 | TIGR02204<br>TIGR02203<br>TIGR00958<br>TIGR03796<br>TIGR03375 | MsbA_rel: ABC transporter, permease/ATP-binding protein<br>MsbA_lipidA: lipid A export permease/ATP-binding protein MsbA<br>3a01208: antigen peptide transporter 2<br>NHLM_micro_ABC1: NHLM bacteriocin system ABC transporter, peptidase/ATP-binding protein<br>type_I_sec_LssB: type I secretion system ATPase | 5.80E-100<br>2.80E-96<br>6.90E-90<br>3.50E-81<br>2.20E-80 |
| SEE50060.1 | 4552385 | 4553870 | + | 494 | TIGR02734<br>TIGR00292<br>PF13450<br>TIGR03467<br>PF01593 | TIGR02734<br>TIGR00292<br>NAD_binding_8<br>TIGR03467<br>Amino_oxidase | crtI_fam: phytoene desaturase<br>TIGR00292: thiazole biosynthesis enzyme<br>NAD(P)-binding Rossmann-like domain<br>HpnE: squalene-associated FAD-dependent desaturase<br>Flavin containing amine oxidoreductase | 8.90E-09<br>4.70E-08<br>5.30E-08<br>7.50E-08<br>3.40E-07 |
| SEE50099.1 | 4554187 | 4554946 | + | 252 | PF13649<br>PF08241<br>PF13489<br>PF08242<br>PF13847 | Methyltransf_25<br>Methyltransf_11<br>Methyltransf_23<br>Methyltransf_12<br>Methyltransf_31 | Methyltransferase domain<br>Methyltransferase domain<br>Methyltransferase domain<br>Methyltransferase domain<br>Methyltransferase domain | 5.70E-17<br>1.90E-12<br>1.30E-10<br>5.60E-10<br>5.60E-09 |
| SEE50142.1 | 4554988 | 4555336 | + | 115 | PF13518<br>PF01527<br>PF13551<br>PF13011<br>PF13384 | HTH_28<br>HTH_Tnp_1<br>HTH_29<br>LZ_Tnp_IS481<br>HTH_23 | Helix-turn-helix domain<br>Transposase<br>Winged helix-turn helix<br>leucine-zipper of insertion element IS481<br>Homeodomain-like domain | 4.20E-09<br>2.50E-06<br>1.30E-05<br>1.00E-04<br>4.20E-04 |
| SEE50185.1 | 4555408 | 4556428 | + | 339 | LinE<br>LinH<br>PF12697<br>TIGR02427<br>PF00561 | LinE<br>LinH<br>Abhydrolase_6<br>TIGR02427<br>Abhydrolase_1 | LinE Homolog<br>LinH (Fusion of LinE/LinG)<br>Alpha/beta hydrolase family<br>protocat_pcaD: 3-oxoadipate enol-lactonase<br>alpha/beta hydrolase fold | 4.70E-104<br>4.10E-53<br>3.10E-15<br>8.90E-12<br>7.20E-10 |
| SEE50226.1 | 4556424 | 4557012 | + | 195 | LinL | LinL | LinL | 2.50E-56 |
| SEE50268.1 | 4557139 | 4557322 | + | 60 | LinA | LinA | LinA (linaridin precursor) | 1.80E-13 |
| SEE50309.1 | 4557467 | 4558322 | + | 284 | LinG<br>LinH<br>TIGR04033 | LinG<br>LinH<br>TIGR04033 | LinG Homolog<br>LinH (Fusion of LinE/LinG)<br>export_SdpB: antimicrobial peptide system protein, SdpB family | 3.00E-111<br>3.70E-79<br>2.10E-04 |
| SEE50354.1 | 4558387 | 4558747 | + | 119 | NO PFAM MATCH | - | - | - |
| SEE50397.1 | 4559423 | 4558928 | - | 164 | PF00719 | Pyrophosphatase | Inorganic pyrophosphatase | 2.50E-53 |
| SEE50437.1 | 4559509 | 4561117 | + | 535 | PF02113<br>TIGR00666 | Peptidase_S13<br>TIGR00666 | D-Ala-D-Ala carboxypeptidase 3 (S13) family<br>PBP4: D-alanyl-D-alanine carboxypeptidase/D-alanyl-D-alanine-endopeptidase | 6.00E-76<br>6.80E-58 |
| SEE50474.1 | 4561665 | 4562784 | + | 372 | TIGR03883<br>TIGR03624<br>PF10103 | TIGR03883<br>TIGR03624<br>Zincin_2 | DUF2342_F420: uncharacterized protein, coenzyme F420 biosynthesis associated<br>TIGR03624: putative hydrolase<br>Zincin-like metallopeptidase | 7.30E-142<br>1.00E-125<br>3.80E-121 |
| SEE50514.1 | 4562976 | 4563984 | + | 335 | TIGR02432<br>PF01171<br>PF09179 | TIGR02432<br>ATP_bind_3<br>TilS | lysidine_TilS_N: tRNA(Ile)-lysidine synthetase<br>PP-loop family<br>TilS substrate binding domain | 3.60E-43<br>6.60E-38<br>5.30E-12 |

| index | leader | core | start | end | dir | score |
|---|---|---|---|---|---|---|
| 1 | MSVVANFANTELADVNPGRIGNDATPTM | LTPLAALATPEGVAVTSATAYALHEVTKDIQR | 4557139 | 4557322 | + | 33 |
